# Tutorial: Assessing metagenomics software with the CAMI benchmarking toolkit

**DOI:** 10.1101/2020.08.11.245712

**Authors:** Fernando Meyer, Till-Robin Lesker, David Koslicki, Adrian Fritz, Alexey Gurevich, Aaron E. Darling, Alexander Sczyrba, Andreas Bremges, Alice C. McHardy

**Affiliations:** Computational Biology of Infection Research, Helmholtz Centre for Infection Research, Braunschweig, Germany; German Center for Infection Research (DZIF), Braunschweig, Germany; Computer Science and Engineering, Biology, and The Huck Institutes of the Life Sciences, Penn State University, State College, PA, USA; Center for Algorithmic Biotechnology, St. Petersburg State University, St. Petersburg, Russia; The ithree institute, University of Technology Sydney, Sydney, Australia; Faculty of Technology and Center for Biotechnology, Bielefeld University, Bielefeld, Germany

## Abstract

Computational methods are key in microbiome research, and obtaining a quantitative and unbiased performance estimate is important for method developers and applied researchers. For meaningful comparisons between methods, to identify best practices, common use cases, and to reduce overhead in benchmarking, it is necessary to have standardized data sets, procedures, and metrics for evaluation. In this tutorial, we describe emerging standards in computational metaomics benchmarking derived and agreed upon by a larger community of researchers. Specifically, we outline recent efforts by the Critical Assessment of Metagenome Interpretation (CAMI) initiative, which supplies method developers and applied researchers with exhaustive quantitative data about software performance in realistic scenarios and organizes community-driven benchmarking challenges. We explain the most relevant evaluation metrics to assess metagenome assembly, binning, and profiling results, and provide step-by-step instructions on how to generate them. The instructions use simulated mouse gut metagenome data released in preparation for the second round of CAMI challenges and showcase the use of a repository of tool results for CAMI data sets. This tutorial will serve as a reference to the community and facilitate informative and reproducible benchmarking in microbiome research.

## Introduction

Since the release of the first shotgun metagenome from the Sargasso Sea by metagenomics (see glossary in Table 1) pioneer Craig Venter^1^, the field has witnessed an explosive growth of data and methods. Microbiome data repositories^2,3^ host hundreds of thousands of data sets and numbers are still rising rapidly.

**Table 1:** Glossary

| Term | Definition |
| --- | --- |
| Metagenomics | A set of techniques for recovering and sequencing of the genetic material of microbial communities and their functional and taxonomic characterization. |
| Benchmarking | Systematic comparison of (computational) techniques using performance metrics in specific scenarios. |
| Assembly | Reconstruction of complete or partial genomes or DNA sequence fragments, often by merging sequence reads into longer pieces called contigs. |
| Binning | Clustering or classification of sequences or contigs into bins representing genomes (genome binning) or taxa (taxonomic binning) of the underlying microbial community. |
| Profiling | Microbial community characterization from a metagenomic sample in terms of presence and absence of taxa and their relative abundances. |
| Coverage | Number of reads that cover a certain genomic position. |
| Docker | A software tool designed to make it easy to distribute and run applications by using software packages (containers) and operating system-level virtualization. |

Metagenomics created new computational challenges, such as reconstructing the genomes of community members from a mixture of reads originating from potentially thousands of microbial, viral, and eukaryotic taxa^4^. These taxa differ in their relatedness to each other, are often absent from sequence databases, and present at varying abundances. Genomes can be reconstructed by metagenome assembly, which creates longer, contiguous sequence fragments, followed by binning, which is usually a clustering method placing fragments into genome bins. There have been spectacular successes in recovering thousands of metagenome assembled genomes, or MAGs, for uncultured taxa^5–7^. Identifying the taxa and their abundances for a community is known as taxonomic profiling, while taxonomic binners assign taxonomic labels to individual sequence fragments. Both tasks are challenging particularly for lower taxonomic ranks^8^. Another challenge is the *de novo* assembly of closely related genomes (>95% average nucleotide identity)^8^. Finally, fragmentary assemblies with many short contigs obtained from short read sequence data in metagenomics have required adaptation of gene finding methods and complicate operon-level functional analyses of genes. The maturation of long-read sequencing technologies^9,10^, which for many years were characterized by low throughput, high cost, and high error rates, has sparked further development and is expected to lead to better solutions for some of these challenges.

### The relevance of standards for performance evaluation and benchmarking

Methodological development is oftentimes accompanied by performance evaluations. This has historically been done on an *ad hoc* basis by developers, often using different data sets and performance metrics, which are both critical choices regarding performance evaluation. This practice made it difficult to compare results across publications and to identify suitable techniques for specific data sets and tasks. It also made performance benchmarking for developers very tedious and ineffective. For instance, performance might differ substantially for reference-based methods using public databases across data sets, depending on evolutionary divergence between the sampled and database taxa^8^. Similarly, organismal complexity, strain-level diversity, realistic community genome abundance distributions, the presence of non-bacterial genomic information, as well as sequencing error profiles of data sets may affect method performances, to list some factors.

It became evident, as in other fields^11–13^, that standards would greatly facilitate comparisons across methods and articles and univocal determination of appropriate solutions and open challenges. To satisfy this need, CAMI, the community-driven initiative for the Critical Assessment of Metagenome Interpretation, was founded in 2014 by A. Sczyrba, T. Rattei, and A.C. McHardy^14^ during the metagenomics programme at the Isaac Newton Institute in Cambridge^15^. CAMI design decisions are based on feedback gathered in community workshops, which ensures inclusion of a wide range of expert inputs and establishes a community consensus. By regularly interacting with scientists in workshops, hackathons and at conferences, such as the Microbiome track of ISMB, CAMI aims to identify and implement best practices for benchmarking in microbiome research, including (i) key properties of benchmark data sets (see also^16,17^ for an overview of general benchmarking practices), (ii) appropriate performance metrics for different tasks, (iii) benchmarking procedures, i.e. how to run benchmarking challenges, and (iv) performance evaluation procedures, to allow the most realistic, fair, and unbiased assessment. Reproducibility and reusability (v) have been identified as the fifth key criterion. We provide further details on these key aspects below.

The first CAMI challenge took place in 2015 and provided an extensive performance overview for commonly used data processing methods, namely assembly, genome and taxonomic binning, and taxonomic profiling^8^. The six benchmark data sets reflecting a range of complexities have since been used extensively for further benchmarking in the field. These include three “toy” data sets created from public data and provided before the challenge, as well as three challenge data sets derived exclusively from genomic data that were not publicly available at the time. These data are now in public databases. Further benchmarking studies have also provided valuable insights^18–21^. The second CAMI challenge (CAMI II) was launched in 2019 and offered challenges for the same tasks on two large, multi-sample data sets reflecting specific environments (marine, rhizosphere) and an extremely high strain diversity data set (strain madness). In addition, a clinical pathogen detection challenge was offered. The challenges are expected to provide insights on important questions such as the potential of long-read data for metagenomics^22^.

### Benchmark data sets

Benchmark data sets should be as realistic and representative for real metaomics data as possible. For CAMI challenges, experimental groups contribute unpublished genomes, including some organisms from poorly characterized phyla without any genomes of close relatives publicly available. These genomes are used for benchmark data creation and published only after the challenge. Because many taxa present in real environmental samples have unknown cultivation conditions and no isolate genomes are available in reference databases, measuring performance on novel organisms is essential. This is particularly true for a comprehensive evaluation of reference-based methods such as taxonomic profilers and binners, which perform best for genomes closely related to those in public databases^8^. The challenge data sets have been created from these (and public, in CAMI II) genomes with the CAMISIM microbial community and metagenome simulator^23^. This allows to incorporate many key properties in data sets, such as varying experimental designs (number of samples, sequencing depth, insert sizes, type of experiment, such as differential abundances, time series), sequencing technologies and community properties (organismal complexity, different genome abundance distributions, strain diversity, taxa from different domains of life, viruses, mobile circular elements). An alternative way to create benchmark data is to sequence lab-created DNA mixtures as in^24^, which would enable a more realistic assessment of technical variation and biases introduced in data generation. However, creating communities with realistic organismal complexities for many environments, with hundreds to thousands of genomes at highly varying abundances, is currently impractical. All CAMI benchmark data sets are made available after the challenges with Digital Object Identifiers^25^ (Table 2).

**Table 2:** CAMI benchmark data sets and respective Digital Object Identifiers (DOI). All data sets are also downloadable from the CAMI portal at https://data.cami-challenge.org/.

| **CAMI benchmark data sets** | **DOI** |
| --- | --- |
| CAMI I: low, medium, high complexity, and “toy” data sets | 10.5524/100344 |
| CAMI II: mouse gut “toy” data set | 10.4126/FRL01-006421672 |
| CAMI II: marine, strain madness, rhizosphere, and pathogen detection challenge data sets | DOI available after challenges |

### Metrics for performance evaluation

Choosing the appropriate (combination) of metrics for comparing method performances is a key task in benchmarking that directly influences the ranking of methods. The metrics used in CAMI challenges^8^ are decided on in public workshops and reassessed regularly. They should be easy to interpret and meaningful to both developers and applied scientists. A comprehensive assessment is achieved by including multiple metrics that highlight strengths of different approaches – see below. Furthermore, assessing properties such as runtime, disk space, and memory consumption is important.

### Advantages of benchmarking challenges

Challenges provide insights into method performances, suggesting best practices as well as identifying open problems in the field. They can also further the development and adoption of standards, such as data input and output formats, or choice of reference data sets, such as the NCBI taxonomy. Once standards are realized, benchmarking competitions offer a low-effort opportunity for extensive benchmarking, as data sets, other method results, and evaluation methods do not have to be created by the developer of a new metagenome analysis method.

Some participants might worry about publishing poor performances, which is why CAMI challenge participants can opt out of result publication and use them only for their own benefit. Defining the evaluation metrics is also open for the field, thus all labs participating in these discussions can contribute to the challenge evaluation. Participants can thus suggest and define metrics that highlight the expected benefits of their techniques with these simultaneously being subjected to peer group review. To ensure a maximum of objectivity in these evaluations, CAMI challenges are performed blinded in two ways. The standard of truth for the challenge data set is only provided after challenges end, preventing performance optimization in any way on these particular data sets. Challenge data sets include many genomes that will only become publicly available after the challenge. “Toy” data sets, where a standard of truth is made available at the outset, are provided before the actual challenges to enable teams to familiarize themselves with the data structure and its properties. The evaluation of the different challenge submissions is also performed blindly, such that the evaluation panel does not know the names and information about the submitted techniques, to tackle evaluator biases. Evaluations are open to anyone wishing to participate and a consensus is reached in a workshop with a group of experts.

### Reproducibility and FAIR principles

Imagine running a benchmarking contest and identifying the top performing technique by key criteria, potentially representing the new state-of-the-art for future studies. However, the submitting team has unfortunately lost track of the software version and parameter settings used, and is unable to reproduce its own results. To avoid such issues, reproducibility has been elected as a core principle in CAMI, for all steps of benchmarking, from data generation with CAMISIM^23^, to running software benchmarked in the contest, and to evaluating results. Evaluation metrics are extensively tested and implemented in the MetaQUAST^26^, AMBER^27^, and OPAL^28^ benchmarking packages (see Table 3) available via Bioconda^29^. All software released by CAMI is open source under appropriate licenses such as Apache 2 or GPL. A key result of the first challenge was that parameter settings substantially affect program performances. A minimal requirement for public CAMI challenge results is therefore documenting the exact program versions and command line calls or, even better, using a workflow manager such as GNU make, Snakemake^30^, Nextflow^31^, or CWL^32^. The ideal, though time-consuming, approach is to containerize the program, e.g. in Docker, Bioboxes^33^, or BioContainers^34^, as well as to document and bundle dependencies to facilitate installation with pip or Bioconda^29^.

**Table 3:** CAMI benchmarking software

| Software | Description |
| --- | --- |
| CAMISIM<sup>23</sup> | A microbial community and metagenome simulator that models different microbial abundance profiles, multi-sample time series, and differential abundance studies, real and simulated strain-level diversity, and generates second and third-generation sequencing data from taxonomic profiles or *de novo*. CAMISIM was used to generate several benchmark data sets for CAMI challenges. |
| MetaQUAST<sup>26</sup> | A quality assessment tool for metagenome assembly evaluation. It computes various quality metrics based on alignment of assemblies to a standard of truth or close reference genomes. The first option is used in CAMI. |
| AMBER<sup>27</sup> | Software for the comparative assessment of genome reconstructions and taxonomic assignments from metagenome benchmark data sets. It calculates performance metrics such as (rank-specific taxon) bin completeness and purity, average Rand index, assignment accuracy, and comparative visualizations used in CAMI challenges. |
| OPAL<sup>28</sup> | A tool for computing performance metrics and creating visualizations for assessing taxonomic metagenome profilers. The metrics include presence-absence metrics (number of true and false positives, false negatives, completeness, purity, F1 score, Jaccard index) as well as abundance metrics such as UniFrac, L1 norm and the Bray-Curtis distance. |
| Bioboxes<sup>33</sup> | Docker containers with standardized interfaces facilitating interchange of software in bioinformatics pipelines, distribution of specific software versions with predefined parameter settings, and therefore reproducibility of results and benchmarking. The Bioboxes standard was used to containerize the methods benchmarked in the CAMI I challenges and are continuously used along with BioContainers<sup>34</sup> and workflow and package managers such as Snakemake<sup>30</sup>, Nextflow<sup>31</sup>, and Bioconda<sup>29</sup>. |

To maximize the scientific value, not only the methods, but also all data required for reproducing and building on the results of a study should be made available. CAMI commits to the FAIR (Findable, Accessible, Interoperable, Reusable) principles for scientific data management and stewardship^35^. CAMI benchmark and reference data sets, program results, and computed metrics are provided with DOIs on Zenodo (https://zenodo.org/communities/cami) and GigaDB^25^. This improves reusability and sustainability of the efforts, as others can directly build on a study, for instance by adding their own method’s results to the existing results of a benchmarking effort, or adding calculation of new metrics to a benchmark study for more sophisticated interpretation. A schematic representation of CAMI’S benchmarking workflow is shown in Fig. 1. In the following, we demonstrate this principle of convenient benchmarking by extending previous results for the four software categories (assembly, genome and taxonomic binning, and profiling) benchmarked on the CAMI II multi-sample mouse gut data set, creating a flexible benchmarking resource for individual studies.

**Fig. 1:**
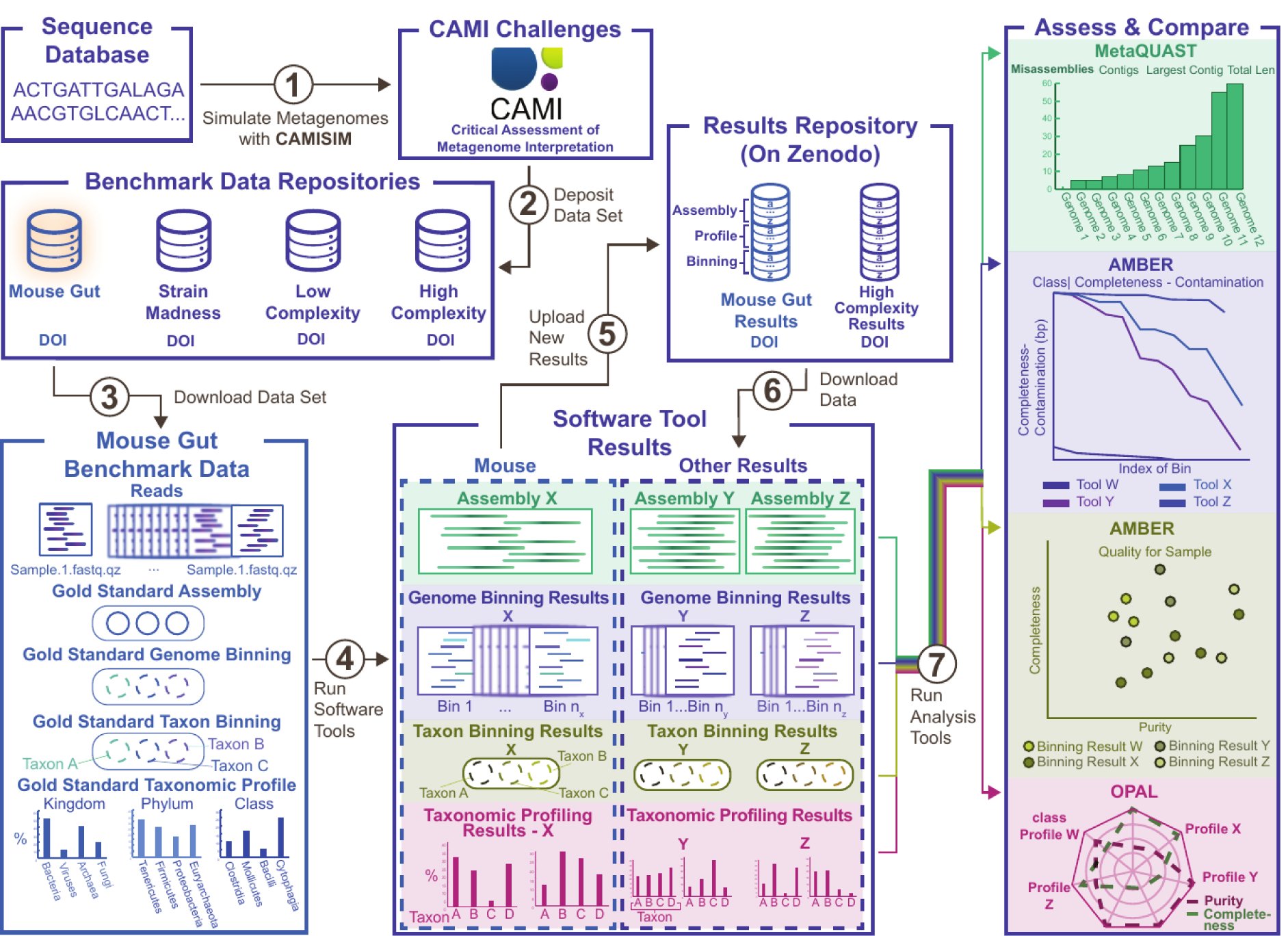
CAMI benchmarking workflow. The initial step is the simulation of metagenome data from a sequence database with CAMISIM^23^ (1), which includes the microbial community design and generation of standards of truth. The simulated metagenome data are stored in benchmark data repositories with Digital Object Identifiers (DOIs) (2) or temporarily without DOIs for ongoing CAMI challenges, as the standards of truth are only provided after the challenges. The data can then be downloaded (3) and software tools like metagenome assemblers, genome and taxonomic binners, and profilers run on the data (4). This leads to the creation of a pool of software tool results. These results can be submitted to an ongoing challenge or uploaded to a public repository, like Zenodo (5). Already existing results can be downloaded (6) and integrated in benchmark analyses with MetaQUAST^26^, AMBER^27^, and OPAL^28^ (7).

## Benchmarking demonstration

We demonstrate how to benchmark in practice using the benchmarking software and standards (Table 3) from previous studies on CAMI data sets for different computational challenges. We analyse the mouse gut metagenome “toy” data set^23^ provided to prepare for CAMI II (Table 2), starting below with a description of its simulation. Analyses of this data set with several taxonomic profiling and assembly methods were previously described^23,28^. The benchmarked assemblers, taxonomic and genome binners, and taxonomic profilers were chosen based on popularity and performance in the first CAMI challenge^8^. All method results for this and other benchmark data sets can be obtained from a new resource on Zenodo at https://zenodo.org/communities/cami, and curated metadata is provided at https://github.com/CAMI-challenge/data. Users can continue to add results to these repositories, thus building a growing method result collection for benchmarking.

### Simulation of benchmark data set

The mouse gut metagenome “toy” data set was generated with CAMISIM version 0.2^23^ (Table 3) using a microbial community genome abundance distribution modelled from 791 public prokaryotic genomes marked as at least “scaffolds” in the NCBI RefSeq^36^. They comprise 8 phyla, 18 classes, 26 orders, 50 families, 157 genera, and 549 species. The community genome abundance distribution matches as close as possible the 16S taxonomic profiles for 64 mouse gut samples. As such, this data set allows us to assess how well sequenced community members can be characterized with different techniques from metagenomes of similar communities. In each of the 64 samples, 91.8 genomes are represented on average. Both long (PacBio) and short-read (Illumina HiSeq 2000) metagenome sequencing data are available, with 5 GB of sequences per sample leading to an average genome coverage of 4.7^23^. The runtime to generate these data was approximately 3 weeks using eight CPU cores of a computer with an AMD Opteron 6378 CPU and 968 GB of main memory.

CAMISIM can be installed according to the instructions at https://github.com/CAMI-challenge/CAMISIM/ or using Docker with the command:

~~~
docker pull cami/camisim
~~~

To generate the mouse gut data set, the following command was used:

~~~
./metagenome_from_profile -p profile.biom -o out/
~~~

`profile.biom` is a BIOM^37^ file storing the microbial community genome abundance distribution for the 64 samples. It can be obtained together with the data set (Table 2). Per default, CAMISIM simulates 5 GB of sequences per sample.

If CAMI benchmark data generated with CAMISIM have been downloaded, the following files and folders should appear:

- One folder per sample
  - Reads (anonymized and shuffled) as FASTQ
  - Contigs (gold standard assembly) as FASTA
  - Gold standard mappings (binning) in BAM and CAMI formats (see format specifications at https://github.com/CAMI-challenge/contest_information)
- For multi-sample simulations:
  - File containing contigs (gold standard assembly) as FASTA
  - File containing gold standard mappings (binning and profiling) in CAMI format
- Profiling gold standard per sample in CAMI format
- One folder (called “source genomes”) containing all reference genome sequences as FASTA
- One folder (called “distributions”) containing files with the absolute abundances per genome for every sampled microbial community
- One folder (called “internal”) containing the input metadata and a list of unused genomes
- Metadata (CAMISIM .ini config file)

### Assembly

Cross-sample co-assemblies of the first 10 of 64 metagenome samples were performed with MEGAHIT^38^ versions 1.0.3, 1.1.3, and 1.2.9, and metaSPAdes^39^ 3.13.0, as the computer main memory was insufficient to run metaSPAdes on more than 10 samples. The choice of the first 10 samples was analogous to the CAMI II challenge specifications. All results and commands used are available on Zenodo (Supplementary Table 1). The computer specifications, memory usage, and runtimes are available in Supplementary Tables 2 and 3.

Assemblies were evaluated by mapping them against the gold standard assembly, defined as the fraction of the genome covered by at least one read in the set of analyzed samples, using MetaQUAST^26^ 5.0.2. The gold standard genomes are known through the simulation with CAMISIM and provided to MetaQUAST for the evaluation. In case the underlying genomes are unknown, such as when assessing *de novo* assemblies from less studied environments, reference-free methods^40–42^ can be considered.

MetaQUAST can be installed with Bioconda using the command:

~~~
conda create --name quast quast
~~~

This requires Conda to be installed and the Bioconda channel configured – see https://bioconda.github.io/user/install.html for details. Other installation methods are described in the MetaQUAST GitHub repository at https://github.com/ablab/quast/. To run MetaQUAST, type:

~~~
conda activate quast
metaquast -r /path/to/set0-9/ref-genomes \
-t 24 --unique-mapping --no-icarus -o /path/to/output_dir \
-l megahit-103-df,megahit-113-df,megahit-113-ml,\
megahit-113-ms,megahit-129-df,metaSPAdes \
/path/to/megahit103-Sample0-9-default/final.contigs.fa \
/path/to/megahit113-Sample0-9-default/final.contigs.fa \
/path/to/megahit113-Sample0-9-meta-large/final.contigs.fa \
/path/to/megahit113-Sample0-9-meta-sensitive/final.contigs.fa \
/path/to/megahit129-Sample0-9-default/final.contigs.fa \
/path/to/metaSPAdes3130-Sample0-9/contigs.fasta
~~~

For evaluating assembly quality, we rely on the metrics provided by MetaQUAST. Table 4 shows the metrics we focus on here, whereas Supplementary results (report.html) shows all metrics computed by MetaQUAST. The **genome fraction** is the total number of aligned bases in the reference, divided by the genome size; **#contigs** is the number of contigs in the assembly; **NG50** is the contig length such that contigs of that length or longer covers half (50%) of the bases of the reference genome; and **NGA50** is NG50 such that the lengths of aligned blocks are counted instead of contig lengths. Performance values are calculated for the whole assembly vs. the combined reference (i.e. concatenation of all provided references).

**Table 4:** MetaQUAST assembly benchmarking metrics

|  |  |  |  |  |  |  |
| --- | --- | --- | --- | --- | --- | --- |
|  | Worst Median Best |  |  |  |  |  |
| **Genome statistics** | **MEGAHIT 1.0.3 df** | **MEGAHIT 1.1.3 df** | **MEGAHIT 1.1.3 ml** | **MEGAHIT 1.1.3 ms** | **MEGAHIT 1.2.9 df** | **metaSPAdes 3.13.0** |
| + Genome fraction (%) | 23.507 | 26.164 | 26.039 | 26.292 | 26.691 | 23.262 |
| + Duplication ratio | 1.023 | 1.037 | 1.046 | 1.05 | 1.034 | 1.017 |
| + Largest alignment | 354 703 | 904 953 | 859 640 | 753 008 | 787 657 | 1 034 619 |
| + Total aligned length | 436 725 459 | 492 514 960 | 493 969 107 | 500 306 789 | 500 856 984 | 429 280 747 |
| + NGA50 | ... | ... | ... | ... | ... | ... |
| + LGA50 | ... | ... | ... | ... | ... | ... |
| **Misassemblies** |  |  |  |  |  |  |
| + # misassemblies | 5770 | 8685 | 5336 | 9381 | 8807 | 3488 |
| + Misassembled contigs length | 10 879 967 | 43 068 359 | 34 576 388 | 56 221 107 | 50 536 067 | 25 409 676 |
| **Mismatches** |  |  |  |  |  |  |
| + # mismatches per 100 kbp | 542.07 | 580.14 | 887.27 | 945.71 | 585.26 | 405.65 |
| + # indels per 100 kbp | 2.39 | 4.17 | 3.92 | 4.75 | 4.3 | 2.57 |
| + # N's per 100 kbp | 0 | 0 | 0 | 0 | 0 | 0 |
| **Statistics without reference** |  |  |  |  |  |  |
| + # contigs | 225 585 | 220 757 | 278 807 | 282 136 | 225 167 | 174 693 |
| + Largest contig | 354 703 | 904 953 | 859 640 | 754 056 | 788 697 | 1 034 619 |
| + Total length | 438 032 656 | 494 653 238 | 496 722 592 | 503 491 159 | 503 073 431 | 430 847 014 |
| + Total length (>= 1000 bp) | 342 669 622 | 399 682 035 | 368 806 791 | 372 764 886 | 405 211 262 | 354 794 894 |
| + Total length (>= 10000 bp) | 154 362 921 | 228 640 882 | 192 818 790 | 198 110 070 | 236 255 195 | 225 930 387 |
| + Total length (>= 50000 bp) | 38 821 616 | 102 990 325 | 82 724 532 | 83 865 010 | 106 551 070 | 119 684 054 |

Overall, the performance of the MEGAHIT and MetaSPAdes assemblers is quite similar. MEGAHIT version 1.0.3 shows poor performance for high coverage (i.e. high abundance) genomes. This effect has been described for earlier versions of MEGAHIT before^8^. The more recent versions of MEGAHIT (1.1.3 and 1.2.9) handle high coverage genomes much better and show similar performance to MetaSPAdes. For coverages of 16 and above, the fraction of the recovered genomes is above 75% with some outliers for coverage higher than 250x. The NGA50 metric shows similar performance for MEGAHIT and metaSPAdes, reaching 32 kb and more for coverage of 32x and above (Fig. 2a-c). MetaSPAdes delivers fewer fragmented assemblies (fewer contigs and higher NGA50, Fig. 2d-e) than the newer MEGAHIT versions with only slightly lower genome fraction (Fig. 2d).

**Fig. 2:**
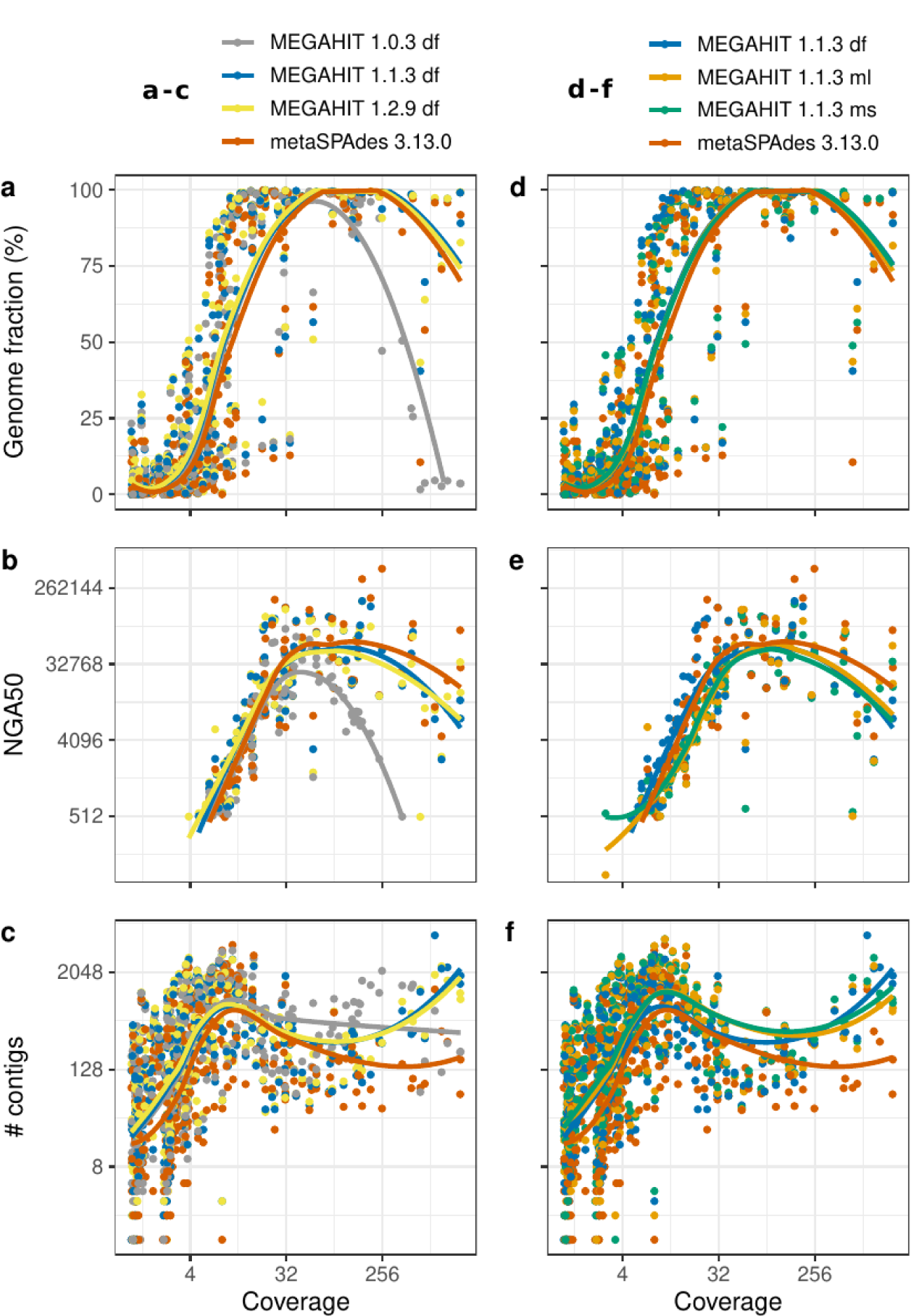
Assessing metagenome cross-sample assembly quality with MetaQUAST for the CAMI II mouse gut data set. **a**-**c** Genome-wide MetaQUAST metrics for assemblies generated with MEGAHIT versions 1.0.3, 1.1.3, 1.2.9 and metaSPAdes 3.13.0 vs. sum of read coverages for individual genomes (dots) in ten cross-sample gold standard assemblies. The higher the genome fraction and NGA50, the better is assembly quality. Higher #contigs can indicate a higher amount of assembled data, but also more fragmented assemblies, whereas lower #contigs can indicate aggressive traversal of repeats by an assembler leading to incorrect junction of sequence fragments and thus misassemblies. **d**-**f** MetaQUAST metrics for assemblies generated with MEGAHIT 1.1.3 and metaSPAdes 3.13.0. All lines are fitted with local regression using the R stats::loess function.

When assessing different settings for MEGAHIT version 1.1.3 (Fig. 2d-f), smaller, but notable differences were found. For instance, the settings “meta-sensitive” (ms) and “meta-large” (ml) delivered higher genome fractions for low coverage genomes, at the cost of higher genome fragmentation rates (decreased NGA50 and more contigs).

### Genome binning

Genome binning can be seen as a clustering problem, where sequences are grouped into bins without taxon labels. We reconstructed genome bins from the cross-sample gold standard assembly with the popular binners MaxBin 2.2.7^43^, MetaBAT 2.12.1^44^, CONCOCT 1.0.0^45^, and DAS Tool 1.1.2^46^. DAS Tool combines the genome bins of individual methods to further improve bin quality. All results and commands used are available on Zenodo (Supplementary Table 4). Runtimes and memory usage are provided in Supplementary Table 5. Binning quality was evaluated with AMBER 2.0.1^27^ (Table 3), which computes binning performance metrics for metagenome data with a ground truth available. To reproduce the evaluation, the binning results must first be downloaded from Zenodo, then AMBER installed using Bioconda:

~~~
conda create --name amber cami-amber
~~~

Other installation methods are described in https://github.com/CAMI-challenge/AMBER/. To run AMBER, type:

~~~
conda activate amber
amber.py --gold_standard_file /path/to/cami2_mouse_gut_gsa_pooled.binning \
/path/to/cami2_mouse_gut_maxbin2.2.7.binning \
/path/to/cami2_mouse_gut_metabat2.12.1.binning \
/path/to/cami2_mouse_gut_concoct1.0.0.binning \
/path/to/cami2_mouse_gut_dastool1.1.2.binning \
--labels “MaxBin 2.2.7, MetaBAT 2.12.1, CONCOCT 1.0.0, DAS Tool 1.1.2” \
--genome_coverage /path/to/cami2_mouse_gut_average_genome_coverage.tsv \
--output_dir /path/to/output_dir
~~~

File `cami2_mouse_gut_average_genome_coverage.tsv` above contains the average coverage of the genomes in the CAMI II mouse gut data set and is also available on Zenodo (Supplementary Table 4). This file is optional and used by AMBER to generate performance plots relative to the average genome coverage (Fig. 3a,b).

**Fig. 3:**
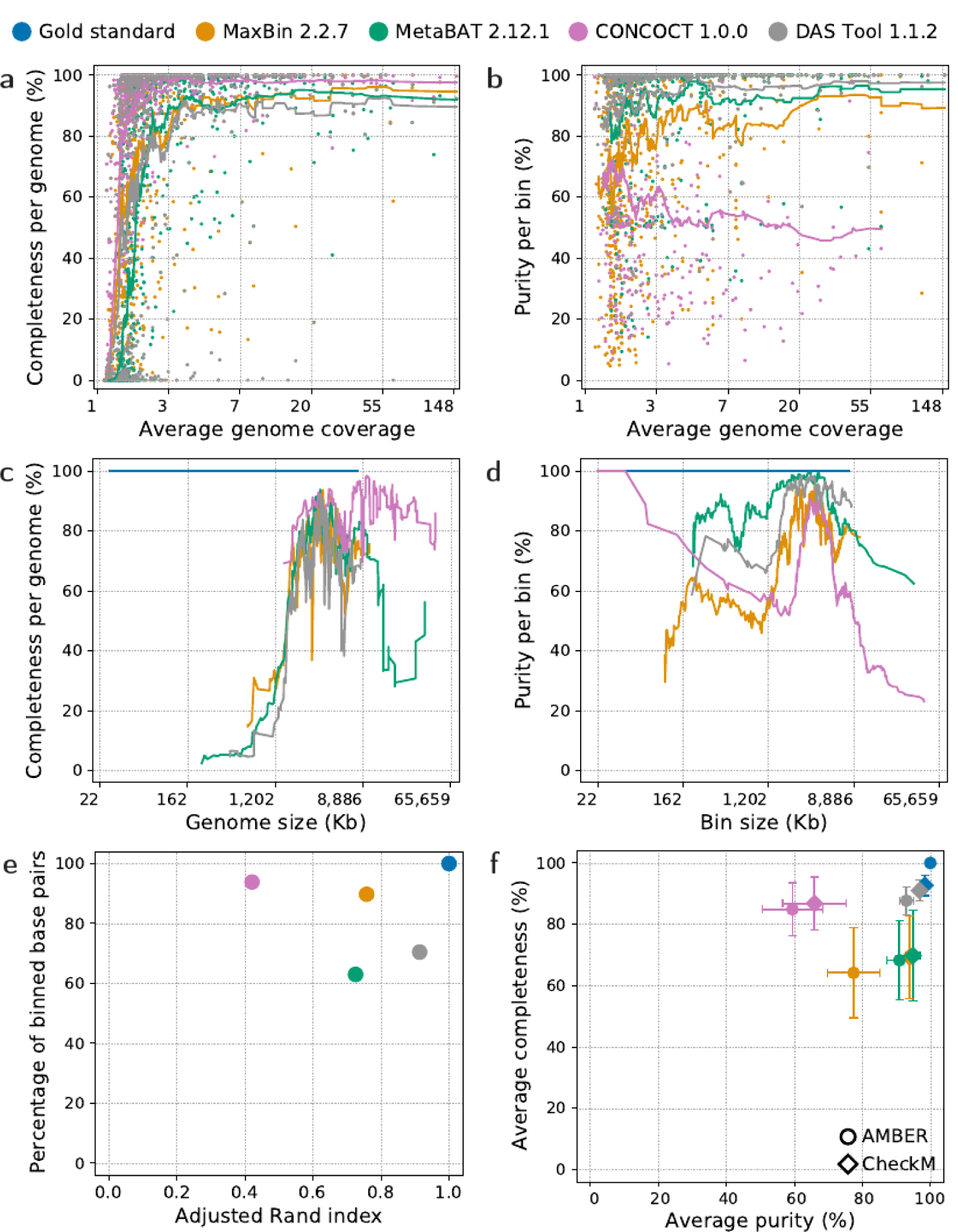
Assessing genome binners on the gold standard assembly of the CAMI II mouse gut data set. **a** Average genome coverage (x axis) vs. completeness per genome (y axis). **b** Average genome coverage (x axis) vs. purity per bin (y axis). The lines in **a** and **b** show the rolling average completeness or purity over 50 bins. **c** Genome size in thousands of bp (x axis) vs. completeness per genome (y axis). **d** Bin size in thousands of bp (x axis) vs. purity per bin (y axis). **e** Adjusted Rand index (x axis) vs. percentage of assigned base pairs (y axis). **f** Average purity (x axis) vs. average completeness (y axis) of all predicted bins per method assessed with AMBER (circles) and CheckM (diamonds), with the whiskers showing the variance. All metrics, except genome and bin sizes, range between 0% (worst) and 100% (best).

In the evaluation of genome binning, several metrics are often jointly assessed. For each genome, **completeness**, or recall, is evaluated from the predicted bin containing the largest number of base pairs (bp) of the genome. It is the number of bp (or contigs) of the genome in that bin divided by the genome size (in bp or contigs). Sequences of that genome assigned to other bins are considered false positives for those bins. Completeness can be zero, in case no part of a genome has been binned by the respective binner. **Purity** denotes how “clean” predicted bins are in terms of their assigned content. It is computed as the fraction of contigs, or bp, coming from one genome, for the most abundant genome in that bin. **Contamination** is defined as 100% minus purity. As genomes can differ in their abundances, it is also common to consider sample-wise metrics, such as the overall **percentage of assigned bp** and the **adjusted Rand index** (ARI) on that assigned fraction. The ARI reflects the overall resolution of the underlying ground truth genomes by a binner on the binned part of the sample. The ARI gives more importance to “large” bins, i.e. bins of large and/or abundant genomes, than averaging over completeness and purity, where each gold standard genome (for completeness) and predicted bin (for purity) contributes the same, irrespective of its size. In the following, all evaluations are based on base pair counts.

Completeness was high for all methods, and highest for CONCOCT. Binners recovered the abundant genomes better, with average completeness above 90% for genomes at more than 3-fold coverage (Fig. 3a). Purity was also high (Fig. 3b), except for CONCOCT, and highest for MetaBAT, which was further improved by DAS Tool. Completeness was above 90% for predicted genomes bins with an average of 3.5 to 4.6 million bp for most binners and 11.4 million bp for CONCOCT, which along with MetaBAT predicted bins that were larger than their true sizes (Fig. 3c,d). Purity was above 90% for predicted genomes bins with an average of 2.6 to 3.5 million bp (Fig. 3d). Both purity and completeness were much lower for smaller and larger bins. CONCOCT assigned the most bp (Fig. 3e), though into fewer bins. Low purity and fewer bins indicate “underbinning”, i.e. multiple genomes being placed together in one bin. The other extreme, “overbinning”, occurs when genomes are split across multiple bins, resulting in low completeness. After DAS Tool, MaxBin predictions had the highest ARI, followed by MetaBAT. DAS Tool substantially improved bin purity and ARI relative to the individual methods, at the cost of completeness and assigning less than two methods. MaxBin and DAS Tool recovered the most high-quality genomes, defined as genomes with more than 50% completeness and less than 10% contamination (Table 5). The total number of predicted bins per method was 867 (MaxBin), 592 (MetaBAT), 344 (CONCOCT), and 577 (DAS Tool).

**Table 5:** Number of high-quality genomes and corresponding percentages recovered from the gold standard assembly of the CAMI II mouse gut data set. The best performing individual method and best performer overall are indicated in bold.

| Genome binner | % contamination | Predicted bins<br>% completeness |  |  |
| --- | --- | --- | --- | --- |
|  |  | >50% | >70% | >90% |
| Gold standard |  | 791 (100%) | 791 (100%) | 791 (100%) |
| MaxBin 2.2.7 | < 10% | **439 (55%)** | **419 (53%)** | **342 (43%)** |
|  | < 5% | **401 (51%)** | **386 (49%)** | **319 (40%)** |
| MetaBAT 2.12.1 | < 10% | 353 (45%) | 318 (40%) | 240 (30%) |
|  | < 5% | 339 (43%) | 309 (39%) | 236 (30%) |
| CONCOCT 1.0.0 | < 10% | 95 (12%) | 95 (12%) | 84 (11%) |
|  | < 5% | 88 (11%) | 88 (11%) | 79 (10%) |
| DAS Tool 1.1.2<br>(ensemble method) | < 10% | **460 (58%)** | **449 (57%)** | **354 (45%)** |
|  | < 5% | **422 (53%)** | **416 (53%)** | **334 (45%)** |

We compared the bin quality metrics to those returned by the commonly used CheckM software version 1.1.2, which assesses bin quality based on the presence of lineage specific marker genes^47^ (Fig. 3f, Supplementary information). Results are largely consistent. CheckM overestimated purity by 4% (MetaBAT and DAS Tool) to 21% (MaxBin) and completeness by 2% (MetaBAT and CONCOCT) to 7% (MaxBin) (Fig. 3f, Supplementary Tables 6 and 7). Due to CheckM’s known bias of overestimating completeness and underestimating contamination^47^, we also computed the averages of only those bins with more than 90% completeness and less than 10% contamination according to AMBER’s assessment. In this case, CheckM’s purity overestimates dropped to only up to 3% for all methods except CONCOCT, for which it increased to 29%. On the other hand, completeness was underestimated for most methods, by 9% (CONCOCT) to 17% (MaxBin).

### Taxonomic binning

A taxon bin is a set of sequences, either contigs or reads, with the same taxonomic label. Taxonomic binning can be evaluated as a multi-class classification problem at individual taxonomic ranks, where one of many possible taxon labels from a reference taxonomy is assigned to every metagenomic sequence. The quality of a taxon binning is assessed by comparing predicted and ground truth taxon bins with each other.

We predicted taxon bins from the cross-sample gold standard assembly with DIAMOND 0.9.24^48^, Kraken 2.0.8 beta^49^, PhyloPythiaS+ 1.4^50^, CAT 4.6^51^, and MEGAN 6.15.2^52^. All results and commands used are available on Zenodo (Supplementary Table 8). Runtimes and memory usage are given in Supplementary Table 9. The release date of the NCBI taxonomy used by each method is indicated on Zenodo and can vary slightly, depending on the reference database of the method. Method performances were assessed with AMBER 2.0.1, for all major taxonomic ranks (Figs. 4 and 5), using the NCBI taxonomy database from 2018/02/26. This reference taxonomy is provided with the mouse gut data set of the CAMI II challenge (Table 2). To run AMBER, type:

**Fig. 4:**
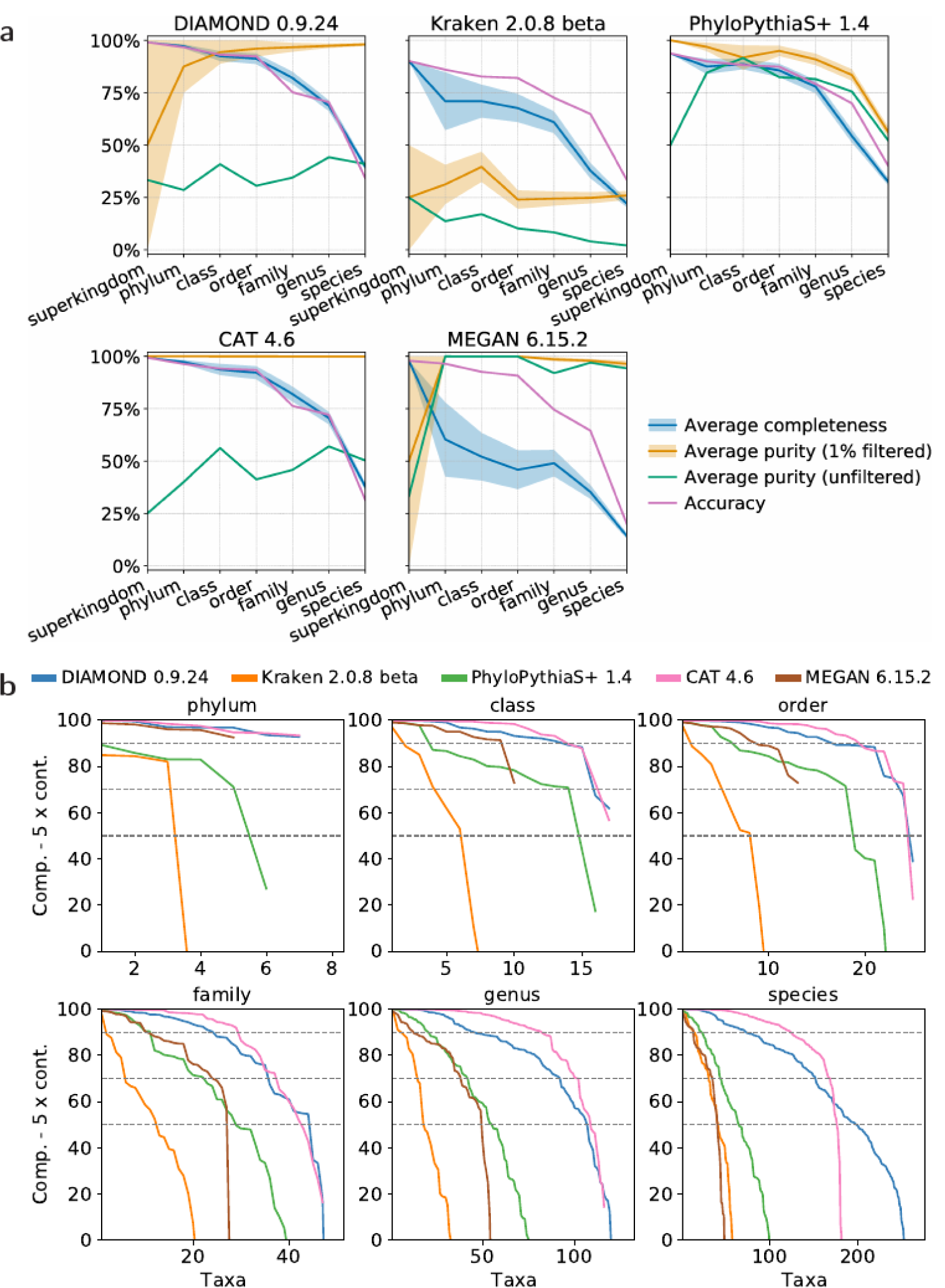
Assessing taxonomic binning results on the CAMI II mouse gut data set. **a** Average completeness and purity (1% filtered and unfiltered, see main text) and accuracy per taxonomic rank for each binner. The shaded bands show the standard deviation of a metric. **b** Score (i.e. completeness - 5 X contamination, y axis) and number of predicted taxon bins (x axis) for the phylum to species ranks. The higher the number of high-scoring bins, the better is the binning performance. Only positive scores are shown. The dotted lines indicate the 90, 70, and 50 score thresholds.

**Fig. 5:**
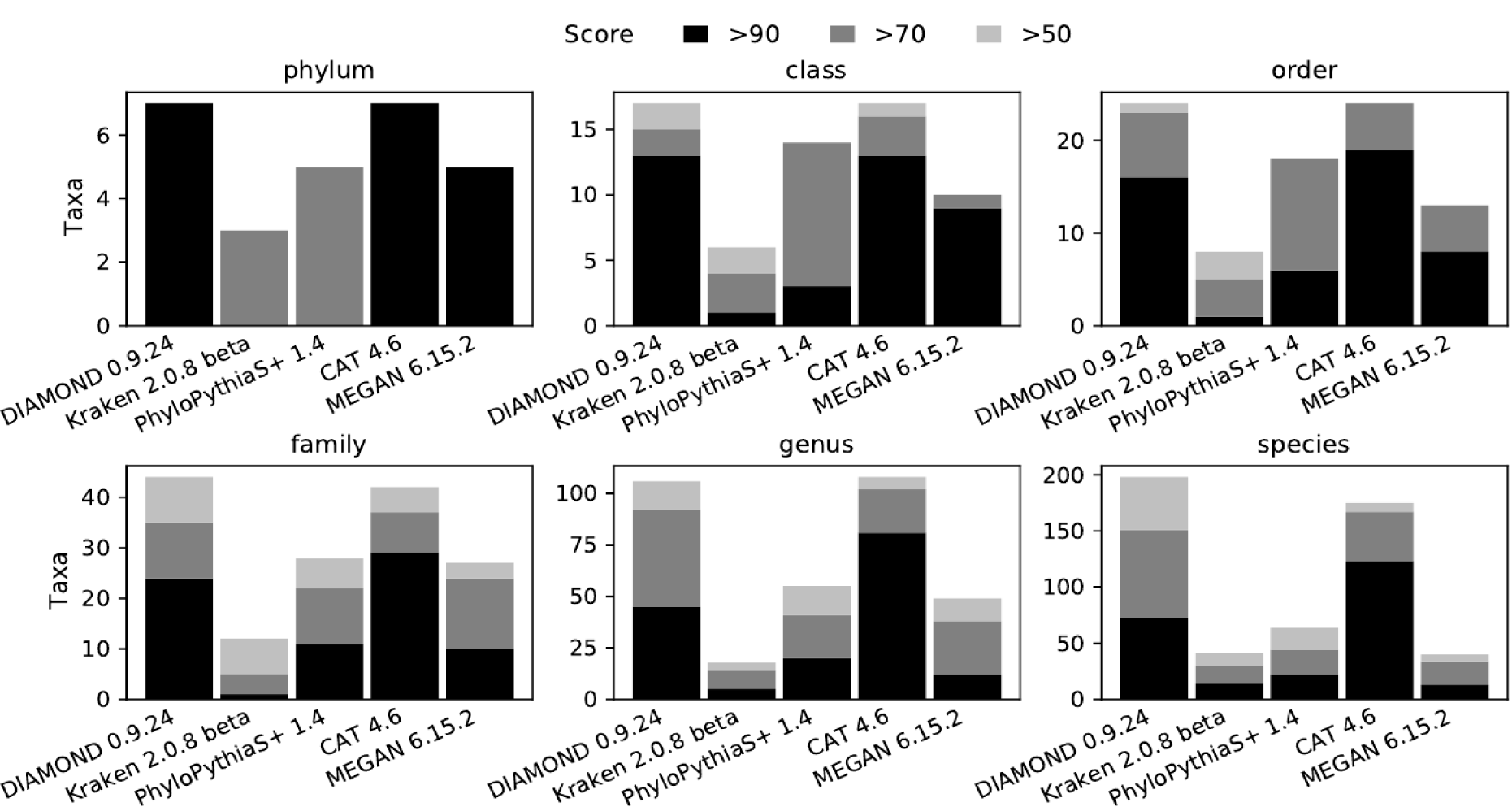
Number of high-quality taxon bins predicted from the CAMI II mouse gut data set for the phylum to species ranks. Counted are the bins with score (i.e. completeness - 5 X contamination) higher than 90, 70, and 50. A number of bins closer to the number of taxa per rank in the gold standard (i.e. 8 phyla, 18 classes, 26 orders, 50 families, 157 genera, and 549 species) is better.

~~~
amber.py --gold_standard_file /path/to/cami2_mouse_gut_gsa_pooled.binning \
--desc “CAMI 2 toy mouse gut data set” \
/path/to/cami2_mouse_gut_diamond0.9.24.binning \
/path/to/cami2_mouse_gut_kraken2.0.8beta.binning \
/path/to/cami2_mouse_gut_ppsp1.4.binning \
/path/to/cami2_mouse_gut_cat4.6.binning \
/path/to/cami2_mouse_gut_megan6.15.2.binning \
--labels “DIAMOND 0.9.24, Kraken 2.0.8 beta, PhyloPythiaS+ 1.4, CAT 4.6, MEGAN 6.15.2”
\
--ncbi_nodes_file /path/to/nodes.dmp \
--ncbi_names_file /path/to/names.dmp \
--ncbi_merged_file /path/to/merged.dmp \
--filter 1 \
--output_dir /path/to/output_dir
~~~

For comparing predicted taxon bins to the ground truth, **completeness** and **purity** can be calculated. The completeness, or recall for a taxon bin found in the ground truth is the fraction of ground truth contigs, or bp, that have been assigned to that taxon by a method. Completeness is averaged over all ground truth taxon bins at a particular rank and undefined for predicted taxon bins not present in the ground truth. The purity of a predicted taxon bin is the fraction of contigs, or bp, belonging to that taxon in the ground truth. Taxon bins without any correctly assigned sequences accordingly have a purity of zero. Purity is averaged over all predicted taxon bins at a particular rank. **Contamination** is defined as 100% minus purity. Finally, the **accuracy** is the fraction of contigs, or bp, that have been assigned by a method to the correct taxa for a rank. Accuracy is a sample-specific metric to which larger taxon bins contribute more strongly than small ones, different from average completeness and purity.

DIAMOND and CAT, which relies on DIAMOND’s output, obtained the highest average completeness for all ranks. This was above 90% from superkingdom to order and continuously dropped at lower ranks (Fig. 4a). MEGAN, which also uses DIAMOND, achieved lower completeness for phylum level and below, but the highest average purity at all ranks, except for superkingdom, at which PhyloPythiaS+ performed best. As purity can be reduced for small bins, we filtered out the smallest predicted bins per method and rank, removing overall 1% of the binned data in bp. This can be done with AMBER (option --filter 1) on the predicted bins, requiring no knowledge of the underlying gold standard. Across all ranks, the average size of the removed taxon bins was 0.35 Mb, whereas the average size of all bins was 235.79 Mb (Supplementary Table 10), with larger bins accumulating at higher ranks. DIAMOND and CAT profited most from this, with CAT reaching almost 100% filtered purity at all ranks. Researchers interested in taxa with small genomes, such as viruses, should keep in mind that filtering could remove these along with false positive bins. Purity and completeness were also influenced by contig length and overall higher for longer contigs (Supplementary Fig. 1). In terms of accuracy, all methods performed similarly well, with PhyloPythiaS+ being the most accurate at the species level.

Based on a quality score defined as completeness - 5 X contamination, as in ^7,53^, we determined the number of high-quality bins found by each method with a score of more than 90, 70, and 50 at different taxonomic ranks (Fig. 5). DIAMOND, CAT, and PhyloPythiaS+, in this order, identified the most high-quality bins (>50) at all taxonomic ranks. CAT, followed by DIAMOND, found the most bins with a score higher than 90.

### Taxonomic profiling

Taxonomic profiling can be considered a multi-label problem at a given rank, where multiple taxon labels are assigned to a single sample and the relative taxon abundances are estimated. Profiling differs from binning in that individual reads are not necessarily assigned taxon labels. We predicted taxonomic identities and relative abundances of microbial community members for the 64 short read samples of the mouse gut data set with MetaPhlAn 2.9.21^54^, mOTUs 2.5.1^55^, and Bracken 2.5^56^. We assessed these together with results for MetaPhlAn 2.2.0, mOTUs 1.1, MetaPalette 1.0.0, MetaPhyler 1.25, FOCUS 0.31, TIPP 2.0.0, and CAMIARKQuikr 1.0.0 from ^28^. The profiling results and commands used can be obtained from Zenodo (Supplementary Table 11). Runtimes and memory usage are given in Supplementary Table 12. Performance metrics and result visualizations were calculated with OPAL^28^

1.0.8 (Table 3), which can be installed with the following command if Bioconda is configured:

~~~
conda create --name opal cami-opal
~~~

Other installation methods are described in the OPAL GitHub repository at https://github.com/CAMI-challenge/OPAL/. We then ran OPAL as:

~~~
conda activate opal
opal.py --gold_standard_file /path/to/cami2_mouse_gut_gs.profile \
/path/to/cami2_mouse_gut_metaphlan2.2.0.profile \
/path/to/cami2_mouse_gut_metaphlan2.9.21.profile \
/path/to/cami2_mouse_gut_motus1.1.profile \
/path/to/cami2_mouse_gut_motus2.5.1.profile \
/path/to/cami2_mouse_gut_bracken2.5.profile \
/path/to/cami2_mouse_gut_metapalette1.0.0.profile \
/path/to/cami2_mouse_gut_metaphyler1.25.profile \
/path/to/cami2_mouse_gut_focus0.31.profile \
/path/to/cami2_mouse_gut_tipp2.0.0.profile \
/path/to/cami2_mouse_gut_camiarkquikr1.0.0.profile \
--labels “MetaPhlAn 2.2.0, MetaPhlAn 2.9.21, mOTUs 1.1, mOTUs 2.5.1, Bracken 2.5,
MetaPalette 1.0.0, MetaPhyler 1.25, FOCUS 0.31, TIPP 2.0.0, CAMIARKQuikr 1.0.0” \
-d “2nd CAMI Challenge Mouse Gut Toy Dataset” \
--metrics_plot c,p,l,w \
--filter 1 \
--output_dir /path/to/output_dir
~~~

OPAL computes performance metrics and creates visualizations for profiling results on a benchmark data set. It also generates weighted summary scores for ranking methods based on these metrics (see^28^ for a complete overview and formal definitions). For a taxonomic rank, the **purity** and **completeness** assess how well a profiler identified the presence and absence of taxa, without considering relative abundances. Purity, or precision, denotes the ratio of correctly predicted taxa to all predicted taxa predicted at a taxonomic rank, whereas completeness, or recall, is the ratio of correctly identified taxa to all ground truth taxa at a taxonomic rank. To explore the effect of heuristic post-processing of predictions on purity, we filtered low abundance taxon predictions as we did for taxonomic binners^8^: by removing predictions with the lowest relative abundances, summing up to one percent of the total predicted organismal abundances per taxonomic rank.

For quantifying relative abundance estimates, the **L1 norm** and **weighted UniFrac** error are determined. The L1 norm assesses relative abundance estimates of taxa at a taxonomic rank, based on the sum of the absolute differences between the true and predicted abundances across all taxa. The weighted UniFrac error computed by OPAL uses a taxonomic tree storing the predicted abundances at the appropriate nodes for eight major taxonomic ranks. The UniFrac error is the total amount of predicted abundances that must be moved along the edges of the tree to cause them to overlap with the true relative abundances. Branch lengths in the taxonomic tree can be set to 1 or any function of the depth of the edge in the taxonomic tree. This choice is motivated by the fact that harmonizing phylogenetic trees (which express evolutionary distance with branch lengths) and taxonomic trees (which do not inherently have branch length information) remains an open problem under active investigation^57–60^. A low UniFrac error indicates good accuracy of abundance estimates. Prior to computing the L1 norm and weighted UniFrac error, OPAL, per default (as used here), normalizes all relative abundance estimates, which may be less than one if some data remains taxonomically unassigned, such that their sum equals 1 at each rank. Normalization can simplify the comparison of the L1 norm between methods (https://github.com/CAMI-challenge/firstchallenge_evaluation/tree/master/profiling), however, may skew results for profilers with low recall that left many taxa unassigned. Assessment results with unnormalized relative abundance estimates are available in the OPAL GitHub repository.

Using all these metrics, OPAL ranks the assessed profilers by their relative performance. For each metric, sample, and major taxonomic rank (from superkingdom to species), the best performing profiler is assigned score 0, the second best, 1, and so on. These scores are then added over the taxonomic ranks and samples to produce a single score per metric for each profiler. OPAL can also assign different weights to the metrics, such that the importance of a metric, defined by the user, is reflected in the overall score and rank of a profiler. In our assessment, all metrics were weighted equally.

mOTUs 2.5.1, Bracken 2.5, MetaPhyler 1.25, and TIPP 2.0.0, in this order, achieved the overall highest completeness (Fig. 6). mOTUs 2.5.1 achieved high completeness up to genus level, whereas the other profilers performed well with this metric up to family level. Along with completeness, purity also drops for lower taxonomic ranks. Filtering low abundant taxon predictions greatly improved purity, most strongly for MetaPhyler and Bracken 2.5, which was ranked 7th instead of last with this metric. MetaPhlAn 2.2.0 and mOTUs 1.1 had the highest filtered purity across ranks, followed by mOTUs 2.5.1 and MetaPhlAn 2.9.21. mOTUs 2.5.1 showed both high (filtered and unfiltered) purity and completeness and improved considerably in terms of completeness compared to its previous version. mOTUs 2.5.1, MetaPhlAn 2.9.21, MetaPhlAn 2.2.0, and MetaPhyler 1.25, in this order, best estimated the relative abundances measured with the L1 norm, with MetaPhlAn 2.9.21 outperforming all methods at the species level. mOTUs 2.5.1 also obtained a low UniFrac error, followed by MetaPhlAn 2.9.21 and MetaPhlAn 2.2.0. Considering all metrics, mOTUs 2.5.1 ranked first, followed by MetaPhlAn 2.2.0 and 2.9.21. Notably, normalization of abundance estimates had almost no effect on the L1 norm error of the methods (Supplementary Fig. 2), as the estimates covered almost 100% of the data (Supplementary Table 13). We note that performance estimates may differ strongly depending on metric definitions. For instance, contrary to the findings reported here, mOTUs and MetaPhlAN were reported to perform poorly in terms of the fraction of sample reads that they classified^21^, which is a task that they were not designed for.

**Fig. 6:**
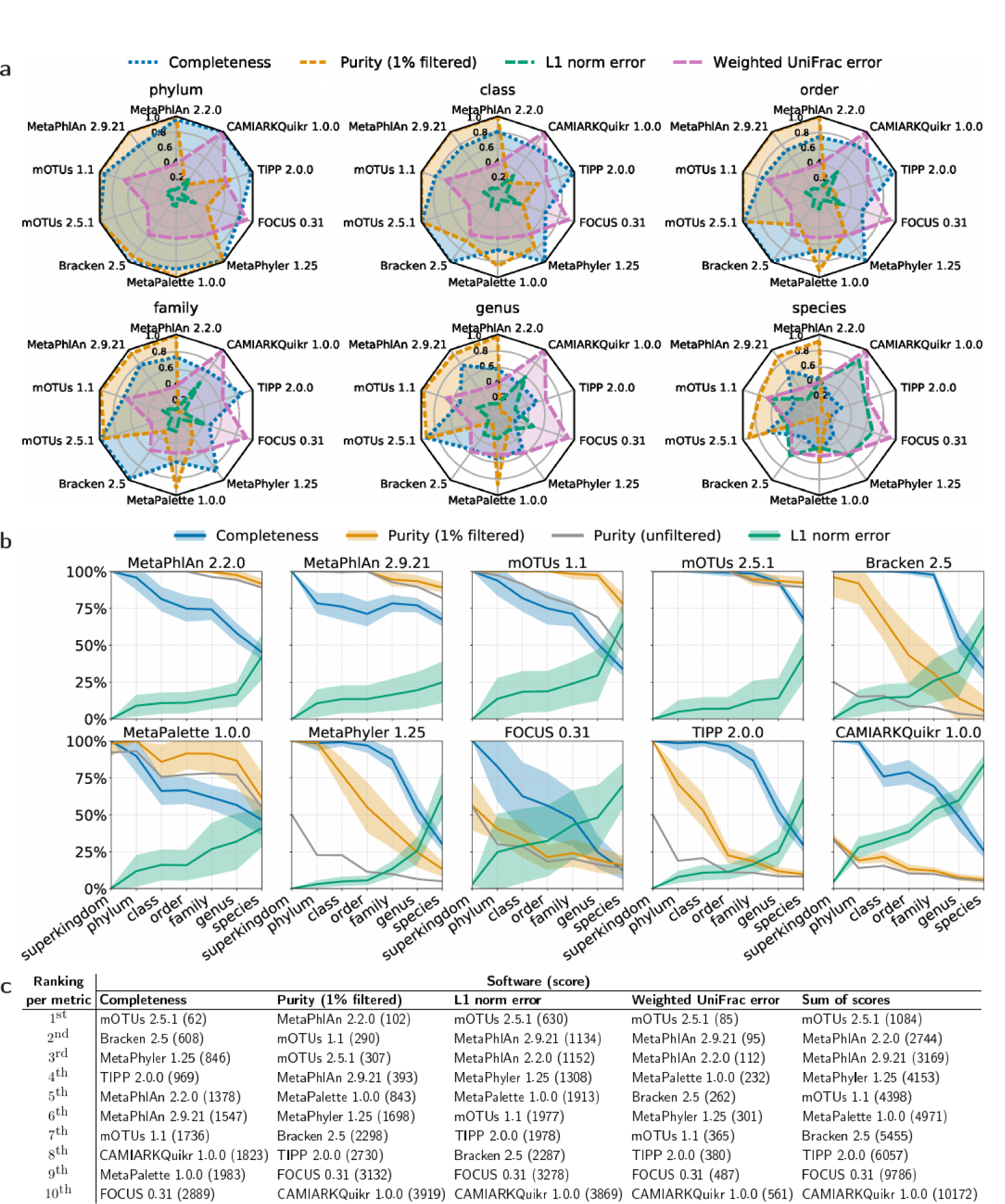
Assessing profiling results on the CAMI II mouse gut data set. **a** Comparison per taxonomic rank of methods in terms of completeness, purity (1% filtered, see main text), L1 norm, and weighted UniFrac error. **b** Performance per method at all major taxonomic ranks, with the shaded bands showing the standard deviation of a metric. In **a** and **b**, completeness, purity, and L1 norm error range between 0 and 1. The L1 norm error is normalized to this range and is also known as Bray-Curtis distance. The weighted UniFrac error is rank-independent and normalized by the maximum value obtained by the profilers. The higher the completeness and purity, and the lower the L1 norm and weighted UniFrac error, the better the profiling performance. **c** Methods rankings and scores obtained for the different metrics over all samples and taxonomic ranks. For score calculation, all metrics were weighted equally.

## Summary and conclusions

Microbiome research using metaomics technologies is a rapidly progressing field producing highly complex and heterogeneous data. For developing and assessing data processing techniques, adoption of benchmarking standards in the field is essential. We here outlined key elements of benchmarking and best practices developed by a larger group of scientists within CAMI for common computational analyses in metagenomics. Community-driven benchmarking challenges are a key component of unbiased performance evaluations, in addition to the assessments by individual developers that are commonly done. To facilitate the latter, we describe a benchmarking tool resource and the mechanisms to use and add to this resource, as indicated in ^8^, in a flexible way. We show how to apply the CAMI standards and data for performance assessment using a benchmarking toolkit developed in large part within CAMI. For profiling methods, we demonstrated the value of incremental benchmarking by reusing and combining tool results from different studies and saving these in the CAMI tool result repositories on Zenodo (https://zenodo.org/communities/cami). Curated metadata and instructions on how to contribute reproducible results are provided at https://github.com/CAMI-challenge/data. As these new resources grow, individual benchmarks of metaomics software will become increasingly more efficient, informative and reproducible.

Using the 64 sample simulated metagenome data set from mouse guts as an example, we performed a comparative evaluation of metagenome assembly (for the first 10 samples), genome binning, taxonomic binning and profiling on these data. Overall, the evaluation included 25 results for 19 computational methods: 2 assemblers, with 6 different settings and versions evaluated, 4 genome and 5 taxon binners, as well as 8 profilers, including 2 different versions. Seven of the profiling results originate from a previous evaluation study on the data, demonstrating the value of incremental data analysis. Notably, as the data set was generated from genomes included in public databases, the results for reference-based methods, such as taxonomic binning and profiling techniques, are to be taken as representative only for microbial community members represented by close relatives in public database content. This is only true for a fraction of most microbial communities, if not considering computationally reconstructed MAGs as a reference. Accordingly, for reference-based techniques, i.e. taxonomic binners and profilers, results were consistent with prior studies on data generated from publicly available genomes^28^, and less congruent with performances on benchmark data including genomes more distantly related to public database content^8^. Performance on species that are distantly to those with genomes in public databases continues to be an important point to keep in mind when selecting the most suitable method for analysis.

With the CAMI benchmarking resources in place, we invite researchers to make full use of these for tackling the big challenges in the field^61^. These include developing strain-resolved assembly, binning and profiling techniques for strain-specific genome reconstructions^62,63^, making use of long-read metagenomic sequencing data^64^, evaluating methods for other metaomics, e.g. metatranscriptomics, metaproteomics^65^, and metametabolomics. The applications of metagenomics are diverse and growing, and the best way to tackle this is via a large collaborative framework supported by good collaborative infrastructure, which CAMI aims to provide.

## Supporting information

Supplementary results: MetaQUAST metrics

## Data availability

The results of all benchmarked methods and gold standards are available at https://zenodo.org/communities/cami. Links to individual results and DOIs are available in Supplementary Tables 1, 4, 8, and 11. The gold standard assembly is provided with the CAMI II mouse gut data set (Table 2). Assembly results and code used to generate Fig. 2 are available at https://github.com/CAMI-challenge/BenchmarkingToolkitTutorial. Genome and taxonomic binning, and taxonomic profiling results used in Figs. 3-6 are available, respectively, in the AMBER and OPAL GitHub repositories at https://github.com/CAMI-challenge/AMBER and https://github.com/CAMI-challenge/OPAL.

## Acknowledgements

The authors thank P.B. Pope for helpful comments. A.E.D.’s contribution was facilitated in part by the Australian Research Council’s Discovery Projects funding scheme (project DP180101506). A.G.’s contribution was facilitated by St. Petersburg State University, Russia (grant ID PURE 51555639).

## Author contributions

F.M. and T.R.L. performed the experiments; F.M., A.F., T.R.L., and A.S. prepared the data; A.C.M., A.B., and A.S. conceived the experiments; A.C.M., F.M., and A.B. wrote the manuscript with comments by others; all authors interpreted the results, and read and approved the final manuscript.

## Supplementary information

Supplementary information: Supplementary Tables 1-13, Supplementary Figs. 1 and 2, Bin quality metrics for CheckM

Supplementary results: MetaQUAST metrics (report.html)

