## Supplementary tables and figures for "Tutorial: Assessing metagenomics software with the CAMI benchmarking toolkit"

**Supplementary Table 1: Digital Object Identifiers (DOIs) for assembly results of the first 10 of 64 short read samples of the CAMI II mouse gut data set and associated links.**

| Assembler | Results DOI | Links for commands used and download |
|---|---|---|
| MEGAHIT 1.0.3 df | 10.5281/zenodo.3663885 | https://doi.org/10.5281/zenodo.3663885 https://zenodo.org/record/3663885/files/megahit103-Sample0-9-df-final.contigs.fa.gz?download=1 |
| MEGAHIT 1.1.3 df | 10.5281/zenodo.3663885 | https://doi.org/10.5281/zenodo.3663885 https://zenodo.org/record/3663885/files/megahit113-Sample0-9-df-final.contigs.fa.gz?download=1 |
| MEGAHIT 1.1.3 ml | 10.5281/zenodo.3663885 | https://doi.org/10.5281/zenodo.3663885 https://zenodo.org/record/3663885/files/megahit113-Sample0-9-ml-final.contigs.fa.gz?download=1 |
| MEGAHIT 1.1.3 ms | 10.5281/zenodo.3663885 | https://doi.org/10.5281/zenodo.3663885 https://zenodo.org/record/3663885/files/megahit113-Sample0-9-ms-final.contigs.fa.gz?download=1 |
| MEGAHIT 1.2.9 df | 10.5281/zenodo.3663885 | https://doi.org/10.5281/zenodo.3663885 https://zenodo.org/record/3663885/files/megahit129-Sample0-9-df-final.contigs.fa.gz?download=1 |
| metaSPAdes 3.13.0 | 10.5281/zenodo.3664090 | https://doi.org/10.5281/zenodo.3664090 https://zenodo.org/record/3664090/files/metaSPAdes3130-Sample0-9-contigs.fasta.gz?download=1 |

**Supplementary Table 2: Elapsed (wall clock) time (h:mm) of assembly methods on the first 10 of 64 short read samples of the CAMI II mouse gut data set.** The best result is shown in bold. The assemblers were run on a computer with several Intel Xeon Gold 6142 CPUs, virtualized to 58 logical cores, and 1.4 TB of main memory.

| Assembler | default (df) | meta-sensitive (ms) | meta-large (ml) |
|---|---:|---:|---:|
| MEGAHIT 1.0.3 | 7:14 | – | – |
| MEGAHIT 1.1.3 | 6:24 | 14:00 | 11:56 |
| MEGAHIT 1.2.9 | **4:11** | – | – |
| metaSPAdes 3.13.0 | 41:06 | – | – |

**Supplementary Table 3: Maximum resident set size (kbytes) of assembly methods on the first 10 of 64 short read samples of the CAMI II mouse gut data set.** The best results are shown in bold.

| Assembler | default (df) | meta-sensitive (ms) | meta-large (ml) |
|---|---:|---:|---:|
| MEGAHIT 1.0.3 | 127,403,256 | – | – |
| MEGAHIT 1.1.3 | **42,278,824** | 186,245,380 | 179,530,964 |
| MEGAHIT 1.2.9 | **42,919,140** | – | – |
| metaSPAdes 3.13.0 | 189,646,496 | – | – |

**Supplementary Table 4: Digital Object Identifiers (DOIs) and associated links of genome binning results and ground truth of the gold standard cross-sample assembly of the CAMI II mouse gut data set.** Also given is the average coverage of the underlying genomes.

| Genome binner | Results DOI | Links for commands used and download |
|---|---|---|
| MaxBin 2.2.7 | 10.5281/zenodo.3629588 | https://doi.org/10.5281/zenodo.3629588 https://zenodo.org/record/3629588/files/cami2_mouse_gut_maxbin2.2.7.binning?download=1 |
| MetaBAT 2.12.1 | 10.5281/zenodo.3629590 | https://doi.org/10.5281/zenodo.3629590 https://zenodo.org/record/3629590/files/cami2_mouse_gut_metabat2.12.1.binning?download=1 |
| CONCOCT 1.0.0 | 10.5281/zenodo.3629592 | https://doi.org/10.5281/zenodo.3629592 https://zenodo.org/record/3629592/files/cami2_mouse_gut_concoct1.0.0.binning?download=1 |
| DAS Tool 1.1.2 | 10.5281/zenodo.3629594 | https://doi.org/10.5281/zenodo.3629594 https://zenodo.org/record/3629594/files/cami2_mouse_gut_dastool1.1.2.binning?download=1 |
| Binning ground truth | 10.5281/zenodo.3632511 | https://doi.org/10.5281/zenodo.3632511 https://zenodo.org/record/3632511/files/cami2_mouse_gut_gsa_pooled.binning?download=1 |
| Average genome coverage | 10.5281/zenodo.3667475 | https://doi.org/10.5281/zenodo.3667475 https://zenodo.org/record/3667475/files/cami2_mouse_gut_average_genome_coverage.tsv?download=1 |

**Supplementary Table 5: Elapsed (wall clock) time (h:mm) and maximum resident set size (kbytes) of genome binning methods on the cross-sample gold standard assembly of the CAMI II mouse gut data set.** The best result is shown in bold. DAS Tool 1.1.2 (refinement only) is the time required to run only DAS Tool 1.1.2 using the output of MaxBin 2.2.7, MetaBAT 2.12.1, and CONCOCT 1.0.0. DAS Tool 1.1.2 (total) is the time required to run all these binners, including DAS Tool 1.1.2. The binners were run on a computer with an Intel Xeon E5-4650 v4 CPU (virtualized to 16 CPU cores, 1 thread per core) and 512 GB (536.870.912 kbytes) of main memory.

| Genome binner | Time (hh:mm) | Memory (kbytes) |
|---|---:|---:|
| MaxBin 2.2.7 | 329:46 | 13,789,512 |
| MetaBAT 2.12.1 | **33:29** | **12,985,728** |
| CONCOCT 1.0.0 | 41:06 | **12,985,728** |
| DAS Tool 1.1.2 (refinement only) | **00:37** | 3,755,972 |
| DAS Tool 1.1.2 (total) | 404:58 | 13,789,512 |

**Supplementary Table 6: CheckM and AMBER average purity assessment of genome binning results and ground truth of the gold standard cross-sample assembly of the CAMI II mouse gut data set**. In parentheses is the average purity of the predicted bins with completeness > 70% and contamination < 10% according to AMBER's assessment. Also shown is the absolute difference between CheckM's and AMBER's assessments.

**Average purity (completeness > 70% and contamination < 10%)**

| Genome binner | CheckM | AMBER | Difference (%) |
|---|---|---|---|
| Binning ground truth | 0.984 (0.984) | 1.000 (1.000) | 1.564 (1.564) |
| MaxBin 2.2.7 | 0.939 (0.957) | 0.774 (0.988) | 21.346 (3.092) |
| MetaBAT 2.12.1 | 0.949 (0.961) | 0.909 (0.994) | 4.400 (3.324) |
| CONCOCT 1.0.0 | 0.659 (0.694) | 0.594 (0.989) | 10.909 (29.829) |
| DAS Tool 1.1.2 | 0.968 (0.988) | 0.929 (0.989) | 4.161 (0.122) |

**Supplementary Table 7: CheckM and AMBER average completeness assessment of genome binning results and ground truth of the gold standard cross-sample assembly of the CAMI II mouse gut data set**. In parentheses is the average completeness of the predicted bins with completeness > 70% and contamination < 10% according to AMBER's assessment. Also shown is the absolute difference between CheckM's and AMBER's assessments.

**Average completeness (completeness > 70% and contamination < 10%)**

| Genome binner | CheckM | AMBER | Difference (%) |
|---|---|---|---|
| Binning ground truth | 0.927 (0.927) | 1.000 (1.000) | 7.270 (7.270) |
| MaxBin 2.2.7 | 0.692 (0.785) | 0.641 (0.954) | 7.949 (17.674) |
| MetaBAT 2.12.1 | 0.698 (0.788) | 0.683 (0.939) | 2.208 (16.105) |
| CONCOCT 1.0.0 | 0.868 (0.874) | 0.848 (0.964) | 2.291 (9.333) |
| DAS Tool 1.1.2 | 0.910 (0.967) | 0.877 (0.949) | 3.770 (1.964) |

### Bin quality metrics for CheckM

The purity for CheckM was calculated as the number of marker genes inferred for the bin lineage divided by the number of markers identified in the bin. An example report from CheckM's output file `bin_stats_ext.tsv` showing the assessments for three bins predicted with CONCOCT (bin IDs 111, 105, and 1), with the respective purity computation shown in bold, is as follows:

111 {'marker lineage': 'root', '# genomes': 5656, '# markers': 56, '# marker sets': 24, '0': 56, '1': 0, '2': 0, '3': 0, '4': 0, '5+': 0, 'Completeness': 0.0, 'Contamination': 0.0, ...
**Purity = 56 / (56 + 0 + 0 * 2 + 0 * 3 + 0 * 4 + 0 * 5) = 1.0**

105 {'marker lineage': 'root', '# genomes': 5656, '# markers': 56, '# marker sets': 24, '0': 0, '1': 0, '2': 7, '3': 46, '4': 3, '5+': 0, 'Completeness': 100.0, 'Contamination': 194.69696969696972, ...
**Purity = 56 / (0 + 0 + 7 * 2 + 46 * 3 + 3 * 4 + 0 + 0 * 5) = 0.341**

1 {'marker lineage': 'root', '# genomes': 5656, '# markers': 56, '# marker sets': 24, '0': 0, '1': 0, '2': 1, '3': 52, '4': 3, '5+': 0, 'Completeness': 100.0, 'Contamination': 206.25, ...
**Purity = 56 / (0 + 0 + 1 * 2 + 52 * 3 + 3 * 4 + 0 + 0 * 5) = 0.329**

As CheckM reports metrics per bin, not per genome, we calculated the average purity and completeness for both CheckM and AMBER as a simple average of these metrics over all bins. For AMBER, the completeness for a bin was then determined as the fraction of bp from the gold standard genome most abundant in a bin.

**Supplementary Table 8: Digital Object Identifiers (DOIs) and associated links of taxonomic binning results and binning ground truth of the gold standard cross-sample assembly of the CAMI II mouse gut data set.**

| Taxonomic binner | Results DOI | Links for commands used and download |
|---|---|---|
| DIAMOND 0.9.24 | 10.5281/zenodo.3629598 | http://doi.org/10.5281/zenodo.3629598 https://zenodo.org/record/3629598/files/cami2_mouse_gut_diamond0.9.24.binning?download=1 |
| Kraken 2.0.8 beta | 10.5281/zenodo.3629600 | http://doi.org/10.5281/zenodo.3629600 https://zenodo.org/record/3629600/files/cami2_mouse_gut_kraken2.0.8beta.binning?download=1 |
| PhyloPythiaS+ 1.4 | 10.5281/zenodo.3629602 | http://doi.org/10.5281/zenodo.3629602 https://zenodo.org/record/3629602/files/cami2_mouse_gut_ppsp1.4.binning?download=1 |
| CAT 4.6 | 10.5281/zenodo.3629604 | http://doi.org/10.5281/zenodo.3629604 https://zenodo.org/record/3629604/files/cami2_mouse_gut_cat4.6.binning?download=1 |
| MEGAN 6.15.2 | 10.5281/zenodo.3629606 | http://doi.org/10.5281/zenodo.3629606 https://zenodo.org/record/3629606/files/cami2_mouse_gut_megan6.15.2.binning?download=1 |
| Binning ground truth | 10.5281/zenodo.3632511 | http://doi.org/10.5281/zenodo.3632511 https://zenodo.org/record/3632511/files/cami2_mouse_gut_gsa_pooled.binning?download=1 |

**Supplementary Table 9: Elapsed (wall clock) time (h:mm) and maximum resident set size (kbytes) of taxonomic binning methods on the cross-sample gold standard assembly of the CAMI II mouse gut data set.** The best result is shown in bold. The time for MEGAN 6.15.2 is the sum of the time to run DIAMOND 0.9.24 and MEGAN's tool daa2rma, which uses DIAMOND's output. The binners were run on a computer with an Intel Xeon E5-4650 v4 CPU (virtualized to 16 CPU cores, 1 thread per core) and 512 GB (536.870.912 kbytes) of main memory.

| Taxonomic binner | Time (hh:mm) | Memory (kbytes) |
|---|---:|---:|
| DIAMOND 0.9.24 | 21:58 | 43,350,528 |
| Kraken 2.0.8 beta | **0:22** | 39,439,795 |
| PhyloPythiaS+ 1.4 | 206:38 | 285,949,912 |
| CAT 4.6 | 49:17 | **19,039,232** |
| MEGAN 6.15.2 | 23:14 | 97,196,924 |

**Supplementary Table 10: Average size in bp of taxonomic bins predicted from the CAMI II mouse gut data set.** For the assessments (see main text), the smallest bins per method and rank were filtered out (overall 1% of the binned data in bp). The average size of those bins is shown in the right-most column.

| Taxonomic binner | Taxonomic rank | Average bin size (#bp) | Average size of removed bins (#bp) |
|---|---|---|---|
| Gold standard | superkingdom | 2,710,998,838.0 | - |
| | phylum | 338,874,854.8 | - |
| | class | 150,611,046.6 | - |
| | order | 104,269,186.1 | - |
| | family | 53,579,524.6 | - |
| | genus | 15,618,714.7 | - |
| | species | 4,938,067.1 | - |
| DIAMOND 0.9.24 | superkingdom | 896,268,023.7 | 597,598.0 |
| | phylum | 93,734,467.0 | 138,154.6 |
| | class | 57,448,709.1 | 101,632.2 |
| | order | 29,488,842.5 | 48,169.9 |
| | family | 14,177,056.6 | 46,205.3 |
| | genus | 5,079,833.3 | 80,259.2 |
| | species | 791,044.8 | 116,384.3 |
| Kraken 2.0.8 beta | superkingdom | 614,745,492.8 | - |
| | phylum | 59,265,069.0 | 60,345.8 |
| | class | 31,313,814.2 | 61,127.0 |
| | order | 13,829,875.4 | 51,040.3 |
| | family | 6,077,181.1 | 49,811.2 |
| | genus | 1,747,856.2 | 35,301.7 |
| | species | 574,999.9 | 24,972.4 |
| PhyloPythiaS+ 1.4 | superkingdom | 1,270,708,176.5 | 218,593.0 |
| | phylum | 272,397,046.0 | 912,405.3 |
| | class | 127,406,374.4 | 1,278,648.5 |
| | order | 79,999,745.2 | 1,064,217.5 |
| | family | 40,439,925.3 | 569,077.8 |
| | genus | 12,216,123.3 | 627,053.6 |
| | species | 4,322,775.1 | 533,396.5 |
| CAT 4.6 | superkingdom | 673,666,601.5 | 218,182.3 |
| | phylum | 130,569,785.9 | 60,631.1 |
| | class | 79,767,843.8 | 54,047.8 |
| | order | 40,230,339.4 | 22,085.7 |
| | family | 18,941,940.6 | 58,487.0 |
| | genus | 6,623,232.4 | 102,320.1 |
| | species | 894,018.2 | 161,734.5 |
| MEGAN 6.15.2 | superkingdom | 885,889,337.7 | 645,792.0 |
| | phylum | 523,054,329.2 | - |
| | class | 251,238,780.9 | - |
| | order | 189,276,427.7 | - |
| | family | 66,919,471.4 | 425,443.0 |
| | genus | 21,543,630.8 | 1,305,936.7 |
| | species | 4,044,983.6 | 1,345,684.4 |
| Average | all ranks | 235,799,604.4 | 355,314.1 |

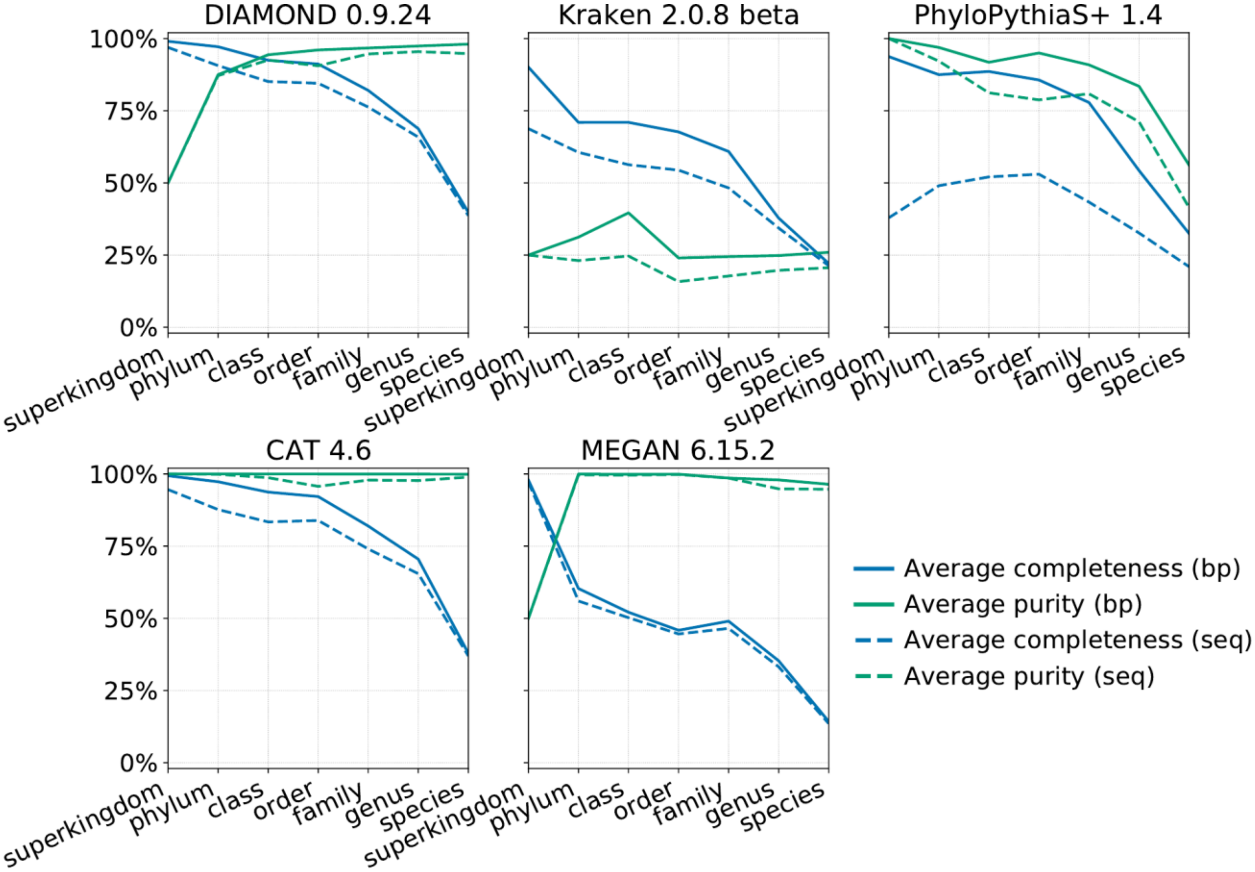


**Supplementary Fig. 1: Average completeness and purity based on base pair (bp, solid lines) and contig (seq, dashed lines) counts assessed for taxonomic bins predicted from the CAMI II mouse gut data set.** In the metrics based on bp counts, longer contigs have higher weight within a bin than shorter contigs, and better performance measured using these counts instead of contig counts indicates more accurate binning of longer contigs than shorter ones. Bp counts are used in the assessments in the main document (Fig. 4) and reproduced here as Average completeness (bp) and Average purity (bp). All bins contribute equally in the average computation. In this assessment, the smallest bins per method and rank were filtered out (overall 1% of the binned data in bp).

**Supplementary Table 11: Digital Object Identifiers (DOIs) and associated links of taxonomic profiling results and profiling ground truth for the 64 short read samples of the CAMI II mouse gut data set.**

| Taxonomic profiler | Results DOI | Links for commands used and download |
|---|---|---|
| MetaPhlAn 2.9.21 | 10.5281/zenodo.3629610 | https://doi.org/10.5281/zenodo.3629610 https://zenodo.org/record/3629610/files/cami2_mouse_gut_metaphlan2.9.21.profile?download=1 |
| MetaPhlAn 2.2.0 | 10.5281/zenodo.3629612 | https://doi.org/10.5281/zenodo.3629612 https://zenodo.org/record/3629612/files/cami2_mouse_gut_metaphlan2.2.0.profile?download=1 |
| Bracken 2.5 | 10.5281/zenodo.3629614 | https://doi.org/10.5281/zenodo.3629614 https://zenodo.org/record/3629614/files/cami2_mouse_gut_bracken2.5.profile?download=1 |
| FOCUS 0.31 | 10.5281/zenodo.3629620 | https://doi.org/10.5281/zenodo.3629620 https://zenodo.org/record/3629620/files/cami2_mouse_gut_focus0.31.profile?download=1 |
| CAMIARKQuikr 1.0.0 | 10.5281/zenodo.3629622 | https://doi.org/10.5281/zenodo.3629622 https://zenodo.org/record/3629622/files/cami2_mouse_gut_camiarkquikr1.0.0.profile?download=1 |
| mOTUs 1.1 | 10.5281/zenodo.3629624 | https://doi.org/10.5281/zenodo.3629624 https://zenodo.org/record/3629624/files/cami2_mouse_gut_motus1.1.profile?download=1 |
| mOTUs 2.5.1 | 10.5281/zenodo.3629626 | https://doi.org/10.5281/zenodo.3629626 https://zenodo.org/record/3629626/files/cami2_mouse_gut_motus2.5.1.profile?download=1 |
| MetaPalette 1.0.0 | 10.5281/zenodo.3629628 | https://doi.org/10.5281/zenodo.3629628 https://zenodo.org/record/3629628/files/cami2_mouse_gut_metapalette1.0.0.profile?download=1 |
| TIPP 2.0.0 | 10.5281/zenodo.3629630 | https://doi.org/10.5281/zenodo.3629630 https://zenodo.org/record/3629630/files/cami2_mouse_gut_tipp2.0.0.profile?download=1 |
| MetaPhyler 1.25 | 10.5281/zenodo.3629632 | https://doi.org/10.5281/zenodo.3629632 https://zenodo.org/record/3629632/files/cami2_mouse_gut_metaphyler1.25.profile?download=1 |
| Profiling ground truth | 10.5281/zenodo.3632528 | https://doi.org/10.5281/zenodo.3632528 https://zenodo.org/record/3632528/files/cami2_mouse_gut_gs.profile?download=1 |

**Supplementary Table 12: Elapsed (wall clock) time (h:mm) and maximum resident set size (kbytes) of taxonomic profiling methods on the 64 short read samples of the CAMI II mouse gut data set.** The best results are shown in bold. Bracken requires to run Kraken, hence the times required to run Bracken and both tools are shown. The taxonomic profilers were run on a computer with an Intel Xeon E5-4650 v4 CPU (virtualized to 16 CPU cores, 1 thread per core) and 512 GB (536.870.912 kbytes) of main memory.

| Taxonomic binner | Time (hh:mm) | Memory (kbytes) |
|---|---:|---:|
| MetaPhlAn 2.9.21 | 18:44 | 5,139,172 |
| MetaPhlAn 2.2.0 | 12:30 | 1,741,304 |
| Bracken 2.5 (only Bracken) | **0:01** | **24,472** |
| Bracken 2.5 (Kraken and Bracken) | **3:03** | 39,439,796 |
| FOCUS 0.31 | 13:27 | 5,236,199 |
| CAMIARKQuikr 1.0.0 | 16:19 | 27,391,555 |
| mOTUs 1.1 | 19:50 | **1,251,296** |
| mOTUs 2.5.1 | 14:29 | 3,922,448 |
| MetaPalette 1.0.0 | 76:49 | 27,297,132 |
| TIPP 2.0.0 | 151:01 | 70,789,939 |
| MetaPhyler 1.25 | 119:30 | 2,684,720 |

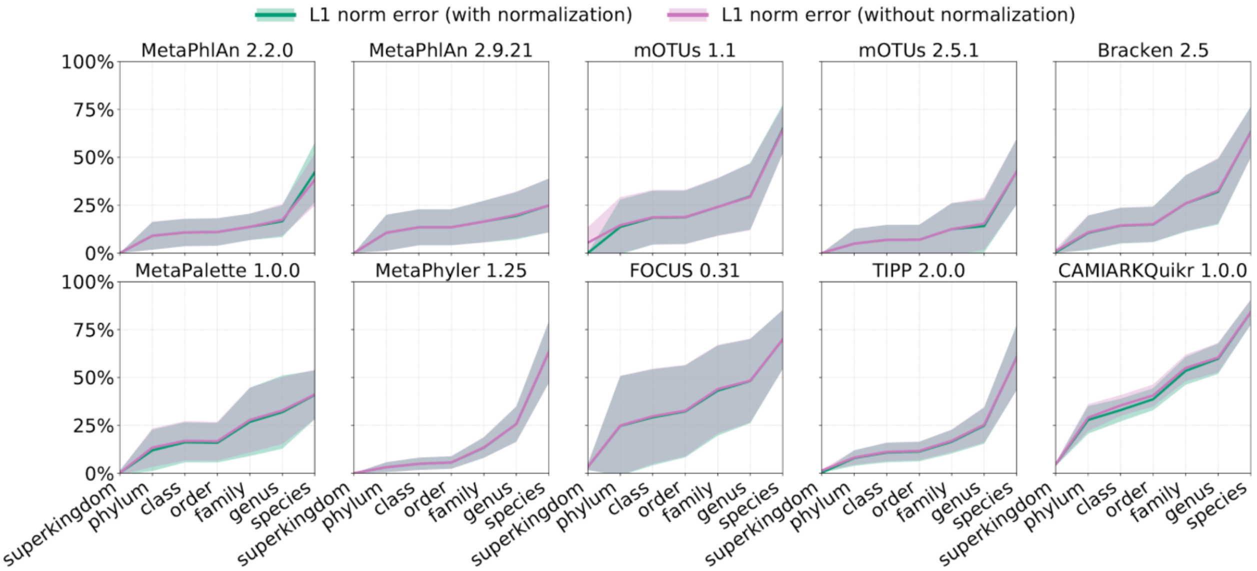


**Supplementary Fig. 2: L1 norm error computed with and without prior normalization of the predicted organismal relative abundances per method and taxonomic rank (such that predicted taxon abundances sum to 100%, dismissing abundances of unassigned taxa at that rank) on the 64 short read samples of the CAMI II mouse gut data set.**

**Supplementary Table 13: Average sum of predicted taxon abundances per taxonomic profiler and rank for the 64 short read samples of the CAMI II mouse gut data set.**

| Taxonomic profiler | superkingdom | phylum | class | order | family | genus | species |
|---|---:|---:|---:|---:|---:|---:|---:|
| MetaPhlAn 2.2.0 | 99.99 | 99.99 | 99.99 | 99.99 | 99.98 | 97.49 | 84.43 |
| MetaPhlAn 2.9.21 | 100.00 | 99.99 | 99.99 | 99.99 | 99.92 | 95.84 | 100.00 |
| mOTUs 1.1 | 90.66 | 90.66 | 90.66 | 90.66 | 90.63 | 86.39 | 90.66 |
| mOTUs 2.5.1 | 100.00 | 99.99 | 99.99 | 99.99 | 99.99 | 99.99 | 99.99 |
| Bracken 2.5 | 98.69 | 98.66 | 98.53 | 98.63 | 98.26 | 97.58 | 98.69 |
| MetaPalette 1.0.0 | 99.99 | 92.12 | 92.11 | 92.12 | 91.25 | 86.10 | 90.98 |
| MetaPhyler 1.25 | 100.00 | 100.00 | 99.85 | 99.96 | 98.67 | 97.84 | 99.99 |
| FOCUS 0.31 | 99.95 | 99.40 | 98.13 | 94.99 | 92.60 | 94.40 | 99.99 |
| TIPP 2.0.0 | 97.83 | 97.83 | 97.60 | 97.71 | 96.58 | 96.35 | 97.83 |
| CAMIARKQuikr 1.0.0 | 99.99 | 95.01 | 90.71 | 92.18 | 90.69 | 91.87 | 97.71 |
