## Supplementary results: MetaQUAST metrics for "Tutorial: Assessing metagenomics software with the CAMI benchmarking toolkit": report.html

### QUAST

**Quality Assessment Tool for Genome Assemblies**
by CAB

Loading...

Aligned to
""

Combined reference
Estimated reference size:
 bp
|
 bp
|
 references
|
 fragments
|
 % G+C
|
 chromosomes
  
 reads
|
% mapped
|
% properly paired
  
 genomic features
|
 operons

Unfortunately, JavaScript in your
browser is disabled or is not supported.
We need JavaScript to build report and plots.

Worst
Median
Best

Show heatmap

|  |
| --- |
| Contigs are ordered from largest (contig #1) to smallest.  FRCurve: Y is the total number of aligned bases divided by the reference length, in the contigs having the total number of  at most X.  Contigs are broken into nonoverlapping 100 bp windows. Plot shows number of windows for each GC percentage.  Plot shows number of contigs with GC percentage in a certain range. |

[{"date":"10 February 2020, Monday, 16:10:03","assembliesNames":["megahit-103-df","megahit-113-df","megahit-113-ml","megahit-113-ms","megahit-129-df","metaSPAdes"],"referenceName":"combined\_reference","order":[0,1,2,3,4,5],"report":[["Genome statistics",[{"metricName":"Genome fraction (%)","quality":"More is better","values":["23.507","26.164","26.039","26.292","26.691","23.262"],"isMain":true},{"metricName":"Duplication ratio","quality":"Less is better","values":["1.023","1.037","1.046","1.050","1.034","1.017"],"isMain":true},{"metricName":"Largest alignment","quality":"More is better","values":[354703,904953,859640,753008,787657,1034619],"isMain":true},{"metricName":"Total aligned length","quality":"More is better","values":[436725459,492514960,493969107,500306789,500856984,429280747],"isMain":true},{"metricName":"NA50","quality":"More is better","values":[4357,7357,3708,3767,7566,11658],"isMain":false},{"metricName":"NA75","quality":"More is better","values":[1126,1319,952,939,1289,1524],"isMain":false},{"metricName":"LA50","quality":"Less is better","values":[16688,9952,16415,16495,9768,5481],"isMain":false},{"metricName":"LA75","quality":"Less is better","values":[71505,55692,94073,95761,57261,36368],"isMain":false}]],["Reads mapping",[]],["Misassemblies",[{"metricName":"# misassemblies","quality":"Less is better","values":[5770,8685,5336,9381,8807,3488],"isMain":true},{"metricName":" # relocations","quality":"Less is better","values":[216,476,437,583,554,142],"isMain":false},{"metricName":" # translocations","quality":"Less is better","values":[913,1881,1715,2115,2147,1063],"isMain":false},{"metricName":" # inversions","quality":"Less is better","values":[174,422,265,313,361,4],"isMain":false},{"metricName":" # interspecies translocations","quality":"Less is better","values":[4467,5906,2919,6370,5745,2279],"isMain":false},{"metricName":"# misassembled contigs","quality":"Less is better","values":[5646,8219,4953,8675,8228,3247],"isMain":false},{"metricName":"Misassembled contigs length","quality":"Less is better","values":[10879967,43068359,34576388,56221107,50536067,25409676],"isMain":true},{"metricName":"# possibly misassembled contigs","quality":"Less is better","values":[30,87,72,122,94,16],"isMain":false},{"metricName":" # possible misassemblies","quality":"Less is better","values":[31,105,79,140,109,19],"isMain":false},{"metricName":"# local misassemblies","quality":"Less is better","values":[384,732,691,834,874,152],"isMain":false},{"metricName":"# scaffold gap ext. mis.","quality":"Less is better","values":[0,0,0,0,0,0],"isMain":false},{"metricName":"# scaffold gap loc. mis.","quality":"Less is better","values":[0,0,0,0,0,0],"isMain":false},{"metricName":"# unaligned mis. contigs","quality":"Less is better","values":[3,16,28,30,19,0],"isMain":false}]],["Unaligned",[{"metricName":"# fully unaligned contigs","quality":"Less is better","values":[185,407,562,620,397,48],"isMain":false},{"metricName":"Fully unaligned length","quality":"Less is better","values":[114573,251600,339109,384200,247781,31569],"isMain":false},{"metricName":"# partially unaligned contigs","quality":"Less is better","values":[30,88,76,124,95,16],"isMain":false},{"metricName":"Partially unaligned length","quality":"Less is better","values":[20997,62787,52777,97345,72561,11336],"isMain":false}]],["Mismatches",[{"metricName":"# mismatches","quality":"Less is better","values":[2321237,2765121,4208738,4529651,2845660,1719000],"isMain":false},{"metricName":"# indels","quality":"Less is better","values":[10227,19864,18616,22735,20925,10895],"isMain":false},{"metricName":"Indels length","quality":"Less is better","values":[32654,67153,61576,80051,89197,33417],"isMain":false},{"metricName":"# mismatches per 100 kbp","quality":"Less is better","values":["542.07","580.14","887.27","945.71","585.26","405.65"],"isMain":true},{"metricName":"# indels per 100 kbp","quality":"Less is better","values":["2.39","4.17","3.92","4.75","4.30","2.57"],"isMain":true},{"metricName":" # indels (<= 5 bp)","quality":"Less is better","values":[9532,18314,17293,21007,18836,10127],"isMain":false},{"metricName":" # indels (> 5 bp)","quality":"Less is better","values":[695,1550,1323,1728,2089,768],"isMain":false},{"metricName":"# N's","quality":"Less is better","values":[0,0,0,0,0,0],"isMain":false},{"metricName":"# N's per 100 kbp","quality":"Less is better","values":["0.00","0.00","0.00","0.00","0.00","0.00"],"isMain":true}]],["Statistics without reference",[{"metricName":"# contigs","quality":"Equal","values":[225585,220757,278807,282136,225167,174693],"isMain":true},{"metricName":"# contigs (>= 0 bp)","quality":"Equal","values":[906440,473755,644349,633018,466764,944240],"isMain":false},{"metricName":"# contigs (>= 1000 bp)","quality":"Equal","values":[83834,78193,87734,87026,78922,61277],"isMain":false},{"metricName":"# contigs (>= 5000 bp)","quality":"Equal","values":[14486,14377,11965,12154,14090,11228],"isMain":false},{"metricName":"# contigs (>= 10000 bp)","quality":"Equal","values":[6562,7268,6227,6367,7374,6183],"isMain":false},{"metricName":"# contigs (>= 25000 bp)","quality":"Equal","values":[1707,2571,2265,2391,2795,2553],"isMain":false},{"metricName":"# contigs (>= 50000 bp)","quality":"Equal","values":[471,1032,863,887,1107,1132],"isMain":false},{"metricName":"Largest contig","quality":"More is better","values":[354703,904953,859640,754056,788697,1034619],"isMain":true},{"metricName":"Total length","quality":"More is better","values":[438032656,494653238,496722592,503491159,503073431,430847014],"isMain":true},{"metricName":"Total length (>= 0 bp)","quality":"More is better","values":[656714148,592169014,635645417,637431469,596517067,649949394],"isMain":false},{"metricName":"Total length (>= 1000 bp)","quality":"More is better","values":[342669622,399682035,368806791,372764886,405211262,354794894],"isMain":true},{"metricName":"Total length (>= 5000 bp)","quality":"More is better","values":[209586919,278087012,232592249,238510195,283234943,261084168],"isMain":false},{"metricName":"Total length (>= 10000 bp)","quality":"More is better","values":[154362921,228640882,192818790,198110070,236255195,225930387],"isMain":true},{"metricName":"Total length (>= 25000 bp)","quality":"More is better","values":[80681912,156259069,131371435,136056759,165014101,169277911],"isMain":false},{"metricName":"Total length (>= 50000 bp)","quality":"More is better","values":[38821616,102990325,82724532,83865010,106551070,119684054],"isMain":true},{"metricName":"N50","quality":"More is better","values":[4406,7710,3836,3964,8087,12077],"isMain":false},{"metricName":"N75","quality":"More is better","values":[1145,1364,972,965,1333,1555],"isMain":false},{"metricName":"L50","quality":"Less is better","values":[16496,9405,15577,15138,9079,5227],"isMain":false},{"metricName":"L75","quality":"Less is better","values":[70586,53392,91524,91969,54548,35419],"isMain":false}]],["K-mer-based statistics",[]],["Predicted genes",[]],["Similarity statistics",[]],["Reference statistics",[{"metricName":"Reference length","quality":"Equal","values":[1821690894,1821690894,1821690894,1821690894,1821690894,1821690894],"isMain":false},{"metricName":"Reference fragments","quality":"Equal","values":[38084,38084,38084,38084,38084,38084],"isMain":false}]]],"subreferences":["GCF\_000006625.1\_ASM662v1","GCF\_000009925.1\_ASM992v1","GCF\_000010505.1\_ASM1050v1","GCF\_000011045.1\_ASM1104v1","GCF\_000011065.1\_ASM1106v1","GCF\_000012845.1\_ASM1284v1","GCF\_000015325.1\_ASM1532v1","GCF\_000018685.1\_ASM1868v1","GCF\_000020225.1\_ASM2022v1","GCF\_000022065.1\_ASM2206v1","GCF\_000026525.1\_ASM2652v1","GCF\_000091345.1\_ASM9134v1","GCF\_000091405.1\_ASM9140v1","GCF\_000142965.1\_ASM14296v1","GCF\_000144405.1\_ASM14440v1","GCF\_000144625.1\_ASM14462v1","GCF\_000145035.1\_ASM14503v1","GCF\_000153905.1\_ASM15390v1","GCF\_000153925.1\_ASM15392v1","GCF\_000154065.1\_ASM15406v1","GCF\_000154105.1\_ASM15410v1","GCF\_000154125.1\_ASM15412v1","GCF\_000154345.1\_ASM15434v1","GCF\_000154385.1\_ASM15438v1","GCF\_000154425.1\_ASM15442v1","GCF\_000154465.1\_ASM15446v1","GCF\_000154505.1\_ASM15450v1","GCF\_000155815.1\_ASM15581v1","GCF\_000155915.2\_Lacto\_jensenii\_1153\_V2","GCF\_000156035.2\_ASM15603v2","GCF\_000156075.1\_ASM15607v1","GCF\_000156175.1\_ASM15617v1","GCF\_000156495.1\_ASM15649v1","GCF\_000156515.1\_ASM15651v1","GCF\_000157015.1\_ASM15701v1","GCF\_000157035.2\_Bacteroides\_sp\_2\_1\_7\_V2","GCF\_000157075.2\_Bact\_sp\_9\_1\_42FAA\_V2","GCF\_000157915.1\_ASM15791v1","GCF\_000157935.1\_ASM15793v1","GCF\_000157955.1\_ASM15795v1","GCF\_000157975.1\_ASM15797v1","GCF\_000158075.1\_ASM15807v1","GCF\_000158555.2\_Erys\_bact\_D7\_V2","GCF\_000158655.1\_ASM15865v1","GCF\_000159215.1\_ASM15921v1","GCF\_000159615.1\_ASM15961v1","GCF\_000159875.2\_Bact\_sp\_1\_1\_6\_V2","GCF\_000159995.1\_ASM15999v1","GCF\_000160035.2\_ASM16003v2","GCF\_000160095.1\_ASM16009v1","GCF\_000160135.1\_ASM16013v1","GCF\_000160675.1\_ASM16067v1","GCF\_000162175.1\_ASM16217v1","GCF\_000162215.1\_ASM16221v1","GCF\_000162275.1\_ASM16227v1","GCF\_000162455.1\_ASM16245v1","GCF\_000163055.2\_ASM16305v2","GCF\_000163655.1\_ASM16365v1","GCF\_000165775.1\_ASM16577v3","GCF\_000177015.3\_Rumi\_bact\_D16\_V2","GCF\_000178115.2\_ASM17811v2","GCF\_000178835.2\_ASM17883v2","GCF\_000184945.1\_ASM18494v1","GCF\_000185525.1\_ASM18552v1","GCF\_000185545.1\_ASM18554v1","GCF\_000185605.1\_ASM18560v1","GCF\_000185705.2\_Bilo\_wads\_3\_1\_6\_V2","GCF\_000185825.1\_ASM18582v1","GCF\_000186105.1\_Bacteroides\_sp\_3\_1\_40A\_V1","GCF\_000187895.1\_ASM18789v1","GCF\_000190535.1\_ASM19053v1","GCF\_000191545.1\_ASM19154v1","GCF\_000191805.1\_Lach\_bact\_4\_1\_37FAA\_V1","GCF\_000195315.1\_ASM19531v1","GCF\_000196175.1\_ASM19617v1","GCF\_000205025.1\_ASM20502v1","GCF\_000205165.1\_ASM20516v1","GCF\_000209385.2\_Lach\_bact\_2\_1\_46\_FAA\_V2","GCF\_000209405.1\_Lach\_bact\_3\_1\_46FAA\_V1","GCF\_000209425.1\_Lach\_bact\_6\_1\_63FAA\_V1","GCF\_000209465.1\_Lach\_bact\_F0167\_V1","GCF\_000214785.1\_ASM21478v1","GCF\_000218385.1\_Lach\_bact\_1\_4\_56FAA\_V1","GCF\_000218405.2\_Lach\_bact\_3\_1\_57FAA\_CT1\_V1","GCF\_000218425.1\_Lach\_bact\_5\_1\_57FAA","GCF\_000218445.1\_Lach\_bact\_1\_1\_57FAA\_V1","GCF\_000220255.1\_ASM22025v1","GCF\_000224615.1\_Prev\_oulo\_F0390\_V1","GCF\_000224655.1\_Bilophila\_sp\_4\_1\_30\_V1","GCF\_000225345.1\_ASM22534v1","GCF\_000225685.1\_Erys\_bact\_2\_2\_44A\_V1","GCF\_000225745.1\_Dorea\_form\_4\_6\_53AFAA\_V1","GCF\_000231275.1\_Alis\_indi\_YIT\_V1","GCF\_000233455.1\_Clos\_citr\_WAL\_17108\_V1","GCF\_000234055.1\_Prev\_hist\_F0411\_V1","GCF\_000234115.1\_Prevotella\_sp\_F0323\_V1","GCF\_000234155.1\_Clos\_clos\_2\_1\_49FAA\_V1","GCF\_000235445.1\_John\_igna\_ATCC\_51276\_V1","GCF\_000236865.1\_ASM23686v1","GCF\_000237085.1\_ASM23708v1","GCF\_000238055.2\_Oribacterium\_sp\_ACB1\_V2","GCF\_000238075.1\_Oribacterium\_sp\_ACB7\_V1","GCF\_000238635.1\_Subdol\_sp\_4\_3\_54A2FAA\_V1","GCF\_000238835.1\_ASM23883v1","GCF\_000242315.1\_ASM24231v1","GCF\_000243015.1\_Prev\_macu\_OT\_289\_V1","GCF\_000243215.1\_Odor\_lane\_YIT\_12061\_V1","GCF\_000245055.1\_ASM24505v1","GCF\_000259235.1\_ASM25923v1","GCF\_000261025.1\_ASM26102v1","GCF\_000265365.1\_ASM26536v1","GCF\_000270225.1\_ASM27022v1","GCF\_000273035.1\_Bact\_dore\_CL02T00C15\_V1","GCF\_000273055.1\_Bact\_dore\_CL02T12C06\_V1","GCF\_000273135.1\_Bact\_frag\_CL05T12C13\_V1","GCF\_000273155.1\_Bact\_frag\_CL07T12C05\_V1","GCF\_000273175.1\_Bact\_nord\_CL02T12C05\_V1","GCF\_000273195.1\_Bact\_ovat\_CL02T12C04\_V1","GCF\_000273235.1\_Bact\_saly\_CL02T12C01\_V1","GCF\_000273295.1\_Bact\_vulg\_CL09T03C04\_V1","GCF\_000273315.1\_Bact\_xyla\_CL03T12C04\_V1","GCF\_000273765.1\_Bact\_frag\_CL05T00C42\_V1","GCF\_000296465.1\_Barn\_inte\_YIT\_11860\_V1","GCF\_000297695.1\_Bact\_frag\_HMW\_610\_V1","GCF\_000297735.1\_Bact\_frag\_HMW\_615\_V1","GCF\_000297755.1\_Bact\_frag\_HMW\_616\_V1","GCF\_000304195.1\_Bact\_thet\_CL09T03C10\_V1","GCF\_000307265.1\_ASM30726v1","GCF\_000307345.1\_Para\_merd\_CL03T12C32\_V1","GCF\_000307455.1\_Para\_dist\_CL03T12C09\_V1","GCF\_000307475.1\_Paraba\_sp\_D25\_V1","GCF\_000307495.1\_Para\_merd\_CL09T00C40\_V1","GCF\_000312145.1\_ASM31214v1","GCF\_000312405.1\_ASM31240v1","GCF\_000318195.2\_ASM31819v2","GCF\_000331715.1\_ASM33171v1","GCF\_000331995.1\_ASM33199v1","GCF\_000332875.2\_ASM33287v2","GCF\_000333275.1\_ASM33327v1","GCF\_000333595.1\_ASM33359v1","GCF\_000345685.2\_ASM34568v2","GCF\_000365025.1\_ASM36502v1","GCF\_000371405.1\_Clos\_clos\_CM201\_V1","GCF\_000371445.1\_Clos\_hath\_12489931\_V1","GCF\_000371485.1\_Clos\_clos\_90B1\_V1","GCF\_000371505.1\_Clos\_clos\_90A8\_V1","GCF\_000371525.1\_Clos\_clos\_90A7\_V1","GCF\_000371545.1\_Clos\_clos\_90A6\_V1","GCF\_000371585.1\_Clos\_clos\_90A3\_V1","GCF\_000371605.1\_Clos\_clos\_90A1\_V1","GCF\_000371685.1\_Clos\_bolt\_90B3\_V1","GCF\_000371705.1\_Clos\_bolt\_90A9\_V1","GCF\_000373885.1\_ASM37388v1","GCF\_000374365.1\_ASM37436v1","GCF\_000374505.1\_ASM37450v1","GCF\_000374585.1\_ASM37458v1","GCF\_000379925.1\_ASM37992v1","GCF\_000379965.1\_ASM37996v1","GCF\_000381365.1\_Bact\_saly\_DSM\_18765\_V1","GCF\_000382385.1\_ASM38238v1","GCF\_000382445.1\_Bact\_mass\_DSM\_17679\_V1","GCF\_000382465.1\_Bact\_sp\_HPS0048\_V1","GCF\_000383295.1\_ASM38329v1","GCF\_000384195.1\_ASM38419v1","GCF\_000391985.1\_Ente\_faec\_9731349\_1\_V1","GCF\_000396085.1\_Ente\_faec\_B1933\_V1","GCF\_000396285.1\_Ente\_faec\_B2867\_V1","GCF\_000398925.1\_Buty\_pull\_1\_20\_V1","GCF\_000403155.2\_Bact\_thet\_dnLKV9\_V1","GCF\_000403235.1\_Bact\_vulg\_dnLKV7\_V1","GCF\_000403255.1\_Lach\_bact\_3\_1\_V1","GCF\_000403275.1\_Lach\_bact\_28\_4\_V1","GCF\_000403355.2\_Ente\_caec\_B7\_V1","GCF\_000403435.2\_Osci\_bact\_1\_3\_V1","GCF\_000403455.2\_Dore\_bact\_5\_2\_V1","GCF\_000404205.1\_ASM40420v1","GCF\_000410615.1\_ASM41061v1","GCF\_000410995.1\_ASM41099v1","GCF\_000411355.1\_Copr\_sp\_HPP0048\_V1","GCF\_000413355.1\_Prev\_oral\_HGA0225\_V1","GCF\_000422625.1\_ASM42262v1","GCF\_000423925.1\_ASM42392v1","GCF\_000423945.1\_ASM42394v1","GCF\_000423985.1\_ASM42398v1","GCF\_000424005.1\_ASM42400v1","GCF\_000424085.1\_ASM42408v1","GCF\_000424105.1\_ASM42410v1","GCF\_000424125.1\_ASM42412v1","GCF\_000424305.1\_ASM42430v1","GCF\_000424365.1\_ASM42436v1","GCF\_000424405.1\_ASM42440v1","GCF\_000424425.1\_ASM42442v1","GCF\_000424465.1\_ASM42446v1","GCF\_000424585.1\_ASM42458v1","GCF\_000424605.1\_ASM42460v1","GCF\_000425525.1\_ASM42552v1","GCF\_000426305.1\_ASM42630v1","GCF\_000426505.1\_ASM42650v1","GCF\_000427365.1\_ASM42736v1","GCF\_000428565.1\_ASM42856v1","GCF\_000429145.1\_ASM42914v1","GCF\_000455285.1\_Sm\_AP9","GCF\_000466505.1\_ASM46650v1","GCF\_000467795.1\_ASM46779v1","GCF\_000467835.1\_ASM46783v1","GCF\_000467855.2\_ASM46785v2","GCF\_000467875.1\_ASM46787v1","GCF\_000468015.1\_ASM46801v1","GCF\_000469425.1\_ASM46942v1","GCF\_000473955.1\_\_oprFasNSB1","GCF\_000487995.1\_Muci\_scha\_ASF457\_V1","GCF\_000493555.1\_LhCIRMBIA101V1","GCF\_000498675.1\_ASM49867v1","GCF\_000507805.1\_Rumi\_gnav\_CC55\_001C\_V1","GCF\_000512915.1\_ASM51291v1","GCF\_000515315.1\_ASM51531v1","GCF\_000539515.1\_Stap\_aure\_LAMC0001\_V1","GCF\_000586615.1\_RaSY3","GCF\_000621465.1\_ASM62146v1","GCF\_000621565.1\_ASM62156v1","GCF\_000621605.1\_ASM62160v1","GCF\_000621765.1\_ASM62176v1","GCF\_000621785.1\_ASM62178v1","GCF\_000621805.1\_ASM62180v1","GCF\_000621825.1\_ASM62182v1","GCF\_000621865.1\_ASM62186v1","GCF\_000621905.1\_ASM62190v1","GCF\_000621925.1\_ASM62192v1","GCF\_000622065.1\_ASM62206v1","GCF\_000622085.1\_ASM62208v1","GCF\_000686665.1\_ASM68666v1","GCF\_000686965.1\_ASM68696v1","GCF\_000700805.1\_ASM70080v1","GCF\_000701545.1\_ASM70154v1","GCF\_000701645.1\_ASM70164v1","GCF\_000701705.1\_ASM70170v1","GCF\_000701765.1\_ASM70176v1","GCF\_000702085.1\_ASM70208v1","GCF\_000702145.1\_ASM70214v1","GCF\_000702245.1\_ASM70224v1","GCF\_000702685.1\_ASM70268v1","GCF\_000702825.1\_ASM70282v1","GCF\_000711825.1\_ASM71182v1","GCF\_000723745.2\_PRJEB6027\_Urmite","GCF\_000739415.1\_ASM73941v1","GCF\_000745455.1\_ASM74545v1","GCF\_000745845.1\_ASM74584v1","GCF\_000765245.1\_ASM76524v1","GCF\_000766005.1\_ASM76600v1","GCF\_000769385.1\_ASM76938v1","GCF\_000789395.1\_ASM78939v1","GCF\_000821285.2\_Alistipes\_jeddahmassiliense\_str.\_AL\_1","GCF\_000875945.1\_TWA4\_1","GCF\_000934625.1\_ASM93462v1","GCF\_000953535.1\_E5B","GCF\_000967245.1\_ASM96724v1","GCF\_000969825.1\_Para\_gord\_MS\_1\_V1","GCF\_000969835.1\_Para\_gold\_DSM\_19448\_V1","GCF\_000969845.1\_Para\_sp\_HGS0025\_V1","GCF\_000970735.1\_ASM97073v1","GCF\_000971515.1\_ASM97151v1","GCF\_001011095.1\_ASM101109v1","GCF\_001012685.1\_IM18\_3v1","GCF\_001012735.1\_IM1v1","GCF\_001034445.1\_Citr\_freu\_complex\_MGH99\_V1","GCF\_001050435.1\_ASM105043v1","GCF\_001056335.1\_ASM105633v1","GCF\_001062165.1\_ASM106216v1","GCF\_001066565.1\_ASM106656v1","GCF\_001073445.1\_ASM107344v1","GCF\_001078425.1\_Clos\_bolt\_WAL\_14578\_V1","GCF\_001115565.1\_6259\_7\_5","GCF\_001185345.1\_ASM118534v2","GCF\_001188985.1\_ASM118898v1","GCF\_001244405.1\_Eubacterium\_massiliense","GCF\_001244995.1\_Intestinimonas\_massiliensis","GCF\_001261775.1\_Anaeromassilibacillus\_senegalense","GCF\_001274615.1\_ASM127461v1","GCF\_001305595.1\_ASM130559v1","GCF\_001403575.1\_Ihuella\_massiliensis","GCF\_001403595.1\_Gabonia\_massiliensis","GCF\_001412315.1\_ASM141231v1","GCF\_001414325.1\_ASM141432v1","GCF\_001429415.1\_Root670","GCF\_001434845.1\_ASM143484v1","GCF\_001435065.1\_ASM143506v1","GCF\_001436015.1\_ASM143601v1","GCF\_001436185.1\_ASM143618v1","GCF\_001436945.1\_ASM143694v1","GCF\_001454945.1\_ASM145494v1","GCF\_001461035.1\_ASM146103v1","GCF\_001487105.1\_Ruminococcus\_phoceensis","GCF\_001487165.1\_Blautia\_massiliensis1","GCF\_001535595.1\_ASM153559v1","GCF\_001535605.1\_ASM153560v1","GCF\_001543345.1\_ASM154334v1","GCF\_001548195.1\_ASM154819v1","GCF\_001552975.1\_ASM155297v1","GCF\_001553145.1\_ASM155314v1","GCF\_001553165.1\_ASM155316v1","GCF\_001553215.1\_ASM155321v1","GCF\_001561955.1\_ASM156195v1","GCF\_001578555.1\_ASM157855v1","GCF\_001578575.1\_ASM157857v1","GCF\_001578635.1\_ASM157863v1","GCF\_001580105.1\_ASM158010v1","GCF\_001580195.1\_ASM158019v1","GCF\_001580455.1\_ASM158045v1","GCF\_001592745.1\_ASM159274v1","GCF\_001594285.1\_ASM159428v1","GCF\_001611075.1\_ASM161107v1","GCF\_001636135.1\_ASM163613v1","GCF\_001647615.1\_ASM164761v1","GCF\_001659745.1\_ASM165974v1","GCF\_001678845.1\_ASM167884v1","GCF\_001682215.1\_ASM168221v1","GCF\_001688625.2\_ASM168862v2","GCF\_001688665.2\_ASM168866v2","GCF\_001688705.2\_ASM168870v2","GCF\_001688845.2\_ASM168884v2","GCF\_001689125.2\_ASM168912v2","GCF\_001692755.1\_ASM169275v1","GCF\_001697245.1\_ASM169724v1","GCF\_001697285.1\_ASM169728v1","GCF\_001702095.1\_ASM170209v1","GCF\_001704315.1\_ASM170431v1","GCF\_001704645.1\_ASM170464v1","GCF\_001715615.1\_ASM171561v1","GCF\_001717125.1\_ASM171712v1","GCF\_001721925.1\_ASM172192v1","GCF\_001746265.1\_ASM174626v2","GCF\_001750685.1\_ASM175068v1","GCF\_001808555.1\_ASM180855v1","GCF\_001811315.1\_ASM181131v1","GCF\_001811695.1\_ASM181169v1","GCF\_001814855.1\_ASM181485v1","GCF\_001815255.1\_ASM181525v1","GCF\_001831275.1\_ASM183127v1","GCF\_001922325.1\_ASM192232v1","GCF\_001938785.1\_ASM193878v1","GCF\_001951175.1\_ASM195117v1","GCF\_001953935.1\_ASM195393v1","GCF\_001981595.1\_ASM198159v1","GCF\_001998765.1\_ASM199876v1","GCF\_002000665.1\_ASM200066v1","GCF\_002003555.1\_ASM200355v1","GCF\_002078855.1\_ASM207885v1","GCF\_002079325.1\_ASM207932v1","GCF\_002079435.1\_ASM207943v1","GCF\_002079585.1\_ASM207958v1","GCF\_002079925.1\_ASM207992v1","GCF\_002091975.1\_ASM209197v1","GCF\_002095655.1\_ASM209565v1","GCF\_002117305.1\_ASM211730v1","GCF\_002119605.1\_ASM211960v1","GCF\_002201475.1\_ASM220147v1","GCF\_002201495.1\_ASM220149v1","GCF\_002204455.1\_ASM220445v1","GCF\_002206085.1\_SJD4","GCF\_002217985.1\_ASM221798v1","GCF\_002218605.1\_ASM221860v1","GCF\_002218725.1\_ASM221872v1","GCF\_002218965.1\_ASM221896v1","GCF\_002221555.2\_ASM222155v2","GCF\_002222595.2\_ASM222259v2","GCF\_002253685.1\_ASM225368v1","GCF\_002254255.1\_ASM225425v1","GCF\_002257605.1\_ASM225760v1","GCF\_002278095.1\_ASM227809v1","GCF\_002442835.1\_ASM244283v1","GCF\_002549775.1\_ASM254977v1","GCF\_002549905.1\_ASM254990v1","GCF\_002549935.1\_ASM254993v1","GCF\_002549945.1\_ASM254994v1","GCF\_002549975.1\_ASM254997v1","GCF\_002550015.1\_ASM255001v1","GCF\_002706705.1\_ASM270670v1","GCF\_002751085.1\_ASM275108v1","GCF\_002753955.1\_ASM275395v1","GCF\_002754015.1\_ASM275401v1","GCF\_002754035.1\_ASM275403v1","GCF\_002754055.1\_ASM275405v1","GCF\_002754095.1\_ASM275409v1","GCF\_002754115.1\_ASM275411v1","GCF\_002763575.1\_ASM276357v1","GCF\_002763745.1\_ASM276374v1","GCF\_002763845.1\_ASM276384v1","GCF\_900018625.1\_13566\_1\_59","GCF\_900021155.1\_CHCKI003","GCF\_900067125.1\_ASM90006712v1","GCF\_900078295.2\_PRJEB13136","GCF\_900081375.1\_12641\_2\_43","GCF\_900083545.1\_PRJEB13794","GCF\_900095495.1\_B\_ovatus\_V975","GCF\_900096735.1\_TFUB20","GCF\_900097105.1\_WK001","GCF\_900099715.1\_IMG\_taxon\_2593339241\_annotated\_assembly","GCF\_900100245.1\_IMG\_taxon\_2606217752\_annotated\_assembly","GCF\_900100465.1\_IMG\_taxon\_2654588146\_annotated\_assembly","GCF\_900100545.1\_IMG\_taxon\_2651870321\_annotated\_assembly","GCF\_900100595.1\_IMG\_taxon\_2593339217\_annotated\_assembly","GCF\_900101295.1\_IMG\_taxon\_2606217764\_annotated\_assembly","GCF\_900101325.1\_IMG\_taxon\_2596583543\_annotated\_assembly","GCF\_900101605.1\_IMG\_taxon\_2595698227\_annotated\_assembly","GCF\_900101965.1\_IMG\_taxon\_2675903020\_annotated\_assembly","GCF\_900103115.1\_IMG\_taxon\_2654588126\_annotated\_assembly","GCF\_900103235.1\_IMG\_taxon\_2593339151\_annotated\_assembly","GCF\_900103835.1\_IMG\_taxon\_2667527408\_annotated\_assembly","GCF\_900104335.1\_IMG\_taxon\_2593339244\_annotated\_assembly","GCF\_900104565.1\_PRJEB16354","GCF\_900104615.1\_PRJEB16353","GCF\_900104665.1\_PRJEB16357","GCF\_900104675.1\_PRJEB16361","GCF\_900106615.1\_IMG\_taxon\_2651870364\_annotated\_assembly","GCF\_900106755.1\_IMG\_taxon\_2693429916\_annotated\_assembly","GCF\_900106845.1\_PRJEB14233","GCF\_900107485.1\_IMG\_taxon\_2599185206\_annotated\_assembly","GCF\_900107545.1\_IMG\_taxon\_2593339240\_annotated\_assembly","GCF\_900107575.1\_IMG\_taxon\_2593339166\_annotated\_assembly","GCF\_900107675.1\_IMG\_taxon\_2693429854\_annotated\_assembly","GCF\_900107785.1\_IMG\_taxon\_2654588140\_annotated\_assembly","GCF\_900107835.1\_IMG\_taxon\_2657244910\_annotated\_assembly","GCF\_900108375.1\_IMG\_taxon\_2596583684\_annotated\_assembly","GCF\_900108895.1\_IMG\_taxon\_2654588144\_annotated\_assembly","GCF\_900109005.1\_IMG\_taxon\_2617270830\_annotated\_assembly","GCF\_900109085.1\_IMG\_taxon\_2616644817\_annotated\_assembly","GCF\_900109215.1\_IMG\_taxon\_2654588124\_annotated\_assembly","GCF\_900109435.1\_IMG\_taxon\_2593339146\_annotated\_assembly","GCF\_900109635.1\_IMG\_taxon\_2623620522\_annotated\_assembly","GCF\_900109715.1\_IMG\_taxon\_2595698239\_annotated\_assembly","GCF\_900109965.1\_IMG\_taxon\_2593339260\_annotated\_assembly","GCF\_900110045.1\_IMG\_taxon\_2667527384\_annotated\_assembly","GCF\_900110805.1\_IMG\_taxon\_2593339242\_annotated\_assembly","GCF\_900111135.1\_IMG\_taxon\_2684622850\_annotated\_assembly","GCF\_900111185.1\_IMG\_taxon\_2623620520\_annotated\_assembly","GCF\_900111355.1\_IMG\_taxon\_2634166323\_annotated\_assembly","GCF\_900111435.1\_IMG\_taxon\_2654588206\_annotated\_assembly","GCF\_900111575.1\_IMG\_taxon\_2602042032\_annotated\_assembly","GCF\_900112775.1\_IMG\_taxon\_2599185221\_annotated\_assembly","GCF\_900112855.1\_IMG\_taxon\_2606217811\_annotated\_assembly","GCF\_900113595.1\_IMG\_taxon\_2657245270\_annotated\_assembly","GCF\_900113655.1\_IMG\_taxon\_2606217763\_annotated\_assembly","GCF\_900113995.1\_IMG\_taxon\_2593339255\_annotated\_assembly","GCF\_900114115.1\_IMG\_taxon\_2622736590\_annotated\_assembly","GCF\_900114365.1\_IMG\_taxon\_2651870357\_annotated\_assembly","GCF\_900114405.1\_IMG\_taxon\_2654588176\_annotated\_assembly","GCF\_900115355.1\_IMG\_taxon\_2623620483\_annotated\_assembly","GCF\_900115365.1\_IMG\_taxon\_2634166351\_annotated\_assembly","GCF\_900115435.1\_IMG\_taxon\_2623620458\_annotated\_assembly","GCF\_900115855.1\_IMG\_taxon\_2623620476\_annotated\_assembly","GCF\_900116075.1\_IMG\_taxon\_2654588129\_annotated\_assembly","GCF\_900116395.1\_IMG\_taxon\_2595698216\_annotated\_assembly","GCF\_900119155.1\_IMG\_taxon\_2593339149\_annotated\_assembly","GCF\_900119285.1\_IMG\_taxon\_2593339271\_annotated\_assembly","GCF\_900119535.1\_IMG\_taxon\_2606217757\_annotated\_assembly","GCF\_900120295.1\_PRJEB18018","GCF\_900120395.1\_PRJEB18035","GCF\_900128445.1\_PRJEB18031","GCF\_900128485.1\_PRJEB18048","GCF\_900128495.1\_PRJEB18049","GCF\_900128505.1\_PRJEB18032","GCF\_900128905.1\_IMG\_taxon\_2695420932\_annotated\_assembly","GCF\_900129025.1\_IMG\_taxon\_2695420943\_annotated\_assembly","GCF\_900129535.1\_IMG\_taxon\_2695420955\_annotated\_assembly","GCF\_900129565.1\_IMG\_taxon\_2675903007\_annotated\_assembly","GCF\_900130025.1\_IMG\_taxon\_2585428081\_annotated\_assembly","GCF\_900130065.1\_IMG\_taxon\_2585428085\_annotated\_assembly","GCF\_900130135.1\_PRJEB18269","GCF\_900141755.1\_IMG\_taxon\_2698536816\_annotated\_assembly","GCF\_900141865.1\_IMG\_taxon\_2585428091\_annotated\_assembly","GCF\_900142115.1\_IMG\_taxon\_2585428154\_annotated\_assembly","GCF\_900142165.1\_IMG\_taxon\_2585428171\_annotated\_assembly","GCF\_900142215.1\_IMG\_taxon\_2585428173\_annotated\_assembly","GCF\_900142265.1\_IMG\_taxon\_2585428190\_annotated\_assembly","GCF\_900142645.1\_IMG\_taxon\_2642422560\_annotated\_assembly","GCF\_900142955.1\_IMG\_taxon\_2588253509\_annotated\_assembly","GCF\_900143195.1\_501571.5","GCF\_900143205.1\_IMG\_taxon\_2582580726\_annotated\_assembly","GCF\_900143645.1\_IMG\_taxon\_2582580727\_annotated\_assembly","GCF\_900155365.1\_PRJEB18748","GCF\_900155425.1\_PRJEB18759","GCF\_900155865.1\_PRJEB18812","GCF\_900157245.1\_15\_9","GCF\_900157255.1\_ATCC49417","GCF\_900157265.1\_A7A1\_28","GCF\_900157275.1\_84\_3","GCF\_900157285.1\_13\_1","GCF\_900157295.1\_11A","GCF\_900157345.1\_AFR\_5B1","GCF\_900167005.1\_IMG\_taxon\_2585428076\_annotated\_assembly","GCF\_900167105.1\_IMG\_taxon\_2582581320\_annotated\_assembly","GCF\_900167555.1\_IMG\_taxon\_2590828806\_annotated\_assembly","GCF\_900168155.1\_IMG\_taxon\_2595698220\_annotated\_assembly","GCF\_900176335.1\_IMG\_taxon\_2588254267\_annotated\_assembly","GCF\_900184885.1\_MOUSE\_M89\_JASPER.fasta","GCF\_900184965.1\_Mouse\_2\_MG.fasta","GCF\_900184975.1\_Chimpanzee\_2\_2.fasta","GCF\_900199595.1\_IMG\_taxon\_2623620511\_annotated\_assembly","GCF\_900234395.1\_JOUY54\_assembly\_v1"],"subreports":[[["Genome statistics",[]],["Reads mapping",[]],["Misassemblies",[{"metricName":"# misassemblies","quality":"Less is better","values":[10,17,4,19,10,15],"isMain":true},{"metricName":" # relocations","quality":"Less is better","values":[0,4,2,2,0,2],"isMain":false},{"metricName":" # translocations","quality":"Less is better","values":[0,0,0,0,0,0],"isMain":false},{"metricName":" # inversions","quality":"Less is better","values":[1,2,2,1,3,0],"isMain":false},{"metricName":" # interspecies translocations","quality":"Less is better","values":[9,11,0,16,7,13],"isMain":false},{"metricName":"# possibly misassembled contigs","quality":"Less is better","values":[0,0,0,0,0,0],"isMain":false},{"metricName":" # possible misassemblies","quality":"Less is better","values":[0,0,0,0,0,0],"isMain":false},{"metricName":"# local misassemblies","quality":"Less is better","values":[0,0,0,0,2,0],"isMain":false},{"metricName":"# scaffold gap ext. mis.","quality":"Less is better","values":[0,0,0,0,0,0],"isMain":false},{"metricName":"# scaffold gap loc. mis.","quality":"Less is better","values":[0,0,0,0,0,0],"isMain":false}]],["Unaligned",[]],["Mismatches",[]],["Statistics without reference",[]],["K-mer-based statistics",[]],["Predicted genes",[]],["Similarity statistics",[]],["Reference statistics",[]]],[["Genome statistics",[]],["Reads mapping",[]],["Misassemblies",[{"metricName":"# misassemblies","quality":"Less is better","values":[27,24,25,33,15,1],"isMain":true},{"metricName":" # relocations","quality":"Less is better","values":[1,1,3,1,4,0],"isMain":false},{"metricName":" # translocations","quality":"Less is better","values":[0,0,0,0,0,0],"isMain":false},{"metricName":" # inversions","quality":"Less is better","values":[0,2,0,1,3,0],"isMain":false},{"metricName":" # interspecies translocations","quality":"Less is better","values":[26,21,22,31,8,1],"isMain":false},{"metricName":"# possibly misassembled contigs","quality":"Less is better","values":[0,0,0,0,1,0],"isMain":false},{"metricName":" # possible misassemblies","quality":"Less is better","values":[0,0,0,0,1,0],"isMain":false},{"metricName":"# local misassemblies","quality":"Less is better","values":[1,3,0,1,2,0],"isMain":false},{"metricName":"# scaffold gap ext. mis.","quality":"Less is better","values":[0,0,0,0,0,0],"isMain":false},{"metricName":"# scaffold gap loc. mis.","quality":"Less is better","values":[0,0,0,0,0,0],"isMain":false}]],["Unaligned",[]],["Mismatches",[]],["Statistics without reference",[]],["K-mer-based statistics",[]],["Predicted genes",[]],["Similarity statistics",[]],["Reference statistics",[]]],[["Genome statistics",[]],["Reads mapping",[]],["Misassemblies",[{"metricName":"# misassemblies","quality":"Less is better","values":[0,1,4,5,8,1],"isMain":true},{"metricName":" # relocations","quality":"Less is better","values":[0,0,1,1,1,0],"isMain":false},{"metricName":" # translocations","quality":"Less is better","values":[0,0,0,0,0,0],"isMain":false},{"metricName":" # inversions","quality":"Less is better","values":[0,0,0,0,0,0],"isMain":false},{"metricName":" # interspecies translocations","quality":"Less is better","values":[0,1,3,4,7,1],"isMain":false},{"metricName":"# possibly misassembled contigs","quality":"Less is better","values":[0,0,0,0,0,0],"isMain":false},{"metricName":" # possible misassemblies","quality":"Less is better","values":[0,0,0,0,0,0],"isMain":false},{"metricName":"# local misassemblies","quality":"Less is better","values":[1,0,0,1,0,0],"isMain":false},{"metricName":"# scaffold gap ext. mis.","quality":"Less is better","values":[0,0,0,0,0,0],"isMain":false},{"metricName":"# scaffold gap loc. mis.","quality":"Less is better","values":[0,0,0,0,0,0],"isMain":false}]],["Unaligned",[]],["Mismatches",[]],["Statistics without reference",[]],["K-mer-based statistics",[]],["Predicted genes",[]],["Similarity statistics",[]],["Reference statistics",[]]],[["Genome statistics",[]],["Reads mapping",[]],["Misassemblies",[{"metricName":"# misassemblies","quality":"Less is better","values":[22,48,20,34,33,21],"isMain":true},{"metricName":" # relocations","quality":"Less is better","values":[4,3,9,9,13,1],"isMain":false},{"metricName":" # translocations","quality":"Less is better","values":[0,0,0,0,0,0],"isMain":false},{"metricName":" # inversions","quality":"Less is better","values":[0,0,0,0,0,0],"isMain":false},{"metricName":" # interspecies translocations","quality":"Less is better","values":[18,45,11,25,20,20],"isMain":false},{"metricName":"# possibly misassembled contigs","quality":"Less is better","values":[0,0,0,0,0,0],"isMain":false},{"metricName":" # possible misassemblies","quality":"Less is better","values":[0,0,0,0,0,0],"isMain":false},{"metricName":"# local misassemblies","quality":"Less is better","values":[3,1,1,5,0,0],"isMain":false},{"metricName":"# scaffold gap ext. mis.","quality":"Less is better","values":[0,0,0,0,0,0],"isMain":false},{"metricName":"# scaffold gap loc. mis.","quality":"Less is better","values":[0,0,0,0,0,0],"isMain":false}]],["Unaligned",[]],["Mismatches",[]],["Statistics without reference",[]],["K-mer-based statistics",[]],["Predicted genes",[]],["Similarity statistics",[]],["Reference statistics",[]]],[["Genome statistics",[]],["Reads mapping",[]],["Misassemblies",[{"metricName":"# misassemblies","quality":"Less is better","values":[26,56,37,63,56,15],"isMain":true},{"metricName":" # relocations","quality":"Less is better","values":[3,4,6,8,1,3],"isMain":false},{"metricName":" # translocations","quality":"Less is better","values":[0,0,0,0,0,0],"isMain":false},{"metricName":" # inversions","quality":"Less is better","values":[1,1,4,1,3,0],"isMain":false},{"metricName":" # interspecies translocations","quality":"Less is better","values":[22,51,27,54,52,12],"isMain":false},{"metricName":"# possibly misassembled contigs","quality":"Less is better","values":[0,0,0,0,0,0],"isMain":false},{"metricName":" # possible misassemblies","quality":"Less is better","values":[0,0,0,0,0,0],"isMain":false},{"metricName":"# local misassemblies","quality":"Less is better","values":[0,3,2,6,1,0],"isMain":false},{"metricName":"# scaffold gap ext. mis.","quality":"Less is better","values":[0,0,0,0,0,0],"isMain":false},{"metricName":"# scaffold gap loc. mis.","quality":"Less is better","values":[0,0,0,0,0,0],"isMain":false}]],["Unaligned",[]],["Mismatches",[]],["Statistics without reference",[]],["K-mer-based statistics",[]],["Predicted genes",[]],["Similarity statistics",[]],["Reference statistics",[]]],[["Genome statistics",[]],["Reads mapping",[]],["Misassemblies",[{"metricName":"# misassemblies","quality":"Less is better","values":[5,14,2,14,10,3],"isMain":true},{"metricName":" # relocations","quality":"Less is better","values":[0,1,0,0,2,0],"isMain":false},{"metricName":" # translocations","quality":"Less is better","values":[0,0,0,0,0,0],"isMain":false},{"metricName":" # inversions","quality":"Less is better","values":[0,0,0,0,1,0],"isMain":false},{"metricName":" # interspecies translocations","quality":"Less is better","values":[5,13,2,14,7,3],"isMain":false},{"metricName":"# possibly misassembled contigs","quality":"Less is better","values":[0,0,0,0,0,0],"isMain":false},{"metricName":" # possible misassemblies","quality":"Less is better","values":[0,0,0,0,0,0],"isMain":false},{"metricName":"# local misassemblies","quality":"Less is better","values":[1,1,1,0,0,1],"isMain":false},{"metricName":"# scaffold gap ext. mis.","quality":"Less is better","values":[0,0,0,0,0,0],"isMain":false},{"metricName":"# scaffold gap loc. mis.","quality":"Less is better","values":[0,0,0,0,0,0],"isMain":false}]],["Unaligned",[]],["Mismatches",[]],["Statistics without reference",[]],["K-mer-based statistics",[]],["Predicted genes",[]],["Similarity statistics",[]],["Reference statistics",[]]],[["Genome statistics",[]],["Reads mapping",[]],["Misassemblies",[{"metricName":"# misassemblies","quality":"Less is better","values":[23,31,14,35,24,22],"isMain":true},{"metricName":" # relocations","quality":"Less is better","values":[2,2,3,6,6,2],"isMain":false},{"metricName":" # translocations","quality":"Less is better","values":[0,0,0,0,0,0],"isMain":false},{"metricName":" # inversions","quality":"Less is better","values":[0,0,0,0,0,0],"isMain":false},{"metricName":" # interspecies translocations","quality":"Less is better","values":[21,29,11,29,18,20],"isMain":false},{"metricName":"# possibly misassembled contigs","quality":"Less is better","values":[0,0,0,0,0,0],"isMain":false},{"metricName":" # possible misassemblies","quality":"Less is better","values":[0,0,0,0,0,0],"isMain":false},{"metricName":"# local misassemblies","quality":"Less is better","values":[0,0,1,1,1,0],"isMain":false},{"metricName":"# scaffold gap ext. mis.","quality":"Less is better","values":[0,0,0,0,0,0],"isMain":false},{"metricName":"# scaffold gap loc. mis.","quality":"Less is better","values":[0,0,0,0,0,0],"isMain":false}]],["Unaligned",[]],["Mismatches",[]],["Statistics without reference",[]],["K-mer-based statistics",[]],["Predicted genes",[]],["Similarity statistics",[]],["Reference statistics",[]]],[["Genome statistics",[]],["Reads mapping",[]],["Misassemblies",[{"metricName":"# misassemblies","quality":"Less is better","values":[86,108,17,90,87,71],"isMain":true},{"metricName":" # relocations","quality":"Less is better","values":[4,12,3,3,7,2],"isMain":false},{"metricName":" # translocations","quality":"Less is better","values":[0,0,0,0,0,0],"isMain":false},{"metricName":" # inversions","quality":"Less is better","values":[0,1,4,5,1,0],"isMain":false},{"metricName":" # interspecies translocations","quality":"Less is better","values":[82,95,10,82,79,69],"isMain":false},{"metricName":"# possibly misassembled contigs","quality":"Less is better","values":[0,0,0,0,0,0],"isMain":false},{"metricName":" # possible misassemblies","quality":"Less is better","values":[0,0,0,0,0,0],"isMain":false},{"metricName":"# local misassemblies","quality":"Less is better","values":[3,3,4,2,3,1],"isMain":false},{"metricName":"# scaffold gap ext. mis.","quality":"Less is better","values":[0,0,0,0,0,0],"isMain":false},{"metricName":"# scaffold gap loc. mis.","quality":"Less is better","values":[0,0,0,0,0,0],"isMain":false}]],["Unaligned",[]],["Mismatches",[]],["Statistics without reference",[]],["K-mer-based statistics",[]],["Predicted genes",[]],["Similarity statistics",[]],["Reference statistics",[]]],[["Genome statistics",[]],["Reads mapping",[]],["Misassemblies",[{"metricName":"# misassemblies","quality":"Less is better","values":[3,14,7,9,5,3],"isMain":true},{"metricName":" # relocations","quality":"Less is better","values":[0,0,0,0,0,0],"isMain":false},{"metricName":" # translocations","quality":"Less is better","values":[0,0,0,0,0,0],"isMain":false},{"metricName":" # inversions","quality":"Less is better","values":[0,1,1,0,0,0],"isMain":false},{"metricName":" # interspecies translocations","quality":"Less is better","values":[3,13,6,9,5,3],"isMain":false},{"metricName":"# possibly misassembled contigs","quality":"Less is better","values":[0,0,0,0,0,0],"isMain":false},{"metricName":" # possible misassemblies","quality":"Less is better","values":[0,0,0,0,0,0],"isMain":false},{"metricName":"# local misassemblies","quality":"Less is better","values":[1,0,2,3,0,0],"isMain":false},{"metricName":"# scaffold gap ext. mis.","quality":"Less is better","values":[0,0,0,0,0,0],"isMain":false},{"metricName":"# scaffold gap loc. mis.","quality":"Less is better","values":[0,0,0,0,0,0],"isMain":false}]],["Unaligned",[]],["Mismatches",[]],["Statistics without reference",[]],["K-mer-based statistics",[]],["Predicted genes",[]],["Similarity statistics",[]],["Reference statistics",[]]],[["Genome statistics",[]],["Reads mapping",[]],["Misassemblies",[{"metricName":"# misassemblies","quality":"Less is better","values":[53,74,26,60,63,49],"isMain":true},{"metricName":" # relocations","quality":"Less is better","values":[6,6,9,8,6,2],"isMain":false},{"metricName":" # translocations","quality":"Less is better","values":[0,0,0,0,0,0],"isMain":false},{"metricName":" # inversions","quality":"Less is better","values":[0,2,2,1,1,0],"isMain":false},{"metricName":" # interspecies translocations","quality":"Less is better","values":[47,66,15,51,56,47],"isMain":false},{"metricName":"# possibly misassembled contigs","quality":"Less is better","values":[0,0,0,0,0,0],"isMain":false},{"metricName":" # possible misassemblies","quality":"Less is better","values":[0,0,0,0,0,0],"isMain":false},{"metricName":"# local misassemblies","quality":"Less is better","values":[2,0,1,0,3,0],"isMain":false},{"metricName":"# scaffold gap ext. mis.","quality":"Less is better","values":[0,0,0,0,0,0],"isMain":false},{"metricName":"# scaffold gap loc. mis.","quality":"Less is better","values":[0,0,0,0,0,0],"isMain":false}]],["Unaligned",[]],["Mismatches",[]],["Statistics without reference",[]],["K-mer-based statistics",[]],["Predicted genes",[]],["Similarity statistics",[]],["Reference statistics",[]]],[["Genome statistics",[]],["Reads mapping",[]],["Misassemblies",[{"metricName":"# misassemblies","quality":"Less is better","values":[2,14,3,15,11,2],"isMain":true},{"metricName":" # relocations","quality":"Less is better","values":[0,0,1,3,1,0],"isMain":false},{"metricName":" # translocations","quality":"Less is better","values":[0,0,0,0,0,0],"isMain":false},{"metricName":" # inversions","quality":"Less is better","values":[0,0,0,0,0,0],"isMain":false},{"metricName":" # interspecies translocations","quality":"Less is better","values":[2,14,2,12,10,2],"isMain":false},{"metricName":"# possibly misassembled contigs","quality":"Less is better","values":[0,0,0,0,0,0],"isMain":false},{"metricName":" # possible misassemblies","quality":"Less is better","values":[0,0,0,0,0,0],"isMain":false},{"metricName":"# local misassemblies","quality":"Less is better","values":[0,1,0,0,2,0],"isMain":false},{"metricName":"# scaffold gap ext. mis.","quality":"Less is better","values":[0,0,0,0,0,0],"isMain":false},{"metricName":"# scaffold gap loc. mis.","quality":"Less is better","values":[0,0,0,0,0,0],"isMain":false}]],["Unaligned",[]],["Mismatches",[]],["Statistics without reference",[]],["K-mer-based statistics",[]],["Predicted genes",[]],["Similarity statistics",[]],["Reference statistics",[]]],[["Genome statistics",[]],["Reads mapping",[]],["Misassemblies",[{"metricName":"# misassemblies","quality":"Less is better","values":[1,1,4,2,1,0],"isMain":true},{"metricName":" # relocations","quality":"Less is better","values":[0,0,0,0,0,0],"isMain":false},{"metricName":" # translocations","quality":"Less is better","values":[0,0,0,0,0,0],"isMain":false},{"metricName":" # inversions","quality":"Less is better","values":[0,0,0,0,0,0],"isMain":false},{"metricName":" # interspecies translocations","quality":"Less is better","values":[1,1,4,2,1,0],"isMain":false},{"metricName":"# possibly misassembled contigs","quality":"Less is better","values":[0,0,0,0,0,0],"isMain":false},{"metricName":" # possible misassemblies","quality":"Less is better","values":[0,0,0,0,0,0],"isMain":false},{"metricName":"# local misassemblies","quality":"Less is better","values":[0,0,0,0,0,0],"isMain":false},{"metricName":"# scaffold gap ext. mis.","quality":"Less is better","values":[0,0,0,0,0,0],"isMain":false},{"metricName":"# scaffold gap loc. mis.","quality":"Less is better","values":[0,0,0,0,0,0],"isMain":false}]],["Unaligned",[]],["Mismatches",[]],["Statistics without reference",[]],["K-mer-based statistics",[]],["Predicted genes",[]],["Similarity statistics",[]],["Reference statistics",[]]],[["Genome statistics",[]],["Reads mapping",[]],["Misassemblies",[{"metricName":"# misassemblies","quality":"Less is better","values":[3,5,5,6,3,2],"isMain":true},{"metricName":" # relocations","quality":"Less is better","values":[0,0,0,0,0,0],"isMain":false},{"metricName":" # translocations","quality":"Less is better","values":[0,0,0,0,0,0],"isMain":false},{"metricName":" # inversions","quality":"Less is better","values":[0,0,0,0,0,0],"isMain":false},{"metricName":" # interspecies translocations","quality":"Less is better","values":[3,5,5,6,3,2],"isMain":false},{"metricName":"# possibly misassembled contigs","quality":"Less is better","values":[0,0,0,0,0,0],"isMain":false},{"metricName":" # possible misassemblies","quality":"Less is better","values":[0,0,0,0,0,0],"isMain":false},{"metricName":"# local misassemblies","quality":"Less is better","values":[0,0,0,0,0,0],"isMain":false},{"metricName":"# scaffold gap ext. mis.","quality":"Less is better","values":[0,0,0,0,0,0],"isMain":false},{"metricName":"# scaffold gap loc. mis.","quality":"Less is better","values":[0,0,0,0,0,0],"isMain":false}]],["Unaligned",[]],["Mismatches",[]],["Statistics without reference",[]],["K-mer-based statistics",[]],["Predicted genes",[]],["Similarity statistics",[]],["Reference statistics",[]]],[["Genome statistics",[]],["Reads mapping",[]],["Misassemblies",[{"metricName":"# misassemblies","quality":"Less is better","values":[26,63,68,88,69,25],"isMain":true},{"metricName":" # relocations","quality":"Less is better","values":[0,1,0,1,0,0],"isMain":false},{"metricName":" # translocations","quality":"Less is better","values":[16,24,21,27,27,11],"isMain":false},{"metricName":" # inversions","quality":"Less is better","values":[0,0,3,2,3,0],"isMain":false},{"metricName":" # interspecies translocations","quality":"Less is better","values":[10,38,44,58,39,14],"isMain":false},{"metricName":"# possibly misassembled contigs","quality":"Less is better","values":[0,2,0,0,0,0],"isMain":false},{"metricName":" # possible misassemblies","quality":"Less is better","values":[0,2,0,0,0,0],"isMain":false},{"metricName":"# local misassemblies","quality":"Less is better","values":[0,2,0,2,4,0],"isMain":false},{"metricName":"# scaffold gap ext. mis.","quality":"Less is better","values":[0,0,0,0,0,0],"isMain":false},{"metricName":"# scaffold gap loc. mis.","quality":"Less is better","values":[0,0,0,0,0,0],"isMain":false}]],["Unaligned",[]],["Mismatches",[]],["Statistics without reference",[]],["K-mer-based statistics",[]],["Predicted genes",[]],["Similarity statistics",[]],["Reference statistics",[]]],[["Genome statistics",[]],["Reads mapping",[]],["Misassemblies",[{"metricName":"# misassemblies","quality":"Less is better","values":[12,55,29,56,51,14],"isMain":true},{"metricName":" # relocations","quality":"Less is better","values":[1,1,0,2,2,0],"isMain":false},{"metricName":" # translocations","quality":"Less is better","values":[0,0,0,0,1,1],"isMain":false},{"metricName":" # inversions","quality":"Less is better","values":[1,0,0,0,0,0],"isMain":false},{"metricName":" # interspecies translocations","quality":"Less is better","values":[10,54,29,54,48,13],"isMain":false},{"metricName":"# possibly misassembled contigs","quality":"Less is better","values":[1,7,0,4,2,0],"isMain":false},{"metricName":" # possible misassemblies","quality":"Less is better","values":[1,7,0,4,2,0],"isMain":false},{"metricName":"# local misassemblies","quality":"Less is better","values":[3,1,3,6,7,0],"isMain":false},{"metricName":"# scaffold gap ext. mis.","quality":"Less is better","values":[0,0,0,0,0,0],"isMain":false},{"metricName":"# scaffold gap loc. mis.","quality":"Less is better","values":[0,0,0,0,0,0],"isMain":false}]],["Unaligned",[]],["Mismatches",[]],["Statistics without reference",[]],["K-mer-based statistics",[]],["Predicted genes",[]],["Similarity statistics",[]],["Reference statistics",[]]],[["Genome statistics",[]],["Reads mapping",[]],["Misassemblies",[{"metricName":"# misassemblies","quality":"Less is better","values":[0,0,0,0,2,0],"isMain":true},{"metricName":" # relocations","quality":"Less is better","values":[0,0,0,0,0,0],"isMain":false},{"metricName":" # translocations","quality":"Less is better","values":[0,0,0,0,0,0],"isMain":false},{"metricName":" # inversions","quality":"Less is better","values":[0,0,0,0,0,0],"isMain":false},{"metricName":" # interspecies translocations","quality":"Less is better","values":[0,0,0,0,2,0],"isMain":false},{"metricName":"# possibly misassembled contigs","quality":"Less is better","values":[0,0,0,0,0,0],"isMain":false},{"metricName":" # possible misassemblies","quality":"Less is better","values":[0,0,0,0,0,0],"isMain":false},{"metricName":"# local misassemblies","quality":"Less is better","values":[0,0,0,0,0,0],"isMain":false},{"metricName":"# scaffold gap ext. mis.","quality":"Less is better","values":[0,0,0,0,0,0],"isMain":false},{"metricName":"# scaffold gap loc. mis.","quality":"Less is better","values":[0,0,0,0,0,0],"isMain":false}]],["Unaligned",[]],["Mismatches",[]],["Statistics without reference",[]],["K-mer-based statistics",[]],["Predicted genes",[]],["Similarity statistics",[]],["Reference statistics",[]]],[["Genome statistics",[]],["Reads mapping",[]],["Misassemblies",[{"metricName":"# misassemblies","quality":"Less is better","values":[71,63,18,61,78,50],"isMain":true},{"metricName":" # relocations","quality":"Less is better","values":[5,5,5,7,6,4],"isMain":false},{"metricName":" # translocations","quality":"Less is better","values":[0,1,0,1,2,1],"isMain":false},{"metricName":" # inversions","quality":"Less is better","values":[1,2,1,1,3,0],"isMain":false},{"metricName":" # interspecies translocations","quality":"Less is better","values":[65,55,12,52,67,45],"isMain":false},{"metricName":"# possibly misassembled contigs","quality":"Less is better","values":[0,0,0,0,0,0],"isMain":false},{"metricName":" # possible misassemblies","quality":"Less is better","values":[0,0,0,0,0,0],"isMain":false},{"metricName":"# local misassemblies","quality":"Less is better","values":[2,2,0,1,2,1],"isMain":false},{"metricName":"# scaffold gap ext. mis.","quality":"Less is better","values":[0,0,0,0,0,0],"isMain":false},{"metricName":"# scaffold gap loc. mis.","quality":"Less is better","values":[0,0,0,0,0,0],"isMain":false}]],["Unaligned",[]],["Mismatches",[]],["Statistics without reference",[]],["K-mer-based statistics",[]],["Predicted genes",[]],["Similarity statistics",[]],["Reference statistics",[]]],[["Genome statistics",[]],["Reads mapping",[]],["Misassemblies",[{"metricName":"# misassemblies","quality":"Less is better","values":[42,64,29,78,69,29],"isMain":true},{"metricName":" # relocations","quality":"Less is better","values":[2,4,3,1,0,2],"isMain":false},{"metricName":" # translocations","quality":"Less is better","values":[2,3,3,2,3,3],"isMain":false},{"metricName":" # inversions","quality":"Less is better","values":[1,1,2,3,0,0],"isMain":false},{"metricName":" # interspecies translocations","quality":"Less is better","values":[37,56,21,72,66,24],"isMain":false},{"metricName":"# possibly misassembled contigs","quality":"Less is better","values":[0,1,0,0,2,0],"isMain":false},{"metricName":" # possible misassemblies","quality":"Less is better","values":[0,1,0,0,2,0],"isMain":false},{"metricName":"# local misassemblies","quality":"Less is better","values":[4,5,5,3,3,4],"isMain":false},{"metricName":"# scaffold gap ext. mis.","quality":"Less is better","values":[0,0,0,0,0,0],"isMain":false},{"metricName":"# scaffold gap loc. mis.","quality":"Less is better","values":[0,0,0,0,0,0],"isMain":false}]],["Unaligned",[]],["Mismatches",[]],["Statistics without reference",[]],["K-mer-based statistics",[]],["Predicted genes",[]],["Similarity statistics",[]],["Reference statistics",[]]],[["Genome statistics",[]],["Reads mapping",[]],["Misassemblies",[{"metricName":"# misassemblies","quality":"Less is better","values":[20,22,11,15,16,12],"isMain":true},{"metricName":" # relocations","quality":"Less is better","values":[0,1,0,0,0,0],"isMain":false},{"metricName":" # translocations","quality":"Less is better","values":[4,1,4,1,1,1],"isMain":false},{"metricName":" # inversions","quality":"Less is better","values":[0,0,1,0,0,0],"isMain":false},{"metricName":" # interspecies translocations","quality":"Less is better","values":[16,20,6,14,15,11],"isMain":false},{"metricName":"# possibly misassembled contigs","quality":"Less is better","values":[0,0,1,0,0,0],"isMain":false},{"metricName":" # possible misassemblies","quality":"Less is better","values":[0,0,1,0,0,0],"isMain":false},{"metricName":"# local misassemblies","quality":"Less is better","values":[1,3,2,1,2,0],"isMain":false},{"metricName":"# scaffold gap ext. mis.","quality":"Less is better","values":[0,0,0,0,0,0],"isMain":false},{"metricName":"# scaffold gap loc. mis.","quality":"Less is better","values":[0,0,0,0,0,0],"isMain":false}]],["Unaligned",[]],["Mismatches",[]],["Statistics without reference",[]],["K-mer-based statistics",[]],["Predicted genes",[]],["Similarity statistics",[]],["Reference statistics",[]]],[["Genome statistics",[]],["Reads mapping",[]],["Misassemblies",[{"metricName":"# misassemblies","quality":"Less is better","values":[41,40,18,49,39,28],"isMain":true},{"metricName":" # relocations","quality":"Less is better","values":[1,0,1,0,0,1],"isMain":false},{"metricName":" # translocations","quality":"Less is better","values":[1,3,2,1,2,0],"isMain":false},{"metricName":" # inversions","quality":"Less is better","values":[0,1,0,0,1,0],"isMain":false},{"metricName":" # interspecies translocations","quality":"Less is better","values":[39,36,15,48,36,27],"isMain":false},{"metricName":"# possibly misassembled contigs","quality":"Less is better","values":[0,0,0,0,0,0],"isMain":false},{"metricName":" # possible misassemblies","quality":"Less is better","values":[0,0,0,0,0,0],"isMain":false},{"metricName":"# local misassemblies","quality":"Less is better","values":[1,1,1,0,1,0],"isMain":false},{"metricName":"# scaffold gap ext. mis.","quality":"Less is better","values":[0,0,0,0,0,0],"isMain":false},{"metricName":"# scaffold gap loc. mis.","quality":"Less is better","values":[0,0,0,0,0,0],"isMain":false}]],["Unaligned",[]],["Mismatches",[]],["Statistics without reference",[]],["K-mer-based statistics",[]],["Predicted genes",[]],["Similarity statistics",[]],["Reference statistics",[]]],[["Genome statistics",[]],["Reads mapping",[]],["Misassemblies",[{"metricName":"# misassemblies","quality":"Less is better","values":[29,46,17,33,29,17],"isMain":true},{"metricName":" # relocations","quality":"Less is better","values":[0,0,0,0,1,0],"isMain":false},{"metricName":" # translocations","quality":"Less is better","values":[2,3,0,0,2,1],"isMain":false},{"metricName":" # inversions","quality":"Less is better","values":[2,0,0,0,0,0],"isMain":false},{"metricName":" # interspecies translocations","quality":"Less is better","values":[25,43,17,33,26,16],"isMain":false},{"metricName":"# possibly misassembled contigs","quality":"Less is better","values":[1,0,1,0,0,0],"isMain":false},{"metricName":" # possible misassemblies","quality":"Less is better","values":[1,0,1,0,0,0],"isMain":false},{"metricName":"# local misassemblies","quality":"Less is better","values":[1,1,0,3,0,0],"isMain":false},{"metricName":"# scaffold gap ext. mis.","quality":"Less is better","values":[0,0,0,0,0,0],"isMain":false},{"metricName":"# scaffold gap loc. mis.","quality":"Less is better","values":[0,0,0,0,0,0],"isMain":false}]],["Unaligned",[]],["Mismatches",[]],["Statistics without reference",[]],["K-mer-based statistics",[]],["Predicted genes",[]],["Similarity statistics",[]],["Reference statistics",[]]],[["Genome statistics",[]],["Reads mapping",[]],["Misassemblies",[{"metricName":"# misassemblies","quality":"Less is better","values":[5,16,10,14,27,1],"isMain":true},{"metricName":" # relocations","quality":"Less is better","values":[0,0,0,0,0,0],"isMain":false},{"metricName":" # translocations","quality":"Less is better","values":[0,0,0,0,2,0],"isMain":false},{"metricName":" # inversions","quality":"Less is better","values":[0,0,0,0,0,0],"isMain":false},{"metricName":" # interspecies translocations","quality":"Less is better","values":[5,16,10,14,25,1],"isMain":false},{"metricName":"# possibly misassembled contigs","quality":"Less is better","values":[0,0,0,1,1,0],"isMain":false},{"metricName":" # possible misassemblies","quality":"Less is better","values":[0,0,0,1,1,0],"isMain":false},{"metricName":"# local misassemblies","quality":"Less is better","values":[0,2,1,0,1,0],"isMain":false},{"metricName":"# scaffold gap ext. mis.","quality":"Less is better","values":[0,0,0,0,0,0],"isMain":false},{"metricName":"# scaffold gap loc. mis.","quality":"Less is better","values":[0,0,0,0,0,0],"isMain":false}]],["Unaligned",[]],["Mismatches",[]],["Statistics without reference",[]],["K-mer-based statistics",[]],["Predicted genes",[]],["Similarity statistics",[]],["Reference statistics",[]]],[["Genome statistics",[]],["Reads mapping",[]],["Misassemblies",[{"metricName":"# misassemblies","quality":"Less is better","values":[14,18,5,17,17,8],"isMain":true},{"metricName":" # relocations","quality":"Less is better","values":[0,0,0,1,0,0],"isMain":false},{"metricName":" # translocations","quality":"Less is better","values":[0,0,1,0,0,0],"isMain":false},{"metricName":" # inversions","quality":"Less is better","values":[0,0,0,0,0,0],"isMain":false},{"metricName":" # interspecies translocations","quality":"Less is better","values":[14,18,4,16,17,8],"isMain":false},{"metricName":"# possibly misassembled contigs","quality":"Less is better","values":[0,0,0,0,0,0],"isMain":false},{"metricName":" # possible misassemblies","quality":"Less is better","values":[0,0,0,0,0,0],"isMain":false},{"metricName":"# local misassemblies","quality":"Less is better","values":[0,0,0,0,0,0],"isMain":false},{"metricName":"# scaffold gap ext. mis.","quality":"Less is better","values":[0,0,0,0,0,0],"isMain":false},{"metricName":"# scaffold gap loc. mis.","quality":"Less is better","values":[0,0,0,0,0,0],"isMain":false}]],["Unaligned",[]],["Mismatches",[]],["Statistics without reference",[]],["K-mer-based statistics",[]],["Predicted genes",[]],["Similarity statistics",[]],["Reference statistics",[]]],[["Genome statistics",[]],["Reads mapping",[]],["Misassemblies",[{"metricName":"# misassemblies","quality":"Less is better","values":[15,45,31,56,84,3],"isMain":true},{"metricName":" # relocations","quality":"Less is better","values":[0,0,1,3,3,0],"isMain":false},{"metricName":" # translocations","quality":"Less is better","values":[2,1,2,5,4,1],"isMain":false},{"metricName":" # inversions","quality":"Less is better","values":[0,1,0,2,2,0],"isMain":false},{"metricName":" # interspecies translocations","quality":"Less is better","values":[13,43,28,46,75,2],"isMain":false},{"metricName":"# possibly misassembled contigs","quality":"Less is better","values":[1,1,1,1,2,0],"isMain":false},{"metricName":" # possible misassemblies","quality":"Less is better","values":[1,1,1,1,2,0],"isMain":false},{"metricName":"# local misassemblies","quality":"Less is better","values":[2,4,1,3,4,0],"isMain":false},{"metricName":"# scaffold gap ext. mis.","quality":"Less is better","values":[0,0,0,0,0,0],"isMain":false},{"metricName":"# scaffold gap loc. mis.","quality":"Less is better","values":[0,0,0,0,0,0],"isMain":false}]],["Unaligned",[]],["Mismatches",[]],["Statistics without reference",[]],["K-mer-based statistics",[]],["Predicted genes",[]],["Similarity statistics",[]],["Reference statistics",[]]],[["Genome statistics",[]],["Reads mapping",[]],["Misassemblies",[{"metricName":"# misassemblies","quality":"Less is better","values":[1,0,6,1,1,0],"isMain":true},{"metricName":" # relocations","quality":"Less is better","values":[0,0,0,0,0,0],"isMain":false},{"metricName":" # translocations","quality":"Less is better","values":[0,0,0,0,0,0],"isMain":false},{"metricName":" # inversions","quality":"Less is better","values":[0,0,0,0,0,0],"isMain":false},{"metricName":" # interspecies translocations","quality":"Less is better","values":[1,0,6,1,1,0],"isMain":false},{"metricName":"# possibly misassembled contigs","quality":"Less is better","values":[0,0,0,0,0,0],"isMain":false},{"metricName":" # possible misassemblies","quality":"Less is better","values":[0,0,0,0,0,0],"isMain":false},{"metricName":"# local misassemblies","quality":"Less is better","values":[0,0,0,0,0,0],"isMain":false},{"metricName":"# scaffold gap ext. mis.","quality":"Less is better","values":[0,0,0,0,0,0],"isMain":false},{"metricName":"# scaffold gap loc. mis.","quality":"Less is better","values":[0,0,0,0,0,0],"isMain":false}]],["Unaligned",[]],["Mismatches",[]],["Statistics without reference",[]],["K-mer-based statistics",[]],["Predicted genes",[]],["Similarity statistics",[]],["Reference statistics",[]]],[["Genome statistics",[]],["Reads mapping",[]],["Misassemblies",[{"metricName":"# misassemblies","quality":"Less is better","values":[4,6,7,12,4,0],"isMain":true},{"metricName":" # relocations","quality":"Less is better","values":[0,0,0,0,1,0],"isMain":false},{"metricName":" # translocations","quality":"Less is better","values":[0,0,0,0,0,0],"isMain":false},{"metricName":" # inversions","quality":"Less is better","values":[0,0,0,0,0,0],"isMain":false},{"metricName":" # interspecies translocations","quality":"Less is better","values":[4,6,7,12,3,0],"isMain":false},{"metricName":"# possibly misassembled contigs","quality":"Less is better","values":[0,0,0,0,0,0],"isMain":false},{"metricName":" # possible misassemblies","quality":"Less is better","values":[0,0,0,0,0,0],"isMain":false},{"metricName":"# local misassemblies","quality":"Less is better","values":[0,1,0,1,1,0],"isMain":false},{"metricName":"# scaffold gap ext. mis.","quality":"Less is better","values":[0,0,0,0,0,0],"isMain":false},{"metricName":"# scaffold gap loc. mis.","quality":"Less is better","values":[0,0,0,0,0,0],"isMain":false}]],["Unaligned",[]],["Mismatches",[]],["Statistics without reference",[]],["K-mer-based statistics",[]],["Predicted genes",[]],["Similarity statistics",[]],["Reference statistics",[]]],[["Genome statistics",[]],["Reads mapping",[]],["Misassemblies",[{"metricName":"# misassemblies","quality":"Less is better","values":[19,18,5,10,12,8],"isMain":true},{"metricName":" # relocations","quality":"Less is better","values":[0,0,0,0,0,0],"isMain":false},{"metricName":" # translocations","quality":"Less is better","values":[1,1,0,0,0,0],"isMain":false},{"metricName":" # inversions","quality":"Less is better","values":[0,0,0,0,0,0],"isMain":false},{"metricName":" # interspecies translocations","quality":"Less is better","values":[18,17,5,10,12,8],"isMain":false},{"metricName":"# possibly misassembled contigs","quality":"Less is better","values":[0,0,0,0,0,0],"isMain":false},{"metricName":" # possible misassemblies","quality":"Less is better","values":[0,0,0,0,0,0],"isMain":false},{"metricName":"# local misassemblies","quality":"Less is better","values":[0,0,0,1,1,0],"isMain":false},{"metricName":"# scaffold gap ext. mis.","quality":"Less is better","values":[0,0,0,0,0,0],"isMain":false},{"metricName":"# scaffold gap loc. mis.","quality":"Less is better","values":[0,0,0,0,0,0],"isMain":false}]],["Unaligned",[]],["Mismatches",[]],["Statistics without reference",[]],["K-mer-based statistics",[]],["Predicted genes",[]],["Similarity statistics",[]],["Reference statistics",[]]],[["Genome statistics",[]],["Reads mapping",[]],["Misassemblies",[{"metricName":"# misassemblies","quality":"Less is better","values":[32,42,18,40,45,21],"isMain":true},{"metricName":" # relocations","quality":"Less is better","values":[0,0,0,0,0,0],"isMain":false},{"metricName":" # translocations","quality":"Less is better","values":[1,2,0,1,0,0],"isMain":false},{"metricName":" # inversions","quality":"Less is better","values":[0,0,0,0,0,0],"isMain":false},{"metricName":" # interspecies translocations","quality":"Less is better","values":[31,40,18,39,45,21],"isMain":false},{"metricName":"# possibly misassembled contigs","quality":"Less is better","values":[0,0,0,1,0,0],"isMain":false},{"metricName":" # possible misassemblies","quality":"Less is better","values":[0,0,0,1,0,0],"isMain":false},{"metricName":"# local misassemblies","quality":"Less is better","values":[0,0,0,0,0,0],"isMain":false},{"metricName":"# scaffold gap ext. mis.","quality":"Less is better","values":[0,0,0,0,0,0],"isMain":false},{"metricName":"# scaffold gap loc. mis.","quality":"Less is better","values":[0,0,0,0,0,0],"isMain":false}]],["Unaligned",[]],["Mismatches",[]],["Statistics without reference",[]],["K-mer-based statistics",[]],["Predicted genes",[]],["Similarity statistics",[]],["Reference statistics",[]]],[["Genome statistics",[]],["Reads mapping",[]],["Misassemblies",[{"metricName":"# misassemblies","quality":"Less is better","values":[8,12,7,14,10,3],"isMain":true},{"metricName":" # relocations","quality":"Less is better","values":[0,0,0,0,1,0],"isMain":false},{"metricName":" # translocations","quality":"Less is better","values":[1,1,0,1,0,0],"isMain":false},{"metricName":" # inversions","quality":"Less is better","values":[1,0,0,0,0,0],"isMain":false},{"metricName":" # interspecies translocations","quality":"Less is better","values":[6,11,7,13,9,3],"isMain":false},{"metricName":"# possibly misassembled contigs","quality":"Less is better","values":[0,0,0,0,0,0],"isMain":false},{"metricName":" # possible misassemblies","quality":"Less is better","values":[0,0,0,0,0,0],"isMain":false},{"metricName":"# local misassemblies","quality":"Less is better","values":[0,0,2,1,0,0],"isMain":false},{"metricName":"# scaffold gap ext. mis.","quality":"Less is better","values":[0,0,0,0,0,0],"isMain":false},{"metricName":"# scaffold gap loc. mis.","quality":"Less is better","values":[0,0,0,0,0,0],"isMain":false}]],["Unaligned",[]],["Mismatches",[]],["Statistics without reference",[]],["K-mer-based statistics",[]],["Predicted genes",[]],["Similarity statistics",[]],["Reference statistics",[]]],[["Genome statistics",[]],["Reads mapping",[]],["Misassemblies",[{"metricName":"# misassemblies","quality":"Less is better","values":[196,181,187,222,190,149],"isMain":true},{"metricName":" # relocations","quality":"Less is better","values":[0,0,1,1,0,6],"isMain":false},{"metricName":" # translocations","quality":"Less is better","values":[93,142,141,147,149,134],"isMain":false},{"metricName":" # inversions","quality":"Less is better","values":[1,0,1,0,1,0],"isMain":false},{"metricName":" # interspecies translocations","quality":"Less is better","values":[102,39,44,74,40,9],"isMain":false},{"metricName":"# possibly misassembled contigs","quality":"Less is better","values":[0,0,1,1,0,0],"isMain":false},{"metricName":" # possible misassemblies","quality":"Less is better","values":[0,0,1,1,0,0],"isMain":false},{"metricName":"# local misassemblies","quality":"Less is better","values":[1,3,2,1,0,2],"isMain":false},{"metricName":"# scaffold gap ext. mis.","quality":"Less is better","values":[0,0,0,0,0,0],"isMain":false},{"metricName":"# scaffold gap loc. mis.","quality":"Less is better","values":[0,0,0,0,0,0],"isMain":false}]],["Unaligned",[]],["Mismatches",[]],["Statistics without reference",[]],["K-mer-based statistics",[]],["Predicted genes",[]],["Similarity statistics",[]],["Reference statistics",[]]],[["Genome statistics",[]],["Reads mapping",[]],["Misassemblies",[{"metricName":"# misassemblies","quality":"Less is better","values":[5,9,2,8,13,3],"isMain":true},{"metricName":" # relocations","quality":"Less is better","values":[0,0,0,0,0,0],"isMain":false},{"metricName":" # translocations","quality":"Less is better","values":[0,0,0,0,0,0],"isMain":false},{"metricName":" # inversions","quality":"Less is better","values":[0,0,0,0,1,0],"isMain":false},{"metricName":" # interspecies translocations","quality":"Less is better","values":[5,9,2,8,12,3],"isMain":false},{"metricName":"# possibly misassembled contigs","quality":"Less is better","values":[0,0,0,0,1,0],"isMain":false},{"metricName":" # possible misassemblies","quality":"Less is better","values":[0,0,0,0,2,0],"isMain":false},{"metricName":"# local misassemblies","quality":"Less is better","values":[0,1,0,1,2,0],"isMain":false},{"metricName":"# scaffold gap ext. mis.","quality":"Less is better","values":[0,0,0,0,0,0],"isMain":false},{"metricName":"# scaffold gap loc. mis.","quality":"Less is better","values":[0,0,0,0,0,0],"isMain":false}]],["Unaligned",[]],["Mismatches",[]],["Statistics without reference",[]],["K-mer-based statistics",[]],["Predicted genes",[]],["Similarity statistics",[]],["Reference statistics",[]]],[["Genome statistics",[]],["Reads mapping",[]],["Misassemblies",[{"metricName":"# misassemblies","quality":"Less is better","values":[35,35,7,27,26,18],"isMain":true},{"metricName":" # relocations","quality":"Less is better","values":[2,1,1,0,1,0],"isMain":false},{"metricName":" # translocations","quality":"Less is better","values":[3,2,1,3,3,2],"isMain":false},{"metricName":" # inversions","quality":"Less is better","values":[1,1,2,1,0,0],"isMain":false},{"metricName":" # interspecies translocations","quality":"Less is better","values":[29,31,3,23,22,16],"isMain":false},{"metricName":"# possibly misassembled contigs","quality":"Less is better","values":[0,0,0,1,0,0],"isMain":false},{"metricName":" # possible misassemblies","quality":"Less is better","values":[0,0,0,1,0,0],"isMain":false},{"metricName":"# local misassemblies","quality":"Less is better","values":[1,2,0,0,1,1],"isMain":false},{"metricName":"# scaffold gap ext. mis.","quality":"Less is better","values":[0,0,0,0,0,0],"isMain":false},{"metricName":"# scaffold gap loc. mis.","quality":"Less is better","values":[0,0,0,0,0,0],"isMain":false}]],["Unaligned",[]],["Mismatches",[]],["Statistics without reference",[]],["K-mer-based statistics",[]],["Predicted genes",[]],["Similarity statistics",[]],["Reference statistics",[]]],[["Genome statistics",[]],["Reads mapping",[]],["Misassemblies",[{"metricName":"# misassemblies","quality":"Less is better","values":[4,10,6,5,11,0],"isMain":true},{"metricName":" # relocations","quality":"Less is better","values":[0,0,0,0,0,0],"isMain":false},{"metricName":" # translocations","quality":"Less is better","values":[0,0,0,0,0,0],"isMain":false},{"metricName":" # inversions","quality":"Less is better","values":[0,0,0,0,0,0],"isMain":false},{"metricName":" # interspecies translocations","quality":"Less is better","values":[4,10,6,5,11,0],"isMain":false},{"metricName":"# possibly misassembled contigs","quality":"Less is better","values":[0,1,2,2,2,0],"isMain":false},{"metricName":" # possible misassemblies","quality":"Less is better","values":[0,1,2,2,2,0],"isMain":false},{"metricName":"# local misassemblies","quality":"Less is better","values":[0,0,0,0,0,0],"isMain":false},{"metricName":"# scaffold gap ext. mis.","quality":"Less is better","values":[0,0,0,0,0,0],"isMain":false},{"metricName":"# scaffold gap loc. mis.","quality":"Less is better","values":[0,0,0,0,0,0],"isMain":false}]],["Unaligned",[]],["Mismatches",[]],["Statistics without reference",[]],["K-mer-based statistics",[]],["Predicted genes",[]],["Similarity statistics",[]],["Reference statistics",[]]],[["Genome statistics",[]],["Reads mapping",[]],["Misassemblies",[{"metricName":"# misassemblies","quality":"Less is better","values":[11,9,14,8,18,6],"isMain":true},{"metricName":" # relocations","quality":"Less is better","values":[1,0,2,0,0,2],"isMain":false},{"metricName":" # translocations","quality":"Less is better","values":[2,5,5,4,4,2],"isMain":false},{"metricName":" # inversions","quality":"Less is better","values":[0,0,0,0,1,0],"isMain":false},{"metricName":" # interspecies translocations","quality":"Less is better","values":[8,4,7,4,13,2],"isMain":false},{"metricName":"# possibly misassembled contigs","quality":"Less is better","values":[0,0,0,0,0,0],"isMain":false},{"metricName":" # possible misassemblies","quality":"Less is better","values":[0,0,0,0,0,0],"isMain":false},{"metricName":"# local misassemblies","quality":"Less is better","values":[0,4,3,3,3,0],"isMain":false},{"metricName":"# scaffold gap ext. mis.","quality":"Less is better","values":[0,0,0,0,0,0],"isMain":false},{"metricName":"# scaffold gap loc. mis.","quality":"Less is better","values":[0,0,0,0,0,0],"isMain":false}]],["Unaligned",[]],["Mismatches",[]],["Statistics without reference",[]],["K-mer-based statistics",[]],["Predicted genes",[]],["Similarity statistics",[]],["Reference statistics",[]]],[["Genome statistics",[]],["Reads mapping",[]],["Misassemblies",[{"metricName":"# misassemblies","quality":"Less is better","values":[19,22,18,16,22,3],"isMain":true},{"metricName":" # relocations","quality":"Less is better","values":[0,1,0,0,0,0],"isMain":false},{"metricName":" # translocations","quality":"Less is better","values":[1,0,1,0,0,1],"isMain":false},{"metricName":" # inversions","quality":"Less is better","values":[0,0,0,0,0,0],"isMain":false},{"metricName":" # interspecies translocations","quality":"Less is better","values":[18,21,17,16,22,2],"isMain":false},{"metricName":"# possibly misassembled contigs","quality":"Less is better","values":[0,0,0,0,0,0],"isMain":false},{"metricName":" # possible misassemblies","quality":"Less is better","values":[0,0,0,0,0,0],"isMain":false},{"metricName":"# local misassemblies","quality":"Less is better","values":[0,1,1,0,2,0],"isMain":false},{"metricName":"# scaffold gap ext. mis.","quality":"Less is better","values":[0,0,0,0,0,0],"isMain":false},{"metricName":"# scaffold gap loc. mis.","quality":"Less is better","values":[0,0,0,0,0,0],"isMain":false}]],["Unaligned",[]],["Mismatches",[]],["Statistics without reference",[]],["K-mer-based statistics",[]],["Predicted genes",[]],["Similarity statistics",[]],["Reference statistics",[]]],[["Genome statistics",[]],["Reads mapping",[]],["Misassemblies",[{"metricName":"# misassemblies","quality":"Less is better","values":[16,19,2,19,19,10],"isMain":true},{"metricName":" # relocations","quality":"Less is better","values":[0,1,0,1,1,0],"isMain":false},{"metricName":" # translocations","quality":"Less is better","values":[0,0,1,0,1,0],"isMain":false},{"metricName":" # inversions","quality":"Less is better","values":[0,0,0,1,0,0],"isMain":false},{"metricName":" # interspecies translocations","quality":"Less is better","values":[16,18,1,17,17,10],"isMain":false},{"metricName":"# possibly misassembled contigs","quality":"Less is better","values":[0,0,0,0,0,0],"isMain":false},{"metricName":" # possible misassemblies","quality":"Less is better","values":[0,0,0,0,0,0],"isMain":false},{"metricName":"# local misassemblies","quality":"Less is better","values":[1,0,0,0,1,0],"isMain":false},{"metricName":"# scaffold gap ext. mis.","quality":"Less is better","values":[0,0,0,0,0,0],"isMain":false},{"metricName":"# scaffold gap loc. mis.","quality":"Less is better","values":[0,0,0,0,0,0],"isMain":false}]],["Unaligned",[]],["Mismatches",[]],["Statistics without reference",[]],["K-mer-based statistics",[]],["Predicted genes",[]],["Similarity statistics",[]],["Reference statistics",[]]],[["Genome statistics",[]],["Reads mapping",[]],["Misassemblies",[{"metricName":"# misassemblies","quality":"Less is better","values":[9,5,7,7,5,4],"isMain":true},{"metricName":" # relocations","quality":"Less is better","values":[0,0,1,1,0,0],"isMain":false},{"metricName":" # translocations","quality":"Less is better","values":[0,0,0,0,0,0],"isMain":false},{"metricName":" # inversions","quality":"Less is better","values":[0,1,0,0,0,0],"isMain":false},{"metricName":" # interspecies translocations","quality":"Less is better","values":[9,4,6,6,5,4],"isMain":false},{"metricName":"# possibly misassembled contigs","quality":"Less is better","values":[0,0,0,0,0,0],"isMain":false},{"metricName":" # possible misassemblies","quality":"Less is better","values":[0,0,0,0,0,0],"isMain":false},{"metricName":"# local misassemblies","quality":"Less is better","values":[0,0,0,0,0,0],"isMain":false},{"metricName":"# scaffold gap ext. mis.","quality":"Less is better","values":[0,0,0,0,0,0],"isMain":false},{"metricName":"# scaffold gap loc. mis.","quality":"Less is better","values":[0,0,0,0,0,0],"isMain":false}]],["Unaligned",[]],["Mismatches",[]],["Statistics without reference",[]],["K-mer-based statistics",[]],["Predicted genes",[]],["Similarity statistics",[]],["Reference statistics",[]]],[["Genome statistics",[]],["Reads mapping",[]],["Misassemblies",[{"metricName":"# misassemblies","quality":"Less is better","values":[1,3,2,0,1,0],"isMain":true},{"metricName":" # relocations","quality":"Less is better","values":[0,0,0,0,0,0],"isMain":false},{"metricName":" # translocations","quality":"Less is better","values":[0,0,0,0,0,0],"isMain":false},{"metricName":" # inversions","quality":"Less is better","values":[0,0,1,0,0,0],"isMain":false},{"metricName":" # interspecies translocations","quality":"Less is better","values":[1,3,1,0,1,0],"isMain":false},{"metricName":"# possibly misassembled contigs","quality":"Less is better","values":[0,0,0,0,0,0],"isMain":false},{"metricName":" # possible misassemblies","quality":"Less is better","values":[0,0,0,0,0,0],"isMain":false},{"metricName":"# local misassemblies","quality":"Less is better","values":[0,0,0,0,0,0],"isMain":false},{"metricName":"# scaffold gap ext. mis.","quality":"Less is better","values":[0,0,0,0,0,0],"isMain":false},{"metricName":"# scaffold gap loc. mis.","quality":"Less is better","values":[0,0,0,0,0,0],"isMain":false}]],["Unaligned",[]],["Mismatches",[]],["Statistics without reference",[]],["K-mer-based statistics",[]],["Predicted genes",[]],["Similarity statistics",[]],["Reference statistics",[]]],[["Genome statistics",[]],["Reads mapping",[]],["Misassemblies",[{"metricName":"# misassemblies","quality":"Less is better","values":[4,10,2,3,5,3],"isMain":true},{"metricName":" # relocations","quality":"Less is better","values":[0,0,0,0,0,0],"isMain":false},{"metricName":" # translocations","quality":"Less is better","values":[0,1,0,0,0,0],"isMain":false},{"metricName":" # inversions","quality":"Less is better","values":[0,0,0,0,0,0],"isMain":false},{"metricName":" # interspecies translocations","quality":"Less is better","values":[4,9,2,3,5,3],"isMain":false},{"metricName":"# possibly misassembled contigs","quality":"Less is better","values":[0,0,0,0,0,0],"isMain":false},{"metricName":" # possible misassemblies","quality":"Less is better","values":[0,0,0,0,0,0],"isMain":false},{"metricName":"# local misassemblies","quality":"Less is better","values":[1,1,0,0,1,0],"isMain":false},{"metricName":"# scaffold gap ext. mis.","quality":"Less is better","values":[0,0,0,0,0,0],"isMain":false},{"metricName":"# scaffold gap loc. mis.","quality":"Less is better","values":[0,0,0,0,0,0],"isMain":false}]],["Unaligned",[]],["Mismatches",[]],["Statistics without reference",[]],["K-mer-based statistics",[]],["Predicted genes",[]],["Similarity statistics",[]],["Reference statistics",[]]],[["Genome statistics",[]],["Reads mapping",[]],["Misassemblies",[{"metricName":"# misassemblies","quality":"Less is better","values":[82,96,44,96,104,54],"isMain":true},{"metricName":" # relocations","quality":"Less is better","values":[1,2,2,3,0,3],"isMain":false},{"metricName":" # translocations","quality":"Less is better","values":[14,13,8,11,15,7],"isMain":false},{"metricName":" # inversions","quality":"Less is better","values":[1,0,2,1,0,0],"isMain":false},{"metricName":" # interspecies translocations","quality":"Less is better","values":[66,81,32,81,89,44],"isMain":false},{"metricName":"# possibly misassembled contigs","quality":"Less is better","values":[0,0,0,1,0,0],"isMain":false},{"metricName":" # possible misassemblies","quality":"Less is better","values":[0,0,0,1,0,0],"isMain":false},{"metricName":"# local misassemblies","quality":"Less is better","values":[1,1,5,4,4,2],"isMain":false},{"metricName":"# scaffold gap ext. mis.","quality":"Less is better","values":[0,0,0,0,0,0],"isMain":false},{"metricName":"# scaffold gap loc. mis.","quality":"Less is better","values":[0,0,0,0,0,0],"isMain":false}]],["Unaligned",[]],["Mismatches",[]],["Statistics without reference",[]],["K-mer-based statistics",[]],["Predicted genes",[]],["Similarity statistics",[]],["Reference statistics",[]]],[["Genome statistics",[]],["Reads mapping",[]],["Misassemblies",[{"metricName":"# misassemblies","quality":"Less is better","values":[21,19,20,26,22,9],"isMain":true},{"metricName":" # relocations","quality":"Less is better","values":[0,0,0,0,0,0],"isMain":false},{"metricName":" # translocations","quality":"Less is better","values":[7,11,9,9,10,9],"isMain":false},{"metricName":" # inversions","quality":"Less is better","values":[0,0,0,1,0,0],"isMain":false},{"metricName":" # interspecies translocations","quality":"Less is better","values":[14,8,11,16,12,0],"isMain":false},{"metricName":"# possibly misassembled contigs","quality":"Less is better","values":[0,0,0,0,0,0],"isMain":false},{"metricName":" # possible misassemblies","quality":"Less is better","values":[0,0,0,0,0,0],"isMain":false},{"metricName":"# local misassemblies","quality":"Less is better","values":[0,0,1,0,0,0],"isMain":false},{"metricName":"# scaffold gap ext. mis.","quality":"Less is better","values":[0,0,0,0,0,0],"isMain":false},{"metricName":"# scaffold gap loc. mis.","quality":"Less is better","values":[0,0,0,0,0,0],"isMain":false}]],["Unaligned",[]],["Mismatches",[]],["Statistics without reference",[]],["K-mer-based statistics",[]],["Predicted genes",[]],["Similarity statistics",[]],["Reference statistics",[]]],[["Genome statistics",[]],["Reads mapping",[]],["Misassemblies",[{"metricName":"# misassemblies","quality":"Less is better","values":[60,40,22,59,41,22],"isMain":true},{"metricName":" # relocations","quality":"Less is better","values":[1,0,0,0,0,0],"isMain":false},{"metricName":" # translocations","quality":"Less is better","values":[11,15,8,17,18,16],"isMain":false},{"metricName":" # inversions","quality":"Less is better","values":[2,0,0,0,0,0],"isMain":false},{"metricName":" # interspecies translocations","quality":"Less is better","values":[46,25,14,42,23,6],"isMain":false},{"metricName":"# possibly misassembled contigs","quality":"Less is better","values":[0,0,1,0,0,0],"isMain":false},{"metricName":" # possible misassemblies","quality":"Less is better","values":[0,0,1,0,0,0],"isMain":false},{"metricName":"# local misassemblies","quality":"Less is better","values":[3,0,1,1,0,1],"isMain":false},{"metricName":"# scaffold gap ext. mis.","quality":"Less is better","values":[0,0,0,0,0,0],"isMain":false},{"metricName":"# scaffold gap loc. mis.","quality":"Less is better","values":[0,0,0,0,0,0],"isMain":false}]],["Unaligned",[]],["Mismatches",[]],["Statistics without reference",[]],["K-mer-based statistics",[]],["Predicted genes",[]],["Similarity statistics",[]],["Reference statistics",[]]],[["Genome statistics",[]],["Reads mapping",[]],["Misassemblies",[{"metricName":"# misassemblies","quality":"Less is better","values":[45,60,9,34,52,28],"isMain":true},{"metricName":" # relocations","quality":"Less is better","values":[0,0,0,0,0,0],"isMain":false},{"metricName":" # translocations","quality":"Less is better","values":[1,3,1,1,3,1],"isMain":false},{"metricName":" # inversions","quality":"Less is better","values":[0,0,0,0,0,0],"isMain":false},{"metricName":" # interspecies translocations","quality":"Less is better","values":[44,57,8,33,49,27],"isMain":false},{"metricName":"# possibly misassembled contigs","quality":"Less is better","values":[0,0,0,0,1,0],"isMain":false},{"metricName":" # possible misassemblies","quality":"Less is better","values":[0,0,0,0,1,0],"isMain":false},{"metricName":"# local misassemblies","quality":"Less is better","values":[0,1,0,1,0,0],"isMain":false},{"metricName":"# scaffold gap ext. mis.","quality":"Less is better","values":[0,0,0,0,0,0],"isMain":false},{"metricName":"# scaffold gap loc. mis.","quality":"Less is better","values":[0,0,0,0,0,0],"isMain":false}]],["Unaligned",[]],["Mismatches",[]],["Statistics without reference",[]],["K-mer-based statistics",[]],["Predicted genes",[]],["Similarity statistics",[]],["Reference statistics",[]]],[["Genome statistics",[]],["Reads mapping",[]],["Misassemblies",[{"metricName":"# misassemblies","quality":"Less is better","values":[43,64,24,37,45,40],"isMain":true},{"metricName":" # relocations","quality":"Less is better","values":[0,1,1,0,1,0],"isMain":false},{"metricName":" # translocations","quality":"Less is better","values":[6,10,3,3,12,12],"isMain":false},{"metricName":" # inversions","quality":"Less is better","values":[0,0,4,0,0,0],"isMain":false},{"metricName":" # interspecies translocations","quality":"Less is better","values":[37,53,16,34,32,28],"isMain":false},{"metricName":"# possibly misassembled contigs","quality":"Less is better","values":[0,0,0,1,0,0],"isMain":false},{"metricName":" # possible misassemblies","quality":"Less is better","values":[0,0,0,1,0,0],"isMain":false},{"metricName":"# local misassemblies","quality":"Less is better","values":[0,4,3,3,3,1],"isMain":false},{"metricName":"# scaffold gap ext. mis.","quality":"Less is better","values":[0,0,0,0,0,0],"isMain":false},{"metricName":"# scaffold gap loc. mis.","quality":"Less is better","values":[0,0,0,0,0,0],"isMain":false}]],["Unaligned",[]],["Mismatches",[]],["Statistics without reference",[]],["K-mer-based statistics",[]],["Predicted genes",[]],["Similarity statistics",[]],["Reference statistics",[]]],[["Genome statistics",[]],["Reads mapping",[]],["Misassemblies",[{"metricName":"# misassemblies","quality":"Less is better","values":[21,16,15,23,14,15],"isMain":true},{"metricName":" # relocations","quality":"Less is better","values":[0,0,0,0,0,1],"isMain":false},{"metricName":" # translocations","quality":"Less is better","values":[3,3,4,4,4,3],"isMain":false},{"metricName":" # inversions","quality":"Less is better","values":[0,0,1,0,0,0],"isMain":false},{"metricName":" # interspecies translocations","quality":"Less is better","values":[18,13,10,19,10,11],"isMain":false},{"metricName":"# possibly misassembled contigs","quality":"Less is better","values":[0,0,0,0,0,0],"isMain":false},{"metricName":" # possible misassemblies","quality":"Less is better","values":[0,0,0,0,0,0],"isMain":false},{"metricName":"# local misassemblies","quality":"Less is better","values":[1,1,6,4,3,1],"isMain":false},{"metricName":"# scaffold gap ext. mis.","quality":"Less is better","values":[0,0,0,0,0,0],"isMain":false},{"metricName":"# scaffold gap loc. mis.","quality":"Less is better","values":[0,0,0,0,0,0],"isMain":false}]],["Unaligned",[]],["Mismatches",[]],["Statistics without reference",[]],["K-mer-based statistics",[]],["Predicted genes",[]],["Similarity statistics",[]],["Reference statistics",[]]],[["Genome statistics",[]],["Reads mapping",[]],["Misassemblies",[{"metricName":"# misassemblies","quality":"Less is better","values":[19,21,14,15,22,13],"isMain":true},{"metricName":" # relocations","quality":"Less is better","values":[0,0,0,0,0,0],"isMain":false},{"metricName":" # translocations","quality":"Less is better","values":[0,1,2,1,2,0],"isMain":false},{"metricName":" # inversions","quality":"Less is better","values":[0,0,0,0,0,0],"isMain":false},{"metricName":" # interspecies translocations","quality":"Less is better","values":[19,20,12,14,20,13],"isMain":false},{"metricName":"# possibly misassembled contigs","quality":"Less is better","values":[0,0,0,0,1,0],"isMain":false},{"metricName":" # possible misassemblies","quality":"Less is better","values":[0,0,0,0,1,0],"isMain":false},{"metricName":"# local misassemblies","quality":"Less is better","values":[0,0,0,1,1,0],"isMain":false},{"metricName":"# scaffold gap ext. mis.","quality":"Less is better","values":[0,0,0,0,0,0],"isMain":false},{"metricName":"# scaffold gap loc. mis.","quality":"Less is better","values":[0,0,0,0,0,0],"isMain":false}]],["Unaligned",[]],["Mismatches",[]],["Statistics without reference",[]],["K-mer-based statistics",[]],["Predicted genes",[]],["Similarity statistics",[]],["Reference statistics",[]]],[["Genome statistics",[]],["Reads mapping",[]],["Misassemblies",[{"metricName":"# misassemblies","quality":"Less is better","values":[15,39,32,37,37,6],"isMain":true},{"metricName":" # relocations","quality":"Less is better","values":[0,0,0,1,0,0],"isMain":false},{"metricName":" # translocations","quality":"Less is better","values":[0,3,2,1,2,0],"isMain":false},{"metricName":" # inversions","quality":"Less is better","values":[0,0,0,0,1,0],"isMain":false},{"metricName":" # interspecies translocations","quality":"Less is better","values":[15,36,30,35,34,6],"isMain":false},{"metricName":"# possibly misassembled contigs","quality":"Less is better","values":[0,0,0,0,0,0],"isMain":false},{"metricName":" # possible misassemblies","quality":"Less is better","values":[0,0,0,0,0,0],"isMain":false},{"metricName":"# local misassemblies","quality":"Less is better","values":[0,6,0,0,3,0],"isMain":false},{"metricName":"# scaffold gap ext. mis.","quality":"Less is better","values":[0,0,0,0,0,0],"isMain":false},{"metricName":"# scaffold gap loc. mis.","quality":"Less is better","values":[0,0,0,0,0,0],"isMain":false}]],["Unaligned",[]],["Mismatches",[]],["Statistics without reference",[]],["K-mer-based statistics",[]],["Predicted genes",[]],["Similarity statistics",[]],["Reference statistics",[]]],[["Genome statistics",[]],["Reads mapping",[]],["Misassemblies",[{"metricName":"# misassemblies","quality":"Less is better","values":[2,8,8,9,3,8],"isMain":true},{"metricName":" # relocations","quality":"Less is better","values":[0,0,0,0,0,1],"isMain":false},{"metricName":" # translocations","quality":"Less is better","values":[0,0,0,0,0,0],"isMain":false},{"metricName":" # inversions","quality":"Less is better","values":[0,0,0,0,0,0],"isMain":false},{"metricName":" # interspecies translocations","quality":"Less is better","values":[2,8,8,9,3,7],"isMain":false},{"metricName":"# possibly misassembled contigs","quality":"Less is better","values":[0,0,0,0,0,0],"isMain":false},{"metricName":" # possible misassemblies","quality":"Less is better","values":[0,0,0,0,0,0],"isMain":false},{"metricName":"# local misassemblies","quality":"Less is better","values":[0,0,0,0,0,0],"isMain":false},{"metricName":"# scaffold gap ext. mis.","quality":"Less is better","values":[0,0,0,0,0,0],"isMain":false},{"metricName":"# scaffold gap loc. mis.","quality":"Less is better","values":[0,0,0,0,0,0],"isMain":false}]],["Unaligned",[]],["Mismatches",[]],["Statistics without reference",[]],["K-mer-based statistics",[]],["Predicted genes",[]],["Similarity statistics",[]],["Reference statistics",[]]],[["Genome statistics",[]],["Reads mapping",[]],["Misassemblies",[{"metricName":"# misassemblies","quality":"Less is better","values":[12,21,16,55,22,1],"isMain":true},{"metricName":" # relocations","quality":"Less is better","values":[2,5,3,2,3,1],"isMain":false},{"metricName":" # translocations","quality":"Less is better","values":[3,5,6,6,7,0],"isMain":false},{"metricName":" # inversions","quality":"Less is better","values":[1,1,0,5,1,0],"isMain":false},{"metricName":" # interspecies translocations","quality":"Less is better","values":[6,10,7,42,11,0],"isMain":false},{"metricName":"# possibly misassembled contigs","quality":"Less is better","values":[0,0,0,0,0,0],"isMain":false},{"metricName":" # possible misassemblies","quality":"Less is better","values":[0,0,0,0,0,0],"isMain":false},{"metricName":"# local misassemblies","quality":"Less is better","values":[0,4,0,1,3,0],"isMain":false},{"metricName":"# scaffold gap ext. mis.","quality":"Less is better","values":[0,0,0,0,0,0],"isMain":false},{"metricName":"# scaffold gap loc. mis.","quality":"Less is better","values":[0,0,0,0,0,0],"isMain":false}]],["Unaligned",[]],["Mismatches",[]],["Statistics without reference",[]],["K-mer-based statistics",[]],["Predicted genes",[]],["Similarity statistics",[]],["Reference statistics",[]]],[["Genome statistics",[]],["Reads mapping",[]],["Misassemblies",[{"metricName":"# misassemblies","quality":"Less is better","values":[154,660,437,602,621,235],"isMain":true},{"metricName":" # relocations","quality":"Less is better","values":[0,3,1,3,3,1],"isMain":false},{"metricName":" # translocations","quality":"Less is better","values":[2,280,258,306,313,189],"isMain":false},{"metricName":" # inversions","quality":"Less is better","values":[16,53,3,6,8,0],"isMain":false},{"metricName":" # interspecies translocations","quality":"Less is better","values":[136,324,175,287,297,45],"isMain":false},{"metricName":"# possibly misassembled contigs","quality":"Less is better","values":[0,20,11,23,14,5],"isMain":false},{"metricName":" # possible misassemblies","quality":"Less is better","values":[0,29,15,33,21,5],"isMain":false},{"metricName":"# local misassemblies","quality":"Less is better","values":[5,23,4,14,8,3],"isMain":false},{"metricName":"# scaffold gap ext. mis.","quality":"Less is better","values":[0,0,0,0,0,0],"isMain":false},{"metricName":"# scaffold gap loc. mis.","quality":"Less is better","values":[0,0,0,0,0,0],"isMain":false}]],["Unaligned",[]],["Mismatches",[]],["Statistics without reference",[]],["K-mer-based statistics",[]],["Predicted genes",[]],["Similarity statistics",[]],["Reference statistics",[]]],[["Genome statistics",[]],["Reads mapping",[]],["Misassemblies",[{"metricName":"# misassemblies","quality":"Less is better","values":[1,1,0,5,1,5],"isMain":true},{"metricName":" # relocations","quality":"Less is better","values":[0,0,0,0,0,0],"isMain":false},{"metricName":" # translocations","quality":"Less is better","values":[0,0,0,0,0,0],"isMain":false},{"metricName":" # inversions","quality":"Less is better","values":[0,0,0,0,0,0],"isMain":false},{"metricName":" # interspecies translocations","quality":"Less is better","values":[1,1,0,5,1,5],"isMain":false},{"metricName":"# possibly misassembled contigs","quality":"Less is better","values":[0,0,0,0,0,0],"isMain":false},{"metricName":" # possible misassemblies","quality":"Less is better","values":[0,0,0,0,0,0],"isMain":false},{"metricName":"# local misassemblies","quality":"Less is better","values":[0,0,0,0,0,0],"isMain":false},{"metricName":"# scaffold gap ext. mis.","quality":"Less is better","values":[0,0,0,0,0,0],"isMain":false},{"metricName":"# scaffold gap loc. mis.","quality":"Less is better","values":[0,0,0,0,0,0],"isMain":false}]],["Unaligned",[]],["Mismatches",[]],["Statistics without reference",[]],["K-mer-based statistics",[]],["Predicted genes",[]],["Similarity statistics",[]],["Reference statistics",[]]],[["Genome statistics",[]],["Reads mapping",[]],["Misassemblies",[{"metricName":"# misassemblies","quality":"Less is better","values":[0,1,0,0,0,0],"isMain":true},{"metricName":" # relocations","quality":"Less is better","values":[0,0,0,0,0,0],"isMain":false},{"metricName":" # translocations","quality":"Less is better","values":[0,0,0,0,0,0],"isMain":false},{"metricName":" # inversions","quality":"Less is better","values":[0,0,0,0,0,0],"isMain":false},{"metricName":" # interspecies translocations","quality":"Less is better","values":[0,1,0,0,0,0],"isMain":false},{"metricName":"# possibly misassembled contigs","quality":"Less is better","values":[0,0,0,0,0,0],"isMain":false},{"metricName":" # possible misassemblies","quality":"Less is better","values":[0,0,0,0,0,0],"isMain":false},{"metricName":"# local misassemblies","quality":"Less is better","values":[0,0,0,0,0,0],"isMain":false},{"metricName":"# scaffold gap ext. mis.","quality":"Less is better","values":[0,0,0,0,0,0],"isMain":false},{"metricName":"# scaffold gap loc. mis.","quality":"Less is better","values":[0,0,0,0,0,0],"isMain":false}]],["Unaligned",[]],["Mismatches",[]],["Statistics without reference",[]],["K-mer-based statistics",[]],["Predicted genes",[]],["Similarity statistics",[]],["Reference statistics",[]]],[["Genome statistics",[]],["Reads mapping",[]],["Misassemblies",[{"metricName":"# misassemblies","quality":"Less is better","values":[46,52,13,33,54,20],"isMain":true},{"metricName":" # relocations","quality":"Less is better","values":[0,1,0,1,2,0],"isMain":false},{"metricName":" # translocations","quality":"Less is better","values":[1,2,1,2,0,2],"isMain":false},{"metricName":" # inversions","quality":"Less is better","values":[1,3,0,1,1,0],"isMain":false},{"metricName":" # interspecies translocations","quality":"Less is better","values":[44,46,12,29,51,18],"isMain":false},{"metricName":"# possibly misassembled contigs","quality":"Less is better","values":[0,0,0,0,0,0],"isMain":false},{"metricName":" # possible misassemblies","quality":"Less is better","values":[0,0,0,0,0,0],"isMain":false},{"metricName":"# local misassemblies","quality":"Less is better","values":[3,1,0,4,2,2],"isMain":false},{"metricName":"# scaffold gap ext. mis.","quality":"Less is better","values":[0,0,0,0,0,0],"isMain":false},{"metricName":"# scaffold gap loc. mis.","quality":"Less is better","values":[0,0,0,0,0,0],"isMain":false}]],["Unaligned",[]],["Mismatches",[]],["Statistics without reference",[]],["K-mer-based statistics",[]],["Predicted genes",[]],["Similarity statistics",[]],["Reference statistics",[]]],[["Genome statistics",[]],["Reads mapping",[]],["Misassemblies",[{"metricName":"# misassemblies","quality":"Less is better","values":[22,25,13,18,20,10],"isMain":true},{"metricName":" # relocations","quality":"Less is better","values":[0,0,0,0,0,0],"isMain":false},{"metricName":" # translocations","quality":"Less is better","values":[0,0,0,0,0,0],"isMain":false},{"metricName":" # inversions","quality":"Less is better","values":[0,0,0,0,0,0],"isMain":false},{"metricName":" # interspecies translocations","quality":"Less is better","values":[22,25,13,18,20,10],"isMain":false},{"metricName":"# possibly misassembled contigs","quality":"Less is better","values":[0,1,0,0,0,0],"isMain":false},{"metricName":" # possible misassemblies","quality":"Less is better","values":[0,1,0,0,0,0],"isMain":false},{"metricName":"# local misassemblies","quality":"Less is better","values":[0,0,1,0,0,0],"isMain":false},{"metricName":"# scaffold gap ext. mis.","quality":"Less is better","values":[0,0,0,0,0,0],"isMain":false},{"metricName":"# scaffold gap loc. mis.","quality":"Less is better","values":[0,0,0,0,0,0],"isMain":false}]],["Unaligned",[]],["Mismatches",[]],["Statistics without reference",[]],["K-mer-based statistics",[]],["Predicted genes",[]],["Similarity statistics",[]],["Reference statistics",[]]],[["Genome statistics",[]],["Reads mapping",[]],["Misassemblies",[{"metricName":"# misassemblies","quality":"Less is better","values":[10,12,4,5,14,4],"isMain":true},{"metricName":" # relocations","quality":"Less is better","values":[0,0,0,0,1,0],"isMain":false},{"metricName":" # translocations","quality":"Less is better","values":[0,1,0,0,0,0],"isMain":false},{"metricName":" # inversions","quality":"Less is better","values":[0,0,0,0,0,0],"isMain":false},{"metricName":" # interspecies translocations","quality":"Less is better","values":[10,11,4,5,13,4],"isMain":false},{"metricName":"# possibly misassembled contigs","quality":"Less is better","values":[0,0,0,0,0,0],"isMain":false},{"metricName":" # possible misassemblies","quality":"Less is better","values":[0,0,0,0,0,0],"isMain":false},{"metricName":"# local misassemblies","quality":"Less is better","values":[2,1,0,0,2,0],"isMain":false},{"metricName":"# scaffold gap ext. mis.","quality":"Less is better","values":[0,0,0,0,0,0],"isMain":false},{"metricName":"# scaffold gap loc. mis.","quality":"Less is better","values":[0,0,0,0,0,0],"isMain":false}]],["Unaligned",[]],["Mismatches",[]],["Statistics without reference",[]],["K-mer-based statistics",[]],["Predicted genes",[]],["Similarity statistics",[]],["Reference statistics",[]]],[["Genome statistics",[]],["Reads mapping",[]],["Misassemblies",[{"metricName":"# misassemblies","quality":"Less is better","values":[19,15,15,24,16,3],"isMain":true},{"metricName":" # relocations","quality":"Less is better","values":[0,0,0,0,0,0],"isMain":false},{"metricName":" # translocations","quality":"Less is better","values":[3,1,2,1,2,0],"isMain":false},{"metricName":" # inversions","quality":"Less is better","values":[0,0,0,1,0,0],"isMain":false},{"metricName":" # interspecies translocations","quality":"Less is better","values":[16,14,13,22,14,3],"isMain":false},{"metricName":"# possibly misassembled contigs","quality":"Less is better","values":[0,0,1,0,0,0],"isMain":false},{"metricName":" # possible misassemblies","quality":"Less is better","values":[0,0,1,0,0,0],"isMain":false},{"metricName":"# local misassemblies","quality":"Less is better","values":[1,1,0,0,0,0],"isMain":false},{"metricName":"# scaffold gap ext. mis.","quality":"Less is better","values":[0,0,0,0,0,0],"isMain":false},{"metricName":"# scaffold gap loc. mis.","quality":"Less is better","values":[0,0,0,0,0,0],"isMain":false}]],["Unaligned",[]],["Mismatches",[]],["Statistics without reference",[]],["K-mer-based statistics",[]],["Predicted genes",[]],["Similarity statistics",[]],["Reference statistics",[]]],[["Genome statistics",[]],["Reads mapping",[]],["Misassemblies",[{"metricName":"# misassemblies","quality":"Less is better","values":[3,3,1,9,2,2],"isMain":true},{"metricName":" # relocations","quality":"Less is better","values":[0,0,0,0,0,0],"isMain":false},{"metricName":" # translocations","quality":"Less is better","values":[0,0,0,0,0,0],"isMain":false},{"metricName":" # inversions","quality":"Less is better","values":[0,0,0,0,0,0],"isMain":false},{"metricName":" # interspecies translocations","quality":"Less is better","values":[3,3,1,9,2,2],"isMain":false},{"metricName":"# possibly misassembled contigs","quality":"Less is better","values":[0,0,0,1,0,0],"isMain":false},{"metricName":" # possible misassemblies","quality":"Less is better","values":[0,0,0,1,0,0],"isMain":false},{"metricName":"# local misassemblies","quality":"Less is better","values":[0,0,0,0,0,0],"isMain":false},{"metricName":"# scaffold gap ext. mis.","quality":"Less is better","values":[0,0,0,0,0,0],"isMain":false},{"metricName":"# scaffold gap loc. mis.","quality":"Less is better","values":[0,0,0,0,0,0],"isMain":false}]],["Unaligned",[]],["Mismatches",[]],["Statistics without reference",[]],["K-mer-based statistics",[]],["Predicted genes",[]],["Similarity statistics",[]],["Reference statistics",[]]],[["Genome statistics",[]],["Reads mapping",[]],["Misassemblies",[{"metricName":"# misassemblies","quality":"Less is better","values":[19,21,10,10,17,7],"isMain":true},{"metricName":" # relocations","quality":"Less is better","values":[0,0,0,1,0,0],"isMain":false},{"metricName":" # translocations","quality":"Less is better","values":[0,0,0,1,1,0],"isMain":false},{"metricName":" # inversions","quality":"Less is better","values":[0,0,0,0,0,0],"isMain":false},{"metricName":" # interspecies translocations","quality":"Less is better","values":[19,21,10,8,16,7],"isMain":false},{"metricName":"# possibly misassembled contigs","quality":"Less is better","values":[0,0,0,0,0,0],"isMain":false},{"metricName":" # possible misassemblies","quality":"Less is better","values":[0,0,0,0,0,0],"isMain":false},{"metricName":"# local misassemblies","quality":"Less is better","values":[0,0,0,0,0,0],"isMain":false},{"metricName":"# scaffold gap ext. mis.","quality":"Less is better","values":[0,0,0,0,0,0],"isMain":false},{"metricName":"# scaffold gap loc. mis.","quality":"Less is better","values":[0,0,0,0,0,0],"isMain":false}]],["Unaligned",[]],["Mismatches",[]],["Statistics without reference",[]],["K-mer-based statistics",[]],["Predicted genes",[]],["Similarity statistics",[]],["Reference statistics",[]]],[["Genome statistics",[]],["Reads mapping",[]],["Misassemblies",[{"metricName":"# misassemblies","quality":"Less is better","values":[6,68,117,150,102,1],"isMain":true},{"metricName":" # relocations","quality":"Less is better","values":[0,6,6,12,10,0],"isMain":false},{"metricName":" # translocations","quality":"Less is better","values":[0,3,3,4,1,0],"isMain":false},{"metricName":" # inversions","quality":"Less is better","values":[0,0,1,2,2,0],"isMain":false},{"metricName":" # interspecies translocations","quality":"Less is better","values":[6,59,107,132,89,1],"isMain":false},{"metricName":"# possibly misassembled contigs","quality":"Less is better","values":[0,0,2,3,1,0],"isMain":false},{"metricName":" # possible misassemblies","quality":"Less is better","values":[0,0,2,3,1,0],"isMain":false},{"metricName":"# local misassemblies","quality":"Less is better","values":[0,6,6,8,7,1],"isMain":false},{"metricName":"# scaffold gap ext. mis.","quality":"Less is better","values":[0,0,0,0,0,0],"isMain":false},{"metricName":"# scaffold gap loc. mis.","quality":"Less is better","values":[0,0,0,0,0,0],"isMain":false}]],["Unaligned",[]],["Mismatches",[]],["Statistics without reference",[]],["K-mer-based statistics",[]],["Predicted genes",[]],["Similarity statistics",[]],["Reference statistics",[]]],[["Genome statistics",[]],["Reads mapping",[]],["Misassemblies",[{"metricName":"# misassemblies","quality":"Less is better","values":[2,9,5,8,10,5],"isMain":true},{"metricName":" # relocations","quality":"Less is better","values":[0,0,0,0,0,0],"isMain":false},{"metricName":" # translocations","quality":"Less is better","values":[0,0,0,0,0,0],"isMain":false},{"metricName":" # inversions","quality":"Less is better","values":[0,0,0,0,0,0],"isMain":false},{"metricName":" # interspecies translocations","quality":"Less is better","values":[2,9,5,8,10,5],"isMain":false},{"metricName":"# possibly misassembled contigs","quality":"Less is better","values":[0,0,0,0,0,0],"isMain":false},{"metricName":" # possible misassemblies","quality":"Less is better","values":[0,0,0,0,0,0],"isMain":false},{"metricName":"# local misassemblies","quality":"Less is better","values":[0,0,0,1,0,0],"isMain":false},{"metricName":"# scaffold gap ext. mis.","quality":"Less is better","values":[0,0,0,0,0,0],"isMain":false},{"metricName":"# scaffold gap loc. mis.","quality":"Less is better","values":[0,0,0,0,0,0],"isMain":false}]],["Unaligned",[]],["Mismatches",[]],["Statistics without reference",[]],["K-mer-based statistics",[]],["Predicted genes",[]],["Similarity statistics",[]],["Reference statistics",[]]],[["Genome statistics",[]],["Reads mapping",[]],["Misassemblies",[{"metricName":"# misassemblies","quality":"Less is better","values":[13,12,4,18,16,3],"isMain":true},{"metricName":" # relocations","quality":"Less is better","values":[0,0,0,0,0,0],"isMain":false},{"metricName":" # translocations","quality":"Less is better","values":[0,0,0,0,0,0],"isMain":false},{"metricName":" # inversions","quality":"Less is better","values":[0,0,0,0,0,0],"isMain":false},{"metricName":" # interspecies translocations","quality":"Less is better","values":[13,12,4,18,16,3],"isMain":false},{"metricName":"# possibly misassembled contigs","quality":"Less is better","values":[0,0,1,0,0,0],"isMain":false},{"metricName":" # possible misassemblies","quality":"Less is better","values":[0,0,1,0,0,0],"isMain":false},{"metricName":"# local misassemblies","quality":"Less is better","values":[0,1,0,0,0,0],"isMain":false},{"metricName":"# scaffold gap ext. mis.","quality":"Less is better","values":[0,0,0,0,0,0],"isMain":false},{"metricName":"# scaffold gap loc. mis.","quality":"Less is better","values":[0,0,0,0,0,0],"isMain":false}]],["Unaligned",[]],["Mismatches",[]],["Statistics without reference",[]],["K-mer-based statistics",[]],["Predicted genes",[]],["Similarity statistics",[]],["Reference statistics",[]]],[["Genome statistics",[]],["Reads mapping",[]],["Misassemblies",[{"metricName":"# misassemblies","quality":"Less is better","values":[1,1,2,7,4,1],"isMain":true},{"metricName":" # relocations","quality":"Less is better","values":[0,0,0,0,0,0],"isMain":false},{"metricName":" # translocations","quality":"Less is better","values":[0,0,0,0,0,0],"isMain":false},{"metricName":" # inversions","quality":"Less is better","values":[0,0,0,0,0,0],"isMain":false},{"metricName":" # interspecies translocations","quality":"Less is better","values":[1,1,2,7,4,1],"isMain":false},{"metricName":"# possibly misassembled contigs","quality":"Less is better","values":[0,0,0,0,0,0],"isMain":false},{"metricName":" # possible misassemblies","quality":"Less is better","values":[0,0,0,0,0,0],"isMain":false},{"metricName":"# local misassemblies","quality":"Less is better","values":[0,0,0,0,0,0],"isMain":false},{"metricName":"# scaffold gap ext. mis.","quality":"Less is better","values":[0,0,0,0,0,0],"isMain":false},{"metricName":"# scaffold gap loc. mis.","quality":"Less is better","values":[0,0,0,0,0,0],"isMain":false}]],["Unaligned",[]],["Mismatches",[]],["Statistics without reference",[]],["K-mer-based statistics",[]],["Predicted genes",[]],["Similarity statistics",[]],["Reference statistics",[]]],[["Genome statistics",[]],["Reads mapping",[]],["Misassemblies",[{"metricName":"# misassemblies","quality":"Less is better","values":[39,45,37,82,62,23],"isMain":true},{"metricName":" # relocations","quality":"Less is better","values":[0,0,0,0,2,0],"isMain":false},{"metricName":" # translocations","quality":"Less is better","values":[2,16,14,14,19,15],"isMain":false},{"metricName":" # inversions","quality":"Less is better","values":[1,1,0,0,0,0],"isMain":false},{"metricName":" # interspecies translocations","quality":"Less is better","values":[36,28,23,68,41,8],"isMain":false},{"metricName":"# possibly misassembled contigs","quality":"Less is better","values":[0,0,0,0,0,0],"isMain":false},{"metricName":" # possible misassemblies","quality":"Less is better","values":[0,0,0,0,0,0],"isMain":false},{"metricName":"# local misassemblies","quality":"Less is better","values":[0,2,1,0,3,0],"isMain":false},{"metricName":"# scaffold gap ext. mis.","quality":"Less is better","values":[0,0,0,0,0,0],"isMain":false},{"metricName":"# scaffold gap loc. mis.","quality":"Less is better","values":[0,0,0,0,0,0],"isMain":false}]],["Unaligned",[]],["Mismatches",[]],["Statistics without reference",[]],["K-mer-based statistics",[]],["Predicted genes",[]],["Similarity statistics",[]],["Reference statistics",[]]],[["Genome statistics",[]],["Reads mapping",[]],["Misassemblies",[{"metricName":"# misassemblies","quality":"Less is better","values":[16,27,30,58,41,10],"isMain":true},{"metricName":" # relocations","quality":"Less is better","values":[0,0,0,0,0,0],"isMain":false},{"metricName":" # translocations","quality":"Less is better","values":[5,8,9,7,6,3],"isMain":false},{"metricName":" # inversions","quality":"Less is better","values":[0,1,1,2,1,0],"isMain":false},{"metricName":" # interspecies translocations","quality":"Less is better","values":[11,18,20,49,34,7],"isMain":false},{"metricName":"# possibly misassembled contigs","quality":"Less is better","values":[0,0,1,0,0,1],"isMain":false},{"metricName":" # possible misassemblies","quality":"Less is better","values":[0,0,2,0,0,2],"isMain":false},{"metricName":"# local misassemblies","quality":"Less is better","values":[1,6,5,3,2,2],"isMain":false},{"metricName":"# scaffold gap ext. mis.","quality":"Less is better","values":[0,0,0,0,0,0],"isMain":false},{"metricName":"# scaffold gap loc. mis.","quality":"Less is better","values":[0,0,0,0,0,0],"isMain":false}]],["Unaligned",[]],["Mismatches",[]],["Statistics without reference",[]],["K-mer-based statistics",[]],["Predicted genes",[]],["Similarity statistics",[]],["Reference statistics",[]]],[["Genome statistics",[]],["Reads mapping",[]],["Misassemblies",[{"metricName":"# misassemblies","quality":"Less is better","values":[23,33,13,27,29,11],"isMain":true},{"metricName":" # relocations","quality":"Less is better","values":[0,0,0,0,0,0],"isMain":false},{"metricName":" # translocations","quality":"Less is better","values":[5,2,5,2,3,1],"isMain":false},{"metricName":" # inversions","quality":"Less is better","values":[0,0,1,0,1,0],"isMain":false},{"metricName":" # interspecies translocations","quality":"Less is better","values":[18,31,7,25,25,10],"isMain":false},{"metricName":"# possibly misassembled contigs","quality":"Less is better","values":[0,0,0,0,1,0],"isMain":false},{"metricName":" # possible misassemblies","quality":"Less is better","values":[0,0,0,0,1,0],"isMain":false},{"metricName":"# local misassemblies","quality":"Less is better","values":[2,1,2,3,1,1],"isMain":false},{"metricName":"# scaffold gap ext. mis.","quality":"Less is better","values":[0,0,0,0,0,0],"isMain":false},{"metricName":"# scaffold gap loc. mis.","quality":"Less is better","values":[0,0,0,0,0,0],"isMain":false}]],["Unaligned",[]],["Mismatches",[]],["Statistics without reference",[]],["K-mer-based statistics",[]],["Predicted genes",[]],["Similarity statistics",[]],["Reference statistics",[]]],[["Genome statistics",[]],["Reads mapping",[]],["Misassemblies",[{"metricName":"# misassemblies","quality":"Less is better","values":[10,23,8,12,16,8],"isMain":true},{"metricName":" # relocations","quality":"Less is better","values":[0,0,0,0,0,0],"isMain":false},{"metricName":" # translocations","quality":"Less is better","values":[0,0,0,0,0,0],"isMain":false},{"metricName":" # inversions","quality":"Less is better","values":[0,0,0,0,0,0],"isMain":false},{"metricName":" # interspecies translocations","quality":"Less is better","values":[10,23,8,12,16,8],"isMain":false},{"metricName":"# possibly misassembled contigs","quality":"Less is better","values":[0,0,0,0,0,0],"isMain":false},{"metricName":" # possible misassemblies","quality":"Less is better","values":[0,0,0,0,0,0],"isMain":false},{"metricName":"# local misassemblies","quality":"Less is better","values":[1,1,1,0,1,0],"isMain":false},{"metricName":"# scaffold gap ext. mis.","quality":"Less is better","values":[0,0,0,0,0,0],"isMain":false},{"metricName":"# scaffold gap loc. mis.","quality":"Less is better","values":[0,0,0,0,0,0],"isMain":false}]],["Unaligned",[]],["Mismatches",[]],["Statistics without reference",[]],["K-mer-based statistics",[]],["Predicted genes",[]],["Similarity statistics",[]],["Reference statistics",[]]],[["Genome statistics",[]],["Reads mapping",[]],["Misassemblies",[{"metricName":"# misassemblies","quality":"Less is better","values":[12,14,8,14,10,5],"isMain":true},{"metricName":" # relocations","quality":"Less is better","values":[0,2,0,1,1,0],"isMain":false},{"metricName":" # translocations","quality":"Less is better","values":[1,1,0,1,2,0],"isMain":false},{"metricName":" # inversions","quality":"Less is better","values":[0,1,0,0,0,0],"isMain":false},{"metricName":" # interspecies translocations","quality":"Less is better","values":[11,10,8,12,7,5],"isMain":false},{"metricName":"# possibly misassembled contigs","quality":"Less is better","values":[0,0,0,0,0,0],"isMain":false},{"metricName":" # possible misassemblies","quality":"Less is better","values":[0,0,0,0,0,0],"isMain":false},{"metricName":"# local misassemblies","quality":"Less is better","values":[3,0,3,2,0,3],"isMain":false},{"metricName":"# scaffold gap ext. mis.","quality":"Less is better","values":[0,0,0,0,0,0],"isMain":false},{"metricName":"# scaffold gap loc. mis.","quality":"Less is better","values":[0,0,0,0,0,0],"isMain":false}]],["Unaligned",[]],["Mismatches",[]],["Statistics without reference",[]],["K-mer-based statistics",[]],["Predicted genes",[]],["Similarity statistics",[]],["Reference statistics",[]]],[["Genome statistics",[]],["Reads mapping",[]],["Misassemblies",[{"metricName":"# misassemblies","quality":"Less is better","values":[10,13,13,20,19,9],"isMain":true},{"metricName":" # relocations","quality":"Less is better","values":[0,0,0,0,0,0],"isMain":false},{"metricName":" # translocations","quality":"Less is better","values":[3,5,4,4,3,6],"isMain":false},{"metricName":" # inversions","quality":"Less is better","values":[0,0,0,0,0,0],"isMain":false},{"metricName":" # interspecies translocations","quality":"Less is better","values":[7,8,9,16,16,3],"isMain":false},{"metricName":"# possibly misassembled contigs","quality":"Less is better","values":[0,0,0,0,0,0],"isMain":false},{"metricName":" # possible misassemblies","quality":"Less is better","values":[0,0,0,0,0,0],"isMain":false},{"metricName":"# local misassemblies","quality":"Less is better","values":[1,1,2,2,2,1],"isMain":false},{"metricName":"# scaffold gap ext. mis.","quality":"Less is better","values":[0,0,0,0,0,0],"isMain":false},{"metricName":"# scaffold gap loc. mis.","quality":"Less is better","values":[0,0,0,0,0,0],"isMain":false}]],["Unaligned",[]],["Mismatches",[]],["Statistics without reference",[]],["K-mer-based statistics",[]],["Predicted genes",[]],["Similarity statistics",[]],["Reference statistics",[]]],[["Genome statistics",[]],["Reads mapping",[]],["Misassemblies",[{"metricName":"# misassemblies","quality":"Less is better","values":[6,14,10,19,15,5],"isMain":true},{"metricName":" # relocations","quality":"Less is better","values":[0,0,0,0,0,0],"isMain":false},{"metricName":" # translocations","quality":"Less is better","values":[0,0,0,1,0,0],"isMain":false},{"metricName":" # inversions","quality":"Less is better","values":[1,0,1,0,0,0],"isMain":false},{"metricName":" # interspecies translocations","quality":"Less is better","values":[5,14,9,18,15,5],"isMain":false},{"metricName":"# possibly misassembled contigs","quality":"Less is better","values":[0,0,0,0,0,0],"isMain":false},{"metricName":" # possible misassemblies","quality":"Less is better","values":[0,0,0,0,0,0],"isMain":false},{"metricName":"# local misassemblies","quality":"Less is better","values":[0,0,0,1,0,0],"isMain":false},{"metricName":"# scaffold gap ext. mis.","quality":"Less is better","values":[0,0,0,0,0,0],"isMain":false},{"metricName":"# scaffold gap loc. mis.","quality":"Less is better","values":[0,0,0,0,0,0],"isMain":false}]],["Unaligned",[]],["Mismatches",[]],["Statistics without reference",[]],["K-mer-based statistics",[]],["Predicted genes",[]],["Similarity statistics",[]],["Reference statistics",[]]],[["Genome statistics",[]],["Reads mapping",[]],["Misassemblies",[{"metricName":"# misassemblies","quality":"Less is better","values":[5,10,4,8,8,3],"isMain":true},{"metricName":" # relocations","quality":"Less is better","values":[0,0,1,0,0,0],"isMain":false},{"metricName":" # translocations","quality":"Less is better","values":[0,1,0,0,0,0],"isMain":false},{"metricName":" # inversions","quality":"Less is better","values":[0,0,0,0,0,0],"isMain":false},{"metricName":" # interspecies translocations","quality":"Less is better","values":[5,9,3,8,8,3],"isMain":false},{"metricName":"# possibly misassembled contigs","quality":"Less is better","values":[0,0,0,0,0,0],"isMain":false},{"metricName":" # possible misassemblies","quality":"Less is better","values":[0,0,0,0,0,0],"isMain":false},{"metricName":"# local misassemblies","quality":"Less is better","values":[0,0,0,0,0,0],"isMain":false},{"metricName":"# scaffold gap ext. mis.","quality":"Less is better","values":[0,0,0,0,0,0],"isMain":false},{"metricName":"# scaffold gap loc. mis.","quality":"Less is better","values":[0,0,0,0,0,0],"isMain":false}]],["Unaligned",[]],["Mismatches",[]],["Statistics without reference",[]],["K-mer-based statistics",[]],["Predicted genes",[]],["Similarity statistics",[]],["Reference statistics",[]]],[["Genome statistics",[]],["Reads mapping",[]],["Misassemblies",[{"metricName":"# misassemblies","quality":"Less is better","values":[19,31,10,30,27,18],"isMain":true},{"metricName":" # relocations","quality":"Less is better","values":[2,5,3,7,2,1],"isMain":false},{"metricName":" # translocations","quality":"Less is better","values":[0,0,0,0,0,0],"isMain":false},{"metricName":" # inversions","quality":"Less is better","values":[0,0,0,0,0,0],"isMain":false},{"metricName":" # interspecies translocations","quality":"Less is better","values":[17,26,7,23,25,17],"isMain":false},{"metricName":"# possibly misassembled contigs","quality":"Less is better","values":[0,0,0,0,0,0],"isMain":false},{"metricName":" # possible misassemblies","quality":"Less is better","values":[0,0,0,0,0,0],"isMain":false},{"metricName":"# local misassemblies","quality":"Less is better","values":[1,0,1,0,1,0],"isMain":false},{"metricName":"# scaffold gap ext. mis.","quality":"Less is better","values":[0,0,0,0,0,0],"isMain":false},{"metricName":"# scaffold gap loc. mis.","quality":"Less is better","values":[0,0,0,0,0,0],"isMain":false}]],["Unaligned",[]],["Mismatches",[]],["Statistics without reference",[]],["K-mer-based statistics",[]],["Predicted genes",[]],["Similarity statistics",[]],["Reference statistics",[]]],[["Genome statistics",[]],["Reads mapping",[]],["Misassemblies",[{"metricName":"# misassemblies","quality":"Less is better","values":[7,45,41,57,56,7],"isMain":true},{"metricName":" # relocations","quality":"Less is better","values":[0,2,4,4,2,0],"isMain":false},{"metricName":" # translocations","quality":"Less is better","values":[0,0,0,0,0,0],"isMain":false},{"metricName":" # inversions","quality":"Less is better","values":[0,0,2,0,0,0],"isMain":false},{"metricName":" # interspecies translocations","quality":"Less is better","values":[7,43,35,53,54,7],"isMain":false},{"metricName":"# possibly misassembled contigs","quality":"Less is better","values":[0,2,0,1,0,1],"isMain":false},{"metricName":" # possible misassemblies","quality":"Less is better","values":[0,2,0,1,0,1],"isMain":false},{"metricName":"# local misassemblies","quality":"Less is better","values":[2,4,3,4,4,0],"isMain":false},{"metricName":"# scaffold gap ext. mis.","quality":"Less is better","values":[0,0,0,0,0,0],"isMain":false},{"metricName":"# scaffold gap loc. mis.","quality":"Less is better","values":[0,0,0,0,0,0],"isMain":false}]],["Unaligned",[]],["Mismatches",[]],["Statistics without reference",[]],["K-mer-based statistics",[]],["Predicted genes",[]],["Similarity statistics",[]],["Reference statistics",[]]],[["Genome statistics",[]],["Reads mapping",[]],["Misassemblies",[{"metricName":"# misassemblies","quality":"Less is better","values":[4,8,3,5,10,0],"isMain":true},{"metricName":" # relocations","quality":"Less is better","values":[0,0,0,0,0,0],"isMain":false},{"metricName":" # translocations","quality":"Less is better","values":[0,0,0,0,0,0],"isMain":false},{"metricName":" # inversions","quality":"Less is better","values":[0,0,0,0,0,0],"isMain":false},{"metricName":" # interspecies translocations","quality":"Less is better","values":[4,8,3,5,10,0],"isMain":false},{"metricName":"# possibly misassembled contigs","quality":"Less is better","values":[0,0,0,0,0,0],"isMain":false},{"metricName":" # possible misassemblies","quality":"Less is better","values":[0,0,0,0,0,0],"isMain":false},{"metricName":"# local misassemblies","quality":"Less is better","values":[0,0,0,0,1,0],"isMain":false},{"metricName":"# scaffold gap ext. mis.","quality":"Less is better","values":[0,0,0,0,0,0],"isMain":false},{"metricName":"# scaffold gap loc. mis.","quality":"Less is better","values":[0,0,0,0,0,0],"isMain":false}]],["Unaligned",[]],["Mismatches",[]],["Statistics without reference",[]],["K-mer-based statistics",[]],["Predicted genes",[]],["Similarity statistics",[]],["Reference statistics",[]]],[["Genome statistics",[]],["Reads mapping",[]],["Misassemblies",[{"metricName":"# misassemblies","quality":"Less is better","values":[1,1,6,5,3,0],"isMain":true},{"metricName":" # relocations","quality":"Less is better","values":[0,1,2,2,0,0],"isMain":false},{"metricName":" # translocations","quality":"Less is better","values":[0,0,0,0,0,0],"isMain":false},{"metricName":" # inversions","quality":"Less is better","values":[0,0,0,0,0,0],"isMain":false},{"metricName":" # interspecies translocations","quality":"Less is better","values":[1,0,4,3,3,0],"isMain":false},{"metricName":"# possibly misassembled contigs","quality":"Less is better","values":[0,0,0,0,0,0],"isMain":false},{"metricName":" # possible misassemblies","quality":"Less is better","values":[0,0,0,0,0,0],"isMain":false},{"metricName":"# local misassemblies","quality":"Less is better","values":[0,0,0,0,1,0],"isMain":false},{"metricName":"# scaffold gap ext. mis.","quality":"Less is better","values":[0,0,0,0,0,0],"isMain":false},{"metricName":"# scaffold gap loc. mis.","quality":"Less is better","values":[0,0,0,0,0,0],"isMain":false}]],["Unaligned",[]],["Mismatches",[]],["Statistics without reference",[]],["K-mer-based statistics",[]],["Predicted genes",[]],["Similarity statistics",[]],["Reference statistics",[]]],[["Genome statistics",[]],["Reads mapping",[]],["Misassemblies",[{"metricName":"# misassemblies","quality":"Less is better","values":[2,4,5,20,6,1],"isMain":true},{"metricName":" # relocations","quality":"Less is better","values":[0,0,1,0,0,0],"isMain":false},{"metricName":" # translocations","quality":"Less is better","values":[0,0,0,0,0,0],"isMain":false},{"metricName":" # inversions","quality":"Less is better","values":[0,0,1,2,0,0],"isMain":false},{"metricName":" # interspecies translocations","quality":"Less is better","values":[2,4,3,18,6,1],"isMain":false},{"metricName":"# possibly misassembled contigs","quality":"Less is better","values":[1,1,0,0,0,0],"isMain":false},{"metricName":" # possible misassemblies","quality":"Less is better","values":[1,1,0,0,0,0],"isMain":false},{"metricName":"# local misassemblies","quality":"Less is better","values":[0,3,3,1,0,0],"isMain":false},{"metricName":"# scaffold gap ext. mis.","quality":"Less is better","values":[0,0,0,0,0,0],"isMain":false},{"metricName":"# scaffold gap loc. mis.","quality":"Less is better","values":[0,0,0,0,0,0],"isMain":false}]],["Unaligned",[]],["Mismatches",[]],["Statistics without reference",[]],["K-mer-based statistics",[]],["Predicted genes",[]],["Similarity statistics",[]],["Reference statistics",[]]],[["Genome statistics",[]],["Reads mapping",[]],["Misassemblies",[{"metricName":"# misassemblies","quality":"Less is better","values":[2,1,1,0,1,0],"isMain":true},{"metricName":" # relocations","quality":"Less is better","values":[0,0,0,0,0,0],"isMain":false},{"metricName":" # translocations","quality":"Less is better","values":[0,0,0,0,0,0],"isMain":false},{"metricName":" # inversions","quality":"Less is better","values":[0,0,0,0,0,0],"isMain":false},{"metricName":" # interspecies translocations","quality":"Less is better","values":[2,1,1,0,1,0],"isMain":false},{"metricName":"# possibly misassembled contigs","quality":"Less is better","values":[0,0,0,0,0,0],"isMain":false},{"metricName":" # possible misassemblies","quality":"Less is better","values":[0,0,0,0,0,0],"isMain":false},{"metricName":"# local misassemblies","quality":"Less is better","values":[0,0,0,0,0,0],"isMain":false},{"metricName":"# scaffold gap ext. mis.","quality":"Less is better","values":[0,0,0,0,0,0],"isMain":false},{"metricName":"# scaffold gap loc. mis.","quality":"Less is better","values":[0,0,0,0,0,0],"isMain":false}]],["Unaligned",[]],["Mismatches",[]],["Statistics without reference",[]],["K-mer-based statistics",[]],["Predicted genes",[]],["Similarity statistics",[]],["Reference statistics",[]]],[["Genome statistics",[]],["Reads mapping",[]],["Misassemblies",[{"metricName":"# misassemblies","quality":"Less is better","values":[0,9,2,5,7,2],"isMain":true},{"metricName":" # relocations","quality":"Less is better","values":[0,0,0,0,0,1],"isMain":false},{"metricName":" # translocations","quality":"Less is better","values":[0,0,0,0,0,0],"isMain":false},{"metricName":" # inversions","quality":"Less is better","values":[0,0,0,0,0,0],"isMain":false},{"metricName":" # interspecies translocations","quality":"Less is better","values":[0,9,2,5,7,1],"isMain":false},{"metricName":"# possibly misassembled contigs","quality":"Less is better","values":[0,0,0,0,0,0],"isMain":false},{"metricName":" # possible misassemblies","quality":"Less is better","values":[0,0,0,0,0,0],"isMain":false},{"metricName":"# local misassemblies","quality":"Less is better","values":[0,0,0,0,0,0],"isMain":false},{"metricName":"# scaffold gap ext. mis.","quality":"Less is better","values":[0,0,0,0,0,0],"isMain":false},{"metricName":"# scaffold gap loc. mis.","quality":"Less is better","values":[0,0,0,0,0,0],"isMain":false}]],["Unaligned",[]],["Mismatches",[]],["Statistics without reference",[]],["K-mer-based statistics",[]],["Predicted genes",[]],["Similarity statistics",[]],["Reference statistics",[]]],[["Genome statistics",[]],["Reads mapping",[]],["Misassemblies",[{"metricName":"# misassemblies","quality":"Less is better","values":[58,13,24,38,15,1],"isMain":true},{"metricName":" # relocations","quality":"Less is better","values":[1,0,0,0,0,0],"isMain":false},{"metricName":" # translocations","quality":"Less is better","values":[0,1,2,4,3,0],"isMain":false},{"metricName":" # inversions","quality":"Less is better","values":[0,0,1,1,0,0],"isMain":false},{"metricName":" # interspecies translocations","quality":"Less is better","values":[57,12,21,33,12,1],"isMain":false},{"metricName":"# possibly misassembled contigs","quality":"Less is better","values":[0,0,0,0,0,0],"isMain":false},{"metricName":" # possible misassemblies","quality":"Less is better","values":[0,0,0,0,0,0],"isMain":false},{"metricName":"# local misassemblies","quality":"Less is better","values":[0,1,2,4,4,0],"isMain":false},{"metricName":"# scaffold gap ext. mis.","quality":"Less is better","values":[0,0,0,0,0,0],"isMain":false},{"metricName":"# scaffold gap loc. mis.","quality":"Less is better","values":[0,0,0,0,0,0],"isMain":false}]],["Unaligned",[]],["Mismatches",[]],["Statistics without reference",[]],["K-mer-based statistics",[]],["Predicted genes",[]],["Similarity statistics",[]],["Reference statistics",[]]],[["Genome statistics",[]],["Reads mapping",[]],["Misassemblies",[{"metricName":"# misassemblies","quality":"Less is better","values":[15,25,8,16,20,18],"isMain":true},{"metricName":" # relocations","quality":"Less is better","values":[0,0,0,0,0,0],"isMain":false},{"metricName":" # translocations","quality":"Less is better","values":[1,4,3,4,5,3],"isMain":false},{"metricName":" # inversions","quality":"Less is better","values":[1,0,0,0,0,0],"isMain":false},{"metricName":" # interspecies translocations","quality":"Less is better","values":[13,21,5,12,15,15],"isMain":false},{"metricName":"# possibly misassembled contigs","quality":"Less is better","values":[0,0,0,0,0,0],"isMain":false},{"metricName":" # possible misassemblies","quality":"Less is better","values":[0,0,0,0,0,0],"isMain":false},{"metricName":"# local misassemblies","quality":"Less is better","values":[0,0,0,2,2,0],"isMain":false},{"metricName":"# scaffold gap ext. mis.","quality":"Less is better","values":[0,0,0,0,0,0],"isMain":false},{"metricName":"# scaffold gap loc. mis.","quality":"Less is better","values":[0,0,0,0,0,0],"isMain":false}]],["Unaligned",[]],["Mismatches",[]],["Statistics without reference",[]],["K-mer-based statistics",[]],["Predicted genes",[]],["Similarity statistics",[]],["Reference statistics",[]]],[["Genome statistics",[]],["Reads mapping",[]],["Misassemblies",[{"metricName":"# misassemblies","quality":"Less is better","values":[6,15,11,30,18,5],"isMain":true},{"metricName":" # relocations","quality":"Less is better","values":[0,0,0,0,0,0],"isMain":false},{"metricName":" # translocations","quality":"Less is better","values":[2,2,3,6,7,3],"isMain":false},{"metricName":" # inversions","quality":"Less is better","values":[0,0,0,0,0,0],"isMain":false},{"metricName":" # interspecies translocations","quality":"Less is better","values":[4,13,8,24,11,2],"isMain":false},{"metricName":"# possibly misassembled contigs","quality":"Less is better","values":[0,0,0,0,1,0],"isMain":false},{"metricName":" # possible misassemblies","quality":"Less is better","values":[0,0,0,0,1,0],"isMain":false},{"metricName":"# local misassemblies","quality":"Less is better","values":[1,1,1,1,1,0],"isMain":false},{"metricName":"# scaffold gap ext. mis.","quality":"Less is better","values":[0,0,0,0,0,0],"isMain":false},{"metricName":"# scaffold gap loc. mis.","quality":"Less is better","values":[0,0,0,0,0,0],"isMain":false}]],["Unaligned",[]],["Mismatches",[]],["Statistics without reference",[]],["K-mer-based statistics",[]],["Predicted genes",[]],["Similarity statistics",[]],["Reference statistics",[]]],[["Genome statistics",[]],["Reads mapping",[]],["Misassemblies",[{"metricName":"# misassemblies","quality":"Less is better","values":[36,55,13,36,49,18],"isMain":true},{"metricName":" # relocations","quality":"Less is better","values":[0,1,0,0,0,0],"isMain":false},{"metricName":" # translocations","quality":"Less is better","values":[4,6,4,3,5,0],"isMain":false},{"metricName":" # inversions","quality":"Less is better","values":[0,0,0,0,0,0],"isMain":false},{"metricName":" # interspecies translocations","quality":"Less is better","values":[32,48,9,33,44,18],"isMain":false},{"metricName":"# possibly misassembled contigs","quality":"Less is better","values":[1,1,0,0,1,0],"isMain":false},{"metricName":" # possible misassemblies","quality":"Less is better","values":[1,1,0,0,1,0],"isMain":false},{"metricName":"# local misassemblies","quality":"Less is better","values":[0,1,0,1,0,0],"isMain":false},{"metricName":"# scaffold gap ext. mis.","quality":"Less is better","values":[0,0,0,0,0,0],"isMain":false},{"metricName":"# scaffold gap loc. mis.","quality":"Less is better","values":[0,0,0,0,0,0],"isMain":false}]],["Unaligned",[]],["Mismatches",[]],["Statistics without reference",[]],["K-mer-based statistics",[]],["Predicted genes",[]],["Similarity statistics",[]],["Reference statistics",[]]],[["Genome statistics",[]],["Reads mapping",[]],["Misassemblies",[{"metricName":"# misassemblies","quality":"Less is better","values":[62,113,143,247,165,3],"isMain":true},{"metricName":" # relocations","quality":"Less is better","values":[0,2,8,12,8,0],"isMain":false},{"metricName":" # translocations","quality":"Less is better","values":[0,4,3,1,3,0],"isMain":false},{"metricName":" # inversions","quality":"Less is better","values":[2,4,4,5,13,0],"isMain":false},{"metricName":" # interspecies translocations","quality":"Less is better","values":[60,103,128,229,141,3],"isMain":false},{"metricName":"# possibly misassembled contigs","quality":"Less is better","values":[0,0,1,2,0,0],"isMain":false},{"metricName":" # possible misassemblies","quality":"Less is better","values":[0,0,1,2,0,0],"isMain":false},{"metricName":"# local misassemblies","quality":"Less is better","values":[0,5,8,9,14,0],"isMain":false},{"metricName":"# scaffold gap ext. mis.","quality":"Less is better","values":[0,0,0,0,0,0],"isMain":false},{"metricName":"# scaffold gap loc. mis.","quality":"Less is better","values":[0,0,0,0,0,0],"isMain":false}]],["Unaligned",[]],["Mismatches",[]],["Statistics without reference",[]],["K-mer-based statistics",[]],["Predicted genes",[]],["Similarity statistics",[]],["Reference statistics",[]]],[["Genome statistics",[]],["Reads mapping",[]],["Misassemblies",[{"metricName":"# misassemblies","quality":"Less is better","values":[63,73,36,80,61,20],"isMain":true},{"metricName":" # relocations","quality":"Less is better","values":[0,0,2,3,1,1],"isMain":false},{"metricName":" # translocations","quality":"Less is better","values":[9,10,11,12,15,4],"isMain":false},{"metricName":" # inversions","quality":"Less is better","values":[2,2,2,2,3,0],"isMain":false},{"metricName":" # interspecies translocations","quality":"Less is better","values":[52,61,21,63,42,15],"isMain":false},{"metricName":"# possibly misassembled contigs","quality":"Less is better","values":[0,0,1,0,0,0],"isMain":false},{"metricName":" # possible misassemblies","quality":"Less is better","values":[0,0,1,0,0,0],"isMain":false},{"metricName":"# local misassemblies","quality":"Less is better","values":[1,9,4,5,2,2],"isMain":false},{"metricName":"# scaffold gap ext. mis.","quality":"Less is better","values":[0,0,0,0,0,0],"isMain":false},{"metricName":"# scaffold gap loc. mis.","quality":"Less is better","values":[0,0,0,0,0,0],"isMain":false}]],["Unaligned",[]],["Mismatches",[]],["Statistics without reference",[]],["K-mer-based statistics",[]],["Predicted genes",[]],["Similarity statistics",[]],["Reference statistics",[]]],[["Genome statistics",[]],["Reads mapping",[]],["Misassemblies",[{"metricName":"# misassemblies","quality":"Less is better","values":[35,52,17,54,50,12],"isMain":true},{"metricName":" # relocations","quality":"Less is better","values":[0,1,2,3,1,1],"isMain":false},{"metricName":" # translocations","quality":"Less is better","values":[1,1,2,3,0,1],"isMain":false},{"metricName":" # inversions","quality":"Less is better","values":[1,2,4,2,5,0],"isMain":false},{"metricName":" # interspecies translocations","quality":"Less is better","values":[33,48,9,46,44,10],"isMain":false},{"metricName":"# possibly misassembled contigs","quality":"Less is better","values":[0,0,0,0,1,0],"isMain":false},{"metricName":" # possible misassemblies","quality":"Less is better","values":[0,0,0,0,1,0],"isMain":false},{"metricName":"# local misassemblies","quality":"Less is better","values":[4,0,3,1,2,1],"isMain":false},{"metricName":"# scaffold gap ext. mis.","quality":"Less is better","values":[0,0,0,0,0,0],"isMain":false},{"metricName":"# scaffold gap loc. mis.","quality":"Less is better","values":[0,0,0,0,0,0],"isMain":false}]],["Unaligned",[]],["Mismatches",[]],["Statistics without reference",[]],["K-mer-based statistics",[]],["Predicted genes",[]],["Similarity statistics",[]],["Reference statistics",[]]],[["Genome statistics",[]],["Reads mapping",[]],["Misassemblies",[{"metricName":"# misassemblies","quality":"Less is better","values":[17,20,6,9,23,5],"isMain":true},{"metricName":" # relocations","quality":"Less is better","values":[0,0,0,0,0,0],"isMain":false},{"metricName":" # translocations","quality":"Less is better","values":[0,1,1,0,1,0],"isMain":false},{"metricName":" # inversions","quality":"Less is better","values":[0,0,0,0,0,0],"isMain":false},{"metricName":" # interspecies translocations","quality":"Less is better","values":[17,19,5,9,22,5],"isMain":false},{"metricName":"# possibly misassembled contigs","quality":"Less is better","values":[0,0,0,0,0,0],"isMain":false},{"metricName":" # possible misassemblies","quality":"Less is better","values":[0,0,0,0,0,0],"isMain":false},{"metricName":"# local misassemblies","quality":"Less is better","values":[0,0,0,0,0,0],"isMain":false},{"metricName":"# scaffold gap ext. mis.","quality":"Less is better","values":[0,0,0,0,0,0],"isMain":false},{"metricName":"# scaffold gap loc. mis.","quality":"Less is better","values":[0,0,0,0,0,0],"isMain":false}]],["Unaligned",[]],["Mismatches",[]],["Statistics without reference",[]],["K-mer-based statistics",[]],["Predicted genes",[]],["Similarity statistics",[]],["Reference statistics",[]]],[["Genome statistics",[]],["Reads mapping",[]],["Misassemblies",[{"metricName":"# misassemblies","quality":"Less is better","values":[22,30,10,26,30,13],"isMain":true},{"metricName":" # relocations","quality":"Less is better","values":[1,0,0,0,0,0],"isMain":false},{"metricName":" # translocations","quality":"Less is better","values":[3,4,4,4,4,2],"isMain":false},{"metricName":" # inversions","quality":"Less is better","values":[0,0,0,1,0,0],"isMain":false},{"metricName":" # interspecies translocations","quality":"Less is better","values":[18,26,6,21,26,11],"isMain":false},{"metricName":"# possibly misassembled contigs","quality":"Less is better","values":[0,0,0,0,0,0],"isMain":false},{"metricName":" # possible misassemblies","quality":"Less is better","values":[0,0,0,0,0,0],"isMain":false},{"metricName":"# local misassemblies","quality":"Less is better","values":[2,2,0,0,0,0],"isMain":false},{"metricName":"# scaffold gap ext. mis.","quality":"Less is better","values":[0,0,0,0,0,0],"isMain":false},{"metricName":"# scaffold gap loc. mis.","quality":"Less is better","values":[0,0,0,0,0,0],"isMain":false}]],["Unaligned",[]],["Mismatches",[]],["Statistics without reference",[]],["K-mer-based statistics",[]],["Predicted genes",[]],["Similarity statistics",[]],["Reference statistics",[]]],[["Genome statistics",[]],["Reads mapping",[]],["Misassemblies",[{"metricName":"# misassemblies","quality":"Less is better","values":[35,69,51,93,68,26],"isMain":true},{"metricName":" # relocations","quality":"Less is better","values":[1,4,3,2,1,3],"isMain":false},{"metricName":" # translocations","quality":"Less is better","values":[6,7,9,6,9,4],"isMain":false},{"metricName":" # inversions","quality":"Less is better","values":[0,0,4,3,2,0],"isMain":false},{"metricName":" # interspecies translocations","quality":"Less is better","values":[28,58,35,82,56,19],"isMain":false},{"metricName":"# possibly misassembled contigs","quality":"Less is better","values":[0,0,0,0,1,0],"isMain":false},{"metricName":" # possible misassemblies","quality":"Less is better","values":[0,0,0,0,1,0],"isMain":false},{"metricName":"# local misassemblies","quality":"Less is better","values":[6,4,2,5,11,3],"isMain":false},{"metricName":"# scaffold gap ext. mis.","quality":"Less is better","values":[0,0,0,0,0,0],"isMain":false},{"metricName":"# scaffold gap loc. mis.","quality":"Less is better","values":[0,0,0,0,0,0],"isMain":false}]],["Unaligned",[]],["Mismatches",[]],["Statistics without reference",[]],["K-mer-based statistics",[]],["Predicted genes",[]],["Similarity statistics",[]],["Reference statistics",[]]],[["Genome statistics",[]],["Reads mapping",[]],["Misassemblies",[{"metricName":"# misassemblies","quality":"Less is better","values":[42,35,23,47,39,25],"isMain":true},{"metricName":" # relocations","quality":"Less is better","values":[1,1,0,1,2,0],"isMain":false},{"metricName":" # translocations","quality":"Less is better","values":[11,13,14,18,10,4],"isMain":false},{"metricName":" # inversions","quality":"Less is better","values":[0,0,0,1,0,0],"isMain":false},{"metricName":" # interspecies translocations","quality":"Less is better","values":[30,21,9,27,27,21],"isMain":false},{"metricName":"# possibly misassembled contigs","quality":"Less is better","values":[0,0,0,0,0,0],"isMain":false},{"metricName":" # possible misassemblies","quality":"Less is better","values":[0,0,0,0,0,0],"isMain":false},{"metricName":"# local misassemblies","quality":"Less is better","values":[1,2,6,6,6,2],"isMain":false},{"metricName":"# scaffold gap ext. mis.","quality":"Less is better","values":[0,0,0,0,0,0],"isMain":false},{"metricName":"# scaffold gap loc. mis.","quality":"Less is better","values":[0,0,0,0,0,0],"isMain":false}]],["Unaligned",[]],["Mismatches",[]],["Statistics without reference",[]],["K-mer-based statistics",[]],["Predicted genes",[]],["Similarity statistics",[]],["Reference statistics",[]]],[["Genome statistics",[]],["Reads mapping",[]],["Misassemblies",[{"metricName":"# misassemblies","quality":"Less is better","values":[5,7,3,3,5,1],"isMain":true},{"metricName":" # relocations","quality":"Less is better","values":[0,0,0,0,0,0],"isMain":false},{"metricName":" # translocations","quality":"Less is better","values":[0,0,0,0,0,0],"isMain":false},{"metricName":" # inversions","quality":"Less is better","values":[0,0,0,0,0,0],"isMain":false},{"metricName":" # interspecies translocations","quality":"Less is better","values":[5,7,3,3,5,1],"isMain":false},{"metricName":"# possibly misassembled contigs","quality":"Less is better","values":[0,0,0,0,0,0],"isMain":false},{"metricName":" # possible misassemblies","quality":"Less is better","values":[0,0,0,0,0,0],"isMain":false},{"metricName":"# local misassemblies","quality":"Less is better","values":[0,0,0,0,0,0],"isMain":false},{"metricName":"# scaffold gap ext. mis.","quality":"Less is better","values":[0,0,0,0,0,0],"isMain":false},{"metricName":"# scaffold gap loc. mis.","quality":"Less is better","values":[0,0,0,0,0,0],"isMain":false}]],["Unaligned",[]],["Mismatches",[]],["Statistics without reference",[]],["K-mer-based statistics",[]],["Predicted genes",[]],["Similarity statistics",[]],["Reference statistics",[]]],[["Genome statistics",[]],["Reads mapping",[]],["Misassemblies",[{"metricName":"# misassemblies","quality":"Less is better","values":[10,9,3,10,13,4],"isMain":true},{"metricName":" # relocations","quality":"Less is better","values":[0,0,0,0,0,0],"isMain":false},{"metricName":" # translocations","quality":"Less is better","values":[0,0,0,0,0,0],"isMain":false},{"metricName":" # inversions","quality":"Less is better","values":[0,0,0,0,0,0],"isMain":false},{"metricName":" # interspecies translocations","quality":"Less is better","values":[10,9,3,10,13,4],"isMain":false},{"metricName":"# possibly misassembled contigs","quality":"Less is better","values":[0,0,0,0,0,0],"isMain":false},{"metricName":" # possible misassemblies","quality":"Less is better","values":[0,0,0,0,0,0],"isMain":false},{"metricName":"# local misassemblies","quality":"Less is better","values":[0,0,0,0,0,0],"isMain":false},{"metricName":"# scaffold gap ext. mis.","quality":"Less is better","values":[0,0,0,0,0,0],"isMain":false},{"metricName":"# scaffold gap loc. mis.","quality":"Less is better","values":[0,0,0,0,0,0],"isMain":false}]],["Unaligned",[]],["Mismatches",[]],["Statistics without reference",[]],["K-mer-based statistics",[]],["Predicted genes",[]],["Similarity statistics",[]],["Reference statistics",[]]],[["Genome statistics",[]],["Reads mapping",[]],["Misassemblies",[{"metricName":"# misassemblies","quality":"Less is better","values":[16,15,8,22,15,8],"isMain":true},{"metricName":" # relocations","quality":"Less is better","values":[0,0,0,0,0,0],"isMain":false},{"metricName":" # translocations","quality":"Less is better","values":[0,0,1,0,0,1],"isMain":false},{"metricName":" # inversions","quality":"Less is better","values":[0,0,0,0,0,0],"isMain":false},{"metricName":" # interspecies translocations","quality":"Less is better","values":[16,15,7,22,15,7],"isMain":false},{"metricName":"# possibly misassembled contigs","quality":"Less is better","values":[0,0,0,0,0,0],"isMain":false},{"metricName":" # possible misassemblies","quality":"Less is better","values":[0,0,0,0,0,0],"isMain":false},{"metricName":"# local misassemblies","quality":"Less is better","values":[0,0,0,0,1,0],"isMain":false},{"metricName":"# scaffold gap ext. mis.","quality":"Less is better","values":[0,0,0,0,0,0],"isMain":false},{"metricName":"# scaffold gap loc. mis.","quality":"Less is better","values":[0,0,0,0,0,0],"isMain":false}]],["Unaligned",[]],["Mismatches",[]],["Statistics without reference",[]],["K-mer-based statistics",[]],["Predicted genes",[]],["Similarity statistics",[]],["Reference statistics",[]]],[["Genome statistics",[]],["Reads mapping",[]],["Misassemblies",[{"metricName":"# misassemblies","quality":"Less is better","values":[5,7,8,7,10,0],"isMain":true},{"metricName":" # relocations","quality":"Less is better","values":[0,0,0,0,0,0],"isMain":false},{"metricName":" # translocations","quality":"Less is better","values":[0,0,0,0,0,0],"isMain":false},{"metricName":" # inversions","quality":"Less is better","values":[0,0,0,0,0,0],"isMain":false},{"metricName":" # interspecies translocations","quality":"Less is better","values":[5,7,8,7,10,0],"isMain":false},{"metricName":"# possibly misassembled contigs","quality":"Less is better","values":[0,0,1,0,0,0],"isMain":false},{"metricName":" # possible misassemblies","quality":"Less is better","values":[0,0,1,0,0,0],"isMain":false},{"metricName":"# local misassemblies","quality":"Less is better","values":[0,0,0,0,0,0],"isMain":false},{"metricName":"# scaffold gap ext. mis.","quality":"Less is better","values":[0,0,0,0,0,0],"isMain":false},{"metricName":"# scaffold gap loc. mis.","quality":"Less is better","values":[0,0,0,0,0,0],"isMain":false}]],["Unaligned",[]],["Mismatches",[]],["Statistics without reference",[]],["K-mer-based statistics",[]],["Predicted genes",[]],["Similarity statistics",[]],["Reference statistics",[]]],[["Genome statistics",[]],["Reads mapping",[]],["Misassemblies",[{"metricName":"# misassemblies","quality":"Less is better","values":[2,1,0,1,1,0],"isMain":true},{"metricName":" # relocations","quality":"Less is better","values":[0,0,0,0,0,0],"isMain":false},{"metricName":" # translocations","quality":"Less is better","values":[0,0,0,0,0,0],"isMain":false},{"metricName":" # inversions","quality":"Less is better","values":[0,0,0,0,0,0],"isMain":false},{"metricName":" # interspecies translocations","quality":"Less is better","values":[2,1,0,1,1,0],"isMain":false},{"metricName":"# possibly misassembled contigs","quality":"Less is better","values":[0,0,0,0,0,0],"isMain":false},{"metricName":" # possible misassemblies","quality":"Less is better","values":[0,0,0,0,0,0],"isMain":false},{"metricName":"# local misassemblies","quality":"Less is better","values":[0,0,0,0,0,0],"isMain":false},{"metricName":"# scaffold gap ext. mis.","quality":"Less is better","values":[0,0,0,0,0,0],"isMain":false},{"metricName":"# scaffold gap loc. mis.","quality":"Less is better","values":[0,0,0,0,0,0],"isMain":false}]],["Unaligned",[]],["Mismatches",[]],["Statistics without reference",[]],["K-mer-based statistics",[]],["Predicted genes",[]],["Similarity statistics",[]],["Reference statistics",[]]],[["Genome statistics",[]],["Reads mapping",[]],["Misassemblies",[{"metricName":"# misassemblies","quality":"Less is better","values":[22,26,7,27,16,7],"isMain":true},{"metricName":" # relocations","quality":"Less is better","values":[0,0,0,0,0,0],"isMain":false},{"metricName":" # translocations","quality":"Less is better","values":[0,0,1,0,0,0],"isMain":false},{"metricName":" # inversions","quality":"Less is better","values":[0,0,0,0,0,0],"isMain":false},{"metricName":" # interspecies translocations","quality":"Less is better","values":[22,26,6,27,16,7],"isMain":false},{"metricName":"# possibly misassembled contigs","quality":"Less is better","values":[0,0,0,1,0,0],"isMain":false},{"metricName":" # possible misassemblies","quality":"Less is better","values":[0,0,0,1,0,0],"isMain":false},{"metricName":"# local misassemblies","quality":"Less is better","values":[0,1,0,1,0,0],"isMain":false},{"metricName":"# scaffold gap ext. mis.","quality":"Less is better","values":[0,0,0,0,0,0],"isMain":false},{"metricName":"# scaffold gap loc. mis.","quality":"Less is better","values":[0,0,0,0,0,0],"isMain":false}]],["Unaligned",[]],["Mismatches",[]],["Statistics without reference",[]],["K-mer-based statistics",[]],["Predicted genes",[]],["Similarity statistics",[]],["Reference statistics",[]]],[["Genome statistics",[]],["Reads mapping",[]],["Misassemblies",[{"metricName":"# misassemblies","quality":"Less is better","values":[3,3,4,4,4,2],"isMain":true},{"metricName":" # relocations","quality":"Less is better","values":[0,0,0,0,0,0],"isMain":false},{"metricName":" # translocations","quality":"Less is better","values":[0,0,0,0,0,0],"isMain":false},{"metricName":" # inversions","quality":"Less is better","values":[0,0,0,0,0,0],"isMain":false},{"metricName":" # interspecies translocations","quality":"Less is better","values":[3,3,4,4,4,2],"isMain":false},{"metricName":"# possibly misassembled contigs","quality":"Less is better","values":[0,0,0,0,0,0],"isMain":false},{"metricName":" # possible misassemblies","quality":"Less is better","values":[0,0,0,0,0,0],"isMain":false},{"metricName":"# local misassemblies","quality":"Less is better","values":[0,0,1,0,0,0],"isMain":false},{"metricName":"# scaffold gap ext. mis.","quality":"Less is better","values":[0,0,0,0,0,0],"isMain":false},{"metricName":"# scaffold gap loc. mis.","quality":"Less is better","values":[0,0,0,0,0,0],"isMain":false}]],["Unaligned",[]],["Mismatches",[]],["Statistics without reference",[]],["K-mer-based statistics",[]],["Predicted genes",[]],["Similarity statistics",[]],["Reference statistics",[]]],[["Genome statistics",[]],["Reads mapping",[]],["Misassemblies",[{"metricName":"# misassemblies","quality":"Less is better","values":[2,3,4,2,3,1],"isMain":true},{"metricName":" # relocations","quality":"Less is better","values":[0,0,0,0,0,0],"isMain":false},{"metricName":" # translocations","quality":"Less is better","values":[0,0,0,0,0,0],"isMain":false},{"metricName":" # inversions","quality":"Less is better","values":[0,0,0,0,0,0],"isMain":false},{"metricName":" # interspecies translocations","quality":"Less is better","values":[2,3,4,2,3,1],"isMain":false},{"metricName":"# possibly misassembled contigs","quality":"Less is better","values":[0,0,0,1,0,0],"isMain":false},{"metricName":" # possible misassemblies","quality":"Less is better","values":[0,0,0,1,0,0],"isMain":false},{"metricName":"# local misassemblies","quality":"Less is better","values":[0,0,0,0,0,0],"isMain":false},{"metricName":"# scaffold gap ext. mis.","quality":"Less is better","values":[0,0,0,0,0,0],"isMain":false},{"metricName":"# scaffold gap loc. mis.","quality":"Less is better","values":[0,0,0,0,0,0],"isMain":false}]],["Unaligned",[]],["Mismatches",[]],["Statistics without reference",[]],["K-mer-based statistics",[]],["Predicted genes",[]],["Similarity statistics",[]],["Reference statistics",[]]],[["Genome statistics",[]],["Reads mapping",[]],["Misassemblies",[{"metricName":"# misassemblies","quality":"Less is better","values":[13,19,15,22,34,3],"isMain":true},{"metricName":" # relocations","quality":"Less is better","values":[0,0,0,0,0,0],"isMain":false},{"metricName":" # translocations","quality":"Less is better","values":[1,0,2,1,2,0],"isMain":false},{"metricName":" # inversions","quality":"Less is better","values":[1,0,0,1,0,0],"isMain":false},{"metricName":" # interspecies translocations","quality":"Less is better","values":[11,19,13,20,32,3],"isMain":false},{"metricName":"# possibly misassembled contigs","quality":"Less is better","values":[0,0,0,0,0,0],"isMain":false},{"metricName":" # possible misassemblies","quality":"Less is better","values":[0,0,0,0,0,0],"isMain":false},{"metricName":"# local misassemblies","quality":"Less is better","values":[2,2,2,4,5,2],"isMain":false},{"metricName":"# scaffold gap ext. mis.","quality":"Less is better","values":[0,0,0,0,0,0],"isMain":false},{"metricName":"# scaffold gap loc. mis.","quality":"Less is better","values":[0,0,0,0,0,0],"isMain":false}]],["Unaligned",[]],["Mismatches",[]],["Statistics without reference",[]],["K-mer-based statistics",[]],["Predicted genes",[]],["Similarity statistics",[]],["Reference statistics",[]]],[["Genome statistics",[]],["Reads mapping",[]],["Misassemblies",[{"metricName":"# misassemblies","quality":"Less is better","values":[0,0,0,1,0,1],"isMain":true},{"metricName":" # relocations","quality":"Less is better","values":[0,0,0,0,0,0],"isMain":false},{"metricName":" # translocations","quality":"Less is better","values":[0,0,0,0,0,0],"isMain":false},{"metricName":" # inversions","quality":"Less is better","values":[0,0,0,0,0,0],"isMain":false},{"metricName":" # interspecies translocations","quality":"Less is better","values":[0,0,0,1,0,1],"isMain":false},{"metricName":"# possibly misassembled contigs","quality":"Less is better","values":[0,0,0,0,0,0],"isMain":false},{"metricName":" # possible misassemblies","quality":"Less is better","values":[0,0,0,0,0,0],"isMain":false},{"metricName":"# local misassemblies","quality":"Less is better","values":[0,0,0,0,0,0],"isMain":false},{"metricName":"# scaffold gap ext. mis.","quality":"Less is better","values":[0,0,0,0,0,0],"isMain":false},{"metricName":"# scaffold gap loc. mis.","quality":"Less is better","values":[0,0,0,0,0,0],"isMain":false}]],["Unaligned",[]],["Mismatches",[]],["Statistics without reference",[]],["K-mer-based statistics",[]],["Predicted genes",[]],["Similarity statistics",[]],["Reference statistics",[]]],[["Genome statistics",[]],["Reads mapping",[]],["Misassemblies",[{"metricName":"# misassemblies","quality":"Less is better","values":[12,7,5,16,10,6],"isMain":true},{"metricName":" # relocations","quality":"Less is better","values":[0,1,0,0,1,0],"isMain":false},{"metricName":" # translocations","quality":"Less is better","values":[0,0,1,0,0,0],"isMain":false},{"metricName":" # inversions","quality":"Less is better","values":[0,0,0,0,0,0],"isMain":false},{"metricName":" # interspecies translocations","quality":"Less is better","values":[12,6,4,16,9,6],"isMain":false},{"metricName":"# possibly misassembled contigs","quality":"Less is better","values":[0,0,0,0,0,0],"isMain":false},{"metricName":" # possible misassemblies","quality":"Less is better","values":[0,0,0,0,0,0],"isMain":false},{"metricName":"# local misassemblies","quality":"Less is better","values":[2,2,0,1,2,0],"isMain":false},{"metricName":"# scaffold gap ext. mis.","quality":"Less is better","values":[0,0,0,0,0,0],"isMain":false},{"metricName":"# scaffold gap loc. mis.","quality":"Less is better","values":[0,0,0,0,0,0],"isMain":false}]],["Unaligned",[]],["Mismatches",[]],["Statistics without reference",[]],["K-mer-based statistics",[]],["Predicted genes",[]],["Similarity statistics",[]],["Reference statistics",[]]],[["Genome statistics",[]],["Reads mapping",[]],["Misassemblies",[{"metricName":"# misassemblies","quality":"Less is better","values":[76,100,16,63,91,61],"isMain":true},{"metricName":" # relocations","quality":"Less is better","values":[12,21,7,9,21,5],"isMain":false},{"metricName":" # translocations","quality":"Less is better","values":[0,0,0,0,0,0],"isMain":false},{"metricName":" # inversions","quality":"Less is better","values":[2,0,1,0,0,0],"isMain":false},{"metricName":" # interspecies translocations","quality":"Less is better","values":[62,79,8,54,70,56],"isMain":false},{"metricName":"# possibly misassembled contigs","quality":"Less is better","values":[0,1,0,1,0,0],"isMain":false},{"metricName":" # possible misassemblies","quality":"Less is better","values":[0,1,0,1,0,0],"isMain":false},{"metricName":"# local misassemblies","quality":"Less is better","values":[2,3,1,3,4,0],"isMain":false},{"metricName":"# scaffold gap ext. mis.","quality":"Less is better","values":[0,0,0,0,0,0],"isMain":false},{"metricName":"# scaffold gap loc. mis.","quality":"Less is better","values":[0,0,0,0,0,0],"isMain":false}]],["Unaligned",[]],["Mismatches",[]],["Statistics without reference",[]],["K-mer-based statistics",[]],["Predicted genes",[]],["Similarity statistics",[]],["Reference statistics",[]]],[["Genome statistics",[]],["Reads mapping",[]],["Misassemblies",[{"metricName":"# misassemblies","quality":"Less is better","values":[2,4,10,23,5,0],"isMain":true},{"metricName":" # relocations","quality":"Less is better","values":[0,0,0,0,0,0],"isMain":false},{"metricName":" # translocations","quality":"Less is better","values":[0,0,2,2,0,0],"isMain":false},{"metricName":" # inversions","quality":"Less is better","values":[0,0,1,0,0,0],"isMain":false},{"metricName":" # interspecies translocations","quality":"Less is better","values":[2,4,7,21,5,0],"isMain":false},{"metricName":"# possibly misassembled contigs","quality":"Less is better","values":[0,0,0,0,0,0],"isMain":false},{"metricName":" # possible misassemblies","quality":"Less is better","values":[0,0,0,0,0,0],"isMain":false},{"metricName":"# local misassemblies","quality":"Less is better","values":[1,1,3,3,1,1],"isMain":false},{"metricName":"# scaffold gap ext. mis.","quality":"Less is better","values":[0,0,0,0,0,0],"isMain":false},{"metricName":"# scaffold gap loc. mis.","quality":"Less is better","values":[0,0,0,0,0,0],"isMain":false}]],["Unaligned",[]],["Mismatches",[]],["Statistics without reference",[]],["K-mer-based statistics",[]],["Predicted genes",[]],["Similarity statistics",[]],["Reference statistics",[]]],[["Genome statistics",[]],["Reads mapping",[]],["Misassemblies",[{"metricName":"# misassemblies","quality":"Less is better","values":[5,7,1,5,3,7],"isMain":true},{"metricName":" # relocations","quality":"Less is better","values":[0,0,0,0,0,0],"isMain":false},{"metricName":" # translocations","quality":"Less is better","values":[0,0,0,0,0,0],"isMain":false},{"metricName":" # inversions","quality":"Less is better","values":[0,0,0,0,0,0],"isMain":false},{"metricName":" # interspecies translocations","quality":"Less is better","values":[5,7,1,5,3,7],"isMain":false},{"metricName":"# possibly misassembled contigs","quality":"Less is better","values":[0,0,0,0,0,0],"isMain":false},{"metricName":" # possible misassemblies","quality":"Less is better","values":[0,0,0,0,0,0],"isMain":false},{"metricName":"# local misassemblies","quality":"Less is better","values":[0,0,0,0,0,0],"isMain":false},{"metricName":"# scaffold gap ext. mis.","quality":"Less is better","values":[0,0,0,0,0,0],"isMain":false},{"metricName":"# scaffold gap loc. mis.","quality":"Less is better","values":[0,0,0,0,0,0],"isMain":false}]],["Unaligned",[]],["Mismatches",[]],["Statistics without reference",[]],["K-mer-based statistics",[]],["Predicted genes",[]],["Similarity statistics",[]],["Reference statistics",[]]],[["Genome statistics",[]],["Reads mapping",[]],["Misassemblies",[{"metricName":"# misassemblies","quality":"Less is better","values":[43,55,16,30,35,23],"isMain":true},{"metricName":" # relocations","quality":"Less is better","values":[1,0,0,1,0,0],"isMain":false},{"metricName":" # translocations","quality":"Less is better","values":[5,9,5,5,5,2],"isMain":false},{"metricName":" # inversions","quality":"Less is better","values":[2,0,0,0,0,0],"isMain":false},{"metricName":" # interspecies translocations","quality":"Less is better","values":[35,46,11,24,30,21],"isMain":false},{"metricName":"# possibly misassembled contigs","quality":"Less is better","values":[0,0,0,2,0,0],"isMain":false},{"metricName":" # possible misassemblies","quality":"Less is better","values":[0,0,0,2,0,0],"isMain":false},{"metricName":"# local misassemblies","quality":"Less is better","values":[1,3,2,0,2,0],"isMain":false},{"metricName":"# scaffold gap ext. mis.","quality":"Less is better","values":[0,0,0,0,0,0],"isMain":false},{"metricName":"# scaffold gap loc. mis.","quality":"Less is better","values":[0,0,0,0,0,0],"isMain":false}]],["Unaligned",[]],["Mismatches",[]],["Statistics without reference",[]],["K-mer-based statistics",[]],["Predicted genes",[]],["Similarity statistics",[]],["Reference statistics",[]]],[["Genome statistics",[]],["Reads mapping",[]],["Misassemblies",[{"metricName":"# misassemblies","quality":"Less is better","values":[9,19,2,15,15,18],"isMain":true},{"metricName":" # relocations","quality":"Less is better","values":[0,0,0,0,0,0],"isMain":false},{"metricName":" # translocations","quality":"Less is better","values":[0,0,0,1,1,0],"isMain":false},{"metricName":" # inversions","quality":"Less is better","values":[1,0,0,0,1,0],"isMain":false},{"metricName":" # interspecies translocations","quality":"Less is better","values":[8,19,2,14,13,18],"isMain":false},{"metricName":"# possibly misassembled contigs","quality":"Less is better","values":[0,0,0,0,0,0],"isMain":false},{"metricName":" # possible misassemblies","quality":"Less is better","values":[0,0,0,0,0,0],"isMain":false},{"metricName":"# local misassemblies","quality":"Less is better","values":[0,0,0,0,0,0],"isMain":false},{"metricName":"# scaffold gap ext. mis.","quality":"Less is better","values":[0,0,0,0,0,0],"isMain":false},{"metricName":"# scaffold gap loc. mis.","quality":"Less is better","values":[0,0,0,0,0,0],"isMain":false}]],["Unaligned",[]],["Mismatches",[]],["Statistics without reference",[]],["K-mer-based statistics",[]],["Predicted genes",[]],["Similarity statistics",[]],["Reference statistics",[]]],[["Genome statistics",[]],["Reads mapping",[]],["Misassemblies",[{"metricName":"# misassemblies","quality":"Less is better","values":[42,62,15,35,53,30],"isMain":true},{"metricName":" # relocations","quality":"Less is better","values":[1,1,1,1,1,1],"isMain":false},{"metricName":" # translocations","quality":"Less is better","values":[0,0,0,0,1,0],"isMain":false},{"metricName":" # inversions","quality":"Less is better","values":[0,1,0,1,1,0],"isMain":false},{"metricName":" # interspecies translocations","quality":"Less is better","values":[41,60,14,33,50,29],"isMain":false},{"metricName":"# possibly misassembled contigs","quality":"Less is better","values":[0,0,0,1,0,0],"isMain":false},{"metricName":" # possible misassemblies","quality":"Less is better","values":[0,0,0,1,0,0],"isMain":false},{"metricName":"# local misassemblies","quality":"Less is better","values":[0,0,0,0,1,1],"isMain":false},{"metricName":"# scaffold gap ext. mis.","quality":"Less is better","values":[0,0,0,0,0,0],"isMain":false},{"metricName":"# scaffold gap loc. mis.","quality":"Less is better","values":[0,0,0,0,0,0],"isMain":false}]],["Unaligned",[]],["Mismatches",[]],["Statistics without reference",[]],["K-mer-based statistics",[]],["Predicted genes",[]],["Similarity statistics",[]],["Reference statistics",[]]],[["Genome statistics",[]],["Reads mapping",[]],["Misassemblies",[{"metricName":"# misassemblies","quality":"Less is better","values":[92,68,79,155,99,14],"isMain":true},{"metricName":" # relocations","quality":"Less is better","values":[0,1,5,3,5,0],"isMain":false},{"metricName":" # translocations","quality":"Less is better","values":[5,17,21,38,16,9],"isMain":false},{"metricName":" # inversions","quality":"Less is better","values":[8,2,0,2,11,0],"isMain":false},{"metricName":" # interspecies translocations","quality":"Less is better","values":[79,48,53,112,67,5],"isMain":false},{"metricName":"# possibly misassembled contigs","quality":"Less is better","values":[1,0,2,1,0,0],"isMain":false},{"metricName":" # possible misassemblies","quality":"Less is better","values":[1,0,2,1,0,0],"isMain":false},{"metricName":"# local misassemblies","quality":"Less is better","values":[2,7,10,9,14,0],"isMain":false},{"metricName":"# scaffold gap ext. mis.","quality":"Less is better","values":[0,0,0,0,0,0],"isMain":false},{"metricName":"# scaffold gap loc. mis.","quality":"Less is better","values":[0,0,0,0,0,0],"isMain":false}]],["Unaligned",[]],["Mismatches",[]],["Statistics without reference",[]],["K-mer-based statistics",[]],["Predicted genes",[]],["Similarity statistics",[]],["Reference statistics",[]]],[["Genome statistics",[]],["Reads mapping",[]],["Misassemblies",[{"metricName":"# misassemblies","quality":"Less is better","values":[25,10,16,20,11,9],"isMain":true},{"metricName":" # relocations","quality":"Less is better","values":[0,0,0,0,0,1],"isMain":false},{"metricName":" # translocations","quality":"Less is better","values":[7,7,7,9,8,7],"isMain":false},{"metricName":" # inversions","quality":"Less is better","values":[0,0,0,0,0,0],"isMain":false},{"metricName":" # interspecies translocations","quality":"Less is better","values":[18,3,9,11,3,1],"isMain":false},{"metricName":"# possibly misassembled contigs","quality":"Less is better","values":[0,0,0,0,0,0],"isMain":false},{"metricName":" # possible misassemblies","quality":"Less is better","values":[0,0,0,0,0,0],"isMain":false},{"metricName":"# local misassemblies","quality":"Less is better","values":[1,1,1,3,1,0],"isMain":false},{"metricName":"# scaffold gap ext. mis.","quality":"Less is better","values":[0,0,0,0,0,0],"isMain":false},{"metricName":"# scaffold gap loc. mis.","quality":"Less is better","values":[0,0,0,0,0,0],"isMain":false}]],["Unaligned",[]],["Mismatches",[]],["Statistics without reference",[]],["K-mer-based statistics",[]],["Predicted genes",[]],["Similarity statistics",[]],["Reference statistics",[]]],[["Genome statistics",[]],["Reads mapping",[]],["Misassemblies",[{"metricName":"# misassemblies","quality":"Less is better","values":[69,16,7,18,5,3],"isMain":true},{"metricName":" # relocations","quality":"Less is better","values":[0,1,1,0,0,0],"isMain":false},{"metricName":" # translocations","quality":"Less is better","values":[2,8,2,9,2,2],"isMain":false},{"metricName":" # inversions","quality":"Less is better","values":[1,1,0,0,2,0],"isMain":false},{"metricName":" # interspecies translocations","quality":"Less is better","values":[66,6,4,9,1,1],"isMain":false},{"metricName":"# possibly misassembled contigs","quality":"Less is better","values":[0,0,0,0,0,0],"isMain":false},{"metricName":" # possible misassemblies","quality":"Less is better","values":[0,0,0,0,0,0],"isMain":false},{"metricName":"# local misassemblies","quality":"Less is better","values":[1,3,0,1,1,0],"isMain":false},{"metricName":"# scaffold gap ext. mis.","quality":"Less is better","values":[0,0,0,0,0,0],"isMain":false},{"metricName":"# scaffold gap loc. mis.","quality":"Less is better","values":[0,0,0,0,0,0],"isMain":false}]],["Unaligned",[]],["Mismatches",[]],["Statistics without reference",[]],["K-mer-based statistics",[]],["Predicted genes",[]],["Similarity statistics",[]],["Reference statistics",[]]],[["Genome statistics",[]],["Reads mapping",[]],["Misassemblies",[{"metricName":"# misassemblies","quality":"Less is better","values":[10,20,24,26,23,5],"isMain":true},{"metricName":" # relocations","quality":"Less is better","values":[1,4,2,5,5,0],"isMain":false},{"metricName":" # translocations","quality":"Less is better","values":[0,0,0,0,0,0],"isMain":false},{"metricName":" # inversions","quality":"Less is better","values":[0,0,2,1,1,0],"isMain":false},{"metricName":" # interspecies translocations","quality":"Less is better","values":[9,16,20,20,17,5],"isMain":false},{"metricName":"# possibly misassembled contigs","quality":"Less is better","values":[0,0,0,0,0,0],"isMain":false},{"metricName":" # possible misassemblies","quality":"Less is better","values":[0,0,0,0,0,0],"isMain":false},{"metricName":"# local misassemblies","quality":"Less is better","values":[5,5,3,5,1,0],"isMain":false},{"metricName":"# scaffold gap ext. mis.","quality":"Less is better","values":[0,0,0,0,0,0],"isMain":false},{"metricName":"# scaffold gap loc. mis.","quality":"Less is better","values":[0,0,0,0,0,0],"isMain":false}]],["Unaligned",[]],["Mismatches",[]],["Statistics without reference",[]],["K-mer-based statistics",[]],["Predicted genes",[]],["Similarity statistics",[]],["Reference statistics",[]]],[["Genome statistics",[]],["Reads mapping",[]],["Misassemblies",[{"metricName":"# misassemblies","quality":"Less is better","values":[6,8,3,11,11,4],"isMain":true},{"metricName":" # relocations","quality":"Less is better","values":[1,0,0,1,0,0],"isMain":false},{"metricName":" # translocations","quality":"Less is better","values":[0,1,0,0,0,0],"isMain":false},{"metricName":" # inversions","quality":"Less is better","values":[0,0,0,0,0,0],"isMain":false},{"metricName":" # interspecies translocations","quality":"Less is better","values":[5,7,3,10,11,4],"isMain":false},{"metricName":"# possibly misassembled contigs","quality":"Less is better","values":[0,0,0,0,0,0],"isMain":false},{"metricName":" # possible misassemblies","quality":"Less is better","values":[0,0,0,0,0,0],"isMain":false},{"metricName":"# local misassemblies","quality":"Less is better","values":[0,0,0,1,0,1],"isMain":false},{"metricName":"# scaffold gap ext. mis.","quality":"Less is better","values":[0,0,0,0,0,0],"isMain":false},{"metricName":"# scaffold gap loc. mis.","quality":"Less is better","values":[0,0,0,0,0,0],"isMain":false}]],["Unaligned",[]],["Mismatches",[]],["Statistics without reference",[]],["K-mer-based statistics",[]],["Predicted genes",[]],["Similarity statistics",[]],["Reference statistics",[]]],[["Genome statistics",[]],["Reads mapping",[]],["Misassemblies",[{"metricName":"# misassemblies","quality":"Less is better","values":[122,36,32,73,36,1],"isMain":true},{"metricName":" # relocations","quality":"Less is better","values":[0,10,12,19,11,0],"isMain":false},{"metricName":" # translocations","quality":"Less is better","values":[0,0,0,0,0,0],"isMain":false},{"metricName":" # inversions","quality":"Less is better","values":[7,1,1,6,4,0],"isMain":false},{"metricName":" # interspecies translocations","quality":"Less is better","values":[115,25,19,48,21,1],"isMain":false},{"metricName":"# possibly misassembled contigs","quality":"Less is better","values":[1,0,0,0,0,0],"isMain":false},{"metricName":" # possible misassemblies","quality":"Less is better","values":[1,0,0,0,0,0],"isMain":false},{"metricName":"# local misassemblies","quality":"Less is better","values":[4,0,4,1,11,0],"isMain":false},{"metricName":"# scaffold gap ext. mis.","quality":"Less is better","values":[0,0,0,0,0,0],"isMain":false},{"metricName":"# scaffold gap loc. mis.","quality":"Less is better","values":[0,0,0,0,0,0],"isMain":false}]],["Unaligned",[]],["Mismatches",[]],["Statistics without reference",[]],["K-mer-based statistics",[]],["Predicted genes",[]],["Similarity statistics",[]],["Reference statistics",[]]],[["Genome statistics",[]],["Reads mapping",[]],["Misassemblies",[{"metricName":"# misassemblies","quality":"Less is better","values":[3,3,3,9,12,3],"isMain":true},{"metricName":" # relocations","quality":"Less is better","values":[0,0,1,0,0,0],"isMain":false},{"metricName":" # translocations","quality":"Less is better","values":[0,0,0,0,0,0],"isMain":false},{"metricName":" # inversions","quality":"Less is better","values":[0,0,0,0,1,0],"isMain":false},{"metricName":" # interspecies translocations","quality":"Less is better","values":[3,3,2,9,11,3],"isMain":false},{"metricName":"# possibly misassembled contigs","quality":"Less is better","values":[0,0,0,0,0,0],"isMain":false},{"metricName":" # possible misassemblies","quality":"Less is better","values":[0,0,0,0,0,0],"isMain":false},{"metricName":"# local misassemblies","quality":"Less is better","values":[0,0,0,0,0,0],"isMain":false},{"metricName":"# scaffold gap ext. mis.","quality":"Less is better","values":[0,0,0,0,0,0],"isMain":false},{"metricName":"# scaffold gap loc. mis.","quality":"Less is better","values":[0,0,0,0,0,0],"isMain":false}]],["Unaligned",[]],["Mismatches",[]],["Statistics without reference",[]],["K-mer-based statistics",[]],["Predicted genes",[]],["Similarity statistics",[]],["Reference statistics",[]]],[["Genome statistics",[]],["Reads mapping",[]],["Misassemblies",[{"metricName":"# misassemblies","quality":"Less is better","values":[28,34,14,33,34,16],"isMain":true},{"metricName":" # relocations","quality":"Less is better","values":[0,1,0,0,0,0],"isMain":false},{"metricName":" # translocations","quality":"Less is better","values":[2,1,1,0,0,0],"isMain":false},{"metricName":" # inversions","quality":"Less is better","values":[0,1,0,1,0,0],"isMain":false},{"metricName":" # interspecies translocations","quality":"Less is better","values":[26,31,13,32,34,16],"isMain":false},{"metricName":"# possibly misassembled contigs","quality":"Less is better","values":[0,1,0,0,1,0],"isMain":false},{"metricName":" # possible misassemblies","quality":"Less is better","values":[0,1,0,0,1,0],"isMain":false},{"metricName":"# local misassemblies","quality":"Less is better","values":[0,1,1,1,2,1],"isMain":false},{"metricName":"# scaffold gap ext. mis.","quality":"Less is better","values":[0,0,0,0,0,0],"isMain":false},{"metricName":"# scaffold gap loc. mis.","quality":"Less is better","values":[0,0,0,0,0,0],"isMain":false}]],["Unaligned",[]],["Mismatches",[]],["Statistics without reference",[]],["K-mer-based statistics",[]],["Predicted genes",[]],["Similarity statistics",[]],["Reference statistics",[]]],[["Genome statistics",[]],["Reads mapping",[]],["Misassemblies",[{"metricName":"# misassemblies","quality":"Less is better","values":[24,24,16,33,28,5],"isMain":true},{"metricName":" # relocations","quality":"Less is better","values":[0,1,0,0,1,0],"isMain":false},{"metricName":" # translocations","quality":"Less is better","values":[3,2,0,1,3,1],"isMain":false},{"metricName":" # inversions","quality":"Less is better","values":[0,0,1,1,0,0],"isMain":false},{"metricName":" # interspecies translocations","quality":"Less is better","values":[21,21,15,31,24,4],"isMain":false},{"metricName":"# possibly misassembled contigs","quality":"Less is better","values":[0,0,0,1,0,0],"isMain":false},{"metricName":" # possible misassemblies","quality":"Less is better","values":[0,0,0,1,0,0],"isMain":false},{"metricName":"# local misassemblies","quality":"Less is better","values":[1,2,0,1,1,0],"isMain":false},{"metricName":"# scaffold gap ext. mis.","quality":"Less is better","values":[0,0,0,0,0,0],"isMain":false},{"metricName":"# scaffold gap loc. mis.","quality":"Less is better","values":[0,0,0,0,0,0],"isMain":false}]],["Unaligned",[]],["Mismatches",[]],["Statistics without reference",[]],["K-mer-based statistics",[]],["Predicted genes",[]],["Similarity statistics",[]],["Reference statistics",[]]],[["Genome statistics",[]],["Reads mapping",[]],["Misassemblies",[{"metricName":"# misassemblies","quality":"Less is better","values":[19,18,13,8,11,0],"isMain":true},{"metricName":" # relocations","quality":"Less is better","values":[0,0,0,0,0,0],"isMain":false},{"metricName":" # translocations","quality":"Less is better","values":[0,0,0,0,0,0],"isMain":false},{"metricName":" # inversions","quality":"Less is better","values":[0,1,0,0,0,0],"isMain":false},{"metricName":" # interspecies translocations","quality":"Less is better","values":[19,17,13,8,11,0],"isMain":false},{"metricName":"# possibly misassembled contigs","quality":"Less is better","values":[0,0,0,0,0,0],"isMain":false},{"metricName":" # possible misassemblies","quality":"Less is better","values":[0,0,0,0,0,0],"isMain":false},{"metricName":"# local misassemblies","quality":"Less is better","values":[0,0,0,0,0,0],"isMain":false},{"metricName":"# scaffold gap ext. mis.","quality":"Less is better","values":[0,0,0,0,0,0],"isMain":false},{"metricName":"# scaffold gap loc. mis.","quality":"Less is better","values":[0,0,0,0,0,0],"isMain":false}]],["Unaligned",[]],["Mismatches",[]],["Statistics without reference",[]],["K-mer-based statistics",[]],["Predicted genes",[]],["Similarity statistics",[]],["Reference statistics",[]]],[["Genome statistics",[]],["Reads mapping",[]],["Misassemblies",[{"metricName":"# misassemblies","quality":"Less is better","values":[19,7,7,9,2,1],"isMain":true},{"metricName":" # relocations","quality":"Less is better","values":[0,0,0,0,0,0],"isMain":false},{"metricName":" # translocations","quality":"Less is better","values":[0,0,0,0,0,0],"isMain":false},{"metricName":" # inversions","quality":"Less is better","values":[0,1,0,0,0,0],"isMain":false},{"metricName":" # interspecies translocations","quality":"Less is better","values":[19,6,7,9,2,1],"isMain":false},{"metricName":"# possibly misassembled contigs","quality":"Less is better","values":[0,0,0,0,0,0],"isMain":false},{"metricName":" # possible misassemblies","quality":"Less is better","values":[0,0,0,0,0,0],"isMain":false},{"metricName":"# local misassemblies","quality":"Less is better","values":[0,1,0,1,0,0],"isMain":false},{"metricName":"# scaffold gap ext. mis.","quality":"Less is better","values":[0,0,0,0,0,0],"isMain":false},{"metricName":"# scaffold gap loc. mis.","quality":"Less is better","values":[0,0,0,0,0,0],"isMain":false}]],["Unaligned",[]],["Mismatches",[]],["Statistics without reference",[]],["K-mer-based statistics",[]],["Predicted genes",[]],["Similarity statistics",[]],["Reference statistics",[]]],[["Genome statistics",[]],["Reads mapping",[]],["Misassemblies",[{"metricName":"# misassemblies","quality":"Less is better","values":[52,67,24,40,68,29],"isMain":true},{"metricName":" # relocations","quality":"Less is better","values":[0,0,0,1,2,0],"isMain":false},{"metricName":" # translocations","quality":"Less is better","values":[3,2,1,1,3,1],"isMain":false},{"metricName":" # inversions","quality":"Less is better","values":[0,0,0,1,0,0],"isMain":false},{"metricName":" # interspecies translocations","quality":"Less is better","values":[49,65,23,37,63,28],"isMain":false},{"metricName":"# possibly misassembled contigs","quality":"Less is better","values":[0,0,0,0,0,0],"isMain":false},{"metricName":" # possible misassemblies","quality":"Less is better","values":[0,0,0,0,0,0],"isMain":false},{"metricName":"# local misassemblies","quality":"Less is better","values":[0,1,1,1,1,0],"isMain":false},{"metricName":"# scaffold gap ext. mis.","quality":"Less is better","values":[0,0,0,0,0,0],"isMain":false},{"metricName":"# scaffold gap loc. mis.","quality":"Less is better","values":[0,0,0,0,0,0],"isMain":false}]],["Unaligned",[]],["Mismatches",[]],["Statistics without reference",[]],["K-mer-based statistics",[]],["Predicted genes",[]],["Similarity statistics",[]],["Reference statistics",[]]],[["Genome statistics",[]],["Reads mapping",[]],["Misassemblies",[{"metricName":"# misassemblies","quality":"Less is better","values":[9,9,9,11,19,4],"isMain":true},{"metricName":" # relocations","quality":"Less is better","values":[0,0,1,1,0,0],"isMain":false},{"metricName":" # translocations","quality":"Less is better","values":[0,0,0,0,0,0],"isMain":false},{"metricName":" # inversions","quality":"Less is better","values":[0,0,0,0,0,0],"isMain":false},{"metricName":" # interspecies translocations","quality":"Less is better","values":[9,9,8,10,19,4],"isMain":false},{"metricName":"# possibly misassembled contigs","quality":"Less is better","values":[0,0,0,0,0,0],"isMain":false},{"metricName":" # possible misassemblies","quality":"Less is better","values":[0,0,0,0,0,0],"isMain":false},{"metricName":"# local misassemblies","quality":"Less is better","values":[0,0,0,1,0,0],"isMain":false},{"metricName":"# scaffold gap ext. mis.","quality":"Less is better","values":[0,0,0,0,0,0],"isMain":false},{"metricName":"# scaffold gap loc. mis.","quality":"Less is better","values":[0,0,0,0,0,0],"isMain":false}]],["Unaligned",[]],["Mismatches",[]],["Statistics without reference",[]],["K-mer-based statistics",[]],["Predicted genes",[]],["Similarity statistics",[]],["Reference statistics",[]]],[["Genome statistics",[]],["Reads mapping",[]],["Misassemblies",[{"metricName":"# misassemblies","quality":"Less is better","values":[21,18,18,30,18,13],"isMain":true},{"metricName":" # relocations","quality":"Less is better","values":[0,0,0,0,0,0],"isMain":false},{"metricName":" # translocations","quality":"Less is better","values":[0,1,0,0,1,0],"isMain":false},{"metricName":" # inversions","quality":"Less is better","values":[0,0,1,0,0,0],"isMain":false},{"metricName":" # interspecies translocations","quality":"Less is better","values":[21,17,17,30,17,13],"isMain":false},{"metricName":"# possibly misassembled contigs","quality":"Less is better","values":[0,0,0,0,0,0],"isMain":false},{"metricName":" # possible misassemblies","quality":"Less is better","values":[0,0,0,0,0,0],"isMain":false},{"metricName":"# local misassemblies","quality":"Less is better","values":[0,0,0,1,0,0],"isMain":false},{"metricName":"# scaffold gap ext. mis.","quality":"Less is better","values":[0,0,0,0,0,0],"isMain":false},{"metricName":"# scaffold gap loc. mis.","quality":"Less is better","values":[0,0,0,0,0,0],"isMain":false}]],["Unaligned",[]],["Mismatches",[]],["Statistics without reference",[]],["K-mer-based statistics",[]],["Predicted genes",[]],["Similarity statistics",[]],["Reference statistics",[]]],[["Genome statistics",[]],["Reads mapping",[]],["Misassemblies",[{"metricName":"# misassemblies","quality":"Less is better","values":[1,9,5,5,4,1],"isMain":true},{"metricName":" # relocations","quality":"Less is better","values":[0,0,0,0,0,0],"isMain":false},{"metricName":" # translocations","quality":"Less is better","values":[0,0,0,0,0,0],"isMain":false},{"metricName":" # inversions","quality":"Less is better","values":[0,0,0,0,0,0],"isMain":false},{"metricName":" # interspecies translocations","quality":"Less is better","values":[1,9,5,5,4,1],"isMain":false},{"metricName":"# possibly misassembled contigs","quality":"Less is better","values":[0,0,0,0,0,0],"isMain":false},{"metricName":" # possible misassemblies","quality":"Less is better","values":[0,0,0,0,0,0],"isMain":false},{"metricName":"# local misassemblies","quality":"Less is better","values":[0,1,0,2,0,0],"isMain":false},{"metricName":"# scaffold gap ext. mis.","quality":"Less is better","values":[0,0,0,0,0,0],"isMain":false},{"metricName":"# scaffold gap loc. mis.","quality":"Less is better","values":[0,0,0,0,0,0],"isMain":false}]],["Unaligned",[]],["Mismatches",[]],["Statistics without reference",[]],["K-mer-based statistics",[]],["Predicted genes",[]],["Similarity statistics",[]],["Reference statistics",[]]],[["Genome statistics",[]],["Reads mapping",[]],["Misassemblies",[{"metricName":"# misassemblies","quality":"Less is better","values":[42,59,37,47,53,18],"isMain":true},{"metricName":" # relocations","quality":"Less is better","values":[2,2,0,0,2,1],"isMain":false},{"metricName":" # translocations","quality":"Less is better","values":[2,3,1,0,2,0],"isMain":false},{"metricName":" # inversions","quality":"Less is better","values":[0,1,2,1,0,0],"isMain":false},{"metricName":" # interspecies translocations","quality":"Less is better","values":[38,53,34,46,49,17],"isMain":false},{"metricName":"# possibly misassembled contigs","quality":"Less is better","values":[0,1,0,0,1,0],"isMain":false},{"metricName":" # possible misassemblies","quality":"Less is better","values":[0,1,0,0,1,0],"isMain":false},{"metricName":"# local misassemblies","quality":"Less is better","values":[0,1,1,3,1,0],"isMain":false},{"metricName":"# scaffold gap ext. mis.","quality":"Less is better","values":[0,0,0,0,0,0],"isMain":false},{"metricName":"# scaffold gap loc. mis.","quality":"Less is better","values":[0,0,0,0,0,0],"isMain":false}]],["Unaligned",[]],["Mismatches",[]],["Statistics without reference",[]],["K-mer-based statistics",[]],["Predicted genes",[]],["Similarity statistics",[]],["Reference statistics",[]]],[["Genome statistics",[]],["Reads mapping",[]],["Misassemblies",[{"metricName":"# misassemblies","quality":"Less is better","values":[26,11,15,12,13,2],"isMain":true},{"metricName":" # relocations","quality":"Less is better","values":[0,0,0,0,0,0],"isMain":false},{"metricName":" # translocations","quality":"Less is better","values":[1,0,0,0,1,0],"isMain":false},{"metricName":" # inversions","quality":"Less is better","values":[0,0,0,0,0,0],"isMain":false},{"metricName":" # interspecies translocations","quality":"Less is better","values":[25,11,15,12,12,2],"isMain":false},{"metricName":"# possibly misassembled contigs","quality":"Less is better","values":[0,0,0,0,0,0],"isMain":false},{"metricName":" # possible misassemblies","quality":"Less is better","values":[0,0,0,0,0,0],"isMain":false},{"metricName":"# local misassemblies","quality":"Less is better","values":[1,0,1,0,1,0],"isMain":false},{"metricName":"# scaffold gap ext. mis.","quality":"Less is better","values":[0,0,0,0,0,0],"isMain":false},{"metricName":"# scaffold gap loc. mis.","quality":"Less is better","values":[0,0,0,0,0,0],"isMain":false}]],["Unaligned",[]],["Mismatches",[]],["Statistics without reference",[]],["K-mer-based statistics",[]],["Predicted genes",[]],["Similarity statistics",[]],["Reference statistics",[]]],[["Genome statistics",[]],["Reads mapping",[]],["Misassemblies",[{"metricName":"# misassemblies","quality":"Less is better","values":[2,1,1,3,0,0],"isMain":true},{"metricName":" # relocations","quality":"Less is better","values":[0,0,0,0,0,0],"isMain":false},{"metricName":" # translocations","quality":"Less is better","values":[0,0,0,0,0,0],"isMain":false},{"metricName":" # inversions","quality":"Less is better","values":[0,0,0,0,0,0],"isMain":false},{"metricName":" # interspecies translocations","quality":"Less is better","values":[2,1,1,3,0,0],"isMain":false},{"metricName":"# possibly misassembled contigs","quality":"Less is better","values":[0,0,0,0,0,0],"isMain":false},{"metricName":" # possible misassemblies","quality":"Less is better","values":[0,0,0,0,0,0],"isMain":false},{"metricName":"# local misassemblies","quality":"Less is better","values":[0,0,0,0,0,0],"isMain":false},{"metricName":"# scaffold gap ext. mis.","quality":"Less is better","values":[0,0,0,0,0,0],"isMain":false},{"metricName":"# scaffold gap loc. mis.","quality":"Less is better","values":[0,0,0,0,0,0],"isMain":false}]],["Unaligned",[]],["Mismatches",[]],["Statistics without reference",[]],["K-mer-based statistics",[]],["Predicted genes",[]],["Similarity statistics",[]],["Reference statistics",[]]],[["Genome statistics",[]],["Reads mapping",[]],["Misassemblies",[{"metricName":"# misassemblies","quality":"Less is better","values":[72,79,32,74,78,33],"isMain":true},{"metricName":" # relocations","quality":"Less is better","values":[0,1,0,0,1,0],"isMain":false},{"metricName":" # translocations","quality":"Less is better","values":[2,1,2,2,2,1],"isMain":false},{"metricName":" # inversions","quality":"Less is better","values":[2,2,0,3,5,0],"isMain":false},{"metricName":" # interspecies translocations","quality":"Less is better","values":[68,75,30,69,70,32],"isMain":false},{"metricName":"# possibly misassembled contigs","quality":"Less is better","values":[0,1,0,0,0,0],"isMain":false},{"metricName":" # possible misassemblies","quality":"Less is better","values":[0,1,0,0,0,0],"isMain":false},{"metricName":"# local misassemblies","quality":"Less is better","values":[1,0,6,3,3,0],"isMain":false},{"metricName":"# scaffold gap ext. mis.","quality":"Less is better","values":[0,0,0,0,0,0],"isMain":false},{"metricName":"# scaffold gap loc. mis.","quality":"Less is better","values":[0,0,0,0,0,0],"isMain":false}]],["Unaligned",[]],["Mismatches",[]],["Statistics without reference",[]],["K-mer-based statistics",[]],["Predicted genes",[]],["Similarity statistics",[]],["Reference statistics",[]]],[["Genome statistics",[]],["Reads mapping",[]],["Misassemblies",[{"metricName":"# misassemblies","quality":"Less is better","values":[22,11,9,10,5,0],"isMain":true},{"metricName":" # relocations","quality":"Less is better","values":[0,0,0,0,0,0],"isMain":false},{"metricName":" # translocations","quality":"Less is better","values":[0,0,0,0,0,0],"isMain":false},{"metricName":" # inversions","quality":"Less is better","values":[0,0,0,0,0,0],"isMain":false},{"metricName":" # interspecies translocations","quality":"Less is better","values":[22,11,9,10,5,0],"isMain":false},{"metricName":"# possibly misassembled contigs","quality":"Less is better","values":[0,0,0,0,0,0],"isMain":false},{"metricName":" # possible misassemblies","quality":"Less is better","values":[0,0,0,0,0,0],"isMain":false},{"metricName":"# local misassemblies","quality":"Less is better","values":[0,1,0,0,0,0],"isMain":false},{"metricName":"# scaffold gap ext. mis.","quality":"Less is better","values":[0,0,0,0,0,0],"isMain":false},{"metricName":"# scaffold gap loc. mis.","quality":"Less is better","values":[0,0,0,0,0,0],"isMain":false}]],["Unaligned",[]],["Mismatches",[]],["Statistics without reference",[]],["K-mer-based statistics",[]],["Predicted genes",[]],["Similarity statistics",[]],["Reference statistics",[]]],[["Genome statistics",[]],["Reads mapping",[]],["Misassemblies",[{"metricName":"# misassemblies","quality":"Less is better","values":[12,28,12,15,15,9],"isMain":true},{"metricName":" # relocations","quality":"Less is better","values":[0,0,0,0,0,0],"isMain":false},{"metricName":" # translocations","quality":"Less is better","values":[0,2,1,0,0,0],"isMain":false},{"metricName":" # inversions","quality":"Less is better","values":[0,0,0,1,0,0],"isMain":false},{"metricName":" # interspecies translocations","quality":"Less is better","values":[12,26,11,14,15,9],"isMain":false},{"metricName":"# possibly misassembled contigs","quality":"Less is better","values":[0,1,0,0,0,0],"isMain":false},{"metricName":" # possible misassemblies","quality":"Less is better","values":[0,1,0,0,0,0],"isMain":false},{"metricName":"# local misassemblies","quality":"Less is better","values":[0,0,0,0,0,0],"isMain":false},{"metricName":"# scaffold gap ext. mis.","quality":"Less is better","values":[0,0,0,0,0,0],"isMain":false},{"metricName":"# scaffold gap loc. mis.","quality":"Less is better","values":[0,0,0,0,0,0],"isMain":false}]],["Unaligned",[]],["Mismatches",[]],["Statistics without reference",[]],["K-mer-based statistics",[]],["Predicted genes",[]],["Similarity statistics",[]],["Reference statistics",[]]],[["Genome statistics",[]],["Reads mapping",[]],["Misassemblies",[{"metricName":"# misassemblies","quality":"Less is better","values":[1,7,4,7,7,2],"isMain":true},{"metricName":" # relocations","quality":"Less is better","values":[0,0,0,0,0,0],"isMain":false},{"metricName":" # translocations","quality":"Less is better","values":[0,0,0,0,0,0],"isMain":false},{"metricName":" # inversions","quality":"Less is better","values":[0,0,0,0,0,0],"isMain":false},{"metricName":" # interspecies translocations","quality":"Less is better","values":[1,7,4,7,7,2],"isMain":false},{"metricName":"# possibly misassembled contigs","quality":"Less is better","values":[0,0,0,0,0,0],"isMain":false},{"metricName":" # possible misassemblies","quality":"Less is better","values":[0,0,0,0,0,0],"isMain":false},{"metricName":"# local misassemblies","quality":"Less is better","values":[0,0,0,0,0,0],"isMain":false},{"metricName":"# scaffold gap ext. mis.","quality":"Less is better","values":[0,0,0,0,0,0],"isMain":false},{"metricName":"# scaffold gap loc. mis.","quality":"Less is better","values":[0,0,0,0,0,0],"isMain":false}]],["Unaligned",[]],["Mismatches",[]],["Statistics without reference",[]],["K-mer-based statistics",[]],["Predicted genes",[]],["Similarity statistics",[]],["Reference statistics",[]]],[["Genome statistics",[]],["Reads mapping",[]],["Misassemblies",[{"metricName":"# misassemblies","quality":"Less is better","values":[27,35,9,28,33,14],"isMain":true},{"metricName":" # relocations","quality":"Less is better","values":[0,0,0,0,0,0],"isMain":false},{"metricName":" # translocations","quality":"Less is better","values":[0,2,0,0,1,0],"isMain":false},{"metricName":" # inversions","quality":"Less is better","values":[0,0,0,0,0,0],"isMain":false},{"metricName":" # interspecies translocations","quality":"Less is better","values":[27,33,9,28,32,14],"isMain":false},{"metricName":"# possibly misassembled contigs","quality":"Less is better","values":[0,0,0,0,0,0],"isMain":false},{"metricName":" # possible misassemblies","quality":"Less is better","values":[0,0,0,0,0,0],"isMain":false},{"metricName":"# local misassemblies","quality":"Less is better","values":[0,0,1,0,0,1],"isMain":false},{"metricName":"# scaffold gap ext. mis.","quality":"Less is better","values":[0,0,0,0,0,0],"isMain":false},{"metricName":"# scaffold gap loc. mis.","quality":"Less is better","values":[0,0,0,0,0,0],"isMain":false}]],["Unaligned",[]],["Mismatches",[]],["Statistics without reference",[]],["K-mer-based statistics",[]],["Predicted genes",[]],["Similarity statistics",[]],["Reference statistics",[]]],[["Genome statistics",[]],["Reads mapping",[]],["Misassemblies",[{"metricName":"# misassemblies","quality":"Less is better","values":[91,131,62,99,133,48],"isMain":true},{"metricName":" # relocations","quality":"Less is better","values":[1,5,1,0,3,1],"isMain":false},{"metricName":" # translocations","quality":"Less is better","values":[6,6,3,8,4,2],"isMain":false},{"metricName":" # inversions","quality":"Less is better","values":[1,0,0,3,2,0],"isMain":false},{"metricName":" # interspecies translocations","quality":"Less is better","values":[83,120,58,88,124,45],"isMain":false},{"metricName":"# possibly misassembled contigs","quality":"Less is better","values":[0,4,1,2,4,1],"isMain":false},{"metricName":" # possible misassemblies","quality":"Less is better","values":[0,4,1,2,4,1],"isMain":false},{"metricName":"# local misassemblies","quality":"Less is better","values":[5,3,3,2,8,1],"isMain":false},{"metricName":"# scaffold gap ext. mis.","quality":"Less is better","values":[0,0,0,0,0,0],"isMain":false},{"metricName":"# scaffold gap loc. mis.","quality":"Less is better","values":[0,0,0,0,0,0],"isMain":false}]],["Unaligned",[]],["Mismatches",[]],["Statistics without reference",[]],["K-mer-based statistics",[]],["Predicted genes",[]],["Similarity statistics",[]],["Reference statistics",[]]],[["Genome statistics",[]],["Reads mapping",[]],["Misassemblies",[{"metricName":"# misassemblies","quality":"Less is better","values":[13,24,6,17,20,12],"isMain":true},{"metricName":" # relocations","quality":"Less is better","values":[0,0,0,0,0,0],"isMain":false},{"metricName":" # translocations","quality":"Less is better","values":[0,0,0,0,0,0],"isMain":false},{"metricName":" # inversions","quality":"Less is better","values":[0,0,1,0,0,0],"isMain":false},{"metricName":" # interspecies translocations","quality":"Less is better","values":[13,24,5,17,20,12],"isMain":false},{"metricName":"# possibly misassembled contigs","quality":"Less is better","values":[1,0,0,0,0,0],"isMain":false},{"metricName":" # possible misassemblies","quality":"Less is better","values":[1,0,0,0,0,0],"isMain":false},{"metricName":"# local misassemblies","quality":"Less is better","values":[0,0,1,1,0,0],"isMain":false},{"metricName":"# scaffold gap ext. mis.","quality":"Less is better","values":[0,0,0,0,0,0],"isMain":false},{"metricName":"# scaffold gap loc. mis.","quality":"Less is better","values":[0,0,0,0,0,0],"isMain":false}]],["Unaligned",[]],["Mismatches",[]],["Statistics without reference",[]],["K-mer-based statistics",[]],["Predicted genes",[]],["Similarity statistics",[]],["Reference statistics",[]]],[["Genome statistics",[]],["Reads mapping",[]],["Misassemblies",[{"metricName":"# misassemblies","quality":"Less is better","values":[16,24,8,15,16,6],"isMain":true},{"metricName":" # relocations","quality":"Less is better","values":[0,0,0,0,0,0],"isMain":false},{"metricName":" # translocations","quality":"Less is better","values":[1,3,0,0,0,0],"isMain":false},{"metricName":" # inversions","quality":"Less is better","values":[0,0,0,0,0,0],"isMain":false},{"metricName":" # interspecies translocations","quality":"Less is better","values":[15,21,8,15,16,6],"isMain":false},{"metricName":"# possibly misassembled contigs","quality":"Less is better","values":[1,1,0,0,0,0],"isMain":false},{"metricName":" # possible misassemblies","quality":"Less is better","values":[1,1,0,0,0,0],"isMain":false},{"metricName":"# local misassemblies","quality":"Less is better","values":[2,0,1,0,1,0],"isMain":false},{"metricName":"# scaffold gap ext. mis.","quality":"Less is better","values":[0,0,0,0,0,0],"isMain":false},{"metricName":"# scaffold gap loc. mis.","quality":"Less is better","values":[0,0,0,0,0,0],"isMain":false}]],["Unaligned",[]],["Mismatches",[]],["Statistics without reference",[]],["K-mer-based statistics",[]],["Predicted genes",[]],["Similarity statistics",[]],["Reference statistics",[]]],[["Genome statistics",[]],["Reads mapping",[]],["Misassemblies",[{"metricName":"# misassemblies","quality":"Less is better","values":[27,31,18,25,39,17],"isMain":true},{"metricName":" # relocations","quality":"Less is better","values":[0,0,0,0,0,0],"isMain":false},{"metricName":" # translocations","quality":"Less is better","values":[0,0,0,1,0,1],"isMain":false},{"metricName":" # inversions","quality":"Less is better","values":[0,1,0,0,0,0],"isMain":false},{"metricName":" # interspecies translocations","quality":"Less is better","values":[27,30,18,24,39,16],"isMain":false},{"metricName":"# possibly misassembled contigs","quality":"Less is better","values":[0,0,2,0,1,0],"isMain":false},{"metricName":" # possible misassemblies","quality":"Less is better","values":[0,0,2,0,1,0],"isMain":false},{"metricName":"# local misassemblies","quality":"Less is better","values":[1,0,0,1,0,0],"isMain":false},{"metricName":"# scaffold gap ext. mis.","quality":"Less is better","values":[0,0,0,0,0,0],"isMain":false},{"metricName":"# scaffold gap loc. mis.","quality":"Less is better","values":[0,0,0,0,0,0],"isMain":false}]],["Unaligned",[]],["Mismatches",[]],["Statistics without reference",[]],["K-mer-based statistics",[]],["Predicted genes",[]],["Similarity statistics",[]],["Reference statistics",[]]],[["Genome statistics",[]],["Reads mapping",[]],["Misassemblies",[{"metricName":"# misassemblies","quality":"Less is better","values":[4,8,6,7,10,2],"isMain":true},{"metricName":" # relocations","quality":"Less is better","values":[0,0,0,0,0,0],"isMain":false},{"metricName":" # translocations","quality":"Less is better","values":[0,0,0,0,0,0],"isMain":false},{"metricName":" # inversions","quality":"Less is better","values":[0,0,0,0,0,0],"isMain":false},{"metricName":" # interspecies translocations","quality":"Less is better","values":[4,8,6,7,10,2],"isMain":false},{"metricName":"# possibly misassembled contigs","quality":"Less is better","values":[0,0,0,0,0,0],"isMain":false},{"metricName":" # possible misassemblies","quality":"Less is better","values":[0,0,0,0,0,0],"isMain":false},{"metricName":"# local misassemblies","quality":"Less is better","values":[0,0,1,0,0,0],"isMain":false},{"metricName":"# scaffold gap ext. mis.","quality":"Less is better","values":[0,0,0,0,0,0],"isMain":false},{"metricName":"# scaffold gap loc. mis.","quality":"Less is better","values":[0,0,0,0,0,0],"isMain":false}]],["Unaligned",[]],["Mismatches",[]],["Statistics without reference",[]],["K-mer-based statistics",[]],["Predicted genes",[]],["Similarity statistics",[]],["Reference statistics",[]]],[["Genome statistics",[]],["Reads mapping",[]],["Misassemblies",[{"metricName":"# misassemblies","quality":"Less is better","values":[0,2,1,1,0,0],"isMain":true},{"metricName":" # relocations","quality":"Less is better","values":[0,0,0,0,0,0],"isMain":false},{"metricName":" # translocations","quality":"Less is better","values":[0,0,0,0,0,0],"isMain":false},{"metricName":" # inversions","quality":"Less is better","values":[0,0,0,0,0,0],"isMain":false},{"metricName":" # interspecies translocations","quality":"Less is better","values":[0,2,1,1,0,0],"isMain":false},{"metricName":"# possibly misassembled contigs","quality":"Less is better","values":[0,0,0,0,0,0],"isMain":false},{"metricName":" # possible misassemblies","quality":"Less is better","values":[0,0,0,0,0,0],"isMain":false},{"metricName":"# local misassemblies","quality":"Less is better","values":[0,0,0,0,0,0],"isMain":false},{"metricName":"# scaffold gap ext. mis.","quality":"Less is better","values":[0,0,0,0,0,0],"isMain":false},{"metricName":"# scaffold gap loc. mis.","quality":"Less is better","values":[0,0,0,0,0,0],"isMain":false}]],["Unaligned",[]],["Mismatches",[]],["Statistics without reference",[]],["K-mer-based statistics",[]],["Predicted genes",[]],["Similarity statistics",[]],["Reference statistics",[]]],[["Genome statistics",[]],["Reads mapping",[]],["Misassemblies",[{"metricName":"# misassemblies","quality":"Less is better","values":[26,26,21,43,28,11],"isMain":true},{"metricName":" # relocations","quality":"Less is better","values":[0,1,0,0,0,0],"isMain":false},{"metricName":" # translocations","quality":"Less is better","values":[2,12,8,17,14,6],"isMain":false},{"metricName":" # inversions","quality":"Less is better","values":[0,0,0,1,0,0],"isMain":false},{"metricName":" # interspecies translocations","quality":"Less is better","values":[24,13,13,25,14,5],"isMain":false},{"metricName":"# possibly misassembled contigs","quality":"Less is better","values":[0,0,0,0,0,0],"isMain":false},{"metricName":" # possible misassemblies","quality":"Less is better","values":[0,0,0,0,0,0],"isMain":false},{"metricName":"# local misassemblies","quality":"Less is better","values":[2,1,0,1,1,0],"isMain":false},{"metricName":"# scaffold gap ext. mis.","quality":"Less is better","values":[0,0,0,0,0,0],"isMain":false},{"metricName":"# scaffold gap loc. mis.","quality":"Less is better","values":[0,0,0,0,0,0],"isMain":false}]],["Unaligned",[]],["Mismatches",[]],["Statistics without reference",[]],["K-mer-based statistics",[]],["Predicted genes",[]],["Similarity statistics",[]],["Reference statistics",[]]],[["Genome statistics",[]],["Reads mapping",[]],["Misassemblies",[{"metricName":"# misassemblies","quality":"Less is better","values":[3,3,0,7,1,2],"isMain":true},{"metricName":" # relocations","quality":"Less is better","values":[0,0,0,0,0,0],"isMain":false},{"metricName":" # translocations","quality":"Less is better","values":[0,0,0,0,0,0],"isMain":false},{"metricName":" # inversions","quality":"Less is better","values":[0,0,0,0,0,0],"isMain":false},{"metricName":" # interspecies translocations","quality":"Less is better","values":[3,3,0,7,1,2],"isMain":false},{"metricName":"# possibly misassembled contigs","quality":"Less is better","values":[0,0,0,0,0,0],"isMain":false},{"metricName":" # possible misassemblies","quality":"Less is better","values":[0,0,0,0,0,0],"isMain":false},{"metricName":"# local misassemblies","quality":"Less is better","values":[0,0,0,1,0,0],"isMain":false},{"metricName":"# scaffold gap ext. mis.","quality":"Less is better","values":[0,0,0,0,0,0],"isMain":false},{"metricName":"# scaffold gap loc. mis.","quality":"Less is better","values":[0,0,0,0,0,0],"isMain":false}]],["Unaligned",[]],["Mismatches",[]],["Statistics without reference",[]],["K-mer-based statistics",[]],["Predicted genes",[]],["Similarity statistics",[]],["Reference statistics",[]]],[["Genome statistics",[]],["Reads mapping",[]],["Misassemblies",[{"metricName":"# misassemblies","quality":"Less is better","values":[4,6,0,5,5,2],"isMain":true},{"metricName":" # relocations","quality":"Less is better","values":[0,0,0,0,0,0],"isMain":false},{"metricName":" # translocations","quality":"Less is better","values":[0,0,0,0,0,0],"isMain":false},{"metricName":" # inversions","quality":"Less is better","values":[0,0,0,0,0,0],"isMain":false},{"metricName":" # interspecies translocations","quality":"Less is better","values":[4,6,0,5,5,2],"isMain":false},{"metricName":"# possibly misassembled contigs","quality":"Less is better","values":[0,0,0,0,0,0],"isMain":false},{"metricName":" # possible misassemblies","quality":"Less is better","values":[0,0,0,0,0,0],"isMain":false},{"metricName":"# local misassemblies","quality":"Less is better","values":[0,0,0,0,0,0],"isMain":false},{"metricName":"# scaffold gap ext. mis.","quality":"Less is better","values":[0,0,0,0,0,0],"isMain":false},{"metricName":"# scaffold gap loc. mis.","quality":"Less is better","values":[0,0,0,0,0,0],"isMain":false}]],["Unaligned",[]],["Mismatches",[]],["Statistics without reference",[]],["K-mer-based statistics",[]],["Predicted genes",[]],["Similarity statistics",[]],["Reference statistics",[]]],[["Genome statistics",[]],["Reads mapping",[]],["Misassemblies",[{"metricName":"# misassemblies","quality":"Less is better","values":[7,11,14,19,12,6],"isMain":true},{"metricName":" # relocations","quality":"Less is better","values":[0,0,0,0,0,0],"isMain":false},{"metricName":" # translocations","quality":"Less is better","values":[0,0,1,0,0,0],"isMain":false},{"metricName":" # inversions","quality":"Less is better","values":[0,0,0,0,0,0],"isMain":false},{"metricName":" # interspecies translocations","quality":"Less is better","values":[7,11,13,19,12,6],"isMain":false},{"metricName":"# possibly misassembled contigs","quality":"Less is better","values":[0,0,0,0,0,0],"isMain":false},{"metricName":" # possible misassemblies","quality":"Less is better","values":[0,0,0,0,0,0],"isMain":false},{"metricName":"# local misassemblies","quality":"Less is better","values":[0,0,0,0,1,0],"isMain":false},{"metricName":"# scaffold gap ext. mis.","quality":"Less is better","values":[0,0,0,0,0,0],"isMain":false},{"metricName":"# scaffold gap loc. mis.","quality":"Less is better","values":[0,0,0,0,0,0],"isMain":false}]],["Unaligned",[]],["Mismatches",[]],["Statistics without reference",[]],["K-mer-based statistics",[]],["Predicted genes",[]],["Similarity statistics",[]],["Reference statistics",[]]],[["Genome statistics",[]],["Reads mapping",[]],["Misassemblies",[{"metricName":"# misassemblies","quality":"Less is better","values":[47,62,57,68,71,27],"isMain":true},{"metricName":" # relocations","quality":"Less is better","values":[0,0,1,0,0,1],"isMain":false},{"metricName":" # translocations","quality":"Less is better","values":[41,51,55,52,61,20],"isMain":false},{"metricName":" # inversions","quality":"Less is better","values":[0,0,0,0,0,0],"isMain":false},{"metricName":" # interspecies translocations","quality":"Less is better","values":[6,11,1,16,10,6],"isMain":false},{"metricName":"# possibly misassembled contigs","quality":"Less is better","values":[0,0,0,0,0,0],"isMain":false},{"metricName":" # possible misassemblies","quality":"Less is better","values":[0,0,0,0,0,0],"isMain":false},{"metricName":"# local misassemblies","quality":"Less is better","values":[2,4,5,4,3,0],"isMain":false},{"metricName":"# scaffold gap ext. mis.","quality":"Less is better","values":[0,0,0,0,0,0],"isMain":false},{"metricName":"# scaffold gap loc. mis.","quality":"Less is better","values":[0,0,0,0,0,0],"isMain":false}]],["Unaligned",[]],["Mismatches",[]],["Statistics without reference",[]],["K-mer-based statistics",[]],["Predicted genes",[]],["Similarity statistics",[]],["Reference statistics",[]]],[["Genome statistics",[]],["Reads mapping",[]],["Misassemblies",[{"metricName":"# misassemblies","quality":"Less is better","values":[56,61,15,56,48,37],"isMain":true},{"metricName":" # relocations","quality":"Less is better","values":[1,1,1,1,1,1],"isMain":false},{"metricName":" # translocations","quality":"Less is better","values":[10,14,9,13,14,6],"isMain":false},{"metricName":" # inversions","quality":"Less is better","values":[0,0,1,2,0,0],"isMain":false},{"metricName":" # interspecies translocations","quality":"Less is better","values":[45,46,4,40,33,30],"isMain":false},{"metricName":"# possibly misassembled contigs","quality":"Less is better","values":[0,0,0,0,0,0],"isMain":false},{"metricName":" # possible misassemblies","quality":"Less is better","values":[0,0,0,0,0,0],"isMain":false},{"metricName":"# local misassemblies","quality":"Less is better","values":[1,1,5,5,2,0],"isMain":false},{"metricName":"# scaffold gap ext. mis.","quality":"Less is better","values":[0,0,0,0,0,0],"isMain":false},{"metricName":"# scaffold gap loc. mis.","quality":"Less is better","values":[0,0,0,0,0,0],"isMain":false}]],["Unaligned",[]],["Mismatches",[]],["Statistics without reference",[]],["K-mer-based statistics",[]],["Predicted genes",[]],["Similarity statistics",[]],["Reference statistics",[]]],[["Genome statistics",[]],["Reads mapping",[]],["Misassemblies",[{"metricName":"# misassemblies","quality":"Less is better","values":[10,23,21,37,23,13],"isMain":true},{"metricName":" # relocations","quality":"Less is better","values":[0,0,0,0,0,0],"isMain":false},{"metricName":" # translocations","quality":"Less is better","values":[1,2,2,8,8,7],"isMain":false},{"metricName":" # inversions","quality":"Less is better","values":[3,4,1,2,1,0],"isMain":false},{"metricName":" # interspecies translocations","quality":"Less is better","values":[6,17,18,27,14,6],"isMain":false},{"metricName":"# possibly misassembled contigs","quality":"Less is better","values":[0,0,0,0,0,0],"isMain":false},{"metricName":" # possible misassemblies","quality":"Less is better","values":[0,0,0,0,0,0],"isMain":false},{"metricName":"# local misassemblies","quality":"Less is better","values":[2,1,1,5,0,1],"isMain":false},{"metricName":"# scaffold gap ext. mis.","quality":"Less is better","values":[0,0,0,0,0,0],"isMain":false},{"metricName":"# scaffold gap loc. mis.","quality":"Less is better","values":[0,0,0,0,0,0],"isMain":false}]],["Unaligned",[]],["Mismatches",[]],["Statistics without reference",[]],["K-mer-based statistics",[]],["Predicted genes",[]],["Similarity statistics",[]],["Reference statistics",[]]],[["Genome statistics",[]],["Reads mapping",[]],["Misassemblies",[{"metricName":"# misassemblies","quality":"Less is better","values":[4,7,6,12,8,3],"isMain":true},{"metricName":" # relocations","quality":"Less is better","values":[0,1,0,0,1,0],"isMain":false},{"metricName":" # translocations","quality":"Less is better","values":[4,5,3,7,6,3],"isMain":false},{"metricName":" # inversions","quality":"Less is better","values":[0,0,1,0,0,0],"isMain":false},{"metricName":" # interspecies translocations","quality":"Less is better","values":[0,1,2,5,1,0],"isMain":false},{"metricName":"# possibly misassembled contigs","quality":"Less is better","values":[0,0,0,0,0,0],"isMain":false},{"metricName":" # possible misassemblies","quality":"Less is better","values":[0,0,0,0,0,0],"isMain":false},{"metricName":"# local misassemblies","quality":"Less is better","values":[0,0,0,0,0,0],"isMain":false},{"metricName":"# scaffold gap ext. mis.","quality":"Less is better","values":[0,0,0,0,0,0],"isMain":false},{"metricName":"# scaffold gap loc. mis.","quality":"Less is better","values":[0,0,0,0,0,0],"isMain":false}]],["Unaligned",[]],["Mismatches",[]],["Statistics without reference",[]],["K-mer-based statistics",[]],["Predicted genes",[]],["Similarity statistics",[]],["Reference statistics",[]]],[["Genome statistics",[]],["Reads mapping",[]],["Misassemblies",[{"metricName":"# misassemblies","quality":"Less is better","values":[3,10,9,15,15,0],"isMain":true},{"metricName":" # relocations","quality":"Less is better","values":[0,1,0,0,1,0],"isMain":false},{"metricName":" # translocations","quality":"Less is better","values":[0,0,0,0,0,0],"isMain":false},{"metricName":" # inversions","quality":"Less is better","values":[0,0,0,0,0,0],"isMain":false},{"metricName":" # interspecies translocations","quality":"Less is better","values":[3,9,9,15,14,0],"isMain":false},{"metricName":"# possibly misassembled contigs","quality":"Less is better","values":[0,0,1,1,0,0],"isMain":false},{"metricName":" # possible misassemblies","quality":"Less is better","values":[0,0,1,1,0,0],"isMain":false},{"metricName":"# local misassemblies","quality":"Less is better","values":[1,0,2,1,6,0],"isMain":false},{"metricName":"# scaffold gap ext. mis.","quality":"Less is better","values":[0,0,0,0,0,0],"isMain":false},{"metricName":"# scaffold gap loc. mis.","quality":"Less is better","values":[0,0,0,0,0,0],"isMain":false}]],["Unaligned",[]],["Mismatches",[]],["Statistics without reference",[]],["K-mer-based statistics",[]],["Predicted genes",[]],["Similarity statistics",[]],["Reference statistics",[]]],[["Genome statistics",[]],["Reads mapping",[]],["Misassemblies",[{"metricName":"# misassemblies","quality":"Less is better","values":[7,197,147,212,224,42],"isMain":true},{"metricName":" # relocations","quality":"Less is better","values":[0,3,1,5,1,0],"isMain":false},{"metricName":" # translocations","quality":"Less is better","values":[2,8,7,9,10,3],"isMain":false},{"metricName":" # inversions","quality":"Less is better","values":[0,3,1,5,1,0],"isMain":false},{"metricName":" # interspecies translocations","quality":"Less is better","values":[5,183,138,193,212,39],"isMain":false},{"metricName":"# possibly misassembled contigs","quality":"Less is better","values":[0,2,1,3,2,1],"isMain":false},{"metricName":" # possible misassemblies","quality":"Less is better","values":[0,2,2,5,4,2],"isMain":false},{"metricName":"# local misassemblies","quality":"Less is better","values":[1,11,9,13,14,4],"isMain":false},{"metricName":"# scaffold gap ext. mis.","quality":"Less is better","values":[0,0,0,0,0,0],"isMain":false},{"metricName":"# scaffold gap loc. mis.","quality":"Less is better","values":[0,0,0,0,0,0],"isMain":false}]],["Unaligned",[]],["Mismatches",[]],["Statistics without reference",[]],["K-mer-based statistics",[]],["Predicted genes",[]],["Similarity statistics",[]],["Reference statistics",[]]],[["Genome statistics",[]],["Reads mapping",[]],["Misassemblies",[{"metricName":"# misassemblies","quality":"Less is better","values":[2,13,10,15,17,1],"isMain":true},{"metricName":" # relocations","quality":"Less is better","values":[0,0,0,0,0,0],"isMain":false},{"metricName":" # translocations","quality":"Less is better","values":[0,1,0,0,1,0],"isMain":false},{"metricName":" # inversions","quality":"Less is better","values":[0,0,0,0,0,0],"isMain":false},{"metricName":" # interspecies translocations","quality":"Less is better","values":[2,12,10,15,16,1],"isMain":false},{"metricName":"# possibly misassembled contigs","quality":"Less is better","values":[0,0,0,1,0,0],"isMain":false},{"metricName":" # possible misassemblies","quality":"Less is better","values":[0,0,0,1,0,0],"isMain":false},{"metricName":"# local misassemblies","quality":"Less is better","values":[1,7,2,5,2,1],"isMain":false},{"metricName":"# scaffold gap ext. mis.","quality":"Less is better","values":[0,0,0,0,0,0],"isMain":false},{"metricName":"# scaffold gap loc. mis.","quality":"Less is better","values":[0,0,0,0,0,0],"isMain":false}]],["Unaligned",[]],["Mismatches",[]],["Statistics without reference",[]],["K-mer-based statistics",[]],["Predicted genes",[]],["Similarity statistics",[]],["Reference statistics",[]]],[["Genome statistics",[]],["Reads mapping",[]],["Misassemblies",[{"metricName":"# misassemblies","quality":"Less is better","values":[7,15,10,21,11,0],"isMain":true},{"metricName":" # relocations","quality":"Less is better","values":[0,0,1,0,0,0],"isMain":false},{"metricName":" # translocations","quality":"Less is better","values":[0,0,0,2,0,0],"isMain":false},{"metricName":" # inversions","quality":"Less is better","values":[0,0,0,0,1,0],"isMain":false},{"metricName":" # interspecies translocations","quality":"Less is better","values":[7,15,9,19,10,0],"isMain":false},{"metricName":"# possibly misassembled contigs","quality":"Less is better","values":[0,0,0,1,0,0],"isMain":false},{"metricName":" # possible misassemblies","quality":"Less is better","values":[0,0,0,1,0,0],"isMain":false},{"metricName":"# local misassemblies","quality":"Less is better","values":[1,1,0,1,1,0],"isMain":false},{"metricName":"# scaffold gap ext. mis.","quality":"Less is better","values":[0,0,0,0,0,0],"isMain":false},{"metricName":"# scaffold gap loc. mis.","quality":"Less is better","values":[0,0,0,0,0,0],"isMain":false}]],["Unaligned",[]],["Mismatches",[]],["Statistics without reference",[]],["K-mer-based statistics",[]],["Predicted genes",[]],["Similarity statistics",[]],["Reference statistics",[]]],[["Genome statistics",[]],["Reads mapping",[]],["Misassemblies",[{"metricName":"# misassemblies","quality":"Less is better","values":[16,45,26,34,29,7],"isMain":true},{"metricName":" # relocations","quality":"Less is better","values":[1,1,0,0,2,0],"isMain":false},{"metricName":" # translocations","quality":"Less is better","values":[0,1,0,0,1,0],"isMain":false},{"metricName":" # inversions","quality":"Less is better","values":[0,2,2,4,0,0],"isMain":false},{"metricName":" # interspecies translocations","quality":"Less is better","values":[15,41,24,30,26,7],"isMain":false},{"metricName":"# possibly misassembled contigs","quality":"Less is better","values":[0,0,0,0,0,0],"isMain":false},{"metricName":" # possible misassemblies","quality":"Less is better","values":[0,0,0,0,0,0],"isMain":false},{"metricName":"# local misassemblies","quality":"Less is better","values":[0,1,1,1,1,0],"isMain":false},{"metricName":"# scaffold gap ext. mis.","quality":"Less is better","values":[0,0,0,0,0,0],"isMain":false},{"metricName":"# scaffold gap loc. mis.","quality":"Less is better","values":[0,0,0,0,0,0],"isMain":false}]],["Unaligned",[]],["Mismatches",[]],["Statistics without reference",[]],["K-mer-based statistics",[]],["Predicted genes",[]],["Similarity statistics",[]],["Reference statistics",[]]],[["Genome statistics",[]],["Reads mapping",[]],["Misassemblies",[{"metricName":"# misassemblies","quality":"Less is better","values":[20,24,29,32,39,6],"isMain":true},{"metricName":" # relocations","quality":"Less is better","values":[0,1,1,0,1,0],"isMain":false},{"metricName":" # translocations","quality":"Less is better","values":[3,3,5,6,3,1],"isMain":false},{"metricName":" # inversions","quality":"Less is better","values":[0,0,0,1,0,0],"isMain":false},{"metricName":" # interspecies translocations","quality":"Less is better","values":[17,20,23,25,35,5],"isMain":false},{"metricName":"# possibly misassembled contigs","quality":"Less is better","values":[0,1,0,2,1,0],"isMain":false},{"metricName":" # possible misassemblies","quality":"Less is better","values":[0,1,0,2,1,0],"isMain":false},{"metricName":"# local misassemblies","quality":"Less is better","values":[2,2,1,6,8,1],"isMain":false},{"metricName":"# scaffold gap ext. mis.","quality":"Less is better","values":[0,0,0,0,0,0],"isMain":false},{"metricName":"# scaffold gap loc. mis.","quality":"Less is better","values":[0,0,0,0,0,0],"isMain":false}]],["Unaligned",[]],["Mismatches",[]],["Statistics without reference",[]],["K-mer-based statistics",[]],["Predicted genes",[]],["Similarity statistics",[]],["Reference statistics",[]]],[["Genome statistics",[]],["Reads mapping",[]],["Misassemblies",[{"metricName":"# misassemblies","quality":"Less is better","values":[4,14,15,26,28,1],"isMain":true},{"metricName":" # relocations","quality":"Less is better","values":[0,0,0,0,1,0],"isMain":false},{"metricName":" # translocations","quality":"Less is better","values":[0,0,3,1,3,0],"isMain":false},{"metricName":" # inversions","quality":"Less is better","values":[0,0,0,1,0,0],"isMain":false},{"metricName":" # interspecies translocations","quality":"Less is better","values":[4,14,12,24,24,1],"isMain":false},{"metricName":"# possibly misassembled contigs","quality":"Less is better","values":[0,0,0,0,0,0],"isMain":false},{"metricName":" # possible misassemblies","quality":"Less is better","values":[0,0,0,0,0,0],"isMain":false},{"metricName":"# local misassemblies","quality":"Less is better","values":[0,4,4,0,2,0],"isMain":false},{"metricName":"# scaffold gap ext. mis.","quality":"Less is better","values":[0,0,0,0,0,0],"isMain":false},{"metricName":"# scaffold gap loc. mis.","quality":"Less is better","values":[0,0,0,0,0,0],"isMain":false}]],["Unaligned",[]],["Mismatches",[]],["Statistics without reference",[]],["K-mer-based statistics",[]],["Predicted genes",[]],["Similarity statistics",[]],["Reference statistics",[]]],[["Genome statistics",[]],["Reads mapping",[]],["Misassemblies",[{"metricName":"# misassemblies","quality":"Less is better","values":[9,16,14,27,23,3],"isMain":true},{"metricName":" # relocations","quality":"Less is better","values":[0,0,0,1,2,0],"isMain":false},{"metricName":" # translocations","quality":"Less is better","values":[0,0,1,1,0,0],"isMain":false},{"metricName":" # inversions","quality":"Less is better","values":[1,0,0,0,0,0],"isMain":false},{"metricName":" # interspecies translocations","quality":"Less is better","values":[8,16,13,25,21,3],"isMain":false},{"metricName":"# possibly misassembled contigs","quality":"Less is better","values":[0,0,0,0,0,0],"isMain":false},{"metricName":" # possible misassemblies","quality":"Less is better","values":[0,0,0,0,0,0],"isMain":false},{"metricName":"# local misassemblies","quality":"Less is better","values":[0,1,2,3,3,0],"isMain":false},{"metricName":"# scaffold gap ext. mis.","quality":"Less is better","values":[0,0,0,0,0,0],"isMain":false},{"metricName":"# scaffold gap loc. mis.","quality":"Less is better","values":[0,0,0,0,0,0],"isMain":false}]],["Unaligned",[]],["Mismatches",[]],["Statistics without reference",[]],["K-mer-based statistics",[]],["Predicted genes",[]],["Similarity statistics",[]],["Reference statistics",[]]],[["Genome statistics",[]],["Reads mapping",[]],["Misassemblies",[{"metricName":"# misassemblies","quality":"Less is better","values":[26,26,12,31,29,14],"isMain":true},{"metricName":" # relocations","quality":"Less is better","values":[0,0,0,0,0,0],"isMain":false},{"metricName":" # translocations","quality":"Less is better","values":[0,0,1,0,0,0],"isMain":false},{"metricName":" # inversions","quality":"Less is better","values":[0,0,1,1,1,0],"isMain":false},{"metricName":" # interspecies translocations","quality":"Less is better","values":[26,26,10,30,28,14],"isMain":false},{"metricName":"# possibly misassembled contigs","quality":"Less is better","values":[0,1,0,0,1,0],"isMain":false},{"metricName":" # possible misassemblies","quality":"Less is better","values":[0,1,0,0,1,0],"isMain":false},{"metricName":"# local misassemblies","quality":"Less is better","values":[1,2,1,0,3,0],"isMain":false},{"metricName":"# scaffold gap ext. mis.","quality":"Less is better","values":[0,0,0,0,0,0],"isMain":false},{"metricName":"# scaffold gap loc. mis.","quality":"Less is better","values":[0,0,0,0,0,0],"isMain":false}]],["Unaligned",[]],["Mismatches",[]],["Statistics without reference",[]],["K-mer-based statistics",[]],["Predicted genes",[]],["Similarity statistics",[]],["Reference statistics",[]]],[["Genome statistics",[]],["Reads mapping",[]],["Misassemblies",[{"metricName":"# misassemblies","quality":"Less is better","values":[19,33,9,24,21,7],"isMain":true},{"metricName":" # relocations","quality":"Less is better","values":[0,0,0,0,0,0],"isMain":false},{"metricName":" # translocations","quality":"Less is better","values":[0,0,0,0,0,0],"isMain":false},{"metricName":" # inversions","quality":"Less is better","values":[0,1,0,0,0,0],"isMain":false},{"metricName":" # interspecies translocations","quality":"Less is better","values":[19,32,9,24,21,7],"isMain":false},{"metricName":"# possibly misassembled contigs","quality":"Less is better","values":[1,0,0,2,0,0],"isMain":false},{"metricName":" # possible misassemblies","quality":"Less is better","values":[1,0,0,2,0,0],"isMain":false},{"metricName":"# local misassemblies","quality":"Less is better","values":[1,0,1,3,2,0],"isMain":false},{"metricName":"# scaffold gap ext. mis.","quality":"Less is better","values":[0,0,0,0,0,0],"isMain":false},{"metricName":"# scaffold gap loc. mis.","quality":"Less is better","values":[0,0,0,0,0,0],"isMain":false}]],["Unaligned",[]],["Mismatches",[]],["Statistics without reference",[]],["K-mer-based statistics",[]],["Predicted genes",[]],["Similarity statistics",[]],["Reference statistics",[]]],[["Genome statistics",[]],["Reads mapping",[]],["Misassemblies",[{"metricName":"# misassemblies","quality":"Less is better","values":[14,25,4,31,22,12],"isMain":true},{"metricName":" # relocations","quality":"Less is better","values":[0,1,0,0,1,0],"isMain":false},{"metricName":" # translocations","quality":"Less is better","values":[1,0,0,3,0,1],"isMain":false},{"metricName":" # inversions","quality":"Less is better","values":[0,0,0,1,1,0],"isMain":false},{"metricName":" # interspecies translocations","quality":"Less is better","values":[13,24,4,27,20,11],"isMain":false},{"metricName":"# possibly misassembled contigs","quality":"Less is better","values":[0,0,0,0,0,0],"isMain":false},{"metricName":" # possible misassemblies","quality":"Less is better","values":[0,0,0,0,0,0],"isMain":false},{"metricName":"# local misassemblies","quality":"Less is better","values":[0,0,0,0,1,0],"isMain":false},{"metricName":"# scaffold gap ext. mis.","quality":"Less is better","values":[0,0,0,0,0,0],"isMain":false},{"metricName":"# scaffold gap loc. mis.","quality":"Less is better","values":[0,0,0,0,0,0],"isMain":false}]],["Unaligned",[]],["Mismatches",[]],["Statistics without reference",[]],["K-mer-based statistics",[]],["Predicted genes",[]],["Similarity statistics",[]],["Reference statistics",[]]],[["Genome statistics",[]],["Reads mapping",[]],["Misassemblies",[{"metricName":"# misassemblies","quality":"Less is better","values":[77,116,61,118,105,34],"isMain":true},{"metricName":" # relocations","quality":"Less is better","values":[0,2,2,0,0,0],"isMain":false},{"metricName":" # translocations","quality":"Less is better","values":[5,5,7,7,9,4],"isMain":false},{"metricName":" # inversions","quality":"Less is better","values":[2,0,4,5,0,1],"isMain":false},{"metricName":" # interspecies translocations","quality":"Less is better","values":[70,109,48,106,96,29],"isMain":false},{"metricName":"# possibly misassembled contigs","quality":"Less is better","values":[0,1,3,1,2,0],"isMain":false},{"metricName":" # possible misassemblies","quality":"Less is better","values":[0,1,3,1,2,0],"isMain":false},{"metricName":"# local misassemblies","quality":"Less is better","values":[2,4,4,6,5,1],"isMain":false},{"metricName":"# scaffold gap ext. mis.","quality":"Less is better","values":[0,0,0,0,0,0],"isMain":false},{"metricName":"# scaffold gap loc. mis.","quality":"Less is better","values":[0,0,0,0,0,0],"isMain":false}]],["Unaligned",[]],["Mismatches",[]],["Statistics without reference",[]],["K-mer-based statistics",[]],["Predicted genes",[]],["Similarity statistics",[]],["Reference statistics",[]]],[["Genome statistics",[]],["Reads mapping",[]],["Misassemblies",[{"metricName":"# misassemblies","quality":"Less is better","values":[65,74,16,47,55,30],"isMain":true},{"metricName":" # relocations","quality":"Less is better","values":[2,2,0,1,2,2],"isMain":false},{"metricName":" # translocations","quality":"Less is better","values":[2,4,0,2,1,1],"isMain":false},{"metricName":" # inversions","quality":"Less is better","values":[2,1,0,0,1,0],"isMain":false},{"metricName":" # interspecies translocations","quality":"Less is better","values":[59,67,16,44,51,27],"isMain":false},{"metricName":"# possibly misassembled contigs","quality":"Less is better","values":[0,0,0,0,0,0],"isMain":false},{"metricName":" # possible misassemblies","quality":"Less is better","values":[0,0,0,0,0,0],"isMain":false},{"metricName":"# local misassemblies","quality":"Less is better","values":[4,4,3,2,1,1],"isMain":false},{"metricName":"# scaffold gap ext. mis.","quality":"Less is better","values":[0,0,0,0,0,0],"isMain":false},{"metricName":"# scaffold gap loc. mis.","quality":"Less is better","values":[0,0,0,0,0,0],"isMain":false}]],["Unaligned",[]],["Mismatches",[]],["Statistics without reference",[]],["K-mer-based statistics",[]],["Predicted genes",[]],["Similarity statistics",[]],["Reference statistics",[]]],[["Genome statistics",[]],["Reads mapping",[]],["Misassemblies",[{"metricName":"# misassemblies","quality":"Less is better","values":[14,20,8,38,21,11],"isMain":true},{"metricName":" # relocations","quality":"Less is better","values":[0,0,0,0,0,1],"isMain":false},{"metricName":" # translocations","quality":"Less is better","values":[0,0,1,1,0,0],"isMain":false},{"metricName":" # inversions","quality":"Less is better","values":[0,0,0,0,0,0],"isMain":false},{"metricName":" # interspecies translocations","quality":"Less is better","values":[14,20,7,37,21,10],"isMain":false},{"metricName":"# possibly misassembled contigs","quality":"Less is better","values":[0,0,0,0,0,0],"isMain":false},{"metricName":" # possible misassemblies","quality":"Less is better","values":[0,0,0,0,0,0],"isMain":false},{"metricName":"# local misassemblies","quality":"Less is better","values":[1,0,0,0,1,0],"isMain":false},{"metricName":"# scaffold gap ext. mis.","quality":"Less is better","values":[0,0,0,0,0,0],"isMain":false},{"metricName":"# scaffold gap loc. mis.","quality":"Less is better","values":[0,0,0,0,0,0],"isMain":false}]],["Unaligned",[]],["Mismatches",[]],["Statistics without reference",[]],["K-mer-based statistics",[]],["Predicted genes",[]],["Similarity statistics",[]],["Reference statistics",[]]],[["Genome statistics",[]],["Reads mapping",[]],["Misassemblies",[{"metricName":"# misassemblies","quality":"Less is better","values":[0,2,0,5,2,2],"isMain":true},{"metricName":" # relocations","quality":"Less is better","values":[0,0,0,0,0,0],"isMain":false},{"metricName":" # translocations","quality":"Less is better","values":[0,0,0,0,0,0],"isMain":false},{"metricName":" # inversions","quality":"Less is better","values":[0,0,0,0,0,0],"isMain":false},{"metricName":" # interspecies translocations","quality":"Less is better","values":[0,2,0,5,2,2],"isMain":false},{"metricName":"# possibly misassembled contigs","quality":"Less is better","values":[0,0,0,0,0,0],"isMain":false},{"metricName":" # possible misassemblies","quality":"Less is better","values":[0,0,0,0,0,0],"isMain":false},{"metricName":"# local misassemblies","quality":"Less is better","values":[0,0,0,0,0,0],"isMain":false},{"metricName":"# scaffold gap ext. mis.","quality":"Less is better","values":[0,0,0,0,0,0],"isMain":false},{"metricName":"# scaffold gap loc. mis.","quality":"Less is better","values":[0,0,0,0,0,0],"isMain":false}]],["Unaligned",[]],["Mismatches",[]],["Statistics without reference",[]],["K-mer-based statistics",[]],["Predicted genes",[]],["Similarity statistics",[]],["Reference statistics",[]]],[["Genome statistics",[]],["Reads mapping",[]],["Misassemblies",[{"metricName":"# misassemblies","quality":"Less is better","values":[5,8,1,5,6,1],"isMain":true},{"metricName":" # relocations","quality":"Less is better","values":[0,0,0,0,0,0],"isMain":false},{"metricName":" # translocations","quality":"Less is better","values":[0,0,0,0,0,0],"isMain":false},{"metricName":" # inversions","quality":"Less is better","values":[0,0,0,0,0,0],"isMain":false},{"metricName":" # interspecies translocations","quality":"Less is better","values":[5,8,1,5,6,1],"isMain":false},{"metricName":"# possibly misassembled contigs","quality":"Less is better","values":[0,0,0,0,0,0],"isMain":false},{"metricName":" # possible misassemblies","quality":"Less is better","values":[0,0,0,0,0,0],"isMain":false},{"metricName":"# local misassemblies","quality":"Less is better","values":[0,0,0,0,0,0],"isMain":false},{"metricName":"# scaffold gap ext. mis.","quality":"Less is better","values":[0,0,0,0,0,0],"isMain":false},{"metricName":"# scaffold gap loc. mis.","quality":"Less is better","values":[0,0,0,0,0,0],"isMain":false}]],["Unaligned",[]],["Mismatches",[]],["Statistics without reference",[]],["K-mer-based statistics",[]],["Predicted genes",[]],["Similarity statistics",[]],["Reference statistics",[]]],[["Genome statistics",[]],["Reads mapping",[]],["Misassemblies",[{"metricName":"# misassemblies","quality":"Less is better","values":[96,134,55,112,120,54],"isMain":true},{"metricName":" # relocations","quality":"Less is better","values":[1,3,3,0,1,1],"isMain":false},{"metricName":" # translocations","quality":"Less is better","values":[2,6,4,4,6,5],"isMain":false},{"metricName":" # inversions","quality":"Less is better","values":[4,0,5,6,1,0],"isMain":false},{"metricName":" # interspecies translocations","quality":"Less is better","values":[89,125,43,102,112,48],"isMain":false},{"metricName":"# possibly misassembled contigs","quality":"Less is better","values":[0,1,0,0,0,0],"isMain":false},{"metricName":" # possible misassemblies","quality":"Less is better","values":[0,1,0,0,0,0],"isMain":false},{"metricName":"# local misassemblies","quality":"Less is better","values":[2,2,5,3,3,0],"isMain":false},{"metricName":"# scaffold gap ext. mis.","quality":"Less is better","values":[0,0,0,0,0,0],"isMain":false},{"metricName":"# scaffold gap loc. mis.","quality":"Less is better","values":[0,0,0,0,0,0],"isMain":false}]],["Unaligned",[]],["Mismatches",[]],["Statistics without reference",[]],["K-mer-based statistics",[]],["Predicted genes",[]],["Similarity statistics",[]],["Reference statistics",[]]],[["Genome statistics",[]],["Reads mapping",[]],["Misassemblies",[{"metricName":"# misassemblies","quality":"Less is better","values":[38,32,31,41,51,10],"isMain":true},{"metricName":" # relocations","quality":"Less is better","values":[2,1,0,1,2,1],"isMain":false},{"metricName":" # translocations","quality":"Less is better","values":[4,3,2,0,6,1],"isMain":false},{"metricName":" # inversions","quality":"Less is better","values":[1,0,0,0,3,0],"isMain":false},{"metricName":" # interspecies translocations","quality":"Less is better","values":[31,28,29,40,40,8],"isMain":false},{"metricName":"# possibly misassembled contigs","quality":"Less is better","values":[0,0,0,0,0,0],"isMain":false},{"metricName":" # possible misassemblies","quality":"Less is better","values":[0,0,0,0,0,0],"isMain":false},{"metricName":"# local misassemblies","quality":"Less is better","values":[2,0,0,1,4,0],"isMain":false},{"metricName":"# scaffold gap ext. mis.","quality":"Less is better","values":[0,0,0,0,0,0],"isMain":false},{"metricName":"# scaffold gap loc. mis.","quality":"Less is better","values":[0,0,0,0,0,0],"isMain":false}]],["Unaligned",[]],["Mismatches",[]],["Statistics without reference",[]],["K-mer-based statistics",[]],["Predicted genes",[]],["Similarity statistics",[]],["Reference statistics",[]]],[["Genome statistics",[]],["Reads mapping",[]],["Misassemblies",[{"metricName":"# misassemblies","quality":"Less is better","values":[6,10,0,9,9,1],"isMain":true},{"metricName":" # relocations","quality":"Less is better","values":[0,0,0,0,0,0],"isMain":false},{"metricName":" # translocations","quality":"Less is better","values":[0,0,0,0,0,0],"isMain":false},{"metricName":" # inversions","quality":"Less is better","values":[0,0,0,0,0,0],"isMain":false},{"metricName":" # interspecies translocations","quality":"Less is better","values":[6,10,0,9,9,1],"isMain":false},{"metricName":"# possibly misassembled contigs","quality":"Less is better","values":[1,0,0,0,0,0],"isMain":false},{"metricName":" # possible misassemblies","quality":"Less is better","values":[1,0,0,0,0,0],"isMain":false},{"metricName":"# local misassemblies","quality":"Less is better","values":[0,0,0,1,0,0],"isMain":false},{"metricName":"# scaffold gap ext. mis.","quality":"Less is better","values":[0,0,0,0,0,0],"isMain":false},{"metricName":"# scaffold gap loc. mis.","quality":"Less is better","values":[0,0,0,0,0,0],"isMain":false}]],["Unaligned",[]],["Mismatches",[]],["Statistics without reference",[]],["K-mer-based statistics",[]],["Predicted genes",[]],["Similarity statistics",[]],["Reference statistics",[]]],[["Genome statistics",[]],["Reads mapping",[]],["Misassemblies",[{"metricName":"# misassemblies","quality":"Less is better","values":[37,50,19,43,40,24],"isMain":true},{"metricName":" # relocations","quality":"Less is better","values":[0,0,0,0,0,0],"isMain":false},{"metricName":" # translocations","quality":"Less is better","values":[2,2,2,0,2,0],"isMain":false},{"metricName":" # inversions","quality":"Less is better","values":[0,1,0,0,0,0],"isMain":false},{"metricName":" # interspecies translocations","quality":"Less is better","values":[35,47,17,43,38,24],"isMain":false},{"metricName":"# possibly misassembled contigs","quality":"Less is better","values":[0,0,0,0,0,0],"isMain":false},{"metricName":" # possible misassemblies","quality":"Less is better","values":[0,0,0,0,0,0],"isMain":false},{"metricName":"# local misassemblies","quality":"Less is better","values":[0,1,0,1,0,0],"isMain":false},{"metricName":"# scaffold gap ext. mis.","quality":"Less is better","values":[0,0,0,0,0,0],"isMain":false},{"metricName":"# scaffold gap loc. mis.","quality":"Less is better","values":[0,0,0,0,0,0],"isMain":false}]],["Unaligned",[]],["Mismatches",[]],["Statistics without reference",[]],["K-mer-based statistics",[]],["Predicted genes",[]],["Similarity statistics",[]],["Reference statistics",[]]],[["Genome statistics",[]],["Reads mapping",[]],["Misassemblies",[{"metricName":"# misassemblies","quality":"Less is better","values":[6,12,2,6,8,6],"isMain":true},{"metricName":" # relocations","quality":"Less is better","values":[0,0,0,0,0,0],"isMain":false},{"metricName":" # translocations","quality":"Less is better","values":[0,0,0,0,0,0],"isMain":false},{"metricName":" # inversions","quality":"Less is better","values":[0,0,0,0,0,0],"isMain":false},{"metricName":" # interspecies translocations","quality":"Less is better","values":[6,12,2,6,8,6],"isMain":false},{"metricName":"# possibly misassembled contigs","quality":"Less is better","values":[0,0,0,0,0,0],"isMain":false},{"metricName":" # possible misassemblies","quality":"Less is better","values":[0,0,0,0,0,0],"isMain":false},{"metricName":"# local misassemblies","quality":"Less is better","values":[0,0,0,0,0,0],"isMain":false},{"metricName":"# scaffold gap ext. mis.","quality":"Less is better","values":[0,0,0,0,0,0],"isMain":false},{"metricName":"# scaffold gap loc. mis.","quality":"Less is better","values":[0,0,0,0,0,0],"isMain":false}]],["Unaligned",[]],["Mismatches",[]],["Statistics without reference",[]],["K-mer-based statistics",[]],["Predicted genes",[]],["Similarity statistics",[]],["Reference statistics",[]]],[["Genome statistics",[]],["Reads mapping",[]],["Misassemblies",[{"metricName":"# misassemblies","quality":"Less is better","values":[0,1,0,1,0,0],"isMain":true},{"metricName":" # relocations","quality":"Less is better","values":[0,0,0,0,0,0],"isMain":false},{"metricName":" # translocations","quality":"Less is better","values":[0,0,0,0,0,0],"isMain":false},{"metricName":" # inversions","quality":"Less is better","values":[0,1,0,0,0,0],"isMain":false},{"metricName":" # interspecies translocations","quality":"Less is better","values":[0,0,0,1,0,0],"isMain":false},{"metricName":"# possibly misassembled contigs","quality":"Less is better","values":[0,0,0,0,0,0],"isMain":false},{"metricName":" # possible misassemblies","quality":"Less is better","values":[0,0,0,0,0,0],"isMain":false},{"metricName":"# local misassemblies","quality":"Less is better","values":[0,0,0,0,0,0],"isMain":false},{"metricName":"# scaffold gap ext. mis.","quality":"Less is better","values":[0,0,0,0,0,0],"isMain":false},{"metricName":"# scaffold gap loc. mis.","quality":"Less is better","values":[0,0,0,0,0,0],"isMain":false}]],["Unaligned",[]],["Mismatches",[]],["Statistics without reference",[]],["K-mer-based statistics",[]],["Predicted genes",[]],["Similarity statistics",[]],["Reference statistics",[]]],[["Genome statistics",[]],["Reads mapping",[]],["Misassemblies",[{"metricName":"# misassemblies","quality":"Less is better","values":[15,33,10,31,21,18],"isMain":true},{"metricName":" # relocations","quality":"Less is better","values":[0,0,0,0,0,0],"isMain":false},{"metricName":" # translocations","quality":"Less is better","values":[2,9,5,5,8,2],"isMain":false},{"metricName":" # inversions","quality":"Less is better","values":[0,0,0,0,0,0],"isMain":false},{"metricName":" # interspecies translocations","quality":"Less is better","values":[13,24,5,26,13,16],"isMain":false},{"metricName":"# possibly misassembled contigs","quality":"Less is better","values":[0,0,0,0,0,0],"isMain":false},{"metricName":" # possible misassemblies","quality":"Less is better","values":[0,0,0,0,0,0],"isMain":false},{"metricName":"# local misassemblies","quality":"Less is better","values":[0,3,0,0,2,0],"isMain":false},{"metricName":"# scaffold gap ext. mis.","quality":"Less is better","values":[0,0,0,0,0,0],"isMain":false},{"metricName":"# scaffold gap loc. mis.","quality":"Less is better","values":[0,0,0,0,0,0],"isMain":false}]],["Unaligned",[]],["Mismatches",[]],["Statistics without reference",[]],["K-mer-based statistics",[]],["Predicted genes",[]],["Similarity statistics",[]],["Reference statistics",[]]],[["Genome statistics",[]],["Reads mapping",[]],["Misassemblies",[{"metricName":"# misassemblies","quality":"Less is better","values":[16,20,3,20,16,15],"isMain":true},{"metricName":" # relocations","quality":"Less is better","values":[0,0,0,0,0,1],"isMain":false},{"metricName":" # translocations","quality":"Less is better","values":[1,1,1,0,1,0],"isMain":false},{"metricName":" # inversions","quality":"Less is better","values":[0,0,0,0,0,0],"isMain":false},{"metricName":" # interspecies translocations","quality":"Less is better","values":[15,19,2,20,15,14],"isMain":false},{"metricName":"# possibly misassembled contigs","quality":"Less is better","values":[0,0,0,0,0,0],"isMain":false},{"metricName":" # possible misassemblies","quality":"Less is better","values":[0,0,0,0,0,0],"isMain":false},{"metricName":"# local misassemblies","quality":"Less is better","values":[0,0,0,0,0,0],"isMain":false},{"metricName":"# scaffold gap ext. mis.","quality":"Less is better","values":[0,0,0,0,0,0],"isMain":false},{"metricName":"# scaffold gap loc. mis.","quality":"Less is better","values":[0,0,0,0,0,0],"isMain":false}]],["Unaligned",[]],["Mismatches",[]],["Statistics without reference",[]],["K-mer-based statistics",[]],["Predicted genes",[]],["Similarity statistics",[]],["Reference statistics",[]]],[["Genome statistics",[]],["Reads mapping",[]],["Misassemblies",[{"metricName":"# misassemblies","quality":"Less is better","values":[18,18,3,17,19,15],"isMain":true},{"metricName":" # relocations","quality":"Less is better","values":[0,0,0,0,0,0],"isMain":false},{"metricName":" # translocations","quality":"Less is better","values":[1,0,0,1,1,0],"isMain":false},{"metricName":" # inversions","quality":"Less is better","values":[0,0,0,0,0,0],"isMain":false},{"metricName":" # interspecies translocations","quality":"Less is better","values":[17,18,3,16,18,15],"isMain":false},{"metricName":"# possibly misassembled contigs","quality":"Less is better","values":[0,0,0,0,0,0],"isMain":false},{"metricName":" # possible misassemblies","quality":"Less is better","values":[0,0,0,0,0,0],"isMain":false},{"metricName":"# local misassemblies","quality":"Less is better","values":[1,1,0,1,2,0],"isMain":false},{"metricName":"# scaffold gap ext. mis.","quality":"Less is better","values":[0,0,0,0,0,0],"isMain":false},{"metricName":"# scaffold gap loc. mis.","quality":"Less is better","values":[0,0,0,0,0,0],"isMain":false}]],["Unaligned",[]],["Mismatches",[]],["Statistics without reference",[]],["K-mer-based statistics",[]],["Predicted genes",[]],["Similarity statistics",[]],["Reference statistics",[]]],[["Genome statistics",[]],["Reads mapping",[]],["Misassemblies",[{"metricName":"# misassemblies","quality":"Less is better","values":[10,19,11,19,21,7],"isMain":true},{"metricName":" # relocations","quality":"Less is better","values":[1,2,1,1,2,1],"isMain":false},{"metricName":" # translocations","quality":"Less is better","values":[3,1,3,3,4,0],"isMain":false},{"metricName":" # inversions","quality":"Less is better","values":[0,0,0,0,1,0],"isMain":false},{"metricName":" # interspecies translocations","quality":"Less is better","values":[6,16,7,15,14,6],"isMain":false},{"metricName":"# possibly misassembled contigs","quality":"Less is better","values":[0,0,0,0,0,0],"isMain":false},{"metricName":" # possible misassemblies","quality":"Less is better","values":[0,0,0,0,0,0],"isMain":false},{"metricName":"# local misassemblies","quality":"Less is better","values":[2,0,0,2,0,0],"isMain":false},{"metricName":"# scaffold gap ext. mis.","quality":"Less is better","values":[0,0,0,0,0,0],"isMain":false},{"metricName":"# scaffold gap loc. mis.","quality":"Less is better","values":[0,0,0,0,0,0],"isMain":false}]],["Unaligned",[]],["Mismatches",[]],["Statistics without reference",[]],["K-mer-based statistics",[]],["Predicted genes",[]],["Similarity statistics",[]],["Reference statistics",[]]],[["Genome statistics",[]],["Reads mapping",[]],["Misassemblies",[{"metricName":"# misassemblies","quality":"Less is better","values":[18,43,44,68,81,7],"isMain":true},{"metricName":" # relocations","quality":"Less is better","values":[0,2,1,1,0,0],"isMain":false},{"metricName":" # translocations","quality":"Less is better","values":[0,1,1,0,1,1],"isMain":false},{"metricName":" # inversions","quality":"Less is better","values":[0,0,2,2,1,0],"isMain":false},{"metricName":" # interspecies translocations","quality":"Less is better","values":[18,40,40,65,79,6],"isMain":false},{"metricName":"# possibly misassembled contigs","quality":"Less is better","values":[0,0,1,1,3,0],"isMain":false},{"metricName":" # possible misassemblies","quality":"Less is better","values":[0,0,1,1,3,0],"isMain":false},{"metricName":"# local misassemblies","quality":"Less is better","values":[2,2,1,3,4,1],"isMain":false},{"metricName":"# scaffold gap ext. mis.","quality":"Less is better","values":[0,0,0,0,0,0],"isMain":false},{"metricName":"# scaffold gap loc. mis.","quality":"Less is better","values":[0,0,0,0,0,0],"isMain":false}]],["Unaligned",[]],["Mismatches",[]],["Statistics without reference",[]],["K-mer-based statistics",[]],["Predicted genes",[]],["Similarity statistics",[]],["Reference statistics",[]]],[["Genome statistics",[]],["Reads mapping",[]],["Misassemblies",[{"metricName":"# misassemblies","quality":"Less is better","values":[12,22,15,30,23,8],"isMain":true},{"metricName":" # relocations","quality":"Less is better","values":[0,0,0,0,0,0],"isMain":false},{"metricName":" # translocations","quality":"Less is better","values":[0,1,0,1,2,0],"isMain":false},{"metricName":" # inversions","quality":"Less is better","values":[0,0,0,3,0,0],"isMain":false},{"metricName":" # interspecies translocations","quality":"Less is better","values":[12,21,15,26,21,8],"isMain":false},{"metricName":"# possibly misassembled contigs","quality":"Less is better","values":[0,0,0,0,0,0],"isMain":false},{"metricName":" # possible misassemblies","quality":"Less is better","values":[0,0,0,0,0,0],"isMain":false},{"metricName":"# local misassemblies","quality":"Less is better","values":[0,1,0,1,2,0],"isMain":false},{"metricName":"# scaffold gap ext. mis.","quality":"Less is better","values":[0,0,0,0,0,0],"isMain":false},{"metricName":"# scaffold gap loc. mis.","quality":"Less is better","values":[0,0,0,0,0,0],"isMain":false}]],["Unaligned",[]],["Mismatches",[]],["Statistics without reference",[]],["K-mer-based statistics",[]],["Predicted genes",[]],["Similarity statistics",[]],["Reference statistics",[]]],[["Genome statistics",[]],["Reads mapping",[]],["Misassemblies",[{"metricName":"# misassemblies","quality":"Less is better","values":[17,42,25,46,38,11],"isMain":true},{"metricName":" # relocations","quality":"Less is better","values":[0,2,1,0,0,0],"isMain":false},{"metricName":" # translocations","quality":"Less is better","values":[5,21,13,19,20,7],"isMain":false},{"metricName":" # inversions","quality":"Less is better","values":[0,0,0,0,1,0],"isMain":false},{"metricName":" # interspecies translocations","quality":"Less is better","values":[12,19,11,27,17,4],"isMain":false},{"metricName":"# possibly misassembled contigs","quality":"Less is better","values":[0,0,0,1,0,0],"isMain":false},{"metricName":" # possible misassemblies","quality":"Less is better","values":[0,0,0,2,0,0],"isMain":false},{"metricName":"# local misassemblies","quality":"Less is better","values":[1,3,3,4,5,0],"isMain":false},{"metricName":"# scaffold gap ext. mis.","quality":"Less is better","values":[0,0,0,0,0,0],"isMain":false},{"metricName":"# scaffold gap loc. mis.","quality":"Less is better","values":[0,0,0,0,0,0],"isMain":false}]],["Unaligned",[]],["Mismatches",[]],["Statistics without reference",[]],["K-mer-based statistics",[]],["Predicted genes",[]],["Similarity statistics",[]],["Reference statistics",[]]],[["Genome statistics",[]],["Reads mapping",[]],["Misassemblies",[{"metricName":"# misassemblies","quality":"Less is better","values":[7,33,32,47,41,6],"isMain":true},{"metricName":" # relocations","quality":"Less is better","values":[0,3,1,1,2,0],"isMain":false},{"metricName":" # translocations","quality":"Less is better","values":[2,10,9,15,7,3],"isMain":false},{"metricName":" # inversions","quality":"Less is better","values":[0,1,1,0,2,0],"isMain":false},{"metricName":" # interspecies translocations","quality":"Less is better","values":[5,19,21,31,30,3],"isMain":false},{"metricName":"# possibly misassembled contigs","quality":"Less is better","values":[0,0,0,0,0,0],"isMain":false},{"metricName":" # possible misassemblies","quality":"Less is better","values":[0,0,0,0,0,0],"isMain":false},{"metricName":"# local misassemblies","quality":"Less is better","values":[1,3,2,2,1,0],"isMain":false},{"metricName":"# scaffold gap ext. mis.","quality":"Less is better","values":[0,0,0,0,0,0],"isMain":false},{"metricName":"# scaffold gap loc. mis.","quality":"Less is better","values":[0,0,0,0,0,0],"isMain":false}]],["Unaligned",[]],["Mismatches",[]],["Statistics without reference",[]],["K-mer-based statistics",[]],["Predicted genes",[]],["Similarity statistics",[]],["Reference statistics",[]]],[["Genome statistics",[]],["Reads mapping",[]],["Misassemblies",[{"metricName":"# misassemblies","quality":"Less is better","values":[1,1,1,3,2,1],"isMain":true},{"metricName":" # relocations","quality":"Less is better","values":[0,0,0,0,0,0],"isMain":false},{"metricName":" # translocations","quality":"Less is better","values":[0,0,0,0,0,0],"isMain":false},{"metricName":" # inversions","quality":"Less is better","values":[0,0,0,0,0,0],"isMain":false},{"metricName":" # interspecies translocations","quality":"Less is better","values":[1,1,1,3,2,1],"isMain":false},{"metricName":"# possibly misassembled contigs","quality":"Less is better","values":[0,0,0,0,0,0],"isMain":false},{"metricName":" # possible misassemblies","quality":"Less is better","values":[0,0,0,0,0,0],"isMain":false},{"metricName":"# local misassemblies","quality":"Less is better","values":[0,0,0,0,0,0],"isMain":false},{"metricName":"# scaffold gap ext. mis.","quality":"Less is better","values":[0,0,0,0,0,0],"isMain":false},{"metricName":"# scaffold gap loc. mis.","quality":"Less is better","values":[0,0,0,0,0,0],"isMain":false}]],["Unaligned",[]],["Mismatches",[]],["Statistics without reference",[]],["K-mer-based statistics",[]],["Predicted genes",[]],["Similarity statistics",[]],["Reference statistics",[]]],[["Genome statistics",[]],["Reads mapping",[]],["Misassemblies",[{"metricName":"# misassemblies","quality":"Less is better","values":[126,145,58,75,134,69],"isMain":true},{"metricName":" # relocations","quality":"Less is better","values":[3,2,1,2,2,1],"isMain":false},{"metricName":" # translocations","quality":"Less is better","values":[20,22,15,16,22,9],"isMain":false},{"metricName":" # inversions","quality":"Less is better","values":[0,1,1,2,3,0],"isMain":false},{"metricName":" # interspecies translocations","quality":"Less is better","values":[103,120,41,55,107,59],"isMain":false},{"metricName":"# possibly misassembled contigs","quality":"Less is better","values":[1,0,0,0,1,0],"isMain":false},{"metricName":" # possible misassemblies","quality":"Less is better","values":[1,0,0,0,1,0],"isMain":false},{"metricName":"# local misassemblies","quality":"Less is better","values":[5,6,4,2,3,1],"isMain":false},{"metricName":"# scaffold gap ext. mis.","quality":"Less is better","values":[0,0,0,0,0,0],"isMain":false},{"metricName":"# scaffold gap loc. mis.","quality":"Less is better","values":[0,0,0,0,0,0],"isMain":false}]],["Unaligned",[]],["Mismatches",[]],["Statistics without reference",[]],["K-mer-based statistics",[]],["Predicted genes",[]],["Similarity statistics",[]],["Reference statistics",[]]],[["Genome statistics",[]],["Reads mapping",[]],["Misassemblies",[{"metricName":"# misassemblies","quality":"Less is better","values":[4,1,2,9,7,1],"isMain":true},{"metricName":" # relocations","quality":"Less is better","values":[0,0,0,0,0,0],"isMain":false},{"metricName":" # translocations","quality":"Less is better","values":[0,0,0,0,0,0],"isMain":false},{"metricName":" # inversions","quality":"Less is better","values":[0,0,0,0,0,0],"isMain":false},{"metricName":" # interspecies translocations","quality":"Less is better","values":[4,1,2,9,7,1],"isMain":false},{"metricName":"# possibly misassembled contigs","quality":"Less is better","values":[0,0,0,0,0,0],"isMain":false},{"metricName":" # possible misassemblies","quality":"Less is better","values":[0,0,0,0,0,0],"isMain":false},{"metricName":"# local misassemblies","quality":"Less is better","values":[0,0,0,0,0,0],"isMain":false},{"metricName":"# scaffold gap ext. mis.","quality":"Less is better","values":[0,0,0,0,0,0],"isMain":false},{"metricName":"# scaffold gap loc. mis.","quality":"Less is better","values":[0,0,0,0,0,0],"isMain":false}]],["Unaligned",[]],["Mismatches",[]],["Statistics without reference",[]],["K-mer-based statistics",[]],["Predicted genes",[]],["Similarity statistics",[]],["Reference statistics",[]]],[["Genome statistics",[]],["Reads mapping",[]],["Misassemblies",[{"metricName":"# misassemblies","quality":"Less is better","values":[26,29,11,27,23,24],"isMain":true},{"metricName":" # relocations","quality":"Less is better","values":[3,4,4,6,5,3],"isMain":false},{"metricName":" # translocations","quality":"Less is better","values":[0,0,0,0,0,0],"isMain":false},{"metricName":" # inversions","quality":"Less is better","values":[0,2,2,2,1,0],"isMain":false},{"metricName":" # interspecies translocations","quality":"Less is better","values":[23,23,5,19,17,21],"isMain":false},{"metricName":"# possibly misassembled contigs","quality":"Less is better","values":[0,0,0,0,0,0],"isMain":false},{"metricName":" # possible misassemblies","quality":"Less is better","values":[0,0,0,0,0,0],"isMain":false},{"metricName":"# local misassemblies","quality":"Less is better","values":[3,2,9,5,4,2],"isMain":false},{"metricName":"# scaffold gap ext. mis.","quality":"Less is better","values":[0,0,0,0,0,0],"isMain":false},{"metricName":"# scaffold gap loc. mis.","quality":"Less is better","values":[0,0,0,0,0,0],"isMain":false}]],["Unaligned",[]],["Mismatches",[]],["Statistics without reference",[]],["K-mer-based statistics",[]],["Predicted genes",[]],["Similarity statistics",[]],["Reference statistics",[]]],[["Genome statistics",[]],["Reads mapping",[]],["Misassemblies",[{"metricName":"# misassemblies","quality":"Less is better","values":[61,78,4,38,65,38],"isMain":true},{"metricName":" # relocations","quality":"Less is better","values":[0,1,0,1,0,0],"isMain":false},{"metricName":" # translocations","quality":"Less is better","values":[6,19,0,4,13,3],"isMain":false},{"metricName":" # inversions","quality":"Less is better","values":[1,0,2,1,0,0],"isMain":false},{"metricName":" # interspecies translocations","quality":"Less is better","values":[54,58,2,32,52,35],"isMain":false},{"metricName":"# possibly misassembled contigs","quality":"Less is better","values":[0,0,0,0,0,0],"isMain":false},{"metricName":" # possible misassemblies","quality":"Less is better","values":[0,0,0,0,0,0],"isMain":false},{"metricName":"# local misassemblies","quality":"Less is better","values":[0,0,2,1,2,0],"isMain":false},{"metricName":"# scaffold gap ext. mis.","quality":"Less is better","values":[0,0,0,0,0,0],"isMain":false},{"metricName":"# scaffold gap loc. mis.","quality":"Less is better","values":[0,0,0,0,0,0],"isMain":false}]],["Unaligned",[]],["Mismatches",[]],["Statistics without reference",[]],["K-mer-based statistics",[]],["Predicted genes",[]],["Similarity statistics",[]],["Reference statistics",[]]],[["Genome statistics",[]],["Reads mapping",[]],["Misassemblies",[{"metricName":"# misassemblies","quality":"Less is better","values":[6,12,8,10,9,3],"isMain":true},{"metricName":" # relocations","quality":"Less is better","values":[0,0,1,0,2,1],"isMain":false},{"metricName":" # translocations","quality":"Less is better","values":[0,0,0,0,0,0],"isMain":false},{"metricName":" # inversions","quality":"Less is better","values":[0,1,0,0,0,0],"isMain":false},{"metricName":" # interspecies translocations","quality":"Less is better","values":[6,11,7,10,7,2],"isMain":false},{"metricName":"# possibly misassembled contigs","quality":"Less is better","values":[0,0,0,0,0,0],"isMain":false},{"metricName":" # possible misassemblies","quality":"Less is better","values":[0,0,0,0,0,0],"isMain":false},{"metricName":"# local misassemblies","quality":"Less is better","values":[0,0,0,1,0,0],"isMain":false},{"metricName":"# scaffold gap ext. mis.","quality":"Less is better","values":[0,0,0,0,0,0],"isMain":false},{"metricName":"# scaffold gap loc. mis.","quality":"Less is better","values":[0,0,0,0,0,0],"isMain":false}]],["Unaligned",[]],["Mismatches",[]],["Statistics without reference",[]],["K-mer-based statistics",[]],["Predicted genes",[]],["Similarity statistics",[]],["Reference statistics",[]]],[["Genome statistics",[]],["Reads mapping",[]],["Misassemblies",[{"metricName":"# misassemblies","quality":"Less is better","values":[21,27,8,15,22,12],"isMain":true},{"metricName":" # relocations","quality":"Less is better","values":[0,0,0,1,0,0],"isMain":false},{"metricName":" # translocations","quality":"Less is better","values":[0,0,0,0,0,0],"isMain":false},{"metricName":" # inversions","quality":"Less is better","values":[0,0,0,0,0,0],"isMain":false},{"metricName":" # interspecies translocations","quality":"Less is better","values":[21,27,8,14,22,12],"isMain":false},{"metricName":"# possibly misassembled contigs","quality":"Less is better","values":[0,0,0,0,0,0],"isMain":false},{"metricName":" # possible misassemblies","quality":"Less is better","values":[0,0,0,0,0,0],"isMain":false},{"metricName":"# local misassemblies","quality":"Less is better","values":[0,0,0,0,0,0],"isMain":false},{"metricName":"# scaffold gap ext. mis.","quality":"Less is better","values":[0,0,0,0,0,0],"isMain":false},{"metricName":"# scaffold gap loc. mis.","quality":"Less is better","values":[0,0,0,0,0,0],"isMain":false}]],["Unaligned",[]],["Mismatches",[]],["Statistics without reference",[]],["K-mer-based statistics",[]],["Predicted genes",[]],["Similarity statistics",[]],["Reference statistics",[]]],[["Genome statistics",[]],["Reads mapping",[]],["Misassemblies",[{"metricName":"# misassemblies","quality":"Less is better","values":[2,6,0,8,5,5],"isMain":true},{"metricName":" # relocations","quality":"Less is better","values":[0,0,0,0,0,0],"isMain":false},{"metricName":" # translocations","quality":"Less is better","values":[0,0,0,0,0,0],"isMain":false},{"metricName":" # inversions","quality":"Less is better","values":[0,0,0,0,0,0],"isMain":false},{"metricName":" # interspecies translocations","quality":"Less is better","values":[2,6,0,8,5,5],"isMain":false},{"metricName":"# possibly misassembled contigs","quality":"Less is better","values":[0,0,0,0,0,0],"isMain":false},{"metricName":" # possible misassemblies","quality":"Less is better","values":[0,0,0,0,0,0],"isMain":false},{"metricName":"# local misassemblies","quality":"Less is better","values":[0,0,0,0,0,0],"isMain":false},{"metricName":"# scaffold gap ext. mis.","quality":"Less is better","values":[0,0,0,0,0,0],"isMain":false},{"metricName":"# scaffold gap loc. mis.","quality":"Less is better","values":[0,0,0,0,0,0],"isMain":false}]],["Unaligned",[]],["Mismatches",[]],["Statistics without reference",[]],["K-mer-based statistics",[]],["Predicted genes",[]],["Similarity statistics",[]],["Reference statistics",[]]],[["Genome statistics",[]],["Reads mapping",[]],["Misassemblies",[{"metricName":"# misassemblies","quality":"Less is better","values":[10,12,5,7,12,5],"isMain":true},{"metricName":" # relocations","quality":"Less is better","values":[0,0,0,0,1,0],"isMain":false},{"metricName":" # translocations","quality":"Less is better","values":[3,6,2,3,4,4],"isMain":false},{"metricName":" # inversions","quality":"Less is better","values":[0,0,0,0,0,0],"isMain":false},{"metricName":" # interspecies translocations","quality":"Less is better","values":[7,6,3,4,7,1],"isMain":false},{"metricName":"# possibly misassembled contigs","quality":"Less is better","values":[0,0,0,0,0,0],"isMain":false},{"metricName":" # possible misassemblies","quality":"Less is better","values":[0,0,0,0,0,0],"isMain":false},{"metricName":"# local misassemblies","quality":"Less is better","values":[0,1,0,1,0,0],"isMain":false},{"metricName":"# scaffold gap ext. mis.","quality":"Less is better","values":[0,0,0,0,0,0],"isMain":false},{"metricName":"# scaffold gap loc. mis.","quality":"Less is better","values":[0,0,0,0,0,0],"isMain":false}]],["Unaligned",[]],["Mismatches",[]],["Statistics without reference",[]],["K-mer-based statistics",[]],["Predicted genes",[]],["Similarity statistics",[]],["Reference statistics",[]]],[["Genome statistics",[]],["Reads mapping",[]],["Misassemblies",[{"metricName":"# misassemblies","quality":"Less is better","values":[19,20,2,20,17,9],"isMain":true},{"metricName":" # relocations","quality":"Less is better","values":[0,0,0,0,0,0],"isMain":false},{"metricName":" # translocations","quality":"Less is better","values":[0,0,0,0,0,0],"isMain":false},{"metricName":" # inversions","quality":"Less is better","values":[0,0,0,0,0,0],"isMain":false},{"metricName":" # interspecies translocations","quality":"Less is better","values":[19,20,2,20,17,9],"isMain":false},{"metricName":"# possibly misassembled contigs","quality":"Less is better","values":[0,0,0,0,0,0],"isMain":false},{"metricName":" # possible misassemblies","quality":"Less is better","values":[0,0,0,0,0,0],"isMain":false},{"metricName":"# local misassemblies","quality":"Less is better","values":[0,0,0,1,0,0],"isMain":false},{"metricName":"# scaffold gap ext. mis.","quality":"Less is better","values":[0,0,0,0,0,0],"isMain":false},{"metricName":"# scaffold gap loc. mis.","quality":"Less is better","values":[0,0,0,0,0,0],"isMain":false}]],["Unaligned",[]],["Mismatches",[]],["Statistics without reference",[]],["K-mer-based statistics",[]],["Predicted genes",[]],["Similarity statistics",[]],["Reference statistics",[]]],[["Genome statistics",[]],["Reads mapping",[]],["Misassemblies",[{"metricName":"# misassemblies","quality":"Less is better","values":[26,5,1,10,7,1],"isMain":true},{"metricName":" # relocations","quality":"Less is better","values":[0,0,0,0,0,0],"isMain":false},{"metricName":" # translocations","quality":"Less is better","values":[0,0,0,0,0,0],"isMain":false},{"metricName":" # inversions","quality":"Less is better","values":[0,0,0,0,0,0],"isMain":false},{"metricName":" # interspecies translocations","quality":"Less is better","values":[26,5,1,10,7,1],"isMain":false},{"metricName":"# possibly misassembled contigs","quality":"Less is better","values":[0,0,0,0,0,0],"isMain":false},{"metricName":" # possible misassemblies","quality":"Less is better","values":[0,0,0,0,0,0],"isMain":false},{"metricName":"# local misassemblies","quality":"Less is better","values":[0,0,0,1,0,0],"isMain":false},{"metricName":"# scaffold gap ext. mis.","quality":"Less is better","values":[0,0,0,0,0,0],"isMain":false},{"metricName":"# scaffold gap loc. mis.","quality":"Less is better","values":[0,0,0,0,0,0],"isMain":false}]],["Unaligned",[]],["Mismatches",[]],["Statistics without reference",[]],["K-mer-based statistics",[]],["Predicted genes",[]],["Similarity statistics",[]],["Reference statistics",[]]],[["Genome statistics",[]],["Reads mapping",[]],["Misassemblies",[{"metricName":"# misassemblies","quality":"Less is better","values":[21,25,23,32,40,8],"isMain":true},{"metricName":" # relocations","quality":"Less is better","values":[1,1,1,0,1,0],"isMain":false},{"metricName":" # translocations","quality":"Less is better","values":[2,0,2,1,2,0],"isMain":false},{"metricName":" # inversions","quality":"Less is better","values":[1,1,1,1,2,0],"isMain":false},{"metricName":" # interspecies translocations","quality":"Less is better","values":[17,23,19,30,35,8],"isMain":false},{"metricName":"# possibly misassembled contigs","quality":"Less is better","values":[0,0,0,0,0,0],"isMain":false},{"metricName":" # possible misassemblies","quality":"Less is better","values":[0,0,0,0,0,0],"isMain":false},{"metricName":"# local misassemblies","quality":"Less is better","values":[1,7,4,4,3,1],"isMain":false},{"metricName":"# scaffold gap ext. mis.","quality":"Less is better","values":[0,0,0,0,0,0],"isMain":false},{"metricName":"# scaffold gap loc. mis.","quality":"Less is better","values":[0,0,0,0,0,0],"isMain":false}]],["Unaligned",[]],["Mismatches",[]],["Statistics without reference",[]],["K-mer-based statistics",[]],["Predicted genes",[]],["Similarity statistics",[]],["Reference statistics",[]]],[["Genome statistics",[]],["Reads mapping",[]],["Misassemblies",[{"metricName":"# misassemblies","quality":"Less is better","values":[60,82,37,68,97,33],"isMain":true},{"metricName":" # relocations","quality":"Less is better","values":[1,0,1,1,2,0],"isMain":false},{"metricName":" # translocations","quality":"Less is better","values":[0,0,0,0,1,0],"isMain":false},{"metricName":" # inversions","quality":"Less is better","values":[1,0,0,0,0,0],"isMain":false},{"metricName":" # interspecies translocations","quality":"Less is better","values":[58,82,36,67,94,33],"isMain":false},{"metricName":"# possibly misassembled contigs","quality":"Less is better","values":[0,0,0,0,0,0],"isMain":false},{"metricName":" # possible misassemblies","quality":"Less is better","values":[0,0,0,0,0,0],"isMain":false},{"metricName":"# local misassemblies","quality":"Less is better","values":[2,1,3,2,1,0],"isMain":false},{"metricName":"# scaffold gap ext. mis.","quality":"Less is better","values":[0,0,0,0,0,0],"isMain":false},{"metricName":"# scaffold gap loc. mis.","quality":"Less is better","values":[0,0,0,0,0,0],"isMain":false}]],["Unaligned",[]],["Mismatches",[]],["Statistics without reference",[]],["K-mer-based statistics",[]],["Predicted genes",[]],["Similarity statistics",[]],["Reference statistics",[]]],[["Genome statistics",[]],["Reads mapping",[]],["Misassemblies",[{"metricName":"# misassemblies","quality":"Less is better","values":[7,7,5,7,8,4],"isMain":true},{"metricName":" # relocations","quality":"Less is better","values":[0,0,0,0,0,0],"isMain":false},{"metricName":" # translocations","quality":"Less is better","values":[1,0,1,2,1,0],"isMain":false},{"metricName":" # inversions","quality":"Less is better","values":[0,0,0,0,0,0],"isMain":false},{"metricName":" # interspecies translocations","quality":"Less is better","values":[6,7,4,5,7,4],"isMain":false},{"metricName":"# possibly misassembled contigs","quality":"Less is better","values":[0,0,0,0,0,0],"isMain":false},{"metricName":" # possible misassemblies","quality":"Less is better","values":[0,0,0,0,0,0],"isMain":false},{"metricName":"# local misassemblies","quality":"Less is better","values":[0,0,0,0,0,0],"isMain":false},{"metricName":"# scaffold gap ext. mis.","quality":"Less is better","values":[0,0,0,0,0,0],"isMain":false},{"metricName":"# scaffold gap loc. mis.","quality":"Less is better","values":[0,0,0,0,0,0],"isMain":false}]],["Unaligned",[]],["Mismatches",[]],["Statistics without reference",[]],["K-mer-based statistics",[]],["Predicted genes",[]],["Similarity statistics",[]],["Reference statistics",[]]],[["Genome statistics",[]],["Reads mapping",[]],["Misassemblies",[{"metricName":"# misassemblies","quality":"Less is better","values":[0,1,1,0,1,0],"isMain":true},{"metricName":" # relocations","quality":"Less is better","values":[0,0,0,0,0,0],"isMain":false},{"metricName":" # translocations","quality":"Less is better","values":[0,0,0,0,0,0],"isMain":false},{"metricName":" # inversions","quality":"Less is better","values":[0,0,0,0,0,0],"isMain":false},{"metricName":" # interspecies translocations","quality":"Less is better","values":[0,1,1,0,1,0],"isMain":false},{"metricName":"# possibly misassembled contigs","quality":"Less is better","values":[0,0,0,0,0,0],"isMain":false},{"metricName":" # possible misassemblies","quality":"Less is better","values":[0,0,0,0,0,0],"isMain":false},{"metricName":"# local misassemblies","quality":"Less is better","values":[0,0,0,0,0,0],"isMain":false},{"metricName":"# scaffold gap ext. mis.","quality":"Less is better","values":[0,0,0,0,0,0],"isMain":false},{"metricName":"# scaffold gap loc. mis.","quality":"Less is better","values":[0,0,0,0,0,0],"isMain":false}]],["Unaligned",[]],["Mismatches",[]],["Statistics without reference",[]],["K-mer-based statistics",[]],["Predicted genes",[]],["Similarity statistics",[]],["Reference statistics",[]]],[["Genome statistics",[]],["Reads mapping",[]],["Misassemblies",[{"metricName":"# misassemblies","quality":"Less is better","values":[97,21,23,41,37,5],"isMain":true},{"metricName":" # relocations","quality":"Less is better","values":[0,1,1,5,3,0],"isMain":false},{"metricName":" # translocations","quality":"Less is better","values":[1,8,6,8,11,3],"isMain":false},{"metricName":" # inversions","quality":"Less is better","values":[0,0,0,1,1,0],"isMain":false},{"metricName":" # interspecies translocations","quality":"Less is better","values":[96,12,16,27,22,2],"isMain":false},{"metricName":"# possibly misassembled contigs","quality":"Less is better","values":[0,0,0,0,0,0],"isMain":false},{"metricName":" # possible misassemblies","quality":"Less is better","values":[0,0,0,0,0,0],"isMain":false},{"metricName":"# local misassemblies","quality":"Less is better","values":[0,1,2,3,2,0],"isMain":false},{"metricName":"# scaffold gap ext. mis.","quality":"Less is better","values":[0,0,0,0,0,0],"isMain":false},{"metricName":"# scaffold gap loc. mis.","quality":"Less is better","values":[0,0,0,0,0,0],"isMain":false}]],["Unaligned",[]],["Mismatches",[]],["Statistics without reference",[]],["K-mer-based statistics",[]],["Predicted genes",[]],["Similarity statistics",[]],["Reference statistics",[]]],[["Genome statistics",[]],["Reads mapping",[]],["Misassemblies",[{"metricName":"# misassemblies","quality":"Less is better","values":[5,1,3,3,3,1],"isMain":true},{"metricName":" # relocations","quality":"Less is better","values":[0,0,0,0,0,0],"isMain":false},{"metricName":" # translocations","quality":"Less is better","values":[0,0,0,0,0,0],"isMain":false},{"metricName":" # inversions","quality":"Less is better","values":[0,0,0,0,0,0],"isMain":false},{"metricName":" # interspecies translocations","quality":"Less is better","values":[5,1,3,3,3,1],"isMain":false},{"metricName":"# possibly misassembled contigs","quality":"Less is better","values":[0,0,0,1,0,0],"isMain":false},{"metricName":" # possible misassemblies","quality":"Less is better","values":[0,0,0,1,0,0],"isMain":false},{"metricName":"# local misassemblies","quality":"Less is better","values":[0,0,0,0,0,0],"isMain":false},{"metricName":"# scaffold gap ext. mis.","quality":"Less is better","values":[0,0,0,0,0,0],"isMain":false},{"metricName":"# scaffold gap loc. mis.","quality":"Less is better","values":[0,0,0,0,0,0],"isMain":false}]],["Unaligned",[]],["Mismatches",[]],["Statistics without reference",[]],["K-mer-based statistics",[]],["Predicted genes",[]],["Similarity statistics",[]],["Reference statistics",[]]],[["Genome statistics",[]],["Reads mapping",[]],["Misassemblies",[{"metricName":"# misassemblies","quality":"Less is better","values":[25,25,20,36,39,13],"isMain":true},{"metricName":" # relocations","quality":"Less is better","values":[0,1,1,1,2,0],"isMain":false},{"metricName":" # translocations","quality":"Less is better","values":[3,14,11,17,28,12],"isMain":false},{"metricName":" # inversions","quality":"Less is better","values":[0,0,0,0,0,0],"isMain":false},{"metricName":" # interspecies translocations","quality":"Less is better","values":[22,10,8,18,9,1],"isMain":false},{"metricName":"# possibly misassembled contigs","quality":"Less is better","values":[0,0,0,0,1,0],"isMain":false},{"metricName":" # possible misassemblies","quality":"Less is better","values":[0,0,0,0,1,0],"isMain":false},{"metricName":"# local misassemblies","quality":"Less is better","values":[0,0,0,2,1,0],"isMain":false},{"metricName":"# scaffold gap ext. mis.","quality":"Less is better","values":[0,0,0,0,0,0],"isMain":false},{"metricName":"# scaffold gap loc. mis.","quality":"Less is better","values":[0,0,0,0,0,0],"isMain":false}]],["Unaligned",[]],["Mismatches",[]],["Statistics without reference",[]],["K-mer-based statistics",[]],["Predicted genes",[]],["Similarity statistics",[]],["Reference statistics",[]]],[["Genome statistics",[]],["Reads mapping",[]],["Misassemblies",[{"metricName":"# misassemblies","quality":"Less is better","values":[49,63,56,70,87,20],"isMain":true},{"metricName":" # relocations","quality":"Less is better","values":[2,5,2,2,4,0],"isMain":false},{"metricName":" # translocations","quality":"Less is better","values":[4,2,8,9,8,2],"isMain":false},{"metricName":" # inversions","quality":"Less is better","values":[2,2,3,4,1,0],"isMain":false},{"metricName":" # interspecies translocations","quality":"Less is better","values":[41,54,43,55,74,18],"isMain":false},{"metricName":"# possibly misassembled contigs","quality":"Less is better","values":[0,0,1,0,0,0],"isMain":false},{"metricName":" # possible misassemblies","quality":"Less is better","values":[0,0,1,0,0,0],"isMain":false},{"metricName":"# local misassemblies","quality":"Less is better","values":[6,19,9,7,17,3],"isMain":false},{"metricName":"# scaffold gap ext. mis.","quality":"Less is better","values":[0,0,0,0,0,0],"isMain":false},{"metricName":"# scaffold gap loc. mis.","quality":"Less is better","values":[0,0,0,0,0,0],"isMain":false}]],["Unaligned",[]],["Mismatches",[]],["Statistics without reference",[]],["K-mer-based statistics",[]],["Predicted genes",[]],["Similarity statistics",[]],["Reference statistics",[]]],[["Genome statistics",[]],["Reads mapping",[]],["Misassemblies",[{"metricName":"# misassemblies","quality":"Less is better","values":[38,64,70,83,90,20],"isMain":true},{"metricName":" # relocations","quality":"Less is better","values":[0,1,1,1,1,1],"isMain":false},{"metricName":" # translocations","quality":"Less is better","values":[4,7,12,10,11,1],"isMain":false},{"metricName":" # inversions","quality":"Less is better","values":[2,4,2,3,3,0],"isMain":false},{"metricName":" # interspecies translocations","quality":"Less is better","values":[32,52,55,69,75,18],"isMain":false},{"metricName":"# possibly misassembled contigs","quality":"Less is better","values":[0,1,0,0,0,1],"isMain":false},{"metricName":" # possible misassemblies","quality":"Less is better","values":[0,2,0,0,0,2],"isMain":false},{"metricName":"# local misassemblies","quality":"Less is better","values":[5,11,10,18,19,4],"isMain":false},{"metricName":"# scaffold gap ext. mis.","quality":"Less is better","values":[0,0,0,0,0,0],"isMain":false},{"metricName":"# scaffold gap loc. mis.","quality":"Less is better","values":[0,0,0,0,0,0],"isMain":false}]],["Unaligned",[]],["Mismatches",[]],["Statistics without reference",[]],["K-mer-based statistics",[]],["Predicted genes",[]],["Similarity statistics",[]],["Reference statistics",[]]],[["Genome statistics",[]],["Reads mapping",[]],["Misassemblies",[{"metricName":"# misassemblies","quality":"Less is better","values":[13,42,25,63,41,2],"isMain":true},{"metricName":" # relocations","quality":"Less is better","values":[0,1,0,2,0,0],"isMain":false},{"metricName":" # translocations","quality":"Less is better","values":[3,4,6,6,5,1],"isMain":false},{"metricName":" # inversions","quality":"Less is better","values":[0,0,0,0,0,0],"isMain":false},{"metricName":" # interspecies translocations","quality":"Less is better","values":[10,37,19,55,36,1],"isMain":false},{"metricName":"# possibly misassembled contigs","quality":"Less is better","values":[0,0,0,0,0,0],"isMain":false},{"metricName":" # possible misassemblies","quality":"Less is better","values":[0,0,0,0,0,0],"isMain":false},{"metricName":"# local misassemblies","quality":"Less is better","values":[1,8,3,3,4,0],"isMain":false},{"metricName":"# scaffold gap ext. mis.","quality":"Less is better","values":[0,0,0,0,0,0],"isMain":false},{"metricName":"# scaffold gap loc. mis.","quality":"Less is better","values":[0,0,0,0,0,0],"isMain":false}]],["Unaligned",[]],["Mismatches",[]],["Statistics without reference",[]],["K-mer-based statistics",[]],["Predicted genes",[]],["Similarity statistics",[]],["Reference statistics",[]]],[["Genome statistics",[]],["Reads mapping",[]],["Misassemblies",[{"metricName":"# misassemblies","quality":"Less is better","values":[1,7,2,9,10,4],"isMain":true},{"metricName":" # relocations","quality":"Less is better","values":[0,0,0,0,0,0],"isMain":false},{"metricName":" # translocations","quality":"Less is better","values":[0,1,0,1,0,1],"isMain":false},{"metricName":" # inversions","quality":"Less is better","values":[0,0,0,0,0,0],"isMain":false},{"metricName":" # interspecies translocations","quality":"Less is better","values":[1,6,2,8,10,3],"isMain":false},{"metricName":"# possibly misassembled contigs","quality":"Less is better","values":[0,0,0,0,0,0],"isMain":false},{"metricName":" # possible misassemblies","quality":"Less is better","values":[0,0,0,0,0,0],"isMain":false},{"metricName":"# local misassemblies","quality":"Less is better","values":[0,0,0,0,0,0],"isMain":false},{"metricName":"# scaffold gap ext. mis.","quality":"Less is better","values":[0,0,0,0,0,0],"isMain":false},{"metricName":"# scaffold gap loc. mis.","quality":"Less is better","values":[0,0,0,0,0,0],"isMain":false}]],["Unaligned",[]],["Mismatches",[]],["Statistics without reference",[]],["K-mer-based statistics",[]],["Predicted genes",[]],["Similarity statistics",[]],["Reference statistics",[]]],[["Genome statistics",[]],["Reads mapping",[]],["Misassemblies",[{"metricName":"# misassemblies","quality":"Less is better","values":[3,3,1,7,2,2],"isMain":true},{"metricName":" # relocations","quality":"Less is better","values":[0,0,0,0,0,0],"isMain":false},{"metricName":" # translocations","quality":"Less is better","values":[0,0,0,0,0,0],"isMain":false},{"metricName":" # inversions","quality":"Less is better","values":[0,0,0,0,0,0],"isMain":false},{"metricName":" # interspecies translocations","quality":"Less is better","values":[3,3,1,7,2,2],"isMain":false},{"metricName":"# possibly misassembled contigs","quality":"Less is better","values":[0,0,0,0,0,0],"isMain":false},{"metricName":" # possible misassemblies","quality":"Less is better","values":[0,0,0,0,0,0],"isMain":false},{"metricName":"# local misassemblies","quality":"Less is better","values":[0,0,0,1,0,0],"isMain":false},{"metricName":"# scaffold gap ext. mis.","quality":"Less is better","values":[0,0,0,0,0,0],"isMain":false},{"metricName":"# scaffold gap loc. mis.","quality":"Less is better","values":[0,0,0,0,0,0],"isMain":false}]],["Unaligned",[]],["Mismatches",[]],["Statistics without reference",[]],["K-mer-based statistics",[]],["Predicted genes",[]],["Similarity statistics",[]],["Reference statistics",[]]],[["Genome statistics",[]],["Reads mapping",[]],["Misassemblies",[{"metricName":"# misassemblies","quality":"Less is better","values":[0,0,0,0,1,1],"isMain":true},{"metricName":" # relocations","quality":"Less is better","values":[0,0,0,0,0,0],"isMain":false},{"metricName":" # translocations","quality":"Less is better","values":[0,0,0,0,0,0],"isMain":false},{"metricName":" # inversions","quality":"Less is better","values":[0,0,0,0,0,0],"isMain":false},{"metricName":" # interspecies translocations","quality":"Less is better","values":[0,0,0,0,1,1],"isMain":false},{"metricName":"# possibly misassembled contigs","quality":"Less is better","values":[0,0,0,0,0,0],"isMain":false},{"metricName":" # possible misassemblies","quality":"Less is better","values":[0,0,0,0,0,0],"isMain":false},{"metricName":"# local misassemblies","quality":"Less is better","values":[0,0,0,0,0,0],"isMain":false},{"metricName":"# scaffold gap ext. mis.","quality":"Less is better","values":[0,0,0,0,0,0],"isMain":false},{"metricName":"# scaffold gap loc. mis.","quality":"Less is better","values":[0,0,0,0,0,0],"isMain":false}]],["Unaligned",[]],["Mismatches",[]],["Statistics without reference",[]],["K-mer-based statistics",[]],["Predicted genes",[]],["Similarity statistics",[]],["Reference statistics",[]]],[["Genome statistics",[]],["Reads mapping",[]],["Misassemblies",[{"metricName":"# misassemblies","quality":"Less is better","values":[89,95,14,64,65,81],"isMain":true},{"metricName":" # relocations","quality":"Less is better","values":[5,1,2,4,2,2],"isMain":false},{"metricName":" # translocations","quality":"Less is better","values":[6,12,7,6,7,7],"isMain":false},{"metricName":" # inversions","quality":"Less is better","values":[0,2,3,2,1,0],"isMain":false},{"metricName":" # interspecies translocations","quality":"Less is better","values":[78,80,2,52,55,72],"isMain":false},{"metricName":"# possibly misassembled contigs","quality":"Less is better","values":[1,0,0,0,0,0],"isMain":false},{"metricName":" # possible misassemblies","quality":"Less is better","values":[1,0,0,0,0,0],"isMain":false},{"metricName":"# local misassemblies","quality":"Less is better","values":[2,2,9,6,2,1],"isMain":false},{"metricName":"# scaffold gap ext. mis.","quality":"Less is better","values":[0,0,0,0,0,0],"isMain":false},{"metricName":"# scaffold gap loc. mis.","quality":"Less is better","values":[0,0,0,0,0,0],"isMain":false}]],["Unaligned",[]],["Mismatches",[]],["Statistics without reference",[]],["K-mer-based statistics",[]],["Predicted genes",[]],["Similarity statistics",[]],["Reference statistics",[]]],[["Genome statistics",[]],["Reads mapping",[]],["Misassemblies",[{"metricName":"# misassemblies","quality":"Less is better","values":[21,18,7,56,17,12],"isMain":true},{"metricName":" # relocations","quality":"Less is better","values":[0,0,0,1,0,0],"isMain":false},{"metricName":" # translocations","quality":"Less is better","values":[5,4,2,5,5,4],"isMain":false},{"metricName":" # inversions","quality":"Less is better","values":[0,2,2,1,0,0],"isMain":false},{"metricName":" # interspecies translocations","quality":"Less is better","values":[16,12,3,49,12,8],"isMain":false},{"metricName":"# possibly misassembled contigs","quality":"Less is better","values":[0,0,0,0,0,0],"isMain":false},{"metricName":" # possible misassemblies","quality":"Less is better","values":[0,0,0,0,0,0],"isMain":false},{"metricName":"# local misassemblies","quality":"Less is better","values":[0,4,6,6,2,0],"isMain":false},{"metricName":"# scaffold gap ext. mis.","quality":"Less is better","values":[0,0,0,0,0,0],"isMain":false},{"metricName":"# scaffold gap loc. mis.","quality":"Less is better","values":[0,0,0,0,0,0],"isMain":false}]],["Unaligned",[]],["Mismatches",[]],["Statistics without reference",[]],["K-mer-based statistics",[]],["Predicted genes",[]],["Similarity statistics",[]],["Reference statistics",[]]],[["Genome statistics",[]],["Reads mapping",[]],["Misassemblies",[{"metricName":"# misassemblies","quality":"Less is better","values":[8,14,9,35,18,0],"isMain":true},{"metricName":" # relocations","quality":"Less is better","values":[2,4,2,14,2,0],"isMain":false},{"metricName":" # translocations","quality":"Less is better","values":[0,6,4,9,4,0],"isMain":false},{"metricName":" # inversions","quality":"Less is better","values":[0,0,0,3,3,0],"isMain":false},{"metricName":" # interspecies translocations","quality":"Less is better","values":[6,4,3,9,9,0],"isMain":false},{"metricName":"# possibly misassembled contigs","quality":"Less is better","values":[0,0,0,0,0,0],"isMain":false},{"metricName":" # possible misassemblies","quality":"Less is better","values":[0,0,0,0,0,0],"isMain":false},{"metricName":"# local misassemblies","quality":"Less is better","values":[0,0,0,4,1,0],"isMain":false},{"metricName":"# scaffold gap ext. mis.","quality":"Less is better","values":[0,0,0,0,0,0],"isMain":false},{"metricName":"# scaffold gap loc. mis.","quality":"Less is better","values":[0,0,0,0,0,0],"isMain":false}]],["Unaligned",[]],["Mismatches",[]],["Statistics without reference",[]],["K-mer-based statistics",[]],["Predicted genes",[]],["Similarity statistics",[]],["Reference statistics",[]]],[["Genome statistics",[]],["Reads mapping",[]],["Misassemblies",[{"metricName":"# misassemblies","quality":"Less is better","values":[19,5,0,3,3,0],"isMain":true},{"metricName":" # relocations","quality":"Less is better","values":[0,0,0,0,0,0],"isMain":false},{"metricName":" # translocations","quality":"Less is better","values":[0,0,0,0,0,0],"isMain":false},{"metricName":" # inversions","quality":"Less is better","values":[0,0,0,0,0,0],"isMain":false},{"metricName":" # interspecies translocations","quality":"Less is better","values":[19,5,0,3,3,0],"isMain":false},{"metricName":"# possibly misassembled contigs","quality":"Less is better","values":[0,0,0,0,0,0],"isMain":false},{"metricName":" # possible misassemblies","quality":"Less is better","values":[0,0,0,0,0,0],"isMain":false},{"metricName":"# local misassemblies","quality":"Less is better","values":[0,0,0,0,0,0],"isMain":false},{"metricName":"# scaffold gap ext. mis.","quality":"Less is better","values":[0,0,0,0,0,0],"isMain":false},{"metricName":"# scaffold gap loc. mis.","quality":"Less is better","values":[0,0,0,0,0,0],"isMain":false}]],["Unaligned",[]],["Mismatches",[]],["Statistics without reference",[]],["K-mer-based statistics",[]],["Predicted genes",[]],["Similarity statistics",[]],["Reference statistics",[]]],[["Genome statistics",[]],["Reads mapping",[]],["Misassemblies",[{"metricName":"# misassemblies","quality":"Less is better","values":[29,31,7,29,23,23],"isMain":true},{"metricName":" # relocations","quality":"Less is better","values":[0,0,0,0,0,0],"isMain":false},{"metricName":" # translocations","quality":"Less is better","values":[0,2,1,1,1,0],"isMain":false},{"metricName":" # inversions","quality":"Less is better","values":[0,0,1,1,0,0],"isMain":false},{"metricName":" # interspecies translocations","quality":"Less is better","values":[29,29,5,27,22,23],"isMain":false},{"metricName":"# possibly misassembled contigs","quality":"Less is better","values":[0,0,0,0,0,0],"isMain":false},{"metricName":" # possible misassemblies","quality":"Less is better","values":[0,0,0,0,0,0],"isMain":false},{"metricName":"# local misassemblies","quality":"Less is better","values":[1,0,1,1,0,0],"isMain":false},{"metricName":"# scaffold gap ext. mis.","quality":"Less is better","values":[0,0,0,0,0,0],"isMain":false},{"metricName":"# scaffold gap loc. mis.","quality":"Less is better","values":[0,0,0,0,0,0],"isMain":false}]],["Unaligned",[]],["Mismatches",[]],["Statistics without reference",[]],["K-mer-based statistics",[]],["Predicted genes",[]],["Similarity statistics",[]],["Reference statistics",[]]],[["Genome statistics",[]],["Reads mapping",[]],["Misassemblies",[{"metricName":"# misassemblies","quality":"Less is better","values":[42,55,7,38,56,47],"isMain":true},{"metricName":" # relocations","quality":"Less is better","values":[0,0,0,0,0,0],"isMain":false},{"metricName":" # translocations","quality":"Less is better","values":[1,4,0,2,3,4],"isMain":false},{"metricName":" # inversions","quality":"Less is better","values":[0,0,0,0,0,0],"isMain":false},{"metricName":" # interspecies translocations","quality":"Less is better","values":[41,51,7,36,53,43],"isMain":false},{"metricName":"# possibly misassembled contigs","quality":"Less is better","values":[0,0,0,0,0,0],"isMain":false},{"metricName":" # possible misassemblies","quality":"Less is better","values":[0,0,0,0,0,0],"isMain":false},{"metricName":"# local misassemblies","quality":"Less is better","values":[1,1,0,1,1,0],"isMain":false},{"metricName":"# scaffold gap ext. mis.","quality":"Less is better","values":[0,0,0,0,0,0],"isMain":false},{"metricName":"# scaffold gap loc. mis.","quality":"Less is better","values":[0,0,0,0,0,0],"isMain":false}]],["Unaligned",[]],["Mismatches",[]],["Statistics without reference",[]],["K-mer-based statistics",[]],["Predicted genes",[]],["Similarity statistics",[]],["Reference statistics",[]]],[["Genome statistics",[]],["Reads mapping",[]],["Misassemblies",[{"metricName":"# misassemblies","quality":"Less is better","values":[2,0,2,7,0,0],"isMain":true},{"metricName":" # relocations","quality":"Less is better","values":[0,0,0,0,0,0],"isMain":false},{"metricName":" # translocations","quality":"Less is better","values":[0,0,0,0,0,0],"isMain":false},{"metricName":" # inversions","quality":"Less is better","values":[0,0,0,0,0,0],"isMain":false},{"metricName":" # interspecies translocations","quality":"Less is better","values":[2,0,2,7,0,0],"isMain":false},{"metricName":"# possibly misassembled contigs","quality":"Less is better","values":[0,0,0,0,0,0],"isMain":false},{"metricName":" # possible misassemblies","quality":"Less is better","values":[0,0,0,0,0,0],"isMain":false},{"metricName":"# local misassemblies","quality":"Less is better","values":[0,0,0,0,0,0],"isMain":false},{"metricName":"# scaffold gap ext. mis.","quality":"Less is better","values":[0,0,0,0,0,0],"isMain":false},{"metricName":"# scaffold gap loc. mis.","quality":"Less is better","values":[0,0,0,0,0,0],"isMain":false}]],["Unaligned",[]],["Mismatches",[]],["Statistics without reference",[]],["K-mer-based statistics",[]],["Predicted genes",[]],["Similarity statistics",[]],["Reference statistics",[]]],[["Genome statistics",[]],["Reads mapping",[]],["Misassemblies",[{"metricName":"# misassemblies","quality":"Less is better","values":[14,18,31,30,42,6],"isMain":true},{"metricName":" # relocations","quality":"Less is better","values":[0,0,1,0,0,0],"isMain":false},{"metricName":" # translocations","quality":"Less is better","values":[3,3,4,4,8,3],"isMain":false},{"metricName":" # inversions","quality":"Less is better","values":[0,1,1,1,1,0],"isMain":false},{"metricName":" # interspecies translocations","quality":"Less is better","values":[11,14,25,25,33,3],"isMain":false},{"metricName":"# possibly misassembled contigs","quality":"Less is better","values":[0,0,0,1,1,0],"isMain":false},{"metricName":" # possible misassemblies","quality":"Less is better","values":[0,0,0,1,1,0],"isMain":false},{"metricName":"# local misassemblies","quality":"Less is better","values":[1,2,1,0,0,0],"isMain":false},{"metricName":"# scaffold gap ext. mis.","quality":"Less is better","values":[0,0,0,0,0,0],"isMain":false},{"metricName":"# scaffold gap loc. mis.","quality":"Less is better","values":[0,0,0,0,0,0],"isMain":false}]],["Unaligned",[]],["Mismatches",[]],["Statistics without reference",[]],["K-mer-based statistics",[]],["Predicted genes",[]],["Similarity statistics",[]],["Reference statistics",[]]],[["Genome statistics",[]],["Reads mapping",[]],["Misassemblies",[{"metricName":"# misassemblies","quality":"Less is better","values":[1,2,0,5,11,0],"isMain":true},{"metricName":" # relocations","quality":"Less is better","values":[0,0,0,0,0,0],"isMain":false},{"metricName":" # translocations","quality":"Less is better","values":[0,0,0,0,1,0],"isMain":false},{"metricName":" # inversions","quality":"Less is better","values":[0,0,0,0,0,0],"isMain":false},{"metricName":" # interspecies translocations","quality":"Less is better","values":[1,2,0,5,10,0],"isMain":false},{"metricName":"# possibly misassembled contigs","quality":"Less is better","values":[0,0,0,0,0,0],"isMain":false},{"metricName":" # possible misassemblies","quality":"Less is better","values":[0,0,0,0,0,0],"isMain":false},{"metricName":"# local misassemblies","quality":"Less is better","values":[1,0,0,1,1,0],"isMain":false},{"metricName":"# scaffold gap ext. mis.","quality":"Less is better","values":[0,0,0,0,0,0],"isMain":false},{"metricName":"# scaffold gap loc. mis.","quality":"Less is better","values":[0,0,0,0,0,0],"isMain":false}]],["Unaligned",[]],["Mismatches",[]],["Statistics without reference",[]],["K-mer-based statistics",[]],["Predicted genes",[]],["Similarity statistics",[]],["Reference statistics",[]]],[["Genome statistics",[]],["Reads mapping",[]],["Misassemblies",[{"metricName":"# misassemblies","quality":"Less is better","values":[4,14,12,18,20,9],"isMain":true},{"metricName":" # relocations","quality":"Less is better","values":[0,0,0,1,0,0],"isMain":false},{"metricName":" # translocations","quality":"Less is better","values":[1,2,3,3,3,3],"isMain":false},{"metricName":" # inversions","quality":"Less is better","values":[0,1,1,1,0,0],"isMain":false},{"metricName":" # interspecies translocations","quality":"Less is better","values":[3,11,8,13,17,6],"isMain":false},{"metricName":"# possibly misassembled contigs","quality":"Less is better","values":[0,0,0,0,0,0],"isMain":false},{"metricName":" # possible misassemblies","quality":"Less is better","values":[0,0,0,0,0,0],"isMain":false},{"metricName":"# local misassemblies","quality":"Less is better","values":[1,3,4,1,3,1],"isMain":false},{"metricName":"# scaffold gap ext. mis.","quality":"Less is better","values":[0,0,0,0,0,0],"isMain":false},{"metricName":"# scaffold gap loc. mis.","quality":"Less is better","values":[0,0,0,0,0,0],"isMain":false}]],["Unaligned",[]],["Mismatches",[]],["Statistics without reference",[]],["K-mer-based statistics",[]],["Predicted genes",[]],["Similarity statistics",[]],["Reference statistics",[]]],[["Genome statistics",[]],["Reads mapping",[]],["Misassemblies",[{"metricName":"# misassemblies","quality":"Less is better","values":[59,226,170,253,193,45],"isMain":true},{"metricName":" # relocations","quality":"Less is better","values":[0,2,3,1,2,0],"isMain":false},{"metricName":" # translocations","quality":"Less is better","values":[0,43,56,68,51,26],"isMain":false},{"metricName":" # inversions","quality":"Less is better","values":[3,36,5,6,19,0],"isMain":false},{"metricName":" # interspecies translocations","quality":"Less is better","values":[56,145,106,178,121,19],"isMain":false},{"metricName":"# possibly misassembled contigs","quality":"Less is better","values":[0,3,1,4,0,0],"isMain":false},{"metricName":" # possible misassemblies","quality":"Less is better","values":[0,4,1,4,0,0],"isMain":false},{"metricName":"# local misassemblies","quality":"Less is better","values":[1,29,5,8,17,1],"isMain":false},{"metricName":"# scaffold gap ext. mis.","quality":"Less is better","values":[0,0,0,0,0,0],"isMain":false},{"metricName":"# scaffold gap loc. mis.","quality":"Less is better","values":[0,0,0,0,0,0],"isMain":false}]],["Unaligned",[]],["Mismatches",[]],["Statistics without reference",[]],["K-mer-based statistics",[]],["Predicted genes",[]],["Similarity statistics",[]],["Reference statistics",[]]],[["Genome statistics",[]],["Reads mapping",[]],["Misassemblies",[{"metricName":"# misassemblies","quality":"Less is better","values":[1,0,1,5,2,3],"isMain":true},{"metricName":" # relocations","quality":"Less is better","values":[0,0,0,0,0,0],"isMain":false},{"metricName":" # translocations","quality":"Less is better","values":[0,0,0,0,0,0],"isMain":false},{"metricName":" # inversions","quality":"Less is better","values":[0,0,0,0,0,0],"isMain":false},{"metricName":" # interspecies translocations","quality":"Less is better","values":[1,0,1,5,2,3],"isMain":false},{"metricName":"# possibly misassembled contigs","quality":"Less is better","values":[0,0,0,0,0,0],"isMain":false},{"metricName":" # possible misassemblies","quality":"Less is better","values":[0,0,0,0,0,0],"isMain":false},{"metricName":"# local misassemblies","quality":"Less is better","values":[0,0,0,0,0,0],"isMain":false},{"metricName":"# scaffold gap ext. mis.","quality":"Less is better","values":[0,0,0,0,0,0],"isMain":false},{"metricName":"# scaffold gap loc. mis.","quality":"Less is better","values":[0,0,0,0,0,0],"isMain":false}]],["Unaligned",[]],["Mismatches",[]],["Statistics without reference",[]],["K-mer-based statistics",[]],["Predicted genes",[]],["Similarity statistics",[]],["Reference statistics",[]]],[["Genome statistics",[]],["Reads mapping",[]],["Misassemblies",[{"metricName":"# misassemblies","quality":"Less is better","values":[97,125,42,57,106,60],"isMain":true},{"metricName":" # relocations","quality":"Less is better","values":[0,0,0,0,0,0],"isMain":false},{"metricName":" # translocations","quality":"Less is better","values":[3,8,5,5,9,2],"isMain":false},{"metricName":" # inversions","quality":"Less is better","values":[1,1,0,2,1,0],"isMain":false},{"metricName":" # interspecies translocations","quality":"Less is better","values":[93,116,37,50,96,58],"isMain":false},{"metricName":"# possibly misassembled contigs","quality":"Less is better","values":[1,1,1,0,4,0],"isMain":false},{"metricName":" # possible misassemblies","quality":"Less is better","values":[1,1,1,0,5,0],"isMain":false},{"metricName":"# local misassemblies","quality":"Less is better","values":[0,2,3,5,1,1],"isMain":false},{"metricName":"# scaffold gap ext. mis.","quality":"Less is better","values":[0,0,0,0,0,0],"isMain":false},{"metricName":"# scaffold gap loc. mis.","quality":"Less is better","values":[0,0,0,0,0,0],"isMain":false}]],["Unaligned",[]],["Mismatches",[]],["Statistics without reference",[]],["K-mer-based statistics",[]],["Predicted genes",[]],["Similarity statistics",[]],["Reference statistics",[]]],[["Genome statistics",[]],["Reads mapping",[]],["Misassemblies",[{"metricName":"# misassemblies","quality":"Less is better","values":[11,20,3,17,19,14],"isMain":true},{"metricName":" # relocations","quality":"Less is better","values":[0,0,0,0,0,0],"isMain":false},{"metricName":" # translocations","quality":"Less is better","values":[0,1,1,1,0,0],"isMain":false},{"metricName":" # inversions","quality":"Less is better","values":[0,0,0,0,1,0],"isMain":false},{"metricName":" # interspecies translocations","quality":"Less is better","values":[11,19,2,16,18,14],"isMain":false},{"metricName":"# possibly misassembled contigs","quality":"Less is better","values":[0,0,0,0,0,0],"isMain":false},{"metricName":" # possible misassemblies","quality":"Less is better","values":[0,0,0,0,0,0],"isMain":false},{"metricName":"# local misassemblies","quality":"Less is better","values":[0,0,0,0,0,0],"isMain":false},{"metricName":"# scaffold gap ext. mis.","quality":"Less is better","values":[0,0,0,0,0,0],"isMain":false},{"metricName":"# scaffold gap loc. mis.","quality":"Less is better","values":[0,0,0,0,0,0],"isMain":false}]],["Unaligned",[]],["Mismatches",[]],["Statistics without reference",[]],["K-mer-based statistics",[]],["Predicted genes",[]],["Similarity statistics",[]],["Reference statistics",[]]],[["Genome statistics",[]],["Reads mapping",[]],["Misassemblies",[{"metricName":"# misassemblies","quality":"Less is better","values":[47,37,21,59,46,38],"isMain":true},{"metricName":" # relocations","quality":"Less is better","values":[2,3,2,2,4,2],"isMain":false},{"metricName":" # translocations","quality":"Less is better","values":[9,12,13,10,20,6],"isMain":false},{"metricName":" # inversions","quality":"Less is better","values":[1,1,5,3,1,0],"isMain":false},{"metricName":" # interspecies translocations","quality":"Less is better","values":[35,21,1,44,21,30],"isMain":false},{"metricName":"# possibly misassembled contigs","quality":"Less is better","values":[0,0,0,0,0,0],"isMain":false},{"metricName":" # possible misassemblies","quality":"Less is better","values":[0,0,0,0,0,0],"isMain":false},{"metricName":"# local misassemblies","quality":"Less is better","values":[6,4,5,8,9,2],"isMain":false},{"metricName":"# scaffold gap ext. mis.","quality":"Less is better","values":[0,0,0,0,0,0],"isMain":false},{"metricName":"# scaffold gap loc. mis.","quality":"Less is better","values":[0,0,0,0,0,0],"isMain":false}]],["Unaligned",[]],["Mismatches",[]],["Statistics without reference",[]],["K-mer-based statistics",[]],["Predicted genes",[]],["Similarity statistics",[]],["Reference statistics",[]]],[["Genome statistics",[]],["Reads mapping",[]],["Misassemblies",[{"metricName":"# misassemblies","quality":"Less is better","values":[21,249,290,392,313,9],"isMain":true},{"metricName":" # relocations","quality":"Less is better","values":[0,0,0,1,4,0],"isMain":false},{"metricName":" # translocations","quality":"Less is better","values":[1,41,60,77,58,3],"isMain":false},{"metricName":" # inversions","quality":"Less is better","values":[0,49,14,14,33,0],"isMain":false},{"metricName":" # interspecies translocations","quality":"Less is better","values":[20,159,216,300,218,6],"isMain":false},{"metricName":"# possibly misassembled contigs","quality":"Less is better","values":[0,1,3,2,0,0],"isMain":false},{"metricName":" # possible misassemblies","quality":"Less is better","values":[0,1,3,2,0,0],"isMain":false},{"metricName":"# local misassemblies","quality":"Less is better","values":[2,17,17,13,24,1],"isMain":false},{"metricName":"# scaffold gap ext. mis.","quality":"Less is better","values":[0,0,0,0,0,0],"isMain":false},{"metricName":"# scaffold gap loc. mis.","quality":"Less is better","values":[0,0,0,0,0,0],"isMain":false}]],["Unaligned",[]],["Mismatches",[]],["Statistics without reference",[]],["K-mer-based statistics",[]],["Predicted genes",[]],["Similarity statistics",[]],["Reference statistics",[]]],[["Genome statistics",[]],["Reads mapping",[]],["Misassemblies",[{"metricName":"# misassemblies","quality":"Less is better","values":[7,28,13,46,26,3],"isMain":true},{"metricName":" # relocations","quality":"Less is better","values":[0,1,2,3,5,0],"isMain":false},{"metricName":" # translocations","quality":"Less is better","values":[0,0,0,0,0,0],"isMain":false},{"metricName":" # inversions","quality":"Less is better","values":[0,0,1,1,0,0],"isMain":false},{"metricName":" # interspecies translocations","quality":"Less is better","values":[7,27,10,42,21,3],"isMain":false},{"metricName":"# possibly misassembled contigs","quality":"Less is better","values":[0,0,0,0,2,0],"isMain":false},{"metricName":" # possible misassemblies","quality":"Less is better","values":[0,0,0,0,2,0],"isMain":false},{"metricName":"# local misassemblies","quality":"Less is better","values":[2,2,3,5,4,2],"isMain":false},{"metricName":"# scaffold gap ext. mis.","quality":"Less is better","values":[0,0,0,0,0,0],"isMain":false},{"metricName":"# scaffold gap loc. mis.","quality":"Less is better","values":[0,0,0,0,0,0],"isMain":false}]],["Unaligned",[]],["Mismatches",[]],["Statistics without reference",[]],["K-mer-based statistics",[]],["Predicted genes",[]],["Similarity statistics",[]],["Reference statistics",[]]],[["Genome statistics",[]],["Reads mapping",[]],["Misassemblies",[{"metricName":"# misassemblies","quality":"Less is better","values":[3,5,2,7,4,2],"isMain":true},{"metricName":" # relocations","quality":"Less is better","values":[0,0,0,0,0,0],"isMain":false},{"metricName":" # translocations","quality":"Less is better","values":[0,0,0,0,0,0],"isMain":false},{"metricName":" # inversions","quality":"Less is better","values":[0,0,0,0,0,0],"isMain":false},{"metricName":" # interspecies translocations","quality":"Less is better","values":[3,5,2,7,4,2],"isMain":false},{"metricName":"# possibly misassembled contigs","quality":"Less is better","values":[0,0,0,0,0,0],"isMain":false},{"metricName":" # possible misassemblies","quality":"Less is better","values":[0,0,0,0,0,0],"isMain":false},{"metricName":"# local misassemblies","quality":"Less is better","values":[1,0,0,1,0,0],"isMain":false},{"metricName":"# scaffold gap ext. mis.","quality":"Less is better","values":[0,0,0,0,0,0],"isMain":false},{"metricName":"# scaffold gap loc. mis.","quality":"Less is better","values":[0,0,0,0,0,0],"isMain":false}]],["Unaligned",[]],["Mismatches",[]],["Statistics without reference",[]],["K-mer-based statistics",[]],["Predicted genes",[]],["Similarity statistics",[]],["Reference statistics",[]]],[["Genome statistics",[]],["Reads mapping",[]],["Misassemblies",[{"metricName":"# misassemblies","quality":"Less is better","values":[34,41,10,28,32,9],"isMain":true},{"metricName":" # relocations","quality":"Less is better","values":[2,4,0,0,4,0],"isMain":false},{"metricName":" # translocations","quality":"Less is better","values":[0,0,0,0,0,0],"isMain":false},{"metricName":" # inversions","quality":"Less is better","values":[0,0,0,0,0,0],"isMain":false},{"metricName":" # interspecies translocations","quality":"Less is better","values":[32,37,10,28,28,9],"isMain":false},{"metricName":"# possibly misassembled contigs","quality":"Less is better","values":[0,0,0,1,1,0],"isMain":false},{"metricName":" # possible misassemblies","quality":"Less is better","values":[0,0,0,1,1,0],"isMain":false},{"metricName":"# local misassemblies","quality":"Less is better","values":[0,2,0,0,1,0],"isMain":false},{"metricName":"# scaffold gap ext. mis.","quality":"Less is better","values":[0,0,0,0,0,0],"isMain":false},{"metricName":"# scaffold gap loc. mis.","quality":"Less is better","values":[0,0,0,0,0,0],"isMain":false}]],["Unaligned",[]],["Mismatches",[]],["Statistics without reference",[]],["K-mer-based statistics",[]],["Predicted genes",[]],["Similarity statistics",[]],["Reference statistics",[]]],[["Genome statistics",[]],["Reads mapping",[]],["Misassemblies",[{"metricName":"# misassemblies","quality":"Less is better","values":[1,1,4,6,3,3],"isMain":true},{"metricName":" # relocations","quality":"Less is better","values":[0,0,1,2,0,1],"isMain":false},{"metricName":" # translocations","quality":"Less is better","values":[1,1,3,3,3,1],"isMain":false},{"metricName":" # inversions","quality":"Less is better","values":[0,0,0,0,0,0],"isMain":false},{"metricName":" # interspecies translocations","quality":"Less is better","values":[0,0,0,1,0,1],"isMain":false},{"metricName":"# possibly misassembled contigs","quality":"Less is better","values":[0,0,0,0,0,0],"isMain":false},{"metricName":" # possible misassemblies","quality":"Less is better","values":[0,0,0,0,0,0],"isMain":false},{"metricName":"# local misassemblies","quality":"Less is better","values":[0,0,1,0,4,0],"isMain":false},{"metricName":"# scaffold gap ext. mis.","quality":"Less is better","values":[0,0,0,0,0,0],"isMain":false},{"metricName":"# scaffold gap loc. mis.","quality":"Less is better","values":[0,0,0,0,0,0],"isMain":false}]],["Unaligned",[]],["Mismatches",[]],["Statistics without reference",[]],["K-mer-based statistics",[]],["Predicted genes",[]],["Similarity statistics",[]],["Reference statistics",[]]],[["Genome statistics",[]],["Reads mapping",[]],["Misassemblies",[{"metricName":"# misassemblies","quality":"Less is better","values":[9,6,5,19,12,1],"isMain":true},{"metricName":" # relocations","quality":"Less is better","values":[1,0,0,0,2,0],"isMain":false},{"metricName":" # translocations","quality":"Less is better","values":[1,0,0,0,0,0],"isMain":false},{"metricName":" # inversions","quality":"Less is better","values":[0,1,1,0,1,0],"isMain":false},{"metricName":" # interspecies translocations","quality":"Less is better","values":[7,5,4,19,9,1],"isMain":false},{"metricName":"# possibly misassembled contigs","quality":"Less is better","values":[0,0,0,0,0,0],"isMain":false},{"metricName":" # possible misassemblies","quality":"Less is better","values":[0,0,0,0,0,0],"isMain":false},{"metricName":"# local misassemblies","quality":"Less is better","values":[2,1,4,3,5,0],"isMain":false},{"metricName":"# scaffold gap ext. mis.","quality":"Less is better","values":[0,0,0,0,0,0],"isMain":false},{"metricName":"# scaffold gap loc. mis.","quality":"Less is better","values":[0,0,0,0,0,0],"isMain":false}]],["Unaligned",[]],["Mismatches",[]],["Statistics without reference",[]],["K-mer-based statistics",[]],["Predicted genes",[]],["Similarity statistics",[]],["Reference statistics",[]]],[["Genome statistics",[]],["Reads mapping",[]],["Misassemblies",[{"metricName":"# misassemblies","quality":"Less is better","values":[4,6,4,16,5,3],"isMain":true},{"metricName":" # relocations","quality":"Less is better","values":[0,1,1,0,0,0],"isMain":false},{"metricName":" # translocations","quality":"Less is better","values":[0,1,0,0,0,1],"isMain":false},{"metricName":" # inversions","quality":"Less is better","values":[0,0,0,0,0,0],"isMain":false},{"metricName":" # interspecies translocations","quality":"Less is better","values":[4,4,3,16,5,2],"isMain":false},{"metricName":"# possibly misassembled contigs","quality":"Less is better","values":[0,0,0,0,0,0],"isMain":false},{"metricName":" # possible misassemblies","quality":"Less is better","values":[0,0,0,0,0,0],"isMain":false},{"metricName":"# local misassemblies","quality":"Less is better","values":[0,0,0,0,0,0],"isMain":false},{"metricName":"# scaffold gap ext. mis.","quality":"Less is better","values":[0,0,0,0,0,0],"isMain":false},{"metricName":"# scaffold gap loc. mis.","quality":"Less is better","values":[0,0,0,0,0,0],"isMain":false}]],["Unaligned",[]],["Mismatches",[]],["Statistics without reference",[]],["K-mer-based statistics",[]],["Predicted genes",[]],["Similarity statistics",[]],["Reference statistics",[]]],[["Genome statistics",[]],["Reads mapping",[]],["Misassemblies",[{"metricName":"# misassemblies","quality":"Less is better","values":[5,9,2,8,12,5],"isMain":true},{"metricName":" # relocations","quality":"Less is better","values":[0,0,0,0,1,0],"isMain":false},{"metricName":" # translocations","quality":"Less is better","values":[0,0,0,0,0,0],"isMain":false},{"metricName":" # inversions","quality":"Less is better","values":[0,0,0,0,0,0],"isMain":false},{"metricName":" # interspecies translocations","quality":"Less is better","values":[5,9,2,8,11,5],"isMain":false},{"metricName":"# possibly misassembled contigs","quality":"Less is better","values":[0,0,0,0,0,0],"isMain":false},{"metricName":" # possible misassemblies","quality":"Less is better","values":[0,0,0,0,0,0],"isMain":false},{"metricName":"# local misassemblies","quality":"Less is better","values":[0,0,0,0,0,0],"isMain":false},{"metricName":"# scaffold gap ext. mis.","quality":"Less is better","values":[0,0,0,0,0,0],"isMain":false},{"metricName":"# scaffold gap loc. mis.","quality":"Less is better","values":[0,0,0,0,0,0],"isMain":false}]],["Unaligned",[]],["Mismatches",[]],["Statistics without reference",[]],["K-mer-based statistics",[]],["Predicted genes",[]],["Similarity statistics",[]],["Reference statistics",[]]],[["Genome statistics",[]],["Reads mapping",[]],["Misassemblies",[{"metricName":"# misassemblies","quality":"Less is better","values":[11,47,33,69,51,3],"isMain":true},{"metricName":" # relocations","quality":"Less is better","values":[1,0,1,0,0,0],"isMain":false},{"metricName":" # translocations","quality":"Less is better","values":[3,7,5,7,9,2],"isMain":false},{"metricName":" # inversions","quality":"Less is better","values":[0,0,0,0,0,0],"isMain":false},{"metricName":" # interspecies translocations","quality":"Less is better","values":[7,40,27,62,42,1],"isMain":false},{"metricName":"# possibly misassembled contigs","quality":"Less is better","values":[0,0,0,0,0,0],"isMain":false},{"metricName":" # possible misassemblies","quality":"Less is better","values":[0,0,0,0,0,0],"isMain":false},{"metricName":"# local misassemblies","quality":"Less is better","values":[1,4,0,5,4,0],"isMain":false},{"metricName":"# scaffold gap ext. mis.","quality":"Less is better","values":[0,0,0,0,0,0],"isMain":false},{"metricName":"# scaffold gap loc. mis.","quality":"Less is better","values":[0,0,0,0,0,0],"isMain":false}]],["Unaligned",[]],["Mismatches",[]],["Statistics without reference",[]],["K-mer-based statistics",[]],["Predicted genes",[]],["Similarity statistics",[]],["Reference statistics",[]]],[["Genome statistics",[]],["Reads mapping",[]],["Misassemblies",[{"metricName":"# misassemblies","quality":"Less is better","values":[12,12,10,20,14,8],"isMain":true},{"metricName":" # relocations","quality":"Less is better","values":[0,0,1,0,0,0],"isMain":false},{"metricName":" # translocations","quality":"Less is better","values":[0,0,0,0,0,0],"isMain":false},{"metricName":" # inversions","quality":"Less is better","values":[0,0,0,0,0,0],"isMain":false},{"metricName":" # interspecies translocations","quality":"Less is better","values":[12,12,9,20,14,8],"isMain":false},{"metricName":"# possibly misassembled contigs","quality":"Less is better","values":[0,0,0,0,0,0],"isMain":false},{"metricName":" # possible misassemblies","quality":"Less is better","values":[0,0,0,0,0,0],"isMain":false},{"metricName":"# local misassemblies","quality":"Less is better","values":[0,0,1,0,0,0],"isMain":false},{"metricName":"# scaffold gap ext. mis.","quality":"Less is better","values":[0,0,0,0,0,0],"isMain":false},{"metricName":"# scaffold gap loc. mis.","quality":"Less is better","values":[0,0,0,0,0,0],"isMain":false}]],["Unaligned",[]],["Mismatches",[]],["Statistics without reference",[]],["K-mer-based statistics",[]],["Predicted genes",[]],["Similarity statistics",[]],["Reference statistics",[]]],[["Genome statistics",[]],["Reads mapping",[]],["Misassemblies",[{"metricName":"# misassemblies","quality":"Less is better","values":[24,5,7,17,12,5],"isMain":true},{"metricName":" # relocations","quality":"Less is better","values":[0,0,0,0,0,0],"isMain":false},{"metricName":" # translocations","quality":"Less is better","values":[2,2,4,5,5,3],"isMain":false},{"metricName":" # inversions","quality":"Less is better","values":[1,0,0,0,1,0],"isMain":false},{"metricName":" # interspecies translocations","quality":"Less is better","values":[21,3,3,12,6,2],"isMain":false},{"metricName":"# possibly misassembled contigs","quality":"Less is better","values":[0,0,0,0,0,0],"isMain":false},{"metricName":" # possible misassemblies","quality":"Less is better","values":[0,0,0,0,0,0],"isMain":false},{"metricName":"# local misassemblies","quality":"Less is better","values":[0,0,1,1,1,0],"isMain":false},{"metricName":"# scaffold gap ext. mis.","quality":"Less is better","values":[0,0,0,0,0,0],"isMain":false},{"metricName":"# scaffold gap loc. mis.","quality":"Less is better","values":[0,0,0,0,0,0],"isMain":false}]],["Unaligned",[]],["Mismatches",[]],["Statistics without reference",[]],["K-mer-based statistics",[]],["Predicted genes",[]],["Similarity statistics",[]],["Reference statistics",[]]],[["Genome statistics",[]],["Reads mapping",[]],["Misassemblies",[{"metricName":"# misassemblies","quality":"Less is better","values":[68,72,17,54,50,30],"isMain":true},{"metricName":" # relocations","quality":"Less is better","values":[1,2,1,3,2,0],"isMain":false},{"metricName":" # translocations","quality":"Less is better","values":[15,16,6,8,13,6],"isMain":false},{"metricName":" # inversions","quality":"Less is better","values":[0,0,0,0,0,0],"isMain":false},{"metricName":" # interspecies translocations","quality":"Less is better","values":[52,54,10,43,35,24],"isMain":false},{"metricName":"# possibly misassembled contigs","quality":"Less is better","values":[0,0,0,0,0,0],"isMain":false},{"metricName":" # possible misassemblies","quality":"Less is better","values":[0,0,0,0,0,0],"isMain":false},{"metricName":"# local misassemblies","quality":"Less is better","values":[4,7,4,6,4,1],"isMain":false},{"metricName":"# scaffold gap ext. mis.","quality":"Less is better","values":[0,0,0,0,0,0],"isMain":false},{"metricName":"# scaffold gap loc. mis.","quality":"Less is better","values":[0,0,0,0,0,0],"isMain":false}]],["Unaligned",[]],["Mismatches",[]],["Statistics without reference",[]],["K-mer-based statistics",[]],["Predicted genes",[]],["Similarity statistics",[]],["Reference statistics",[]]],[["Genome statistics",[]],["Reads mapping",[]],["Misassemblies",[{"metricName":"# misassemblies","quality":"Less is better","values":[11,21,12,37,20,6],"isMain":true},{"metricName":" # relocations","quality":"Less is better","values":[1,0,0,2,0,1],"isMain":false},{"metricName":" # translocations","quality":"Less is better","values":[5,9,9,13,11,4],"isMain":false},{"metricName":" # inversions","quality":"Less is better","values":[0,0,0,0,1,0],"isMain":false},{"metricName":" # interspecies translocations","quality":"Less is better","values":[5,12,3,22,8,1],"isMain":false},{"metricName":"# possibly misassembled contigs","quality":"Less is better","values":[0,0,0,0,0,0],"isMain":false},{"metricName":" # possible misassemblies","quality":"Less is better","values":[0,0,0,0,0,0],"isMain":false},{"metricName":"# local misassemblies","quality":"Less is better","values":[3,1,4,1,1,1],"isMain":false},{"metricName":"# scaffold gap ext. mis.","quality":"Less is better","values":[0,0,0,0,0,0],"isMain":false},{"metricName":"# scaffold gap loc. mis.","quality":"Less is better","values":[0,0,0,0,0,0],"isMain":false}]],["Unaligned",[]],["Mismatches",[]],["Statistics without reference",[]],["K-mer-based statistics",[]],["Predicted genes",[]],["Similarity statistics",[]],["Reference statistics",[]]],[["Genome statistics",[]],["Reads mapping",[]],["Misassemblies",[{"metricName":"# misassemblies","quality":"Less is better","values":[31,38,26,57,37,10],"isMain":true},{"metricName":" # relocations","quality":"Less is better","values":[1,1,2,2,1,0],"isMain":false},{"metricName":" # translocations","quality":"Less is better","values":[10,11,13,11,10,5],"isMain":false},{"metricName":" # inversions","quality":"Less is better","values":[1,0,2,0,1,0],"isMain":false},{"metricName":" # interspecies translocations","quality":"Less is better","values":[19,26,9,44,25,5],"isMain":false},{"metricName":"# possibly misassembled contigs","quality":"Less is better","values":[0,0,0,0,0,0],"isMain":false},{"metricName":" # possible misassemblies","quality":"Less is better","values":[0,0,0,0,0,0],"isMain":false},{"metricName":"# local misassemblies","quality":"Less is better","values":[2,5,6,3,3,0],"isMain":false},{"metricName":"# scaffold gap ext. mis.","quality":"Less is better","values":[0,0,0,0,0,0],"isMain":false},{"metricName":"# scaffold gap loc. mis.","quality":"Less is better","values":[0,0,0,0,0,0],"isMain":false}]],["Unaligned",[]],["Mismatches",[]],["Statistics without reference",[]],["K-mer-based statistics",[]],["Predicted genes",[]],["Similarity statistics",[]],["Reference statistics",[]]],[["Genome statistics",[]],["Reads mapping",[]],["Misassemblies",[{"metricName":"# misassemblies","quality":"Less is better","values":[88,125,57,86,137,50],"isMain":true},{"metricName":" # relocations","quality":"Less is better","values":[2,2,2,2,3,2],"isMain":false},{"metricName":" # translocations","quality":"Less is better","values":[5,5,7,3,10,1],"isMain":false},{"metricName":" # inversions","quality":"Less is better","values":[1,2,2,1,3,0],"isMain":false},{"metricName":" # interspecies translocations","quality":"Less is better","values":[80,116,46,80,121,47],"isMain":false},{"metricName":"# possibly misassembled contigs","quality":"Less is better","values":[0,0,0,1,1,0],"isMain":false},{"metricName":" # possible misassemblies","quality":"Less is better","values":[0,0,0,1,1,0],"isMain":false},{"metricName":"# local misassemblies","quality":"Less is better","values":[4,3,4,4,3,2],"isMain":false},{"metricName":"# scaffold gap ext. mis.","quality":"Less is better","values":[0,0,0,0,0,0],"isMain":false},{"metricName":"# scaffold gap loc. mis.","quality":"Less is better","values":[0,0,0,0,0,0],"isMain":false}]],["Unaligned",[]],["Mismatches",[]],["Statistics without reference",[]],["K-mer-based statistics",[]],["Predicted genes",[]],["Similarity statistics",[]],["Reference statistics",[]]],[["Genome statistics",[]],["Reads mapping",[]],["Misassemblies",[{"metricName":"# misassemblies","quality":"Less is better","values":[5,11,3,15,12,8],"isMain":true},{"metricName":" # relocations","quality":"Less is better","values":[0,0,0,1,0,0],"isMain":false},{"metricName":" # translocations","quality":"Less is better","values":[0,0,0,2,0,1],"isMain":false},{"metricName":" # inversions","quality":"Less is better","values":[0,0,0,0,0,0],"isMain":false},{"metricName":" # interspecies translocations","quality":"Less is better","values":[5,11,3,12,12,7],"isMain":false},{"metricName":"# possibly misassembled contigs","quality":"Less is better","values":[0,1,0,0,0,0],"isMain":false},{"metricName":" # possible misassemblies","quality":"Less is better","values":[0,1,0,0,0,0],"isMain":false},{"metricName":"# local misassemblies","quality":"Less is better","values":[0,0,0,0,0,0],"isMain":false},{"metricName":"# scaffold gap ext. mis.","quality":"Less is better","values":[0,0,0,0,0,0],"isMain":false},{"metricName":"# scaffold gap loc. mis.","quality":"Less is better","values":[0,0,0,0,0,0],"isMain":false}]],["Unaligned",[]],["Mismatches",[]],["Statistics without reference",[]],["K-mer-based statistics",[]],["Predicted genes",[]],["Similarity statistics",[]],["Reference statistics",[]]],[["Genome statistics",[]],["Reads mapping",[]],["Misassemblies",[{"metricName":"# misassemblies","quality":"Less is better","values":[2,4,0,3,7,3],"isMain":true},{"metricName":" # relocations","quality":"Less is better","values":[1,1,0,0,1,0],"isMain":false},{"metricName":" # translocations","quality":"Less is better","values":[0,1,0,0,0,0],"isMain":false},{"metricName":" # inversions","quality":"Less is better","values":[0,0,0,0,0,0],"isMain":false},{"metricName":" # interspecies translocations","quality":"Less is better","values":[1,2,0,3,6,3],"isMain":false},{"metricName":"# possibly misassembled contigs","quality":"Less is better","values":[0,0,0,0,0,0],"isMain":false},{"metricName":" # possible misassemblies","quality":"Less is better","values":[0,0,0,0,0,0],"isMain":false},{"metricName":"# local misassemblies","quality":"Less is better","values":[1,4,0,1,2,0],"isMain":false},{"metricName":"# scaffold gap ext. mis.","quality":"Less is better","values":[0,0,0,0,0,0],"isMain":false},{"metricName":"# scaffold gap loc. mis.","quality":"Less is better","values":[0,0,0,0,0,0],"isMain":false}]],["Unaligned",[]],["Mismatches",[]],["Statistics without reference",[]],["K-mer-based statistics",[]],["Predicted genes",[]],["Similarity statistics",[]],["Reference statistics",[]]],[["Genome statistics",[]],["Reads mapping",[]],["Misassemblies",[{"metricName":"# misassemblies","quality":"Less is better","values":[70,88,29,55,71,47],"isMain":true},{"metricName":" # relocations","quality":"Less is better","values":[3,4,2,1,3,1],"isMain":false},{"metricName":" # translocations","quality":"Less is better","values":[9,10,11,6,10,6],"isMain":false},{"metricName":" # inversions","quality":"Less is better","values":[0,0,2,1,0,0],"isMain":false},{"metricName":" # interspecies translocations","quality":"Less is better","values":[58,74,14,47,58,40],"isMain":false},{"metricName":"# possibly misassembled contigs","quality":"Less is better","values":[1,0,0,0,0,1],"isMain":false},{"metricName":" # possible misassemblies","quality":"Less is better","values":[1,0,0,0,0,1],"isMain":false},{"metricName":"# local misassemblies","quality":"Less is better","values":[3,2,6,5,4,0],"isMain":false},{"metricName":"# scaffold gap ext. mis.","quality":"Less is better","values":[0,0,0,0,0,0],"isMain":false},{"metricName":"# scaffold gap loc. mis.","quality":"Less is better","values":[0,0,0,0,0,0],"isMain":false}]],["Unaligned",[]],["Mismatches",[]],["Statistics without reference",[]],["K-mer-based statistics",[]],["Predicted genes",[]],["Similarity statistics",[]],["Reference statistics",[]]],[["Genome statistics",[]],["Reads mapping",[]],["Misassemblies",[{"metricName":"# misassemblies","quality":"Less is better","values":[22,20,11,30,29,6],"isMain":true},{"metricName":" # relocations","quality":"Less is better","values":[0,1,0,0,1,0],"isMain":false},{"metricName":" # translocations","quality":"Less is better","values":[4,7,5,9,9,3],"isMain":false},{"metricName":" # inversions","quality":"Less is better","values":[0,0,0,0,0,0],"isMain":false},{"metricName":" # interspecies translocations","quality":"Less is better","values":[18,12,6,21,19,3],"isMain":false},{"metricName":"# possibly misassembled contigs","quality":"Less is better","values":[0,0,0,0,0,0],"isMain":false},{"metricName":" # possible misassemblies","quality":"Less is better","values":[0,0,0,0,0,0],"isMain":false},{"metricName":"# local misassemblies","quality":"Less is better","values":[1,0,0,1,0,0],"isMain":false},{"metricName":"# scaffold gap ext. mis.","quality":"Less is better","values":[0,0,0,0,0,0],"isMain":false},{"metricName":"# scaffold gap loc. mis.","quality":"Less is better","values":[0,0,0,0,0,0],"isMain":false}]],["Unaligned",[]],["Mismatches",[]],["Statistics without reference",[]],["K-mer-based statistics",[]],["Predicted genes",[]],["Similarity statistics",[]],["Reference statistics",[]]],[["Genome statistics",[]],["Reads mapping",[]],["Misassemblies",[{"metricName":"# misassemblies","quality":"Less is better","values":[78,109,16,72,95,52],"isMain":true},{"metricName":" # relocations","quality":"Less is better","values":[1,2,1,0,1,0],"isMain":false},{"metricName":" # translocations","quality":"Less is better","values":[9,15,4,17,16,6],"isMain":false},{"metricName":" # inversions","quality":"Less is better","values":[0,1,1,1,1,0],"isMain":false},{"metricName":" # interspecies translocations","quality":"Less is better","values":[68,91,10,54,77,46],"isMain":false},{"metricName":"# possibly misassembled contigs","quality":"Less is better","values":[0,0,1,0,0,0],"isMain":false},{"metricName":" # possible misassemblies","quality":"Less is better","values":[0,0,1,0,0,0],"isMain":false},{"metricName":"# local misassemblies","quality":"Less is better","values":[2,3,3,1,2,0],"isMain":false},{"metricName":"# scaffold gap ext. mis.","quality":"Less is better","values":[0,0,0,0,0,0],"isMain":false},{"metricName":"# scaffold gap loc. mis.","quality":"Less is better","values":[0,0,0,0,0,0],"isMain":false}]],["Unaligned",[]],["Mismatches",[]],["Statistics without reference",[]],["K-mer-based statistics",[]],["Predicted genes",[]],["Similarity statistics",[]],["Reference statistics",[]]],[["Genome statistics",[]],["Reads mapping",[]],["Misassemblies",[{"metricName":"# misassemblies","quality":"Less is better","values":[5,7,10,11,12,1],"isMain":true},{"metricName":" # relocations","quality":"Less is better","values":[0,0,0,0,0,0],"isMain":false},{"metricName":" # translocations","quality":"Less is better","values":[1,0,1,1,1,0],"isMain":false},{"metricName":" # inversions","quality":"Less is better","values":[0,0,0,0,0,0],"isMain":false},{"metricName":" # interspecies translocations","quality":"Less is better","values":[4,7,9,10,11,1],"isMain":false},{"metricName":"# possibly misassembled contigs","quality":"Less is better","values":[0,0,0,0,0,0],"isMain":false},{"metricName":" # possible misassemblies","quality":"Less is better","values":[0,0,0,0,0,0],"isMain":false},{"metricName":"# local misassemblies","quality":"Less is better","values":[0,1,0,0,2,0],"isMain":false},{"metricName":"# scaffold gap ext. mis.","quality":"Less is better","values":[0,0,0,0,0,0],"isMain":false},{"metricName":"# scaffold gap loc. mis.","quality":"Less is better","values":[0,0,0,0,0,0],"isMain":false}]],["Unaligned",[]],["Mismatches",[]],["Statistics without reference",[]],["K-mer-based statistics",[]],["Predicted genes",[]],["Similarity statistics",[]],["Reference statistics",[]]],[["Genome statistics",[]],["Reads mapping",[]],["Misassemblies",[{"metricName":"# misassemblies","quality":"Less is better","values":[2,5,5,5,6,2],"isMain":true},{"metricName":" # relocations","quality":"Less is better","values":[0,0,0,0,0,0],"isMain":false},{"metricName":" # translocations","quality":"Less is better","values":[0,0,0,0,0,0],"isMain":false},{"metricName":" # inversions","quality":"Less is better","values":[0,0,0,1,0,0],"isMain":false},{"metricName":" # interspecies translocations","quality":"Less is better","values":[2,5,5,4,6,2],"isMain":false},{"metricName":"# possibly misassembled contigs","quality":"Less is better","values":[0,0,0,0,0,0],"isMain":false},{"metricName":" # possible misassemblies","quality":"Less is better","values":[0,0,0,0,0,0],"isMain":false},{"metricName":"# local misassemblies","quality":"Less is better","values":[0,0,0,0,0,0],"isMain":false},{"metricName":"# scaffold gap ext. mis.","quality":"Less is better","values":[0,0,0,0,0,0],"isMain":false},{"metricName":"# scaffold gap loc. mis.","quality":"Less is better","values":[0,0,0,0,0,0],"isMain":false}]],["Unaligned",[]],["Mismatches",[]],["Statistics without reference",[]],["K-mer-based statistics",[]],["Predicted genes",[]],["Similarity statistics",[]],["Reference statistics",[]]],[["Genome statistics",[]],["Reads mapping",[]],["Misassemblies",[{"metricName":"# misassemblies","quality":"Less is better","values":[67,74,21,52,72,34],"isMain":true},{"metricName":" # relocations","quality":"Less is better","values":[1,0,1,2,3,0],"isMain":false},{"metricName":" # translocations","quality":"Less is better","values":[4,2,8,6,6,1],"isMain":false},{"metricName":" # inversions","quality":"Less is better","values":[2,1,0,1,0,0],"isMain":false},{"metricName":" # interspecies translocations","quality":"Less is better","values":[60,71,12,43,63,33],"isMain":false},{"metricName":"# possibly misassembled contigs","quality":"Less is better","values":[0,0,0,0,0,0],"isMain":false},{"metricName":" # possible misassemblies","quality":"Less is better","values":[0,0,0,0,0,0],"isMain":false},{"metricName":"# local misassemblies","quality":"Less is better","values":[2,2,1,0,2,0],"isMain":false},{"metricName":"# scaffold gap ext. mis.","quality":"Less is better","values":[0,0,0,0,0,0],"isMain":false},{"metricName":"# scaffold gap loc. mis.","quality":"Less is better","values":[0,0,0,0,0,0],"isMain":false}]],["Unaligned",[]],["Mismatches",[]],["Statistics without reference",[]],["K-mer-based statistics",[]],["Predicted genes",[]],["Similarity statistics",[]],["Reference statistics",[]]],[["Genome statistics",[]],["Reads mapping",[]],["Misassemblies",[{"metricName":"# misassemblies","quality":"Less is better","values":[16,32,7,22,21,17],"isMain":true},{"metricName":" # relocations","quality":"Less is better","values":[0,1,1,1,1,0],"isMain":false},{"metricName":" # translocations","quality":"Less is better","values":[1,3,0,1,3,2],"isMain":false},{"metricName":" # inversions","quality":"Less is better","values":[0,0,0,0,0,0],"isMain":false},{"metricName":" # interspecies translocations","quality":"Less is better","values":[15,28,6,20,17,15],"isMain":false},{"metricName":"# possibly misassembled contigs","quality":"Less is better","values":[0,1,0,0,0,0],"isMain":false},{"metricName":" # possible misassemblies","quality":"Less is better","values":[0,1,0,0,0,0],"isMain":false},{"metricName":"# local misassemblies","quality":"Less is better","values":[1,0,1,1,0,0],"isMain":false},{"metricName":"# scaffold gap ext. mis.","quality":"Less is better","values":[0,0,0,0,0,0],"isMain":false},{"metricName":"# scaffold gap loc. mis.","quality":"Less is better","values":[0,0,0,0,0,0],"isMain":false}]],["Unaligned",[]],["Mismatches",[]],["Statistics without reference",[]],["K-mer-based statistics",[]],["Predicted genes",[]],["Similarity statistics",[]],["Reference statistics",[]]],[["Genome statistics",[]],["Reads mapping",[]],["Misassemblies",[{"metricName":"# misassemblies","quality":"Less is better","values":[32,33,34,37,40,29],"isMain":true},{"metricName":" # relocations","quality":"Less is better","values":[0,0,1,0,2,0],"isMain":false},{"metricName":" # translocations","quality":"Less is better","values":[16,28,22,29,31,27],"isMain":false},{"metricName":" # inversions","quality":"Less is better","values":[0,0,0,1,0,0],"isMain":false},{"metricName":" # interspecies translocations","quality":"Less is better","values":[16,5,11,7,7,2],"isMain":false},{"metricName":"# possibly misassembled contigs","quality":"Less is better","values":[0,0,0,0,0,0],"isMain":false},{"metricName":" # possible misassemblies","quality":"Less is better","values":[0,0,0,0,0,0],"isMain":false},{"metricName":"# local misassemblies","quality":"Less is better","values":[1,4,5,3,3,1],"isMain":false},{"metricName":"# scaffold gap ext. mis.","quality":"Less is better","values":[0,0,0,0,0,0],"isMain":false},{"metricName":"# scaffold gap loc. mis.","quality":"Less is better","values":[0,0,0,0,0,0],"isMain":false}]],["Unaligned",[]],["Mismatches",[]],["Statistics without reference",[]],["K-mer-based statistics",[]],["Predicted genes",[]],["Similarity statistics",[]],["Reference statistics",[]]],[["Genome statistics",[]],["Reads mapping",[]],["Misassemblies",[{"metricName":"# misassemblies","quality":"Less is better","values":[7,9,8,11,11,3],"isMain":true},{"metricName":" # relocations","quality":"Less is better","values":[0,0,0,0,1,0],"isMain":false},{"metricName":" # translocations","quality":"Less is better","values":[0,0,0,0,0,0],"isMain":false},{"metricName":" # inversions","quality":"Less is better","values":[1,0,0,0,0,0],"isMain":false},{"metricName":" # interspecies translocations","quality":"Less is better","values":[6,9,8,11,10,3],"isMain":false},{"metricName":"# possibly misassembled contigs","quality":"Less is better","values":[0,0,1,0,0,0],"isMain":false},{"metricName":" # possible misassemblies","quality":"Less is better","values":[0,0,1,0,0,0],"isMain":false},{"metricName":"# local misassemblies","quality":"Less is better","values":[0,0,0,0,1,0],"isMain":false},{"metricName":"# scaffold gap ext. mis.","quality":"Less is better","values":[0,0,0,0,0,0],"isMain":false},{"metricName":"# scaffold gap loc. mis.","quality":"Less is better","values":[0,0,0,0,0,0],"isMain":false}]],["Unaligned",[]],["Mismatches",[]],["Statistics without reference",[]],["K-mer-based statistics",[]],["Predicted genes",[]],["Similarity statistics",[]],["Reference statistics",[]]],[["Genome statistics",[]],["Reads mapping",[]],["Misassemblies",[{"metricName":"# misassemblies","quality":"Less is better","values":[5,1,0,8,5,2],"isMain":true},{"metricName":" # relocations","quality":"Less is better","values":[0,0,0,0,0,0],"isMain":false},{"metricName":" # translocations","quality":"Less is better","values":[0,0,0,0,0,0],"isMain":false},{"metricName":" # inversions","quality":"Less is better","values":[0,0,0,0,0,0],"isMain":false},{"metricName":" # interspecies translocations","quality":"Less is better","values":[5,1,0,8,5,2],"isMain":false},{"metricName":"# possibly misassembled contigs","quality":"Less is better","values":[0,0,0,0,0,0],"isMain":false},{"metricName":" # possible misassemblies","quality":"Less is better","values":[0,0,0,0,0,0],"isMain":false},{"metricName":"# local misassemblies","quality":"Less is better","values":[0,0,0,0,0,0],"isMain":false},{"metricName":"# scaffold gap ext. mis.","quality":"Less is better","values":[0,0,0,0,0,0],"isMain":false},{"metricName":"# scaffold gap loc. mis.","quality":"Less is better","values":[0,0,0,0,0,0],"isMain":false}]],["Unaligned",[]],["Mismatches",[]],["Statistics without reference",[]],["K-mer-based statistics",[]],["Predicted genes",[]],["Similarity statistics",[]],["Reference statistics",[]]],[["Genome statistics",[]],["Reads mapping",[]],["Misassemblies",[{"metricName":"# misassemblies","quality":"Less is better","values":[4,1,4,8,1,3],"isMain":true},{"metricName":" # relocations","quality":"Less is better","values":[0,0,0,0,0,0],"isMain":false},{"metricName":" # translocations","quality":"Less is better","values":[0,0,0,0,0,0],"isMain":false},{"metricName":" # inversions","quality":"Less is better","values":[0,1,0,0,0,0],"isMain":false},{"metricName":" # interspecies translocations","quality":"Less is better","values":[4,0,4,8,1,3],"isMain":false},{"metricName":"# possibly misassembled contigs","quality":"Less is better","values":[0,0,0,0,0,0],"isMain":false},{"metricName":" # possible misassemblies","quality":"Less is better","values":[0,0,0,0,0,0],"isMain":false},{"metricName":"# local misassemblies","quality":"Less is better","values":[0,0,1,0,0,0],"isMain":false},{"metricName":"# scaffold gap ext. mis.","quality":"Less is better","values":[0,0,0,0,0,0],"isMain":false},{"metricName":"# scaffold gap loc. mis.","quality":"Less is better","values":[0,0,0,0,0,0],"isMain":false}]],["Unaligned",[]],["Mismatches",[]],["Statistics without reference",[]],["K-mer-based statistics",[]],["Predicted genes",[]],["Similarity statistics",[]],["Reference statistics",[]]],[["Genome statistics",[]],["Reads mapping",[]],["Misassemblies",[{"metricName":"# misassemblies","quality":"Less is better","values":[7,16,9,46,20,9],"isMain":true},{"metricName":" # relocations","quality":"Less is better","values":[0,1,1,1,0,0],"isMain":false},{"metricName":" # translocations","quality":"Less is better","values":[2,3,3,2,1,1],"isMain":false},{"metricName":" # inversions","quality":"Less is better","values":[0,0,0,1,0,0],"isMain":false},{"metricName":" # interspecies translocations","quality":"Less is better","values":[5,12,5,42,19,8],"isMain":false},{"metricName":"# possibly misassembled contigs","quality":"Less is better","values":[0,0,0,0,0,0],"isMain":false},{"metricName":" # possible misassemblies","quality":"Less is better","values":[0,0,0,0,0,0],"isMain":false},{"metricName":"# local misassemblies","quality":"Less is better","values":[1,2,2,5,4,1],"isMain":false},{"metricName":"# scaffold gap ext. mis.","quality":"Less is better","values":[0,0,0,0,0,0],"isMain":false},{"metricName":"# scaffold gap loc. mis.","quality":"Less is better","values":[0,0,0,0,0,0],"isMain":false}]],["Unaligned",[]],["Mismatches",[]],["Statistics without reference",[]],["K-mer-based statistics",[]],["Predicted genes",[]],["Similarity statistics",[]],["Reference statistics",[]]],[["Genome statistics",[]],["Reads mapping",[]],["Misassemblies",[{"metricName":"# misassemblies","quality":"Less is better","values":[11,19,7,22,21,10],"isMain":true},{"metricName":" # relocations","quality":"Less is better","values":[0,0,0,0,0,0],"isMain":false},{"metricName":" # translocations","quality":"Less is better","values":[0,0,0,0,0,0],"isMain":false},{"metricName":" # inversions","quality":"Less is better","values":[0,0,0,0,0,0],"isMain":false},{"metricName":" # interspecies translocations","quality":"Less is better","values":[11,19,7,22,21,10],"isMain":false},{"metricName":"# possibly misassembled contigs","quality":"Less is better","values":[0,0,0,0,0,0],"isMain":false},{"metricName":" # possible misassemblies","quality":"Less is better","values":[0,0,0,0,0,0],"isMain":false},{"metricName":"# local misassemblies","quality":"Less is better","values":[0,0,0,1,0,0],"isMain":false},{"metricName":"# scaffold gap ext. mis.","quality":"Less is better","values":[0,0,0,0,0,0],"isMain":false},{"metricName":"# scaffold gap loc. mis.","quality":"Less is better","values":[0,0,0,0,0,0],"isMain":false}]],["Unaligned",[]],["Mismatches",[]],["Statistics without reference",[]],["K-mer-based statistics",[]],["Predicted genes",[]],["Similarity statistics",[]],["Reference statistics",[]]],[["Genome statistics",[]],["Reads mapping",[]],["Misassemblies",[{"metricName":"# misassemblies","quality":"Less is better","values":[1,0,1,3,0,0],"isMain":true},{"metricName":" # relocations","quality":"Less is better","values":[0,0,0,0,0,0],"isMain":false},{"metricName":" # translocations","quality":"Less is better","values":[0,0,0,0,0,0],"isMain":false},{"metricName":" # inversions","quality":"Less is better","values":[0,0,0,0,0,0],"isMain":false},{"metricName":" # interspecies translocations","quality":"Less is better","values":[1,0,1,3,0,0],"isMain":false},{"metricName":"# possibly misassembled contigs","quality":"Less is better","values":[0,0,0,0,0,0],"isMain":false},{"metricName":" # possible misassemblies","quality":"Less is better","values":[0,0,0,0,0,0],"isMain":false},{"metricName":"# local misassemblies","quality":"Less is better","values":[0,0,0,0,0,0],"isMain":false},{"metricName":"# scaffold gap ext. mis.","quality":"Less is better","values":[0,0,0,0,0,0],"isMain":false},{"metricName":"# scaffold gap loc. mis.","quality":"Less is better","values":[0,0,0,0,0,0],"isMain":false}]],["Unaligned",[]],["Mismatches",[]],["Statistics without reference",[]],["K-mer-based statistics",[]],["Predicted genes",[]],["Similarity statistics",[]],["Reference statistics",[]]],[["Genome statistics",[]],["Reads mapping",[]],["Misassemblies",[{"metricName":"# misassemblies","quality":"Less is better","values":[55,59,10,39,53,27],"isMain":true},{"metricName":" # relocations","quality":"Less is better","values":[0,1,1,1,0,0],"isMain":false},{"metricName":" # translocations","quality":"Less is better","values":[1,2,2,3,2,3],"isMain":false},{"metricName":" # inversions","quality":"Less is better","values":[0,0,0,1,1,0],"isMain":false},{"metricName":" # interspecies translocations","quality":"Less is better","values":[54,56,7,34,50,24],"isMain":false},{"metricName":"# possibly misassembled contigs","quality":"Less is better","values":[0,0,0,0,0,0],"isMain":false},{"metricName":" # possible misassemblies","quality":"Less is better","values":[0,0,0,0,0,0],"isMain":false},{"metricName":"# local misassemblies","quality":"Less is better","values":[5,4,6,3,4,2],"isMain":false},{"metricName":"# scaffold gap ext. mis.","quality":"Less is better","values":[0,0,0,0,0,0],"isMain":false},{"metricName":"# scaffold gap loc. mis.","quality":"Less is better","values":[0,0,0,0,0,0],"isMain":false}]],["Unaligned",[]],["Mismatches",[]],["Statistics without reference",[]],["K-mer-based statistics",[]],["Predicted genes",[]],["Similarity statistics",[]],["Reference statistics",[]]],[["Genome statistics",[]],["Reads mapping",[]],["Misassemblies",[{"metricName":"# misassemblies","quality":"Less is better","values":[16,18,18,16,14,2],"isMain":true},{"metricName":" # relocations","quality":"Less is better","values":[1,0,0,0,2,0],"isMain":false},{"metricName":" # translocations","quality":"Less is better","values":[0,0,0,0,0,0],"isMain":false},{"metricName":" # inversions","quality":"Less is better","values":[0,0,3,2,2,0],"isMain":false},{"metricName":" # interspecies translocations","quality":"Less is better","values":[15,18,15,14,10,2],"isMain":false},{"metricName":"# possibly misassembled contigs","quality":"Less is better","values":[0,0,1,0,1,0],"isMain":false},{"metricName":" # possible misassemblies","quality":"Less is better","values":[0,0,1,0,1,0],"isMain":false},{"metricName":"# local misassemblies","quality":"Less is better","values":[1,0,2,2,2,0],"isMain":false},{"metricName":"# scaffold gap ext. mis.","quality":"Less is better","values":[0,0,0,0,0,0],"isMain":false},{"metricName":"# scaffold gap loc. mis.","quality":"Less is better","values":[0,0,0,0,0,0],"isMain":false}]],["Unaligned",[]],["Mismatches",[]],["Statistics without reference",[]],["K-mer-based statistics",[]],["Predicted genes",[]],["Similarity statistics",[]],["Reference statistics",[]]],[["Genome statistics",[]],["Reads mapping",[]],["Misassemblies",[{"metricName":"# misassemblies","quality":"Less is better","values":[1,2,4,5,7,0],"isMain":true},{"metricName":" # relocations","quality":"Less is better","values":[0,1,0,1,0,0],"isMain":false},{"metricName":" # translocations","quality":"Less is better","values":[0,0,0,0,0,0],"isMain":false},{"metricName":" # inversions","quality":"Less is better","values":[0,0,0,0,1,0],"isMain":false},{"metricName":" # interspecies translocations","quality":"Less is better","values":[1,1,4,4,6,0],"isMain":false},{"metricName":"# possibly misassembled contigs","quality":"Less is better","values":[0,0,0,0,0,0],"isMain":false},{"metricName":" # possible misassemblies","quality":"Less is better","values":[0,0,0,0,0,0],"isMain":false},{"metricName":"# local misassemblies","quality":"Less is better","values":[0,0,0,0,1,0],"isMain":false},{"metricName":"# scaffold gap ext. mis.","quality":"Less is better","values":[0,0,0,0,0,0],"isMain":false},{"metricName":"# scaffold gap loc. mis.","quality":"Less is better","values":[0,0,0,0,0,0],"isMain":false}]],["Unaligned",[]],["Mismatches",[]],["Statistics without reference",[]],["K-mer-based statistics",[]],["Predicted genes",[]],["Similarity statistics",[]],["Reference statistics",[]]],[["Genome statistics",[]],["Reads mapping",[]],["Misassemblies",[{"metricName":"# misassemblies","quality":"Less is better","values":[30,42,23,43,46,30],"isMain":true},{"metricName":" # relocations","quality":"Less is better","values":[1,4,1,1,3,2],"isMain":false},{"metricName":" # translocations","quality":"Less is better","values":[12,15,19,15,34,9],"isMain":false},{"metricName":" # inversions","quality":"Less is better","values":[0,1,1,0,0,0],"isMain":false},{"metricName":" # interspecies translocations","quality":"Less is better","values":[17,22,2,27,9,19],"isMain":false},{"metricName":"# possibly misassembled contigs","quality":"Less is better","values":[0,0,0,0,0,0],"isMain":false},{"metricName":" # possible misassemblies","quality":"Less is better","values":[0,0,0,0,0,0],"isMain":false},{"metricName":"# local misassemblies","quality":"Less is better","values":[3,2,4,2,7,1],"isMain":false},{"metricName":"# scaffold gap ext. mis.","quality":"Less is better","values":[0,0,0,0,0,0],"isMain":false},{"metricName":"# scaffold gap loc. mis.","quality":"Less is better","values":[0,0,0,0,0,0],"isMain":false}]],["Unaligned",[]],["Mismatches",[]],["Statistics without reference",[]],["K-mer-based statistics",[]],["Predicted genes",[]],["Similarity statistics",[]],["Reference statistics",[]]],[["Genome statistics",[]],["Reads mapping",[]],["Misassemblies",[{"metricName":"# misassemblies","quality":"Less is better","values":[45,44,14,43,38,33],"isMain":true},{"metricName":" # relocations","quality":"Less is better","values":[1,1,2,1,1,1],"isMain":false},{"metricName":" # translocations","quality":"Less is better","values":[6,6,8,6,3,3],"isMain":false},{"metricName":" # inversions","quality":"Less is better","values":[0,0,0,0,0,0],"isMain":false},{"metricName":" # interspecies translocations","quality":"Less is better","values":[38,37,4,36,34,29],"isMain":false},{"metricName":"# possibly misassembled contigs","quality":"Less is better","values":[0,0,1,0,0,0],"isMain":false},{"metricName":" # possible misassemblies","quality":"Less is better","values":[0,0,1,0,0,0],"isMain":false},{"metricName":"# local misassemblies","quality":"Less is better","values":[1,0,0,0,2,0],"isMain":false},{"metricName":"# scaffold gap ext. mis.","quality":"Less is better","values":[0,0,0,0,0,0],"isMain":false},{"metricName":"# scaffold gap loc. mis.","quality":"Less is better","values":[0,0,0,0,0,0],"isMain":false}]],["Unaligned",[]],["Mismatches",[]],["Statistics without reference",[]],["K-mer-based statistics",[]],["Predicted genes",[]],["Similarity statistics",[]],["Reference statistics",[]]],[["Genome statistics",[]],["Reads mapping",[]],["Misassemblies",[{"metricName":"# misassemblies","quality":"Less is better","values":[7,21,8,21,23,7],"isMain":true},{"metricName":" # relocations","quality":"Less is better","values":[0,0,0,0,0,0],"isMain":false},{"metricName":" # translocations","quality":"Less is better","values":[0,0,0,0,0,0],"isMain":false},{"metricName":" # inversions","quality":"Less is better","values":[1,0,0,0,0,0],"isMain":false},{"metricName":" # interspecies translocations","quality":"Less is better","values":[6,21,8,21,23,7],"isMain":false},{"metricName":"# possibly misassembled contigs","quality":"Less is better","values":[0,0,0,0,0,0],"isMain":false},{"metricName":" # possible misassemblies","quality":"Less is better","values":[0,0,0,0,0,0],"isMain":false},{"metricName":"# local misassemblies","quality":"Less is better","values":[0,0,0,0,0,0],"isMain":false},{"metricName":"# scaffold gap ext. mis.","quality":"Less is better","values":[0,0,0,0,0,0],"isMain":false},{"metricName":"# scaffold gap loc. mis.","quality":"Less is better","values":[0,0,0,0,0,0],"isMain":false}]],["Unaligned",[]],["Mismatches",[]],["Statistics without reference",[]],["K-mer-based statistics",[]],["Predicted genes",[]],["Similarity statistics",[]],["Reference statistics",[]]],[["Genome statistics",[]],["Reads mapping",[]],["Misassemblies",[{"metricName":"# misassemblies","quality":"Less is better","values":[9,15,10,16,9,7],"isMain":true},{"metricName":" # relocations","quality":"Less is better","values":[0,0,0,0,0,0],"isMain":false},{"metricName":" # translocations","quality":"Less is better","values":[5,7,4,5,3,4],"isMain":false},{"metricName":" # inversions","quality":"Less is better","values":[0,0,0,0,0,0],"isMain":false},{"metricName":" # interspecies translocations","quality":"Less is better","values":[4,8,6,11,6,3],"isMain":false},{"metricName":"# possibly misassembled contigs","quality":"Less is better","values":[0,0,0,0,0,0],"isMain":false},{"metricName":" # possible misassemblies","quality":"Less is better","values":[0,0,0,0,0,0],"isMain":false},{"metricName":"# local misassemblies","quality":"Less is better","values":[0,0,0,1,0,0],"isMain":false},{"metricName":"# scaffold gap ext. mis.","quality":"Less is better","values":[0,0,0,0,0,0],"isMain":false},{"metricName":"# scaffold gap loc. mis.","quality":"Less is better","values":[0,0,0,0,0,0],"isMain":false}]],["Unaligned",[]],["Mismatches",[]],["Statistics without reference",[]],["K-mer-based statistics",[]],["Predicted genes",[]],["Similarity statistics",[]],["Reference statistics",[]]],[["Genome statistics",[]],["Reads mapping",[]],["Misassemblies",[{"metricName":"# misassemblies","quality":"Less is better","values":[44,46,43,65,69,18],"isMain":true},{"metricName":" # relocations","quality":"Less is better","values":[0,1,0,0,1,2],"isMain":false},{"metricName":" # translocations","quality":"Less is better","values":[10,6,11,9,16,7],"isMain":false},{"metricName":" # inversions","quality":"Less is better","values":[1,0,2,2,1,0],"isMain":false},{"metricName":" # interspecies translocations","quality":"Less is better","values":[33,39,30,54,51,9],"isMain":false},{"metricName":"# possibly misassembled contigs","quality":"Less is better","values":[0,0,0,0,1,0],"isMain":false},{"metricName":" # possible misassemblies","quality":"Less is better","values":[0,0,0,0,1,0],"isMain":false},{"metricName":"# local misassemblies","quality":"Less is better","values":[3,4,6,3,6,0],"isMain":false},{"metricName":"# scaffold gap ext. mis.","quality":"Less is better","values":[0,0,0,0,0,0],"isMain":false},{"metricName":"# scaffold gap loc. mis.","quality":"Less is better","values":[0,0,0,0,0,0],"isMain":false}]],["Unaligned",[]],["Mismatches",[]],["Statistics without reference",[]],["K-mer-based statistics",[]],["Predicted genes",[]],["Similarity statistics",[]],["Reference statistics",[]]],[["Genome statistics",[]],["Reads mapping",[]],["Misassemblies",[{"metricName":"# misassemblies","quality":"Less is better","values":[72,86,7,62,72,78],"isMain":true},{"metricName":" # relocations","quality":"Less is better","values":[4,7,2,3,4,1],"isMain":false},{"metricName":" # translocations","quality":"Less is better","values":[1,0,0,0,0,0],"isMain":false},{"metricName":" # inversions","quality":"Less is better","values":[2,1,0,2,1,0],"isMain":false},{"metricName":" # interspecies translocations","quality":"Less is better","values":[65,78,5,57,67,77],"isMain":false},{"metricName":"# possibly misassembled contigs","quality":"Less is better","values":[0,0,0,0,0,0],"isMain":false},{"metricName":" # possible misassemblies","quality":"Less is better","values":[0,0,0,0,0,0],"isMain":false},{"metricName":"# local misassemblies","quality":"Less is better","values":[1,2,7,5,7,1],"isMain":false},{"metricName":"# scaffold gap ext. mis.","quality":"Less is better","values":[0,0,0,0,0,0],"isMain":false},{"metricName":"# scaffold gap loc. mis.","quality":"Less is better","values":[0,0,0,0,0,0],"isMain":false}]],["Unaligned",[]],["Mismatches",[]],["Statistics without reference",[]],["K-mer-based statistics",[]],["Predicted genes",[]],["Similarity statistics",[]],["Reference statistics",[]]],[["Genome statistics",[]],["Reads mapping",[]],["Misassemblies",[{"metricName":"# misassemblies","quality":"Less is better","values":[8,7,5,7,11,2],"isMain":true},{"metricName":" # relocations","quality":"Less is better","values":[0,0,0,0,0,0],"isMain":false},{"metricName":" # translocations","quality":"Less is better","values":[0,0,0,0,0,0],"isMain":false},{"metricName":" # inversions","quality":"Less is better","values":[0,0,0,0,0,0],"isMain":false},{"metricName":" # interspecies translocations","quality":"Less is better","values":[8,7,5,7,11,2],"isMain":false},{"metricName":"# possibly misassembled contigs","quality":"Less is better","values":[0,0,0,0,0,0],"isMain":false},{"metricName":" # possible misassemblies","quality":"Less is better","values":[0,0,0,0,0,0],"isMain":false},{"metricName":"# local misassemblies","quality":"Less is better","values":[0,0,0,0,1,0],"isMain":false},{"metricName":"# scaffold gap ext. mis.","quality":"Less is better","values":[0,0,0,0,0,0],"isMain":false},{"metricName":"# scaffold gap loc. mis.","quality":"Less is better","values":[0,0,0,0,0,0],"isMain":false}]],["Unaligned",[]],["Mismatches",[]],["Statistics without reference",[]],["K-mer-based statistics",[]],["Predicted genes",[]],["Similarity statistics",[]],["Reference statistics",[]]],[["Genome statistics",[]],["Reads mapping",[]],["Misassemblies",[{"metricName":"# misassemblies","quality":"Less is better","values":[0,1,1,0,0,0],"isMain":true},{"metricName":" # relocations","quality":"Less is better","values":[0,0,0,0,0,0],"isMain":false},{"metricName":" # translocations","quality":"Less is better","values":[0,0,0,0,0,0],"isMain":false},{"metricName":" # inversions","quality":"Less is better","values":[0,0,0,0,0,0],"isMain":false},{"metricName":" # interspecies translocations","quality":"Less is better","values":[0,1,1,0,0,0],"isMain":false},{"metricName":"# possibly misassembled contigs","quality":"Less is better","values":[0,0,0,0,0,0],"isMain":false},{"metricName":" # possible misassemblies","quality":"Less is better","values":[0,0,0,0,0,0],"isMain":false},{"metricName":"# local misassemblies","quality":"Less is better","values":[0,0,0,0,0,0],"isMain":false},{"metricName":"# scaffold gap ext. mis.","quality":"Less is better","values":[0,0,0,0,0,0],"isMain":false},{"metricName":"# scaffold gap loc. mis.","quality":"Less is better","values":[0,0,0,0,0,0],"isMain":false}]],["Unaligned",[]],["Mismatches",[]],["Statistics without reference",[]],["K-mer-based statistics",[]],["Predicted genes",[]],["Similarity statistics",[]],["Reference statistics",[]]],[["Genome statistics",[]],["Reads mapping",[]],["Misassemblies",[{"metricName":"# misassemblies","quality":"Less is better","values":[8,36,28,54,36,7],"isMain":true},{"metricName":" # relocations","quality":"Less is better","values":[0,0,1,3,0,0],"isMain":false},{"metricName":" # translocations","quality":"Less is better","values":[0,0,0,0,0,0],"isMain":false},{"metricName":" # inversions","quality":"Less is better","values":[0,3,1,1,1,0],"isMain":false},{"metricName":" # interspecies translocations","quality":"Less is better","values":[8,33,26,50,35,7],"isMain":false},{"metricName":"# possibly misassembled contigs","quality":"Less is better","values":[0,0,0,1,3,0],"isMain":false},{"metricName":" # possible misassemblies","quality":"Less is better","values":[0,0,0,1,3,0],"isMain":false},{"metricName":"# local misassemblies","quality":"Less is better","values":[1,2,3,1,1,0],"isMain":false},{"metricName":"# scaffold gap ext. mis.","quality":"Less is better","values":[0,0,0,0,0,0],"isMain":false},{"metricName":"# scaffold gap loc. mis.","quality":"Less is better","values":[0,0,0,0,0,0],"isMain":false}]],["Unaligned",[]],["Mismatches",[]],["Statistics without reference",[]],["K-mer-based statistics",[]],["Predicted genes",[]],["Similarity statistics",[]],["Reference statistics",[]]],[["Genome statistics",[]],["Reads mapping",[]],["Misassemblies",[{"metricName":"# misassemblies","quality":"Less is better","values":[5,6,0,7,3,6],"isMain":true},{"metricName":" # relocations","quality":"Less is better","values":[0,0,0,0,0,0],"isMain":false},{"metricName":" # translocations","quality":"Less is better","values":[0,0,0,0,0,0],"isMain":false},{"metricName":" # inversions","quality":"Less is better","values":[0,0,0,0,0,0],"isMain":false},{"metricName":" # interspecies translocations","quality":"Less is better","values":[5,6,0,7,3,6],"isMain":false},{"metricName":"# possibly misassembled contigs","quality":"Less is better","values":[0,0,0,0,0,0],"isMain":false},{"metricName":" # possible misassemblies","quality":"Less is better","values":[0,0,0,0,0,0],"isMain":false},{"metricName":"# local misassemblies","quality":"Less is better","values":[0,0,0,0,0,0],"isMain":false},{"metricName":"# scaffold gap ext. mis.","quality":"Less is better","values":[0,0,0,0,0,0],"isMain":false},{"metricName":"# scaffold gap loc. mis.","quality":"Less is better","values":[0,0,0,0,0,0],"isMain":false}]],["Unaligned",[]],["Mismatches",[]],["Statistics without reference",[]],["K-mer-based statistics",[]],["Predicted genes",[]],["Similarity statistics",[]],["Reference statistics",[]]],[["Genome statistics",[]],["Reads mapping",[]],["Misassemblies",[{"metricName":"# misassemblies","quality":"Less is better","values":[3,6,0,11,6,7],"isMain":true},{"metricName":" # relocations","quality":"Less is better","values":[0,0,0,0,0,0],"isMain":false},{"metricName":" # translocations","quality":"Less is better","values":[0,0,0,1,1,0],"isMain":false},{"metricName":" # inversions","quality":"Less is better","values":[1,0,0,0,0,0],"isMain":false},{"metricName":" # interspecies translocations","quality":"Less is better","values":[2,6,0,10,5,7],"isMain":false},{"metricName":"# possibly misassembled contigs","quality":"Less is better","values":[0,0,0,0,0,0],"isMain":false},{"metricName":" # possible misassemblies","quality":"Less is better","values":[0,0,0,0,0,0],"isMain":false},{"metricName":"# local misassemblies","quality":"Less is better","values":[0,0,0,1,0,0],"isMain":false},{"metricName":"# scaffold gap ext. mis.","quality":"Less is better","values":[0,0,0,0,0,0],"isMain":false},{"metricName":"# scaffold gap loc. mis.","quality":"Less is better","values":[0,0,0,0,0,0],"isMain":false}]],["Unaligned",[]],["Mismatches",[]],["Statistics without reference",[]],["K-mer-based statistics",[]],["Predicted genes",[]],["Similarity statistics",[]],["Reference statistics",[]]],[["Genome statistics",[]],["Reads mapping",[]],["Misassemblies",[{"metricName":"# misassemblies","quality":"Less is better","values":[23,7,5,9,8,3],"isMain":true},{"metricName":" # relocations","quality":"Less is better","values":[0,0,0,0,0,0],"isMain":false},{"metricName":" # translocations","quality":"Less is better","values":[1,0,0,0,0,0],"isMain":false},{"metricName":" # inversions","quality":"Less is better","values":[0,0,0,0,0,0],"isMain":false},{"metricName":" # interspecies translocations","quality":"Less is better","values":[22,7,5,9,8,3],"isMain":false},{"metricName":"# possibly misassembled contigs","quality":"Less is better","values":[1,0,0,1,0,0],"isMain":false},{"metricName":" # possible misassemblies","quality":"Less is better","values":[1,0,0,1,0,0],"isMain":false},{"metricName":"# local misassemblies","quality":"Less is better","values":[1,0,0,0,0,0],"isMain":false},{"metricName":"# scaffold gap ext. mis.","quality":"Less is better","values":[0,0,0,0,0,0],"isMain":false},{"metricName":"# scaffold gap loc. mis.","quality":"Less is better","values":[0,0,0,0,0,0],"isMain":false}]],["Unaligned",[]],["Mismatches",[]],["Statistics without reference",[]],["K-mer-based statistics",[]],["Predicted genes",[]],["Similarity statistics",[]],["Reference statistics",[]]],[["Genome statistics",[]],["Reads mapping",[]],["Misassemblies",[{"metricName":"# misassemblies","quality":"Less is better","values":[30,32,7,32,32,14],"isMain":true},{"metricName":" # relocations","quality":"Less is better","values":[1,1,2,1,1,0],"isMain":false},{"metricName":" # translocations","quality":"Less is better","values":[1,1,1,1,0,0],"isMain":false},{"metricName":" # inversions","quality":"Less is better","values":[0,0,0,0,0,0],"isMain":false},{"metricName":" # interspecies translocations","quality":"Less is better","values":[28,30,4,30,31,14],"isMain":false},{"metricName":"# possibly misassembled contigs","quality":"Less is better","values":[0,0,0,0,0,0],"isMain":false},{"metricName":" # possible misassemblies","quality":"Less is better","values":[0,0,0,0,0,0],"isMain":false},{"metricName":"# local misassemblies","quality":"Less is better","values":[0,0,0,0,0,0],"isMain":false},{"metricName":"# scaffold gap ext. mis.","quality":"Less is better","values":[0,0,0,0,0,0],"isMain":false},{"metricName":"# scaffold gap loc. mis.","quality":"Less is better","values":[0,0,0,0,0,0],"isMain":false}]],["Unaligned",[]],["Mismatches",[]],["Statistics without reference",[]],["K-mer-based statistics",[]],["Predicted genes",[]],["Similarity statistics",[]],["Reference statistics",[]]],[["Genome statistics",[]],["Reads mapping",[]],["Misassemblies",[{"metricName":"# misassemblies","quality":"Less is better","values":[96,24,24,55,26,6],"isMain":true},{"metricName":" # relocations","quality":"Less is better","values":[0,2,1,3,0,0],"isMain":false},{"metricName":" # translocations","quality":"Less is better","values":[1,8,5,10,6,5],"isMain":false},{"metricName":" # inversions","quality":"Less is better","values":[0,0,1,1,1,0],"isMain":false},{"metricName":" # interspecies translocations","quality":"Less is better","values":[95,14,17,41,19,1],"isMain":false},{"metricName":"# possibly misassembled contigs","quality":"Less is better","values":[1,0,0,0,0,0],"isMain":false},{"metricName":" # possible misassemblies","quality":"Less is better","values":[1,0,0,0,0,0],"isMain":false},{"metricName":"# local misassemblies","quality":"Less is better","values":[2,1,0,1,1,0],"isMain":false},{"metricName":"# scaffold gap ext. mis.","quality":"Less is better","values":[0,0,0,0,0,0],"isMain":false},{"metricName":"# scaffold gap loc. mis.","quality":"Less is better","values":[0,0,0,0,0,0],"isMain":false}]],["Unaligned",[]],["Mismatches",[]],["Statistics without reference",[]],["K-mer-based statistics",[]],["Predicted genes",[]],["Similarity statistics",[]],["Reference statistics",[]]],[["Genome statistics",[]],["Reads mapping",[]],["Misassemblies",[{"metricName":"# misassemblies","quality":"Less is better","values":[1,1,2,1,1,0],"isMain":true},{"metricName":" # relocations","quality":"Less is better","values":[0,0,0,0,0,0],"isMain":false},{"metricName":" # translocations","quality":"Less is better","values":[0,0,0,0,0,0],"isMain":false},{"metricName":" # inversions","quality":"Less is better","values":[0,0,0,0,0,0],"isMain":false},{"metricName":" # interspecies translocations","quality":"Less is better","values":[1,1,2,1,1,0],"isMain":false},{"metricName":"# possibly misassembled contigs","quality":"Less is better","values":[0,1,0,0,0,0],"isMain":false},{"metricName":" # possible misassemblies","quality":"Less is better","values":[0,1,0,0,0,0],"isMain":false},{"metricName":"# local misassemblies","quality":"Less is better","values":[0,0,0,0,0,0],"isMain":false},{"metricName":"# scaffold gap ext. mis.","quality":"Less is better","values":[0,0,0,0,0,0],"isMain":false},{"metricName":"# scaffold gap loc. mis.","quality":"Less is better","values":[0,0,0,0,0,0],"isMain":false}]],["Unaligned",[]],["Mismatches",[]],["Statistics without reference",[]],["K-mer-based statistics",[]],["Predicted genes",[]],["Similarity statistics",[]],["Reference statistics",[]]],[["Genome statistics",[]],["Reads mapping",[]],["Misassemblies",[{"metricName":"# misassemblies","quality":"Less is better","values":[9,17,18,18,22,7],"isMain":true},{"metricName":" # relocations","quality":"Less is better","values":[0,0,0,1,0,0],"isMain":false},{"metricName":" # translocations","quality":"Less is better","values":[1,1,3,1,0,0],"isMain":false},{"metricName":" # inversions","quality":"Less is better","values":[0,2,0,0,1,0],"isMain":false},{"metricName":" # interspecies translocations","quality":"Less is better","values":[8,14,15,16,21,7],"isMain":false},{"metricName":"# possibly misassembled contigs","quality":"Less is better","values":[0,0,0,0,0,0],"isMain":false},{"metricName":" # possible misassemblies","quality":"Less is better","values":[0,0,0,0,0,0],"isMain":false},{"metricName":"# local misassemblies","quality":"Less is better","values":[0,1,0,0,0,0],"isMain":false},{"metricName":"# scaffold gap ext. mis.","quality":"Less is better","values":[0,0,0,0,0,0],"isMain":false},{"metricName":"# scaffold gap loc. mis.","quality":"Less is better","values":[0,0,0,0,0,0],"isMain":false}]],["Unaligned",[]],["Mismatches",[]],["Statistics without reference",[]],["K-mer-based statistics",[]],["Predicted genes",[]],["Similarity statistics",[]],["Reference statistics",[]]],[["Genome statistics",[]],["Reads mapping",[]],["Misassemblies",[{"metricName":"# misassemblies","quality":"Less is better","values":[6,14,7,11,9,7],"isMain":true},{"metricName":" # relocations","quality":"Less is better","values":[0,0,1,0,0,0],"isMain":false},{"metricName":" # translocations","quality":"Less is better","values":[0,0,0,0,0,1],"isMain":false},{"metricName":" # inversions","quality":"Less is better","values":[0,0,0,0,0,0],"isMain":false},{"metricName":" # interspecies translocations","quality":"Less is better","values":[6,14,6,11,9,6],"isMain":false},{"metricName":"# possibly misassembled contigs","quality":"Less is better","values":[0,0,0,0,0,0],"isMain":false},{"metricName":" # possible misassemblies","quality":"Less is better","values":[0,0,0,0,0,0],"isMain":false},{"metricName":"# local misassemblies","quality":"Less is better","values":[0,1,0,0,1,1],"isMain":false},{"metricName":"# scaffold gap ext. mis.","quality":"Less is better","values":[0,0,0,0,0,0],"isMain":false},{"metricName":"# scaffold gap loc. mis.","quality":"Less is better","values":[0,0,0,0,0,0],"isMain":false}]],["Unaligned",[]],["Mismatches",[]],["Statistics without reference",[]],["K-mer-based statistics",[]],["Predicted genes",[]],["Similarity statistics",[]],["Reference statistics",[]]],[["Genome statistics",[]],["Reads mapping",[]],["Misassemblies",[{"metricName":"# misassemblies","quality":"Less is better","values":[11,11,3,16,9,3],"isMain":true},{"metricName":" # relocations","quality":"Less is better","values":[0,0,0,0,0,0],"isMain":false},{"metricName":" # translocations","quality":"Less is better","values":[0,0,1,1,0,0],"isMain":false},{"metricName":" # inversions","quality":"Less is better","values":[0,0,0,0,0,0],"isMain":false},{"metricName":" # interspecies translocations","quality":"Less is better","values":[11,11,2,15,9,3],"isMain":false},{"metricName":"# possibly misassembled contigs","quality":"Less is better","values":[0,0,0,0,0,0],"isMain":false},{"metricName":" # possible misassemblies","quality":"Less is better","values":[0,0,0,0,0,0],"isMain":false},{"metricName":"# local misassemblies","quality":"Less is better","values":[0,0,2,1,1,0],"isMain":false},{"metricName":"# scaffold gap ext. mis.","quality":"Less is better","values":[0,0,0,0,0,0],"isMain":false},{"metricName":"# scaffold gap loc. mis.","quality":"Less is better","values":[0,0,0,0,0,0],"isMain":false}]],["Unaligned",[]],["Mismatches",[]],["Statistics without reference",[]],["K-mer-based statistics",[]],["Predicted genes",[]],["Similarity statistics",[]],["Reference statistics",[]]],[["Genome statistics",[]],["Reads mapping",[]],["Misassemblies",[{"metricName":"# misassemblies","quality":"Less is better","values":[11,19,12,30,29,10],"isMain":true},{"metricName":" # relocations","quality":"Less is better","values":[0,0,0,0,0,0],"isMain":false},{"metricName":" # translocations","quality":"Less is better","values":[5,4,0,3,6,5],"isMain":false},{"metricName":" # inversions","quality":"Less is better","values":[0,1,0,0,0,0],"isMain":false},{"metricName":" # interspecies translocations","quality":"Less is better","values":[6,14,12,27,23,5],"isMain":false},{"metricName":"# possibly misassembled contigs","quality":"Less is better","values":[3,2,1,0,1,0],"isMain":false},{"metricName":" # possible misassemblies","quality":"Less is better","values":[3,2,1,0,1,0],"isMain":false},{"metricName":"# local misassemblies","quality":"Less is better","values":[0,0,2,0,1,0],"isMain":false},{"metricName":"# scaffold gap ext. mis.","quality":"Less is better","values":[0,0,0,0,0,0],"isMain":false},{"metricName":"# scaffold gap loc. mis.","quality":"Less is better","values":[0,0,0,0,0,0],"isMain":false}]],["Unaligned",[]],["Mismatches",[]],["Statistics without reference",[]],["K-mer-based statistics",[]],["Predicted genes",[]],["Similarity statistics",[]],["Reference statistics",[]]],[["Genome statistics",[]],["Reads mapping",[]],["Misassemblies",[{"metricName":"# misassemblies","quality":"Less is better","values":[1,1,0,0,0,0],"isMain":true},{"metricName":" # relocations","quality":"Less is better","values":[0,0,0,0,0,0],"isMain":false},{"metricName":" # translocations","quality":"Less is better","values":[0,0,0,0,0,0],"isMain":false},{"metricName":" # inversions","quality":"Less is better","values":[0,0,0,0,0,0],"isMain":false},{"metricName":" # interspecies translocations","quality":"Less is better","values":[1,1,0,0,0,0],"isMain":false},{"metricName":"# possibly misassembled contigs","quality":"Less is better","values":[0,0,0,0,0,0],"isMain":false},{"metricName":" # possible misassemblies","quality":"Less is better","values":[0,0,0,0,0,0],"isMain":false},{"metricName":"# local misassemblies","quality":"Less is better","values":[0,0,0,0,0,0],"isMain":false},{"metricName":"# scaffold gap ext. mis.","quality":"Less is better","values":[0,0,0,0,0,0],"isMain":false},{"metricName":"# scaffold gap loc. mis.","quality":"Less is better","values":[0,0,0,0,0,0],"isMain":false}]],["Unaligned",[]],["Mismatches",[]],["Statistics without reference",[]],["K-mer-based statistics",[]],["Predicted genes",[]],["Similarity statistics",[]],["Reference statistics",[]]],[["Genome statistics",[]],["Reads mapping",[]],["Misassemblies",[{"metricName":"# misassemblies","quality":"Less is better","values":[1,3,0,4,1,0],"isMain":true},{"metricName":" # relocations","quality":"Less is better","values":[0,0,0,0,0,0],"isMain":false},{"metricName":" # translocations","quality":"Less is better","values":[0,0,0,0,0,0],"isMain":false},{"metricName":" # inversions","quality":"Less is better","values":[0,0,0,0,0,0],"isMain":false},{"metricName":" # interspecies translocations","quality":"Less is better","values":[1,3,0,4,1,0],"isMain":false},{"metricName":"# possibly misassembled contigs","quality":"Less is better","values":[0,0,0,0,0,0],"isMain":false},{"metricName":" # possible misassemblies","quality":"Less is better","values":[0,0,0,0,0,0],"isMain":false},{"metricName":"# local misassemblies","quality":"Less is better","values":[0,0,0,0,0,0],"isMain":false},{"metricName":"# scaffold gap ext. mis.","quality":"Less is better","values":[0,0,0,0,0,0],"isMain":false},{"metricName":"# scaffold gap loc. mis.","quality":"Less is better","values":[0,0,0,0,0,0],"isMain":false}]],["Unaligned",[]],["Mismatches",[]],["Statistics without reference",[]],["K-mer-based statistics",[]],["Predicted genes",[]],["Similarity statistics",[]],["Reference statistics",[]]],[["Genome statistics",[]],["Reads mapping",[]],["Misassemblies",[{"metricName":"# misassemblies","quality":"Less is better","values":[36,29,14,23,23,3],"isMain":true},{"metricName":" # relocations","quality":"Less is better","values":[0,0,0,0,0,0],"isMain":false},{"metricName":" # translocations","quality":"Less is better","values":[2,3,2,2,1,1],"isMain":false},{"metricName":" # inversions","quality":"Less is better","values":[3,2,0,0,1,0],"isMain":false},{"metricName":" # interspecies translocations","quality":"Less is better","values":[31,24,12,21,21,2],"isMain":false},{"metricName":"# possibly misassembled contigs","quality":"Less is better","values":[0,1,0,0,0,0],"isMain":false},{"metricName":" # possible misassemblies","quality":"Less is better","values":[0,1,0,0,0,0],"isMain":false},{"metricName":"# local misassemblies","quality":"Less is better","values":[1,1,1,3,2,0],"isMain":false},{"metricName":"# scaffold gap ext. mis.","quality":"Less is better","values":[0,0,0,0,0,0],"isMain":false},{"metricName":"# scaffold gap loc. mis.","quality":"Less is better","values":[0,0,0,0,0,0],"isMain":false}]],["Unaligned",[]],["Mismatches",[]],["Statistics without reference",[]],["K-mer-based statistics",[]],["Predicted genes",[]],["Similarity statistics",[]],["Reference statistics",[]]],[["Genome statistics",[]],["Reads mapping",[]],["Misassemblies",[{"metricName":"# misassemblies","quality":"Less is better","values":[3,4,0,11,6,1],"isMain":true},{"metricName":" # relocations","quality":"Less is better","values":[0,0,0,0,0,0],"isMain":false},{"metricName":" # translocations","quality":"Less is better","values":[0,0,0,0,0,0],"isMain":false},{"metricName":" # inversions","quality":"Less is better","values":[0,0,0,0,0,0],"isMain":false},{"metricName":" # interspecies translocations","quality":"Less is better","values":[3,4,0,11,6,1],"isMain":false},{"metricName":"# possibly misassembled contigs","quality":"Less is better","values":[0,0,0,0,0,0],"isMain":false},{"metricName":" # possible misassemblies","quality":"Less is better","values":[0,0,0,0,0,0],"isMain":false},{"metricName":"# local misassemblies","quality":"Less is better","values":[0,0,0,0,0,0],"isMain":false},{"metricName":"# scaffold gap ext. mis.","quality":"Less is better","values":[0,0,0,0,0,0],"isMain":false},{"metricName":"# scaffold gap loc. mis.","quality":"Less is better","values":[0,0,0,0,0,0],"isMain":false}]],["Unaligned",[]],["Mismatches",[]],["Statistics without reference",[]],["K-mer-based statistics",[]],["Predicted genes",[]],["Similarity statistics",[]],["Reference statistics",[]]],[["Genome statistics",[]],["Reads mapping",[]],["Misassemblies",[{"metricName":"# misassemblies","quality":"Less is better","values":[30,182,108,160,87,26],"isMain":true},{"metricName":" # relocations","quality":"Less is better","values":[0,0,0,3,0,0],"isMain":false},{"metricName":" # translocations","quality":"Less is better","values":[0,28,47,48,30,21],"isMain":false},{"metricName":" # inversions","quality":"Less is better","values":[0,29,3,3,13,0],"isMain":false},{"metricName":" # interspecies translocations","quality":"Less is better","values":[30,125,58,106,44,5],"isMain":false},{"metricName":"# possibly misassembled contigs","quality":"Less is better","values":[0,0,0,0,1,0],"isMain":false},{"metricName":" # possible misassemblies","quality":"Less is better","values":[0,0,0,0,1,0],"isMain":false},{"metricName":"# local misassemblies","quality":"Less is better","values":[1,19,7,12,7,0],"isMain":false},{"metricName":"# scaffold gap ext. mis.","quality":"Less is better","values":[0,0,0,0,0,0],"isMain":false},{"metricName":"# scaffold gap loc. mis.","quality":"Less is better","values":[0,0,0,0,0,0],"isMain":false}]],["Unaligned",[]],["Mismatches",[]],["Statistics without reference",[]],["K-mer-based statistics",[]],["Predicted genes",[]],["Similarity statistics",[]],["Reference statistics",[]]],[["Genome statistics",[]],["Reads mapping",[]],["Misassemblies",[{"metricName":"# misassemblies","quality":"Less is better","values":[2,3,0,3,0,0],"isMain":true},{"metricName":" # relocations","quality":"Less is better","values":[0,0,0,0,0,0],"isMain":false},{"metricName":" # translocations","quality":"Less is better","values":[0,0,0,0,0,0],"isMain":false},{"metricName":" # inversions","quality":"Less is better","values":[0,0,0,0,0,0],"isMain":false},{"metricName":" # interspecies translocations","quality":"Less is better","values":[2,3,0,3,0,0],"isMain":false},{"metricName":"# possibly misassembled contigs","quality":"Less is better","values":[0,0,0,0,0,0],"isMain":false},{"metricName":" # possible misassemblies","quality":"Less is better","values":[0,0,0,0,0,0],"isMain":false},{"metricName":"# local misassemblies","quality":"Less is better","values":[0,0,0,0,0,0],"isMain":false},{"metricName":"# scaffold gap ext. mis.","quality":"Less is better","values":[0,0,0,0,0,0],"isMain":false},{"metricName":"# scaffold gap loc. mis.","quality":"Less is better","values":[0,0,0,0,0,0],"isMain":false}]],["Unaligned",[]],["Mismatches",[]],["Statistics without reference",[]],["K-mer-based statistics",[]],["Predicted genes",[]],["Similarity statistics",[]],["Reference statistics",[]]],[["Genome statistics",[]],["Reads mapping",[]],["Misassemblies",[{"metricName":"# misassemblies","quality":"Less is better","values":[30,47,25,37,64,13],"isMain":true},{"metricName":" # relocations","quality":"Less is better","values":[0,0,0,0,0,0],"isMain":false},{"metricName":" # translocations","quality":"Less is better","values":[3,2,0,1,0,1],"isMain":false},{"metricName":" # inversions","quality":"Less is better","values":[2,0,0,2,3,0],"isMain":false},{"metricName":" # interspecies translocations","quality":"Less is better","values":[25,45,25,34,61,12],"isMain":false},{"metricName":"# possibly misassembled contigs","quality":"Less is better","values":[0,0,0,2,0,1],"isMain":false},{"metricName":" # possible misassemblies","quality":"Less is better","values":[0,0,0,2,0,1],"isMain":false},{"metricName":"# local misassemblies","quality":"Less is better","values":[2,2,3,1,2,0],"isMain":false},{"metricName":"# scaffold gap ext. mis.","quality":"Less is better","values":[0,0,0,0,0,0],"isMain":false},{"metricName":"# scaffold gap loc. mis.","quality":"Less is better","values":[0,0,0,0,0,0],"isMain":false}]],["Unaligned",[]],["Mismatches",[]],["Statistics without reference",[]],["K-mer-based statistics",[]],["Predicted genes",[]],["Similarity statistics",[]],["Reference statistics",[]]],[["Genome statistics",[]],["Reads mapping",[]],["Misassemblies",[{"metricName":"# misassemblies","quality":"Less is better","values":[5,14,13,21,12,7],"isMain":true},{"metricName":" # relocations","quality":"Less is better","values":[0,1,0,0,0,1],"isMain":false},{"metricName":" # translocations","quality":"Less is better","values":[2,11,8,13,9,6],"isMain":false},{"metricName":" # inversions","quality":"Less is better","values":[0,0,0,0,0,0],"isMain":false},{"metricName":" # interspecies translocations","quality":"Less is better","values":[3,2,5,8,3,0],"isMain":false},{"metricName":"# possibly misassembled contigs","quality":"Less is better","values":[0,0,0,0,0,0],"isMain":false},{"metricName":" # possible misassemblies","quality":"Less is better","values":[0,0,0,0,0,0],"isMain":false},{"metricName":"# local misassemblies","quality":"Less is better","values":[1,2,1,2,1,1],"isMain":false},{"metricName":"# scaffold gap ext. mis.","quality":"Less is better","values":[0,0,0,0,0,0],"isMain":false},{"metricName":"# scaffold gap loc. mis.","quality":"Less is better","values":[0,0,0,0,0,0],"isMain":false}]],["Unaligned",[]],["Mismatches",[]],["Statistics without reference",[]],["K-mer-based statistics",[]],["Predicted genes",[]],["Similarity statistics",[]],["Reference statistics",[]]],[["Genome statistics",[]],["Reads mapping",[]],["Misassemblies",[{"metricName":"# misassemblies","quality":"Less is better","values":[32,38,10,26,39,16],"isMain":true},{"metricName":" # relocations","quality":"Less is better","values":[2,1,1,0,1,0],"isMain":false},{"metricName":" # translocations","quality":"Less is better","values":[3,8,2,5,8,2],"isMain":false},{"metricName":" # inversions","quality":"Less is better","values":[0,0,0,0,0,0],"isMain":false},{"metricName":" # interspecies translocations","quality":"Less is better","values":[27,29,7,21,30,14],"isMain":false},{"metricName":"# possibly misassembled contigs","quality":"Less is better","values":[0,0,0,0,0,0],"isMain":false},{"metricName":" # possible misassemblies","quality":"Less is better","values":[0,0,0,0,0,0],"isMain":false},{"metricName":"# local misassemblies","quality":"Less is better","values":[2,0,1,0,1,0],"isMain":false},{"metricName":"# scaffold gap ext. mis.","quality":"Less is better","values":[0,0,0,0,0,0],"isMain":false},{"metricName":"# scaffold gap loc. mis.","quality":"Less is better","values":[0,0,0,0,0,0],"isMain":false}]],["Unaligned",[]],["Mismatches",[]],["Statistics without reference",[]],["K-mer-based statistics",[]],["Predicted genes",[]],["Similarity statistics",[]],["Reference statistics",[]]],[["Genome statistics",[]],["Reads mapping",[]],["Misassemblies",[{"metricName":"# misassemblies","quality":"Less is better","values":[30,32,14,30,33,16],"isMain":true},{"metricName":" # relocations","quality":"Less is better","values":[1,2,0,1,2,0],"isMain":false},{"metricName":" # translocations","quality":"Less is better","values":[3,6,7,5,10,1],"isMain":false},{"metricName":" # inversions","quality":"Less is better","values":[0,0,0,0,0,0],"isMain":false},{"metricName":" # interspecies translocations","quality":"Less is better","values":[26,24,7,24,21,15],"isMain":false},{"metricName":"# possibly misassembled contigs","quality":"Less is better","values":[0,0,0,0,0,0],"isMain":false},{"metricName":" # possible misassemblies","quality":"Less is better","values":[0,0,0,0,0,0],"isMain":false},{"metricName":"# local misassemblies","quality":"Less is better","values":[2,1,1,1,1,0],"isMain":false},{"metricName":"# scaffold gap ext. mis.","quality":"Less is better","values":[0,0,0,0,0,0],"isMain":false},{"metricName":"# scaffold gap loc. mis.","quality":"Less is better","values":[0,0,0,0,0,0],"isMain":false}]],["Unaligned",[]],["Mismatches",[]],["Statistics without reference",[]],["K-mer-based statistics",[]],["Predicted genes",[]],["Similarity statistics",[]],["Reference statistics",[]]],[["Genome statistics",[]],["Reads mapping",[]],["Misassemblies",[{"metricName":"# misassemblies","quality":"Less is better","values":[0,1,2,2,2,0],"isMain":true},{"metricName":" # relocations","quality":"Less is better","values":[0,0,0,0,0,0],"isMain":false},{"metricName":" # translocations","quality":"Less is better","values":[0,0,0,0,0,0],"isMain":false},{"metricName":" # inversions","quality":"Less is better","values":[0,0,0,0,0,0],"isMain":false},{"metricName":" # interspecies translocations","quality":"Less is better","values":[0,1,2,2,2,0],"isMain":false},{"metricName":"# possibly misassembled contigs","quality":"Less is better","values":[0,0,0,0,0,0],"isMain":false},{"metricName":" # possible misassemblies","quality":"Less is better","values":[0,0,0,0,0,0],"isMain":false},{"metricName":"# local misassemblies","quality":"Less is better","values":[0,0,0,0,0,0],"isMain":false},{"metricName":"# scaffold gap ext. mis.","quality":"Less is better","values":[0,0,0,0,0,0],"isMain":false},{"metricName":"# scaffold gap loc. mis.","quality":"Less is better","values":[0,0,0,0,0,0],"isMain":false}]],["Unaligned",[]],["Mismatches",[]],["Statistics without reference",[]],["K-mer-based statistics",[]],["Predicted genes",[]],["Similarity statistics",[]],["Reference statistics",[]]],[["Genome statistics",[]],["Reads mapping",[]],["Misassemblies",[{"metricName":"# misassemblies","quality":"Less is better","values":[0,0,0,2,1,0],"isMain":true},{"metricName":" # relocations","quality":"Less is better","values":[0,0,0,0,0,0],"isMain":false},{"metricName":" # translocations","quality":"Less is better","values":[0,0,0,0,0,0],"isMain":false},{"metricName":" # inversions","quality":"Less is better","values":[0,0,0,0,0,0],"isMain":false},{"metricName":" # interspecies translocations","quality":"Less is better","values":[0,0,0,2,1,0],"isMain":false},{"metricName":"# possibly misassembled contigs","quality":"Less is better","values":[0,0,0,0,0,0],"isMain":false},{"metricName":" # possible misassemblies","quality":"Less is better","values":[0,0,0,0,0,0],"isMain":false},{"metricName":"# local misassemblies","quality":"Less is better","values":[0,0,0,0,0,0],"isMain":false},{"metricName":"# scaffold gap ext. mis.","quality":"Less is better","values":[0,0,0,0,0,0],"isMain":false},{"metricName":"# scaffold gap loc. mis.","quality":"Less is better","values":[0,0,0,0,0,0],"isMain":false}]],["Unaligned",[]],["Mismatches",[]],["Statistics without reference",[]],["K-mer-based statistics",[]],["Predicted genes",[]],["Similarity statistics",[]],["Reference statistics",[]]],[["Genome statistics",[]],["Reads mapping",[]],["Misassemblies",[{"metricName":"# misassemblies","quality":"Less is better","values":[12,20,6,24,25,14],"isMain":true},{"metricName":" # relocations","quality":"Less is better","values":[0,1,1,1,1,0],"isMain":false},{"metricName":" # translocations","quality":"Less is better","values":[0,2,2,6,3,1],"isMain":false},{"metricName":" # inversions","quality":"Less is better","values":[0,0,0,0,0,0],"isMain":false},{"metricName":" # interspecies translocations","quality":"Less is better","values":[12,17,3,17,21,13],"isMain":false},{"metricName":"# possibly misassembled contigs","quality":"Less is better","values":[0,0,0,1,0,0],"isMain":false},{"metricName":" # possible misassemblies","quality":"Less is better","values":[0,0,0,1,0,0],"isMain":false},{"metricName":"# local misassemblies","quality":"Less is better","values":[2,3,1,2,1,0],"isMain":false},{"metricName":"# scaffold gap ext. mis.","quality":"Less is better","values":[0,0,0,0,0,0],"isMain":false},{"metricName":"# scaffold gap loc. mis.","quality":"Less is better","values":[0,0,0,0,0,0],"isMain":false}]],["Unaligned",[]],["Mismatches",[]],["Statistics without reference",[]],["K-mer-based statistics",[]],["Predicted genes",[]],["Similarity statistics",[]],["Reference statistics",[]]],[["Genome statistics",[]],["Reads mapping",[]],["Misassemblies",[{"metricName":"# misassemblies","quality":"Less is better","values":[5,8,9,11,12,4],"isMain":true},{"metricName":" # relocations","quality":"Less is better","values":[0,0,0,0,1,0],"isMain":false},{"metricName":" # translocations","quality":"Less is better","values":[0,0,0,0,0,0],"isMain":false},{"metricName":" # inversions","quality":"Less is better","values":[0,0,1,0,0,0],"isMain":false},{"metricName":" # interspecies translocations","quality":"Less is better","values":[5,8,8,11,11,4],"isMain":false},{"metricName":"# possibly misassembled contigs","quality":"Less is better","values":[0,0,0,0,0,0],"isMain":false},{"metricName":" # possible misassemblies","quality":"Less is better","values":[0,0,0,0,0,0],"isMain":false},{"metricName":"# local misassemblies","quality":"Less is better","values":[1,2,1,1,2,0],"isMain":false},{"metricName":"# scaffold gap ext. mis.","quality":"Less is better","values":[0,0,0,0,0,0],"isMain":false},{"metricName":"# scaffold gap loc. mis.","quality":"Less is better","values":[0,0,0,0,0,0],"isMain":false}]],["Unaligned",[]],["Mismatches",[]],["Statistics without reference",[]],["K-mer-based statistics",[]],["Predicted genes",[]],["Similarity statistics",[]],["Reference statistics",[]]],[["Genome statistics",[]],["Reads mapping",[]],["Misassemblies",[{"metricName":"# misassemblies","quality":"Less is better","values":[1,0,1,0,0,0],"isMain":true},{"metricName":" # relocations","quality":"Less is better","values":[0,0,0,0,0,0],"isMain":false},{"metricName":" # translocations","quality":"Less is better","values":[0,0,0,0,0,0],"isMain":false},{"metricName":" # inversions","quality":"Less is better","values":[0,0,0,0,0,0],"isMain":false},{"metricName":" # interspecies translocations","quality":"Less is better","values":[1,0,1,0,0,0],"isMain":false},{"metricName":"# possibly misassembled contigs","quality":"Less is better","values":[0,0,0,0,0,0],"isMain":false},{"metricName":" # possible misassemblies","quality":"Less is better","values":[0,0,0,0,0,0],"isMain":false},{"metricName":"# local misassemblies","quality":"Less is better","values":[0,0,0,0,0,0],"isMain":false},{"metricName":"# scaffold gap ext. mis.","quality":"Less is better","values":[0,0,0,0,0,0],"isMain":false},{"metricName":"# scaffold gap loc. mis.","quality":"Less is better","values":[0,0,0,0,0,0],"isMain":false}]],["Unaligned",[]],["Mismatches",[]],["Statistics without reference",[]],["K-mer-based statistics",[]],["Predicted genes",[]],["Similarity statistics",[]],["Reference statistics",[]]],[["Genome statistics",[]],["Reads mapping",[]],["Misassemblies",[{"metricName":"# misassemblies","quality":"Less is better","values":[11,17,9,12,10,7],"isMain":true},{"metricName":" # relocations","quality":"Less is better","values":[0,0,1,0,0,1],"isMain":false},{"metricName":" # translocations","quality":"Less is better","values":[3,4,1,2,3,1],"isMain":false},{"metricName":" # inversions","quality":"Less is better","values":[0,0,0,0,0,0],"isMain":false},{"metricName":" # interspecies translocations","quality":"Less is better","values":[8,13,7,10,7,5],"isMain":false},{"metricName":"# possibly misassembled contigs","quality":"Less is better","values":[0,0,0,0,0,0],"isMain":false},{"metricName":" # possible misassemblies","quality":"Less is better","values":[0,0,0,0,0,0],"isMain":false},{"metricName":"# local misassemblies","quality":"Less is better","values":[0,0,0,0,0,0],"isMain":false},{"metricName":"# scaffold gap ext. mis.","quality":"Less is better","values":[0,0,0,0,0,0],"isMain":false},{"metricName":"# scaffold gap loc. mis.","quality":"Less is better","values":[0,0,0,0,0,0],"isMain":false}]],["Unaligned",[]],["Mismatches",[]],["Statistics without reference",[]],["K-mer-based statistics",[]],["Predicted genes",[]],["Similarity statistics",[]],["Reference statistics",[]]],[["Genome statistics",[]],["Reads mapping",[]],["Misassemblies",[{"metricName":"# misassemblies","quality":"Less is better","values":[2,1,2,6,1,2],"isMain":true},{"metricName":" # relocations","quality":"Less is better","values":[0,0,0,0,0,0],"isMain":false},{"metricName":" # translocations","quality":"Less is better","values":[0,0,0,0,0,0],"isMain":false},{"metricName":" # inversions","quality":"Less is better","values":[0,0,0,0,0,0],"isMain":false},{"metricName":" # interspecies translocations","quality":"Less is better","values":[2,1,2,6,1,2],"isMain":false},{"metricName":"# possibly misassembled contigs","quality":"Less is better","values":[0,0,0,0,0,0],"isMain":false},{"metricName":" # possible misassemblies","quality":"Less is better","values":[0,0,0,0,0,0],"isMain":false},{"metricName":"# local misassemblies","quality":"Less is better","values":[0,0,0,0,0,0],"isMain":false},{"metricName":"# scaffold gap ext. mis.","quality":"Less is better","values":[0,0,0,0,0,0],"isMain":false},{"metricName":"# scaffold gap loc. mis.","quality":"Less is better","values":[0,0,0,0,0,0],"isMain":false}]],["Unaligned",[]],["Mismatches",[]],["Statistics without reference",[]],["K-mer-based statistics",[]],["Predicted genes",[]],["Similarity statistics",[]],["Reference statistics",[]]],[["Genome statistics",[]],["Reads mapping",[]],["Misassemblies",[{"metricName":"# misassemblies","quality":"Less is better","values":[6,17,5,13,7,2],"isMain":true},{"metricName":" # relocations","quality":"Less is better","values":[0,0,0,1,0,0],"isMain":false},{"metricName":" # translocations","quality":"Less is better","values":[0,0,0,0,0,0],"isMain":false},{"metricName":" # inversions","quality":"Less is better","values":[0,0,0,0,0,0],"isMain":false},{"metricName":" # interspecies translocations","quality":"Less is better","values":[6,17,5,12,7,2],"isMain":false},{"metricName":"# possibly misassembled contigs","quality":"Less is better","values":[0,0,0,0,0,0],"isMain":false},{"metricName":" # possible misassemblies","quality":"Less is better","values":[0,0,0,0,0,0],"isMain":false},{"metricName":"# local misassemblies","quality":"Less is better","values":[0,0,0,0,0,0],"isMain":false},{"metricName":"# scaffold gap ext. mis.","quality":"Less is better","values":[0,0,0,0,0,0],"isMain":false},{"metricName":"# scaffold gap loc. mis.","quality":"Less is better","values":[0,0,0,0,0,0],"isMain":false}]],["Unaligned",[]],["Mismatches",[]],["Statistics without reference",[]],["K-mer-based statistics",[]],["Predicted genes",[]],["Similarity statistics",[]],["Reference statistics",[]]],[["Genome statistics",[]],["Reads mapping",[]],["Misassemblies",[{"metricName":"# misassemblies","quality":"Less is better","values":[50,75,20,49,70,49],"isMain":true},{"metricName":" # relocations","quality":"Less is better","values":[1,1,3,2,3,2],"isMain":false},{"metricName":" # translocations","quality":"Less is better","values":[12,19,5,11,19,2],"isMain":false},{"metricName":" # inversions","quality":"Less is better","values":[2,2,1,0,1,0],"isMain":false},{"metricName":" # interspecies translocations","quality":"Less is better","values":[35,53,11,36,47,45],"isMain":false},{"metricName":"# possibly misassembled contigs","quality":"Less is better","values":[0,0,0,0,0,0],"isMain":false},{"metricName":" # possible misassemblies","quality":"Less is better","values":[0,0,0,0,0,0],"isMain":false},{"metricName":"# local misassemblies","quality":"Less is better","values":[1,0,2,3,1,1],"isMain":false},{"metricName":"# scaffold gap ext. mis.","quality":"Less is better","values":[0,0,0,0,0,0],"isMain":false},{"metricName":"# scaffold gap loc. mis.","quality":"Less is better","values":[0,0,0,0,0,0],"isMain":false}]],["Unaligned",[]],["Mismatches",[]],["Statistics without reference",[]],["K-mer-based statistics",[]],["Predicted genes",[]],["Similarity statistics",[]],["Reference statistics",[]]],[["Genome statistics",[]],["Reads mapping",[]],["Misassemblies",[{"metricName":"# misassemblies","quality":"Less is better","values":[7,14,17,21,18,9],"isMain":true},{"metricName":" # relocations","quality":"Less is better","values":[0,0,0,0,0,0],"isMain":false},{"metricName":" # translocations","quality":"Less is better","values":[7,11,16,12,11,7],"isMain":false},{"metricName":" # inversions","quality":"Less is better","values":[0,0,0,0,0,0],"isMain":false},{"metricName":" # interspecies translocations","quality":"Less is better","values":[0,3,1,9,7,2],"isMain":false},{"metricName":"# possibly misassembled contigs","quality":"Less is better","values":[0,0,0,0,0,0],"isMain":false},{"metricName":" # possible misassemblies","quality":"Less is better","values":[0,0,0,0,0,0],"isMain":false},{"metricName":"# local misassemblies","quality":"Less is better","values":[0,1,2,1,1,0],"isMain":false},{"metricName":"# scaffold gap ext. mis.","quality":"Less is better","values":[0,0,0,0,0,0],"isMain":false},{"metricName":"# scaffold gap loc. mis.","quality":"Less is better","values":[0,0,0,0,0,0],"isMain":false}]],["Unaligned",[]],["Mismatches",[]],["Statistics without reference",[]],["K-mer-based statistics",[]],["Predicted genes",[]],["Similarity statistics",[]],["Reference statistics",[]]],[["Genome statistics",[]],["Reads mapping",[]],["Misassemblies",[{"metricName":"# misassemblies","quality":"Less is better","values":[10,17,8,35,14,24],"isMain":true},{"metricName":" # relocations","quality":"Less is better","values":[0,0,0,0,0,0],"isMain":false},{"metricName":" # translocations","quality":"Less is better","values":[0,1,1,1,0,0],"isMain":false},{"metricName":" # inversions","quality":"Less is better","values":[1,1,0,0,0,0],"isMain":false},{"metricName":" # interspecies translocations","quality":"Less is better","values":[9,15,7,34,14,24],"isMain":false},{"metricName":"# possibly misassembled contigs","quality":"Less is better","values":[0,0,0,0,0,0],"isMain":false},{"metricName":" # possible misassemblies","quality":"Less is better","values":[0,0,0,0,0,0],"isMain":false},{"metricName":"# local misassemblies","quality":"Less is better","values":[0,0,0,1,0,0],"isMain":false},{"metricName":"# scaffold gap ext. mis.","quality":"Less is better","values":[0,0,0,0,0,0],"isMain":false},{"metricName":"# scaffold gap loc. mis.","quality":"Less is better","values":[0,0,0,0,0,0],"isMain":false}]],["Unaligned",[]],["Mismatches",[]],["Statistics without reference",[]],["K-mer-based statistics",[]],["Predicted genes",[]],["Similarity statistics",[]],["Reference statistics",[]]],[["Genome statistics",[]],["Reads mapping",[]],["Misassemblies",[{"metricName":"# misassemblies","quality":"Less is better","values":[26,26,6,24,16,10],"isMain":true},{"metricName":" # relocations","quality":"Less is better","values":[1,1,0,1,1,0],"isMain":false},{"metricName":" # translocations","quality":"Less is better","values":[3,7,3,4,4,0],"isMain":false},{"metricName":" # inversions","quality":"Less is better","values":[1,0,0,1,0,0],"isMain":false},{"metricName":" # interspecies translocations","quality":"Less is better","values":[21,18,3,18,11,10],"isMain":false},{"metricName":"# possibly misassembled contigs","quality":"Less is better","values":[0,0,0,0,0,0],"isMain":false},{"metricName":" # possible misassemblies","quality":"Less is better","values":[0,0,0,0,0,0],"isMain":false},{"metricName":"# local misassemblies","quality":"Less is better","values":[2,2,1,2,4,0],"isMain":false},{"metricName":"# scaffold gap ext. mis.","quality":"Less is better","values":[0,0,0,0,0,0],"isMain":false},{"metricName":"# scaffold gap loc. mis.","quality":"Less is better","values":[0,0,0,0,0,0],"isMain":false}]],["Unaligned",[]],["Mismatches",[]],["Statistics without reference",[]],["K-mer-based statistics",[]],["Predicted genes",[]],["Similarity statistics",[]],["Reference statistics",[]]],[["Genome statistics",[]],["Reads mapping",[]],["Misassemblies",[{"metricName":"# misassemblies","quality":"Less is better","values":[0,1,2,1,0,0],"isMain":true},{"metricName":" # relocations","quality":"Less is better","values":[0,0,0,0,0,0],"isMain":false},{"metricName":" # translocations","quality":"Less is better","values":[0,0,0,0,0,0],"isMain":false},{"metricName":" # inversions","quality":"Less is better","values":[0,0,0,0,0,0],"isMain":false},{"metricName":" # interspecies translocations","quality":"Less is better","values":[0,1,2,1,0,0],"isMain":false},{"metricName":"# possibly misassembled contigs","quality":"Less is better","values":[0,0,0,0,0,0],"isMain":false},{"metricName":" # possible misassemblies","quality":"Less is better","values":[0,0,0,0,0,0],"isMain":false},{"metricName":"# local misassemblies","quality":"Less is better","values":[0,0,0,0,0,0],"isMain":false},{"metricName":"# scaffold gap ext. mis.","quality":"Less is better","values":[0,0,0,0,0,0],"isMain":false},{"metricName":"# scaffold gap loc. mis.","quality":"Less is better","values":[0,0,0,0,0,0],"isMain":false}]],["Unaligned",[]],["Mismatches",[]],["Statistics without reference",[]],["K-mer-based statistics",[]],["Predicted genes",[]],["Similarity statistics",[]],["Reference statistics",[]]],[["Genome statistics",[]],["Reads mapping",[]],["Misassemblies",[{"metricName":"# misassemblies","quality":"Less is better","values":[0,0,3,3,1,1],"isMain":true},{"metricName":" # relocations","quality":"Less is better","values":[0,0,0,0,0,0],"isMain":false},{"metricName":" # translocations","quality":"Less is better","values":[0,0,0,0,0,0],"isMain":false},{"metricName":" # inversions","quality":"Less is better","values":[0,0,0,0,0,0],"isMain":false},{"metricName":" # interspecies translocations","quality":"Less is better","values":[0,0,3,3,1,1],"isMain":false},{"metricName":"# possibly misassembled contigs","quality":"Less is better","values":[0,0,0,0,0,0],"isMain":false},{"metricName":" # possible misassemblies","quality":"Less is better","values":[0,0,0,0,0,0],"isMain":false},{"metricName":"# local misassemblies","quality":"Less is better","values":[0,0,0,0,0,0],"isMain":false},{"metricName":"# scaffold gap ext. mis.","quality":"Less is better","values":[0,0,0,0,0,0],"isMain":false},{"metricName":"# scaffold gap loc. mis.","quality":"Less is better","values":[0,0,0,0,0,0],"isMain":false}]],["Unaligned",[]],["Mismatches",[]],["Statistics without reference",[]],["K-mer-based statistics",[]],["Predicted genes",[]],["Similarity statistics",[]],["Reference statistics",[]]],[["Genome statistics",[]],["Reads mapping",[]],["Misassemblies",[{"metricName":"# misassemblies","quality":"Less is better","values":[59,35,44,68,40,20],"isMain":true},{"metricName":" # relocations","quality":"Less is better","values":[0,0,0,1,0,0],"isMain":false},{"metricName":" # translocations","quality":"Less is better","values":[10,20,21,29,22,18],"isMain":false},{"metricName":" # inversions","quality":"Less is better","values":[2,0,0,0,1,0],"isMain":false},{"metricName":" # interspecies translocations","quality":"Less is better","values":[47,15,23,38,17,2],"isMain":false},{"metricName":"# possibly misassembled contigs","quality":"Less is better","values":[0,0,0,0,0,0],"isMain":false},{"metricName":" # possible misassemblies","quality":"Less is better","values":[0,0,0,0,0,0],"isMain":false},{"metricName":"# local misassemblies","quality":"Less is better","values":[1,1,1,4,1,0],"isMain":false},{"metricName":"# scaffold gap ext. mis.","quality":"Less is better","values":[0,0,0,0,0,0],"isMain":false},{"metricName":"# scaffold gap loc. mis.","quality":"Less is better","values":[0,0,0,0,0,0],"isMain":false}]],["Unaligned",[]],["Mismatches",[]],["Statistics without reference",[]],["K-mer-based statistics",[]],["Predicted genes",[]],["Similarity statistics",[]],["Reference statistics",[]]],[["Genome statistics",[]],["Reads mapping",[]],["Misassemblies",[{"metricName":"# misassemblies","quality":"Less is better","values":[7,7,5,3,9,6],"isMain":true},{"metricName":" # relocations","quality":"Less is better","values":[0,0,0,0,0,0],"isMain":false},{"metricName":" # translocations","quality":"Less is better","values":[0,0,0,0,0,0],"isMain":false},{"metricName":" # inversions","quality":"Less is better","values":[0,0,0,0,0,0],"isMain":false},{"metricName":" # interspecies translocations","quality":"Less is better","values":[7,7,5,3,9,6],"isMain":false},{"metricName":"# possibly misassembled contigs","quality":"Less is better","values":[0,0,0,0,0,0],"isMain":false},{"metricName":" # possible misassemblies","quality":"Less is better","values":[0,0,0,0,0,0],"isMain":false},{"metricName":"# local misassemblies","quality":"Less is better","values":[0,0,0,0,0,0],"isMain":false},{"metricName":"# scaffold gap ext. mis.","quality":"Less is better","values":[0,0,0,0,0,0],"isMain":false},{"metricName":"# scaffold gap loc. mis.","quality":"Less is better","values":[0,0,0,0,0,0],"isMain":false}]],["Unaligned",[]],["Mismatches",[]],["Statistics without reference",[]],["K-mer-based statistics",[]],["Predicted genes",[]],["Similarity statistics",[]],["Reference statistics",[]]],[["Genome statistics",[]],["Reads mapping",[]],["Misassemblies",[{"metricName":"# misassemblies","quality":"Less is better","values":[10,13,3,17,8,7],"isMain":true},{"metricName":" # relocations","quality":"Less is better","values":[0,0,0,0,0,0],"isMain":false},{"metricName":" # translocations","quality":"Less is better","values":[0,0,0,1,0,0],"isMain":false},{"metricName":" # inversions","quality":"Less is better","values":[0,0,0,0,0,0],"isMain":false},{"metricName":" # interspecies translocations","quality":"Less is better","values":[10,13,3,16,8,7],"isMain":false},{"metricName":"# possibly misassembled contigs","quality":"Less is better","values":[0,0,0,0,0,0],"isMain":false},{"metricName":" # possible misassemblies","quality":"Less is better","values":[0,0,0,0,0,0],"isMain":false},{"metricName":"# local misassemblies","quality":"Less is better","values":[0,0,0,1,0,0],"isMain":false},{"metricName":"# scaffold gap ext. mis.","quality":"Less is better","values":[0,0,0,0,0,0],"isMain":false},{"metricName":"# scaffold gap loc. mis.","quality":"Less is better","values":[0,0,0,0,0,0],"isMain":false}]],["Unaligned",[]],["Mismatches",[]],["Statistics without reference",[]],["K-mer-based statistics",[]],["Predicted genes",[]],["Similarity statistics",[]],["Reference statistics",[]]],[["Genome statistics",[]],["Reads mapping",[]],["Misassemblies",[{"metricName":"# misassemblies","quality":"Less is better","values":[6,17,8,19,20,12],"isMain":true},{"metricName":" # relocations","quality":"Less is better","values":[0,0,0,0,0,0],"isMain":false},{"metricName":" # translocations","quality":"Less is better","values":[1,1,1,2,1,1],"isMain":false},{"metricName":" # inversions","quality":"Less is better","values":[0,0,0,0,0,0],"isMain":false},{"metricName":" # interspecies translocations","quality":"Less is better","values":[5,16,7,17,19,11],"isMain":false},{"metricName":"# possibly misassembled contigs","quality":"Less is better","values":[0,0,0,0,0,0],"isMain":false},{"metricName":" # possible misassemblies","quality":"Less is better","values":[0,0,0,0,0,0],"isMain":false},{"metricName":"# local misassemblies","quality":"Less is better","values":[0,0,0,0,0,0],"isMain":false},{"metricName":"# scaffold gap ext. mis.","quality":"Less is better","values":[0,0,0,0,0,0],"isMain":false},{"metricName":"# scaffold gap loc. mis.","quality":"Less is better","values":[0,0,0,0,0,0],"isMain":false}]],["Unaligned",[]],["Mismatches",[]],["Statistics without reference",[]],["K-mer-based statistics",[]],["Predicted genes",[]],["Similarity statistics",[]],["Reference statistics",[]]],[["Genome statistics",[]],["Reads mapping",[]],["Misassemblies",[{"metricName":"# misassemblies","quality":"Less is better","values":[12,17,3,17,14,10],"isMain":true},{"metricName":" # relocations","quality":"Less is better","values":[0,0,0,1,0,0],"isMain":false},{"metricName":" # translocations","quality":"Less is better","values":[0,0,0,1,0,0],"isMain":false},{"metricName":" # inversions","quality":"Less is better","values":[0,0,0,0,0,0],"isMain":false},{"metricName":" # interspecies translocations","quality":"Less is better","values":[12,17,3,15,14,10],"isMain":false},{"metricName":"# possibly misassembled contigs","quality":"Less is better","values":[0,0,0,0,0,0],"isMain":false},{"metricName":" # possible misassemblies","quality":"Less is better","values":[0,0,0,0,0,0],"isMain":false},{"metricName":"# local misassemblies","quality":"Less is better","values":[1,0,0,1,0,0],"isMain":false},{"metricName":"# scaffold gap ext. mis.","quality":"Less is better","values":[0,0,0,0,0,0],"isMain":false},{"metricName":"# scaffold gap loc. mis.","quality":"Less is better","values":[0,0,0,0,0,0],"isMain":false}]],["Unaligned",[]],["Mismatches",[]],["Statistics without reference",[]],["K-mer-based statistics",[]],["Predicted genes",[]],["Similarity statistics",[]],["Reference statistics",[]]],[["Genome statistics",[]],["Reads mapping",[]],["Misassemblies",[{"metricName":"# misassemblies","quality":"Less is better","values":[4,13,5,20,6,3],"isMain":true},{"metricName":" # relocations","quality":"Less is better","values":[0,0,0,0,0,0],"isMain":false},{"metricName":" # translocations","quality":"Less is better","values":[0,0,0,0,0,0],"isMain":false},{"metricName":" # inversions","quality":"Less is better","values":[0,0,0,0,0,0],"isMain":false},{"metricName":" # interspecies translocations","quality":"Less is better","values":[4,13,5,20,6,3],"isMain":false},{"metricName":"# possibly misassembled contigs","quality":"Less is better","values":[0,0,0,0,0,0],"isMain":false},{"metricName":" # possible misassemblies","quality":"Less is better","values":[0,0,0,0,0,0],"isMain":false},{"metricName":"# local misassemblies","quality":"Less is better","values":[0,0,0,0,0,0],"isMain":false},{"metricName":"# scaffold gap ext. mis.","quality":"Less is better","values":[0,0,0,0,0,0],"isMain":false},{"metricName":"# scaffold gap loc. mis.","quality":"Less is better","values":[0,0,0,0,0,0],"isMain":false}]],["Unaligned",[]],["Mismatches",[]],["Statistics without reference",[]],["K-mer-based statistics",[]],["Predicted genes",[]],["Similarity statistics",[]],["Reference statistics",[]]],[["Genome statistics",[]],["Reads mapping",[]],["Misassemblies",[{"metricName":"# misassemblies","quality":"Less is better","values":[37,55,29,40,46,21],"isMain":true},{"metricName":" # relocations","quality":"Less is better","values":[0,0,0,0,0,0],"isMain":false},{"metricName":" # translocations","quality":"Less is better","values":[1,0,1,2,0,0],"isMain":false},{"metricName":" # inversions","quality":"Less is better","values":[0,0,0,0,0,0],"isMain":false},{"metricName":" # interspecies translocations","quality":"Less is better","values":[36,55,28,38,46,21],"isMain":false},{"metricName":"# possibly misassembled contigs","quality":"Less is better","values":[0,0,0,2,0,0],"isMain":false},{"metricName":" # possible misassemblies","quality":"Less is better","values":[0,0,0,2,0,0],"isMain":false},{"metricName":"# local misassemblies","quality":"Less is better","values":[0,0,0,0,0,0],"isMain":false},{"metricName":"# scaffold gap ext. mis.","quality":"Less is better","values":[0,0,0,0,0,0],"isMain":false},{"metricName":"# scaffold gap loc. mis.","quality":"Less is better","values":[0,0,0,0,0,0],"isMain":false}]],["Unaligned",[]],["Mismatches",[]],["Statistics without reference",[]],["K-mer-based statistics",[]],["Predicted genes",[]],["Similarity statistics",[]],["Reference statistics",[]]],[["Genome statistics",[]],["Reads mapping",[]],["Misassemblies",[{"metricName":"# misassemblies","quality":"Less is better","values":[18,14,8,14,15,6],"isMain":true},{"metricName":" # relocations","quality":"Less is better","values":[0,1,1,0,1,0],"isMain":false},{"metricName":" # translocations","quality":"Less is better","values":[0,0,0,0,0,0],"isMain":false},{"metricName":" # inversions","quality":"Less is better","values":[0,0,0,0,0,0],"isMain":false},{"metricName":" # interspecies translocations","quality":"Less is better","values":[18,13,7,14,14,6],"isMain":false},{"metricName":"# possibly misassembled contigs","quality":"Less is better","values":[0,0,0,0,0,0],"isMain":false},{"metricName":" # possible misassemblies","quality":"Less is better","values":[0,0,0,0,0,0],"isMain":false},{"metricName":"# local misassemblies","quality":"Less is better","values":[0,0,2,0,0,1],"isMain":false},{"metricName":"# scaffold gap ext. mis.","quality":"Less is better","values":[0,0,0,0,0,0],"isMain":false},{"metricName":"# scaffold gap loc. mis.","quality":"Less is better","values":[0,0,0,0,0,0],"isMain":false}]],["Unaligned",[]],["Mismatches",[]],["Statistics without reference",[]],["K-mer-based statistics",[]],["Predicted genes",[]],["Similarity statistics",[]],["Reference statistics",[]]],[["Genome statistics",[]],["Reads mapping",[]],["Misassemblies",[{"metricName":"# misassemblies","quality":"Less is better","values":[11,10,14,8,15,7],"isMain":true},{"metricName":" # relocations","quality":"Less is better","values":[3,2,4,1,4,1],"isMain":false},{"metricName":" # translocations","quality":"Less is better","values":[1,0,0,0,0,0],"isMain":false},{"metricName":" # inversions","quality":"Less is better","values":[1,2,0,1,0,0],"isMain":false},{"metricName":" # interspecies translocations","quality":"Less is better","values":[6,6,10,6,11,6],"isMain":false},{"metricName":"# possibly misassembled contigs","quality":"Less is better","values":[0,0,0,0,0,0],"isMain":false},{"metricName":" # possible misassemblies","quality":"Less is better","values":[0,0,0,0,0,0],"isMain":false},{"metricName":"# local misassemblies","quality":"Less is better","values":[0,0,0,0,0,0],"isMain":false},{"metricName":"# scaffold gap ext. mis.","quality":"Less is better","values":[0,0,0,0,0,0],"isMain":false},{"metricName":"# scaffold gap loc. mis.","quality":"Less is better","values":[0,0,0,0,0,0],"isMain":false}]],["Unaligned",[]],["Mismatches",[]],["Statistics without reference",[]],["K-mer-based statistics",[]],["Predicted genes",[]],["Similarity statistics",[]],["Reference statistics",[]]],[["Genome statistics",[]],["Reads mapping",[]],["Misassemblies",[{"metricName":"# misassemblies","quality":"Less is better","values":[7,21,4,9,18,16],"isMain":true},{"metricName":" # relocations","quality":"Less is better","values":[0,0,0,0,0,0],"isMain":false},{"metricName":" # translocations","quality":"Less is better","values":[0,0,1,0,0,0],"isMain":false},{"metricName":" # inversions","quality":"Less is better","values":[0,1,0,0,0,0],"isMain":false},{"metricName":" # interspecies translocations","quality":"Less is better","values":[7,20,3,9,18,16],"isMain":false},{"metricName":"# possibly misassembled contigs","quality":"Less is better","values":[0,0,0,0,1,0],"isMain":false},{"metricName":" # possible misassemblies","quality":"Less is better","values":[0,0,0,0,1,0],"isMain":false},{"metricName":"# local misassemblies","quality":"Less is better","values":[0,0,1,0,0,0],"isMain":false},{"metricName":"# scaffold gap ext. mis.","quality":"Less is better","values":[0,0,0,0,0,0],"isMain":false},{"metricName":"# scaffold gap loc. mis.","quality":"Less is better","values":[0,0,0,0,0,0],"isMain":false}]],["Unaligned",[]],["Mismatches",[]],["Statistics without reference",[]],["K-mer-based statistics",[]],["Predicted genes",[]],["Similarity statistics",[]],["Reference statistics",[]]],[["Genome statistics",[]],["Reads mapping",[]],["Misassemblies",[{"metricName":"# misassemblies","quality":"Less is better","values":[1,0,0,0,0,0],"isMain":true},{"metricName":" # relocations","quality":"Less is better","values":[0,0,0,0,0,0],"isMain":false},{"metricName":" # translocations","quality":"Less is better","values":[0,0,0,0,0,0],"isMain":false},{"metricName":" # inversions","quality":"Less is better","values":[0,0,0,0,0,0],"isMain":false},{"metricName":" # interspecies translocations","quality":"Less is better","values":[1,0,0,0,0,0],"isMain":false},{"metricName":"# possibly misassembled contigs","quality":"Less is better","values":[0,0,0,0,0,0],"isMain":false},{"metricName":" # possible misassemblies","quality":"Less is better","values":[0,0,0,0,0,0],"isMain":false},{"metricName":"# local misassemblies","quality":"Less is better","values":[0,0,0,0,0,0],"isMain":false},{"metricName":"# scaffold gap ext. mis.","quality":"Less is better","values":[0,0,0,0,0,0],"isMain":false},{"metricName":"# scaffold gap loc. mis.","quality":"Less is better","values":[0,0,0,0,0,0],"isMain":false}]],["Unaligned",[]],["Mismatches",[]],["Statistics without reference",[]],["K-mer-based statistics",[]],["Predicted genes",[]],["Similarity statistics",[]],["Reference statistics",[]]],[["Genome statistics",[]],["Reads mapping",[]],["Misassemblies",[{"metricName":"# misassemblies","quality":"Less is better","values":[5,10,9,6,12,5],"isMain":true},{"metricName":" # relocations","quality":"Less is better","values":[0,0,1,0,0,0],"isMain":false},{"metricName":" # translocations","quality":"Less is better","values":[3,4,3,3,7,5],"isMain":false},{"metricName":" # inversions","quality":"Less is better","values":[0,0,0,0,0,0],"isMain":false},{"metricName":" # interspecies translocations","quality":"Less is better","values":[2,6,5,3,5,0],"isMain":false},{"metricName":"# possibly misassembled contigs","quality":"Less is better","values":[0,0,0,0,0,0],"isMain":false},{"metricName":" # possible misassemblies","quality":"Less is better","values":[0,0,0,0,0,0],"isMain":false},{"metricName":"# local misassemblies","quality":"Less is better","values":[0,0,0,0,1,0],"isMain":false},{"metricName":"# scaffold gap ext. mis.","quality":"Less is better","values":[0,0,0,0,0,0],"isMain":false},{"metricName":"# scaffold gap loc. mis.","quality":"Less is better","values":[0,0,0,0,0,0],"isMain":false}]],["Unaligned",[]],["Mismatches",[]],["Statistics without reference",[]],["K-mer-based statistics",[]],["Predicted genes",[]],["Similarity statistics",[]],["Reference statistics",[]]],[["Genome statistics",[]],["Reads mapping",[]],["Misassemblies",[{"metricName":"# misassemblies","quality":"Less is better","values":[2,8,5,12,12,9],"isMain":true},{"metricName":" # relocations","quality":"Less is better","values":[0,0,0,0,0,0],"isMain":false},{"metricName":" # translocations","quality":"Less is better","values":[2,3,0,4,5,5],"isMain":false},{"metricName":" # inversions","quality":"Less is better","values":[0,0,0,0,0,0],"isMain":false},{"metricName":" # interspecies translocations","quality":"Less is better","values":[0,5,5,8,7,4],"isMain":false},{"metricName":"# possibly misassembled contigs","quality":"Less is better","values":[0,0,0,0,0,0],"isMain":false},{"metricName":" # possible misassemblies","quality":"Less is better","values":[0,0,0,0,0,0],"isMain":false},{"metricName":"# local misassemblies","quality":"Less is better","values":[0,1,2,0,0,0],"isMain":false},{"metricName":"# scaffold gap ext. mis.","quality":"Less is better","values":[0,0,0,0,0,0],"isMain":false},{"metricName":"# scaffold gap loc. mis.","quality":"Less is better","values":[0,0,0,0,0,0],"isMain":false}]],["Unaligned",[]],["Mismatches",[]],["Statistics without reference",[]],["K-mer-based statistics",[]],["Predicted genes",[]],["Similarity statistics",[]],["Reference statistics",[]]],[["Genome statistics",[]],["Reads mapping",[]],["Misassemblies",[{"metricName":"# misassemblies","quality":"Less is better","values":[19,24,8,53,21,12],"isMain":true},{"metricName":" # relocations","quality":"Less is better","values":[0,2,2,5,2,1],"isMain":false},{"metricName":" # translocations","quality":"Less is better","values":[0,0,0,0,0,0],"isMain":false},{"metricName":" # inversions","quality":"Less is better","values":[0,0,0,2,1,0],"isMain":false},{"metricName":" # interspecies translocations","quality":"Less is better","values":[19,22,6,46,18,11],"isMain":false},{"metricName":"# possibly misassembled contigs","quality":"Less is better","values":[0,0,1,1,0,0],"isMain":false},{"metricName":" # possible misassemblies","quality":"Less is better","values":[0,0,1,1,0,0],"isMain":false},{"metricName":"# local misassemblies","quality":"Less is better","values":[1,2,4,6,5,1],"isMain":false},{"metricName":"# scaffold gap ext. mis.","quality":"Less is better","values":[0,0,0,0,0,0],"isMain":false},{"metricName":"# scaffold gap loc. mis.","quality":"Less is better","values":[0,0,0,0,0,0],"isMain":false}]],["Unaligned",[]],["Mismatches",[]],["Statistics without reference",[]],["K-mer-based statistics",[]],["Predicted genes",[]],["Similarity statistics",[]],["Reference statistics",[]]],[["Genome statistics",[]],["Reads mapping",[]],["Misassemblies",[{"metricName":"# misassemblies","quality":"Less is better","values":[7,12,6,14,14,6],"isMain":true},{"metricName":" # relocations","quality":"Less is better","values":[0,0,0,0,0,0],"isMain":false},{"metricName":" # translocations","quality":"Less is better","values":[0,0,0,0,0,0],"isMain":false},{"metricName":" # inversions","quality":"Less is better","values":[0,0,1,0,0,0],"isMain":false},{"metricName":" # interspecies translocations","quality":"Less is better","values":[7,12,5,14,14,6],"isMain":false},{"metricName":"# possibly misassembled contigs","quality":"Less is better","values":[0,0,0,0,0,0],"isMain":false},{"metricName":" # possible misassemblies","quality":"Less is better","values":[0,0,0,0,0,0],"isMain":false},{"metricName":"# local misassemblies","quality":"Less is better","values":[0,2,0,0,0,0],"isMain":false},{"metricName":"# scaffold gap ext. mis.","quality":"Less is better","values":[0,0,0,0,0,0],"isMain":false},{"metricName":"# scaffold gap loc. mis.","quality":"Less is better","values":[0,0,0,0,0,0],"isMain":false}]],["Unaligned",[]],["Mismatches",[]],["Statistics without reference",[]],["K-mer-based statistics",[]],["Predicted genes",[]],["Similarity statistics",[]],["Reference statistics",[]]],[["Genome statistics",[]],["Reads mapping",[]],["Misassemblies",[{"metricName":"# misassemblies","quality":"Less is better","values":[6,8,6,14,10,2],"isMain":true},{"metricName":" # relocations","quality":"Less is better","values":[0,0,0,0,0,0],"isMain":false},{"metricName":" # translocations","quality":"Less is better","values":[0,0,0,0,0,0],"isMain":false},{"metricName":" # inversions","quality":"Less is better","values":[1,0,0,0,0,0],"isMain":false},{"metricName":" # interspecies translocations","quality":"Less is better","values":[5,8,6,14,10,2],"isMain":false},{"metricName":"# possibly misassembled contigs","quality":"Less is better","values":[0,0,0,0,0,0],"isMain":false},{"metricName":" # possible misassemblies","quality":"Less is better","values":[0,0,0,0,0,0],"isMain":false},{"metricName":"# local misassemblies","quality":"Less is better","values":[0,0,0,0,0,0],"isMain":false},{"metricName":"# scaffold gap ext. mis.","quality":"Less is better","values":[0,0,0,0,0,0],"isMain":false},{"metricName":"# scaffold gap loc. mis.","quality":"Less is better","values":[0,0,0,0,0,0],"isMain":false}]],["Unaligned",[]],["Mismatches",[]],["Statistics without reference",[]],["K-mer-based statistics",[]],["Predicted genes",[]],["Similarity statistics",[]],["Reference statistics",[]]],[["Genome statistics",[]],["Reads mapping",[]],["Misassemblies",[{"metricName":"# misassemblies","quality":"Less is better","values":[1,3,2,3,3,2],"isMain":true},{"metricName":" # relocations","quality":"Less is better","values":[0,0,0,0,0,0],"isMain":false},{"metricName":" # translocations","quality":"Less is better","values":[0,0,0,0,0,0],"isMain":false},{"metricName":" # inversions","quality":"Less is better","values":[0,0,0,0,0,0],"isMain":false},{"metricName":" # interspecies translocations","quality":"Less is better","values":[1,3,2,3,3,2],"isMain":false},{"metricName":"# possibly misassembled contigs","quality":"Less is better","values":[0,0,0,0,0,0],"isMain":false},{"metricName":" # possible misassemblies","quality":"Less is better","values":[0,0,0,0,0,0],"isMain":false},{"metricName":"# local misassemblies","quality":"Less is better","values":[0,0,0,0,0,0],"isMain":false},{"metricName":"# scaffold gap ext. mis.","quality":"Less is better","values":[0,0,0,0,0,0],"isMain":false},{"metricName":"# scaffold gap loc. mis.","quality":"Less is better","values":[0,0,0,0,0,0],"isMain":false}]],["Unaligned",[]],["Mismatches",[]],["Statistics without reference",[]],["K-mer-based statistics",[]],["Predicted genes",[]],["Similarity statistics",[]],["Reference statistics",[]]],[["Genome statistics",[]],["Reads mapping",[]],["Misassemblies",[{"metricName":"# misassemblies","quality":"Less is better","values":[26,14,13,17,16,1],"isMain":true},{"metricName":" # relocations","quality":"Less is better","values":[0,0,0,0,0,0],"isMain":false},{"metricName":" # translocations","quality":"Less is better","values":[0,1,1,0,1,0],"isMain":false},{"metricName":" # inversions","quality":"Less is better","values":[0,0,0,0,0,0],"isMain":false},{"metricName":" # interspecies translocations","quality":"Less is better","values":[26,13,12,17,15,1],"isMain":false},{"metricName":"# possibly misassembled contigs","quality":"Less is better","values":[0,0,0,1,0,0],"isMain":false},{"metricName":" # possible misassemblies","quality":"Less is better","values":[0,0,0,1,0,0],"isMain":false},{"metricName":"# local misassemblies","quality":"Less is better","values":[2,0,1,1,0,0],"isMain":false},{"metricName":"# scaffold gap ext. mis.","quality":"Less is better","values":[0,0,0,0,0,0],"isMain":false},{"metricName":"# scaffold gap loc. mis.","quality":"Less is better","values":[0,0,0,0,0,0],"isMain":false}]],["Unaligned",[]],["Mismatches",[]],["Statistics without reference",[]],["K-mer-based statistics",[]],["Predicted genes",[]],["Similarity statistics",[]],["Reference statistics",[]]],[["Genome statistics",[]],["Reads mapping",[]],["Misassemblies",[{"metricName":"# misassemblies","quality":"Less is better","values":[24,50,10,22,15,12],"isMain":true},{"metricName":" # relocations","quality":"Less is better","values":[0,0,0,0,0,1],"isMain":false},{"metricName":" # translocations","quality":"Less is better","values":[1,1,1,2,0,2],"isMain":false},{"metricName":" # inversions","quality":"Less is better","values":[0,0,0,2,0,0],"isMain":false},{"metricName":" # interspecies translocations","quality":"Less is better","values":[23,49,9,18,15,9],"isMain":false},{"metricName":"# possibly misassembled contigs","quality":"Less is better","values":[0,0,0,2,0,0],"isMain":false},{"metricName":" # possible misassemblies","quality":"Less is better","values":[0,0,0,2,0,0],"isMain":false},{"metricName":"# local misassemblies","quality":"Less is better","values":[1,0,1,0,0,1],"isMain":false},{"metricName":"# scaffold gap ext. mis.","quality":"Less is better","values":[0,0,0,0,0,0],"isMain":false},{"metricName":"# scaffold gap loc. mis.","quality":"Less is better","values":[0,0,0,0,0,0],"isMain":false}]],["Unaligned",[]],["Mismatches",[]],["Statistics without reference",[]],["K-mer-based statistics",[]],["Predicted genes",[]],["Similarity statistics",[]],["Reference statistics",[]]],[["Genome statistics",[]],["Reads mapping",[]],["Misassemblies",[{"metricName":"# misassemblies","quality":"Less is better","values":[12,19,11,17,19,11],"isMain":true},{"metricName":" # relocations","quality":"Less is better","values":[1,3,0,0,3,0],"isMain":false},{"metricName":" # translocations","quality":"Less is better","values":[0,0,0,0,0,0],"isMain":false},{"metricName":" # inversions","quality":"Less is better","values":[0,0,0,0,0,0],"isMain":false},{"metricName":" # interspecies translocations","quality":"Less is better","values":[11,16,11,17,16,11],"isMain":false},{"metricName":"# possibly misassembled contigs","quality":"Less is better","values":[0,0,0,0,0,0],"isMain":false},{"metricName":" # possible misassemblies","quality":"Less is better","values":[0,0,0,0,0,0],"isMain":false},{"metricName":"# local misassemblies","quality":"Less is better","values":[0,1,0,0,1,0],"isMain":false},{"metricName":"# scaffold gap ext. mis.","quality":"Less is better","values":[0,0,0,0,0,0],"isMain":false},{"metricName":"# scaffold gap loc. mis.","quality":"Less is better","values":[0,0,0,0,0,0],"isMain":false}]],["Unaligned",[]],["Mismatches",[]],["Statistics without reference",[]],["K-mer-based statistics",[]],["Predicted genes",[]],["Similarity statistics",[]],["Reference statistics",[]]],[["Genome statistics",[]],["Reads mapping",[]],["Misassemblies",[{"metricName":"# misassemblies","quality":"Less is better","values":[3,4,5,35,7,1],"isMain":true},{"metricName":" # relocations","quality":"Less is better","values":[0,0,0,0,0,0],"isMain":false},{"metricName":" # translocations","quality":"Less is better","values":[0,1,1,0,1,0],"isMain":false},{"metricName":" # inversions","quality":"Less is better","values":[0,0,1,1,1,0],"isMain":false},{"metricName":" # interspecies translocations","quality":"Less is better","values":[3,3,3,34,5,1],"isMain":false},{"metricName":"# possibly misassembled contigs","quality":"Less is better","values":[0,0,0,0,0,0],"isMain":false},{"metricName":" # possible misassemblies","quality":"Less is better","values":[0,0,0,0,0,0],"isMain":false},{"metricName":"# local misassemblies","quality":"Less is better","values":[2,0,3,2,2,0],"isMain":false},{"metricName":"# scaffold gap ext. mis.","quality":"Less is better","values":[0,0,0,0,0,0],"isMain":false},{"metricName":"# scaffold gap loc. mis.","quality":"Less is better","values":[0,0,0,0,0,0],"isMain":false}]],["Unaligned",[]],["Mismatches",[]],["Statistics without reference",[]],["K-mer-based statistics",[]],["Predicted genes",[]],["Similarity statistics",[]],["Reference statistics",[]]],[["Genome statistics",[]],["Reads mapping",[]],["Misassemblies",[{"metricName":"# misassemblies","quality":"Less is better","values":[26,46,64,124,87,11],"isMain":true},{"metricName":" # relocations","quality":"Less is better","values":[0,0,1,2,3,0],"isMain":false},{"metricName":" # translocations","quality":"Less is better","values":[4,8,13,18,20,5],"isMain":false},{"metricName":" # inversions","quality":"Less is better","values":[1,1,0,0,1,0],"isMain":false},{"metricName":" # interspecies translocations","quality":"Less is better","values":[21,37,50,104,63,6],"isMain":false},{"metricName":"# possibly misassembled contigs","quality":"Less is better","values":[1,1,0,2,1,0],"isMain":false},{"metricName":" # possible misassemblies","quality":"Less is better","values":[1,1,0,2,1,0],"isMain":false},{"metricName":"# local misassemblies","quality":"Less is better","values":[2,7,7,6,7,0],"isMain":false},{"metricName":"# scaffold gap ext. mis.","quality":"Less is better","values":[0,0,0,0,0,0],"isMain":false},{"metricName":"# scaffold gap loc. mis.","quality":"Less is better","values":[0,0,0,0,0,0],"isMain":false}]],["Unaligned",[]],["Mismatches",[]],["Statistics without reference",[]],["K-mer-based statistics",[]],["Predicted genes",[]],["Similarity statistics",[]],["Reference statistics",[]]],[["Genome statistics",[]],["Reads mapping",[]],["Misassemblies",[{"metricName":"# misassemblies","quality":"Less is better","values":[5,9,5,13,10,4],"isMain":true},{"metricName":" # relocations","quality":"Less is better","values":[0,0,0,0,0,0],"isMain":false},{"metricName":" # translocations","quality":"Less is better","values":[0,0,0,0,0,0],"isMain":false},{"metricName":" # inversions","quality":"Less is better","values":[0,0,0,0,0,0],"isMain":false},{"metricName":" # interspecies translocations","quality":"Less is better","values":[5,9,5,13,10,4],"isMain":false},{"metricName":"# possibly misassembled contigs","quality":"Less is better","values":[0,0,0,0,0,0],"isMain":false},{"metricName":" # possible misassemblies","quality":"Less is better","values":[0,0,0,0,0,0],"isMain":false},{"metricName":"# local misassemblies","quality":"Less is better","values":[0,0,1,0,1,0],"isMain":false},{"metricName":"# scaffold gap ext. mis.","quality":"Less is better","values":[0,0,0,0,0,0],"isMain":false},{"metricName":"# scaffold gap loc. mis.","quality":"Less is better","values":[0,0,0,0,0,0],"isMain":false}]],["Unaligned",[]],["Mismatches",[]],["Statistics without reference",[]],["K-mer-based statistics",[]],["Predicted genes",[]],["Similarity statistics",[]],["Reference statistics",[]]],[["Genome statistics",[]],["Reads mapping",[]],["Misassemblies",[{"metricName":"# misassemblies","quality":"Less is better","values":[5,9,4,9,8,2],"isMain":true},{"metricName":" # relocations","quality":"Less is better","values":[0,0,0,0,0,0],"isMain":false},{"metricName":" # translocations","quality":"Less is better","values":[0,0,0,0,0,0],"isMain":false},{"metricName":" # inversions","quality":"Less is better","values":[0,0,0,0,0,0],"isMain":false},{"metricName":" # interspecies translocations","quality":"Less is better","values":[5,9,4,9,8,2],"isMain":false},{"metricName":"# possibly misassembled contigs","quality":"Less is better","values":[0,0,0,0,0,0],"isMain":false},{"metricName":" # possible misassemblies","quality":"Less is better","values":[0,0,0,0,0,0],"isMain":false},{"metricName":"# local misassemblies","quality":"Less is better","values":[0,0,0,1,0,0],"isMain":false},{"metricName":"# scaffold gap ext. mis.","quality":"Less is better","values":[0,0,0,0,0,0],"isMain":false},{"metricName":"# scaffold gap loc. mis.","quality":"Less is better","values":[0,0,0,0,0,0],"isMain":false}]],["Unaligned",[]],["Mismatches",[]],["Statistics without reference",[]],["K-mer-based statistics",[]],["Predicted genes",[]],["Similarity statistics",[]],["Reference statistics",[]]],[["Genome statistics",[]],["Reads mapping",[]],["Misassemblies",[{"metricName":"# misassemblies","quality":"Less is better","values":[29,52,15,22,25,8],"isMain":true},{"metricName":" # relocations","quality":"Less is better","values":[0,0,0,0,0,0],"isMain":false},{"metricName":" # translocations","quality":"Less is better","values":[2,3,1,0,2,1],"isMain":false},{"metricName":" # inversions","quality":"Less is better","values":[1,0,0,1,0,0],"isMain":false},{"metricName":" # interspecies translocations","quality":"Less is better","values":[26,49,14,21,23,7],"isMain":false},{"metricName":"# possibly misassembled contigs","quality":"Less is better","values":[0,0,0,0,0,0],"isMain":false},{"metricName":" # possible misassemblies","quality":"Less is better","values":[0,0,0,0,0,0],"isMain":false},{"metricName":"# local misassemblies","quality":"Less is better","values":[5,0,0,0,2,0],"isMain":false},{"metricName":"# scaffold gap ext. mis.","quality":"Less is better","values":[0,0,0,0,0,0],"isMain":false},{"metricName":"# scaffold gap loc. mis.","quality":"Less is better","values":[0,0,0,0,0,0],"isMain":false}]],["Unaligned",[]],["Mismatches",[]],["Statistics without reference",[]],["K-mer-based statistics",[]],["Predicted genes",[]],["Similarity statistics",[]],["Reference statistics",[]]],[["Genome statistics",[]],["Reads mapping",[]],["Misassemblies",[{"metricName":"# misassemblies","quality":"Less is better","values":[10,6,1,8,4,1],"isMain":true},{"metricName":" # relocations","quality":"Less is better","values":[0,0,0,0,0,0],"isMain":false},{"metricName":" # translocations","quality":"Less is better","values":[0,0,0,0,0,0],"isMain":false},{"metricName":" # inversions","quality":"Less is better","values":[0,0,0,0,0,0],"isMain":false},{"metricName":" # interspecies translocations","quality":"Less is better","values":[10,6,1,8,4,1],"isMain":false},{"metricName":"# possibly misassembled contigs","quality":"Less is better","values":[0,0,0,1,0,0],"isMain":false},{"metricName":" # possible misassemblies","quality":"Less is better","values":[0,0,0,1,0,0],"isMain":false},{"metricName":"# local misassemblies","quality":"Less is better","values":[1,1,0,0,1,0],"isMain":false},{"metricName":"# scaffold gap ext. mis.","quality":"Less is better","values":[0,0,0,0,0,0],"isMain":false},{"metricName":"# scaffold gap loc. mis.","quality":"Less is better","values":[0,0,0,0,0,0],"isMain":false}]],["Unaligned",[]],["Mismatches",[]],["Statistics without reference",[]],["K-mer-based statistics",[]],["Predicted genes",[]],["Similarity statistics",[]],["Reference statistics",[]]],[["Genome statistics",[]],["Reads mapping",[]],["Misassemblies",[{"metricName":"# misassemblies","quality":"Less is better","values":[33,37,3,25,34,27],"isMain":true},{"metricName":" # relocations","quality":"Less is better","values":[0,3,0,1,2,0],"isMain":false},{"metricName":" # translocations","quality":"Less is better","values":[0,4,2,2,3,0],"isMain":false},{"metricName":" # inversions","quality":"Less is better","values":[0,0,0,0,0,0],"isMain":false},{"metricName":" # interspecies translocations","quality":"Less is better","values":[33,30,1,22,29,27],"isMain":false},{"metricName":"# possibly misassembled contigs","quality":"Less is better","values":[0,0,0,0,0,0],"isMain":false},{"metricName":" # possible misassemblies","quality":"Less is better","values":[0,0,0,0,0,0],"isMain":false},{"metricName":"# local misassemblies","quality":"Less is better","values":[3,1,6,5,5,0],"isMain":false},{"metricName":"# scaffold gap ext. mis.","quality":"Less is better","values":[0,0,0,0,0,0],"isMain":false},{"metricName":"# scaffold gap loc. mis.","quality":"Less is better","values":[0,0,0,0,0,0],"isMain":false}]],["Unaligned",[]],["Mismatches",[]],["Statistics without reference",[]],["K-mer-based statistics",[]],["Predicted genes",[]],["Similarity statistics",[]],["Reference statistics",[]]],[["Genome statistics",[]],["Reads mapping",[]],["Misassemblies",[{"metricName":"# misassemblies","quality":"Less is better","values":[66,28,40,70,49,2],"isMain":true},{"metricName":" # relocations","quality":"Less is better","values":[0,2,4,6,4,0],"isMain":false},{"metricName":" # translocations","quality":"Less is better","values":[0,0,0,2,1,2],"isMain":false},{"metricName":" # inversions","quality":"Less is better","values":[1,1,0,4,2,0],"isMain":false},{"metricName":" # interspecies translocations","quality":"Less is better","values":[65,25,36,58,42,0],"isMain":false},{"metricName":"# possibly misassembled contigs","quality":"Less is better","values":[0,0,0,1,0,0],"isMain":false},{"metricName":" # possible misassemblies","quality":"Less is better","values":[0,0,0,1,0,0],"isMain":false},{"metricName":"# local misassemblies","quality":"Less is better","values":[3,1,1,5,3,0],"isMain":false},{"metricName":"# scaffold gap ext. mis.","quality":"Less is better","values":[0,0,0,0,0,0],"isMain":false},{"metricName":"# scaffold gap loc. mis.","quality":"Less is better","values":[0,0,0,0,0,0],"isMain":false}]],["Unaligned",[]],["Mismatches",[]],["Statistics without reference",[]],["K-mer-based statistics",[]],["Predicted genes",[]],["Similarity statistics",[]],["Reference statistics",[]]],[["Genome statistics",[]],["Reads mapping",[]],["Misassemblies",[{"metricName":"# misassemblies","quality":"Less is better","values":[6,6,3,6,7,5],"isMain":true},{"metricName":" # relocations","quality":"Less is better","values":[1,1,0,0,1,0],"isMain":false},{"metricName":" # translocations","quality":"Less is better","values":[0,0,0,0,0,0],"isMain":false},{"metricName":" # inversions","quality":"Less is better","values":[0,0,0,0,0,0],"isMain":false},{"metricName":" # interspecies translocations","quality":"Less is better","values":[5,5,3,6,6,5],"isMain":false},{"metricName":"# possibly misassembled contigs","quality":"Less is better","values":[0,0,0,0,0,0],"isMain":false},{"metricName":" # possible misassemblies","quality":"Less is better","values":[0,0,0,0,0,0],"isMain":false},{"metricName":"# local misassemblies","quality":"Less is better","values":[0,0,0,0,0,0],"isMain":false},{"metricName":"# scaffold gap ext. mis.","quality":"Less is better","values":[0,0,0,0,0,0],"isMain":false},{"metricName":"# scaffold gap loc. mis.","quality":"Less is better","values":[0,0,0,0,0,0],"isMain":false}]],["Unaligned",[]],["Mismatches",[]],["Statistics without reference",[]],["K-mer-based statistics",[]],["Predicted genes",[]],["Similarity statistics",[]],["Reference statistics",[]]],[["Genome statistics",[]],["Reads mapping",[]],["Misassemblies",[{"metricName":"# misassemblies","quality":"Less is better","values":[60,105,61,109,115,31],"isMain":true},{"metricName":" # relocations","quality":"Less is better","values":[16,26,23,22,26,6],"isMain":false},{"metricName":" # translocations","quality":"Less is better","values":[0,0,0,0,0,0],"isMain":false},{"metricName":" # inversions","quality":"Less is better","values":[1,0,2,1,1,0],"isMain":false},{"metricName":" # interspecies translocations","quality":"Less is better","values":[43,79,36,86,88,25],"isMain":false},{"metricName":"# possibly misassembled contigs","quality":"Less is better","values":[1,1,1,0,0,0],"isMain":false},{"metricName":" # possible misassemblies","quality":"Less is better","values":[1,1,1,0,0,0],"isMain":false},{"metricName":"# local misassemblies","quality":"Less is better","values":[11,9,16,11,15,1],"isMain":false},{"metricName":"# scaffold gap ext. mis.","quality":"Less is better","values":[0,0,0,0,0,0],"isMain":false},{"metricName":"# scaffold gap loc. mis.","quality":"Less is better","values":[0,0,0,0,0,0],"isMain":false}]],["Unaligned",[]],["Mismatches",[]],["Statistics without reference",[]],["K-mer-based statistics",[]],["Predicted genes",[]],["Similarity statistics",[]],["Reference statistics",[]]],[["Genome statistics",[]],["Reads mapping",[]],["Misassemblies",[{"metricName":"# misassemblies","quality":"Less is better","values":[30,31,9,28,24,19],"isMain":true},{"metricName":" # relocations","quality":"Less is better","values":[3,5,2,2,4,2],"isMain":false},{"metricName":" # translocations","quality":"Less is better","values":[0,0,0,0,0,0],"isMain":false},{"metricName":" # inversions","quality":"Less is better","values":[2,0,0,0,0,0],"isMain":false},{"metricName":" # interspecies translocations","quality":"Less is better","values":[25,26,7,26,20,17],"isMain":false},{"metricName":"# possibly misassembled contigs","quality":"Less is better","values":[0,0,0,1,0,0],"isMain":false},{"metricName":" # possible misassemblies","quality":"Less is better","values":[0,0,0,1,0,0],"isMain":false},{"metricName":"# local misassemblies","quality":"Less is better","values":[0,0,2,2,2,1],"isMain":false},{"metricName":"# scaffold gap ext. mis.","quality":"Less is better","values":[0,0,0,0,0,0],"isMain":false},{"metricName":"# scaffold gap loc. mis.","quality":"Less is better","values":[0,0,0,0,0,0],"isMain":false}]],["Unaligned",[]],["Mismatches",[]],["Statistics without reference",[]],["K-mer-based statistics",[]],["Predicted genes",[]],["Similarity statistics",[]],["Reference statistics",[]]],[["Genome statistics",[]],["Reads mapping",[]],["Misassemblies",[{"metricName":"# misassemblies","quality":"Less is better","values":[0,0,0,0,0,0],"isMain":true},{"metricName":" # relocations","quality":"Less is better","values":[0,0,0,0,0,0],"isMain":false},{"metricName":" # translocations","quality":"Less is better","values":[0,0,0,0,0,0],"isMain":false},{"metricName":" # inversions","quality":"Less is better","values":[0,0,0,0,0,0],"isMain":false},{"metricName":" # interspecies translocations","quality":"Less is better","values":[0,0,0,0,0,0],"isMain":false},{"metricName":"# possibly misassembled contigs","quality":"Less is better","values":[0,0,0,0,0,0],"isMain":false},{"metricName":" # possible misassemblies","quality":"Less is better","values":[0,0,0,0,0,0],"isMain":false},{"metricName":"# local misassemblies","quality":"Less is better","values":[0,0,0,0,0,0],"isMain":false},{"metricName":"# scaffold gap ext. mis.","quality":"Less is better","values":[0,0,0,0,0,0],"isMain":false},{"metricName":"# scaffold gap loc. mis.","quality":"Less is better","values":[0,0,0,0,0,0],"isMain":false}]],["Unaligned",[]],["Mismatches",[]],["Statistics without reference",[]],["K-mer-based statistics",[]],["Predicted genes",[]],["Similarity statistics",[]],["Reference statistics",[]]],[["Genome statistics",[]],["Reads mapping",[]],["Misassemblies",[{"metricName":"# misassemblies","quality":"Less is better","values":[4,8,3,14,13,0],"isMain":true},{"metricName":" # relocations","quality":"Less is better","values":[0,1,1,2,4,0],"isMain":false},{"metricName":" # translocations","quality":"Less is better","values":[0,0,0,0,0,0],"isMain":false},{"metricName":" # inversions","quality":"Less is better","values":[0,0,0,0,0,0],"isMain":false},{"metricName":" # interspecies translocations","quality":"Less is better","values":[4,7,2,12,9,0],"isMain":false},{"metricName":"# possibly misassembled contigs","quality":"Less is better","values":[0,0,0,0,0,0],"isMain":false},{"metricName":" # possible misassemblies","quality":"Less is better","values":[0,0,0,0,0,0],"isMain":false},{"metricName":"# local misassemblies","quality":"Less is better","values":[0,0,5,0,2,0],"isMain":false},{"metricName":"# scaffold gap ext. mis.","quality":"Less is better","values":[0,0,0,0,0,0],"isMain":false},{"metricName":"# scaffold gap loc. mis.","quality":"Less is better","values":[0,0,0,0,0,0],"isMain":false}]],["Unaligned",[]],["Mismatches",[]],["Statistics without reference",[]],["K-mer-based statistics",[]],["Predicted genes",[]],["Similarity statistics",[]],["Reference statistics",[]]],[["Genome statistics",[]],["Reads mapping",[]],["Misassemblies",[{"metricName":"# misassemblies","quality":"Less is better","values":[0,0,0,2,2,0],"isMain":true},{"metricName":" # relocations","quality":"Less is better","values":[0,0,0,0,0,0],"isMain":false},{"metricName":" # translocations","quality":"Less is better","values":[0,0,0,0,0,0],"isMain":false},{"metricName":" # inversions","quality":"Less is better","values":[0,0,0,0,0,0],"isMain":false},{"metricName":" # interspecies translocations","quality":"Less is better","values":[0,0,0,2,2,0],"isMain":false},{"metricName":"# possibly misassembled contigs","quality":"Less is better","values":[0,0,0,0,0,0],"isMain":false},{"metricName":" # possible misassemblies","quality":"Less is better","values":[0,0,0,0,0,0],"isMain":false},{"metricName":"# local misassemblies","quality":"Less is better","values":[0,0,0,0,0,0],"isMain":false},{"metricName":"# scaffold gap ext. mis.","quality":"Less is better","values":[0,0,0,0,0,0],"isMain":false},{"metricName":"# scaffold gap loc. mis.","quality":"Less is better","values":[0,0,0,0,0,0],"isMain":false}]],["Unaligned",[]],["Mismatches",[]],["Statistics without reference",[]],["K-mer-based statistics",[]],["Predicted genes",[]],["Similarity statistics",[]],["Reference statistics",[]]],[["Genome statistics",[]],["Reads mapping",[]],["Misassemblies",[{"metricName":"# misassemblies","quality":"Less is better","values":[2,7,1,8,8,2],"isMain":true},{"metricName":" # relocations","quality":"Less is better","values":[0,0,1,1,1,0],"isMain":false},{"metricName":" # translocations","quality":"Less is better","values":[0,0,0,0,0,0],"isMain":false},{"metricName":" # inversions","quality":"Less is better","values":[0,0,0,0,0,0],"isMain":false},{"metricName":" # interspecies translocations","quality":"Less is better","values":[2,7,0,7,7,2],"isMain":false},{"metricName":"# possibly misassembled contigs","quality":"Less is better","values":[0,0,0,0,0,0],"isMain":false},{"metricName":" # possible misassemblies","quality":"Less is better","values":[0,0,0,0,0,0],"isMain":false},{"metricName":"# local misassemblies","quality":"Less is better","values":[0,1,1,0,0,0],"isMain":false},{"metricName":"# scaffold gap ext. mis.","quality":"Less is better","values":[0,0,0,0,0,0],"isMain":false},{"metricName":"# scaffold gap loc. mis.","quality":"Less is better","values":[0,0,0,0,0,0],"isMain":false}]],["Unaligned",[]],["Mismatches",[]],["Statistics without reference",[]],["K-mer-based statistics",[]],["Predicted genes",[]],["Similarity statistics",[]],["Reference statistics",[]]],[["Genome statistics",[]],["Reads mapping",[]],["Misassemblies",[{"metricName":"# misassemblies","quality":"Less is better","values":[7,3,0,6,2,7],"isMain":true},{"metricName":" # relocations","quality":"Less is better","values":[0,0,0,0,0,0],"isMain":false},{"metricName":" # translocations","quality":"Less is better","values":[0,0,0,0,0,0],"isMain":false},{"metricName":" # inversions","quality":"Less is better","values":[0,0,0,0,0,0],"isMain":false},{"metricName":" # interspecies translocations","quality":"Less is better","values":[7,3,0,6,2,7],"isMain":false},{"metricName":"# possibly misassembled contigs","quality":"Less is better","values":[0,0,0,0,0,0],"isMain":false},{"metricName":" # possible misassemblies","quality":"Less is better","values":[0,0,0,0,0,0],"isMain":false},{"metricName":"# local misassemblies","quality":"Less is better","values":[0,0,0,0,0,0],"isMain":false},{"metricName":"# scaffold gap ext. mis.","quality":"Less is better","values":[0,0,0,0,0,0],"isMain":false},{"metricName":"# scaffold gap loc. mis.","quality":"Less is better","values":[0,0,0,0,0,0],"isMain":false}]],["Unaligned",[]],["Mismatches",[]],["Statistics without reference",[]],["K-mer-based statistics",[]],["Predicted genes",[]],["Similarity statistics",[]],["Reference statistics",[]]],[["Genome statistics",[]],["Reads mapping",[]],["Misassemblies",[{"metricName":"# misassemblies","quality":"Less is better","values":[18,104,176,234,186,1],"isMain":true},{"metricName":" # relocations","quality":"Less is better","values":[0,22,57,80,58,1],"isMain":false},{"metricName":" # translocations","quality":"Less is better","values":[0,0,0,0,0,0],"isMain":false},{"metricName":" # inversions","quality":"Less is better","values":[3,10,11,6,25,0],"isMain":false},{"metricName":" # interspecies translocations","quality":"Less is better","values":[15,72,108,148,103,0],"isMain":false},{"metricName":"# possibly misassembled contigs","quality":"Less is better","values":[1,1,2,1,0,0],"isMain":false},{"metricName":" # possible misassemblies","quality":"Less is better","values":[1,1,2,1,0,0],"isMain":false},{"metricName":"# local misassemblies","quality":"Less is better","values":[0,11,6,14,16,0],"isMain":false},{"metricName":"# scaffold gap ext. mis.","quality":"Less is better","values":[0,0,0,0,0,0],"isMain":false},{"metricName":"# scaffold gap loc. mis.","quality":"Less is better","values":[0,0,0,0,0,0],"isMain":false}]],["Unaligned",[]],["Mismatches",[]],["Statistics without reference",[]],["K-mer-based statistics",[]],["Predicted genes",[]],["Similarity statistics",[]],["Reference statistics",[]]],[["Genome statistics",[]],["Reads mapping",[]],["Misassemblies",[{"metricName":"# misassemblies","quality":"Less is better","values":[1,1,0,2,3,2],"isMain":true},{"metricName":" # relocations","quality":"Less is better","values":[0,0,0,0,0,0],"isMain":false},{"metricName":" # translocations","quality":"Less is better","values":[0,0,0,0,0,0],"isMain":false},{"metricName":" # inversions","quality":"Less is better","values":[0,0,0,0,0,0],"isMain":false},{"metricName":" # interspecies translocations","quality":"Less is better","values":[1,1,0,2,3,2],"isMain":false},{"metricName":"# possibly misassembled contigs","quality":"Less is better","values":[0,0,0,0,0,0],"isMain":false},{"metricName":" # possible misassemblies","quality":"Less is better","values":[0,0,0,0,0,0],"isMain":false},{"metricName":"# local misassemblies","quality":"Less is better","values":[0,0,0,0,0,0],"isMain":false},{"metricName":"# scaffold gap ext. mis.","quality":"Less is better","values":[0,0,0,0,0,0],"isMain":false},{"metricName":"# scaffold gap loc. mis.","quality":"Less is better","values":[0,0,0,0,0,0],"isMain":false}]],["Unaligned",[]],["Mismatches",[]],["Statistics without reference",[]],["K-mer-based statistics",[]],["Predicted genes",[]],["Similarity statistics",[]],["Reference statistics",[]]],[["Genome statistics",[]],["Reads mapping",[]],["Misassemblies",[{"metricName":"# misassemblies","quality":"Less is better","values":[10,5,2,7,7,5],"isMain":true},{"metricName":" # relocations","quality":"Less is better","values":[0,0,0,0,0,0],"isMain":false},{"metricName":" # translocations","quality":"Less is better","values":[1,0,0,0,0,0],"isMain":false},{"metricName":" # inversions","quality":"Less is better","values":[0,0,0,0,0,0],"isMain":false},{"metricName":" # interspecies translocations","quality":"Less is better","values":[9,5,2,7,7,5],"isMain":false},{"metricName":"# possibly misassembled contigs","quality":"Less is better","values":[0,0,0,0,0,0],"isMain":false},{"metricName":" # possible misassemblies","quality":"Less is better","values":[0,0,0,0,0,0],"isMain":false},{"metricName":"# local misassemblies","quality":"Less is better","values":[1,0,0,0,0,0],"isMain":false},{"metricName":"# scaffold gap ext. mis.","quality":"Less is better","values":[0,0,0,0,0,0],"isMain":false},{"metricName":"# scaffold gap loc. mis.","quality":"Less is better","values":[0,0,0,0,0,0],"isMain":false}]],["Unaligned",[]],["Mismatches",[]],["Statistics without reference",[]],["K-mer-based statistics",[]],["Predicted genes",[]],["Similarity statistics",[]],["Reference statistics",[]]],[["Genome statistics",[]],["Reads mapping",[]],["Misassemblies",[{"metricName":"# misassemblies","quality":"Less is better","values":[1,7,2,9,2,0],"isMain":true},{"metricName":" # relocations","quality":"Less is better","values":[0,0,0,0,0,0],"isMain":false},{"metricName":" # translocations","quality":"Less is better","values":[0,0,0,0,0,0],"isMain":false},{"metricName":" # inversions","quality":"Less is better","values":[0,0,0,0,0,0],"isMain":false},{"metricName":" # interspecies translocations","quality":"Less is better","values":[1,7,2,9,2,0],"isMain":false},{"metricName":"# possibly misassembled contigs","quality":"Less is better","values":[0,0,0,0,0,0],"isMain":false},{"metricName":" # possible misassemblies","quality":"Less is better","values":[0,0,0,0,0,0],"isMain":false},{"metricName":"# local misassemblies","quality":"Less is better","values":[0,0,0,0,0,0],"isMain":false},{"metricName":"# scaffold gap ext. mis.","quality":"Less is better","values":[0,0,0,0,0,0],"isMain":false},{"metricName":"# scaffold gap loc. mis.","quality":"Less is better","values":[0,0,0,0,0,0],"isMain":false}]],["Unaligned",[]],["Mismatches",[]],["Statistics without reference",[]],["K-mer-based statistics",[]],["Predicted genes",[]],["Similarity statistics",[]],["Reference statistics",[]]],[["Genome statistics",[]],["Reads mapping",[]],["Misassemblies",[{"metricName":"# misassemblies","quality":"Less is better","values":[3,20,5,10,18,8],"isMain":true},{"metricName":" # relocations","quality":"Less is better","values":[0,0,0,0,0,0],"isMain":false},{"metricName":" # translocations","quality":"Less is better","values":[0,1,0,0,0,0],"isMain":false},{"metricName":" # inversions","quality":"Less is better","values":[0,1,1,0,0,0],"isMain":false},{"metricName":" # interspecies translocations","quality":"Less is better","values":[3,18,4,10,18,8],"isMain":false},{"metricName":"# possibly misassembled contigs","quality":"Less is better","values":[0,0,0,0,0,0],"isMain":false},{"metricName":" # possible misassemblies","quality":"Less is better","values":[0,0,0,0,0,0],"isMain":false},{"metricName":"# local misassemblies","quality":"Less is better","values":[0,1,0,0,1,0],"isMain":false},{"metricName":"# scaffold gap ext. mis.","quality":"Less is better","values":[0,0,0,0,0,0],"isMain":false},{"metricName":"# scaffold gap loc. mis.","quality":"Less is better","values":[0,0,0,0,0,0],"isMain":false}]],["Unaligned",[]],["Mismatches",[]],["Statistics without reference",[]],["K-mer-based statistics",[]],["Predicted genes",[]],["Similarity statistics",[]],["Reference statistics",[]]],[["Genome statistics",[]],["Reads mapping",[]],["Misassemblies",[{"metricName":"# misassemblies","quality":"Less is better","values":[21,46,12,32,23,10],"isMain":true},{"metricName":" # relocations","quality":"Less is better","values":[0,0,1,0,1,1],"isMain":false},{"metricName":" # translocations","quality":"Less is better","values":[0,0,0,0,0,0],"isMain":false},{"metricName":" # inversions","quality":"Less is better","values":[0,1,0,0,0,0],"isMain":false},{"metricName":" # interspecies translocations","quality":"Less is better","values":[21,45,11,32,22,9],"isMain":false},{"metricName":"# possibly misassembled contigs","quality":"Less is better","values":[0,1,0,0,0,0],"isMain":false},{"metricName":" # possible misassemblies","quality":"Less is better","values":[0,2,0,0,0,0],"isMain":false},{"metricName":"# local misassemblies","quality":"Less is better","values":[2,3,2,3,0,2],"isMain":false},{"metricName":"# scaffold gap ext. mis.","quality":"Less is better","values":[0,0,0,0,0,0],"isMain":false},{"metricName":"# scaffold gap loc. mis.","quality":"Less is better","values":[0,0,0,0,0,0],"isMain":false}]],["Unaligned",[]],["Mismatches",[]],["Statistics without reference",[]],["K-mer-based statistics",[]],["Predicted genes",[]],["Similarity statistics",[]],["Reference statistics",[]]],[["Genome statistics",[]],["Reads mapping",[]],["Misassemblies",[{"metricName":"# misassemblies","quality":"Less is better","values":[15,119,138,203,141,2],"isMain":true},{"metricName":" # relocations","quality":"Less is better","values":[0,22,26,48,35,0],"isMain":false},{"metricName":" # translocations","quality":"Less is better","values":[0,0,0,0,0,0],"isMain":false},{"metricName":" # inversions","quality":"Less is better","values":[0,17,11,8,17,0],"isMain":false},{"metricName":" # interspecies translocations","quality":"Less is better","values":[15,80,101,147,89,2],"isMain":false},{"metricName":"# possibly misassembled contigs","quality":"Less is better","values":[0,1,0,1,0,0],"isMain":false},{"metricName":" # possible misassemblies","quality":"Less is better","values":[0,1,0,1,0,0],"isMain":false},{"metricName":"# local misassemblies","quality":"Less is better","values":[0,12,11,15,17,0],"isMain":false},{"metricName":"# scaffold gap ext. mis.","quality":"Less is better","values":[0,0,0,0,0,0],"isMain":false},{"metricName":"# scaffold gap loc. mis.","quality":"Less is better","values":[0,0,0,0,0,0],"isMain":false}]],["Unaligned",[]],["Mismatches",[]],["Statistics without reference",[]],["K-mer-based statistics",[]],["Predicted genes",[]],["Similarity statistics",[]],["Reference statistics",[]]],[["Genome statistics",[]],["Reads mapping",[]],["Misassemblies",[{"metricName":"# misassemblies","quality":"Less is better","values":[51,56,20,84,50,39],"isMain":true},{"metricName":" # relocations","quality":"Less is better","values":[7,14,12,14,14,7],"isMain":false},{"metricName":" # translocations","quality":"Less is better","values":[0,0,0,0,0,0],"isMain":false},{"metricName":" # inversions","quality":"Less is better","values":[3,1,1,6,1,0],"isMain":false},{"metricName":" # interspecies translocations","quality":"Less is better","values":[41,41,7,64,35,32],"isMain":false},{"metricName":"# possibly misassembled contigs","quality":"Less is better","values":[0,0,0,0,0,0],"isMain":false},{"metricName":" # possible misassemblies","quality":"Less is better","values":[0,0,0,0,0,0],"isMain":false},{"metricName":"# local misassemblies","quality":"Less is better","values":[5,5,5,4,7,4],"isMain":false},{"metricName":"# scaffold gap ext. mis.","quality":"Less is better","values":[0,0,0,0,0,0],"isMain":false},{"metricName":"# scaffold gap loc. mis.","quality":"Less is better","values":[0,0,0,0,0,0],"isMain":false}]],["Unaligned",[]],["Mismatches",[]],["Statistics without reference",[]],["K-mer-based statistics",[]],["Predicted genes",[]],["Similarity statistics",[]],["Reference statistics",[]]],[["Genome statistics",[]],["Reads mapping",[]],["Misassemblies",[{"metricName":"# misassemblies","quality":"Less is better","values":[21,26,30,45,41,14],"isMain":true},{"metricName":" # relocations","quality":"Less is better","values":[0,0,0,0,0,0],"isMain":false},{"metricName":" # translocations","quality":"Less is better","values":[4,21,24,34,29,14],"isMain":false},{"metricName":" # inversions","quality":"Less is better","values":[0,0,1,0,1,0],"isMain":false},{"metricName":" # interspecies translocations","quality":"Less is better","values":[17,5,5,11,11,0],"isMain":false},{"metricName":"# possibly misassembled contigs","quality":"Less is better","values":[0,0,0,0,0,0],"isMain":false},{"metricName":" # possible misassemblies","quality":"Less is better","values":[0,0,0,0,0,0],"isMain":false},{"metricName":"# local misassemblies","quality":"Less is better","values":[1,1,1,2,1,0],"isMain":false},{"metricName":"# scaffold gap ext. mis.","quality":"Less is better","values":[0,0,0,0,0,0],"isMain":false},{"metricName":"# scaffold gap loc. mis.","quality":"Less is better","values":[0,0,0,0,0,0],"isMain":false}]],["Unaligned",[]],["Mismatches",[]],["Statistics without reference",[]],["K-mer-based statistics",[]],["Predicted genes",[]],["Similarity statistics",[]],["Reference statistics",[]]],[["Genome statistics",[]],["Reads mapping",[]],["Misassemblies",[{"metricName":"# misassemblies","quality":"Less is better","values":[27,41,14,34,41,13],"isMain":true},{"metricName":" # relocations","quality":"Less is better","values":[0,0,0,0,0,0],"isMain":false},{"metricName":" # translocations","quality":"Less is better","values":[1,1,0,1,0,0],"isMain":false},{"metricName":" # inversions","quality":"Less is better","values":[0,0,1,0,0,0],"isMain":false},{"metricName":" # interspecies translocations","quality":"Less is better","values":[26,40,13,33,41,13],"isMain":false},{"metricName":"# possibly misassembled contigs","quality":"Less is better","values":[0,0,0,0,1,0],"isMain":false},{"metricName":" # possible misassemblies","quality":"Less is better","values":[0,0,0,0,1,0],"isMain":false},{"metricName":"# local misassemblies","quality":"Less is better","values":[1,1,1,1,1,0],"isMain":false},{"metricName":"# scaffold gap ext. mis.","quality":"Less is better","values":[0,0,0,0,0,0],"isMain":false},{"metricName":"# scaffold gap loc. mis.","quality":"Less is better","values":[0,0,0,0,0,0],"isMain":false}]],["Unaligned",[]],["Mismatches",[]],["Statistics without reference",[]],["K-mer-based statistics",[]],["Predicted genes",[]],["Similarity statistics",[]],["Reference statistics",[]]],[["Genome statistics",[]],["Reads mapping",[]],["Misassemblies",[{"metricName":"# misassemblies","quality":"Less is better","values":[27,48,26,58,54,21],"isMain":true},{"metricName":" # relocations","quality":"Less is better","values":[0,0,0,0,0,0],"isMain":false},{"metricName":" # translocations","quality":"Less is better","values":[3,2,0,3,1,1],"isMain":false},{"metricName":" # inversions","quality":"Less is better","values":[1,0,0,0,0,0],"isMain":false},{"metricName":" # interspecies translocations","quality":"Less is better","values":[23,46,26,55,53,20],"isMain":false},{"metricName":"# possibly misassembled contigs","quality":"Less is better","values":[2,0,0,0,0,0],"isMain":false},{"metricName":" # possible misassemblies","quality":"Less is better","values":[2,0,0,0,0,0],"isMain":false},{"metricName":"# local misassemblies","quality":"Less is better","values":[0,0,3,2,0,0],"isMain":false},{"metricName":"# scaffold gap ext. mis.","quality":"Less is better","values":[0,0,0,0,0,0],"isMain":false},{"metricName":"# scaffold gap loc. mis.","quality":"Less is better","values":[0,0,0,0,0,0],"isMain":false}]],["Unaligned",[]],["Mismatches",[]],["Statistics without reference",[]],["K-mer-based statistics",[]],["Predicted genes",[]],["Similarity statistics",[]],["Reference statistics",[]]],[["Genome statistics",[]],["Reads mapping",[]],["Misassemblies",[{"metricName":"# misassemblies","quality":"Less is better","values":[12,29,31,44,33,7],"isMain":true},{"metricName":" # relocations","quality":"Less is better","values":[1,0,0,0,0,0],"isMain":false},{"metricName":" # translocations","quality":"Less is better","values":[1,0,4,6,2,0],"isMain":false},{"metricName":" # inversions","quality":"Less is better","values":[0,0,1,0,1,0],"isMain":false},{"metricName":" # interspecies translocations","quality":"Less is better","values":[10,29,26,38,30,7],"isMain":false},{"metricName":"# possibly misassembled contigs","quality":"Less is better","values":[0,0,1,1,2,0],"isMain":false},{"metricName":" # possible misassemblies","quality":"Less is better","values":[0,0,1,2,2,0],"isMain":false},{"metricName":"# local misassemblies","quality":"Less is better","values":[0,2,1,1,0,0],"isMain":false},{"metricName":"# scaffold gap ext. mis.","quality":"Less is better","values":[0,0,0,0,0,0],"isMain":false},{"metricName":"# scaffold gap loc. mis.","quality":"Less is better","values":[0,0,0,0,0,0],"isMain":false}]],["Unaligned",[]],["Mismatches",[]],["Statistics without reference",[]],["K-mer-based statistics",[]],["Predicted genes",[]],["Similarity statistics",[]],["Reference statistics",[]]],[["Genome statistics",[]],["Reads mapping",[]],["Misassemblies",[{"metricName":"# misassemblies","quality":"Less is better","values":[33,38,18,40,43,14],"isMain":true},{"metricName":" # relocations","quality":"Less is better","values":[0,2,0,2,1,0],"isMain":false},{"metricName":" # translocations","quality":"Less is better","values":[1,0,0,1,1,0],"isMain":false},{"metricName":" # inversions","quality":"Less is better","values":[0,1,2,1,0,0],"isMain":false},{"metricName":" # interspecies translocations","quality":"Less is better","values":[32,35,16,36,41,14],"isMain":false},{"metricName":"# possibly misassembled contigs","quality":"Less is better","values":[1,0,0,0,1,0],"isMain":false},{"metricName":" # possible misassemblies","quality":"Less is better","values":[1,0,0,0,1,0],"isMain":false},{"metricName":"# local misassemblies","quality":"Less is better","values":[0,1,0,0,2,0],"isMain":false},{"metricName":"# scaffold gap ext. mis.","quality":"Less is better","values":[0,0,0,0,0,0],"isMain":false},{"metricName":"# scaffold gap loc. mis.","quality":"Less is better","values":[0,0,0,0,0,0],"isMain":false}]],["Unaligned",[]],["Mismatches",[]],["Statistics without reference",[]],["K-mer-based statistics",[]],["Predicted genes",[]],["Similarity statistics",[]],["Reference statistics",[]]],[["Genome statistics",[]],["Reads mapping",[]],["Misassemblies",[{"metricName":"# misassemblies","quality":"Less is better","values":[7,15,6,12,4,4],"isMain":true},{"metricName":" # relocations","quality":"Less is better","values":[0,0,0,0,0,0],"isMain":false},{"metricName":" # translocations","quality":"Less is better","values":[0,1,0,0,0,0],"isMain":false},{"metricName":" # inversions","quality":"Less is better","values":[0,0,0,0,0,0],"isMain":false},{"metricName":" # interspecies translocations","quality":"Less is better","values":[7,14,6,12,4,4],"isMain":false},{"metricName":"# possibly misassembled contigs","quality":"Less is better","values":[0,0,0,0,0,0],"isMain":false},{"metricName":" # possible misassemblies","quality":"Less is better","values":[0,0,0,0,0,0],"isMain":false},{"metricName":"# local misassemblies","quality":"Less is better","values":[0,0,0,0,0,0],"isMain":false},{"metricName":"# scaffold gap ext. mis.","quality":"Less is better","values":[0,0,0,0,0,0],"isMain":false},{"metricName":"# scaffold gap loc. mis.","quality":"Less is better","values":[0,0,0,0,0,0],"isMain":false}]],["Unaligned",[]],["Mismatches",[]],["Statistics without reference",[]],["K-mer-based statistics",[]],["Predicted genes",[]],["Similarity statistics",[]],["Reference statistics",[]]],[["Genome statistics",[]],["Reads mapping",[]],["Misassemblies",[{"metricName":"# misassemblies","quality":"Less is better","values":[2,5,4,7,1,0],"isMain":true},{"metricName":" # relocations","quality":"Less is better","values":[0,0,0,0,0,0],"isMain":false},{"metricName":" # translocations","quality":"Less is better","values":[0,0,0,0,0,0],"isMain":false},{"metricName":" # inversions","quality":"Less is better","values":[0,0,0,0,0,0],"isMain":false},{"metricName":" # interspecies translocations","quality":"Less is better","values":[2,5,4,7,1,0],"isMain":false},{"metricName":"# possibly misassembled contigs","quality":"Less is better","values":[0,0,0,0,0,0],"isMain":false},{"metricName":" # possible misassemblies","quality":"Less is better","values":[0,0,0,0,0,0],"isMain":false},{"metricName":"# local misassemblies","quality":"Less is better","values":[0,1,0,0,0,0],"isMain":false},{"metricName":"# scaffold gap ext. mis.","quality":"Less is better","values":[0,0,0,0,0,0],"isMain":false},{"metricName":"# scaffold gap loc. mis.","quality":"Less is better","values":[0,0,0,0,0,0],"isMain":false}]],["Unaligned",[]],["Mismatches",[]],["Statistics without reference",[]],["K-mer-based statistics",[]],["Predicted genes",[]],["Similarity statistics",[]],["Reference statistics",[]]],[["Genome statistics",[]],["Reads mapping",[]],["Misassemblies",[{"metricName":"# misassemblies","quality":"Less is better","values":[13,18,10,26,22,9],"isMain":true},{"metricName":" # relocations","quality":"Less is better","values":[0,0,0,0,0,0],"isMain":false},{"metricName":" # translocations","quality":"Less is better","values":[2,2,2,5,2,2],"isMain":false},{"metricName":" # inversions","quality":"Less is better","values":[0,0,0,0,0,0],"isMain":false},{"metricName":" # interspecies translocations","quality":"Less is better","values":[11,16,8,21,20,7],"isMain":false},{"metricName":"# possibly misassembled contigs","quality":"Less is better","values":[0,0,0,0,0,0],"isMain":false},{"metricName":" # possible misassemblies","quality":"Less is better","values":[0,0,0,0,0,0],"isMain":false},{"metricName":"# local misassemblies","quality":"Less is better","values":[0,0,2,0,1,0],"isMain":false},{"metricName":"# scaffold gap ext. mis.","quality":"Less is better","values":[0,0,0,0,0,0],"isMain":false},{"metricName":"# scaffold gap loc. mis.","quality":"Less is better","values":[0,0,0,0,0,0],"isMain":false}]],["Unaligned",[]],["Mismatches",[]],["Statistics without reference",[]],["K-mer-based statistics",[]],["Predicted genes",[]],["Similarity statistics",[]],["Reference statistics",[]]],[["Genome statistics",[]],["Reads mapping",[]],["Misassemblies",[{"metricName":"# misassemblies","quality":"Less is better","values":[5,13,17,24,19,3],"isMain":true},{"metricName":" # relocations","quality":"Less is better","values":[0,2,1,1,4,2],"isMain":false},{"metricName":" # translocations","quality":"Less is better","values":[0,1,0,1,0,0],"isMain":false},{"metricName":" # inversions","quality":"Less is better","values":[0,0,0,0,0,0],"isMain":false},{"metricName":" # interspecies translocations","quality":"Less is better","values":[5,10,16,22,15,1],"isMain":false},{"metricName":"# possibly misassembled contigs","quality":"Less is better","values":[0,0,0,0,0,0],"isMain":false},{"metricName":" # possible misassemblies","quality":"Less is better","values":[0,0,0,0,0,0],"isMain":false},{"metricName":"# local misassemblies","quality":"Less is better","values":[0,1,1,2,0,0],"isMain":false},{"metricName":"# scaffold gap ext. mis.","quality":"Less is better","values":[0,0,0,0,0,0],"isMain":false},{"metricName":"# scaffold gap loc. mis.","quality":"Less is better","values":[0,0,0,0,0,0],"isMain":false}]],["Unaligned",[]],["Mismatches",[]],["Statistics without reference",[]],["K-mer-based statistics",[]],["Predicted genes",[]],["Similarity statistics",[]],["Reference statistics",[]]],[["Genome statistics",[]],["Reads mapping",[]],["Misassemblies",[{"metricName":"# misassemblies","quality":"Less is better","values":[9,10,5,16,8,3],"isMain":true},{"metricName":" # relocations","quality":"Less is better","values":[1,1,0,0,1,2],"isMain":false},{"metricName":" # translocations","quality":"Less is better","values":[0,0,0,0,0,0],"isMain":false},{"metricName":" # inversions","quality":"Less is better","values":[0,0,0,0,0,0],"isMain":false},{"metricName":" # interspecies translocations","quality":"Less is better","values":[8,9,5,16,7,1],"isMain":false},{"metricName":"# possibly misassembled contigs","quality":"Less is better","values":[0,0,0,0,0,0],"isMain":false},{"metricName":" # possible misassemblies","quality":"Less is better","values":[0,0,0,0,0,0],"isMain":false},{"metricName":"# local misassemblies","quality":"Less is better","values":[1,0,0,0,1,0],"isMain":false},{"metricName":"# scaffold gap ext. mis.","quality":"Less is better","values":[0,0,0,0,0,0],"isMain":false},{"metricName":"# scaffold gap loc. mis.","quality":"Less is better","values":[0,0,0,0,0,0],"isMain":false}]],["Unaligned",[]],["Mismatches",[]],["Statistics without reference",[]],["K-mer-based statistics",[]],["Predicted genes",[]],["Similarity statistics",[]],["Reference statistics",[]]],[["Genome statistics",[]],["Reads mapping",[]],["Misassemblies",[{"metricName":"# misassemblies","quality":"Less is better","values":[23,25,39,54,37,14],"isMain":true},{"metricName":" # relocations","quality":"Less is better","values":[0,0,0,0,0,0],"isMain":false},{"metricName":" # translocations","quality":"Less is better","values":[2,11,17,15,18,12],"isMain":false},{"metricName":" # inversions","quality":"Less is better","values":[0,0,0,1,1,0],"isMain":false},{"metricName":" # interspecies translocations","quality":"Less is better","values":[21,14,22,38,18,2],"isMain":false},{"metricName":"# possibly misassembled contigs","quality":"Less is better","values":[0,0,0,0,0,0],"isMain":false},{"metricName":" # possible misassemblies","quality":"Less is better","values":[0,0,0,0,0,0],"isMain":false},{"metricName":"# local misassemblies","quality":"Less is better","values":[2,1,1,4,2,0],"isMain":false},{"metricName":"# scaffold gap ext. mis.","quality":"Less is better","values":[0,0,0,0,0,0],"isMain":false},{"metricName":"# scaffold gap loc. mis.","quality":"Less is better","values":[0,0,0,0,0,0],"isMain":false}]],["Unaligned",[]],["Mismatches",[]],["Statistics without reference",[]],["K-mer-based statistics",[]],["Predicted genes",[]],["Similarity statistics",[]],["Reference statistics",[]]],[["Genome statistics",[]],["Reads mapping",[]],["Misassemblies",[{"metricName":"# misassemblies","quality":"Less is better","values":[3,7,9,17,9,0],"isMain":true},{"metricName":" # relocations","quality":"Less is better","values":[0,1,1,3,1,0],"isMain":false},{"metricName":" # translocations","quality":"Less is better","values":[0,0,0,0,0,0],"isMain":false},{"metricName":" # inversions","quality":"Less is better","values":[0,0,0,0,0,0],"isMain":false},{"metricName":" # interspecies translocations","quality":"Less is better","values":[3,6,8,14,8,0],"isMain":false},{"metricName":"# possibly misassembled contigs","quality":"Less is better","values":[0,0,0,0,0,0],"isMain":false},{"metricName":" # possible misassemblies","quality":"Less is better","values":[0,0,0,0,0,0],"isMain":false},{"metricName":"# local misassemblies","quality":"Less is better","values":[0,0,0,1,1,0],"isMain":false},{"metricName":"# scaffold gap ext. mis.","quality":"Less is better","values":[0,0,0,0,0,0],"isMain":false},{"metricName":"# scaffold gap loc. mis.","quality":"Less is better","values":[0,0,0,0,0,0],"isMain":false}]],["Unaligned",[]],["Mismatches",[]],["Statistics without reference",[]],["K-mer-based statistics",[]],["Predicted genes",[]],["Similarity statistics",[]],["Reference statistics",[]]],[["Genome statistics",[]],["Reads mapping",[]],["Misassemblies",[{"metricName":"# misassemblies","quality":"Less is better","values":[4,12,12,27,17,2],"isMain":true},{"metricName":" # relocations","quality":"Less is better","values":[0,5,3,4,4,0],"isMain":false},{"metricName":" # translocations","quality":"Less is better","values":[0,0,0,0,0,0],"isMain":false},{"metricName":" # inversions","quality":"Less is better","values":[1,0,1,0,0,0],"isMain":false},{"metricName":" # interspecies translocations","quality":"Less is better","values":[3,7,8,23,13,2],"isMain":false},{"metricName":"# possibly misassembled contigs","quality":"Less is better","values":[0,0,0,0,0,0],"isMain":false},{"metricName":" # possible misassemblies","quality":"Less is better","values":[0,0,0,0,0,0],"isMain":false},{"metricName":"# local misassemblies","quality":"Less is better","values":[0,2,2,3,1,1],"isMain":false},{"metricName":"# scaffold gap ext. mis.","quality":"Less is better","values":[0,0,0,0,0,0],"isMain":false},{"metricName":"# scaffold gap loc. mis.","quality":"Less is better","values":[0,0,0,0,0,0],"isMain":false}]],["Unaligned",[]],["Mismatches",[]],["Statistics without reference",[]],["K-mer-based statistics",[]],["Predicted genes",[]],["Similarity statistics",[]],["Reference statistics",[]]],[["Genome statistics",[]],["Reads mapping",[]],["Misassemblies",[{"metricName":"# misassemblies","quality":"Less is better","values":[4,10,4,11,4,5],"isMain":true},{"metricName":" # relocations","quality":"Less is better","values":[0,0,0,0,0,0],"isMain":false},{"metricName":" # translocations","quality":"Less is better","values":[0,0,0,0,0,0],"isMain":false},{"metricName":" # inversions","quality":"Less is better","values":[0,0,0,0,0,0],"isMain":false},{"metricName":" # interspecies translocations","quality":"Less is better","values":[4,10,4,11,4,5],"isMain":false},{"metricName":"# possibly misassembled contigs","quality":"Less is better","values":[0,0,0,0,0,0],"isMain":false},{"metricName":" # possible misassemblies","quality":"Less is better","values":[0,0,0,0,0,0],"isMain":false},{"metricName":"# local misassemblies","quality":"Less is better","values":[0,0,0,0,0,0],"isMain":false},{"metricName":"# scaffold gap ext. mis.","quality":"Less is better","values":[0,0,0,0,0,0],"isMain":false},{"metricName":"# scaffold gap loc. mis.","quality":"Less is better","values":[0,0,0,0,0,0],"isMain":false}]],["Unaligned",[]],["Mismatches",[]],["Statistics without reference",[]],["K-mer-based statistics",[]],["Predicted genes",[]],["Similarity statistics",[]],["Reference statistics",[]]],[["Genome statistics",[]],["Reads mapping",[]],["Misassemblies",[{"metricName":"# misassemblies","quality":"Less is better","values":[6,18,12,38,19,2],"isMain":true},{"metricName":" # relocations","quality":"Less is better","values":[5,10,8,7,7,0],"isMain":false},{"metricName":" # translocations","quality":"Less is better","values":[0,1,0,1,0,0],"isMain":false},{"metricName":" # inversions","quality":"Less is better","values":[0,0,0,2,0,0],"isMain":false},{"metricName":" # interspecies translocations","quality":"Less is better","values":[1,7,4,28,12,2],"isMain":false},{"metricName":"# possibly misassembled contigs","quality":"Less is better","values":[0,0,0,0,0,0],"isMain":false},{"metricName":" # possible misassemblies","quality":"Less is better","values":[0,0,0,0,0,0],"isMain":false},{"metricName":"# local misassemblies","quality":"Less is better","values":[1,4,6,4,6,0],"isMain":false},{"metricName":"# scaffold gap ext. mis.","quality":"Less is better","values":[0,0,0,0,0,0],"isMain":false},{"metricName":"# scaffold gap loc. mis.","quality":"Less is better","values":[0,0,0,0,0,0],"isMain":false}]],["Unaligned",[]],["Mismatches",[]],["Statistics without reference",[]],["K-mer-based statistics",[]],["Predicted genes",[]],["Similarity statistics",[]],["Reference statistics",[]]],[["Genome statistics",[]],["Reads mapping",[]],["Misassemblies",[{"metricName":"# misassemblies","quality":"Less is better","values":[3,5,1,4,4,4],"isMain":true},{"metricName":" # relocations","quality":"Less is better","values":[0,0,0,0,0,0],"isMain":false},{"metricName":" # translocations","quality":"Less is better","values":[0,0,0,0,0,0],"isMain":false},{"metricName":" # inversions","quality":"Less is better","values":[0,0,0,0,0,0],"isMain":false},{"metricName":" # interspecies translocations","quality":"Less is better","values":[3,5,1,4,4,4],"isMain":false},{"metricName":"# possibly misassembled contigs","quality":"Less is better","values":[0,0,1,0,0,0],"isMain":false},{"metricName":" # possible misassemblies","quality":"Less is better","values":[0,0,1,0,0,0],"isMain":false},{"metricName":"# local misassemblies","quality":"Less is better","values":[0,0,0,1,0,0],"isMain":false},{"metricName":"# scaffold gap ext. mis.","quality":"Less is better","values":[0,0,0,0,0,0],"isMain":false},{"metricName":"# scaffold gap loc. mis.","quality":"Less is better","values":[0,0,0,0,0,0],"isMain":false}]],["Unaligned",[]],["Mismatches",[]],["Statistics without reference",[]],["K-mer-based statistics",[]],["Predicted genes",[]],["Similarity statistics",[]],["Reference statistics",[]]],[["Genome statistics",[]],["Reads mapping",[]],["Misassemblies",[{"metricName":"# misassemblies","quality":"Less is better","values":[6,7,3,4,8,6],"isMain":true},{"metricName":" # relocations","quality":"Less is better","values":[0,0,0,0,0,0],"isMain":false},{"metricName":" # translocations","quality":"Less is better","values":[0,0,0,1,0,0],"isMain":false},{"metricName":" # inversions","quality":"Less is better","values":[0,0,0,0,0,0],"isMain":false},{"metricName":" # interspecies translocations","quality":"Less is better","values":[6,7,3,3,8,6],"isMain":false},{"metricName":"# possibly misassembled contigs","quality":"Less is better","values":[0,0,0,0,0,0],"isMain":false},{"metricName":" # possible misassemblies","quality":"Less is better","values":[0,0,0,0,0,0],"isMain":false},{"metricName":"# local misassemblies","quality":"Less is better","values":[0,1,0,0,1,0],"isMain":false},{"metricName":"# scaffold gap ext. mis.","quality":"Less is better","values":[0,0,0,0,0,0],"isMain":false},{"metricName":"# scaffold gap loc. mis.","quality":"Less is better","values":[0,0,0,0,0,0],"isMain":false}]],["Unaligned",[]],["Mismatches",[]],["Statistics without reference",[]],["K-mer-based statistics",[]],["Predicted genes",[]],["Similarity statistics",[]],["Reference statistics",[]]],[["Genome statistics",[]],["Reads mapping",[]],["Misassemblies",[{"metricName":"# misassemblies","quality":"Less is better","values":[5,8,4,4,7,9],"isMain":true},{"metricName":" # relocations","quality":"Less is better","values":[0,0,0,0,0,0],"isMain":false},{"metricName":" # translocations","quality":"Less is better","values":[0,0,0,0,0,0],"isMain":false},{"metricName":" # inversions","quality":"Less is better","values":[0,0,0,0,0,0],"isMain":false},{"metricName":" # interspecies translocations","quality":"Less is better","values":[5,8,4,4,7,9],"isMain":false},{"metricName":"# possibly misassembled contigs","quality":"Less is better","values":[0,0,0,0,0,0],"isMain":false},{"metricName":" # possible misassemblies","quality":"Less is better","values":[0,0,0,0,0,0],"isMain":false},{"metricName":"# local misassemblies","quality":"Less is better","values":[0,0,0,0,0,0],"isMain":false},{"metricName":"# scaffold gap ext. mis.","quality":"Less is better","values":[0,0,0,0,0,0],"isMain":false},{"metricName":"# scaffold gap loc. mis.","quality":"Less is better","values":[0,0,0,0,0,0],"isMain":false}]],["Unaligned",[]],["Mismatches",[]],["Statistics without reference",[]],["K-mer-based statistics",[]],["Predicted genes",[]],["Similarity statistics",[]],["Reference statistics",[]]],[["Genome statistics",[]],["Reads mapping",[]],["Misassemblies",[{"metricName":"# misassemblies","quality":"Less is better","values":[10,12,5,13,15,13],"isMain":true},{"metricName":" # relocations","quality":"Less is better","values":[0,0,0,0,0,1],"isMain":false},{"metricName":" # translocations","quality":"Less is better","values":[0,0,0,0,0,1],"isMain":false},{"metricName":" # inversions","quality":"Less is better","values":[0,1,1,0,1,0],"isMain":false},{"metricName":" # interspecies translocations","quality":"Less is better","values":[10,11,4,13,14,11],"isMain":false},{"metricName":"# possibly misassembled contigs","quality":"Less is better","values":[0,0,0,0,0,0],"isMain":false},{"metricName":" # possible misassemblies","quality":"Less is better","values":[0,0,0,0,0,0],"isMain":false},{"metricName":"# local misassemblies","quality":"Less is better","values":[0,0,0,0,0,1],"isMain":false},{"metricName":"# scaffold gap ext. mis.","quality":"Less is better","values":[0,0,0,0,0,0],"isMain":false},{"metricName":"# scaffold gap loc. mis.","quality":"Less is better","values":[0,0,0,0,0,0],"isMain":false}]],["Unaligned",[]],["Mismatches",[]],["Statistics without reference",[]],["K-mer-based statistics",[]],["Predicted genes",[]],["Similarity statistics",[]],["Reference statistics",[]]],[["Genome statistics",[]],["Reads mapping",[]],["Misassemblies",[{"metricName":"# misassemblies","quality":"Less is better","values":[7,29,20,24,15,2],"isMain":true},{"metricName":" # relocations","quality":"Less is better","values":[0,0,0,0,0,0],"isMain":false},{"metricName":" # translocations","quality":"Less is better","values":[0,1,2,0,1,0],"isMain":false},{"metricName":" # inversions","quality":"Less is better","values":[2,2,0,0,0,0],"isMain":false},{"metricName":" # interspecies translocations","quality":"Less is better","values":[5,26,18,24,14,2],"isMain":false},{"metricName":"# possibly misassembled contigs","quality":"Less is better","values":[0,0,0,0,0,0],"isMain":false},{"metricName":" # possible misassemblies","quality":"Less is better","values":[0,0,0,0,0,0],"isMain":false},{"metricName":"# local misassemblies","quality":"Less is better","values":[0,3,1,5,4,1],"isMain":false},{"metricName":"# scaffold gap ext. mis.","quality":"Less is better","values":[0,0,0,0,0,0],"isMain":false},{"metricName":"# scaffold gap loc. mis.","quality":"Less is better","values":[0,0,0,0,0,0],"isMain":false}]],["Unaligned",[]],["Mismatches",[]],["Statistics without reference",[]],["K-mer-based statistics",[]],["Predicted genes",[]],["Similarity statistics",[]],["Reference statistics",[]]],[["Genome statistics",[]],["Reads mapping",[]],["Misassemblies",[{"metricName":"# misassemblies","quality":"Less is better","values":[9,22,7,13,15,12],"isMain":true},{"metricName":" # relocations","quality":"Less is better","values":[0,0,0,0,0,0],"isMain":false},{"metricName":" # translocations","quality":"Less is better","values":[0,1,1,0,0,0],"isMain":false},{"metricName":" # inversions","quality":"Less is better","values":[0,0,0,0,0,0],"isMain":false},{"metricName":" # interspecies translocations","quality":"Less is better","values":[9,21,6,13,15,12],"isMain":false},{"metricName":"# possibly misassembled contigs","quality":"Less is better","values":[0,0,0,0,0,0],"isMain":false},{"metricName":" # possible misassemblies","quality":"Less is better","values":[0,0,0,0,0,0],"isMain":false},{"metricName":"# local misassemblies","quality":"Less is better","values":[1,0,1,0,0,0],"isMain":false},{"metricName":"# scaffold gap ext. mis.","quality":"Less is better","values":[0,0,0,0,0,0],"isMain":false},{"metricName":"# scaffold gap loc. mis.","quality":"Less is better","values":[0,0,0,0,0,0],"isMain":false}]],["Unaligned",[]],["Mismatches",[]],["Statistics without reference",[]],["K-mer-based statistics",[]],["Predicted genes",[]],["Similarity statistics",[]],["Reference statistics",[]]],[["Genome statistics",[]],["Reads mapping",[]],["Misassemblies",[{"metricName":"# misassemblies","quality":"Less is better","values":[4,1,2,6,4,1],"isMain":true},{"metricName":" # relocations","quality":"Less is better","values":[0,0,0,0,0,0],"isMain":false},{"metricName":" # translocations","quality":"Less is better","values":[0,0,0,0,0,0],"isMain":false},{"metricName":" # inversions","quality":"Less is better","values":[0,0,0,0,0,0],"isMain":false},{"metricName":" # interspecies translocations","quality":"Less is better","values":[4,1,2,6,4,1],"isMain":false},{"metricName":"# possibly misassembled contigs","quality":"Less is better","values":[0,0,0,0,0,0],"isMain":false},{"metricName":" # possible misassemblies","quality":"Less is better","values":[0,0,0,0,0,0],"isMain":false},{"metricName":"# local misassemblies","quality":"Less is better","values":[0,0,0,0,0,0],"isMain":false},{"metricName":"# scaffold gap ext. mis.","quality":"Less is better","values":[0,0,0,0,0,0],"isMain":false},{"metricName":"# scaffold gap loc. mis.","quality":"Less is better","values":[0,0,0,0,0,0],"isMain":false}]],["Unaligned",[]],["Mismatches",[]],["Statistics without reference",[]],["K-mer-based statistics",[]],["Predicted genes",[]],["Similarity statistics",[]],["Reference statistics",[]]],[["Genome statistics",[]],["Reads mapping",[]],["Misassemblies",[{"metricName":"# misassemblies","quality":"Less is better","values":[0,0,0,1,0,0],"isMain":true},{"metricName":" # relocations","quality":"Less is better","values":[0,0,0,0,0,0],"isMain":false},{"metricName":" # translocations","quality":"Less is better","values":[0,0,0,0,0,0],"isMain":false},{"metricName":" # inversions","quality":"Less is better","values":[0,0,0,0,0,0],"isMain":false},{"metricName":" # interspecies translocations","quality":"Less is better","values":[0,0,0,1,0,0],"isMain":false},{"metricName":"# possibly misassembled contigs","quality":"Less is better","values":[0,0,0,0,0,0],"isMain":false},{"metricName":" # possible misassemblies","quality":"Less is better","values":[0,0,0,0,0,0],"isMain":false},{"metricName":"# local misassemblies","quality":"Less is better","values":[0,0,0,0,0,0],"isMain":false},{"metricName":"# scaffold gap ext. mis.","quality":"Less is better","values":[0,0,0,0,0,0],"isMain":false},{"metricName":"# scaffold gap loc. mis.","quality":"Less is better","values":[0,0,0,0,0,0],"isMain":false}]],["Unaligned",[]],["Mismatches",[]],["Statistics without reference",[]],["K-mer-based statistics",[]],["Predicted genes",[]],["Similarity statistics",[]],["Reference statistics",[]]],[["Genome statistics",[]],["Reads mapping",[]],["Misassemblies",[{"metricName":"# misassemblies","quality":"Less is better","values":[30,50,20,31,34,21],"isMain":true},{"metricName":" # relocations","quality":"Less is better","values":[8,14,12,9,12,7],"isMain":false},{"metricName":" # translocations","quality":"Less is better","values":[0,0,0,0,0,0],"isMain":false},{"metricName":" # inversions","quality":"Less is better","values":[0,0,0,0,0,0],"isMain":false},{"metricName":" # interspecies translocations","quality":"Less is better","values":[22,36,8,22,22,14],"isMain":false},{"metricName":"# possibly misassembled contigs","quality":"Less is better","values":[0,2,0,0,0,0],"isMain":false},{"metricName":" # possible misassemblies","quality":"Less is better","values":[0,2,0,0,0,0],"isMain":false},{"metricName":"# local misassemblies","quality":"Less is better","values":[0,0,1,3,2,1],"isMain":false},{"metricName":"# scaffold gap ext. mis.","quality":"Less is better","values":[0,0,0,0,0,0],"isMain":false},{"metricName":"# scaffold gap loc. mis.","quality":"Less is better","values":[0,0,0,0,0,0],"isMain":false}]],["Unaligned",[]],["Mismatches",[]],["Statistics without reference",[]],["K-mer-based statistics",[]],["Predicted genes",[]],["Similarity statistics",[]],["Reference statistics",[]]],[["Genome statistics",[]],["Reads mapping",[]],["Misassemblies",[{"metricName":"# misassemblies","quality":"Less is better","values":[7,14,6,13,10,0],"isMain":true},{"metricName":" # relocations","quality":"Less is better","values":[1,0,0,0,0,0],"isMain":false},{"metricName":" # translocations","quality":"Less is better","values":[0,0,0,0,0,0],"isMain":false},{"metricName":" # inversions","quality":"Less is better","values":[0,1,1,0,1,0],"isMain":false},{"metricName":" # interspecies translocations","quality":"Less is better","values":[6,13,5,13,9,0],"isMain":false},{"metricName":"# possibly misassembled contigs","quality":"Less is better","values":[0,0,1,0,0,0],"isMain":false},{"metricName":" # possible misassemblies","quality":"Less is better","values":[0,0,1,0,0,0],"isMain":false},{"metricName":"# local misassemblies","quality":"Less is better","values":[0,0,0,0,0,0],"isMain":false},{"metricName":"# scaffold gap ext. mis.","quality":"Less is better","values":[0,0,0,0,0,0],"isMain":false},{"metricName":"# scaffold gap loc. mis.","quality":"Less is better","values":[0,0,0,0,0,0],"isMain":false}]],["Unaligned",[]],["Mismatches",[]],["Statistics without reference",[]],["K-mer-based statistics",[]],["Predicted genes",[]],["Similarity statistics",[]],["Reference statistics",[]]],[["Genome statistics",[]],["Reads mapping",[]],["Misassemblies",[{"metricName":"# misassemblies","quality":"Less is better","values":[7,3,4,4,8,1],"isMain":true},{"metricName":" # relocations","quality":"Less is better","values":[0,0,0,0,0,0],"isMain":false},{"metricName":" # translocations","quality":"Less is better","values":[0,1,1,0,0,0],"isMain":false},{"metricName":" # inversions","quality":"Less is better","values":[0,0,0,0,0,0],"isMain":false},{"metricName":" # interspecies translocations","quality":"Less is better","values":[7,2,3,4,8,1],"isMain":false},{"metricName":"# possibly misassembled contigs","quality":"Less is better","values":[0,0,0,1,0,0],"isMain":false},{"metricName":" # possible misassemblies","quality":"Less is better","values":[0,0,0,1,0,0],"isMain":false},{"metricName":"# local misassemblies","quality":"Less is better","values":[0,0,0,0,1,0],"isMain":false},{"metricName":"# scaffold gap ext. mis.","quality":"Less is better","values":[0,0,0,0,0,0],"isMain":false},{"metricName":"# scaffold gap loc. mis.","quality":"Less is better","values":[0,0,0,0,0,0],"isMain":false}]],["Unaligned",[]],["Mismatches",[]],["Statistics without reference",[]],["K-mer-based statistics",[]],["Predicted genes",[]],["Similarity statistics",[]],["Reference statistics",[]]],[["Genome statistics",[]],["Reads mapping",[]],["Misassemblies",[{"metricName":"# misassemblies","quality":"Less is better","values":[3,5,2,3,5,0],"isMain":true},{"metricName":" # relocations","quality":"Less is better","values":[0,0,0,0,0,0],"isMain":false},{"metricName":" # translocations","quality":"Less is better","values":[0,0,0,0,0,0],"isMain":false},{"metricName":" # inversions","quality":"Less is better","values":[0,0,0,0,0,0],"isMain":false},{"metricName":" # interspecies translocations","quality":"Less is better","values":[3,5,2,3,5,0],"isMain":false},{"metricName":"# possibly misassembled contigs","quality":"Less is better","values":[0,0,0,0,0,0],"isMain":false},{"metricName":" # possible misassemblies","quality":"Less is better","values":[0,0,0,0,0,0],"isMain":false},{"metricName":"# local misassemblies","quality":"Less is better","values":[0,0,0,0,0,0],"isMain":false},{"metricName":"# scaffold gap ext. mis.","quality":"Less is better","values":[0,0,0,0,0,0],"isMain":false},{"metricName":"# scaffold gap loc. mis.","quality":"Less is better","values":[0,0,0,0,0,0],"isMain":false}]],["Unaligned",[]],["Mismatches",[]],["Statistics without reference",[]],["K-mer-based statistics",[]],["Predicted genes",[]],["Similarity statistics",[]],["Reference statistics",[]]],[["Genome statistics",[]],["Reads mapping",[]],["Misassemblies",[{"metricName":"# misassemblies","quality":"Less is better","values":[37,143,68,133,57,6],"isMain":true},{"metricName":" # relocations","quality":"Less is better","values":[1,2,3,6,4,0],"isMain":false},{"metricName":" # translocations","quality":"Less is better","values":[0,14,11,28,9,5],"isMain":false},{"metricName":" # inversions","quality":"Less is better","values":[4,30,7,10,23,0],"isMain":false},{"metricName":" # interspecies translocations","quality":"Less is better","values":[32,97,47,89,21,1],"isMain":false},{"metricName":"# possibly misassembled contigs","quality":"Less is better","values":[0,0,1,0,0,0],"isMain":false},{"metricName":" # possible misassemblies","quality":"Less is better","values":[0,0,1,0,0,0],"isMain":false},{"metricName":"# local misassemblies","quality":"Less is better","values":[1,11,15,16,22,0],"isMain":false},{"metricName":"# scaffold gap ext. mis.","quality":"Less is better","values":[0,0,0,0,0,0],"isMain":false},{"metricName":"# scaffold gap loc. mis.","quality":"Less is better","values":[0,0,0,0,0,0],"isMain":false}]],["Unaligned",[]],["Mismatches",[]],["Statistics without reference",[]],["K-mer-based statistics",[]],["Predicted genes",[]],["Similarity statistics",[]],["Reference statistics",[]]],[["Genome statistics",[]],["Reads mapping",[]],["Misassemblies",[{"metricName":"# misassemblies","quality":"Less is better","values":[31,29,29,54,30,2],"isMain":true},{"metricName":" # relocations","quality":"Less is better","values":[0,0,0,0,0,0],"isMain":false},{"metricName":" # translocations","quality":"Less is better","values":[1,10,10,11,9,2],"isMain":false},{"metricName":" # inversions","quality":"Less is better","values":[4,2,0,1,1,0],"isMain":false},{"metricName":" # interspecies translocations","quality":"Less is better","values":[26,17,19,42,20,0],"isMain":false},{"metricName":"# possibly misassembled contigs","quality":"Less is better","values":[0,0,0,1,1,0],"isMain":false},{"metricName":" # possible misassemblies","quality":"Less is better","values":[0,0,0,1,1,0],"isMain":false},{"metricName":"# local misassemblies","quality":"Less is better","values":[0,5,6,2,8,0],"isMain":false},{"metricName":"# scaffold gap ext. mis.","quality":"Less is better","values":[0,0,0,0,0,0],"isMain":false},{"metricName":"# scaffold gap loc. mis.","quality":"Less is better","values":[0,0,0,0,0,0],"isMain":false}]],["Unaligned",[]],["Mismatches",[]],["Statistics without reference",[]],["K-mer-based statistics",[]],["Predicted genes",[]],["Similarity statistics",[]],["Reference statistics",[]]],[["Genome statistics",[]],["Reads mapping",[]],["Misassemblies",[{"metricName":"# misassemblies","quality":"Less is better","values":[36,26,36,72,35,3],"isMain":true},{"metricName":" # relocations","quality":"Less is better","values":[0,0,0,1,0,0],"isMain":false},{"metricName":" # translocations","quality":"Less is better","values":[0,4,7,16,14,2],"isMain":false},{"metricName":" # inversions","quality":"Less is better","values":[1,2,1,2,2,0],"isMain":false},{"metricName":" # interspecies translocations","quality":"Less is better","values":[35,20,28,53,19,1],"isMain":false},{"metricName":"# possibly misassembled contigs","quality":"Less is better","values":[0,0,1,1,0,0],"isMain":false},{"metricName":" # possible misassemblies","quality":"Less is better","values":[0,0,1,1,0,0],"isMain":false},{"metricName":"# local misassemblies","quality":"Less is better","values":[1,5,8,13,6,2],"isMain":false},{"metricName":"# scaffold gap ext. mis.","quality":"Less is better","values":[0,0,0,0,0,0],"isMain":false},{"metricName":"# scaffold gap loc. mis.","quality":"Less is better","values":[0,0,0,0,0,0],"isMain":false}]],["Unaligned",[]],["Mismatches",[]],["Statistics without reference",[]],["K-mer-based statistics",[]],["Predicted genes",[]],["Similarity statistics",[]],["Reference statistics",[]]],[["Genome statistics",[]],["Reads mapping",[]],["Misassemblies",[{"metricName":"# misassemblies","quality":"Less is better","values":[3,2,3,12,3,0],"isMain":true},{"metricName":" # relocations","quality":"Less is better","values":[0,0,0,0,0,0],"isMain":false},{"metricName":" # translocations","quality":"Less is better","values":[0,0,0,0,0,0],"isMain":false},{"metricName":" # inversions","quality":"Less is better","values":[0,0,0,0,0,0],"isMain":false},{"metricName":" # interspecies translocations","quality":"Less is better","values":[3,2,3,12,3,0],"isMain":false},{"metricName":"# possibly misassembled contigs","quality":"Less is better","values":[0,0,0,0,0,0],"isMain":false},{"metricName":" # possible misassemblies","quality":"Less is better","values":[0,0,0,0,0,0],"isMain":false},{"metricName":"# local misassemblies","quality":"Less is better","values":[0,0,0,0,0,0],"isMain":false},{"metricName":"# scaffold gap ext. mis.","quality":"Less is better","values":[0,0,0,0,0,0],"isMain":false},{"metricName":"# scaffold gap loc. mis.","quality":"Less is better","values":[0,0,0,0,0,0],"isMain":false}]],["Unaligned",[]],["Mismatches",[]],["Statistics without reference",[]],["K-mer-based statistics",[]],["Predicted genes",[]],["Similarity statistics",[]],["Reference statistics",[]]],[["Genome statistics",[]],["Reads mapping",[]],["Misassemblies",[{"metricName":"# misassemblies","quality":"Less is better","values":[0,7,2,15,7,2],"isMain":true},{"metricName":" # relocations","quality":"Less is better","values":[0,0,0,0,0,0],"isMain":false},{"metricName":" # translocations","quality":"Less is better","values":[0,0,0,0,0,0],"isMain":false},{"metricName":" # inversions","quality":"Less is better","values":[0,0,0,0,0,0],"isMain":false},{"metricName":" # interspecies translocations","quality":"Less is better","values":[0,7,2,15,7,2],"isMain":false},{"metricName":"# possibly misassembled contigs","quality":"Less is better","values":[0,0,0,0,0,0],"isMain":false},{"metricName":" # possible misassemblies","quality":"Less is better","values":[0,0,0,0,0,0],"isMain":false},{"metricName":"# local misassemblies","quality":"Less is better","values":[0,5,1,3,4,0],"isMain":false},{"metricName":"# scaffold gap ext. mis.","quality":"Less is better","values":[0,0,0,0,0,0],"isMain":false},{"metricName":"# scaffold gap loc. mis.","quality":"Less is better","values":[0,0,0,0,0,0],"isMain":false}]],["Unaligned",[]],["Mismatches",[]],["Statistics without reference",[]],["K-mer-based statistics",[]],["Predicted genes",[]],["Similarity statistics",[]],["Reference statistics",[]]],[["Genome statistics",[]],["Reads mapping",[]],["Misassemblies",[{"metricName":"# misassemblies","quality":"Less is better","values":[2,3,1,7,4,2],"isMain":true},{"metricName":" # relocations","quality":"Less is better","values":[1,0,0,0,0,0],"isMain":false},{"metricName":" # translocations","quality":"Less is better","values":[0,0,0,0,0,0],"isMain":false},{"metricName":" # inversions","quality":"Less is better","values":[0,0,0,0,0,0],"isMain":false},{"metricName":" # interspecies translocations","quality":"Less is better","values":[1,3,1,7,4,2],"isMain":false},{"metricName":"# possibly misassembled contigs","quality":"Less is better","values":[0,0,0,0,0,0],"isMain":false},{"metricName":" # possible misassemblies","quality":"Less is better","values":[0,0,0,0,0,0],"isMain":false},{"metricName":"# local misassemblies","quality":"Less is better","values":[0,0,0,0,0,0],"isMain":false},{"metricName":"# scaffold gap ext. mis.","quality":"Less is better","values":[0,0,0,0,0,0],"isMain":false},{"metricName":"# scaffold gap loc. mis.","quality":"Less is better","values":[0,0,0,0,0,0],"isMain":false}]],["Unaligned",[]],["Mismatches",[]],["Statistics without reference",[]],["K-mer-based statistics",[]],["Predicted genes",[]],["Similarity statistics",[]],["Reference statistics",[]]],[["Genome statistics",[]],["Reads mapping",[]],["Misassemblies",[{"metricName":"# misassemblies","quality":"Less is better","values":[37,50,14,30,38,32],"isMain":true},{"metricName":" # relocations","quality":"Less is better","values":[0,0,0,0,1,0],"isMain":false},{"metricName":" # translocations","quality":"Less is better","values":[7,9,2,4,5,10],"isMain":false},{"metricName":" # inversions","quality":"Less is better","values":[1,0,2,2,1,0],"isMain":false},{"metricName":" # interspecies translocations","quality":"Less is better","values":[29,41,10,24,31,22],"isMain":false},{"metricName":"# possibly misassembled contigs","quality":"Less is better","values":[0,0,0,0,0,0],"isMain":false},{"metricName":" # possible misassemblies","quality":"Less is better","values":[0,0,0,0,0,0],"isMain":false},{"metricName":"# local misassemblies","quality":"Less is better","values":[0,0,2,3,2,0],"isMain":false},{"metricName":"# scaffold gap ext. mis.","quality":"Less is better","values":[0,0,0,0,0,0],"isMain":false},{"metricName":"# scaffold gap loc. mis.","quality":"Less is better","values":[0,0,0,0,0,0],"isMain":false}]],["Unaligned",[]],["Mismatches",[]],["Statistics without reference",[]],["K-mer-based statistics",[]],["Predicted genes",[]],["Similarity statistics",[]],["Reference statistics",[]]],[["Genome statistics",[]],["Reads mapping",[]],["Misassemblies",[{"metricName":"# misassemblies","quality":"Less is better","values":[17,18,6,18,22,6],"isMain":true},{"metricName":" # relocations","quality":"Less is better","values":[0,0,0,0,0,0],"isMain":false},{"metricName":" # translocations","quality":"Less is better","values":[1,1,0,3,0,0],"isMain":false},{"metricName":" # inversions","quality":"Less is better","values":[0,0,0,0,0,0],"isMain":false},{"metricName":" # interspecies translocations","quality":"Less is better","values":[16,17,6,15,22,6],"isMain":false},{"metricName":"# possibly misassembled contigs","quality":"Less is better","values":[0,0,0,0,0,0],"isMain":false},{"metricName":" # possible misassemblies","quality":"Less is better","values":[0,0,0,0,0,0],"isMain":false},{"metricName":"# local misassemblies","quality":"Less is better","values":[2,1,0,2,2,0],"isMain":false},{"metricName":"# scaffold gap ext. mis.","quality":"Less is better","values":[0,0,0,0,0,0],"isMain":false},{"metricName":"# scaffold gap loc. mis.","quality":"Less is better","values":[0,0,0,0,0,0],"isMain":false}]],["Unaligned",[]],["Mismatches",[]],["Statistics without reference",[]],["K-mer-based statistics",[]],["Predicted genes",[]],["Similarity statistics",[]],["Reference statistics",[]]],[["Genome statistics",[]],["Reads mapping",[]],["Misassemblies",[{"metricName":"# misassemblies","quality":"Less is better","values":[17,14,10,15,15,9],"isMain":true},{"metricName":" # relocations","quality":"Less is better","values":[1,2,1,2,1,0],"isMain":false},{"metricName":" # translocations","quality":"Less is better","values":[0,0,0,0,0,0],"isMain":false},{"metricName":" # inversions","quality":"Less is better","values":[0,1,0,0,1,0],"isMain":false},{"metricName":" # interspecies translocations","quality":"Less is better","values":[16,11,9,13,13,9],"isMain":false},{"metricName":"# possibly misassembled contigs","quality":"Less is better","values":[0,0,0,0,1,0],"isMain":false},{"metricName":" # possible misassemblies","quality":"Less is better","values":[0,0,0,0,1,0],"isMain":false},{"metricName":"# local misassemblies","quality":"Less is better","values":[0,0,1,1,0,1],"isMain":false},{"metricName":"# scaffold gap ext. mis.","quality":"Less is better","values":[0,0,0,0,0,0],"isMain":false},{"metricName":"# scaffold gap loc. mis.","quality":"Less is better","values":[0,0,0,0,0,0],"isMain":false}]],["Unaligned",[]],["Mismatches",[]],["Statistics without reference",[]],["K-mer-based statistics",[]],["Predicted genes",[]],["Similarity statistics",[]],["Reference statistics",[]]],[["Genome statistics",[]],["Reads mapping",[]],["Misassemblies",[{"metricName":"# misassemblies","quality":"Less is better","values":[0,4,6,3,3,0],"isMain":true},{"metricName":" # relocations","quality":"Less is better","values":[0,0,0,0,0,0],"isMain":false},{"metricName":" # translocations","quality":"Less is better","values":[0,0,0,0,0,0],"isMain":false},{"metricName":" # inversions","quality":"Less is better","values":[0,0,0,0,0,0],"isMain":false},{"metricName":" # interspecies translocations","quality":"Less is better","values":[0,4,6,3,3,0],"isMain":false},{"metricName":"# possibly misassembled contigs","quality":"Less is better","values":[0,0,0,0,0,0],"isMain":false},{"metricName":" # possible misassemblies","quality":"Less is better","values":[0,0,0,0,0,0],"isMain":false},{"metricName":"# local misassemblies","quality":"Less is better","values":[0,0,0,0,0,0],"isMain":false},{"metricName":"# scaffold gap ext. mis.","quality":"Less is better","values":[0,0,0,0,0,0],"isMain":false},{"metricName":"# scaffold gap loc. mis.","quality":"Less is better","values":[0,0,0,0,0,0],"isMain":false}]],["Unaligned",[]],["Mismatches",[]],["Statistics without reference",[]],["K-mer-based statistics",[]],["Predicted genes",[]],["Similarity statistics",[]],["Reference statistics",[]]],[["Genome statistics",[]],["Reads mapping",[]],["Misassemblies",[{"metricName":"# misassemblies","quality":"Less is better","values":[11,55,44,60,57,6],"isMain":true},{"metricName":" # relocations","quality":"Less is better","values":[2,3,4,8,6,1],"isMain":false},{"metricName":" # translocations","quality":"Less is better","values":[0,4,1,2,3,0],"isMain":false},{"metricName":" # inversions","quality":"Less is better","values":[0,0,0,0,0,0],"isMain":false},{"metricName":" # interspecies translocations","quality":"Less is better","values":[9,48,39,50,48,5],"isMain":false},{"metricName":"# possibly misassembled contigs","quality":"Less is better","values":[0,0,0,0,0,0],"isMain":false},{"metricName":" # possible misassemblies","quality":"Less is better","values":[0,0,0,0,0,0],"isMain":false},{"metricName":"# local misassemblies","quality":"Less is better","values":[1,2,5,4,3,0],"isMain":false},{"metricName":"# scaffold gap ext. mis.","quality":"Less is better","values":[0,0,0,0,0,0],"isMain":false},{"metricName":"# scaffold gap loc. mis.","quality":"Less is better","values":[0,0,0,0,0,0],"isMain":false}]],["Unaligned",[]],["Mismatches",[]],["Statistics without reference",[]],["K-mer-based statistics",[]],["Predicted genes",[]],["Similarity statistics",[]],["Reference statistics",[]]],[["Genome statistics",[]],["Reads mapping",[]],["Misassemblies",[{"metricName":"# misassemblies","quality":"Less is better","values":[5,14,12,18,14,4],"isMain":true},{"metricName":" # relocations","quality":"Less is better","values":[0,0,0,0,1,0],"isMain":false},{"metricName":" # translocations","quality":"Less is better","values":[0,0,0,0,0,0],"isMain":false},{"metricName":" # inversions","quality":"Less is better","values":[0,0,0,0,0,0],"isMain":false},{"metricName":" # interspecies translocations","quality":"Less is better","values":[5,14,12,18,13,4],"isMain":false},{"metricName":"# possibly misassembled contigs","quality":"Less is better","values":[0,0,0,0,0,0],"isMain":false},{"metricName":" # possible misassemblies","quality":"Less is better","values":[0,0,0,0,0,0],"isMain":false},{"metricName":"# local misassemblies","quality":"Less is better","values":[0,0,0,0,0,0],"isMain":false},{"metricName":"# scaffold gap ext. mis.","quality":"Less is better","values":[0,0,0,0,0,0],"isMain":false},{"metricName":"# scaffold gap loc. mis.","quality":"Less is better","values":[0,0,0,0,0,0],"isMain":false}]],["Unaligned",[]],["Mismatches",[]],["Statistics without reference",[]],["K-mer-based statistics",[]],["Predicted genes",[]],["Similarity statistics",[]],["Reference statistics",[]]],[["Genome statistics",[]],["Reads mapping",[]],["Misassemblies",[{"metricName":"# misassemblies","quality":"Less is better","values":[58,88,43,75,97,25],"isMain":true},{"metricName":" # relocations","quality":"Less is better","values":[1,1,1,1,1,2],"isMain":false},{"metricName":" # translocations","quality":"Less is better","values":[5,6,6,8,9,2],"isMain":false},{"metricName":" # inversions","quality":"Less is better","values":[1,1,2,0,3,0],"isMain":false},{"metricName":" # interspecies translocations","quality":"Less is better","values":[51,80,34,66,84,21],"isMain":false},{"metricName":"# possibly misassembled contigs","quality":"Less is better","values":[0,0,1,0,0,0],"isMain":false},{"metricName":" # possible misassemblies","quality":"Less is better","values":[0,0,1,0,0,0],"isMain":false},{"metricName":"# local misassemblies","quality":"Less is better","values":[3,2,1,3,1,2],"isMain":false},{"metricName":"# scaffold gap ext. mis.","quality":"Less is better","values":[0,0,0,0,0,0],"isMain":false},{"metricName":"# scaffold gap loc. mis.","quality":"Less is better","values":[0,0,0,0,0,0],"isMain":false}]],["Unaligned",[]],["Mismatches",[]],["Statistics without reference",[]],["K-mer-based statistics",[]],["Predicted genes",[]],["Similarity statistics",[]],["Reference statistics",[]]],[["Genome statistics",[]],["Reads mapping",[]],["Misassemblies",[{"metricName":"# misassemblies","quality":"Less is better","values":[33,38,25,40,56,17],"isMain":true},{"metricName":" # relocations","quality":"Less is better","values":[0,0,0,0,0,1],"isMain":false},{"metricName":" # translocations","quality":"Less is better","values":[3,4,7,3,6,5],"isMain":false},{"metricName":" # inversions","quality":"Less is better","values":[0,1,0,1,0,0],"isMain":false},{"metricName":" # interspecies translocations","quality":"Less is better","values":[30,33,18,36,50,11],"isMain":false},{"metricName":"# possibly misassembled contigs","quality":"Less is better","values":[1,2,2,0,0,0],"isMain":false},{"metricName":" # possible misassemblies","quality":"Less is better","values":[1,2,2,0,0,0],"isMain":false},{"metricName":"# local misassemblies","quality":"Less is better","values":[0,4,1,1,1,1],"isMain":false},{"metricName":"# scaffold gap ext. mis.","quality":"Less is better","values":[0,0,0,0,0,0],"isMain":false},{"metricName":"# scaffold gap loc. mis.","quality":"Less is better","values":[0,0,0,0,0,0],"isMain":false}]],["Unaligned",[]],["Mismatches",[]],["Statistics without reference",[]],["K-mer-based statistics",[]],["Predicted genes",[]],["Similarity statistics",[]],["Reference statistics",[]]],[["Genome statistics",[]],["Reads mapping",[]],["Misassemblies",[{"metricName":"# misassemblies","quality":"Less is better","values":[3,11,8,19,13,1],"isMain":true},{"metricName":" # relocations","quality":"Less is better","values":[0,0,0,0,0,0],"isMain":false},{"metricName":" # translocations","quality":"Less is better","values":[0,0,0,1,2,0],"isMain":false},{"metricName":" # inversions","quality":"Less is better","values":[0,0,1,0,0,0],"isMain":false},{"metricName":" # interspecies translocations","quality":"Less is better","values":[3,11,7,18,11,1],"isMain":false},{"metricName":"# possibly misassembled contigs","quality":"Less is better","values":[0,1,0,2,0,0],"isMain":false},{"metricName":" # possible misassemblies","quality":"Less is better","values":[0,1,0,2,0,0],"isMain":false},{"metricName":"# local misassemblies","quality":"Less is better","values":[0,0,0,1,1,0],"isMain":false},{"metricName":"# scaffold gap ext. mis.","quality":"Less is better","values":[0,0,0,0,0,0],"isMain":false},{"metricName":"# scaffold gap loc. mis.","quality":"Less is better","values":[0,0,0,0,0,0],"isMain":false}]],["Unaligned",[]],["Mismatches",[]],["Statistics without reference",[]],["K-mer-based statistics",[]],["Predicted genes",[]],["Similarity statistics",[]],["Reference statistics",[]]],[["Genome statistics",[]],["Reads mapping",[]],["Misassemblies",[{"metricName":"# misassemblies","quality":"Less is better","values":[13,35,36,48,70,1],"isMain":true},{"metricName":" # relocations","quality":"Less is better","values":[0,1,2,2,1,0],"isMain":false},{"metricName":" # translocations","quality":"Less is better","values":[3,5,4,4,5,0],"isMain":false},{"metricName":" # inversions","quality":"Less is better","values":[0,0,0,1,0,0],"isMain":false},{"metricName":" # interspecies translocations","quality":"Less is better","values":[10,29,30,41,64,1],"isMain":false},{"metricName":"# possibly misassembled contigs","quality":"Less is better","values":[0,1,0,0,2,0],"isMain":false},{"metricName":" # possible misassemblies","quality":"Less is better","values":[0,1,0,0,2,0],"isMain":false},{"metricName":"# local misassemblies","quality":"Less is better","values":[1,0,3,4,5,1],"isMain":false},{"metricName":"# scaffold gap ext. mis.","quality":"Less is better","values":[0,0,0,0,0,0],"isMain":false},{"metricName":"# scaffold gap loc. mis.","quality":"Less is better","values":[0,0,0,0,0,0],"isMain":false}]],["Unaligned",[]],["Mismatches",[]],["Statistics without reference",[]],["K-mer-based statistics",[]],["Predicted genes",[]],["Similarity statistics",[]],["Reference statistics",[]]],[["Genome statistics",[]],["Reads mapping",[]],["Misassemblies",[{"metricName":"# misassemblies","quality":"Less is better","values":[3,3,4,5,8,1],"isMain":true},{"metricName":" # relocations","quality":"Less is better","values":[0,0,0,0,0,0],"isMain":false},{"metricName":" # translocations","quality":"Less is better","values":[0,0,0,0,0,0],"isMain":false},{"metricName":" # inversions","quality":"Less is better","values":[0,0,0,0,1,0],"isMain":false},{"metricName":" # interspecies translocations","quality":"Less is better","values":[3,3,4,5,7,1],"isMain":false},{"metricName":"# possibly misassembled contigs","quality":"Less is better","values":[0,0,0,1,0,0],"isMain":false},{"metricName":" # possible misassemblies","quality":"Less is better","values":[0,0,0,1,0,0],"isMain":false},{"metricName":"# local misassemblies","quality":"Less is better","values":[0,0,0,0,0,0],"isMain":false},{"metricName":"# scaffold gap ext. mis.","quality":"Less is better","values":[0,0,0,0,0,0],"isMain":false},{"metricName":"# scaffold gap loc. mis.","quality":"Less is better","values":[0,0,0,0,0,0],"isMain":false}]],["Unaligned",[]],["Mismatches",[]],["Statistics without reference",[]],["K-mer-based statistics",[]],["Predicted genes",[]],["Similarity statistics",[]],["Reference statistics",[]]],[["Genome statistics",[]],["Reads mapping",[]],["Misassemblies",[{"metricName":"# misassemblies","quality":"Less is better","values":[40,36,51,71,41,3],"isMain":true},{"metricName":" # relocations","quality":"Less is better","values":[0,1,6,11,6,1],"isMain":false},{"metricName":" # translocations","quality":"Less is better","values":[0,3,0,2,4,0],"isMain":false},{"metricName":" # inversions","quality":"Less is better","values":[0,3,1,0,3,0],"isMain":false},{"metricName":" # interspecies translocations","quality":"Less is better","values":[40,29,44,58,28,2],"isMain":false},{"metricName":"# possibly misassembled contigs","quality":"Less is better","values":[0,0,1,0,0,0],"isMain":false},{"metricName":" # possible misassemblies","quality":"Less is better","values":[0,0,1,0,0,0],"isMain":false},{"metricName":"# local misassemblies","quality":"Less is better","values":[1,9,17,9,11,1],"isMain":false},{"metricName":"# scaffold gap ext. mis.","quality":"Less is better","values":[0,0,0,0,0,0],"isMain":false},{"metricName":"# scaffold gap loc. mis.","quality":"Less is better","values":[0,0,0,0,0,0],"isMain":false}]],["Unaligned",[]],["Mismatches",[]],["Statistics without reference",[]],["K-mer-based statistics",[]],["Predicted genes",[]],["Similarity statistics",[]],["Reference statistics",[]]],[["Genome statistics",[]],["Reads mapping",[]],["Misassemblies",[{"metricName":"# misassemblies","quality":"Less is better","values":[4,8,5,4,5,0],"isMain":true},{"metricName":" # relocations","quality":"Less is better","values":[0,0,0,0,0,0],"isMain":false},{"metricName":" # translocations","quality":"Less is better","values":[0,0,0,0,0,0],"isMain":false},{"metricName":" # inversions","quality":"Less is better","values":[0,0,0,0,0,0],"isMain":false},{"metricName":" # interspecies translocations","quality":"Less is better","values":[4,8,5,4,5,0],"isMain":false},{"metricName":"# possibly misassembled contigs","quality":"Less is better","values":[0,0,0,0,0,0],"isMain":false},{"metricName":" # possible misassemblies","quality":"Less is better","values":[0,0,0,0,0,0],"isMain":false},{"metricName":"# local misassemblies","quality":"Less is better","values":[0,0,0,0,0,0],"isMain":false},{"metricName":"# scaffold gap ext. mis.","quality":"Less is better","values":[0,0,0,0,0,0],"isMain":false},{"metricName":"# scaffold gap loc. mis.","quality":"Less is better","values":[0,0,0,0,0,0],"isMain":false}]],["Unaligned",[]],["Mismatches",[]],["Statistics without reference",[]],["K-mer-based statistics",[]],["Predicted genes",[]],["Similarity statistics",[]],["Reference statistics",[]]],[["Genome statistics",[]],["Reads mapping",[]],["Misassemblies",[{"metricName":"# misassemblies","quality":"Less is better","values":[5,8,11,18,15,0],"isMain":true},{"metricName":" # relocations","quality":"Less is better","values":[0,0,1,0,2,0],"isMain":false},{"metricName":" # translocations","quality":"Less is better","values":[0,0,0,0,0,0],"isMain":false},{"metricName":" # inversions","quality":"Less is better","values":[1,0,0,0,0,0],"isMain":false},{"metricName":" # interspecies translocations","quality":"Less is better","values":[4,8,10,18,13,0],"isMain":false},{"metricName":"# possibly misassembled contigs","quality":"Less is better","values":[0,0,0,0,0,0],"isMain":false},{"metricName":" # possible misassemblies","quality":"Less is better","values":[0,0,0,0,0,0],"isMain":false},{"metricName":"# local misassemblies","quality":"Less is better","values":[0,0,0,0,1,0],"isMain":false},{"metricName":"# scaffold gap ext. mis.","quality":"Less is better","values":[0,0,0,0,0,0],"isMain":false},{"metricName":"# scaffold gap loc. mis.","quality":"Less is better","values":[0,0,0,0,0,0],"isMain":false}]],["Unaligned",[]],["Mismatches",[]],["Statistics without reference",[]],["K-mer-based statistics",[]],["Predicted genes",[]],["Similarity statistics",[]],["Reference statistics",[]]],[["Genome statistics",[]],["Reads mapping",[]],["Misassemblies",[{"metricName":"# misassemblies","quality":"Less is better","values":[1,5,7,4,8,0],"isMain":true},{"metricName":" # relocations","quality":"Less is better","values":[0,0,1,0,0,0],"isMain":false},{"metricName":" # translocations","quality":"Less is better","values":[0,0,0,0,0,0],"isMain":false},{"metricName":" # inversions","quality":"Less is better","values":[0,0,0,0,0,0],"isMain":false},{"metricName":" # interspecies translocations","quality":"Less is better","values":[1,5,6,4,8,0],"isMain":false},{"metricName":"# possibly misassembled contigs","quality":"Less is better","values":[0,0,0,0,0,0],"isMain":false},{"metricName":" # possible misassemblies","quality":"Less is better","values":[0,0,0,0,0,0],"isMain":false},{"metricName":"# local misassemblies","quality":"Less is better","values":[0,0,0,1,0,0],"isMain":false},{"metricName":"# scaffold gap ext. mis.","quality":"Less is better","values":[0,0,0,0,0,0],"isMain":false},{"metricName":"# scaffold gap loc. mis.","quality":"Less is better","values":[0,0,0,0,0,0],"isMain":false}]],["Unaligned",[]],["Mismatches",[]],["Statistics without reference",[]],["K-mer-based statistics",[]],["Predicted genes",[]],["Similarity statistics",[]],["Reference statistics",[]]],[["Genome statistics",[]],["Reads mapping",[]],["Misassemblies",[{"metricName":"# misassemblies","quality":"Less is better","values":[0,4,2,0,11,2],"isMain":true},{"metricName":" # relocations","quality":"Less is better","values":[0,0,0,0,0,0],"isMain":false},{"metricName":" # translocations","quality":"Less is better","values":[0,0,0,0,0,0],"isMain":false},{"metricName":" # inversions","quality":"Less is better","values":[0,0,0,0,0,0],"isMain":false},{"metricName":" # interspecies translocations","quality":"Less is better","values":[0,4,2,0,11,2],"isMain":false},{"metricName":"# possibly misassembled contigs","quality":"Less is better","values":[0,0,0,0,0,0],"isMain":false},{"metricName":" # possible misassemblies","quality":"Less is better","values":[0,0,0,0,0,0],"isMain":false},{"metricName":"# local misassemblies","quality":"Less is better","values":[0,0,0,0,0,0],"isMain":false},{"metricName":"# scaffold gap ext. mis.","quality":"Less is better","values":[0,0,0,0,0,0],"isMain":false},{"metricName":"# scaffold gap loc. mis.","quality":"Less is better","values":[0,0,0,0,0,0],"isMain":false}]],["Unaligned",[]],["Mismatches",[]],["Statistics without reference",[]],["K-mer-based statistics",[]],["Predicted genes",[]],["Similarity statistics",[]],["Reference statistics",[]]],[["Genome statistics",[]],["Reads mapping",[]],["Misassemblies",[{"metricName":"# misassemblies","quality":"Less is better","values":[5,0,3,1,7,0],"isMain":true},{"metricName":" # relocations","quality":"Less is better","values":[1,0,0,0,0,0],"isMain":false},{"metricName":" # translocations","quality":"Less is better","values":[0,0,0,0,0,0],"isMain":false},{"metricName":" # inversions","quality":"Less is better","values":[1,0,0,0,0,0],"isMain":false},{"metricName":" # interspecies translocations","quality":"Less is better","values":[3,0,3,1,7,0],"isMain":false},{"metricName":"# possibly misassembled contigs","quality":"Less is better","values":[0,0,0,0,0,0],"isMain":false},{"metricName":" # possible misassemblies","quality":"Less is better","values":[0,0,0,0,0,0],"isMain":false},{"metricName":"# local misassemblies","quality":"Less is better","values":[0,0,0,1,1,0],"isMain":false},{"metricName":"# scaffold gap ext. mis.","quality":"Less is better","values":[0,0,0,0,0,0],"isMain":false},{"metricName":"# scaffold gap loc. mis.","quality":"Less is better","values":[0,0,0,0,0,0],"isMain":false}]],["Unaligned",[]],["Mismatches",[]],["Statistics without reference",[]],["K-mer-based statistics",[]],["Predicted genes",[]],["Similarity statistics",[]],["Reference statistics",[]]],[["Genome statistics",[]],["Reads mapping",[]],["Misassemblies",[{"metricName":"# misassemblies","quality":"Less is better","values":[8,8,5,8,19,3],"isMain":true},{"metricName":" # relocations","quality":"Less is better","values":[0,0,0,0,0,0],"isMain":false},{"metricName":" # translocations","quality":"Less is better","values":[0,0,0,0,0,0],"isMain":false},{"metricName":" # inversions","quality":"Less is better","values":[0,0,0,0,1,0],"isMain":false},{"metricName":" # interspecies translocations","quality":"Less is better","values":[8,8,5,8,18,3],"isMain":false},{"metricName":"# possibly misassembled contigs","quality":"Less is better","values":[0,0,0,0,0,0],"isMain":false},{"metricName":" # possible misassemblies","quality":"Less is better","values":[0,0,0,0,0,0],"isMain":false},{"metricName":"# local misassemblies","quality":"Less is better","values":[0,3,0,3,2,1],"isMain":false},{"metricName":"# scaffold gap ext. mis.","quality":"Less is better","values":[0,0,0,0,0,0],"isMain":false},{"metricName":"# scaffold gap loc. mis.","quality":"Less is better","values":[0,0,0,0,0,0],"isMain":false}]],["Unaligned",[]],["Mismatches",[]],["Statistics without reference",[]],["K-mer-based statistics",[]],["Predicted genes",[]],["Similarity statistics",[]],["Reference statistics",[]]],[["Genome statistics",[]],["Reads mapping",[]],["Misassemblies",[{"metricName":"# misassemblies","quality":"Less is better","values":[17,11,15,17,21,3],"isMain":true},{"metricName":" # relocations","quality":"Less is better","values":[1,2,1,4,3,0],"isMain":false},{"metricName":" # translocations","quality":"Less is better","values":[0,0,0,0,0,0],"isMain":false},{"metricName":" # inversions","quality":"Less is better","values":[1,0,0,2,0,0],"isMain":false},{"metricName":" # interspecies translocations","quality":"Less is better","values":[15,9,14,11,18,3],"isMain":false},{"metricName":"# possibly misassembled contigs","quality":"Less is better","values":[0,0,0,0,0,0],"isMain":false},{"metricName":" # possible misassemblies","quality":"Less is better","values":[0,0,0,0,0,0],"isMain":false},{"metricName":"# local misassemblies","quality":"Less is better","values":[2,3,2,1,3,0],"isMain":false},{"metricName":"# scaffold gap ext. mis.","quality":"Less is better","values":[0,0,0,0,0,0],"isMain":false},{"metricName":"# scaffold gap loc. mis.","quality":"Less is better","values":[0,0,0,0,0,0],"isMain":false}]],["Unaligned",[]],["Mismatches",[]],["Statistics without reference",[]],["K-mer-based statistics",[]],["Predicted genes",[]],["Similarity statistics",[]],["Reference statistics",[]]],[["Genome statistics",[]],["Reads mapping",[]],["Misassemblies",[{"metricName":"# misassemblies","quality":"Less is better","values":[7,14,10,19,19,3],"isMain":true},{"metricName":" # relocations","quality":"Less is better","values":[1,0,1,0,1,0],"isMain":false},{"metricName":" # translocations","quality":"Less is better","values":[0,0,0,0,0,0],"isMain":false},{"metricName":" # inversions","quality":"Less is better","values":[1,0,0,0,0,0],"isMain":false},{"metricName":" # interspecies translocations","quality":"Less is better","values":[5,14,9,19,18,3],"isMain":false},{"metricName":"# possibly misassembled contigs","quality":"Less is better","values":[0,1,0,1,0,0],"isMain":false},{"metricName":" # possible misassemblies","quality":"Less is better","values":[0,1,0,1,0,0],"isMain":false},{"metricName":"# local misassemblies","quality":"Less is better","values":[0,0,0,0,0,0],"isMain":false},{"metricName":"# scaffold gap ext. mis.","quality":"Less is better","values":[0,0,0,0,0,0],"isMain":false},{"metricName":"# scaffold gap loc. mis.","quality":"Less is better","values":[0,0,0,0,0,0],"isMain":false}]],["Unaligned",[]],["Mismatches",[]],["Statistics without reference",[]],["K-mer-based statistics",[]],["Predicted genes",[]],["Similarity statistics",[]],["Reference statistics",[]]],[["Genome statistics",[]],["Reads mapping",[]],["Misassemblies",[{"metricName":"# misassemblies","quality":"Less is better","values":[8,9,9,12,11,1],"isMain":true},{"metricName":" # relocations","quality":"Less is better","values":[0,1,0,0,0,0],"isMain":false},{"metricName":" # translocations","quality":"Less is better","values":[1,0,0,0,0,0],"isMain":false},{"metricName":" # inversions","quality":"Less is better","values":[0,0,0,0,0,0],"isMain":false},{"metricName":" # interspecies translocations","quality":"Less is better","values":[7,8,9,12,11,1],"isMain":false},{"metricName":"# possibly misassembled contigs","quality":"Less is better","values":[0,0,0,0,1,0],"isMain":false},{"metricName":" # possible misassemblies","quality":"Less is better","values":[0,0,0,0,1,0],"isMain":false},{"metricName":"# local misassemblies","quality":"Less is better","values":[0,0,1,0,4,0],"isMain":false},{"metricName":"# scaffold gap ext. mis.","quality":"Less is better","values":[0,0,0,0,0,0],"isMain":false},{"metricName":"# scaffold gap loc. mis.","quality":"Less is better","values":[0,0,0,0,0,0],"isMain":false}]],["Unaligned",[]],["Mismatches",[]],["Statistics without reference",[]],["K-mer-based statistics",[]],["Predicted genes",[]],["Similarity statistics",[]],["Reference statistics",[]]],[["Genome statistics",[]],["Reads mapping",[]],["Misassemblies",[{"metricName":"# misassemblies","quality":"Less is better","values":[15,25,30,29,35,2],"isMain":true},{"metricName":" # relocations","quality":"Less is better","values":[3,3,3,1,2,0],"isMain":false},{"metricName":" # translocations","quality":"Less is better","values":[0,0,0,0,0,0],"isMain":false},{"metricName":" # inversions","quality":"Less is better","values":[0,1,2,1,2,0],"isMain":false},{"metricName":" # interspecies translocations","quality":"Less is better","values":[12,21,25,27,31,2],"isMain":false},{"metricName":"# possibly misassembled contigs","quality":"Less is better","values":[0,0,2,2,0,0],"isMain":false},{"metricName":" # possible misassemblies","quality":"Less is better","values":[0,0,2,3,0,0],"isMain":false},{"metricName":"# local misassemblies","quality":"Less is better","values":[1,2,2,4,2,0],"isMain":false},{"metricName":"# scaffold gap ext. mis.","quality":"Less is better","values":[0,0,0,0,0,0],"isMain":false},{"metricName":"# scaffold gap loc. mis.","quality":"Less is better","values":[0,0,0,0,0,0],"isMain":false}]],["Unaligned",[]],["Mismatches",[]],["Statistics without reference",[]],["K-mer-based statistics",[]],["Predicted genes",[]],["Similarity statistics",[]],["Reference statistics",[]]],[["Genome statistics",[]],["Reads mapping",[]],["Misassemblies",[{"metricName":"# misassemblies","quality":"Less is better","values":[2,1,0,6,1,2],"isMain":true},{"metricName":" # relocations","quality":"Less is better","values":[0,0,0,0,0,0],"isMain":false},{"metricName":" # translocations","quality":"Less is better","values":[0,0,0,1,0,0],"isMain":false},{"metricName":" # inversions","quality":"Less is better","values":[0,0,0,0,0,0],"isMain":false},{"metricName":" # interspecies translocations","quality":"Less is better","values":[2,1,0,5,1,2],"isMain":false},{"metricName":"# possibly misassembled contigs","quality":"Less is better","values":[0,0,0,0,0,0],"isMain":false},{"metricName":" # possible misassemblies","quality":"Less is better","values":[0,0,0,0,0,0],"isMain":false},{"metricName":"# local misassemblies","quality":"Less is better","values":[0,0,0,0,0,0],"isMain":false},{"metricName":"# scaffold gap ext. mis.","quality":"Less is better","values":[0,0,0,0,0,0],"isMain":false},{"metricName":"# scaffold gap loc. mis.","quality":"Less is better","values":[0,0,0,0,0,0],"isMain":false}]],["Unaligned",[]],["Mismatches",[]],["Statistics without reference",[]],["K-mer-based statistics",[]],["Predicted genes",[]],["Similarity statistics",[]],["Reference statistics",[]]],[["Genome statistics",[]],["Reads mapping",[]],["Misassemblies",[{"metricName":"# misassemblies","quality":"Less is better","values":[58,71,47,82,66,34],"isMain":true},{"metricName":" # relocations","quality":"Less is better","values":[1,2,2,3,4,0],"isMain":false},{"metricName":" # translocations","quality":"Less is better","values":[20,24,24,25,18,16],"isMain":false},{"metricName":" # inversions","quality":"Less is better","values":[1,3,4,5,2,0],"isMain":false},{"metricName":" # interspecies translocations","quality":"Less is better","values":[36,42,17,49,42,18],"isMain":false},{"metricName":"# possibly misassembled contigs","quality":"Less is better","values":[0,1,0,0,1,0],"isMain":false},{"metricName":" # possible misassemblies","quality":"Less is better","values":[0,1,0,0,1,0],"isMain":false},{"metricName":"# local misassemblies","quality":"Less is better","values":[7,10,7,8,18,8],"isMain":false},{"metricName":"# scaffold gap ext. mis.","quality":"Less is better","values":[0,0,0,0,0,0],"isMain":false},{"metricName":"# scaffold gap loc. mis.","quality":"Less is better","values":[0,0,0,0,0,0],"isMain":false}]],["Unaligned",[]],["Mismatches",[]],["Statistics without reference",[]],["K-mer-based statistics",[]],["Predicted genes",[]],["Similarity statistics",[]],["Reference statistics",[]]],[["Genome statistics",[]],["Reads mapping",[]],["Misassemblies",[{"metricName":"# misassemblies","quality":"Less is better","values":[12,8,3,12,5,4],"isMain":true},{"metricName":" # relocations","quality":"Less is better","values":[0,0,0,0,0,0],"isMain":false},{"metricName":" # translocations","quality":"Less is better","values":[1,2,1,3,1,2],"isMain":false},{"metricName":" # inversions","quality":"Less is better","values":[0,0,0,0,0,0],"isMain":false},{"metricName":" # interspecies translocations","quality":"Less is better","values":[11,6,2,9,4,2],"isMain":false},{"metricName":"# possibly misassembled contigs","quality":"Less is better","values":[0,0,0,0,0,0],"isMain":false},{"metricName":" # possible misassemblies","quality":"Less is better","values":[0,0,0,0,0,0],"isMain":false},{"metricName":"# local misassemblies","quality":"Less is better","values":[0,1,0,0,0,0],"isMain":false},{"metricName":"# scaffold gap ext. mis.","quality":"Less is better","values":[0,0,0,0,0,0],"isMain":false},{"metricName":"# scaffold gap loc. mis.","quality":"Less is better","values":[0,0,0,0,0,0],"isMain":false}]],["Unaligned",[]],["Mismatches",[]],["Statistics without reference",[]],["K-mer-based statistics",[]],["Predicted genes",[]],["Similarity statistics",[]],["Reference statistics",[]]],[["Genome statistics",[]],["Reads mapping",[]],["Misassemblies",[{"metricName":"# misassemblies","quality":"Less is better","values":[7,7,7,11,11,11],"isMain":true},{"metricName":" # relocations","quality":"Less is better","values":[0,0,0,0,0,0],"isMain":false},{"metricName":" # translocations","quality":"Less is better","values":[0,0,1,0,0,0],"isMain":false},{"metricName":" # inversions","quality":"Less is better","values":[0,0,1,0,0,0],"isMain":false},{"metricName":" # interspecies translocations","quality":"Less is better","values":[7,7,5,11,11,11],"isMain":false},{"metricName":"# possibly misassembled contigs","quality":"Less is better","values":[0,0,0,0,0,0],"isMain":false},{"metricName":" # possible misassemblies","quality":"Less is better","values":[0,0,0,0,0,0],"isMain":false},{"metricName":"# local misassemblies","quality":"Less is better","values":[0,0,0,0,0,0],"isMain":false},{"metricName":"# scaffold gap ext. mis.","quality":"Less is better","values":[0,0,0,0,0,0],"isMain":false},{"metricName":"# scaffold gap loc. mis.","quality":"Less is better","values":[0,0,0,0,0,0],"isMain":false}]],["Unaligned",[]],["Mismatches",[]],["Statistics without reference",[]],["K-mer-based statistics",[]],["Predicted genes",[]],["Similarity statistics",[]],["Reference statistics",[]]],[["Genome statistics",[]],["Reads mapping",[]],["Misassemblies",[{"metricName":"# misassemblies","quality":"Less is better","values":[14,14,11,20,14,11],"isMain":true},{"metricName":" # relocations","quality":"Less is better","values":[1,2,0,0,2,1],"isMain":false},{"metricName":" # translocations","quality":"Less is better","values":[3,1,2,5,1,3],"isMain":false},{"metricName":" # inversions","quality":"Less is better","values":[0,0,1,0,0,0],"isMain":false},{"metricName":" # interspecies translocations","quality":"Less is better","values":[10,11,8,15,11,7],"isMain":false},{"metricName":"# possibly misassembled contigs","quality":"Less is better","values":[0,0,0,0,0,0],"isMain":false},{"metricName":" # possible misassemblies","quality":"Less is better","values":[0,0,0,0,0,0],"isMain":false},{"metricName":"# local misassemblies","quality":"Less is better","values":[2,2,2,3,2,0],"isMain":false},{"metricName":"# scaffold gap ext. mis.","quality":"Less is better","values":[0,0,0,0,0,0],"isMain":false},{"metricName":"# scaffold gap loc. mis.","quality":"Less is better","values":[0,0,0,0,0,0],"isMain":false}]],["Unaligned",[]],["Mismatches",[]],["Statistics without reference",[]],["K-mer-based statistics",[]],["Predicted genes",[]],["Similarity statistics",[]],["Reference statistics",[]]],[["Genome statistics",[]],["Reads mapping",[]],["Misassemblies",[{"metricName":"# misassemblies","quality":"Less is better","values":[38,56,23,40,49,21],"isMain":true},{"metricName":" # relocations","quality":"Less is better","values":[5,3,0,1,3,0],"isMain":false},{"metricName":" # translocations","quality":"Less is better","values":[1,2,1,1,1,2],"isMain":false},{"metricName":" # inversions","quality":"Less is better","values":[0,0,0,1,0,0],"isMain":false},{"metricName":" # interspecies translocations","quality":"Less is better","values":[32,51,22,37,45,19],"isMain":false},{"metricName":"# possibly misassembled contigs","quality":"Less is better","values":[0,0,1,1,0,0],"isMain":false},{"metricName":" # possible misassemblies","quality":"Less is better","values":[0,0,1,1,0,0],"isMain":false},{"metricName":"# local misassemblies","quality":"Less is better","values":[0,0,1,1,0,0],"isMain":false},{"metricName":"# scaffold gap ext. mis.","quality":"Less is better","values":[0,0,0,0,0,0],"isMain":false},{"metricName":"# scaffold gap loc. mis.","quality":"Less is better","values":[0,0,0,0,0,0],"isMain":false}]],["Unaligned",[]],["Mismatches",[]],["Statistics without reference",[]],["K-mer-based statistics",[]],["Predicted genes",[]],["Similarity statistics",[]],["Reference statistics",[]]],[["Genome statistics",[]],["Reads mapping",[]],["Misassemblies",[{"metricName":"# misassemblies","quality":"Less is better","values":[8,21,11,31,37,5],"isMain":true},{"metricName":" # relocations","quality":"Less is better","values":[1,5,3,3,2,0],"isMain":false},{"metricName":" # translocations","quality":"Less is better","values":[0,0,0,0,0,0],"isMain":false},{"metricName":" # inversions","quality":"Less is better","values":[0,0,0,1,0,0],"isMain":false},{"metricName":" # interspecies translocations","quality":"Less is better","values":[7,16,8,27,35,5],"isMain":false},{"metricName":"# possibly misassembled contigs","quality":"Less is better","values":[0,1,0,0,2,0],"isMain":false},{"metricName":" # possible misassemblies","quality":"Less is better","values":[0,1,0,0,2,0],"isMain":false},{"metricName":"# local misassemblies","quality":"Less is better","values":[0,0,0,1,1,0],"isMain":false},{"metricName":"# scaffold gap ext. mis.","quality":"Less is better","values":[0,0,0,0,0,0],"isMain":false},{"metricName":"# scaffold gap loc. mis.","quality":"Less is better","values":[0,0,0,0,0,0],"isMain":false}]],["Unaligned",[]],["Mismatches",[]],["Statistics without reference",[]],["K-mer-based statistics",[]],["Predicted genes",[]],["Similarity statistics",[]],["Reference statistics",[]]],[["Genome statistics",[]],["Reads mapping",[]],["Misassemblies",[{"metricName":"# misassemblies","quality":"Less is better","values":[6,20,5,10,12,4],"isMain":true},{"metricName":" # relocations","quality":"Less is better","values":[0,0,0,2,2,0],"isMain":false},{"metricName":" # translocations","quality":"Less is better","values":[1,4,2,2,2,0],"isMain":false},{"metricName":" # inversions","quality":"Less is better","values":[0,0,0,0,0,0],"isMain":false},{"metricName":" # interspecies translocations","quality":"Less is better","values":[5,16,3,6,8,4],"isMain":false},{"metricName":"# possibly misassembled contigs","quality":"Less is better","values":[0,0,0,0,0,0],"isMain":false},{"metricName":" # possible misassemblies","quality":"Less is better","values":[0,0,0,0,0,0],"isMain":false},{"metricName":"# local misassemblies","quality":"Less is better","values":[0,0,0,0,0,1],"isMain":false},{"metricName":"# scaffold gap ext. mis.","quality":"Less is better","values":[0,0,0,0,0,0],"isMain":false},{"metricName":"# scaffold gap loc. mis.","quality":"Less is better","values":[0,0,0,0,0,0],"isMain":false}]],["Unaligned",[]],["Mismatches",[]],["Statistics without reference",[]],["K-mer-based statistics",[]],["Predicted genes",[]],["Similarity statistics",[]],["Reference statistics",[]]],[["Genome statistics",[]],["Reads mapping",[]],["Misassemblies",[{"metricName":"# misassemblies","quality":"Less is better","values":[3,7,1,10,2,3],"isMain":true},{"metricName":" # relocations","quality":"Less is better","values":[0,0,0,0,0,0],"isMain":false},{"metricName":" # translocations","quality":"Less is better","values":[0,0,0,0,0,0],"isMain":false},{"metricName":" # inversions","quality":"Less is better","values":[0,0,0,0,0,0],"isMain":false},{"metricName":" # interspecies translocations","quality":"Less is better","values":[3,7,1,10,2,3],"isMain":false},{"metricName":"# possibly misassembled contigs","quality":"Less is better","values":[0,0,0,0,0,0],"isMain":false},{"metricName":" # possible misassemblies","quality":"Less is better","values":[0,0,0,0,0,0],"isMain":false},{"metricName":"# local misassemblies","quality":"Less is better","values":[0,0,0,0,0,0],"isMain":false},{"metricName":"# scaffold gap ext. mis.","quality":"Less is better","values":[0,0,0,0,0,0],"isMain":false},{"metricName":"# scaffold gap loc. mis.","quality":"Less is better","values":[0,0,0,0,0,0],"isMain":false}]],["Unaligned",[]],["Mismatches",[]],["Statistics without reference",[]],["K-mer-based statistics",[]],["Predicted genes",[]],["Similarity statistics",[]],["Reference statistics",[]]],[["Genome statistics",[]],["Reads mapping",[]],["Misassemblies",[{"metricName":"# misassemblies","quality":"Less is better","values":[5,3,1,7,4,5],"isMain":true},{"metricName":" # relocations","quality":"Less is better","values":[0,0,0,0,0,0],"isMain":false},{"metricName":" # translocations","quality":"Less is better","values":[1,1,0,0,1,0],"isMain":false},{"metricName":" # inversions","quality":"Less is better","values":[0,0,0,0,0,0],"isMain":false},{"metricName":" # interspecies translocations","quality":"Less is better","values":[4,2,1,7,3,5],"isMain":false},{"metricName":"# possibly misassembled contigs","quality":"Less is better","values":[0,0,0,0,0,0],"isMain":false},{"metricName":" # possible misassemblies","quality":"Less is better","values":[0,0,0,0,0,0],"isMain":false},{"metricName":"# local misassemblies","quality":"Less is better","values":[0,0,0,0,0,0],"isMain":false},{"metricName":"# scaffold gap ext. mis.","quality":"Less is better","values":[0,0,0,0,0,0],"isMain":false},{"metricName":"# scaffold gap loc. mis.","quality":"Less is better","values":[0,0,0,0,0,0],"isMain":false}]],["Unaligned",[]],["Mismatches",[]],["Statistics without reference",[]],["K-mer-based statistics",[]],["Predicted genes",[]],["Similarity statistics",[]],["Reference statistics",[]]],[["Genome statistics",[]],["Reads mapping",[]],["Misassemblies",[{"metricName":"# misassemblies","quality":"Less is better","values":[38,38,8,33,37,30],"isMain":true},{"metricName":" # relocations","quality":"Less is better","values":[1,1,0,0,1,0],"isMain":false},{"metricName":" # translocations","quality":"Less is better","values":[1,2,1,1,3,0],"isMain":false},{"metricName":" # inversions","quality":"Less is better","values":[1,1,1,1,1,0],"isMain":false},{"metricName":" # interspecies translocations","quality":"Less is better","values":[35,34,6,31,32,30],"isMain":false},{"metricName":"# possibly misassembled contigs","quality":"Less is better","values":[0,0,0,0,0,0],"isMain":false},{"metricName":" # possible misassemblies","quality":"Less is better","values":[0,0,0,0,0,0],"isMain":false},{"metricName":"# local misassemblies","quality":"Less is better","values":[1,1,0,1,2,0],"isMain":false},{"metricName":"# scaffold gap ext. mis.","quality":"Less is better","values":[0,0,0,0,0,0],"isMain":false},{"metricName":"# scaffold gap loc. mis.","quality":"Less is better","values":[0,0,0,0,0,0],"isMain":false}]],["Unaligned",[]],["Mismatches",[]],["Statistics without reference",[]],["K-mer-based statistics",[]],["Predicted genes",[]],["Similarity statistics",[]],["Reference statistics",[]]],[["Genome statistics",[]],["Reads mapping",[]],["Misassemblies",[{"metricName":"# misassemblies","quality":"Less is better","values":[9,10,6,10,10,5],"isMain":true},{"metricName":" # relocations","quality":"Less is better","values":[0,0,0,0,0,0],"isMain":false},{"metricName":" # translocations","quality":"Less is better","values":[0,0,0,0,1,0],"isMain":false},{"metricName":" # inversions","quality":"Less is better","values":[0,0,0,0,0,0],"isMain":false},{"metricName":" # interspecies translocations","quality":"Less is better","values":[9,10,6,10,9,5],"isMain":false},{"metricName":"# possibly misassembled contigs","quality":"Less is better","values":[0,0,0,0,0,0],"isMain":false},{"metricName":" # possible misassemblies","quality":"Less is better","values":[0,0,0,0,0,0],"isMain":false},{"metricName":"# local misassemblies","quality":"Less is better","values":[0,1,0,0,0,0],"isMain":false},{"metricName":"# scaffold gap ext. mis.","quality":"Less is better","values":[0,0,0,0,0,0],"isMain":false},{"metricName":"# scaffold gap loc. mis.","quality":"Less is better","values":[0,0,0,0,0,0],"isMain":false}]],["Unaligned",[]],["Mismatches",[]],["Statistics without reference",[]],["K-mer-based statistics",[]],["Predicted genes",[]],["Similarity statistics",[]],["Reference statistics",[]]],[["Genome statistics",[]],["Reads mapping",[]],["Misassemblies",[{"metricName":"# misassemblies","quality":"Less is better","values":[4,3,6,4,4,0],"isMain":true},{"metricName":" # relocations","quality":"Less is better","values":[0,0,0,0,0,0],"isMain":false},{"metricName":" # translocations","quality":"Less is better","values":[0,0,0,0,0,0],"isMain":false},{"metricName":" # inversions","quality":"Less is better","values":[0,0,0,0,0,0],"isMain":false},{"metricName":" # interspecies translocations","quality":"Less is better","values":[4,3,6,4,4,0],"isMain":false},{"metricName":"# possibly misassembled contigs","quality":"Less is better","values":[0,0,0,0,0,0],"isMain":false},{"metricName":" # possible misassemblies","quality":"Less is better","values":[0,0,0,0,0,0],"isMain":false},{"metricName":"# local misassemblies","quality":"Less is better","values":[0,0,0,0,0,0],"isMain":false},{"metricName":"# scaffold gap ext. mis.","quality":"Less is better","values":[0,0,0,0,0,0],"isMain":false},{"metricName":"# scaffold gap loc. mis.","quality":"Less is better","values":[0,0,0,0,0,0],"isMain":false}]],["Unaligned",[]],["Mismatches",[]],["Statistics without reference",[]],["K-mer-based statistics",[]],["Predicted genes",[]],["Similarity statistics",[]],["Reference statistics",[]]],[["Genome statistics",[]],["Reads mapping",[]],["Misassemblies",[{"metricName":"# misassemblies","quality":"Less is better","values":[61,78,27,49,73,46],"isMain":true},{"metricName":" # relocations","quality":"Less is better","values":[4,7,3,3,7,2],"isMain":false},{"metricName":" # translocations","quality":"Less is better","values":[14,24,12,6,18,13],"isMain":false},{"metricName":" # inversions","quality":"Less is better","values":[2,0,0,1,0,0],"isMain":false},{"metricName":" # interspecies translocations","quality":"Less is better","values":[41,47,12,39,48,31],"isMain":false},{"metricName":"# possibly misassembled contigs","quality":"Less is better","values":[0,0,0,0,0,0],"isMain":false},{"metricName":" # possible misassemblies","quality":"Less is better","values":[0,0,0,0,0,0],"isMain":false},{"metricName":"# local misassemblies","quality":"Less is better","values":[2,2,2,2,3,0],"isMain":false},{"metricName":"# scaffold gap ext. mis.","quality":"Less is better","values":[0,0,0,0,0,0],"isMain":false},{"metricName":"# scaffold gap loc. mis.","quality":"Less is better","values":[0,0,0,0,0,0],"isMain":false}]],["Unaligned",[]],["Mismatches",[]],["Statistics without reference",[]],["K-mer-based statistics",[]],["Predicted genes",[]],["Similarity statistics",[]],["Reference statistics",[]]],[["Genome statistics",[]],["Reads mapping",[]],["Misassemblies",[{"metricName":"# misassemblies","quality":"Less is better","values":[0,2,3,7,4,3],"isMain":true},{"metricName":" # relocations","quality":"Less is better","values":[0,0,0,0,0,0],"isMain":false},{"metricName":" # translocations","quality":"Less is better","values":[0,0,0,0,0,0],"isMain":false},{"metricName":" # inversions","quality":"Less is better","values":[0,0,0,1,0,0],"isMain":false},{"metricName":" # interspecies translocations","quality":"Less is better","values":[0,2,3,6,4,3],"isMain":false},{"metricName":"# possibly misassembled contigs","quality":"Less is better","values":[0,0,0,0,0,0],"isMain":false},{"metricName":" # possible misassemblies","quality":"Less is better","values":[0,0,0,0,0,0],"isMain":false},{"metricName":"# local misassemblies","quality":"Less is better","values":[0,0,0,0,0,0],"isMain":false},{"metricName":"# scaffold gap ext. mis.","quality":"Less is better","values":[0,0,0,0,0,0],"isMain":false},{"metricName":"# scaffold gap loc. mis.","quality":"Less is better","values":[0,0,0,0,0,0],"isMain":false}]],["Unaligned",[]],["Mismatches",[]],["Statistics without reference",[]],["K-mer-based statistics",[]],["Predicted genes",[]],["Similarity statistics",[]],["Reference statistics",[]]],[["Genome statistics",[]],["Reads mapping",[]],["Misassemblies",[{"metricName":"# misassemblies","quality":"Less is better","values":[80,106,38,76,96,40],"isMain":true},{"metricName":" # relocations","quality":"Less is better","values":[0,2,1,0,0,1],"isMain":false},{"metricName":" # translocations","quality":"Less is better","values":[4,5,4,4,8,4],"isMain":false},{"metricName":" # inversions","quality":"Less is better","values":[0,1,0,2,0,0],"isMain":false},{"metricName":" # interspecies translocations","quality":"Less is better","values":[76,98,33,70,88,35],"isMain":false},{"metricName":"# possibly misassembled contigs","quality":"Less is better","values":[0,0,0,1,0,1],"isMain":false},{"metricName":" # possible misassemblies","quality":"Less is better","values":[0,0,0,1,0,1],"isMain":false},{"metricName":"# local misassemblies","quality":"Less is better","values":[3,1,3,7,0,1],"isMain":false},{"metricName":"# scaffold gap ext. mis.","quality":"Less is better","values":[0,0,0,0,0,0],"isMain":false},{"metricName":"# scaffold gap loc. mis.","quality":"Less is better","values":[0,0,0,0,0,0],"isMain":false}]],["Unaligned",[]],["Mismatches",[]],["Statistics without reference",[]],["K-mer-based statistics",[]],["Predicted genes",[]],["Similarity statistics",[]],["Reference statistics",[]]],[["Genome statistics",[]],["Reads mapping",[]],["Misassemblies",[{"metricName":"# misassemblies","quality":"Less is better","values":[5,7,1,12,7,1],"isMain":true},{"metricName":" # relocations","quality":"Less is better","values":[0,0,0,0,0,0],"isMain":false},{"metricName":" # translocations","quality":"Less is better","values":[0,0,0,0,0,0],"isMain":false},{"metricName":" # inversions","quality":"Less is better","values":[0,0,0,0,0,0],"isMain":false},{"metricName":" # interspecies translocations","quality":"Less is better","values":[5,7,1,12,7,1],"isMain":false},{"metricName":"# possibly misassembled contigs","quality":"Less is better","values":[0,0,0,0,0,0],"isMain":false},{"metricName":" # possible misassemblies","quality":"Less is better","values":[0,0,0,0,0,0],"isMain":false},{"metricName":"# local misassemblies","quality":"Less is better","values":[0,0,0,0,1,0],"isMain":false},{"metricName":"# scaffold gap ext. mis.","quality":"Less is better","values":[0,0,0,0,0,0],"isMain":false},{"metricName":"# scaffold gap loc. mis.","quality":"Less is better","values":[0,0,0,0,0,0],"isMain":false}]],["Unaligned",[]],["Mismatches",[]],["Statistics without reference",[]],["K-mer-based statistics",[]],["Predicted genes",[]],["Similarity statistics",[]],["Reference statistics",[]]],[["Genome statistics",[]],["Reads mapping",[]],["Misassemblies",[{"metricName":"# misassemblies","quality":"Less is better","values":[1,1,1,1,0,0],"isMain":true},{"metricName":" # relocations","quality":"Less is better","values":[0,0,0,0,0,0],"isMain":false},{"metricName":" # translocations","quality":"Less is better","values":[0,0,0,0,0,0],"isMain":false},{"metricName":" # inversions","quality":"Less is better","values":[0,0,0,0,0,0],"isMain":false},{"metricName":" # interspecies translocations","quality":"Less is better","values":[1,1,1,1,0,0],"isMain":false},{"metricName":"# possibly misassembled contigs","quality":"Less is better","values":[0,0,0,0,0,0],"isMain":false},{"metricName":" # possible misassemblies","quality":"Less is better","values":[0,0,0,0,0,0],"isMain":false},{"metricName":"# local misassemblies","quality":"Less is better","values":[0,0,0,0,0,0],"isMain":false},{"metricName":"# scaffold gap ext. mis.","quality":"Less is better","values":[0,0,0,0,0,0],"isMain":false},{"metricName":"# scaffold gap loc. mis.","quality":"Less is better","values":[0,0,0,0,0,0],"isMain":false}]],["Unaligned",[]],["Mismatches",[]],["Statistics without reference",[]],["K-mer-based statistics",[]],["Predicted genes",[]],["Similarity statistics",[]],["Reference statistics",[]]],[["Genome statistics",[]],["Reads mapping",[]],["Misassemblies",[{"metricName":"# misassemblies","quality":"Less is better","values":[9,9,10,14,12,4],"isMain":true},{"metricName":" # relocations","quality":"Less is better","values":[0,0,0,0,0,0],"isMain":false},{"metricName":" # translocations","quality":"Less is better","values":[0,2,2,2,0,1],"isMain":false},{"metricName":" # inversions","quality":"Less is better","values":[0,0,1,0,0,0],"isMain":false},{"metricName":" # interspecies translocations","quality":"Less is better","values":[9,7,7,12,12,3],"isMain":false},{"metricName":"# possibly misassembled contigs","quality":"Less is better","values":[0,1,0,0,0,0],"isMain":false},{"metricName":" # possible misassemblies","quality":"Less is better","values":[0,1,0,0,0,0],"isMain":false},{"metricName":"# local misassemblies","quality":"Less is better","values":[0,0,1,1,0,1],"isMain":false},{"metricName":"# scaffold gap ext. mis.","quality":"Less is better","values":[0,0,0,0,0,0],"isMain":false},{"metricName":"# scaffold gap loc. mis.","quality":"Less is better","values":[0,0,0,0,0,0],"isMain":false}]],["Unaligned",[]],["Mismatches",[]],["Statistics without reference",[]],["K-mer-based statistics",[]],["Predicted genes",[]],["Similarity statistics",[]],["Reference statistics",[]]],[["Genome statistics",[]],["Reads mapping",[]],["Misassemblies",[{"metricName":"# misassemblies","quality":"Less is better","values":[2,10,10,17,10,4],"isMain":true},{"metricName":" # relocations","quality":"Less is better","values":[0,1,1,1,1,0],"isMain":false},{"metricName":" # translocations","quality":"Less is better","values":[0,3,3,3,5,0],"isMain":false},{"metricName":" # inversions","quality":"Less is better","values":[0,0,0,0,0,0],"isMain":false},{"metricName":" # interspecies translocations","quality":"Less is better","values":[2,6,6,13,4,4],"isMain":false},{"metricName":"# possibly misassembled contigs","quality":"Less is better","values":[0,0,0,0,0,0],"isMain":false},{"metricName":" # possible misassemblies","quality":"Less is better","values":[0,0,0,0,0,0],"isMain":false},{"metricName":"# local misassemblies","quality":"Less is better","values":[0,0,2,2,0,0],"isMain":false},{"metricName":"# scaffold gap ext. mis.","quality":"Less is better","values":[0,0,0,0,0,0],"isMain":false},{"metricName":"# scaffold gap loc. mis.","quality":"Less is better","values":[0,0,0,0,0,0],"isMain":false}]],["Unaligned",[]],["Mismatches",[]],["Statistics without reference",[]],["K-mer-based statistics",[]],["Predicted genes",[]],["Similarity statistics",[]],["Reference statistics",[]]],[["Genome statistics",[]],["Reads mapping",[]],["Misassemblies",[{"metricName":"# misassemblies","quality":"Less is better","values":[11,10,6,18,12,10],"isMain":true},{"metricName":" # relocations","quality":"Less is better","values":[0,0,0,0,0,0],"isMain":false},{"metricName":" # translocations","quality":"Less is better","values":[2,0,2,1,1,1],"isMain":false},{"metricName":" # inversions","quality":"Less is better","values":[0,0,0,0,0,0],"isMain":false},{"metricName":" # interspecies translocations","quality":"Less is better","values":[9,10,4,17,11,9],"isMain":false},{"metricName":"# possibly misassembled contigs","quality":"Less is better","values":[0,0,0,0,0,0],"isMain":false},{"metricName":" # possible misassemblies","quality":"Less is better","values":[0,0,0,0,0,0],"isMain":false},{"metricName":"# local misassemblies","quality":"Less is better","values":[0,0,0,0,0,0],"isMain":false},{"metricName":"# scaffold gap ext. mis.","quality":"Less is better","values":[0,0,0,0,0,0],"isMain":false},{"metricName":"# scaffold gap loc. mis.","quality":"Less is better","values":[0,0,0,0,0,0],"isMain":false}]],["Unaligned",[]],["Mismatches",[]],["Statistics without reference",[]],["K-mer-based statistics",[]],["Predicted genes",[]],["Similarity statistics",[]],["Reference statistics",[]]],[["Genome statistics",[]],["Reads mapping",[]],["Misassemblies",[{"metricName":"# misassemblies","quality":"Less is better","values":[14,18,10,11,23,10],"isMain":true},{"metricName":" # relocations","quality":"Less is better","values":[0,0,0,0,0,0],"isMain":false},{"metricName":" # translocations","quality":"Less is better","values":[0,0,0,0,0,0],"isMain":false},{"metricName":" # inversions","quality":"Less is better","values":[0,0,0,0,0,0],"isMain":false},{"metricName":" # interspecies translocations","quality":"Less is better","values":[14,18,10,11,23,10],"isMain":false},{"metricName":"# possibly misassembled contigs","quality":"Less is better","values":[0,0,2,0,0,0],"isMain":false},{"metricName":" # possible misassemblies","quality":"Less is better","values":[0,0,2,0,0,0],"isMain":false},{"metricName":"# local misassemblies","quality":"Less is better","values":[0,0,0,0,1,0],"isMain":false},{"metricName":"# scaffold gap ext. mis.","quality":"Less is better","values":[0,0,0,0,0,0],"isMain":false},{"metricName":"# scaffold gap loc. mis.","quality":"Less is better","values":[0,0,0,0,0,0],"isMain":false}]],["Unaligned",[]],["Mismatches",[]],["Statistics without reference",[]],["K-mer-based statistics",[]],["Predicted genes",[]],["Similarity statistics",[]],["Reference statistics",[]]],[["Genome statistics",[]],["Reads mapping",[]],["Misassemblies",[{"metricName":"# misassemblies","quality":"Less is better","values":[0,0,0,1,1,0],"isMain":true},{"metricName":" # relocations","quality":"Less is better","values":[0,0,0,0,0,0],"isMain":false},{"metricName":" # translocations","quality":"Less is better","values":[0,0,0,0,0,0],"isMain":false},{"metricName":" # inversions","quality":"Less is better","values":[0,0,0,0,0,0],"isMain":false},{"metricName":" # interspecies translocations","quality":"Less is better","values":[0,0,0,1,1,0],"isMain":false},{"metricName":"# possibly misassembled contigs","quality":"Less is better","values":[0,0,0,0,0,0],"isMain":false},{"metricName":" # possible misassemblies","quality":"Less is better","values":[0,0,0,0,0,0],"isMain":false},{"metricName":"# local misassemblies","quality":"Less is better","values":[0,0,0,0,0,0],"isMain":false},{"metricName":"# scaffold gap ext. mis.","quality":"Less is better","values":[0,0,0,0,0,0],"isMain":false},{"metricName":"# scaffold gap loc. mis.","quality":"Less is better","values":[0,0,0,0,0,0],"isMain":false}]],["Unaligned",[]],["Mismatches",[]],["Statistics without reference",[]],["K-mer-based statistics",[]],["Predicted genes",[]],["Similarity statistics",[]],["Reference statistics",[]]],[["Genome statistics",[]],["Reads mapping",[]],["Misassemblies",[{"metricName":"# misassemblies","quality":"Less is better","values":[0,0,1,0,0,0],"isMain":true},{"metricName":" # relocations","quality":"Less is better","values":[0,0,0,0,0,0],"isMain":false},{"metricName":" # translocations","quality":"Less is better","values":[0,0,0,0,0,0],"isMain":false},{"metricName":" # inversions","quality":"Less is better","values":[0,0,0,0,0,0],"isMain":false},{"metricName":" # interspecies translocations","quality":"Less is better","values":[0,0,1,0,0,0],"isMain":false},{"metricName":"# possibly misassembled contigs","quality":"Less is better","values":[0,0,0,0,0,0],"isMain":false},{"metricName":" # possible misassemblies","quality":"Less is better","values":[0,0,0,0,0,0],"isMain":false},{"metricName":"# local misassemblies","quality":"Less is better","values":[0,0,0,0,0,0],"isMain":false},{"metricName":"# scaffold gap ext. mis.","quality":"Less is better","values":[0,0,0,0,0,0],"isMain":false},{"metricName":"# scaffold gap loc. mis.","quality":"Less is better","values":[0,0,0,0,0,0],"isMain":false}]],["Unaligned",[]],["Mismatches",[]],["Statistics without reference",[]],["K-mer-based statistics",[]],["Predicted genes",[]],["Similarity statistics",[]],["Reference statistics",[]]],[["Genome statistics",[]],["Reads mapping",[]],["Misassemblies",[{"metricName":"# misassemblies","quality":"Less is better","values":[49,56,11,31,45,50],"isMain":true},{"metricName":" # relocations","quality":"Less is better","values":[0,3,1,2,3,0],"isMain":false},{"metricName":" # translocations","quality":"Less is better","values":[4,7,4,6,10,5],"isMain":false},{"metricName":" # inversions","quality":"Less is better","values":[2,0,0,0,1,0],"isMain":false},{"metricName":" # interspecies translocations","quality":"Less is better","values":[43,46,6,23,31,45],"isMain":false},{"metricName":"# possibly misassembled contigs","quality":"Less is better","values":[0,0,0,0,0,0],"isMain":false},{"metricName":" # possible misassemblies","quality":"Less is better","values":[0,0,0,0,0,0],"isMain":false},{"metricName":"# local misassemblies","quality":"Less is better","values":[3,3,2,2,1,0],"isMain":false},{"metricName":"# scaffold gap ext. mis.","quality":"Less is better","values":[0,0,0,0,0,0],"isMain":false},{"metricName":"# scaffold gap loc. mis.","quality":"Less is better","values":[0,0,0,0,0,0],"isMain":false}]],["Unaligned",[]],["Mismatches",[]],["Statistics without reference",[]],["K-mer-based statistics",[]],["Predicted genes",[]],["Similarity statistics",[]],["Reference statistics",[]]],[["Genome statistics",[]],["Reads mapping",[]],["Misassemblies",[{"metricName":"# misassemblies","quality":"Less is better","values":[6,18,10,13,13,4],"isMain":true},{"metricName":" # relocations","quality":"Less is better","values":[1,0,0,0,0,0],"isMain":false},{"metricName":" # translocations","quality":"Less is better","values":[0,1,1,0,1,0],"isMain":false},{"metricName":" # inversions","quality":"Less is better","values":[0,0,0,0,0,0],"isMain":false},{"metricName":" # interspecies translocations","quality":"Less is better","values":[5,17,9,13,12,4],"isMain":false},{"metricName":"# possibly misassembled contigs","quality":"Less is better","values":[0,0,0,0,0,0],"isMain":false},{"metricName":" # possible misassemblies","quality":"Less is better","values":[0,0,0,0,0,0],"isMain":false},{"metricName":"# local misassemblies","quality":"Less is better","values":[1,0,0,0,0,0],"isMain":false},{"metricName":"# scaffold gap ext. mis.","quality":"Less is better","values":[0,0,0,0,0,0],"isMain":false},{"metricName":"# scaffold gap loc. mis.","quality":"Less is better","values":[0,0,0,0,0,0],"isMain":false}]],["Unaligned",[]],["Mismatches",[]],["Statistics without reference",[]],["K-mer-based statistics",[]],["Predicted genes",[]],["Similarity statistics",[]],["Reference statistics",[]]],[["Genome statistics",[]],["Reads mapping",[]],["Misassemblies",[{"metricName":"# misassemblies","quality":"Less is better","values":[3,1,3,3,2,0],"isMain":true},{"metricName":" # relocations","quality":"Less is better","values":[0,0,0,0,0,0],"isMain":false},{"metricName":" # translocations","quality":"Less is better","values":[0,0,0,0,0,0],"isMain":false},{"metricName":" # inversions","quality":"Less is better","values":[0,0,0,0,0,0],"isMain":false},{"metricName":" # interspecies translocations","quality":"Less is better","values":[3,1,3,3,2,0],"isMain":false},{"metricName":"# possibly misassembled contigs","quality":"Less is better","values":[0,0,0,0,0,0],"isMain":false},{"metricName":" # possible misassemblies","quality":"Less is better","values":[0,0,0,0,0,0],"isMain":false},{"metricName":"# local misassemblies","quality":"Less is better","values":[0,0,0,0,0,0],"isMain":false},{"metricName":"# scaffold gap ext. mis.","quality":"Less is better","values":[0,0,0,0,0,0],"isMain":false},{"metricName":"# scaffold gap loc. mis.","quality":"Less is better","values":[0,0,0,0,0,0],"isMain":false}]],["Unaligned",[]],["Mismatches",[]],["Statistics without reference",[]],["K-mer-based statistics",[]],["Predicted genes",[]],["Similarity statistics",[]],["Reference statistics",[]]],[["Genome statistics",[]],["Reads mapping",[]],["Misassemblies",[{"metricName":"# misassemblies","quality":"Less is better","values":[64,58,32,41,72,15],"isMain":true},{"metricName":" # relocations","quality":"Less is better","values":[2,0,1,0,1,0],"isMain":false},{"metricName":" # translocations","quality":"Less is better","values":[1,2,2,2,2,1],"isMain":false},{"metricName":" # inversions","quality":"Less is better","values":[0,1,3,0,3,0],"isMain":false},{"metricName":" # interspecies translocations","quality":"Less is better","values":[61,55,26,39,66,14],"isMain":false},{"metricName":"# possibly misassembled contigs","quality":"Less is better","values":[0,0,0,0,1,0],"isMain":false},{"metricName":" # possible misassemblies","quality":"Less is better","values":[0,0,0,0,1,0],"isMain":false},{"metricName":"# local misassemblies","quality":"Less is better","values":[0,0,2,2,1,0],"isMain":false},{"metricName":"# scaffold gap ext. mis.","quality":"Less is better","values":[0,0,0,0,0,0],"isMain":false},{"metricName":"# scaffold gap loc. mis.","quality":"Less is better","values":[0,0,0,0,0,0],"isMain":false}]],["Unaligned",[]],["Mismatches",[]],["Statistics without reference",[]],["K-mer-based statistics",[]],["Predicted genes",[]],["Similarity statistics",[]],["Reference statistics",[]]],[["Genome statistics",[]],["Reads mapping",[]],["Misassemblies",[{"metricName":"# misassemblies","quality":"Less is better","values":[8,15,3,10,13,4],"isMain":true},{"metricName":" # relocations","quality":"Less is better","values":[0,0,0,0,0,0],"isMain":false},{"metricName":" # translocations","quality":"Less is better","values":[1,3,2,0,2,0],"isMain":false},{"metricName":" # inversions","quality":"Less is better","values":[0,0,0,0,0,0],"isMain":false},{"metricName":" # interspecies translocations","quality":"Less is better","values":[7,12,1,10,11,4],"isMain":false},{"metricName":"# possibly misassembled contigs","quality":"Less is better","values":[0,0,0,0,0,0],"isMain":false},{"metricName":" # possible misassemblies","quality":"Less is better","values":[0,0,0,0,0,0],"isMain":false},{"metricName":"# local misassemblies","quality":"Less is better","values":[0,0,0,0,0,0],"isMain":false},{"metricName":"# scaffold gap ext. mis.","quality":"Less is better","values":[0,0,0,0,0,0],"isMain":false},{"metricName":"# scaffold gap loc. mis.","quality":"Less is better","values":[0,0,0,0,0,0],"isMain":false}]],["Unaligned",[]],["Mismatches",[]],["Statistics without reference",[]],["K-mer-based statistics",[]],["Predicted genes",[]],["Similarity statistics",[]],["Reference statistics",[]]],[["Genome statistics",[]],["Reads mapping",[]],["Misassemblies",[{"metricName":"# misassemblies","quality":"Less is better","values":[0,1,0,2,0,0],"isMain":true},{"metricName":" # relocations","quality":"Less is better","values":[0,0,0,0,0,0],"isMain":false},{"metricName":" # translocations","quality":"Less is better","values":[0,0,0,0,0,0],"isMain":false},{"metricName":" # inversions","quality":"Less is better","values":[0,0,0,0,0,0],"isMain":false},{"metricName":" # interspecies translocations","quality":"Less is better","values":[0,1,0,2,0,0],"isMain":false},{"metricName":"# possibly misassembled contigs","quality":"Less is better","values":[0,0,0,0,0,0],"isMain":false},{"metricName":" # possible misassemblies","quality":"Less is better","values":[0,0,0,0,0,0],"isMain":false},{"metricName":"# local misassemblies","quality":"Less is better","values":[0,0,0,0,0,0],"isMain":false},{"metricName":"# scaffold gap ext. mis.","quality":"Less is better","values":[0,0,0,0,0,0],"isMain":false},{"metricName":"# scaffold gap loc. mis.","quality":"Less is better","values":[0,0,0,0,0,0],"isMain":false}]],["Unaligned",[]],["Mismatches",[]],["Statistics without reference",[]],["K-mer-based statistics",[]],["Predicted genes",[]],["Similarity statistics",[]],["Reference statistics",[]]],[["Genome statistics",[]],["Reads mapping",[]],["Misassemblies",[{"metricName":"# misassemblies","quality":"Less is better","values":[28,48,47,77,60,12],"isMain":true},{"metricName":" # relocations","quality":"Less is better","values":[1,1,2,4,0,0],"isMain":false},{"metricName":" # translocations","quality":"Less is better","values":[6,8,9,6,6,5],"isMain":false},{"metricName":" # inversions","quality":"Less is better","values":[0,3,2,1,1,0],"isMain":false},{"metricName":" # interspecies translocations","quality":"Less is better","values":[21,36,34,66,53,7],"isMain":false},{"metricName":"# possibly misassembled contigs","quality":"Less is better","values":[0,1,1,0,2,0],"isMain":false},{"metricName":" # possible misassemblies","quality":"Less is better","values":[0,1,1,0,2,0],"isMain":false},{"metricName":"# local misassemblies","quality":"Less is better","values":[2,1,2,2,6,2],"isMain":false},{"metricName":"# scaffold gap ext. mis.","quality":"Less is better","values":[0,0,0,0,0,0],"isMain":false},{"metricName":"# scaffold gap loc. mis.","quality":"Less is better","values":[0,0,0,0,0,0],"isMain":false}]],["Unaligned",[]],["Mismatches",[]],["Statistics without reference",[]],["K-mer-based statistics",[]],["Predicted genes",[]],["Similarity statistics",[]],["Reference statistics",[]]],[["Genome statistics",[]],["Reads mapping",[]],["Misassemblies",[{"metricName":"# misassemblies","quality":"Less is better","values":[15,20,2,15,21,10],"isMain":true},{"metricName":" # relocations","quality":"Less is better","values":[0,1,0,0,1,0],"isMain":false},{"metricName":" # translocations","quality":"Less is better","values":[2,1,2,0,1,1],"isMain":false},{"metricName":" # inversions","quality":"Less is better","values":[0,0,0,0,0,0],"isMain":false},{"metricName":" # interspecies translocations","quality":"Less is better","values":[13,18,0,15,19,9],"isMain":false},{"metricName":"# possibly misassembled contigs","quality":"Less is better","values":[0,0,0,0,0,0],"isMain":false},{"metricName":" # possible misassemblies","quality":"Less is better","values":[0,0,0,0,0,0],"isMain":false},{"metricName":"# local misassemblies","quality":"Less is better","values":[1,1,0,0,0,1],"isMain":false},{"metricName":"# scaffold gap ext. mis.","quality":"Less is better","values":[0,0,0,0,0,0],"isMain":false},{"metricName":"# scaffold gap loc. mis.","quality":"Less is better","values":[0,0,0,0,0,0],"isMain":false}]],["Unaligned",[]],["Mismatches",[]],["Statistics without reference",[]],["K-mer-based statistics",[]],["Predicted genes",[]],["Similarity statistics",[]],["Reference statistics",[]]],[["Genome statistics",[]],["Reads mapping",[]],["Misassemblies",[{"metricName":"# misassemblies","quality":"Less is better","values":[29,32,16,28,32,12],"isMain":true},{"metricName":" # relocations","quality":"Less is better","values":[0,1,0,0,1,0],"isMain":false},{"metricName":" # translocations","quality":"Less is better","values":[0,0,0,2,0,1],"isMain":false},{"metricName":" # inversions","quality":"Less is better","values":[0,0,0,0,0,0],"isMain":false},{"metricName":" # interspecies translocations","quality":"Less is better","values":[29,31,16,26,31,11],"isMain":false},{"metricName":"# possibly misassembled contigs","quality":"Less is better","values":[0,2,0,0,1,0],"isMain":false},{"metricName":" # possible misassemblies","quality":"Less is better","values":[0,2,0,0,1,0],"isMain":false},{"metricName":"# local misassemblies","quality":"Less is better","values":[0,0,0,1,0,1],"isMain":false},{"metricName":"# scaffold gap ext. mis.","quality":"Less is better","values":[0,0,0,0,0,0],"isMain":false},{"metricName":"# scaffold gap loc. mis.","quality":"Less is better","values":[0,0,0,0,0,0],"isMain":false}]],["Unaligned",[]],["Mismatches",[]],["Statistics without reference",[]],["K-mer-based statistics",[]],["Predicted genes",[]],["Similarity statistics",[]],["Reference statistics",[]]],[["Genome statistics",[]],["Reads mapping",[]],["Misassemblies",[{"metricName":"# misassemblies","quality":"Less is better","values":[9,14,6,12,12,11],"isMain":true},{"metricName":" # relocations","quality":"Less is better","values":[0,0,0,0,0,0],"isMain":false},{"metricName":" # translocations","quality":"Less is better","values":[1,2,0,0,1,1],"isMain":false},{"metricName":" # inversions","quality":"Less is better","values":[0,0,0,1,0,0],"isMain":false},{"metricName":" # interspecies translocations","quality":"Less is better","values":[8,12,6,11,11,10],"isMain":false},{"metricName":"# possibly misassembled contigs","quality":"Less is better","values":[0,0,0,0,0,0],"isMain":false},{"metricName":" # possible misassemblies","quality":"Less is better","values":[0,0,0,0,0,0],"isMain":false},{"metricName":"# local misassemblies","quality":"Less is better","values":[0,0,0,0,0,0],"isMain":false},{"metricName":"# scaffold gap ext. mis.","quality":"Less is better","values":[0,0,0,0,0,0],"isMain":false},{"metricName":"# scaffold gap loc. mis.","quality":"Less is better","values":[0,0,0,0,0,0],"isMain":false}]],["Unaligned",[]],["Mismatches",[]],["Statistics without reference",[]],["K-mer-based statistics",[]],["Predicted genes",[]],["Similarity statistics",[]],["Reference statistics",[]]],[["Genome statistics",[]],["Reads mapping",[]],["Misassemblies",[{"metricName":"# misassemblies","quality":"Less is better","values":[34,57,12,47,46,27],"isMain":true},{"metricName":" # relocations","quality":"Less is better","values":[1,1,1,2,0,1],"isMain":false},{"metricName":" # translocations","quality":"Less is better","values":[2,5,5,4,4,1],"isMain":false},{"metricName":" # inversions","quality":"Less is better","values":[0,2,0,0,2,0],"isMain":false},{"metricName":" # interspecies translocations","quality":"Less is better","values":[31,49,6,41,40,25],"isMain":false},{"metricName":"# possibly misassembled contigs","quality":"Less is better","values":[0,0,0,0,0,0],"isMain":false},{"metricName":" # possible misassemblies","quality":"Less is better","values":[0,0,0,0,0,0],"isMain":false},{"metricName":"# local misassemblies","quality":"Less is better","values":[1,4,0,0,5,1],"isMain":false},{"metricName":"# scaffold gap ext. mis.","quality":"Less is better","values":[0,0,0,0,0,0],"isMain":false},{"metricName":"# scaffold gap loc. mis.","quality":"Less is better","values":[0,0,0,0,0,0],"isMain":false}]],["Unaligned",[]],["Mismatches",[]],["Statistics without reference",[]],["K-mer-based statistics",[]],["Predicted genes",[]],["Similarity statistics",[]],["Reference statistics",[]]],[["Genome statistics",[]],["Reads mapping",[]],["Misassemblies",[{"metricName":"# misassemblies","quality":"Less is better","values":[9,6,0,16,5,3],"isMain":true},{"metricName":" # relocations","quality":"Less is better","values":[1,0,0,0,0,0],"isMain":false},{"metricName":" # translocations","quality":"Less is better","values":[0,0,0,1,0,0],"isMain":false},{"metricName":" # inversions","quality":"Less is better","values":[0,0,0,0,0,0],"isMain":false},{"metricName":" # interspecies translocations","quality":"Less is better","values":[8,6,0,15,5,3],"isMain":false},{"metricName":"# possibly misassembled contigs","quality":"Less is better","values":[0,0,0,0,0,0],"isMain":false},{"metricName":" # possible misassemblies","quality":"Less is better","values":[0,0,0,0,0,0],"isMain":false},{"metricName":"# local misassemblies","quality":"Less is better","values":[0,0,0,0,0,0],"isMain":false},{"metricName":"# scaffold gap ext. mis.","quality":"Less is better","values":[0,0,0,0,0,0],"isMain":false},{"metricName":"# scaffold gap loc. mis.","quality":"Less is better","values":[0,0,0,0,0,0],"isMain":false}]],["Unaligned",[]],["Mismatches",[]],["Statistics without reference",[]],["K-mer-based statistics",[]],["Predicted genes",[]],["Similarity statistics",[]],["Reference statistics",[]]],[["Genome statistics",[]],["Reads mapping",[]],["Misassemblies",[{"metricName":"# misassemblies","quality":"Less is better","values":[5,3,9,16,13,4],"isMain":true},{"metricName":" # relocations","quality":"Less is better","values":[0,0,0,0,0,0],"isMain":false},{"metricName":" # translocations","quality":"Less is better","values":[1,0,0,0,2,0],"isMain":false},{"metricName":" # inversions","quality":"Less is better","values":[0,0,0,0,0,0],"isMain":false},{"metricName":" # interspecies translocations","quality":"Less is better","values":[4,3,9,16,11,4],"isMain":false},{"metricName":"# possibly misassembled contigs","quality":"Less is better","values":[0,0,0,0,0,0],"isMain":false},{"metricName":" # possible misassemblies","quality":"Less is better","values":[0,0,0,0,0,0],"isMain":false},{"metricName":"# local misassemblies","quality":"Less is better","values":[0,0,0,1,1,0],"isMain":false},{"metricName":"# scaffold gap ext. mis.","quality":"Less is better","values":[0,0,0,0,0,0],"isMain":false},{"metricName":"# scaffold gap loc. mis.","quality":"Less is better","values":[0,0,0,0,0,0],"isMain":false}]],["Unaligned",[]],["Mismatches",[]],["Statistics without reference",[]],["K-mer-based statistics",[]],["Predicted genes",[]],["Similarity statistics",[]],["Reference statistics",[]]],[["Genome statistics",[]],["Reads mapping",[]],["Misassemblies",[{"metricName":"# misassemblies","quality":"Less is better","values":[55,56,22,50,58,32],"isMain":true},{"metricName":" # relocations","quality":"Less is better","values":[1,1,1,0,1,1],"isMain":false},{"metricName":" # translocations","quality":"Less is better","values":[3,2,1,2,1,1],"isMain":false},{"metricName":" # inversions","quality":"Less is better","values":[1,0,1,5,2,0],"isMain":false},{"metricName":" # interspecies translocations","quality":"Less is better","values":[50,53,19,43,54,30],"isMain":false},{"metricName":"# possibly misassembled contigs","quality":"Less is better","values":[0,0,0,0,0,0],"isMain":false},{"metricName":" # possible misassemblies","quality":"Less is better","values":[0,0,0,0,0,0],"isMain":false},{"metricName":"# local misassemblies","quality":"Less is better","values":[1,0,0,1,0,0],"isMain":false},{"metricName":"# scaffold gap ext. mis.","quality":"Less is better","values":[0,0,0,0,0,0],"isMain":false},{"metricName":"# scaffold gap loc. mis.","quality":"Less is better","values":[0,0,0,0,0,0],"isMain":false}]],["Unaligned",[]],["Mismatches",[]],["Statistics without reference",[]],["K-mer-based statistics",[]],["Predicted genes",[]],["Similarity statistics",[]],["Reference statistics",[]]],[["Genome statistics",[]],["Reads mapping",[]],["Misassemblies",[{"metricName":"# misassemblies","quality":"Less is better","values":[2,1,0,4,1,5],"isMain":true},{"metricName":" # relocations","quality":"Less is better","values":[0,0,0,0,0,0],"isMain":false},{"metricName":" # translocations","quality":"Less is better","values":[0,0,0,0,0,0],"isMain":false},{"metricName":" # inversions","quality":"Less is better","values":[0,0,0,0,0,0],"isMain":false},{"metricName":" # interspecies translocations","quality":"Less is better","values":[2,1,0,4,1,5],"isMain":false},{"metricName":"# possibly misassembled contigs","quality":"Less is better","values":[0,0,0,0,0,0],"isMain":false},{"metricName":" # possible misassemblies","quality":"Less is better","values":[0,0,0,0,0,0],"isMain":false},{"metricName":"# local misassemblies","quality":"Less is better","values":[0,0,0,0,0,0],"isMain":false},{"metricName":"# scaffold gap ext. mis.","quality":"Less is better","values":[0,0,0,0,0,0],"isMain":false},{"metricName":"# scaffold gap loc. mis.","quality":"Less is better","values":[0,0,0,0,0,0],"isMain":false}]],["Unaligned",[]],["Mismatches",[]],["Statistics without reference",[]],["K-mer-based statistics",[]],["Predicted genes",[]],["Similarity statistics",[]],["Reference statistics",[]]],[["Genome statistics",[]],["Reads mapping",[]],["Misassemblies",[{"metricName":"# misassemblies","quality":"Less is better","values":[9,15,11,13,15,4],"isMain":true},{"metricName":" # relocations","quality":"Less is better","values":[0,1,0,2,1,1],"isMain":false},{"metricName":" # translocations","quality":"Less is better","values":[1,5,3,2,2,2],"isMain":false},{"metricName":" # inversions","quality":"Less is better","values":[0,0,0,0,0,0],"isMain":false},{"metricName":" # interspecies translocations","quality":"Less is better","values":[8,9,8,9,12,1],"isMain":false},{"metricName":"# possibly misassembled contigs","quality":"Less is better","values":[0,0,0,0,0,0],"isMain":false},{"metricName":" # possible misassemblies","quality":"Less is better","values":[0,0,0,0,0,0],"isMain":false},{"metricName":"# local misassemblies","quality":"Less is better","values":[0,1,5,4,2,0],"isMain":false},{"metricName":"# scaffold gap ext. mis.","quality":"Less is better","values":[0,0,0,0,0,0],"isMain":false},{"metricName":"# scaffold gap loc. mis.","quality":"Less is better","values":[0,0,0,0,0,0],"isMain":false}]],["Unaligned",[]],["Mismatches",[]],["Statistics without reference",[]],["K-mer-based statistics",[]],["Predicted genes",[]],["Similarity statistics",[]],["Reference statistics",[]]],[["Genome statistics",[]],["Reads mapping",[]],["Misassemblies",[{"metricName":"# misassemblies","quality":"Less is better","values":[0,2,2,4,3,0],"isMain":true},{"metricName":" # relocations","quality":"Less is better","values":[0,0,0,0,0,0],"isMain":false},{"metricName":" # translocations","quality":"Less is better","values":[0,0,0,0,0,0],"isMain":false},{"metricName":" # inversions","quality":"Less is better","values":[0,0,0,0,0,0],"isMain":false},{"metricName":" # interspecies translocations","quality":"Less is better","values":[0,2,2,4,3,0],"isMain":false},{"metricName":"# possibly misassembled contigs","quality":"Less is better","values":[0,0,0,0,0,0],"isMain":false},{"metricName":" # possible misassemblies","quality":"Less is better","values":[0,0,0,0,0,0],"isMain":false},{"metricName":"# local misassemblies","quality":"Less is better","values":[0,0,0,0,0,0],"isMain":false},{"metricName":"# scaffold gap ext. mis.","quality":"Less is better","values":[0,0,0,0,0,0],"isMain":false},{"metricName":"# scaffold gap loc. mis.","quality":"Less is better","values":[0,0,0,0,0,0],"isMain":false}]],["Unaligned",[]],["Mismatches",[]],["Statistics without reference",[]],["K-mer-based statistics",[]],["Predicted genes",[]],["Similarity statistics",[]],["Reference statistics",[]]],[["Genome statistics",[]],["Reads mapping",[]],["Misassemblies",[{"metricName":"# misassemblies","quality":"Less is better","values":[25,49,35,54,60,14],"isMain":true},{"metricName":" # relocations","quality":"Less is better","values":[0,0,0,0,0,0],"isMain":false},{"metricName":" # translocations","quality":"Less is better","values":[4,4,3,1,6,2],"isMain":false},{"metricName":" # inversions","quality":"Less is better","values":[2,1,2,3,1,0],"isMain":false},{"metricName":" # interspecies translocations","quality":"Less is better","values":[19,44,30,50,53,12],"isMain":false},{"metricName":"# possibly misassembled contigs","quality":"Less is better","values":[0,0,2,0,0,0],"isMain":false},{"metricName":" # possible misassemblies","quality":"Less is better","values":[0,0,2,0,0,0],"isMain":false},{"metricName":"# local misassemblies","quality":"Less is better","values":[1,2,3,4,2,1],"isMain":false},{"metricName":"# scaffold gap ext. mis.","quality":"Less is better","values":[0,0,0,0,0,0],"isMain":false},{"metricName":"# scaffold gap loc. mis.","quality":"Less is better","values":[0,0,0,0,0,0],"isMain":false}]],["Unaligned",[]],["Mismatches",[]],["Statistics without reference",[]],["K-mer-based statistics",[]],["Predicted genes",[]],["Similarity statistics",[]],["Reference statistics",[]]],[["Genome statistics",[]],["Reads mapping",[]],["Misassemblies",[{"metricName":"# misassemblies","quality":"Less is better","values":[20,28,17,30,21,5],"isMain":true},{"metricName":" # relocations","quality":"Less is better","values":[0,0,0,0,0,0],"isMain":false},{"metricName":" # translocations","quality":"Less is better","values":[2,0,1,2,0,0],"isMain":false},{"metricName":" # inversions","quality":"Less is better","values":[0,0,0,1,0,0],"isMain":false},{"metricName":" # interspecies translocations","quality":"Less is better","values":[18,28,16,27,21,5],"isMain":false},{"metricName":"# possibly misassembled contigs","quality":"Less is better","values":[0,0,0,0,0,0],"isMain":false},{"metricName":" # possible misassemblies","quality":"Less is better","values":[0,0,0,0,0,0],"isMain":false},{"metricName":"# local misassemblies","quality":"Less is better","values":[0,1,0,2,0,0],"isMain":false},{"metricName":"# scaffold gap ext. mis.","quality":"Less is better","values":[0,0,0,0,0,0],"isMain":false},{"metricName":"# scaffold gap loc. mis.","quality":"Less is better","values":[0,0,0,0,0,0],"isMain":false}]],["Unaligned",[]],["Mismatches",[]],["Statistics without reference",[]],["K-mer-based statistics",[]],["Predicted genes",[]],["Similarity statistics",[]],["Reference statistics",[]]],[["Genome statistics",[]],["Reads mapping",[]],["Misassemblies",[{"metricName":"# misassemblies","quality":"Less is better","values":[8,14,9,17,22,3],"isMain":true},{"metricName":" # relocations","quality":"Less is better","values":[0,0,0,0,0,0],"isMain":false},{"metricName":" # translocations","quality":"Less is better","values":[0,0,0,0,0,0],"isMain":false},{"metricName":" # inversions","quality":"Less is better","values":[0,0,0,0,0,0],"isMain":false},{"metricName":" # interspecies translocations","quality":"Less is better","values":[8,14,9,17,22,3],"isMain":false},{"metricName":"# possibly misassembled contigs","quality":"Less is better","values":[0,0,0,0,1,0],"isMain":false},{"metricName":" # possible misassemblies","quality":"Less is better","values":[0,0,0,0,1,0],"isMain":false},{"metricName":"# local misassemblies","quality":"Less is better","values":[0,0,2,1,1,0],"isMain":false},{"metricName":"# scaffold gap ext. mis.","quality":"Less is better","values":[0,0,0,0,0,0],"isMain":false},{"metricName":"# scaffold gap loc. mis.","quality":"Less is better","values":[0,0,0,0,0,0],"isMain":false}]],["Unaligned",[]],["Mismatches",[]],["Statistics without reference",[]],["K-mer-based statistics",[]],["Predicted genes",[]],["Similarity statistics",[]],["Reference statistics",[]]],[["Genome statistics",[]],["Reads mapping",[]],["Misassemblies",[{"metricName":"# misassemblies","quality":"Less is better","values":[4,3,2,4,4,3],"isMain":true},{"metricName":" # relocations","quality":"Less is better","values":[0,0,0,0,0,0],"isMain":false},{"metricName":" # translocations","quality":"Less is better","values":[0,0,0,0,0,0],"isMain":false},{"metricName":" # inversions","quality":"Less is better","values":[0,0,0,0,0,0],"isMain":false},{"metricName":" # interspecies translocations","quality":"Less is better","values":[4,3,2,4,4,3],"isMain":false},{"metricName":"# possibly misassembled contigs","quality":"Less is better","values":[0,0,0,0,0,0],"isMain":false},{"metricName":" # possible misassemblies","quality":"Less is better","values":[0,0,0,0,0,0],"isMain":false},{"metricName":"# local misassemblies","quality":"Less is better","values":[0,0,0,0,0,0],"isMain":false},{"metricName":"# scaffold gap ext. mis.","quality":"Less is better","values":[0,0,0,0,0,0],"isMain":false},{"metricName":"# scaffold gap loc. mis.","quality":"Less is better","values":[0,0,0,0,0,0],"isMain":false}]],["Unaligned",[]],["Mismatches",[]],["Statistics without reference",[]],["K-mer-based statistics",[]],["Predicted genes",[]],["Similarity statistics",[]],["Reference statistics",[]]],[["Genome statistics",[]],["Reads mapping",[]],["Misassemblies",[{"metricName":"# misassemblies","quality":"Less is better","values":[8,15,11,14,25,5],"isMain":true},{"metricName":" # relocations","quality":"Less is better","values":[0,0,0,0,0,0],"isMain":false},{"metricName":" # translocations","quality":"Less is better","values":[1,0,0,1,0,0],"isMain":false},{"metricName":" # inversions","quality":"Less is better","values":[0,0,0,0,1,0],"isMain":false},{"metricName":" # interspecies translocations","quality":"Less is better","values":[7,15,11,13,24,5],"isMain":false},{"metricName":"# possibly misassembled contigs","quality":"Less is better","values":[0,0,0,0,0,0],"isMain":false},{"metricName":" # possible misassemblies","quality":"Less is better","values":[0,0,0,0,0,0],"isMain":false},{"metricName":"# local misassemblies","quality":"Less is better","values":[2,3,1,5,5,0],"isMain":false},{"metricName":"# scaffold gap ext. mis.","quality":"Less is better","values":[0,0,0,0,0,0],"isMain":false},{"metricName":"# scaffold gap loc. mis.","quality":"Less is better","values":[0,0,0,0,0,0],"isMain":false}]],["Unaligned",[]],["Mismatches",[]],["Statistics without reference",[]],["K-mer-based statistics",[]],["Predicted genes",[]],["Similarity statistics",[]],["Reference statistics",[]]],[["Genome statistics",[]],["Reads mapping",[]],["Misassemblies",[{"metricName":"# misassemblies","quality":"Less is better","values":[2,6,3,11,9,3],"isMain":true},{"metricName":" # relocations","quality":"Less is better","values":[0,0,0,0,0,0],"isMain":false},{"metricName":" # translocations","quality":"Less is better","values":[0,0,1,1,1,0],"isMain":false},{"metricName":" # inversions","quality":"Less is better","values":[0,0,0,0,0,0],"isMain":false},{"metricName":" # interspecies translocations","quality":"Less is better","values":[2,6,2,10,8,3],"isMain":false},{"metricName":"# possibly misassembled contigs","quality":"Less is better","values":[0,0,0,0,0,0],"isMain":false},{"metricName":" # possible misassemblies","quality":"Less is better","values":[0,0,0,0,0,0],"isMain":false},{"metricName":"# local misassemblies","quality":"Less is better","values":[0,0,0,0,0,0],"isMain":false},{"metricName":"# scaffold gap ext. mis.","quality":"Less is better","values":[0,0,0,0,0,0],"isMain":false},{"metricName":"# scaffold gap loc. mis.","quality":"Less is better","values":[0,0,0,0,0,0],"isMain":false}]],["Unaligned",[]],["Mismatches",[]],["Statistics without reference",[]],["K-mer-based statistics",[]],["Predicted genes",[]],["Similarity statistics",[]],["Reference statistics",[]]],[["Genome statistics",[]],["Reads mapping",[]],["Misassemblies",[{"metricName":"# misassemblies","quality":"Less is better","values":[10,18,14,29,31,6],"isMain":true},{"metricName":" # relocations","quality":"Less is better","values":[0,0,1,0,0,0],"isMain":false},{"metricName":" # translocations","quality":"Less is better","values":[0,1,0,0,0,0],"isMain":false},{"metricName":" # inversions","quality":"Less is better","values":[0,0,0,0,0,0],"isMain":false},{"metricName":" # interspecies translocations","quality":"Less is better","values":[10,17,13,29,31,6],"isMain":false},{"metricName":"# possibly misassembled contigs","quality":"Less is better","values":[0,0,0,0,1,0],"isMain":false},{"metricName":" # possible misassemblies","quality":"Less is better","values":[0,0,0,0,1,0],"isMain":false},{"metricName":"# local misassemblies","quality":"Less is better","values":[2,3,5,7,7,0],"isMain":false},{"metricName":"# scaffold gap ext. mis.","quality":"Less is better","values":[0,0,0,0,0,0],"isMain":false},{"metricName":"# scaffold gap loc. mis.","quality":"Less is better","values":[0,0,0,0,0,0],"isMain":false}]],["Unaligned",[]],["Mismatches",[]],["Statistics without reference",[]],["K-mer-based statistics",[]],["Predicted genes",[]],["Similarity statistics",[]],["Reference statistics",[]]],[["Genome statistics",[]],["Reads mapping",[]],["Misassemblies",[{"metricName":"# misassemblies","quality":"Less is better","values":[48,58,7,38,42,33],"isMain":true},{"metricName":" # relocations","quality":"Less is better","values":[0,1,0,0,1,0],"isMain":false},{"metricName":" # translocations","quality":"Less is better","values":[4,6,3,6,9,2],"isMain":false},{"metricName":" # inversions","quality":"Less is better","values":[1,1,0,0,1,0],"isMain":false},{"metricName":" # interspecies translocations","quality":"Less is better","values":[43,50,4,32,31,31],"isMain":false},{"metricName":"# possibly misassembled contigs","quality":"Less is better","values":[0,0,0,0,0,0],"isMain":false},{"metricName":" # possible misassemblies","quality":"Less is better","values":[0,0,0,0,0,0],"isMain":false},{"metricName":"# local misassemblies","quality":"Less is better","values":[4,4,1,5,2,0],"isMain":false},{"metricName":"# scaffold gap ext. mis.","quality":"Less is better","values":[0,0,0,0,0,0],"isMain":false},{"metricName":"# scaffold gap loc. mis.","quality":"Less is better","values":[0,0,0,0,0,0],"isMain":false}]],["Unaligned",[]],["Mismatches",[]],["Statistics without reference",[]],["K-mer-based statistics",[]],["Predicted genes",[]],["Similarity statistics",[]],["Reference statistics",[]]],[["Genome statistics",[]],["Reads mapping",[]],["Misassemblies",[{"metricName":"# misassemblies","quality":"Less is better","values":[11,16,9,40,18,4],"isMain":true},{"metricName":" # relocations","quality":"Less is better","values":[1,1,2,1,0,0],"isMain":false},{"metricName":" # translocations","quality":"Less is better","values":[2,3,1,4,5,2],"isMain":false},{"metricName":" # inversions","quality":"Less is better","values":[0,0,0,2,0,0],"isMain":false},{"metricName":" # interspecies translocations","quality":"Less is better","values":[8,12,6,33,13,2],"isMain":false},{"metricName":"# possibly misassembled contigs","quality":"Less is better","values":[0,0,0,0,0,0],"isMain":false},{"metricName":" # possible misassemblies","quality":"Less is better","values":[0,0,0,0,0,0],"isMain":false},{"metricName":"# local misassemblies","quality":"Less is better","values":[1,1,1,7,3,2],"isMain":false},{"metricName":"# scaffold gap ext. mis.","quality":"Less is better","values":[0,0,0,0,0,0],"isMain":false},{"metricName":"# scaffold gap loc. mis.","quality":"Less is better","values":[0,0,0,0,0,0],"isMain":false}]],["Unaligned",[]],["Mismatches",[]],["Statistics without reference",[]],["K-mer-based statistics",[]],["Predicted genes",[]],["Similarity statistics",[]],["Reference statistics",[]]],[["Genome statistics",[]],["Reads mapping",[]],["Misassemblies",[{"metricName":"# misassemblies","quality":"Less is better","values":[4,8,0,4,6,3],"isMain":true},{"metricName":" # relocations","quality":"Less is better","values":[0,0,0,0,0,0],"isMain":false},{"metricName":" # translocations","quality":"Less is better","values":[0,1,0,0,0,0],"isMain":false},{"metricName":" # inversions","quality":"Less is better","values":[0,0,0,0,0,0],"isMain":false},{"metricName":" # interspecies translocations","quality":"Less is better","values":[4,7,0,4,6,3],"isMain":false},{"metricName":"# possibly misassembled contigs","quality":"Less is better","values":[0,0,0,0,0,0],"isMain":false},{"metricName":" # possible misassemblies","quality":"Less is better","values":[0,0,0,0,0,0],"isMain":false},{"metricName":"# local misassemblies","quality":"Less is better","values":[0,0,0,0,0,0],"isMain":false},{"metricName":"# scaffold gap ext. mis.","quality":"Less is better","values":[0,0,0,0,0,0],"isMain":false},{"metricName":"# scaffold gap loc. mis.","quality":"Less is better","values":[0,0,0,0,0,0],"isMain":false}]],["Unaligned",[]],["Mismatches",[]],["Statistics without reference",[]],["K-mer-based statistics",[]],["Predicted genes",[]],["Similarity statistics",[]],["Reference statistics",[]]],[["Genome statistics",[]],["Reads mapping",[]],["Misassemblies",[{"metricName":"# misassemblies","quality":"Less is better","values":[67,75,23,73,65,29],"isMain":true},{"metricName":" # relocations","quality":"Less is better","values":[0,1,2,0,0,1],"isMain":false},{"metricName":" # translocations","quality":"Less is better","values":[4,3,9,7,6,3],"isMain":false},{"metricName":" # inversions","quality":"Less is better","values":[0,2,4,5,3,0],"isMain":false},{"metricName":" # interspecies translocations","quality":"Less is better","values":[63,69,8,61,56,25],"isMain":false},{"metricName":"# possibly misassembled contigs","quality":"Less is better","values":[1,0,0,0,0,0],"isMain":false},{"metricName":" # possible misassemblies","quality":"Less is better","values":[1,0,0,0,0,0],"isMain":false},{"metricName":"# local misassemblies","quality":"Less is better","values":[3,2,5,6,5,2],"isMain":false},{"metricName":"# scaffold gap ext. mis.","quality":"Less is better","values":[0,0,0,0,0,0],"isMain":false},{"metricName":"# scaffold gap loc. mis.","quality":"Less is better","values":[0,0,0,0,0,0],"isMain":false}]],["Unaligned",[]],["Mismatches",[]],["Statistics without reference",[]],["K-mer-based statistics",[]],["Predicted genes",[]],["Similarity statistics",[]],["Reference statistics",[]]],[["Genome statistics",[]],["Reads mapping",[]],["Misassemblies",[{"metricName":"# misassemblies","quality":"Less is better","values":[6,7,6,11,14,4],"isMain":true},{"metricName":" # relocations","quality":"Less is better","values":[0,0,0,0,0,0],"isMain":false},{"metricName":" # translocations","quality":"Less is better","values":[0,0,0,0,0,0],"isMain":false},{"metricName":" # inversions","quality":"Less is better","values":[0,0,0,0,0,1],"isMain":false},{"metricName":" # interspecies translocations","quality":"Less is better","values":[6,7,6,11,14,3],"isMain":false},{"metricName":"# possibly misassembled contigs","quality":"Less is better","values":[0,0,0,0,0,0],"isMain":false},{"metricName":" # possible misassemblies","quality":"Less is better","values":[0,0,0,0,0,0],"isMain":false},{"metricName":"# local misassemblies","quality":"Less is better","values":[0,1,0,0,1,0],"isMain":false},{"metricName":"# scaffold gap ext. mis.","quality":"Less is better","values":[0,0,0,0,0,0],"isMain":false},{"metricName":"# scaffold gap loc. mis.","quality":"Less is better","values":[0,0,0,0,0,0],"isMain":false}]],["Unaligned",[]],["Mismatches",[]],["Statistics without reference",[]],["K-mer-based statistics",[]],["Predicted genes",[]],["Similarity statistics",[]],["Reference statistics",[]]],[["Genome statistics",[]],["Reads mapping",[]],["Misassemblies",[{"metricName":"# misassemblies","quality":"Less is better","values":[9,20,10,28,30,2],"isMain":true},{"metricName":" # relocations","quality":"Less is better","values":[0,1,0,0,1,0],"isMain":false},{"metricName":" # translocations","quality":"Less is better","values":[3,6,4,7,5,0],"isMain":false},{"metricName":" # inversions","quality":"Less is better","values":[1,0,1,1,0,0],"isMain":false},{"metricName":" # interspecies translocations","quality":"Less is better","values":[5,13,5,20,24,2],"isMain":false},{"metricName":"# possibly misassembled contigs","quality":"Less is better","values":[0,0,0,0,0,0],"isMain":false},{"metricName":" # possible misassemblies","quality":"Less is better","values":[0,0,0,0,0,0],"isMain":false},{"metricName":"# local misassemblies","quality":"Less is better","values":[0,9,4,6,6,0],"isMain":false},{"metricName":"# scaffold gap ext. mis.","quality":"Less is better","values":[0,0,0,0,0,0],"isMain":false},{"metricName":"# scaffold gap loc. mis.","quality":"Less is better","values":[0,0,0,0,0,0],"isMain":false}]],["Unaligned",[]],["Mismatches",[]],["Statistics without reference",[]],["K-mer-based statistics",[]],["Predicted genes",[]],["Similarity statistics",[]],["Reference statistics",[]]],[["Genome statistics",[]],["Reads mapping",[]],["Misassemblies",[{"metricName":"# misassemblies","quality":"Less is better","values":[58,99,34,69,108,25],"isMain":true},{"metricName":" # relocations","quality":"Less is better","values":[0,1,0,2,3,0],"isMain":false},{"metricName":" # translocations","quality":"Less is better","values":[0,6,5,6,4,1],"isMain":false},{"metricName":" # inversions","quality":"Less is better","values":[1,1,1,0,1,0],"isMain":false},{"metricName":" # interspecies translocations","quality":"Less is better","values":[57,91,28,61,100,24],"isMain":false},{"metricName":"# possibly misassembled contigs","quality":"Less is better","values":[0,1,0,0,0,0],"isMain":false},{"metricName":" # possible misassemblies","quality":"Less is better","values":[0,1,0,0,0,0],"isMain":false},{"metricName":"# local misassemblies","quality":"Less is better","values":[1,1,0,0,1,0],"isMain":false},{"metricName":"# scaffold gap ext. mis.","quality":"Less is better","values":[0,0,0,0,0,0],"isMain":false},{"metricName":"# scaffold gap loc. mis.","quality":"Less is better","values":[0,0,0,0,0,0],"isMain":false}]],["Unaligned",[]],["Mismatches",[]],["Statistics without reference",[]],["K-mer-based statistics",[]],["Predicted genes",[]],["Similarity statistics",[]],["Reference statistics",[]]],[["Genome statistics",[]],["Reads mapping",[]],["Misassemblies",[{"metricName":"# misassemblies","quality":"Less is better","values":[2,2,1,1,1,1],"isMain":true},{"metricName":" # relocations","quality":"Less is better","values":[0,0,0,0,0,0],"isMain":false},{"metricName":" # translocations","quality":"Less is better","values":[0,0,0,0,0,0],"isMain":false},{"metricName":" # inversions","quality":"Less is better","values":[0,0,0,0,0,0],"isMain":false},{"metricName":" # interspecies translocations","quality":"Less is better","values":[2,2,1,1,1,1],"isMain":false},{"metricName":"# possibly misassembled contigs","quality":"Less is better","values":[0,0,0,0,0,0],"isMain":false},{"metricName":" # possible misassemblies","quality":"Less is better","values":[0,0,0,0,0,0],"isMain":false},{"metricName":"# local misassemblies","quality":"Less is better","values":[0,0,0,0,0,0],"isMain":false},{"metricName":"# scaffold gap ext. mis.","quality":"Less is better","values":[0,0,0,0,0,0],"isMain":false},{"metricName":"# scaffold gap loc. mis.","quality":"Less is better","values":[0,0,0,0,0,0],"isMain":false}]],["Unaligned",[]],["Mismatches",[]],["Statistics without reference",[]],["K-mer-based statistics",[]],["Predicted genes",[]],["Similarity statistics",[]],["Reference statistics",[]]],[["Genome statistics",[]],["Reads mapping",[]],["Misassemblies",[{"metricName":"# misassemblies","quality":"Less is better","values":[31,34,6,25,29,25],"isMain":true},{"metricName":" # relocations","quality":"Less is better","values":[0,1,2,0,1,1],"isMain":false},{"metricName":" # translocations","quality":"Less is better","values":[3,5,0,5,3,3],"isMain":false},{"metricName":" # inversions","quality":"Less is better","values":[1,1,0,0,2,0],"isMain":false},{"metricName":" # interspecies translocations","quality":"Less is better","values":[27,27,4,20,23,21],"isMain":false},{"metricName":"# possibly misassembled contigs","quality":"Less is better","values":[0,0,0,0,0,0],"isMain":false},{"metricName":" # possible misassemblies","quality":"Less is better","values":[0,0,0,0,0,0],"isMain":false},{"metricName":"# local misassemblies","quality":"Less is better","values":[0,1,0,1,0,0],"isMain":false},{"metricName":"# scaffold gap ext. mis.","quality":"Less is better","values":[0,0,0,0,0,0],"isMain":false},{"metricName":"# scaffold gap loc. mis.","quality":"Less is better","values":[0,0,0,0,0,0],"isMain":false}]],["Unaligned",[]],["Mismatches",[]],["Statistics without reference",[]],["K-mer-based statistics",[]],["Predicted genes",[]],["Similarity statistics",[]],["Reference statistics",[]]],[["Genome statistics",[]],["Reads mapping",[]],["Misassemblies",[{"metricName":"# misassemblies","quality":"Less is better","values":[0,3,2,0,4,2],"isMain":true},{"metricName":" # relocations","quality":"Less is better","values":[0,0,0,0,0,0],"isMain":false},{"metricName":" # translocations","quality":"Less is better","values":[0,0,0,0,1,0],"isMain":false},{"metricName":" # inversions","quality":"Less is better","values":[0,0,0,0,0,0],"isMain":false},{"metricName":" # interspecies translocations","quality":"Less is better","values":[0,3,2,0,3,2],"isMain":false},{"metricName":"# possibly misassembled contigs","quality":"Less is better","values":[0,0,0,0,0,0],"isMain":false},{"metricName":" # possible misassemblies","quality":"Less is better","values":[0,0,0,0,0,0],"isMain":false},{"metricName":"# local misassemblies","quality":"Less is better","values":[0,0,0,0,0,0],"isMain":false},{"metricName":"# scaffold gap ext. mis.","quality":"Less is better","values":[0,0,0,0,0,0],"isMain":false},{"metricName":"# scaffold gap loc. mis.","quality":"Less is better","values":[0,0,0,0,0,0],"isMain":false}]],["Unaligned",[]],["Mismatches",[]],["Statistics without reference",[]],["K-mer-based statistics",[]],["Predicted genes",[]],["Similarity statistics",[]],["Reference statistics",[]]],[["Genome statistics",[]],["Reads mapping",[]],["Misassemblies",[{"metricName":"# misassemblies","quality":"Less is better","values":[10,9,3,14,8,4],"isMain":true},{"metricName":" # relocations","quality":"Less is better","values":[0,0,0,0,0,0],"isMain":false},{"metricName":" # translocations","quality":"Less is better","values":[0,0,0,1,0,0],"isMain":false},{"metricName":" # inversions","quality":"Less is better","values":[0,0,0,0,0,0],"isMain":false},{"metricName":" # interspecies translocations","quality":"Less is better","values":[10,9,3,13,8,4],"isMain":false},{"metricName":"# possibly misassembled contigs","quality":"Less is better","values":[0,0,0,0,0,0],"isMain":false},{"metricName":" # possible misassemblies","quality":"Less is better","values":[0,0,0,0,0,0],"isMain":false},{"metricName":"# local misassemblies","quality":"Less is better","values":[0,0,0,0,0,0],"isMain":false},{"metricName":"# scaffold gap ext. mis.","quality":"Less is better","values":[0,0,0,0,0,0],"isMain":false},{"metricName":"# scaffold gap loc. mis.","quality":"Less is better","values":[0,0,0,0,0,0],"isMain":false}]],["Unaligned",[]],["Mismatches",[]],["Statistics without reference",[]],["K-mer-based statistics",[]],["Predicted genes",[]],["Similarity statistics",[]],["Reference statistics",[]]],[["Genome statistics",[]],["Reads mapping",[]],["Misassemblies",[{"metricName":"# misassemblies","quality":"Less is better","values":[11,42,32,45,42,4],"isMain":true},{"metricName":" # relocations","quality":"Less is better","values":[0,0,2,1,0,0],"isMain":false},{"metricName":" # translocations","quality":"Less is better","values":[1,1,2,1,3,1],"isMain":false},{"metricName":" # inversions","quality":"Less is better","values":[0,1,1,0,1,0],"isMain":false},{"metricName":" # interspecies translocations","quality":"Less is better","values":[10,40,27,43,38,3],"isMain":false},{"metricName":"# possibly misassembled contigs","quality":"Less is better","values":[1,0,0,1,0,0],"isMain":false},{"metricName":" # possible misassemblies","quality":"Less is better","values":[1,0,0,1,0,0],"isMain":false},{"metricName":"# local misassemblies","quality":"Less is better","values":[0,2,0,1,6,0],"isMain":false},{"metricName":"# scaffold gap ext. mis.","quality":"Less is better","values":[0,0,0,0,0,0],"isMain":false},{"metricName":"# scaffold gap loc. mis.","quality":"Less is better","values":[0,0,0,0,0,0],"isMain":false}]],["Unaligned",[]],["Mismatches",[]],["Statistics without reference",[]],["K-mer-based statistics",[]],["Predicted genes",[]],["Similarity statistics",[]],["Reference statistics",[]]],[["Genome statistics",[]],["Reads mapping",[]],["Misassemblies",[{"metricName":"# misassemblies","quality":"Less is better","values":[5,11,5,22,7,8],"isMain":true},{"metricName":" # relocations","quality":"Less is better","values":[0,0,0,0,0,0],"isMain":false},{"metricName":" # translocations","quality":"Less is better","values":[0,0,1,0,0,0],"isMain":false},{"metricName":" # inversions","quality":"Less is better","values":[0,0,0,0,0,0],"isMain":false},{"metricName":" # interspecies translocations","quality":"Less is better","values":[5,11,4,22,7,8],"isMain":false},{"metricName":"# possibly misassembled contigs","quality":"Less is better","values":[0,0,0,0,0,0],"isMain":false},{"metricName":" # possible misassemblies","quality":"Less is better","values":[0,0,0,0,0,0],"isMain":false},{"metricName":"# local misassemblies","quality":"Less is better","values":[0,0,0,1,1,0],"isMain":false},{"metricName":"# scaffold gap ext. mis.","quality":"Less is better","values":[0,0,0,0,0,0],"isMain":false},{"metricName":"# scaffold gap loc. mis.","quality":"Less is better","values":[0,0,0,0,0,0],"isMain":false}]],["Unaligned",[]],["Mismatches",[]],["Statistics without reference",[]],["K-mer-based statistics",[]],["Predicted genes",[]],["Similarity statistics",[]],["Reference statistics",[]]],[["Genome statistics",[]],["Reads mapping",[]],["Misassemblies",[{"metricName":"# misassemblies","quality":"Less is better","values":[1,1,2,4,3,0],"isMain":true},{"metricName":" # relocations","quality":"Less is better","values":[0,0,0,0,0,0],"isMain":false},{"metricName":" # translocations","quality":"Less is better","values":[0,0,0,0,0,0],"isMain":false},{"metricName":" # inversions","quality":"Less is better","values":[0,0,0,0,0,0],"isMain":false},{"metricName":" # interspecies translocations","quality":"Less is better","values":[1,1,2,4,3,0],"isMain":false},{"metricName":"# possibly misassembled contigs","quality":"Less is better","values":[1,0,0,0,0,0],"isMain":false},{"metricName":" # possible misassemblies","quality":"Less is better","values":[1,0,0,0,0,0],"isMain":false},{"metricName":"# local misassemblies","quality":"Less is better","values":[0,0,0,0,0,0],"isMain":false},{"metricName":"# scaffold gap ext. mis.","quality":"Less is better","values":[0,0,0,0,0,0],"isMain":false},{"metricName":"# scaffold gap loc. mis.","quality":"Less is better","values":[0,0,0,0,0,0],"isMain":false}]],["Unaligned",[]],["Mismatches",[]],["Statistics without reference",[]],["K-mer-based statistics",[]],["Predicted genes",[]],["Similarity statistics",[]],["Reference statistics",[]]],[["Genome statistics",[]],["Reads mapping",[]],["Misassemblies",[{"metricName":"# misassemblies","quality":"Less is better","values":[47,64,12,49,55,19],"isMain":true},{"metricName":" # relocations","quality":"Less is better","values":[1,2,0,0,1,0],"isMain":false},{"metricName":" # translocations","quality":"Less is better","values":[1,2,0,2,2,1],"isMain":false},{"metricName":" # inversions","quality":"Less is better","values":[0,1,0,1,1,0],"isMain":false},{"metricName":" # interspecies translocations","quality":"Less is better","values":[45,59,12,46,51,18],"isMain":false},{"metricName":"# possibly misassembled contigs","quality":"Less is better","values":[0,0,0,0,0,0],"isMain":false},{"metricName":" # possible misassemblies","quality":"Less is better","values":[0,0,0,0,0,0],"isMain":false},{"metricName":"# local misassemblies","quality":"Less is better","values":[3,3,1,2,0,1],"isMain":false},{"metricName":"# scaffold gap ext. mis.","quality":"Less is better","values":[0,0,0,0,0,0],"isMain":false},{"metricName":"# scaffold gap loc. mis.","quality":"Less is better","values":[0,0,0,0,0,0],"isMain":false}]],["Unaligned",[]],["Mismatches",[]],["Statistics without reference",[]],["K-mer-based statistics",[]],["Predicted genes",[]],["Similarity statistics",[]],["Reference statistics",[]]],[["Genome statistics",[]],["Reads mapping",[]],["Misassemblies",[{"metricName":"# misassemblies","quality":"Less is better","values":[44,54,14,33,43,34],"isMain":true},{"metricName":" # relocations","quality":"Less is better","values":[1,2,1,0,2,0],"isMain":false},{"metricName":" # translocations","quality":"Less is better","values":[6,7,3,2,7,2],"isMain":false},{"metricName":" # inversions","quality":"Less is better","values":[1,1,2,0,1,0],"isMain":false},{"metricName":" # interspecies translocations","quality":"Less is better","values":[36,44,8,31,33,32],"isMain":false},{"metricName":"# possibly misassembled contigs","quality":"Less is better","values":[0,0,1,0,0,1],"isMain":false},{"metricName":" # possible misassemblies","quality":"Less is better","values":[0,0,1,0,0,1],"isMain":false},{"metricName":"# local misassemblies","quality":"Less is better","values":[0,0,0,1,1,0],"isMain":false},{"metricName":"# scaffold gap ext. mis.","quality":"Less is better","values":[0,0,0,0,0,0],"isMain":false},{"metricName":"# scaffold gap loc. mis.","quality":"Less is better","values":[0,0,0,0,0,0],"isMain":false}]],["Unaligned",[]],["Mismatches",[]],["Statistics without reference",[]],["K-mer-based statistics",[]],["Predicted genes",[]],["Similarity statistics",[]],["Reference statistics",[]]],[["Genome statistics",[]],["Reads mapping",[]],["Misassemblies",[{"metricName":"# misassemblies","quality":"Less is better","values":[0,0,0,1,0,0],"isMain":true},{"metricName":" # relocations","quality":"Less is better","values":[0,0,0,0,0,0],"isMain":false},{"metricName":" # translocations","quality":"Less is better","values":[0,0,0,0,0,0],"isMain":false},{"metricName":" # inversions","quality":"Less is better","values":[0,0,0,0,0,0],"isMain":false},{"metricName":" # interspecies translocations","quality":"Less is better","values":[0,0,0,1,0,0],"isMain":false},{"metricName":"# possibly misassembled contigs","quality":"Less is better","values":[0,0,0,0,0,0],"isMain":false},{"metricName":" # possible misassemblies","quality":"Less is better","values":[0,0,0,0,0,0],"isMain":false},{"metricName":"# local misassemblies","quality":"Less is better","values":[0,0,0,0,0,0],"isMain":false},{"metricName":"# scaffold gap ext. mis.","quality":"Less is better","values":[0,0,0,0,0,0],"isMain":false},{"metricName":"# scaffold gap loc. mis.","quality":"Less is better","values":[0,0,0,0,0,0],"isMain":false}]],["Unaligned",[]],["Mismatches",[]],["Statistics without reference",[]],["K-mer-based statistics",[]],["Predicted genes",[]],["Similarity statistics",[]],["Reference statistics",[]]],[["Genome statistics",[]],["Reads mapping",[]],["Misassemblies",[{"metricName":"# misassemblies","quality":"Less is better","values":[6,14,11,20,19,6],"isMain":true},{"metricName":" # relocations","quality":"Less is better","values":[0,1,2,1,2,0],"isMain":false},{"metricName":" # translocations","quality":"Less is better","values":[3,7,5,5,9,3],"isMain":false},{"metricName":" # inversions","quality":"Less is better","values":[0,0,0,0,0,0],"isMain":false},{"metricName":" # interspecies translocations","quality":"Less is better","values":[3,6,4,14,8,3],"isMain":false},{"metricName":"# possibly misassembled contigs","quality":"Less is better","values":[0,0,0,0,0,0],"isMain":false},{"metricName":" # possible misassemblies","quality":"Less is better","values":[0,0,0,0,0,0],"isMain":false},{"metricName":"# local misassemblies","quality":"Less is better","values":[0,2,2,1,1,0],"isMain":false},{"metricName":"# scaffold gap ext. mis.","quality":"Less is better","values":[0,0,0,0,0,0],"isMain":false},{"metricName":"# scaffold gap loc. mis.","quality":"Less is better","values":[0,0,0,0,0,0],"isMain":false}]],["Unaligned",[]],["Mismatches",[]],["Statistics without reference",[]],["K-mer-based statistics",[]],["Predicted genes",[]],["Similarity statistics",[]],["Reference statistics",[]]],[["Genome statistics",[]],["Reads mapping",[]],["Misassemblies",[{"metricName":"# misassemblies","quality":"Less is better","values":[8,15,5,10,10,3],"isMain":true},{"metricName":" # relocations","quality":"Less is better","values":[0,0,0,1,0,0],"isMain":false},{"metricName":" # translocations","quality":"Less is better","values":[0,3,3,1,2,1],"isMain":false},{"metricName":" # inversions","quality":"Less is better","values":[0,0,0,0,0,0],"isMain":false},{"metricName":" # interspecies translocations","quality":"Less is better","values":[8,12,2,8,8,2],"isMain":false},{"metricName":"# possibly misassembled contigs","quality":"Less is better","values":[0,0,0,0,0,0],"isMain":false},{"metricName":" # possible misassemblies","quality":"Less is better","values":[0,0,0,0,0,0],"isMain":false},{"metricName":"# local misassemblies","quality":"Less is better","values":[0,0,0,0,0,0],"isMain":false},{"metricName":"# scaffold gap ext. mis.","quality":"Less is better","values":[0,0,0,0,0,0],"isMain":false},{"metricName":"# scaffold gap loc. mis.","quality":"Less is better","values":[0,0,0,0,0,0],"isMain":false}]],["Unaligned",[]],["Mismatches",[]],["Statistics without reference",[]],["K-mer-based statistics",[]],["Predicted genes",[]],["Similarity statistics",[]],["Reference statistics",[]]],[["Genome statistics",[]],["Reads mapping",[]],["Misassemblies",[{"metricName":"# misassemblies","quality":"Less is better","values":[12,22,15,36,26,8],"isMain":true},{"metricName":" # relocations","quality":"Less is better","values":[0,0,0,1,1,0],"isMain":false},{"metricName":" # translocations","quality":"Less is better","values":[8,14,12,11,18,4],"isMain":false},{"metricName":" # inversions","quality":"Less is better","values":[0,2,0,0,0,1],"isMain":false},{"metricName":" # interspecies translocations","quality":"Less is better","values":[4,6,3,24,7,3],"isMain":false},{"metricName":"# possibly misassembled contigs","quality":"Less is better","values":[0,0,0,0,0,0],"isMain":false},{"metricName":" # possible misassemblies","quality":"Less is better","values":[0,0,0,0,0,0],"isMain":false},{"metricName":"# local misassemblies","quality":"Less is better","values":[4,8,6,6,10,1],"isMain":false},{"metricName":"# scaffold gap ext. mis.","quality":"Less is better","values":[0,0,0,0,0,0],"isMain":false},{"metricName":"# scaffold gap loc. mis.","quality":"Less is better","values":[0,0,0,0,0,0],"isMain":false}]],["Unaligned",[]],["Mismatches",[]],["Statistics without reference",[]],["K-mer-based statistics",[]],["Predicted genes",[]],["Similarity statistics",[]],["Reference statistics",[]]],[["Genome statistics",[]],["Reads mapping",[]],["Misassemblies",[{"metricName":"# misassemblies","quality":"Less is better","values":[23,31,3,23,25,21],"isMain":true},{"metricName":" # relocations","quality":"Less is better","values":[0,1,0,1,0,0],"isMain":false},{"metricName":" # translocations","quality":"Less is better","values":[0,0,1,0,0,0],"isMain":false},{"metricName":" # inversions","quality":"Less is better","values":[0,0,0,0,2,0],"isMain":false},{"metricName":" # interspecies translocations","quality":"Less is better","values":[23,30,2,22,23,21],"isMain":false},{"metricName":"# possibly misassembled contigs","quality":"Less is better","values":[0,0,0,0,0,0],"isMain":false},{"metricName":" # possible misassemblies","quality":"Less is better","values":[0,0,0,0,0,0],"isMain":false},{"metricName":"# local misassemblies","quality":"Less is better","values":[0,0,0,1,0,0],"isMain":false},{"metricName":"# scaffold gap ext. mis.","quality":"Less is better","values":[0,0,0,0,0,0],"isMain":false},{"metricName":"# scaffold gap loc. mis.","quality":"Less is better","values":[0,0,0,0,0,0],"isMain":false}]],["Unaligned",[]],["Mismatches",[]],["Statistics without reference",[]],["K-mer-based statistics",[]],["Predicted genes",[]],["Similarity statistics",[]],["Reference statistics",[]]],[["Genome statistics",[]],["Reads mapping",[]],["Misassemblies",[{"metricName":"# misassemblies","quality":"Less is better","values":[3,3,2,7,2,0],"isMain":true},{"metricName":" # relocations","quality":"Less is better","values":[0,0,0,0,0,0],"isMain":false},{"metricName":" # translocations","quality":"Less is better","values":[0,0,2,1,0,0],"isMain":false},{"metricName":" # inversions","quality":"Less is better","values":[0,0,0,0,0,0],"isMain":false},{"metricName":" # interspecies translocations","quality":"Less is better","values":[3,3,0,6,2,0],"isMain":false},{"metricName":"# possibly misassembled contigs","quality":"Less is better","values":[0,0,0,0,0,0],"isMain":false},{"metricName":" # possible misassemblies","quality":"Less is better","values":[0,0,0,0,0,0],"isMain":false},{"metricName":"# local misassemblies","quality":"Less is better","values":[0,0,0,0,0,0],"isMain":false},{"metricName":"# scaffold gap ext. mis.","quality":"Less is better","values":[0,0,0,0,0,0],"isMain":false},{"metricName":"# scaffold gap loc. mis.","quality":"Less is better","values":[0,0,0,0,0,0],"isMain":false}]],["Unaligned",[]],["Mismatches",[]],["Statistics without reference",[]],["K-mer-based statistics",[]],["Predicted genes",[]],["Similarity statistics",[]],["Reference statistics",[]]],[["Genome statistics",[]],["Reads mapping",[]],["Misassemblies",[{"metricName":"# misassemblies","quality":"Less is better","values":[19,24,21,38,28,12],"isMain":true},{"metricName":" # relocations","quality":"Less is better","values":[0,1,4,1,0,1],"isMain":false},{"metricName":" # translocations","quality":"Less is better","values":[0,1,1,2,3,0],"isMain":false},{"metricName":" # inversions","quality":"Less is better","values":[0,0,7,2,0,0],"isMain":false},{"metricName":" # interspecies translocations","quality":"Less is better","values":[19,22,9,33,25,11],"isMain":false},{"metricName":"# possibly misassembled contigs","quality":"Less is better","values":[0,0,0,0,0,0],"isMain":false},{"metricName":" # possible misassemblies","quality":"Less is better","values":[0,0,0,0,0,0],"isMain":false},{"metricName":"# local misassemblies","quality":"Less is better","values":[2,1,4,3,0,0],"isMain":false},{"metricName":"# scaffold gap ext. mis.","quality":"Less is better","values":[0,0,0,0,0,0],"isMain":false},{"metricName":"# scaffold gap loc. mis.","quality":"Less is better","values":[0,0,0,0,0,0],"isMain":false}]],["Unaligned",[]],["Mismatches",[]],["Statistics without reference",[]],["K-mer-based statistics",[]],["Predicted genes",[]],["Similarity statistics",[]],["Reference statistics",[]]],[["Genome statistics",[]],["Reads mapping",[]],["Misassemblies",[{"metricName":"# misassemblies","quality":"Less is better","values":[39,32,20,48,34,15],"isMain":true},{"metricName":" # relocations","quality":"Less is better","values":[0,0,2,0,0,0],"isMain":false},{"metricName":" # translocations","quality":"Less is better","values":[15,15,14,18,21,5],"isMain":false},{"metricName":" # inversions","quality":"Less is better","values":[1,0,3,3,1,0],"isMain":false},{"metricName":" # interspecies translocations","quality":"Less is better","values":[23,17,1,27,12,10],"isMain":false},{"metricName":"# possibly misassembled contigs","quality":"Less is better","values":[0,0,0,0,0,0],"isMain":false},{"metricName":" # possible misassemblies","quality":"Less is better","values":[0,0,0,0,0,0],"isMain":false},{"metricName":"# local misassemblies","quality":"Less is better","values":[1,5,4,2,2,1],"isMain":false},{"metricName":"# scaffold gap ext. mis.","quality":"Less is better","values":[0,0,0,0,0,0],"isMain":false},{"metricName":"# scaffold gap loc. mis.","quality":"Less is better","values":[0,0,0,0,0,0],"isMain":false}]],["Unaligned",[]],["Mismatches",[]],["Statistics without reference",[]],["K-mer-based statistics",[]],["Predicted genes",[]],["Similarity statistics",[]],["Reference statistics",[]]],[["Genome statistics",[]],["Reads mapping",[]],["Misassemblies",[{"metricName":"# misassemblies","quality":"Less is better","values":[26,41,16,28,35,18],"isMain":true},{"metricName":" # relocations","quality":"Less is better","values":[1,0,2,1,0,0],"isMain":false},{"metricName":" # translocations","quality":"Less is better","values":[1,1,2,2,1,2],"isMain":false},{"metricName":" # inversions","quality":"Less is better","values":[0,0,0,0,0,0],"isMain":false},{"metricName":" # interspecies translocations","quality":"Less is better","values":[24,40,12,25,34,16],"isMain":false},{"metricName":"# possibly misassembled contigs","quality":"Less is better","values":[0,0,0,0,0,0],"isMain":false},{"metricName":" # possible misassemblies","quality":"Less is better","values":[0,0,0,0,0,0],"isMain":false},{"metricName":"# local misassemblies","quality":"Less is better","values":[0,0,1,0,0,0],"isMain":false},{"metricName":"# scaffold gap ext. mis.","quality":"Less is better","values":[0,0,0,0,0,0],"isMain":false},{"metricName":"# scaffold gap loc. mis.","quality":"Less is better","values":[0,0,0,0,0,0],"isMain":false}]],["Unaligned",[]],["Mismatches",[]],["Statistics without reference",[]],["K-mer-based statistics",[]],["Predicted genes",[]],["Similarity statistics",[]],["Reference statistics",[]]],[["Genome statistics",[]],["Reads mapping",[]],["Misassemblies",[{"metricName":"# misassemblies","quality":"Less is better","values":[33,44,17,51,42,11],"isMain":true},{"metricName":" # relocations","quality":"Less is better","values":[0,1,0,2,1,0],"isMain":false},{"metricName":" # translocations","quality":"Less is better","values":[2,3,2,1,3,0],"isMain":false},{"metricName":" # inversions","quality":"Less is better","values":[0,0,0,0,0,0],"isMain":false},{"metricName":" # interspecies translocations","quality":"Less is better","values":[31,40,15,48,38,11],"isMain":false},{"metricName":"# possibly misassembled contigs","quality":"Less is better","values":[0,0,0,2,0,0],"isMain":false},{"metricName":" # possible misassemblies","quality":"Less is better","values":[0,0,0,2,0,0],"isMain":false},{"metricName":"# local misassemblies","quality":"Less is better","values":[0,0,0,0,1,0],"isMain":false},{"metricName":"# scaffold gap ext. mis.","quality":"Less is better","values":[0,0,0,0,0,0],"isMain":false},{"metricName":"# scaffold gap loc. mis.","quality":"Less is better","values":[0,0,0,0,0,0],"isMain":false}]],["Unaligned",[]],["Mismatches",[]],["Statistics without reference",[]],["K-mer-based statistics",[]],["Predicted genes",[]],["Similarity statistics",[]],["Reference statistics",[]]],[["Genome statistics",[]],["Reads mapping",[]],["Misassemblies",[{"metricName":"# misassemblies","quality":"Less is better","values":[3,5,0,7,4,1],"isMain":true},{"metricName":" # relocations","quality":"Less is better","values":[0,0,0,0,0,0],"isMain":false},{"metricName":" # translocations","quality":"Less is better","values":[0,0,0,0,0,0],"isMain":false},{"metricName":" # inversions","quality":"Less is better","values":[0,0,0,0,0,0],"isMain":false},{"metricName":" # interspecies translocations","quality":"Less is better","values":[3,5,0,7,4,1],"isMain":false},{"metricName":"# possibly misassembled contigs","quality":"Less is better","values":[0,0,0,0,0,0],"isMain":false},{"metricName":" # possible misassemblies","quality":"Less is better","values":[0,0,0,0,0,0],"isMain":false},{"metricName":"# local misassemblies","quality":"Less is better","values":[0,0,0,0,0,0],"isMain":false},{"metricName":"# scaffold gap ext. mis.","quality":"Less is better","values":[0,0,0,0,0,0],"isMain":false},{"metricName":"# scaffold gap loc. mis.","quality":"Less is better","values":[0,0,0,0,0,0],"isMain":false}]],["Unaligned",[]],["Mismatches",[]],["Statistics without reference",[]],["K-mer-based statistics",[]],["Predicted genes",[]],["Similarity statistics",[]],["Reference statistics",[]]],[["Genome statistics",[]],["Reads mapping",[]],["Misassemblies",[{"metricName":"# misassemblies","quality":"Less is better","values":[67,80,9,68,54,51],"isMain":true},{"metricName":" # relocations","quality":"Less is better","values":[2,4,1,0,4,2],"isMain":false},{"metricName":" # translocations","quality":"Less is better","values":[5,6,5,12,6,4],"isMain":false},{"metricName":" # inversions","quality":"Less is better","values":[0,0,1,6,0,0],"isMain":false},{"metricName":" # interspecies translocations","quality":"Less is better","values":[60,70,2,50,44,45],"isMain":false},{"metricName":"# possibly misassembled contigs","quality":"Less is better","values":[0,0,1,0,0,0],"isMain":false},{"metricName":" # possible misassemblies","quality":"Less is better","values":[0,0,1,0,0,0],"isMain":false},{"metricName":"# local misassemblies","quality":"Less is better","values":[4,4,5,3,5,4],"isMain":false},{"metricName":"# scaffold gap ext. mis.","quality":"Less is better","values":[0,0,0,0,0,0],"isMain":false},{"metricName":"# scaffold gap loc. mis.","quality":"Less is better","values":[0,0,0,0,0,0],"isMain":false}]],["Unaligned",[]],["Mismatches",[]],["Statistics without reference",[]],["K-mer-based statistics",[]],["Predicted genes",[]],["Similarity statistics",[]],["Reference statistics",[]]],[["Genome statistics",[]],["Reads mapping",[]],["Misassemblies",[{"metricName":"# misassemblies","quality":"Less is better","values":[39,86,48,88,68,23],"isMain":true},{"metricName":" # relocations","quality":"Less is better","values":[1,3,0,1,1,1],"isMain":false},{"metricName":" # translocations","quality":"Less is better","values":[10,8,16,16,11,6],"isMain":false},{"metricName":" # inversions","quality":"Less is better","values":[0,1,4,4,0,0],"isMain":false},{"metricName":" # interspecies translocations","quality":"Less is better","values":[28,74,28,67,56,16],"isMain":false},{"metricName":"# possibly misassembled contigs","quality":"Less is better","values":[0,2,0,1,1,0],"isMain":false},{"metricName":" # possible misassemblies","quality":"Less is better","values":[0,2,0,1,1,0],"isMain":false},{"metricName":"# local misassemblies","quality":"Less is better","values":[4,6,5,6,6,4],"isMain":false},{"metricName":"# scaffold gap ext. mis.","quality":"Less is better","values":[0,0,0,0,0,0],"isMain":false},{"metricName":"# scaffold gap loc. mis.","quality":"Less is better","values":[0,0,0,0,0,0],"isMain":false}]],["Unaligned",[]],["Mismatches",[]],["Statistics without reference",[]],["K-mer-based statistics",[]],["Predicted genes",[]],["Similarity statistics",[]],["Reference statistics",[]]],[["Genome statistics",[]],["Reads mapping",[]],["Misassemblies",[{"metricName":"# misassemblies","quality":"Less is better","values":[59,60,11,49,54,44],"isMain":true},{"metricName":" # relocations","quality":"Less is better","values":[1,3,1,2,3,2],"isMain":false},{"metricName":" # translocations","quality":"Less is better","values":[5,6,3,6,7,5],"isMain":false},{"metricName":" # inversions","quality":"Less is better","values":[0,1,0,1,2,1],"isMain":false},{"metricName":" # interspecies translocations","quality":"Less is better","values":[53,50,7,40,42,36],"isMain":false},{"metricName":"# possibly misassembled contigs","quality":"Less is better","values":[0,0,0,0,0,0],"isMain":false},{"metricName":" # possible misassemblies","quality":"Less is better","values":[0,0,0,0,0,0],"isMain":false},{"metricName":"# local misassemblies","quality":"Less is better","values":[3,5,2,2,4,0],"isMain":false},{"metricName":"# scaffold gap ext. mis.","quality":"Less is better","values":[0,0,0,0,0,0],"isMain":false},{"metricName":"# scaffold gap loc. mis.","quality":"Less is better","values":[0,0,0,0,0,0],"isMain":false}]],["Unaligned",[]],["Mismatches",[]],["Statistics without reference",[]],["K-mer-based statistics",[]],["Predicted genes",[]],["Similarity statistics",[]],["Reference statistics",[]]],[["Genome statistics",[]],["Reads mapping",[]],["Misassemblies",[{"metricName":"# misassemblies","quality":"Less is better","values":[24,36,1,38,24,38],"isMain":true},{"metricName":" # relocations","quality":"Less is better","values":[0,0,0,0,0,0],"isMain":false},{"metricName":" # translocations","quality":"Less is better","values":[0,0,0,1,0,1],"isMain":false},{"metricName":" # inversions","quality":"Less is better","values":[0,0,0,0,0,0],"isMain":false},{"metricName":" # interspecies translocations","quality":"Less is better","values":[24,36,1,37,24,37],"isMain":false},{"metricName":"# possibly misassembled contigs","quality":"Less is better","values":[0,0,0,0,0,0],"isMain":false},{"metricName":" # possible misassemblies","quality":"Less is better","values":[0,0,0,0,0,0],"isMain":false},{"metricName":"# local misassemblies","quality":"Less is better","values":[1,2,0,1,2,0],"isMain":false},{"metricName":"# scaffold gap ext. mis.","quality":"Less is better","values":[0,0,0,0,0,0],"isMain":false},{"metricName":"# scaffold gap loc. mis.","quality":"Less is better","values":[0,0,0,0,0,0],"isMain":false}]],["Unaligned",[]],["Mismatches",[]],["Statistics without reference",[]],["K-mer-based statistics",[]],["Predicted genes",[]],["Similarity statistics",[]],["Reference statistics",[]]],[["Genome statistics",[]],["Reads mapping",[]],["Misassemblies",[{"metricName":"# misassemblies","quality":"Less is better","values":[40,52,17,42,42,29],"isMain":true},{"metricName":" # relocations","quality":"Less is better","values":[0,0,1,2,1,1],"isMain":false},{"metricName":" # translocations","quality":"Less is better","values":[4,9,8,11,9,2],"isMain":false},{"metricName":" # inversions","quality":"Less is better","values":[0,0,1,2,0,0],"isMain":false},{"metricName":" # interspecies translocations","quality":"Less is better","values":[36,43,7,27,32,26],"isMain":false},{"metricName":"# possibly misassembled contigs","quality":"Less is better","values":[0,0,0,0,0,0],"isMain":false},{"metricName":" # possible misassemblies","quality":"Less is better","values":[0,0,0,0,0,0],"isMain":false},{"metricName":"# local misassemblies","quality":"Less is better","values":[2,1,1,3,2,0],"isMain":false},{"metricName":"# scaffold gap ext. mis.","quality":"Less is better","values":[0,0,0,0,0,0],"isMain":false},{"metricName":"# scaffold gap loc. mis.","quality":"Less is better","values":[0,0,0,0,0,0],"isMain":false}]],["Unaligned",[]],["Mismatches",[]],["Statistics without reference",[]],["K-mer-based statistics",[]],["Predicted genes",[]],["Similarity statistics",[]],["Reference statistics",[]]],[["Genome statistics",[]],["Reads mapping",[]],["Misassemblies",[{"metricName":"# misassemblies","quality":"Less is better","values":[54,67,24,50,71,22],"isMain":true},{"metricName":" # relocations","quality":"Less is better","values":[0,0,1,0,0,1],"isMain":false},{"metricName":" # translocations","quality":"Less is better","values":[3,3,3,4,5,2],"isMain":false},{"metricName":" # inversions","quality":"Less is better","values":[0,0,1,0,0,0],"isMain":false},{"metricName":" # interspecies translocations","quality":"Less is better","values":[51,64,19,46,66,19],"isMain":false},{"metricName":"# possibly misassembled contigs","quality":"Less is better","values":[0,0,0,1,1,0],"isMain":false},{"metricName":" # possible misassemblies","quality":"Less is better","values":[0,0,0,1,1,0],"isMain":false},{"metricName":"# local misassemblies","quality":"Less is better","values":[0,1,2,2,1,0],"isMain":false},{"metricName":"# scaffold gap ext. mis.","quality":"Less is better","values":[0,0,0,0,0,0],"isMain":false},{"metricName":"# scaffold gap loc. mis.","quality":"Less is better","values":[0,0,0,0,0,0],"isMain":false}]],["Unaligned",[]],["Mismatches",[]],["Statistics without reference",[]],["K-mer-based statistics",[]],["Predicted genes",[]],["Similarity statistics",[]],["Reference statistics",[]]],[["Genome statistics",[]],["Reads mapping",[]],["Misassemblies",[{"metricName":"# misassemblies","quality":"Less is better","values":[6,9,4,19,5,7],"isMain":true},{"metricName":" # relocations","quality":"Less is better","values":[2,2,3,5,1,1],"isMain":false},{"metricName":" # translocations","quality":"Less is better","values":[0,0,0,0,0,0],"isMain":false},{"metricName":" # inversions","quality":"Less is better","values":[0,0,0,0,0,0],"isMain":false},{"metricName":" # interspecies translocations","quality":"Less is better","values":[4,7,1,14,4,6],"isMain":false},{"metricName":"# possibly misassembled contigs","quality":"Less is better","values":[0,0,0,0,0,0],"isMain":false},{"metricName":" # possible misassemblies","quality":"Less is better","values":[0,0,0,0,0,0],"isMain":false},{"metricName":"# local misassemblies","quality":"Less is better","values":[0,0,0,0,0,0],"isMain":false},{"metricName":"# scaffold gap ext. mis.","quality":"Less is better","values":[0,0,0,0,0,0],"isMain":false},{"metricName":"# scaffold gap loc. mis.","quality":"Less is better","values":[0,0,0,0,0,0],"isMain":false}]],["Unaligned",[]],["Mismatches",[]],["Statistics without reference",[]],["K-mer-based statistics",[]],["Predicted genes",[]],["Similarity statistics",[]],["Reference statistics",[]]],[["Genome statistics",[]],["Reads mapping",[]],["Misassemblies",[{"metricName":"# misassemblies","quality":"Less is better","values":[0,0,0,2,0,0],"isMain":true},{"metricName":" # relocations","quality":"Less is better","values":[0,0,0,0,0,0],"isMain":false},{"metricName":" # translocations","quality":"Less is better","values":[0,0,0,0,0,0],"isMain":false},{"metricName":" # inversions","quality":"Less is better","values":[0,0,0,0,0,0],"isMain":false},{"metricName":" # interspecies translocations","quality":"Less is better","values":[0,0,0,2,0,0],"isMain":false},{"metricName":"# possibly misassembled contigs","quality":"Less is better","values":[0,0,0,0,0,0],"isMain":false},{"metricName":" # possible misassemblies","quality":"Less is better","values":[0,0,0,0,0,0],"isMain":false},{"metricName":"# local misassemblies","quality":"Less is better","values":[0,0,0,0,0,0],"isMain":false},{"metricName":"# scaffold gap ext. mis.","quality":"Less is better","values":[0,0,0,0,0,0],"isMain":false},{"metricName":"# scaffold gap loc. mis.","quality":"Less is better","values":[0,0,0,0,0,0],"isMain":false}]],["Unaligned",[]],["Mismatches",[]],["Statistics without reference",[]],["K-mer-based statistics",[]],["Predicted genes",[]],["Similarity statistics",[]],["Reference statistics",[]]],[["Genome statistics",[]],["Reads mapping",[]],["Misassemblies",[{"metricName":"# misassemblies","quality":"Less is better","values":[7,10,4,11,5,5],"isMain":true},{"metricName":" # relocations","quality":"Less is better","values":[0,1,0,2,0,0],"isMain":false},{"metricName":" # translocations","quality":"Less is better","values":[4,4,2,5,3,4],"isMain":false},{"metricName":" # inversions","quality":"Less is better","values":[0,0,0,0,0,0],"isMain":false},{"metricName":" # interspecies translocations","quality":"Less is better","values":[3,5,2,4,2,1],"isMain":false},{"metricName":"# possibly misassembled contigs","quality":"Less is better","values":[0,0,0,0,0,0],"isMain":false},{"metricName":" # possible misassemblies","quality":"Less is better","values":[0,0,0,0,0,0],"isMain":false},{"metricName":"# local misassemblies","quality":"Less is better","values":[1,1,1,1,1,1],"isMain":false},{"metricName":"# scaffold gap ext. mis.","quality":"Less is better","values":[0,0,0,0,0,0],"isMain":false},{"metricName":"# scaffold gap loc. mis.","quality":"Less is better","values":[0,0,0,0,0,0],"isMain":false}]],["Unaligned",[]],["Mismatches",[]],["Statistics without reference",[]],["K-mer-based statistics",[]],["Predicted genes",[]],["Similarity statistics",[]],["Reference statistics",[]]],[["Genome statistics",[]],["Reads mapping",[]],["Misassemblies",[{"metricName":"# misassemblies","quality":"Less is better","values":[84,26,25,49,34,5],"isMain":true},{"metricName":" # relocations","quality":"Less is better","values":[0,0,0,0,0,0],"isMain":false},{"metricName":" # translocations","quality":"Less is better","values":[4,7,7,17,16,3],"isMain":false},{"metricName":" # inversions","quality":"Less is better","values":[2,0,0,0,1,0],"isMain":false},{"metricName":" # interspecies translocations","quality":"Less is better","values":[78,19,18,32,17,2],"isMain":false},{"metricName":"# possibly misassembled contigs","quality":"Less is better","values":[0,0,0,1,0,0],"isMain":false},{"metricName":" # possible misassemblies","quality":"Less is better","values":[0,0,0,1,0,0],"isMain":false},{"metricName":"# local misassemblies","quality":"Less is better","values":[0,4,3,6,2,0],"isMain":false},{"metricName":"# scaffold gap ext. mis.","quality":"Less is better","values":[0,0,0,0,0,0],"isMain":false},{"metricName":"# scaffold gap loc. mis.","quality":"Less is better","values":[0,0,0,0,0,0],"isMain":false}]],["Unaligned",[]],["Mismatches",[]],["Statistics without reference",[]],["K-mer-based statistics",[]],["Predicted genes",[]],["Similarity statistics",[]],["Reference statistics",[]]],[["Genome statistics",[]],["Reads mapping",[]],["Misassemblies",[{"metricName":"# misassemblies","quality":"Less is better","values":[2,4,2,13,4,1],"isMain":true},{"metricName":" # relocations","quality":"Less is better","values":[0,0,0,0,0,0],"isMain":false},{"metricName":" # translocations","quality":"Less is better","values":[0,0,0,0,0,0],"isMain":false},{"metricName":" # inversions","quality":"Less is better","values":[0,0,0,0,0,0],"isMain":false},{"metricName":" # interspecies translocations","quality":"Less is better","values":[2,4,2,13,4,1],"isMain":false},{"metricName":"# possibly misassembled contigs","quality":"Less is better","values":[0,0,0,0,0,0],"isMain":false},{"metricName":" # possible misassemblies","quality":"Less is better","values":[0,0,0,0,0,0],"isMain":false},{"metricName":"# local misassemblies","quality":"Less is better","values":[0,0,0,0,0,0],"isMain":false},{"metricName":"# scaffold gap ext. mis.","quality":"Less is better","values":[0,0,0,0,0,0],"isMain":false},{"metricName":"# scaffold gap loc. mis.","quality":"Less is better","values":[0,0,0,0,0,0],"isMain":false}]],["Unaligned",[]],["Mismatches",[]],["Statistics without reference",[]],["K-mer-based statistics",[]],["Predicted genes",[]],["Similarity statistics",[]],["Reference statistics",[]]],[["Genome statistics",[]],["Reads mapping",[]],["Misassemblies",[{"metricName":"# misassemblies","quality":"Less is better","values":[96,97,89,171,137,9],"isMain":true},{"metricName":" # relocations","quality":"Less is better","values":[0,2,3,5,2,0],"isMain":false},{"metricName":" # translocations","quality":"Less is better","values":[0,34,36,69,67,6],"isMain":false},{"metricName":" # inversions","quality":"Less is better","values":[7,7,4,10,12,0],"isMain":false},{"metricName":" # interspecies translocations","quality":"Less is better","values":[89,54,46,87,56,3],"isMain":false},{"metricName":"# possibly misassembled contigs","quality":"Less is better","values":[0,1,0,1,1,0],"isMain":false},{"metricName":" # possible misassemblies","quality":"Less is better","values":[0,1,0,1,1,0],"isMain":false},{"metricName":"# local misassemblies","quality":"Less is better","values":[5,10,7,16,15,0],"isMain":false},{"metricName":"# scaffold gap ext. mis.","quality":"Less is better","values":[0,0,0,0,0,0],"isMain":false},{"metricName":"# scaffold gap loc. mis.","quality":"Less is better","values":[0,0,0,0,0,0],"isMain":false}]],["Unaligned",[]],["Mismatches",[]],["Statistics without reference",[]],["K-mer-based statistics",[]],["Predicted genes",[]],["Similarity statistics",[]],["Reference statistics",[]]],[["Genome statistics",[]],["Reads mapping",[]],["Misassemblies",[{"metricName":"# misassemblies","quality":"Less is better","values":[26,27,20,54,22,13],"isMain":true},{"metricName":" # relocations","quality":"Less is better","values":[0,1,0,0,0,0],"isMain":false},{"metricName":" # translocations","quality":"Less is better","values":[7,7,6,6,9,5],"isMain":false},{"metricName":" # inversions","quality":"Less is better","values":[0,0,4,1,0,0],"isMain":false},{"metricName":" # interspecies translocations","quality":"Less is better","values":[19,19,10,47,13,8],"isMain":false},{"metricName":"# possibly misassembled contigs","quality":"Less is better","values":[0,0,0,1,0,0],"isMain":false},{"metricName":" # possible misassemblies","quality":"Less is better","values":[0,0,0,1,0,0],"isMain":false},{"metricName":"# local misassemblies","quality":"Less is better","values":[3,1,2,3,4,0],"isMain":false},{"metricName":"# scaffold gap ext. mis.","quality":"Less is better","values":[0,0,0,0,0,0],"isMain":false},{"metricName":"# scaffold gap loc. mis.","quality":"Less is better","values":[0,0,0,0,0,0],"isMain":false}]],["Unaligned",[]],["Mismatches",[]],["Statistics without reference",[]],["K-mer-based statistics",[]],["Predicted genes",[]],["Similarity statistics",[]],["Reference statistics",[]]],[["Genome statistics",[]],["Reads mapping",[]],["Misassemblies",[{"metricName":"# misassemblies","quality":"Less is better","values":[103,125,19,111,89,79],"isMain":true},{"metricName":" # relocations","quality":"Less is better","values":[6,4,2,2,3,1],"isMain":false},{"metricName":" # translocations","quality":"Less is better","values":[10,17,5,9,15,8],"isMain":false},{"metricName":" # inversions","quality":"Less is better","values":[1,2,2,2,1,0],"isMain":false},{"metricName":" # interspecies translocations","quality":"Less is better","values":[86,102,10,98,70,70],"isMain":false},{"metricName":"# possibly misassembled contigs","quality":"Less is better","values":[0,1,0,0,0,0],"isMain":false},{"metricName":" # possible misassemblies","quality":"Less is better","values":[0,1,0,0,0,0],"isMain":false},{"metricName":"# local misassemblies","quality":"Less is better","values":[2,2,3,2,2,0],"isMain":false},{"metricName":"# scaffold gap ext. mis.","quality":"Less is better","values":[0,0,0,0,0,0],"isMain":false},{"metricName":"# scaffold gap loc. mis.","quality":"Less is better","values":[0,0,0,0,0,0],"isMain":false}]],["Unaligned",[]],["Mismatches",[]],["Statistics without reference",[]],["K-mer-based statistics",[]],["Predicted genes",[]],["Similarity statistics",[]],["Reference statistics",[]]],[["Genome statistics",[]],["Reads mapping",[]],["Misassemblies",[{"metricName":"# misassemblies","quality":"Less is better","values":[13,19,9,25,26,8],"isMain":true},{"metricName":" # relocations","quality":"Less is better","values":[0,1,0,0,1,0],"isMain":false},{"metricName":" # translocations","quality":"Less is better","values":[2,5,2,8,7,1],"isMain":false},{"metricName":" # inversions","quality":"Less is better","values":[0,0,0,1,0,0],"isMain":false},{"metricName":" # interspecies translocations","quality":"Less is better","values":[11,13,7,16,18,7],"isMain":false},{"metricName":"# possibly misassembled contigs","quality":"Less is better","values":[0,0,0,0,1,0],"isMain":false},{"metricName":" # possible misassemblies","quality":"Less is better","values":[0,0,0,0,1,0],"isMain":false},{"metricName":"# local misassemblies","quality":"Less is better","values":[0,0,3,2,1,1],"isMain":false},{"metricName":"# scaffold gap ext. mis.","quality":"Less is better","values":[0,0,0,0,0,0],"isMain":false},{"metricName":"# scaffold gap loc. mis.","quality":"Less is better","values":[0,0,0,0,0,0],"isMain":false}]],["Unaligned",[]],["Mismatches",[]],["Statistics without reference",[]],["K-mer-based statistics",[]],["Predicted genes",[]],["Similarity statistics",[]],["Reference statistics",[]]],[["Genome statistics",[]],["Reads mapping",[]],["Misassemblies",[{"metricName":"# misassemblies","quality":"Less is better","values":[3,12,11,16,12,4],"isMain":true},{"metricName":" # relocations","quality":"Less is better","values":[0,0,0,0,0,0],"isMain":false},{"metricName":" # translocations","quality":"Less is better","values":[0,0,1,1,0,0],"isMain":false},{"metricName":" # inversions","quality":"Less is better","values":[0,0,0,0,0,0],"isMain":false},{"metricName":" # interspecies translocations","quality":"Less is better","values":[3,12,10,15,12,4],"isMain":false},{"metricName":"# possibly misassembled contigs","quality":"Less is better","values":[0,0,0,0,0,0],"isMain":false},{"metricName":" # possible misassemblies","quality":"Less is better","values":[0,0,0,0,0,0],"isMain":false},{"metricName":"# local misassemblies","quality":"Less is better","values":[0,0,0,0,0,0],"isMain":false},{"metricName":"# scaffold gap ext. mis.","quality":"Less is better","values":[0,0,0,0,0,0],"isMain":false},{"metricName":"# scaffold gap loc. mis.","quality":"Less is better","values":[0,0,0,0,0,0],"isMain":false}]],["Unaligned",[]],["Mismatches",[]],["Statistics without reference",[]],["K-mer-based statistics",[]],["Predicted genes",[]],["Similarity statistics",[]],["Reference statistics",[]]],[["Genome statistics",[]],["Reads mapping",[]],["Misassemblies",[{"metricName":"# misassemblies","quality":"Less is better","values":[29,31,9,39,30,22],"isMain":true},{"metricName":" # relocations","quality":"Less is better","values":[0,0,0,0,0,0],"isMain":false},{"metricName":" # translocations","quality":"Less is better","values":[1,2,0,0,3,2],"isMain":false},{"metricName":" # inversions","quality":"Less is better","values":[0,0,0,0,0,0],"isMain":false},{"metricName":" # interspecies translocations","quality":"Less is better","values":[28,29,9,39,27,20],"isMain":false},{"metricName":"# possibly misassembled contigs","quality":"Less is better","values":[0,0,0,0,0,0],"isMain":false},{"metricName":" # possible misassemblies","quality":"Less is better","values":[0,0,0,0,0,0],"isMain":false},{"metricName":"# local misassemblies","quality":"Less is better","values":[0,1,1,0,2,0],"isMain":false},{"metricName":"# scaffold gap ext. mis.","quality":"Less is better","values":[0,0,0,0,0,0],"isMain":false},{"metricName":"# scaffold gap loc. mis.","quality":"Less is better","values":[0,0,0,0,0,0],"isMain":false}]],["Unaligned",[]],["Mismatches",[]],["Statistics without reference",[]],["K-mer-based statistics",[]],["Predicted genes",[]],["Similarity statistics",[]],["Reference statistics",[]]],[["Genome statistics",[]],["Reads mapping",[]],["Misassemblies",[{"metricName":"# misassemblies","quality":"Less is better","values":[11,15,11,28,14,13],"isMain":true},{"metricName":" # relocations","quality":"Less is better","values":[0,0,0,0,0,0],"isMain":false},{"metricName":" # translocations","quality":"Less is better","values":[10,11,9,10,11,11],"isMain":false},{"metricName":" # inversions","quality":"Less is better","values":[1,0,1,0,0,0],"isMain":false},{"metricName":" # interspecies translocations","quality":"Less is better","values":[0,4,1,18,3,2],"isMain":false},{"metricName":"# possibly misassembled contigs","quality":"Less is better","values":[0,0,0,0,0,0],"isMain":false},{"metricName":" # possible misassemblies","quality":"Less is better","values":[0,0,0,0,0,0],"isMain":false},{"metricName":"# local misassemblies","quality":"Less is better","values":[0,0,0,4,1,0],"isMain":false},{"metricName":"# scaffold gap ext. mis.","quality":"Less is better","values":[0,0,0,0,0,0],"isMain":false},{"metricName":"# scaffold gap loc. mis.","quality":"Less is better","values":[0,0,0,0,0,0],"isMain":false}]],["Unaligned",[]],["Mismatches",[]],["Statistics without reference",[]],["K-mer-based statistics",[]],["Predicted genes",[]],["Similarity statistics",[]],["Reference statistics",[]]],[["Genome statistics",[]],["Reads mapping",[]],["Misassemblies",[{"metricName":"# misassemblies","quality":"Less is better","values":[111,133,54,95,129,58],"isMain":true},{"metricName":" # relocations","quality":"Less is better","values":[0,4,2,5,4,1],"isMain":false},{"metricName":" # translocations","quality":"Less is better","values":[6,5,2,7,4,1],"isMain":false},{"metricName":" # inversions","quality":"Less is better","values":[1,3,0,0,1,0],"isMain":false},{"metricName":" # interspecies translocations","quality":"Less is better","values":[104,121,50,83,120,56],"isMain":false},{"metricName":"# possibly misassembled contigs","quality":"Less is better","values":[1,4,1,2,6,2],"isMain":false},{"metricName":" # possible misassemblies","quality":"Less is better","values":[1,4,1,2,6,2],"isMain":false},{"metricName":"# local misassemblies","quality":"Less is better","values":[1,2,3,0,0,0],"isMain":false},{"metricName":"# scaffold gap ext. mis.","quality":"Less is better","values":[0,0,0,0,0,0],"isMain":false},{"metricName":"# scaffold gap loc. mis.","quality":"Less is better","values":[0,0,0,0,0,0],"isMain":false}]],["Unaligned",[]],["Mismatches",[]],["Statistics without reference",[]],["K-mer-based statistics",[]],["Predicted genes",[]],["Similarity statistics",[]],["Reference statistics",[]]],[["Genome statistics",[]],["Reads mapping",[]],["Misassemblies",[{"metricName":"# misassemblies","quality":"Less is better","values":[10,11,11,21,9,5],"isMain":true},{"metricName":" # relocations","quality":"Less is better","values":[0,0,0,0,0,0],"isMain":false},{"metricName":" # translocations","quality":"Less is better","values":[0,0,1,0,0,0],"isMain":false},{"metricName":" # inversions","quality":"Less is better","values":[0,0,0,0,0,0],"isMain":false},{"metricName":" # interspecies translocations","quality":"Less is better","values":[10,11,10,21,9,5],"isMain":false},{"metricName":"# possibly misassembled contigs","quality":"Less is better","values":[0,0,0,1,0,0],"isMain":false},{"metricName":" # possible misassemblies","quality":"Less is better","values":[0,0,0,1,0,0],"isMain":false},{"metricName":"# local misassemblies","quality":"Less is better","values":[0,0,0,1,1,0],"isMain":false},{"metricName":"# scaffold gap ext. mis.","quality":"Less is better","values":[0,0,0,0,0,0],"isMain":false},{"metricName":"# scaffold gap loc. mis.","quality":"Less is better","values":[0,0,0,0,0,0],"isMain":false}]],["Unaligned",[]],["Mismatches",[]],["Statistics without reference",[]],["K-mer-based statistics",[]],["Predicted genes",[]],["Similarity statistics",[]],["Reference statistics",[]]],[["Genome statistics",[]],["Reads mapping",[]],["Misassemblies",[{"metricName":"# misassemblies","quality":"Less is better","values":[2,4,8,5,6,1],"isMain":true},{"metricName":" # relocations","quality":"Less is better","values":[0,0,0,0,0,0],"isMain":false},{"metricName":" # translocations","quality":"Less is better","values":[0,0,0,0,0,0],"isMain":false},{"metricName":" # inversions","quality":"Less is better","values":[1,0,0,1,0,0],"isMain":false},{"metricName":" # interspecies translocations","quality":"Less is better","values":[1,4,8,4,6,1],"isMain":false},{"metricName":"# possibly misassembled contigs","quality":"Less is better","values":[0,0,0,0,0,0],"isMain":false},{"metricName":" # possible misassemblies","quality":"Less is better","values":[0,0,0,0,0,0],"isMain":false},{"metricName":"# local misassemblies","quality":"Less is better","values":[0,0,0,1,0,0],"isMain":false},{"metricName":"# scaffold gap ext. mis.","quality":"Less is better","values":[0,0,0,0,0,0],"isMain":false},{"metricName":"# scaffold gap loc. mis.","quality":"Less is better","values":[0,0,0,0,0,0],"isMain":false}]],["Unaligned",[]],["Mismatches",[]],["Statistics without reference",[]],["K-mer-based statistics",[]],["Predicted genes",[]],["Similarity statistics",[]],["Reference statistics",[]]],[["Genome statistics",[]],["Reads mapping",[]],["Misassemblies",[{"metricName":"# misassemblies","quality":"Less is better","values":[8,20,13,8,18,6],"isMain":true},{"metricName":" # relocations","quality":"Less is better","values":[0,0,0,0,0,0],"isMain":false},{"metricName":" # translocations","quality":"Less is better","values":[0,3,1,1,0,0],"isMain":false},{"metricName":" # inversions","quality":"Less is better","values":[0,0,0,0,0,0],"isMain":false},{"metricName":" # interspecies translocations","quality":"Less is better","values":[8,17,12,7,18,6],"isMain":false},{"metricName":"# possibly misassembled contigs","quality":"Less is better","values":[0,0,0,0,0,0],"isMain":false},{"metricName":" # possible misassemblies","quality":"Less is better","values":[0,0,0,0,0,0],"isMain":false},{"metricName":"# local misassemblies","quality":"Less is better","values":[0,1,1,1,1,0],"isMain":false},{"metricName":"# scaffold gap ext. mis.","quality":"Less is better","values":[0,0,0,0,0,0],"isMain":false},{"metricName":"# scaffold gap loc. mis.","quality":"Less is better","values":[0,0,0,0,0,0],"isMain":false}]],["Unaligned",[]],["Mismatches",[]],["Statistics without reference",[]],["K-mer-based statistics",[]],["Predicted genes",[]],["Similarity statistics",[]],["Reference statistics",[]]],[["Genome statistics",[]],["Reads mapping",[]],["Misassemblies",[{"metricName":"# misassemblies","quality":"Less is better","values":[3,6,7,9,10,1],"isMain":true},{"metricName":" # relocations","quality":"Less is better","values":[0,0,0,0,0,0],"isMain":false},{"metricName":" # translocations","quality":"Less is better","values":[0,0,0,0,0,0],"isMain":false},{"metricName":" # inversions","quality":"Less is better","values":[0,0,0,0,0,0],"isMain":false},{"metricName":" # interspecies translocations","quality":"Less is better","values":[3,6,7,9,10,1],"isMain":false},{"metricName":"# possibly misassembled contigs","quality":"Less is better","values":[0,0,0,0,0,0],"isMain":false},{"metricName":" # possible misassemblies","quality":"Less is better","values":[0,0,0,0,0,0],"isMain":false},{"metricName":"# local misassemblies","quality":"Less is better","values":[0,0,0,1,0,0],"isMain":false},{"metricName":"# scaffold gap ext. mis.","quality":"Less is better","values":[0,0,0,0,0,0],"isMain":false},{"metricName":"# scaffold gap loc. mis.","quality":"Less is better","values":[0,0,0,0,0,0],"isMain":false}]],["Unaligned",[]],["Mismatches",[]],["Statistics without reference",[]],["K-mer-based statistics",[]],["Predicted genes",[]],["Similarity statistics",[]],["Reference statistics",[]]],[["Genome statistics",[]],["Reads mapping",[]],["Misassemblies",[{"metricName":"# misassemblies","quality":"Less is better","values":[4,7,10,11,17,2],"isMain":true},{"metricName":" # relocations","quality":"Less is better","values":[0,0,0,0,0,0],"isMain":false},{"metricName":" # translocations","quality":"Less is better","values":[0,0,2,0,0,0],"isMain":false},{"metricName":" # inversions","quality":"Less is better","values":[0,0,0,0,0,0],"isMain":false},{"metricName":" # interspecies translocations","quality":"Less is better","values":[4,7,8,11,17,2],"isMain":false},{"metricName":"# possibly misassembled contigs","quality":"Less is better","values":[0,0,0,0,0,0],"isMain":false},{"metricName":" # possible misassemblies","quality":"Less is better","values":[0,0,0,0,0,0],"isMain":false},{"metricName":"# local misassemblies","quality":"Less is better","values":[0,0,0,0,1,0],"isMain":false},{"metricName":"# scaffold gap ext. mis.","quality":"Less is better","values":[0,0,0,0,0,0],"isMain":false},{"metricName":"# scaffold gap loc. mis.","quality":"Less is better","values":[0,0,0,0,0,0],"isMain":false}]],["Unaligned",[]],["Mismatches",[]],["Statistics without reference",[]],["K-mer-based statistics",[]],["Predicted genes",[]],["Similarity statistics",[]],["Reference statistics",[]]],[["Genome statistics",[]],["Reads mapping",[]],["Misassemblies",[{"metricName":"# misassemblies","quality":"Less is better","values":[2,4,2,8,4,0],"isMain":true},{"metricName":" # relocations","quality":"Less is better","values":[0,0,0,0,0,0],"isMain":false},{"metricName":" # translocations","quality":"Less is better","values":[0,0,0,0,0,0],"isMain":false},{"metricName":" # inversions","quality":"Less is better","values":[0,1,0,0,0,0],"isMain":false},{"metricName":" # interspecies translocations","quality":"Less is better","values":[2,3,2,8,4,0],"isMain":false},{"metricName":"# possibly misassembled contigs","quality":"Less is better","values":[0,0,0,0,0,0],"isMain":false},{"metricName":" # possible misassemblies","quality":"Less is better","values":[0,0,0,0,0,0],"isMain":false},{"metricName":"# local misassemblies","quality":"Less is better","values":[1,0,1,0,0,0],"isMain":false},{"metricName":"# scaffold gap ext. mis.","quality":"Less is better","values":[0,0,0,0,0,0],"isMain":false},{"metricName":"# scaffold gap loc. mis.","quality":"Less is better","values":[0,0,0,0,0,0],"isMain":false}]],["Unaligned",[]],["Mismatches",[]],["Statistics without reference",[]],["K-mer-based statistics",[]],["Predicted genes",[]],["Similarity statistics",[]],["Reference statistics",[]]],[["Genome statistics",[]],["Reads mapping",[]],["Misassemblies",[{"metricName":"# misassemblies","quality":"Less is better","values":[2,3,4,6,9,1],"isMain":true},{"metricName":" # relocations","quality":"Less is better","values":[0,0,0,0,0,0],"isMain":false},{"metricName":" # translocations","quality":"Less is better","values":[0,0,0,0,0,0],"isMain":false},{"metricName":" # inversions","quality":"Less is better","values":[0,0,0,0,0,0],"isMain":false},{"metricName":" # interspecies translocations","quality":"Less is better","values":[2,3,4,6,9,1],"isMain":false},{"metricName":"# possibly misassembled contigs","quality":"Less is better","values":[0,0,0,0,0,0],"isMain":false},{"metricName":" # possible misassemblies","quality":"Less is better","values":[0,0,0,0,0,0],"isMain":false},{"metricName":"# local misassemblies","quality":"Less is better","values":[1,0,0,0,1,0],"isMain":false},{"metricName":"# scaffold gap ext. mis.","quality":"Less is better","values":[0,0,0,0,0,0],"isMain":false},{"metricName":"# scaffold gap loc. mis.","quality":"Less is better","values":[0,0,0,0,0,0],"isMain":false}]],["Unaligned",[]],["Mismatches",[]],["Statistics without reference",[]],["K-mer-based statistics",[]],["Predicted genes",[]],["Similarity statistics",[]],["Reference statistics",[]]],[["Genome statistics",[]],["Reads mapping",[]],["Misassemblies",[{"metricName":"# misassemblies","quality":"Less is better","values":[4,2,5,6,7,1],"isMain":true},{"metricName":" # relocations","quality":"Less is better","values":[0,0,0,0,0,0],"isMain":false},{"metricName":" # translocations","quality":"Less is better","values":[0,0,0,0,0,0],"isMain":false},{"metricName":" # inversions","quality":"Less is better","values":[0,0,1,0,0,0],"isMain":false},{"metricName":" # interspecies translocations","quality":"Less is better","values":[4,2,4,6,7,1],"isMain":false},{"metricName":"# possibly misassembled contigs","quality":"Less is better","values":[0,0,0,0,0,0],"isMain":false},{"metricName":" # possible misassemblies","quality":"Less is better","values":[0,0,0,0,0,0],"isMain":false},{"metricName":"# local misassemblies","quality":"Less is better","values":[0,1,0,1,0,0],"isMain":false},{"metricName":"# scaffold gap ext. mis.","quality":"Less is better","values":[0,0,0,0,0,0],"isMain":false},{"metricName":"# scaffold gap loc. mis.","quality":"Less is better","values":[0,0,0,0,0,0],"isMain":false}]],["Unaligned",[]],["Mismatches",[]],["Statistics without reference",[]],["K-mer-based statistics",[]],["Predicted genes",[]],["Similarity statistics",[]],["Reference statistics",[]]],[["Genome statistics",[]],["Reads mapping",[]],["Misassemblies",[{"metricName":"# misassemblies","quality":"Less is better","values":[18,18,9,11,18,12],"isMain":true},{"metricName":" # relocations","quality":"Less is better","values":[0,0,0,0,0,0],"isMain":false},{"metricName":" # translocations","quality":"Less is better","values":[3,4,4,3,3,1],"isMain":false},{"metricName":" # inversions","quality":"Less is better","values":[1,0,0,0,1,0],"isMain":false},{"metricName":" # interspecies translocations","quality":"Less is better","values":[14,14,5,8,14,11],"isMain":false},{"metricName":"# possibly misassembled contigs","quality":"Less is better","values":[0,0,0,0,0,0],"isMain":false},{"metricName":" # possible misassemblies","quality":"Less is better","values":[0,0,0,0,0,0],"isMain":false},{"metricName":"# local misassemblies","quality":"Less is better","values":[0,1,0,2,1,0],"isMain":false},{"metricName":"# scaffold gap ext. mis.","quality":"Less is better","values":[0,0,0,0,0,0],"isMain":false},{"metricName":"# scaffold gap loc. mis.","quality":"Less is better","values":[0,0,0,0,0,0],"isMain":false}]],["Unaligned",[]],["Mismatches",[]],["Statistics without reference",[]],["K-mer-based statistics",[]],["Predicted genes",[]],["Similarity statistics",[]],["Reference statistics",[]]],[["Genome statistics",[]],["Reads mapping",[]],["Misassemblies",[{"metricName":"# misassemblies","quality":"Less is better","values":[19,22,5,14,14,3],"isMain":true},{"metricName":" # relocations","quality":"Less is better","values":[0,1,0,1,1,0],"isMain":false},{"metricName":" # translocations","quality":"Less is better","values":[1,3,1,2,4,0],"isMain":false},{"metricName":" # inversions","quality":"Less is better","values":[0,0,0,0,0,0],"isMain":false},{"metricName":" # interspecies translocations","quality":"Less is better","values":[18,18,4,11,9,3],"isMain":false},{"metricName":"# possibly misassembled contigs","quality":"Less is better","values":[0,0,0,0,0,0],"isMain":false},{"metricName":" # possible misassemblies","quality":"Less is better","values":[0,0,0,0,0,0],"isMain":false},{"metricName":"# local misassemblies","quality":"Less is better","values":[0,0,1,0,1,0],"isMain":false},{"metricName":"# scaffold gap ext. mis.","quality":"Less is better","values":[0,0,0,0,0,0],"isMain":false},{"metricName":"# scaffold gap loc. mis.","quality":"Less is better","values":[0,0,0,0,0,0],"isMain":false}]],["Unaligned",[]],["Mismatches",[]],["Statistics without reference",[]],["K-mer-based statistics",[]],["Predicted genes",[]],["Similarity statistics",[]],["Reference statistics",[]]],[["Genome statistics",[]],["Reads mapping",[]],["Misassemblies",[{"metricName":"# misassemblies","quality":"Less is better","values":[102,116,47,99,125,43],"isMain":true},{"metricName":" # relocations","quality":"Less is better","values":[3,5,2,4,0,0],"isMain":false},{"metricName":" # translocations","quality":"Less is better","values":[26,26,24,24,30,10],"isMain":false},{"metricName":" # inversions","quality":"Less is better","values":[0,2,0,2,3,0],"isMain":false},{"metricName":" # interspecies translocations","quality":"Less is better","values":[73,83,21,69,92,33],"isMain":false},{"metricName":"# possibly misassembled contigs","quality":"Less is better","values":[0,0,1,5,1,0],"isMain":false},{"metricName":" # possible misassemblies","quality":"Less is better","values":[0,0,1,5,1,0],"isMain":false},{"metricName":"# local misassemblies","quality":"Less is better","values":[2,2,0,3,1,2],"isMain":false},{"metricName":"# scaffold gap ext. mis.","quality":"Less is better","values":[0,0,0,0,0,0],"isMain":false},{"metricName":"# scaffold gap loc. mis.","quality":"Less is better","values":[0,0,0,0,0,0],"isMain":false}]],["Unaligned",[]],["Mismatches",[]],["Statistics without reference",[]],["K-mer-based statistics",[]],["Predicted genes",[]],["Similarity statistics",[]],["Reference statistics",[]]],[["Genome statistics",[]],["Reads mapping",[]],["Misassemblies",[{"metricName":"# misassemblies","quality":"Less is better","values":[0,0,0,0,0,0],"isMain":true},{"metricName":" # relocations","quality":"Less is better","values":[0,0,0,0,0,0],"isMain":false},{"metricName":" # translocations","quality":"Less is better","values":[0,0,0,0,0,0],"isMain":false},{"metricName":" # inversions","quality":"Less is better","values":[0,0,0,0,0,0],"isMain":false},{"metricName":" # interspecies translocations","quality":"Less is better","values":[0,0,0,0,0,0],"isMain":false},{"metricName":"# possibly misassembled contigs","quality":"Less is better","values":[0,0,0,0,0,0],"isMain":false},{"metricName":" # possible misassemblies","quality":"Less is better","values":[0,0,0,0,0,0],"isMain":false},{"metricName":"# local misassemblies","quality":"Less is better","values":[0,0,0,0,0,0],"isMain":false},{"metricName":"# scaffold gap ext. mis.","quality":"Less is better","values":[0,0,0,0,0,0],"isMain":false},{"metricName":"# scaffold gap loc. mis.","quality":"Less is better","values":[0,0,0,0,0,0],"isMain":false}]],["Unaligned",[]],["Mismatches",[]],["Statistics without reference",[]],["K-mer-based statistics",[]],["Predicted genes",[]],["Similarity statistics",[]],["Reference statistics",[]]],[["Genome statistics",[]],["Reads mapping",[]],["Misassemblies",[{"metricName":"# misassemblies","quality":"Less is better","values":[12,20,11,48,21,20],"isMain":true},{"metricName":" # relocations","quality":"Less is better","values":[1,3,0,1,0,1],"isMain":false},{"metricName":" # translocations","quality":"Less is better","values":[4,8,5,4,12,9],"isMain":false},{"metricName":" # inversions","quality":"Less is better","values":[0,0,2,1,0,0],"isMain":false},{"metricName":" # interspecies translocations","quality":"Less is better","values":[7,9,4,42,9,10],"isMain":false},{"metricName":"# possibly misassembled contigs","quality":"Less is better","values":[0,0,0,0,0,0],"isMain":false},{"metricName":" # possible misassemblies","quality":"Less is better","values":[0,0,0,0,0,0],"isMain":false},{"metricName":"# local misassemblies","quality":"Less is better","values":[3,5,7,7,5,4],"isMain":false},{"metricName":"# scaffold gap ext. mis.","quality":"Less is better","values":[0,0,0,0,0,0],"isMain":false},{"metricName":"# scaffold gap loc. mis.","quality":"Less is better","values":[0,0,0,0,0,0],"isMain":false}]],["Unaligned",[]],["Mismatches",[]],["Statistics without reference",[]],["K-mer-based statistics",[]],["Predicted genes",[]],["Similarity statistics",[]],["Reference statistics",[]]],[["Genome statistics",[]],["Reads mapping",[]],["Misassemblies",[{"metricName":"# misassemblies","quality":"Less is better","values":[27,101,41,56,38,3],"isMain":true},{"metricName":" # relocations","quality":"Less is better","values":[0,1,2,3,5,0],"isMain":false},{"metricName":" # translocations","quality":"Less is better","values":[0,7,12,18,13,3],"isMain":false},{"metricName":" # inversions","quality":"Less is better","values":[1,20,6,3,4,0],"isMain":false},{"metricName":" # interspecies translocations","quality":"Less is better","values":[26,73,21,32,16,0],"isMain":false},{"metricName":"# possibly misassembled contigs","quality":"Less is better","values":[0,0,0,0,0,0],"isMain":false},{"metricName":" # possible misassemblies","quality":"Less is better","values":[0,0,0,0,0,0],"isMain":false},{"metricName":"# local misassemblies","quality":"Less is better","values":[2,9,3,10,4,0],"isMain":false},{"metricName":"# scaffold gap ext. mis.","quality":"Less is better","values":[0,0,0,0,0,0],"isMain":false},{"metricName":"# scaffold gap loc. mis.","quality":"Less is better","values":[0,0,0,0,0,0],"isMain":false}]],["Unaligned",[]],["Mismatches",[]],["Statistics without reference",[]],["K-mer-based statistics",[]],["Predicted genes",[]],["Similarity statistics",[]],["Reference statistics",[]]],[["Genome statistics",[]],["Reads mapping",[]],["Misassemblies",[{"metricName":"# misassemblies","quality":"Less is better","values":[30,106,27,54,25,5],"isMain":true},{"metricName":" # relocations","quality":"Less is better","values":[0,2,1,3,1,0],"isMain":false},{"metricName":" # translocations","quality":"Less is better","values":[1,9,12,25,9,4],"isMain":false},{"metricName":" # inversions","quality":"Less is better","values":[0,19,1,3,5,0],"isMain":false},{"metricName":" # interspecies translocations","quality":"Less is better","values":[29,76,13,23,10,1],"isMain":false},{"metricName":"# possibly misassembled contigs","quality":"Less is better","values":[0,1,0,2,0,0],"isMain":false},{"metricName":" # possible misassemblies","quality":"Less is better","values":[0,1,0,2,0,0],"isMain":false},{"metricName":"# local misassemblies","quality":"Less is better","values":[0,15,4,4,3,0],"isMain":false},{"metricName":"# scaffold gap ext. mis.","quality":"Less is better","values":[0,0,0,0,0,0],"isMain":false},{"metricName":"# scaffold gap loc. mis.","quality":"Less is better","values":[0,0,0,0,0,0],"isMain":false}]],["Unaligned",[]],["Mismatches",[]],["Statistics without reference",[]],["K-mer-based statistics",[]],["Predicted genes",[]],["Similarity statistics",[]],["Reference statistics",[]]],[["Genome statistics",[]],["Reads mapping",[]],["Misassemblies",[{"metricName":"# misassemblies","quality":"Less is better","values":[7,19,9,11,13,5],"isMain":true},{"metricName":" # relocations","quality":"Less is better","values":[0,0,0,0,0,0],"isMain":false},{"metricName":" # translocations","quality":"Less is better","values":[0,0,0,0,0,1],"isMain":false},{"metricName":" # inversions","quality":"Less is better","values":[0,1,0,1,0,0],"isMain":false},{"metricName":" # interspecies translocations","quality":"Less is better","values":[7,18,9,10,13,4],"isMain":false},{"metricName":"# possibly misassembled contigs","quality":"Less is better","values":[0,0,0,0,0,0],"isMain":false},{"metricName":" # possible misassemblies","quality":"Less is better","values":[0,0,0,0,0,0],"isMain":false},{"metricName":"# local misassemblies","quality":"Less is better","values":[0,1,0,0,1,0],"isMain":false},{"metricName":"# scaffold gap ext. mis.","quality":"Less is better","values":[0,0,0,0,0,0],"isMain":false},{"metricName":"# scaffold gap loc. mis.","quality":"Less is better","values":[0,0,0,0,0,0],"isMain":false}]],["Unaligned",[]],["Mismatches",[]],["Statistics without reference",[]],["K-mer-based statistics",[]],["Predicted genes",[]],["Similarity statistics",[]],["Reference statistics",[]]],[["Genome statistics",[]],["Reads mapping",[]],["Misassemblies",[{"metricName":"# misassemblies","quality":"Less is better","values":[54,57,17,55,46,55],"isMain":true},{"metricName":" # relocations","quality":"Less is better","values":[0,3,2,2,1,0],"isMain":false},{"metricName":" # translocations","quality":"Less is better","values":[4,5,5,11,9,2],"isMain":false},{"metricName":" # inversions","quality":"Less is better","values":[0,0,0,0,0,0],"isMain":false},{"metricName":" # interspecies translocations","quality":"Less is better","values":[50,49,10,42,36,53],"isMain":false},{"metricName":"# possibly misassembled contigs","quality":"Less is better","values":[0,0,0,1,0,0],"isMain":false},{"metricName":" # possible misassemblies","quality":"Less is better","values":[0,0,0,1,0,0],"isMain":false},{"metricName":"# local misassemblies","quality":"Less is better","values":[0,0,1,1,2,1],"isMain":false},{"metricName":"# scaffold gap ext. mis.","quality":"Less is better","values":[0,0,0,0,0,0],"isMain":false},{"metricName":"# scaffold gap loc. mis.","quality":"Less is better","values":[0,0,0,0,0,0],"isMain":false}]],["Unaligned",[]],["Mismatches",[]],["Statistics without reference",[]],["K-mer-based statistics",[]],["Predicted genes",[]],["Similarity statistics",[]],["Reference statistics",[]]],[["Genome statistics",[]],["Reads mapping",[]],["Misassemblies",[{"metricName":"# misassemblies","quality":"Less is better","values":[17,13,19,35,24,0],"isMain":true},{"metricName":" # relocations","quality":"Less is better","values":[1,1,1,2,2,0],"isMain":false},{"metricName":" # translocations","quality":"Less is better","values":[0,0,0,2,0,0],"isMain":false},{"metricName":" # inversions","quality":"Less is better","values":[0,0,0,0,0,0],"isMain":false},{"metricName":" # interspecies translocations","quality":"Less is better","values":[16,12,18,31,22,0],"isMain":false},{"metricName":"# possibly misassembled contigs","quality":"Less is better","values":[0,0,0,0,0,0],"isMain":false},{"metricName":" # possible misassemblies","quality":"Less is better","values":[0,0,0,0,0,0],"isMain":false},{"metricName":"# local misassemblies","quality":"Less is better","values":[0,0,2,2,2,0],"isMain":false},{"metricName":"# scaffold gap ext. mis.","quality":"Less is better","values":[0,0,0,0,0,0],"isMain":false},{"metricName":"# scaffold gap loc. mis.","quality":"Less is better","values":[0,0,0,0,0,0],"isMain":false}]],["Unaligned",[]],["Mismatches",[]],["Statistics without reference",[]],["K-mer-based statistics",[]],["Predicted genes",[]],["Similarity statistics",[]],["Reference statistics",[]]]],"minContig":500},{"date":"10 February 2020, Monday, 18:00:13","assembliesNames":["megahit-103-df","megahit-113-df","megahit-113-ml","megahit-113-ms","megahit-129-df","metaSPAdes"],"referenceName":"GCF\_000006625.1\_ASM662v1","order":[0,1,2,3,4,5],"report":[["Genome statistics",[{"metricName":"Genome fraction (%)","quality":"More is better","values":["75.493","81.493","78.159","78.308","84.272","69.616"],"isMain":true},{"metricName":"Duplication ratio","quality":"Less is better","values":["1.052","1.044","1.075","1.078","1.054","1.055"],"isMain":true},{"metricName":"Largest alignment","quality":"More is better","values":[2984,3992,3733,3506,4707,3277],"isMain":true},{"metricName":"Total aligned length","quality":"More is better","values":[591964,634501,626898,627244,663206,545317],"isMain":true},{"metricName":"NG50","quality":"More is better","values":[864,1100,851,858,1147,720],"isMain":false},{"metricName":"NG75","quality":"More is better","values":[544,640,578,595,696,null],"isMain":false},{"metricName":"NA50","quality":"More is better","values":[1013,1249,959,931,1231,841],"isMain":false},{"metricName":"NA75","quality":"More is better","values":[701,786,681,678,820,638],"isMain":false},{"metricName":"NGA50","quality":"More is better","values":[851,1072,843,842,1133,691],"isMain":true},{"metricName":"NGA75","quality":"More is better","values":[534,614,562,571,682,null],"isMain":false},{"metricName":"LG50","quality":"Less is better","values":[280,219,297,291,211,353],"isMain":false},{"metricName":"LG75","quality":"Less is better","values":[558,444,567,559,421,null],"isMain":false},{"metricName":"LA50","quality":"Less is better","values":[199,174,234,229,178,235],"isMain":false},{"metricName":"LA75","quality":"Less is better","values":[375,336,430,430,342,427],"isMain":false},{"metricName":"LGA50","quality":"Less is better","values":[282,221,301,295,214,362],"isMain":true},{"metricName":"LGA75","quality":"Less is better","values":[565,454,576,570,427,null],"isMain":false}]],["Reads mapping",[]],["Misassemblies",[{"metricName":"# misassemblies","quality":"Less is better","values":[1,6,4,3,3,2],"isMain":true},{"metricName":" # relocations","quality":"Less is better","values":[0,4,2,2,0,2],"isMain":false},{"metricName":" # translocations","quality":"Less is better","values":[0,0,0,0,0,0],"isMain":false},{"metricName":" # inversions","quality":"Less is better","values":[1,2,2,1,3,0],"isMain":false},{"metricName":" # interspecies translocations","quality":"Less is better","values":[9,11,0,16,7,13],"isMain":false},{"metricName":"# misassembled contigs","quality":"Less is better","values":[1,6,4,3,3,2],"isMain":false},{"metricName":"Misassembled contigs length","quality":"Less is better","values":[554,7051,5468,2944,4546,1966],"isMain":true},{"metricName":"# possibly misassembled contigs","quality":"Less is better","values":[0,0,0,0,0,0],"isMain":false},{"metricName":" # possible misassemblies","quality":"Less is better","values":[0,0,0,0,0,0],"isMain":false},{"metricName":"# local misassemblies","quality":"Less is better","values":[0,0,0,0,2,0],"isMain":false},{"metricName":"# scaffold gap ext. mis.","quality":"Less is better","values":[0,0,0,0,0,0],"isMain":false},{"metricName":"# scaffold gap loc. mis.","quality":"Less is better","values":[0,0,0,0,0,0],"isMain":false},{"metricName":"# unaligned mis. contigs","quality":"Less is better","values":[0,0,0,0,0,0],"isMain":false}]],["Unaligned",[{"metricName":"# fully unaligned contigs","quality":"Less is better","values":[0,0,0,0,0,0],"isMain":false},{"metricName":"Fully unaligned length","quality":"Less is better","values":[0,0,0,0,0,0],"isMain":false},{"metricName":"# partially unaligned contigs","quality":"Less is better","values":[0,2,0,2,1,9],"isMain":false},{"metricName":"Partially unaligned length","quality":"Less is better","values":[0,1764,0,1500,577,6120],"isMain":false}]],["Mismatches",[{"metricName":"# mismatches","quality":"Less is better","values":[6183,6207,10286,10423,6579,6376],"isMain":false},{"metricName":"# indels","quality":"Less is better","values":[6,13,14,14,17,11],"isMain":false},{"metricName":"Indels length","quality":"Less is better","values":[20,13,17,38,72,13],"isMain":false},{"metricName":"# mismatches per 100 kbp","quality":"Less is better","values":["1089.53","1013.23","1750.69","1770.64","1038.53","1218.38"],"isMain":true},{"metricName":"# indels per 100 kbp","quality":"Less is better","values":["1.06","2.12","2.38","2.38","2.68","2.10"],"isMain":true},{"metricName":" # indels (<= 5 bp)","quality":"Less is better","values":[5,13,14,13,14,11],"isMain":false},{"metricName":" # indels (> 5 bp)","quality":"Less is better","values":[1,0,0,1,3,0],"isMain":false},{"metricName":"# N's","quality":"Less is better","values":[0,0,0,0,0,0],"isMain":false},{"metricName":"# N's per 100 kbp","quality":"Less is better","values":["0.00","0.00","0.00","0.00","0.00","0.00"],"isMain":true}]],["Statistics without reference",[{"metricName":"# contigs","quality":"Equal","values":[621,582,693,691,599,657],"isMain":true},{"metricName":"# contigs (>= 1000 bp)","quality":"Equal","values":[207,245,209,193,265,160],"isMain":false},{"metricName":"# contigs (>= 5000 bp)","quality":"Equal","values":[0,0,0,0,0,0],"isMain":false},{"metricName":"# contigs (>= 10000 bp)","quality":"Equal","values":[0,0,0,0,0,0],"isMain":false},{"metricName":"# contigs (>= 25000 bp)","quality":"Equal","values":[0,0,0,0,0,0],"isMain":false},{"metricName":"# contigs (>= 50000 bp)","quality":"Equal","values":[0,0,0,0,0,0],"isMain":false},{"metricName":"Largest contig","quality":"More is better","values":[2998,3993,3733,3506,4725,3285],"isMain":true},{"metricName":"Total length","quality":"More is better","values":[597165,641571,631444,636209,668370,558295],"isMain":true},{"metricName":"Total length (>= 1000 bp)","quality":"More is better","values":[308444,403924,294367,285569,434419,215059],"isMain":true},{"metricName":"Total length (>= 5000 bp)","quality":"More is better","values":[0,0,0,0,0,0],"isMain":false},{"metricName":"Total length (>= 10000 bp)","quality":"More is better","values":[0,0,0,0,0,0],"isMain":true},{"metricName":"Total length (>= 25000 bp)","quality":"More is better","values":[0,0,0,0,0,0],"isMain":false},{"metricName":"Total length (>= 50000 bp)","quality":"More is better","values":[0,0,0,0,0,0],"isMain":true},{"metricName":"N50","quality":"More is better","values":[1022,1265,970,942,1255,865],"isMain":false},{"metricName":"N75","quality":"More is better","values":[719,829,691,696,836,670],"isMain":false},{"metricName":"L50","quality":"Less is better","values":[198,172,231,227,176,230],"isMain":false},{"metricName":"L75","quality":"Less is better","values":[372,330,425,423,339,415],"isMain":false},{"metricName":"GC (%)","quality":"Equal","values":["25.81","25.69","25.91","26.08","25.69","26.01"],"isMain":false}]],["K-mer-based statistics",[]],["Predicted genes",[]],["Similarity statistics",[]],["Reference statistics",[{"metricName":"Reference length","quality":"Equal","values":[751719,751719,751719,751719,751719,751719],"isMain":false},{"metricName":"Reference fragments","quality":"Equal","values":[1,1,1,1,1,1],"isMain":false},{"metricName":"Reference GC (%)","quality":"Equal","values":["25.50","25.50","25.50","25.50","25.50","25.50"],"isMain":false}]]],"subreferences":[],"subreports":[],"minContig":500},{"date":"10 February 2020, Monday, 18:00:20","assembliesNames":["megahit-103-df","megahit-113-df","megahit-113-ml","megahit-113-ms","megahit-129-df","metaSPAdes"],"referenceName":"GCF\_000009925.1\_ASM992v1","order":[0,1,2,3,4,5],"report":[["Genome statistics",[{"metricName":"Genome fraction (%)","quality":"More is better","values":["17.970","19.147","19.817","21.261","19.160","12.810"],"isMain":true},{"metricName":"Duplication ratio","quality":"Less is better","values":["1.018","1.025","1.025","1.033","1.025","1.002"],"isMain":true},{"metricName":"Largest alignment","quality":"More is better","values":[23951,66317,55765,97687,72610,26812],"isMain":true},{"metricName":"Total aligned length","quality":"More is better","values":[965009,1038675,1074530,1160268,1039594,681719],"isMain":true},{"metricName":"NA50","quality":"More is better","values":[3463,11133,11611,9771,11391,10224],"isMain":false},{"metricName":"NA75","quality":"More is better","values":[1127,2547,3193,2360,4175,3913],"isMain":false},{"metricName":"NGA50","quality":"More is better","values":[null,null,null,null,null,null],"isMain":true},{"metricName":"LA50","quality":"Less is better","values":[62,27,27,27,27,22],"isMain":false},{"metricName":"LA75","quality":"Less is better","values":[202,87,78,101,69,53],"isMain":false}]],["Reads mapping",[]],["Misassemblies",[{"metricName":"# misassemblies","quality":"Less is better","values":[0,10,9,8,10,0],"isMain":true},{"metricName":" # relocations","quality":"Less is better","values":[0,8,9,6,6,0],"isMain":false},{"metricName":" # translocations","quality":"Less is better","values":[0,0,0,0,0,0],"isMain":false},{"metricName":" # inversions","quality":"Less is better","values":[0,2,0,2,4,0],"isMain":false},{"metricName":" # interspecies translocations","quality":"Less is better","values":[26,21,22,31,8,1],"isMain":false},{"metricName":"# misassembled contigs","quality":"Less is better","values":[0,8,8,7,9,0],"isMain":false},{"metricName":"Misassembled contigs length","quality":"Less is better","values":[0,279791,265719,286762,137431,0],"isMain":true},{"metricName":"# possibly misassembled contigs","quality":"Less is better","values":[0,0,0,0,1,0],"isMain":false},{"metricName":" # possible misassemblies","quality":"Less is better","values":[0,0,0,0,1,0],"isMain":false},{"metricName":"# local misassemblies","quality":"Less is better","values":[1,6,3,2,2,0],"isMain":false},{"metricName":"# scaffold gap ext. mis.","quality":"Less is better","values":[0,0,0,0,0,0],"isMain":false},{"metricName":"# scaffold gap loc. mis.","quality":"Less is better","values":[0,0,0,0,0,0],"isMain":false},{"metricName":"# unaligned mis. contigs","quality":"Less is better","values":[0,1,1,1,0,0],"isMain":false}]],["Unaligned",[{"metricName":"# fully unaligned contigs","quality":"Less is better","values":[0,1,0,2,0,0],"isMain":false},{"metricName":"Fully unaligned length","quality":"Less is better","values":[0,563,0,1237,0,0],"isMain":false},{"metricName":"# partially unaligned contigs","quality":"Less is better","values":[12,10,13,17,4,1],"isMain":false},{"metricName":"Partially unaligned length","quality":"Less is better","values":[9315,100403,92411,156616,24209,871],"isMain":false}]],["Mismatches",[{"metricName":"# mismatches","quality":"Less is better","values":[1766,4289,4504,5653,3971,389],"isMain":false},{"metricName":"# indels","quality":"Less is better","values":[15,112,85,81,59,14],"isMain":false},{"metricName":"Indels length","quality":"Less is better","values":[71,506,418,370,178,60],"isMain":false},{"metricName":"# mismatches per 100 kbp","quality":"Less is better","values":["185.04","421.77","427.94","500.64","390.24","57.18"],"isMain":true},{"metricName":"# indels per 100 kbp","quality":"Less is better","values":["1.57","11.01","8.08","7.17","5.80","2.06"],"isMain":true},{"metricName":" # indels (<= 5 bp)","quality":"Less is better","values":[13,92,71,69,55,13],"isMain":false},{"metricName":" # indels (> 5 bp)","quality":"Less is better","values":[2,20,14,12,4,1],"isMain":false},{"metricName":"# N's","quality":"Less is better","values":[0,0,0,0,0,0],"isMain":false},{"metricName":"# N's per 100 kbp","quality":"Less is better","values":["0.00","0.00","0.00","0.00","0.00","0.00"],"isMain":true}]],["Statistics without reference",[{"metricName":"# contigs","quality":"Equal","values":[534,260,251,280,263,147],"isMain":true},{"metricName":"# contigs (>= 1000 bp)","quality":"Equal","values":[242,149,150,158,135,124],"isMain":false},{"metricName":"# contigs (>= 5000 bp)","quality":"Equal","values":[36,46,50,48,54,39],"isMain":false},{"metricName":"# contigs (>= 10000 bp)","quality":"Equal","values":[19,25,30,25,26,22],"isMain":false},{"metricName":"# contigs (>= 25000 bp)","quality":"Equal","values":[0,5,6,9,4,2],"isMain":false},{"metricName":"# contigs (>= 50000 bp)","quality":"Equal","values":[0,3,3,4,3,0],"isMain":false},{"metricName":"Largest contig","quality":"More is better","values":[23951,137238,123777,188932,80023,26812],"isMain":true},{"metricName":"Total length","quality":"More is better","values":[981291,1143088,1170970,1324020,1067568,682892],"isMain":true},{"metricName":"Total length (>= 1000 bp)","quality":"More is better","values":[781194,1071410,1104576,1244536,983245,666554],"isMain":true},{"metricName":"Total length (>= 5000 bp)","quality":"More is better","values":[385392,823433,869095,976240,775219,453046],"isMain":false},{"metricName":"Total length (>= 10000 bp)","quality":"More is better","values":[273523,688115,734178,814808,587873,342644],"isMain":true},{"metricName":"Total length (>= 25000 bp)","quality":"More is better","values":[0,374331,357746,579260,229718,51956],"isMain":false},{"metricName":"Total length (>= 50000 bp)","quality":"More is better","values":[0,309728,250515,403965,203993,0],"isMain":true},{"metricName":"N50","quality":"More is better","values":[3463,14257,14412,17232,13604,10224],"isMain":false},{"metricName":"N75","quality":"More is better","values":[1140,4163,4872,4873,4380,3913],"isMain":false},{"metricName":"L50","quality":"Less is better","values":[62,16,18,14,22,22],"isMain":false},{"metricName":"L75","quality":"Less is better","values":[200,54,52,52,60,53],"isMain":false},{"metricName":"GC (%)","quality":"Equal","values":["40.51","40.11","40.15","40.70","39.92","39.25"],"isMain":false}]],["K-mer-based statistics",[]],["Predicted genes",[]],["Similarity statistics",[]],["Reference statistics",[{"metricName":"Reference length","quality":"Equal","values":[5310990,5310990,5310990,5310990,5310990,5310990],"isMain":false},{"metricName":"Reference fragments","quality":"Equal","values":[2,2,2,2,2,2],"isMain":false},{"metricName":"Reference GC (%)","quality":"Equal","values":["43.20","43.20","43.20","43.20","43.20","43.20"],"isMain":false}]]],"subreferences":[],"subreports":[],"minContig":500},{"date":"10 February 2020, Monday, 18:00:20","assembliesNames":["megahit-103-df","megahit-113-df","megahit-113-ml","megahit-113-ms","megahit-129-df","metaSPAdes"],"referenceName":"GCF\_000010505.1\_ASM1050v1","order":[0,1,2,3,4,5],"report":[["Genome statistics",[{"metricName":"Genome fraction (%)","quality":"More is better","values":["0.285","0.956","1.496","2.502","2.057","0.253"],"isMain":true},{"metricName":"Duplication ratio","quality":"Less is better","values":["1.004","1.004","1.009","1.020","1.040","1.003"],"isMain":true},{"metricName":"Largest alignment","quality":"More is better","values":[1157,9757,3516,12621,38038,4695],"isMain":true},{"metricName":"Total aligned length","quality":"More is better","values":[6727,22531,35242,58927,48748,5970],"isMain":true},{"metricName":"NA50","quality":"More is better","values":[747,3679,1111,8463,38038,568],"isMain":false},{"metricName":"NA75","quality":"More is better","values":[555,743,614,1765,716,null],"isMain":false},{"metricName":"NGA50","quality":"More is better","values":[null,null,null,null,null,null],"isMain":true},{"metricName":"LA50","quality":"Less is better","values":[4,2,9,3,1,2],"isMain":false},{"metricName":"LA75","quality":"Less is better","values":[7,6,22,8,4,null],"isMain":false}]],["Reads mapping",[]],["Misassemblies",[{"metricName":"# misassemblies","quality":"Less is better","values":[0,0,2,4,2,0],"isMain":true},{"metricName":" # relocations","quality":"Less is better","values":[0,0,2,4,2,0],"isMain":false},{"metricName":" # translocations","quality":"Less is better","values":[0,0,0,0,0,0],"isMain":false},{"metricName":" # inversions","quality":"Less is better","values":[0,0,0,0,0,0],"isMain":false},{"metricName":" # interspecies translocations","quality":"Less is better","values":[0,1,3,4,7,1],"isMain":false},{"metricName":"# misassembled contigs","quality":"Less is better","values":[0,0,2,4,2,0],"isMain":false},{"metricName":"Misassembled contigs length","quality":"Less is better","values":[0,0,3966,17565,1162,0],"isMain":true},{"metricName":"# possibly misassembled contigs","quality":"Less is better","values":[0,0,0,0,0,0],"isMain":false},{"metricName":" # possible misassemblies","quality":"Less is better","values":[0,0,0,0,0,0],"isMain":false},{"metricName":"# local misassemblies","quality":"Less is better","values":[0,0,0,1,0,0],"isMain":false},{"metricName":"# scaffold gap ext. mis.","quality":"Less is better","values":[0,0,0,0,0,0],"isMain":false},{"metricName":"# scaffold gap loc. mis.","quality":"Less is better","values":[0,0,0,0,0,0],"isMain":false},{"metricName":"# unaligned mis. contigs","quality":"Less is better","values":[0,0,0,0,0,0],"isMain":false}]],["Unaligned",[{"metricName":"# fully unaligned contigs","quality":"Less is better","values":[1,0,2,0,3,0],"isMain":false},{"metricName":"Fully unaligned length","quality":"Less is better","values":[700,0,2483,0,2574,0],"isMain":false},{"metricName":"# partially unaligned contigs","quality":"Less is better","values":[0,1,1,1,0,1],"isMain":false},{"metricName":"Partially unaligned length","quality":"Less is better","values":[0,1222,839,1106,0,3540],"isMain":false}]],["Mismatches",[{"metricName":"# mismatches","quality":"Less is better","values":[118,217,688,791,698,45],"isMain":false},{"metricName":"# indels","quality":"Less is better","values":[3,2,15,48,11,5],"isMain":false},{"metricName":"Indels length","quality":"Less is better","values":[15,15,59,102,22,14],"isMain":false},{"metricName":"# mismatches per 100 kbp","quality":"Less is better","values":["1757.52","963.50","1952.49","1342.66","1441.23","755.29"],"isMain":true},{"metricName":"# indels per 100 kbp","quality":"Less is better","values":["44.68","8.88","42.57","81.48","22.71","83.92"],"isMain":true},{"metricName":" # indels (<= 5 bp)","quality":"Less is better","values":[2,1,13,45,10,4],"isMain":false},{"metricName":" # indels (> 5 bp)","quality":"Less is better","values":[1,1,2,3,1,1],"isMain":false},{"metricName":"# N's","quality":"Less is better","values":[0,0,0,0,0,0],"isMain":false},{"metricName":"# N's per 100 kbp","quality":"Less is better","values":["0.00","0.00","0.00","0.00","0.00","0.00"],"isMain":true}]],["Statistics without reference",[{"metricName":"# contigs","quality":"Equal","values":[10,14,35,27,24,4],"isMain":true},{"metricName":"# contigs (>= 1000 bp)","quality":"Equal","values":[2,5,11,9,2,2],"isMain":false},{"metricName":"# contigs (>= 5000 bp)","quality":"Equal","values":[0,1,0,4,1,0],"isMain":false},{"metricName":"# contigs (>= 10000 bp)","quality":"Equal","values":[0,0,0,2,1,0],"isMain":false},{"metricName":"# contigs (>= 25000 bp)","quality":"Equal","values":[0,0,0,0,1,0],"isMain":false},{"metricName":"# contigs (>= 50000 bp)","quality":"Equal","values":[0,0,0,0,0,0],"isMain":false},{"metricName":"Largest contig","quality":"More is better","values":[1157,9757,3516,15902,38038,4695],"isMain":true},{"metricName":"Total length","quality":"More is better","values":[7443,23836,38864,61178,52920,9514],"isMain":true},{"metricName":"Total length (>= 1000 bp)","quality":"More is better","values":[2164,18528,23566,50488,39136,8426],"isMain":true},{"metricName":"Total length (>= 5000 bp)","quality":"More is better","values":[0,9757,0,40067,38038,0],"isMain":false},{"metricName":"Total length (>= 10000 bp)","quality":"More is better","values":[0,0,0,26455,38038,0],"isMain":true},{"metricName":"Total length (>= 25000 bp)","quality":"More is better","values":[0,0,0,0,38038,0],"isMain":false},{"metricName":"Total length (>= 50000 bp)","quality":"More is better","values":[0,0,0,0,0,0],"isMain":true},{"metricName":"N50","quality":"More is better","values":[747,3679,1406,8493,38038,3731],"isMain":false},{"metricName":"N75","quality":"More is better","values":[638,1279,681,1765,907,3731],"isMain":false},{"metricName":"L50","quality":"Less is better","values":[4,2,8,3,1,2],"isMain":false},{"metricName":"L75","quality":"Less is better","values":[7,5,19,7,3,2],"isMain":false},{"metricName":"GC (%)","quality":"Equal","values":["45.09","45.66","46.11","48.43","49.28","45.84"],"isMain":false}]],["K-mer-based statistics",[]],["Predicted genes",[]],["Similarity statistics",[]],["Reference statistics",[{"metricName":"Reference length","quality":"Equal","values":[2354886,2354886,2354886,2354886,2354886,2354886],"isMain":false},{"metricName":"Reference fragments","quality":"Equal","values":[1,1,1,1,1,1],"isMain":false},{"metricName":"Reference GC (%)","quality":"Equal","values":["48.36","48.36","48.36","48.36","48.36","48.36"],"isMain":false}]]],"subreferences":[],"subreports":[],"minContig":500},{"date":"10 February 2020, Monday, 18:00:21","assembliesNames":["megahit-103-df","megahit-113-df","megahit-113-ml","megahit-113-ms","megahit-129-df","metaSPAdes"],"referenceName":"GCF\_000011045.1\_ASM1104v1","order":[0,1,2,3,4,5],"report":[["Genome statistics",[{"metricName":"Genome fraction (%)","quality":"More is better","values":["36.836","38.841","34.048","35.440","41.151","29.700"],"isMain":true},{"metricName":"Duplication ratio","quality":"Less is better","values":["1.031","1.031","1.040","1.044","1.031","1.032"],"isMain":true},{"metricName":"Largest alignment","quality":"More is better","values":[4011,8336,3267,2729,7990,2665],"isMain":true},{"metricName":"Total aligned length","quality":"More is better","values":[1131874,1187821,1052832,1097356,1266946,907792],"isMain":true},{"metricName":"NA50","quality":"More is better","values":[955,1085,824,849,1090,818],"isMain":false},{"metricName":"NA75","quality":"More is better","values":[697,756,632,639,759,624],"isMain":false},{"metricName":"NGA50","quality":"More is better","values":[null,null,null,null,null,null],"isMain":true},{"metricName":"LA50","quality":"Less is better","values":[418,378,474,466,413,401],"isMain":false},{"metricName":"LA75","quality":"Less is better","values":[772,716,853,844,770,725],"isMain":false}]],["Reads mapping",[]],["Misassemblies",[{"metricName":"# misassemblies","quality":"Less is better","values":[4,7,8,13,16,1],"isMain":true},{"metricName":" # relocations","quality":"Less is better","values":[4,7,8,13,16,1],"isMain":false},{"metricName":" # translocations","quality":"Less is better","values":[0,0,0,0,0,0],"isMain":false},{"metricName":" # inversions","quality":"Less is better","values":[0,0,0,0,0,0],"isMain":false},{"metricName":" # interspecies translocations","quality":"Less is better","values":[18,45,11,25,20,20],"isMain":false},{"metricName":"# misassembled contigs","quality":"Less is better","values":[4,7,8,13,15,1],"isMain":false},{"metricName":"Misassembled contigs length","quality":"Less is better","values":[5669,11758,7948,13913,19151,766],"isMain":true},{"metricName":"# possibly misassembled contigs","quality":"Less is better","values":[0,0,0,0,0,0],"isMain":false},{"metricName":" # possible misassemblies","quality":"Less is better","values":[0,0,0,0,0,0],"isMain":false},{"metricName":"# local misassemblies","quality":"Less is better","values":[4,1,1,5,0,0],"isMain":false},{"metricName":"# scaffold gap ext. mis.","quality":"Less is better","values":[0,0,0,0,0,0],"isMain":false},{"metricName":"# scaffold gap loc. mis.","quality":"Less is better","values":[0,0,0,0,0,0],"isMain":false},{"metricName":"# unaligned mis. contigs","quality":"Less is better","values":[0,1,1,0,1,0],"isMain":false}]],["Unaligned",[{"metricName":"# fully unaligned contigs","quality":"Less is better","values":[1,0,1,1,0,0],"isMain":false},{"metricName":"Fully unaligned length","quality":"Less is better","values":[1137,0,553,675,0,0],"isMain":false},{"metricName":"# partially unaligned contigs","quality":"Less is better","values":[1,8,2,5,5,1],"isMain":false},{"metricName":"Partially unaligned length","quality":"Less is better","values":[1747,22598,23477,4603,36383,1048],"isMain":false}]],["Mismatches",[{"metricName":"# mismatches","quality":"Less is better","values":[14327,14970,19583,21066,16197,11819],"isMain":false},{"metricName":"# indels","quality":"Less is better","values":[42,94,58,68,63,34],"isMain":false},{"metricName":"Indels length","quality":"Less is better","values":[101,150,155,133,138,38],"isMain":false},{"metricName":"# mismatches per 100 kbp","quality":"Less is better","values":["1294.28","1282.57","1913.98","1978.05","1309.80","1324.27"],"isMain":true},{"metricName":"# indels per 100 kbp","quality":"Less is better","values":["3.79","8.05","5.67","6.39","5.09","3.81"],"isMain":true},{"metricName":" # indels (<= 5 bp)","quality":"Less is better","values":[39,91,56,67,57,34],"isMain":false},{"metricName":" # indels (> 5 bp)","quality":"Less is better","values":[3,3,2,1,6,0],"isMain":false},{"metricName":"# N's","quality":"Less is better","values":[0,0,0,0,0,0],"isMain":false},{"metricName":"# N's per 100 kbp","quality":"Less is better","values":["0.00","0.00","0.00","0.00","0.00","0.00"],"isMain":true}]],["Statistics without reference",[{"metricName":"# contigs","quality":"Equal","values":[1238,1154,1279,1309,1227,1120],"isMain":true},{"metricName":"# contigs (>= 1000 bp)","quality":"Equal","values":[377,469,291,311,505,236],"isMain":false},{"metricName":"# contigs (>= 5000 bp)","quality":"Equal","values":[0,4,1,0,3,0],"isMain":false},{"metricName":"# contigs (>= 10000 bp)","quality":"Equal","values":[0,1,1,0,2,0],"isMain":false},{"metricName":"# contigs (>= 25000 bp)","quality":"Equal","values":[0,0,0,0,0,0],"isMain":false},{"metricName":"# contigs (>= 50000 bp)","quality":"Equal","values":[0,0,0,0,0,0],"isMain":false},{"metricName":"Largest contig","quality":"More is better","values":[4098,13337,23737,3384,20578,2669],"isMain":true},{"metricName":"Total length","quality":"More is better","values":[1143729,1226518,1088373,1116831,1311929,922273],"isMain":true},{"metricName":"Total length (>= 1000 bp)","quality":"More is better","values":[538098,737660,405906,424431,793462,316529],"isMain":true},{"metricName":"Total length (>= 5000 bp)","quality":"More is better","values":[0,35339,23737,0,43561,0],"isMain":false},{"metricName":"Total length (>= 10000 bp)","quality":"More is better","values":[0,13337,23737,0,35516,0],"isMain":true},{"metricName":"Total length (>= 25000 bp)","quality":"More is better","values":[0,0,0,0,0,0],"isMain":false},{"metricName":"Total length (>= 50000 bp)","quality":"More is better","values":[0,0,0,0,0,0],"isMain":true},{"metricName":"N50","quality":"More is better","values":[971,1139,863,870,1138,831],"isMain":false},{"metricName":"N75","quality":"More is better","values":[712,807,658,662,817,639],"isMain":false},{"metricName":"L50","quality":"Less is better","values":[412,353,442,456,377,396],"isMain":false},{"metricName":"L75","quality":"Less is better","values":[759,673,804,823,715,712],"isMain":false},{"metricName":"GC (%)","quality":"Equal","values":["46.53","46.27","46.63","46.45","46.59","46.51"],"isMain":false}]],["K-mer-based statistics",[]],["Predicted genes",[]],["Similarity statistics",[]],["Reference statistics",[{"metricName":"Reference length","quality":"Equal","values":[3005051,3005051,3005051,3005051,3005051,3005051],"isMain":false},{"metricName":"Reference fragments","quality":"Equal","values":[1,1,1,1,1,1],"isMain":false},{"metricName":"Reference GC (%)","quality":"Equal","values":["46.69","46.69","46.69","46.69","46.69","46.69"],"isMain":false}]]],"subreferences":[],"subreports":[],"minContig":500},{"date":"10 February 2020, Monday, 18:00:39","assembliesNames":["megahit-103-df","megahit-113-df","megahit-113-ml","megahit-113-ms","megahit-129-df","metaSPAdes"],"referenceName":"GCF\_000011065.1\_ASM1106v1","order":[0,1,2,3,4,5],"report":[["Genome statistics",[{"metricName":"Genome fraction (%)","quality":"More is better","values":["25.979","30.303","31.437","31.190","28.073","15.665"],"isMain":true},{"metricName":"Duplication ratio","quality":"Less is better","values":["1.035","1.037","1.057","1.062","1.047","1.036"],"isMain":true},{"metricName":"Largest alignment","quality":"More is better","values":[22683,27420,11105,10798,19833,16699],"isMain":true},{"metricName":"Total aligned length","quality":"More is better","values":[1680509,1960625,2076894,2063915,1830216,1011368],"isMain":true},{"metricName":"NA50","quality":"More is better","values":[1713,2552,1350,1506,1898,1386],"isMain":false},{"metricName":"NA75","quality":"More is better","values":[970,1225,793,830,913,860],"isMain":false},{"metricName":"NGA50","quality":"More is better","values":[null,null,null,null,null,null],"isMain":true},{"metricName":"LA50","quality":"Less is better","values":[269,194,446,390,251,211],"isMain":false},{"metricName":"LA75","quality":"Less is better","values":[602,476,972,877,620,446],"isMain":false}]],["Reads mapping",[]],["Misassemblies",[{"metricName":"# misassemblies","quality":"Less is better","values":[6,14,13,16,7,3],"isMain":true},{"metricName":" # relocations","quality":"Less is better","values":[5,13,9,14,4,3],"isMain":false},{"metricName":" # translocations","quality":"Less is better","values":[0,0,0,0,0,0],"isMain":false},{"metricName":" # inversions","quality":"Less is better","values":[1,1,4,2,3,0],"isMain":false},{"metricName":" # interspecies translocations","quality":"Less is better","values":[22,51,27,54,52,12],"isMain":false},{"metricName":"# misassembled contigs","quality":"Less is better","values":[6,13,13,15,7,3],"isMain":false},{"metricName":"Misassembled contigs length","quality":"Less is better","values":[18037,69572,31797,44981,16972,9567],"isMain":true},{"metricName":"# possibly misassembled contigs","quality":"Less is better","values":[0,0,0,0,0,0],"isMain":false},{"metricName":" # possible misassemblies","quality":"Less is better","values":[0,0,0,0,0,0],"isMain":false},{"metricName":"# local misassemblies","quality":"Less is better","values":[1,5,3,8,1,0],"isMain":false},{"metricName":"# scaffold gap ext. mis.","quality":"Less is better","values":[0,0,0,0,0,0],"isMain":false},{"metricName":"# scaffold gap loc. mis.","quality":"Less is better","values":[0,0,0,0,0,0],"isMain":false},{"metricName":"# unaligned mis. contigs","quality":"Less is better","values":[0,1,1,0,1,0],"isMain":false}]],["Unaligned",[{"metricName":"# fully unaligned contigs","quality":"Less is better","values":[1,3,2,0,1,0],"isMain":false},{"metricName":"Fully unaligned length","quality":"Less is better","values":[621,3931,2276,0,509,0],"isMain":false},{"metricName":"# partially unaligned contigs","quality":"Less is better","values":[4,18,8,18,17,5],"isMain":false},{"metricName":"Partially unaligned length","quality":"Less is better","values":[11075,31904,56703,50672,103280,4609],"isMain":false}]],["Mismatches",[{"metricName":"# mismatches","quality":"Less is better","values":[14607,18659,29002,29578,19162,8619],"isMain":false},{"metricName":"# indels","quality":"Less is better","values":[96,134,137,190,126,44],"isMain":false},{"metricName":"Indels length","quality":"Less is better","values":[377,303,214,481,316,94],"isMain":false},{"metricName":"# mismatches per 100 kbp","quality":"Less is better","values":["893.41","978.41","1465.88","1506.84","1084.60","874.28"],"isMain":true},{"metricName":"# indels per 100 kbp","quality":"Less is better","values":["5.87","7.03","6.92","9.68","7.13","4.46"],"isMain":true},{"metricName":" # indels (<= 5 bp)","quality":"Less is better","values":[87,128,132,177,114,41],"isMain":false},{"metricName":" # indels (> 5 bp)","quality":"Less is better","values":[9,6,5,13,12,3],"isMain":false},{"metricName":"# N's","quality":"Less is better","values":[0,0,0,0,0,0],"isMain":false},{"metricName":"# N's per 100 kbp","quality":"Less is better","values":["0.00","0.00","0.00","0.00","0.00","0.00"],"isMain":true}]],["Statistics without reference",[{"metricName":"# contigs","quality":"Equal","values":[1194,1061,1724,1610,1181,818],"isMain":true},{"metricName":"# contigs (>= 1000 bp)","quality":"Equal","values":[585,601,735,696,583,383],"isMain":false},{"metricName":"# contigs (>= 5000 bp)","quality":"Equal","values":[33,73,22,33,59,12],"isMain":false},{"metricName":"# contigs (>= 10000 bp)","quality":"Equal","values":[3,17,4,5,12,2],"isMain":false},{"metricName":"# contigs (>= 25000 bp)","quality":"Equal","values":[0,2,2,0,2,0],"isMain":false},{"metricName":"# contigs (>= 50000 bp)","quality":"Equal","values":[0,0,0,0,1,0],"isMain":false},{"metricName":"Largest contig","quality":"More is better","values":[22683,27990,30188,24857,53082,16707],"isMain":true},{"metricName":"Total length","quality":"More is better","values":[1703181,2012993,2150813,2135855,1952742,1025524],"isMain":true},{"metricName":"Total length (>= 1000 bp)","quality":"More is better","values":[1277763,1691750,1472244,1500923,1542058,720382],"isMain":true},{"metricName":"Total length (>= 5000 bp)","quality":"More is better","values":[244360,623681,202167,257386,512520,92548],"isMain":false},{"metricName":"Total length (>= 10000 bp)","quality":"More is better","values":[48388,246692,82335,71631,209542,28271],"isMain":true},{"metricName":"Total length (>= 25000 bp)","quality":"More is better","values":[0,55424,58412,0,86492,0],"isMain":false},{"metricName":"Total length (>= 50000 bp)","quality":"More is better","values":[0,0,0,0,53082,0],"isMain":true},{"metricName":"N50","quality":"More is better","values":[1785,2688,1432,1633,2269,1422],"isMain":false},{"metricName":"N75","quality":"More is better","values":[1002,1352,862,902,1124,898],"isMain":false},{"metricName":"L50","quality":"Less is better","values":[261,182,401,354,202,207],"isMain":false},{"metricName":"L75","quality":"Less is better","values":[585,444,889,803,511,435],"isMain":false},{"metricName":"GC (%)","quality":"Equal","values":["41.40","41.92","41.99","41.89","41.53","40.48"],"isMain":false}]],["K-mer-based statistics",[]],["Predicted genes",[]],["Similarity statistics",[]],["Reference statistics",[{"metricName":"Reference length","quality":"Equal","values":[6293399,6293399,6293399,6293399,6293399,6293399],"isMain":false},{"metricName":"Reference fragments","quality":"Equal","values":[2,2,2,2,2,2],"isMain":false},{"metricName":"Reference GC (%)","quality":"Equal","values":["42.86","42.86","42.86","42.86","42.86","42.86"],"isMain":false}]]],"subreferences":[],"subreports":[],"minContig":500},{"date":"10 February 2020, Monday, 18:00:28","assembliesNames":["megahit-103-df","megahit-113-df","megahit-113-ml","megahit-113-ms","megahit-129-df","metaSPAdes"],"referenceName":"GCF\_000012845.1\_ASM1284v1","order":[0,1,2,3,4,5],"report":[["Genome statistics",[{"metricName":"Genome fraction (%)","quality":"More is better","values":["4.161","4.776","4.638","4.624","5.172","3.640"],"isMain":true},{"metricName":"Duplication ratio","quality":"Less is better","values":["1.010","1.024","1.019","1.028","1.013","1.017"],"isMain":true},{"metricName":"Largest alignment","quality":"More is better","values":[2921,2567,1822,2246,2567,1599],"isMain":true},{"metricName":"Total aligned length","quality":"More is better","values":[200562,230301,224972,224256,249848,175490],"isMain":true},{"metricName":"NA50","quality":"More is better","values":[797,888,710,724,917,689],"isMain":false},{"metricName":"NA75","quality":"More is better","values":[618,629,588,580,659,579],"isMain":false},{"metricName":"NGA50","quality":"More is better","values":[null,null,null,null,null,null],"isMain":true},{"metricName":"LA50","quality":"Less is better","values":[93,93,122,118,96,98],"isMain":false},{"metricName":"LA75","quality":"Less is better","values":[166,172,212,209,177,169],"isMain":false}]],["Reads mapping",[]],["Misassemblies",[{"metricName":"# misassemblies","quality":"Less is better","values":[0,2,0,1,5,0],"isMain":true},{"metricName":" # relocations","quality":"Less is better","values":[0,2,0,1,4,0],"isMain":false},{"metricName":" # translocations","quality":"Less is better","values":[0,0,0,0,0,0],"isMain":false},{"metricName":" # inversions","quality":"Less is better","values":[0,0,0,0,1,0],"isMain":false},{"metricName":" # interspecies translocations","quality":"Less is better","values":[5,13,2,14,7,3],"isMain":false},{"metricName":"# misassembled contigs","quality":"Less is better","values":[0,2,0,1,5,0],"isMain":false},{"metricName":"Misassembled contigs length","quality":"Less is better","values":[0,1674,0,665,3501,0],"isMain":true},{"metricName":"# possibly misassembled contigs","quality":"Less is better","values":[0,0,0,0,0,0],"isMain":false},{"metricName":" # possible misassemblies","quality":"Less is better","values":[0,0,0,0,0,0],"isMain":false},{"metricName":"# local misassemblies","quality":"Less is better","values":[1,1,1,0,0,1],"isMain":false},{"metricName":"# scaffold gap ext. mis.","quality":"Less is better","values":[0,0,0,0,0,0],"isMain":false},{"metricName":"# scaffold gap loc. mis.","quality":"Less is better","values":[0,0,0,0,0,0],"isMain":false},{"metricName":"# unaligned mis. contigs","quality":"Less is better","values":[0,0,0,2,0,0],"isMain":false}]],["Unaligned",[{"metricName":"# fully unaligned contigs","quality":"Less is better","values":[0,0,0,0,0,0],"isMain":false},{"metricName":"Fully unaligned length","quality":"Less is better","values":[0,0,0,0,0,0],"isMain":false},{"metricName":"# partially unaligned contigs","quality":"Less is better","values":[1,2,0,5,1,1],"isMain":false},{"metricName":"Partially unaligned length","quality":"Less is better","values":[593,2012,0,4725,511,593],"isMain":false}]],["Mismatches",[{"metricName":"# mismatches","quality":"Less is better","values":[3200,3854,4904,4898,4298,2910],"isMain":false},{"metricName":"# indels","quality":"Less is better","values":[11,14,10,15,18,9],"isMain":false},{"metricName":"Indels length","quality":"Less is better","values":[79,63,10,18,146,37],"isMain":false},{"metricName":"# mismatches per 100 kbp","quality":"Less is better","values":["1598.56","1677.20","2197.56","2201.35","1727.21","1661.42"],"isMain":true},{"metricName":"# indels per 100 kbp","quality":"Less is better","values":["5.50","6.09","4.48","6.74","7.23","5.14"],"isMain":true},{"metricName":" # indels (<= 5 bp)","quality":"Less is better","values":[8,12,10,15,15,5],"isMain":false},{"metricName":" # indels (> 5 bp)","quality":"Less is better","values":[3,2,0,0,3,4],"isMain":false},{"metricName":"# N's","quality":"Less is better","values":[0,0,0,0,0,0],"isMain":false},{"metricName":"# N's per 100 kbp","quality":"Less is better","values":["0.00","0.00","0.00","0.00","0.00","0.00"],"isMain":true}]],["Statistics without reference",[{"metricName":"# contigs","quality":"Equal","values":[255,268,312,305,284,248],"isMain":true},{"metricName":"# contigs (>= 1000 bp)","quality":"Equal","values":[44,71,33,43,80,28],"isMain":false},{"metricName":"# contigs (>= 5000 bp)","quality":"Equal","values":[0,0,0,0,0,0],"isMain":false},{"metricName":"# contigs (>= 10000 bp)","quality":"Equal","values":[0,0,0,0,0,0],"isMain":false},{"metricName":"# contigs (>= 25000 bp)","quality":"Equal","values":[0,0,0,0,0,0],"isMain":false},{"metricName":"# contigs (>= 50000 bp)","quality":"Equal","values":[0,0,0,0,0,0],"isMain":false},{"metricName":"Largest contig","quality":"More is better","values":[2921,2567,1903,2246,2567,1617],"isMain":true},{"metricName":"Total length","quality":"More is better","values":[202746,237391,227494,233408,252653,178737],"isMain":true},{"metricName":"Total length (>= 1000 bp)","quality":"More is better","values":[58947,101081,40714,55304,111701,33423],"isMain":true},{"metricName":"Total length (>= 5000 bp)","quality":"More is better","values":[0,0,0,0,0,0],"isMain":false},{"metricName":"Total length (>= 10000 bp)","quality":"More is better","values":[0,0,0,0,0,0],"isMain":true},{"metricName":"Total length (>= 25000 bp)","quality":"More is better","values":[0,0,0,0,0,0],"isMain":false},{"metricName":"Total length (>= 50000 bp)","quality":"More is better","values":[0,0,0,0,0,0],"isMain":true},{"metricName":"N50","quality":"More is better","values":[805,932,715,746,932,700],"isMain":false},{"metricName":"N75","quality":"More is better","values":[625,692,598,607,679,596],"isMain":false},{"metricName":"L50","quality":"Less is better","values":[92,90,121,114,96,97],"isMain":false},{"metricName":"L75","quality":"Less is better","values":[164,164,209,201,174,166],"isMain":false},{"metricName":"GC (%)","quality":"Equal","values":["45.05","45.31","45.52","44.79","45.07","45.30"],"isMain":false}]],["K-mer-based statistics",[]],["Predicted genes",[]],["Similarity statistics",[]],["Reference statistics",[{"metricName":"Reference length","quality":"Equal","values":[4811379,4811379,4811379,4811379,4811379,4811379],"isMain":false},{"metricName":"Reference fragments","quality":"Equal","values":[1,1,1,1,1,1],"isMain":false},{"metricName":"Reference GC (%)","quality":"Equal","values":["45.06","45.06","45.06","45.06","45.06","45.06"],"isMain":false}]]],"subreferences":[],"subreports":[],"minContig":500},{"date":"10 February 2020, Monday, 18:00:43","assembliesNames":["megahit-103-df","megahit-113-df","megahit-113-ml","megahit-113-ms","megahit-129-df","metaSPAdes"],"referenceName":"GCF\_000015325.1\_ASM1532v1","order":[0,1,2,3,4,5],"report":[["Genome statistics",[{"metricName":"Genome fraction (%)","quality":"More is better","values":["2.113","2.816","3.205","3.828","3.238","1.303"],"isMain":true},{"metricName":"Duplication ratio","quality":"Less is better","values":["1.047","1.055","1.030","1.045","1.026","1.088"],"isMain":true},{"metricName":"Largest alignment","quality":"More is better","values":[1664,2030,2019,2288,2168,2097],"isMain":true},{"metricName":"Total aligned length","quality":"More is better","values":[113085,150824,171965,205809,173610,69726],"isMain":true},{"metricName":"NA50","quality":"More is better","values":[577,578,576,550,598,551],"isMain":false},{"metricName":"NA75","quality":"More is better","values":[516,515,527,514,531,506],"isMain":false},{"metricName":"NGA50","quality":"More is better","values":[null,null,null,null,null,null],"isMain":true},{"metricName":"LA50","quality":"Less is better","values":[84,113,128,169,121,56],"isMain":false},{"metricName":"LA75","quality":"Less is better","values":[141,190,209,276,202,94],"isMain":false}]],["Reads mapping",[]],["Misassemblies",[{"metricName":"# misassemblies","quality":"Less is better","values":[0,1,2,6,5,2],"isMain":true},{"metricName":" # relocations","quality":"Less is better","values":[0,1,2,6,5,2],"isMain":false},{"metricName":" # translocations","quality":"Less is better","values":[0,0,0,0,0,0],"isMain":false},{"metricName":" # inversions","quality":"Less is better","values":[0,0,0,0,0,0],"isMain":false},{"metricName":" # interspecies translocations","quality":"Less is better","values":[21,29,11,29,18,20],"isMain":false},{"metricName":"# misassembled contigs","quality":"Less is better","values":[0,1,2,6,5,2],"isMain":false},{"metricName":"Misassembled contigs length","quality":"Less is better","values":[0,1142,2197,6488,5869,1894],"isMain":true},{"metricName":"# possibly misassembled contigs","quality":"Less is better","values":[0,0,0,0,0,0],"isMain":false},{"metricName":" # possible misassemblies","quality":"Less is better","values":[0,0,0,0,0,0],"isMain":false},{"metricName":"# local misassemblies","quality":"Less is better","values":[0,0,1,0,1,0],"isMain":false},{"metricName":"# scaffold gap ext. mis.","quality":"Less is better","values":[0,0,0,0,0,0],"isMain":false},{"metricName":"# scaffold gap loc. mis.","quality":"Less is better","values":[0,0,0,0,0,0],"isMain":false},{"metricName":"# unaligned mis. contigs","quality":"Less is better","values":[0,0,0,0,0,0],"isMain":false}]],["Unaligned",[{"metricName":"# fully unaligned contigs","quality":"Less is better","values":[0,1,1,0,1,0],"isMain":false},{"metricName":"Fully unaligned length","quality":"Less is better","values":[0,580,600,0,661,0],"isMain":false},{"metricName":"# partially unaligned contigs","quality":"Less is better","values":[4,7,1,10,3,5],"isMain":false},{"metricName":"Partially unaligned length","quality":"Less is better","values":[3661,7101,603,12259,3720,4456],"isMain":false}]],["Mismatches",[{"metricName":"# mismatches","quality":"Less is better","values":[1834,2525,3447,4345,3026,1235],"isMain":false},{"metricName":"# indels","quality":"Less is better","values":[0,6,9,11,8,4],"isMain":false},{"metricName":"Indels length","quality":"Less is better","values":[0,6,12,37,9,13],"isMain":false},{"metricName":"# mismatches per 100 kbp","quality":"Less is better","values":["1621.79","1675.19","2009.35","2120.66","1745.85","1770.94"],"isMain":true},{"metricName":"# indels per 100 kbp","quality":"Less is better","values":["0.00","3.98","5.25","5.37","4.62","5.74"],"isMain":true},{"metricName":" # indels (<= 5 bp)","quality":"Less is better","values":[0,6,9,10,8,3],"isMain":false},{"metricName":" # indels (> 5 bp)","quality":"Less is better","values":[0,0,0,1,0,1],"isMain":false},{"metricName":"# N's","quality":"Less is better","values":[0,0,0,0,0,0],"isMain":false},{"metricName":"# N's per 100 kbp","quality":"Less is better","values":["0.00","0.00","0.00","0.00","0.00","0.00"],"isMain":true}]],["Statistics without reference",[{"metricName":"# contigs","quality":"Equal","values":[190,253,289,360,279,122],"isMain":true},{"metricName":"# contigs (>= 1000 bp)","quality":"Equal","values":[8,16,6,10,14,8],"isMain":false},{"metricName":"# contigs (>= 5000 bp)","quality":"Equal","values":[0,0,0,1,0,0],"isMain":false},{"metricName":"# contigs (>= 10000 bp)","quality":"Equal","values":[0,0,0,0,0,0],"isMain":false},{"metricName":"# contigs (>= 25000 bp)","quality":"Equal","values":[0,0,0,0,0,0],"isMain":false},{"metricName":"# contigs (>= 50000 bp)","quality":"Equal","values":[0,0,0,0,0,0],"isMain":false},{"metricName":"Largest contig","quality":"More is better","values":[1732,2216,2169,5572,2174,2234],"isMain":true},{"metricName":"Total length","quality":"More is better","values":[122010,166769,177941,226361,182174,80296],"isMain":true},{"metricName":"Total length (>= 1000 bp)","quality":"More is better","values":[11330,21619,9061,18746,19688,12199],"isMain":true},{"metricName":"Total length (>= 5000 bp)","quality":"More is better","values":[0,0,0,5572,0,0],"isMain":false},{"metricName":"Total length (>= 10000 bp)","quality":"More is better","values":[0,0,0,0,0,0],"isMain":true},{"metricName":"Total length (>= 25000 bp)","quality":"More is better","values":[0,0,0,0,0,0],"isMain":false},{"metricName":"Total length (>= 50000 bp)","quality":"More is better","values":[0,0,0,0,0,0],"isMain":true},{"metricName":"N50","quality":"More is better","values":[615,613,592,586,623,604],"isMain":false},{"metricName":"N75","quality":"More is better","values":[540,546,539,531,545,543],"isMain":false},{"metricName":"L50","quality":"Less is better","values":[78,102,125,149,114,49],"isMain":false},{"metricName":"L75","quality":"Less is better","values":[132,173,204,251,192,84],"isMain":false},{"metricName":"GC (%)","quality":"Equal","values":["66.78","66.75","67.43","66.78","66.93","67.16"],"isMain":false}]],["K-mer-based statistics",[]],["Predicted genes",[]],["Similarity statistics",[]],["Reference statistics",[{"metricName":"Reference length","quality":"Equal","values":[5352772,5352772,5352772,5352772,5352772,5352772],"isMain":false},{"metricName":"Reference fragments","quality":"Equal","values":[1,1,1,1,1,1],"isMain":false},{"metricName":"Reference GC (%)","quality":"Equal","values":["68.53","68.53","68.53","68.53","68.53","68.53"],"isMain":false}]]],"subreferences":[],"subreports":[],"minContig":500},{"date":"10 February 2020, Monday, 18:00:38","assembliesNames":["megahit-103-df","megahit-113-df","megahit-113-ml","megahit-113-ms","megahit-129-df","metaSPAdes"],"referenceName":"GCF\_000018685.1\_ASM1868v1","order":[0,1,2,3,4,5],"report":[["Genome statistics",[{"metricName":"Genome fraction (%)","quality":"More is better","values":["41.423","47.511","43.043","46.277","50.595","32.122"],"isMain":true},{"metricName":"Duplication ratio","quality":"Less is better","values":["1.031","1.029","1.037","1.048","1.029","1.036"],"isMain":true},{"metricName":"Largest alignment","quality":"More is better","values":[2404,2719,2284,2604,2704,2088],"isMain":true},{"metricName":"Total aligned length","quality":"More is better","values":[2044220,2339076,2143531,2314876,2495745,1581673],"isMain":true},{"metricName":"NG50","quality":"More is better","values":[null,null,null,null,529,null],"isMain":false},{"metricName":"NA50","quality":"More is better","values":[739,806,712,719,825,671],"isMain":false},{"metricName":"NA75","quality":"More is better","values":[593,622,581,580,629,560],"isMain":false},{"metricName":"NGA50","quality":"More is better","values":[null,null,null,null,508,null],"isMain":true},{"metricName":"LG50","quality":"Less is better","values":[null,null,null,null,2813,null],"isMain":false},{"metricName":"LA50","quality":"Less is better","values":[1051,1088,1152,1229,1124,938],"isMain":false},{"metricName":"LA75","quality":"Less is better","values":[1842,1939,2000,2148,2015,1605],"isMain":false},{"metricName":"LGA50","quality":"Less is better","values":[null,null,null,null,2911,null],"isMain":true}]],["Reads mapping",[]],["Misassemblies",[{"metricName":"# misassemblies","quality":"Less is better","values":[4,14,7,8,8,2],"isMain":true},{"metricName":" # relocations","quality":"Less is better","values":[4,13,3,3,7,2],"isMain":false},{"metricName":" # translocations","quality":"Less is better","values":[0,0,0,0,0,0],"isMain":false},{"metricName":" # inversions","quality":"Less is better","values":[0,1,4,5,1,0],"isMain":false},{"metricName":" # interspecies translocations","quality":"Less is better","values":[82,95,10,82,79,69],"isMain":false},{"metricName":"# misassembled contigs","quality":"Less is better","values":[4,14,7,8,8,2],"isMain":false},{"metricName":"Misassembled contigs length","quality":"Less is better","values":[3505,13674,7281,8047,7688,2728],"isMain":true},{"metricName":"# possibly misassembled contigs","quality":"Less is better","values":[0,0,0,0,0,0],"isMain":false},{"metricName":" # possible misassemblies","quality":"Less is better","values":[0,0,0,0,0,0],"isMain":false},{"metricName":"# local misassemblies","quality":"Less is better","values":[2,4,4,2,3,1],"isMain":false},{"metricName":"# scaffold gap ext. mis.","quality":"Less is better","values":[0,0,0,0,0,0],"isMain":false},{"metricName":"# scaffold gap loc. mis.","quality":"Less is better","values":[0,0,0,0,0,0],"isMain":false},{"metricName":"# unaligned mis. contigs","quality":"Less is better","values":[0,0,1,0,0,0],"isMain":false}]],["Unaligned",[{"metricName":"# fully unaligned contigs","quality":"Less is better","values":[2,2,1,1,3,0],"isMain":false},{"metricName":"Fully unaligned length","quality":"Less is better","values":[1540,1652,796,513,3625,0],"isMain":false},{"metricName":"# partially unaligned contigs","quality":"Less is better","values":[9,20,1,15,23,16],"isMain":false},{"metricName":"Partially unaligned length","quality":"Less is better","values":[7975,34323,672,13703,31477,12013],"isMain":false}]],["Mismatches",[{"metricName":"# mismatches","quality":"Less is better","values":[26436,29224,39098,43978,31774,22197],"isMain":false},{"metricName":"# indels","quality":"Less is better","values":[37,49,67,74,74,29],"isMain":false},{"metricName":"Indels length","quality":"Less is better","values":[317,489,407,434,783,170],"isMain":false},{"metricName":"# mismatches per 100 kbp","quality":"Less is better","values":["1316.52","1268.88","1873.80","1960.38","1295.51","1425.49"],"isMain":true},{"metricName":"# indels per 100 kbp","quality":"Less is better","values":["1.84","2.13","3.21","3.30","3.02","1.86"],"isMain":true},{"metricName":" # indels (<= 5 bp)","quality":"Less is better","values":[30,37,61,65,53,25],"isMain":false},{"metricName":" # indels (> 5 bp)","quality":"Less is better","values":[7,12,6,9,21,4],"isMain":false},{"metricName":"# N's","quality":"Less is better","values":[0,0,0,0,0,0],"isMain":false},{"metricName":"# N's per 100 kbp","quality":"Less is better","values":["0.00","0.00","0.00","0.00","0.00","0.00"],"isMain":true}]],["Statistics without reference",[{"metricName":"# contigs","quality":"Equal","values":[2758,2923,2972,3182,3073,2318],"isMain":true},{"metricName":"# contigs (>= 1000 bp)","quality":"Equal","values":[388,600,341,400,654,197],"isMain":false},{"metricName":"# contigs (>= 5000 bp)","quality":"Equal","values":[0,1,0,0,1,0],"isMain":false},{"metricName":"# contigs (>= 10000 bp)","quality":"Equal","values":[0,1,0,0,1,0],"isMain":false},{"metricName":"# contigs (>= 25000 bp)","quality":"Equal","values":[0,0,0,0,0,0],"isMain":false},{"metricName":"# contigs (>= 50000 bp)","quality":"Equal","values":[0,0,0,0,0,0],"isMain":false},{"metricName":"Largest contig","quality":"More is better","values":[2700,15327,2738,2669,12761,2232],"isMain":true},{"metricName":"Total length","quality":"More is better","values":[2079516,2405005,2166230,2365066,2557763,1625153],"isMain":true},{"metricName":"Total length (>= 1000 bp)","quality":"More is better","values":[485728,798424,416875,505062,881478,239611],"isMain":true},{"metricName":"Total length (>= 5000 bp)","quality":"More is better","values":[0,15327,0,0,12761,0],"isMain":false},{"metricName":"Total length (>= 10000 bp)","quality":"More is better","values":[0,15327,0,0,12761,0],"isMain":true},{"metricName":"Total length (>= 25000 bp)","quality":"More is better","values":[0,0,0,0,0,0],"isMain":false},{"metricName":"Total length (>= 50000 bp)","quality":"More is better","values":[0,0,0,0,0,0],"isMain":true},{"metricName":"N50","quality":"More is better","values":[752,839,720,738,847,686],"isMain":false},{"metricName":"N75","quality":"More is better","values":[604,646,591,598,653,581],"isMain":false},{"metricName":"L50","quality":"Less is better","values":[1035,1043,1140,1202,1088,914],"isMain":false},{"metricName":"L75","quality":"Less is better","values":[1811,1864,1975,2095,1951,1562],"isMain":false},{"metricName":"GC (%)","quality":"Equal","values":["35.54","35.58","35.61","35.65","35.54","35.56"],"isMain":false}]],["K-mer-based statistics",[]],["Predicted genes",[]],["Similarity statistics",[]],["Reference statistics",[{"metricName":"Reference length","quality":"Equal","values":[4847594,4847594,4847594,4847594,4847594,4847594],"isMain":false},{"metricName":"Reference fragments","quality":"Equal","values":[1,1,1,1,1,1],"isMain":false},{"metricName":"Reference GC (%)","quality":"Equal","values":["35.35","35.35","35.35","35.35","35.35","35.35"],"isMain":false}]]],"subreferences":[],"subreports":[],"minContig":500},{"date":"10 February 2020, Monday, 18:00:05","assembliesNames":["megahit-103-df","megahit-113-df","megahit-113-ml","megahit-113-ms","megahit-129-df","metaSPAdes"],"referenceName":"GCF\_000020225.1\_ASM2022v1","order":[0,1,2,3,4,5],"report":[["Genome statistics",[{"metricName":"Genome fraction (%)","quality":"More is better","values":["3.588","3.198","5.465","3.980","2.572","1.207"],"isMain":true},{"metricName":"Duplication ratio","quality":"Less is better","values":["1.033","1.056","1.045","1.044","1.048","1.039"],"isMain":true},{"metricName":"Largest alignment","quality":"More is better","values":[1577,7492,2341,2287,2148,3067],"isMain":true},{"metricName":"Total aligned length","quality":"More is better","values":[96482,87246,148413,107704,69938,32662],"isMain":true},{"metricName":"NA50","quality":"More is better","values":[641,769,706,736,728,711],"isMain":false},{"metricName":"NA75","quality":"More is better","values":[548,575,579,585,574,575],"isMain":false},{"metricName":"NGA50","quality":"More is better","values":[null,null,null,null,null,null],"isMain":true},{"metricName":"LA50","quality":"Less is better","values":[55,38,80,57,35,16],"isMain":false},{"metricName":"LA75","quality":"Less is better","values":[97,73,141,101,63,30],"isMain":false}]],["Reads mapping",[]],["Misassemblies",[{"metricName":"# misassemblies","quality":"Less is better","values":[0,1,2,0,0,0],"isMain":true},{"metricName":" # relocations","quality":"Less is better","values":[0,0,1,0,0,0],"isMain":false},{"metricName":" # translocations","quality":"Less is better","values":[0,0,0,0,0,0],"isMain":false},{"metricName":" # inversions","quality":"Less is better","values":[0,1,1,0,0,0],"isMain":false},{"metricName":" # interspecies translocations","quality":"Less is better","values":[3,13,6,9,5,3],"isMain":false},{"metricName":"# misassembled contigs","quality":"Less is better","values":[0,1,2,0,0,0],"isMain":false},{"metricName":"Misassembled contigs length","quality":"Less is better","values":[0,984,1131,0,0,0],"isMain":true},{"metricName":"# possibly misassembled contigs","quality":"Less is better","values":[0,0,0,0,0,0],"isMain":false},{"metricName":" # possible misassemblies","quality":"Less is better","values":[0,0,0,0,0,0],"isMain":false},{"metricName":"# local misassemblies","quality":"Less is better","values":[1,0,3,3,0,0],"isMain":false},{"metricName":"# scaffold gap ext. mis.","quality":"Less is better","values":[0,0,0,0,0,0],"isMain":false},{"metricName":"# scaffold gap loc. mis.","quality":"Less is better","values":[0,0,0,0,0,0],"isMain":false},{"metricName":"# unaligned mis. contigs","quality":"Less is better","values":[0,0,0,0,0,0],"isMain":false}]],["Unaligned",[{"metricName":"# fully unaligned contigs","quality":"Less is better","values":[0,1,0,1,0,0],"isMain":false},{"metricName":"Fully unaligned length","quality":"Less is better","values":[0,1967,0,559,0,0],"isMain":false},{"metricName":"# partially unaligned contigs","quality":"Less is better","values":[0,4,1,3,2,2],"isMain":false},{"metricName":"Partially unaligned length","quality":"Less is better","values":[0,2447,972,2881,1126,1289],"isMain":false}]],["Mismatches",[{"metricName":"# mismatches","quality":"Less is better","values":[1649,2233,3760,2663,1384,501],"isMain":false},{"metricName":"# indels","quality":"Less is better","values":[29,33,31,36,15,7],"isMain":false},{"metricName":"Indels length","quality":"Less is better","values":[33,40,49,136,17,7],"isMain":false},{"metricName":"# mismatches per 100 kbp","quality":"Less is better","values":["1724.99","2620.98","2582.59","2511.70","2019.61","1557.59"],"isMain":true},{"metricName":"# indels per 100 kbp","quality":"Less is better","values":["30.34","38.73","21.29","33.95","21.89","21.76"],"isMain":true},{"metricName":" # indels (<= 5 bp)","quality":"Less is better","values":[29,33,29,34,15,7],"isMain":false},{"metricName":" # indels (> 5 bp)","quality":"Less is better","values":[0,0,2,2,0,0],"isMain":false},{"metricName":"# N's","quality":"Less is better","values":[0,0,0,0,0,0],"isMain":false},{"metricName":"# N's per 100 kbp","quality":"Less is better","values":["0.00","0.00","0.00","0.00","0.00","0.00"],"isMain":true}]],["Statistics without reference",[{"metricName":"# contigs","quality":"Equal","values":[142,109,207,146,93,43],"isMain":true},{"metricName":"# contigs (>= 1000 bp)","quality":"Equal","values":[15,19,26,21,16,6],"isMain":false},{"metricName":"# contigs (>= 5000 bp)","quality":"Equal","values":[0,1,0,0,0,0],"isMain":false},{"metricName":"# contigs (>= 10000 bp)","quality":"Equal","values":[0,0,0,0,0,0],"isMain":false},{"metricName":"# contigs (>= 25000 bp)","quality":"Equal","values":[0,0,0,0,0,0],"isMain":false},{"metricName":"# contigs (>= 50000 bp)","quality":"Equal","values":[0,0,0,0,0,0],"isMain":false},{"metricName":"Largest contig","quality":"More is better","values":[1577,7533,2341,2287,2148,3067],"isMain":true},{"metricName":"Total length","quality":"More is better","values":[98787,94354,153123,114132,72936,34695],"isMain":true},{"metricName":"Total length (>= 1000 bp)","quality":"More is better","values":[18738,32675,33255,28278,21559,9352],"isMain":true},{"metricName":"Total length (>= 5000 bp)","quality":"More is better","values":[0,7533,0,0,0,0],"isMain":false},{"metricName":"Total length (>= 10000 bp)","quality":"More is better","values":[0,0,0,0,0,0],"isMain":true},{"metricName":"Total length (>= 25000 bp)","quality":"More is better","values":[0,0,0,0,0,0],"isMain":false},{"metricName":"Total length (>= 50000 bp)","quality":"More is better","values":[0,0,0,0,0,0],"isMain":true},{"metricName":"N50","quality":"More is better","values":[647,827,724,782,789,798],"isMain":false},{"metricName":"N75","quality":"More is better","values":[561,625,597,621,608,626],"isMain":false},{"metricName":"L50","quality":"Less is better","values":[55,36,79,54,34,16],"isMain":false},{"metricName":"L75","quality":"Less is better","values":[96,68,138,96,60,28],"isMain":false},{"metricName":"GC (%)","quality":"Equal","values":["53.87","54.20","53.72","53.73","52.34","52.70"],"isMain":false}]],["K-mer-based statistics",[]],["Predicted genes",[]],["Similarity statistics",[]],["Reference statistics",[{"metricName":"Reference length","quality":"Equal","values":[2664102,2664102,2664102,2664102,2664102,2664102],"isMain":false},{"metricName":"Reference fragments","quality":"Equal","values":[1,1,1,1,1,1],"isMain":false},{"metricName":"Reference GC (%)","quality":"Equal","values":["55.76","55.76","55.76","55.76","55.76","55.76"],"isMain":false}]]],"subreferences":[],"subreports":[],"minContig":500},{"date":"10 February 2020, Monday, 18:00:24","assembliesNames":["megahit-103-df","megahit-113-df","megahit-113-ml","megahit-113-ms","megahit-129-df","metaSPAdes"],"referenceName":"GCF\_000022065.1\_ASM2206v1","order":[0,1,2,3,4,5],"report":[["Genome statistics",[{"metricName":"Genome fraction (%)","quality":"More is better","values":["29.520","35.144","32.340","35.114","37.469","22.271"],"isMain":true},{"metricName":"Duplication ratio","quality":"Less is better","values":["1.026","1.028","1.028","1.038","1.028","1.036"],"isMain":true},{"metricName":"Largest alignment","quality":"More is better","values":[3050,5226,2575,3254,5393,5267],"isMain":true},{"metricName":"Total aligned length","quality":"More is better","values":[1217306,1448425,1337289,1459562,1545820,917439],"isMain":true},{"metricName":"NA50","quality":"More is better","values":[697,740,672,689,763,630],"isMain":false},{"metricName":"NA75","quality":"More is better","values":[577,591,569,564,598,546],"isMain":false},{"metricName":"NGA50","quality":"More is better","values":[null,null,null,null,null,null],"isMain":true},{"metricName":"LA50","quality":"Less is better","values":[675,734,769,835,762,578],"isMain":false},{"metricName":"LA75","quality":"Less is better","values":[1166,1302,1323,1450,1350,983],"isMain":false}]],["Reads mapping",[]],["Misassemblies",[{"metricName":"# misassemblies","quality":"Less is better","values":[5,7,9,9,4,2],"isMain":true},{"metricName":" # relocations","quality":"Less is better","values":[5,5,7,8,3,2],"isMain":false},{"metricName":" # translocations","quality":"Less is better","values":[0,0,0,0,0,0],"isMain":false},{"metricName":" # inversions","quality":"Less is better","values":[0,2,2,1,1,0],"isMain":false},{"metricName":" # interspecies translocations","quality":"Less is better","values":[47,66,15,51,56,47],"isMain":false},{"metricName":"# misassembled contigs","quality":"Less is better","values":[5,7,9,9,4,2],"isMain":false},{"metricName":"Misassembled contigs length","quality":"Less is better","values":[5405,10666,11709,12802,3834,2110],"isMain":true},{"metricName":"# possibly misassembled contigs","quality":"Less is better","values":[0,0,0,0,0,0],"isMain":false},{"metricName":" # possible misassemblies","quality":"Less is better","values":[0,0,0,0,0,0],"isMain":false},{"metricName":"# local misassemblies","quality":"Less is better","values":[3,0,2,0,2,0],"isMain":false},{"metricName":"# scaffold gap ext. mis.","quality":"Less is better","values":[0,0,0,0,0,0],"isMain":false},{"metricName":"# scaffold gap loc. mis.","quality":"Less is better","values":[0,0,0,0,0,0],"isMain":false},{"metricName":"# unaligned mis. contigs","quality":"Less is better","values":[0,0,0,0,0,0],"isMain":false}]],["Unaligned",[{"metricName":"# fully unaligned contigs","quality":"Less is better","values":[1,0,0,0,0,0],"isMain":false},{"metricName":"Fully unaligned length","quality":"Less is better","values":[525,0,0,0,0,0],"isMain":false},{"metricName":"# partially unaligned contigs","quality":"Less is better","values":[10,15,3,12,12,11],"isMain":false},{"metricName":"Partially unaligned length","quality":"Less is better","values":[7213,20798,11551,44142,14572,7103],"isMain":false}]],["Mismatches",[{"metricName":"# mismatches","quality":"Less is better","values":[16821,19267,25099,28463,21057,13550],"isMain":false},{"metricName":"# indels","quality":"Less is better","values":[21,32,23,21,39,14],"isMain":false},{"metricName":"Indels length","quality":"Less is better","values":[96,166,55,28,176,38],"isMain":false},{"metricName":"# mismatches per 100 kbp","quality":"Less is better","values":["1400.46","1347.43","1907.46","1992.22","1381.24","1495.33"],"isMain":true},{"metricName":"# indels per 100 kbp","quality":"Less is better","values":["1.75","2.24","1.75","1.47","2.56","1.54"],"isMain":true},{"metricName":" # indels (<= 5 bp)","quality":"Less is better","values":[19,28,22,21,35,13],"isMain":false},{"metricName":" # indels (> 5 bp)","quality":"Less is better","values":[2,4,1,0,4,1],"isMain":false},{"metricName":"# N's","quality":"Less is better","values":[0,0,0,0,0,0],"isMain":false},{"metricName":"# N's per 100 kbp","quality":"Less is better","values":["0.00","0.00","0.00","0.00","0.00","0.00"],"isMain":true}]],["Statistics without reference",[{"metricName":"# contigs","quality":"Equal","values":[1713,1927,1918,2059,2027,1399],"isMain":true},{"metricName":"# contigs (>= 1000 bp)","quality":"Equal","values":[177,307,168,220,320,83],"isMain":false},{"metricName":"# contigs (>= 5000 bp)","quality":"Equal","values":[0,2,1,1,1,1],"isMain":false},{"metricName":"# contigs (>= 10000 bp)","quality":"Equal","values":[0,0,1,1,0,0],"isMain":false},{"metricName":"# contigs (>= 25000 bp)","quality":"Equal","values":[0,0,0,1,0,0],"isMain":false},{"metricName":"# contigs (>= 50000 bp)","quality":"Equal","values":[0,0,0,0,0,0],"isMain":false},{"metricName":"Largest contig","quality":"More is better","values":[3063,7777,10347,33053,5416,5302],"isMain":true},{"metricName":"Total length","quality":"More is better","values":[1240341,1490776,1364254,1526805,1581659,945958],"isMain":true},{"metricName":"Total length (>= 1000 bp)","quality":"More is better","values":[220928,401308,220287,310020,422078,105106],"isMain":true},{"metricName":"Total length (>= 5000 bp)","quality":"More is better","values":[0,13003,10347,33053,5416,5302],"isMain":false},{"metricName":"Total length (>= 10000 bp)","quality":"More is better","values":[0,0,10347,33053,0,0],"isMain":true},{"metricName":"Total length (>= 25000 bp)","quality":"More is better","values":[0,0,0,33053,0,0],"isMain":false},{"metricName":"Total length (>= 50000 bp)","quality":"More is better","values":[0,0,0,0,0,0],"isMain":true},{"metricName":"N50","quality":"More is better","values":[709,766,687,719,780,648],"isMain":false},{"metricName":"N75","quality":"More is better","values":[591,614,580,590,617,562],"isMain":false},{"metricName":"L50","quality":"Less is better","values":[662,704,743,767,743,562],"isMain":false},{"metricName":"L75","quality":"Less is better","values":[1142,1252,1285,1354,1313,955],"isMain":false},{"metricName":"GC (%)","quality":"Equal","values":["37.64","37.66","37.73","37.87","37.64","37.55"],"isMain":false}]],["K-mer-based statistics",[]],["Predicted genes",[]],["Similarity statistics",[]],["Reference statistics",[{"metricName":"Reference length","quality":"Equal","values":[4068724,4068724,4068724,4068724,4068724,4068724],"isMain":false},{"metricName":"Reference fragments","quality":"Equal","values":[1,1,1,1,1,1],"isMain":false},{"metricName":"Reference GC (%)","quality":"Equal","values":["37.40","37.40","37.40","37.40","37.40","37.40"],"isMain":false}]]],"subreferences":[],"subreports":[],"minContig":500},{"date":"10 February 2020, Monday, 18:00:36","assembliesNames":["megahit-103-df","megahit-113-df","megahit-113-ml","megahit-113-ms","megahit-129-df","metaSPAdes"],"referenceName":"GCF\_000026525.1\_ASM2652v1","order":[0,1,2,3,4,5],"report":[["Genome statistics",[{"metricName":"Genome fraction (%)","quality":"More is better","values":["2.392","3.536","3.436","4.355","2.928","2.236"],"isMain":true},{"metricName":"Duplication ratio","quality":"Less is better","values":["1.015","1.028","1.016","1.045","1.032","1.020"],"isMain":true},{"metricName":"Largest alignment","quality":"More is better","values":[1558,2559,1445,2045,2314,1766],"isMain":true},{"metricName":"Total aligned length","quality":"More is better","values":[72687,107356,104391,134423,89183,68199],"isMain":true},{"metricName":"NA50","quality":"More is better","values":[720,708,679,685,629,674],"isMain":false},{"metricName":"NA75","quality":"More is better","values":[559,554,573,545,284,561],"isMain":false},{"metricName":"NGA50","quality":"More is better","values":[null,null,null,null,null,null],"isMain":true},{"metricName":"LA50","quality":"Less is better","values":[39,57,60,76,61,40],"isMain":false},{"metricName":"LA75","quality":"Less is better","values":[69,106,103,134,116,68],"isMain":false}]],["Reads mapping",[]],["Misassemblies",[{"metricName":"# misassemblies","quality":"Less is better","values":[0,3,2,4,4,2],"isMain":true},{"metricName":" # relocations","quality":"Less is better","values":[0,3,2,4,4,2],"isMain":false},{"metricName":" # translocations","quality":"Less is better","values":[0,0,0,0,0,0],"isMain":false},{"metricName":" # inversions","quality":"Less is better","values":[0,0,0,0,0,0],"isMain":false},{"metricName":" # interspecies translocations","quality":"Less is better","values":[2,14,2,12,10,2],"isMain":false},{"metricName":"# misassembled contigs","quality":"Less is better","values":[0,3,2,4,2,2],"isMain":false},{"metricName":"Misassembled contigs length","quality":"Less is better","values":[0,2502,1608,4514,1776,3161],"isMain":true},{"metricName":"# possibly misassembled contigs","quality":"Less is better","values":[0,0,0,0,0,0],"isMain":false},{"metricName":" # possible misassemblies","quality":"Less is better","values":[0,0,0,0,0,0],"isMain":false},{"metricName":"# local misassemblies","quality":"Less is better","values":[0,2,0,0,3,0],"isMain":false},{"metricName":"# scaffold gap ext. mis.","quality":"Less is better","values":[0,0,0,0,0,0],"isMain":false},{"metricName":"# scaffold gap loc. mis.","quality":"Less is better","values":[0,0,0,0,0,0],"isMain":false},{"metricName":"# unaligned mis. contigs","quality":"Less is better","values":[0,1,0,0,0,0],"isMain":false}]],["Unaligned",[{"metricName":"# fully unaligned contigs","quality":"Less is better","values":[1,1,0,1,0,0],"isMain":false},{"metricName":"Fully unaligned length","quality":"Less is better","values":[569,1199,0,630,0,0],"isMain":false},{"metricName":"# partially unaligned contigs","quality":"Less is better","values":[0,3,1,3,5,0],"isMain":false},{"metricName":"Partially unaligned length","quality":"Less is better","values":[0,10806,1275,2724,25718,0],"isMain":false}]],["Mismatches",[{"metricName":"# mismatches","quality":"Less is better","values":[1538,2414,2627,3385,1949,1530],"isMain":false},{"metricName":"# indels","quality":"Less is better","values":[7,18,15,15,12,8],"isMain":false},{"metricName":"Indels length","quality":"Less is better","values":[7,114,23,84,15,8],"isMain":false},{"metricName":"# mismatches per 100 kbp","quality":"Less is better","values":["2120.00","2250.65","2520.39","2562.38","2194.50","2255.51"],"isMain":true},{"metricName":"# indels per 100 kbp","quality":"Less is better","values":["9.65","16.78","14.39","11.35","13.51","11.79"],"isMain":true},{"metricName":" # indels (<= 5 bp)","quality":"Less is better","values":[7,15,15,13,12,8],"isMain":false},{"metricName":" # indels (> 5 bp)","quality":"Less is better","values":[0,3,0,2,0,0],"isMain":false},{"metricName":"# N's","quality":"Less is better","values":[0,0,0,0,0,0],"isMain":false},{"metricName":"# N's per 100 kbp","quality":"Less is better","values":["0.00","0.00","0.00","0.00","0.00","0.00"],"isMain":true}]],["Statistics without reference",[{"metricName":"# contigs","quality":"Equal","values":[103,137,148,191,118,98],"isMain":true},{"metricName":"# contigs (>= 1000 bp)","quality":"Equal","values":[14,26,16,27,23,8],"isMain":false},{"metricName":"# contigs (>= 5000 bp)","quality":"Equal","values":[0,1,0,0,1,0],"isMain":false},{"metricName":"# contigs (>= 10000 bp)","quality":"Equal","values":[0,0,0,0,1,0],"isMain":false},{"metricName":"# contigs (>= 25000 bp)","quality":"Equal","values":[0,0,0,0,0,0],"isMain":false},{"metricName":"# contigs (>= 50000 bp)","quality":"Equal","values":[0,0,0,0,0,0],"isMain":false},{"metricName":"Largest contig","quality":"More is better","values":[1558,8522,2063,2075,21637,1771],"isMain":true},{"metricName":"Total length","quality":"More is better","values":[74228,122315,107142,141360,117375,69220],"isMain":true},{"metricName":"Total length (>= 1000 bp)","quality":"More is better","values":[16494,47128,19771,34217,55334,10507],"isMain":true},{"metricName":"Total length (>= 5000 bp)","quality":"More is better","values":[0,8522,0,0,21637,0],"isMain":false},{"metricName":"Total length (>= 10000 bp)","quality":"More is better","values":[0,0,0,0,21637,0],"isMain":true},{"metricName":"Total length (>= 25000 bp)","quality":"More is better","values":[0,0,0,0,0,0],"isMain":false},{"metricName":"Total length (>= 50000 bp)","quality":"More is better","values":[0,0,0,0,0,0],"isMain":true},{"metricName":"N50","quality":"More is better","values":[725,866,709,727,919,676],"isMain":false},{"metricName":"N75","quality":"More is better","values":[569,644,593,580,657,571],"isMain":false},{"metricName":"L50","quality":"Less is better","values":[39,41,57,71,27,39],"isMain":false},{"metricName":"L75","quality":"Less is better","values":[69,83,99,125,66,66],"isMain":false},{"metricName":"GC (%)","quality":"Equal","values":["46.72","46.79","46.76","46.83","46.83","47.32"],"isMain":false}]],["K-mer-based statistics",[]],["Predicted genes",[]],["Similarity statistics",[]],["Reference statistics",[{"metricName":"Reference length","quality":"Equal","values":[3033106,3033106,3033106,3033106,3033106,3033106],"isMain":false},{"metricName":"Reference fragments","quality":"Equal","values":[2,2,2,2,2,2],"isMain":false},{"metricName":"Reference GC (%)","quality":"Equal","values":["46.68","46.68","46.68","46.68","46.68","46.68"],"isMain":false}]]],"subreferences":[],"subreports":[],"minContig":500},{"date":"10 February 2020, Monday, 18:00:02","assembliesNames":["megahit-103-df","megahit-113-df","megahit-113-ml","megahit-113-ms","megahit-129-df","metaSPAdes"],"referenceName":"GCF\_000091345.1\_ASM9134v1","order":[0,1,2,3,4,5],"report":[["Genome statistics",[{"metricName":"Genome fraction (%)","quality":"More is better","values":["0.205","0.014","0.155","0.722","0.291","0.051"],"isMain":true},{"metricName":"Duplication ratio","quality":"Less is better","values":["1.000","2.908","1.409","1.055","1.089","1.001"],"isMain":true},{"metricName":"Largest alignment","quality":"More is better","values":[2516,113,558,2308,1320,807],"isMain":true},{"metricName":"Total aligned length","quality":"More is better","values":[3237,217,2444,11520,4595,807],"isMain":true},{"metricName":"NA50","quality":"More is better","values":[2516,null,330,1031,590,807],"isMain":false},{"metricName":"NA75","quality":"More is better","values":[721,null,null,611,534,807],"isMain":false},{"metricName":"NGA50","quality":"More is better","values":[null,null,null,null,null,null],"isMain":true},{"metricName":"LA50","quality":"Less is better","values":[1,null,5,4,3,1],"isMain":false},{"metricName":"LA75","quality":"Less is better","values":[2,null,null,8,5,1],"isMain":false}]],["Reads mapping",[]],["Misassemblies",[{"metricName":"# misassemblies","quality":"Less is better","values":[0,0,1,1,0,0],"isMain":true},{"metricName":" # relocations","quality":"Less is better","values":[0,0,1,0,0,0],"isMain":false},{"metricName":" # translocations","quality":"Less is better","values":[0,0,0,0,0,0],"isMain":false},{"metricName":" # inversions","quality":"Less is better","values":[0,0,0,1,0,0],"isMain":false},{"metricName":" # interspecies translocations","quality":"Less is better","values":[1,1,4,2,1,0],"isMain":false},{"metricName":"# misassembled contigs","quality":"Less is better","values":[0,0,1,1,0,0],"isMain":false},{"metricName":"Misassembled contigs length","quality":"Less is better","values":[0,0,939,1508,0,0],"isMain":true},{"metricName":"# possibly misassembled contigs","quality":"Less is better","values":[0,0,0,0,0,0],"isMain":false},{"metricName":" # possible misassemblies","quality":"Less is better","values":[0,0,0,0,0,0],"isMain":false},{"metricName":"# local misassemblies","quality":"Less is better","values":[0,0,0,0,0,0],"isMain":false},{"metricName":"# scaffold gap ext. mis.","quality":"Less is better","values":[0,0,0,0,0,0],"isMain":false},{"metricName":"# scaffold gap loc. mis.","quality":"Less is better","values":[0,0,0,0,0,0],"isMain":false},{"metricName":"# unaligned mis. contigs","quality":"Less is better","values":[0,0,0,0,0,0],"isMain":false}]],["Unaligned",[{"metricName":"# fully unaligned contigs","quality":"Less is better","values":[0,0,0,0,0,0],"isMain":false},{"metricName":"Fully unaligned length","quality":"Less is better","values":[0,0,0,0,0,0],"isMain":false},{"metricName":"# partially unaligned contigs","quality":"Less is better","values":[1,1,1,0,0,0],"isMain":false},{"metricName":"Partially unaligned length","quality":"Less is better","values":[503,1022,826,0,0,0],"isMain":false}]],["Mismatches",[{"metricName":"# mismatches","quality":"Less is better","values":[103,6,79,472,163,20],"isMain":false},{"metricName":"# indels","quality":"Less is better","values":[0,0,3,8,0,5],"isMain":false},{"metricName":"Indels length","quality":"Less is better","values":[0,0,4,14,0,5],"isMain":false},{"metricName":"# mismatches per 100 kbp","quality":"Less is better","values":["3181.96","2764.98","3235.05","4143.99","3547.33","2481.39"],"isMain":true},{"metricName":"# indels per 100 kbp","quality":"Less is better","values":["0.00","0.00","122.85","70.24","0.00","620.35"],"isMain":true},{"metricName":" # indels (<= 5 bp)","quality":"Less is better","values":[0,0,3,8,0,5],"isMain":false},{"metricName":" # indels (> 5 bp)","quality":"Less is better","values":[0,0,0,0,0,0],"isMain":false},{"metricName":"# N's","quality":"Less is better","values":[0,0,0,0,0,0],"isMain":false},{"metricName":"# N's per 100 kbp","quality":"Less is better","values":["0.00","0.00","0.00","0.00","0.00","0.00"],"isMain":true}]],["Statistics without reference",[{"metricName":"# contigs","quality":"Equal","values":[2,2,6,12,7,1],"isMain":true},{"metricName":"# contigs (>= 1000 bp)","quality":"Equal","values":[2,1,1,4,1,0],"isMain":false},{"metricName":"# contigs (>= 5000 bp)","quality":"Equal","values":[0,0,0,0,0,0],"isMain":false},{"metricName":"# contigs (>= 10000 bp)","quality":"Equal","values":[0,0,0,0,0,0],"isMain":false},{"metricName":"# contigs (>= 25000 bp)","quality":"Equal","values":[0,0,0,0,0,0],"isMain":false},{"metricName":"# contigs (>= 50000 bp)","quality":"Equal","values":[0,0,0,0,0,0],"isMain":false},{"metricName":"Largest contig","quality":"More is better","values":[2516,1126,1156,2308,1324,807],"isMain":true},{"metricName":"Total length","quality":"More is better","values":[3740,1653,4266,12019,5002,807],"isMain":true},{"metricName":"Total length (>= 1000 bp)","quality":"More is better","values":[3740,1126,1156,6441,1324,0],"isMain":true},{"metricName":"Total length (>= 5000 bp)","quality":"More is better","values":[0,0,0,0,0,0],"isMain":false},{"metricName":"Total length (>= 10000 bp)","quality":"More is better","values":[0,0,0,0,0,0],"isMain":true},{"metricName":"Total length (>= 25000 bp)","quality":"More is better","values":[0,0,0,0,0,0],"isMain":false},{"metricName":"Total length (>= 50000 bp)","quality":"More is better","values":[0,0,0,0,0,0],"isMain":true},{"metricName":"N50","quality":"More is better","values":[2516,1126,576,1091,650,807],"isMain":false},{"metricName":"N75","quality":"More is better","values":[1224,527,562,762,553,807],"isMain":false},{"metricName":"L50","quality":"Less is better","values":[1,1,3,4,3,1],"isMain":false},{"metricName":"L75","quality":"Less is better","values":[2,2,4,8,5,1],"isMain":false},{"metricName":"GC (%)","quality":"Equal","values":["38.16","33.76","38.05","40.47","38.86","40.52"],"isMain":false}]],["K-mer-based statistics",[]],["Predicted genes",[]],["Similarity statistics",[]],["Reference statistics",[{"metricName":"Reference length","quality":"Equal","values":[1576758,1576758,1576758,1576758,1576758,1576758],"isMain":false},{"metricName":"Reference fragments","quality":"Equal","values":[1,1,1,1,1,1],"isMain":false},{"metricName":"Reference GC (%)","quality":"Equal","values":["39.16","39.16","39.16","39.16","39.16","39.16"],"isMain":false}]]],"subreferences":[],"subreports":[],"minContig":500},{"date":"10 February 2020, Monday, 18:00:29","assembliesNames":["megahit-103-df","megahit-113-df","megahit-113-ml","megahit-113-ms","megahit-129-df","metaSPAdes"],"referenceName":"GCF\_000091405.1\_ASM9140v1","order":[0,1,2,3,4,5],"report":[["Genome statistics",[{"metricName":"Genome fraction (%)","quality":"More is better","values":["0.249","0.414","0.605","0.517","0.972","0.298"],"isMain":true},{"metricName":"Duplication ratio","quality":"Less is better","values":["1.187","1.191","1.173","1.157","1.018","1.076"],"isMain":true},{"metricName":"Largest alignment","quality":"More is better","values":[1458,2785,1793,1629,7047,1720],"isMain":true},{"metricName":"Total aligned length","quality":"More is better","values":[4438,7397,10804,9364,17366,5325],"isMain":true},{"metricName":"NA50","quality":"More is better","values":[810,1123,623,null,5031,517],"isMain":false},{"metricName":"NA75","quality":"More is better","values":[190,225,null,null,940,null],"isMain":false},{"metricName":"NGA50","quality":"More is better","values":[null,null,null,null,null,null],"isMain":true},{"metricName":"LA50","quality":"Less is better","values":[3,3,10,null,2,6],"isMain":false},{"metricName":"LA75","quality":"Less is better","values":[7,8,null,null,4,null],"isMain":false}]],["Reads mapping",[]],["Misassemblies",[{"metricName":"# misassemblies","quality":"Less is better","values":[0,2,0,0,2,0],"isMain":true},{"metricName":" # relocations","quality":"Less is better","values":[0,2,0,0,1,0],"isMain":false},{"metricName":" # translocations","quality":"Less is better","values":[0,0,0,0,0,0],"isMain":false},{"metricName":" # inversions","quality":"Less is better","values":[0,0,0,0,1,0],"isMain":false},{"metricName":" # interspecies translocations","quality":"Less is better","values":[3,5,5,6,3,2],"isMain":false},{"metricName":"# misassembled contigs","quality":"Less is better","values":[0,1,0,0,1,0],"isMain":false},{"metricName":"Misassembled contigs length","quality":"Less is better","values":[0,4863,0,0,9490,0],"isMain":true},{"metricName":"# possibly misassembled contigs","quality":"Less is better","values":[0,0,0,0,0,0],"isMain":false},{"metricName":" # possible misassemblies","quality":"Less is better","values":[0,0,0,0,0,0],"isMain":false},{"metricName":"# local misassemblies","quality":"Less is better","values":[0,0,0,0,0,0],"isMain":false},{"metricName":"# scaffold gap ext. mis.","quality":"Less is better","values":[0,0,0,0,0,0],"isMain":false},{"metricName":"# scaffold gap loc. mis.","quality":"Less is better","values":[0,0,0,0,0,0],"isMain":false},{"metricName":"# unaligned mis. contigs","quality":"Less is better","values":[0,0,1,0,0,1],"isMain":false}]],["Unaligned",[{"metricName":"# fully unaligned contigs","quality":"Less is better","values":[0,0,0,2,0,0],"isMain":false},{"metricName":"Fully unaligned length","quality":"Less is better","values":[0,0,0,1588,0,0],"isMain":false},{"metricName":"# partially unaligned contigs","quality":"Less is better","values":[1,1,2,4,2,1],"isMain":false},{"metricName":"Partially unaligned length","quality":"Less is better","values":[555,540,3333,39521,1088,3812],"isMain":false}]],["Mismatches",[{"metricName":"# mismatches","quality":"Less is better","values":[45,157,303,258,452,61],"isMain":false},{"metricName":"# indels","quality":"Less is better","values":[0,1,8,5,9,4],"isMain":false},{"metricName":"Indels length","quality":"Less is better","values":[0,1,8,5,11,4],"isMain":false},{"metricName":"# mismatches per 100 kbp","quality":"Less is better","values":["1013.97","2122.77","2805.04","2797.96","2604.89","1145.11"],"isMain":true},{"metricName":"# indels per 100 kbp","quality":"Less is better","values":["0.00","13.52","74.06","54.22","51.87","75.09"],"isMain":true},{"metricName":" # indels (<= 5 bp)","quality":"Less is better","values":[0,1,8,5,9,4],"isMain":false},{"metricName":" # indels (> 5 bp)","quality":"Less is better","values":[0,0,0,0,0,0],"isMain":false},{"metricName":"# N's","quality":"Less is better","values":[0,0,0,0,0,0],"isMain":false},{"metricName":"# N's per 100 kbp","quality":"Less is better","values":["0.00","0.00","0.00","0.00","0.00","0.00"],"isMain":true}]],["Statistics without reference",[{"metricName":"# contigs","quality":"Equal","values":[7,7,18,18,8,6],"isMain":true},{"metricName":"# contigs (>= 1000 bp)","quality":"Equal","values":[1,2,3,5,2,3],"isMain":false},{"metricName":"# contigs (>= 5000 bp)","quality":"Equal","values":[0,0,0,2,2,0],"isMain":false},{"metricName":"# contigs (>= 10000 bp)","quality":"Equal","values":[0,0,0,1,0,0],"isMain":false},{"metricName":"# contigs (>= 25000 bp)","quality":"Equal","values":[0,0,0,1,0,0],"isMain":false},{"metricName":"# contigs (>= 50000 bp)","quality":"Equal","values":[0,0,0,0,0,0],"isMain":false},{"metricName":"Largest contig","quality":"More is better","values":[1458,4863,2554,33159,9490,4980],"isMain":true},{"metricName":"Total length","quality":"More is better","values":[5821,9350,15999,51779,18753,9542],"isMain":true},{"metricName":"Total length (>= 1000 bp)","quality":"More is better","values":[1458,6005,5923,43440,14532,7719],"isMain":true},{"metricName":"Total length (>= 5000 bp)","quality":"More is better","values":[0,0,0,39618,14532,0],"isMain":false},{"metricName":"Total length (>= 10000 bp)","quality":"More is better","values":[0,0,0,33159,0,0],"isMain":true},{"metricName":"Total length (>= 25000 bp)","quality":"More is better","values":[0,0,0,33159,0,0],"isMain":false},{"metricName":"Total length (>= 50000 bp)","quality":"More is better","values":[0,0,0,0,0,0],"isMain":true},{"metricName":"N50","quality":"More is better","values":[810,4863,746,33159,9490,4980],"isMain":false},{"metricName":"N75","quality":"More is better","values":[745,730,662,6459,5042,1012],"isMain":false},{"metricName":"L50","quality":"Less is better","values":[3,1,6,1,1,1],"isMain":false},{"metricName":"L75","quality":"Less is better","values":[5,4,12,2,2,3],"isMain":false},{"metricName":"GC (%)","quality":"Equal","values":["37.42","35.48","37.21","44.83","34.45","35.07"],"isMain":false}]],["K-mer-based statistics",[]],["Predicted genes",[]],["Similarity statistics",[]],["Reference statistics",[{"metricName":"Reference length","quality":"Equal","values":[1785116,1785116,1785116,1785116,1785116,1785116],"isMain":false},{"metricName":"Reference fragments","quality":"Equal","values":[3,3,3,3,3,3],"isMain":false},{"metricName":"Reference GC (%)","quality":"Equal","values":["34.43","34.43","34.43","34.43","34.43","34.43"],"isMain":false}]]],"subreferences":[],"subreports":[],"minContig":500},{"date":"10 February 2020, Monday, 18:00:51","assembliesNames":["megahit-103-df","megahit-113-df","megahit-113-ml","megahit-113-ms","megahit-129-df","metaSPAdes"],"referenceName":"GCF\_000142965.1\_ASM14296v1","order":[0,1,2,3,4,5],"report":[["Genome statistics",[{"metricName":"Genome fraction (%)","quality":"More is better","values":["89.271","88.411","90.441","89.066","91.401","85.770"],"isMain":true},{"metricName":"Duplication ratio","quality":"Less is better","values":["1.025","1.035","1.072","1.064","1.036","1.010"],"isMain":true},{"metricName":"Largest alignment","quality":"More is better","values":[41681,56402,23360,24066,57157,52106],"isMain":true},{"metricName":"Total aligned length","quality":"More is better","values":[3031410,3025575,3202095,3126851,3129332,2872308],"isMain":true},{"metricName":"NG50","quality":"More is better","values":[8373,10126,5425,6070,11418,13077],"isMain":false},{"metricName":"NG75","quality":"More is better","values":[3327,3727,2923,3326,5256,5365],"isMain":false},{"metricName":"NA50","quality":"More is better","values":[9198,9917,4471,4814,10776,13890],"isMain":false},{"metricName":"NA75","quality":"More is better","values":[4233,4172,2074,2224,4806,6391],"isMain":false},{"metricName":"NGA50","quality":"More is better","values":[8097,9170,4601,4964,10517,11368],"isMain":true},{"metricName":"NGA75","quality":"More is better","values":[3115,3064,2282,2412,4307,4052],"isMain":false},{"metricName":"LG50","quality":"Less is better","values":[114,89,179,148,83,68],"isMain":false},{"metricName":"LG75","quality":"Less is better","values":[268,223,389,334,189,168],"isMain":false},{"metricName":"LA50","quality":"Less is better","values":[100,87,233,199,85,64],"isMain":false},{"metricName":"LA75","quality":"Less is better","values":[218,207,510,452,195,145],"isMain":false},{"metricName":"LGA50","quality":"Less is better","values":[116,98,222,192,89,78],"isMain":true},{"metricName":"LGA75","quality":"Less is better","values":[275,250,475,429,211,194],"isMain":false}]],["Reads mapping",[]],["Misassemblies",[{"metricName":"# misassemblies","quality":"Less is better","values":[17,37,30,39,41,11],"isMain":true},{"metricName":" # relocations","quality":"Less is better","values":[0,3,0,1,0,0],"isMain":false},{"metricName":" # translocations","quality":"Less is better","values":[17,34,27,36,38,11],"isMain":false},{"metricName":" # inversions","quality":"Less is better","values":[0,0,3,2,3,0],"isMain":false},{"metricName":" # interspecies translocations","quality":"Less is better","values":[10,38,44,58,39,14],"isMain":false},{"metricName":"# misassembled contigs","quality":"Less is better","values":[16,33,24,32,36,9],"isMain":false},{"metricName":"Misassembled contigs length","quality":"Less is better","values":[106338,249908,85750,170114,258787,174299],"isMain":true},{"metricName":"# possibly misassembled contigs","quality":"Less is better","values":[0,2,0,0,0,0],"isMain":false},{"metricName":" # possible misassemblies","quality":"Less is better","values":[0,2,0,0,0,0],"isMain":false},{"metricName":"# local misassemblies","quality":"Less is better","values":[1,2,2,2,3,1],"isMain":false},{"metricName":"# scaffold gap ext. mis.","quality":"Less is better","values":[0,0,0,0,0,0],"isMain":false},{"metricName":"# scaffold gap loc. mis.","quality":"Less is better","values":[0,0,0,0,0,0],"isMain":false},{"metricName":"# unaligned mis. contigs","quality":"Less is better","values":[1,1,3,0,1,0],"isMain":false}]],["Unaligned",[{"metricName":"# fully unaligned contigs","quality":"Less is better","values":[0,1,5,5,4,0],"isMain":false},{"metricName":"Fully unaligned length","quality":"Less is better","values":[0,687,7175,2947,21667,0],"isMain":false},{"metricName":"# partially unaligned contigs","quality":"Less is better","values":[5,14,13,20,11,8],"isMain":false},{"metricName":"Partially unaligned length","quality":"Less is better","values":[5515,71874,195594,240413,60705,99383],"isMain":false}]],["Mismatches",[{"metricName":"# mismatches","quality":"Less is better","values":[6665,11906,21339,20617,10440,3824],"isMain":false},{"metricName":"# indels","quality":"Less is better","values":[59,135,157,171,108,46],"isMain":false},{"metricName":"Indels length","quality":"Less is better","values":[337,380,461,472,383,149],"isMain":false},{"metricName":"# mismatches per 100 kbp","quality":"Less is better","values":["224.75","405.39","710.27","696.84","343.85","134.22"],"isMain":true},{"metricName":"# indels per 100 kbp","quality":"Less is better","values":["1.99","4.60","5.23","5.78","3.56","1.61"],"isMain":true},{"metricName":" # indels (<= 5 bp)","quality":"Less is better","values":[53,127,146,161,102,42],"isMain":false},{"metricName":" # indels (> 5 bp)","quality":"Less is better","values":[6,8,11,10,6,4],"isMain":false},{"metricName":"# N's","quality":"Less is better","values":[0,0,0,0,0,0],"isMain":false},{"metricName":"# N's per 100 kbp","quality":"Less is better","values":["0.00","0.00","0.00","0.00","0.00","0.00"],"isMain":true}]],["Statistics without reference",[{"metricName":"# contigs","quality":"Equal","values":[680,648,1093,957,645,414],"isMain":true},{"metricName":"# contigs (>= 1000 bp)","quality":"Equal","values":[499,459,783,718,447,338],"isMain":false},{"metricName":"# contigs (>= 5000 bp)","quality":"Equal","values":[197,178,196,200,193,177],"isMain":false},{"metricName":"# contigs (>= 10000 bp)","quality":"Equal","values":[83,89,41,46,98,95],"isMain":false},{"metricName":"# contigs (>= 25000 bp)","quality":"Equal","values":[10,15,3,3,17,25],"isMain":false},{"metricName":"# contigs (>= 50000 bp)","quality":"Equal","values":[0,2,1,1,2,2],"isMain":false},{"metricName":"Largest contig","quality":"More is better","values":[41681,62299,113298,183133,58272,89862],"isMain":true},{"metricName":"Total length","quality":"More is better","values":[3043951,3112492,3422765,3391801,3227594,2977317],"isMain":true},{"metricName":"Total length (>= 1000 bp)","quality":"More is better","values":[2921397,2979384,3202515,3224875,3090374,2923641],"isMain":true},{"metricName":"Total length (>= 5000 bp)","quality":"More is better","values":[2209186,2298652,1750414,1949981,2512641,2539730],"isMain":false},{"metricName":"Total length (>= 10000 bp)","quality":"More is better","values":[1379633,1664890,706972,863108,1829453,1968149],"isMain":true},{"metricName":"Total length (>= 25000 bp)","quality":"More is better","values":[297080,538979,205733,245203,593934,904377],"isMain":false},{"metricName":"Total length (>= 50000 bp)","quality":"More is better","values":[0,118709,113298,183133,108893,141971],"isMain":true},{"metricName":"N50","quality":"More is better","values":[9222,10929,5201,5892,12067,14818],"isMain":false},{"metricName":"N75","quality":"More is better","values":[4454,4709,2739,3110,5581,7381],"isMain":false},{"metricName":"L50","quality":"Less is better","values":[98,79,189,154,79,56],"isMain":false},{"metricName":"L75","quality":"Less is better","values":[213,186,415,350,176,127],"isMain":false},{"metricName":"GC (%)","quality":"Equal","values":["43.85","43.92","44.24","44.49","43.89","43.88"],"isMain":false}]],["K-mer-based statistics",[]],["Predicted genes",[]],["Similarity statistics",[]],["Reference statistics",[{"metricName":"Reference length","quality":"Equal","values":[3321873,3321873,3321873,3321873,3321873,3321873],"isMain":false},{"metricName":"Reference fragments","quality":"Equal","values":[132,132,132,132,132,132],"isMain":false},{"metricName":"Reference GC (%)","quality":"Equal","values":["43.91","43.91","43.91","43.91","43.91","43.91"],"isMain":false}]]],"subreferences":[],"subreports":[],"minContig":500},{"date":"10 February 2020, Monday, 18:00:41","assembliesNames":["megahit-103-df","megahit-113-df","megahit-113-ml","megahit-113-ms","megahit-129-df","metaSPAdes"],"referenceName":"GCF\_000144405.1\_ASM14440v1","order":[0,1,2,3,4,5],"report":[["Genome statistics",[{"metricName":"Genome fraction (%)","quality":"More is better","values":["96.485","95.638","96.905","96.594","97.131","94.364"],"isMain":true},{"metricName":"Duplication ratio","quality":"Less is better","values":["1.034","1.060","1.068","1.068","1.055","1.013"],"isMain":true},{"metricName":"Largest alignment","quality":"More is better","values":[33368,38241,37918,23875,51286,36490],"isMain":true},{"metricName":"Total aligned length","quality":"More is better","values":[3154377,3196825,3268344,3248098,3233559,3025790],"isMain":true},{"metricName":"NG50","quality":"More is better","values":[8932,9199,5977,6839,11372,8309],"isMain":false},{"metricName":"NG75","quality":"More is better","values":[3952,4533,3350,3738,5962,4096],"isMain":false},{"metricName":"NA50","quality":"More is better","values":[8773,8102,5552,6149,10188,8573],"isMain":false},{"metricName":"NA75","quality":"More is better","values":[3903,3785,2803,3248,4640,4319],"isMain":false},{"metricName":"NGA50","quality":"More is better","values":[8773,8488,5826,6545,10959,8090],"isMain":true},{"metricName":"NGA75","quality":"More is better","values":[3903,4278,3169,3536,5263,3945],"isMain":false},{"metricName":"LG50","quality":"Less is better","values":[108,101,150,144,85,107],"isMain":false},{"metricName":"LG75","quality":"Less is better","values":[242,223,327,300,180,242],"isMain":false},{"metricName":"LA50","quality":"Less is better","values":[109,115,171,160,98,102],"isMain":false},{"metricName":"LA75","quality":"Less is better","values":[244,265,384,347,219,226],"isMain":false},{"metricName":"LGA50","quality":"Less is better","values":[109,107,156,148,90,108],"isMain":true},{"metricName":"LGA75","quality":"Less is better","values":[244,240,341,313,194,247],"isMain":false}]],["Reads mapping",[]],["Misassemblies",[{"metricName":"# misassemblies","quality":"Less is better","values":[2,1,2,2,6,1],"isMain":true},{"metricName":" # relocations","quality":"Less is better","values":[1,1,1,2,4,0],"isMain":false},{"metricName":" # translocations","quality":"Less is better","values":[0,0,1,0,2,1],"isMain":false},{"metricName":" # inversions","quality":"Less is better","values":[1,0,0,0,0,0],"isMain":false},{"metricName":" # interspecies translocations","quality":"Less is better","values":[10,54,29,54,48,13],"isMain":false},{"metricName":"# misassembled contigs","quality":"Less is better","values":[2,1,2,2,6,1],"isMain":false},{"metricName":"Misassembled contigs length","quality":"Less is better","values":[11641,1002,1254,12886,66874,11308],"isMain":true},{"metricName":"# possibly misassembled contigs","quality":"Less is better","values":[1,7,0,4,2,0],"isMain":false},{"metricName":" # possible misassemblies","quality":"Less is better","values":[1,7,0,4,2,0],"isMain":false},{"metricName":"# local misassemblies","quality":"Less is better","values":[3,4,6,6,7,0],"isMain":false},{"metricName":"# scaffold gap ext. mis.","quality":"Less is better","values":[0,0,0,0,0,0],"isMain":false},{"metricName":"# scaffold gap loc. mis.","quality":"Less is better","values":[0,0,0,0,0,0],"isMain":false},{"metricName":"# unaligned mis. contigs","quality":"Less is better","values":[0,1,1,1,0,0],"isMain":false}]],["Unaligned",[{"metricName":"# fully unaligned contigs","quality":"Less is better","values":[0,2,4,3,0,0],"isMain":false},{"metricName":"Fully unaligned length","quality":"Less is better","values":[0,2267,4578,2279,0,0],"isMain":false},{"metricName":"# partially unaligned contigs","quality":"Less is better","values":[8,36,14,23,31,10],"isMain":false},{"metricName":"Partially unaligned length","quality":"Less is better","values":[7598,86823,52055,49567,89262,25474],"isMain":false}]],["Mismatches",[{"metricName":"# mismatches","quality":"Less is better","values":[7513,16948,19428,20131,12460,3473],"isMain":false},{"metricName":"# indels","quality":"Less is better","values":[79,255,106,147,156,57],"isMain":false},{"metricName":"Indels length","quality":"Less is better","values":[566,754,556,572,702,260],"isMain":false},{"metricName":"# mismatches per 100 kbp","quality":"Less is better","values":["245.77","559.32","632.79","657.80","404.89","116.17"],"isMain":true},{"metricName":"# indels per 100 kbp","quality":"Less is better","values":["2.58","8.42","3.45","4.80","5.07","1.91"],"isMain":true},{"metricName":" # indels (<= 5 bp)","quality":"Less is better","values":[71,236,99,137,142,51],"isMain":false},{"metricName":" # indels (> 5 bp)","quality":"Less is better","values":[8,19,7,10,14,6],"isMain":false},{"metricName":"# N's","quality":"Less is better","values":[0,0,0,0,0,0],"isMain":false},{"metricName":"# N's per 100 kbp","quality":"Less is better","values":["0.00","0.00","0.00","0.00","0.00","0.00"],"isMain":true}]],["Statistics without reference",[{"metricName":"# contigs","quality":"Equal","values":[722,718,944,875,644,595],"isMain":true},{"metricName":"# contigs (>= 1000 bp)","quality":"Equal","values":[556,558,740,679,489,503],"isMain":false},{"metricName":"# contigs (>= 5000 bp)","quality":"Equal","values":[195,207,201,221,211,197],"isMain":false},{"metricName":"# contigs (>= 10000 bp)","quality":"Equal","values":[89,87,50,69,102,84],"isMain":false},{"metricName":"# contigs (>= 25000 bp)","quality":"Equal","values":[6,10,5,0,16,9],"isMain":false},{"metricName":"# contigs (>= 50000 bp)","quality":"Equal","values":[0,0,1,0,1,0],"isMain":false},{"metricName":"Largest contig","quality":"More is better","values":[33368,40885,59338,23875,51286,36490],"isMain":true},{"metricName":"Total length","quality":"More is better","values":[3167005,3301800,3337024,3319526,3335354,3054241],"isMain":true},{"metricName":"Total length (>= 1000 bp)","quality":"More is better","values":[3049882,3189264,3194926,3183567,3226991,2987396],"isMain":true},{"metricName":"Total length (>= 5000 bp)","quality":"More is better","values":[2169244,2301834,1868518,2037454,2543549,2178291],"isMain":false},{"metricName":"Total length (>= 10000 bp)","quality":"More is better","values":[1406608,1455113,823697,969146,1770049,1378972],"isMain":true},{"metricName":"Total length (>= 25000 bp)","quality":"More is better","values":[180262,312089,181298,0,502730,271922],"isMain":false},{"metricName":"Total length (>= 50000 bp)","quality":"More is better","values":[0,0,59338,0,51286,0],"isMain":true},{"metricName":"N50","quality":"More is better","values":[8932,8554,5641,6545,10892,8756],"isMain":false},{"metricName":"N75","quality":"More is better","values":[3952,4224,2998,3375,5214,4428],"isMain":false},{"metricName":"L50","quality":"Less is better","values":[108,109,164,155,93,100],"isMain":false},{"metricName":"L75","quality":"Less is better","values":[242,246,367,333,203,221],"isMain":false},{"metricName":"GC (%)","quality":"Equal","values":["40.92","40.97","40.97","40.98","41.01","40.85"],"isMain":false}]],["K-mer-based statistics",[]],["Predicted genes",[]],["Similarity statistics",[]],["Reference statistics",[{"metricName":"Reference length","quality":"Equal","values":[3168282,3168282,3168282,3168282,3168282,3168282],"isMain":false},{"metricName":"Reference fragments","quality":"Equal","values":[2,2,2,2,2,2],"isMain":false},{"metricName":"Reference GC (%)","quality":"Equal","values":["40.98","40.98","40.98","40.98","40.98","40.98"],"isMain":false}]]],"subreferences":[],"subreports":[],"minContig":500},{"date":"10 February 2020, Monday, 17:34:56","assembliesNames":["megahit-113-ml","megahit-113-ms","megahit-129-df"],"referenceName":"GCF\_000144625.1\_ASM14462v1","order":[0,1,2],"report":[["Genome statistics",[{"metricName":"Genome fraction (%)","quality":"More is better","values":["0.010","0.011","0.008"],"isMain":true},{"metricName":"Duplication ratio","quality":"Less is better","values":["1.337","1.049","1.184"],"isMain":true},{"metricName":"Largest alignment","quality":"More is better","values":[459,509,206],"isMain":true},{"metricName":"Total aligned length","quality":"More is better","values":[459,509,431],"isMain":true},{"metricName":"NA50","quality":"More is better","values":[459,509,null],"isMain":false},{"metricName":"NA75","quality":"More is better","values":[null,509,null],"isMain":false},{"metricName":"NGA50","quality":"More is better","values":[null,null,null],"isMain":true},{"metricName":"LA50","quality":"Less is better","values":[1,1,null],"isMain":false},{"metricName":"LA75","quality":"Less is better","values":[null,1,null],"isMain":false}]],["Reads mapping",[]],["Misassemblies",[{"metricName":"# misassemblies","quality":"Less is better","values":[0,0,0],"isMain":true},{"metricName":" # relocations","quality":"Less is better","values":[0,0,0],"isMain":false},{"metricName":" # translocations","quality":"Less is better","values":[0,0,0],"isMain":false},{"metricName":" # inversions","quality":"Less is better","values":[0,0,0],"isMain":false},{"metricName":" # interspecies translocations","quality":"Less is better","values":[0,0,2],"isMain":false},{"metricName":"# misassembled contigs","quality":"Less is better","values":[0,0,0],"isMain":false},{"metricName":"Misassembled contigs length","quality":"Less is better","values":[0,0,0],"isMain":true},{"metricName":"# possibly misassembled contigs","quality":"Less is better","values":[0,0,0],"isMain":false},{"metricName":" # possible misassemblies","quality":"Less is better","values":[0,0,0],"isMain":false},{"metricName":"# local misassemblies","quality":"Less is better","values":[0,0,0],"isMain":false},{"metricName":"# scaffold gap ext. mis.","quality":"Less is better","values":[0,0,0],"isMain":false},{"metricName":"# scaffold gap loc. mis.","quality":"Less is better","values":[0,0,0],"isMain":false},{"metricName":"# unaligned mis. contigs","quality":"Less is better","values":[0,0,1],"isMain":false}]],["Unaligned",[{"metricName":"# fully unaligned contigs","quality":"Less is better","values":[0,0,0],"isMain":false},{"metricName":"Fully unaligned length","quality":"Less is better","values":[0,0,0],"isMain":false},{"metricName":"# partially unaligned contigs","quality":"Less is better","values":[0,0,2],"isMain":false},{"metricName":"Partially unaligned length","quality":"Less is better","values":[0,0,1712],"isMain":false}]],["Mismatches",[{"metricName":"# mismatches","quality":"Less is better","values":[21,24,16],"isMain":false},{"metricName":"# indels","quality":"Less is better","values":[1,0,0],"isMain":false},{"metricName":"Indels length","quality":"Less is better","values":[1,0,0],"isMain":false},{"metricName":"# mismatches per 100 kbp","quality":"Less is better","values":["4565.22","4715.13","4395.60"],"isMain":true},{"metricName":"# indels per 100 kbp","quality":"Less is better","values":["217.39","0.00","0.00"],"isMain":true},{"metricName":" # indels (<= 5 bp)","quality":"Less is better","values":[1,0,0],"isMain":false},{"metricName":" # indels (> 5 bp)","quality":"Less is better","values":[0,0,0],"isMain":false},{"metricName":"# N's","quality":"Less is better","values":[0,0,0],"isMain":false},{"metricName":"# N's per 100 kbp","quality":"Less is better","values":["0.00","0.00","0.00"],"isMain":true}]],["Statistics without reference",[{"metricName":"# contigs","quality":"Equal","values":[1,1,2],"isMain":true},{"metricName":"# contigs (>= 1000 bp)","quality":"Equal","values":[0,0,2],"isMain":false},{"metricName":"# contigs (>= 5000 bp)","quality":"Equal","values":[0,0,0],"isMain":false},{"metricName":"# contigs (>= 10000 bp)","quality":"Equal","values":[0,0,0],"isMain":false},{"metricName":"# contigs (>= 25000 bp)","quality":"Equal","values":[0,0,0],"isMain":false},{"metricName":"# contigs (>= 50000 bp)","quality":"Equal","values":[0,0,0],"isMain":false},{"metricName":"Largest contig","quality":"More is better","values":[615,534,1119],"isMain":true},{"metricName":"Total length","quality":"More is better","values":[615,534,2143],"isMain":true},{"metricName":"Total length (>= 1000 bp)","quality":"More is better","values":[0,0,2143],"isMain":true},{"metricName":"Total length (>= 5000 bp)","quality":"More is better","values":[0,0,0],"isMain":false},{"metricName":"Total length (>= 10000 bp)","quality":"More is better","values":[0,0,0],"isMain":true},{"metricName":"Total length (>= 25000 bp)","quality":"More is better","values":[0,0,0],"isMain":false},{"metricName":"Total length (>= 50000 bp)","quality":"More is better","values":[0,0,0],"isMain":true},{"metricName":"N50","quality":"More is better","values":[615,534,1119],"isMain":false},{"metricName":"N75","quality":"More is better","values":[615,534,1024],"isMain":false},{"metricName":"L50","quality":"Less is better","values":[1,1,1],"isMain":false},{"metricName":"L75","quality":"Less is better","values":[1,1,2],"isMain":false},{"metricName":"GC (%)","quality":"Equal","values":["52.36","48.31","47.32"],"isMain":false}]],["K-mer-based statistics",[]],["Predicted genes",[]],["Similarity statistics",[]],["Reference statistics",[{"metricName":"Reference length","quality":"Equal","values":[4662871,4662871,4662871],"isMain":false},{"metricName":"Reference fragments","quality":"Equal","values":[1,1,1],"isMain":false},{"metricName":"Reference GC (%)","quality":"Equal","values":["45.00","45.00","45.00"],"isMain":false}]]],"subreferences":[],"subreports":[],"minContig":500},{"date":"10 February 2020, Monday, 18:00:52","assembliesNames":["megahit-103-df","megahit-113-df","megahit-113-ml","megahit-113-ms","megahit-129-df","metaSPAdes"],"referenceName":"GCF\_000145035.1\_ASM14503v1","order":[0,1,2,3,4,5],"report":[["Genome statistics",[{"metricName":"Genome fraction (%)","quality":"More is better","values":["24.185","28.712","27.033","28.883","31.434","16.713"],"isMain":true},{"metricName":"Duplication ratio","quality":"Less is better","values":["1.024","1.022","1.025","1.035","1.027","1.036"],"isMain":true},{"metricName":"Largest alignment","quality":"More is better","values":[2432,3496,1646,2318,2205,1914],"isMain":true},{"metricName":"Total aligned length","quality":"More is better","values":[1074765,1276085,1207637,1293582,1402128,743694],"isMain":true},{"metricName":"NA50","quality":"More is better","values":[670,692,658,668,702,616],"isMain":false},{"metricName":"NA75","quality":"More is better","values":[558,569,559,559,570,538],"isMain":false},{"metricName":"NGA50","quality":"More is better","values":[null,null,null,null,null,null],"isMain":true},{"metricName":"LA50","quality":"Less is better","values":[642,714,730,787,794,504],"isMain":false},{"metricName":"LA75","quality":"Less is better","values":[1093,1245,1235,1340,1381,842],"isMain":false}]],["Reads mapping",[]],["Misassemblies",[{"metricName":"# misassemblies","quality":"Less is better","values":[6,8,6,10,10,5],"isMain":true},{"metricName":" # relocations","quality":"Less is better","values":[4,5,5,7,6,4],"isMain":false},{"metricName":" # translocations","quality":"Less is better","values":[1,1,0,1,2,1],"isMain":false},{"metricName":" # inversions","quality":"Less is better","values":[1,2,1,2,2,0],"isMain":false},{"metricName":" # interspecies translocations","quality":"Less is better","values":[65,55,12,52,67,45],"isMain":false},{"metricName":"# misassembled contigs","quality":"Less is better","values":[6,8,6,10,9,5],"isMain":false},{"metricName":"Misassembled contigs length","quality":"Less is better","values":[6478,8776,5166,7355,8189,3913],"isMain":true},{"metricName":"# possibly misassembled contigs","quality":"Less is better","values":[0,0,0,0,0,0],"isMain":false},{"metricName":" # possible misassemblies","quality":"Less is better","values":[0,0,0,0,0,0],"isMain":false},{"metricName":"# local misassemblies","quality":"Less is better","values":[2,2,0,1,2,1],"isMain":false},{"metricName":"# scaffold gap ext. mis.","quality":"Less is better","values":[0,0,0,0,0,0],"isMain":false},{"metricName":"# scaffold gap loc. mis.","quality":"Less is better","values":[0,0,0,0,0,0],"isMain":false},{"metricName":"# unaligned mis. contigs","quality":"Less is better","values":[0,0,0,0,0,0],"isMain":false}]],["Unaligned",[{"metricName":"# fully unaligned contigs","quality":"Less is better","values":[0,0,0,0,0,0],"isMain":false},{"metricName":"Fully unaligned length","quality":"Less is better","values":[0,0,0,0,0,0],"isMain":false},{"metricName":"# partially unaligned contigs","quality":"Less is better","values":[9,20,2,12,19,12],"isMain":false},{"metricName":"Partially unaligned length","quality":"Less is better","values":[11071,36577,1269,32083,58198,12608],"isMain":false}]],["Mismatches",[{"metricName":"# mismatches","quality":"Less is better","values":[15934,18526,23633,26392,20688,11699],"isMain":false},{"metricName":"# indels","quality":"Less is better","values":[31,54,32,34,60,28],"isMain":false},{"metricName":"Indels length","quality":"Less is better","values":[103,121,83,78,247,133],"isMain":false},{"metricName":"# mismatches per 100 kbp","quality":"Less is better","values":["1495.71","1464.84","1984.71","2074.44","1494.13","1589.13"],"isMain":true},{"metricName":"# indels per 100 kbp","quality":"Less is better","values":["2.91","4.27","2.69","2.67","4.33","3.80"],"isMain":true},{"metricName":" # indels (<= 5 bp)","quality":"Less is better","values":[28,52,30,33,55,25],"isMain":false},{"metricName":" # indels (> 5 bp)","quality":"Less is better","values":[3,2,2,1,5,3],"isMain":false},{"metricName":"# N's","quality":"Less is better","values":[0,0,0,0,0,0],"isMain":false},{"metricName":"# N's per 100 kbp","quality":"Less is better","values":["0.00","0.00","0.00","0.00","0.00","0.00"],"isMain":true}]],["Statistics without reference",[{"metricName":"# contigs","quality":"Equal","values":[1575,1784,1792,1892,1944,1171],"isMain":true},{"metricName":"# contigs (>= 1000 bp)","quality":"Equal","values":[125,208,117,151,229,50],"isMain":false},{"metricName":"# contigs (>= 5000 bp)","quality":"Equal","values":[0,2,0,1,2,0],"isMain":false},{"metricName":"# contigs (>= 10000 bp)","quality":"Equal","values":[0,0,0,1,2,0],"isMain":false},{"metricName":"# contigs (>= 25000 bp)","quality":"Equal","values":[0,0,0,0,1,0],"isMain":false},{"metricName":"# contigs (>= 50000 bp)","quality":"Equal","values":[0,0,0,0,0,0],"isMain":false},{"metricName":"Largest contig","quality":"More is better","values":[3734,9859,1661,21210,30080,2507],"isMain":true},{"metricName":"Total length","quality":"More is better","values":[1101703,1329710,1221982,1349293,1480380,775655],"isMain":true},{"metricName":"Total length (>= 1000 bp)","quality":"More is better","values":[156343,284041,137443,202521,332708,61112],"isMain":true},{"metricName":"Total length (>= 5000 bp)","quality":"More is better","values":[0,16292,0,21210,40879,0],"isMain":false},{"metricName":"Total length (>= 10000 bp)","quality":"More is better","values":[0,0,0,21210,40879,0],"isMain":true},{"metricName":"Total length (>= 25000 bp)","quality":"More is better","values":[0,0,0,0,30080,0],"isMain":false},{"metricName":"Total length (>= 50000 bp)","quality":"More is better","values":[0,0,0,0,0,0],"isMain":true},{"metricName":"N50","quality":"More is better","values":[685,727,668,692,741,643],"isMain":false},{"metricName":"N75","quality":"More is better","values":[576,595,569,585,602,561],"isMain":false},{"metricName":"L50","quality":"Less is better","values":[621,665,721,735,712,480],"isMain":false},{"metricName":"L75","quality":"Less is better","values":[1060,1173,1219,1266,1269,805],"isMain":false},{"metricName":"GC (%)","quality":"Equal","values":["40.12","40.05","40.16","40.19","39.98","40.04"],"isMain":false}]],["K-mer-based statistics",[]],["Predicted genes",[]],["Similarity statistics",[]],["Reference statistics",[{"metricName":"Reference length","quality":"Equal","values":[4404877,4404877,4404877,4404877,4404877,4404877],"isMain":false},{"metricName":"Reference fragments","quality":"Equal","values":[13,13,13,13,13,13],"isMain":false},{"metricName":"Reference GC (%)","quality":"Equal","values":["40.01","40.01","40.01","40.01","40.01","40.01"],"isMain":false}]]],"subreferences":[],"subreports":[],"minContig":500},{"date":"10 February 2020, Monday, 18:00:46","assembliesNames":["megahit-103-df","megahit-113-df","megahit-113-ml","megahit-113-ms","megahit-129-df","metaSPAdes"],"referenceName":"GCF\_000153905.1\_ASM15390v1","order":[0,1,2,3,4,5],"report":[["Genome statistics",[{"metricName":"Genome fraction (%)","quality":"More is better","values":["89.856","90.497","87.210","87.601","91.606","84.563"],"isMain":true},{"metricName":"Duplication ratio","quality":"Less is better","values":["1.055","1.045","1.102","1.106","1.056","1.060"],"isMain":true},{"metricName":"Largest alignment","quality":"More is better","values":[8553,15038,5539,6451,15237,5824],"isMain":true},{"metricName":"Total aligned length","quality":"More is better","values":[3415829,3405552,3459059,3478839,3481194,3223887],"isMain":true},{"metricName":"NG50","quality":"More is better","values":[1818,2475,1210,1211,2462,1359],"isMain":false},{"metricName":"NG75","quality":"More is better","values":[1106,1379,794,813,1420,801],"isMain":false},{"metricName":"NA50","quality":"More is better","values":[1815,2436,1218,1196,2397,1459],"isMain":false},{"metricName":"NA75","quality":"More is better","values":[1129,1366,805,805,1353,935],"isMain":false},{"metricName":"NGA50","quality":"More is better","values":[1765,2347,1181,1178,2381,1331],"isMain":true},{"metricName":"NGA75","quality":"More is better","values":[1051,1279,772,784,1316,777],"isMain":false},{"metricName":"LG50","quality":"Less is better","values":[629,454,959,965,454,842],"isMain":false},{"metricName":"LG75","quality":"Less is better","values":[1261,939,1889,1878,938,1704],"isMain":false},{"metricName":"LA50","quality":"Less is better","values":[620,457,931,960,469,728],"isMain":false},{"metricName":"LA75","quality":"Less is better","values":[1223,930,1821,1862,966,1423],"isMain":false},{"metricName":"LGA50","quality":"Less is better","values":[659,483,982,993,478,859],"isMain":true},{"metricName":"LGA75","quality":"Less is better","values":[1319,1000,1935,1936,991,1744],"isMain":false}]],["Reads mapping",[]],["Misassemblies",[{"metricName":"# misassemblies","quality":"Less is better","values":[5,8,9,7,5,6],"isMain":true},{"metricName":" # relocations","quality":"Less is better","values":[2,2,3,1,0,3],"isMain":false},{"metricName":" # translocations","quality":"Less is better","values":[3,5,4,3,4,3],"isMain":false},{"metricName":" # inversions","quality":"Less is better","values":[0,1,2,3,1,0],"isMain":false},{"metricName":" # interspecies translocations","quality":"Less is better","values":[37,56,21,72,66,24],"isMain":false},{"metricName":"# misassembled contigs","quality":"Less is better","values":[5,8,9,7,5,6],"isMain":false},{"metricName":"Misassembled contigs length","quality":"Less is better","values":[12283,19342,13740,9921,7681,13843],"isMain":true},{"metricName":"# possibly misassembled contigs","quality":"Less is better","values":[0,1,0,0,2,0],"isMain":false},{"metricName":" # possible misassemblies","quality":"Less is better","values":[0,1,0,0,2,0],"isMain":false},{"metricName":"# local misassemblies","quality":"Less is better","values":[3,5,5,3,5,1],"isMain":false},{"metricName":"# scaffold gap ext. mis.","quality":"Less is better","values":[0,0,0,0,0,0],"isMain":false},{"metricName":"# scaffold gap loc. mis.","quality":"Less is better","values":[0,0,0,0,0,0],"isMain":false},{"metricName":"# unaligned mis. contigs","quality":"Less is better","values":[0,0,1,0,0,0],"isMain":false}]],["Unaligned",[{"metricName":"# fully unaligned contigs","quality":"Less is better","values":[0,0,0,1,2,0],"isMain":false},{"metricName":"Fully unaligned length","quality":"Less is better","values":[0,0,0,1253,4172,0],"isMain":false},{"metricName":"# partially unaligned contigs","quality":"Less is better","values":[7,17,5,14,22,5],"isMain":false},{"metricName":"Partially unaligned length","quality":"Less is better","values":[50922,72933,20309,31700,68852,11143],"isMain":false}]],["Mismatches",[{"metricName":"# mismatches","quality":"Less is better","values":[24669,22907,49402,51116,24719,25532],"isMain":false},{"metricName":"# indels","quality":"Less is better","values":[50,55,54,87,69,49],"isMain":false},{"metricName":"Indels length","quality":"Less is better","values":[104,112,81,154,165,95],"isMain":false},{"metricName":"# mismatches per 100 kbp","quality":"Less is better","values":["757.41","698.33","1562.80","1609.81","744.45","832.98"],"isMain":true},{"metricName":"# indels per 100 kbp","quality":"Less is better","values":["1.54","1.68","1.71","2.74","2.08","1.60"],"isMain":true},{"metricName":" # indels (<= 5 bp)","quality":"Less is better","values":[47,52,53,82,63,45],"isMain":false},{"metricName":" # indels (> 5 bp)","quality":"Less is better","values":[3,3,1,5,6,4],"isMain":false},{"metricName":"# N's","quality":"Less is better","values":[0,0,0,0,0,0],"isMain":false},{"metricName":"# N's per 100 kbp","quality":"Less is better","values":["0.00","0.00","0.00","0.00","0.00","0.00"],"isMain":true}]],["Statistics without reference",[{"metricName":"# contigs","quality":"Equal","values":[2250,1827,3119,3161,1892,2551],"isMain":true},{"metricName":"# contigs (>= 1000 bp)","quality":"Equal","values":[1391,1247,1339,1381,1286,1342],"isMain":false},{"metricName":"# contigs (>= 5000 bp)","quality":"Equal","values":[29,75,5,6,73,7],"isMain":false},{"metricName":"# contigs (>= 10000 bp)","quality":"Equal","values":[2,8,1,1,7,0],"isMain":false},{"metricName":"# contigs (>= 25000 bp)","quality":"Equal","values":[1,1,0,0,0,0],"isMain":false},{"metricName":"# contigs (>= 50000 bp)","quality":"Equal","values":[0,0,0,0,0,0],"isMain":false},{"metricName":"Largest contig","quality":"More is better","values":[28478,44027,11686,13095,21212,7777],"isMain":true},{"metricName":"Total length","quality":"More is better","values":[3486316,3501519,3504111,3546389,3579611,3260682],"isMain":true},{"metricName":"Total length (>= 1000 bp)","quality":"More is better","values":[2855364,3083934,2231464,2268442,3138548,2395573],"isMain":true},{"metricName":"Total length (>= 5000 bp)","quality":"More is better","values":[210378,533184,33945,44933,524183,40873],"isMain":false},{"metricName":"Total length (>= 10000 bp)","quality":"More is better","values":[45420,124670,11686,13095,106826,0],"isMain":true},{"metricName":"Total length (>= 25000 bp)","quality":"More is better","values":[28478,44027,0,0,0,0],"isMain":false},{"metricName":"Total length (>= 50000 bp)","quality":"More is better","values":[0,0,0,0,0,0],"isMain":true},{"metricName":"N50","quality":"More is better","values":[1871,2536,1243,1237,2479,1488],"isMain":false},{"metricName":"N75","quality":"More is better","values":[1187,1491,831,843,1437,966],"isMain":false},{"metricName":"L50","quality":"Less is better","values":[591,430,909,933,445,714],"isMain":false},{"metricName":"L75","quality":"Less is better","values":[1171,874,1777,1807,914,1393],"isMain":false},{"metricName":"GC (%)","quality":"Equal","values":["41.63","41.71","41.72","41.70","41.63","41.64"],"isMain":false}]],["K-mer-based statistics",[]],["Predicted genes",[]],["Similarity statistics",[]],["Reference statistics",[{"metricName":"Reference length","quality":"Equal","values":[3624708,3624708,3624708,3624708,3624708,3624708],"isMain":false},{"metricName":"Reference fragments","quality":"Equal","values":[62,62,62,62,62,62],"isMain":false},{"metricName":"Reference GC (%)","quality":"Equal","values":["41.59","41.59","41.59","41.59","41.59","41.59"],"isMain":false}]]],"subreferences":[],"subreports":[],"minContig":500},{"date":"10 February 2020, Monday, 18:00:09","assembliesNames":["megahit-103-df","megahit-113-df","megahit-113-ml","megahit-113-ms","megahit-129-df","metaSPAdes"],"referenceName":"GCF\_000153925.1\_ASM15392v1","order":[0,1,2,3,4,5],"report":[["Genome statistics",[{"metricName":"Genome fraction (%)","quality":"More is better","values":["9.684","10.017","10.525","10.358","10.354","7.751"],"isMain":true},{"metricName":"Duplication ratio","quality":"Less is better","values":["1.016","1.017","1.029","1.033","1.023","1.025"],"isMain":true},{"metricName":"Largest alignment","quality":"More is better","values":[2938,3007,3078,3367,3047,1912],"isMain":true},{"metricName":"Total aligned length","quality":"More is better","values":[266724,275254,292704,287682,285231,213444],"isMain":true},{"metricName":"NA50","quality":"More is better","values":[832,984,811,786,897,730],"isMain":false},{"metricName":"NA75","quality":"More is better","values":[613,666,612,618,636,595],"isMain":false},{"metricName":"NGA50","quality":"More is better","values":[null,null,null,null,null,null],"isMain":true},{"metricName":"LA50","quality":"Less is better","values":[114,105,136,133,116,108],"isMain":false},{"metricName":"LA75","quality":"Less is better","values":[214,196,242,240,215,192],"isMain":false}]],["Reads mapping",[]],["Misassemblies",[{"metricName":"# misassemblies","quality":"Less is better","values":[5,2,3,3,3,3],"isMain":true},{"metricName":" # relocations","quality":"Less is better","values":[0,1,0,0,0,0],"isMain":false},{"metricName":" # translocations","quality":"Less is better","values":[5,1,2,3,3,3],"isMain":false},{"metricName":" # inversions","quality":"Less is better","values":[0,0,1,0,0,0],"isMain":false},{"metricName":" # interspecies translocations","quality":"Less is better","values":[16,20,6,14,15,11],"isMain":false},{"metricName":"# misassembled contigs","quality":"Less is better","values":[5,2,3,3,3,3],"isMain":false},{"metricName":"Misassembled contigs length","quality":"Less is better","values":[7603,2255,3311,4502,5071,4469],"isMain":true},{"metricName":"# possibly misassembled contigs","quality":"Less is better","values":[0,0,1,0,0,0],"isMain":false},{"metricName":" # possible misassemblies","quality":"Less is better","values":[0,0,1,0,0,0],"isMain":false},{"metricName":"# local misassemblies","quality":"Less is better","values":[1,3,2,1,2,0],"isMain":false},{"metricName":"# scaffold gap ext. mis.","quality":"Less is better","values":[0,0,0,0,0,0],"isMain":false},{"metricName":"# scaffold gap loc. mis.","quality":"Less is better","values":[0,0,0,0,0,0],"isMain":false},{"metricName":"# unaligned mis. contigs","quality":"Less is better","values":[0,1,0,0,0,0],"isMain":false}]],["Unaligned",[{"metricName":"# fully unaligned contigs","quality":"Less is better","values":[0,0,0,0,0,0],"isMain":false},{"metricName":"Fully unaligned length","quality":"Less is better","values":[0,0,0,0,0,0],"isMain":false},{"metricName":"# partially unaligned contigs","quality":"Less is better","values":[7,11,3,3,5,1],"isMain":false},{"metricName":"Partially unaligned length","quality":"Less is better","values":[12180,14086,3994,3817,10539,777],"isMain":false}]],["Mismatches",[{"metricName":"# mismatches","quality":"Less is better","values":[3124,2999,5373,5119,3465,2724],"isMain":false},{"metricName":"# indels","quality":"Less is better","values":[8,8,16,11,11,2],"isMain":false},{"metricName":"Indels length","quality":"Less is better","values":[46,46,117,46,111,2],"isMain":false},{"metricName":"# mismatches per 100 kbp","quality":"Less is better","values":["1177.60","1092.94","1863.52","1803.98","1221.64","1282.91"],"isMain":true},{"metricName":"# indels per 100 kbp","quality":"Less is better","values":["3.02","2.92","5.55","3.88","3.88","0.94"],"isMain":true},{"metricName":" # indels (<= 5 bp)","quality":"Less is better","values":[7,7,13,10,8,2],"isMain":false},{"metricName":" # indels (> 5 bp)","quality":"Less is better","values":[1,1,3,1,3,0],"isMain":false},{"metricName":"# N's","quality":"Less is better","values":[0,0,0,0,0,0],"isMain":false},{"metricName":"# N's per 100 kbp","quality":"Less is better","values":["0.00","0.00","0.00","0.00","0.00","0.00"],"isMain":true}]],["Statistics without reference",[{"metricName":"# contigs","quality":"Equal","values":[315,298,364,359,322,284],"isMain":true},{"metricName":"# contigs (>= 1000 bp)","quality":"Equal","values":[84,108,84,74,100,44],"isMain":false},{"metricName":"# contigs (>= 5000 bp)","quality":"Equal","values":[1,0,0,0,1,0],"isMain":false},{"metricName":"# contigs (>= 10000 bp)","quality":"Equal","values":[0,0,0,0,0,0],"isMain":false},{"metricName":"# contigs (>= 25000 bp)","quality":"Equal","values":[0,0,0,0,0,0],"isMain":false},{"metricName":"# contigs (>= 50000 bp)","quality":"Equal","values":[0,0,0,0,0,0],"isMain":false},{"metricName":"Largest contig","quality":"More is better","values":[5501,3741,3078,3367,5302,2880],"isMain":true},{"metricName":"Total length","quality":"More is better","values":[281833,293077,300789,296967,300660,218343],"isMain":true},{"metricName":"Total length (>= 1000 bp)","quality":"More is better","values":[126176,160455,108328,101611,147741,57914],"isMain":true},{"metricName":"Total length (>= 5000 bp)","quality":"More is better","values":[5501,0,0,0,5302,0],"isMain":false},{"metricName":"Total length (>= 10000 bp)","quality":"More is better","values":[0,0,0,0,0,0],"isMain":true},{"metricName":"Total length (>= 25000 bp)","quality":"More is better","values":[0,0,0,0,0,0],"isMain":false},{"metricName":"Total length (>= 50000 bp)","quality":"More is better","values":[0,0,0,0,0,0],"isMain":true},{"metricName":"N50","quality":"More is better","values":[932,1069,859,823,994,753],"isMain":false},{"metricName":"N75","quality":"More is better","values":[654,734,650,652,695,616],"isMain":false},{"metricName":"L50","quality":"Less is better","values":[100,95,131,126,103,104],"isMain":false},{"metricName":"L75","quality":"Less is better","values":[192,177,232,229,194,185],"isMain":false},{"metricName":"GC (%)","quality":"Equal","values":["42.38","42.22","42.30","42.36","42.35","42.33"],"isMain":false}]],["K-mer-based statistics",[]],["Predicted genes",[]],["Similarity statistics",[]],["Reference statistics",[{"metricName":"Reference length","quality":"Equal","values":[2739406,2739406,2739406,2739406,2739406,2739406],"isMain":false},{"metricName":"Reference fragments","quality":"Equal","values":[64,64,64,64,64,64],"isMain":false},{"metricName":"Reference GC (%)","quality":"Equal","values":["41.97","41.97","41.97","41.97","41.97","41.97"],"isMain":false}]]],"subreferences":[],"subreports":[],"minContig":500},{"date":"10 February 2020, Monday, 18:00:24","assembliesNames":["megahit-103-df","megahit-113-df","megahit-113-ml","megahit-113-ms","megahit-129-df","metaSPAdes"],"referenceName":"GCF\_000154065.1\_ASM15406v1","order":[0,1,2,3,4,5],"report":[["Genome statistics",[{"metricName":"Genome fraction (%)","quality":"More is better","values":["10.240","13.317","13.927","14.188","14.450","6.627"],"isMain":true},{"metricName":"Duplication ratio","quality":"Less is better","values":["1.033","1.031","1.024","1.040","1.029","1.044"],"isMain":true},{"metricName":"Largest alignment","quality":"More is better","values":[1917,2714,1741,1495,1955,2003],"isMain":true},{"metricName":"Total aligned length","quality":"More is better","values":[299598,389534,409160,416054,423486,193555],"isMain":true},{"metricName":"NA50","quality":"More is better","values":[617,632,597,607,642,573],"isMain":false},{"metricName":"NA75","quality":"More is better","values":[532,534,525,531,540,521],"isMain":false},{"metricName":"NGA50","quality":"More is better","values":[null,null,null,null,null,null],"isMain":true},{"metricName":"LA50","quality":"Less is better","values":[200,258,293,293,271,146],"isMain":false},{"metricName":"LA75","quality":"Less is better","values":[339,441,493,491,459,241],"isMain":false}]],["Reads mapping",[]],["Misassemblies",[{"metricName":"# misassemblies","quality":"Less is better","values":[2,4,4,1,3,1],"isMain":true},{"metricName":" # relocations","quality":"Less is better","values":[1,0,1,0,0,1],"isMain":false},{"metricName":" # translocations","quality":"Less is better","values":[1,3,3,1,2,0],"isMain":false},{"metricName":" # inversions","quality":"Less is better","values":[0,1,0,0,1,0],"isMain":false},{"metricName":" # interspecies translocations","quality":"Less is better","values":[39,36,15,48,36,27],"isMain":false},{"metricName":"# misassembled contigs","quality":"Less is better","values":[2,4,4,1,3,1],"isMain":false},{"metricName":"Misassembled contigs length","quality":"Less is better","values":[1439,5146,3661,854,4023,968],"isMain":true},{"metricName":"# possibly misassembled contigs","quality":"Less is better","values":[0,0,0,0,0,0],"isMain":false},{"metricName":" # possible misassemblies","quality":"Less is better","values":[0,0,0,0,0,0],"isMain":false},{"metricName":"# local misassemblies","quality":"Less is better","values":[1,1,1,0,1,0],"isMain":false},{"metricName":"# scaffold gap ext. mis.","quality":"Less is better","values":[0,0,0,0,0,0],"isMain":false},{"metricName":"# scaffold gap loc. mis.","quality":"Less is better","values":[0,0,0,0,0,0],"isMain":false},{"metricName":"# unaligned mis. contigs","quality":"Less is better","values":[0,0,0,0,0,0],"isMain":false}]],["Unaligned",[{"metricName":"# fully unaligned contigs","quality":"Less is better","values":[1,0,0,0,0,0],"isMain":false},{"metricName":"Fully unaligned length","quality":"Less is better","values":[734,0,0,0,0,0],"isMain":false},{"metricName":"# partially unaligned contigs","quality":"Less is better","values":[8,10,3,11,7,7],"isMain":false},{"metricName":"Partially unaligned length","quality":"Less is better","values":[6003,22750,33132,17760,9534,5076],"isMain":false}]],["Mismatches",[{"metricName":"# mismatches","quality":"Less is better","values":[4625,5978,7958,8484,6708,3190],"isMain":false},{"metricName":"# indels","quality":"Less is better","values":[17,14,14,6,9,11],"isMain":false},{"metricName":"Indels length","quality":"Less is better","values":[35,14,14,8,62,55],"isMain":false},{"metricName":"# mismatches per 100 kbp","quality":"Less is better","values":["1549.98","1540.56","1961.01","2052.23","1593.11","1651.99"],"isMain":true},{"metricName":"# indels per 100 kbp","quality":"Less is better","values":["5.70","3.61","3.45","1.45","2.14","5.70"],"isMain":true},{"metricName":" # indels (<= 5 bp)","quality":"Less is better","values":[16,14,14,6,8,10],"isMain":false},{"metricName":" # indels (> 5 bp)","quality":"Less is better","values":[1,0,0,0,1,1],"isMain":false},{"metricName":"# N's","quality":"Less is better","values":[0,0,0,0,0,0],"isMain":false},{"metricName":"# N's per 100 kbp","quality":"Less is better","values":["0.00","0.00","0.00","0.00","0.00","0.00"],"isMain":true}]],["Statistics without reference",[{"metricName":"# contigs","quality":"Equal","values":[473,591,638,664,645,326],"isMain":true},{"metricName":"# contigs (>= 1000 bp)","quality":"Equal","values":[24,44,27,33,40,9],"isMain":false},{"metricName":"# contigs (>= 5000 bp)","quality":"Equal","values":[0,3,1,0,1,0],"isMain":false},{"metricName":"# contigs (>= 10000 bp)","quality":"Equal","values":[0,0,1,0,0,0],"isMain":false},{"metricName":"# contigs (>= 25000 bp)","quality":"Equal","values":[0,0,1,0,0,0],"isMain":false},{"metricName":"# contigs (>= 50000 bp)","quality":"Equal","values":[0,0,0,0,0,0],"isMain":false},{"metricName":"Largest contig","quality":"More is better","values":[2425,6114,31963,4337,6310,2061],"isMain":true},{"metricName":"Total length","quality":"More is better","values":[314951,422856,448676,447744,442989,206649],"isMain":true},{"metricName":"Total length (>= 1000 bp)","quality":"More is better","values":[30744,71140,62347,47257,54313,12067],"isMain":true},{"metricName":"Total length (>= 5000 bp)","quality":"More is better","values":[0,17300,31963,0,6310,0],"isMain":false},{"metricName":"Total length (>= 10000 bp)","quality":"More is better","values":[0,0,31963,0,0,0],"isMain":true},{"metricName":"Total length (>= 25000 bp)","quality":"More is better","values":[0,0,31963,0,0,0],"isMain":false},{"metricName":"Total length (>= 50000 bp)","quality":"More is better","values":[0,0,0,0,0,0],"isMain":true},{"metricName":"N50","quality":"More is better","values":[648,679,641,640,662,610],"isMain":false},{"metricName":"N75","quality":"More is better","values":[555,571,559,562,566,541],"isMain":false},{"metricName":"L50","quality":"Less is better","values":[192,222,237,265,254,137],"isMain":false},{"metricName":"L75","quality":"Less is better","values":[324,392,425,453,435,227],"isMain":false},{"metricName":"GC (%)","quality":"Equal","values":["41.88","41.51","41.95","41.52","41.40","41.19"],"isMain":false}]],["K-mer-based statistics",[]],["Predicted genes",[]],["Similarity statistics",[]],["Reference statistics",[{"metricName":"Reference length","quality":"Equal","values":[2913833,2913833,2913833,2913833,2913833,2913833],"isMain":false},{"metricName":"Reference fragments","quality":"Equal","values":[52,52,52,52,52,52],"isMain":false},{"metricName":"Reference GC (%)","quality":"Equal","values":["41.44","41.44","41.44","41.44","41.44","41.44"],"isMain":false}]]],"subreferences":[],"subreports":[],"minContig":500},{"date":"10 February 2020, Monday, 18:00:43","assembliesNames":["megahit-103-df","megahit-113-df","megahit-113-ml","megahit-113-ms","megahit-129-df","metaSPAdes"],"referenceName":"GCF\_000154105.1\_ASM15410v1","order":[0,1,2,3,4,5],"report":[["Genome statistics",[{"metricName":"Genome fraction (%)","quality":"More is better","values":["8.663","9.568","8.839","10.681","10.200","6.989"],"isMain":true},{"metricName":"Duplication ratio","quality":"Less is better","values":["1.032","1.040","1.033","1.045","1.033","1.035"],"isMain":true},{"metricName":"Largest alignment","quality":"More is better","values":[3059,2981,2543,2363,4035,2337],"isMain":true},{"metricName":"Total aligned length","quality":"More is better","values":[387092,426884,395375,480552,455530,311992],"isMain":true},{"metricName":"NA50","quality":"More is better","values":[712,797,691,695,807,708],"isMain":false},{"metricName":"NA75","quality":"More is better","values":[563,602,568,550,602,565],"isMain":false},{"metricName":"NGA50","quality":"More is better","values":[null,null,null,null,null,null],"isMain":true},{"metricName":"LA50","quality":"Less is better","values":[207,191,217,283,197,171],"isMain":false},{"metricName":"LA75","quality":"Less is better","values":[368,355,381,502,366,300],"isMain":false}]],["Reads mapping",[]],["Misassemblies",[{"metricName":"# misassemblies","quality":"Less is better","values":[6,8,2,1,6,1],"isMain":true},{"metricName":" # relocations","quality":"Less is better","values":[1,2,1,0,1,0],"isMain":false},{"metricName":" # translocations","quality":"Less is better","values":[5,5,1,1,5,1],"isMain":false},{"metricName":" # inversions","quality":"Less is better","values":[0,1,0,0,0,0],"isMain":false},{"metricName":" # interspecies translocations","quality":"Less is better","values":[25,43,17,33,26,16],"isMain":false},{"metricName":"# misassembled contigs","quality":"Less is better","values":[6,7,2,1,6,1],"isMain":false},{"metricName":"Misassembled contigs length","quality":"Less is better","values":[5149,7823,1296,525,8060,608],"isMain":true},{"metricName":"# possibly misassembled contigs","quality":"Less is better","values":[1,0,1,0,0,0],"isMain":false},{"metricName":" # possible misassemblies","quality":"Less is better","values":[1,0,1,0,0,0],"isMain":false},{"metricName":"# local misassemblies","quality":"Less is better","values":[2,1,0,3,0,0],"isMain":false},{"metricName":"# scaffold gap ext. mis.","quality":"Less is better","values":[0,0,0,0,0,0],"isMain":false},{"metricName":"# scaffold gap loc. mis.","quality":"Less is better","values":[0,0,0,0,0,0],"isMain":false},{"metricName":"# unaligned mis. contigs","quality":"Less is better","values":[0,0,0,0,0,0],"isMain":false}]],["Unaligned",[{"metricName":"# fully unaligned contigs","quality":"Less is better","values":[4,4,2,2,3,0],"isMain":false},{"metricName":"Fully unaligned length","quality":"Less is better","values":[2905,2932,1454,1675,2417,0],"isMain":false},{"metricName":"# partially unaligned contigs","quality":"Less is better","values":[4,10,2,7,5,4],"isMain":false},{"metricName":"Partially unaligned length","quality":"Less is better","values":[8987,7908,1509,39388,3617,2426],"isMain":false}]],["Mismatches",[{"metricName":"# mismatches","quality":"Less is better","values":[6965,8079,8704,10876,8327,5433],"isMain":false},{"metricName":"# indels","quality":"Less is better","values":[47,68,48,53,68,42],"isMain":false},{"metricName":"Indels length","quality":"Less is better","values":[77,101,76,82,137,83],"isMain":false},{"metricName":"# mismatches per 100 kbp","quality":"Less is better","values":["1814.12","1905.18","2221.93","2297.55","1842.09","1753.94"],"isMain":true},{"metricName":"# indels per 100 kbp","quality":"Less is better","values":["12.24","16.04","12.25","11.20","15.04","13.56"],"isMain":true},{"metricName":" # indels (<= 5 bp)","quality":"Less is better","values":[44,67,46,51,66,39],"isMain":false},{"metricName":" # indels (> 5 bp)","quality":"Less is better","values":[3,1,2,2,2,3],"isMain":false},{"metricName":"# N's","quality":"Less is better","values":[0,0,0,0,0,0],"isMain":false},{"metricName":"# N's per 100 kbp","quality":"Less is better","values":["0.00","0.00","0.00","0.00","0.00","0.00"],"isMain":true}]],["Statistics without reference",[{"metricName":"# contigs","quality":"Equal","values":[527,527,555,665,555,438],"isMain":true},{"metricName":"# contigs (>= 1000 bp)","quality":"Equal","values":[81,118,66,91,116,54],"isMain":false},{"metricName":"# contigs (>= 5000 bp)","quality":"Equal","values":[1,0,0,1,0,0],"isMain":false},{"metricName":"# contigs (>= 10000 bp)","quality":"Equal","values":[0,0,0,1,0,0],"isMain":false},{"metricName":"# contigs (>= 25000 bp)","quality":"Equal","values":[0,0,0,1,0,0],"isMain":false},{"metricName":"# contigs (>= 50000 bp)","quality":"Equal","values":[0,0,0,0,0,0],"isMain":false},{"metricName":"Largest contig","quality":"More is better","values":[6522,2981,2545,35116,4035,2445],"isMain":true},{"metricName":"Total length","quality":"More is better","values":[408242,451940,407716,535604,472821,323081],"isMain":true},{"metricName":"Total length (>= 1000 bp)","quality":"More is better","values":[111014,172113,85804,149920,170462,68118],"isMain":true},{"metricName":"Total length (>= 5000 bp)","quality":"More is better","values":[6522,0,0,35116,0,0],"isMain":false},{"metricName":"Total length (>= 10000 bp)","quality":"More is better","values":[0,0,0,35116,0,0],"isMain":true},{"metricName":"Total length (>= 25000 bp)","quality":"More is better","values":[0,0,0,35116,0,0],"isMain":false},{"metricName":"Total length (>= 50000 bp)","quality":"More is better","values":[0,0,0,0,0,0],"isMain":true},{"metricName":"N50","quality":"More is better","values":[764,852,714,767,857,734],"isMain":false},{"metricName":"N75","quality":"More is better","values":[599,654,588,610,648,593],"isMain":false},{"metricName":"L50","quality":"Less is better","values":[191,177,209,228,187,165],"isMain":false},{"metricName":"L75","quality":"Less is better","values":[342,330,367,423,346,289],"isMain":false},{"metricName":"GC (%)","quality":"Equal","values":["45.43","45.28","45.76","45.10","45.42","45.76"],"isMain":false}]],["K-mer-based statistics",[]],["Predicted genes",[]],["Similarity statistics",[]],["Reference statistics",[{"metricName":"Reference length","quality":"Equal","values":[4431877,4431877,4431877,4431877,4431877,4431877],"isMain":false},{"metricName":"Reference fragments","quality":"Equal","values":[118,118,118,118,118,118],"isMain":false},{"metricName":"Reference GC (%)","quality":"Equal","values":["45.30","45.30","45.30","45.30","45.30","45.30"],"isMain":false}]]],"subreferences":[],"subreports":[],"minContig":500},{"date":"10 February 2020, Monday, 18:01:30","assembliesNames":["megahit-103-df","megahit-113-df","megahit-113-ml","megahit-113-ms","megahit-129-df","metaSPAdes"],"referenceName":"GCF\_000154125.1\_ASM15412v1","order":[0,1,2,3,4,5],"report":[["Genome statistics",[{"metricName":"Genome fraction (%)","quality":"More is better","values":["22.934","15.225","16.023","19.746","16.099","10.733"],"isMain":true},{"metricName":"Duplication ratio","quality":"Less is better","values":["1.006","1.012","1.011","1.015","1.022","1.001"],"isMain":true},{"metricName":"Largest alignment","quality":"More is better","values":[94586,43249,63215,60452,50038,82969],"isMain":true},{"metricName":"Total aligned length","quality":"More is better","values":[1489405,991326,1043861,1289919,1055892,694201],"isMain":true},{"metricName":"NA50","quality":"More is better","values":[14445,16047,15705,17824,12142,20077],"isMain":false},{"metricName":"NA75","quality":"More is better","values":[7550,9855,8757,8241,6407,13036],"isMain":false},{"metricName":"NGA50","quality":"More is better","values":[null,null,null,null,null,null],"isMain":true},{"metricName":"LA50","quality":"Less is better","values":[33,21,20,24,28,9],"isMain":false},{"metricName":"LA75","quality":"Less is better","values":[68,42,43,52,60,20],"isMain":false}]],["Reads mapping",[]],["Misassemblies",[{"metricName":"# misassemblies","quality":"Less is better","values":[0,0,0,2,3,0],"isMain":true},{"metricName":" # relocations","quality":"Less is better","values":[0,0,0,0,0,0],"isMain":false},{"metricName":" # translocations","quality":"Less is better","values":[0,0,0,2,2,0],"isMain":false},{"metricName":" # inversions","quality":"Less is better","values":[0,0,0,0,1,0],"isMain":false},{"metricName":" # interspecies translocations","quality":"Less is better","values":[5,16,10,14,25,1],"isMain":false},{"metricName":"# misassembled contigs","quality":"Less is better","values":[0,0,0,1,3,0],"isMain":false},{"metricName":"Misassembled contigs length","quality":"Less is better","values":[0,0,0,36105,75266,0],"isMain":true},{"metricName":"# possibly misassembled contigs","quality":"Less is better","values":[0,0,0,1,1,0],"isMain":false},{"metricName":" # possible misassemblies","quality":"Less is better","values":[0,0,0,1,1,0],"isMain":false},{"metricName":"# local misassemblies","quality":"Less is better","values":[0,2,1,1,1,0],"isMain":false},{"metricName":"# scaffold gap ext. mis.","quality":"Less is better","values":[0,0,0,0,0,0],"isMain":false},{"metricName":"# scaffold gap loc. mis.","quality":"Less is better","values":[0,0,0,0,0,0],"isMain":false},{"metricName":"# unaligned mis. contigs","quality":"Less is better","values":[0,0,0,0,0,0],"isMain":false}]],["Unaligned",[{"metricName":"# fully unaligned contigs","quality":"Less is better","values":[0,0,0,0,0,0],"isMain":false},{"metricName":"Fully unaligned length","quality":"Less is better","values":[0,0,0,0,0,0],"isMain":false},{"metricName":"# partially unaligned contigs","quality":"Less is better","values":[1,7,6,8,13,1],"isMain":false},{"metricName":"Partially unaligned length","quality":"Less is better","values":[1419,44621,26050,62612,69071,749],"isMain":false}]],["Mismatches",[{"metricName":"# mismatches","quality":"Less is better","values":[1621,2566,2228,3714,3091,422],"isMain":false},{"metricName":"# indels","quality":"Less is better","values":[22,42,55,40,65,21],"isMain":false},{"metricName":"Indels length","quality":"Less is better","values":[23,84,175,110,115,27],"isMain":false},{"metricName":"# mismatches per 100 kbp","quality":"Less is better","values":["109.36","260.77","215.14","291.02","297.07","60.83"],"isMain":true},{"metricName":"# indels per 100 kbp","quality":"Less is better","values":["1.48","4.27","5.31","3.13","6.25","3.03"],"isMain":true},{"metricName":" # indels (<= 5 bp)","quality":"Less is better","values":[22,40,51,37,63,21],"isMain":false},{"metricName":" # indels (> 5 bp)","quality":"Less is better","values":[0,2,4,3,2,0],"isMain":false},{"metricName":"# N's","quality":"Less is better","values":[0,0,0,0,0,0],"isMain":false},{"metricName":"# N's per 100 kbp","quality":"Less is better","values":["0.00","0.00","0.00","0.00","0.00","0.00"],"isMain":true}]],["Statistics without reference",[{"metricName":"# contigs","quality":"Equal","values":[187,120,123,145,155,76],"isMain":true},{"metricName":"# contigs (>= 1000 bp)","quality":"Equal","values":[169,93,103,120,121,58],"isMain":false},{"metricName":"# contigs (>= 5000 bp)","quality":"Equal","values":[91,57,62,73,66,32],"isMain":false},{"metricName":"# contigs (>= 10000 bp)","quality":"Equal","values":[51,42,37,45,38,22],"isMain":false},{"metricName":"# contigs (>= 25000 bp)","quality":"Equal","values":[6,9,7,13,6,7],"isMain":false},{"metricName":"# contigs (>= 50000 bp)","quality":"Equal","values":[1,0,2,1,2,1],"isMain":false},{"metricName":"Largest contig","quality":"More is better","values":[94590,43249,63215,60523,56868,82969],"isMain":true},{"metricName":"Total length","quality":"More is better","values":[1492951,1040920,1072649,1357839,1132211,694962],"isMain":true},{"metricName":"Total length (>= 1000 bp)","quality":"More is better","values":[1480431,1023202,1059269,1340489,1110563,680872],"isMain":true},{"metricName":"Total length (>= 5000 bp)","quality":"More is better","values":[1264089,937445,960676,1225406,964612,626647],"isMain":false},{"metricName":"Total length (>= 10000 bp)","quality":"More is better","values":[974209,821052,778773,1014686,753623,552001],"isMain":true},{"metricName":"Total length (>= 25000 bp)","quality":"More is better","values":[251070,287511,302853,472430,246932,309742],"isMain":false},{"metricName":"Total length (>= 50000 bp)","quality":"More is better","values":[94590,0,125005,60523,107041,82969],"isMain":true},{"metricName":"N50","quality":"More is better","values":[14708,17597,17247,20327,14993,20077],"isMain":false},{"metricName":"N75","quality":"More is better","values":[7550,11183,9221,9917,8505,13036],"isMain":false},{"metricName":"L50","quality":"Less is better","values":[32,20,19,22,23,9],"isMain":false},{"metricName":"L75","quality":"Less is better","values":[68,39,40,46,49,20],"isMain":false},{"metricName":"GC (%)","quality":"Equal","values":["41.72","41.82","42.46","41.56","42.42","42.21"],"isMain":false}]],["K-mer-based statistics",[]],["Predicted genes",[]],["Similarity statistics",[]],["Reference statistics",[{"metricName":"Reference length","quality":"Equal","values":[6463169,6463169,6463169,6463169,6463169,6463169],"isMain":false},{"metricName":"Reference fragments","quality":"Equal","values":[54,54,54,54,54,54],"isMain":false},{"metricName":"Reference GC (%)","quality":"Equal","values":["41.86","41.86","41.86","41.86","41.86","41.86"],"isMain":false}]]],"subreferences":[],"subreports":[],"minContig":500},{"date":"10 February 2020, Monday, 18:00:22","assembliesNames":["megahit-103-df","megahit-113-df","megahit-113-ml","megahit-113-ms","megahit-129-df","metaSPAdes"],"referenceName":"GCF\_000154345.1\_ASM15434v1","order":[0,1,2,3,4,5],"report":[["Genome statistics",[{"metricName":"Genome fraction (%)","quality":"More is better","values":["1.048","1.163","1.561","2.346","1.391","0.715"],"isMain":true},{"metricName":"Duplication ratio","quality":"Less is better","values":["1.105","1.093","1.037","1.048","1.096","1.103"],"isMain":true},{"metricName":"Largest alignment","quality":"More is better","values":[991,967,907,1861,1267,1651],"isMain":true},{"metricName":"Total aligned length","quality":"More is better","values":[34265,38045,51060,76759,45454,23383],"isMain":true},{"metricName":"NA50","quality":"More is better","values":[556,543,552,528,554,563],"isMain":false},{"metricName":"NA75","quality":"More is better","values":[506,326,521,null,501,493],"isMain":false},{"metricName":"NGA50","quality":"More is better","values":[null,null,null,null,null,null],"isMain":true},{"metricName":"LA50","quality":"Less is better","values":[29,38,42,83,41,20],"isMain":false},{"metricName":"LA75","quality":"Less is better","values":[47,62,67,null,66,34],"isMain":false}]],["Reads mapping",[]],["Misassemblies",[{"metricName":"# misassemblies","quality":"Less is better","values":[0,0,1,1,0,0],"isMain":true},{"metricName":" # relocations","quality":"Less is better","values":[0,0,0,1,0,0],"isMain":false},{"metricName":" # translocations","quality":"Less is better","values":[0,0,1,0,0,0],"isMain":false},{"metricName":" # inversions","quality":"Less is better","values":[0,0,0,0,0,0],"isMain":false},{"metricName":" # interspecies translocations","quality":"Less is better","values":[14,18,4,16,17,8],"isMain":false},{"metricName":"# misassembled contigs","quality":"Less is better","values":[0,0,1,1,0,0],"isMain":false},{"metricName":"Misassembled contigs length","quality":"Less is better","values":[0,0,511,711,0,0],"isMain":true},{"metricName":"# possibly misassembled contigs","quality":"Less is better","values":[0,0,0,0,0,0],"isMain":false},{"metricName":" # possible misassemblies","quality":"Less is better","values":[0,0,0,0,0,0],"isMain":false},{"metricName":"# local misassemblies","quality":"Less is better","values":[0,0,0,0,0,0],"isMain":false},{"metricName":"# scaffold gap ext. mis.","quality":"Less is better","values":[0,0,0,0,0,0],"isMain":false},{"metricName":"# scaffold gap loc. mis.","quality":"Less is better","values":[0,0,0,0,0,0],"isMain":false},{"metricName":"# unaligned mis. contigs","quality":"Less is better","values":[0,0,0,1,0,0],"isMain":false}]],["Unaligned",[{"metricName":"# fully unaligned contigs","quality":"Less is better","values":[0,0,1,0,1,0],"isMain":false},{"metricName":"Fully unaligned length","quality":"Less is better","values":[0,0,568,0,646,0],"isMain":false},{"metricName":"# partially unaligned contigs","quality":"Less is better","values":[1,7,0,8,6,1],"isMain":false},{"metricName":"Partially unaligned length","quality":"Less is better","values":[588,5191,0,25378,3892,1619],"isMain":false}]],["Mismatches",[{"metricName":"# mismatches","quality":"Less is better","values":[652,779,1069,1653,912,429],"isMain":false},{"metricName":"# indels","quality":"Less is better","values":[3,3,3,0,2,3],"isMain":false},{"metricName":"Indels length","quality":"Less is better","values":[3,3,3,0,42,3],"isMain":false},{"metricName":"# mismatches per 100 kbp","quality":"Less is better","values":["1902.76","2047.52","2094.23","2154.53","2005.10","1834.59"],"isMain":true},{"metricName":"# indels per 100 kbp","quality":"Less is better","values":["8.76","7.89","5.88","0.00","4.40","12.83"],"isMain":true},{"metricName":" # indels (<= 5 bp)","quality":"Less is better","values":[3,3,3,0,0,3],"isMain":false},{"metricName":" # indels (> 5 bp)","quality":"Less is better","values":[0,0,0,0,2,0],"isMain":false},{"metricName":"# N's","quality":"Less is better","values":[0,0,0,0,0,0],"isMain":false},{"metricName":"# N's per 100 kbp","quality":"Less is better","values":["0.00","0.00","0.00","0.00","0.00","0.00"],"isMain":true}]],["Statistics without reference",[{"metricName":"# contigs","quality":"Equal","values":[62,73,91,132,84,42],"isMain":true},{"metricName":"# contigs (>= 1000 bp)","quality":"Equal","values":[0,4,0,5,3,2],"isMain":false},{"metricName":"# contigs (>= 5000 bp)","quality":"Equal","values":[0,0,0,1,0,0],"isMain":false},{"metricName":"# contigs (>= 10000 bp)","quality":"Equal","values":[0,0,0,1,0,0],"isMain":false},{"metricName":"# contigs (>= 25000 bp)","quality":"Equal","values":[0,0,0,0,0,0],"isMain":false},{"metricName":"# contigs (>= 50000 bp)","quality":"Equal","values":[0,0,0,0,0,0],"isMain":false},{"metricName":"Largest contig","quality":"More is better","values":[991,1151,907,24289,1276,2220],"isMain":true},{"metricName":"Total length","quality":"More is better","values":[38441,46788,53496,105801,54379,27422],"isMain":true},{"metricName":"Total length (>= 1000 bp)","quality":"More is better","values":[0,4320,0,30053,3533,3871],"isMain":true},{"metricName":"Total length (>= 5000 bp)","quality":"More is better","values":[0,0,0,24289,0,0],"isMain":false},{"metricName":"Total length (>= 10000 bp)","quality":"More is better","values":[0,0,0,24289,0,0],"isMain":true},{"metricName":"Total length (>= 25000 bp)","quality":"More is better","values":[0,0,0,0,0,0],"isMain":false},{"metricName":"Total length (>= 50000 bp)","quality":"More is better","values":[0,0,0,0,0,0],"isMain":true},{"metricName":"N50","quality":"More is better","values":[579,590,568,618,614,606],"isMain":false},{"metricName":"N75","quality":"More is better","values":[541,553,532,545,556,521],"isMain":false},{"metricName":"L50","quality":"Less is better","values":[27,31,41,36,36,17],"isMain":false},{"metricName":"L75","quality":"Less is better","values":[44,51,66,82,59,29],"isMain":false},{"metricName":"GC (%)","quality":"Equal","values":["48.57","50.78","49.94","47.43","50.80","47.97"],"isMain":false}]],["K-mer-based statistics",[]],["Predicted genes",[]],["Similarity statistics",[]],["Reference statistics",[{"metricName":"Reference length","quality":"Equal","values":[3270109,3270109,3270109,3270109,3270109,3270109],"isMain":false},{"metricName":"Reference fragments","quality":"Equal","values":[22,22,22,22,22,22],"isMain":false},{"metricName":"Reference GC (%)","quality":"Equal","values":["50.19","50.19","50.19","50.19","50.19","50.19"],"isMain":false}]]],"subreferences":[],"subreports":[],"minContig":500},{"date":"10 February 2020, Monday, 18:00:33","assembliesNames":["megahit-103-df","megahit-113-df","megahit-113-ml","megahit-113-ms","megahit-129-df","metaSPAdes"],"referenceName":"GCF\_000154385.1\_ASM15438v1","order":[0,1,2,3,4,5],"report":[["Genome statistics",[{"metricName":"Genome fraction (%)","quality":"More is better","values":["91.911","91.429","94.798","94.088","92.700","91.910"],"isMain":true},{"metricName":"Duplication ratio","quality":"Less is better","values":["1.022","1.046","1.037","1.049","1.055","1.003"],"isMain":true},{"metricName":"Largest alignment","quality":"More is better","values":[47965,57214,89416,76434,82825,160312],"isMain":true},{"metricName":"Total aligned length","quality":"More is better","values":[2934176,2977766,3064887,3070532,3042359,2881386],"isMain":true},{"metricName":"NG50","quality":"More is better","values":[8572,12940,16020,17089,19821,28826],"isMain":false},{"metricName":"NG75","quality":"More is better","values":[3174,6090,7548,8339,9836,8312],"isMain":false},{"metricName":"NA50","quality":"More is better","values":[9487,12798,15010,15034,12531,30057],"isMain":false},{"metricName":"NA75","quality":"More is better","values":[3917,5979,6904,6672,3857,11623],"isMain":false},{"metricName":"NGA50","quality":"More is better","values":[8556,12336,15151,15650,13602,28826],"isMain":true},{"metricName":"NGA75","quality":"More is better","values":[3153,5723,6928,7125,5853,8072],"isMain":false},{"metricName":"LG50","quality":"Less is better","values":[105,72,52,54,51,33],"isMain":false},{"metricName":"LG75","quality":"Less is better","values":[255,158,121,119,108,87],"isMain":false},{"metricName":"LA50","quality":"Less is better","values":[95,72,55,62,76,29],"isMain":false},{"metricName":"LA75","quality":"Less is better","values":[216,159,131,141,192,70],"isMain":false},{"metricName":"LGA50","quality":"Less is better","values":[105,75,54,60,66,33],"isMain":true},{"metricName":"LGA75","quality":"Less is better","values":[256,167,130,134,149,89],"isMain":false}]],["Reads mapping",[]],["Misassemblies",[{"metricName":"# misassemblies","quality":"Less is better","values":[4,7,5,17,14,2],"isMain":true},{"metricName":" # relocations","quality":"Less is better","values":[0,1,2,5,4,0],"isMain":false},{"metricName":" # translocations","quality":"Less is better","values":[4,5,2,10,8,2],"isMain":false},{"metricName":" # inversions","quality":"Less is better","values":[0,1,1,2,2,0],"isMain":false},{"metricName":" # interspecies translocations","quality":"Less is better","values":[13,43,28,46,75,2],"isMain":false},{"metricName":"# misassembled contigs","quality":"Less is better","values":[4,7,5,17,14,2],"isMain":false},{"metricName":"Misassembled contigs length","quality":"Less is better","values":[7415,46224,54519,215009,172970,40719],"isMain":true},{"metricName":"# possibly misassembled contigs","quality":"Less is better","values":[1,1,1,1,2,0],"isMain":false},{"metricName":" # possible misassemblies","quality":"Less is better","values":[1,1,1,1,2,0],"isMain":false},{"metricName":"# local misassemblies","quality":"Less is better","values":[2,6,2,5,7,0],"isMain":false},{"metricName":"# scaffold gap ext. mis.","quality":"Less is better","values":[0,0,0,0,0,0],"isMain":false},{"metricName":"# scaffold gap loc. mis.","quality":"Less is better","values":[0,0,0,0,0,0],"isMain":false},{"metricName":"# unaligned mis. contigs","quality":"Less is better","values":[0,0,0,1,4,0],"isMain":false}]],["Unaligned",[{"metricName":"# fully unaligned contigs","quality":"Less is better","values":[3,2,0,1,2,0],"isMain":false},{"metricName":"Fully unaligned length","quality":"Less is better","values":[4529,2891,0,1925,1374,0],"isMain":false},{"metricName":"# partially unaligned contigs","quality":"Less is better","values":[2,19,15,21,47,1],"isMain":false},{"metricName":"Partially unaligned length","quality":"Less is better","values":[2227,65451,66630,104947,343933,1565],"isMain":false}]],["Mismatches",[{"metricName":"# mismatches","quality":"Less is better","values":[4457,11650,7844,11714,11656,3393],"isMain":false},{"metricName":"# indels","quality":"Less is better","values":[42,203,153,203,201,128],"isMain":false},{"metricName":"Indels length","quality":"Less is better","values":[103,330,526,546,508,232],"isMain":false},{"metricName":"# mismatches per 100 kbp","quality":"Less is better","values":["155.08","407.49","264.61","398.15","402.11","118.06"],"isMain":true},{"metricName":"# indels per 100 kbp","quality":"Less is better","values":["1.46","7.10","5.16","6.90","6.93","4.45"],"isMain":true},{"metricName":" # indels (<= 5 bp)","quality":"Less is better","values":[40,200,139,187,192,123],"isMain":false},{"metricName":" # indels (> 5 bp)","quality":"Less is better","values":[2,3,14,16,9,5],"isMain":false},{"metricName":"# N's","quality":"Less is better","values":[0,0,0,0,0,0],"isMain":false},{"metricName":"# N's per 100 kbp","quality":"Less is better","values":["0.00","0.00","0.00","0.00","0.00","0.00"],"isMain":true}]],["Statistics without reference",[{"metricName":"# contigs","quality":"Equal","values":[702,530,485,470,562,302],"isMain":true},{"metricName":"# contigs (>= 1000 bp)","quality":"Equal","values":[512,389,360,342,373,235],"isMain":false},{"metricName":"# contigs (>= 5000 bp)","quality":"Equal","values":[179,184,155,171,172,120],"isMain":false},{"metricName":"# contigs (>= 10000 bp)","quality":"Equal","values":[83,100,97,100,105,75],"isMain":false},{"metricName":"# contigs (>= 25000 bp)","quality":"Equal","values":[6,16,27,26,32,35],"isMain":false},{"metricName":"# contigs (>= 50000 bp)","quality":"Equal","values":[0,2,4,5,2,11],"isMain":false},{"metricName":"Largest contig","quality":"More is better","values":[47965,57214,89416,76491,82825,160314],"isMain":true},{"metricName":"Total length","quality":"More is better","values":[2945287,3058334,3139517,3192729,3402870,2883482],"isMain":true},{"metricName":"Total length (>= 1000 bp)","quality":"More is better","values":[2812008,2959345,3053452,3106294,3275692,2836257],"isMain":true},{"metricName":"Total length (>= 5000 bp)","quality":"More is better","values":[2043596,2492981,2571554,2684252,2810646,2562050],"isMain":false},{"metricName":"Total length (>= 10000 bp)","quality":"More is better","values":[1361629,1881820,2143280,2174391,2319584,2235071],"isMain":true},{"metricName":"Total length (>= 25000 bp)","quality":"More is better","values":[219704,600916,1063486,990197,1160795,1624543],"isMain":false},{"metricName":"Total length (>= 50000 bp)","quality":"More is better","values":[0,111715,266851,296030,148327,784483],"isMain":true},{"metricName":"N50","quality":"More is better","values":[9500,13149,16020,16895,16583,30057],"isMain":false},{"metricName":"N75","quality":"More is better","values":[3917,6711,7541,7983,8172,11726],"isMain":false},{"metricName":"L50","quality":"Less is better","values":[95,70,52,56,58,29],"isMain":false},{"metricName":"L75","quality":"Less is better","values":[216,150,122,125,131,69],"isMain":false},{"metricName":"GC (%)","quality":"Equal","values":["56.51","56.38","56.49","56.48","56.10","56.57"],"isMain":false}]],["K-mer-based statistics",[]],["Predicted genes",[]],["Similarity statistics",[]],["Reference statistics",[{"metricName":"Reference length","quality":"Equal","values":[3126983,3126983,3126983,3126983,3126983,3126983],"isMain":false},{"metricName":"Reference fragments","quality":"Equal","values":[29,29,29,29,29,29],"isMain":false},{"metricName":"Reference GC (%)","quality":"Equal","values":["56.24","56.24","56.24","56.24","56.24","56.24"],"isMain":false}]]],"subreferences":[],"subreports":[],"minContig":500},{"date":"10 February 2020, Monday, 17:51:39","assembliesNames":["megahit-103-df","megahit-113-df","megahit-113-ml","megahit-113-ms","megahit-129-df"],"referenceName":"GCF\_000154425.1\_ASM15442v1","order":[0,1,2,3,4],"report":[["Genome statistics",[{"metricName":"Genome fraction (%)","quality":"More is better","values":["0.040","0.022","0.076","0.255","0.132"],"isMain":true},{"metricName":"Duplication ratio","quality":"Less is better","values":["1.379","1.000","1.337","1.035","1.142"],"isMain":true},{"metricName":"Largest alignment","quality":"More is better","values":[542,678,586,4445,1400],"isMain":true},{"metricName":"Total aligned length","quality":"More is better","values":[1256,678,2370,7910,4325],"isMain":true},{"metricName":"NA50","quality":"More is better","values":[541,678,null,4445,1099],"isMain":false},{"metricName":"NA75","quality":"More is better","values":[null,678,null,599,511],"isMain":false},{"metricName":"NGA50","quality":"More is better","values":[null,null,null,null,null],"isMain":true},{"metricName":"LA50","quality":"Less is better","values":[2,1,null,1,2],"isMain":false},{"metricName":"LA75","quality":"Less is better","values":[null,1,null,4,4],"isMain":false}]],["Reads mapping",[]],["Misassemblies",[{"metricName":"# misassemblies","quality":"Less is better","values":[0,0,0,0,0],"isMain":true},{"metricName":" # relocations","quality":"Less is better","values":[0,0,0,0,0],"isMain":false},{"metricName":" # translocations","quality":"Less is better","values":[0,0,0,0,0],"isMain":false},{"metricName":" # inversions","quality":"Less is better","values":[0,0,0,0,0],"isMain":false},{"metricName":" # interspecies translocations","quality":"Less is better","values":[1,0,6,1,1],"isMain":false},{"metricName":"# misassembled contigs","quality":"Less is better","values":[0,0,0,0,0],"isMain":false},{"metricName":"Misassembled contigs length","quality":"Less is better","values":[0,0,0,0,0],"isMain":true},{"metricName":"# possibly misassembled contigs","quality":"Less is better","values":[0,0,0,0,0],"isMain":false},{"metricName":" # possible misassemblies","quality":"Less is better","values":[0,0,0,0,0],"isMain":false},{"metricName":"# local misassemblies","quality":"Less is better","values":[0,0,0,0,0],"isMain":false},{"metricName":"# scaffold gap ext. mis.","quality":"Less is better","values":[0,0,0,0,0],"isMain":false},{"metricName":"# scaffold gap loc. mis.","quality":"Less is better","values":[0,0,0,0,0],"isMain":false},{"metricName":"# unaligned mis. contigs","quality":"Less is better","values":[0,0,0,0,0],"isMain":false}]],["Unaligned",[{"metricName":"# fully unaligned contigs","quality":"Less is better","values":[0,0,1,0,0],"isMain":false},{"metricName":"Fully unaligned length","quality":"Less is better","values":[0,0,569,0,0],"isMain":false},{"metricName":"# partially unaligned contigs","quality":"Less is better","values":[0,0,3,0,0],"isMain":false},{"metricName":"Partially unaligned length","quality":"Less is better","values":[0,0,8852,0,0],"isMain":false}]],["Mismatches",[{"metricName":"# mismatches","quality":"Less is better","values":[18,14,41,179,121],"isMain":false},{"metricName":"# indels","quality":"Less is better","values":[0,0,1,2,3],"isMain":false},{"metricName":"Indels length","quality":"Less is better","values":[0,0,1,3,3],"isMain":false},{"metricName":"# mismatches per 100 kbp","quality":"Less is better","values":["1433.12","2064.90","1729.23","2263.24","2965.69"],"isMain":true},{"metricName":"# indels per 100 kbp","quality":"Less is better","values":["0.00","0.00","42.18","25.29","73.53"],"isMain":true},{"metricName":" # indels (<= 5 bp)","quality":"Less is better","values":[0,0,1,2,3],"isMain":false},{"metricName":" # indels (> 5 bp)","quality":"Less is better","values":[0,0,0,0,0],"isMain":false},{"metricName":"# N's","quality":"Less is better","values":[0,0,0,0,0],"isMain":false},{"metricName":"# N's per 100 kbp","quality":"Less is better","values":["0.00","0.00","0.00","0.00","0.00"],"isMain":true}]],["Statistics without reference",[{"metricName":"# contigs","quality":"Equal","values":[3,1,8,7,6],"isMain":true},{"metricName":"# contigs (>= 1000 bp)","quality":"Equal","values":[0,0,3,1,2],"isMain":false},{"metricName":"# contigs (>= 5000 bp)","quality":"Equal","values":[0,0,0,0,0],"isMain":false},{"metricName":"# contigs (>= 10000 bp)","quality":"Equal","values":[0,0,0,0,0],"isMain":false},{"metricName":"# contigs (>= 25000 bp)","quality":"Equal","values":[0,0,0,0,0],"isMain":false},{"metricName":"# contigs (>= 50000 bp)","quality":"Equal","values":[0,0,0,0,0],"isMain":false},{"metricName":"Largest contig","quality":"More is better","values":[649,678,4161,4445,1400],"isMain":true},{"metricName":"Total length","quality":"More is better","values":[1732,678,12592,8182,4658],"isMain":true},{"metricName":"Total length (>= 1000 bp)","quality":"More is better","values":[0,0,9795,4445,2499],"isMain":true},{"metricName":"Total length (>= 5000 bp)","quality":"More is better","values":[0,0,0,0,0],"isMain":false},{"metricName":"Total length (>= 10000 bp)","quality":"More is better","values":[0,0,0,0,0],"isMain":true},{"metricName":"Total length (>= 25000 bp)","quality":"More is better","values":[0,0,0,0,0],"isMain":false},{"metricName":"Total length (>= 50000 bp)","quality":"More is better","values":[0,0,0,0,0],"isMain":true},{"metricName":"N50","quality":"More is better","values":[542,678,3544,4445,1099],"isMain":false},{"metricName":"N75","quality":"More is better","values":[541,678,2090,599,559],"isMain":false},{"metricName":"L50","quality":"Less is better","values":[2,1,2,1,2],"isMain":false},{"metricName":"L75","quality":"Less is better","values":[3,1,3,4,4],"isMain":false},{"metricName":"GC (%)","quality":"Equal","values":["48.44","34.81","44.93","52.49","52.00"],"isMain":false}]],["K-mer-based statistics",[]],["Predicted genes",[]],["Similarity statistics",[]],["Reference statistics",[{"metricName":"Reference length","quality":"Equal","values":[3102087,3102087,3102087,3102087,3102087],"isMain":false},{"metricName":"Reference fragments","quality":"Equal","values":[32,32,32,32,32],"isMain":false},{"metricName":"Reference GC (%)","quality":"Equal","values":["43.09","43.09","43.09","43.09","43.09"],"isMain":false}]]],"subreferences":[],"subreports":[],"minContig":500},{"date":"10 February 2020, Monday, 18:00:57","assembliesNames":["megahit-103-df","megahit-113-df","megahit-113-ml","megahit-113-ms","megahit-129-df","metaSPAdes"],"referenceName":"GCF\_000154465.1\_ASM15446v1","order":[0,1,2,3,4,5],"report":[["Genome statistics",[{"metricName":"Genome fraction (%)","quality":"More is better","values":["0.634","0.334","0.475","0.741","0.533","0.453"],"isMain":true},{"metricName":"Duplication ratio","quality":"Less is better","values":["1.081","1.199","1.161","1.088","1.073","1.035"],"isMain":true},{"metricName":"Largest alignment","quality":"More is better","values":[2660,1964,3254,2363,2392,2117],"isMain":true},{"metricName":"Total aligned length","quality":"More is better","values":[16158,8511,12633,19130,13694,11768],"isMain":true},{"metricName":"NA50","quality":"More is better","values":[917,576,567,561,752,731],"isMain":false},{"metricName":"NA75","quality":"More is better","values":[545,246,null,null,559,602],"isMain":false},{"metricName":"NGA50","quality":"More is better","values":[null,null,null,null,null,null],"isMain":true},{"metricName":"LA50","quality":"Less is better","values":[6,7,9,16,7,6],"isMain":false},{"metricName":"LA75","quality":"Less is better","values":[13,12,null,null,13,10],"isMain":false}]],["Reads mapping",[]],["Misassemblies",[{"metricName":"# misassemblies","quality":"Less is better","values":[0,0,0,0,1,0],"isMain":true},{"metricName":" # relocations","quality":"Less is better","values":[0,0,0,0,1,0],"isMain":false},{"metricName":" # translocations","quality":"Less is better","values":[0,0,0,0,0,0],"isMain":false},{"metricName":" # inversions","quality":"Less is better","values":[0,0,0,0,0,0],"isMain":false},{"metricName":" # interspecies translocations","quality":"Less is better","values":[4,6,7,12,3,0],"isMain":false},{"metricName":"# misassembled contigs","quality":"Less is better","values":[0,0,0,0,1,0],"isMain":false},{"metricName":"Misassembled contigs length","quality":"Less is better","values":[0,0,0,0,528,0],"isMain":true},{"metricName":"# possibly misassembled contigs","quality":"Less is better","values":[0,0,0,0,0,0],"isMain":false},{"metricName":" # possible misassemblies","quality":"Less is better","values":[0,0,0,0,0,0],"isMain":false},{"metricName":"# local misassemblies","quality":"Less is better","values":[0,0,0,0,0,0],"isMain":false},{"metricName":"# scaffold gap ext. mis.","quality":"Less is better","values":[0,0,0,0,0,0],"isMain":false},{"metricName":"# scaffold gap loc. mis.","quality":"Less is better","values":[0,0,0,0,0,0],"isMain":false},{"metricName":"# unaligned mis. contigs","quality":"Less is better","values":[0,0,0,0,0,0],"isMain":false}]],["Unaligned",[{"metricName":"# fully unaligned contigs","quality":"Less is better","values":[0,0,0,0,0,0],"isMain":false},{"metricName":"Fully unaligned length","quality":"Less is better","values":[0,0,0,0,0,0],"isMain":false},{"metricName":"# partially unaligned contigs","quality":"Less is better","values":[1,1,4,8,1,0],"isMain":false},{"metricName":"Partially unaligned length","quality":"Less is better","values":[678,678,5394,6011,1404,0],"isMain":false}]],["Mismatches",[{"metricName":"# mismatches","quality":"Less is better","values":[155,160,216,421,202,151],"isMain":false},{"metricName":"# indels","quality":"Less is better","values":[0,2,0,1,0,1],"isMain":false},{"metricName":"Indels length","quality":"Less is better","values":[0,2,0,1,0,1],"isMain":false},{"metricName":"# mismatches per 100 kbp","quality":"Less is better","values":["959.28","1879.48","1782.47","2227.87","1487.04","1307.02"],"isMain":true},{"metricName":"# indels per 100 kbp","quality":"Less is better","values":["0.00","23.49","0.00","5.29","0.00","8.66"],"isMain":true},{"metricName":" # indels (<= 5 bp)","quality":"Less is better","values":[0,2,0,1,0,1],"isMain":false},{"metricName":" # indels (> 5 bp)","quality":"Less is better","values":[0,0,0,0,0,0],"isMain":false},{"metricName":"# N's","quality":"Less is better","values":[0,0,0,0,0,0],"isMain":false},{"metricName":"# N's per 100 kbp","quality":"Less is better","values":["0.00","0.00","0.00","0.00","0.00","0.00"],"isMain":true}]],["Statistics without reference",[{"metricName":"# contigs","quality":"Equal","values":[20,14,19,34,18,15],"isMain":true},{"metricName":"# contigs (>= 1000 bp)","quality":"Equal","values":[5,2,6,5,4,2],"isMain":false},{"metricName":"# contigs (>= 5000 bp)","quality":"Equal","values":[0,0,0,0,0,0],"isMain":false},{"metricName":"# contigs (>= 10000 bp)","quality":"Equal","values":[0,0,0,0,0,0],"isMain":false},{"metricName":"# contigs (>= 25000 bp)","quality":"Equal","values":[0,0,0,0,0,0],"isMain":false},{"metricName":"# contigs (>= 50000 bp)","quality":"Equal","values":[0,0,0,0,0,0],"isMain":false},{"metricName":"Largest contig","quality":"More is better","values":[2660,1981,3271,2380,2392,2117],"isMain":true},{"metricName":"Total length","quality":"More is better","values":[18151,10888,19459,26575,15975,11961],"isMain":true},{"metricName":"Total length (>= 1000 bp)","quality":"More is better","values":[8279,3038,11434,7853,6591,3251],"isMain":true},{"metricName":"Total length (>= 5000 bp)","quality":"More is better","values":[0,0,0,0,0,0],"isMain":false},{"metricName":"Total length (>= 10000 bp)","quality":"More is better","values":[0,0,0,0,0,0],"isMain":true},{"metricName":"Total length (>= 25000 bp)","quality":"More is better","values":[0,0,0,0,0,0],"isMain":false},{"metricName":"Total length (>= 50000 bp)","quality":"More is better","values":[0,0,0,0,0,0],"isMain":true},{"metricName":"N50","quality":"More is better","values":[948,734,1438,719,793,747],"isMain":false},{"metricName":"N75","quality":"More is better","values":[615,586,636,605,626,603],"isMain":false},{"metricName":"L50","quality":"Less is better","values":[6,6,5,12,6,6],"isMain":false},{"metricName":"L75","quality":"Less is better","values":[12,10,11,22,12,10],"isMain":false},{"metricName":"GC (%)","quality":"Equal","values":["48.80","46.79","49.28","48.90","51.35","49.09"],"isMain":false}]],["K-mer-based statistics",[]],["Predicted genes",[]],["Similarity statistics",[]],["Reference statistics",[{"metricName":"Reference length","quality":"Equal","values":[2549878,2549878,2549878,2549878,2549878,2549878],"isMain":false},{"metricName":"Reference fragments","quality":"Equal","values":[20,20,20,20,20,20],"isMain":false},{"metricName":"Reference GC (%)","quality":"Equal","values":["53.27","53.27","53.27","53.27","53.27","53.27"],"isMain":false}]]],"subreferences":[],"subreports":[],"minContig":500},{"date":"10 February 2020, Monday, 18:00:20","assembliesNames":["megahit-103-df","megahit-113-df","megahit-113-ml","megahit-113-ms","megahit-129-df","metaSPAdes"],"referenceName":"GCF\_000154505.1\_ASM15450v1","order":[0,1,2,3,4,5],"report":[["Genome statistics",[{"metricName":"Genome fraction (%)","quality":"More is better","values":["1.325","1.926","2.236","2.058","2.090","0.866"],"isMain":true},{"metricName":"Duplication ratio","quality":"Less is better","values":["1.104","1.065","1.021","1.046","1.054","1.077"],"isMain":true},{"metricName":"Largest alignment","quality":"More is better","values":[990,1984,2036,1015,2215,1000],"isMain":true},{"metricName":"Total aligned length","quality":"More is better","values":[48104,69978,80946,74699,77060,31467],"isMain":true},{"metricName":"NA50","quality":"More is better","values":[574,562,583,588,576,593],"isMain":false},{"metricName":"NA75","quality":"More is better","values":[506,509,530,520,517,522],"isMain":false},{"metricName":"NGA50","quality":"More is better","values":[null,null,null,null,null,null],"isMain":true},{"metricName":"LA50","quality":"Less is better","values":[41,56,59,59,61,25],"isMain":false},{"metricName":"LA75","quality":"Less is better","values":[67,94,96,95,101,41],"isMain":false}]],["Reads mapping",[]],["Misassemblies",[{"metricName":"# misassemblies","quality":"Less is better","values":[2,2,0,0,2,0],"isMain":true},{"metricName":" # relocations","quality":"Less is better","values":[0,0,0,0,0,0],"isMain":false},{"metricName":" # translocations","quality":"Less is better","values":[2,2,0,0,1,0],"isMain":false},{"metricName":" # inversions","quality":"Less is better","values":[0,0,0,0,1,0],"isMain":false},{"metricName":" # interspecies translocations","quality":"Less is better","values":[18,17,5,10,12,8],"isMain":false},{"metricName":"# misassembled contigs","quality":"Less is better","values":[2,2,0,0,2,0],"isMain":false},{"metricName":"Misassembled contigs length","quality":"Less is better","values":[1164,1170,0,0,1265,0],"isMain":true},{"metricName":"# possibly misassembled contigs","quality":"Less is better","values":[0,0,0,0,0,0],"isMain":false},{"metricName":" # possible misassemblies","quality":"Less is better","values":[0,0,0,0,0,0],"isMain":false},{"metricName":"# local misassemblies","quality":"Less is better","values":[0,0,0,1,1,0],"isMain":false},{"metricName":"# scaffold gap ext. mis.","quality":"Less is better","values":[0,0,0,0,0,0],"isMain":false},{"metricName":"# scaffold gap loc. mis.","quality":"Less is better","values":[0,0,0,0,0,0],"isMain":false},{"metricName":"# unaligned mis. contigs","quality":"Less is better","values":[0,0,0,0,0,0],"isMain":false}]],["Unaligned",[{"metricName":"# fully unaligned contigs","quality":"Less is better","values":[0,0,0,0,0,0],"isMain":false},{"metricName":"Fully unaligned length","quality":"Less is better","values":[0,0,0,0,0,0],"isMain":false},{"metricName":"# partially unaligned contigs","quality":"Less is better","values":[4,6,2,3,6,2],"isMain":false},{"metricName":"Partially unaligned length","quality":"Less is better","values":[2793,5766,1080,2336,6748,1218],"isMain":false}]],["Mismatches",[{"metricName":"# mismatches","quality":"Less is better","values":[779,1122,1662,1651,1386,521],"isMain":false},{"metricName":"# indels","quality":"Less is better","values":[1,2,4,2,13,1],"isMain":false},{"metricName":"Indels length","quality":"Less is better","values":[1,2,5,2,15,1],"isMain":false},{"metricName":"# mismatches per 100 kbp","quality":"Less is better","values":["1624.20","1609.18","2053.22","2216.14","1832.29","1662.57"],"isMain":true},{"metricName":"# indels per 100 kbp","quality":"Less is better","values":["2.08","2.87","4.94","2.68","17.19","3.19"],"isMain":true},{"metricName":" # indels (<= 5 bp)","quality":"Less is better","values":[1,2,4,2,13,1],"isMain":false},{"metricName":" # indels (> 5 bp)","quality":"Less is better","values":[0,0,0,0,0,0],"isMain":false},{"metricName":"# N's","quality":"Less is better","values":[0,0,0,0,0,0],"isMain":false},{"metricName":"# N's per 100 kbp","quality":"Less is better","values":["0.00","0.00","0.00","0.00","0.00","0.00"],"isMain":true}]],["Statistics without reference",[{"metricName":"# contigs","quality":"Equal","values":[87,121,134,129,130,55],"isMain":true},{"metricName":"# contigs (>= 1000 bp)","quality":"Equal","values":[2,8,1,2,8,2],"isMain":false},{"metricName":"# contigs (>= 5000 bp)","quality":"Equal","values":[0,0,0,0,0,0],"isMain":false},{"metricName":"# contigs (>= 10000 bp)","quality":"Equal","values":[0,0,0,0,0,0],"isMain":false},{"metricName":"# contigs (>= 25000 bp)","quality":"Equal","values":[0,0,0,0,0,0],"isMain":false},{"metricName":"# contigs (>= 50000 bp)","quality":"Equal","values":[0,0,0,0,0,0],"isMain":false},{"metricName":"Largest contig","quality":"More is better","values":[1136,1984,2036,1318,2238,1059],"isMain":true},{"metricName":"Total length","quality":"More is better","values":[55732,80023,83688,80272,86491,34981],"isMain":true},{"metricName":"Total length (>= 1000 bp)","quality":"More is better","values":[2229,10775,2036,2436,11293,2062],"isMain":true},{"metricName":"Total length (>= 5000 bp)","quality":"More is better","values":[0,0,0,0,0,0],"isMain":false},{"metricName":"Total length (>= 10000 bp)","quality":"More is better","values":[0,0,0,0,0,0],"isMain":true},{"metricName":"Total length (>= 25000 bp)","quality":"More is better","values":[0,0,0,0,0,0],"isMain":false},{"metricName":"Total length (>= 50000 bp)","quality":"More is better","values":[0,0,0,0,0,0],"isMain":true},{"metricName":"N50","quality":"More is better","values":[628,620,595,608,625,617],"isMain":false},{"metricName":"N75","quality":"More is better","values":[563,543,551,541,547,556],"isMain":false},{"metricName":"L50","quality":"Less is better","values":[38,48,58,56,52,24],"isMain":false},{"metricName":"L75","quality":"Less is better","values":[61,83,94,91,89,39],"isMain":false},{"metricName":"GC (%)","quality":"Equal","values":["46.60","46.74","46.62","45.76","47.36","45.17"],"isMain":false}]],["K-mer-based statistics",[]],["Predicted genes",[]],["Similarity statistics",[]],["Reference statistics",[{"metricName":"Reference length","quality":"Equal","values":[3619905,3619905,3619905,3619905,3619905,3619905],"isMain":false},{"metricName":"Reference fragments","quality":"Equal","values":[68,68,68,68,68,68],"isMain":false},{"metricName":"Reference GC (%)","quality":"Equal","values":["46.35","46.35","46.35","46.35","46.35","46.35"],"isMain":false}]]],"subreferences":[],"subreports":[],"minContig":500},{"date":"10 February 2020, Monday, 18:00:17","assembliesNames":["megahit-103-df","megahit-113-df","megahit-113-ml","megahit-113-ms","megahit-129-df","metaSPAdes"],"referenceName":"GCF\_000155815.1\_ASM15581v1","order":[0,1,2,3,4,5],"report":[["Genome statistics",[{"metricName":"Genome fraction (%)","quality":"More is better","values":["3.986","5.023","6.036","6.866","5.663","2.301"],"isMain":true},{"metricName":"Duplication ratio","quality":"Less is better","values":["1.047","1.046","1.034","1.047","1.046","1.052"],"isMain":true},{"metricName":"Largest alignment","quality":"More is better","values":[11666,2433,2615,1955,4496,10456],"isMain":true},{"metricName":"Total aligned length","quality":"More is better","values":[166195,209554,251957,286297,236132,95949],"isMain":true},{"metricName":"NA50","quality":"More is better","values":[596,582,593,598,586,596],"isMain":false},{"metricName":"NA75","quality":"More is better","values":[521,509,528,519,512,515],"isMain":false},{"metricName":"NGA50","quality":"More is better","values":[null,null,null,null,null,null],"isMain":true},{"metricName":"LA50","quality":"Less is better","values":[109,161,179,211,174,58],"isMain":false},{"metricName":"LA75","quality":"Less is better","values":[190,272,299,353,297,106],"isMain":false}]],["Reads mapping",[]],["Misassemblies",[{"metricName":"# misassemblies","quality":"Less is better","values":[1,2,0,1,0,0],"isMain":true},{"metricName":" # relocations","quality":"Less is better","values":[0,0,0,0,0,0],"isMain":false},{"metricName":" # translocations","quality":"Less is better","values":[1,2,0,1,0,0],"isMain":false},{"metricName":" # inversions","quality":"Less is better","values":[0,0,0,0,0,0],"isMain":false},{"metricName":" # interspecies translocations","quality":"Less is better","values":[31,40,18,39,45,21],"isMain":false},{"metricName":"# misassembled contigs","quality":"Less is better","values":[1,2,0,1,0,0],"isMain":false},{"metricName":"Misassembled contigs length","quality":"Less is better","values":[611,1140,0,600,0,0],"isMain":true},{"metricName":"# possibly misassembled contigs","quality":"Less is better","values":[0,0,0,1,0,0],"isMain":false},{"metricName":" # possible misassemblies","quality":"Less is better","values":[0,0,0,1,0,0],"isMain":false},{"metricName":"# local misassemblies","quality":"Less is better","values":[0,0,1,1,0,0],"isMain":false},{"metricName":"# scaffold gap ext. mis.","quality":"Less is better","values":[0,0,0,0,0,0],"isMain":false},{"metricName":"# scaffold gap loc. mis.","quality":"Less is better","values":[0,0,0,0,0,0],"isMain":false},{"metricName":"# unaligned mis. contigs","quality":"Less is better","values":[0,0,0,0,2,0],"isMain":false}]],["Unaligned",[{"metricName":"# fully unaligned contigs","quality":"Less is better","values":[1,2,1,1,1,0],"isMain":false},{"metricName":"Fully unaligned length","quality":"Less is better","values":[769,1307,1011,910,744,0],"isMain":false},{"metricName":"# partially unaligned contigs","quality":"Less is better","values":[5,13,6,13,18,6],"isMain":false},{"metricName":"Partially unaligned length","quality":"Less is better","values":[4924,20837,6078,15803,20979,5195],"isMain":false}]],["Mismatches",[{"metricName":"# mismatches","quality":"Less is better","values":[2714,3589,5381,6070,4219,1572],"isMain":false},{"metricName":"# indels","quality":"Less is better","values":[6,8,16,13,18,3],"isMain":false},{"metricName":"Indels length","quality":"Less is better","values":[6,26,39,15,33,3],"isMain":false},{"metricName":"# mismatches per 100 kbp","quality":"Less is better","values":["1637.39","1718.52","2143.99","2126.04","1791.88","1642.86"],"isMain":true},{"metricName":"# indels per 100 kbp","quality":"Less is better","values":["3.62","3.83","6.37","4.55","7.64","3.14"],"isMain":true},{"metricName":" # indels (<= 5 bp)","quality":"Less is better","values":[6,6,15,13,17,3],"isMain":false},{"metricName":" # indels (> 5 bp)","quality":"Less is better","values":[0,2,1,0,1,0],"isMain":false},{"metricName":"# N's","quality":"Less is better","values":[0,0,0,0,0,0],"isMain":false},{"metricName":"# N's per 100 kbp","quality":"Less is better","values":["0.00","0.00","0.00","0.00","0.00","0.00"],"isMain":true}]],["Statistics without reference",[{"metricName":"# contigs","quality":"Equal","values":[263,345,409,468,383,147],"isMain":true},{"metricName":"# contigs (>= 1000 bp)","quality":"Equal","values":[11,23,21,30,25,9],"isMain":false},{"metricName":"# contigs (>= 5000 bp)","quality":"Equal","values":[1,1,0,0,0,1],"isMain":false},{"metricName":"# contigs (>= 10000 bp)","quality":"Equal","values":[1,0,0,0,0,1],"isMain":false},{"metricName":"# contigs (>= 25000 bp)","quality":"Equal","values":[0,0,0,0,0,0],"isMain":false},{"metricName":"# contigs (>= 50000 bp)","quality":"Equal","values":[0,0,0,0,0,0],"isMain":false},{"metricName":"Largest contig","quality":"More is better","values":[11666,6789,2624,4215,4821,10456],"isMain":true},{"metricName":"Total length","quality":"More is better","values":[179286,240566,266541,315696,268045,105850],"isMain":true},{"metricName":"Total length (>= 1000 bp)","quality":"More is better","values":[25120,40375,28414,42473,43686,21008],"isMain":true},{"metricName":"Total length (>= 5000 bp)","quality":"More is better","values":[11666,6789,0,0,0,10456],"isMain":false},{"metricName":"Total length (>= 10000 bp)","quality":"More is better","values":[11666,0,0,0,0,10456],"isMain":true},{"metricName":"Total length (>= 25000 bp)","quality":"More is better","values":[0,0,0,0,0,0],"isMain":false},{"metricName":"Total length (>= 50000 bp)","quality":"More is better","values":[0,0,0,0,0,0],"isMain":true},{"metricName":"N50","quality":"More is better","values":[626,639,621,635,648,653],"isMain":false},{"metricName":"N75","quality":"More is better","values":[549,555,549,553,564,548],"isMain":false},{"metricName":"L50","quality":"Less is better","values":[101,131,167,186,145,52],"isMain":false},{"metricName":"L75","quality":"Less is better","values":[178,231,282,318,256,97],"isMain":false},{"metricName":"GC (%)","quality":"Equal","values":["45.67","45.07","45.25","45.08","45.10","44.67"],"isMain":false}]],["K-mer-based statistics",[]],["Predicted genes",[]],["Similarity statistics",[]],["Reference statistics",[{"metricName":"Reference length","quality":"Equal","values":[4157980,4157980,4157980,4157980,4157980,4157980],"isMain":false},{"metricName":"Reference fragments","quality":"Equal","values":[70,70,70,70,70,70],"isMain":false},{"metricName":"Reference GC (%)","quality":"Equal","values":["44.58","44.58","44.58","44.58","44.58","44.58"],"isMain":false}]]],"subreferences":[],"subreports":[],"minContig":500},{"date":"10 February 2020, Monday, 18:00:52","assembliesNames":["megahit-103-df","megahit-113-df","megahit-113-ml","megahit-113-ms","megahit-129-df","metaSPAdes"],"referenceName":"GCF\_000155915.2\_Lacto\_jensenii\_1153\_V2","order":[0,1,2,3,4,5],"report":[["Genome statistics",[{"metricName":"Genome fraction (%)","quality":"More is better","values":["7.891","8.708","11.594","10.540","9.563","2.902"],"isMain":true},{"metricName":"Duplication ratio","quality":"Less is better","values":["1.015","1.027","1.030","1.039","1.025","1.029"],"isMain":true},{"metricName":"Largest alignment","quality":"More is better","values":[1628,1775,1686,1848,1468,1266],"isMain":true},{"metricName":"Total aligned length","quality":"More is better","values":[137602,152331,202890,186385,166874,50693],"isMain":true},{"metricName":"NA50","quality":"More is better","values":[611,636,653,652,648,586],"isMain":false},{"metricName":"NA75","quality":"More is better","values":[518,528,553,541,545,532],"isMain":false},{"metricName":"NGA50","quality":"More is better","values":[null,null,null,null,null,null],"isMain":true},{"metricName":"LA50","quality":"Less is better","values":[103,101,129,117,107,37],"isMain":false},{"metricName":"LA75","quality":"Less is better","values":[175,176,218,200,183,60],"isMain":false}]],["Reads mapping",[]],["Misassemblies",[{"metricName":"# misassemblies","quality":"Less is better","values":[2,1,0,1,2,0],"isMain":true},{"metricName":" # relocations","quality":"Less is better","values":[0,0,0,0,1,0],"isMain":false},{"metricName":" # translocations","quality":"Less is better","values":[1,1,0,1,1,0],"isMain":false},{"metricName":" # inversions","quality":"Less is better","values":[1,0,0,0,0,0],"isMain":false},{"metricName":" # interspecies translocations","quality":"Less is better","values":[6,11,7,13,9,3],"isMain":false},{"metricName":"# misassembled contigs","quality":"Less is better","values":[2,1,0,1,2,0],"isMain":false},{"metricName":"Misassembled contigs length","quality":"Less is better","values":[2019,744,0,889,1960,0],"isMain":true},{"metricName":"# possibly misassembled contigs","quality":"Less is better","values":[0,0,0,0,0,0],"isMain":false},{"metricName":" # possible misassemblies","quality":"Less is better","values":[0,0,0,0,0,0],"isMain":false},{"metricName":"# local misassemblies","quality":"Less is better","values":[0,0,1,1,0,0],"isMain":false},{"metricName":"# scaffold gap ext. mis.","quality":"Less is better","values":[0,0,0,0,0,0],"isMain":false},{"metricName":"# scaffold gap loc. mis.","quality":"Less is better","values":[0,0,0,0,0,0],"isMain":false},{"metricName":"# unaligned mis. contigs","quality":"Less is better","values":[2,1,2,1,1,0],"isMain":false}]],["Unaligned",[{"metricName":"# fully unaligned contigs","quality":"Less is better","values":[0,0,1,1,1,0],"isMain":false},{"metricName":"Fully unaligned length","quality":"Less is better","values":[0,0,515,608,730,0],"isMain":false},{"metricName":"# partially unaligned contigs","quality":"Less is better","values":[3,6,4,4,3,1],"isMain":false},{"metricName":"Partially unaligned length","quality":"Less is better","values":[19971,18317,5308,5274,6800,1161],"isMain":false}]],["Mismatches",[{"metricName":"# mismatches","quality":"Less is better","values":[2022,2205,3753,3555,2542,839],"isMain":false},{"metricName":"# indels","quality":"Less is better","values":[7,9,26,19,18,3],"isMain":false},{"metricName":"Indels length","quality":"Less is better","values":[7,36,29,19,31,3],"isMain":false},{"metricName":"# mismatches per 100 kbp","quality":"Less is better","values":["1474.41","1456.98","1862.54","1940.87","1529.59","1663.40"],"isMain":true},{"metricName":"# indels per 100 kbp","quality":"Less is better","values":["5.10","5.95","12.90","10.37","10.83","5.95"],"isMain":true},{"metricName":" # indels (<= 5 bp)","quality":"Less is better","values":[7,8,26,19,17,3],"isMain":false},{"metricName":" # indels (> 5 bp)","quality":"Less is better","values":[0,1,0,0,1,0],"isMain":false},{"metricName":"# N's","quality":"Less is better","values":[0,0,0,0,0,0],"isMain":false},{"metricName":"# N's per 100 kbp","quality":"Less is better","values":["0.00","0.00","0.00","0.00","0.00","0.00"],"isMain":true}]],["Statistics without reference",[{"metricName":"# contigs","quality":"Equal","values":[210,222,306,280,250,83],"isMain":true},{"metricName":"# contigs (>= 1000 bp)","quality":"Equal","values":[13,24,25,21,22,4],"isMain":false},{"metricName":"# contigs (>= 5000 bp)","quality":"Equal","values":[1,1,0,0,0,0],"isMain":false},{"metricName":"# contigs (>= 10000 bp)","quality":"Equal","values":[1,0,0,0,0,0],"isMain":false},{"metricName":"# contigs (>= 25000 bp)","quality":"Equal","values":[0,0,0,0,0,0],"isMain":false},{"metricName":"# contigs (>= 50000 bp)","quality":"Equal","values":[0,0,0,0,0,0],"isMain":false},{"metricName":"Largest contig","quality":"More is better","values":[14263,8883,3322,3686,4654,1394],"isMain":true},{"metricName":"Total length","quality":"More is better","values":[159129,173706,213391,196271,177939,53075],"isMain":true},{"metricName":"Total length (>= 1000 bp)","quality":"More is better","values":[34059,45034,31438,28957,29991,4778],"isMain":true},{"metricName":"Total length (>= 5000 bp)","quality":"More is better","values":[14263,8883,0,0,0,0],"isMain":false},{"metricName":"Total length (>= 10000 bp)","quality":"More is better","values":[14263,0,0,0,0,0],"isMain":true},{"metricName":"Total length (>= 25000 bp)","quality":"More is better","values":[0,0,0,0,0,0],"isMain":false},{"metricName":"Total length (>= 50000 bp)","quality":"More is better","values":[0,0,0,0,0,0],"isMain":true},{"metricName":"N50","quality":"More is better","values":[676,741,674,671,691,605],"isMain":false},{"metricName":"N75","quality":"More is better","values":[573,583,578,573,579,552],"isMain":false},{"metricName":"L50","quality":"Less is better","values":[71,75,123,109,96,35],"isMain":false},{"metricName":"L75","quality":"Less is better","values":[135,142,208,188,167,58],"isMain":false},{"metricName":"GC (%)","quality":"Equal","values":["34.46","34.57","35.01","34.20","34.26","34.48"],"isMain":false}]],["K-mer-based statistics",[]],["Predicted genes",[]],["Similarity statistics",[]],["Reference statistics",[{"metricName":"Reference length","quality":"Equal","values":[1737886,1737886,1737886,1737886,1737886,1737886],"isMain":false},{"metricName":"Reference fragments","quality":"Equal","values":[9,9,9,9,9,9],"isMain":false},{"metricName":"Reference GC (%)","quality":"Equal","values":["34.54","34.54","34.54","34.54","34.54","34.54"],"isMain":false}]]],"subreferences":[],"subreports":[],"minContig":500},{"date":"10 February 2020, Monday, 18:01:05","assembliesNames":["megahit-103-df","megahit-113-df","megahit-113-ml","megahit-113-ms","megahit-129-df","metaSPAdes"],"referenceName":"GCF\_000156035.2\_ASM15603v2","order":[0,1,2,3,4,5],"report":[["Genome statistics",[{"metricName":"Genome fraction (%)","quality":"More is better","values":["92.272","94.643","95.114","95.544","94.762","92.615"],"isMain":true},{"metricName":"Duplication ratio","quality":"Less is better","values":["1.021","1.014","1.018","1.025","1.020","1.006"],"isMain":true},{"metricName":"Largest alignment","quality":"More is better","values":[35351,83697,68148,83774,83369,95900],"isMain":true},{"metricName":"Total aligned length","quality":"More is better","values":[3612275,3679438,3710220,3751898,3710261,3581755],"isMain":true},{"metricName":"NG50","quality":"More is better","values":[7820,53070,48201,73936,44449,46199],"isMain":false},{"metricName":"NG75","quality":"More is better","values":[3534,25861,24204,41318,18327,22404],"isMain":false},{"metricName":"NA50","quality":"More is better","values":[6882,20083,20010,14871,20336,22404],"isMain":false},{"metricName":"NA75","quality":"More is better","values":[3526,8198,7204,965,8017,11087],"isMain":false},{"metricName":"NGA50","quality":"More is better","values":[6575,20521,20374,20812,20336,21193],"isMain":true},{"metricName":"NGA75","quality":"More is better","values":[3103,8734,8276,8276,8225,9179],"isMain":false},{"metricName":"LG50","quality":"Less is better","values":[139,23,26,15,25,25],"isMain":false},{"metricName":"LG75","quality":"Less is better","values":[321,48,53,32,57,53],"isMain":false},{"metricName":"LA50","quality":"Less is better","values":[151,57,62,88,57,49],"isMain":false},{"metricName":"LA75","quality":"Less is better","values":[331,131,141,342,131,105],"isMain":false},{"metricName":"LGA50","quality":"Less is better","values":[164,55,58,60,57,53],"isMain":true},{"metricName":"LGA75","quality":"Less is better","values":[372,124,127,130,129,118],"isMain":false}]],["Reads mapping",[]],["Misassemblies",[{"metricName":"# misassemblies","quality":"Less is better","values":[101,149,152,164,157,143],"isMain":true},{"metricName":" # relocations","quality":"Less is better","values":[0,1,2,3,0,6],"isMain":false},{"metricName":" # translocations","quality":"Less is better","values":[100,148,149,161,156,137],"isMain":false},{"metricName":" # inversions","quality":"Less is better","values":[1,0,1,0,1,0],"isMain":false},{"metricName":" # interspecies translocations","quality":"Less is better","values":[102,39,44,74,40,9],"isMain":false},{"metricName":"# misassembled contigs","quality":"Less is better","values":[84,65,78,80,77,65],"isMain":false},{"metricName":"Misassembled contigs length","quality":"Less is better","values":[832284,2605405,2630902,2705688,2630376,2788657],"isMain":true},{"metricName":"# possibly misassembled contigs","quality":"Less is better","values":[0,0,1,1,0,0],"isMain":false},{"metricName":" # possible misassemblies","quality":"Less is better","values":[0,0,1,1,0,0],"isMain":false},{"metricName":"# local misassemblies","quality":"Less is better","values":[1,4,4,6,0,1],"isMain":false},{"metricName":"# scaffold gap ext. mis.","quality":"Less is better","values":[0,0,0,0,0,0],"isMain":false},{"metricName":"# scaffold gap loc. mis.","quality":"Less is better","values":[0,0,0,0,0,0],"isMain":false},{"metricName":"# unaligned mis. contigs","quality":"Less is better","values":[0,2,2,3,5,1],"isMain":false}]],["Unaligned",[{"metricName":"# fully unaligned contigs","quality":"Less is better","values":[1,0,4,2,0,0],"isMain":false},{"metricName":"Fully unaligned length","quality":"Less is better","values":[523,0,3364,1162,0,0],"isMain":false},{"metricName":"# partially unaligned contigs","quality":"Less is better","values":[43,18,18,45,21,6],"isMain":false},{"metricName":"Partially unaligned length","quality":"Less is better","values":[37431,235862,265638,1069920,158072,92947],"isMain":false}]],["Mismatches",[{"metricName":"# mismatches","quality":"Less is better","values":[6387,3104,4888,6449,4186,3846],"isMain":false},{"metricName":"# indels","quality":"Less is better","values":[32,138,167,216,137,154],"isMain":false},{"metricName":"Indels length","quality":"Less is better","values":[34,305,480,681,470,233],"isMain":false},{"metricName":"# mismatches per 100 kbp","quality":"Less is better","values":["179.28","84.94","133.10","174.82","114.41","107.55"],"isMain":true},{"metricName":"# indels per 100 kbp","quality":"Less is better","values":["0.90","3.78","4.55","5.86","3.74","4.31"],"isMain":true},{"metricName":" # indels (<= 5 bp)","quality":"Less is better","values":[32,133,157,204,127,151],"isMain":false},{"metricName":" # indels (> 5 bp)","quality":"Less is better","values":[0,5,10,12,10,3],"isMain":false},{"metricName":"# N's","quality":"Less is better","values":[0,0,0,0,0,0],"isMain":false},{"metricName":"# N's per 100 kbp","quality":"Less is better","values":["0.00","0.00","0.00","0.00","0.00","0.00"],"isMain":true}]],["Statistics without reference",[{"metricName":"# contigs","quality":"Equal","values":[873,302,361,374,355,197],"isMain":true},{"metricName":"# contigs (>= 1000 bp)","quality":"Equal","values":[625,208,224,218,212,167],"isMain":false},{"metricName":"# contigs (>= 5000 bp)","quality":"Equal","values":[237,112,121,132,127,109],"isMain":false},{"metricName":"# contigs (>= 10000 bp)","quality":"Equal","values":[95,79,86,90,84,77],"isMain":false},{"metricName":"# contigs (>= 25000 bp)","quality":"Equal","values":[8,48,49,55,46,46],"isMain":false},{"metricName":"# contigs (>= 50000 bp)","quality":"Equal","values":[0,26,23,23,20,23],"isMain":false},{"metricName":"Largest contig","quality":"More is better","values":[43528,128658,128645,609940,139032,139787],"isMain":true},{"metricName":"Total length","quality":"More is better","values":[3676077,3942603,4005866,4850964,3891397,3691089],"isMain":true},{"metricName":"Total length (>= 1000 bp)","quality":"More is better","values":[3510597,3879966,3916960,4748312,3801779,3669163],"isMain":true},{"metricName":"Total length (>= 5000 bp)","quality":"More is better","values":[2544926,3656070,3687679,4551391,3602369,3526211],"isMain":false},{"metricName":"Total length (>= 10000 bp)","quality":"More is better","values":[1534460,3420818,3443991,4251943,3289425,3313035],"isMain":true},{"metricName":"Total length (>= 25000 bp)","quality":"More is better","values":[254669,2919690,2814461,3660867,2668548,2747464],"isMain":false},{"metricName":"Total length (>= 50000 bp)","quality":"More is better","values":[0,2103957,1828602,2505745,1715838,1875325],"isMain":true},{"metricName":"N50","quality":"More is better","values":[8565,52725,47790,57449,44449,50070],"isMain":false},{"metricName":"N75","quality":"More is better","values":[4053,22886,22962,25522,18261,24249],"isMain":false},{"metricName":"L50","quality":"Less is better","values":[128,24,27,22,25,23],"isMain":false},{"metricName":"L75","quality":"Less is better","values":[284,50,57,55,58,47],"isMain":false},{"metricName":"GC (%)","quality":"Equal","values":["39.96","40.15","40.21","40.57","40.26","40.05"],"isMain":false}]],["K-mer-based statistics",[]],["Predicted genes",[]],["Similarity statistics",[]],["Reference statistics",[{"metricName":"Reference length","quality":"Equal","values":[3861016,3861016,3861016,3861016,3861016,3861016],"isMain":false},{"metricName":"Reference fragments","quality":"Equal","values":[316,316,316,316,316,316],"isMain":false},{"metricName":"Reference GC (%)","quality":"Equal","values":["40.09","40.09","40.09","40.09","40.09","40.09"],"isMain":false}]]],"subreferences":[],"subreports":[],"minContig":500},{"date":"10 February 2020, Monday, 18:00:26","assembliesNames":["megahit-103-df","megahit-113-df","megahit-113-ml","megahit-113-ms","megahit-129-df","metaSPAdes"],"referenceName":"GCF\_000156075.1\_ASM15607v1","order":[0,1,2,3,4,5],"report":[["Genome statistics",[{"metricName":"Genome fraction (%)","quality":"More is better","values":["1.735","2.593","2.944","2.685","1.978","1.236"],"isMain":true},{"metricName":"Duplication ratio","quality":"Less is better","values":["1.023","1.023","1.012","1.027","1.039","1.016"],"isMain":true},{"metricName":"Largest alignment","quality":"More is better","values":[2159,7300,5599,4973,6721,5709],"isMain":true},{"metricName":"Total aligned length","quality":"More is better","values":[95885,142356,162215,148407,110014,68084],"isMain":true},{"metricName":"NA50","quality":"More is better","values":[703,1085,774,722,728,766],"isMain":false},{"metricName":"NA75","quality":"More is better","values":[583,653,589,565,573,648],"isMain":false},{"metricName":"NGA50","quality":"More is better","values":[null,null,null,null,null,null],"isMain":true},{"metricName":"LA50","quality":"Less is better","values":[49,32,69,76,49,23],"isMain":false},{"metricName":"LA75","quality":"Less is better","values":[87,76,132,135,96,48],"isMain":false}]],["Reads mapping",[]],["Misassemblies",[{"metricName":"# misassemblies","quality":"Less is better","values":[0,1,0,1,1,2],"isMain":true},{"metricName":" # relocations","quality":"Less is better","values":[0,0,0,0,0,0],"isMain":false},{"metricName":" # translocations","quality":"Less is better","values":[0,1,0,1,0,2],"isMain":false},{"metricName":" # inversions","quality":"Less is better","values":[0,0,0,0,1,0],"isMain":false},{"metricName":" # interspecies translocations","quality":"Less is better","values":[5,9,2,8,12,3],"isMain":false},{"metricName":"# misassembled contigs","quality":"Less is better","values":[0,1,0,1,1,1],"isMain":false},{"metricName":"Misassembled contigs length","quality":"Less is better","values":[0,1237,0,648,552,2232],"isMain":true},{"metricName":"# possibly misassembled contigs","quality":"Less is better","values":[0,0,0,0,1,0],"isMain":false},{"metricName":" # possible misassemblies","quality":"Less is better","values":[0,0,0,0,2,0],"isMain":false},{"metricName":"# local misassemblies","quality":"Less is better","values":[0,1,0,1,2,0],"isMain":false},{"metricName":"# scaffold gap ext. mis.","quality":"Less is better","values":[0,0,0,0,0,0],"isMain":false},{"metricName":"# scaffold gap loc. mis.","quality":"Less is better","values":[0,0,0,0,0,0],"isMain":false},{"metricName":"# unaligned mis. contigs","quality":"Less is better","values":[0,0,1,0,0,0],"isMain":false}]],["Unaligned",[{"metricName":"# fully unaligned contigs","quality":"Less is better","values":[0,0,0,0,0,0],"isMain":false},{"metricName":"Fully unaligned length","quality":"Less is better","values":[0,0,0,0,0,0],"isMain":false},{"metricName":"# partially unaligned contigs","quality":"Less is better","values":[1,2,2,2,4,1],"isMain":false},{"metricName":"Partially unaligned length","quality":"Less is better","values":[936,1708,2558,1845,4189,858],"isMain":false}]],["Mismatches",[{"metricName":"# mismatches","quality":"Less is better","values":[1543,2626,3442,3316,1874,1130],"isMain":false},{"metricName":"# indels","quality":"Less is better","values":[4,24,19,11,19,14],"isMain":false},{"metricName":"Indels length","quality":"Less is better","values":[4,50,34,12,98,19],"isMain":false},{"metricName":"# mismatches per 100 kbp","quality":"Less is better","values":["1620.41","1845.19","2130.69","2250.33","1726.03","1665.86"],"isMain":true},{"metricName":"# indels per 100 kbp","quality":"Less is better","values":["4.20","16.86","11.76","7.46","17.50","20.64"],"isMain":true},{"metricName":" # indels (<= 5 bp)","quality":"Less is better","values":[4,21,18,11,18,14],"isMain":false},{"metricName":" # indels (> 5 bp)","quality":"Less is better","values":[0,3,1,0,1,0],"isMain":false},{"metricName":"# N's","quality":"Less is better","values":[0,0,0,0,0,0],"isMain":false},{"metricName":"# N's per 100 kbp","quality":"Less is better","values":["0.00","0.00","0.00","0.00","0.00","0.00"],"isMain":true}]],["Statistics without reference",[{"metricName":"# contigs","quality":"Equal","values":[130,135,203,200,142,75],"isMain":true},{"metricName":"# contigs (>= 1000 bp)","quality":"Equal","values":[19,39,31,22,17,14],"isMain":false},{"metricName":"# contigs (>= 5000 bp)","quality":"Equal","values":[0,3,1,0,2,1],"isMain":false},{"metricName":"# contigs (>= 10000 bp)","quality":"Equal","values":[0,0,0,0,0,0],"isMain":false},{"metricName":"# contigs (>= 25000 bp)","quality":"Equal","values":[0,0,0,0,0,0],"isMain":false},{"metricName":"# contigs (>= 50000 bp)","quality":"Equal","values":[0,0,0,0,0,0],"isMain":false},{"metricName":"Largest contig","quality":"More is better","values":[2160,7305,5599,4973,6721,5709],"isMain":true},{"metricName":"Total length","quality":"More is better","values":[98340,147259,166050,153152,117048,69782],"isMain":true},{"metricName":"Total length (>= 1000 bp)","quality":"More is better","values":[25607,83036,52297,34250,34581,29357],"isMain":true},{"metricName":"Total length (>= 5000 bp)","quality":"More is better","values":[0,19201,5599,0,12621,5709],"isMain":false},{"metricName":"Total length (>= 10000 bp)","quality":"More is better","values":[0,0,0,0,0,0],"isMain":true},{"metricName":"Total length (>= 25000 bp)","quality":"More is better","values":[0,0,0,0,0,0],"isMain":false},{"metricName":"Total length (>= 50000 bp)","quality":"More is better","values":[0,0,0,0,0,0],"isMain":true},{"metricName":"N50","quality":"More is better","values":[733,1152,791,728,770,780],"isMain":false},{"metricName":"N75","quality":"More is better","values":[605,678,604,604,599,661],"isMain":false},{"metricName":"L50","quality":"Less is better","values":[48,31,66,73,45,21],"isMain":false},{"metricName":"L75","quality":"Less is better","values":[85,73,127,130,89,45],"isMain":false},{"metricName":"GC (%)","quality":"Equal","values":["41.01","41.12","41.37","41.56","40.97","40.96"],"isMain":false}]],["K-mer-based statistics",[]],["Predicted genes",[]],["Similarity statistics",[]],["Reference statistics",[{"metricName":"Reference length","quality":"Equal","values":[5487733,5487733,5487733,5487733,5487733,5487733],"isMain":false},{"metricName":"Reference fragments","quality":"Equal","values":[132,132,132,132,132,132],"isMain":false},{"metricName":"Reference GC (%)","quality":"Equal","values":["41.99","41.99","41.99","41.99","41.99","41.99"],"isMain":false}]]],"subreferences":[],"subreports":[],"minContig":500},{"date":"10 February 2020, Monday, 18:00:26","assembliesNames":["megahit-103-df","megahit-113-df","megahit-113-ml","megahit-113-ms","megahit-129-df","metaSPAdes"],"referenceName":"GCF\_000156175.1\_ASM15617v1","order":[0,1,2,3,4,5],"report":[["Genome statistics",[{"metricName":"Genome fraction (%)","quality":"More is better","values":["56.609","62.993","57.152","59.356","65.471","45.536"],"isMain":true},{"metricName":"Duplication ratio","quality":"Less is better","values":["1.033","1.028","1.047","1.050","1.032","1.037"],"isMain":true},{"metricName":"Largest alignment","quality":"More is better","values":[3180,4321,2444,3201,3788,2225],"isMain":true},{"metricName":"Total aligned length","quality":"More is better","values":[1046877,1158468,1070607,1111528,1210052,841958],"isMain":true},{"metricName":"NG50","quality":"More is better","values":[579,689,573,599,749,null],"isMain":false},{"metricName":"NA50","quality":"More is better","values":[826,956,776,788,967,745],"isMain":false},{"metricName":"NA75","quality":"More is better","values":[632,677,601,607,687,599],"isMain":false},{"metricName":"NGA50","quality":"More is better","values":[563,668,565,582,709,null],"isMain":true},{"metricName":"LG50","quality":"Less is better","values":[973,792,1038,984,732,null],"isMain":false},{"metricName":"LA50","quality":"Less is better","values":[450,427,503,514,444,427],"isMain":false},{"metricName":"LA75","quality":"Less is better","values":[821,797,900,923,827,751],"isMain":false},{"metricName":"LGA50","quality":"Less is better","values":[997,819,1054,1011,777,null],"isMain":true}]],["Reads mapping",[]],["Misassemblies",[{"metricName":"# misassemblies","quality":"Less is better","values":[6,4,4,4,4,2],"isMain":true},{"metricName":" # relocations","quality":"Less is better","values":[2,1,1,0,1,0],"isMain":false},{"metricName":" # translocations","quality":"Less is better","values":[3,2,1,3,3,2],"isMain":false},{"metricName":" # inversions","quality":"Less is better","values":[1,1,2,1,0,0],"isMain":false},{"metricName":" # interspecies translocations","quality":"Less is better","values":[29,31,3,23,22,16],"isMain":false},{"metricName":"# misassembled contigs","quality":"Less is better","values":[6,4,4,4,4,2],"isMain":false},{"metricName":"Misassembled contigs length","quality":"Less is better","values":[4382,3693,3519,3374,6496,1701],"isMain":true},{"metricName":"# possibly misassembled contigs","quality":"Less is better","values":[0,0,0,1,0,0],"isMain":false},{"metricName":" # possible misassemblies","quality":"Less is better","values":[0,0,0,1,0,0],"isMain":false},{"metricName":"# local misassemblies","quality":"Less is better","values":[1,2,0,0,1,1],"isMain":false},{"metricName":"# scaffold gap ext. mis.","quality":"Less is better","values":[0,0,0,0,0,0],"isMain":false},{"metricName":"# scaffold gap loc. mis.","quality":"Less is better","values":[0,0,0,0,0,0],"isMain":false},{"metricName":"# unaligned mis. contigs","quality":"Less is better","values":[0,0,0,0,0,0],"isMain":false}]],["Unaligned",[{"metricName":"# fully unaligned contigs","quality":"Less is better","values":[0,0,2,1,0,0],"isMain":false},{"metricName":"Fully unaligned length","quality":"Less is better","values":[0,0,1971,1224,0,0],"isMain":false},{"metricName":"# partially unaligned contigs","quality":"Less is better","values":[7,10,1,8,11,6],"isMain":false},{"metricName":"Partially unaligned length","quality":"Less is better","values":[8201,13873,636,7873,31150,5118],"isMain":false}]],["Mismatches",[{"metricName":"# mismatches","quality":"Less is better","values":[12277,12862,18915,20585,13979,10795],"isMain":false},{"metricName":"# indels","quality":"Less is better","values":[18,24,21,24,21,10],"isMain":false},{"metricName":"Indels length","quality":"Less is better","values":[28,25,74,25,49,14],"isMain":false},{"metricName":"# mismatches per 100 kbp","quality":"Less is better","values":["1201.99","1131.64","1834.28","1922.10","1183.36","1313.90"],"isMain":true},{"metricName":"# indels per 100 kbp","quality":"Less is better","values":["1.76","2.11","2.04","2.24","1.78","1.22"],"isMain":true},{"metricName":" # indels (<= 5 bp)","quality":"Less is better","values":[18,24,20,24,19,10],"isMain":false},{"metricName":" # indels (> 5 bp)","quality":"Less is better","values":[0,0,1,0,2,0],"isMain":false},{"metricName":"# N's","quality":"Less is better","values":[0,0,0,0,0,0],"isMain":false},{"metricName":"# N's per 100 kbp","quality":"Less is better","values":["0.00","0.00","0.00","0.00","0.00","0.00"],"isMain":true}]],["Statistics without reference",[{"metricName":"# contigs","quality":"Equal","values":[1272,1268,1374,1406,1299,1119],"isMain":true},{"metricName":"# contigs (>= 1000 bp)","quality":"Equal","values":[283,396,240,278,424,194],"isMain":false},{"metricName":"# contigs (>= 5000 bp)","quality":"Equal","values":[0,0,0,0,2,0],"isMain":false},{"metricName":"# contigs (>= 10000 bp)","quality":"Equal","values":[0,0,0,0,1,0],"isMain":false},{"metricName":"# contigs (>= 25000 bp)","quality":"Equal","values":[0,0,0,0,0,0],"isMain":false},{"metricName":"# contigs (>= 50000 bp)","quality":"Equal","values":[0,0,0,0,0,0],"isMain":false},{"metricName":"Largest contig","quality":"More is better","values":[3193,4610,2444,3201,16989,2232],"isMain":true},{"metricName":"Total length","quality":"More is better","values":[1063048,1182592,1082685,1133814,1249800,857331],"isMain":true},{"metricName":"Total length (>= 1000 bp)","quality":"More is better","values":[387888,575490,315340,365575,634859,237835],"isMain":true},{"metricName":"Total length (>= 5000 bp)","quality":"More is better","values":[0,0,0,0,23809,0],"isMain":false},{"metricName":"Total length (>= 10000 bp)","quality":"More is better","values":[0,0,0,0,16989,0],"isMain":true},{"metricName":"Total length (>= 25000 bp)","quality":"More is better","values":[0,0,0,0,0,0],"isMain":false},{"metricName":"Total length (>= 50000 bp)","quality":"More is better","values":[0,0,0,0,0,0],"isMain":true},{"metricName":"N50","quality":"More is better","values":[854,977,788,808,1018,763],"isMain":false},{"metricName":"N75","quality":"More is better","values":[646,704,616,629,720,614],"isMain":false},{"metricName":"L50","quality":"Less is better","values":[440,412,496,502,415,417],"isMain":false},{"metricName":"L75","quality":"Less is better","values":[803,770,886,900,779,733],"isMain":false},{"metricName":"GC (%)","quality":"Equal","values":["61.94","62.07","61.96","61.95","62.03","61.88"],"isMain":false}]],["K-mer-based statistics",[]],["Predicted genes",[]],["Similarity statistics",[]],["Reference statistics",[{"metricName":"Reference length","quality":"Equal","values":[1804297,1804297,1804297,1804297,1804297,1804297],"isMain":false},{"metricName":"Reference fragments","quality":"Equal","values":[55,55,55,55,55,55],"isMain":false},{"metricName":"Reference GC (%)","quality":"Equal","values":["62.47","62.47","62.47","62.47","62.47","62.47"],"isMain":false}]]],"subreferences":[],"subreports":[],"minContig":500},{"date":"10 February 2020, Monday, 18:00:21","assembliesNames":["megahit-103-df","megahit-113-df","megahit-113-ml","megahit-113-ms","megahit-129-df","metaSPAdes"],"referenceName":"GCF\_000156495.1\_ASM15649v1","order":[0,1,2,3,4,5],"report":[["Genome statistics",[{"metricName":"Genome fraction (%)","quality":"More is better","values":["0.370","0.365","0.398","0.273","0.264","0.129"],"isMain":true},{"metricName":"Duplication ratio","quality":"Less is better","values":["1.059","1.167","1.081","1.146","1.174","1.036"],"isMain":true},{"metricName":"Largest alignment","quality":"More is better","values":[1760,1673,2012,1322,1295,1120],"isMain":true},{"metricName":"Total aligned length","quality":"More is better","values":[17282,16847,18762,12779,12164,6033],"isMain":true},{"metricName":"NA50","quality":"More is better","values":[762,549,630,569,435,650],"isMain":false},{"metricName":"NA75","quality":"More is better","values":[517,null,351,null,null,558],"isMain":false},{"metricName":"NGA50","quality":"More is better","values":[null,null,null,null,null,null],"isMain":true},{"metricName":"LA50","quality":"Less is better","values":[11,14,13,11,15,4],"isMain":false},{"metricName":"LA75","quality":"Less is better","values":[19,null,24,null,null,7],"isMain":false}]],["Reads mapping",[]],["Misassemblies",[{"metricName":"# misassemblies","quality":"Less is better","values":[0,0,1,0,0,0],"isMain":true},{"metricName":" # relocations","quality":"Less is better","values":[0,0,0,0,0,0],"isMain":false},{"metricName":" # translocations","quality":"Less is better","values":[0,0,1,0,0,0],"isMain":false},{"metricName":" # inversions","quality":"Less is better","values":[0,0,0,0,0,0],"isMain":false},{"metricName":" # interspecies translocations","quality":"Less is better","values":[4,10,6,5,11,0],"isMain":false},{"metricName":"# misassembled contigs","quality":"Less is better","values":[0,0,1,0,0,0],"isMain":false},{"metricName":"Misassembled contigs length","quality":"Less is better","values":[0,0,784,0,0,0],"isMain":true},{"metricName":"# possibly misassembled contigs","quality":"Less is better","values":[0,1,2,2,2,0],"isMain":false},{"metricName":" # possible misassemblies","quality":"Less is better","values":[0,1,2,2,2,0],"isMain":false},{"metricName":"# local misassemblies","quality":"Less is better","values":[0,0,0,0,0,0],"isMain":false},{"metricName":"# scaffold gap ext. mis.","quality":"Less is better","values":[0,0,0,0,0,0],"isMain":false},{"metricName":"# scaffold gap loc. mis.","quality":"Less is better","values":[0,0,0,0,0,0],"isMain":false},{"metricName":"# unaligned mis. contigs","quality":"Less is better","values":[1,1,1,0,0,0],"isMain":false}]],["Unaligned",[{"metricName":"# fully unaligned contigs","quality":"Less is better","values":[0,0,2,2,2,0],"isMain":false},{"metricName":"Fully unaligned length","quality":"Less is better","values":[0,0,1259,2362,1711,0],"isMain":false},{"metricName":"# partially unaligned contigs","quality":"Less is better","values":[3,4,3,1,5,0],"isMain":false},{"metricName":"Partially unaligned length","quality":"Less is better","values":[2685,3864,2841,694,4539,0],"isMain":false}]],["Mismatches",[{"metricName":"# mismatches","quality":"Less is better","values":[374,430,431,347,284,135],"isMain":false},{"metricName":"# indels","quality":"Less is better","values":[1,4,6,4,3,0],"isMain":false},{"metricName":"Indels length","quality":"Less is better","values":[1,4,11,5,3,0],"isMain":false},{"metricName":"# mismatches per 100 kbp","quality":"Less is better","values":["2189.82","2552.08","2349.41","2754.84","2334.57","2265.86"],"isMain":true},{"metricName":"# indels per 100 kbp","quality":"Less is better","values":["5.86","23.74","32.71","31.76","24.66","0.00"],"isMain":true},{"metricName":" # indels (<= 5 bp)","quality":"Less is better","values":[1,4,6,4,3,0],"isMain":false},{"metricName":" # indels (> 5 bp)","quality":"Less is better","values":[0,0,0,0,0,0],"isMain":false},{"metricName":"# N's","quality":"Less is better","values":[0,0,0,0,0,0],"isMain":false},{"metricName":"# N's per 100 kbp","quality":"Less is better","values":["0.00","0.00","0.00","0.00","0.00","0.00"],"isMain":true}]],["Statistics without reference",[{"metricName":"# contigs","quality":"Equal","values":[25,28,29,22,26,9],"isMain":true},{"metricName":"# contigs (>= 1000 bp)","quality":"Equal","values":[5,6,6,6,6,1],"isMain":false},{"metricName":"# contigs (>= 5000 bp)","quality":"Equal","values":[0,0,0,0,0,0],"isMain":false},{"metricName":"# contigs (>= 10000 bp)","quality":"Equal","values":[0,0,0,0,0,0],"isMain":false},{"metricName":"# contigs (>= 25000 bp)","quality":"Equal","values":[0,0,0,0,0,0],"isMain":false},{"metricName":"# contigs (>= 50000 bp)","quality":"Equal","values":[0,0,0,0,0,0],"isMain":false},{"metricName":"Largest contig","quality":"More is better","values":[1760,2032,2135,1341,1501,1120],"isMain":true},{"metricName":"Total length","quality":"More is better","values":[20769,23520,23938,17488,20535,6172],"isMain":true},{"metricName":"Total length (>= 1000 bp)","quality":"More is better","values":[6562,8765,8748,7117,7690,1120],"isMain":true},{"metricName":"Total length (>= 5000 bp)","quality":"More is better","values":[0,0,0,0,0,0],"isMain":false},{"metricName":"Total length (>= 10000 bp)","quality":"More is better","values":[0,0,0,0,0,0],"isMain":true},{"metricName":"Total length (>= 25000 bp)","quality":"More is better","values":[0,0,0,0,0,0],"isMain":false},{"metricName":"Total length (>= 50000 bp)","quality":"More is better","values":[0,0,0,0,0,0],"isMain":true},{"metricName":"N50","quality":"More is better","values":[859,858,799,843,818,669],"isMain":false},{"metricName":"N75","quality":"More is better","values":[692,649,614,626,582,569],"isMain":false},{"metricName":"L50","quality":"Less is better","values":[10,10,10,8,9,4],"isMain":false},{"metricName":"L75","quality":"Less is better","values":[16,18,19,15,17,7],"isMain":false},{"metricName":"GC (%)","quality":"Equal","values":["44.71","43.10","43.85","44.00","44.77","44.67"],"isMain":false}]],["K-mer-based statistics",[]],["Predicted genes",[]],["Similarity statistics",[]],["Reference statistics",[{"metricName":"Reference length","quality":"Equal","values":[4612238,4612238,4612238,4612238,4612238,4612238],"isMain":false},{"metricName":"Reference fragments","quality":"Equal","values":[527,527,527,527,527,527],"isMain":false},{"metricName":"Reference GC (%)","quality":"Equal","values":["45.20","45.20","45.20","45.20","45.20","45.20"],"isMain":false}]]],"subreferences":[],"subreports":[],"minContig":500},{"date":"10 February 2020, Monday, 18:00:58","assembliesNames":["megahit-103-df","megahit-113-df","megahit-113-ml","megahit-113-ms","megahit-129-df","metaSPAdes"],"referenceName":"GCF\_000156515.1\_ASM15651v1","order":[0,1,2,3,4,5],"report":[["Genome statistics",[{"metricName":"Genome fraction (%)","quality":"More is better","values":["97.784","97.915","97.877","97.960","97.901","97.716"],"isMain":true},{"metricName":"Duplication ratio","quality":"Less is better","values":["1.005","1.002","1.003","1.002","1.003","1.004"],"isMain":true},{"metricName":"Largest alignment","quality":"More is better","values":[178976,270164,335264,321453,335252,335264],"isMain":true},{"metricName":"Total aligned length","quality":"More is better","values":[3819042,3808180,3809888,3811116,3812438,3811356],"isMain":true},{"metricName":"NG50","quality":"More is better","values":[51776,117116,113041,120071,109588,113041],"isMain":false},{"metricName":"NG75","quality":"More is better","values":[29425,80139,74312,90673,68788,60924],"isMain":false},{"metricName":"NA50","quality":"More is better","values":[52365,116955,109184,120071,109538,113041],"isMain":false},{"metricName":"NA75","quality":"More is better","values":[30651,80139,69610,80083,67533,63219],"isMain":false},{"metricName":"NGA50","quality":"More is better","values":[51776,116955,109184,120071,109538,113041],"isMain":true},{"metricName":"NGA75","quality":"More is better","values":[29425,80139,69610,80083,67533,60548],"isMain":false},{"metricName":"LG50","quality":"Less is better","values":[21,12,12,11,13,10],"isMain":false},{"metricName":"LG75","quality":"Less is better","values":[45,21,22,20,24,22],"isMain":false},{"metricName":"LA50","quality":"Less is better","values":[20,12,12,11,13,10],"isMain":false},{"metricName":"LA75","quality":"Less is better","values":[43,21,23,20,24,21],"isMain":false},{"metricName":"LGA50","quality":"Less is better","values":[21,12,12,11,13,10],"isMain":true},{"metricName":"LGA75","quality":"Less is better","values":[45,21,23,20,24,22],"isMain":false}]],["Reads mapping",[]],["Misassemblies",[{"metricName":"# misassemblies","quality":"Less is better","values":[3,6,6,5,4,3],"isMain":true},{"metricName":" # relocations","quality":"Less is better","values":[1,0,2,0,0,2],"isMain":false},{"metricName":" # translocations","quality":"Less is better","values":[2,6,4,5,3,1],"isMain":false},{"metricName":" # inversions","quality":"Less is better","values":[0,0,0,0,1,0],"isMain":false},{"metricName":" # interspecies translocations","quality":"Less is better","values":[8,4,7,4,13,2],"isMain":false},{"metricName":"# misassembled contigs","quality":"Less is better","values":[3,4,6,5,4,3],"isMain":false},{"metricName":"Misassembled contigs length","quality":"Less is better","values":[27961,63596,114241,161589,107105,64160],"isMain":true},{"metricName":"# possibly misassembled contigs","quality":"Less is better","values":[0,0,0,0,0,0],"isMain":false},{"metricName":" # possible misassemblies","quality":"Less is better","values":[0,0,0,0,0,0],"isMain":false},{"metricName":"# local misassemblies","quality":"Less is better","values":[0,4,3,2,3,0],"isMain":false},{"metricName":"# scaffold gap ext. mis.","quality":"Less is better","values":[0,0,0,0,0,0],"isMain":false},{"metricName":"# scaffold gap loc. mis.","quality":"Less is better","values":[0,0,0,0,0,0],"isMain":false},{"metricName":"# unaligned mis. contigs","quality":"Less is better","values":[0,0,0,0,1,0],"isMain":false}]],["Unaligned",[{"metricName":"# fully unaligned contigs","quality":"Less is better","values":[0,0,0,0,0,1],"isMain":false},{"metricName":"Fully unaligned length","quality":"Less is better","values":[0,0,0,0,0,620],"isMain":false},{"metricName":"# partially unaligned contigs","quality":"Less is better","values":[6,3,3,1,7,1],"isMain":false},{"metricName":"Partially unaligned length","quality":"Less is better","values":[3974,2407,75330,631,45462,1172],"isMain":false}]],["Mismatches",[{"metricName":"# mismatches","quality":"Less is better","values":[3040,407,510,549,626,245],"isMain":false},{"metricName":"# indels","quality":"Less is better","values":[3,21,20,17,23,14],"isMain":false},{"metricName":"Indels length","quality":"Less is better","values":[3,233,239,220,225,19],"isMain":false},{"metricName":"# mismatches per 100 kbp","quality":"Less is better","values":["80.01","10.70","13.41","14.42","16.46","6.45"],"isMain":true},{"metricName":"# indels per 100 kbp","quality":"Less is better","values":["0.08","0.55","0.53","0.45","0.60","0.37"],"isMain":true},{"metricName":" # indels (<= 5 bp)","quality":"Less is better","values":[3,17,14,12,19,14],"isMain":false},{"metricName":" # indels (> 5 bp)","quality":"Less is better","values":[0,4,6,5,4,0],"isMain":false},{"metricName":"# N's","quality":"Less is better","values":[0,0,0,0,0,0],"isMain":false},{"metricName":"# N's per 100 kbp","quality":"Less is better","values":["0.00","0.00","0.00","0.00","0.00","0.00"],"isMain":true}]],["Statistics without reference",[{"metricName":"# contigs","quality":"Equal","values":[137,66,76,71,82,71],"isMain":true},{"metricName":"# contigs (>= 1000 bp)","quality":"Equal","values":[118,57,62,55,64,63],"isMain":false},{"metricName":"# contigs (>= 5000 bp)","quality":"Equal","values":[99,46,52,48,49,50],"isMain":false},{"metricName":"# contigs (>= 10000 bp)","quality":"Equal","values":[81,45,48,45,46,45],"isMain":false},{"metricName":"# contigs (>= 25000 bp)","quality":"Equal","values":[53,36,38,33,39,36],"isMain":false},{"metricName":"# contigs (>= 50000 bp)","quality":"Equal","values":[22,26,27,24,31,29],"isMain":false},{"metricName":"Largest contig","quality":"More is better","values":[178976,270164,335264,321479,335252,335264],"isMain":true},{"metricName":"Total length","quality":"More is better","values":[3823793,3812715,3889116,3814375,3861911,3813917],"isMain":true},{"metricName":"Total length (>= 1000 bp)","quality":"More is better","values":[3810186,3806237,3879650,3803115,3849822,3808280],"isMain":true},{"metricName":"Total length (>= 5000 bp)","quality":"More is better","values":[3768399,3781159,3858695,3784948,3812216,3779443],"isMain":false},{"metricName":"Total length (>= 10000 bp)","quality":"More is better","values":[3637117,3774060,3827121,3763379,3789869,3744128],"isMain":true},{"metricName":"Total length (>= 25000 bp)","quality":"More is better","values":[3158894,3617508,3656494,3562266,3668633,3594535],"isMain":false},{"metricName":"Total length (>= 50000 bp)","quality":"More is better","values":[2026508,3231073,3253389,3220464,3347877,3338497],"isMain":true},{"metricName":"N50","quality":"More is better","values":[52365,117116,113041,120071,109588,113041],"isMain":false},{"metricName":"N75","quality":"More is better","values":[30651,80139,74312,90673,68788,63219],"isMain":false},{"metricName":"L50","quality":"Less is better","values":[20,12,12,11,13,10],"isMain":false},{"metricName":"L75","quality":"Less is better","values":[43,21,22,20,24,21],"isMain":false},{"metricName":"GC (%)","quality":"Equal","values":["48.84","48.84","48.65","48.85","48.79","48.85"],"isMain":false}]],["K-mer-based statistics",[]],["Predicted genes",[]],["Similarity statistics",[]],["Reference statistics",[{"metricName":"Reference length","quality":"Equal","values":[3885459,3885459,3885459,3885459,3885459,3885459],"isMain":false},{"metricName":"Reference fragments","quality":"Equal","values":[167,167,167,167,167,167],"isMain":false},{"metricName":"Reference GC (%)","quality":"Equal","values":["48.85","48.85","48.85","48.85","48.85","48.85"],"isMain":false}]]],"subreferences":[],"subreports":[],"minContig":500},{"date":"10 February 2020, Monday, 18:00:18","assembliesNames":["megahit-103-df","megahit-113-df","megahit-113-ml","megahit-113-ms","megahit-129-df","metaSPAdes"],"referenceName":"GCF\_000157015.1\_ASM15701v1","order":[0,1,2,3,4,5],"report":[["Genome statistics",[{"metricName":"Genome fraction (%)","quality":"More is better","values":["4.639","4.361","5.784","5.122","4.963","2.091"],"isMain":true},{"metricName":"Duplication ratio","quality":"Less is better","values":["1.035","1.026","1.027","1.024","1.029","1.019"],"isMain":true},{"metricName":"Largest alignment","quality":"More is better","values":[2067,7741,5961,2141,25906,2798],"isMain":true},{"metricName":"Total aligned length","quality":"More is better","values":[256001,240004,319269,282176,273273,115271],"isMain":true},{"metricName":"NA50","quality":"More is better","values":[678,685,601,662,729,648],"isMain":false},{"metricName":"NA75","quality":"More is better","values":[570,547,507,557,561,541],"isMain":false},{"metricName":"NGA50","quality":"More is better","values":[null,null,null,null,null,null],"isMain":true},{"metricName":"LA50","quality":"Less is better","values":[146,126,236,170,119,66],"isMain":false},{"metricName":"LA75","quality":"Less is better","values":[253,234,412,291,237,116],"isMain":false}]],["Reads mapping",[]],["Misassemblies",[{"metricName":"# misassemblies","quality":"Less is better","values":[1,9,1,1,5,1],"isMain":true},{"metricName":" # relocations","quality":"Less is better","values":[0,2,0,0,0,0],"isMain":false},{"metricName":" # translocations","quality":"Less is better","values":[1,7,1,1,5,1],"isMain":false},{"metricName":" # inversions","quality":"Less is better","values":[0,0,0,0,0,0],"isMain":false},{"metricName":" # interspecies translocations","quality":"Less is better","values":[18,21,17,16,22,2],"isMain":false},{"metricName":"# misassembled contigs","quality":"Less is better","values":[1,4,1,1,3,1],"isMain":false},{"metricName":"Misassembled contigs length","quality":"Less is better","values":[804,22601,647,953,20868,559],"isMain":true},{"metricName":"# possibly misassembled contigs","quality":"Less is better","values":[0,0,0,0,0,0],"isMain":false},{"metricName":" # possible misassemblies","quality":"Less is better","values":[0,0,0,0,0,0],"isMain":false},{"metricName":"# local misassemblies","quality":"Less is better","values":[0,2,1,0,3,0],"isMain":false},{"metricName":"# scaffold gap ext. mis.","quality":"Less is better","values":[0,0,0,0,0,0],"isMain":false},{"metricName":"# scaffold gap loc. mis.","quality":"Less is better","values":[0,0,0,0,0,0],"isMain":false},{"metricName":"# unaligned mis. contigs","quality":"Less is better","values":[0,0,1,1,0,0],"isMain":false}]],["Unaligned",[{"metricName":"# fully unaligned contigs","quality":"Less is better","values":[1,0,1,0,0,0],"isMain":false},{"metricName":"Fully unaligned length","quality":"Less is better","values":[787,0,757,0,0,0],"isMain":false},{"metricName":"# partially unaligned contigs","quality":"Less is better","values":[2,9,5,5,7,1],"isMain":false},{"metricName":"Partially unaligned length","quality":"Less is better","values":[1046,21295,63070,5018,18578,585],"isMain":false}]],["Mismatches",[{"metricName":"# mismatches","quality":"Less is better","values":[4267,3707,6508,6205,3983,1759],"isMain":false},{"metricName":"# indels","quality":"Less is better","values":[26,9,35,31,13,4],"isMain":false},{"metricName":"Indels length","quality":"Less is better","values":[53,20,61,43,60,4],"isMain":false},{"metricName":"# mismatches per 100 kbp","quality":"Less is better","values":["1676.66","1549.40","2050.92","2208.04","1462.70","1533.67"],"isMain":true},{"metricName":"# indels per 100 kbp","quality":"Less is better","values":["10.22","3.76","11.03","11.03","4.77","3.49"],"isMain":true},{"metricName":" # indels (<= 5 bp)","quality":"Less is better","values":[23,8,32,30,12,4],"isMain":false},{"metricName":" # indels (> 5 bp)","quality":"Less is better","values":[3,1,3,1,1,0],"isMain":false},{"metricName":"# N's","quality":"Less is better","values":[0,0,0,0,0,0],"isMain":false},{"metricName":"# N's per 100 kbp","quality":"Less is better","values":["0.00","0.00","0.00","0.00","0.00","0.00"],"isMain":true}]],["Statistics without reference",[{"metricName":"# contigs","quality":"Equal","values":[367,311,470,415,335,169],"isMain":true},{"metricName":"# contigs (>= 1000 bp)","quality":"Equal","values":[40,41,41,34,49,16],"isMain":false},{"metricName":"# contigs (>= 5000 bp)","quality":"Equal","values":[0,3,3,0,3,0],"isMain":false},{"metricName":"# contigs (>= 10000 bp)","quality":"Equal","values":[0,1,2,0,2,0],"isMain":false},{"metricName":"# contigs (>= 25000 bp)","quality":"Equal","values":[0,0,1,0,1,0],"isMain":false},{"metricName":"# contigs (>= 50000 bp)","quality":"Equal","values":[0,0,0,0,0,0],"isMain":false},{"metricName":"Largest contig","quality":"More is better","values":[2067,18666,35975,2278,25906,2804],"isMain":true},{"metricName":"Total length","quality":"More is better","values":[265182,266722,389693,292890,298724,117509],"isMain":true},{"metricName":"Total length (>= 1000 bp)","quality":"More is better","values":[49024,86944,115741,43360,109305,21885],"isMain":true},{"metricName":"Total length (>= 5000 bp)","quality":"More is better","values":[0,33510,66656,0,50732,0],"isMain":false},{"metricName":"Total length (>= 10000 bp)","quality":"More is better","values":[0,18666,60695,0,45111,0],"isMain":true},{"metricName":"Total length (>= 25000 bp)","quality":"More is better","values":[0,0,35975,0,25906,0],"isMain":false},{"metricName":"Total length (>= 50000 bp)","quality":"More is better","values":[0,0,0,0,0,0],"isMain":true},{"metricName":"N50","quality":"More is better","values":[707,805,711,682,813,653],"isMain":false},{"metricName":"N75","quality":"More is better","values":[595,618,586,578,618,559],"isMain":false},{"metricName":"L50","quality":"Less is better","values":[143,94,137,163,94,65],"isMain":false},{"metricName":"L75","quality":"Less is better","values":[246,191,289,280,201,114],"isMain":false},{"metricName":"GC (%)","quality":"Equal","values":["44.16","44.03","44.35","44.04","45.02","44.20"],"isMain":false}]],["K-mer-based statistics",[]],["Predicted genes",[]],["Similarity statistics",[]],["Reference statistics",[{"metricName":"Reference length","quality":"Equal","values":[5486239,5486239,5486239,5486239,5486239,5486239],"isMain":false},{"metricName":"Reference fragments","quality":"Equal","values":[105,105,105,105,105,105],"isMain":false},{"metricName":"Reference GC (%)","quality":"Equal","values":["43.62","43.62","43.62","43.62","43.62","43.62"],"isMain":false}]]],"subreferences":[],"subreports":[],"minContig":500},{"date":"10 February 2020, Monday, 18:00:38","assembliesNames":["megahit-103-df","megahit-113-df","megahit-113-ml","megahit-113-ms","megahit-129-df","metaSPAdes"],"referenceName":"GCF\_000157035.2\_Bacteroides\_sp\_2\_1\_7\_V2","order":[0,1,2,3,4,5],"report":[["Genome statistics",[{"metricName":"Genome fraction (%)","quality":"More is better","values":["5.559","6.371","5.154","6.224","7.033","4.009"],"isMain":true},{"metricName":"Duplication ratio","quality":"Less is better","values":["1.023","1.019","1.017","1.028","1.017","1.024"],"isMain":true},{"metricName":"Largest alignment","quality":"More is better","values":[2601,3220,4741,2040,3118,2636],"isMain":true},{"metricName":"Total aligned length","quality":"More is better","values":[288211,330015,267184,323293,363831,207930],"isMain":true},{"metricName":"NA50","quality":"More is better","values":[761,895,720,727,930,713],"isMain":false},{"metricName":"NA75","quality":"More is better","values":[583,645,578,592,644,558],"isMain":false},{"metricName":"NGA50","quality":"More is better","values":[null,null,null,null,null,null],"isMain":true},{"metricName":"LA50","quality":"Less is better","values":[133,128,136,169,135,106],"isMain":false},{"metricName":"LA75","quality":"Less is better","values":[243,241,241,296,258,191],"isMain":false}]],["Reads mapping",[]],["Misassemblies",[{"metricName":"# misassemblies","quality":"Less is better","values":[1,2,1,2,3,0],"isMain":true},{"metricName":" # relocations","quality":"Less is better","values":[1,1,0,0,1,0],"isMain":false},{"metricName":" # translocations","quality":"Less is better","values":[0,1,1,1,2,0],"isMain":false},{"metricName":" # inversions","quality":"Less is better","values":[0,0,0,1,0,0],"isMain":false},{"metricName":" # interspecies translocations","quality":"Less is better","values":[16,18,1,17,17,10],"isMain":false},{"metricName":"# misassembled contigs","quality":"Less is better","values":[1,2,1,2,3,0],"isMain":false},{"metricName":"Misassembled contigs length","quality":"Less is better","values":[1214,2369,1187,1378,2120,0],"isMain":true},{"metricName":"# possibly misassembled contigs","quality":"Less is better","values":[0,0,0,0,0,0],"isMain":false},{"metricName":" # possible misassemblies","quality":"Less is better","values":[0,0,0,0,0,0],"isMain":false},{"metricName":"# local misassemblies","quality":"Less is better","values":[1,0,0,0,1,0],"isMain":false},{"metricName":"# scaffold gap ext. mis.","quality":"Less is better","values":[0,0,0,0,0,0],"isMain":false},{"metricName":"# scaffold gap loc. mis.","quality":"Less is better","values":[0,0,0,0,0,0],"isMain":false},{"metricName":"# unaligned mis. contigs","quality":"Less is better","values":[0,0,0,0,0,0],"isMain":false}]],["Unaligned",[{"metricName":"# fully unaligned contigs","quality":"Less is better","values":[0,2,1,0,2,0],"isMain":false},{"metricName":"Fully unaligned length","quality":"Less is better","values":[0,1495,700,0,1593,0],"isMain":false},{"metricName":"# partially unaligned contigs","quality":"Less is better","values":[2,4,0,3,2,1],"isMain":false},{"metricName":"Partially unaligned length","quality":"Less is better","values":[1625,4780,0,2417,1457,550],"isMain":false}]],["Mismatches",[{"metricName":"# mismatches","quality":"Less is better","values":[4740,5241,5658,7278,6003,3442],"isMain":false},{"metricName":"# indels","quality":"Less is better","values":[18,17,17,26,24,18],"isMain":false},{"metricName":"Indels length","quality":"Less is better","values":[89,43,43,128,49,40],"isMain":false},{"metricName":"# mismatches per 100 kbp","quality":"Less is better","values":["1651.58","1593.36","2126.24","2264.83","1653.16","1663.05"],"isMain":true},{"metricName":"# indels per 100 kbp","quality":"Less is better","values":["6.27","5.17","6.39","8.09","6.61","8.70"],"isMain":true},{"metricName":" # indels (<= 5 bp)","quality":"Less is better","values":[16,16,16,22,23,17],"isMain":false},{"metricName":" # indels (> 5 bp)","quality":"Less is better","values":[2,1,1,4,1,1],"isMain":false},{"metricName":"# N's","quality":"Less is better","values":[0,0,0,0,0,0],"isMain":false},{"metricName":"# N's per 100 kbp","quality":"Less is better","values":["0.00","0.00","0.00","0.00","0.00","0.00"],"isMain":true}]],["Statistics without reference",[{"metricName":"# contigs","quality":"Equal","values":[371,377,361,438,411,285],"isMain":true},{"metricName":"# contigs (>= 1000 bp)","quality":"Equal","values":[70,102,44,56,117,41],"isMain":false},{"metricName":"# contigs (>= 5000 bp)","quality":"Equal","values":[0,0,0,0,0,0],"isMain":false},{"metricName":"# contigs (>= 10000 bp)","quality":"Equal","values":[0,0,0,0,0,0],"isMain":false},{"metricName":"# contigs (>= 25000 bp)","quality":"Equal","values":[0,0,0,0,0,0],"isMain":false},{"metricName":"# contigs (>= 50000 bp)","quality":"Equal","values":[0,0,0,0,0,0],"isMain":false},{"metricName":"Largest contig","quality":"More is better","values":[2601,4300,4741,2040,3138,2636],"isMain":true},{"metricName":"Total length","quality":"More is better","values":[295112,341432,271351,332717,372352,212560],"isMain":true},{"metricName":"Total length (>= 1000 bp)","quality":"More is better","values":[95472,151287,59544,72279,170543,54045],"isMain":true},{"metricName":"Total length (>= 5000 bp)","quality":"More is better","values":[0,0,0,0,0,0],"isMain":false},{"metricName":"Total length (>= 10000 bp)","quality":"More is better","values":[0,0,0,0,0,0],"isMain":true},{"metricName":"Total length (>= 25000 bp)","quality":"More is better","values":[0,0,0,0,0,0],"isMain":false},{"metricName":"Total length (>= 50000 bp)","quality":"More is better","values":[0,0,0,0,0,0],"isMain":true},{"metricName":"N50","quality":"More is better","values":[779,930,740,761,949,733],"isMain":false},{"metricName":"N75","quality":"More is better","values":[609,680,594,613,674,577],"isMain":false},{"metricName":"L50","quality":"Less is better","values":[130,123,134,165,134,105],"isMain":false},{"metricName":"L75","quality":"Less is better","values":[237,231,237,288,251,187],"isMain":false},{"metricName":"GC (%)","quality":"Equal","values":["45.90","45.54","45.47","45.43","45.86","45.64"],"isMain":false}]],["K-mer-based statistics",[]],["Predicted genes",[]],["Similarity statistics",[]],["Reference statistics",[{"metricName":"Reference length","quality":"Equal","values":[5163202,5163202,5163202,5163202,5163202,5163202],"isMain":false},{"metricName":"Reference fragments","quality":"Equal","values":[41,41,41,41,41,41],"isMain":false},{"metricName":"Reference GC (%)","quality":"Equal","values":["45.08","45.08","45.08","45.08","45.08","45.08"],"isMain":false}]]],"subreferences":[],"subreports":[],"minContig":500},{"date":"10 February 2020, Monday, 18:00:36","assembliesNames":["megahit-103-df","megahit-113-df","megahit-113-ml","megahit-113-ms","megahit-129-df","metaSPAdes"],"referenceName":"GCF\_000157075.2\_Bact\_sp\_9\_1\_42FAA\_V2","order":[0,1,2,3,4,5],"report":[["Genome statistics",[{"metricName":"Genome fraction (%)","quality":"More is better","values":["2.039","2.314","3.234","3.143","2.724","1.328"],"isMain":true},{"metricName":"Duplication ratio","quality":"Less is better","values":["1.027","1.011","1.021","1.018","1.020","1.024"],"isMain":true},{"metricName":"Largest alignment","quality":"More is better","values":[4500,6889,2581,2760,5194,3223],"isMain":true},{"metricName":"Total aligned length","quality":"More is better","values":[115099,130006,182521,177483,153493,74905],"isMain":true},{"metricName":"NA50","quality":"More is better","values":[740,847,747,724,836,726],"isMain":false},{"metricName":"NA75","quality":"More is better","values":[599,631,576,576,593,585],"isMain":false},{"metricName":"NGA50","quality":"More is better","values":[null,null,null,null,null,null],"isMain":true},{"metricName":"LA50","quality":"Less is better","values":[51,43,84,88,64,36],"isMain":false},{"metricName":"LA75","quality":"Less is better","values":[97,89,155,159,122,66],"isMain":false}]],["Reads mapping",[]],["Misassemblies",[{"metricName":"# misassemblies","quality":"Less is better","values":[1,1,1,1,0,0],"isMain":true},{"metricName":" # relocations","quality":"Less is better","values":[0,0,1,1,0,0],"isMain":false},{"metricName":" # translocations","quality":"Less is better","values":[0,0,0,0,0,0],"isMain":false},{"metricName":" # inversions","quality":"Less is better","values":[1,1,0,0,0,0],"isMain":false},{"metricName":" # interspecies translocations","quality":"Less is better","values":[9,4,6,6,5,4],"isMain":false},{"metricName":"# misassembled contigs","quality":"Less is better","values":[1,1,1,1,0,0],"isMain":false},{"metricName":"Misassembled contigs length","quality":"Less is better","values":[640,524,1742,612,0,0],"isMain":true},{"metricName":"# possibly misassembled contigs","quality":"Less is better","values":[0,0,0,0,0,0],"isMain":false},{"metricName":" # possible misassemblies","quality":"Less is better","values":[0,0,0,0,0,0],"isMain":false},{"metricName":"# local misassemblies","quality":"Less is better","values":[0,1,0,0,0,0],"isMain":false},{"metricName":"# scaffold gap ext. mis.","quality":"Less is better","values":[0,0,0,0,0,0],"isMain":false},{"metricName":"# scaffold gap loc. mis.","quality":"Less is better","values":[0,0,0,0,0,0],"isMain":false},{"metricName":"# unaligned mis. contigs","quality":"Less is better","values":[0,0,0,0,0,0],"isMain":false}]],["Unaligned",[{"metricName":"# fully unaligned contigs","quality":"Less is better","values":[0,0,0,0,0,0],"isMain":false},{"metricName":"Fully unaligned length","quality":"Less is better","values":[0,0,0,0,0,0],"isMain":false},{"metricName":"# partially unaligned contigs","quality":"Less is better","values":[2,2,1,3,2,2],"isMain":false},{"metricName":"Partially unaligned length","quality":"Less is better","values":[3222,1644,725,3360,5067,1634],"isMain":false}]],["Mismatches",[{"metricName":"# mismatches","quality":"Less is better","values":[1574,2102,3694,3839,2733,1264],"isMain":false},{"metricName":"# indels","quality":"Less is better","values":[6,9,29,10,20,8],"isMain":false},{"metricName":"Indels length","quality":"Less is better","values":[17,10,44,51,29,14],"isMain":false},{"metricName":"# mismatches per 100 kbp","quality":"Less is better","values":["1375.31","1617.76","2034.30","2175.28","1787.12","1695.12"],"isMain":true},{"metricName":"# indels per 100 kbp","quality":"Less is better","values":["5.24","6.93","15.97","5.67","13.08","10.73"],"isMain":true},{"metricName":" # indels (<= 5 bp)","quality":"Less is better","values":[5,9,29,9,20,7],"isMain":false},{"metricName":" # indels (> 5 bp)","quality":"Less is better","values":[1,0,0,1,0,1],"isMain":false},{"metricName":"# N's","quality":"Less is better","values":[0,0,0,0,0,0],"isMain":false},{"metricName":"# N's per 100 kbp","quality":"Less is better","values":["0.00","0.00","0.00","0.00","0.00","0.00"],"isMain":true}]],["Statistics without reference",[{"metricName":"# contigs","quality":"Equal","values":[144,142,236,236,183,98],"isMain":true},{"metricName":"# contigs (>= 1000 bp)","quality":"Equal","values":[28,33,40,38,40,19],"isMain":false},{"metricName":"# contigs (>= 5000 bp)","quality":"Equal","values":[0,2,0,0,1,0],"isMain":false},{"metricName":"# contigs (>= 10000 bp)","quality":"Equal","values":[0,0,0,0,0,0],"isMain":false},{"metricName":"# contigs (>= 25000 bp)","quality":"Equal","values":[0,0,0,0,0,0],"isMain":false},{"metricName":"# contigs (>= 50000 bp)","quality":"Equal","values":[0,0,0,0,0,0],"isMain":false},{"metricName":"Largest contig","quality":"More is better","values":[4500,7870,2607,2784,5207,3240],"isMain":true},{"metricName":"Total length","quality":"More is better","values":[120761,132961,186207,183077,161010,77965],"isMain":true},{"metricName":"Total length (>= 1000 bp)","quality":"More is better","values":[44682,60199,57848,51974,64875,26057],"isMain":true},{"metricName":"Total length (>= 5000 bp)","quality":"More is better","values":[0,13077,0,0,5207,0],"isMain":false},{"metricName":"Total length (>= 10000 bp)","quality":"More is better","values":[0,0,0,0,0,0],"isMain":true},{"metricName":"Total length (>= 25000 bp)","quality":"More is better","values":[0,0,0,0,0,0],"isMain":false},{"metricName":"Total length (>= 50000 bp)","quality":"More is better","values":[0,0,0,0,0,0],"isMain":true},{"metricName":"N50","quality":"More is better","values":[759,913,774,757,856,751],"isMain":false},{"metricName":"N75","quality":"More is better","values":[630,661,593,608,640,609],"isMain":false},{"metricName":"L50","quality":"Less is better","values":[47,40,82,85,58,35],"isMain":false},{"metricName":"L75","quality":"Less is better","values":[90,84,151,154,112,64],"isMain":false},{"metricName":"GC (%)","quality":"Equal","values":["41.36","41.66","42.15","41.65","41.22","41.93"],"isMain":false}]],["K-mer-based statistics",[]],["Predicted genes",[]],["Similarity statistics",[]],["Reference statistics",[{"metricName":"Reference length","quality":"Equal","values":[5614268,5614268,5614268,5614268,5614268,5614268],"isMain":false},{"metricName":"Reference fragments","quality":"Equal","values":[60,60,60,60,60,60],"isMain":false},{"metricName":"Reference GC (%)","quality":"Equal","values":["42.33","42.33","42.33","42.33","42.33","42.33"],"isMain":false}]]],"subreferences":[],"subreports":[],"minContig":500},{"date":"10 February 2020, Monday, 18:00:41","assembliesNames":["megahit-103-df","megahit-113-df","megahit-113-ml","megahit-113-ms","megahit-129-df","metaSPAdes"],"referenceName":"GCF\_000157915.1\_ASM15791v1","order":[0,1,2,3,4,5],"report":[["Genome statistics",[{"metricName":"Genome fraction (%)","quality":"More is better","values":["0.455","0.220","0.399","0.485","0.181","0.064"],"isMain":true},{"metricName":"Duplication ratio","quality":"Less is better","values":["1.027","1.144","1.038","1.059","1.034","1.014"],"isMain":true},{"metricName":"Largest alignment","quality":"More is better","values":[3978,2813,2051,3133,1820,859],"isMain":true},{"metricName":"Total aligned length","quality":"More is better","values":[17548,8615,15516,19691,6977,2470],"isMain":true},{"metricName":"NA50","quality":"More is better","values":[1563,1058,1373,1316,1805,591],"isMain":false},{"metricName":"NA75","quality":"More is better","values":[750,611,1097,742,808,525],"isMain":false},{"metricName":"NGA50","quality":"More is better","values":[null,null,null,null,null,null],"isMain":true},{"metricName":"LA50","quality":"Less is better","values":[4,3,5,6,2,2],"isMain":false},{"metricName":"LA75","quality":"Less is better","values":[8,6,8,11,4,3],"isMain":false}]],["Reads mapping",[]],["Misassemblies",[{"metricName":"# misassemblies","quality":"Less is better","values":[0,0,1,0,1,0],"isMain":true},{"metricName":" # relocations","quality":"Less is better","values":[0,0,0,0,0,0],"isMain":false},{"metricName":" # translocations","quality":"Less is better","values":[0,0,0,0,1,0],"isMain":false},{"metricName":" # inversions","quality":"Less is better","values":[0,0,1,0,0,0],"isMain":false},{"metricName":" # interspecies translocations","quality":"Less is better","values":[1,3,1,0,1,0],"isMain":false},{"metricName":"# misassembled contigs","quality":"Less is better","values":[0,0,1,0,1,0],"isMain":false},{"metricName":"Misassembled contigs length","quality":"Less is better","values":[0,0,681,0,539,0],"isMain":true},{"metricName":"# possibly misassembled contigs","quality":"Less is better","values":[0,0,0,0,0,0],"isMain":false},{"metricName":" # possible misassemblies","quality":"Less is better","values":[0,0,0,0,0,0],"isMain":false},{"metricName":"# local misassemblies","quality":"Less is better","values":[0,0,0,0,0,0],"isMain":false},{"metricName":"# scaffold gap ext. mis.","quality":"Less is better","values":[0,0,0,0,0,0],"isMain":false},{"metricName":"# scaffold gap loc. mis.","quality":"Less is better","values":[0,0,0,0,0,0],"isMain":false},{"metricName":"# unaligned mis. contigs","quality":"Less is better","values":[0,0,0,0,0,0],"isMain":false}]],["Unaligned",[{"metricName":"# fully unaligned contigs","quality":"Less is better","values":[0,0,0,0,0,0],"isMain":false},{"metricName":"Fully unaligned length","quality":"Less is better","values":[0,0,0,0,0,0],"isMain":false},{"metricName":"# partially unaligned contigs","quality":"Less is better","values":[0,0,0,0,0,0],"isMain":false},{"metricName":"Partially unaligned length","quality":"Less is better","values":[0,0,0,0,0,0],"isMain":false}]],["Mismatches",[{"metricName":"# mismatches","quality":"Less is better","values":[141,83,189,281,72,21],"isMain":false},{"metricName":"# indels","quality":"Less is better","values":[0,0,1,1,6,0],"isMain":false},{"metricName":"Indels length","quality":"Less is better","values":[0,0,1,1,19,0],"isMain":false},{"metricName":"# mismatches per 100 kbp","quality":"Less is better","values":["803.51","978.89","1227.67","1501.71","1033.89","850.20"],"isMain":true},{"metricName":"# indels per 100 kbp","quality":"Less is better","values":["0.00","0.00","6.50","5.34","86.16","0.00"],"isMain":true},{"metricName":" # indels (<= 5 bp)","quality":"Less is better","values":[0,0,1,1,5,0],"isMain":false},{"metricName":" # indels (> 5 bp)","quality":"Less is better","values":[0,0,0,0,1,0],"isMain":false},{"metricName":"# N's","quality":"Less is better","values":[0,0,0,0,0,0],"isMain":false},{"metricName":"# N's per 100 kbp","quality":"Less is better","values":["0.00","0.00","0.00","0.00","0.00","0.00"],"isMain":true}]],["Statistics without reference",[{"metricName":"# contigs","quality":"Equal","values":[15,10,14,18,7,4],"isMain":true},{"metricName":"# contigs (>= 1000 bp)","quality":"Equal","values":[5,3,8,7,3,0],"isMain":false},{"metricName":"# contigs (>= 5000 bp)","quality":"Equal","values":[0,0,0,0,0,0],"isMain":false},{"metricName":"# contigs (>= 10000 bp)","quality":"Equal","values":[0,0,0,0,0,0],"isMain":false},{"metricName":"# contigs (>= 25000 bp)","quality":"Equal","values":[0,0,0,0,0,0],"isMain":false},{"metricName":"# contigs (>= 50000 bp)","quality":"Equal","values":[0,0,0,0,0,0],"isMain":false},{"metricName":"Largest contig","quality":"More is better","values":[3978,2817,2051,3133,1820,859],"isMain":true},{"metricName":"Total length","quality":"More is better","values":[18025,9700,15978,19815,7199,2505],"isMain":true},{"metricName":"Total length (>= 1000 bp)","quality":"More is better","values":[11472,5182,12161,11832,4691,0],"isMain":true},{"metricName":"Total length (>= 5000 bp)","quality":"More is better","values":[0,0,0,0,0,0],"isMain":false},{"metricName":"Total length (>= 10000 bp)","quality":"More is better","values":[0,0,0,0,0,0],"isMain":true},{"metricName":"Total length (>= 25000 bp)","quality":"More is better","values":[0,0,0,0,0,0],"isMain":false},{"metricName":"Total length (>= 50000 bp)","quality":"More is better","values":[0,0,0,0,0,0],"isMain":true},{"metricName":"N50","quality":"More is better","values":[1563,1064,1375,1316,1805,596],"isMain":false},{"metricName":"N75","quality":"More is better","values":[750,626,1097,752,814,545],"isMain":false},{"metricName":"L50","quality":"Less is better","values":[4,3,5,6,2,2],"isMain":false},{"metricName":"L75","quality":"Less is better","values":[8,6,8,11,4,3],"isMain":false},{"metricName":"GC (%)","quality":"Equal","values":["47.30","43.61","48.61","50.11","45.87","48.50"],"isMain":false}]],["K-mer-based statistics",[]],["Predicted genes",[]],["Similarity statistics",[]],["Reference statistics",[{"metricName":"Reference length","quality":"Equal","values":[3855443,3855443,3855443,3855443,3855443,3855443],"isMain":false},{"metricName":"Reference fragments","quality":"Equal","values":[285,285,285,285,285,285],"isMain":false},{"metricName":"Reference GC (%)","quality":"Equal","values":["45.72","45.72","45.72","45.72","45.72","45.72"],"isMain":false}]]],"subreferences":[],"subreports":[],"minContig":500},{"date":"10 February 2020, Monday, 18:00:15","assembliesNames":["megahit-103-df","megahit-113-df","megahit-113-ml","megahit-113-ms","megahit-129-df","metaSPAdes"],"referenceName":"GCF\_000157935.1\_ASM15793v1","order":[0,1,2,3,4,5],"report":[["Genome statistics",[{"metricName":"Genome fraction (%)","quality":"More is better","values":["0.102","0.214","0.087","0.201","0.168","0.077"],"isMain":true},{"metricName":"Duplication ratio","quality":"Less is better","values":["1.275","1.233","1.138","1.178","1.211","1.333"],"isMain":true},{"metricName":"Largest alignment","quality":"More is better","values":[780,725,579,573,767,640],"isMain":true},{"metricName":"Total aligned length","quality":"More is better","values":[3571,7513,3067,7052,5904,2700],"isMain":true},{"metricName":"NA50","quality":"More is better","values":[508,335,518,501,508,480],"isMain":false},{"metricName":"NA75","quality":"More is better","values":[null,null,null,209,147,null],"isMain":false},{"metricName":"NGA50","quality":"More is better","values":[null,null,null,null,null,null],"isMain":true},{"metricName":"LA50","quality":"Less is better","values":[5,12,4,9,7,4],"isMain":false},{"metricName":"LA75","quality":"Less is better","values":[null,null,null,14,13,null],"isMain":false}]],["Reads mapping",[]],["Misassemblies",[{"metricName":"# misassemblies","quality":"Less is better","values":[0,1,0,0,0,0],"isMain":true},{"metricName":" # relocations","quality":"Less is better","values":[0,0,0,0,0,0],"isMain":false},{"metricName":" # translocations","quality":"Less is better","values":[0,1,0,0,0,0],"isMain":false},{"metricName":" # inversions","quality":"Less is better","values":[0,0,0,0,0,0],"isMain":false},{"metricName":" # interspecies translocations","quality":"Less is better","values":[4,9,2,3,5,3],"isMain":false},{"metricName":"# misassembled contigs","quality":"Less is better","values":[0,1,0,0,0,0],"isMain":false},{"metricName":"Misassembled contigs length","quality":"Less is better","values":[0,532,0,0,0,0],"isMain":true},{"metricName":"# possibly misassembled contigs","quality":"Less is better","values":[0,0,0,0,0,0],"isMain":false},{"metricName":" # possible misassemblies","quality":"Less is better","values":[0,0,0,0,0,0],"isMain":false},{"metricName":"# local misassemblies","quality":"Less is better","values":[1,1,0,0,1,0],"isMain":false},{"metricName":"# scaffold gap ext. mis.","quality":"Less is better","values":[0,0,0,0,0,0],"isMain":false},{"metricName":"# scaffold gap loc. mis.","quality":"Less is better","values":[0,0,0,0,0,0],"isMain":false},{"metricName":"# unaligned mis. contigs","quality":"Less is better","values":[0,0,0,1,0,0],"isMain":false}]],["Unaligned",[{"metricName":"# fully unaligned contigs","quality":"Less is better","values":[1,0,0,0,0,0],"isMain":false},{"metricName":"Fully unaligned length","quality":"Less is better","values":[513,0,0,0,0,0],"isMain":false},{"metricName":"# partially unaligned contigs","quality":"Less is better","values":[0,3,1,1,1,1],"isMain":false},{"metricName":"Partially unaligned length","quality":"Less is better","values":[0,2905,617,640,623,823],"isMain":false}]],["Mismatches",[{"metricName":"# mismatches","quality":"Less is better","values":[67,175,70,168,135,87],"isMain":false},{"metricName":"# indels","quality":"Less is better","values":[1,3,4,1,5,0],"isMain":false},{"metricName":"Indels length","quality":"Less is better","values":[1,3,4,1,5,0],"isMain":false},{"metricName":"# mismatches per 100 kbp","quality":"Less is better","values":["1876.75","2329.61","2285.34","2382.64","2286.97","3222.22"],"isMain":true},{"metricName":"# indels per 100 kbp","quality":"Less is better","values":["28.01","39.94","130.59","14.18","84.70","0.00"],"isMain":true},{"metricName":" # indels (<= 5 bp)","quality":"Less is better","values":[1,3,4,1,5,0],"isMain":false},{"metricName":" # indels (> 5 bp)","quality":"Less is better","values":[0,0,0,0,0,0],"isMain":false},{"metricName":"# N's","quality":"Less is better","values":[0,0,0,0,0,0],"isMain":false},{"metricName":"# N's per 100 kbp","quality":"Less is better","values":["0.00","0.00","0.00","0.00","0.00","0.00"],"isMain":true}]],["Statistics without reference",[{"metricName":"# contigs","quality":"Equal","values":[9,18,7,16,13,7],"isMain":true},{"metricName":"# contigs (>= 1000 bp)","quality":"Equal","values":[0,2,0,0,0,0],"isMain":false},{"metricName":"# contigs (>= 5000 bp)","quality":"Equal","values":[0,0,0,0,0,0],"isMain":false},{"metricName":"# contigs (>= 10000 bp)","quality":"Equal","values":[0,0,0,0,0,0],"isMain":false},{"metricName":"# contigs (>= 25000 bp)","quality":"Equal","values":[0,0,0,0,0,0],"isMain":false},{"metricName":"# contigs (>= 50000 bp)","quality":"Equal","values":[0,0,0,0,0,0],"isMain":false},{"metricName":"Largest contig","quality":"More is better","values":[784,1533,783,820,770,959],"isMain":true},{"metricName":"Total length","quality":"More is better","values":[5063,12168,4103,8947,7773,4422],"isMain":true},{"metricName":"Total length (>= 1000 bp)","quality":"More is better","values":[0,2576,0,0,0,0],"isMain":true},{"metricName":"Total length (>= 5000 bp)","quality":"More is better","values":[0,0,0,0,0,0],"isMain":false},{"metricName":"Total length (>= 10000 bp)","quality":"More is better","values":[0,0,0,0,0,0],"isMain":true},{"metricName":"Total length (>= 25000 bp)","quality":"More is better","values":[0,0,0,0,0,0],"isMain":false},{"metricName":"Total length (>= 50000 bp)","quality":"More is better","values":[0,0,0,0,0,0],"isMain":true},{"metricName":"N50","quality":"More is better","values":[532,615,566,531,571,644],"isMain":false},{"metricName":"N75","quality":"More is better","values":[509,532,519,511,520,555],"isMain":false},{"metricName":"L50","quality":"Less is better","values":[5,7,4,8,6,3],"isMain":false},{"metricName":"L75","quality":"Less is better","values":[7,13,6,12,10,5],"isMain":false},{"metricName":"GC (%)","quality":"Equal","values":["45.92","43.33","43.58","44.77","45.04","46.99"],"isMain":false}]],["K-mer-based statistics",[]],["Predicted genes",[]],["Similarity statistics",[]],["Reference statistics",[{"metricName":"Reference length","quality":"Equal","values":[3507873,3507873,3507873,3507873,3507873,3507873],"isMain":false},{"metricName":"Reference fragments","quality":"Equal","values":[73,73,73,73,73,73],"isMain":false},{"metricName":"Reference GC (%)","quality":"Equal","values":["44.85","44.85","44.85","44.85","44.85","44.85"],"isMain":false}]]],"subreferences":[],"subreports":[],"minContig":500},{"date":"10 February 2020, Monday, 18:00:49","assembliesNames":["megahit-103-df","megahit-113-df","megahit-113-ml","megahit-113-ms","megahit-129-df","metaSPAdes"],"referenceName":"GCF\_000157955.1\_ASM15795v1","order":[0,1,2,3,4,5],"report":[["Genome statistics",[{"metricName":"Genome fraction (%)","quality":"More is better","values":["76.342","80.075","75.796","75.693","82.512","68.179"],"isMain":true},{"metricName":"Duplication ratio","quality":"Less is better","values":["1.053","1.047","1.075","1.081","1.057","1.056"],"isMain":true},{"metricName":"Largest alignment","quality":"More is better","values":[5336,7018,6656,4128,5857,5524],"isMain":true},{"metricName":"Total aligned length","quality":"More is better","values":[2579587,2682984,2614238,2616694,2793224,2300103],"isMain":true},{"metricName":"NG50","quality":"More is better","values":[967,1250,841,892,1338,752],"isMain":false},{"metricName":"NG75","quality":"More is better","values":[579,697,569,597,803,null],"isMain":false},{"metricName":"NA50","quality":"More is better","values":[1119,1361,944,949,1332,900],"isMain":false},{"metricName":"NA75","quality":"More is better","values":[755,854,685,673,837,677],"isMain":false},{"metricName":"NGA50","quality":"More is better","values":[946,1178,821,834,1208,727],"isMain":true},{"metricName":"NGA75","quality":"More is better","values":[556,646,548,549,709,null],"isMain":false},{"metricName":"LG50","quality":"Less is better","values":[1070,815,1268,1163,743,1404],"isMain":false},{"metricName":"LG75","quality":"Less is better","values":[2156,1686,2439,2273,1528,null],"isMain":false},{"metricName":"LA50","quality":"Less is better","values":[789,669,966,983,718,890],"isMain":false},{"metricName":"LA75","quality":"Less is better","values":[1497,1309,1794,1842,1418,1641],"isMain":false},{"metricName":"LGA50","quality":"Less is better","values":[1090,858,1298,1265,835,1443],"isMain":true},{"metricName":"LGA75","quality":"Less is better","values":[2213,1793,2506,2460,1704,null],"isMain":false}]],["Reads mapping",[]],["Misassemblies",[{"metricName":"# misassemblies","quality":"Less is better","values":[16,16,12,15,14,10],"isMain":true},{"metricName":" # relocations","quality":"Less is better","values":[1,3,2,3,1,3],"isMain":false},{"metricName":" # translocations","quality":"Less is better","values":[14,13,8,11,13,7],"isMain":false},{"metricName":" # inversions","quality":"Less is better","values":[1,0,2,1,0,0],"isMain":false},{"metricName":" # interspecies translocations","quality":"Less is better","values":[66,81,32,81,89,44],"isMain":false},{"metricName":"# misassembled contigs","quality":"Less is better","values":[16,16,12,15,14,10],"isMain":false},{"metricName":"Misassembled contigs length","quality":"Less is better","values":[20457,26313,16145,18009,23233,15208],"isMain":true},{"metricName":"# possibly misassembled contigs","quality":"Less is better","values":[0,0,0,1,0,0],"isMain":false},{"metricName":" # possible misassemblies","quality":"Less is better","values":[0,0,0,1,0,0],"isMain":false},{"metricName":"# local misassemblies","quality":"Less is better","values":[1,1,6,3,3,1],"isMain":false},{"metricName":"# scaffold gap ext. mis.","quality":"Less is better","values":[0,0,0,0,0,0],"isMain":false},{"metricName":"# scaffold gap loc. mis.","quality":"Less is better","values":[0,0,0,0,0,0],"isMain":false},{"metricName":"# unaligned mis. contigs","quality":"Less is better","values":[0,0,0,1,0,0],"isMain":false}]],["Unaligned",[{"metricName":"# fully unaligned contigs","quality":"Less is better","values":[2,3,1,4,3,1],"isMain":false},{"metricName":"Fully unaligned length","quality":"Less is better","values":[2658,2631,586,3761,4024,678],"isMain":false},{"metricName":"# partially unaligned contigs","quality":"Less is better","values":[14,21,11,25,30,7],"isMain":false},{"metricName":"Partially unaligned length","quality":"Less is better","values":[11062,43427,14400,82635,114621,10913],"isMain":false}]],["Mismatches",[{"metricName":"# mismatches","quality":"Less is better","values":[26269,25507,42784,44381,27760,25133],"isMain":false},{"metricName":"# indels","quality":"Less is better","values":[57,68,52,70,78,52],"isMain":false},{"metricName":"Indels length","quality":"Less is better","values":[110,127,101,156,244,147],"isMain":false},{"metricName":"# mismatches per 100 kbp","quality":"Less is better","values":["1062.86","983.91","1743.53","1811.07","1039.19","1138.64"],"isMain":true},{"metricName":"# indels per 100 kbp","quality":"Less is better","values":["2.31","2.62","2.12","2.86","2.92","2.36"],"isMain":true},{"metricName":" # indels (<= 5 bp)","quality":"Less is better","values":[56,66,51,67,71,50],"isMain":false},{"metricName":" # indels (> 5 bp)","quality":"Less is better","values":[1,2,1,3,7,2],"isMain":false},{"metricName":"# N's","quality":"Less is better","values":[0,0,0,0,0,0],"isMain":false},{"metricName":"# N's per 100 kbp","quality":"Less is better","values":["0.00","0.00","0.00","0.00","0.00","0.00"],"isMain":true}]],["Statistics without reference",[{"metricName":"# contigs","quality":"Equal","values":[2503,2244,2864,2836,2331,2587],"isMain":true},{"metricName":"# contigs (>= 1000 bp)","quality":"Equal","values":[1029,1110,868,917,1166,724],"isMain":false},{"metricName":"# contigs (>= 5000 bp)","quality":"Equal","values":[1,5,2,6,10,1],"isMain":false},{"metricName":"# contigs (>= 10000 bp)","quality":"Equal","values":[0,1,0,3,3,0],"isMain":false},{"metricName":"# contigs (>= 25000 bp)","quality":"Equal","values":[0,0,0,1,1,0],"isMain":false},{"metricName":"# contigs (>= 50000 bp)","quality":"Equal","values":[0,0,0,0,0,0],"isMain":false},{"metricName":"Largest contig","quality":"More is better","values":[5360,14682,7803,27897,27759,5527],"isMain":true},{"metricName":"Total length","quality":"More is better","values":[2616165,2759714,2653965,2736330,2942268,2343410],"isMain":true},{"metricName":"Total length (>= 1000 bp)","quality":"More is better","values":[1578596,1948853,1253748,1387072,2105854,1033804],"isMain":true},{"metricName":"Total length (>= 5000 bp)","quality":"More is better","values":[5360,43116,15450,70851,107426,5527],"isMain":false},{"metricName":"Total length (>= 10000 bp)","quality":"More is better","values":[0,14682,0,53154,60130,0],"isMain":true},{"metricName":"Total length (>= 25000 bp)","quality":"More is better","values":[0,0,0,27897,27759,0],"isMain":false},{"metricName":"Total length (>= 50000 bp)","quality":"More is better","values":[0,0,0,0,0,0],"isMain":true},{"metricName":"N50","quality":"More is better","values":[1143,1427,959,1009,1477,929],"isMain":false},{"metricName":"N75","quality":"More is better","values":[782,915,704,724,931,703],"isMain":false},{"metricName":"L50","quality":"Less is better","values":[777,636,943,899,638,868],"isMain":false},{"metricName":"L75","quality":"Less is better","values":[1465,1237,1750,1702,1271,1595],"isMain":false},{"metricName":"GC (%)","quality":"Equal","values":["58.02","58.01","57.85","57.69","58.03","57.99"],"isMain":false}]],["K-mer-based statistics",[]],["Predicted genes",[]],["Similarity statistics",[]],["Reference statistics",[{"metricName":"Reference length","quality":"Equal","values":[3237471,3237471,3237471,3237471,3237471,3237471],"isMain":false},{"metricName":"Reference fragments","quality":"Equal","values":[91,91,91,91,91,91],"isMain":false},{"metricName":"Reference GC (%)","quality":"Equal","values":["57.92","57.92","57.92","57.92","57.92","57.92"],"isMain":false}]]],"subreferences":[],"subreports":[],"minContig":500},{"date":"10 February 2020, Monday, 18:00:48","assembliesNames":["megahit-103-df","megahit-113-df","megahit-113-ml","megahit-113-ms","megahit-129-df","metaSPAdes"],"referenceName":"GCF\_000157975.1\_ASM15797v1","order":[0,1,2,3,4,5],"report":[["Genome statistics",[{"metricName":"Genome fraction (%)","quality":"More is better","values":["98.877","99.164","99.498","99.347","99.509","99.025"],"isMain":true},{"metricName":"Duplication ratio","quality":"Less is better","values":["1.004","1.002","1.003","1.002","1.003","1.000"],"isMain":true},{"metricName":"Largest alignment","quality":"More is better","values":[72981,128761,128763,80422,128571,127467],"isMain":true},{"metricName":"Total aligned length","quality":"More is better","values":[3534138,3540041,3554654,3546565,3555064,3531154],"isMain":true},{"metricName":"NG50","quality":"More is better","values":[20533,24974,26480,29486,24964,24737],"isMain":false},{"metricName":"NG75","quality":"More is better","values":[10686,14902,14907,15996,14904,14032],"isMain":false},{"metricName":"NA50","quality":"More is better","values":[20284,24862,24862,23562,24862,24514],"isMain":false},{"metricName":"NA75","quality":"More is better","values":[10550,14261,14011,11725,14021,13835],"isMain":false},{"metricName":"NGA50","quality":"More is better","values":[20284,24862,24862,24097,24862,24124],"isMain":true},{"metricName":"NGA75","quality":"More is better","values":[10525,14069,14077,12857,14021,13683],"isMain":false},{"metricName":"LG50","quality":"Less is better","values":[53,39,38,37,39,40],"isMain":false},{"metricName":"LG75","quality":"Less is better","values":[113,82,81,79,82,85],"isMain":false},{"metricName":"LA50","quality":"Less is better","values":[54,40,40,46,40,41],"isMain":false},{"metricName":"LA75","quality":"Less is better","values":[114,83,85,99,85,87],"isMain":false},{"metricName":"LGA50","quality":"Less is better","values":[54,40,40,43,40,42],"isMain":true},{"metricName":"LGA75","quality":"Less is better","values":[115,84,84,90,85,88],"isMain":false}]],["Reads mapping",[]],["Misassemblies",[{"metricName":"# misassemblies","quality":"Less is better","values":[7,12,10,10,11,9],"isMain":true},{"metricName":" # relocations","quality":"Less is better","values":[0,0,1,0,0,0],"isMain":false},{"metricName":" # translocations","quality":"Less is better","values":[7,11,9,9,10,9],"isMain":false},{"metricName":" # inversions","quality":"Less is better","values":[0,1,0,1,1,0],"isMain":false},{"metricName":" # interspecies translocations","quality":"Less is better","values":[14,8,11,16,12,0],"isMain":false},{"metricName":"# misassembled contigs","quality":"Less is better","values":[7,12,10,10,10,8],"isMain":false},{"metricName":"Misassembled contigs length","quality":"Less is better","values":[158448,231998,279485,197607,195034,232636],"isMain":true},{"metricName":"# possibly misassembled contigs","quality":"Less is better","values":[0,0,0,0,0,0],"isMain":false},{"metricName":" # possible misassemblies","quality":"Less is better","values":[0,0,0,0,0,0],"isMain":false},{"metricName":"# local misassemblies","quality":"Less is better","values":[0,0,3,0,0,0],"isMain":false},{"metricName":"# scaffold gap ext. mis.","quality":"Less is better","values":[0,0,0,0,0,0],"isMain":false},{"metricName":"# scaffold gap loc. mis.","quality":"Less is better","values":[0,0,0,0,0,0],"isMain":false},{"metricName":"# unaligned mis. contigs","quality":"Less is better","values":[0,0,0,0,0,0],"isMain":false}]],["Unaligned",[{"metricName":"# fully unaligned contigs","quality":"Less is better","values":[0,0,0,0,1,0],"isMain":false},{"metricName":"Fully unaligned length","quality":"Less is better","values":[0,0,0,0,580,0],"isMain":false},{"metricName":"# partially unaligned contigs","quality":"Less is better","values":[5,3,5,13,5,0],"isMain":false},{"metricName":"Partially unaligned length","quality":"Less is better","values":[4271,2707,17000,156524,3190,0],"isMain":false}]],["Mismatches",[{"metricName":"# mismatches","quality":"Less is better","values":[3788,435,691,727,534,288],"isMain":false},{"metricName":"# indels","quality":"Less is better","values":[0,4,18,18,5,14],"isMain":false},{"metricName":"Indels length","quality":"Less is better","values":[0,81,22,208,95,82],"isMain":false},{"metricName":"# mismatches per 100 kbp","quality":"Less is better","values":["107.45","12.30","19.48","20.52","15.05","8.16"],"isMain":true},{"metricName":"# indels per 100 kbp","quality":"Less is better","values":["0.00","0.11","0.51","0.51","0.14","0.40"],"isMain":true},{"metricName":" # indels (<= 5 bp)","quality":"Less is better","values":[0,2,18,15,2,12],"isMain":false},{"metricName":" # indels (> 5 bp)","quality":"Less is better","values":[0,2,0,3,3,2],"isMain":false},{"metricName":"# N's","quality":"Less is better","values":[0,0,0,0,0,0],"isMain":false},{"metricName":"# N's per 100 kbp","quality":"Less is better","values":["0.00","0.00","0.00","0.00","0.00","0.00"],"isMain":true}]],["Statistics without reference",[{"metricName":"# contigs","quality":"Equal","values":[338,252,258,258,260,248],"isMain":true},{"metricName":"# contigs (>= 1000 bp)","quality":"Equal","values":[291,229,232,238,231,234],"isMain":false},{"metricName":"# contigs (>= 5000 bp)","quality":"Equal","values":[189,156,158,167,159,160],"isMain":false},{"metricName":"# contigs (>= 10000 bp)","quality":"Equal","values":[122,111,111,116,111,111],"isMain":false},{"metricName":"# contigs (>= 25000 bp)","quality":"Equal","values":[34,38,38,40,38,37],"isMain":false},{"metricName":"# contigs (>= 50000 bp)","quality":"Equal","values":[9,13,13,15,14,13],"isMain":false},{"metricName":"Largest contig","quality":"More is better","values":[72982,128761,128763,102797,128571,127467],"isMain":true},{"metricName":"Total length","quality":"More is better","values":[3542507,3544803,3574840,3705378,3561627,3531535],"isMain":true},{"metricName":"Total length (>= 1000 bp)","quality":"More is better","values":[3510698,3527584,3556908,3690769,3541856,3520857],"isMain":true},{"metricName":"Total length (>= 5000 bp)","quality":"More is better","values":[3254333,3352263,3380789,3507867,3364323,3338714],"isMain":false},{"metricName":"Total length (>= 10000 bp)","quality":"More is better","values":[2772072,3027603,3034313,3134547,3020343,2978214],"isMain":true},{"metricName":"Total length (>= 25000 bp)","quality":"More is better","values":[1374905,1782664,1795516,1869688,1779852,1719445],"isMain":false},{"metricName":"Total length (>= 50000 bp)","quality":"More is better","values":[548314,869192,881079,952800,905343,868607],"isMain":true},{"metricName":"N50","quality":"More is better","values":[20558,26480,26480,25947,24964,24798],"isMain":false},{"metricName":"N75","quality":"More is better","values":[10704,15717,14285,13708,14904,14077],"isMain":false},{"metricName":"L50","quality":"Less is better","values":[52,38,38,40,39,39],"isMain":false},{"metricName":"L75","quality":"Less is better","values":[112,81,82,86,82,84],"isMain":false},{"metricName":"GC (%)","quality":"Equal","values":["45.03","45.03","45.04","45.02","45.06","45.01"],"isMain":false}]],["K-mer-based statistics",[]],["Predicted genes",[]],["Similarity statistics",[]],["Reference statistics",[{"metricName":"Reference length","quality":"Equal","values":[3565428,3565428,3565428,3565428,3565428,3565428],"isMain":false},{"metricName":"Reference fragments","quality":"Equal","values":[233,233,233,233,233,233],"isMain":false},{"metricName":"Reference GC (%)","quality":"Equal","values":["45.07","45.07","45.07","45.07","45.07","45.07"],"isMain":false}]]],"subreferences":[],"subreports":[],"minContig":500},{"date":"10 February 2020, Monday, 18:02:04","assembliesNames":["megahit-103-df","megahit-113-df","megahit-113-ml","megahit-113-ms","megahit-129-df","metaSPAdes"],"referenceName":"GCF\_000158075.1\_ASM15807v1","order":[0,1,2,3,4,5],"report":[["Genome statistics",[{"metricName":"Genome fraction (%)","quality":"More is better","values":["99.105","99.390","99.467","99.459","99.542","99.043"],"isMain":true},{"metricName":"Duplication ratio","quality":"Less is better","values":["1.009","1.003","1.002","1.005","1.003","1.001"],"isMain":true},{"metricName":"Largest alignment","quality":"More is better","values":[58574,140988,140992,140992,140992,140992],"isMain":true},{"metricName":"Total aligned length","quality":"More is better","values":[6212638,6194646,6199635,6210049,6206270,6170951],"isMain":true},{"metricName":"NG50","quality":"More is better","values":[11637,17356,17456,19198,17393,17366],"isMain":false},{"metricName":"NG75","quality":"More is better","values":[6182,9140,9140,10339,9138,8987],"isMain":false},{"metricName":"NA50","quality":"More is better","values":[11595,17131,17040,15795,16839,16924],"isMain":false},{"metricName":"NA75","quality":"More is better","values":[5864,8852,8799,7367,8575,8594],"isMain":false},{"metricName":"NGA50","quality":"More is better","values":[11602,17228,17203,17131,16948,17006],"isMain":true},{"metricName":"NGA75","quality":"More is better","values":[5906,8874,8860,8582,8706,8630],"isMain":false},{"metricName":"LG50","quality":"Less is better","values":[157,98,95,87,95,96],"isMain":false},{"metricName":"LG75","quality":"Less is better","values":[341,220,217,198,218,221],"isMain":false},{"metricName":"LA50","quality":"Less is better","values":[159,99,99,109,100,101],"isMain":false},{"metricName":"LA75","quality":"Less is better","values":[347,225,227,261,231,232],"isMain":false},{"metricName":"LGA50","quality":"Less is better","values":[158,98,97,99,98,100],"isMain":true},{"metricName":"LGA75","quality":"Less is better","values":[345,224,224,229,226,231],"isMain":false}]],["Reads mapping",[]],["Misassemblies",[{"metricName":"# misassemblies","quality":"Less is better","values":[14,15,8,17,18,16],"isMain":true},{"metricName":" # relocations","quality":"Less is better","values":[1,0,0,0,0,0],"isMain":false},{"metricName":" # translocations","quality":"Less is better","values":[11,15,8,17,18,16],"isMain":false},{"metricName":" # inversions","quality":"Less is better","values":[2,0,0,0,0,0],"isMain":false},{"metricName":" # interspecies translocations","quality":"Less is better","values":[46,25,14,42,23,6],"isMain":false},{"metricName":"# misassembled contigs","quality":"Less is better","values":[14,14,7,16,17,15],"isMain":false},{"metricName":"Misassembled contigs length","quality":"Less is better","values":[99046,127361,111751,178157,176718,202237],"isMain":true},{"metricName":"# possibly misassembled contigs","quality":"Less is better","values":[0,0,1,0,0,0],"isMain":false},{"metricName":" # possible misassemblies","quality":"Less is better","values":[0,0,1,0,0,0],"isMain":false},{"metricName":"# local misassemblies","quality":"Less is better","values":[3,0,1,3,0,3],"isMain":false},{"metricName":"# scaffold gap ext. mis.","quality":"Less is better","values":[0,0,0,0,0,0],"isMain":false},{"metricName":"# scaffold gap loc. mis.","quality":"Less is better","values":[0,0,0,0,0,0],"isMain":false},{"metricName":"# unaligned mis. contigs","quality":"Less is better","values":[0,0,0,1,0,0],"isMain":false}]],["Unaligned",[{"metricName":"# fully unaligned contigs","quality":"Less is better","values":[0,0,0,1,0,0],"isMain":false},{"metricName":"Fully unaligned length","quality":"Less is better","values":[0,0,0,595,0,0],"isMain":false},{"metricName":"# partially unaligned contigs","quality":"Less is better","values":[19,11,8,25,11,4],"isMain":false},{"metricName":"Partially unaligned length","quality":"Less is better","values":[14564,31781,56204,337816,65318,63159],"isMain":false}]],["Mismatches",[{"metricName":"# mismatches","quality":"Less is better","values":[10387,916,749,2186,1092,850],"isMain":false},{"metricName":"# indels","quality":"Less is better","values":[1,8,23,63,20,19],"isMain":false},{"metricName":"Indels length","quality":"Less is better","values":[1,194,184,430,599,43],"isMain":false},{"metricName":"# mismatches per 100 kbp","quality":"Less is better","values":["168.38","14.81","12.10","35.31","17.62","13.79"],"isMain":true},{"metricName":"# indels per 100 kbp","quality":"Less is better","values":["0.02","0.13","0.37","1.02","0.32","0.31"],"isMain":true},{"metricName":" # indels (<= 5 bp)","quality":"Less is better","values":[1,3,18,54,5,18],"isMain":false},{"metricName":" # indels (> 5 bp)","quality":"Less is better","values":[0,5,5,9,15,1],"isMain":false},{"metricName":"# N's","quality":"Less is better","values":[0,0,0,0,0,0],"isMain":false},{"metricName":"# N's per 100 kbp","quality":"Less is better","values":["0.00","0.00","0.00","0.00","0.00","0.00"],"isMain":true}]],["Statistics without reference",[{"metricName":"# contigs","quality":"Equal","values":[978,661,666,678,667,647],"isMain":true},{"metricName":"# contigs (>= 1000 bp)","quality":"Equal","values":[830,585,595,603,588,600],"isMain":false},{"metricName":"# contigs (>= 5000 bp)","quality":"Equal","values":[397,334,333,349,340,336],"isMain":false},{"metricName":"# contigs (>= 10000 bp)","quality":"Equal","values":[200,206,204,207,205,201],"isMain":false},{"metricName":"# contigs (>= 25000 bp)","quality":"Equal","values":[27,53,55,59,54,50],"isMain":false},{"metricName":"# contigs (>= 50000 bp)","quality":"Equal","values":[4,11,11,14,10,12],"isMain":false},{"metricName":"Largest contig","quality":"More is better","values":[58574,140991,140995,140995,140995,168904],"isMain":true},{"metricName":"Total length","quality":"More is better","values":[6239564,6233789,6261124,6562410,6279542,6236841],"isMain":true},{"metricName":"Total length (>= 1000 bp)","quality":"More is better","values":[6137365,6180135,6211980,6509533,6226163,6203018],"isMain":true},{"metricName":"Total length (>= 5000 bp)","quality":"More is better","values":[4982777,5473010,5479637,5795979,5531423,5464700],"isMain":false},{"metricName":"Total length (>= 10000 bp)","quality":"More is better","values":[3579178,4538911,4547811,4766396,4547431,4482353],"isMain":true},{"metricName":"Total length (>= 25000 bp)","quality":"More is better","values":[959315,2215846,2310836,2503042,2266810,2197924],"isMain":false},{"metricName":"Total length (>= 50000 bp)","quality":"More is better","values":[228015,854867,869148,1073994,810993,984506],"isMain":true},{"metricName":"N50","quality":"More is better","values":[11612,17356,17395,18093,17320,17366],"isMain":false},{"metricName":"N75","quality":"More is better","values":[6154,9088,9059,9077,8979,8955],"isMain":false},{"metricName":"L50","quality":"Less is better","values":[158,98,96,96,97,96],"isMain":false},{"metricName":"L75","quality":"Less is better","values":[343,221,220,224,223,222],"isMain":false},{"metricName":"GC (%)","quality":"Equal","values":["55.60","55.64","55.57","55.34","55.53","55.55"],"isMain":false}]],["K-mer-based statistics",[]],["Predicted genes",[]],["Similarity statistics",[]],["Reference statistics",[{"metricName":"Reference length","quality":"Equal","values":[6224391,6224391,6224391,6224391,6224391,6224391],"isMain":false},{"metricName":"Reference fragments","quality":"Equal","values":[640,640,640,640,640,640],"isMain":false},{"metricName":"Reference GC (%)","quality":"Equal","values":["55.60","55.60","55.60","55.60","55.60","55.60"],"isMain":false}]]],"subreferences":[],"subreports":[],"minContig":500},{"date":"10 February 2020, Monday, 18:00:15","assembliesNames":["megahit-103-df","megahit-113-df","megahit-113-ml","megahit-113-ms","megahit-129-df","metaSPAdes"],"referenceName":"GCF\_000158555.2\_Erys\_bact\_D7\_V2","order":[0,1,2,3,4,5],"report":[["Genome statistics",[{"metricName":"Genome fraction (%)","quality":"More is better","values":["3.239","4.519","5.163","6.230","5.013","2.437"],"isMain":true},{"metricName":"Duplication ratio","quality":"Less is better","values":["1.106","1.090","1.024","1.046","1.075","1.073"],"isMain":true},{"metricName":"Largest alignment","quality":"More is better","values":[984,1150,1146,1499,1102,1330],"isMain":true},{"metricName":"Total aligned length","quality":"More is better","values":[113868,159388,181935,219650,176555,85709],"isMain":true},{"metricName":"NA50","quality":"More is better","values":[566,566,582,594,569,540],"isMain":false},{"metricName":"NA75","quality":"More is better","values":[501,501,530,522,505,497],"isMain":false},{"metricName":"NGA50","quality":"More is better","values":[null,null,null,null,null,null],"isMain":true},{"metricName":"LA50","quality":"Less is better","values":[98,136,133,162,147,78],"isMain":false},{"metricName":"LA75","quality":"Less is better","values":[158,223,217,270,240,126],"isMain":false}]],["Reads mapping",[]],["Misassemblies",[{"metricName":"# misassemblies","quality":"Less is better","values":[1,4,2,1,3,1],"isMain":true},{"metricName":" # relocations","quality":"Less is better","values":[0,0,1,0,0,0],"isMain":false},{"metricName":" # translocations","quality":"Less is better","values":[1,4,1,1,3,1],"isMain":false},{"metricName":" # inversions","quality":"Less is better","values":[0,0,0,0,0,0],"isMain":false},{"metricName":" # interspecies translocations","quality":"Less is better","values":[44,57,8,33,49,27],"isMain":false},{"metricName":"# misassembled contigs","quality":"Less is better","values":[1,4,2,1,3,1],"isMain":false},{"metricName":"Misassembled contigs length","quality":"Less is better","values":[607,2534,1315,1045,1780,620],"isMain":true},{"metricName":"# possibly misassembled contigs","quality":"Less is better","values":[0,0,0,0,1,0],"isMain":false},{"metricName":" # possible misassemblies","quality":"Less is better","values":[0,0,0,0,1,0],"isMain":false},{"metricName":"# local misassemblies","quality":"Less is better","values":[0,1,0,1,0,0],"isMain":false},{"metricName":"# scaffold gap ext. mis.","quality":"Less is better","values":[0,0,0,0,0,0],"isMain":false},{"metricName":"# scaffold gap loc. mis.","quality":"Less is better","values":[0,0,0,0,0,0],"isMain":false},{"metricName":"# unaligned mis. contigs","quality":"Less is better","values":[0,1,0,0,0,0],"isMain":false}]],["Unaligned",[{"metricName":"# fully unaligned contigs","quality":"Less is better","values":[0,1,0,1,3,0],"isMain":false},{"metricName":"Fully unaligned length","quality":"Less is better","values":[0,569,0,883,1925,0],"isMain":false},{"metricName":"# partially unaligned contigs","quality":"Less is better","values":[4,13,0,7,13,10],"isMain":false},{"metricName":"Partially unaligned length","quality":"Less is better","values":[2612,10057,0,9947,10415,7568],"isMain":false}]],["Mismatches",[{"metricName":"# mismatches","quality":"Less is better","values":[2033,2777,3530,4534,3102,1583],"isMain":false},{"metricName":"# indels","quality":"Less is better","values":[6,9,5,13,4,0],"isMain":false},{"metricName":"Indels length","quality":"Less is better","values":[11,10,5,13,4,0],"isMain":false},{"metricName":"# mismatches per 100 kbp","quality":"Less is better","values":["1785.89","1748.32","1945.43","2070.65","1760.78","1848.54"],"isMain":true},{"metricName":"# indels per 100 kbp","quality":"Less is better","values":["5.27","5.67","2.76","5.94","2.27","0.00"],"isMain":true},{"metricName":" # indels (<= 5 bp)","quality":"Less is better","values":[6,9,5,13,4,0],"isMain":false},{"metricName":" # indels (> 5 bp)","quality":"Less is better","values":[0,0,0,0,0,0],"isMain":false},{"metricName":"# N's","quality":"Less is better","values":[0,0,0,0,0,0],"isMain":false},{"metricName":"# N's per 100 kbp","quality":"Less is better","values":["0.00","0.00","0.00","0.00","0.00","0.00"],"isMain":true}]],["Statistics without reference",[{"metricName":"# contigs","quality":"Equal","values":[209,288,303,363,313,161],"isMain":true},{"metricName":"# contigs (>= 1000 bp)","quality":"Equal","values":[2,9,4,13,8,5],"isMain":false},{"metricName":"# contigs (>= 5000 bp)","quality":"Equal","values":[0,0,0,1,0,0],"isMain":false},{"metricName":"# contigs (>= 10000 bp)","quality":"Equal","values":[0,0,0,0,0,0],"isMain":false},{"metricName":"# contigs (>= 25000 bp)","quality":"Equal","values":[0,0,0,0,0,0],"isMain":false},{"metricName":"# contigs (>= 50000 bp)","quality":"Equal","values":[0,0,0,0,0,0],"isMain":false},{"metricName":"Largest contig","quality":"More is better","values":[1341,2289,1146,5435,1483,1780],"isMain":true},{"metricName":"Total length","quality":"More is better","values":[128572,183735,185738,239823,201675,99436],"isMain":true},{"metricName":"Total length (>= 1000 bp)","quality":"More is better","values":[2368,11326,4188,19906,9623,6435],"isMain":true},{"metricName":"Total length (>= 5000 bp)","quality":"More is better","values":[0,0,0,5435,0,0],"isMain":false},{"metricName":"Total length (>= 10000 bp)","quality":"More is better","values":[0,0,0,0,0,0],"isMain":true},{"metricName":"Total length (>= 25000 bp)","quality":"More is better","values":[0,0,0,0,0,0],"isMain":false},{"metricName":"Total length (>= 50000 bp)","quality":"More is better","values":[0,0,0,0,0,0],"isMain":true},{"metricName":"N50","quality":"More is better","values":[592,621,600,622,627,571],"isMain":false},{"metricName":"N75","quality":"More is better","values":[546,543,539,553,560,534],"isMain":false},{"metricName":"L50","quality":"Less is better","values":[91,121,132,147,132,68],"isMain":false},{"metricName":"L75","quality":"Less is better","values":[148,200,214,250,218,113],"isMain":false},{"metricName":"GC (%)","quality":"Equal","values":["32.56","32.55","32.08","32.38","32.40","32.24"],"isMain":false}]],["K-mer-based statistics",[]],["Predicted genes",[]],["Similarity statistics",[]],["Reference statistics",[{"metricName":"Reference length","quality":"Equal","values":[3514630,3514630,3514630,3514630,3514630,3514630],"isMain":false},{"metricName":"Reference fragments","quality":"Equal","values":[168,168,168,168,168,168],"isMain":false},{"metricName":"Reference GC (%)","quality":"Equal","values":["31.40","31.40","31.40","31.40","31.40","31.40"],"isMain":false}]]],"subreferences":[],"subreports":[],"minContig":500},{"date":"10 February 2020, Monday, 18:01:49","assembliesNames":["megahit-103-df","megahit-113-df","megahit-113-ml","megahit-113-ms","megahit-129-df","metaSPAdes"],"referenceName":"GCF\_000158655.1\_ASM15865v1","order":[0,1,2,3,4,5],"report":[["Genome statistics",[{"metricName":"Genome fraction (%)","quality":"More is better","values":["64.585","69.699","62.368","64.757","72.558","55.481"],"isMain":true},{"metricName":"Duplication ratio","quality":"Less is better","values":["1.038","1.036","1.058","1.060","1.036","1.043"],"isMain":true},{"metricName":"Largest alignment","quality":"More is better","values":[3202,4158,3007,3806,4348,2689],"isMain":true},{"metricName":"Total aligned length","quality":"More is better","values":[2267466,2438788,2226085,2312914,2544377,1946634],"isMain":true},{"metricName":"NG50","quality":"More is better","values":[691,821,634,672,879,563],"isMain":false},{"metricName":"NG75","quality":"More is better","values":[null,null,null,null,516,null],"isMain":false},{"metricName":"NA50","quality":"More is better","values":[910,1007,803,823,1065,773],"isMain":false},{"metricName":"NA75","quality":"More is better","values":[674,734,626,632,747,608],"isMain":false},{"metricName":"NGA50","quality":"More is better","values":[677,800,621,654,851,547],"isMain":true},{"metricName":"LG50","quality":"Less is better","values":[1575,1309,1806,1683,1211,1988],"isMain":false},{"metricName":"LG75","quality":"Less is better","values":[null,null,null,null,2469,null],"isMain":false},{"metricName":"LA50","quality":"Less is better","values":[883,824,1010,1015,820,923],"isMain":false},{"metricName":"LA75","quality":"Less is better","values":[1617,1545,1805,1838,1549,1643],"isMain":false},{"metricName":"LGA50","quality":"Less is better","values":[1596,1341,1831,1732,1251,2029],"isMain":true}]],["Reads mapping",[]],["Misassemblies",[{"metricName":"# misassemblies","quality":"Less is better","values":[6,11,8,3,13,12],"isMain":true},{"metricName":" # relocations","quality":"Less is better","values":[0,1,1,0,1,0],"isMain":false},{"metricName":" # translocations","quality":"Less is better","values":[6,10,3,3,12,12],"isMain":false},{"metricName":" # inversions","quality":"Less is better","values":[0,0,4,0,0,0],"isMain":false},{"metricName":" # interspecies translocations","quality":"Less is better","values":[37,53,16,34,32,28],"isMain":false},{"metricName":"# misassembled contigs","quality":"Less is better","values":[6,11,8,3,13,12],"isMain":false},{"metricName":"Misassembled contigs length","quality":"Less is better","values":[6935,16483,6015,2583,17730,9780],"isMain":true},{"metricName":"# possibly misassembled contigs","quality":"Less is better","values":[0,0,0,1,0,0],"isMain":false},{"metricName":" # possible misassemblies","quality":"Less is better","values":[0,0,0,1,0,0],"isMain":false},{"metricName":"# local misassemblies","quality":"Less is better","values":[0,4,3,3,3,1],"isMain":false},{"metricName":"# scaffold gap ext. mis.","quality":"Less is better","values":[0,0,0,0,0,0],"isMain":false},{"metricName":"# scaffold gap loc. mis.","quality":"Less is better","values":[0,0,0,0,0,0],"isMain":false},{"metricName":"# unaligned mis. contigs","quality":"Less is better","values":[0,0,0,0,0,0],"isMain":false}]],["Unaligned",[{"metricName":"# fully unaligned contigs","quality":"Less is better","values":[0,1,0,0,0,0],"isMain":false},{"metricName":"Fully unaligned length","quality":"Less is better","values":[0,588,0,0,0,0],"isMain":false},{"metricName":"# partially unaligned contigs","quality":"Less is better","values":[6,9,3,8,6,4],"isMain":false},{"metricName":"Partially unaligned length","quality":"Less is better","values":[5523,17599,1723,22747,25900,3285],"isMain":false}]],["Mismatches",[{"metricName":"# mismatches","quality":"Less is better","values":[25300,25531,39070,41544,26903,23674],"isMain":false},{"metricName":"# indels","quality":"Less is better","values":[32,37,43,51,41,31],"isMain":false},{"metricName":"Indels length","quality":"Less is better","values":[43,61,68,83,240,49],"isMain":false},{"metricName":"# mismatches per 100 kbp","quality":"Less is better","values":["1150.02","1075.36","1839.06","1883.36","1088.51","1252.68"],"isMain":true},{"metricName":"# indels per 100 kbp","quality":"Less is better","values":["1.45","1.56","2.02","2.31","1.66","1.64"],"isMain":true},{"metricName":" # indels (<= 5 bp)","quality":"Less is better","values":[31,34,41,49,35,29],"isMain":false},{"metricName":" # indels (> 5 bp)","quality":"Less is better","values":[1,3,2,2,6,2],"isMain":false},{"metricName":"# N's","quality":"Less is better","values":[0,0,0,0,0,0],"isMain":false},{"metricName":"# N's per 100 kbp","quality":"Less is better","values":["0.00","0.00","0.00","0.00","0.00","0.00"],"isMain":true}]],["Statistics without reference",[{"metricName":"# contigs","quality":"Equal","values":[2571,2506,2779,2825,2533,2501],"isMain":true},{"metricName":"# contigs (>= 1000 bp)","quality":"Equal","values":[694,856,556,600,935,450],"isMain":false},{"metricName":"# contigs (>= 5000 bp)","quality":"Equal","values":[0,1,0,1,1,0],"isMain":false},{"metricName":"# contigs (>= 10000 bp)","quality":"Equal","values":[0,0,0,1,1,0],"isMain":false},{"metricName":"# contigs (>= 25000 bp)","quality":"Equal","values":[0,0,0,0,0,0],"isMain":false},{"metricName":"# contigs (>= 50000 bp)","quality":"Equal","values":[0,0,0,0,0,0],"isMain":false},{"metricName":"Largest contig","quality":"More is better","values":[4223,9458,3018,18084,19074,2689],"isMain":true},{"metricName":"Total length","quality":"More is better","values":[2289503,2478796,2249620,2361536,2587400,1974944],"isMain":true},{"metricName":"Total length (>= 1000 bp)","quality":"More is better","values":[973113,1293349,727695,828639,1444954,581918],"isMain":true},{"metricName":"Total length (>= 5000 bp)","quality":"More is better","values":[0,9458,0,18084,19074,0],"isMain":false},{"metricName":"Total length (>= 10000 bp)","quality":"More is better","values":[0,0,0,18084,19074,0],"isMain":true},{"metricName":"Total length (>= 25000 bp)","quality":"More is better","values":[0,0,0,0,0,0],"isMain":false},{"metricName":"Total length (>= 50000 bp)","quality":"More is better","values":[0,0,0,0,0,0],"isMain":true},{"metricName":"N50","quality":"More is better","values":[924,1030,813,845,1090,792],"isMain":false},{"metricName":"N75","quality":"More is better","values":[685,757,639,653,769,622],"isMain":false},{"metricName":"L50","quality":"Less is better","values":[873,803,1000,984,791,909],"isMain":false},{"metricName":"L75","quality":"Less is better","values":[1595,1507,1781,1786,1500,1613],"isMain":false},{"metricName":"GC (%)","quality":"Equal","values":["51.84","51.87","51.88","51.88","51.88","51.87"],"isMain":false}]],["K-mer-based statistics",[]],["Predicted genes",[]],["Similarity statistics",[]],["Reference statistics",[{"metricName":"Reference length","quality":"Equal","values":[3406326,3406326,3406326,3406326,3406326,3406326],"isMain":false},{"metricName":"Reference fragments","quality":"Equal","values":[146,146,146,146,146,146],"isMain":false},{"metricName":"Reference GC (%)","quality":"Equal","values":["51.82","51.82","51.82","51.82","51.82","51.82"],"isMain":false}]]],"subreferences":[],"subreports":[],"minContig":500},{"date":"10 February 2020, Monday, 18:01:12","assembliesNames":["megahit-103-df","megahit-113-df","megahit-113-ml","megahit-113-ms","megahit-129-df","metaSPAdes"],"referenceName":"GCF\_000159215.1\_ASM15921v1","order":[0,1,2,3,4,5],"report":[["Genome statistics",[{"metricName":"Genome fraction (%)","quality":"More is better","values":["90.545","92.023","87.594","87.510","93.159","87.088"],"isMain":true},{"metricName":"Duplication ratio","quality":"Less is better","values":["1.053","1.036","1.094","1.100","1.043","1.058"],"isMain":true},{"metricName":"Largest alignment","quality":"More is better","values":[8797,11637,5143,5465,9556,6006],"isMain":true},{"metricName":"Total aligned length","quality":"More is better","values":[1706062,1705503,1709478,1714538,1739279,1642065],"isMain":true},{"metricName":"NG50","quality":"More is better","values":[1553,2259,1135,1130,2400,1291],"isMain":false},{"metricName":"NG75","quality":"More is better","values":[967,1201,757,743,1297,795],"isMain":false},{"metricName":"NA50","quality":"More is better","values":[1610,2316,1159,1153,2400,1387],"isMain":false},{"metricName":"NA75","quality":"More is better","values":[1026,1301,774,764,1310,874],"isMain":false},{"metricName":"NGA50","quality":"More is better","values":[1551,2230,1123,1109,2346,1271],"isMain":true},{"metricName":"NGA75","quality":"More is better","values":[953,1182,741,731,1262,780],"isMain":false},{"metricName":"LG50","quality":"Less is better","values":[359,255,523,524,240,439],"isMain":false},{"metricName":"LG75","quality":"Less is better","values":[726,528,1016,1019,496,884],"isMain":false},{"metricName":"LA50","quality":"Less is better","values":[334,238,495,500,237,390],"isMain":false},{"metricName":"LA75","quality":"Less is better","values":[669,482,956,967,483,768],"isMain":false},{"metricName":"LGA50","quality":"Less is better","values":[360,256,527,528,245,443],"isMain":true},{"metricName":"LGA75","quality":"Less is better","values":[731,533,1028,1032,507,897],"isMain":false}]],["Reads mapping",[]],["Misassemblies",[{"metricName":"# misassemblies","quality":"Less is better","values":[3,5,5,4,6,4],"isMain":true},{"metricName":" # relocations","quality":"Less is better","values":[0,0,0,0,0,1],"isMain":false},{"metricName":" # translocations","quality":"Less is better","values":[3,5,4,4,6,3],"isMain":false},{"metricName":" # inversions","quality":"Less is better","values":[0,0,1,0,0,0],"isMain":false},{"metricName":" # interspecies translocations","quality":"Less is better","values":[18,13,10,19,10,11],"isMain":false},{"metricName":"# misassembled contigs","quality":"Less is better","values":[3,5,5,4,6,4],"isMain":false},{"metricName":"Misassembled contigs length","quality":"Less is better","values":[2737,6254,6001,3016,18102,6502],"isMain":true},{"metricName":"# possibly misassembled contigs","quality":"Less is better","values":[0,0,0,0,0,0],"isMain":false},{"metricName":" # possible misassemblies","quality":"Less is better","values":[0,0,0,0,0,0],"isMain":false},{"metricName":"# local misassemblies","quality":"Less is better","values":[2,1,6,4,3,1],"isMain":false},{"metricName":"# scaffold gap ext. mis.","quality":"Less is better","values":[0,0,0,0,0,0],"isMain":false},{"metricName":"# scaffold gap loc. mis.","quality":"Less is better","values":[0,0,0,0,0,0],"isMain":false},{"metricName":"# unaligned mis. contigs","quality":"Less is better","values":[0,0,1,0,0,0],"isMain":false}]],["Unaligned",[{"metricName":"# fully unaligned contigs","quality":"Less is better","values":[0,0,0,0,0,0],"isMain":false},{"metricName":"Fully unaligned length","quality":"Less is better","values":[0,0,0,0,0,0],"isMain":false},{"metricName":"# partially unaligned contigs","quality":"Less is better","values":[1,1,4,3,3,0],"isMain":false},{"metricName":"Partially unaligned length","quality":"Less is better","values":[828,537,2368,2540,8921,0],"isMain":false}]],["Mismatches",[{"metricName":"# mismatches","quality":"Less is better","values":[12543,10834,24604,25819,11401,13049],"isMain":false},{"metricName":"# indels","quality":"Less is better","values":[18,13,38,40,24,26],"isMain":false},{"metricName":"Indels length","quality":"Less is better","values":[82,13,123,120,99,85],"isMain":false},{"metricName":"# mismatches per 100 kbp","quality":"Less is better","values":["770.91","655.18","1563.14","1641.89","681.06","833.84"],"isMain":true},{"metricName":"# indels per 100 kbp","quality":"Less is better","values":["1.11","0.79","2.41","2.54","1.43","1.66"],"isMain":true},{"metricName":" # indels (<= 5 bp)","quality":"Less is better","values":[16,13,36,38,21,24],"isMain":false},{"metricName":" # indels (> 5 bp)","quality":"Less is better","values":[2,0,2,2,3,2],"isMain":false},{"metricName":"# N's","quality":"Less is better","values":[0,0,0,0,0,0],"isMain":false},{"metricName":"# N's per 100 kbp","quality":"Less is better","values":["0.00","0.00","0.00","0.00","0.00","0.00"],"isMain":true}]],["Statistics without reference",[{"metricName":"# contigs","quality":"Equal","values":[1237,979,1619,1644,970,1364],"isMain":true},{"metricName":"# contigs (>= 1000 bp)","quality":"Equal","values":[696,629,642,642,637,659],"isMain":false},{"metricName":"# contigs (>= 5000 bp)","quality":"Equal","values":[17,31,2,2,38,2],"isMain":false},{"metricName":"# contigs (>= 10000 bp)","quality":"Equal","values":[0,1,0,0,0,0],"isMain":false},{"metricName":"# contigs (>= 25000 bp)","quality":"Equal","values":[0,0,0,0,0,0],"isMain":false},{"metricName":"# contigs (>= 50000 bp)","quality":"Equal","values":[0,0,0,0,0,0],"isMain":false},{"metricName":"Largest contig","quality":"More is better","values":[8801,11637,5143,5474,9558,6007],"isMain":true},{"metricName":"Total length","quality":"More is better","values":[1714634,1713715,1723758,1732607,1755584,1655740],"isMain":true},{"metricName":"Total length (>= 1000 bp)","quality":"More is better","values":[1319190,1458712,1025453,1024262,1508928,1148526],"isMain":true},{"metricName":"Total length (>= 5000 bp)","quality":"More is better","values":[99734,191836,10144,10729,238925,11658],"isMain":false},{"metricName":"Total length (>= 10000 bp)","quality":"More is better","values":[0,11637,0,0,0,0],"isMain":true},{"metricName":"Total length (>= 25000 bp)","quality":"More is better","values":[0,0,0,0,0,0],"isMain":false},{"metricName":"Total length (>= 50000 bp)","quality":"More is better","values":[0,0,0,0,0,0],"isMain":true},{"metricName":"N50","quality":"More is better","values":[1624,2338,1178,1173,2430,1403],"isMain":false},{"metricName":"N75","quality":"More is better","values":[1038,1335,790,776,1363,897],"isMain":false},{"metricName":"L50","quality":"Less is better","values":[333,237,491,496,231,386],"isMain":false},{"metricName":"L75","quality":"Less is better","values":[664,479,944,956,472,758],"isMain":false},{"metricName":"GC (%)","quality":"Equal","values":["53.00","53.01","52.98","52.93","52.97","53.07"],"isMain":false}]],["K-mer-based statistics",[]],["Predicted genes",[]],["Similarity statistics",[]],["Reference statistics",[{"metricName":"Reference length","quality":"Equal","values":[1796948,1796948,1796948,1796948,1796948,1796948],"isMain":false},{"metricName":"Reference fragments","quality":"Equal","values":[146,146,146,146,146,146],"isMain":false},{"metricName":"Reference GC (%)","quality":"Equal","values":["52.78","52.78","52.78","52.78","52.78","52.78"],"isMain":false}]]],"subreferences":[],"subreports":[],"minContig":500},{"date":"10 February 2020, Monday, 18:00:15","assembliesNames":["megahit-103-df","megahit-113-df","megahit-113-ml","megahit-113-ms","megahit-129-df","metaSPAdes"],"referenceName":"GCF\_000159615.1\_ASM15961v1","order":[0,1,2,3,4,5],"report":[["Genome statistics",[{"metricName":"Genome fraction (%)","quality":"More is better","values":["7.276","8.698","9.561","10.718","9.624","4.467"],"isMain":true},{"metricName":"Duplication ratio","quality":"Less is better","values":["1.041","1.034","1.027","1.043","1.035","1.056"],"isMain":true},{"metricName":"Largest alignment","quality":"More is better","values":[1661,2361,1758,1485,1938,1534],"isMain":true},{"metricName":"Total aligned length","quality":"More is better","values":[149494,178553,195462,221273,197357,90948],"isMain":true},{"metricName":"NA50","quality":"More is better","values":[647,677,627,641,717,642],"isMain":false},{"metricName":"NA75","quality":"More is better","values":[544,545,542,552,561,537],"isMain":false},{"metricName":"NGA50","quality":"More is better","values":[null,null,null,null,null,null],"isMain":true},{"metricName":"LA50","quality":"Less is better","values":[96,100,124,141,103,59],"isMain":false},{"metricName":"LA75","quality":"Less is better","values":[162,180,210,239,186,101],"isMain":false}]],["Reads mapping",[]],["Misassemblies",[{"metricName":"# misassemblies","quality":"Less is better","values":[2,4,5,1,4,0],"isMain":true},{"metricName":" # relocations","quality":"Less is better","values":[0,0,0,0,0,0],"isMain":false},{"metricName":" # translocations","quality":"Less is better","values":[2,4,5,1,4,0],"isMain":false},{"metricName":" # inversions","quality":"Less is better","values":[0,0,0,0,0,0],"isMain":false},{"metricName":" # interspecies translocations","quality":"Less is better","values":[19,20,12,14,20,13],"isMain":false},{"metricName":"# misassembled contigs","quality":"Less is better","values":[2,3,4,1,4,0],"isMain":false},{"metricName":"Misassembled contigs length","quality":"Less is better","values":[2329,3765,3010,645,3068,0],"isMain":true},{"metricName":"# possibly misassembled contigs","quality":"Less is better","values":[0,0,0,0,1,0],"isMain":false},{"metricName":" # possible misassemblies","quality":"Less is better","values":[0,0,0,0,1,0],"isMain":false},{"metricName":"# local misassemblies","quality":"Less is better","values":[0,1,0,1,1,0],"isMain":false},{"metricName":"# scaffold gap ext. mis.","quality":"Less is better","values":[0,0,0,0,0,0],"isMain":false},{"metricName":"# scaffold gap loc. mis.","quality":"Less is better","values":[0,0,0,0,0,0],"isMain":false},{"metricName":"# unaligned mis. contigs","quality":"Less is better","values":[0,2,0,0,0,1],"isMain":false}]],["Unaligned",[{"metricName":"# fully unaligned contigs","quality":"Less is better","values":[1,1,0,1,0,1],"isMain":false},{"metricName":"Fully unaligned length","quality":"Less is better","values":[618,861,0,570,0,631],"isMain":false},{"metricName":"# partially unaligned contigs","quality":"Less is better","values":[5,10,3,5,7,1],"isMain":false},{"metricName":"Partially unaligned length","quality":"Less is better","values":[3466,8002,2044,3570,4555,1302],"isMain":false}]],["Mismatches",[{"metricName":"# mismatches","quality":"Less is better","values":[3696,4558,5225,6002,5002,2290],"isMain":false},{"metricName":"# indels","quality":"Less is better","values":[38,28,26,34,27,16],"isMain":false},{"metricName":"Indels length","quality":"Less is better","values":[84,40,55,53,46,35],"isMain":false},{"metricName":"# mismatches per 100 kbp","quality":"Less is better","values":["2499.22","2577.98","2688.55","2755.18","2557.10","2522.03"],"isMain":true},{"metricName":"# indels per 100 kbp","quality":"Less is better","values":["25.70","15.84","13.38","15.61","13.80","17.62"],"isMain":true},{"metricName":" # indels (<= 5 bp)","quality":"Less is better","values":[36,28,24,33,25,15],"isMain":false},{"metricName":" # indels (> 5 bp)","quality":"Less is better","values":[2,0,2,1,2,1],"isMain":false},{"metricName":"# N's","quality":"Less is better","values":[0,0,0,0,0,0],"isMain":false},{"metricName":"# N's per 100 kbp","quality":"Less is better","values":["0.00","0.00","0.00","0.00","0.00","0.00"],"isMain":true}]],["Statistics without reference",[{"metricName":"# contigs","quality":"Equal","values":[227,251,297,339,270,140],"isMain":true},{"metricName":"# contigs (>= 1000 bp)","quality":"Equal","values":[20,39,19,22,44,11],"isMain":false},{"metricName":"# contigs (>= 5000 bp)","quality":"Equal","values":[0,0,0,0,0,0],"isMain":false},{"metricName":"# contigs (>= 10000 bp)","quality":"Equal","values":[0,0,0,0,0,0],"isMain":false},{"metricName":"# contigs (>= 25000 bp)","quality":"Equal","values":[0,0,0,0,0,0],"isMain":false},{"metricName":"# contigs (>= 50000 bp)","quality":"Equal","values":[0,0,0,0,0,0],"isMain":false},{"metricName":"Largest contig","quality":"More is better","values":[1702,2407,1769,1506,2014,1646],"isMain":true},{"metricName":"Total length","quality":"More is better","values":[158062,191762,201553,231268,207080,97839],"isMain":true},{"metricName":"Total length (>= 1000 bp)","quality":"More is better","values":[23483,51807,24147,25343,58604,13639],"isMain":true},{"metricName":"Total length (>= 5000 bp)","quality":"More is better","values":[0,0,0,0,0,0],"isMain":false},{"metricName":"Total length (>= 10000 bp)","quality":"More is better","values":[0,0,0,0,0,0],"isMain":true},{"metricName":"Total length (>= 25000 bp)","quality":"More is better","values":[0,0,0,0,0,0],"isMain":false},{"metricName":"Total length (>= 50000 bp)","quality":"More is better","values":[0,0,0,0,0,0],"isMain":true},{"metricName":"N50","quality":"More is better","values":[681,756,655,665,762,680],"isMain":false},{"metricName":"N75","quality":"More is better","values":[589,587,562,570,590,580],"isMain":false},{"metricName":"L50","quality":"Less is better","values":[91,90,119,137,97,56],"isMain":false},{"metricName":"L75","quality":"Less is better","values":[154,163,202,231,175,95],"isMain":false},{"metricName":"GC (%)","quality":"Equal","values":["38.97","38.97","39.05","39.14","38.80","38.94"],"isMain":false}]],["K-mer-based statistics",[]],["Predicted genes",[]],["Similarity statistics",[]],["Reference statistics",[{"metricName":"Reference length","quality":"Equal","values":[2032595,2032595,2032595,2032595,2032595,2032595],"isMain":false},{"metricName":"Reference fragments","quality":"Equal","values":[244,244,244,244,244,244],"isMain":false},{"metricName":"Reference GC (%)","quality":"Equal","values":["38.72","38.72","38.72","38.72","38.72","38.72"],"isMain":false}]]],"subreferences":[],"subreports":[],"minContig":500},{"date":"10 February 2020, Monday, 18:01:13","assembliesNames":["megahit-103-df","megahit-113-df","megahit-113-ml","megahit-113-ms","megahit-129-df","metaSPAdes"],"referenceName":"GCF\_000159875.2\_Bact\_sp\_1\_1\_6\_V2","order":[0,1,2,3,4,5],"report":[["Genome statistics",[{"metricName":"Genome fraction (%)","quality":"More is better","values":["8.435","9.246","10.760","11.121","9.112","4.353"],"isMain":true},{"metricName":"Duplication ratio","quality":"Less is better","values":["1.021","1.025","1.042","1.039","1.033","1.023"],"isMain":true},{"metricName":"Largest alignment","quality":"More is better","values":[14106,24853,8462,20105,20795,12653],"isMain":true},{"metricName":"Total aligned length","quality":"More is better","values":[576478,629815,746053,769013,624159,297504],"isMain":true},{"metricName":"NA50","quality":"More is better","values":[1626,2315,1268,1275,1735,989],"isMain":false},{"metricName":"NA75","quality":"More is better","values":[825,886,716,717,689,null],"isMain":false},{"metricName":"NGA50","quality":"More is better","values":[null,null,null,null,null,null],"isMain":true},{"metricName":"LA50","quality":"Less is better","values":[99,71,172,165,94,110],"isMain":false},{"metricName":"LA75","quality":"Less is better","values":[228,186,377,376,262,null],"isMain":false}]],["Reads mapping",[]],["Misassemblies",[{"metricName":"# misassemblies","quality":"Less is better","values":[1,7,8,6,6,0],"isMain":true},{"metricName":" # relocations","quality":"Less is better","values":[0,2,0,1,1,0],"isMain":false},{"metricName":" # translocations","quality":"Less is better","values":[1,5,7,5,4,0],"isMain":false},{"metricName":" # inversions","quality":"Less is better","values":[0,0,1,0,1,0],"isMain":false},{"metricName":" # interspecies translocations","quality":"Less is better","values":[15,36,30,35,34,6],"isMain":false},{"metricName":"# misassembled contigs","quality":"Less is better","values":[1,7,8,6,6,0],"isMain":false},{"metricName":"Misassembled contigs length","quality":"Less is better","values":[638,17075,27553,13098,55133,0],"isMain":true},{"metricName":"# possibly misassembled contigs","quality":"Less is better","values":[0,0,0,0,0,0],"isMain":false},{"metricName":" # possible misassemblies","quality":"Less is better","values":[0,0,0,0,0,0],"isMain":false},{"metricName":"# local misassemblies","quality":"Less is better","values":[0,6,1,1,4,0],"isMain":false},{"metricName":"# scaffold gap ext. mis.","quality":"Less is better","values":[0,0,0,0,0,0],"isMain":false},{"metricName":"# scaffold gap loc. mis.","quality":"Less is better","values":[0,0,0,0,0,0],"isMain":false},{"metricName":"# unaligned mis. contigs","quality":"Less is better","values":[0,4,2,3,1,0],"isMain":false}]],["Unaligned",[{"metricName":"# fully unaligned contigs","quality":"Less is better","values":[2,1,2,0,0,0],"isMain":false},{"metricName":"Fully unaligned length","quality":"Less is better","values":[1823,573,1599,0,0,0],"isMain":false},{"metricName":"# partially unaligned contigs","quality":"Less is better","values":[5,12,9,13,10,2],"isMain":false},{"metricName":"Partially unaligned length","quality":"Less is better","values":[3571,32199,14588,22978,57431,127995],"isMain":false}]],["Mismatches",[{"metricName":"# mismatches","quality":"Less is better","values":[5586,7073,10942,11733,7353,2823],"isMain":false},{"metricName":"# indels","quality":"Less is better","values":[40,68,65,55,57,21],"isMain":false},{"metricName":"Indels length","quality":"Less is better","values":[122,107,131,165,173,40],"isMain":false},{"metricName":"# mismatches per 100 kbp","quality":"Less is better","values":["980.68","1132.93","1505.96","1562.40","1195.08","960.42"],"isMain":true},{"metricName":"# indels per 100 kbp","quality":"Less is better","values":["7.02","10.89","8.95","7.32","9.26","7.14"],"isMain":true},{"metricName":" # indels (<= 5 bp)","quality":"Less is better","values":[37,66,62,49,52,19],"isMain":false},{"metricName":" # indels (> 5 bp)","quality":"Less is better","values":[3,2,3,6,5,2],"isMain":false},{"metricName":"# N's","quality":"Less is better","values":[0,0,0,0,0,0],"isMain":false},{"metricName":"# N's per 100 kbp","quality":"Less is better","values":["0.00","0.00","0.00","0.00","0.00","0.00"],"isMain":true}]],["Statistics without reference",[{"metricName":"# contigs","quality":"Equal","values":[455,386,668,665,448,232],"isMain":true},{"metricName":"# contigs (>= 1000 bp)","quality":"Equal","values":[187,174,256,252,171,108],"isMain":false},{"metricName":"# contigs (>= 5000 bp)","quality":"Equal","values":[8,21,9,10,13,4],"isMain":false},{"metricName":"# contigs (>= 10000 bp)","quality":"Equal","values":[2,6,1,2,5,2],"isMain":false},{"metricName":"# contigs (>= 25000 bp)","quality":"Equal","values":[0,0,0,0,1,1],"isMain":false},{"metricName":"# contigs (>= 50000 bp)","quality":"Equal","values":[0,0,0,0,0,1],"isMain":false},{"metricName":"Largest contig","quality":"More is better","values":[14106,24853,17220,20105,46298,130043],"isMain":true},{"metricName":"Total length","quality":"More is better","values":[587020,672615,773092,803342,692782,428557],"isMain":true},{"metricName":"Total length (>= 1000 bp)","quality":"More is better","values":[406358,531082,496430,524772,507094,341676],"isMain":true},{"metricName":"Total length (>= 5000 bp)","quality":"More is better","values":[61112,204378,69191,86122,172283,157943],"isMain":false},{"metricName":"Total length (>= 10000 bp)","quality":"More is better","values":[26327,97638,17220,31396,120571,142714],"isMain":true},{"metricName":"Total length (>= 25000 bp)","quality":"More is better","values":[0,0,0,0,46298,130043],"isMain":false},{"metricName":"Total length (>= 50000 bp)","quality":"More is better","values":[0,0,0,0,0,130043],"isMain":true},{"metricName":"N50","quality":"More is better","values":[1630,2640,1367,1423,2260,2606],"isMain":false},{"metricName":"N75","quality":"More is better","values":[850,1215,780,812,945,1160],"isMain":false},{"metricName":"L50","quality":"Less is better","values":[99,58,160,149,65,20],"isMain":false},{"metricName":"L75","quality":"Less is better","values":[224,150,350,339,184,89],"isMain":false},{"metricName":"GC (%)","quality":"Equal","values":["41.85","41.78","42.34","42.28","41.36","41.15"],"isMain":false}]],["K-mer-based statistics",[]],["Predicted genes",[]],["Similarity statistics",[]],["Reference statistics",[{"metricName":"Reference length","quality":"Equal","values":[6752535,6752535,6752535,6752535,6752535,6752535],"isMain":false},{"metricName":"Reference fragments","quality":"Equal","values":[97,97,97,97,97,97],"isMain":false},{"metricName":"Reference GC (%)","quality":"Equal","values":["43.02","43.02","43.02","43.02","43.02","43.02"],"isMain":false}]]],"subreferences":[],"subreports":[],"minContig":500},{"date":"10 February 2020, Monday, 18:00:13","assembliesNames":["megahit-103-df","megahit-113-df","megahit-113-ml","megahit-113-ms","megahit-129-df","metaSPAdes"],"referenceName":"GCF\_000159995.1\_ASM15999v1","order":[0,1,2,3,4,5],"report":[["Genome statistics",[{"metricName":"Genome fraction (%)","quality":"More is better","values":["0.176","0.444","0.515","0.652","0.453","0.252"],"isMain":true},{"metricName":"Duplication ratio","quality":"Less is better","values":["1.054","1.081","1.077","1.138","1.015","1.204"],"isMain":true},{"metricName":"Largest alignment","quality":"More is better","values":[812,3827,3640,819,4181,683],"isMain":true},{"metricName":"Total aligned length","quality":"More is better","values":[4556,11473,13309,17289,11686,6695],"isMain":true},{"metricName":"NA50","quality":"More is better","values":[526,536,524,518,546,null],"isMain":false},{"metricName":"NA75","quality":"More is better","values":[null,null,null,274,207,null],"isMain":false},{"metricName":"NGA50","quality":"More is better","values":[null,null,null,null,null,null],"isMain":true},{"metricName":"LA50","quality":"Less is better","values":[5,9,12,18,7,null],"isMain":false},{"metricName":"LA75","quality":"Less is better","values":[null,null,null,29,15,null],"isMain":false}]],["Reads mapping",[]],["Misassemblies",[{"metricName":"# misassemblies","quality":"Less is better","values":[0,2,1,1,0,1],"isMain":true},{"metricName":" # relocations","quality":"Less is better","values":[0,0,1,0,0,1],"isMain":false},{"metricName":" # translocations","quality":"Less is better","values":[0,2,0,1,0,0],"isMain":false},{"metricName":" # inversions","quality":"Less is better","values":[0,0,0,0,0,0],"isMain":false},{"metricName":" # interspecies translocations","quality":"Less is better","values":[2,8,8,9,3,7],"isMain":false},{"metricName":"# misassembled contigs","quality":"Less is better","values":[0,1,1,1,0,1],"isMain":false},{"metricName":"Misassembled contigs length","quality":"Less is better","values":[0,654,943,557,0,502],"isMain":true},{"metricName":"# possibly misassembled contigs","quality":"Less is better","values":[0,0,0,0,0,0],"isMain":false},{"metricName":" # possible misassemblies","quality":"Less is better","values":[0,0,0,0,0,0],"isMain":false},{"metricName":"# local misassemblies","quality":"Less is better","values":[0,0,0,0,0,0],"isMain":false},{"metricName":"# scaffold gap ext. mis.","quality":"Less is better","values":[0,0,0,0,0,0],"isMain":false},{"metricName":"# scaffold gap loc. mis.","quality":"Less is better","values":[0,0,0,0,0,0],"isMain":false},{"metricName":"# unaligned mis. contigs","quality":"Less is better","values":[0,0,0,0,1,2],"isMain":false}]],["Unaligned",[{"metricName":"# fully unaligned contigs","quality":"Less is better","values":[0,0,1,1,0,0],"isMain":false},{"metricName":"Fully unaligned length","quality":"Less is better","values":[0,0,1182,1344,0,0],"isMain":false},{"metricName":"# partially unaligned contigs","quality":"Less is better","values":[1,5,4,2,3,4],"isMain":false},{"metricName":"Partially unaligned length","quality":"Less is better","values":[1289,3491,5124,1220,3071,114808],"isMain":false}]],["Mismatches",[{"metricName":"# mismatches","quality":"Less is better","values":[114,337,364,475,361,172],"isMain":false},{"metricName":"# indels","quality":"Less is better","values":[0,3,4,3,3,0],"isMain":false},{"metricName":"Indels length","quality":"Less is better","values":[0,17,4,4,3,0],"isMain":false},{"metricName":"# mismatches per 100 kbp","quality":"Less is better","values":["2502.19","2937.33","2734.58","2819.33","3088.37","2642.09"],"isMain":true},{"metricName":"# indels per 100 kbp","quality":"Less is better","values":["0.00","26.15","30.05","17.81","25.67","0.00"],"isMain":true},{"metricName":" # indels (<= 5 bp)","quality":"Less is better","values":[0,1,4,3,3,0],"isMain":false},{"metricName":" # indels (> 5 bp)","quality":"Less is better","values":[0,2,0,0,0,0],"isMain":false},{"metricName":"# N's","quality":"Less is better","values":[0,0,0,0,0,0],"isMain":false},{"metricName":"# N's per 100 kbp","quality":"Less is better","values":["0.00","0.00","0.00","0.00","0.00","0.00"],"isMain":true}]],["Statistics without reference",[{"metricName":"# contigs","quality":"Equal","values":[9,19,21,34,16,14],"isMain":true},{"metricName":"# contigs (>= 1000 bp)","quality":"Equal","values":[1,2,5,2,3,3],"isMain":false},{"metricName":"# contigs (>= 5000 bp)","quality":"Equal","values":[0,0,0,0,0,2],"isMain":false},{"metricName":"# contigs (>= 10000 bp)","quality":"Equal","values":[0,0,0,0,0,2],"isMain":false},{"metricName":"# contigs (>= 25000 bp)","quality":"Equal","values":[0,0,0,0,0,2],"isMain":false},{"metricName":"# contigs (>= 50000 bp)","quality":"Equal","values":[0,0,0,0,0,1],"isMain":false},{"metricName":"Largest contig","quality":"More is better","values":[1407,3827,3640,1344,4345,88424],"isMain":true},{"metricName":"Total length","quality":"More is better","values":[6090,15893,20636,21744,14930,122649],"isMain":true},{"metricName":"Total length (>= 1000 bp)","quality":"More is better","values":[1407,5065,10401,2385,7533,116062],"isMain":true},{"metricName":"Total length (>= 5000 bp)","quality":"More is better","values":[0,0,0,0,0,114655],"isMain":false},{"metricName":"Total length (>= 10000 bp)","quality":"More is better","values":[0,0,0,0,0,114655],"isMain":true},{"metricName":"Total length (>= 25000 bp)","quality":"More is better","values":[0,0,0,0,0,114655],"isMain":false},{"metricName":"Total length (>= 50000 bp)","quality":"More is better","values":[0,0,0,0,0,88424],"isMain":true},{"metricName":"N50","quality":"More is better","values":[549,717,1182,606,1108,88424],"isMain":false},{"metricName":"N75","quality":"More is better","values":[526,601,620,545,566,26231],"isMain":false},{"metricName":"L50","quality":"Less is better","values":[4,6,5,14,3,1],"isMain":false},{"metricName":"L75","quality":"Less is better","values":[7,12,12,24,9,2],"isMain":false},{"metricName":"GC (%)","quality":"Equal","values":["45.17","44.62","43.41","46.16","43.64","46.32"],"isMain":false}]],["K-mer-based statistics",[]],["Predicted genes",[]],["Similarity statistics",[]],["Reference statistics",[{"metricName":"Reference length","quality":"Equal","values":[2582919,2582919,2582919,2582919,2582919,2582919],"isMain":false},{"metricName":"Reference fragments","quality":"Equal","values":[34,34,34,34,34,34],"isMain":false},{"metricName":"Reference GC (%)","quality":"Equal","values":["46.56","46.56","46.56","46.56","46.56","46.56"],"isMain":false}]]],"subreferences":[],"subreports":[],"minContig":500},{"date":"10 February 2020, Monday, 18:26:56","assembliesNames":["megahit-103-df","megahit-113-df","megahit-113-ml","megahit-113-ms","megahit-129-df","metaSPAdes"],"referenceName":"GCF\_000160035.2\_ASM16003v2","order":[0,1,2,3,4,5],"report":[["Genome statistics",[{"metricName":"Genome fraction (%)","quality":"More is better","values":["98.681","98.865","98.431","98.403","98.806","97.617"],"isMain":true},{"metricName":"Duplication ratio","quality":"Less is better","values":["1.020","1.014","1.073","1.070","1.019","1.019"],"isMain":true},{"metricName":"Largest alignment","quality":"More is better","values":[39441,93222,17609,20592,58804,40312],"isMain":true},{"metricName":"Total aligned length","quality":"More is better","values":[3497902,3482082,3662902,3642277,3493645,3453532],"isMain":true},{"metricName":"NG50","quality":"More is better","values":[11261,19215,3859,4346,18966,9494],"isMain":false},{"metricName":"NG75","quality":"More is better","values":[6563,11194,2428,2706,11246,4998],"isMain":false},{"metricName":"NA50","quality":"More is better","values":[11054,17788,3582,3885,18526,9495],"isMain":false},{"metricName":"NA75","quality":"More is better","values":[6377,9935,2135,2201,10489,5062],"isMain":false},{"metricName":"NGA50","quality":"More is better","values":[11112,17853,3779,4216,18641,9491],"isMain":true},{"metricName":"NGA75","quality":"More is better","values":[6447,10529,2358,2565,10711,4994],"isMain":false},{"metricName":"LG50","quality":"Less is better","values":[96,59,281,258,59,109],"isMain":false},{"metricName":"LG75","quality":"Less is better","values":[198,120,562,516,120,230],"isMain":false},{"metricName":"LA50","quality":"Less is better","values":[98,63,321,299,61,108],"isMain":false},{"metricName":"LA75","quality":"Less is better","values":[203,130,652,615,127,228],"isMain":false},{"metricName":"LGA50","quality":"Less is better","values":[97,62,290,269,60,109],"isMain":true},{"metricName":"LGA75","quality":"Less is better","values":[200,126,578,538,124,230],"isMain":false}]],["Reads mapping",[]],["Misassemblies",[{"metricName":"# misassemblies","quality":"Less is better","values":[6,10,10,13,10,1],"isMain":true},{"metricName":" # relocations","quality":"Less is better","values":[2,4,3,2,3,1],"isMain":false},{"metricName":" # translocations","quality":"Less is better","values":[3,5,7,6,6,0],"isMain":false},{"metricName":" # inversions","quality":"Less is better","values":[1,1,0,5,1,0],"isMain":false},{"metricName":" # interspecies translocations","quality":"Less is better","values":[6,10,7,42,11,0],"isMain":false},{"metricName":"# misassembled contigs","quality":"Less is better","values":[6,10,10,13,10,1],"isMain":false},{"metricName":"Misassembled contigs length","quality":"Less is better","values":[81457,166067,48978,41539,176788,1218],"isMain":true},{"metricName":"# possibly misassembled contigs","quality":"Less is better","values":[0,0,0,0,0,0],"isMain":false},{"metricName":" # possible misassemblies","quality":"Less is better","values":[0,0,0,0,0,0],"isMain":false},{"metricName":"# local misassemblies","quality":"Less is better","values":[0,4,0,1,2,0],"isMain":false},{"metricName":"# scaffold gap ext. mis.","quality":"Less is better","values":[0,0,0,0,0,0],"isMain":false},{"metricName":"# scaffold gap loc. mis.","quality":"Less is better","values":[0,0,0,0,0,0],"isMain":false},{"metricName":"# unaligned mis. contigs","quality":"Less is better","values":[0,0,0,0,1,0],"isMain":false}]],["Unaligned",[{"metricName":"# fully unaligned contigs","quality":"Less is better","values":[0,2,0,0,0,0],"isMain":false},{"metricName":"Fully unaligned length","quality":"Less is better","values":[0,2647,0,0,0,0],"isMain":false},{"metricName":"# partially unaligned contigs","quality":"Less is better","values":[2,5,2,14,6,0],"isMain":false},{"metricName":"Partially unaligned length","quality":"Less is better","values":[1430,42436,27282,60546,17641,0],"isMain":false}]],["Mismatches",[{"metricName":"# mismatches","quality":"Less is better","values":[6125,5000,26109,26064,5197,5277],"isMain":false},{"metricName":"# indels","quality":"Less is better","values":[12,29,62,50,27,18],"isMain":false},{"metricName":"Indels length","quality":"Less is better","values":[12,137,240,90,302,126],"isMain":false},{"metricName":"# mismatches per 100 kbp","quality":"Less is better","values":["178.49","145.44","762.79","761.69","151.26","155.46"],"isMain":true},{"metricName":"# indels per 100 kbp","quality":"Less is better","values":["0.35","0.84","1.81","1.46","0.79","0.53"],"isMain":true},{"metricName":" # indels (<= 5 bp)","quality":"Less is better","values":[12,25,57,48,19,14],"isMain":false},{"metricName":" # indels (> 5 bp)","quality":"Less is better","values":[0,4,5,2,8,4],"isMain":false},{"metricName":"# N's","quality":"Less is better","values":[0,0,0,0,0,0],"isMain":false},{"metricName":"# N's per 100 kbp","quality":"Less is better","values":["0.00","0.00","0.00","0.00","0.00","0.00"],"isMain":true}]],["Statistics without reference",[{"metricName":"# contigs","quality":"Equal","values":[526,348,1413,1323,376,585],"isMain":true},{"metricName":"# contigs (>= 1000 bp)","quality":"Equal","values":[459,305,1074,1035,317,520],"isMain":false},{"metricName":"# contigs (>= 5000 bp)","quality":"Equal","values":[244,194,176,196,191,229],"isMain":false},{"metricName":"# contigs (>= 10000 bp)","quality":"Equal","values":[114,134,18,21,131,98],"isMain":false},{"metricName":"# contigs (>= 25000 bp)","quality":"Equal","values":[14,29,1,1,34,14],"isMain":false},{"metricName":"# contigs (>= 50000 bp)","quality":"Equal","values":[0,4,0,0,4,0],"isMain":false},{"metricName":"Largest contig","quality":"More is better","values":[39445,93222,34007,47906,78856,40323],"isMain":true},{"metricName":"Total length","quality":"More is better","values":[3503275,3531881,3700480,3720763,3517589,3459056],"isMain":true},{"metricName":"Total length (>= 1000 bp)","quality":"More is better","values":[3456307,3501903,3453466,3512294,3475756,3410938],"isMain":true},{"metricName":"Total length (>= 5000 bp)","quality":"More is better","values":[2871914,3212342,1282731,1450290,3161138,2606741],"isMain":false},{"metricName":"Total length (>= 10000 bp)","quality":"More is better","values":[1934335,2766324,249529,296075,2729140,1632695],"isMain":true},{"metricName":"Total length (>= 25000 bp)","quality":"More is better","values":[450339,1092881,34007,47906,1223828,422786],"isMain":false},{"metricName":"Total length (>= 50000 bp)","quality":"More is better","values":[0,289935,0,0,245815,0],"isMain":true},{"metricName":"N50","quality":"More is better","values":[11150,18679,3673,4044,18921,9500],"isMain":false},{"metricName":"N75","quality":"More is better","values":[6441,11001,2191,2352,10769,5164],"isMain":false},{"metricName":"L50","quality":"Less is better","values":[97,61,310,287,60,108],"isMain":false},{"metricName":"L75","quality":"Less is better","values":[201,123,634,588,123,227],"isMain":false},{"metricName":"GC (%)","quality":"Equal","values":["37.11","37.23","37.21","37.28","37.14","37.07"],"isMain":false}]],["K-mer-based statistics",[]],["Predicted genes",[]],["Similarity statistics",[]],["Reference statistics",[{"metricName":"Reference length","quality":"Equal","values":[3477404,3477404,3477404,3477404,3477404,3477404],"isMain":false},{"metricName":"Reference fragments","quality":"Equal","values":[26,26,26,26,26,26],"isMain":false},{"metricName":"Reference GC (%)","quality":"Equal","values":["37.14","37.14","37.14","37.14","37.14","37.14"],"isMain":false}]]],"subreferences":[],"subreports":[],"minContig":500},{"date":"10 February 2020, Monday, 18:43:47","assembliesNames":["megahit-103-df","megahit-113-df","megahit-113-ml","megahit-113-ms","megahit-129-df","metaSPAdes"],"referenceName":"GCF\_000160095.1\_ASM16009v1","order":[0,1,2,3,4,5],"report":[["Genome statistics",[{"metricName":"Genome fraction (%)","quality":"More is better","values":["4.628","97.058","96.984","97.257","97.450","95.833"],"isMain":true},{"metricName":"Duplication ratio","quality":"Less is better","values":["1.130","1.431","1.040","1.057","1.084","1.005"],"isMain":true},{"metricName":"Largest alignment","quality":"More is better","values":[747,49472,55241,55241,54267,55241],"isMain":true},{"metricName":"Total aligned length","quality":"More is better","values":[318373,9123815,6650738,6771288,6953112,6366597],"isMain":true},{"metricName":"NG50","quality":"More is better","values":[null,12038,19486,23861,20370,13189],"isMain":false},{"metricName":"NG75","quality":"More is better","values":[null,6568,10227,13480,10996,4515],"isMain":false},{"metricName":"NA50","quality":"More is better","values":[514,3015,7284,5753,6277,10189],"isMain":false},{"metricName":"NA75","quality":"More is better","values":[258,572,1872,648,1045,3324],"isMain":false},{"metricName":"NGA50","quality":"More is better","values":[null,6830,9721,9516,9347,10189],"isMain":true},{"metricName":"NGA75","quality":"More is better","values":[null,3261,4079,3953,4012,3285],"isMain":false},{"metricName":"LG50","quality":"Less is better","values":[null,151,89,82,93,130],"isMain":false},{"metricName":"LG75","quality":"Less is better","values":[null,340,210,174,203,340],"isMain":false},{"metricName":"LA50","quality":"Less is better","values":[359,660,250,316,302,171],"isMain":false},{"metricName":"LA75","quality":"Less is better","values":[577,3438,759,1300,1027,479],"isMain":false},{"metricName":"LGA50","quality":"Less is better","values":[null,262,176,181,189,171],"isMain":true},{"metricName":"LGA75","quality":"Less is better","values":[null,615,441,458,461,482],"isMain":false}]],["Reads mapping",[]],["Misassemblies",[{"metricName":"# misassemblies","quality":"Less is better","values":[21,372,296,334,366,205],"isMain":true},{"metricName":" # relocations","quality":"Less is better","values":[0,3,2,3,4,1],"isMain":false},{"metricName":" # translocations","quality":"Less is better","values":[4,316,290,323,354,204],"isMain":false},{"metricName":" # inversions","quality":"Less is better","values":[17,53,4,8,8,0],"isMain":false},{"metricName":" # interspecies translocations","quality":"Less is better","values":[136,324,175,287,297,45],"isMain":false},{"metricName":"# misassembled contigs","quality":"Less is better","values":[20,305,226,256,272,159],"isMain":false},{"metricName":"Misassembled contigs length","quality":"Less is better","values":[10933,2072780,2452750,2871649,2944025,1955819],"isMain":true},{"metricName":"# possibly misassembled contigs","quality":"Less is better","values":[0,20,11,23,14,5],"isMain":false},{"metricName":" # possible misassemblies","quality":"Less is better","values":[0,29,15,33,21,5],"isMain":false},{"metricName":"# local misassemblies","quality":"Less is better","values":[6,24,7,11,7,1],"isMain":false},{"metricName":"# scaffold gap ext. mis.","quality":"Less is better","values":[0,0,0,0,0,0],"isMain":false},{"metricName":"# scaffold gap loc. mis.","quality":"Less is better","values":[0,0,0,0,0,0],"isMain":false},{"metricName":"# unaligned mis. contigs","quality":"Less is better","values":[0,13,13,33,15,2],"isMain":false}]],["Unaligned",[{"metricName":"# fully unaligned contigs","quality":"Less is better","values":[0,5,3,7,5,0],"isMain":false},{"metricName":"Fully unaligned length","quality":"Less is better","values":[0,4848,1995,7530,2703,0],"isMain":false},{"metricName":"# partially unaligned contigs","quality":"Less is better","values":[63,214,128,225,195,36],"isMain":false},{"metricName":"Partially unaligned length","quality":"Less is better","values":[50409,1014708,1157546,1772967,1322567,231023],"isMain":false}]],["Mismatches",[{"metricName":"# mismatches","quality":"Less is better","values":[7715,83858,9994,13735,16656,1056],"isMain":false},{"metricName":"# indels","quality":"Less is better","values":[4,215,93,171,135,40],"isMain":false},{"metricName":"Indels length","quality":"Less is better","values":[4,902,642,1245,826,88],"isMain":false},{"metricName":"# mismatches per 100 kbp","quality":"Less is better","values":["2515.73","1303.91","155.52","213.13","257.94","16.63"],"isMain":true},{"metricName":"# indels per 100 kbp","quality":"Less is better","values":["1.30","3.34","1.45","2.65","2.09","0.63"],"isMain":true},{"metricName":" # indels (<= 5 bp)","quality":"Less is better","values":[4,192,82,142,115,36],"isMain":false},{"metricName":" # indels (> 5 bp)","quality":"Less is better","values":[0,23,11,29,20,4],"isMain":false},{"metricName":"# N's","quality":"Less is better","values":[0,0,0,0,0,0],"isMain":false},{"metricName":"# N's per 100 kbp","quality":"Less is better","values":["0.00","0.00","0.00","0.00","0.00","0.00"],"isMain":true}]],["Statistics without reference",[{"metricName":"# contigs","quality":"Equal","values":[680,5972,1436,1585,1885,1246],"isMain":true},{"metricName":"# contigs (>= 1000 bp)","quality":"Equal","values":[18,1163,864,879,867,983],"isMain":false},{"metricName":"# contigs (>= 5000 bp)","quality":"Equal","values":[0,437,397,422,421,314],"isMain":false},{"metricName":"# contigs (>= 10000 bp)","quality":"Equal","values":[0,209,216,232,225,187],"isMain":false},{"metricName":"# contigs (>= 25000 bp)","quality":"Equal","values":[0,31,58,74,68,46],"isMain":false},{"metricName":"# contigs (>= 50000 bp)","quality":"Equal","values":[0,6,10,16,11,7],"isMain":false},{"metricName":"Largest contig","quality":"More is better","values":[2322,184174,365478,117383,112101,168904],"isMain":true},{"metricName":"Total length","quality":"More is better","values":[396893,10223458,7841637,8592786,8324911,6613689],"isMain":true},{"metricName":"Total length (>= 1000 bp)","quality":"More is better","values":[25281,7388449,7487421,8149444,7715816,6423994],"isMain":true},{"metricName":"Total length (>= 5000 bp)","quality":"More is better","values":[0,5537705,6303266,6982412,6589598,4850079],"isMain":false},{"metricName":"Total length (>= 10000 bp)","quality":"More is better","values":[0,3949640,5033489,5661865,5208392,3986488],"isMain":true},{"metricName":"Total length (>= 25000 bp)","quality":"More is better","values":[0,1263468,2637270,3142310,2749742,1834771],"isMain":false},{"metricName":"Total length (>= 50000 bp)","quality":"More is better","values":[0,508211,1017597,1134658,857202,510068],"isMain":true},{"metricName":"N50","quality":"More is better","values":[543,6205,14854,17066,14896,13229],"isMain":false},{"metricName":"N75","quality":"More is better","values":[516,734,6546,6747,6176,4546],"isMain":false},{"metricName":"L50","quality":"Less is better","values":[297,362,126,129,141,129],"isMain":false},{"metricName":"L75","quality":"Less is better","values":[485,1507,324,331,360,338],"isMain":false},{"metricName":"GC (%)","quality":"Equal","values":["47.78","48.35","47.92","48.14","48.25","48.22"],"isMain":false}]],["K-mer-based statistics",[]],["Predicted genes",[]],["Similarity statistics",[]],["Reference statistics",[{"metricName":"Reference length","quality":"Equal","values":[6626223,6626223,6626223,6626223,6626223,6626223],"isMain":false},{"metricName":"Reference fragments","quality":"Equal","values":[1295,1295,1295,1295,1295,1295],"isMain":false},{"metricName":"Reference GC (%)","quality":"Equal","values":["48.12","48.12","48.12","48.12","48.12","48.12"],"isMain":false}]]],"subreferences":[],"subreports":[],"minContig":500},{"date":"10 February 2020, Monday, 18:42:39","assembliesNames":["megahit-103-df","megahit-113-df","megahit-113-ms","megahit-129-df","metaSPAdes"],"referenceName":"GCF\_000160135.1\_ASM16013v1","order":[0,1,2,3,4],"report":[["Genome statistics",[{"metricName":"Genome fraction (%)","quality":"More is better","values":["0.052","0.004","0.056","0.004","0.054"],"isMain":true},{"metricName":"Duplication ratio","quality":"Less is better","values":["1.334","5.336","1.103","5.419","1.535"],"isMain":true},{"metricName":"Largest alignment","quality":"More is better","values":[735,107,502,105,564],"isMain":true},{"metricName":"Total aligned length","quality":"More is better","values":[1391,107,1505,105,1436],"isMain":true},{"metricName":"NA50","quality":"More is better","values":[551,null,null,null,null],"isMain":false},{"metricName":"NGA50","quality":"More is better","values":[null,null,null,null,null],"isMain":true},{"metricName":"LA50","quality":"Less is better","values":[2,null,null,null,null],"isMain":false}]],["Reads mapping",[]],["Misassemblies",[{"metricName":"# misassemblies","quality":"Less is better","values":[0,0,0,0,0],"isMain":true},{"metricName":" # relocations","quality":"Less is better","values":[0,0,0,0,0],"isMain":false},{"metricName":" # translocations","quality":"Less is better","values":[0,0,0,0,0],"isMain":false},{"metricName":" # inversions","quality":"Less is better","values":[0,0,0,0,0],"isMain":false},{"metricName":" # interspecies translocations","quality":"Less is better","values":[1,1,5,1,5],"isMain":false},{"metricName":"# misassembled contigs","quality":"Less is better","values":[0,0,0,0,0],"isMain":false},{"metricName":"Misassembled contigs length","quality":"Less is better","values":[0,0,0,0,0],"isMain":true},{"metricName":"# possibly misassembled contigs","quality":"Less is better","values":[0,0,0,0,0],"isMain":false},{"metricName":" # possible misassemblies","quality":"Less is better","values":[0,0,0,0,0],"isMain":false},{"metricName":"# local misassemblies","quality":"Less is better","values":[0,0,0,0,0],"isMain":false},{"metricName":"# scaffold gap ext. mis.","quality":"Less is better","values":[0,0,0,0,0],"isMain":false},{"metricName":"# scaffold gap loc. mis.","quality":"Less is better","values":[0,0,0,0,0],"isMain":false},{"metricName":"# unaligned mis. contigs","quality":"Less is better","values":[0,0,0,0,0],"isMain":false}]],["Unaligned",[{"metricName":"# fully unaligned contigs","quality":"Less is better","values":[0,0,0,0,0],"isMain":false},{"metricName":"Fully unaligned length","quality":"Less is better","values":[0,0,0,0,0],"isMain":false},{"metricName":"# partially unaligned contigs","quality":"Less is better","values":[0,0,4,0,3],"isMain":false},{"metricName":"Partially unaligned length","quality":"Less is better","values":[0,0,3128,0,2046],"isMain":false}]],["Mismatches",[{"metricName":"# mismatches","quality":"Less is better","values":[10,2,39,2,38],"isMain":false},{"metricName":"# indels","quality":"Less is better","values":[0,0,0,0,0],"isMain":false},{"metricName":"Indels length","quality":"Less is better","values":[0,0,0,0,0],"isMain":false},{"metricName":"# mismatches per 100 kbp","quality":"Less is better","values":["718.91","1869.16","2591.36","1904.76","2646.24"],"isMain":true},{"metricName":"# indels per 100 kbp","quality":"Less is better","values":["0.00","0.00","0.00","0.00","0.00"],"isMain":true},{"metricName":" # indels (<= 5 bp)","quality":"Less is better","values":[0,0,0,0,0],"isMain":false},{"metricName":" # indels (> 5 bp)","quality":"Less is better","values":[0,0,0,0,0],"isMain":false},{"metricName":"# N's","quality":"Less is better","values":[0,0,0,0,0],"isMain":false},{"metricName":"# N's per 100 kbp","quality":"Less is better","values":["0.00","0.00","0.00","0.00","0.00"],"isMain":true}]],["Statistics without reference",[{"metricName":"# contigs","quality":"Equal","values":[3,1,6,1,6],"isMain":true},{"metricName":"# contigs (>= 1000 bp)","quality":"Equal","values":[0,0,1,0,0],"isMain":false},{"metricName":"# contigs (>= 5000 bp)","quality":"Equal","values":[0,0,0,0,0],"isMain":false},{"metricName":"# contigs (>= 10000 bp)","quality":"Equal","values":[0,0,0,0,0],"isMain":false},{"metricName":"# contigs (>= 25000 bp)","quality":"Equal","values":[0,0,0,0,0],"isMain":false},{"metricName":"# contigs (>= 50000 bp)","quality":"Equal","values":[0,0,0,0,0],"isMain":false},{"metricName":"Largest contig","quality":"More is better","values":[735,571,1408,569,948],"isMain":true},{"metricName":"Total length","quality":"More is better","values":[1855,571,4788,569,4250],"isMain":true},{"metricName":"Total length (>= 1000 bp)","quality":"More is better","values":[0,0,1408,0,0],"isMain":true},{"metricName":"Total length (>= 5000 bp)","quality":"More is better","values":[0,0,0,0,0],"isMain":false},{"metricName":"Total length (>= 10000 bp)","quality":"More is better","values":[0,0,0,0,0],"isMain":true},{"metricName":"Total length (>= 25000 bp)","quality":"More is better","values":[0,0,0,0,0],"isMain":false},{"metricName":"Total length (>= 50000 bp)","quality":"More is better","values":[0,0,0,0,0],"isMain":true},{"metricName":"N50","quality":"More is better","values":[569,571,801,569,798],"isMain":false},{"metricName":"N75","quality":"More is better","values":[551,571,693,569,564],"isMain":false},{"metricName":"L50","quality":"Less is better","values":[2,1,3,1,3],"isMain":false},{"metricName":"L75","quality":"Less is better","values":[3,1,4,1,5],"isMain":false},{"metricName":"GC (%)","quality":"Equal","values":["44.80","45.71","45.66","45.69","46.73"],"isMain":false}]],["K-mer-based statistics",[]],["Predicted genes",[]],["Similarity statistics",[]],["Reference statistics",[{"metricName":"Reference length","quality":"Equal","values":[2672954,2672954,2672954,2672954,2672954],"isMain":false},{"metricName":"Reference fragments","quality":"Equal","values":[48,48,48,48,48],"isMain":false},{"metricName":"Reference GC (%)","quality":"Equal","values":["54.68","54.68","54.68","54.68","54.68"],"isMain":false}]]],"subreferences":[],"subreports":[],"minContig":500},{"date":"10 February 2020, Monday, 18:42:36","assembliesNames":["megahit-113-df","megahit-113-ml","megahit-113-ms","megahit-129-df","metaSPAdes"],"referenceName":"GCF\_000160675.1\_ASM16067v1","order":[0,1,2,3,4],"report":[["Genome statistics",[{"metricName":"Genome fraction (%)","quality":"More is better","values":["0.071","0.058","0.057","0.054","0.027"],"isMain":true},{"metricName":"Duplication ratio","quality":"Less is better","values":["1.200","1.001","1.004","1.004","1.000"],"isMain":true},{"metricName":"Largest alignment","quality":"More is better","values":[534,609,599,520,523],"isMain":true},{"metricName":"Total aligned length","quality":"More is better","values":[1367,1121,1105,1029,523],"isMain":true},{"metricName":"NA50","quality":"More is better","values":[509,609,599,520,523],"isMain":false},{"metricName":"NA75","quality":"More is better","values":[324,512,506,509,523],"isMain":false},{"metricName":"NGA50","quality":"More is better","values":[null,null,null,null,null],"isMain":true},{"metricName":"LA50","quality":"Less is better","values":[2,1,1,1,1],"isMain":false},{"metricName":"LA75","quality":"Less is better","values":[3,2,2,2,1],"isMain":false}]],["Reads mapping",[]],["Misassemblies",[{"metricName":"# misassemblies","quality":"Less is better","values":[0,0,0,0,0],"isMain":true},{"metricName":" # relocations","quality":"Less is better","values":[0,0,0,0,0],"isMain":false},{"metricName":" # translocations","quality":"Less is better","values":[0,0,0,0,0],"isMain":false},{"metricName":" # inversions","quality":"Less is better","values":[0,0,0,0,0],"isMain":false},{"metricName":" # interspecies translocations","quality":"Less is better","values":[1,0,0,0,0],"isMain":false},{"metricName":"# misassembled contigs","quality":"Less is better","values":[0,0,0,0,0],"isMain":false},{"metricName":"Misassembled contigs length","quality":"Less is better","values":[0,0,0,0,0],"isMain":true},{"metricName":"# possibly misassembled contigs","quality":"Less is better","values":[0,0,0,0,0],"isMain":false},{"metricName":" # possible misassemblies","quality":"Less is better","values":[0,0,0,0,0],"isMain":false},{"metricName":"# local misassemblies","quality":"Less is better","values":[0,0,0,0,0],"isMain":false},{"metricName":"# scaffold gap ext. mis.","quality":"Less is better","values":[0,0,0,0,0],"isMain":false},{"metricName":"# scaffold gap loc. mis.","quality":"Less is better","values":[0,0,0,0,0],"isMain":false},{"metricName":"# unaligned mis. contigs","quality":"Less is better","values":[0,0,0,0,0],"isMain":false}]],["Unaligned",[{"metricName":"# fully unaligned contigs","quality":"Less is better","values":[0,0,0,0,0],"isMain":false},{"metricName":"Fully unaligned length","quality":"Less is better","values":[0,0,0,0,0],"isMain":false},{"metricName":"# partially unaligned contigs","quality":"Less is better","values":[0,0,0,0,0],"isMain":false},{"metricName":"Partially unaligned length","quality":"Less is better","values":[0,0,0,0,0],"isMain":false}]],["Mismatches",[{"metricName":"# mismatches","quality":"Less is better","values":[28,18,13,13,6],"isMain":false},{"metricName":"# indels","quality":"Less is better","values":[0,0,0,0,0],"isMain":false},{"metricName":"Indels length","quality":"Less is better","values":[0,0,0,0,0],"isMain":false},{"metricName":"# mismatches per 100 kbp","quality":"Less is better","values":["2048.28","1605.71","1176.47","1263.36","1147.23"],"isMain":true},{"metricName":"# indels per 100 kbp","quality":"Less is better","values":["0.00","0.00","0.00","0.00","0.00"],"isMain":true},{"metricName":" # indels (<= 5 bp)","quality":"Less is better","values":[0,0,0,0,0],"isMain":false},{"metricName":" # indels (> 5 bp)","quality":"Less is better","values":[0,0,0,0,0],"isMain":false},{"metricName":"# N's","quality":"Less is better","values":[0,0,0,0,0],"isMain":false},{"metricName":"# N's per 100 kbp","quality":"Less is better","values":["0.00","0.00","0.00","0.00","0.00"],"isMain":true}]],["Statistics without reference",[{"metricName":"# contigs","quality":"Equal","values":[3,2,2,2,1],"isMain":true},{"metricName":"# contigs (>= 1000 bp)","quality":"Equal","values":[0,0,0,0,0],"isMain":false},{"metricName":"# contigs (>= 5000 bp)","quality":"Equal","values":[0,0,0,0,0],"isMain":false},{"metricName":"# contigs (>= 10000 bp)","quality":"Equal","values":[0,0,0,0,0],"isMain":false},{"metricName":"# contigs (>= 25000 bp)","quality":"Equal","values":[0,0,0,0,0],"isMain":false},{"metricName":"# contigs (>= 50000 bp)","quality":"Equal","values":[0,0,0,0,0],"isMain":false},{"metricName":"Largest contig","quality":"More is better","values":[581,609,603,520,523],"isMain":true},{"metricName":"Total length","quality":"More is better","values":[1641,1122,1109,1033,523],"isMain":true},{"metricName":"Total length (>= 1000 bp)","quality":"More is better","values":[0,0,0,0,0],"isMain":true},{"metricName":"Total length (>= 5000 bp)","quality":"More is better","values":[0,0,0,0,0],"isMain":false},{"metricName":"Total length (>= 10000 bp)","quality":"More is better","values":[0,0,0,0,0],"isMain":true},{"metricName":"Total length (>= 25000 bp)","quality":"More is better","values":[0,0,0,0,0],"isMain":false},{"metricName":"Total length (>= 50000 bp)","quality":"More is better","values":[0,0,0,0,0],"isMain":true},{"metricName":"N50","quality":"More is better","values":[547,609,603,520,523],"isMain":false},{"metricName":"N75","quality":"More is better","values":[513,513,506,513,523],"isMain":false},{"metricName":"L50","quality":"Less is better","values":[2,1,1,1,1],"isMain":false},{"metricName":"L75","quality":"Less is better","values":[3,2,2,2,1],"isMain":false},{"metricName":"GC (%)","quality":"Equal","values":["41.93","38.86","38.41","37.08","38.43"],"isMain":false}]],["K-mer-based statistics",[]],["Predicted genes",[]],["Similarity statistics",[]],["Reference statistics",[{"metricName":"Reference length","quality":"Equal","values":[1923088,1923088,1923088,1923088,1923088],"isMain":false},{"metricName":"Reference fragments","quality":"Equal","values":[31,31,31,31,31],"isMain":false},{"metricName":"Reference GC (%)","quality":"Equal","values":["37.65","37.65","37.65","37.65","37.65"],"isMain":false}]]],"subreferences":[],"subreports":[],"minContig":500},{"date":"10 February 2020, Monday, 18:51:24","assembliesNames":["megahit-103-df","megahit-113-df","megahit-113-ml","megahit-113-ms","megahit-129-df","metaSPAdes"],"referenceName":"GCF\_000162175.1\_ASM16217v1","order":[0,1,2,3,4,5],"report":[["Genome statistics",[{"metricName":"Genome fraction (%)","quality":"More is better","values":["16.490","18.375","15.967","16.653","19.972","11.637"],"isMain":true},{"metricName":"Duplication ratio","quality":"Less is better","values":["1.029","1.024","1.028","1.036","1.027","1.024"],"isMain":true},{"metricName":"Largest alignment","quality":"More is better","values":[2783,2622,2529,2191,2804,2185],"isMain":true},{"metricName":"Total aligned length","quality":"More is better","values":[814878,907469,792614,830334,988139,573117],"isMain":true},{"metricName":"NA50","quality":"More is better","values":[759,830,721,710,849,690],"isMain":false},{"metricName":"NA75","quality":"More is better","values":[605,631,582,573,626,570],"isMain":false},{"metricName":"NGA50","quality":"More is better","values":[null,null,null,null,null,null],"isMain":true},{"metricName":"LA50","quality":"Less is better","values":[393,390,411,446,425,323],"isMain":false},{"metricName":"LA75","quality":"Less is better","values":[704,714,721,780,782,557],"isMain":false}]],["Reads mapping",[]],["Misassemblies",[{"metricName":"# misassemblies","quality":"Less is better","values":[3,10,1,6,9,2],"isMain":true},{"metricName":" # relocations","quality":"Less is better","values":[0,1,0,1,2,0],"isMain":false},{"metricName":" # translocations","quality":"Less is better","values":[2,6,1,4,6,2],"isMain":false},{"metricName":" # inversions","quality":"Less is better","values":[1,3,0,1,1,0],"isMain":false},{"metricName":" # interspecies translocations","quality":"Less is better","values":[44,46,12,29,51,18],"isMain":false},{"metricName":"# misassembled contigs","quality":"Less is better","values":[3,10,1,6,8,2],"isMain":false},{"metricName":"Misassembled contigs length","quality":"Less is better","values":[2252,10321,600,4975,9328,1704],"isMain":true},{"metricName":"# possibly misassembled contigs","quality":"Less is better","values":[0,0,0,0,0,0],"isMain":false},{"metricName":" # possible misassemblies","quality":"Less is better","values":[0,0,0,0,0,0],"isMain":false},{"metricName":"# local misassemblies","quality":"Less is better","values":[3,1,0,4,2,2],"isMain":false},{"metricName":"# scaffold gap ext. mis.","quality":"Less is better","values":[0,0,0,0,0,0],"isMain":false},{"metricName":"# scaffold gap loc. mis.","quality":"Less is better","values":[0,0,0,0,0,0],"isMain":false},{"metricName":"# unaligned mis. contigs","quality":"Less is better","values":[0,0,0,0,0,1],"isMain":false}]],["Unaligned",[{"metricName":"# fully unaligned contigs","quality":"Less is better","values":[0,2,2,1,2,0],"isMain":false},{"metricName":"Fully unaligned length","quality":"Less is better","values":[0,1857,1534,561,1877,0],"isMain":false},{"metricName":"# partially unaligned contigs","quality":"Less is better","values":[3,6,0,5,15,4],"isMain":false},{"metricName":"Partially unaligned length","quality":"Less is better","values":[3064,11811,0,5410,36063,2708],"isMain":false}]],["Mismatches",[{"metricName":"# mismatches","quality":"Less is better","values":[13036,14403,16706,18474,16440,9359],"isMain":false},{"metricName":"# indels","quality":"Less is better","values":[38,51,58,56,52,32],"isMain":false},{"metricName":"Indels length","quality":"Less is better","values":[72,69,86,135,82,36],"isMain":false},{"metricName":"# mismatches per 100 kbp","quality":"Less is better","values":["1616.10","1602.36","2138.94","2267.87","1682.74","1644.11"],"isMain":true},{"metricName":"# indels per 100 kbp","quality":"Less is better","values":["4.71","5.67","7.43","6.87","5.32","5.62"],"isMain":true},{"metricName":" # indels (<= 5 bp)","quality":"Less is better","values":[36,50,57,53,50,32],"isMain":false},{"metricName":" # indels (> 5 bp)","quality":"Less is better","values":[2,1,1,3,2,0],"isMain":false},{"metricName":"# N's","quality":"Less is better","values":[0,0,0,0,0,0],"isMain":false},{"metricName":"# N's per 100 kbp","quality":"Less is better","values":["0.00","0.00","0.00","0.00","0.00","0.00"],"isMain":true}]],["Statistics without reference",[{"metricName":"# contigs","quality":"Equal","values":[1060,1090,1080,1150,1165,814],"isMain":true},{"metricName":"# contigs (>= 1000 bp)","quality":"Equal","values":[176,259,132,142,303,82],"isMain":false},{"metricName":"# contigs (>= 5000 bp)","quality":"Equal","values":[0,1,0,0,2,0],"isMain":false},{"metricName":"# contigs (>= 10000 bp)","quality":"Equal","values":[0,0,0,0,1,0],"isMain":false},{"metricName":"# contigs (>= 25000 bp)","quality":"Equal","values":[0,0,0,0,0,0],"isMain":false},{"metricName":"# contigs (>= 50000 bp)","quality":"Equal","values":[0,0,0,0,0,0],"isMain":false},{"metricName":"Largest contig","quality":"More is better","values":[2806,7670,2533,2194,21892,2194],"isMain":true},{"metricName":"Total length","quality":"More is better","values":[832824,934444,804385,849782,1040928,585555],"isMain":true},{"metricName":"Total length (>= 1000 bp)","quality":"More is better","values":[232799,362405,170041,179427,445866,101198],"isMain":true},{"metricName":"Total length (>= 5000 bp)","quality":"More is better","values":[0,7670,0,0,27182,0],"isMain":false},{"metricName":"Total length (>= 10000 bp)","quality":"More is better","values":[0,0,0,0,21892,0],"isMain":true},{"metricName":"Total length (>= 25000 bp)","quality":"More is better","values":[0,0,0,0,0,0],"isMain":false},{"metricName":"Total length (>= 50000 bp)","quality":"More is better","values":[0,0,0,0,0,0],"isMain":true},{"metricName":"N50","quality":"More is better","values":[785,868,733,728,903,704],"isMain":false},{"metricName":"N75","quality":"More is better","values":[623,658,596,595,678,587],"isMain":false},{"metricName":"L50","quality":"Less is better","values":[386,372,406,434,382,316],"isMain":false},{"metricName":"L75","quality":"Less is better","values":[687,683,711,758,715,544],"isMain":false},{"metricName":"GC (%)","quality":"Equal","values":["45.27","45.50","45.60","45.36","45.59","45.30"],"isMain":false}]],["K-mer-based statistics",[]],["Predicted genes",[]],["Similarity statistics",[]],["Reference statistics",[{"metricName":"Reference length","quality":"Equal","values":[4891696,4891696,4891696,4891696,4891696,4891696],"isMain":false},{"metricName":"Reference fragments","quality":"Equal","values":[71,71,71,71,71,71],"isMain":false},{"metricName":"Reference GC (%)","quality":"Equal","values":["44.87","44.87","44.87","44.87","44.87","44.87"],"isMain":false}]]],"subreferences":[],"subreports":[],"minContig":500},{"date":"10 February 2020, Monday, 18:51:22","assembliesNames":["megahit-103-df","megahit-113-df","megahit-113-ml","megahit-113-ms","megahit-129-df","metaSPAdes"],"referenceName":"GCF\_000162215.1\_ASM16221v1","order":[0,1,2,3,4,5],"report":[["Genome statistics",[{"metricName":"Genome fraction (%)","quality":"More is better","values":["1.025","1.356","1.551","2.196","1.473","0.455"],"isMain":true},{"metricName":"Duplication ratio","quality":"Less is better","values":["1.119","1.100","1.061","1.058","1.065","1.162"],"isMain":true},{"metricName":"Largest alignment","quality":"More is better","values":[792,1131,979,1064,960,975],"isMain":true},{"metricName":"Total aligned length","quality":"More is better","values":[45689,60361,69142,97928,65608,20275],"isMain":true},{"metricName":"NA50","quality":"More is better","values":[524,542,539,565,514,542],"isMain":false},{"metricName":"NA75","quality":"More is better","values":[345,477,505,513,null,397],"isMain":false},{"metricName":"NGA50","quality":"More is better","values":[null,null,null,null,null,null],"isMain":true},{"metricName":"LA50","quality":"Less is better","values":[47,57,60,81,87,19],"isMain":false},{"metricName":"LA75","quality":"Less is better","values":[76,92,96,130,null,31],"isMain":false}]],["Reads mapping",[]],["Misassemblies",[{"metricName":"# misassemblies","quality":"Less is better","values":[0,0,0,0,1,0],"isMain":true},{"metricName":" # relocations","quality":"Less is better","values":[0,0,0,0,0,0],"isMain":false},{"metricName":" # translocations","quality":"Less is better","values":[0,0,0,0,1,0],"isMain":false},{"metricName":" # inversions","quality":"Less is better","values":[0,0,0,0,0,0],"isMain":false},{"metricName":" # interspecies translocations","quality":"Less is better","values":[22,25,13,18,20,10],"isMain":false},{"metricName":"# misassembled contigs","quality":"Less is better","values":[0,0,0,0,1,0],"isMain":false},{"metricName":"Misassembled contigs length","quality":"Less is better","values":[0,0,0,0,729,0],"isMain":true},{"metricName":"# possibly misassembled contigs","quality":"Less is better","values":[0,1,0,0,0,0],"isMain":false},{"metricName":" # possible misassemblies","quality":"Less is better","values":[0,1,0,0,0,0],"isMain":false},{"metricName":"# local misassemblies","quality":"Less is better","values":[0,0,1,0,0,0],"isMain":false},{"metricName":"# scaffold gap ext. mis.","quality":"Less is better","values":[0,0,0,0,0,0],"isMain":false},{"metricName":"# scaffold gap loc. mis.","quality":"Less is better","values":[0,0,0,0,0,0],"isMain":false},{"metricName":"# unaligned mis. contigs","quality":"Less is better","values":[0,1,0,0,0,1],"isMain":false}]],["Unaligned",[{"metricName":"# fully unaligned contigs","quality":"Less is better","values":[0,1,1,0,0,1],"isMain":false},{"metricName":"Fully unaligned length","quality":"Less is better","values":[0,923,977,0,0,656],"isMain":false},{"metricName":"# partially unaligned contigs","quality":"Less is better","values":[5,6,1,3,10,0],"isMain":false},{"metricName":"Partially unaligned length","quality":"Less is better","values":[4766,4632,777,2630,35844,0],"isMain":false}]],["Mismatches",[{"metricName":"# mismatches","quality":"Less is better","values":[837,1198,1435,2178,1309,384],"isMain":false},{"metricName":"# indels","quality":"Less is better","values":[1,3,5,5,8,0],"isMain":false},{"metricName":"Indels length","quality":"Less is better","values":[2,3,11,9,12,0],"isMain":false},{"metricName":"# mismatches per 100 kbp","quality":"Less is better","values":["1833.48","1984.69","2078.08","2227.11","1995.31","1893.96"],"isMain":true},{"metricName":"# indels per 100 kbp","quality":"Less is better","values":["2.19","4.97","7.24","5.11","12.19","0.00"],"isMain":true},{"metricName":" # indels (<= 5 bp)","quality":"Less is better","values":[1,3,4,5,8,0],"isMain":false},{"metricName":" # indels (> 5 bp)","quality":"Less is better","values":[0,0,1,0,0,0],"isMain":false},{"metricName":"# N's","quality":"Less is better","values":[0,0,0,0,0,0],"isMain":false},{"metricName":"# N's per 100 kbp","quality":"Less is better","values":["0.00","0.00","0.00","0.00","0.00","0.00"],"isMain":true}]],["Statistics without reference",[{"metricName":"# contigs","quality":"Equal","values":[92,114,127,174,120,40],"isMain":true},{"metricName":"# contigs (>= 1000 bp)","quality":"Equal","values":[4,4,0,3,5,0],"isMain":false},{"metricName":"# contigs (>= 5000 bp)","quality":"Equal","values":[0,0,0,0,1,0],"isMain":false},{"metricName":"# contigs (>= 10000 bp)","quality":"Equal","values":[0,0,0,0,1,0],"isMain":false},{"metricName":"# contigs (>= 25000 bp)","quality":"Equal","values":[0,0,0,0,1,0],"isMain":false},{"metricName":"# contigs (>= 50000 bp)","quality":"Equal","values":[0,0,0,0,0,0],"isMain":false},{"metricName":"Largest contig","quality":"More is better","values":[1539,1691,983,1625,30143,987],"isMain":true},{"metricName":"Total length","quality":"More is better","values":[55849,71976,75032,106056,105729,24218],"isMain":true},{"metricName":"Total length (>= 1000 bp)","quality":"More is better","values":[4785,5302,0,3817,35234,0],"isMain":true},{"metricName":"Total length (>= 5000 bp)","quality":"More is better","values":[0,0,0,0,30143,0],"isMain":false},{"metricName":"Total length (>= 10000 bp)","quality":"More is better","values":[0,0,0,0,30143,0],"isMain":true},{"metricName":"Total length (>= 25000 bp)","quality":"More is better","values":[0,0,0,0,30143,0],"isMain":false},{"metricName":"Total length (>= 50000 bp)","quality":"More is better","values":[0,0,0,0,0,0],"isMain":true},{"metricName":"N50","quality":"More is better","values":[579,599,571,588,705,575],"isMain":false},{"metricName":"N75","quality":"More is better","values":[536,541,523,531,554,530],"isMain":false},{"metricName":"L50","quality":"Less is better","values":[40,48,56,75,27,18],"isMain":false},{"metricName":"L75","quality":"Less is better","values":[65,80,91,123,70,29],"isMain":false},{"metricName":"GC (%)","quality":"Equal","values":["46.04","47.16","47.05","46.21","46.37","46.51"],"isMain":false}]],["K-mer-based statistics",[]],["Predicted genes",[]],["Similarity statistics",[]],["Reference statistics",[{"metricName":"Reference length","quality":"Equal","values":[4452513,4452513,4452513,4452513,4452513,4452513],"isMain":false},{"metricName":"Reference fragments","quality":"Equal","values":[63,63,63,63,63,63],"isMain":false},{"metricName":"Reference GC (%)","quality":"Equal","values":["46.38","46.38","46.38","46.38","46.38","46.38"],"isMain":false}]]],"subreferences":[],"subreports":[],"minContig":500},{"date":"10 February 2020, Monday, 18:51:31","assembliesNames":["megahit-103-df","megahit-113-df","megahit-113-ml","megahit-113-ms","megahit-129-df","metaSPAdes"],"referenceName":"GCF\_000162275.1\_ASM16227v1","order":[0,1,2,3,4,5],"report":[["Genome statistics",[{"metricName":"Genome fraction (%)","quality":"More is better","values":["3.208","3.140","3.473","3.505","3.688","2.051"],"isMain":true},{"metricName":"Duplication ratio","quality":"Less is better","values":["1.023","1.025","1.020","1.016","1.020","1.016"],"isMain":true},{"metricName":"Largest alignment","quality":"More is better","values":[1903,2070,1996,2101,2438,1600],"isMain":true},{"metricName":"Total aligned length","quality":"More is better","values":[171522,167894,185908,187239,197195,109478],"isMain":true},{"metricName":"NA50","quality":"More is better","values":[732,758,751,708,771,655],"isMain":false},{"metricName":"NA75","quality":"More is better","values":[594,592,593,586,589,571],"isMain":false},{"metricName":"NGA50","quality":"More is better","values":[null,null,null,null,null,null],"isMain":true},{"metricName":"LA50","quality":"Less is better","values":[88,84,95,103,96,69],"isMain":false},{"metricName":"LA75","quality":"Less is better","values":[156,151,166,178,174,115],"isMain":false}]],["Reads mapping",[]],["Misassemblies",[{"metricName":"# misassemblies","quality":"Less is better","values":[0,2,0,0,1,0],"isMain":true},{"metricName":" # relocations","quality":"Less is better","values":[0,0,0,0,1,0],"isMain":false},{"metricName":" # translocations","quality":"Less is better","values":[0,2,0,0,0,0],"isMain":false},{"metricName":" # inversions","quality":"Less is better","values":[0,0,0,0,0,0],"isMain":false},{"metricName":" # interspecies translocations","quality":"Less is better","values":[10,11,4,5,13,4],"isMain":false},{"metricName":"# misassembled contigs","quality":"Less is better","values":[0,2,0,0,1,0],"isMain":false},{"metricName":"Misassembled contigs length","quality":"Less is better","values":[0,2233,0,0,741,0],"isMain":true},{"metricName":"# possibly misassembled contigs","quality":"Less is better","values":[0,0,0,0,0,0],"isMain":false},{"metricName":" # possible misassemblies","quality":"Less is better","values":[0,0,0,0,0,0],"isMain":false},{"metricName":"# local misassemblies","quality":"Less is better","values":[2,1,0,0,2,0],"isMain":false},{"metricName":"# scaffold gap ext. mis.","quality":"Less is better","values":[0,0,0,0,0,0],"isMain":false},{"metricName":"# scaffold gap loc. mis.","quality":"Less is better","values":[0,0,0,0,0,0],"isMain":false},{"metricName":"# unaligned mis. contigs","quality":"Less is better","values":[0,1,0,0,0,0],"isMain":false}]],["Unaligned",[{"metricName":"# fully unaligned contigs","quality":"Less is better","values":[0,0,0,2,0,0],"isMain":false},{"metricName":"Fully unaligned length","quality":"Less is better","values":[0,0,0,2273,0,0],"isMain":false},{"metricName":"# partially unaligned contigs","quality":"Less is better","values":[1,3,2,0,7,2],"isMain":false},{"metricName":"Partially unaligned length","quality":"Less is better","values":[809,6051,1537,0,9504,1161],"isMain":false}]],["Mismatches",[{"metricName":"# mismatches","quality":"Less is better","values":[2755,2762,3845,4121,3332,1819],"isMain":false},{"metricName":"# indels","quality":"Less is better","values":[20,19,17,18,22,6],"isMain":false},{"metricName":"Indels length","quality":"Less is better","values":[30,27,21,30,99,8],"isMain":false},{"metricName":"# mismatches per 100 kbp","quality":"Less is better","values":["1612.22","1650.97","2077.99","2206.91","1696.11","1664.73"],"isMain":true},{"metricName":"# indels per 100 kbp","quality":"Less is better","values":["11.70","11.36","9.19","9.64","11.20","5.49"],"isMain":true},{"metricName":" # indels (<= 5 bp)","quality":"Less is better","values":[19,19,17,17,20,6],"isMain":false},{"metricName":" # indels (> 5 bp)","quality":"Less is better","values":[1,0,0,1,2,0],"isMain":false},{"metricName":"# N's","quality":"Less is better","values":[0,0,0,0,0,0],"isMain":false},{"metricName":"# N's per 100 kbp","quality":"Less is better","values":["0.00","0.00","0.00","0.00","0.00","0.00"],"isMain":true}]],["Statistics without reference",[{"metricName":"# contigs","quality":"Equal","values":[232,217,248,261,250,164],"isMain":true},{"metricName":"# contigs (>= 1000 bp)","quality":"Equal","values":[37,44,34,30,56,12],"isMain":false},{"metricName":"# contigs (>= 5000 bp)","quality":"Equal","values":[0,0,0,0,0,0],"isMain":false},{"metricName":"# contigs (>= 10000 bp)","quality":"Equal","values":[0,0,0,0,0,0],"isMain":false},{"metricName":"# contigs (>= 25000 bp)","quality":"Equal","values":[0,0,0,0,0,0],"isMain":false},{"metricName":"# contigs (>= 50000 bp)","quality":"Equal","values":[0,0,0,0,0,0],"isMain":false},{"metricName":"Largest contig","quality":"More is better","values":[1903,3708,2023,2104,3525,1604],"isMain":true},{"metricName":"Total length","quality":"More is better","values":[175574,177599,190266,192053,209828,112193],"isMain":true},{"metricName":"Total length (>= 1000 bp)","quality":"More is better","values":[45112,60282,44944,37585,78119,13888],"isMain":true},{"metricName":"Total length (>= 5000 bp)","quality":"More is better","values":[0,0,0,0,0,0],"isMain":false},{"metricName":"Total length (>= 10000 bp)","quality":"More is better","values":[0,0,0,0,0,0],"isMain":true},{"metricName":"Total length (>= 25000 bp)","quality":"More is better","values":[0,0,0,0,0,0],"isMain":false},{"metricName":"Total length (>= 50000 bp)","quality":"More is better","values":[0,0,0,0,0,0],"isMain":true},{"metricName":"N50","quality":"More is better","values":[755,818,759,718,829,668],"isMain":false},{"metricName":"N75","quality":"More is better","values":[607,629,609,597,649,583],"isMain":false},{"metricName":"L50","quality":"Less is better","values":[87,76,93,100,86,67],"isMain":false},{"metricName":"L75","quality":"Less is better","values":[153,139,163,174,158,112],"isMain":false},{"metricName":"GC (%)","quality":"Equal","values":["45.13","45.36","45.63","45.43","45.69","45.60"],"isMain":false}]],["K-mer-based statistics",[]],["Predicted genes",[]],["Similarity statistics",[]],["Reference statistics",[{"metricName":"Reference length","quality":"Equal","values":[5327234,5327234,5327234,5327234,5327234,5327234],"isMain":false},{"metricName":"Reference fragments","quality":"Equal","values":[74,74,74,74,74,74],"isMain":false},{"metricName":"Reference GC (%)","quality":"Equal","values":["45.18","45.18","45.18","45.18","45.18","45.18"],"isMain":false}]]],"subreferences":[],"subreports":[],"minContig":500},{"date":"10 February 2020, Monday, 18:51:28","assembliesNames":["megahit-103-df","megahit-113-df","megahit-113-ml","megahit-113-ms","megahit-129-df","metaSPAdes"],"referenceName":"GCF\_000162455.1\_ASM16245v1","order":[0,1,2,3,4,5],"report":[["Genome statistics",[{"metricName":"Genome fraction (%)","quality":"More is better","values":["9.580","14.650","8.900","10.653","10.016","5.666"],"isMain":true},{"metricName":"Duplication ratio","quality":"Less is better","values":["1.040","1.021","1.063","1.059","1.041","1.034"],"isMain":true},{"metricName":"Largest alignment","quality":"More is better","values":[19868,24869,19106,19852,19106,8647],"isMain":true},{"metricName":"Total aligned length","quality":"More is better","values":[324993,488955,307520,364616,338853,191201],"isMain":true},{"metricName":"NA50","quality":"More is better","values":[963,1976,807,976,1154,862],"isMain":false},{"metricName":"NA75","quality":"More is better","values":[682,833,608,626,695,634],"isMain":false},{"metricName":"NGA50","quality":"More is better","values":[null,null,null,null,null,null],"isMain":true},{"metricName":"LA50","quality":"Less is better","values":[90,44,117,87,75,74],"isMain":false},{"metricName":"LA75","quality":"Less is better","values":[195,153,233,216,175,139],"isMain":false}]],["Reads mapping",[]],["Misassemblies",[{"metricName":"# misassemblies","quality":"Less is better","values":[5,28,2,13,3,0],"isMain":true},{"metricName":" # relocations","quality":"Less is better","values":[0,2,0,1,0,0],"isMain":false},{"metricName":" # translocations","quality":"Less is better","values":[5,26,2,11,2,0],"isMain":false},{"metricName":" # inversions","quality":"Less is better","values":[0,0,0,1,1,0],"isMain":false},{"metricName":" # interspecies translocations","quality":"Less is better","values":[16,14,13,22,14,3],"isMain":false},{"metricName":"# misassembled contigs","quality":"Less is better","values":[5,2,2,3,2,0],"isMain":false},{"metricName":"Misassembled contigs length","quality":"Less is better","values":[8153,204322,22274,86184,16573,0],"isMain":true},{"metricName":"# possibly misassembled contigs","quality":"Less is better","values":[0,0,1,0,0,0],"isMain":false},{"metricName":" # possible misassemblies","quality":"Less is better","values":[0,0,1,0,0,0],"isMain":false},{"metricName":"# local misassemblies","quality":"Less is better","values":[1,3,0,1,1,0],"isMain":false},{"metricName":"# scaffold gap ext. mis.","quality":"Less is better","values":[0,0,0,0,0,0],"isMain":false},{"metricName":"# scaffold gap loc. mis.","quality":"Less is better","values":[0,0,0,0,0,0],"isMain":false},{"metricName":"# unaligned mis. contigs","quality":"Less is better","values":[0,1,1,1,0,1],"isMain":false}]],["Unaligned",[{"metricName":"# fully unaligned contigs","quality":"Less is better","values":[0,1,0,4,0,0],"isMain":false},{"metricName":"Fully unaligned length","quality":"Less is better","values":[0,566,0,2630,0,0],"isMain":false},{"metricName":"# partially unaligned contigs","quality":"Less is better","values":[6,7,7,7,5,2],"isMain":false},{"metricName":"Partially unaligned length","quality":"Less is better","values":[5165,25524,12221,23092,7363,2899],"isMain":false}]],["Mismatches",[{"metricName":"# mismatches","quality":"Less is better","values":[3865,5270,5227,6002,3970,2132],"isMain":false},{"metricName":"# indels","quality":"Less is better","values":[28,95,24,47,21,6],"isMain":false},{"metricName":"Indels length","quality":"Less is better","values":[67,571,32,212,28,53],"isMain":false},{"metricName":"# mismatches per 100 kbp","quality":"Less is better","values":["1223.37","1090.79","1780.71","1708.28","1201.84","1140.98"],"isMain":true},{"metricName":"# indels per 100 kbp","quality":"Less is better","values":["8.86","19.66","8.18","13.38","6.36","3.21"],"isMain":true},{"metricName":" # indels (<= 5 bp)","quality":"Less is better","values":[26,75,24,42,21,5],"isMain":false},{"metricName":" # indels (> 5 bp)","quality":"Less is better","values":[2,20,0,5,0,1],"isMain":false},{"metricName":"# N's","quality":"Less is better","values":[0,0,0,0,0,0],"isMain":false},{"metricName":"# N's per 100 kbp","quality":"Less is better","values":["0.00","0.00","0.00","0.00","0.00","0.00"],"isMain":true}]],["Statistics without reference",[{"metricName":"# contigs","quality":"Equal","values":[327,287,355,336,307,218],"isMain":true},{"metricName":"# contigs (>= 1000 bp)","quality":"Equal","values":[87,97,64,79,94,53],"isMain":false},{"metricName":"# contigs (>= 5000 bp)","quality":"Equal","values":[4,5,3,4,5,1],"isMain":false},{"metricName":"# contigs (>= 10000 bp)","quality":"Equal","values":[1,3,2,2,3,0],"isMain":false},{"metricName":"# contigs (>= 25000 bp)","quality":"Equal","values":[0,1,0,1,0,0],"isMain":false},{"metricName":"# contigs (>= 50000 bp)","quality":"Equal","values":[0,1,0,1,0,0],"isMain":false},{"metricName":"Largest contig","quality":"More is better","values":[19868,202920,19106,84674,19106,8663],"isMain":true},{"metricName":"Total length","quality":"More is better","values":[333810,519576,324277,397652,351232,196083],"isMain":true},{"metricName":"Total length (>= 1000 bp)","quality":"More is better","values":[168479,390754,126301,222792,206448,82593],"isMain":true},{"metricName":"Total length (>= 5000 bp)","quality":"More is better","values":[42388,245814,41380,117321,60832,8663],"isMain":false},{"metricName":"Total length (>= 10000 bp)","quality":"More is better","values":[19868,233922,32903,103763,48849,0],"isMain":true},{"metricName":"Total length (>= 25000 bp)","quality":"More is better","values":[0,202920,0,84674,0,0],"isMain":false},{"metricName":"Total length (>= 50000 bp)","quality":"More is better","values":[0,202920,0,84674,0,0],"isMain":true},{"metricName":"N50","quality":"More is better","values":[1006,3440,858,1100,1199,881],"isMain":false},{"metricName":"N75","quality":"More is better","values":[704,1007,650,736,752,663],"isMain":false},{"metricName":"L50","quality":"Less is better","values":[86,9,103,57,67,70],"isMain":false},{"metricName":"L75","quality":"Less is better","values":[187,96,212,169,161,133],"isMain":false},{"metricName":"GC (%)","quality":"Equal","values":["45.86","49.20","45.23","47.08","45.56","44.07"],"isMain":false}]],["K-mer-based statistics",[]],["Predicted genes",[]],["Similarity statistics",[]],["Reference statistics",[{"metricName":"Reference length","quality":"Equal","values":[3297977,3297977,3297977,3297977,3297977,3297977],"isMain":false},{"metricName":"Reference fragments","quality":"Equal","values":[267,267,267,267,267,267],"isMain":false},{"metricName":"Reference GC (%)","quality":"Equal","values":["50.95","50.95","50.95","50.95","50.95","50.95"],"isMain":false}]]],"subreferences":[],"subreports":[],"minContig":500},{"date":"10 February 2020, Monday, 18:51:09","assembliesNames":["megahit-103-df","megahit-113-df","megahit-113-ml","megahit-113-ms","megahit-129-df","metaSPAdes"],"referenceName":"GCF\_000163055.2\_ASM16305v2","order":[0,1,2,3,4,5],"report":[["Genome statistics",[{"metricName":"Genome fraction (%)","quality":"More is better","values":["0.116","0.137","0.199","0.199","0.136","0.090"],"isMain":true},{"metricName":"Duplication ratio","quality":"Less is better","values":["1.259","1.274","1.168","1.293","1.064","1.218"],"isMain":true},{"metricName":"Largest alignment","quality":"More is better","values":[659,703,1030,654,974,962],"isMain":true},{"metricName":"Total aligned length","quality":"More is better","values":[2888,3390,4925,4938,3383,2236],"isMain":true},{"metricName":"NA50","quality":"More is better","values":[577,346,531,null,506,487],"isMain":false},{"metricName":"NA75","quality":"More is better","values":[232,null,246,null,null,null],"isMain":false},{"metricName":"NGA50","quality":"More is better","values":[null,null,null,null,null,null],"isMain":true},{"metricName":"LA50","quality":"Less is better","values":[3,5,5,null,4,3],"isMain":false},{"metricName":"LA75","quality":"Less is better","values":[6,null,9,null,null,null],"isMain":false}]],["Reads mapping",[]],["Misassemblies",[{"metricName":"# misassemblies","quality":"Less is better","values":[0,0,0,0,0,0],"isMain":true},{"metricName":" # relocations","quality":"Less is better","values":[0,0,0,0,0,0],"isMain":false},{"metricName":" # translocations","quality":"Less is better","values":[0,0,0,0,0,0],"isMain":false},{"metricName":" # inversions","quality":"Less is better","values":[0,0,0,0,0,0],"isMain":false},{"metricName":" # interspecies translocations","quality":"Less is better","values":[3,3,1,9,2,2],"isMain":false},{"metricName":"# misassembled contigs","quality":"Less is better","values":[0,0,0,0,0,0],"isMain":false},{"metricName":"Misassembled contigs length","quality":"Less is better","values":[0,0,0,0,0,0],"isMain":true},{"metricName":"# possibly misassembled contigs","quality":"Less is better","values":[0,0,0,1,0,0],"isMain":false},{"metricName":" # possible misassemblies","quality":"Less is better","values":[0,0,0,1,0,0],"isMain":false},{"metricName":"# local misassemblies","quality":"Less is better","values":[0,0,0,0,0,0],"isMain":false},{"metricName":"# scaffold gap ext. mis.","quality":"Less is better","values":[0,0,0,0,0,0],"isMain":false},{"metricName":"# scaffold gap loc. mis.","quality":"Less is better","values":[0,0,0,0,0,0],"isMain":false},{"metricName":"# unaligned mis. contigs","quality":"Less is better","values":[0,1,0,1,0,0],"isMain":false}]],["Unaligned",[{"metricName":"# fully unaligned contigs","quality":"Less is better","values":[0,1,1,3,0,0],"isMain":false},{"metricName":"Fully unaligned length","quality":"Less is better","values":[0,580,544,18464,0,0],"isMain":false},{"metricName":"# partially unaligned contigs","quality":"Less is better","values":[0,0,0,3,1,1],"isMain":false},{"metricName":"Partially unaligned length","quality":"Less is better","values":[0,0,0,2571,1493,1025],"isMain":false}]],["Mismatches",[{"metricName":"# mismatches","quality":"Less is better","values":[60,80,145,147,130,75],"isMain":false},{"metricName":"# indels","quality":"Less is better","values":[2,2,0,0,0,1],"isMain":false},{"metricName":"Indels length","quality":"Less is better","values":[2,2,0,0,0,1],"isMain":false},{"metricName":"# mismatches per 100 kbp","quality":"Less is better","values":["2077.56","2359.88","2944.16","2976.91","3842.74","3355.70"],"isMain":true},{"metricName":"# indels per 100 kbp","quality":"Less is better","values":["69.25","59.00","0.00","0.00","0.00","44.74"],"isMain":true},{"metricName":" # indels (<= 5 bp)","quality":"Less is better","values":[2,2,0,0,0,1],"isMain":false},{"metricName":" # indels (> 5 bp)","quality":"Less is better","values":[0,0,0,0,0,0],"isMain":false},{"metricName":"# N's","quality":"Less is better","values":[0,0,0,0,0,0],"isMain":false},{"metricName":"# N's per 100 kbp","quality":"Less is better","values":["0.00","0.00","0.00","0.00","0.00","0.00"],"isMain":true}]],["Statistics without reference",[{"metricName":"# contigs","quality":"Equal","values":[6,8,10,16,6,5],"isMain":true},{"metricName":"# contigs (>= 1000 bp)","quality":"Equal","values":[0,0,1,2,1,1],"isMain":false},{"metricName":"# contigs (>= 5000 bp)","quality":"Equal","values":[0,0,0,1,0,0],"isMain":false},{"metricName":"# contigs (>= 10000 bp)","quality":"Equal","values":[0,0,0,1,0,0],"isMain":false},{"metricName":"# contigs (>= 25000 bp)","quality":"Equal","values":[0,0,0,0,0,0],"isMain":false},{"metricName":"# contigs (>= 50000 bp)","quality":"Equal","values":[0,0,0,0,0,0],"isMain":false},{"metricName":"Largest contig","quality":"More is better","values":[659,930,1030,17284,1814,1145],"isMain":true},{"metricName":"Total length","quality":"More is better","values":[3635,4900,6295,27422,5091,3748],"isMain":true},{"metricName":"Total length (>= 1000 bp)","quality":"More is better","values":[0,0,1030,18891,1814,1145],"isMain":true},{"metricName":"Total length (>= 5000 bp)","quality":"More is better","values":[0,0,0,17284,0,0],"isMain":false},{"metricName":"Total length (>= 10000 bp)","quality":"More is better","values":[0,0,0,17284,0,0],"isMain":true},{"metricName":"Total length (>= 25000 bp)","quality":"More is better","values":[0,0,0,0,0,0],"isMain":false},{"metricName":"Total length (>= 50000 bp)","quality":"More is better","values":[0,0,0,0,0,0],"isMain":true},{"metricName":"N50","quality":"More is better","values":[631,580,548,17284,977,989],"isMain":false},{"metricName":"N75","quality":"More is better","values":[577,516,539,657,602,544],"isMain":false},{"metricName":"L50","quality":"Less is better","values":[3,4,5,1,2,2],"isMain":false},{"metricName":"L75","quality":"Less is better","values":[5,6,7,5,4,4],"isMain":false},{"metricName":"GC (%)","quality":"Equal","values":["40.03","43.06","39.38","41.45","44.96","39.65"],"isMain":false}]],["K-mer-based statistics",[]],["Predicted genes",[]],["Similarity statistics",[]],["Reference statistics",[{"metricName":"Reference length","quality":"Equal","values":[2480269,2480269,2480269,2480269,2480269,2480269],"isMain":false},{"metricName":"Reference fragments","quality":"Equal","values":[2,2,2,2,2,2],"isMain":false},{"metricName":"Reference GC (%)","quality":"Equal","values":["37.50","37.50","37.50","37.50","37.50","37.50"],"isMain":false}]]],"subreferences":[],"subreports":[],"minContig":500},{"date":"10 February 2020, Monday, 18:51:32","assembliesNames":["megahit-103-df","megahit-113-df","megahit-113-ml","megahit-113-ms","megahit-129-df","metaSPAdes"],"referenceName":"GCF\_000163655.1\_ASM16365v1","order":[0,1,2,3,4,5],"report":[["Genome statistics",[{"metricName":"Genome fraction (%)","quality":"More is better","values":["4.680","5.538","5.112","5.558","5.443","3.619"],"isMain":true},{"metricName":"Duplication ratio","quality":"Less is better","values":["1.024","1.022","1.024","1.017","1.020","1.020"],"isMain":true},{"metricName":"Largest alignment","quality":"More is better","values":[2305,3211,1786,2259,2806,9760],"isMain":true},{"metricName":"Total aligned length","quality":"More is better","values":[245377,290743,269343,291738,285453,189602],"isMain":true},{"metricName":"NA50","quality":"More is better","values":[748,802,725,743,799,688],"isMain":false},{"metricName":"NA75","quality":"More is better","values":[599,610,584,595,612,582],"isMain":false},{"metricName":"NGA50","quality":"More is better","values":[null,null,null,null,null,null],"isMain":true},{"metricName":"LA50","quality":"Less is better","values":[123,132,144,149,135,95],"isMain":false},{"metricName":"LA75","quality":"Less is better","values":[217,243,251,261,241,173],"isMain":false}]],["Reads mapping",[]],["Misassemblies",[{"metricName":"# misassemblies","quality":"Less is better","values":[0,2,1,2,3,0],"isMain":true},{"metricName":" # relocations","quality":"Less is better","values":[0,0,0,1,0,0],"isMain":false},{"metricName":" # translocations","quality":"Less is better","values":[0,2,1,1,3,0],"isMain":false},{"metricName":" # inversions","quality":"Less is better","values":[0,0,0,0,0,0],"isMain":false},{"metricName":" # interspecies translocations","quality":"Less is better","values":[19,21,10,8,16,7],"isMain":false},{"metricName":"# misassembled contigs","quality":"Less is better","values":[0,2,1,2,3,0],"isMain":false},{"metricName":"Misassembled contigs length","quality":"Less is better","values":[0,2562,536,1542,1985,0],"isMain":true},{"metricName":"# possibly misassembled contigs","quality":"Less is better","values":[0,0,0,0,0,0],"isMain":false},{"metricName":" # possible misassemblies","quality":"Less is better","values":[0,0,0,0,0,0],"isMain":false},{"metricName":"# local misassemblies","quality":"Less is better","values":[0,1,0,0,0,0],"isMain":false},{"metricName":"# scaffold gap ext. mis.","quality":"Less is better","values":[0,0,0,0,0,0],"isMain":false},{"metricName":"# scaffold gap loc. mis.","quality":"Less is better","values":[0,0,0,0,0,0],"isMain":false},{"metricName":"# unaligned mis. contigs","quality":"Less is better","values":[0,0,2,1,1,0],"isMain":false}]],["Unaligned",[{"metricName":"# fully unaligned contigs","quality":"Less is better","values":[0,2,1,1,0,0],"isMain":false},{"metricName":"Fully unaligned length","quality":"Less is better","values":[0,1094,901,748,0,0],"isMain":false},{"metricName":"# partially unaligned contigs","quality":"Less is better","values":[3,7,1,1,6,2],"isMain":false},{"metricName":"Partially unaligned length","quality":"Less is better","values":[2460,9835,590,609,4810,2930],"isMain":false}]],["Mismatches",[{"metricName":"# mismatches","quality":"Less is better","values":[4070,4800,5798,6410,4840,3392],"isMain":false},{"metricName":"# indels","quality":"Less is better","values":[16,23,22,13,31,29],"isMain":false},{"metricName":"Indels length","quality":"Less is better","values":[16,30,113,138,73,59],"isMain":false},{"metricName":"# mismatches per 100 kbp","quality":"Less is better","values":["1663.50","1657.76","2169.63","2206.14","1700.75","1792.76"],"isMain":true},{"metricName":"# indels per 100 kbp","quality":"Less is better","values":["6.54","7.94","8.23","4.47","10.89","15.33"],"isMain":true},{"metricName":" # indels (<= 5 bp)","quality":"Less is better","values":[16,22,19,10,30,27],"isMain":false},{"metricName":" # indels (> 5 bp)","quality":"Less is better","values":[0,1,3,3,1,2],"isMain":false},{"metricName":"# N's","quality":"Less is better","values":[0,0,0,0,0,0],"isMain":false},{"metricName":"# N's per 100 kbp","quality":"Less is better","values":["0.00","0.00","0.00","0.00","0.00","0.00"],"isMain":true}]],["Statistics without reference",[{"metricName":"# contigs","quality":"Equal","values":[325,358,372,391,363,256],"isMain":true},{"metricName":"# contigs (>= 1000 bp)","quality":"Equal","values":[49,81,47,52,73,35],"isMain":false},{"metricName":"# contigs (>= 5000 bp)","quality":"Equal","values":[0,1,0,0,0,1],"isMain":false},{"metricName":"# contigs (>= 10000 bp)","quality":"Equal","values":[0,0,0,0,0,0],"isMain":false},{"metricName":"# contigs (>= 25000 bp)","quality":"Equal","values":[0,0,0,0,0,0],"isMain":false},{"metricName":"# contigs (>= 50000 bp)","quality":"Equal","values":[0,0,0,0,0,0],"isMain":false},{"metricName":"Largest contig","quality":"More is better","values":[2314,5318,1786,2289,2806,9775],"isMain":true},{"metricName":"Total length","quality":"More is better","values":[252945,306951,275223,296786,295188,195995],"isMain":true},{"metricName":"Total length (>= 1000 bp)","quality":"More is better","values":[64734,117791,57551,66715,95592,53295],"isMain":true},{"metricName":"Total length (>= 5000 bp)","quality":"More is better","values":[0,5318,0,0,0,9775],"isMain":false},{"metricName":"Total length (>= 10000 bp)","quality":"More is better","values":[0,0,0,0,0,0],"isMain":true},{"metricName":"Total length (>= 25000 bp)","quality":"More is better","values":[0,0,0,0,0,0],"isMain":false},{"metricName":"Total length (>= 50000 bp)","quality":"More is better","values":[0,0,0,0,0,0],"isMain":true},{"metricName":"N50","quality":"More is better","values":[771,849,733,752,837,709],"isMain":false},{"metricName":"N75","quality":"More is better","values":[619,648,598,615,643,609],"isMain":false},{"metricName":"L50","quality":"Less is better","values":[121,121,142,147,131,91],"isMain":false},{"metricName":"L75","quality":"Less is better","values":[212,225,247,256,232,167],"isMain":false},{"metricName":"GC (%)","quality":"Equal","values":["45.42","45.13","45.87","45.91","45.38","46.05"],"isMain":false}]],["K-mer-based statistics",[]],["Predicted genes",[]],["Similarity statistics",[]],["Reference statistics",[{"metricName":"Reference length","quality":"Equal","values":[5228026,5228026,5228026,5228026,5228026,5228026],"isMain":false},{"metricName":"Reference fragments","quality":"Equal","values":[86,86,86,86,86,86],"isMain":false},{"metricName":"Reference GC (%)","quality":"Equal","values":["44.99","44.99","44.99","44.99","44.99","44.99"],"isMain":false}]]],"subreferences":[],"subreports":[],"minContig":500},{"date":"10 February 2020, Monday, 18:51:16","assembliesNames":["megahit-103-df","megahit-113-df","megahit-113-ml","megahit-113-ms","megahit-129-df","metaSPAdes"],"referenceName":"GCF\_000165775.1\_ASM16577v3","order":[0,1,2,3,4,5],"report":[["Genome statistics",[{"metricName":"Genome fraction (%)","quality":"More is better","values":["0.207","7.818","7.810","10.225","10.993","2.798"],"isMain":true},{"metricName":"Duplication ratio","quality":"Less is better","values":["1.332","1.160","1.287","1.319","1.148","1.005"],"isMain":true},{"metricName":"Largest alignment","quality":"More is better","values":[598,13184,20108,15913,19281,37160],"isMain":true},{"metricName":"Total aligned length","quality":"More is better","values":[4516,178135,184744,253336,249005,59592],"isMain":true},{"metricName":"NA50","quality":"More is better","values":[182,695,175,null,545,37160],"isMain":false},{"metricName":"NA75","quality":"More is better","values":[null,196,null,null,null,18371],"isMain":false},{"metricName":"NGA50","quality":"More is better","values":[null,null,null,null,null,null],"isMain":true},{"metricName":"LA50","quality":"Less is better","values":[10,43,239,null,81,1],"isMain":false},{"metricName":"LA75","quality":"Less is better","values":[null,165,null,null,null,2],"isMain":false}]],["Reads mapping",[]],["Misassemblies",[{"metricName":"# misassemblies","quality":"Less is better","values":[1,26,45,63,54,1],"isMain":true},{"metricName":" # relocations","quality":"Less is better","values":[0,17,30,39,34,1],"isMain":false},{"metricName":" # translocations","quality":"Less is better","values":[0,8,14,21,16,0],"isMain":false},{"metricName":" # inversions","quality":"Less is better","values":[1,1,1,3,4,0],"isMain":false},{"metricName":" # interspecies translocations","quality":"Less is better","values":[6,59,107,132,89,1],"isMain":false},{"metricName":"# misassembled contigs","quality":"Less is better","values":[1,25,40,57,44,1],"isMain":false},{"metricName":"Misassembled contigs length","quality":"Less is better","values":[526,26081,35200,64690,68758,1105],"isMain":true},{"metricName":"# possibly misassembled contigs","quality":"Less is better","values":[0,0,2,3,1,0],"isMain":false},{"metricName":" # possible misassemblies","quality":"Less is better","values":[0,0,2,3,1,0],"isMain":false},{"metricName":"# local misassemblies","quality":"Less is better","values":[0,7,5,17,9,2],"isMain":false},{"metricName":"# scaffold gap ext. mis.","quality":"Less is better","values":[0,0,0,0,0,0],"isMain":false},{"metricName":"# scaffold gap loc. mis.","quality":"Less is better","values":[0,0,0,0,0,0],"isMain":false},{"metricName":"# unaligned mis. contigs","quality":"Less is better","values":[0,8,16,23,11,0],"isMain":false}]],["Unaligned",[{"metricName":"# fully unaligned contigs","quality":"Less is better","values":[0,3,9,5,4,0],"isMain":false},{"metricName":"Fully unaligned length","quality":"Less is better","values":[0,1859,5792,3227,2260,0],"isMain":false},{"metricName":"# partially unaligned contigs","quality":"Less is better","values":[2,16,24,35,18,0],"isMain":false},{"metricName":"Partially unaligned length","quality":"Less is better","values":[2112,34520,134761,286764,111032,0],"isMain":false}]],["Mismatches",[{"metricName":"# mismatches","quality":"Less is better","values":[121,3481,4474,6605,4739,513],"isMain":false},{"metricName":"# indels","quality":"Less is better","values":[0,116,127,194,176,32],"isMain":false},{"metricName":"Indels length","quality":"Less is better","values":[0,459,410,654,654,80],"isMain":false},{"metricName":"# mismatches per 100 kbp","quality":"Less is better","values":["2743.76","2091.21","2690.61","3033.87","2024.62","861.24"],"isMain":true},{"metricName":"# indels per 100 kbp","quality":"Less is better","values":["0.00","69.69","76.38","89.11","75.19","53.72"],"isMain":true},{"metricName":" # indels (<= 5 bp)","quality":"Less is better","values":[0,93,114,168,151,26],"isMain":false},{"metricName":" # indels (> 5 bp)","quality":"Less is better","values":[0,23,13,26,25,6],"isMain":false},{"metricName":"# N's","quality":"Less is better","values":[0,0,0,0,0,0],"isMain":false},{"metricName":"# N's per 100 kbp","quality":"Less is better","values":["0.00","0.00","0.00","0.00","0.00","0.00"],"isMain":true}]],["Statistics without reference",[{"metricName":"# contigs","quality":"Equal","values":[12,177,247,333,205,6],"isMain":true},{"metricName":"# contigs (>= 1000 bp)","quality":"Equal","values":[1,32,27,45,42,5],"isMain":false},{"metricName":"# contigs (>= 5000 bp)","quality":"Equal","values":[0,8,7,12,17,2],"isMain":false},{"metricName":"# contigs (>= 10000 bp)","quality":"Equal","values":[0,2,4,8,7,2],"isMain":false},{"metricName":"# contigs (>= 25000 bp)","quality":"Equal","values":[0,0,3,4,1,1],"isMain":false},{"metricName":"# contigs (>= 50000 bp)","quality":"Equal","values":[0,0,1,1,1,0],"isMain":false},{"metricName":"Largest contig","quality":"More is better","values":[1559,20438,53474,121821,63870,37160],"isMain":true},{"metricName":"Total length","quality":"More is better","values":[7987,229442,354618,577057,382039,59873],"isMain":true},{"metricName":"Total length (>= 1000 bp)","quality":"More is better","values":[1559,140187,215363,391638,285145,59335],"isMain":true},{"metricName":"Total length (>= 5000 bp)","quality":"More is better","values":[0,77156,175124,327450,227173,55542],"isMain":false},{"metricName":"Total length (>= 10000 bp)","quality":"More is better","values":[0,33709,155452,298242,152046,55542],"isMain":true},{"metricName":"Total length (>= 25000 bp)","quality":"More is better","values":[0,0,135088,240584,63870,37160],"isMain":false},{"metricName":"Total length (>= 50000 bp)","quality":"More is better","values":[0,0,53474,121821,63870,0],"isMain":true},{"metricName":"N50","quality":"More is better","values":[598,2787,3287,10093,7722,37160],"isMain":false},{"metricName":"N75","quality":"More is better","values":[526,654,669,760,975,18382],"isMain":false},{"metricName":"L50","quality":"Less is better","values":[5,19,8,8,12,1],"isMain":false},{"metricName":"L75","quality":"Less is better","values":[9,75,92,95,44,2],"isMain":false},{"metricName":"GC (%)","quality":"Equal","values":["39.10","38.06","38.49","39.49","37.58","36.56"],"isMain":false}]],["K-mer-based statistics",[]],["Predicted genes",[]],["Similarity statistics",[]],["Reference statistics",[{"metricName":"Reference length","quality":"Equal","values":[2129203,2129203,2129203,2129203,2129203,2129203],"isMain":false},{"metricName":"Reference fragments","quality":"Equal","values":[4,4,4,4,4,4],"isMain":false},{"metricName":"Reference GC (%)","quality":"Equal","values":["36.80","36.80","36.80","36.80","36.80","36.80"],"isMain":false}]]],"subreferences":[],"subreports":[],"minContig":500},{"date":"10 February 2020, Monday, 18:51:29","assembliesNames":["megahit-103-df","megahit-113-df","megahit-113-ml","megahit-113-ms","megahit-129-df","metaSPAdes"],"referenceName":"GCF\_000177015.3\_Rumi\_bact\_D16\_V2","order":[0,1,2,3,4,5],"report":[["Genome statistics",[{"metricName":"Genome fraction (%)","quality":"More is better","values":["0.069","0.449","0.509","0.293","0.455","0.367"],"isMain":true},{"metricName":"Duplication ratio","quality":"Less is better","values":["1.116","1.132","1.086","1.309","1.157","1.078"],"isMain":true},{"metricName":"Largest alignment","quality":"More is better","values":[596,2586,2931,1884,2649,3984],"isMain":true},{"metricName":"Total aligned length","quality":"More is better","values":[2220,14365,16279,10417,14560,11734],"isMain":true},{"metricName":"NA50","quality":"More is better","values":[505,586,1135,517,521,2561],"isMain":false},{"metricName":"NA75","quality":"More is better","values":[null,null,486,null,null,null],"isMain":false},{"metricName":"NGA50","quality":"More is better","values":[null,null,null,null,null,null],"isMain":true},{"metricName":"LA50","quality":"Less is better","values":[3,10,5,9,10,3],"isMain":false},{"metricName":"LA75","quality":"Less is better","values":[null,null,13,null,null,null],"isMain":false}]],["Reads mapping",[]],["Misassemblies",[{"metricName":"# misassemblies","quality":"Less is better","values":[0,0,0,0,0,0],"isMain":true},{"metricName":" # relocations","quality":"Less is better","values":[0,0,0,0,0,0],"isMain":false},{"metricName":" # translocations","quality":"Less is better","values":[0,0,0,0,0,0],"isMain":false},{"metricName":" # inversions","quality":"Less is better","values":[0,0,0,0,0,0],"isMain":false},{"metricName":" # interspecies translocations","quality":"Less is better","values":[2,9,5,8,10,5],"isMain":false},{"metricName":"# misassembled contigs","quality":"Less is better","values":[0,0,0,0,0,0],"isMain":false},{"metricName":"Misassembled contigs length","quality":"Less is better","values":[0,0,0,0,0,0],"isMain":true},{"metricName":"# possibly misassembled contigs","quality":"Less is better","values":[0,0,0,0,0,0],"isMain":false},{"metricName":" # possible misassemblies","quality":"Less is better","values":[0,0,0,0,0,0],"isMain":false},{"metricName":"# local misassemblies","quality":"Less is better","values":[0,0,0,1,0,0],"isMain":false},{"metricName":"# scaffold gap ext. mis.","quality":"Less is better","values":[0,0,0,0,0,0],"isMain":false},{"metricName":"# scaffold gap loc. mis.","quality":"Less is better","values":[0,0,0,0,0,0],"isMain":false},{"metricName":"# unaligned mis. contigs","quality":"Less is better","values":[0,1,0,0,0,0],"isMain":false}]],["Unaligned",[{"metricName":"# fully unaligned contigs","quality":"Less is better","values":[0,1,2,1,1,0],"isMain":false},{"metricName":"Fully unaligned length","quality":"Less is better","values":[0,514,1064,1373,871,0],"isMain":false},{"metricName":"# partially unaligned contigs","quality":"Less is better","values":[1,3,1,2,3,3],"isMain":false},{"metricName":"Partially unaligned length","quality":"Less is better","values":[633,5407,1374,1214,4253,4119],"isMain":false}]],["Mismatches",[{"metricName":"# mismatches","quality":"Less is better","values":[49,355,434,240,388,153],"isMain":false},{"metricName":"# indels","quality":"Less is better","values":[1,5,5,2,4,1],"isMain":false},{"metricName":"Indels length","quality":"Less is better","values":[1,6,6,3,4,1],"isMain":false},{"metricName":"# mismatches per 100 kbp","quality":"Less is better","values":["2208.20","2471.63","2666.34","2564.65","2664.47","1303.79"],"isMain":true},{"metricName":"# indels per 100 kbp","quality":"Less is better","values":["45.07","34.81","30.72","21.37","27.47","8.52"],"isMain":true},{"metricName":" # indels (<= 5 bp)","quality":"Less is better","values":[1,5,5,2,4,1],"isMain":false},{"metricName":" # indels (> 5 bp)","quality":"Less is better","values":[0,0,0,0,0,0],"isMain":false},{"metricName":"# N's","quality":"Less is better","values":[0,0,0,0,0,0],"isMain":false},{"metricName":"# N's per 100 kbp","quality":"Less is better","values":["0.00","0.00","0.00","0.00","0.00","0.00"],"isMain":true}]],["Statistics without reference",[{"metricName":"# contigs","quality":"Equal","values":[5,20,19,19,21,8],"isMain":true},{"metricName":"# contigs (>= 1000 bp)","quality":"Equal","values":[0,6,7,3,4,4],"isMain":false},{"metricName":"# contigs (>= 5000 bp)","quality":"Equal","values":[0,0,0,0,1,1],"isMain":false},{"metricName":"# contigs (>= 10000 bp)","quality":"Equal","values":[0,0,0,0,0,0],"isMain":false},{"metricName":"# contigs (>= 25000 bp)","quality":"Equal","values":[0,0,0,0,0,0],"isMain":false},{"metricName":"# contigs (>= 50000 bp)","quality":"Equal","values":[0,0,0,0,0,0],"isMain":false},{"metricName":"Largest contig","quality":"More is better","values":[881,4020,3043,1936,5001,6618],"isMain":true},{"metricName":"Total length","quality":"More is better","values":[3109,22174,20115,14839,21977,16764],"isMain":true},{"metricName":"Total length (>= 1000 bp)","quality":"More is better","values":[0,13458,13367,5193,10971,14227],"isMain":true},{"metricName":"Total length (>= 5000 bp)","quality":"More is better","values":[0,0,0,0,5001,6618],"isMain":false},{"metricName":"Total length (>= 10000 bp)","quality":"More is better","values":[0,0,0,0,0,0],"isMain":true},{"metricName":"Total length (>= 25000 bp)","quality":"More is better","values":[0,0,0,0,0,0],"isMain":false},{"metricName":"Total length (>= 50000 bp)","quality":"More is better","values":[0,0,0,0,0,0],"isMain":true},{"metricName":"N50","quality":"More is better","values":[596,1940,1254,716,880,3538],"isMain":false},{"metricName":"N75","quality":"More is better","values":[516,652,577,544,651,3003],"isMain":false},{"metricName":"L50","quality":"Less is better","values":[3,4,5,6,5,2],"isMain":false},{"metricName":"L75","quality":"Less is better","values":[4,11,10,12,11,3],"isMain":false},{"metricName":"GC (%)","quality":"Equal","values":["54.42","49.13","50.58","54.94","52.98","48.78"],"isMain":false}]],["K-mer-based statistics",[]],["Predicted genes",[]],["Similarity statistics",[]],["Reference statistics",[{"metricName":"Reference length","quality":"Equal","values":[3197155,3197155,3197155,3197155,3197155,3197155],"isMain":false},{"metricName":"Reference fragments","quality":"Equal","values":[8,8,8,8,8,8],"isMain":false},{"metricName":"Reference GC (%)","quality":"Equal","values":["56.80","56.80","56.80","56.80","56.80","56.80"],"isMain":false}]]],"subreferences":[],"subreports":[],"minContig":500},{"date":"10 February 2020, Monday, 18:51:30","assembliesNames":["megahit-103-df","megahit-113-df","megahit-113-ml","megahit-113-ms","megahit-129-df","metaSPAdes"],"referenceName":"GCF\_000178115.2\_ASM17811v2","order":[0,1,2,3,4,5],"report":[["Genome statistics",[{"metricName":"Genome fraction (%)","quality":"More is better","values":["0.815","0.892","1.222","1.602","0.963","0.619"],"isMain":true},{"metricName":"Duplication ratio","quality":"Less is better","values":["1.124","1.095","1.015","1.063","1.129","1.026"],"isMain":true},{"metricName":"Largest alignment","quality":"More is better","values":[1658,1997,1517,1607,1666,1602],"isMain":true},{"metricName":"Total aligned length","quality":"More is better","values":[24530,26984,36765,48428,28973,18620],"isMain":true},{"metricName":"NA50","quality":"More is better","values":[528,535,567,547,533,566],"isMain":false},{"metricName":"NA75","quality":"More is better","values":[388,355,523,244,287,513],"isMain":false},{"metricName":"NGA50","quality":"More is better","values":[null,null,null,null,null,null],"isMain":true},{"metricName":"LA50","quality":"Less is better","values":[22,23,29,48,26,14],"isMain":false},{"metricName":"LA75","quality":"Less is better","values":[36,40,47,81,46,24],"isMain":false}]],["Reads mapping",[]],["Misassemblies",[{"metricName":"# misassemblies","quality":"Less is better","values":[0,0,0,0,0,0],"isMain":true},{"metricName":" # relocations","quality":"Less is better","values":[0,0,0,0,0,0],"isMain":false},{"metricName":" # translocations","quality":"Less is better","values":[0,0,0,0,0,0],"isMain":false},{"metricName":" # inversions","quality":"Less is better","values":[0,0,0,0,0,0],"isMain":false},{"metricName":" # interspecies translocations","quality":"Less is better","values":[13,12,4,18,16,3],"isMain":false},{"metricName":"# misassembled contigs","quality":"Less is better","values":[0,0,0,0,0,0],"isMain":false},{"metricName":"Misassembled contigs length","quality":"Less is better","values":[0,0,0,0,0,0],"isMain":true},{"metricName":"# possibly misassembled contigs","quality":"Less is better","values":[0,0,1,0,0,0],"isMain":false},{"metricName":" # possible misassemblies","quality":"Less is better","values":[0,0,1,0,0,0],"isMain":false},{"metricName":"# local misassemblies","quality":"Less is better","values":[0,1,0,0,0,0],"isMain":false},{"metricName":"# scaffold gap ext. mis.","quality":"Less is better","values":[0,0,0,0,0,0],"isMain":false},{"metricName":"# scaffold gap loc. mis.","quality":"Less is better","values":[0,0,0,0,0,0],"isMain":false},{"metricName":"# unaligned mis. contigs","quality":"Less is better","values":[0,0,0,0,0,0],"isMain":false}]],["Unaligned",[{"metricName":"# fully unaligned contigs","quality":"Less is better","values":[0,0,0,0,0,0],"isMain":false},{"metricName":"Fully unaligned length","quality":"Less is better","values":[0,0,0,0,0,0],"isMain":false},{"metricName":"# partially unaligned contigs","quality":"Less is better","values":[2,5,4,10,5,2],"isMain":false},{"metricName":"Partially unaligned length","quality":"Less is better","values":[1790,3893,2708,11029,3633,1382],"isMain":false}]],["Mismatches",[{"metricName":"# mismatches","quality":"Less is better","values":[367,418,697,1055,473,311],"isMain":false},{"metricName":"# indels","quality":"Less is better","values":[2,1,3,6,2,0],"isMain":false},{"metricName":"Indels length","quality":"Less is better","values":[2,1,3,7,2,0],"isMain":false},{"metricName":"# mismatches per 100 kbp","quality":"Less is better","values":["1496.13","1557.14","1896.19","2188.62","1632.55","1670.25"],"isMain":true},{"metricName":"# indels per 100 kbp","quality":"Less is better","values":["8.15","3.73","8.16","12.45","6.90","0.00"],"isMain":true},{"metricName":" # indels (<= 5 bp)","quality":"Less is better","values":[2,1,3,6,2,0],"isMain":false},{"metricName":" # indels (> 5 bp)","quality":"Less is better","values":[0,0,0,0,0,0],"isMain":false},{"metricName":"# N's","quality":"Less is better","values":[0,0,0,0,0,0],"isMain":false},{"metricName":"# N's per 100 kbp","quality":"Less is better","values":["0.00","0.00","0.00","0.00","0.00","0.00"],"isMain":true}]],["Statistics without reference",[{"metricName":"# contigs","quality":"Equal","values":[46,49,63,92,55,31],"isMain":true},{"metricName":"# contigs (>= 1000 bp)","quality":"Equal","values":[5,5,4,8,6,3],"isMain":false},{"metricName":"# contigs (>= 5000 bp)","quality":"Equal","values":[0,0,0,0,0,0],"isMain":false},{"metricName":"# contigs (>= 10000 bp)","quality":"Equal","values":[0,0,0,0,0,0],"isMain":false},{"metricName":"# contigs (>= 25000 bp)","quality":"Equal","values":[0,0,0,0,0,0],"isMain":false},{"metricName":"# contigs (>= 50000 bp)","quality":"Equal","values":[0,0,0,0,0,0],"isMain":false},{"metricName":"Largest contig","quality":"More is better","values":[1658,2013,1520,2918,1666,1602],"isMain":true},{"metricName":"Total length","quality":"More is better","values":[29356,33295,40013,62249,36331,20481],"isMain":true},{"metricName":"Total length (>= 1000 bp)","quality":"More is better","values":[6225,6774,5033,12228,7155,4010],"isMain":true},{"metricName":"Total length (>= 5000 bp)","quality":"More is better","values":[0,0,0,0,0,0],"isMain":false},{"metricName":"Total length (>= 10000 bp)","quality":"More is better","values":[0,0,0,0,0,0],"isMain":true},{"metricName":"Total length (>= 25000 bp)","quality":"More is better","values":[0,0,0,0,0,0],"isMain":false},{"metricName":"Total length (>= 50000 bp)","quality":"More is better","values":[0,0,0,0,0,0],"isMain":true},{"metricName":"N50","quality":"More is better","values":[566,614,598,597,586,609],"isMain":false},{"metricName":"N75","quality":"More is better","values":[527,543,546,548,543,538],"isMain":false},{"metricName":"L50","quality":"Less is better","values":[19,19,27,35,22,13],"isMain":false},{"metricName":"L75","quality":"Less is better","values":[32,34,44,62,38,22],"isMain":false},{"metricName":"GC (%)","quality":"Equal","values":["52.20","53.32","54.60","52.39","54.20","53.88"],"isMain":false}]],["K-mer-based statistics",[]],["Predicted genes",[]],["Similarity statistics",[]],["Reference statistics",[{"metricName":"Reference length","quality":"Equal","values":[3008576,3008576,3008576,3008576,3008576,3008576],"isMain":false},{"metricName":"Reference fragments","quality":"Equal","values":[1,1,1,1,1,1],"isMain":false},{"metricName":"Reference GC (%)","quality":"Equal","values":["55.56","55.56","55.56","55.56","55.56","55.56"],"isMain":false}]]],"subreferences":[],"subreports":[],"minContig":500},{"date":"10 February 2020, Monday, 18:51:25","assembliesNames":["megahit-103-df","megahit-113-df","megahit-113-ml","megahit-113-ms","megahit-129-df","metaSPAdes"],"referenceName":"GCF\_000178835.2\_ASM17883v2","order":[0,1,2,3,4,5],"report":[["Genome statistics",[{"metricName":"Genome fraction (%)","quality":"More is better","values":["0.008","0.034","0.024","0.053","0.037","0.003"],"isMain":true},{"metricName":"Duplication ratio","quality":"Less is better","values":["1.403","1.021","1.382","1.629","1.497","3.725"],"isMain":true},{"metricName":"Largest alignment","quality":"More is better","values":[377,719,544,586,651,153],"isMain":true},{"metricName":"Total aligned length","quality":"More is better","values":[377,1600,1140,2505,1759,153],"isMain":true},{"metricName":"NA50","quality":"More is better","values":[377,586,502,null,129,null],"isMain":false},{"metricName":"NGA50","quality":"More is better","values":[null,null,null,null,null,null],"isMain":true},{"metricName":"LA50","quality":"Less is better","values":[1,2,2,null,5,null],"isMain":false}]],["Reads mapping",[]],["Misassemblies",[{"metricName":"# misassemblies","quality":"Less is better","values":[0,0,0,0,0,0],"isMain":true},{"metricName":" # relocations","quality":"Less is better","values":[0,0,0,0,0,0],"isMain":false},{"metricName":" # translocations","quality":"Less is better","values":[0,0,0,0,0,0],"isMain":false},{"metricName":" # inversions","quality":"Less is better","values":[0,0,0,0,0,0],"isMain":false},{"metricName":" # interspecies translocations","quality":"Less is better","values":[1,1,2,7,4,1],"isMain":false},{"metricName":"# misassembled contigs","quality":"Less is better","values":[0,0,0,0,0,0],"isMain":false},{"metricName":"Misassembled contigs length","quality":"Less is better","values":[0,0,0,0,0,0],"isMain":true},{"metricName":"# possibly misassembled contigs","quality":"Less is better","values":[0,0,0,0,0,0],"isMain":false},{"metricName":" # possible misassemblies","quality":"Less is better","values":[0,0,0,0,0,0],"isMain":false},{"metricName":"# local misassemblies","quality":"Less is better","values":[0,0,0,0,0,0],"isMain":false},{"metricName":"# scaffold gap ext. mis.","quality":"Less is better","values":[0,0,0,0,0,0],"isMain":false},{"metricName":"# scaffold gap loc. mis.","quality":"Less is better","values":[0,0,0,0,0,0],"isMain":false},{"metricName":"# unaligned mis. contigs","quality":"Less is better","values":[0,0,0,0,0,0],"isMain":false}]],["Unaligned",[{"metricName":"# fully unaligned contigs","quality":"Less is better","values":[0,0,0,0,0,0],"isMain":false},{"metricName":"Fully unaligned length","quality":"Less is better","values":[0,0,0,0,0,0],"isMain":false},{"metricName":"# partially unaligned contigs","quality":"Less is better","values":[0,1,0,3,1,0],"isMain":false},{"metricName":"Partially unaligned length","quality":"Less is better","values":[0,654,0,3152,634,0],"isMain":false}]],["Mismatches",[{"metricName":"# mismatches","quality":"Less is better","values":[7,44,30,66,34,2],"isMain":false},{"metricName":"# indels","quality":"Less is better","values":[0,1,0,2,1,0],"isMain":false},{"metricName":"Indels length","quality":"Less is better","values":[0,1,0,2,1,0],"isMain":false},{"metricName":"# mismatches per 100 kbp","quality":"Less is better","values":["1856.76","2748.28","2631.58","2632.63","1931.82","1307.19"],"isMain":true},{"metricName":"# indels per 100 kbp","quality":"Less is better","values":["0.00","62.46","0.00","79.78","56.82","0.00"],"isMain":true},{"metricName":" # indels (<= 5 bp)","quality":"Less is better","values":[0,1,0,2,1,0],"isMain":false},{"metricName":" # indels (> 5 bp)","quality":"Less is better","values":[0,0,0,0,0,0],"isMain":false},{"metricName":"# N's","quality":"Less is better","values":[0,0,0,0,0,0],"isMain":false},{"metricName":"# N's per 100 kbp","quality":"Less is better","values":["0.00","0.00","0.00","0.00","0.00","0.00"],"isMain":true}]],["Statistics without reference",[{"metricName":"# contigs","quality":"Equal","values":[1,3,3,9,5,1],"isMain":true},{"metricName":"# contigs (>= 1000 bp)","quality":"Equal","values":[0,0,0,1,0,0],"isMain":false},{"metricName":"# contigs (>= 5000 bp)","quality":"Equal","values":[0,0,0,0,0,0],"isMain":false},{"metricName":"# contigs (>= 10000 bp)","quality":"Equal","values":[0,0,0,0,0,0],"isMain":false},{"metricName":"# contigs (>= 25000 bp)","quality":"Equal","values":[0,0,0,0,0,0],"isMain":false},{"metricName":"# contigs (>= 50000 bp)","quality":"Equal","values":[0,0,0,0,0,0],"isMain":false},{"metricName":"Largest contig","quality":"More is better","values":[529,949,544,2046,763,570],"isMain":true},{"metricName":"Total length","quality":"More is better","values":[529,2288,1575,7237,3269,570],"isMain":true},{"metricName":"Total length (>= 1000 bp)","quality":"More is better","values":[0,0,0,2046,0,0],"isMain":true},{"metricName":"Total length (>= 5000 bp)","quality":"More is better","values":[0,0,0,0,0,0],"isMain":false},{"metricName":"Total length (>= 10000 bp)","quality":"More is better","values":[0,0,0,0,0,0],"isMain":true},{"metricName":"Total length (>= 25000 bp)","quality":"More is better","values":[0,0,0,0,0,0],"isMain":false},{"metricName":"Total length (>= 50000 bp)","quality":"More is better","values":[0,0,0,0,0,0],"isMain":true},{"metricName":"N50","quality":"More is better","values":[529,753,524,860,723,570],"isMain":false},{"metricName":"N75","quality":"More is better","values":[529,586,507,586,548,570],"isMain":false},{"metricName":"L50","quality":"Less is better","values":[1,2,2,3,3,1],"isMain":false},{"metricName":"L75","quality":"Less is better","values":[1,3,3,6,4,1],"isMain":false},{"metricName":"GC (%)","quality":"Equal","values":["38.00","38.55","41.02","41.30","40.65","35.44"],"isMain":false}]],["K-mer-based statistics",[]],["Predicted genes",[]],["Similarity statistics",[]],["Reference statistics",[{"metricName":"Reference length","quality":"Equal","values":[4714237,4714237,4714237,4714237,4714237,4714237],"isMain":false},{"metricName":"Reference fragments","quality":"Equal","values":[1,1,1,1,1,1],"isMain":false},{"metricName":"Reference GC (%)","quality":"Equal","values":["34.32","34.32","34.32","34.32","34.32","34.32"],"isMain":false}]]],"subreferences":[],"subreports":[],"minContig":500},{"date":"10 February 2020, Monday, 18:51:53","assembliesNames":["megahit-103-df","megahit-113-df","megahit-113-ml","megahit-113-ms","megahit-129-df","metaSPAdes"],"referenceName":"GCF\_000184945.1\_ASM18494v1","order":[0,1,2,3,4,5],"report":[["Genome statistics",[{"metricName":"Genome fraction (%)","quality":"More is better","values":["95.286","98.718","98.418","98.503","97.885","98.294"],"isMain":true},{"metricName":"Duplication ratio","quality":"Less is better","values":["1.018","1.010","1.010","1.022","1.015","1.002"],"isMain":true},{"metricName":"Largest alignment","quality":"More is better","values":[29305,453589,453523,274963,453533,374881],"isMain":true},{"metricName":"Total aligned length","quality":"More is better","values":[3118731,3208598,3197977,3235054,3193601,3170702],"isMain":true},{"metricName":"NG50","quality":"More is better","values":[5794,160867,148214,122122,106474,103571],"isMain":false},{"metricName":"NG75","quality":"More is better","values":[3307,90263,90288,89334,44558,70236],"isMain":false},{"metricName":"NA50","quality":"More is better","values":[5960,90263,98092,48569,84766,89588],"isMain":false},{"metricName":"NA75","quality":"More is better","values":[3593,33477,41763,2928,39074,54367],"isMain":false},{"metricName":"NGA50","quality":"More is better","values":[5778,92449,105684,68247,84766,89588],"isMain":true},{"metricName":"NGA75","quality":"More is better","values":[3304,42875,42875,32996,40622,54367],"isMain":false},{"metricName":"LG50","quality":"Less is better","values":[162,7,7,9,8,9],"isMain":false},{"metricName":"LG75","quality":"Less is better","values":[339,14,14,17,19,18],"isMain":false},{"metricName":"LA50","quality":"Less is better","values":[157,12,11,23,12,12],"isMain":false},{"metricName":"LA75","quality":"Less is better","values":[324,27,23,80,27,23],"isMain":false},{"metricName":"LGA50","quality":"Less is better","values":[163,11,10,16,12,12],"isMain":true},{"metricName":"LGA75","quality":"Less is better","values":[341,22,21,32,26,23],"isMain":false}]],["Reads mapping",[]],["Misassemblies",[{"metricName":"# misassemblies","quality":"Less is better","values":[3,22,15,20,24,16],"isMain":true},{"metricName":" # relocations","quality":"Less is better","values":[0,0,0,4,1,0],"isMain":false},{"metricName":" # translocations","quality":"Less is better","values":[2,21,15,16,23,16],"isMain":false},{"metricName":" # inversions","quality":"Less is better","values":[1,1,0,0,0,0],"isMain":false},{"metricName":" # interspecies translocations","quality":"Less is better","values":[36,28,23,68,41,8],"isMain":false},{"metricName":"# misassembled contigs","quality":"Less is better","values":[3,12,11,13,17,13],"isMain":false},{"metricName":"Misassembled contigs length","quality":"Less is better","values":[24505,1632110,1461763,907991,1403336,1369569],"isMain":true},{"metricName":"# possibly misassembled contigs","quality":"Less is better","values":[0,0,0,0,0,0],"isMain":false},{"metricName":" # possible misassemblies","quality":"Less is better","values":[0,0,0,0,0,0],"isMain":false},{"metricName":"# local misassemblies","quality":"Less is better","values":[0,3,3,0,4,2],"isMain":false},{"metricName":"# scaffold gap ext. mis.","quality":"Less is better","values":[0,0,0,0,0,0],"isMain":false},{"metricName":"# scaffold gap loc. mis.","quality":"Less is better","values":[0,0,0,0,0,0],"isMain":false},{"metricName":"# unaligned mis. contigs","quality":"Less is better","values":[0,0,0,1,1,0],"isMain":false}]],["Unaligned",[{"metricName":"# fully unaligned contigs","quality":"Less is better","values":[1,1,2,3,0,0],"isMain":false},{"metricName":"Fully unaligned length","quality":"Less is better","values":[640,603,9635,1738,0,0],"isMain":false},{"metricName":"# partially unaligned contigs","quality":"Less is better","values":[17,17,11,36,28,5],"isMain":false},{"metricName":"Partially unaligned length","quality":"Less is better","values":[14687,230724,138245,879739,56461,59854],"isMain":false}]],["Mismatches",[{"metricName":"# mismatches","quality":"Less is better","values":[4995,1998,2452,4717,2374,1094],"isMain":false},{"metricName":"# indels","quality":"Less is better","values":[6,51,42,120,26,48],"isMain":false},{"metricName":"Indels length","quality":"Less is better","values":[6,223,560,934,420,137],"isMain":false},{"metricName":"# mismatches per 100 kbp","quality":"Less is better","values":["162.66","62.80","77.31","148.59","75.25","34.53"],"isMain":true},{"metricName":"# indels per 100 kbp","quality":"Less is better","values":["0.20","1.60","1.32","3.78","0.82","1.52"],"isMain":true},{"metricName":" # indels (<= 5 bp)","quality":"Less is better","values":[6,47,35,106,19,45],"isMain":false},{"metricName":" # indels (> 5 bp)","quality":"Less is better","values":[0,4,7,14,7,3],"isMain":false},{"metricName":"# N's","quality":"Less is better","values":[0,0,0,0,0,0],"isMain":false},{"metricName":"# N's per 100 kbp","quality":"Less is better","values":["0.00","0.00","0.00","0.00","0.00","0.00"],"isMain":true}]],["Statistics without reference",[{"metricName":"# contigs","quality":"Equal","values":[824,137,145,211,176,103],"isMain":true},{"metricName":"# contigs (>= 1000 bp)","quality":"Equal","values":[627,88,90,117,106,90],"isMain":false},{"metricName":"# contigs (>= 5000 bp)","quality":"Equal","values":[217,49,49,61,58,49],"isMain":false},{"metricName":"# contigs (>= 10000 bp)","quality":"Equal","values":[63,39,35,56,44,40],"isMain":false},{"metricName":"# contigs (>= 25000 bp)","quality":"Equal","values":[1,26,25,39,25,26],"isMain":false},{"metricName":"# contigs (>= 50000 bp)","quality":"Equal","values":[0,19,17,30,18,21],"isMain":false},{"metricName":"Largest contig","quality":"More is better","values":[29305,521349,630620,295363,698428,436970],"isMain":true},{"metricName":"Total length","quality":"More is better","values":[3142106,3445248,3353007,4127427,3257217,3233293],"isMain":true},{"metricName":"Total length (>= 1000 bp)","quality":"More is better","values":[3004952,3413455,3315878,4066996,3212340,3223592],"isMain":true},{"metricName":"Total length (>= 5000 bp)","quality":"More is better","values":[1910422,3327204,3218574,3956071,3098683,3132798],"isMain":false},{"metricName":"Total length (>= 10000 bp)","quality":"More is better","values":[871035,3250790,3111402,3913098,2997228,3065448],"isMain":true},{"metricName":"Total length (>= 25000 bp)","quality":"More is better","values":[29305,3033456,2937350,3609012,2669622,2831181],"isMain":false},{"metricName":"Total length (>= 50000 bp)","quality":"More is better","values":[0,2815206,2633289,3288476,2413381,2641888],"isMain":true},{"metricName":"N50","quality":"More is better","values":[6075,151768,148214,95529,106474,103571],"isMain":false},{"metricName":"N75","quality":"More is better","values":[3656,72543,74460,57893,44558,70236],"isMain":false},{"metricName":"L50","quality":"Less is better","values":[155,8,7,13,8,9],"isMain":false},{"metricName":"L75","quality":"Less is better","values":[321,16,15,27,19,18],"isMain":false},{"metricName":"GC (%)","quality":"Equal","values":["51.05","50.72","50.66","50.50","50.82","50.86"],"isMain":false}]],["K-mer-based statistics",[]],["Predicted genes",[]],["Similarity statistics",[]],["Reference statistics",[{"metricName":"Reference length","quality":"Equal","values":[3222824,3222824,3222824,3222824,3222824,3222824],"isMain":false},{"metricName":"Reference fragments","quality":"Equal","values":[69,69,69,69,69,69],"isMain":false},{"metricName":"Reference GC (%)","quality":"Equal","values":["51.02","51.02","51.02","51.02","51.02","51.02"],"isMain":false}]]],"subreferences":[],"subreports":[],"minContig":500},{"date":"10 February 2020, Monday, 18:52:03","assembliesNames":["megahit-103-df","megahit-113-df","megahit-113-ml","megahit-113-ms","megahit-129-df","metaSPAdes"],"referenceName":"GCF\_000185525.1\_ASM18552v1","order":[0,1,2,3,4,5],"report":[["Genome statistics",[{"metricName":"Genome fraction (%)","quality":"More is better","values":["94.533","93.399","92.930","92.633","94.123","92.933"],"isMain":true},{"metricName":"Duplication ratio","quality":"Less is better","values":["1.027","1.020","1.075","1.076","1.026","1.024"],"isMain":true},{"metricName":"Largest alignment","quality":"More is better","values":[29680,41499,12529,13799,57926,21024],"isMain":true},{"metricName":"Total aligned length","quality":"More is better","values":[2935418,2877643,3016234,3003988,2916117,2876576],"isMain":true},{"metricName":"NG50","quality":"More is better","values":[7563,12555,3172,3248,12138,6337],"isMain":false},{"metricName":"NG75","quality":"More is better","values":[4000,6105,1799,1958,5743,3363],"isMain":false},{"metricName":"NA50","quality":"More is better","values":[7621,12991,3135,3153,11339,6503],"isMain":false},{"metricName":"NA75","quality":"More is better","values":[4275,6828,1770,1889,5576,3806],"isMain":false},{"metricName":"NGA50","quality":"More is better","values":[7304,12434,3135,3157,10842,6207],"isMain":true},{"metricName":"NGA75","quality":"More is better","values":[3831,5740,1776,1895,5291,3355],"isMain":false},{"metricName":"LG50","quality":"Less is better","values":[117,77,303,291,73,150],"isMain":false},{"metricName":"LG75","quality":"Less is better","values":[253,166,617,588,168,311],"isMain":false},{"metricName":"LA50","quality":"Less is better","values":[113,73,308,299,74,140],"isMain":false},{"metricName":"LA75","quality":"Less is better","values":[240,153,629,608,171,283],"isMain":false},{"metricName":"LGA50","quality":"Less is better","values":[118,79,308,298,77,151],"isMain":true},{"metricName":"LGA75","quality":"Less is better","values":[256,169,628,604,179,314],"isMain":false}]],["Reads mapping",[]],["Misassemblies",[{"metricName":"# misassemblies","quality":"Less is better","values":[8,20,20,17,17,9],"isMain":true},{"metricName":" # relocations","quality":"Less is better","values":[0,2,1,0,1,0],"isMain":false},{"metricName":" # translocations","quality":"Less is better","values":[8,17,18,15,15,9],"isMain":false},{"metricName":" # inversions","quality":"Less is better","values":[0,1,1,2,1,0],"isMain":false},{"metricName":" # interspecies translocations","quality":"Less is better","values":[11,18,20,49,34,7],"isMain":false},{"metricName":"# misassembled contigs","quality":"Less is better","values":[8,17,18,17,16,8],"isMain":false},{"metricName":"Misassembled contigs length","quality":"Less is better","values":[65897,162043,71392,69314,157041,39667],"isMain":true},{"metricName":"# possibly misassembled contigs","quality":"Less is better","values":[0,0,1,0,0,1],"isMain":false},{"metricName":" # possible misassemblies","quality":"Less is better","values":[0,0,2,0,0,2],"isMain":false},{"metricName":"# local misassemblies","quality":"Less is better","values":[1,4,4,2,3,1],"isMain":false},{"metricName":"# scaffold gap ext. mis.","quality":"Less is better","values":[0,0,0,0,0,0],"isMain":false},{"metricName":"# scaffold gap loc. mis.","quality":"Less is better","values":[0,0,0,0,0,0],"isMain":false},{"metricName":"# unaligned mis. contigs","quality":"Less is better","values":[0,1,1,1,1,0],"isMain":false}]],["Unaligned",[{"metricName":"# fully unaligned contigs","quality":"Less is better","values":[0,1,3,2,1,0],"isMain":false},{"metricName":"Fully unaligned length","quality":"Less is better","values":[0,1354,2731,1091,650,0],"isMain":false},{"metricName":"# partially unaligned contigs","quality":"Less is better","values":[3,3,3,10,7,1],"isMain":false},{"metricName":"Partially unaligned length","quality":"Less is better","values":[3232,2310,2597,15925,44216,925],"isMain":false}]],["Mismatches",[{"metricName":"# mismatches","quality":"Less is better","values":[7053,6215,24614,25202,6910,5806],"isMain":false},{"metricName":"# indels","quality":"Less is better","values":[52,78,110,114,71,56],"isMain":false},{"metricName":"Indels length","quality":"Less is better","values":[133,331,279,282,221,59],"isMain":false},{"metricName":"# mismatches per 100 kbp","quality":"Less is better","values":["246.27","219.64","874.27","898.03","242.33","206.22"],"isMain":true},{"metricName":"# indels per 100 kbp","quality":"Less is better","values":["1.82","2.76","3.91","4.06","2.49","1.99"],"isMain":true},{"metricName":" # indels (<= 5 bp)","quality":"Less is better","values":[50,70,105,108,67,56],"isMain":false},{"metricName":" # indels (> 5 bp)","quality":"Less is better","values":[2,8,5,6,4,0],"isMain":false},{"metricName":"# N's","quality":"Less is better","values":[0,0,0,0,0,0],"isMain":false},{"metricName":"# N's per 100 kbp","quality":"Less is better","values":["0.00","0.00","0.00","0.00","0.00","0.00"],"isMain":true}]],["Statistics without reference",[{"metricName":"# contigs","quality":"Equal","values":[629,422,1346,1293,471,647],"isMain":true},{"metricName":"# contigs (>= 1000 bp)","quality":"Equal","values":[513,356,971,932,376,568],"isMain":false},{"metricName":"# contigs (>= 5000 bp)","quality":"Equal","values":[207,191,112,122,190,208],"isMain":false},{"metricName":"# contigs (>= 10000 bp)","quality":"Equal","values":[80,94,6,12,90,62],"isMain":false},{"metricName":"# contigs (>= 25000 bp)","quality":"Equal","values":[7,15,0,0,17,0],"isMain":false},{"metricName":"# contigs (>= 50000 bp)","quality":"Equal","values":[0,0,0,0,1,0],"isMain":false},{"metricName":"Largest contig","quality":"More is better","values":[29699,49807,12933,14212,57942,21117],"isMain":true},{"metricName":"Total length","quality":"More is better","values":[2944128,2888734,3032741,3037996,2971346,2883523],"isMain":true},{"metricName":"Total length (>= 1000 bp)","quality":"More is better","values":[2861675,2842585,2758871,2775012,2902561,2825987],"isMain":true},{"metricName":"Total length (>= 5000 bp)","quality":"More is better","values":[2068373,2414450,755458,850209,2394369,1843363],"isMain":false},{"metricName":"Total length (>= 10000 bp)","quality":"More is better","values":[1206707,1714671,70348,143136,1700552,831770],"isMain":true},{"metricName":"Total length (>= 25000 bp)","quality":"More is better","values":[191576,469052,0,0,566416,0],"isMain":false},{"metricName":"Total length (>= 50000 bp)","quality":"More is better","values":[0,0,0,0,57942,0],"isMain":true},{"metricName":"N50","quality":"More is better","values":[7898,13445,3172,3246,12494,6565],"isMain":false},{"metricName":"N75","quality":"More is better","values":[4326,6931,1799,1946,5887,3881],"isMain":false},{"metricName":"L50","quality":"Less is better","values":[111,71,303,292,71,139],"isMain":false},{"metricName":"L75","quality":"Less is better","values":[237,150,618,592,160,280],"isMain":false},{"metricName":"GC (%)","quality":"Equal","values":["37.16","37.15","37.29","37.28","37.25","37.03"],"isMain":false}]],["K-mer-based statistics",[]],["Predicted genes",[]],["Similarity statistics",[]],["Reference statistics",[{"metricName":"Reference length","quality":"Equal","values":[3029555,3029555,3029555,3029555,3029555,3029555],"isMain":false},{"metricName":"Reference fragments","quality":"Equal","values":[96,96,96,96,96,96],"isMain":false},{"metricName":"Reference GC (%)","quality":"Equal","values":["37.21","37.21","37.21","37.21","37.21","37.21"],"isMain":false}]]],"subreferences":[],"subreports":[],"minContig":500},{"date":"10 February 2020, Monday, 18:51:29","assembliesNames":["megahit-103-df","megahit-113-df","megahit-113-ml","megahit-113-ms","megahit-129-df","metaSPAdes"],"referenceName":"GCF\_000185545.1\_ASM18554v1","order":[0,1,2,3,4,5],"report":[["Genome statistics",[{"metricName":"Genome fraction (%)","quality":"More is better","values":["19.547","21.504","20.765","21.374","25.001","15.880"],"isMain":true},{"metricName":"Duplication ratio","quality":"Less is better","values":["1.022","1.026","1.029","1.032","1.024","1.023"],"isMain":true},{"metricName":"Largest alignment","quality":"More is better","values":[2759,3793,3194,2976,6739,2465],"isMain":true},{"metricName":"Total aligned length","quality":"More is better","values":[574411,631498,615330,632266,735872,466370],"isMain":true},{"metricName":"NA50","quality":"More is better","values":[915,1007,839,853,1039,797],"isMain":false},{"metricName":"NA75","quality":"More is better","values":[668,699,629,618,689,614],"isMain":false},{"metricName":"NGA50","quality":"More is better","values":[null,null,null,null,null,null],"isMain":true},{"metricName":"LA50","quality":"Less is better","values":[228,222,262,270,248,215],"isMain":false},{"metricName":"LA75","quality":"Less is better","values":[417,415,478,496,482,384],"isMain":false}]],["Reads mapping",[]],["Misassemblies",[{"metricName":"# misassemblies","quality":"Less is better","values":[6,4,6,5,6,4],"isMain":true},{"metricName":" # relocations","quality":"Less is better","values":[0,0,0,0,0,1],"isMain":false},{"metricName":" # translocations","quality":"Less is better","values":[6,4,5,5,5,3],"isMain":false},{"metricName":" # inversions","quality":"Less is better","values":[0,0,1,0,1,0],"isMain":false},{"metricName":" # interspecies translocations","quality":"Less is better","values":[18,31,7,25,25,10],"isMain":false},{"metricName":"# misassembled contigs","quality":"Less is better","values":[6,4,6,4,5,4],"isMain":false},{"metricName":"Misassembled contigs length","quality":"Less is better","values":[7994,6440,5781,6044,8044,5546],"isMain":true},{"metricName":"# possibly misassembled contigs","quality":"Less is better","values":[0,0,0,0,1,0],"isMain":false},{"metricName":" # possible misassemblies","quality":"Less is better","values":[0,0,0,0,1,0],"isMain":false},{"metricName":"# local misassemblies","quality":"Less is better","values":[2,1,2,3,1,1],"isMain":false},{"metricName":"# scaffold gap ext. mis.","quality":"Less is better","values":[0,0,0,0,0,0],"isMain":false},{"metricName":"# scaffold gap loc. mis.","quality":"Less is better","values":[0,0,0,0,0,0],"isMain":false},{"metricName":"# unaligned mis. contigs","quality":"Less is better","values":[0,1,0,1,0,0],"isMain":false}]],["Unaligned",[{"metricName":"# fully unaligned contigs","quality":"Less is better","values":[0,1,1,0,0,0],"isMain":false},{"metricName":"Fully unaligned length","quality":"Less is better","values":[0,588,594,0,0,0],"isMain":false},{"metricName":"# partially unaligned contigs","quality":"Less is better","values":[4,8,2,4,7,2],"isMain":false},{"metricName":"Partially unaligned length","quality":"Less is better","values":[5047,9212,1075,11085,48918,1880],"isMain":false}]],["Mismatches",[{"metricName":"# mismatches","quality":"Less is better","values":[6504,7030,10692,11414,8535,6025],"isMain":false},{"metricName":"# indels","quality":"Less is better","values":[17,18,39,24,16,7],"isMain":false},{"metricName":"Indels length","quality":"Less is better","values":[21,18,130,37,38,87],"isMain":false},{"metricName":"# mismatches per 100 kbp","quality":"Less is better","values":["1143.73","1123.76","1769.95","1835.63","1173.48","1304.18"],"isMain":true},{"metricName":"# indels per 100 kbp","quality":"Less is better","values":["2.99","2.88","6.46","3.86","2.20","1.52"],"isMain":true},{"metricName":" # indels (<= 5 bp)","quality":"Less is better","values":[17,18,36,23,15,5],"isMain":false},{"metricName":" # indels (> 5 bp)","quality":"Less is better","values":[0,0,3,1,1,2],"isMain":false},{"metricName":"# N's","quality":"Less is better","values":[0,0,0,0,0,0],"isMain":false},{"metricName":"# N's per 100 kbp","quality":"Less is better","values":["0.00","0.00","0.00","0.00","0.00","0.00"],"isMain":true}]],["Statistics without reference",[{"metricName":"# contigs","quality":"Equal","values":[653,664,743,754,723,585],"isMain":true},{"metricName":"# contigs (>= 1000 bp)","quality":"Equal","values":[188,240,176,187,275,123],"isMain":false},{"metricName":"# contigs (>= 5000 bp)","quality":"Equal","values":[0,0,0,1,2,0],"isMain":false},{"metricName":"# contigs (>= 10000 bp)","quality":"Equal","values":[0,0,0,0,1,0],"isMain":false},{"metricName":"# contigs (>= 25000 bp)","quality":"Equal","values":[0,0,0,0,1,0],"isMain":false},{"metricName":"# contigs (>= 50000 bp)","quality":"Equal","values":[0,0,0,0,0,0],"isMain":false},{"metricName":"Largest contig","quality":"More is better","values":[3319,3973,3312,8229,45111,2465],"isMain":true},{"metricName":"Total length","quality":"More is better","values":[585958,651479,623316,652907,793650,474491],"isMain":true},{"metricName":"Total length (>= 1000 bp)","quality":"More is better","values":[261528,355220,235201,263296,475091,158905],"isMain":true},{"metricName":"Total length (>= 5000 bp)","quality":"More is better","values":[0,0,0,8229,51850,0],"isMain":false},{"metricName":"Total length (>= 10000 bp)","quality":"More is better","values":[0,0,0,0,45111,0],"isMain":true},{"metricName":"Total length (>= 25000 bp)","quality":"More is better","values":[0,0,0,0,45111,0],"isMain":false},{"metricName":"Total length (>= 50000 bp)","quality":"More is better","values":[0,0,0,0,0,0],"isMain":true},{"metricName":"N50","quality":"More is better","values":[944,1039,850,894,1136,826],"isMain":false},{"metricName":"N75","quality":"More is better","values":[692,745,646,654,795,634],"isMain":false},{"metricName":"L50","quality":"Less is better","values":[221,212,259,254,202,210],"isMain":false},{"metricName":"L75","quality":"Less is better","values":[404,395,470,469,409,374],"isMain":false},{"metricName":"GC (%)","quality":"Equal","values":["41.98","41.80","41.99","42.06","42.14","41.89"],"isMain":false}]],["K-mer-based statistics",[]],["Predicted genes",[]],["Similarity statistics",[]],["Reference statistics",[{"metricName":"Reference length","quality":"Equal","values":[2909168,2909168,2909168,2909168,2909168,2909168],"isMain":false},{"metricName":"Reference fragments","quality":"Equal","values":[117,117,117,117,117,117],"isMain":false},{"metricName":"Reference GC (%)","quality":"Equal","values":["41.86","41.86","41.86","41.86","41.86","41.86"],"isMain":false}]]],"subreferences":[],"subreports":[],"minContig":500},{"date":"10 February 2020, Monday, 18:51:19","assembliesNames":["megahit-103-df","megahit-113-df","megahit-113-ml","megahit-113-ms","megahit-129-df","metaSPAdes"],"referenceName":"GCF\_000185605.1\_ASM18560v1","order":[0,1,2,3,4,5],"report":[["Genome statistics",[{"metricName":"Genome fraction (%)","quality":"More is better","values":["0.825","1.315","1.505","1.608","1.598","0.816"],"isMain":true},{"metricName":"Duplication ratio","quality":"Less is better","values":["1.062","1.090","1.060","1.051","1.046","1.062"],"isMain":true},{"metricName":"Largest alignment","quality":"More is better","values":[2332,6893,2303,2437,8006,3436],"isMain":true},{"metricName":"Total aligned length","quality":"More is better","values":[37894,60115,68921,73824,73034,37622],"isMain":true},{"metricName":"NA50","quality":"More is better","values":[595,621,603,647,659,593],"isMain":false},{"metricName":"NA75","quality":"More is better","values":[504,null,514,527,519,511],"isMain":false},{"metricName":"NGA50","quality":"More is better","values":[null,null,null,null,null,null],"isMain":true},{"metricName":"LA50","quality":"Less is better","values":[23,37,44,41,32,18],"isMain":false},{"metricName":"LA75","quality":"Less is better","values":[43,null,79,76,69,38],"isMain":false}]],["Reads mapping",[]],["Misassemblies",[{"metricName":"# misassemblies","quality":"Less is better","values":[0,0,0,0,1,0],"isMain":true},{"metricName":" # relocations","quality":"Less is better","values":[0,0,0,0,0,0],"isMain":false},{"metricName":" # translocations","quality":"Less is better","values":[0,0,0,0,1,0],"isMain":false},{"metricName":" # inversions","quality":"Less is better","values":[0,0,0,0,0,0],"isMain":false},{"metricName":" # interspecies translocations","quality":"Less is better","values":[10,23,8,12,16,8],"isMain":false},{"metricName":"# misassembled contigs","quality":"Less is better","values":[0,0,0,0,1,0],"isMain":false},{"metricName":"Misassembled contigs length","quality":"Less is better","values":[0,0,0,0,521,0],"isMain":true},{"metricName":"# possibly misassembled contigs","quality":"Less is better","values":[0,0,0,0,0,0],"isMain":false},{"metricName":" # possible misassemblies","quality":"Less is better","values":[0,0,0,0,0,0],"isMain":false},{"metricName":"# local misassemblies","quality":"Less is better","values":[1,0,1,0,1,0],"isMain":false},{"metricName":"# scaffold gap ext. mis.","quality":"Less is better","values":[0,0,0,0,0,0],"isMain":false},{"metricName":"# scaffold gap loc. mis.","quality":"Less is better","values":[0,0,0,0,0,0],"isMain":false},{"metricName":"# unaligned mis. contigs","quality":"Less is better","values":[0,1,0,0,0,0],"isMain":false}]],["Unaligned",[{"metricName":"# fully unaligned contigs","quality":"Less is better","values":[0,1,0,0,2,0],"isMain":false},{"metricName":"Fully unaligned length","quality":"Less is better","values":[0,642,0,0,1444,0],"isMain":false},{"metricName":"# partially unaligned contigs","quality":"Less is better","values":[3,8,3,4,5,4],"isMain":false},{"metricName":"Partially unaligned length","quality":"Less is better","values":[4347,14600,3485,2856,7334,4500],"isMain":false}]],["Mismatches",[{"metricName":"# mismatches","quality":"Less is better","values":[565,879,1622,1437,1167,468],"isMain":false},{"metricName":"# indels","quality":"Less is better","values":[5,3,20,11,17,0],"isMain":false},{"metricName":"Indels length","quality":"Less is better","values":[6,3,77,29,36,0],"isMain":false},{"metricName":"# mismatches per 100 kbp","quality":"Less is better","values":["1498.40","1462.12","2357.94","1955.31","1597.49","1254.76"],"isMain":true},{"metricName":"# indels per 100 kbp","quality":"Less is better","values":["13.26","4.99","29.07","14.97","23.27","0.00"],"isMain":true},{"metricName":" # indels (<= 5 bp)","quality":"Less is better","values":[5,3,17,9,16,0],"isMain":false},{"metricName":" # indels (> 5 bp)","quality":"Less is better","values":[0,0,3,2,1,0],"isMain":false},{"metricName":"# N's","quality":"Less is better","values":[0,0,0,0,0,0],"isMain":false},{"metricName":"# N's per 100 kbp","quality":"Less is better","values":["0.00","0.00","0.00","0.00","0.00","0.00"],"isMain":true}]],["Statistics without reference",[{"metricName":"# contigs","quality":"Equal","values":[55,84,107,107,93,52],"isMain":true},{"metricName":"# contigs (>= 1000 bp)","quality":"Equal","values":[7,10,11,16,11,8],"isMain":false},{"metricName":"# contigs (>= 5000 bp)","quality":"Equal","values":[0,3,0,0,1,0],"isMain":false},{"metricName":"# contigs (>= 10000 bp)","quality":"Equal","values":[0,0,0,0,0,0],"isMain":false},{"metricName":"# contigs (>= 25000 bp)","quality":"Equal","values":[0,0,0,0,0,0],"isMain":false},{"metricName":"# contigs (>= 50000 bp)","quality":"Equal","values":[0,0,0,0,0,0],"isMain":false},{"metricName":"Largest contig","quality":"More is better","values":[3666,7067,2303,2439,8010,3436],"isMain":true},{"metricName":"Total length","quality":"More is better","values":[44403,80748,76416,80088,85161,44122],"isMain":true},{"metricName":"Total length (>= 1000 bp)","quality":"More is better","values":[14421,33837,15913,23866,33152,17326],"isMain":true},{"metricName":"Total length (>= 5000 bp)","quality":"More is better","values":[0,19473,0,0,8010,0],"isMain":false},{"metricName":"Total length (>= 10000 bp)","quality":"More is better","values":[0,0,0,0,0,0],"isMain":true},{"metricName":"Total length (>= 25000 bp)","quality":"More is better","values":[0,0,0,0,0,0],"isMain":false},{"metricName":"Total length (>= 50000 bp)","quality":"More is better","values":[0,0,0,0,0,0],"isMain":true},{"metricName":"N50","quality":"More is better","values":[680,748,652,702,762,741],"isMain":false},{"metricName":"N75","quality":"More is better","values":[578,624,556,566,608,570],"isMain":false},{"metricName":"L50","quality":"Less is better","values":[17,19,39,37,23,14],"isMain":false},{"metricName":"L75","quality":"Less is better","values":[35,48,71,70,54,32],"isMain":false},{"metricName":"GC (%)","quality":"Equal","values":["44.93","45.84","44.80","43.79","44.42","41.63"],"isMain":false}]],["K-mer-based statistics",[]],["Predicted genes",[]],["Similarity statistics",[]],["Reference statistics",[{"metricName":"Reference length","quality":"Equal","values":[4571407,4571407,4571407,4571407,4571407,4571407],"isMain":false},{"metricName":"Reference fragments","quality":"Equal","values":[85,85,85,85,85,85],"isMain":false},{"metricName":"Reference GC (%)","quality":"Equal","values":["44.65","44.65","44.65","44.65","44.65","44.65"],"isMain":false}]]],"subreferences":[],"subreports":[],"minContig":500},{"date":"10 February 2020, Monday, 18:51:37","assembliesNames":["megahit-103-df","megahit-113-df","megahit-113-ml","megahit-113-ms","megahit-129-df","metaSPAdes"],"referenceName":"GCF\_000185705.2\_Bilo\_wads\_3\_1\_6\_V2","order":[0,1,2,3,4,5],"report":[["Genome statistics",[{"metricName":"Genome fraction (%)","quality":"More is better","values":["8.742","10.737","13.485","11.902","12.129","14.273"],"isMain":true},{"metricName":"Duplication ratio","quality":"Less is better","values":["1.014","1.012","1.006","1.009","1.011","1.007"],"isMain":true},{"metricName":"Largest alignment","quality":"More is better","values":[34492,58444,60106,66431,66478,66468],"isMain":true},{"metricName":"Total aligned length","quality":"More is better","values":[385244,473045,593398,523275,535191,627285],"isMain":true},{"metricName":"NA50","quality":"More is better","values":[7162,16539,26452,27031,17431,21739],"isMain":false},{"metricName":"NA75","quality":"More is better","values":[849,993,7200,1891,1178,13993],"isMain":false},{"metricName":"NGA50","quality":"More is better","values":[null,null,null,null,null,null],"isMain":true},{"metricName":"LA50","quality":"Less is better","values":[15,7,9,7,8,9],"isMain":false},{"metricName":"LA75","quality":"Less is better","values":[51,39,18,22,30,17],"isMain":false}]],["Reads mapping",[]],["Misassemblies",[{"metricName":"# misassemblies","quality":"Less is better","values":[2,5,6,4,4,2],"isMain":true},{"metricName":" # relocations","quality":"Less is better","values":[0,2,4,1,1,1],"isMain":false},{"metricName":" # translocations","quality":"Less is better","values":[2,2,2,3,3,1],"isMain":false},{"metricName":" # inversions","quality":"Less is better","values":[0,1,0,0,0,0],"isMain":false},{"metricName":" # interspecies translocations","quality":"Less is better","values":[11,10,8,12,7,5],"isMain":false},{"metricName":"# misassembled contigs","quality":"Less is better","values":[2,5,6,3,4,2],"isMain":false},{"metricName":"Misassembled contigs length","quality":"Less is better","values":[2687,4821,237630,50006,56877,87106],"isMain":true},{"metricName":"# possibly misassembled contigs","quality":"Less is better","values":[0,0,0,0,0,0],"isMain":false},{"metricName":" # possible misassemblies","quality":"Less is better","values":[0,0,0,0,0,0],"isMain":false},{"metricName":"# local misassemblies","quality":"Less is better","values":[3,0,5,3,1,3],"isMain":false},{"metricName":"# scaffold gap ext. mis.","quality":"Less is better","values":[0,0,0,0,0,0],"isMain":false},{"metricName":"# scaffold gap loc. mis.","quality":"Less is better","values":[0,0,0,0,0,0],"isMain":false},{"metricName":"# unaligned mis. contigs","quality":"Less is better","values":[0,0,0,1,1,0],"isMain":false}]],["Unaligned",[{"metricName":"# fully unaligned contigs","quality":"Less is better","values":[1,0,0,1,0,0],"isMain":false},{"metricName":"Fully unaligned length","quality":"Less is better","values":[792,0,0,531,0,0],"isMain":false},{"metricName":"# partially unaligned contigs","quality":"Less is better","values":[4,2,2,3,4,0],"isMain":false},{"metricName":"Partially unaligned length","quality":"Less is better","values":[4063,2164,1816,10780,4723,0],"isMain":false}]],["Mismatches",[{"metricName":"# mismatches","quality":"Less is better","values":[3883,4125,5302,5054,4413,4615],"isMain":false},{"metricName":"# indels","quality":"Less is better","values":[133,106,163,143,132,222],"isMain":false},{"metricName":"Indels length","quality":"Less is better","values":[519,306,614,511,419,740],"isMain":false},{"metricName":"# mismatches per 100 kbp","quality":"Less is better","values":["1011.50","874.89","895.42","967.05","828.58","736.35"],"isMain":true},{"metricName":"# indels per 100 kbp","quality":"Less is better","values":["34.65","22.48","27.53","27.36","24.78","35.42"],"isMain":true},{"metricName":" # indels (<= 5 bp)","quality":"Less is better","values":[115,97,146,125,119,195],"isMain":false},{"metricName":" # indels (> 5 bp)","quality":"Less is better","values":[18,9,17,18,13,27],"isMain":false},{"metricName":"# N's","quality":"Less is better","values":[0,0,0,0,0,0],"isMain":false},{"metricName":"# N's per 100 kbp","quality":"Less is better","values":["0.00","0.00","0.00","0.00","0.00","0.00"],"isMain":true}]],["Statistics without reference",[{"metricName":"# contigs","quality":"Equal","values":[203,215,183,202,225,142],"isMain":true},{"metricName":"# contigs (>= 1000 bp)","quality":"Equal","values":[50,43,27,33,40,34],"isMain":false},{"metricName":"# contigs (>= 5000 bp)","quality":"Equal","values":[20,15,17,16,15,23],"isMain":false},{"metricName":"# contigs (>= 10000 bp)","quality":"Equal","values":[10,10,13,12,13,17],"isMain":false},{"metricName":"# contigs (>= 25000 bp)","quality":"Equal","values":[1,5,9,7,5,8],"isMain":false},{"metricName":"# contigs (>= 50000 bp)","quality":"Equal","values":[0,2,2,2,2,3],"isMain":false},{"metricName":"Largest contig","quality":"More is better","values":[34492,58603,89527,66499,66478,66468],"isMain":true},{"metricName":"Total length","quality":"More is better","values":[394036,479271,597593,538572,543423,630953],"isMain":true},{"metricName":"Total length (>= 1000 bp)","quality":"More is better","values":[297897,366494,497750,432934,422651,561133],"isMain":true},{"metricName":"Total length (>= 5000 bp)","quality":"More is better","values":[230584,318901,474717,403711,372461,540532],"isMain":false},{"metricName":"Total length (>= 10000 bp)","quality":"More is better","values":[161577,283077,445568,379038,357498,495915],"isMain":true},{"metricName":"Total length (>= 25000 bp)","quality":"More is better","values":[34492,210066,376851,295552,231621,339362],"isMain":false},{"metricName":"Total length (>= 50000 bp)","quality":"More is better","values":[0,116353,148505,116861,120514,183561],"isMain":true},{"metricName":"N50","quality":"More is better","values":[7240,16586,30574,27031,17638,25565],"isMain":false},{"metricName":"N75","quality":"More is better","values":[1030,1043,9252,4785,1391,14039],"isMain":false},{"metricName":"L50","quality":"Less is better","values":[15,7,7,7,8,8],"isMain":false},{"metricName":"L75","quality":"Less is better","values":[48,37,14,17,28,16],"isMain":false},{"metricName":"GC (%)","quality":"Equal","values":["58.91","58.84","59.31","58.73","58.83","59.97"],"isMain":false}]],["K-mer-based statistics",[]],["Predicted genes",[]],["Similarity statistics",[]],["Reference statistics",[{"metricName":"Reference length","quality":"Equal","values":[4391094,4391094,4391094,4391094,4391094,4391094],"isMain":false},{"metricName":"Reference fragments","quality":"Equal","values":[5,5,5,5,5,5],"isMain":false},{"metricName":"Reference GC (%)","quality":"Equal","values":["59.31","59.31","59.31","59.31","59.31","59.31"],"isMain":false}]]],"subreferences":[],"subreports":[],"minContig":500},{"date":"10 February 2020, Monday, 18:51:58","assembliesNames":["megahit-103-df","megahit-113-df","megahit-113-ml","megahit-113-ms","megahit-129-df","metaSPAdes"],"referenceName":"GCF\_000185825.1\_ASM18582v1","order":[0,1,2,3,4,5],"report":[["Genome statistics",[{"metricName":"Genome fraction (%)","quality":"More is better","values":["99.223","99.343","99.374","99.454","99.409","98.795"],"isMain":true},{"metricName":"Duplication ratio","quality":"Less is better","values":["1.003","1.005","1.005","1.007","1.005","1.001"],"isMain":true},{"metricName":"Largest alignment","quality":"More is better","values":[159679,124152,123247,107026,124152,128078],"isMain":true},{"metricName":"Total aligned length","quality":"More is better","values":[3235800,3246713,3246799,3256466,3248686,3216723],"isMain":true},{"metricName":"NG50","quality":"More is better","values":[34675,36813,34096,34133,35800,42303],"isMain":false},{"metricName":"NG75","quality":"More is better","values":[19151,22490,18924,21047,22373,24467],"isMain":false},{"metricName":"NA50","quality":"More is better","values":[33982,36476,33233,32899,35516,36419],"isMain":false},{"metricName":"NA75","quality":"More is better","values":[19156,22490,18924,19156,19173,22304],"isMain":false},{"metricName":"NGA50","quality":"More is better","values":[33209,36476,33233,32937,35516,37407],"isMain":true},{"metricName":"NGA75","quality":"More is better","values":[19151,22490,18924,19654,19642,22490],"isMain":false},{"metricName":"LG50","quality":"Less is better","values":[26,27,29,32,28,25],"isMain":false},{"metricName":"LG75","quality":"Less is better","values":[58,54,60,61,57,51],"isMain":false},{"metricName":"LA50","quality":"Less is better","values":[26,28,29,35,28,28],"isMain":false},{"metricName":"LA75","quality":"Less is better","values":[57,55,60,67,59,56],"isMain":false},{"metricName":"LGA50","quality":"Less is better","values":[27,28,29,34,28,27],"isMain":true},{"metricName":"LGA75","quality":"Less is better","values":[58,55,60,65,58,55],"isMain":false}]],["Reads mapping",[]],["Misassemblies",[{"metricName":"# misassemblies","quality":"Less is better","values":[3,7,5,6,4,6],"isMain":true},{"metricName":" # relocations","quality":"Less is better","values":[0,0,0,0,0,0],"isMain":false},{"metricName":" # translocations","quality":"Less is better","values":[3,7,5,6,4,6],"isMain":false},{"metricName":" # inversions","quality":"Less is better","values":[0,0,0,0,0,0],"isMain":false},{"metricName":" # interspecies translocations","quality":"Less is better","values":[7,8,9,16,16,3],"isMain":false},{"metricName":"# misassembled contigs","quality":"Less is better","values":[3,7,5,6,4,5],"isMain":false},{"metricName":"Misassembled contigs length","quality":"Less is better","values":[60219,191771,122532,160247,65003,221346],"isMain":true},{"metricName":"# possibly misassembled contigs","quality":"Less is better","values":[0,0,0,0,0,0],"isMain":false},{"metricName":" # possible misassemblies","quality":"Less is better","values":[0,0,0,0,0,0],"isMain":false},{"metricName":"# local misassemblies","quality":"Less is better","values":[1,1,3,4,2,1],"isMain":false},{"metricName":"# scaffold gap ext. mis.","quality":"Less is better","values":[0,0,0,0,0,0],"isMain":false},{"metricName":"# scaffold gap loc. mis.","quality":"Less is better","values":[0,0,0,0,0,0],"isMain":false},{"metricName":"# unaligned mis. contigs","quality":"Less is better","values":[0,0,0,0,0,0],"isMain":false}]],["Unaligned",[{"metricName":"# fully unaligned contigs","quality":"Less is better","values":[0,0,2,1,0,0],"isMain":false},{"metricName":"Fully unaligned length","quality":"Less is better","values":[0,0,1342,555,0,0],"isMain":false},{"metricName":"# partially unaligned contigs","quality":"Less is better","values":[1,3,0,9,8,2],"isMain":false},{"metricName":"Partially unaligned length","quality":"Less is better","values":[543,12981,0,53268,35522,61310],"isMain":false}]],["Mismatches",[{"metricName":"# mismatches","quality":"Less is better","values":[1238,1434,1990,2847,1343,336],"isMain":false},{"metricName":"# indels","quality":"Less is better","values":[17,25,21,20,38,26],"isMain":false},{"metricName":"Indels length","quality":"Less is better","values":[24,45,86,23,374,54],"isMain":false},{"metricName":"# mismatches per 100 kbp","quality":"Less is better","values":["38.33","44.35","61.53","87.95","41.51","10.45"],"isMain":true},{"metricName":"# indels per 100 kbp","quality":"Less is better","values":["0.53","0.77","0.65","0.62","1.17","0.81"],"isMain":true},{"metricName":" # indels (<= 5 bp)","quality":"Less is better","values":[17,23,20,20,31,24],"isMain":false},{"metricName":" # indels (> 5 bp)","quality":"Less is better","values":[0,2,1,0,7,2],"isMain":false},{"metricName":"# N's","quality":"Less is better","values":[0,0,0,0,0,0],"isMain":false},{"metricName":"# N's per 100 kbp","quality":"Less is better","values":["0.00","0.00","0.00","0.00","0.00","0.00"],"isMain":true}]],["Statistics without reference",[{"metricName":"# contigs","quality":"Equal","values":[173,165,185,198,185,147],"isMain":true},{"metricName":"# contigs (>= 1000 bp)","quality":"Equal","values":[153,145,161,167,155,135],"isMain":false},{"metricName":"# contigs (>= 5000 bp)","quality":"Equal","values":[118,110,122,123,117,107],"isMain":false},{"metricName":"# contigs (>= 10000 bp)","quality":"Equal","values":[94,93,94,99,94,90],"isMain":false},{"metricName":"# contigs (>= 25000 bp)","quality":"Equal","values":[43,49,45,52,47,48],"isMain":false},{"metricName":"# contigs (>= 50000 bp)","quality":"Equal","values":[15,16,15,14,13,20],"isMain":false},{"metricName":"Largest contig","quality":"More is better","values":[159679,124152,123247,107186,124152,128078],"isMain":true},{"metricName":"Total length","quality":"More is better","values":[3239049,3262813,3251219,3314580,3287593,3279365],"isMain":true},{"metricName":"Total length (>= 1000 bp)","quality":"More is better","values":[3226073,3248793,3234719,3294092,3267161,3269677],"isMain":true},{"metricName":"Total length (>= 5000 bp)","quality":"More is better","values":[3140300,3166225,3136356,3184834,3173896,3189192],"isMain":false},{"metricName":"Total length (>= 10000 bp)","quality":"More is better","values":[2966299,3050320,2932666,3015980,3019865,3072921],"isMain":true},{"metricName":"Total length (>= 25000 bp)","quality":"More is better","values":[2127511,2333123,2121472,2238664,2216376,2375482],"isMain":false},{"metricName":"Total length (>= 50000 bp)","quality":"More is better","values":[1183159,1203160,1094495,926221,1032323,1407142],"isMain":true},{"metricName":"N50","quality":"More is better","values":[34675,36813,34096,33957,35800,39796],"isMain":false},{"metricName":"N75","quality":"More is better","values":[19156,22490,18924,19654,21511,24197],"isMain":false},{"metricName":"L50","quality":"Less is better","values":[26,27,29,33,28,26],"isMain":false},{"metricName":"L75","quality":"Less is better","values":[57,54,60,64,58,52],"isMain":false},{"metricName":"GC (%)","quality":"Equal","values":["44.43","44.40","44.43","44.38","44.44","44.51"],"isMain":false}]],["K-mer-based statistics",[]],["Predicted genes",[]],["Similarity statistics",[]],["Reference statistics",[{"metricName":"Reference length","quality":"Equal","values":[3254802,3254802,3254802,3254802,3254802,3254802],"isMain":false},{"metricName":"Reference fragments","quality":"Equal","values":[116,116,116,116,116,116],"isMain":false},{"metricName":"Reference GC (%)","quality":"Equal","values":["44.43","44.43","44.43","44.43","44.43","44.43"],"isMain":false}]]],"subreferences":[],"subreports":[],"minContig":500},{"date":"10 February 2020, Monday, 18:51:32","assembliesNames":["megahit-103-df","megahit-113-df","megahit-113-ml","megahit-113-ms","megahit-129-df","metaSPAdes"],"referenceName":"GCF\_000186105.1\_Bacteroides\_sp\_3\_1\_40A\_V1","order":[0,1,2,3,4,5],"report":[["Genome statistics",[{"metricName":"Genome fraction (%)","quality":"More is better","values":["2.956","3.297","4.709","4.037","2.170","1.604"],"isMain":true},{"metricName":"Duplication ratio","quality":"Less is better","values":["1.022","1.028","1.037","1.036","1.043","1.019"],"isMain":true},{"metricName":"Largest alignment","quality":"More is better","values":[5824,10056,7040,4259,9487,5326],"isMain":true},{"metricName":"Total aligned length","quality":"More is better","values":[163313,181231,263329,222864,119482,87545],"isMain":true},{"metricName":"NA50","quality":"More is better","values":[1834,2426,1296,1351,1402,1661],"isMain":false},{"metricName":"NA75","quality":"More is better","values":[962,1115,794,821,535,889],"isMain":false},{"metricName":"NGA50","quality":"More is better","values":[null,null,null,null,null,null],"isMain":true},{"metricName":"LA50","quality":"Less is better","values":[29,21,61,57,29,16],"isMain":false},{"metricName":"LA75","quality":"Less is better","values":[62,51,129,114,74,34],"isMain":false}]],["Reads mapping",[]],["Misassemblies",[{"metricName":"# misassemblies","quality":"Less is better","values":[3,4,2,5,4,2],"isMain":true},{"metricName":" # relocations","quality":"Less is better","values":[0,0,0,1,1,0],"isMain":false},{"metricName":" # translocations","quality":"Less is better","values":[1,4,1,4,2,2],"isMain":false},{"metricName":" # inversions","quality":"Less is better","values":[2,0,1,0,1,0],"isMain":false},{"metricName":" # interspecies translocations","quality":"Less is better","values":[5,14,9,18,15,5],"isMain":false},{"metricName":"# misassembled contigs","quality":"Less is better","values":[3,3,2,5,4,2],"isMain":false},{"metricName":"Misassembled contigs length","quality":"Less is better","values":[5934,10197,2380,7437,5469,4469],"isMain":true},{"metricName":"# possibly misassembled contigs","quality":"Less is better","values":[0,0,0,0,0,0],"isMain":false},{"metricName":" # possible misassemblies","quality":"Less is better","values":[0,0,0,0,0,0],"isMain":false},{"metricName":"# local misassemblies","quality":"Less is better","values":[0,0,0,1,0,0],"isMain":false},{"metricName":"# scaffold gap ext. mis.","quality":"Less is better","values":[0,0,0,0,0,0],"isMain":false},{"metricName":"# scaffold gap loc. mis.","quality":"Less is better","values":[0,0,0,0,0,0],"isMain":false},{"metricName":"# unaligned mis. contigs","quality":"Less is better","values":[0,0,0,0,1,0],"isMain":false}]],["Unaligned",[{"metricName":"# fully unaligned contigs","quality":"Less is better","values":[0,0,3,3,0,1],"isMain":false},{"metricName":"Fully unaligned length","quality":"Less is better","values":[0,0,2294,2143,0,1175],"isMain":false},{"metricName":"# partially unaligned contigs","quality":"Less is better","values":[2,6,3,5,2,1],"isMain":false},{"metricName":"Partially unaligned length","quality":"Less is better","values":[2149,11544,2720,4963,26575,880],"isMain":false}]],["Mismatches",[{"metricName":"# mismatches","quality":"Less is better","values":[1542,1884,3944,3447,1485,656],"isMain":false},{"metricName":"# indels","quality":"Less is better","values":[12,20,31,21,42,14],"isMain":false},{"metricName":"Indels length","quality":"Less is better","values":[23,71,57,84,80,19],"isMain":false},{"metricName":"# mismatches per 100 kbp","quality":"Less is better","values":["958.94","1050.53","1539.64","1569.67","1257.81","751.91"],"isMain":true},{"metricName":"# indels per 100 kbp","quality":"Less is better","values":["7.46","11.15","12.10","9.56","35.57","16.05"],"isMain":true},{"metricName":" # indels (<= 5 bp)","quality":"Less is better","values":[10,16,29,17,39,14],"isMain":false},{"metricName":" # indels (> 5 bp)","quality":"Less is better","values":[2,4,2,4,3,0],"isMain":false},{"metricName":"# N's","quality":"Less is better","values":[0,0,0,0,0,0],"isMain":false},{"metricName":"# N's per 100 kbp","quality":"Less is better","values":["0.00","0.00","0.00","0.00","0.00","0.00"],"isMain":true}]],["Statistics without reference",[{"metricName":"# contigs","quality":"Equal","values":[118,99,228,197,87,63],"isMain":true},{"metricName":"# contigs (>= 1000 bp)","quality":"Equal","values":[60,58,95,86,42,34],"isMain":false},{"metricName":"# contigs (>= 5000 bp)","quality":"Equal","values":[1,9,1,0,5,1],"isMain":false},{"metricName":"# contigs (>= 10000 bp)","quality":"Equal","values":[0,1,0,0,2,0],"isMain":false},{"metricName":"# contigs (>= 25000 bp)","quality":"Equal","values":[0,0,0,0,0,0],"isMain":false},{"metricName":"# contigs (>= 50000 bp)","quality":"Equal","values":[0,0,0,0,0,0],"isMain":false},{"metricName":"Largest contig","quality":"More is better","values":[5840,10191,7049,4259,19590,5337],"isMain":true},{"metricName":"Total length","quality":"More is better","values":[166531,195972,270620,234613,149770,90957],"isMain":true},{"metricName":"Total length (>= 1000 bp)","quality":"More is better","values":[125532,167222,176930,155477,118754,71309],"isMain":true},{"metricName":"Total length (>= 5000 bp)","quality":"More is better","values":[5840,63303,7049,0,50784,5337],"isMain":false},{"metricName":"Total length (>= 10000 bp)","quality":"More is better","values":[0,10191,0,0,30292,0],"isMain":true},{"metricName":"Total length (>= 25000 bp)","quality":"More is better","values":[0,0,0,0,0,0],"isMain":false},{"metricName":"Total length (>= 50000 bp)","quality":"More is better","values":[0,0,0,0,0,0],"isMain":true},{"metricName":"N50","quality":"More is better","values":[1851,3142,1360,1387,2379,2232],"isMain":false},{"metricName":"N75","quality":"More is better","values":[1001,1470,836,885,1108,1026],"isMain":false},{"metricName":"L50","quality":"Less is better","values":[28,18,59,54,13,15],"isMain":false},{"metricName":"L75","quality":"Less is better","values":[60,41,124,108,37,31],"isMain":false},{"metricName":"GC (%)","quality":"Equal","values":["42.58","42.67","42.51","42.56","41.98","42.73"],"isMain":false}]],["K-mer-based statistics",[]],["Predicted genes",[]],["Similarity statistics",[]],["Reference statistics",[{"metricName":"Reference length","quality":"Equal","values":[5439699,5439699,5439699,5439699,5439699,5439699],"isMain":false},{"metricName":"Reference fragments","quality":"Equal","values":[195,195,195,195,195,195],"isMain":false},{"metricName":"Reference GC (%)","quality":"Equal","values":["42.58","42.58","42.58","42.58","42.58","42.58"],"isMain":false}]]],"subreferences":[],"subreports":[],"minContig":500},{"date":"10 February 2020, Monday, 18:51:38","assembliesNames":["megahit-103-df","megahit-113-df","megahit-113-ml","megahit-113-ms","megahit-129-df","metaSPAdes"],"referenceName":"GCF\_000187895.1\_ASM18789v1","order":[0,1,2,3,4,5],"report":[["Genome statistics",[{"metricName":"Genome fraction (%)","quality":"More is better","values":["0.822","0.774","0.464","0.535","0.271","0.221"],"isMain":true},{"metricName":"Duplication ratio","quality":"Less is better","values":["1.037","1.091","1.045","1.133","1.164","1.097"],"isMain":true},{"metricName":"Largest alignment","quality":"More is better","values":[12745,13916,6614,8225,1928,2535],"isMain":true},{"metricName":"Total aligned length","quality":"More is better","values":[36353,34187,20536,23892,11962,9778],"isMain":true},{"metricName":"NA50","quality":"More is better","values":[11216,10353,742,1003,613,890],"isMain":false},{"metricName":"NA75","quality":"More is better","values":[1586,601,519,517,116,680],"isMain":false},{"metricName":"NGA50","quality":"More is better","values":[null,null,null,null,null,null],"isMain":true},{"metricName":"LA50","quality":"Less is better","values":[2,2,4,4,8,4],"isMain":false},{"metricName":"LA75","quality":"Less is better","values":[5,9,13,15,19,7],"isMain":false}]],["Reads mapping",[]],["Misassemblies",[{"metricName":"# misassemblies","quality":"Less is better","values":[1,0,1,0,1,0],"isMain":true},{"metricName":" # relocations","quality":"Less is better","values":[1,0,1,0,1,0],"isMain":false},{"metricName":" # translocations","quality":"Less is better","values":[0,0,0,0,0,0],"isMain":false},{"metricName":" # inversions","quality":"Less is better","values":[0,0,0,0,0,0],"isMain":false},{"metricName":" # interspecies translocations","quality":"Less is better","values":[5,9,3,8,8,3],"isMain":false},{"metricName":"# misassembled contigs","quality":"Less is better","values":[1,0,1,0,1,0],"isMain":false},{"metricName":"Misassembled contigs length","quality":"Less is better","values":[1190,0,985,0,1305,0],"isMain":true},{"metricName":"# possibly misassembled contigs","quality":"Less is better","values":[0,0,0,0,0,0],"isMain":false},{"metricName":" # possible misassemblies","quality":"Less is better","values":[0,0,0,0,0,0],"isMain":false},{"metricName":"# local misassemblies","quality":"Less is better","values":[0,0,0,0,0,0],"isMain":false},{"metricName":"# scaffold gap ext. mis.","quality":"Less is better","values":[0,0,0,0,0,0],"isMain":false},{"metricName":"# scaffold gap loc. mis.","quality":"Less is better","values":[0,0,0,0,0,0],"isMain":false},{"metricName":"# unaligned mis. contigs","quality":"Less is better","values":[0,0,0,0,0,0],"isMain":false}]],["Unaligned",[{"metricName":"# fully unaligned contigs","quality":"Less is better","values":[0,0,0,0,0,0],"isMain":false},{"metricName":"Fully unaligned length","quality":"Less is better","values":[0,0,0,0,0,0],"isMain":false},{"metricName":"# partially unaligned contigs","quality":"Less is better","values":[1,2,1,2,3,0],"isMain":false},{"metricName":"Partially unaligned length","quality":"Less is better","values":[2051,2753,1001,1388,1976,0],"isMain":false}]],["Mismatches",[{"metricName":"# mismatches","quality":"Less is better","values":[174,323,339,360,201,164],"isMain":false},{"metricName":"# indels","quality":"Less is better","values":[9,5,7,3,4,1],"isMain":false},{"metricName":"Indels length","quality":"Less is better","values":[14,60,9,3,9,2],"isMain":false},{"metricName":"# mismatches per 100 kbp","quality":"Less is better","values":["478.59","943.31","1650.84","1522.33","1680.18","1676.89"],"isMain":true},{"metricName":"# indels per 100 kbp","quality":"Less is better","values":["24.75","14.60","34.09","12.69","33.44","10.22"],"isMain":true},{"metricName":" # indels (<= 5 bp)","quality":"Less is better","values":[9,4,7,3,4,1],"isMain":false},{"metricName":" # indels (> 5 bp)","quality":"Less is better","values":[0,1,0,0,0,0],"isMain":false},{"metricName":"# N's","quality":"Less is better","values":[0,0,0,0,0,0],"isMain":false},{"metricName":"# N's per 100 kbp","quality":"Less is better","values":["0.00","0.00","0.00","0.00","0.00","0.00"],"isMain":true}]],["Statistics without reference",[{"metricName":"# contigs","quality":"Equal","values":[15,22,21,25,18,11],"isMain":true},{"metricName":"# contigs (>= 1000 bp)","quality":"Equal","values":[9,5,4,4,4,2],"isMain":false},{"metricName":"# contigs (>= 5000 bp)","quality":"Equal","values":[2,2,1,1,0,0],"isMain":false},{"metricName":"# contigs (>= 10000 bp)","quality":"Equal","values":[2,2,0,0,0,0],"isMain":false},{"metricName":"# contigs (>= 25000 bp)","quality":"Equal","values":[0,0,0,0,0,0],"isMain":false},{"metricName":"# contigs (>= 50000 bp)","quality":"Equal","values":[0,0,0,0,0,0],"isMain":false},{"metricName":"Largest contig","quality":"More is better","values":[12836,13943,6620,8248,2134,2548],"isMain":true},{"metricName":"Total length","quality":"More is better","values":[39740,40111,22451,28175,15899,10733],"isMain":true},{"metricName":"Total length (>= 1000 bp)","quality":"More is better","values":[35831,29621,12411,14943,6276,4157],"isMain":true},{"metricName":"Total length (>= 5000 bp)","quality":"More is better","values":[24052,24313,6620,8248,0,0],"isMain":false},{"metricName":"Total length (>= 10000 bp)","quality":"More is better","values":[24052,24313,0,0,0,0],"isMain":true},{"metricName":"Total length (>= 25000 bp)","quality":"More is better","values":[0,0,0,0,0,0],"isMain":false},{"metricName":"Total length (>= 50000 bp)","quality":"More is better","values":[0,0,0,0,0,0],"isMain":true},{"metricName":"N50","quality":"More is better","values":[11216,10370,1545,1007,892,901],"isMain":false},{"metricName":"N75","quality":"More is better","values":[1648,982,562,648,718,740],"isMain":false},{"metricName":"L50","quality":"Less is better","values":[2,2,3,4,6,4],"isMain":false},{"metricName":"L75","quality":"Less is better","values":[5,6,11,13,11,7],"isMain":false},{"metricName":"GC (%)","quality":"Equal","values":["38.70","36.97","38.34","41.83","42.99","43.48"],"isMain":false}]],["K-mer-based statistics",[]],["Predicted genes",[]],["Similarity statistics",[]],["Reference statistics",[{"metricName":"Reference length","quality":"Equal","values":[4421324,4421324,4421324,4421324,4421324,4421324],"isMain":false},{"metricName":"Reference fragments","quality":"Equal","values":[25,25,25,25,25,25],"isMain":false},{"metricName":"Reference GC (%)","quality":"Equal","values":["44.31","44.31","44.31","44.31","44.31","44.31"],"isMain":false}]]],"subreferences":[],"subreports":[],"minContig":500},{"date":"10 February 2020, Monday, 18:51:34","assembliesNames":["megahit-103-df","megahit-113-df","megahit-113-ml","megahit-113-ms","megahit-129-df","metaSPAdes"],"referenceName":"GCF\_000190535.1\_ASM19053v1","order":[0,1,2,3,4,5],"report":[["Genome statistics",[{"metricName":"Genome fraction (%)","quality":"More is better","values":["2.656","3.715","3.371","4.242","3.931","1.459"],"isMain":true},{"metricName":"Duplication ratio","quality":"Less is better","values":["1.048","1.041","1.030","1.035","1.027","1.067"],"isMain":true},{"metricName":"Largest alignment","quality":"More is better","values":[1319,2131,1180,1493,2505,1051],"isMain":true},{"metricName":"Total aligned length","quality":"More is better","values":[117154,163684,148568,186896,173160,64184],"isMain":true},{"metricName":"NA50","quality":"More is better","values":[566,572,559,567,574,540],"isMain":false},{"metricName":"NA75","quality":"More is better","values":[510,513,520,514,515,503],"isMain":false},{"metricName":"NGA50","quality":"More is better","values":[null,null,null,null,null,null],"isMain":true},{"metricName":"LA50","quality":"Less is better","values":[89,128,115,148,133,57],"isMain":false},{"metricName":"LA75","quality":"Less is better","values":[148,211,187,241,222,92],"isMain":false}]],["Reads mapping",[]],["Misassemblies",[{"metricName":"# misassemblies","quality":"Less is better","values":[2,6,3,5,2,1],"isMain":true},{"metricName":" # relocations","quality":"Less is better","values":[2,6,3,5,2,1],"isMain":false},{"metricName":" # translocations","quality":"Less is better","values":[0,0,0,0,0,0],"isMain":false},{"metricName":" # inversions","quality":"Less is better","values":[0,0,0,0,0,0],"isMain":false},{"metricName":" # interspecies translocations","quality":"Less is better","values":[17,26,7,23,25,17],"isMain":false},{"metricName":"# misassembled contigs","quality":"Less is better","values":[2,6,3,5,2,1],"isMain":false},{"metricName":"Misassembled contigs length","quality":"Less is better","values":[1490,4075,2137,3453,1473,569],"isMain":true},{"metricName":"# possibly misassembled contigs","quality":"Less is better","values":[0,0,0,0,0,0],"isMain":false},{"metricName":" # possible misassemblies","quality":"Less is better","values":[0,0,0,0,0,0],"isMain":false},{"metricName":"# local misassemblies","quality":"Less is better","values":[1,0,1,0,1,0],"isMain":false},{"metricName":"# scaffold gap ext. mis.","quality":"Less is better","values":[0,0,0,0,0,0],"isMain":false},{"metricName":"# scaffold gap loc. mis.","quality":"Less is better","values":[0,0,0,0,0,0],"isMain":false},{"metricName":"# unaligned mis. contigs","quality":"Less is better","values":[0,0,0,0,0,0],"isMain":false}]],["Unaligned",[{"metricName":"# fully unaligned contigs","quality":"Less is better","values":[0,0,1,0,3,0],"isMain":false},{"metricName":"Fully unaligned length","quality":"Less is better","values":[0,0,929,0,2231,0],"isMain":false},{"metricName":"# partially unaligned contigs","quality":"Less is better","values":[2,10,2,9,13,6],"isMain":false},{"metricName":"Partially unaligned length","quality":"Less is better","values":[2459,11720,1075,8694,14263,4577],"isMain":false}]],["Mismatches",[{"metricName":"# mismatches","quality":"Less is better","values":[2054,2880,2921,3841,3002,1214],"isMain":false},{"metricName":"# indels","quality":"Less is better","values":[9,12,7,17,9,5],"isMain":false},{"metricName":"Indels length","quality":"Less is better","values":[9,12,7,20,9,5],"isMain":false},{"metricName":"# mismatches per 100 kbp","quality":"Less is better","values":["1760.99","1764.88","1972.86","2061.52","1738.59","1894.63"],"isMain":true},{"metricName":"# indels per 100 kbp","quality":"Less is better","values":["7.72","7.35","4.73","9.12","5.21","7.80"],"isMain":true},{"metricName":" # indels (<= 5 bp)","quality":"Less is better","values":[9,12,7,17,9,5],"isMain":false},{"metricName":" # indels (> 5 bp)","quality":"Less is better","values":[0,0,0,0,0,0],"isMain":false},{"metricName":"# N's","quality":"Less is better","values":[0,0,0,0,0,0],"isMain":false},{"metricName":"# N's per 100 kbp","quality":"Less is better","values":["0.00","0.00","0.00","0.00","0.00","0.00"],"isMain":true}]],["Statistics without reference",[{"metricName":"# contigs","quality":"Equal","values":[201,277,254,321,292,119],"isMain":true},{"metricName":"# contigs (>= 1000 bp)","quality":"Equal","values":[7,12,4,11,14,5],"isMain":false},{"metricName":"# contigs (>= 5000 bp)","quality":"Equal","values":[0,0,0,0,0,0],"isMain":false},{"metricName":"# contigs (>= 10000 bp)","quality":"Equal","values":[0,0,0,0,0,0],"isMain":false},{"metricName":"# contigs (>= 25000 bp)","quality":"Equal","values":[0,0,0,0,0,0],"isMain":false},{"metricName":"# contigs (>= 50000 bp)","quality":"Equal","values":[0,0,0,0,0,0],"isMain":false},{"metricName":"Largest contig","quality":"More is better","values":[1674,3148,1329,1823,3271,1312],"isMain":true},{"metricName":"Total length","quality":"More is better","values":[124704,181604,154489,201470,193782,72947],"isMain":true},{"metricName":"Total length (>= 1000 bp)","quality":"More is better","values":[8655,19093,4632,15011,21945,5453],"isMain":true},{"metricName":"Total length (>= 5000 bp)","quality":"More is better","values":[0,0,0,0,0,0],"isMain":false},{"metricName":"Total length (>= 10000 bp)","quality":"More is better","values":[0,0,0,0,0,0],"isMain":true},{"metricName":"Total length (>= 25000 bp)","quality":"More is better","values":[0,0,0,0,0,0],"isMain":false},{"metricName":"Total length (>= 50000 bp)","quality":"More is better","values":[0,0,0,0,0,0],"isMain":true},{"metricName":"N50","quality":"More is better","values":[586,611,578,595,618,586],"isMain":false},{"metricName":"N75","quality":"More is better","values":[527,544,531,539,547,537],"isMain":false},{"metricName":"L50","quality":"Less is better","values":[85,111,110,135,116,51],"isMain":false},{"metricName":"L75","quality":"Less is better","values":[141,190,180,224,199,84],"isMain":false},{"metricName":"GC (%)","quality":"Equal","values":["44.09","43.86","43.82","43.74","44.00","44.12"],"isMain":false}]],["K-mer-based statistics",[]],["Predicted genes",[]],["Similarity statistics",[]],["Reference statistics",[{"metricName":"Reference length","quality":"Equal","values":[4392288,4392288,4392288,4392288,4392288,4392288],"isMain":false},{"metricName":"Reference fragments","quality":"Equal","values":[1,1,1,1,1,1],"isMain":false},{"metricName":"Reference GC (%)","quality":"Equal","values":["43.36","43.36","43.36","43.36","43.36","43.36"],"isMain":false}]]],"subreferences":[],"subreports":[],"minContig":500},{"date":"10 February 2020, Monday, 18:51:56","assembliesNames":["megahit-103-df","megahit-113-df","megahit-113-ml","megahit-113-ms","megahit-129-df","metaSPAdes"],"referenceName":"GCF\_000191545.1\_ASM19154v1","order":[0,1,2,3,4,5],"report":[["Genome statistics",[{"metricName":"Genome fraction (%)","quality":"More is better","values":["92.236","91.437","93.792","93.462","93.370","83.957"],"isMain":true},{"metricName":"Duplication ratio","quality":"Less is better","values":["1.015","1.051","1.054","1.061","1.059","1.010"],"isMain":true},{"metricName":"Largest alignment","quality":"More is better","values":[54480,45651,32565,48336,44425,37749],"isMain":true},{"metricName":"Total aligned length","quality":"More is better","values":[1961178,2001828,2059987,2062184,2062371,1776242],"isMain":true},{"metricName":"NG50","quality":"More is better","values":[10495,9019,8090,7701,8603,6185],"isMain":false},{"metricName":"NG75","quality":"More is better","values":[4450,4470,4025,4364,4672,2262],"isMain":false},{"metricName":"NA50","quality":"More is better","values":[11076,6882,7980,6761,6904,7561],"isMain":false},{"metricName":"NA75","quality":"More is better","values":[5631,2869,3891,3256,2898,3663],"isMain":false},{"metricName":"NGA50","quality":"More is better","values":[10495,7627,7980,6978,7491,6161],"isMain":true},{"metricName":"NGA75","quality":"More is better","values":[4450,3313,3931,3672,3515,2178],"isMain":false},{"metricName":"LG50","quality":"Less is better","values":[54,63,82,76,69,90],"isMain":false},{"metricName":"LG75","quality":"Less is better","values":[129,144,176,166,149,229],"isMain":false},{"metricName":"LA50","quality":"Less is better","values":[48,82,84,92,89,67],"isMain":false},{"metricName":"LA75","quality":"Less is better","values":[109,201,181,207,206,151],"isMain":false},{"metricName":"LGA50","quality":"Less is better","values":[54,77,84,86,80,90],"isMain":true},{"metricName":"LGA75","quality":"Less is better","values":[129,181,180,189,179,231],"isMain":false}]],["Reads mapping",[]],["Misassemblies",[{"metricName":"# misassemblies","quality":"Less is better","values":[2,4,7,9,7,0],"isMain":true},{"metricName":" # relocations","quality":"Less is better","values":[1,4,5,9,7,0],"isMain":false},{"metricName":" # translocations","quality":"Less is better","values":[0,0,0,0,0,0],"isMain":false},{"metricName":" # inversions","quality":"Less is better","values":[1,0,2,0,0,0],"isMain":false},{"metricName":" # interspecies translocations","quality":"Less is better","values":[7,43,35,53,54,7],"isMain":false},{"metricName":"# misassembled contigs","quality":"Less is better","values":[2,4,7,9,7,0],"isMain":false},{"metricName":"Misassembled contigs length","quality":"Less is better","values":[1044,52751,30765,47626,46080,0],"isMain":true},{"metricName":"# possibly misassembled contigs","quality":"Less is better","values":[0,2,0,1,0,1],"isMain":false},{"metricName":" # possible misassemblies","quality":"Less is better","values":[0,2,0,1,0,1],"isMain":false},{"metricName":"# local misassemblies","quality":"Less is better","values":[2,5,3,4,6,0],"isMain":false},{"metricName":"# scaffold gap ext. mis.","quality":"Less is better","values":[0,0,0,0,0,0],"isMain":false},{"metricName":"# scaffold gap loc. mis.","quality":"Less is better","values":[0,0,0,0,0,0],"isMain":false},{"metricName":"# unaligned mis. contigs","quality":"Less is better","values":[0,1,0,5,1,0],"isMain":false}]],["Unaligned",[{"metricName":"# fully unaligned contigs","quality":"Less is better","values":[1,1,3,0,3,1],"isMain":false},{"metricName":"Fully unaligned length","quality":"Less is better","values":[552,506,2011,0,1725,1275],"isMain":false},{"metricName":"# partially unaligned contigs","quality":"Less is better","values":[0,22,15,23,29,4],"isMain":false},{"metricName":"Partially unaligned length","quality":"Less is better","values":[0,162119,25196,102979,137587,4922],"isMain":false}]],["Mismatches",[{"metricName":"# mismatches","quality":"Less is better","values":[2499,9526,10171,12214,9066,2828],"isMain":false},{"metricName":"# indels","quality":"Less is better","values":[29,93,95,145,76,54],"isMain":false},{"metricName":"Indels length","quality":"Less is better","values":[39,232,431,432,313,345],"isMain":false},{"metricName":"# mismatches per 100 kbp","quality":"Less is better","values":["129.15","496.63","516.94","622.97","462.86","160.57"],"isMain":true},{"metricName":"# indels per 100 kbp","quality":"Less is better","values":["1.50","4.85","4.83","7.40","3.88","3.07"],"isMain":true},{"metricName":" # indels (<= 5 bp)","quality":"Less is better","values":[28,85,86,135,68,39],"isMain":false},{"metricName":" # indels (> 5 bp)","quality":"Less is better","values":[1,8,9,10,8,15],"isMain":false},{"metricName":"# N's","quality":"Less is better","values":[0,0,0,0,0,0],"isMain":false},{"metricName":"# N's per 100 kbp","quality":"Less is better","values":["0.00","0.00","0.00","0.00","0.00","0.00"],"isMain":true}]],["Statistics without reference",[{"metricName":"# contigs","quality":"Equal","values":[366,519,539,559,568,403],"isMain":true},{"metricName":"# contigs (>= 1000 bp)","quality":"Equal","values":[264,345,362,378,360,335],"isMain":false},{"metricName":"# contigs (>= 5000 bp)","quality":"Equal","values":[118,129,132,141,143,113],"isMain":false},{"metricName":"# contigs (>= 10000 bp)","quality":"Equal","values":[56,46,52,43,57,38],"isMain":false},{"metricName":"# contigs (>= 25000 bp)","quality":"Equal","values":[11,8,3,8,8,4],"isMain":false},{"metricName":"# contigs (>= 50000 bp)","quality":"Equal","values":[1,1,0,0,0,0],"isMain":false},{"metricName":"Largest contig","quality":"More is better","values":[54480,107760,32565,48336,44425,37749],"isMain":true},{"metricName":"Total length","quality":"More is better","values":[1964840,2177894,2100026,2182464,2213745,1785228],"isMain":true},{"metricName":"Total length (>= 1000 bp)","quality":"More is better","values":[1894690,2057755,1976278,2054166,2074653,1736394],"isMain":true},{"metricName":"Total length (>= 5000 bp)","quality":"More is better","values":[1523623,1506901,1380911,1460046,1546520,1180442],"isMain":false},{"metricName":"Total length (>= 10000 bp)","quality":"More is better","values":[1077107,893923,782592,766901,945626,650127],"isMain":true},{"metricName":"Total length (>= 25000 bp)","quality":"More is better","values":[383308,324041,82914,259069,257550,136930],"isMain":false},{"metricName":"Total length (>= 50000 bp)","quality":"More is better","values":[54480,107760,0,0,0,0],"isMain":true},{"metricName":"N50","quality":"More is better","values":[11081,8875,8090,7542,8053,7561],"isMain":false},{"metricName":"N75","quality":"More is better","values":[5631,4064,4025,3961,3971,3766],"isMain":false},{"metricName":"L50","quality":"Less is better","values":[48,67,82,81,76,67],"isMain":false},{"metricName":"L75","quality":"Less is better","values":[109,158,176,181,169,150],"isMain":false},{"metricName":"GC (%)","quality":"Equal","values":["37.90","37.97","37.96","38.02","38.04","37.73"],"isMain":false}]],["K-mer-based statistics",[]],["Predicted genes",[]],["Similarity statistics",[]],["Reference statistics",[{"metricName":"Reference length","quality":"Equal","values":[2097766,2097766,2097766,2097766,2097766,2097766],"isMain":false},{"metricName":"Reference fragments","quality":"Equal","values":[3,3,3,3,3,3],"isMain":false},{"metricName":"Reference GC (%)","quality":"Equal","values":["38.13","38.13","38.13","38.13","38.13","38.13"],"isMain":false}]]],"subreferences":[],"subreports":[],"minContig":500},{"date":"10 February 2020, Monday, 18:51:33","assembliesNames":["megahit-103-df","megahit-113-df","megahit-113-ml","megahit-113-ms","megahit-129-df","metaSPAdes"],"referenceName":"GCF\_000191805.1\_Lach\_bact\_4\_1\_37FAA\_V1","order":[0,1,2,3,4,5],"report":[["Genome statistics",[{"metricName":"Genome fraction (%)","quality":"More is better","values":["0.343","0.631","0.666","0.767","0.862","0.202"],"isMain":true},{"metricName":"Duplication ratio","quality":"Less is better","values":["1.075","1.087","1.070","1.054","1.107","1.011"],"isMain":true},{"metricName":"Largest alignment","quality":"More is better","values":[629,1390,990,728,4290,633],"isMain":true},{"metricName":"Total aligned length","quality":"More is better","values":[10498,19296,20360,23465,26539,6185],"isMain":true},{"metricName":"NA50","quality":"More is better","values":[540,557,539,530,556,554],"isMain":false},{"metricName":"NA75","quality":"More is better","values":[504,489,513,413,272,516],"isMain":false},{"metricName":"NGA50","quality":"More is better","values":[null,null,null,null,null,null],"isMain":true},{"metricName":"LA50","quality":"Less is better","values":[11,17,18,25,20,6],"isMain":false},{"metricName":"LA75","quality":"Less is better","values":[17,28,28,40,38,9],"isMain":false}]],["Reads mapping",[]],["Misassemblies",[{"metricName":"# misassemblies","quality":"Less is better","values":[0,0,0,0,2,0],"isMain":true},{"metricName":" # relocations","quality":"Less is better","values":[0,0,0,0,1,0],"isMain":false},{"metricName":" # translocations","quality":"Less is better","values":[0,0,0,0,1,0],"isMain":false},{"metricName":" # inversions","quality":"Less is better","values":[0,0,0,0,0,0],"isMain":false},{"metricName":" # interspecies translocations","quality":"Less is better","values":[4,8,3,5,10,0],"isMain":false},{"metricName":"# misassembled contigs","quality":"Less is better","values":[0,0,0,0,1,0],"isMain":false},{"metricName":"Misassembled contigs length","quality":"Less is better","values":[0,0,0,0,6907,0],"isMain":true},{"metricName":"# possibly misassembled contigs","quality":"Less is better","values":[0,0,0,0,0,0],"isMain":false},{"metricName":" # possible misassemblies","quality":"Less is better","values":[0,0,0,0,0,0],"isMain":false},{"metricName":"# local misassemblies","quality":"Less is better","values":[0,0,0,0,2,0],"isMain":false},{"metricName":"# scaffold gap ext. mis.","quality":"Less is better","values":[0,0,0,0,0,0],"isMain":false},{"metricName":"# scaffold gap loc. mis.","quality":"Less is better","values":[0,0,0,0,0,0],"isMain":false},{"metricName":"# unaligned mis. contigs","quality":"Less is better","values":[0,1,0,0,0,0],"isMain":false}]],["Unaligned",[{"metricName":"# fully unaligned contigs","quality":"Less is better","values":[0,0,0,0,0,0],"isMain":false},{"metricName":"Fully unaligned length","quality":"Less is better","values":[0,0,0,0,0,0],"isMain":false},{"metricName":"# partially unaligned contigs","quality":"Less is better","values":[1,3,0,2,3,0],"isMain":false},{"metricName":"Partially unaligned length","quality":"Less is better","values":[861,2398,0,5193,4933,0],"isMain":false}]],["Mismatches",[{"metricName":"# mismatches","quality":"Less is better","values":[210,387,518,571,615,107],"isMain":false},{"metricName":"# indels","quality":"Less is better","values":[0,2,2,1,4,0],"isMain":false},{"metricName":"Indels length","quality":"Less is better","values":[0,2,2,1,4,0],"isMain":false},{"metricName":"# mismatches per 100 kbp","quality":"Less is better","values":["2000.38","2005.60","2544.45","2433.41","2334.59","1729.99"],"isMain":true},{"metricName":"# indels per 100 kbp","quality":"Less is better","values":["0.00","10.36","9.82","4.26","15.18","0.00"],"isMain":true},{"metricName":" # indels (<= 5 bp)","quality":"Less is better","values":[0,2,2,1,4,0],"isMain":false},{"metricName":" # indels (> 5 bp)","quality":"Less is better","values":[0,0,0,0,0,0],"isMain":false},{"metricName":"# N's","quality":"Less is better","values":[0,0,0,0,0,0],"isMain":false},{"metricName":"# N's per 100 kbp","quality":"Less is better","values":["0.00","0.00","0.00","0.00","0.00","0.00"],"isMain":true}]],["Statistics without reference",[{"metricName":"# contigs","quality":"Equal","values":[21,35,37,43,41,11],"isMain":true},{"metricName":"# contigs (>= 1000 bp)","quality":"Equal","values":[1,4,1,2,4,0],"isMain":false},{"metricName":"# contigs (>= 5000 bp)","quality":"Equal","values":[0,0,0,0,1,0],"isMain":false},{"metricName":"# contigs (>= 10000 bp)","quality":"Equal","values":[0,0,0,0,0,0],"isMain":false},{"metricName":"# contigs (>= 25000 bp)","quality":"Equal","values":[0,0,0,0,0,0],"isMain":false},{"metricName":"# contigs (>= 50000 bp)","quality":"Equal","values":[0,0,0,0,0,0],"isMain":false},{"metricName":"Largest contig","quality":"More is better","values":[1032,1390,1013,3316,6907,633],"isMain":true},{"metricName":"Total length","quality":"More is better","values":[12149,23368,21782,29933,34107,6250],"isMain":true},{"metricName":"Total length (>= 1000 bp)","quality":"More is better","values":[1032,4709,1013,5885,12127,0],"isMain":true},{"metricName":"Total length (>= 5000 bp)","quality":"More is better","values":[0,0,0,0,6907,0],"isMain":false},{"metricName":"Total length (>= 10000 bp)","quality":"More is better","values":[0,0,0,0,0,0],"isMain":true},{"metricName":"Total length (>= 25000 bp)","quality":"More is better","values":[0,0,0,0,0,0],"isMain":false},{"metricName":"Total length (>= 50000 bp)","quality":"More is better","values":[0,0,0,0,0,0],"isMain":true},{"metricName":"N50","quality":"More is better","values":[554,644,558,628,657,556],"isMain":false},{"metricName":"N75","quality":"More is better","values":[523,557,525,532,550,531],"isMain":false},{"metricName":"L50","quality":"Less is better","values":[10,14,17,16,11,6],"isMain":false},{"metricName":"L75","quality":"Less is better","values":[16,24,27,29,25,8],"isMain":false},{"metricName":"GC (%)","quality":"Equal","values":["42.88","42.15","42.25","40.32","41.01","42.56"],"isMain":false}]],["K-mer-based statistics",[]],["Predicted genes",[]],["Similarity statistics",[]],["Reference statistics",[{"metricName":"Reference length","quality":"Equal","values":[3057754,3057754,3057754,3057754,3057754,3057754],"isMain":false},{"metricName":"Reference fragments","quality":"Equal","values":[40,40,40,40,40,40],"isMain":false},{"metricName":"Reference GC (%)","quality":"Equal","values":["41.05","41.05","41.05","41.05","41.05","41.05"],"isMain":false}]]],"subreferences":[],"subreports":[],"minContig":500},{"date":"10 February 2020, Monday, 18:52:19","assembliesNames":["megahit-103-df","megahit-113-df","megahit-113-ml","megahit-113-ms","megahit-129-df","metaSPAdes"],"referenceName":"GCF\_000195315.1\_ASM19531v1","order":[0,1,2,3,4,5],"report":[["Genome statistics",[{"metricName":"Genome fraction (%)","quality":"More is better","values":["99.615","99.607","99.590","99.624","99.615","99.497"],"isMain":true},{"metricName":"Duplication ratio","quality":"Less is better","values":["1.004","1.003","1.018","1.011","1.004","1.001"],"isMain":true},{"metricName":"Largest alignment","quality":"More is better","values":[180889,296624,71702,75893,521347,292476],"isMain":true},{"metricName":"Total aligned length","quality":"More is better","values":[2115613,2113077,2143566,2128471,2115658,2106474],"isMain":true},{"metricName":"NG50","quality":"More is better","values":[68270,142261,24746,40376,107710,190645],"isMain":false},{"metricName":"NG75","quality":"More is better","values":[48748,61478,12095,23546,57636,100483],"isMain":false},{"metricName":"NA50","quality":"More is better","values":[68270,142255,24720,37554,107708,190643],"isMain":false},{"metricName":"NA75","quality":"More is better","values":[48748,61478,11451,20506,51969,120580],"isMain":false},{"metricName":"NGA50","quality":"More is better","values":[68270,142255,24746,37554,107708,190643],"isMain":true},{"metricName":"NGA75","quality":"More is better","values":[48748,61478,11965,23546,57618,100472],"isMain":false},{"metricName":"LG50","quality":"Less is better","values":[10,6,29,21,5,5],"isMain":false},{"metricName":"LG75","quality":"Less is better","values":[19,12,59,39,11,9],"isMain":false},{"metricName":"LA50","quality":"Less is better","values":[10,6,30,22,5,5],"isMain":false},{"metricName":"LA75","quality":"Less is better","values":[19,12,62,40,12,8],"isMain":false},{"metricName":"LGA50","quality":"Less is better","values":[10,6,29,22,5,5],"isMain":true},{"metricName":"LGA75","quality":"Less is better","values":[19,12,60,39,11,9],"isMain":false}]],["Reads mapping",[]],["Misassemblies",[{"metricName":"# misassemblies","quality":"Less is better","values":[0,1,2,2,0,0],"isMain":true},{"metricName":" # relocations","quality":"Less is better","values":[0,1,2,2,0,0],"isMain":false},{"metricName":" # translocations","quality":"Less is better","values":[0,0,0,0,0,0],"isMain":false},{"metricName":" # inversions","quality":"Less is better","values":[0,0,0,0,0,0],"isMain":false},{"metricName":" # interspecies translocations","quality":"Less is better","values":[1,0,4,3,3,0],"isMain":false},{"metricName":"# misassembled contigs","quality":"Less is better","values":[0,1,2,2,0,0],"isMain":false},{"metricName":"Misassembled contigs length","quality":"Less is better","values":[0,10045,34901,77748,0,0],"isMain":true},{"metricName":"# possibly misassembled contigs","quality":"Less is better","values":[0,0,0,0,0,0],"isMain":false},{"metricName":" # possible misassemblies","quality":"Less is better","values":[0,0,0,0,0,0],"isMain":false},{"metricName":"# local misassemblies","quality":"Less is better","values":[0,0,0,0,1,0],"isMain":false},{"metricName":"# scaffold gap ext. mis.","quality":"Less is better","values":[0,0,0,0,0,0],"isMain":false},{"metricName":"# scaffold gap loc. mis.","quality":"Less is better","values":[0,0,0,0,0,0],"isMain":false},{"metricName":"# unaligned mis. contigs","quality":"Less is better","values":[0,0,0,0,0,0],"isMain":false}]],["Unaligned",[{"metricName":"# fully unaligned contigs","quality":"Less is better","values":[0,0,0,1,0,0],"isMain":false},{"metricName":"Fully unaligned length","quality":"Less is better","values":[0,0,0,681,0,0],"isMain":false},{"metricName":"# partially unaligned contigs","quality":"Less is better","values":[1,0,1,1,2,0],"isMain":false},{"metricName":"Partially unaligned length","quality":"Less is better","values":[1547,0,1118,1488,16532,0],"isMain":false}]],["Mismatches",[{"metricName":"# mismatches","quality":"Less is better","values":[816,638,4380,3598,596,138],"isMain":false},{"metricName":"# indels","quality":"Less is better","values":[5,10,13,11,18,8],"isMain":false},{"metricName":"Indels length","quality":"Less is better","values":[5,31,68,77,124,15],"isMain":false},{"metricName":"# mismatches per 100 kbp","quality":"Less is better","values":["38.72","30.27","207.88","170.70","28.28","6.56"],"isMain":true},{"metricName":"# indels per 100 kbp","quality":"Less is better","values":["0.24","0.47","0.62","0.52","0.85","0.38"],"isMain":true},{"metricName":" # indels (<= 5 bp)","quality":"Less is better","values":[5,8,10,7,13,8],"isMain":false},{"metricName":" # indels (> 5 bp)","quality":"Less is better","values":[0,2,3,4,5,0],"isMain":false},{"metricName":"# N's","quality":"Less is better","values":[0,0,0,0,0,0],"isMain":false},{"metricName":"# N's per 100 kbp","quality":"Less is better","values":["0.00","0.00","0.00","0.00","0.00","0.00"],"isMain":true}]],["Statistics without reference",[{"metricName":"# contigs","quality":"Equal","values":[69,37,173,109,45,23],"isMain":true},{"metricName":"# contigs (>= 1000 bp)","quality":"Equal","values":[61,34,161,101,40,21],"isMain":false},{"metricName":"# contigs (>= 5000 bp)","quality":"Equal","values":[44,29,110,77,31,17],"isMain":false},{"metricName":"# contigs (>= 10000 bp)","quality":"Equal","values":[37,27,73,61,28,16],"isMain":false},{"metricName":"# contigs (>= 25000 bp)","quality":"Equal","values":[27,19,28,37,23,14],"isMain":false},{"metricName":"# contigs (>= 50000 bp)","quality":"Equal","values":[18,15,3,8,12,13],"isMain":false},{"metricName":"Largest contig","quality":"More is better","values":[180889,296627,71702,75902,521347,292482],"isMain":true},{"metricName":"Total length","quality":"More is better","values":[2117330,2113223,2146863,2132068,2132896,2106524],"isMain":true},{"metricName":"Total length (>= 1000 bp)","quality":"More is better","values":[2111888,2111061,2139196,2126388,2129716,2105287],"isMain":true},{"metricName":"Total length (>= 5000 bp)","quality":"More is better","values":[2074671,2098591,1998242,2053124,2107789,2095882],"isMain":false},{"metricName":"Total length (>= 10000 bp)","quality":"More is better","values":[2023269,2083767,1742366,1935844,2085435,2087300],"isMain":true},{"metricName":"Total length (>= 25000 bp)","quality":"More is better","values":[1860491,1941265,1033524,1559553,1996684,2042946],"isMain":false},{"metricName":"Total length (>= 50000 bp)","quality":"More is better","values":[1554260,1805920,184153,499932,1639972,2006965],"isMain":true},{"metricName":"N50","quality":"More is better","values":[68270,142261,24720,37554,107710,190645],"isMain":false},{"metricName":"N75","quality":"More is better","values":[48748,61478,11657,23546,51993,120587],"isMain":false},{"metricName":"L50","quality":"Less is better","values":[10,6,30,22,5,5],"isMain":false},{"metricName":"L75","quality":"Less is better","values":[19,12,61,39,12,8],"isMain":false},{"metricName":"GC (%)","quality":"Equal","values":["60.34","60.37","60.32","60.34","60.42","60.35"],"isMain":false}]],["K-mer-based statistics",[]],["Predicted genes",[]],["Similarity statistics",[]],["Reference statistics",[{"metricName":"Reference length","quality":"Equal","values":[2115681,2115681,2115681,2115681,2115681,2115681],"isMain":false},{"metricName":"Reference fragments","quality":"Equal","values":[1,1,1,1,1,1],"isMain":false},{"metricName":"Reference GC (%)","quality":"Equal","values":["60.39","60.39","60.39","60.39","60.39","60.39"],"isMain":false}]]],"subreferences":[],"subreports":[],"minContig":500},{"date":"10 February 2020, Monday, 18:52:15","assembliesNames":["megahit-103-df","megahit-113-df","megahit-113-ml","megahit-113-ms","megahit-129-df","metaSPAdes"],"referenceName":"GCF\_000196175.1\_ASM19617v1","order":[0,1,2,3,4,5],"report":[["Genome statistics",[{"metricName":"Genome fraction (%)","quality":"More is better","values":["99.776","99.845","99.735","99.772","99.872","99.805"],"isMain":true},{"metricName":"Duplication ratio","quality":"Less is better","values":["1.009","1.004","1.048","1.042","1.005","1.006"],"isMain":true},{"metricName":"Largest alignment","quality":"More is better","values":[124336,204447,30936,45938,272009,104275],"isMain":true},{"metricName":"Total aligned length","quality":"More is better","values":[3807048,3791347,3948458,3923128,3795454,3794733],"isMain":true},{"metricName":"NG50","quality":"More is better","values":[34984,73487,6682,8695,91542,37357],"isMain":false},{"metricName":"NG75","quality":"More is better","values":[18238,46723,4031,5190,46771,22850],"isMain":false},{"metricName":"NA50","quality":"More is better","values":[34964,73486,6455,8305,91542,37357],"isMain":false},{"metricName":"NA75","quality":"More is better","values":[17980,46713,3655,4690,45953,22844],"isMain":false},{"metricName":"NGA50","quality":"More is better","values":[34964,73486,6628,8672,91542,37357],"isMain":true},{"metricName":"NGA75","quality":"More is better","values":[18238,46713,4026,5099,46767,22844],"isMain":false},{"metricName":"LG50","quality":"Less is better","values":[37,16,173,131,15,31],"isMain":false},{"metricName":"LG75","quality":"Less is better","values":[75,32,356,274,30,63],"isMain":false},{"metricName":"LA50","quality":"Less is better","values":[37,16,187,141,15,31],"isMain":false},{"metricName":"LA75","quality":"Less is better","values":[76,32,391,299,31,63],"isMain":false},{"metricName":"LGA50","quality":"Less is better","values":[37,16,174,132,15,31],"isMain":true},{"metricName":"LGA75","quality":"Less is better","values":[75,32,357,275,30,63],"isMain":false}]],["Reads mapping",[]],["Misassemblies",[{"metricName":"# misassemblies","quality":"Less is better","values":[0,0,2,2,0,0],"isMain":true},{"metricName":" # relocations","quality":"Less is better","values":[0,0,1,0,0,0],"isMain":false},{"metricName":" # translocations","quality":"Less is better","values":[0,0,0,0,0,0],"isMain":false},{"metricName":" # inversions","quality":"Less is better","values":[0,0,1,2,0,0],"isMain":false},{"metricName":" # interspecies translocations","quality":"Less is better","values":[2,4,3,18,6,1],"isMain":false},{"metricName":"# misassembled contigs","quality":"Less is better","values":[0,0,2,2,0,0],"isMain":false},{"metricName":"Misassembled contigs length","quality":"Less is better","values":[0,0,6606,1326,0,0],"isMain":true},{"metricName":"# possibly misassembled contigs","quality":"Less is better","values":[1,1,0,0,0,0],"isMain":false},{"metricName":" # possible misassemblies","quality":"Less is better","values":[1,1,0,0,0,0],"isMain":false},{"metricName":"# local misassemblies","quality":"Less is better","values":[0,3,3,1,1,0],"isMain":false},{"metricName":"# scaffold gap ext. mis.","quality":"Less is better","values":[0,0,0,0,0,0],"isMain":false},{"metricName":"# scaffold gap loc. mis.","quality":"Less is better","values":[0,0,0,0,0,0],"isMain":false},{"metricName":"# unaligned mis. contigs","quality":"Less is better","values":[0,0,0,0,0,0],"isMain":false}]],["Unaligned",[{"metricName":"# fully unaligned contigs","quality":"Less is better","values":[0,0,0,0,0,0],"isMain":false},{"metricName":"Fully unaligned length","quality":"Less is better","values":[0,0,0,0,0,0],"isMain":false},{"metricName":"# partially unaligned contigs","quality":"Less is better","values":[2,4,3,9,5,0],"isMain":false},{"metricName":"Partially unaligned length","quality":"Less is better","values":[1473,5045,4003,8761,13055,0],"isMain":false}]],["Mismatches",[{"metricName":"# mismatches","quality":"Less is better","values":[3067,1787,19211,18081,1641,1467],"isMain":false},{"metricName":"# indels","quality":"Less is better","values":[3,13,15,28,6,27],"isMain":false},{"metricName":"Indels length","quality":"Less is better","values":[3,31,18,41,19,50],"isMain":false},{"metricName":"# mismatches per 100 kbp","quality":"Less is better","values":["81.26","47.31","509.18","479.05","43.43","38.85"],"isMain":true},{"metricName":"# indels per 100 kbp","quality":"Less is better","values":["0.08","0.34","0.40","0.74","0.16","0.72"],"isMain":true},{"metricName":" # indels (<= 5 bp)","quality":"Less is better","values":[3,12,15,28,5,26],"isMain":false},{"metricName":" # indels (> 5 bp)","quality":"Less is better","values":[0,1,0,0,1,1],"isMain":false},{"metricName":"# N's","quality":"Less is better","values":[0,0,0,0,0,0],"isMain":false},{"metricName":"# N's per 100 kbp","quality":"Less is better","values":["0.00","0.00","0.00","0.00","0.00","0.00"],"isMain":true}]],["Statistics without reference",[{"metricName":"# contigs","quality":"Equal","values":[206,88,909,757,95,168],"isMain":true},{"metricName":"# contigs (>= 1000 bp)","quality":"Equal","values":[191,83,797,649,88,162],"isMain":false},{"metricName":"# contigs (>= 5000 bp)","quality":"Equal","values":[152,67,272,282,69,131],"isMain":false},{"metricName":"# contigs (>= 10000 bp)","quality":"Equal","values":[116,66,87,99,62,104],"isMain":false},{"metricName":"# contigs (>= 25000 bp)","quality":"Equal","values":[52,47,1,11,49,54],"isMain":false},{"metricName":"# contigs (>= 50000 bp)","quality":"Equal","values":[15,28,0,0,28,20],"isMain":false},{"metricName":"Largest contig","quality":"More is better","values":[124345,204447,30936,45938,272012,104275],"isMain":true},{"metricName":"Total length","quality":"More is better","values":[3809397,3797426,3957782,3940504,3809293,3796454],"isMain":true},{"metricName":"Total length (>= 1000 bp)","quality":"More is better","values":[3799225,3794203,3874304,3861551,3804352,3792154],"isMain":true},{"metricName":"Total length (>= 5000 bp)","quality":"More is better","values":[3683041,3754481,2464903,2880611,3758138,3714098],"isMain":false},{"metricName":"Total length (>= 10000 bp)","quality":"More is better","values":[3414742,3748406,1189298,1595834,3707517,3511012],"isMain":true},{"metricName":"Total length (>= 25000 bp)","quality":"More is better","values":[2381561,3439074,30936,340016,3490768,2640531],"isMain":false},{"metricName":"Total length (>= 50000 bp)","quality":"More is better","values":[1037331,2678484,0,0,2752323,1427523],"isMain":true},{"metricName":"N50","quality":"More is better","values":[34984,73487,6468,8323,91542,37357],"isMain":false},{"metricName":"N75","quality":"More is better","values":[17984,46723,3713,4717,45984,22850],"isMain":false},{"metricName":"L50","quality":"Less is better","values":[37,16,187,141,15,31],"isMain":false},{"metricName":"L75","quality":"Less is better","values":[76,32,389,298,31,63],"isMain":false},{"metricName":"GC (%)","quality":"Equal","values":["50.64","50.64","50.64","50.63","50.63","50.64"],"isMain":false}]],["K-mer-based statistics",[]],["Predicted genes",[]],["Similarity statistics",[]],["Reference statistics",[{"metricName":"Reference length","quality":"Equal","values":[3782950,3782950,3782950,3782950,3782950,3782950],"isMain":false},{"metricName":"Reference fragments","quality":"Equal","values":[1,1,1,1,1,1],"isMain":false},{"metricName":"Reference GC (%)","quality":"Equal","values":["50.65","50.65","50.65","50.65","50.65","50.65"],"isMain":false}]]],"subreferences":[],"subreports":[],"minContig":500},{"date":"10 February 2020, Monday, 18:43:09","assembliesNames":["megahit-103-df","megahit-113-df","megahit-113-ml","megahit-129-df","metaSPAdes"],"referenceName":"GCF\_000205025.1\_ASM20502v1","order":[0,1,2,3,4],"report":[["Genome statistics",[{"metricName":"Genome fraction (%)","quality":"More is better","values":["0.043","0.043","0.049","0.009","0.039"],"isMain":true},{"metricName":"Duplication ratio","quality":"Less is better","values":["1.354","1.290","1.000","1.000","1.013"],"isMain":true},{"metricName":"Largest alignment","quality":"More is better","values":[519,519,595,262,556],"isMain":true},{"metricName":"Total aligned length","quality":"More is better","values":[1212,1212,1381,262,1100],"isMain":true},{"metricName":"NA50","quality":"More is better","values":[442,442,null,null,544],"isMain":false},{"metricName":"NA75","quality":"More is better","values":[null,251,null,null,544],"isMain":false},{"metricName":"NGA50","quality":"More is better","values":[null,null,null,null,null],"isMain":true},{"metricName":"LA50","quality":"Less is better","values":[2,2,null,null,2],"isMain":false},{"metricName":"LA75","quality":"Less is better","values":[null,3,null,null,2],"isMain":false}]],["Reads mapping",[]],["Misassemblies",[{"metricName":"# misassemblies","quality":"Less is better","values":[0,0,0,0,0],"isMain":true},{"metricName":" # relocations","quality":"Less is better","values":[0,0,0,0,0],"isMain":false},{"metricName":" # translocations","quality":"Less is better","values":[0,0,0,0,0],"isMain":false},{"metricName":" # inversions","quality":"Less is better","values":[0,0,0,0,0],"isMain":false},{"metricName":" # interspecies translocations","quality":"Less is better","values":[2,1,1,1,0],"isMain":false},{"metricName":"# misassembled contigs","quality":"Less is better","values":[0,0,0,0,0],"isMain":false},{"metricName":"Misassembled contigs length","quality":"Less is better","values":[0,0,0,0,0],"isMain":true},{"metricName":"# possibly misassembled contigs","quality":"Less is better","values":[0,0,0,0,0],"isMain":false},{"metricName":" # possible misassemblies","quality":"Less is better","values":[0,0,0,0,0],"isMain":false},{"metricName":"# local misassemblies","quality":"Less is better","values":[0,0,0,0,0],"isMain":false},{"metricName":"# scaffold gap ext. mis.","quality":"Less is better","values":[0,0,0,0,0],"isMain":false},{"metricName":"# scaffold gap loc. mis.","quality":"Less is better","values":[0,0,0,0,0],"isMain":false},{"metricName":"# unaligned mis. contigs","quality":"Less is better","values":[0,0,0,0,0],"isMain":false}]],["Unaligned",[{"metricName":"# fully unaligned contigs","quality":"Less is better","values":[0,0,0,0,0],"isMain":false},{"metricName":"Fully unaligned length","quality":"Less is better","values":[0,0,0,0,0],"isMain":false},{"metricName":"# partially unaligned contigs","quality":"Less is better","values":[0,0,1,1,0],"isMain":false},{"metricName":"Partially unaligned length","quality":"Less is better","values":[0,0,1434,621,0],"isMain":false}]],["Mismatches",[{"metricName":"# mismatches","quality":"Less is better","values":[19,19,34,7,22],"isMain":false},{"metricName":"# indels","quality":"Less is better","values":[0,0,0,0,0],"isMain":false},{"metricName":"Indels length","quality":"Less is better","values":[0,0,0,0,0],"isMain":false},{"metricName":"# mismatches per 100 kbp","quality":"Less is better","values":["1567.66","1567.66","2461.98","2671.76","2000.00"],"isMain":true},{"metricName":"# indels per 100 kbp","quality":"Less is better","values":["0.00","0.00","0.00","0.00","0.00"],"isMain":true},{"metricName":" # indels (<= 5 bp)","quality":"Less is better","values":[0,0,0,0,0],"isMain":false},{"metricName":" # indels (> 5 bp)","quality":"Less is better","values":[0,0,0,0,0],"isMain":false},{"metricName":"# N's","quality":"Less is better","values":[0,0,0,0,0],"isMain":false},{"metricName":"# N's per 100 kbp","quality":"Less is better","values":["0.00","0.00","0.00","0.00","0.00"],"isMain":true}]],["Statistics without reference",[{"metricName":"# contigs","quality":"Equal","values":[3,3,3,1,2],"isMain":true},{"metricName":"# contigs (>= 1000 bp)","quality":"Equal","values":[0,0,1,0,0],"isMain":false},{"metricName":"# contigs (>= 5000 bp)","quality":"Equal","values":[0,0,0,0,0],"isMain":false},{"metricName":"# contigs (>= 10000 bp)","quality":"Equal","values":[0,0,0,0,0],"isMain":false},{"metricName":"# contigs (>= 25000 bp)","quality":"Equal","values":[0,0,0,0,0],"isMain":false},{"metricName":"# contigs (>= 50000 bp)","quality":"Equal","values":[0,0,0,0,0],"isMain":false},{"metricName":"Largest contig","quality":"More is better","values":[564,534,1719,883,570],"isMain":true},{"metricName":"Total length","quality":"More is better","values":[1641,1563,2815,883,1114],"isMain":true},{"metricName":"Total length (>= 1000 bp)","quality":"More is better","values":[0,0,1719,0,0],"isMain":true},{"metricName":"Total length (>= 5000 bp)","quality":"More is better","values":[0,0,0,0,0],"isMain":false},{"metricName":"Total length (>= 10000 bp)","quality":"More is better","values":[0,0,0,0,0],"isMain":true},{"metricName":"Total length (>= 25000 bp)","quality":"More is better","values":[0,0,0,0,0],"isMain":false},{"metricName":"Total length (>= 50000 bp)","quality":"More is better","values":[0,0,0,0,0],"isMain":true},{"metricName":"N50","quality":"More is better","values":[558,519,1719,883,570],"isMain":false},{"metricName":"N75","quality":"More is better","values":[519,510,595,883,544],"isMain":false},{"metricName":"L50","quality":"Less is better","values":[2,2,1,1,1],"isMain":false},{"metricName":"L75","quality":"Less is better","values":[3,3,2,1,2],"isMain":false},{"metricName":"GC (%)","quality":"Equal","values":["43.69","43.63","50.91","40.32","47.58"],"isMain":false}]],["K-mer-based statistics",[]],["Predicted genes",[]],["Similarity statistics",[]],["Reference statistics",[{"metricName":"Reference length","quality":"Equal","values":[2831540,2831540,2831540,2831540,2831540],"isMain":false},{"metricName":"Reference fragments","quality":"Equal","values":[103,103,103,103,103],"isMain":false},{"metricName":"Reference GC (%)","quality":"Equal","values":["48.14","48.14","48.14","48.14","48.14"],"isMain":false}]]],"subreferences":[],"subreports":[],"minContig":500},{"date":"10 February 2020, Monday, 18:51:39","assembliesNames":["megahit-103-df","megahit-113-df","megahit-113-ml","megahit-113-ms","megahit-129-df","metaSPAdes"],"referenceName":"GCF\_000205165.1\_ASM20516v1","order":[0,1,2,3,4,5],"report":[["Genome statistics",[{"metricName":"Genome fraction (%)","quality":"More is better","values":["0.071","0.239","0.197","0.274","0.247","0.078"],"isMain":true},{"metricName":"Duplication ratio","quality":"Less is better","values":["1.040","1.270","1.132","1.084","1.193","1.206"],"isMain":true},{"metricName":"Largest alignment","quality":"More is better","values":[643,682,659,763,1001,708],"isMain":true},{"metricName":"Total aligned length","quality":"More is better","values":[2701,9040,7476,10397,9741,2941],"isMain":true},{"metricName":"NA50","quality":"More is better","values":[541,334,535,517,495,523],"isMain":false},{"metricName":"NA75","quality":"More is better","values":[498,null,505,132,null,418],"isMain":false},{"metricName":"NGA50","quality":"More is better","values":[null,null,null,null,null,null],"isMain":true},{"metricName":"LA50","quality":"Less is better","values":[3,14,8,12,13,3],"isMain":false},{"metricName":"LA75","quality":"Less is better","values":[4,null,12,21,null,5],"isMain":false}]],["Reads mapping",[]],["Misassemblies",[{"metricName":"# misassemblies","quality":"Less is better","values":[0,0,0,0,0,0],"isMain":true},{"metricName":" # relocations","quality":"Less is better","values":[0,0,0,0,0,0],"isMain":false},{"metricName":" # translocations","quality":"Less is better","values":[0,0,0,0,0,0],"isMain":false},{"metricName":" # inversions","quality":"Less is better","values":[0,0,0,0,0,0],"isMain":false},{"metricName":" # interspecies translocations","quality":"Less is better","values":[0,9,2,5,7,1],"isMain":false},{"metricName":"# misassembled contigs","quality":"Less is better","values":[0,0,0,0,0,0],"isMain":false},{"metricName":"Misassembled contigs length","quality":"Less is better","values":[0,0,0,0,0,0],"isMain":true},{"metricName":"# possibly misassembled contigs","quality":"Less is better","values":[0,0,0,0,0,0],"isMain":false},{"metricName":" # possible misassemblies","quality":"Less is better","values":[0,0,0,0,0,0],"isMain":false},{"metricName":"# local misassemblies","quality":"Less is better","values":[0,0,0,0,0,0],"isMain":false},{"metricName":"# scaffold gap ext. mis.","quality":"Less is better","values":[0,0,0,0,0,0],"isMain":false},{"metricName":"# scaffold gap loc. mis.","quality":"Less is better","values":[0,0,0,0,0,0],"isMain":false},{"metricName":"# unaligned mis. contigs","quality":"Less is better","values":[0,0,0,0,0,0],"isMain":false}]],["Unaligned",[{"metricName":"# fully unaligned contigs","quality":"Less is better","values":[0,0,0,0,0,0],"isMain":false},{"metricName":"Fully unaligned length","quality":"Less is better","values":[0,0,0,0,0,0],"isMain":false},{"metricName":"# partially unaligned contigs","quality":"Less is better","values":[0,3,0,2,2,0],"isMain":false},{"metricName":"Partially unaligned length","quality":"Less is better","values":[0,2990,0,2567,3086,0],"isMain":false}]],["Mismatches",[{"metricName":"# mismatches","quality":"Less is better","values":[52,198,154,276,186,69],"isMain":false},{"metricName":"# indels","quality":"Less is better","values":[0,0,0,0,1,0],"isMain":false},{"metricName":"Indels length","quality":"Less is better","values":[0,0,0,0,54,0],"isMain":false},{"metricName":"# mismatches per 100 kbp","quality":"Less is better","values":["1925.21","2190.27","2059.93","2654.61","1984.85","2346.14"],"isMain":true},{"metricName":"# indels per 100 kbp","quality":"Less is better","values":["0.00","0.00","0.00","0.00","10.67","0.00"],"isMain":true},{"metricName":" # indels (<= 5 bp)","quality":"Less is better","values":[0,0,0,0,0,0],"isMain":false},{"metricName":" # indels (> 5 bp)","quality":"Less is better","values":[0,0,0,0,1,0],"isMain":false},{"metricName":"# N's","quality":"Less is better","values":[0,0,0,0,0,0],"isMain":false},{"metricName":"# N's per 100 kbp","quality":"Less is better","values":["0.00","0.00","0.00","0.00","0.00","0.00"],"isMain":true}]],["Statistics without reference",[{"metricName":"# contigs","quality":"Equal","values":[5,22,15,21,21,6],"isMain":true},{"metricName":"# contigs (>= 1000 bp)","quality":"Equal","values":[0,2,0,1,3,0],"isMain":false},{"metricName":"# contigs (>= 5000 bp)","quality":"Equal","values":[0,0,0,0,0,0],"isMain":false},{"metricName":"# contigs (>= 10000 bp)","quality":"Equal","values":[0,0,0,0,0,0],"isMain":false},{"metricName":"# contigs (>= 25000 bp)","quality":"Equal","values":[0,0,0,0,0,0],"isMain":false},{"metricName":"# contigs (>= 50000 bp)","quality":"Equal","values":[0,0,0,0,0,0],"isMain":false},{"metricName":"Largest contig","quality":"More is better","values":[643,1288,672,2153,2201,713],"isMain":true},{"metricName":"Total length","quality":"More is better","values":[2809,14469,8460,13836,14267,3547],"isMain":true},{"metricName":"Total length (>= 1000 bp)","quality":"More is better","values":[0,2520,0,2153,4490,0],"isMain":true},{"metricName":"Total length (>= 5000 bp)","quality":"More is better","values":[0,0,0,0,0,0],"isMain":false},{"metricName":"Total length (>= 10000 bp)","quality":"More is better","values":[0,0,0,0,0,0],"isMain":true},{"metricName":"Total length (>= 25000 bp)","quality":"More is better","values":[0,0,0,0,0,0],"isMain":false},{"metricName":"Total length (>= 50000 bp)","quality":"More is better","values":[0,0,0,0,0,0],"isMain":true},{"metricName":"N50","quality":"More is better","values":[541,613,554,600,551,613],"isMain":false},{"metricName":"N75","quality":"More is better","values":[534,546,531,551,519,535],"isMain":false},{"metricName":"L50","quality":"Less is better","values":[3,9,7,9,8,3],"isMain":false},{"metricName":"L75","quality":"Less is better","values":[4,16,11,15,14,5],"isMain":false},{"metricName":"GC (%)","quality":"Equal","values":["48.42","48.05","48.40","45.37","47.04","41.67"],"isMain":false}]],["K-mer-based statistics",[]],["Predicted genes",[]],["Similarity statistics",[]],["Reference statistics",[{"metricName":"Reference length","quality":"Equal","values":[3789431,3789431,3789431,3789431,3789431,3789431],"isMain":false},{"metricName":"Reference fragments","quality":"Equal","values":[99,99,99,99,99,99],"isMain":false},{"metricName":"Reference GC (%)","quality":"Equal","values":["48.52","48.52","48.52","48.52","48.52","48.52"],"isMain":false}]]],"subreferences":[],"subreports":[],"minContig":500},{"date":"10 February 2020, Monday, 18:51:53","assembliesNames":["megahit-103-df","megahit-113-df","megahit-113-ml","megahit-113-ms","megahit-129-df","metaSPAdes"],"referenceName":"GCF\_000209385.2\_Lach\_bact\_2\_1\_46\_FAA\_V2","order":[0,1,2,3,4,5],"report":[["Genome statistics",[{"metricName":"Genome fraction (%)","quality":"More is better","values":["95.860","97.935","97.886","98.054","97.824","97.591"],"isMain":true},{"metricName":"Duplication ratio","quality":"Less is better","values":["1.018","1.012","1.014","1.019","1.015","1.001"],"isMain":true},{"metricName":"Largest alignment","quality":"More is better","values":[35541,141866,141959,141498,142079,251872],"isMain":true},{"metricName":"Total aligned length","quality":"More is better","values":[2154883,2196910,2198275,2211156,2199766,2166436],"isMain":true},{"metricName":"NG50","quality":"More is better","values":[8681,70767,69194,70789,65509,60716],"isMain":false},{"metricName":"NG75","quality":"More is better","values":[4901,48093,50445,54924,32157,35758],"isMain":false},{"metricName":"NA50","quality":"More is better","values":[8709,64200,56202,53789,65176,63710],"isMain":false},{"metricName":"NA75","quality":"More is better","values":[5004,32027,22948,15556,32157,42353],"isMain":false},{"metricName":"NGA50","quality":"More is better","values":[8681,65191,63597,61515,65176,60716],"isMain":true},{"metricName":"NGA75","quality":"More is better","values":[4901,32431,31999,32027,32157,35758],"isMain":false},{"metricName":"LG50","quality":"Less is better","values":[79,12,12,11,11,12],"isMain":false},{"metricName":"LG75","quality":"Less is better","values":[164,21,21,19,22,22],"isMain":false},{"metricName":"LA50","quality":"Less is better","values":[78,13,15,17,11,11],"isMain":false},{"metricName":"LA75","quality":"Less is better","values":[161,24,31,38,22,21],"isMain":false},{"metricName":"LGA50","quality":"Less is better","values":[79,12,13,13,11,12],"isMain":true},{"metricName":"LGA75","quality":"Less is better","values":[164,23,25,25,22,22],"isMain":false}]],["Reads mapping",[]],["Misassemblies",[{"metricName":"# misassemblies","quality":"Less is better","values":[1,1,4,4,3,0],"isMain":true},{"metricName":" # relocations","quality":"Less is better","values":[1,0,0,0,0,0],"isMain":false},{"metricName":" # translocations","quality":"Less is better","values":[0,1,3,3,3,0],"isMain":false},{"metricName":" # inversions","quality":"Less is better","values":[0,0,1,1,0,0],"isMain":false},{"metricName":" # interspecies translocations","quality":"Less is better","values":[57,12,21,33,12,1],"isMain":false},{"metricName":"# misassembled contigs","quality":"Less is better","values":[1,1,4,4,3,0],"isMain":false},{"metricName":"Misassembled contigs length","quality":"Less is better","values":[520,14953,68333,214790,18419,0],"isMain":true},{"metricName":"# possibly misassembled contigs","quality":"Less is better","values":[0,0,0,0,0,0],"isMain":false},{"metricName":" # possible misassemblies","quality":"Less is better","values":[0,0,0,0,0,0],"isMain":false},{"metricName":"# local misassemblies","quality":"Less is better","values":[0,1,3,4,5,0],"isMain":false},{"metricName":"# scaffold gap ext. mis.","quality":"Less is better","values":[0,0,0,0,0,0],"isMain":false},{"metricName":"# scaffold gap loc. mis.","quality":"Less is better","values":[0,0,0,0,0,0],"isMain":false},{"metricName":"# unaligned mis. contigs","quality":"Less is better","values":[0,1,1,3,1,0],"isMain":false}]],["Unaligned",[{"metricName":"# fully unaligned contigs","quality":"Less is better","values":[0,2,3,3,1,0],"isMain":false},{"metricName":"Fully unaligned length","quality":"Less is better","values":[0,2469,1895,3584,600,0],"isMain":false},{"metricName":"# partially unaligned contigs","quality":"Less is better","values":[30,6,12,18,6,1],"isMain":false},{"metricName":"Partially unaligned length","quality":"Less is better","values":[29102,91191,214977,412963,22685,907],"isMain":false}]],["Mismatches",[{"metricName":"# mismatches","quality":"Less is better","values":[3763,1747,2017,2178,1763,975],"isMain":false},{"metricName":"# indels","quality":"Less is better","values":[1,48,83,75,34,18],"isMain":false},{"metricName":"Indels length","quality":"Less is better","values":[2,223,380,268,57,23],"isMain":false},{"metricName":"# mismatches per 100 kbp","quality":"Less is better","values":["176.97","80.42","92.89","100.13","81.25","45.04"],"isMain":true},{"metricName":"# indels per 100 kbp","quality":"Less is better","values":["0.05","2.21","3.82","3.45","1.57","0.83"],"isMain":true},{"metricName":" # indels (<= 5 bp)","quality":"Less is better","values":[1,44,74,71,33,18],"isMain":false},{"metricName":" # indels (> 5 bp)","quality":"Less is better","values":[0,4,9,4,1,0],"isMain":false},{"metricName":"# N's","quality":"Less is better","values":[0,0,0,0,0,0],"isMain":false},{"metricName":"# N's per 100 kbp","quality":"Less is better","values":["0.00","0.00","0.00","0.00","0.00","0.00"],"isMain":true}]],["Statistics without reference",[{"metricName":"# contigs","quality":"Equal","values":[457,117,133,140,130,67],"isMain":true},{"metricName":"# contigs (>= 1000 bp)","quality":"Equal","values":[344,80,84,80,71,65],"isMain":false},{"metricName":"# contigs (>= 5000 bp)","quality":"Equal","values":[161,49,49,53,50,45],"isMain":false},{"metricName":"# contigs (>= 10000 bp)","quality":"Equal","values":[58,43,44,49,45,40],"isMain":false},{"metricName":"# contigs (>= 25000 bp)","quality":"Equal","values":[6,30,32,33,25,30],"isMain":false},{"metricName":"# contigs (>= 50000 bp)","quality":"Equal","values":[0,19,21,20,19,19],"isMain":false},{"metricName":"Largest contig","quality":"More is better","values":[35541,141866,141959,170434,142079,251872],"isMain":true},{"metricName":"Total length","quality":"More is better","values":[2194169,2292770,2417859,2632364,2225982,2167358],"isMain":true},{"metricName":"Total length (>= 1000 bp)","quality":"More is better","values":[2118356,2269493,2387475,2592977,2190160,2165667],"isMain":true},{"metricName":"Total length (>= 5000 bp)","quality":"More is better","values":[1649729,2198587,2311142,2525114,2132129,2125207],"isMain":false},{"metricName":"Total length (>= 10000 bp)","quality":"More is better","values":[913432,2156739,2273458,2498120,2102895,2094775],"isMain":true},{"metricName":"Total length (>= 25000 bp)","quality":"More is better","values":[176566,1950102,2065227,2219135,1769420,1917616],"isMain":false},{"metricName":"Total length (>= 50000 bp)","quality":"More is better","values":[0,1579439,1709068,1744484,1570341,1546971],"isMain":true},{"metricName":"N50","quality":"More is better","values":[8709,70767,65841,65868,65509,63710],"isMain":false},{"metricName":"N75","quality":"More is better","values":[5004,35040,34834,45465,32157,42353],"isMain":false},{"metricName":"L50","quality":"Less is better","values":[78,12,13,14,11,11],"isMain":false},{"metricName":"L75","quality":"Less is better","values":[161,23,24,25,22,21],"isMain":false},{"metricName":"GC (%)","quality":"Equal","values":["38.77","38.94","39.30","39.10","38.79","38.74"],"isMain":false}]],["K-mer-based statistics",[]],["Predicted genes",[]],["Similarity statistics",[]],["Reference statistics",[{"metricName":"Reference length","quality":"Equal","values":[2218229,2218229,2218229,2218229,2218229,2218229],"isMain":false},{"metricName":"Reference fragments","quality":"Equal","values":[9,9,9,9,9,9],"isMain":false},{"metricName":"Reference GC (%)","quality":"Equal","values":["38.77","38.77","38.77","38.77","38.77","38.77"],"isMain":false}]]],"subreferences":[],"subreports":[],"minContig":500},{"date":"10 February 2020, Monday, 18:51:49","assembliesNames":["megahit-103-df","megahit-113-df","megahit-113-ml","megahit-113-ms","megahit-129-df","metaSPAdes"],"referenceName":"GCF\_000209405.1\_Lach\_bact\_3\_1\_46FAA\_V1","order":[0,1,2,3,4,5],"report":[["Genome statistics",[{"metricName":"Genome fraction (%)","quality":"More is better","values":["10.496","12.432","11.356","11.304","11.574","9.835"],"isMain":true},{"metricName":"Duplication ratio","quality":"Less is better","values":["1.018","1.019","1.020","1.025","1.017","1.025"],"isMain":true},{"metricName":"Largest alignment","quality":"More is better","values":[5265,17282,17268,17282,18414,17196],"isMain":true},{"metricName":"Total aligned length","quality":"More is better","values":[326584,386741,354490,354914,360053,306119],"isMain":true},{"metricName":"NA50","quality":"More is better","values":[859,1005,822,848,1005,775],"isMain":false},{"metricName":"NA75","quality":"More is better","values":[640,689,631,628,711,590],"isMain":false},{"metricName":"NGA50","quality":"More is better","values":[null,null,null,null,null,null],"isMain":true},{"metricName":"LA50","quality":"Less is better","values":[135,127,144,130,112,123],"isMain":false},{"metricName":"LA75","quality":"Less is better","values":[252,254,271,256,221,240],"isMain":false}]],["Reads mapping",[]],["Misassemblies",[{"metricName":"# misassemblies","quality":"Less is better","values":[2,6,3,8,7,4],"isMain":true},{"metricName":" # relocations","quality":"Less is better","values":[0,0,0,0,0,0],"isMain":false},{"metricName":" # translocations","quality":"Less is better","values":[1,6,3,8,7,4],"isMain":false},{"metricName":" # inversions","quality":"Less is better","values":[1,0,0,0,0,0],"isMain":false},{"metricName":" # interspecies translocations","quality":"Less is better","values":[13,21,5,12,15,15],"isMain":false},{"metricName":"# misassembled contigs","quality":"Less is better","values":[2,6,3,6,7,4],"isMain":false},{"metricName":"Misassembled contigs length","quality":"Less is better","values":[1859,33286,25351,37808,34507,25412],"isMain":true},{"metricName":"# possibly misassembled contigs","quality":"Less is better","values":[0,0,0,0,0,0],"isMain":false},{"metricName":" # possible misassemblies","quality":"Less is better","values":[0,0,0,0,0,0],"isMain":false},{"metricName":"# local misassemblies","quality":"Less is better","values":[0,0,0,3,2,0],"isMain":false},{"metricName":"# scaffold gap ext. mis.","quality":"Less is better","values":[0,0,0,0,0,0],"isMain":false},{"metricName":"# scaffold gap loc. mis.","quality":"Less is better","values":[0,0,0,0,0,0],"isMain":false},{"metricName":"# unaligned mis. contigs","quality":"Less is better","values":[0,0,0,0,0,0],"isMain":false}]],["Unaligned",[{"metricName":"# fully unaligned contigs","quality":"Less is better","values":[0,0,0,0,1,0],"isMain":false},{"metricName":"Fully unaligned length","quality":"Less is better","values":[0,0,0,0,778,0],"isMain":false},{"metricName":"# partially unaligned contigs","quality":"Less is better","values":[4,6,2,1,5,3],"isMain":false},{"metricName":"Partially unaligned length","quality":"Less is better","values":[9412,22017,2073,504,4989,2573],"isMain":false}]],["Mismatches",[{"metricName":"# mismatches","quality":"Less is better","values":[3916,4188,5732,5874,4003,3806],"isMain":false},{"metricName":"# indels","quality":"Less is better","values":[9,15,6,12,20,6],"isMain":false},{"metricName":"Indels length","quality":"Less is better","values":[26,101,23,55,43,23],"isMain":false},{"metricName":"# mismatches per 100 kbp","quality":"Less is better","values":["1206.48","1089.33","1632.30","1680.33","1118.41","1251.40"],"isMain":true},{"metricName":"# indels per 100 kbp","quality":"Less is better","values":["2.77","3.90","1.71","3.43","5.59","1.97"],"isMain":true},{"metricName":" # indels (<= 5 bp)","quality":"Less is better","values":[8,11,5,9,19,5],"isMain":false},{"metricName":" # indels (> 5 bp)","quality":"Less is better","values":[1,4,1,3,1,1],"isMain":false},{"metricName":"# N's","quality":"Less is better","values":[0,0,0,0,0,0],"isMain":false},{"metricName":"# N's per 100 kbp","quality":"Less is better","values":["0.00","0.00","0.00","0.00","0.00","0.00"],"isMain":true}]],["Statistics without reference",[{"metricName":"# contigs","quality":"Equal","values":[381,382,423,406,357,373],"isMain":true},{"metricName":"# contigs (>= 1000 bp)","quality":"Equal","values":[101,134,76,84,124,63],"isMain":false},{"metricName":"# contigs (>= 5000 bp)","quality":"Equal","values":[2,2,1,1,1,1],"isMain":false},{"metricName":"# contigs (>= 10000 bp)","quality":"Equal","values":[0,2,1,1,1,1],"isMain":false},{"metricName":"# contigs (>= 25000 bp)","quality":"Equal","values":[0,0,0,1,0,0],"isMain":false},{"metricName":"# contigs (>= 50000 bp)","quality":"Equal","values":[0,0,0,0,0,0],"isMain":false},{"metricName":"Largest contig","quality":"More is better","values":[7968,22801,22873,31565,24148,22550],"isMain":true},{"metricName":"Total length","quality":"More is better","values":[339873,413826,360118,358986,369758,314261],"isMain":true},{"metricName":"Total length (>= 1000 bp)","quality":"More is better","values":[147337,239550,122389,142066,205290,107998],"isMain":true},{"metricName":"Total length (>= 5000 bp)","quality":"More is better","values":[13233,39902,22873,31565,24148,22550],"isMain":false},{"metricName":"Total length (>= 10000 bp)","quality":"More is better","values":[0,39902,22873,31565,24148,22550],"isMain":true},{"metricName":"Total length (>= 25000 bp)","quality":"More is better","values":[0,0,0,31565,0,0],"isMain":false},{"metricName":"Total length (>= 50000 bp)","quality":"More is better","values":[0,0,0,0,0,0],"isMain":true},{"metricName":"N50","quality":"More is better","values":[906,1119,843,868,1095,793],"isMain":false},{"metricName":"N75","quality":"More is better","values":[662,749,645,640,772,608],"isMain":false},{"metricName":"L50","quality":"Less is better","values":[125,104,139,125,105,119],"isMain":false},{"metricName":"L75","quality":"Less is better","values":[236,216,263,247,207,231],"isMain":false},{"metricName":"GC (%)","quality":"Equal","values":["42.30","41.93","42.33","42.55","42.07","42.29"],"isMain":false}]],["K-mer-based statistics",[]],["Predicted genes",[]],["Similarity statistics",[]],["Reference statistics",[{"metricName":"Reference length","quality":"Equal","values":[3092383,3092383,3092383,3092383,3092383,3092383],"isMain":false},{"metricName":"Reference fragments","quality":"Equal","values":[83,83,83,83,83,83],"isMain":false},{"metricName":"Reference GC (%)","quality":"Equal","values":["42.03","42.03","42.03","42.03","42.03","42.03"],"isMain":false}]]],"subreferences":[],"subreports":[],"minContig":500},{"date":"10 February 2020, Monday, 18:52:36","assembliesNames":["megahit-103-df","megahit-113-df","megahit-113-ml","megahit-113-ms","megahit-129-df","metaSPAdes"],"referenceName":"GCF\_000209425.1\_Lach\_bact\_6\_1\_63FAA\_V1","order":[0,1,2,3,4,5],"report":[["Genome statistics",[{"metricName":"Genome fraction (%)","quality":"More is better","values":["96.933","97.377","97.705","95.788","96.387","96.338"],"isMain":true},{"metricName":"Duplication ratio","quality":"Less is better","values":["1.005","1.007","1.012","1.013","1.009","1.002"],"isMain":true},{"metricName":"Largest alignment","quality":"More is better","values":[124884,159342,114929,125137,123841,159458],"isMain":true},{"metricName":"Total aligned length","quality":"More is better","values":[2649966,2666138,2689158,2634501,2643221,2626631],"isMain":true},{"metricName":"NG50","quality":"More is better","values":[42225,57085,29256,39158,47170,64043],"isMain":false},{"metricName":"NG75","quality":"More is better","values":[26236,34040,15866,21731,24049,39295],"isMain":false},{"metricName":"NA50","quality":"More is better","values":[42225,53065,28314,31074,44477,64043],"isMain":false},{"metricName":"NA75","quality":"More is better","values":[28080,25898,15166,15175,24049,40823],"isMain":false},{"metricName":"NGA50","quality":"More is better","values":[40289,53630,28314,31960,43932,63234],"isMain":true},{"metricName":"NGA75","quality":"More is better","values":[26209,29832,14861,17670,23795,37136],"isMain":false},{"metricName":"LG50","quality":"Less is better","values":[22,16,28,22,21,14],"isMain":false},{"metricName":"LG75","quality":"Less is better","values":[42,32,58,46,40,27],"isMain":false},{"metricName":"LA50","quality":"Less is better","values":[22,19,29,29,21,14],"isMain":false},{"metricName":"LA75","quality":"Less is better","values":[41,38,60,62,41,27],"isMain":false},{"metricName":"LGA50","quality":"Less is better","values":[23,18,29,27,22,15],"isMain":true},{"metricName":"LGA75","quality":"Less is better","values":[43,36,61,56,42,29],"isMain":false}]],["Reads mapping",[]],["Misassemblies",[{"metricName":"# misassemblies","quality":"Less is better","values":[2,4,4,7,8,3],"isMain":true},{"metricName":" # relocations","quality":"Less is better","values":[0,0,1,0,0,0],"isMain":false},{"metricName":" # translocations","quality":"Less is better","values":[2,4,3,7,8,3],"isMain":false},{"metricName":" # inversions","quality":"Less is better","values":[0,0,0,0,0,0],"isMain":false},{"metricName":" # interspecies translocations","quality":"Less is better","values":[4,13,8,24,11,2],"isMain":false},{"metricName":"# misassembled contigs","quality":"Less is better","values":[2,3,3,6,8,3],"isMain":false},{"metricName":"Misassembled contigs length","quality":"Less is better","values":[185770,137563,92614,225287,296046,443455],"isMain":true},{"metricName":"# possibly misassembled contigs","quality":"Less is better","values":[0,0,0,0,1,0],"isMain":false},{"metricName":" # possible misassemblies","quality":"Less is better","values":[0,0,0,0,1,0],"isMain":false},{"metricName":"# local misassemblies","quality":"Less is better","values":[1,1,1,1,1,0],"isMain":false},{"metricName":"# scaffold gap ext. mis.","quality":"Less is better","values":[0,0,0,0,0,0],"isMain":false},{"metricName":"# scaffold gap loc. mis.","quality":"Less is better","values":[0,0,0,0,0,0],"isMain":false},{"metricName":"# unaligned mis. contigs","quality":"Less is better","values":[0,0,0,2,0,0],"isMain":false}]],["Unaligned",[{"metricName":"# fully unaligned contigs","quality":"Less is better","values":[0,0,0,0,0,0],"isMain":false},{"metricName":"Fully unaligned length","quality":"Less is better","values":[0,0,0,0,0,0],"isMain":false},{"metricName":"# partially unaligned contigs","quality":"Less is better","values":[0,6,2,15,5,1],"isMain":false},{"metricName":"Partially unaligned length","quality":"Less is better","values":[0,140863,24795,207496,30056,1294],"isMain":false}]],["Mismatches",[{"metricName":"# mismatches","quality":"Less is better","values":[1353,1901,3611,4431,2053,493],"isMain":false},{"metricName":"# indels","quality":"Less is better","values":[5,25,26,55,19,16],"isMain":false},{"metricName":"Indels length","quality":"Less is better","values":[66,164,89,267,234,21],"isMain":false},{"metricName":"# mismatches per 100 kbp","quality":"Less is better","values":["51.25","71.68","135.71","169.86","78.21","18.79"],"isMain":true},{"metricName":"# indels per 100 kbp","quality":"Less is better","values":["0.19","0.94","0.98","2.11","0.72","0.61"],"isMain":true},{"metricName":" # indels (<= 5 bp)","quality":"Less is better","values":[4,22,25,50,15,16],"isMain":false},{"metricName":" # indels (> 5 bp)","quality":"Less is better","values":[1,3,1,5,4,0],"isMain":false},{"metricName":"# N's","quality":"Less is better","values":[0,0,0,0,0,0],"isMain":false},{"metricName":"# N's per 100 kbp","quality":"Less is better","values":["0.00","0.00","0.00","0.00","0.00","0.00"],"isMain":true}]],["Statistics without reference",[{"metricName":"# contigs","quality":"Equal","values":[124,129,174,158,127,80],"isMain":true},{"metricName":"# contigs (>= 1000 bp)","quality":"Equal","values":[106,101,148,127,100,75],"isMain":false},{"metricName":"# contigs (>= 5000 bp)","quality":"Equal","values":[83,71,110,103,81,58],"isMain":false},{"metricName":"# contigs (>= 10000 bp)","quality":"Equal","values":[69,60,89,83,68,48],"isMain":false},{"metricName":"# contigs (>= 25000 bp)","quality":"Equal","values":[43,39,36,38,39,33],"isMain":false},{"metricName":"# contigs (>= 50000 bp)","quality":"Equal","values":[14,20,10,13,19,20],"isMain":false},{"metricName":"Largest contig","quality":"More is better","values":[124884,159342,114929,146667,123841,232049],"isMain":true},{"metricName":"Total length","quality":"More is better","values":[2652021,2811018,2717616,2849497,2678039,2628907],"isMain":true},{"metricName":"Total length (>= 1000 bp)","quality":"More is better","values":[2640170,2790369,2699769,2829683,2659830,2625289],"isMain":true},{"metricName":"Total length (>= 5000 bp)","quality":"More is better","values":[2592685,2717338,2602326,2770119,2613304,2585290],"isMain":false},{"metricName":"Total length (>= 10000 bp)","quality":"More is better","values":[2500314,2639713,2439549,2619995,2518726,2510188],"isMain":true},{"metricName":"Total length (>= 25000 bp)","quality":"More is better","values":[2076638,2280751,1598505,1869445,2036220,2249711],"isMain":false},{"metricName":"Total length (>= 50000 bp)","quality":"More is better","values":[1012968,1597328,705528,984722,1306351,1750567],"isMain":true},{"metricName":"N50","quality":"More is better","values":[42669,55260,29256,36864,49127,65884],"isMain":false},{"metricName":"N75","quality":"More is better","values":[28080,31616,15866,20737,28254,42139],"isMain":false},{"metricName":"L50","quality":"Less is better","values":[21,17,28,24,20,13],"isMain":false},{"metricName":"L75","quality":"Less is better","values":[40,34,58,50,38,25],"isMain":false},{"metricName":"GC (%)","quality":"Equal","values":["38.72","39.08","38.78","39.22","38.90","38.73"],"isMain":false}]],["K-mer-based statistics",[]],["Predicted genes",[]],["Similarity statistics",[]],["Reference statistics",[{"metricName":"Reference length","quality":"Equal","values":[2723336,2723336,2723336,2723336,2723336,2723336],"isMain":false},{"metricName":"Reference fragments","quality":"Equal","values":[37,37,37,37,37,37],"isMain":false},{"metricName":"Reference GC (%)","quality":"Equal","values":["38.83","38.83","38.83","38.83","38.83","38.83"],"isMain":false}]]],"subreferences":[],"subreports":[],"minContig":500},{"date":"10 February 2020, Monday, 18:51:49","assembliesNames":["megahit-103-df","megahit-113-df","megahit-113-ml","megahit-113-ms","megahit-129-df","metaSPAdes"],"referenceName":"GCF\_000209465.1\_Lach\_bact\_F0167\_V1","order":[0,1,2,3,4,5],"report":[["Genome statistics",[{"metricName":"Genome fraction (%)","quality":"More is better","values":["2.076","3.088","3.309","3.936","3.150","1.410"],"isMain":true},{"metricName":"Duplication ratio","quality":"Less is better","values":["1.118","1.120","1.032","1.089","1.101","1.087"],"isMain":true},{"metricName":"Largest alignment","quality":"More is better","values":[1145,1174,2741,2440,1373,1962],"isMain":true},{"metricName":"Total aligned length","quality":"More is better","values":[68089,101524,108632,129242,103654,46238],"isMain":true},{"metricName":"NA50","quality":"More is better","values":[529,517,569,555,521,536],"isMain":false},{"metricName":"NA75","quality":"More is better","values":[393,166,523,499,250,459],"isMain":false},{"metricName":"NGA50","quality":"More is better","values":[null,null,null,null,null,null],"isMain":true},{"metricName":"LA50","quality":"Less is better","values":[63,107,82,107,106,45],"isMain":false},{"metricName":"LA75","quality":"Less is better","values":[105,189,134,177,182,73],"isMain":false}]],["Reads mapping",[]],["Misassemblies",[{"metricName":"# misassemblies","quality":"Less is better","values":[3,9,3,2,4,0],"isMain":true},{"metricName":" # relocations","quality":"Less is better","values":[0,1,0,0,0,0],"isMain":false},{"metricName":" # translocations","quality":"Less is better","values":[3,8,3,2,4,0],"isMain":false},{"metricName":" # inversions","quality":"Less is better","values":[0,0,0,0,0,0],"isMain":false},{"metricName":" # interspecies translocations","quality":"Less is better","values":[32,48,9,33,44,18],"isMain":false},{"metricName":"# misassembled contigs","quality":"Less is better","values":[3,9,3,2,4,0],"isMain":false},{"metricName":"Misassembled contigs length","quality":"Less is better","values":[2622,8332,2573,1712,3500,0],"isMain":true},{"metricName":"# possibly misassembled contigs","quality":"Less is better","values":[1,1,0,0,1,0],"isMain":false},{"metricName":" # possible misassemblies","quality":"Less is better","values":[1,1,0,0,1,0],"isMain":false},{"metricName":"# local misassemblies","quality":"Less is better","values":[0,1,0,1,1,0],"isMain":false},{"metricName":"# scaffold gap ext. mis.","quality":"Less is better","values":[0,0,0,0,0,0],"isMain":false},{"metricName":"# scaffold gap loc. mis.","quality":"Less is better","values":[0,0,0,0,0,0],"isMain":false},{"metricName":"# unaligned mis. contigs","quality":"Less is better","values":[0,0,0,0,0,0],"isMain":false}]],["Unaligned",[{"metricName":"# fully unaligned contigs","quality":"Less is better","values":[0,0,1,0,1,0],"isMain":false},{"metricName":"Fully unaligned length","quality":"Less is better","values":[0,0,584,0,508,0],"isMain":false},{"metricName":"# partially unaligned contigs","quality":"Less is better","values":[7,14,1,4,15,7],"isMain":false},{"metricName":"Partially unaligned length","quality":"Less is better","values":[5959,20387,571,4469,21251,5883],"isMain":false}]],["Mismatches",[{"metricName":"# mismatches","quality":"Less is better","values":[1258,1949,2401,2836,2068,921],"isMain":false},{"metricName":"# indels","quality":"Less is better","values":[12,14,14,17,14,4],"isMain":false},{"metricName":"Indels length","quality":"Less is better","values":[15,21,105,21,16,4],"isMain":false},{"metricName":"# mismatches per 100 kbp","quality":"Less is better","values":["1847.50","1924.05","2212.33","2196.63","2001.55","1991.95"],"isMain":true},{"metricName":"# indels per 100 kbp","quality":"Less is better","values":["17.62","13.82","12.90","13.17","13.55","8.65"],"isMain":true},{"metricName":" # indels (<= 5 bp)","quality":"Less is better","values":[12,14,13,17,14,4],"isMain":false},{"metricName":" # indels (> 5 bp)","quality":"Less is better","values":[0,0,1,0,0,0],"isMain":false},{"metricName":"# N's","quality":"Less is better","values":[0,0,0,0,0,0],"isMain":false},{"metricName":"# N's per 100 kbp","quality":"Less is better","values":["0.00","0.00","0.00","0.00","0.00","0.00"],"isMain":true}]],["Statistics without reference",[{"metricName":"# contigs","quality":"Equal","values":[126,188,183,230,190,89],"isMain":true},{"metricName":"# contigs (>= 1000 bp)","quality":"Equal","values":[7,13,1,7,16,4],"isMain":false},{"metricName":"# contigs (>= 5000 bp)","quality":"Equal","values":[0,1,0,0,1,0],"isMain":false},{"metricName":"# contigs (>= 10000 bp)","quality":"Equal","values":[0,1,0,0,1,0],"isMain":false},{"metricName":"# contigs (>= 25000 bp)","quality":"Equal","values":[0,0,0,0,0,0],"isMain":false},{"metricName":"# contigs (>= 50000 bp)","quality":"Equal","values":[0,0,0,0,0,0],"isMain":false},{"metricName":"Largest contig","quality":"More is better","values":[1815,10731,2765,2440,11193,2001],"isMain":true},{"metricName":"Total length","quality":"More is better","values":[82089,133830,113161,145011,135543,56143],"isMain":true},{"metricName":"Total length (>= 1000 bp)","quality":"More is better","values":[8669,25112,2765,10842,29110,6255],"isMain":true},{"metricName":"Total length (>= 5000 bp)","quality":"More is better","values":[0,10731,0,0,11193,0],"isMain":false},{"metricName":"Total length (>= 10000 bp)","quality":"More is better","values":[0,10731,0,0,11193,0],"isMain":true},{"metricName":"Total length (>= 25000 bp)","quality":"More is better","values":[0,0,0,0,0,0],"isMain":false},{"metricName":"Total length (>= 50000 bp)","quality":"More is better","values":[0,0,0,0,0,0],"isMain":true},{"metricName":"N50","quality":"More is better","values":[622,647,589,590,652,590],"isMain":false},{"metricName":"N75","quality":"More is better","values":[554,555,537,543,559,536],"isMain":false},{"metricName":"L50","quality":"Less is better","values":[52,68,78,96,69,37],"isMain":false},{"metricName":"L75","quality":"Less is better","values":[87,124,129,161,126,62],"isMain":false},{"metricName":"GC (%)","quality":"Equal","values":["36.14","36.69","35.82","36.36","36.29","36.01"],"isMain":false}]],["K-mer-based statistics",[]],["Predicted genes",[]],["Similarity statistics",[]],["Reference statistics",[{"metricName":"Reference length","quality":"Equal","values":[3279854,3279854,3279854,3279854,3279854,3279854],"isMain":false},{"metricName":"Reference fragments","quality":"Equal","values":[105,105,105,105,105,105],"isMain":false},{"metricName":"Reference GC (%)","quality":"Equal","values":["35.69","35.69","35.69","35.69","35.69","35.69"],"isMain":false}]]],"subreferences":[],"subreports":[],"minContig":500},{"date":"10 February 2020, Monday, 18:52:43","assembliesNames":["megahit-103-df","megahit-113-df","megahit-113-ml","megahit-113-ms","megahit-129-df","metaSPAdes"],"referenceName":"GCF\_000214785.1\_ASM21478v1","order":[0,1,2,3,4,5],"report":[["Genome statistics",[{"metricName":"Genome fraction (%)","quality":"More is better","values":["50.505","94.440","95.699","96.389","95.492","86.201"],"isMain":true},{"metricName":"Duplication ratio","quality":"Less is better","values":["1.032","1.105","1.111","1.176","1.162","1.001"],"isMain":true},{"metricName":"Largest alignment","quality":"More is better","values":[4427,115333,77047,67517,64252,124344],"isMain":true},{"metricName":"Total aligned length","quality":"More is better","values":[1212876,2429260,2474807,2618383,2578976,2031246],"isMain":true},{"metricName":"NG50","quality":"More is better","values":[531,20438,23979,29488,20456,27763],"isMain":false},{"metricName":"NG75","quality":"More is better","values":[null,9409,10811,18057,9896,9237],"isMain":false},{"metricName":"NA50","quality":"More is better","values":[891,15567,18284,8003,13703,34759],"isMain":false},{"metricName":"NA75","quality":"More is better","values":[649,4648,3477,272,3351,13917],"isMain":false},{"metricName":"NGA50","quality":"More is better","values":[508,18048,20796,15667,18284,27763],"isMain":true},{"metricName":"NGA75","quality":"More is better","values":[null,7797,8636,7844,8253,9237],"isMain":false},{"metricName":"LG50","quality":"Less is better","values":[1269,31,31,25,35,23],"isMain":false},{"metricName":"LG75","quality":"Less is better","values":[null,75,68,50,78,60],"isMain":false},{"metricName":"LA50","quality":"Less is better","values":[476,39,40,94,48,18],"isMain":false},{"metricName":"LA75","quality":"Less is better","values":[886,111,117,916,139,40],"isMain":false},{"metricName":"LGA50","quality":"Less is better","values":[1302,33,33,43,36,23],"isMain":true},{"metricName":"LGA75","quality":"Less is better","values":[null,84,77,98,84,60],"isMain":false}]],["Reads mapping",[]],["Misassemblies",[{"metricName":"# misassemblies","quality":"Less is better","values":[2,20,32,37,51,0],"isMain":true},{"metricName":" # relocations","quality":"Less is better","values":[0,9,21,27,30,0],"isMain":false},{"metricName":" # translocations","quality":"Less is better","values":[0,6,7,4,6,0],"isMain":false},{"metricName":" # inversions","quality":"Less is better","values":[2,5,4,6,15,0],"isMain":false},{"metricName":" # interspecies translocations","quality":"Less is better","values":[60,103,128,229,141,3],"isMain":false},{"metricName":"# misassembled contigs","quality":"Less is better","values":[2,20,32,36,45,0],"isMain":false},{"metricName":"Misassembled contigs length","quality":"Less is better","values":[1014,63624,69248,197043,142375,0],"isMain":true},{"metricName":"# possibly misassembled contigs","quality":"Less is better","values":[0,0,1,2,0,0],"isMain":false},{"metricName":" # possible misassemblies","quality":"Less is better","values":[0,0,1,2,0,0],"isMain":false},{"metricName":"# local misassemblies","quality":"Less is better","values":[0,8,9,15,15,0],"isMain":false},{"metricName":"# scaffold gap ext. mis.","quality":"Less is better","values":[0,0,0,0,0,0],"isMain":false},{"metricName":"# scaffold gap loc. mis.","quality":"Less is better","values":[0,0,0,0,0,0],"isMain":false},{"metricName":"# unaligned mis. contigs","quality":"Less is better","values":[1,4,2,12,6,0],"isMain":false}]],["Unaligned",[{"metricName":"# fully unaligned contigs","quality":"Less is better","values":[0,5,8,11,6,0],"isMain":false},{"metricName":"Fully unaligned length","quality":"Less is better","values":[0,2846,5620,7864,3517,0],"isMain":false},{"metricName":"# partially unaligned contigs","quality":"Less is better","values":[26,40,46,96,44,3],"isMain":false},{"metricName":"Partially unaligned length","quality":"Less is better","values":[24078,114027,131005,796756,132643,6410],"isMain":false}]],["Mismatches",[{"metricName":"# mismatches","quality":"Less is better","values":[2595,10409,13404,20700,15564,3416],"isMain":false},{"metricName":"# indels","quality":"Less is better","values":[0,200,296,459,329,155],"isMain":false},{"metricName":"Indels length","quality":"Less is better","values":[0,640,895,1404,1329,440],"isMain":false},{"metricName":"# mismatches per 100 kbp","quality":"Less is better","values":["218.26","468.20","594.99","912.26","692.36","168.34"],"isMain":true},{"metricName":"# indels per 100 kbp","quality":"Less is better","values":["0.00","9.00","13.14","20.23","14.64","7.64"],"isMain":true},{"metricName":" # indels (<= 5 bp)","quality":"Less is better","values":[0,183,266,417,289,142],"isMain":false},{"metricName":" # indels (> 5 bp)","quality":"Less is better","values":[0,17,30,42,40,13],"isMain":false},{"metricName":"# N's","quality":"Less is better","values":[0,0,0,0,0,0],"isMain":false},{"metricName":"# N's per 100 kbp","quality":"Less is better","values":["0.00","0.00","0.00","0.00","0.00","0.00"],"isMain":true}]],["Statistics without reference",[{"metricName":"# contigs","quality":"Equal","values":[1412,663,747,954,871,149],"isMain":true},{"metricName":"# contigs (>= 1000 bp)","quality":"Equal","values":[372,238,228,272,230,130],"isMain":false},{"metricName":"# contigs (>= 5000 bp)","quality":"Equal","values":[0,111,103,144,123,77],"isMain":false},{"metricName":"# contigs (>= 10000 bp)","quality":"Equal","values":[0,72,73,101,77,59],"isMain":false},{"metricName":"# contigs (>= 25000 bp)","quality":"Equal","values":[0,24,27,32,23,27],"isMain":false},{"metricName":"# contigs (>= 50000 bp)","quality":"Equal","values":[0,5,7,7,5,10],"isMain":false},{"metricName":"Largest contig","quality":"More is better","values":[4427,115333,77350,125146,64272,124344],"isMain":true},{"metricName":"Total length","quality":"More is better","values":[1250711,2572764,2639267,3473029,2747350,2037882],"isMain":true},{"metricName":"Total length (>= 1000 bp)","quality":"More is better","values":[534795,2313078,2320087,3045922,2367433,2024469],"isMain":true},{"metricName":"Total length (>= 5000 bp)","quality":"More is better","values":[0,2024905,2036899,2760467,2099951,1897571],"isMain":false},{"metricName":"Total length (>= 10000 bp)","quality":"More is better","values":[0,1738041,1821546,2434520,1762098,1762420],"isMain":true},{"metricName":"Total length (>= 25000 bp)","quality":"More is better","values":[0,1015169,1099298,1389308,914776,1290062],"isMain":false},{"metricName":"Total length (>= 50000 bp)","quality":"More is better","values":[0,410568,445009,500847,306076,699504],"isMain":true},{"metricName":"N50","quality":"More is better","values":[908,18243,20626,18353,15801,34759],"isMain":false},{"metricName":"N75","quality":"More is better","values":[667,6925,6277,7867,5541,13917],"isMain":false},{"metricName":"L50","quality":"Less is better","values":[467,37,37,49,46,18],"isMain":false},{"metricName":"L75","quality":"Less is better","values":[869,96,93,120,116,40],"isMain":false},{"metricName":"GC (%)","quality":"Equal","values":["37.22","37.23","37.40","38.55","37.28","37.00"],"isMain":false}]],["K-mer-based statistics",[]],["Predicted genes",[]],["Similarity statistics",[]],["Reference statistics",[{"metricName":"Reference length","quality":"Equal","values":[2354088,2354088,2354088,2354088,2354088,2354088],"isMain":false},{"metricName":"Reference fragments","quality":"Equal","values":[3,3,3,3,3,3],"isMain":false},{"metricName":"Reference GC (%)","quality":"Equal","values":["37.36","37.36","37.36","37.36","37.36","37.36"],"isMain":false}]]],"subreferences":[],"subreports":[],"minContig":500},{"date":"10 February 2020, Monday, 18:52:20","assembliesNames":["megahit-103-df","megahit-113-df","megahit-113-ml","megahit-113-ms","megahit-129-df","metaSPAdes"],"referenceName":"GCF\_000218385.1\_Lach\_bact\_1\_4\_56FAA\_V1","order":[0,1,2,3,4,5],"report":[["Genome statistics",[{"metricName":"Genome fraction (%)","quality":"More is better","values":["90.028","90.963","88.052","87.885","92.277","84.678"],"isMain":true},{"metricName":"Duplication ratio","quality":"Less is better","values":["1.056","1.049","1.100","1.105","1.058","1.059"],"isMain":true},{"metricName":"Largest alignment","quality":"More is better","values":[8883,16935,5179,5455,10036,6942],"isMain":true},{"metricName":"Total aligned length","quality":"More is better","values":[3050276,3054092,3101786,3101933,3130922,2869725],"isMain":true},{"metricName":"NG50","quality":"More is better","values":[1684,2145,1167,1251,2241,1277],"isMain":false},{"metricName":"NG75","quality":"More is better","values":[1030,1217,796,839,1312,771],"isMain":false},{"metricName":"NA50","quality":"More is better","values":[1732,2170,1180,1191,2181,1377],"isMain":false},{"metricName":"NA75","quality":"More is better","values":[1079,1250,807,785,1253,893],"isMain":false},{"metricName":"NGA50","quality":"More is better","values":[1658,2096,1153,1186,2162,1259],"isMain":true},{"metricName":"NGA75","quality":"More is better","values":[997,1144,781,780,1228,753],"isMain":false},{"metricName":"LG50","quality":"Less is better","values":[616,475,914,851,457,809],"isMain":false},{"metricName":"LG75","quality":"Less is better","values":[1232,973,1750,1641,927,1624],"isMain":false},{"metricName":"LA50","quality":"Less is better","values":[578,456,882,914,468,691],"isMain":false},{"metricName":"LA75","quality":"Less is better","values":[1145,926,1685,1745,948,1346],"isMain":false},{"metricName":"LGA50","quality":"Less is better","values":[622,485,923,921,477,817],"isMain":true},{"metricName":"LGA75","quality":"Less is better","values":[1253,1003,1776,1760,970,1650],"isMain":false}]],["Reads mapping",[]],["Misassemblies",[{"metricName":"# misassemblies","quality":"Less is better","values":[12,14,15,18,21,5],"isMain":true},{"metricName":" # relocations","quality":"Less is better","values":[0,0,2,3,1,1],"isMain":false},{"metricName":" # translocations","quality":"Less is better","values":[10,12,11,13,17,4],"isMain":false},{"metricName":" # inversions","quality":"Less is better","values":[2,2,2,2,3,0],"isMain":false},{"metricName":" # interspecies translocations","quality":"Less is better","values":[52,61,21,63,42,15],"isMain":false},{"metricName":"# misassembled contigs","quality":"Less is better","values":[11,13,15,18,20,5],"isMain":false},{"metricName":"Misassembled contigs length","quality":"Less is better","values":[23108,30590,24761,28220,50091,11723],"isMain":true},{"metricName":"# possibly misassembled contigs","quality":"Less is better","values":[0,0,1,0,0,0],"isMain":false},{"metricName":" # possible misassemblies","quality":"Less is better","values":[0,0,1,0,0,0],"isMain":false},{"metricName":"# local misassemblies","quality":"Less is better","values":[1,7,4,6,2,2],"isMain":false},{"metricName":"# scaffold gap ext. mis.","quality":"Less is better","values":[0,0,0,0,0,0],"isMain":false},{"metricName":"# scaffold gap loc. mis.","quality":"Less is better","values":[0,0,0,0,0,0],"isMain":false},{"metricName":"# unaligned mis. contigs","quality":"Less is better","values":[0,0,1,0,0,0],"isMain":false}]],["Unaligned",[{"metricName":"# fully unaligned contigs","quality":"Less is better","values":[0,0,2,0,0,0],"isMain":false},{"metricName":"Fully unaligned length","quality":"Less is better","values":[0,0,1968,0,0,0],"isMain":false},{"metricName":"# partially unaligned contigs","quality":"Less is better","values":[8,17,5,10,15,2],"isMain":false},{"metricName":"Partially unaligned length","quality":"Less is better","values":[7659,25239,3765,76512,40237,1795],"isMain":false}]],["Mismatches",[{"metricName":"# mismatches","quality":"Less is better","values":[23055,21826,44900,47022,23148,23617],"isMain":false},{"metricName":"# indels","quality":"Less is better","values":[58,50,77,64,77,43],"isMain":false},{"metricName":"Indels length","quality":"Less is better","values":[92,242,142,76,214,45],"isMain":false},{"metricName":"# mismatches per 100 kbp","quality":"Less is better","values":["793.43","743.41","1579.89","1657.70","777.21","864.11"],"isMain":true},{"metricName":"# indels per 100 kbp","quality":"Less is better","values":["2.00","1.70","2.71","2.26","2.59","1.57"],"isMain":true},{"metricName":" # indels (<= 5 bp)","quality":"Less is better","values":[57,46,75,63,72,43],"isMain":false},{"metricName":" # indels (> 5 bp)","quality":"Less is better","values":[1,4,2,1,5,0],"isMain":false},{"metricName":"# N's","quality":"Less is better","values":[0,0,0,0,0,0],"isMain":false},{"metricName":"# N's per 100 kbp","quality":"Less is better","values":["0.00","0.00","0.00","0.00","0.00","0.00"],"isMain":true}]],["Statistics without reference",[{"metricName":"# contigs","quality":"Equal","values":[2124,1816,2871,2853,1834,2391],"isMain":true},{"metricName":"# contigs (>= 1000 bp)","quality":"Equal","values":[1267,1158,1237,1257,1187,1175],"isMain":false},{"metricName":"# contigs (>= 5000 bp)","quality":"Equal","values":[19,53,2,2,56,3],"isMain":false},{"metricName":"# contigs (>= 10000 bp)","quality":"Equal","values":[0,2,0,1,3,0],"isMain":false},{"metricName":"# contigs (>= 25000 bp)","quality":"Equal","values":[0,0,0,1,0,0],"isMain":false},{"metricName":"# contigs (>= 50000 bp)","quality":"Equal","values":[0,0,0,1,0,0],"isMain":false},{"metricName":"Largest contig","quality":"More is better","values":[8890,16935,5179,67847,22450,6960],"isMain":true},{"metricName":"Total length","quality":"More is better","values":[3077324,3103792,3130989,3211815,3190697,2895361],"isMain":true},{"metricName":"Total length (>= 1000 bp)","quality":"More is better","values":[2456225,2625923,1962644,2070436,2720078,2026653],"isMain":true},{"metricName":"Total length (>= 5000 bp)","quality":"More is better","values":[112131,349027,10187,73306,382644,18959],"isMain":false},{"metricName":"Total length (>= 10000 bp)","quality":"More is better","values":[0,31085,0,67847,42945,0],"isMain":true},{"metricName":"Total length (>= 25000 bp)","quality":"More is better","values":[0,0,0,67847,0,0],"isMain":false},{"metricName":"Total length (>= 50000 bp)","quality":"More is better","values":[0,0,0,67847,0,0],"isMain":true},{"metricName":"N50","quality":"More is better","values":[1758,2211,1188,1254,2265,1397],"isMain":false},{"metricName":"N75","quality":"More is better","values":[1103,1301,828,847,1334,914],"isMain":false},{"metricName":"L50","quality":"Less is better","values":[572,446,873,844,449,684],"isMain":false},{"metricName":"L75","quality":"Less is better","values":[1127,899,1661,1627,906,1327],"isMain":false},{"metricName":"GC (%)","quality":"Equal","values":["45.57","45.60","45.66","45.72","45.60","45.67"],"isMain":false}]],["K-mer-based statistics",[]],["Predicted genes",[]],["Similarity statistics",[]],["Reference statistics",[{"metricName":"Reference length","quality":"Equal","values":[3227612,3227612,3227612,3227612,3227612,3227612],"isMain":false},{"metricName":"Reference fragments","quality":"Equal","values":[46,46,46,46,46,46],"isMain":false},{"metricName":"Reference GC (%)","quality":"Equal","values":["45.46","45.46","45.46","45.46","45.46","45.46"],"isMain":false}]]],"subreferences":[],"subreports":[],"minContig":500},{"date":"10 February 2020, Monday, 18:52:20","assembliesNames":["megahit-103-df","megahit-113-df","megahit-113-ml","megahit-113-ms","megahit-129-df","metaSPAdes"],"referenceName":"GCF\_000218405.2\_Lach\_bact\_3\_1\_57FAA\_CT1\_V1","order":[0,1,2,3,4,5],"report":[["Genome statistics",[{"metricName":"Genome fraction (%)","quality":"More is better","values":["11.260","14.068","13.386","14.282","15.230","8.653"],"isMain":true},{"metricName":"Duplication ratio","quality":"Less is better","values":["1.036","1.045","1.056","1.064","1.040","1.041"],"isMain":true},{"metricName":"Largest alignment","quality":"More is better","values":[19167,18192,23334,30455,26492,14565],"isMain":true},{"metricName":"Total aligned length","quality":"More is better","values":[886648,1116043,1076758,1153074,1206061,684193],"isMain":true},{"metricName":"NA50","quality":"More is better","values":[827,966,745,776,945,744],"isMain":false},{"metricName":"NA75","quality":"More is better","values":[626,632,578,583,612,591],"isMain":false},{"metricName":"NGA50","quality":"More is better","values":[null,null,null,null,null,null],"isMain":true},{"metricName":"LA50","quality":"Less is better","values":[359,358,488,462,390,291],"isMain":false},{"metricName":"LA75","quality":"Less is better","values":[673,743,910,915,826,556],"isMain":false}]],["Reads mapping",[]],["Misassemblies",[{"metricName":"# misassemblies","quality":"Less is better","values":[3,9,10,12,15,3],"isMain":true},{"metricName":" # relocations","quality":"Less is better","values":[0,5,3,7,5,2],"isMain":false},{"metricName":" # translocations","quality":"Less is better","values":[1,3,3,3,4,1],"isMain":false},{"metricName":" # inversions","quality":"Less is better","values":[2,1,4,2,6,0],"isMain":false},{"metricName":" # interspecies translocations","quality":"Less is better","values":[33,48,9,46,44,10],"isMain":false},{"metricName":"# misassembled contigs","quality":"Less is better","values":[3,7,8,8,13,1],"isMain":false},{"metricName":"Misassembled contigs length","quality":"Less is better","values":[2396,54246,11537,93252,93985,22412],"isMain":true},{"metricName":"# possibly misassembled contigs","quality":"Less is better","values":[0,0,0,0,1,0],"isMain":false},{"metricName":" # possible misassemblies","quality":"Less is better","values":[0,0,0,0,1,0],"isMain":false},{"metricName":"# local misassemblies","quality":"Less is better","values":[4,2,3,3,5,1],"isMain":false},{"metricName":"# scaffold gap ext. mis.","quality":"Less is better","values":[0,0,0,0,0,0],"isMain":false},{"metricName":"# scaffold gap loc. mis.","quality":"Less is better","values":[0,0,0,0,0,0],"isMain":false},{"metricName":"# unaligned mis. contigs","quality":"Less is better","values":[0,0,0,2,0,0],"isMain":false}]],["Unaligned",[{"metricName":"# fully unaligned contigs","quality":"Less is better","values":[1,0,2,0,3,0],"isMain":false},{"metricName":"Fully unaligned length","quality":"Less is better","values":[539,0,1470,0,1641,0],"isMain":false},{"metricName":"# partially unaligned contigs","quality":"Less is better","values":[3,11,1,12,12,1],"isMain":false},{"metricName":"Partially unaligned length","quality":"Less is better","values":[2573,50842,11913,39899,96038,7188],"isMain":false}]],["Mismatches",[{"metricName":"# mismatches","quality":"Less is better","values":[11163,14561,19060,21327,15509,8632],"isMain":false},{"metricName":"# indels","quality":"Less is better","values":[27,89,88,71,67,35],"isMain":false},{"metricName":"Indels length","quality":"Less is better","values":[99,248,252,183,353,214],"isMain":false},{"metricName":"# mismatches per 100 kbp","quality":"Less is better","values":["1287.86","1344.51","1849.63","1939.85","1322.79","1295.85"],"isMain":true},{"metricName":"# indels per 100 kbp","quality":"Less is better","values":["3.11","8.22","8.54","6.46","5.71","5.25"],"isMain":true},{"metricName":" # indels (<= 5 bp)","quality":"Less is better","values":[26,80,80,64,61,30],"isMain":false},{"metricName":" # indels (> 5 bp)","quality":"Less is better","values":[1,9,8,7,6,5],"isMain":false},{"metricName":"# N's","quality":"Less is better","values":[0,0,0,0,0,0],"isMain":false},{"metricName":"# N's per 100 kbp","quality":"Less is better","values":["0.00","0.00","0.00","0.00","0.00","0.00"],"isMain":true}]],["Statistics without reference",[{"metricName":"# contigs","quality":"Equal","values":[1062,1159,1381,1375,1231,851],"isMain":true},{"metricName":"# contigs (>= 1000 bp)","quality":"Equal","values":[215,345,211,230,357,128],"isMain":false},{"metricName":"# contigs (>= 5000 bp)","quality":"Equal","values":[2,11,4,5,9,5],"isMain":false},{"metricName":"# contigs (>= 10000 bp)","quality":"Equal","values":[1,6,2,4,6,3],"isMain":false},{"metricName":"# contigs (>= 25000 bp)","quality":"Equal","values":[0,1,0,2,5,0],"isMain":false},{"metricName":"# contigs (>= 50000 bp)","quality":"Equal","values":[0,0,0,1,1,0],"isMain":false},{"metricName":"Largest contig","quality":"More is better","values":[19167,32025,23396,79807,50925,22412],"isMain":true},{"metricName":"Total length","quality":"More is better","values":[900725,1182425,1101817,1209318,1316497,700481],"isMain":true},{"metricName":"Total length (>= 1000 bp)","quality":"More is better","values":[324931,635067,333604,449139,726628,223703],"isMain":true},{"metricName":"Total length (>= 5000 bp)","quality":"More is better","values":[26124,149405,49321,138731,220105,64371],"isMain":false},{"metricName":"Total length (>= 10000 bp)","quality":"More is better","values":[19167,112336,38682,132141,196635,51015],"isMain":true},{"metricName":"Total length (>= 25000 bp)","quality":"More is better","values":[0,32025,0,106580,176511,0],"isMain":false},{"metricName":"Total length (>= 50000 bp)","quality":"More is better","values":[0,0,0,79807,50925,0],"isMain":true},{"metricName":"N50","quality":"More is better","values":[845,1058,768,827,1091,770],"isMain":false},{"metricName":"N75","quality":"More is better","values":[637,693,596,624,716,608],"isMain":false},{"metricName":"L50","quality":"Less is better","values":[352,303,463,403,292,276],"isMain":false},{"metricName":"L75","quality":"Less is better","values":[661,652,873,829,671,534],"isMain":false},{"metricName":"GC (%)","quality":"Equal","values":["43.21","43.88","44.49","44.23","44.18","43.00"],"isMain":false}]],["K-mer-based statistics",[]],["Predicted genes",[]],["Similarity statistics",[]],["Reference statistics",[{"metricName":"Reference length","quality":"Equal","values":[7698054,7698054,7698054,7698054,7698054,7698054],"isMain":false},{"metricName":"Reference fragments","quality":"Equal","values":[23,23,23,23,23,23],"isMain":false},{"metricName":"Reference GC (%)","quality":"Equal","values":["46.76","46.76","46.76","46.76","46.76","46.76"],"isMain":false}]]],"subreferences":[],"subreports":[],"minContig":500},{"date":"10 February 2020, Monday, 18:51:53","assembliesNames":["megahit-103-df","megahit-113-df","megahit-113-ml","megahit-113-ms","megahit-129-df","metaSPAdes"],"referenceName":"GCF\_000218425.1\_Lach\_bact\_5\_1\_57FAA","order":[0,1,2,3,4,5],"report":[["Genome statistics",[{"metricName":"Genome fraction (%)","quality":"More is better","values":["2.015","2.563","2.912","3.525","2.982","1.016"],"isMain":true},{"metricName":"Duplication ratio","quality":"Less is better","values":["1.054","1.045","1.030","1.026","1.051","1.057"],"isMain":true},{"metricName":"Largest alignment","quality":"More is better","values":[874,919,1107,1499,1641,954],"isMain":true},{"metricName":"Total aligned length","quality":"More is better","values":[67239,85741,97413,117648,99705,33891],"isMain":true},{"metricName":"NA50","quality":"More is better","values":[557,563,579,591,568,562],"isMain":false},{"metricName":"NA75","quality":"More is better","values":[512,507,528,520,514,514],"isMain":false},{"metricName":"NGA50","quality":"More is better","values":[null,null,null,null,null,null],"isMain":true},{"metricName":"LA50","quality":"Less is better","values":[57,74,72,89,80,27],"isMain":false},{"metricName":"LA75","quality":"Less is better","values":[92,119,117,145,130,44],"isMain":false}]],["Reads mapping",[]],["Misassemblies",[{"metricName":"# misassemblies","quality":"Less is better","values":[0,1,1,0,1,0],"isMain":true},{"metricName":" # relocations","quality":"Less is better","values":[0,0,0,0,0,0],"isMain":false},{"metricName":" # translocations","quality":"Less is better","values":[0,1,1,0,1,0],"isMain":false},{"metricName":" # inversions","quality":"Less is better","values":[0,0,0,0,0,0],"isMain":false},{"metricName":" # interspecies translocations","quality":"Less is better","values":[17,19,5,9,22,5],"isMain":false},{"metricName":"# misassembled contigs","quality":"Less is better","values":[0,1,1,0,1,0],"isMain":false},{"metricName":"Misassembled contigs length","quality":"Less is better","values":[0,559,772,0,559,0],"isMain":true},{"metricName":"# possibly misassembled contigs","quality":"Less is better","values":[0,0,0,0,0,0],"isMain":false},{"metricName":" # possible misassemblies","quality":"Less is better","values":[0,0,0,0,0,0],"isMain":false},{"metricName":"# local misassemblies","quality":"Less is better","values":[0,0,0,0,0,0],"isMain":false},{"metricName":"# scaffold gap ext. mis.","quality":"Less is better","values":[0,0,0,0,0,0],"isMain":false},{"metricName":"# scaffold gap loc. mis.","quality":"Less is better","values":[0,0,0,0,0,0],"isMain":false},{"metricName":"# unaligned mis. contigs","quality":"Less is better","values":[0,0,0,0,0,0],"isMain":false}]],["Unaligned",[{"metricName":"# fully unaligned contigs","quality":"Less is better","values":[0,0,0,0,0,0],"isMain":false},{"metricName":"Fully unaligned length","quality":"Less is better","values":[0,0,0,0,0,0],"isMain":false},{"metricName":"# partially unaligned contigs","quality":"Less is better","values":[3,7,0,2,7,0],"isMain":false},{"metricName":"Partially unaligned length","quality":"Less is better","values":[3301,7095,0,1730,4478,0],"isMain":false}]],["Mismatches",[{"metricName":"# mismatches","quality":"Less is better","values":[1182,1502,2005,2582,1715,585],"isMain":false},{"metricName":"# indels","quality":"Less is better","values":[0,4,2,6,1,1],"isMain":false},{"metricName":"Indels length","quality":"Less is better","values":[0,4,2,6,1,1],"isMain":false},{"metricName":"# mismatches per 100 kbp","quality":"Less is better","values":["1757.91","1755.84","2063.52","2194.83","1723.62","1726.17"],"isMain":true},{"metricName":"# indels per 100 kbp","quality":"Less is better","values":["0.00","4.68","2.06","5.10","1.01","2.95"],"isMain":true},{"metricName":" # indels (<= 5 bp)","quality":"Less is better","values":[0,4,2,6,1,1],"isMain":false},{"metricName":" # indels (> 5 bp)","quality":"Less is better","values":[0,0,0,0,0,0],"isMain":false},{"metricName":"# N's","quality":"Less is better","values":[0,0,0,0,0,0],"isMain":false},{"metricName":"# N's per 100 kbp","quality":"Less is better","values":["0.00","0.00","0.00","0.00","0.00","0.00"],"isMain":true}]],["Statistics without reference",[{"metricName":"# contigs","quality":"Equal","values":[120,152,163,199,172,59],"isMain":true},{"metricName":"# contigs (>= 1000 bp)","quality":"Equal","values":[3,4,1,2,4,1],"isMain":false},{"metricName":"# contigs (>= 5000 bp)","quality":"Equal","values":[0,0,0,0,0,0],"isMain":false},{"metricName":"# contigs (>= 10000 bp)","quality":"Equal","values":[0,0,0,0,0,0],"isMain":false},{"metricName":"# contigs (>= 25000 bp)","quality":"Equal","values":[0,0,0,0,0,0],"isMain":false},{"metricName":"# contigs (>= 50000 bp)","quality":"Equal","values":[0,0,0,0,0,0],"isMain":false},{"metricName":"Largest contig","quality":"More is better","values":[1615,2252,1118,1521,1766,1069],"isMain":true},{"metricName":"Total length","quality":"More is better","values":[74151,96479,100121,122428,109096,35830],"isMain":true},{"metricName":"Total length (>= 1000 bp)","quality":"More is better","values":[4048,6493,1118,2785,5559,1069],"isMain":true},{"metricName":"Total length (>= 5000 bp)","quality":"More is better","values":[0,0,0,0,0,0],"isMain":false},{"metricName":"Total length (>= 10000 bp)","quality":"More is better","values":[0,0,0,0,0,0],"isMain":true},{"metricName":"Total length (>= 25000 bp)","quality":"More is better","values":[0,0,0,0,0,0],"isMain":false},{"metricName":"Total length (>= 50000 bp)","quality":"More is better","values":[0,0,0,0,0,0],"isMain":true},{"metricName":"N50","quality":"More is better","values":[595,617,598,602,604,585],"isMain":false},{"metricName":"N75","quality":"More is better","values":[539,541,535,538,541,536],"isMain":false},{"metricName":"L50","quality":"Less is better","values":[52,64,70,86,72,26],"isMain":false},{"metricName":"L75","quality":"Less is better","values":[85,106,115,140,120,42],"isMain":false},{"metricName":"GC (%)","quality":"Equal","values":["46.55","45.97","46.49","46.85","46.18","46.15"],"isMain":false}]],["K-mer-based statistics",[]],["Predicted genes",[]],["Similarity statistics",[]],["Reference statistics",[{"metricName":"Reference length","quality":"Equal","values":[3336970,3336970,3336970,3336970,3336970,3336970],"isMain":false},{"metricName":"Reference fragments","quality":"Equal","values":[67,67,67,67,67,67],"isMain":false},{"metricName":"Reference GC (%)","quality":"Equal","values":["46.34","46.34","46.34","46.34","46.34","46.34"],"isMain":false}]]],"subreferences":[],"subreports":[],"minContig":500},{"date":"10 February 2020, Monday, 18:52:02","assembliesNames":["megahit-103-df","megahit-113-df","megahit-113-ml","megahit-113-ms","megahit-129-df","metaSPAdes"],"referenceName":"GCF\_000218445.1\_Lach\_bact\_1\_1\_57FAA\_V1","order":[0,1,2,3,4,5],"report":[["Genome statistics",[{"metricName":"Genome fraction (%)","quality":"More is better","values":["21.777","22.708","22.010","21.889","23.394","17.146"],"isMain":true},{"metricName":"Duplication ratio","quality":"Less is better","values":["1.022","1.021","1.026","1.029","1.019","1.024"],"isMain":true},{"metricName":"Largest alignment","quality":"More is better","values":[3204,3944,2347,2821,3089,2616],"isMain":true},{"metricName":"Total aligned length","quality":"More is better","values":[642095,666726,650687,647837,686587,503669],"isMain":true},{"metricName":"NA50","quality":"More is better","values":[877,1007,843,830,1055,794],"isMain":false},{"metricName":"NA75","quality":"More is better","values":[661,709,645,629,742,615],"isMain":false},{"metricName":"NGA50","quality":"More is better","values":[null,null,null,null,null,null],"isMain":true},{"metricName":"LA50","quality":"Less is better","values":[257,227,286,286,229,233],"isMain":false},{"metricName":"LA75","quality":"Less is better","values":[471,428,511,516,427,419],"isMain":false}]],["Reads mapping",[]],["Misassemblies",[{"metricName":"# misassemblies","quality":"Less is better","values":[4,5,4,7,6,4],"isMain":true},{"metricName":" # relocations","quality":"Less is better","values":[1,0,0,0,0,0],"isMain":false},{"metricName":" # translocations","quality":"Less is better","values":[3,5,4,6,6,4],"isMain":false},{"metricName":" # inversions","quality":"Less is better","values":[0,0,0,1,0,0],"isMain":false},{"metricName":" # interspecies translocations","quality":"Less is better","values":[18,26,6,21,26,11],"isMain":false},{"metricName":"# misassembled contigs","quality":"Less is better","values":[4,5,4,7,6,4],"isMain":false},{"metricName":"Misassembled contigs length","quality":"Less is better","values":[4197,8551,4683,8626,9509,7873],"isMain":true},{"metricName":"# possibly misassembled contigs","quality":"Less is better","values":[0,0,0,0,0,0],"isMain":false},{"metricName":" # possible misassemblies","quality":"Less is better","values":[0,0,0,0,0,0],"isMain":false},{"metricName":"# local misassemblies","quality":"Less is better","values":[2,2,0,0,2,0],"isMain":false},{"metricName":"# scaffold gap ext. mis.","quality":"Less is better","values":[0,0,0,0,0,0],"isMain":false},{"metricName":"# scaffold gap loc. mis.","quality":"Less is better","values":[0,0,0,0,0,0],"isMain":false},{"metricName":"# unaligned mis. contigs","quality":"Less is better","values":[1,1,0,0,0,0],"isMain":false}]],["Unaligned",[{"metricName":"# fully unaligned contigs","quality":"Less is better","values":[0,0,0,3,1,0],"isMain":false},{"metricName":"Fully unaligned length","quality":"Less is better","values":[0,0,0,1987,564,0],"isMain":false},{"metricName":"# partially unaligned contigs","quality":"Less is better","values":[5,5,1,5,5,2],"isMain":false},{"metricName":"Partially unaligned length","quality":"Less is better","values":[6646,6180,7681,4129,4017,2653],"isMain":false}]],["Mismatches",[{"metricName":"# mismatches","quality":"Less is better","values":[7638,7518,11331,11976,7977,6196],"isMain":false},{"metricName":"# indels","quality":"Less is better","values":[17,18,17,19,26,16],"isMain":false},{"metricName":"Indels length","quality":"Less is better","values":[107,64,37,61,94,154],"isMain":false},{"metricName":"# mismatches per 100 kbp","quality":"Less is better","values":["1205.10","1137.53","1768.85","1879.81","1171.55","1241.63"],"isMain":true},{"metricName":"# indels per 100 kbp","quality":"Less is better","values":["2.68","2.72","2.65","2.98","3.82","3.21"],"isMain":true},{"metricName":" # indels (<= 5 bp)","quality":"Less is better","values":[14,17,16,18,24,14],"isMain":false},{"metricName":" # indels (> 5 bp)","quality":"Less is better","values":[3,1,1,1,2,2],"isMain":false},{"metricName":"# N's","quality":"Less is better","values":[0,0,0,0,0,0],"isMain":false},{"metricName":"# N's per 100 kbp","quality":"Less is better","values":["0.00","0.00","0.00","0.00","0.00","0.00"],"isMain":true}]],["Statistics without reference",[{"metricName":"# contigs","quality":"Equal","values":[736,690,781,794,699,632],"isMain":true},{"metricName":"# contigs (>= 1000 bp)","quality":"Equal","values":[193,239,187,173,272,121],"isMain":false},{"metricName":"# contigs (>= 5000 bp)","quality":"Equal","values":[0,0,1,0,0,0],"isMain":false},{"metricName":"# contigs (>= 10000 bp)","quality":"Equal","values":[0,0,0,0,0,0],"isMain":false},{"metricName":"# contigs (>= 25000 bp)","quality":"Equal","values":[0,0,0,0,0,0],"isMain":false},{"metricName":"# contigs (>= 50000 bp)","quality":"Equal","values":[0,0,0,0,0,0],"isMain":false},{"metricName":"Largest contig","quality":"More is better","values":[3204,3944,8500,2821,3089,2880],"isMain":true},{"metricName":"Total length","quality":"More is better","values":[654557,680806,664851,661972,698683,513787],"isMain":true},{"metricName":"Total length (>= 1000 bp)","quality":"More is better","values":[276606,361028,252497,234169,399210,162852],"isMain":true},{"metricName":"Total length (>= 5000 bp)","quality":"More is better","values":[0,0,8500,0,0,0],"isMain":false},{"metricName":"Total length (>= 10000 bp)","quality":"More is better","values":[0,0,0,0,0,0],"isMain":true},{"metricName":"Total length (>= 25000 bp)","quality":"More is better","values":[0,0,0,0,0,0],"isMain":false},{"metricName":"Total length (>= 50000 bp)","quality":"More is better","values":[0,0,0,0,0,0],"isMain":true},{"metricName":"N50","quality":"More is better","values":[895,1043,870,856,1095,820],"isMain":false},{"metricName":"N75","quality":"More is better","values":[680,753,663,653,773,632],"isMain":false},{"metricName":"L50","quality":"Less is better","values":[247,219,273,279,225,226],"isMain":false},{"metricName":"L75","quality":"Less is better","values":[455,412,491,502,415,405],"isMain":false},{"metricName":"GC (%)","quality":"Equal","values":["41.85","41.95","42.10","42.04","41.63","41.90"],"isMain":false}]],["K-mer-based statistics",[]],["Predicted genes",[]],["Similarity statistics",[]],["Reference statistics",[{"metricName":"Reference length","quality":"Equal","values":[2910478,2910478,2910478,2910478,2910478,2910478],"isMain":false},{"metricName":"Reference fragments","quality":"Equal","values":[93,93,93,93,93,93],"isMain":false},{"metricName":"Reference GC (%)","quality":"Equal","values":["41.85","41.85","41.85","41.85","41.85","41.85"],"isMain":false}]]],"subreferences":[],"subreports":[],"minContig":500},{"date":"10 February 2020, Monday, 18:53:06","assembliesNames":["megahit-103-df","megahit-113-df","megahit-113-ml","megahit-113-ms","megahit-129-df","metaSPAdes"],"referenceName":"GCF\_000220255.1\_ASM22025v1","order":[0,1,2,3,4,5],"report":[["Genome statistics",[{"metricName":"Genome fraction (%)","quality":"More is better","values":["87.529","86.985","85.747","84.858","88.777","76.629"],"isMain":true},{"metricName":"Duplication ratio","quality":"Less is better","values":["1.059","1.054","1.103","1.105","1.064","1.055"],"isMain":true},{"metricName":"Largest alignment","quality":"More is better","values":[6604,9264,6693,6466,11530,5697],"isMain":true},{"metricName":"Total aligned length","quality":"More is better","values":[2808096,2772971,2858855,2824568,2856911,2444281],"isMain":true},{"metricName":"NG50","quality":"More is better","values":[1674,2269,1207,1318,2281,1236],"isMain":false},{"metricName":"NG75","quality":"More is better","values":[974,1309,794,856,1323,656],"isMain":false},{"metricName":"NA50","quality":"More is better","values":[1799,1931,1199,1196,2027,1409],"isMain":false},{"metricName":"NA75","quality":"More is better","values":[1087,1032,795,747,1086,902],"isMain":false},{"metricName":"NGA50","quality":"More is better","values":[1644,1910,1166,1191,2027,1179],"isMain":true},{"metricName":"NGA75","quality":"More is better","values":[948,1008,751,739,1088,601],"isMain":false},{"metricName":"LG50","quality":"Less is better","values":[563,371,776,706,375,735],"isMain":false},{"metricName":"LG75","quality":"Less is better","values":[1161,815,1555,1419,813,1562],"isMain":false},{"metricName":"LA50","quality":"Less is better","values":[503,469,767,829,450,574],"isMain":false},{"metricName":"LA75","quality":"Less is better","values":[1011,996,1516,1623,965,1130],"isMain":false},{"metricName":"LGA50","quality":"Less is better","values":[566,476,816,838,450,782],"isMain":true},{"metricName":"LGA75","quality":"Less is better","values":[1173,1017,1630,1644,963,1668],"isMain":false}]],["Reads mapping",[]],["Misassemblies",[{"metricName":"# misassemblies","quality":"Less is better","values":[9,15,18,14,17,8],"isMain":true},{"metricName":" # relocations","quality":"Less is better","values":[3,3,3,3,3,3],"isMain":false},{"metricName":" # translocations","quality":"Less is better","values":[6,11,11,8,12,5],"isMain":false},{"metricName":" # inversions","quality":"Less is better","values":[0,1,4,3,2,0],"isMain":false},{"metricName":" # interspecies translocations","quality":"Less is better","values":[28,58,35,82,56,19],"isMain":false},{"metricName":"# misassembled contigs","quality":"Less is better","values":[9,14,17,14,15,8],"isMain":false},{"metricName":"Misassembled contigs length","quality":"Less is better","values":[12317,34678,25613,17874,22451,10397],"isMain":true},{"metricName":"# possibly misassembled contigs","quality":"Less is better","values":[0,0,0,0,1,0],"isMain":false},{"metricName":" # possible misassemblies","quality":"Less is better","values":[0,0,0,0,1,0],"isMain":false},{"metricName":"# local misassemblies","quality":"Less is better","values":[4,4,2,6,12,2],"isMain":false},{"metricName":"# scaffold gap ext. mis.","quality":"Less is better","values":[0,0,0,0,0,0],"isMain":false},{"metricName":"# scaffold gap loc. mis.","quality":"Less is better","values":[0,0,0,0,0,0],"isMain":false},{"metricName":"# unaligned mis. contigs","quality":"Less is better","values":[0,1,2,2,1,1],"isMain":false}]],["Unaligned",[{"metricName":"# fully unaligned contigs","quality":"Less is better","values":[0,4,3,5,1,0],"isMain":false},{"metricName":"Fully unaligned length","quality":"Less is better","values":[0,4219,3866,42421,614,0],"isMain":false},{"metricName":"# partially unaligned contigs","quality":"Less is better","values":[3,27,14,27,27,9],"isMain":false},{"metricName":"Partially unaligned length","quality":"Less is better","values":[2142,222144,43715,128282,171198,49429],"isMain":false}]],["Mismatches",[{"metricName":"# mismatches","quality":"Less is better","values":[20047,20994,39996,42136,22827,18432],"isMain":false},{"metricName":"# indels","quality":"Less is better","values":[52,91,67,78,104,57],"isMain":false},{"metricName":"Indels length","quality":"Less is better","values":[104,384,438,410,868,165],"isMain":false},{"metricName":"# mismatches per 100 kbp","quality":"Less is better","values":["751.56","791.99","1530.62","1629.40","843.76","789.31"],"isMain":true},{"metricName":"# indels per 100 kbp","quality":"Less is better","values":["1.95","3.43","2.56","3.02","3.84","2.44"],"isMain":true},{"metricName":" # indels (<= 5 bp)","quality":"Less is better","values":[50,83,57,67,82,53],"isMain":false},{"metricName":" # indels (> 5 bp)","quality":"Less is better","values":[2,8,10,11,22,4],"isMain":false},{"metricName":"# N's","quality":"Less is better","values":[0,0,0,0,0,0],"isMain":false},{"metricName":"# N's per 100 kbp","quality":"Less is better","values":["0.00","0.00","0.00","0.00","0.00","0.00"],"isMain":true}]],["Statistics without reference",[{"metricName":"# contigs","quality":"Equal","values":[1919,1701,2566,2548,1740,1958],"isMain":true},{"metricName":"# contigs (>= 1000 bp)","quality":"Equal","values":[1132,1041,1103,1163,1056,1012],"isMain":false},{"metricName":"# contigs (>= 5000 bp)","quality":"Equal","values":[17,57,5,14,68,4],"isMain":false},{"metricName":"# contigs (>= 10000 bp)","quality":"Equal","values":[0,3,1,4,8,1],"isMain":false},{"metricName":"# contigs (>= 25000 bp)","quality":"Equal","values":[0,2,0,2,2,1],"isMain":false},{"metricName":"# contigs (>= 50000 bp)","quality":"Equal","values":[0,1,0,1,0,0],"isMain":false},{"metricName":"Largest contig","quality":"More is better","values":[6604,131657,22202,53167,33835,46797],"isMain":true},{"metricName":"Total length","quality":"More is better","values":[2827734,3019192,2929765,3026975,3050441,2513686],"isMain":true},{"metricName":"Total length (>= 1000 bp)","quality":"More is better","values":[2257608,2544931,1885173,2049642,2563947,1834356],"isMain":true},{"metricName":"Total length (>= 5000 bp)","quality":"More is better","values":[95927,519059,47615,186116,529518,62724],"isMain":false},{"metricName":"Total length (>= 10000 bp)","quality":"More is better","values":[0,181792,22202,126080,153057,46797],"isMain":true},{"metricName":"Total length (>= 25000 bp)","quality":"More is better","values":[0,165874,0,92191,63920,46797],"isMain":false},{"metricName":"Total length (>= 50000 bp)","quality":"More is better","values":[0,131657,0,53167,0,0],"isMain":true},{"metricName":"N50","quality":"More is better","values":[1815,2281,1257,1330,2278,1483],"isMain":false},{"metricName":"N75","quality":"More is better","values":[1102,1337,835,863,1321,960],"isMain":false},{"metricName":"L50","quality":"Less is better","values":[500,365,728,698,376,537],"isMain":false},{"metricName":"L75","quality":"Less is better","values":[1003,799,1446,1401,815,1064],"isMain":false},{"metricName":"GC (%)","quality":"Equal","values":["37.33","37.69","37.60","37.77","37.59","37.18"],"isMain":false}]],["K-mer-based statistics",[]],["Predicted genes",[]],["Similarity statistics",[]],["Reference statistics",[{"metricName":"Reference length","quality":"Equal","values":[3047416,3047416,3047416,3047416,3047416,3047416],"isMain":false},{"metricName":"Reference fragments","quality":"Equal","values":[169,169,169,169,169,169],"isMain":false},{"metricName":"Reference GC (%)","quality":"Equal","values":["37.46","37.46","37.46","37.46","37.46","37.46"],"isMain":false}]]],"subreferences":[],"subreports":[],"minContig":500},{"date":"10 February 2020, Monday, 18:52:34","assembliesNames":["megahit-103-df","megahit-113-df","megahit-113-ml","megahit-113-ms","megahit-129-df","metaSPAdes"],"referenceName":"GCF\_000224615.1\_Prev\_oulo\_F0390\_V1","order":[0,1,2,3,4,5],"report":[["Genome statistics",[{"metricName":"Genome fraction (%)","quality":"More is better","values":["12.042","14.790","15.017","17.604","16.530","7.319"],"isMain":true},{"metricName":"Duplication ratio","quality":"Less is better","values":["1.029","1.023","1.026","1.032","1.036","1.039"],"isMain":true},{"metricName":"Largest alignment","quality":"More is better","values":[1596,2022,1804,1949,2022,1292],"isMain":true},{"metricName":"Total aligned length","quality":"More is better","values":[337196,414864,421449,494805,466402,205296],"isMain":true},{"metricName":"NA50","quality":"More is better","values":[618,630,600,617,630,534],"isMain":false},{"metricName":"NA75","quality":"More is better","values":[538,544,522,536,545,null],"isMain":false},{"metricName":"NGA50","quality":"More is better","values":[null,null,null,null,null,null],"isMain":true},{"metricName":"LA50","quality":"Less is better","values":[225,271,318,334,312,232],"isMain":false},{"metricName":"LA75","quality":"Less is better","values":[376,456,527,559,523,null],"isMain":false}]],["Reads mapping",[]],["Misassemblies",[{"metricName":"# misassemblies","quality":"Less is better","values":[9,12,12,19,10,4],"isMain":true},{"metricName":" # relocations","quality":"Less is better","values":[1,1,0,1,2,0],"isMain":false},{"metricName":" # translocations","quality":"Less is better","values":[8,11,12,17,8,4],"isMain":false},{"metricName":" # inversions","quality":"Less is better","values":[0,0,0,1,0,0],"isMain":false},{"metricName":" # interspecies translocations","quality":"Less is better","values":[30,21,9,27,27,21],"isMain":false},{"metricName":"# misassembled contigs","quality":"Less is better","values":[9,12,11,19,10,4],"isMain":false},{"metricName":"Misassembled contigs length","quality":"Less is better","values":[7023,11071,7923,15963,8028,2590],"isMain":true},{"metricName":"# possibly misassembled contigs","quality":"Less is better","values":[0,0,0,0,0,0],"isMain":false},{"metricName":" # possible misassemblies","quality":"Less is better","values":[0,0,0,0,0,0],"isMain":false},{"metricName":"# local misassemblies","quality":"Less is better","values":[1,2,6,5,5,1],"isMain":false},{"metricName":"# scaffold gap ext. mis.","quality":"Less is better","values":[0,0,0,0,0,0],"isMain":false},{"metricName":"# scaffold gap loc. mis.","quality":"Less is better","values":[0,0,0,0,0,0],"isMain":false},{"metricName":"# unaligned mis. contigs","quality":"Less is better","values":[0,0,0,0,1,1],"isMain":false}]],["Unaligned",[{"metricName":"# fully unaligned contigs","quality":"Less is better","values":[0,1,0,2,0,0],"isMain":false},{"metricName":"Fully unaligned length","quality":"Less is better","values":[0,2384,0,1304,0,0],"isMain":false},{"metricName":"# partially unaligned contigs","quality":"Less is better","values":[2,9,4,8,6,5],"isMain":false},{"metricName":"Partially unaligned length","quality":"Less is better","values":[1677,7391,37624,6191,17494,91237],"isMain":false}]],["Mismatches",[{"metricName":"# mismatches","quality":"Less is better","values":[5380,6418,8478,10425,7419,3591],"isMain":false},{"metricName":"# indels","quality":"Less is better","values":[17,11,19,32,10,11],"isMain":false},{"metricName":"Indels length","quality":"Less is better","values":[184,177,108,152,93,34],"isMain":false},{"metricName":"# mismatches per 100 kbp","quality":"Less is better","values":["1602.26","1556.24","2024.60","2123.73","1609.56","1759.57"],"isMain":true},{"metricName":"# indels per 100 kbp","quality":"Less is better","values":["5.06","2.67","4.54","6.52","2.17","5.39"],"isMain":true},{"metricName":" # indels (<= 5 bp)","quality":"Less is better","values":[14,9,17,29,8,9],"isMain":false},{"metricName":" # indels (> 5 bp)","quality":"Less is better","values":[3,2,2,3,2,2],"isMain":false},{"metricName":"# N's","quality":"Less is better","values":[0,0,0,0,0,0],"isMain":false},{"metricName":"# N's per 100 kbp","quality":"Less is better","values":["0.00","0.00","0.00","0.00","0.00","0.00"],"isMain":true}]],["Statistics without reference",[{"metricName":"# contigs","quality":"Equal","values":[527,635,669,780,712,340],"isMain":true},{"metricName":"# contigs (>= 1000 bp)","quality":"Equal","values":[22,39,26,38,40,12],"isMain":false},{"metricName":"# contigs (>= 5000 bp)","quality":"Equal","values":[0,0,2,0,1,1],"isMain":false},{"metricName":"# contigs (>= 10000 bp)","quality":"Equal","values":[0,0,2,0,1,1],"isMain":false},{"metricName":"# contigs (>= 25000 bp)","quality":"Equal","values":[0,0,1,0,0,1],"isMain":false},{"metricName":"# contigs (>= 50000 bp)","quality":"Equal","values":[0,0,0,0,0,1],"isMain":false},{"metricName":"Largest contig","quality":"More is better","values":[1596,2384,26921,1951,13983,88424],"isMain":true},{"metricName":"Total length","quality":"More is better","values":[347357,431591,467288,514002,495111,303242],"isMain":true},{"metricName":"Total length (>= 1000 bp)","quality":"More is better","values":[26029,49631,64958,44387,61408,101783],"isMain":true},{"metricName":"Total length (>= 5000 bp)","quality":"More is better","values":[0,0,37589,0,13983,88424],"isMain":false},{"metricName":"Total length (>= 10000 bp)","quality":"More is better","values":[0,0,37589,0,13983,88424],"isMain":true},{"metricName":"Total length (>= 25000 bp)","quality":"More is better","values":[0,0,26921,0,0,88424],"isMain":false},{"metricName":"Total length (>= 50000 bp)","quality":"More is better","values":[0,0,0,0,0,88424],"isMain":true},{"metricName":"N50","quality":"More is better","values":[641,658,649,640,671,701],"isMain":false},{"metricName":"N75","quality":"More is better","values":[559,564,559,557,573,566],"isMain":false},{"metricName":"L50","quality":"Less is better","values":[217,255,252,319,280,75],"isMain":false},{"metricName":"L75","quality":"Less is better","values":[363,433,447,536,481,198],"isMain":false},{"metricName":"GC (%)","quality":"Equal","values":["47.21","46.91","46.92","47.15","47.10","46.46"],"isMain":false}]],["K-mer-based statistics",[]],["Predicted genes",[]],["Similarity statistics",[]],["Reference statistics",[{"metricName":"Reference length","quality":"Equal","values":[2788453,2788453,2788453,2788453,2788453,2788453],"isMain":false},{"metricName":"Reference fragments","quality":"Equal","values":[83,83,83,83,83,83],"isMain":false},{"metricName":"Reference GC (%)","quality":"Equal","values":["46.78","46.78","46.78","46.78","46.78","46.78"],"isMain":false}]]],"subreferences":[],"subreports":[],"minContig":500},{"date":"10 February 2020, Monday, 18:52:43","assembliesNames":["megahit-103-df","megahit-113-df","megahit-113-ml","megahit-113-ms","megahit-129-df","metaSPAdes"],"referenceName":"GCF\_000224655.1\_Bilophila\_sp\_4\_1\_30\_V1","order":[0,1,2,3,4,5],"report":[["Genome statistics",[{"metricName":"Genome fraction (%)","quality":"More is better","values":["0.220","1.207","0.804","0.473","1.371","0.139"],"isMain":true},{"metricName":"Duplication ratio","quality":"Less is better","values":["1.160","1.027","1.010","1.029","1.012","1.052"],"isMain":true},{"metricName":"Largest alignment","quality":"More is better","values":[2345,13711,12320,2201,13711,2354],"isMain":true},{"metricName":"Total aligned length","quality":"More is better","values":[9830,54400,36007,21276,61348,6209],"isMain":true},{"metricName":"NA50","quality":"More is better","values":[554,6112,801,513,3360,685],"isMain":false},{"metricName":"NA75","quality":"More is better","values":[133,675,null,null,502,564],"isMain":false},{"metricName":"NGA50","quality":"More is better","values":[null,null,null,null,null,null],"isMain":true},{"metricName":"LA50","quality":"Less is better","values":[8,3,9,22,5,3],"isMain":false},{"metricName":"LA75","quality":"Less is better","values":[16,11,null,null,32,5],"isMain":false}]],["Reads mapping",[]],["Misassemblies",[{"metricName":"# misassemblies","quality":"Less is better","values":[0,5,3,0,5,0],"isMain":true},{"metricName":" # relocations","quality":"Less is better","values":[0,0,0,0,0,0],"isMain":false},{"metricName":" # translocations","quality":"Less is better","values":[0,5,3,0,5,0],"isMain":false},{"metricName":" # inversions","quality":"Less is better","values":[0,0,0,0,0,0],"isMain":false},{"metricName":" # interspecies translocations","quality":"Less is better","values":[5,7,3,3,5,1],"isMain":false},{"metricName":"# misassembled contigs","quality":"Less is better","values":[0,1,1,0,1,0],"isMain":false},{"metricName":"Misassembled contigs length","quality":"Less is better","values":[0,41587,21636,0,41231,0],"isMain":true},{"metricName":"# possibly misassembled contigs","quality":"Less is better","values":[0,0,0,0,0,0],"isMain":false},{"metricName":" # possible misassemblies","quality":"Less is better","values":[0,0,0,0,0,0],"isMain":false},{"metricName":"# local misassemblies","quality":"Less is better","values":[0,1,2,0,1,0],"isMain":false},{"metricName":"# scaffold gap ext. mis.","quality":"Less is better","values":[0,0,0,0,0,0],"isMain":false},{"metricName":"# scaffold gap loc. mis.","quality":"Less is better","values":[0,0,0,0,0,0],"isMain":false},{"metricName":"# unaligned mis. contigs","quality":"Less is better","values":[0,0,1,1,1,0],"isMain":false}]],["Unaligned",[{"metricName":"# fully unaligned contigs","quality":"Less is better","values":[0,0,0,0,1,0],"isMain":false},{"metricName":"Fully unaligned length","quality":"Less is better","values":[0,0,0,0,2490,0],"isMain":false},{"metricName":"# partially unaligned contigs","quality":"Less is better","values":[1,5,3,2,3,0],"isMain":false},{"metricName":"Partially unaligned length","quality":"Less is better","values":[1422,5302,14105,12276,13074,0],"isMain":false}]],["Mismatches",[{"metricName":"# mismatches","quality":"Less is better","values":[198,1059,813,544,1265,199],"isMain":false},{"metricName":"# indels","quality":"Less is better","values":[6,69,27,10,73,4],"isMain":false},{"metricName":"Indels length","quality":"Less is better","values":[16,368,76,21,372,5],"isMain":false},{"metricName":"# mismatches per 100 kbp","quality":"Less is better","values":["2013.01","1962.64","2261.41","2574.90","2064.53","3205.54"],"isMain":true},{"metricName":"# indels per 100 kbp","quality":"Less is better","values":["61.00","127.88","75.10","47.33","119.14","64.43"],"isMain":true},{"metricName":" # indels (<= 5 bp)","quality":"Less is better","values":[5,63,24,9,67,4],"isMain":false},{"metricName":" # indels (> 5 bp)","quality":"Less is better","values":[1,6,3,1,6,0],"isMain":false},{"metricName":"# N's","quality":"Less is better","values":[0,0,0,0,0,0],"isMain":false},{"metricName":"# N's per 100 kbp","quality":"Less is better","values":["0.00","0.00","0.00","0.00","0.00","0.00"],"isMain":true}]],["Statistics without reference",[{"metricName":"# contigs","quality":"Equal","values":[17,25,19,25,29,8],"isMain":true},{"metricName":"# contigs (>= 1000 bp)","quality":"Equal","values":[2,6,5,5,5,1],"isMain":false},{"metricName":"# contigs (>= 5000 bp)","quality":"Equal","values":[0,1,2,1,2,0],"isMain":false},{"metricName":"# contigs (>= 10000 bp)","quality":"Equal","values":[0,1,2,1,2,0],"isMain":false},{"metricName":"# contigs (>= 25000 bp)","quality":"Equal","values":[0,1,0,0,1,0],"isMain":false},{"metricName":"# contigs (>= 50000 bp)","quality":"Equal","values":[0,0,0,0,0,0],"isMain":false},{"metricName":"Largest contig","quality":"More is better","values":[2348,41587,21636,15797,41231,2360],"isMain":true},{"metricName":"Total length","quality":"More is better","values":[12833,60694,50415,34023,77577,6528],"isMain":true},{"metricName":"Total length (>= 1000 bp)","quality":"More is better","values":[4005,49340,41931,21555,62879,2360],"isMain":true},{"metricName":"Total length (>= 5000 bp)","quality":"More is better","values":[0,41587,37419,15797,56936,0],"isMain":false},{"metricName":"Total length (>= 10000 bp)","quality":"More is better","values":[0,41587,37419,15797,56936,0],"isMain":true},{"metricName":"Total length (>= 25000 bp)","quality":"More is better","values":[0,41587,0,0,41231,0],"isMain":false},{"metricName":"Total length (>= 50000 bp)","quality":"More is better","values":[0,0,0,0,0,0],"isMain":true},{"metricName":"N50","quality":"More is better","values":[631,41587,15783,2201,41231,685],"isMain":false},{"metricName":"N75","quality":"More is better","values":[568,1657,2319,685,2490,582],"isMain":false},{"metricName":"L50","quality":"Less is better","values":[6,1,2,2,1,3],"isMain":false},{"metricName":"L75","quality":"Less is better","values":[11,3,3,11,3,5],"isMain":false},{"metricName":"GC (%)","quality":"Equal","values":["53.70","58.48","54.14","51.95","56.75","56.07"],"isMain":false}]],["K-mer-based statistics",[]],["Predicted genes",[]],["Similarity statistics",[]],["Reference statistics",[{"metricName":"Reference length","quality":"Equal","values":[4470163,4470163,4470163,4470163,4470163,4470163],"isMain":false},{"metricName":"Reference fragments","quality":"Equal","values":[425,425,425,425,425,425],"isMain":false},{"metricName":"Reference GC (%)","quality":"Equal","values":["59.40","59.40","59.40","59.40","59.40","59.40"],"isMain":false}]]],"subreferences":[],"subreports":[],"minContig":500},{"date":"10 February 2020, Monday, 18:52:07","assembliesNames":["megahit-103-df","megahit-113-df","megahit-113-ml","megahit-113-ms","megahit-129-df","metaSPAdes"],"referenceName":"GCF\_000225345.1\_ASM22534v1","order":[0,1,2,3,4,5],"report":[["Genome statistics",[{"metricName":"Genome fraction (%)","quality":"More is better","values":["0.391","0.407","0.524","0.593","0.475","0.117"],"isMain":true},{"metricName":"Duplication ratio","quality":"Less is better","values":["1.166","1.132","1.041","1.129","1.161","1.250"],"isMain":true},{"metricName":"Largest alignment","quality":"More is better","values":[3503,4407,5392,3495,2627,2063],"isMain":true},{"metricName":"Total aligned length","quality":"More is better","values":[14041,14624,18820,22374,17068,4190],"isMain":true},{"metricName":"NA50","quality":"More is better","values":[553,544,674,515,513,752],"isMain":false},{"metricName":"NA75","quality":"More is better","values":[null,null,512,null,null,140],"isMain":false},{"metricName":"NGA50","quality":"More is better","values":[null,null,null,null,null,null],"isMain":true},{"metricName":"LA50","quality":"Less is better","values":[9,11,7,22,17,2],"isMain":false},{"metricName":"LA75","quality":"Less is better","values":[null,null,16,null,null,5],"isMain":false}]],["Reads mapping",[]],["Misassemblies",[{"metricName":"# misassemblies","quality":"Less is better","values":[1,2,1,0,1,1],"isMain":true},{"metricName":" # relocations","quality":"Less is better","values":[1,2,1,0,1,1],"isMain":false},{"metricName":" # translocations","quality":"Less is better","values":[0,0,0,0,0,0],"isMain":false},{"metricName":" # inversions","quality":"Less is better","values":[0,0,0,0,0,0],"isMain":false},{"metricName":" # interspecies translocations","quality":"Less is better","values":[10,9,3,10,13,4],"isMain":false},{"metricName":"# misassembled contigs","quality":"Less is better","values":[1,1,1,0,1,1],"isMain":false},{"metricName":"Misassembled contigs length","quality":"Less is better","values":[3414,6112,3953,0,3493,2203],"isMain":true},{"metricName":"# possibly misassembled contigs","quality":"Less is better","values":[0,0,0,0,0,0],"isMain":false},{"metricName":" # possible misassemblies","quality":"Less is better","values":[0,0,0,0,0,0],"isMain":false},{"metricName":"# local misassemblies","quality":"Less is better","values":[0,0,0,0,0,0],"isMain":false},{"metricName":"# scaffold gap ext. mis.","quality":"Less is better","values":[0,0,0,0,0,0],"isMain":false},{"metricName":"# scaffold gap loc. mis.","quality":"Less is better","values":[0,0,0,0,0,0],"isMain":false},{"metricName":"# unaligned mis. contigs","quality":"Less is better","values":[0,0,0,0,1,0],"isMain":false}]],["Unaligned",[{"metricName":"# fully unaligned contigs","quality":"Less is better","values":[0,0,0,0,0,0],"isMain":false},{"metricName":"Fully unaligned length","quality":"Less is better","values":[0,0,0,0,0,0],"isMain":false},{"metricName":"# partially unaligned contigs","quality":"Less is better","values":[2,3,2,7,5,0],"isMain":false},{"metricName":"Partially unaligned length","quality":"Less is better","values":[4419,5906,1820,9670,9038,0],"isMain":false}]],["Mismatches",[{"metricName":"# mismatches","quality":"Less is better","values":[188,336,306,604,324,63],"isMain":false},{"metricName":"# indels","quality":"Less is better","values":[0,7,6,1,5,0],"isMain":false},{"metricName":"Indels length","quality":"Less is better","values":[0,8,6,6,5,0],"isMain":false},{"metricName":"# mismatches per 100 kbp","quality":"Less is better","values":["1338.94","2297.59","1625.93","2836.35","1898.40","1503.58"],"isMain":true},{"metricName":"# indels per 100 kbp","quality":"Less is better","values":["0.00","47.87","31.88","4.70","29.30","0.00"],"isMain":true},{"metricName":" # indels (<= 5 bp)","quality":"Less is better","values":[0,7,6,0,5,0],"isMain":false},{"metricName":" # indels (> 5 bp)","quality":"Less is better","values":[0,0,0,1,0,0],"isMain":false},{"metricName":"# N's","quality":"Less is better","values":[0,0,0,0,0,0],"isMain":false},{"metricName":"# N's per 100 kbp","quality":"Less is better","values":["0.00","0.00","0.00","0.00","0.00","0.00"],"isMain":true}]],["Statistics without reference",[{"metricName":"# contigs","quality":"Equal","values":[19,19,21,38,25,6],"isMain":true},{"metricName":"# contigs (>= 1000 bp)","quality":"Equal","values":[3,5,3,6,6,1],"isMain":false},{"metricName":"# contigs (>= 5000 bp)","quality":"Equal","values":[0,1,1,1,1,0],"isMain":false},{"metricName":"# contigs (>= 10000 bp)","quality":"Equal","values":[0,0,0,0,0,0],"isMain":false},{"metricName":"# contigs (>= 25000 bp)","quality":"Equal","values":[0,0,0,0,0,0],"isMain":false},{"metricName":"# contigs (>= 50000 bp)","quality":"Equal","values":[0,0,0,0,0,0],"isMain":false},{"metricName":"Largest contig","quality":"More is better","values":[3930,6112,5392,6609,5425,2203],"isMain":true},{"metricName":"Total length","quality":"More is better","values":[20793,22454,21415,33703,28854,5236],"isMain":true},{"metricName":"Total length (>= 1000 bp)","quality":"More is better","values":[10854,14133,10428,14819,16868,2203],"isMain":true},{"metricName":"Total length (>= 5000 bp)","quality":"More is better","values":[0,6112,5392,6609,5425,0],"isMain":false},{"metricName":"Total length (>= 10000 bp)","quality":"More is better","values":[0,0,0,0,0,0],"isMain":true},{"metricName":"Total length (>= 25000 bp)","quality":"More is better","values":[0,0,0,0,0,0],"isMain":false},{"metricName":"Total length (>= 50000 bp)","quality":"More is better","values":[0,0,0,0,0,0],"isMain":true},{"metricName":"N50","quality":"More is better","values":[3414,1721,861,769,1382,760],"isMain":false},{"metricName":"N75","quality":"More is better","values":[643,644,599,563,684,608],"isMain":false},{"metricName":"L50","quality":"Less is better","values":[3,3,4,9,5,2],"isMain":false},{"metricName":"L75","quality":"Less is better","values":[10,9,12,22,13,4],"isMain":false},{"metricName":"GC (%)","quality":"Equal","values":["44.31","44.17","43.00","46.57","44.76","42.55"],"isMain":false}]],["K-mer-based statistics",[]],["Predicted genes",[]],["Similarity statistics",[]],["Reference statistics",[{"metricName":"Reference length","quality":"Equal","values":[3592125,3592125,3592125,3592125,3592125,3592125],"isMain":false},{"metricName":"Reference fragments","quality":"Equal","values":[1,1,1,1,1,1],"isMain":false},{"metricName":"Reference GC (%)","quality":"Equal","values":["48.50","48.50","48.50","48.50","48.50","48.50"],"isMain":false}]]],"subreferences":[],"subreports":[],"minContig":500},{"date":"10 February 2020, Monday, 18:52:19","assembliesNames":["megahit-103-df","megahit-113-df","megahit-113-ml","megahit-113-ms","megahit-129-df","metaSPAdes"],"referenceName":"GCF\_000225685.1\_Erys\_bact\_2\_2\_44A\_V1","order":[0,1,2,3,4,5],"report":[["Genome statistics",[{"metricName":"Genome fraction (%)","quality":"More is better","values":["0.712","0.934","0.994","1.730","1.086","0.530"],"isMain":true},{"metricName":"Duplication ratio","quality":"Less is better","values":["1.099","1.073","1.028","1.045","1.075","1.038"],"isMain":true},{"metricName":"Largest alignment","quality":"More is better","values":[1933,1054,2361,6557,2362,1981],"isMain":true},{"metricName":"Total aligned length","quality":"More is better","values":[35349,46946,49639,85922,54304,26313],"isMain":true},{"metricName":"NA50","quality":"More is better","values":[531,532,537,543,532,538],"isMain":false},{"metricName":"NA75","quality":"More is better","values":[495,499,511,504,502,498],"isMain":false},{"metricName":"NGA50","quality":"More is better","values":[null,null,null,null,null,null],"isMain":true},{"metricName":"LA50","quality":"Less is better","values":[32,46,42,63,46,23],"isMain":false},{"metricName":"LA75","quality":"Less is better","values":[53,74,68,110,77,38],"isMain":false}]],["Reads mapping",[]],["Misassemblies",[{"metricName":"# misassemblies","quality":"Less is better","values":[0,0,1,3,0,1],"isMain":true},{"metricName":" # relocations","quality":"Less is better","values":[0,0,0,1,0,0],"isMain":false},{"metricName":" # translocations","quality":"Less is better","values":[0,0,1,2,0,1],"isMain":false},{"metricName":" # inversions","quality":"Less is better","values":[0,0,0,0,0,0],"isMain":false},{"metricName":" # interspecies translocations","quality":"Less is better","values":[16,15,7,22,15,7],"isMain":false},{"metricName":"# misassembled contigs","quality":"Less is better","values":[0,0,1,2,0,1],"isMain":false},{"metricName":"Misassembled contigs length","quality":"Less is better","values":[0,0,1304,1663,0,672],"isMain":true},{"metricName":"# possibly misassembled contigs","quality":"Less is better","values":[0,0,0,0,0,0],"isMain":false},{"metricName":" # possible misassemblies","quality":"Less is better","values":[0,0,0,0,0,0],"isMain":false},{"metricName":"# local misassemblies","quality":"Less is better","values":[0,0,0,0,1,0],"isMain":false},{"metricName":"# scaffold gap ext. mis.","quality":"Less is better","values":[0,0,0,0,0,0],"isMain":false},{"metricName":"# scaffold gap loc. mis.","quality":"Less is better","values":[0,0,0,0,0,0],"isMain":false},{"metricName":"# unaligned mis. contigs","quality":"Less is better","values":[0,0,0,1,0,0],"isMain":false}]],["Unaligned",[{"metricName":"# fully unaligned contigs","quality":"Less is better","values":[0,0,0,0,2,1],"isMain":false},{"metricName":"Fully unaligned length","quality":"Less is better","values":[0,0,0,0,1109,578],"isMain":false},{"metricName":"# partially unaligned contigs","quality":"Less is better","values":[5,9,3,9,6,2],"isMain":false},{"metricName":"Partially unaligned length","quality":"Less is better","values":[3314,7824,4002,7093,4113,4354],"isMain":false}]],["Mismatches",[{"metricName":"# mismatches","quality":"Less is better","values":[653,938,1067,1801,1076,521],"isMain":false},{"metricName":"# indels","quality":"Less is better","values":[0,2,2,7,4,5],"isMain":false},{"metricName":"Indels length","quality":"Less is better","values":[0,2,2,35,27,27],"isMain":false},{"metricName":"# mismatches per 100 kbp","quality":"Less is better","values":["1847.29","2021.16","2161.71","2095.92","1994.96","1980.09"],"isMain":true},{"metricName":"# indels per 100 kbp","quality":"Less is better","values":["0.00","4.31","4.05","8.15","7.42","19.00"],"isMain":true},{"metricName":" # indels (<= 5 bp)","quality":"Less is better","values":[0,2,2,6,3,3],"isMain":false},{"metricName":" # indels (> 5 bp)","quality":"Less is better","values":[0,0,0,1,1,2],"isMain":false},{"metricName":"# N's","quality":"Less is better","values":[0,0,0,0,0,0],"isMain":false},{"metricName":"# N's per 100 kbp","quality":"Less is better","values":["0.00","0.00","0.00","0.00","0.00","0.00"],"isMain":true}]],["Statistics without reference",[{"metricName":"# contigs","quality":"Equal","values":[68,89,86,144,100,44],"isMain":true},{"metricName":"# contigs (>= 1000 bp)","quality":"Equal","values":[2,4,4,11,5,3],"isMain":false},{"metricName":"# contigs (>= 5000 bp)","quality":"Equal","values":[0,0,0,1,0,0],"isMain":false},{"metricName":"# contigs (>= 10000 bp)","quality":"Equal","values":[0,0,0,0,0,0],"isMain":false},{"metricName":"# contigs (>= 25000 bp)","quality":"Equal","values":[0,0,0,0,0,0],"isMain":false},{"metricName":"# contigs (>= 50000 bp)","quality":"Equal","values":[0,0,0,0,0,0],"isMain":false},{"metricName":"Largest contig","quality":"More is better","values":[1933,2120,2745,6557,2362,4542],"isMain":true},{"metricName":"Total length","quality":"More is better","values":[42165,57608,54740,96905,63229,32245],"isMain":true},{"metricName":"Total length (>= 1000 bp)","quality":"More is better","values":[3174,5460,7446,18430,7897,7727],"isMain":true},{"metricName":"Total length (>= 5000 bp)","quality":"More is better","values":[0,0,0,6557,0,0],"isMain":false},{"metricName":"Total length (>= 10000 bp)","quality":"More is better","values":[0,0,0,0,0,0],"isMain":true},{"metricName":"Total length (>= 25000 bp)","quality":"More is better","values":[0,0,0,0,0,0],"isMain":false},{"metricName":"Total length (>= 50000 bp)","quality":"More is better","values":[0,0,0,0,0,0],"isMain":true},{"metricName":"N50","quality":"More is better","values":[575,603,563,609,583,621],"isMain":false},{"metricName":"N75","quality":"More is better","values":[518,532,526,528,528,542],"isMain":false},{"metricName":"L50","quality":"Less is better","values":[28,36,35,54,41,15],"isMain":false},{"metricName":"L75","quality":"Less is better","values":[48,62,60,97,70,29],"isMain":false},{"metricName":"GC (%)","quality":"Equal","values":["44.38","44.84","45.14","43.70","43.37","43.68"],"isMain":false}]],["K-mer-based statistics",[]],["Predicted genes",[]],["Similarity statistics",[]],["Reference statistics",[{"metricName":"Reference length","quality":"Equal","values":[4966372,4966372,4966372,4966372,4966372,4966372],"isMain":false},{"metricName":"Reference fragments","quality":"Equal","values":[94,94,94,94,94,94],"isMain":false},{"metricName":"Reference GC (%)","quality":"Equal","values":["43.34","43.34","43.34","43.34","43.34","43.34"],"isMain":false}]]],"subreferences":[],"subreports":[],"minContig":500},{"date":"10 February 2020, Monday, 18:52:51","assembliesNames":["megahit-103-df","megahit-113-df","megahit-113-ml","megahit-113-ms","megahit-129-df","metaSPAdes"],"referenceName":"GCF\_000225745.1\_Dorea\_form\_4\_6\_53AFAA\_V1","order":[0,1,2,3,4,5],"report":[["Genome statistics",[{"metricName":"Genome fraction (%)","quality":"More is better","values":["0.245","0.586","0.319","0.380","0.549","0.050"],"isMain":true},{"metricName":"Duplication ratio","quality":"Less is better","values":["1.076","1.088","1.119","1.062","1.186","1.002"],"isMain":true},{"metricName":"Largest alignment","quality":"More is better","values":[4932,7947,5820,5960,5960,1896],"isMain":true},{"metricName":"Total aligned length","quality":"More is better","values":[9379,22414,12207,14558,22032,1896],"isMain":true},{"metricName":"NA50","quality":"More is better","values":[578,null,null,null,502,1896],"isMain":false},{"metricName":"NA75","quality":"More is better","values":[null,null,null,null,null,1896],"isMain":false},{"metricName":"NGA50","quality":"More is better","values":[null,null,null,null,null,null],"isMain":true},{"metricName":"LA50","quality":"Less is better","values":[3,null,null,null,11,1],"isMain":false},{"metricName":"LA75","quality":"Less is better","values":[null,null,null,null,null,1],"isMain":false}]],["Reads mapping",[]],["Misassemblies",[{"metricName":"# misassemblies","quality":"Less is better","values":[1,1,1,1,1,0],"isMain":true},{"metricName":" # relocations","quality":"Less is better","values":[0,0,0,0,1,0],"isMain":false},{"metricName":" # translocations","quality":"Less is better","values":[1,1,1,1,0,0],"isMain":false},{"metricName":" # inversions","quality":"Less is better","values":[0,0,0,0,0,0],"isMain":false},{"metricName":" # interspecies translocations","quality":"Less is better","values":[5,7,8,7,10,0],"isMain":false},{"metricName":"# misassembled contigs","quality":"Less is better","values":[1,1,1,1,1,0],"isMain":false},{"metricName":"Misassembled contigs length","quality":"Less is better","values":[6177,14338,11877,11532,538,0],"isMain":true},{"metricName":"# possibly misassembled contigs","quality":"Less is better","values":[0,0,1,0,0,0],"isMain":false},{"metricName":" # possible misassemblies","quality":"Less is better","values":[0,0,1,0,0,0],"isMain":false},{"metricName":"# local misassemblies","quality":"Less is better","values":[0,0,0,0,0,0],"isMain":false},{"metricName":"# scaffold gap ext. mis.","quality":"Less is better","values":[0,0,0,0,0,0],"isMain":false},{"metricName":"# scaffold gap loc. mis.","quality":"Less is better","values":[0,0,0,0,0,0],"isMain":false},{"metricName":"# unaligned mis. contigs","quality":"Less is better","values":[0,0,0,0,1,0],"isMain":false}]],["Unaligned",[{"metricName":"# fully unaligned contigs","quality":"Less is better","values":[0,0,0,0,1,0],"isMain":false},{"metricName":"Fully unaligned length","quality":"Less is better","values":[0,0,0,0,743,0],"isMain":false},{"metricName":"# partially unaligned contigs","quality":"Less is better","values":[3,4,6,4,2,0],"isMain":false},{"metricName":"Partially unaligned length","quality":"Less is better","values":[2678,26976,31626,32802,13419,0],"isMain":false}]],["Mismatches",[{"metricName":"# mismatches","quality":"Less is better","values":[110,292,194,280,332,32],"isMain":false},{"metricName":"# indels","quality":"Less is better","values":[2,11,8,8,9,1],"isMain":false},{"metricName":"Indels length","quality":"Less is better","values":[2,14,8,8,11,1],"isMain":false},{"metricName":"# mismatches per 100 kbp","quality":"Less is better","values":["1173.08","1302.52","1589.51","1923.61","1580.95","1686.87"],"isMain":true},{"metricName":"# indels per 100 kbp","quality":"Less is better","values":["21.33","49.07","65.55","54.96","42.86","52.71"],"isMain":true},{"metricName":" # indels (<= 5 bp)","quality":"Less is better","values":[2,11,8,8,9,1],"isMain":false},{"metricName":" # indels (> 5 bp)","quality":"Less is better","values":[0,0,0,0,0,0],"isMain":false},{"metricName":"# N's","quality":"Less is better","values":[0,0,0,0,0,0],"isMain":false},{"metricName":"# N's per 100 kbp","quality":"Less is better","values":["0.00","0.00","0.00","0.00","0.00","0.00"],"isMain":true}]],["Statistics without reference",[{"metricName":"# contigs","quality":"Equal","values":[10,16,14,15,20,1],"isMain":true},{"metricName":"# contigs (>= 1000 bp)","quality":"Equal","values":[2,5,3,4,5,1],"isMain":false},{"metricName":"# contigs (>= 5000 bp)","quality":"Equal","values":[1,3,2,2,2,0],"isMain":false},{"metricName":"# contigs (>= 10000 bp)","quality":"Equal","values":[0,2,2,2,1,0],"isMain":false},{"metricName":"# contigs (>= 25000 bp)","quality":"Equal","values":[0,0,0,1,0,0],"isMain":false},{"metricName":"# contigs (>= 50000 bp)","quality":"Equal","values":[0,0,0,0,0,0],"isMain":false},{"metricName":"Largest contig","quality":"More is better","values":[6177,19384,24957,27897,18261,1900],"isMain":true},{"metricName":"Total length","quality":"More is better","values":[12772,51377,45284,48258,39064,1900],"isMain":true},{"metricName":"Total length (>= 1000 bp)","quality":"More is better","values":[7954,44443,38230,41767,30265,1900],"isMain":true},{"metricName":"Total length (>= 5000 bp)","quality":"More is better","values":[6177,41669,36834,39429,23387,0],"isMain":false},{"metricName":"Total length (>= 10000 bp)","quality":"More is better","values":[0,33722,36834,39429,18261,0],"isMain":true},{"metricName":"Total length (>= 25000 bp)","quality":"More is better","values":[0,0,0,27897,0,0],"isMain":false},{"metricName":"Total length (>= 50000 bp)","quality":"More is better","values":[0,0,0,0,0,0],"isMain":true},{"metricName":"N50","quality":"More is better","values":[1777,14338,24957,27897,5126,1900],"isMain":false},{"metricName":"N75","quality":"More is better","values":[589,7947,11877,11532,1589,1900],"isMain":false},{"metricName":"L50","quality":"Less is better","values":[2,2,1,1,2,1],"isMain":false},{"metricName":"L75","quality":"Less is better","values":[5,3,2,2,5,1],"isMain":false},{"metricName":"GC (%)","quality":"Equal","values":["35.21","36.49","35.65","35.77","36.89","34.84"],"isMain":false}]],["K-mer-based statistics",[]],["Predicted genes",[]],["Similarity statistics",[]],["Reference statistics",[{"metricName":"Reference length","quality":"Equal","values":[3825692,3825692,3825692,3825692,3825692,3825692],"isMain":false},{"metricName":"Reference fragments","quality":"Equal","values":[151,151,151,151,151,151],"isMain":false},{"metricName":"Reference GC (%)","quality":"Equal","values":["40.34","40.34","40.34","40.34","40.34","40.34"],"isMain":false}]]],"subreferences":[],"subreports":[],"minContig":500},{"date":"10 February 2020, Monday, 18:35:43","assembliesNames":["megahit-103-df","megahit-113-df","megahit-113-ms","megahit-129-df"],"referenceName":"GCF\_000231275.1\_Alis\_indi\_YIT\_V1","order":[0,1,2,3],"report":[["Genome statistics",[{"metricName":"Genome fraction (%)","quality":"More is better","values":["0.017","0.013","0.009","0.010"],"isMain":true},{"metricName":"Duplication ratio","quality":"Less is better","values":["1.907","2.173","1.000","2.470"],"isMain":true},{"metricName":"Largest alignment","quality":"More is better","values":[371,371,261,296],"isMain":true},{"metricName":"Total aligned length","quality":"More is better","values":[471,371,261,296],"isMain":true},{"metricName":"NGA50","quality":"More is better","values":[null,null,null,null],"isMain":true}]],["Reads mapping",[]],["Misassemblies",[{"metricName":"# misassemblies","quality":"Less is better","values":[0,0,0,0],"isMain":true},{"metricName":" # relocations","quality":"Less is better","values":[0,0,0,0],"isMain":false},{"metricName":" # translocations","quality":"Less is better","values":[0,0,0,0],"isMain":false},{"metricName":" # inversions","quality":"Less is better","values":[0,0,0,0],"isMain":false},{"metricName":" # interspecies translocations","quality":"Less is better","values":[2,1,1,1],"isMain":false},{"metricName":"# misassembled contigs","quality":"Less is better","values":[0,0,0,0],"isMain":false},{"metricName":"Misassembled contigs length","quality":"Less is better","values":[0,0,0,0],"isMain":true},{"metricName":"# possibly misassembled contigs","quality":"Less is better","values":[0,0,0,0],"isMain":false},{"metricName":" # possible misassemblies","quality":"Less is better","values":[0,0,0,0],"isMain":false},{"metricName":"# local misassemblies","quality":"Less is better","values":[0,0,0,0],"isMain":false},{"metricName":"# scaffold gap ext. mis.","quality":"Less is better","values":[0,0,0,0],"isMain":false},{"metricName":"# scaffold gap loc. mis.","quality":"Less is better","values":[0,0,0,0],"isMain":false},{"metricName":"# unaligned mis. contigs","quality":"Less is better","values":[0,0,0,0],"isMain":false}]],["Unaligned",[{"metricName":"# fully unaligned contigs","quality":"Less is better","values":[0,0,0,0],"isMain":false},{"metricName":"Fully unaligned length","quality":"Less is better","values":[0,0,0,0],"isMain":false},{"metricName":"# partially unaligned contigs","quality":"Less is better","values":[1,0,1,0],"isMain":false},{"metricName":"Partially unaligned length","quality":"Less is better","values":[609,0,816,0],"isMain":false}]],["Mismatches",[{"metricName":"# mismatches","quality":"Less is better","values":[1,1,6,5],"isMain":false},{"metricName":"# indels","quality":"Less is better","values":[0,0,0,0],"isMain":false},{"metricName":"Indels length","quality":"Less is better","values":[0,0,0,0],"isMain":false},{"metricName":"# mismatches per 100 kbp","quality":"Less is better","values":["212.31","269.54","2298.85","1689.19"],"isMain":true},{"metricName":"# indels per 100 kbp","quality":"Less is better","values":["0.00","0.00","0.00","0.00"],"isMain":true},{"metricName":" # indels (<= 5 bp)","quality":"Less is better","values":[0,0,0,0],"isMain":false},{"metricName":" # indels (> 5 bp)","quality":"Less is better","values":[0,0,0,0],"isMain":false},{"metricName":"# N's","quality":"Less is better","values":[0,0,0,0],"isMain":false},{"metricName":"# N's per 100 kbp","quality":"Less is better","values":["0.00","0.00","0.00","0.00"],"isMain":true}]],["Statistics without reference",[{"metricName":"# contigs","quality":"Equal","values":[2,1,1,1],"isMain":true},{"metricName":"# contigs (>= 1000 bp)","quality":"Equal","values":[0,0,1,0],"isMain":false},{"metricName":"# contigs (>= 5000 bp)","quality":"Equal","values":[0,0,0,0],"isMain":false},{"metricName":"# contigs (>= 10000 bp)","quality":"Equal","values":[0,0,0,0],"isMain":false},{"metricName":"# contigs (>= 25000 bp)","quality":"Equal","values":[0,0,0,0],"isMain":false},{"metricName":"# contigs (>= 50000 bp)","quality":"Equal","values":[0,0,0,0],"isMain":false},{"metricName":"Largest contig","quality":"More is better","values":[798,806,1077,731],"isMain":true},{"metricName":"Total length","quality":"More is better","values":[1507,806,1077,731],"isMain":true},{"metricName":"Total length (>= 1000 bp)","quality":"More is better","values":[0,0,1077,0],"isMain":true},{"metricName":"Total length (>= 5000 bp)","quality":"More is better","values":[0,0,0,0],"isMain":false},{"metricName":"Total length (>= 10000 bp)","quality":"More is better","values":[0,0,0,0],"isMain":true},{"metricName":"Total length (>= 25000 bp)","quality":"More is better","values":[0,0,0,0],"isMain":false},{"metricName":"Total length (>= 50000 bp)","quality":"More is better","values":[0,0,0,0],"isMain":true},{"metricName":"N50","quality":"More is better","values":[798,806,1077,731],"isMain":false},{"metricName":"N75","quality":"More is better","values":[709,806,1077,731],"isMain":false},{"metricName":"L50","quality":"Less is better","values":[1,1,1,1],"isMain":false},{"metricName":"L75","quality":"Less is better","values":[2,1,1,1],"isMain":false},{"metricName":"GC (%)","quality":"Equal","values":["57.73","60.79","47.54","61.83"],"isMain":false}]],["K-mer-based statistics",[]],["Predicted genes",[]],["Similarity statistics",[]],["Reference statistics",[{"metricName":"Reference length","quality":"Equal","values":[2850978,2850978,2850978,2850978],"isMain":false},{"metricName":"Reference fragments","quality":"Equal","values":[18,18,18,18],"isMain":false},{"metricName":"Reference GC (%)","quality":"Equal","values":["54.75","54.75","54.75","54.75"],"isMain":false}]]],"subreferences":[],"subreports":[],"minContig":500},{"date":"10 February 2020, Monday, 18:53:23","assembliesNames":["megahit-103-df","megahit-113-df","megahit-113-ml","megahit-113-ms","megahit-129-df","metaSPAdes"],"referenceName":"GCF\_000233455.1\_Clos\_citr\_WAL\_17108\_V1","order":[0,1,2,3,4,5],"report":[["Genome statistics",[{"metricName":"Genome fraction (%)","quality":"More is better","values":["2.040","2.422","2.787","2.862","2.447","1.356"],"isMain":true},{"metricName":"Duplication ratio","quality":"Less is better","values":["1.044","1.035","1.020","1.045","1.036","1.036"],"isMain":true},{"metricName":"Largest alignment","quality":"More is better","values":[1157,1425,2676,1889,2676,2676],"isMain":true},{"metricName":"Total aligned length","quality":"More is better","values":[135646,160927,184856,190056,162836,89935],"isMain":true},{"metricName":"NA50","quality":"More is better","values":[564,592,615,578,617,563],"isMain":false},{"metricName":"NA75","quality":"More is better","values":[516,526,537,518,525,517],"isMain":false},{"metricName":"NGA50","quality":"More is better","values":[null,null,null,null,null,null],"isMain":true},{"metricName":"LA50","quality":"Less is better","values":[111,122,122,142,112,69],"isMain":false},{"metricName":"LA75","quality":"Less is better","values":[180,202,205,236,190,112],"isMain":false}]],["Reads mapping",[]],["Misassemblies",[{"metricName":"# misassemblies","quality":"Less is better","values":[1,0,1,0,0,0],"isMain":true},{"metricName":" # relocations","quality":"Less is better","values":[0,0,0,0,0,0],"isMain":false},{"metricName":" # translocations","quality":"Less is better","values":[1,0,1,0,0,0],"isMain":false},{"metricName":" # inversions","quality":"Less is better","values":[0,0,0,0,0,0],"isMain":false},{"metricName":" # interspecies translocations","quality":"Less is better","values":[22,26,6,27,16,7],"isMain":false},{"metricName":"# misassembled contigs","quality":"Less is better","values":[1,0,1,0,0,0],"isMain":false},{"metricName":"Misassembled contigs length","quality":"Less is better","values":[1063,0,552,0,0,0],"isMain":true},{"metricName":"# possibly misassembled contigs","quality":"Less is better","values":[0,0,0,1,0,0],"isMain":false},{"metricName":" # possible misassemblies","quality":"Less is better","values":[0,0,0,1,0,0],"isMain":false},{"metricName":"# local misassemblies","quality":"Less is better","values":[0,1,0,1,0,0],"isMain":false},{"metricName":"# scaffold gap ext. mis.","quality":"Less is better","values":[0,0,0,0,0,0],"isMain":false},{"metricName":"# scaffold gap loc. mis.","quality":"Less is better","values":[0,0,0,0,0,0],"isMain":false},{"metricName":"# unaligned mis. contigs","quality":"Less is better","values":[0,0,0,0,0,0],"isMain":false}]],["Unaligned",[{"metricName":"# fully unaligned contigs","quality":"Less is better","values":[0,0,0,0,0,0],"isMain":false},{"metricName":"Fully unaligned length","quality":"Less is better","values":[0,0,0,0,0,0],"isMain":false},{"metricName":"# partially unaligned contigs","quality":"Less is better","values":[5,12,2,7,5,1],"isMain":false},{"metricName":"Partially unaligned length","quality":"Less is better","values":[10046,10750,1183,6805,6389,968],"isMain":false}]],["Mismatches",[{"metricName":"# mismatches","quality":"Less is better","values":[2085,2614,3591,3962,2459,1424],"isMain":false},{"metricName":"# indels","quality":"Less is better","values":[9,1,3,6,7,4],"isMain":false},{"metricName":"Indels length","quality":"Less is better","values":[36,27,34,6,7,4],"isMain":false},{"metricName":"# mismatches per 100 kbp","quality":"Less is better","values":["1542.66","1629.06","1945.23","2089.57","1517.08","1585.53"],"isMain":true},{"metricName":"# indels per 100 kbp","quality":"Less is better","values":["6.66","0.62","1.63","3.16","4.32","4.45"],"isMain":true},{"metricName":" # indels (<= 5 bp)","quality":"Less is better","values":[8,0,2,6,7,4],"isMain":false},{"metricName":" # indels (> 5 bp)","quality":"Less is better","values":[1,1,1,0,0,0],"isMain":false},{"metricName":"# N's","quality":"Less is better","values":[0,0,0,0,0,0],"isMain":false},{"metricName":"# N's per 100 kbp","quality":"Less is better","values":["0.00","0.00","0.00","0.00","0.00","0.00"],"isMain":true}]],["Statistics without reference",[{"metricName":"# contigs","quality":"Equal","values":[231,267,291,317,258,153],"isMain":true},{"metricName":"# contigs (>= 1000 bp)","quality":"Equal","values":[8,13,9,11,18,2],"isMain":false},{"metricName":"# contigs (>= 5000 bp)","quality":"Equal","values":[1,0,0,0,0,0],"isMain":false},{"metricName":"# contigs (>= 10000 bp)","quality":"Equal","values":[0,0,0,0,0,0],"isMain":false},{"metricName":"# contigs (>= 25000 bp)","quality":"Equal","values":[0,0,0,0,0,0],"isMain":false},{"metricName":"# contigs (>= 50000 bp)","quality":"Equal","values":[0,0,0,0,0,0],"isMain":false},{"metricName":"Largest contig","quality":"More is better","values":[7438,1824,2676,2010,3255,2676],"isMain":true},{"metricName":"Total length","quality":"More is better","values":[151210,176802,189500,204851,174264,94050],"isMain":true},{"metricName":"Total length (>= 1000 bp)","quality":"More is better","values":[15560,16345,12219,15149,24506,3949],"isMain":true},{"metricName":"Total length (>= 5000 bp)","quality":"More is better","values":[7438,0,0,0,0,0],"isMain":false},{"metricName":"Total length (>= 10000 bp)","quality":"More is better","values":[0,0,0,0,0,0],"isMain":true},{"metricName":"Total length (>= 25000 bp)","quality":"More is better","values":[0,0,0,0,0,0],"isMain":false},{"metricName":"Total length (>= 50000 bp)","quality":"More is better","values":[0,0,0,0,0,0],"isMain":true},{"metricName":"N50","quality":"More is better","values":[612,636,627,617,650,586],"isMain":false},{"metricName":"N75","quality":"More is better","values":[543,556,550,546,551,534],"isMain":false},{"metricName":"L50","quality":"Less is better","values":[93,109,120,130,102,66],"isMain":false},{"metricName":"L75","quality":"Less is better","values":[159,183,201,219,175,108],"isMain":false},{"metricName":"GC (%)","quality":"Equal","values":["49.23","50.10","50.00","50.14","50.40","50.27"],"isMain":false}]],["K-mer-based statistics",[]],["Predicted genes",[]],["Similarity statistics",[]],["Reference statistics",[{"metricName":"Reference length","quality":"Equal","values":[6624793,6624793,6624793,6624793,6624793,6624793],"isMain":false},{"metricName":"Reference fragments","quality":"Equal","values":[90,90,90,90,90,90],"isMain":false},{"metricName":"Reference GC (%)","quality":"Equal","values":["48.92","48.92","48.92","48.92","48.92","48.92"],"isMain":false}]]],"subreferences":[],"subreports":[],"minContig":500},{"date":"10 February 2020, Monday, 18:53:40","assembliesNames":["megahit-103-df","megahit-113-df","megahit-113-ml","megahit-113-ms","megahit-129-df","metaSPAdes"],"referenceName":"GCF\_000234055.1\_Prev\_hist\_F0411\_V1","order":[0,1,2,3,4,5],"report":[["Genome statistics",[{"metricName":"Genome fraction (%)","quality":"More is better","values":["0.122","0.088","0.213","0.147","0.326","0.066"],"isMain":true},{"metricName":"Duplication ratio","quality":"Less is better","values":["1.197","1.172","1.069","1.170","1.095","1.147"],"isMain":true},{"metricName":"Largest alignment","quality":"More is better","values":[1342,1903,2599,1209,3168,884],"isMain":true},{"metricName":"Total aligned length","quality":"More is better","values":[3768,2723,6364,4379,10617,1965],"isMain":true},{"metricName":"NA50","quality":"More is better","values":[627,null,537,null,1033,369],"isMain":false},{"metricName":"NA75","quality":"More is better","values":[110,null,null,null,null,null],"isMain":false},{"metricName":"NGA50","quality":"More is better","values":[null,null,null,null,null,null],"isMain":true},{"metricName":"LA50","quality":"Less is better","values":[3,null,5,null,5,3],"isMain":false},{"metricName":"LA75","quality":"Less is better","values":[7,null,null,null,null,null],"isMain":false}]],["Reads mapping",[]],["Misassemblies",[{"metricName":"# misassemblies","quality":"Less is better","values":[1,0,0,0,0,0],"isMain":true},{"metricName":" # relocations","quality":"Less is better","values":[0,0,0,0,0,0],"isMain":false},{"metricName":" # translocations","quality":"Less is better","values":[0,0,0,0,0,0],"isMain":false},{"metricName":" # inversions","quality":"Less is better","values":[1,0,0,0,0,0],"isMain":false},{"metricName":" # interspecies translocations","quality":"Less is better","values":[3,3,4,4,4,2],"isMain":false},{"metricName":"# misassembled contigs","quality":"Less is better","values":[1,0,0,0,0,0],"isMain":false},{"metricName":"Misassembled contigs length","quality":"Less is better","values":[589,0,0,0,0,0],"isMain":true},{"metricName":"# possibly misassembled contigs","quality":"Less is better","values":[0,0,0,0,0,0],"isMain":false},{"metricName":" # possible misassemblies","quality":"Less is better","values":[0,0,0,0,0,0],"isMain":false},{"metricName":"# local misassemblies","quality":"Less is better","values":[0,0,1,0,0,0],"isMain":false},{"metricName":"# scaffold gap ext. mis.","quality":"Less is better","values":[0,0,0,0,0,0],"isMain":false},{"metricName":"# scaffold gap loc. mis.","quality":"Less is better","values":[0,0,0,0,0,0],"isMain":false},{"metricName":"# unaligned mis. contigs","quality":"Less is better","values":[0,2,0,0,1,1],"isMain":false}]],["Unaligned",[{"metricName":"# fully unaligned contigs","quality":"Less is better","values":[1,0,0,1,0,0],"isMain":false},{"metricName":"Fully unaligned length","quality":"Less is better","values":[522,0,0,702,0,0],"isMain":false},{"metricName":"# partially unaligned contigs","quality":"Less is better","values":[0,2,3,1,4,1],"isMain":false},{"metricName":"Partially unaligned length","quality":"Less is better","values":[0,3592,3098,6336,6886,646],"isMain":false}]],["Mismatches",[{"metricName":"# mismatches","quality":"Less is better","values":[29,59,140,133,278,35],"isMain":false},{"metricName":"# indels","quality":"Less is better","values":[0,0,6,5,6,1],"isMain":false},{"metricName":"Indels length","quality":"Less is better","values":[0,0,8,7,11,1],"isMain":false},{"metricName":"# mismatches per 100 kbp","quality":"Less is better","values":["792.78","2252.77","2199.87","3037.22","2854.80","1780.26"],"isMain":true},{"metricName":"# indels per 100 kbp","quality":"Less is better","values":["0.00","0.00","94.28","114.18","61.61","50.86"],"isMain":true},{"metricName":" # indels (<= 5 bp)","quality":"Less is better","values":[0,0,6,5,6,1],"isMain":false},{"metricName":" # indels (> 5 bp)","quality":"Less is better","values":[0,0,0,0,0,0],"isMain":false},{"metricName":"# N's","quality":"Less is better","values":[0,0,0,0,0,0],"isMain":false},{"metricName":"# N's per 100 kbp","quality":"Less is better","values":["0.00","0.00","0.00","0.00","0.00","0.00"],"isMain":true}]],["Statistics without reference",[{"metricName":"# contigs","quality":"Equal","values":[7,4,9,8,8,4],"isMain":true},{"metricName":"# contigs (>= 1000 bp)","quality":"Equal","values":[1,2,3,2,4,0],"isMain":false},{"metricName":"# contigs (>= 5000 bp)","quality":"Equal","values":[0,0,0,1,1,0],"isMain":false},{"metricName":"# contigs (>= 10000 bp)","quality":"Equal","values":[0,0,0,0,0,0],"isMain":false},{"metricName":"# contigs (>= 25000 bp)","quality":"Equal","values":[0,0,0,0,0,0],"isMain":false},{"metricName":"# contigs (>= 50000 bp)","quality":"Equal","values":[0,0,0,0,0,0],"isMain":false},{"metricName":"Largest contig","quality":"More is better","values":[1342,3278,4237,6909,9958,884],"isMain":true},{"metricName":"Total length","quality":"More is better","values":[4901,6661,9900,12162,17551,2901],"isMain":true},{"metricName":"Total length (>= 1000 bp)","quality":"More is better","values":[1342,5181,6364,8296,14820,0],"isMain":true},{"metricName":"Total length (>= 5000 bp)","quality":"More is better","values":[0,0,0,6909,9958,0],"isMain":false},{"metricName":"Total length (>= 10000 bp)","quality":"More is better","values":[0,0,0,0,0,0],"isMain":true},{"metricName":"Total length (>= 25000 bp)","quality":"More is better","values":[0,0,0,0,0,0],"isMain":false},{"metricName":"Total length (>= 50000 bp)","quality":"More is better","values":[0,0,0,0,0,0],"isMain":true},{"metricName":"N50","quality":"More is better","values":[627,1903,1097,6909,9958,872],"isMain":false},{"metricName":"N75","quality":"More is better","values":[539,1903,608,933,1725,616],"isMain":false},{"metricName":"L50","quality":"Less is better","values":[3,2,2,1,1,2],"isMain":false},{"metricName":"L75","quality":"Less is better","values":[5,2,5,3,3,3],"isMain":false},{"metricName":"GC (%)","quality":"Equal","values":["42.71","43.16","45.37","42.00","43.64","36.81"],"isMain":false}]],["K-mer-based statistics",[]],["Predicted genes",[]],["Similarity statistics",[]],["Reference statistics",[{"metricName":"Reference length","quality":"Equal","values":[2987441,2987441,2987441,2987441,2987441,2987441],"isMain":false},{"metricName":"Reference fragments","quality":"Equal","values":[39,39,39,39,39,39],"isMain":false},{"metricName":"Reference GC (%)","quality":"Equal","values":["41.18","41.18","41.18","41.18","41.18","41.18"],"isMain":false}]]],"subreferences":[],"subreports":[],"minContig":500},{"date":"10 February 2020, Monday, 18:53:54","assembliesNames":["megahit-103-df","megahit-113-df","megahit-113-ml","megahit-113-ms","megahit-129-df","metaSPAdes"],"referenceName":"GCF\_000234115.1\_Prevotella\_sp\_F0323\_V1","order":[0,1,2,3,4,5],"report":[["Genome statistics",[{"metricName":"Genome fraction (%)","quality":"More is better","values":["0.164","0.310","0.119","0.418","0.136","0.068"],"isMain":true},{"metricName":"Duplication ratio","quality":"Less is better","values":["1.217","1.044","1.264","1.196","1.084","1.208"],"isMain":true},{"metricName":"Largest alignment","quality":"More is better","values":[1080,3446,676,2110,761,1035],"isMain":true},{"metricName":"Total aligned length","quality":"More is better","values":[4337,8033,3084,11197,3518,1751],"isMain":true},{"metricName":"NA50","quality":"More is better","values":[588,1781,431,546,251,500],"isMain":false},{"metricName":"NA75","quality":"More is better","values":[506,550,null,null,null,216],"isMain":false},{"metricName":"NGA50","quality":"More is better","values":[null,null,null,null,null,null],"isMain":true},{"metricName":"LA50","quality":"Less is better","values":[4,2,5,8,6,2],"isMain":false},{"metricName":"LA75","quality":"Less is better","values":[6,5,null,null,null,3],"isMain":false}]],["Reads mapping",[]],["Misassemblies",[{"metricName":"# misassemblies","quality":"Less is better","values":[0,0,0,0,0,0],"isMain":true},{"metricName":" # relocations","quality":"Less is better","values":[0,0,0,0,0,0],"isMain":false},{"metricName":" # translocations","quality":"Less is better","values":[0,0,0,0,0,0],"isMain":false},{"metricName":" # inversions","quality":"Less is better","values":[0,0,0,0,0,0],"isMain":false},{"metricName":" # interspecies translocations","quality":"Less is better","values":[2,3,4,2,3,1],"isMain":false},{"metricName":"# misassembled contigs","quality":"Less is better","values":[0,0,0,0,0,0],"isMain":false},{"metricName":"Misassembled contigs length","quality":"Less is better","values":[0,0,0,0,0,0],"isMain":true},{"metricName":"# possibly misassembled contigs","quality":"Less is better","values":[0,0,0,1,0,0],"isMain":false},{"metricName":" # possible misassemblies","quality":"Less is better","values":[0,0,0,1,0,0],"isMain":false},{"metricName":"# local misassemblies","quality":"Less is better","values":[0,0,0,0,0,0],"isMain":false},{"metricName":"# scaffold gap ext. mis.","quality":"Less is better","values":[0,0,0,0,0,0],"isMain":false},{"metricName":"# scaffold gap loc. mis.","quality":"Less is better","values":[0,0,0,0,0,0],"isMain":false},{"metricName":"# unaligned mis. contigs","quality":"Less is better","values":[0,0,1,1,1,0],"isMain":false}]],["Unaligned",[{"metricName":"# fully unaligned contigs","quality":"Less is better","values":[0,0,2,1,0,0],"isMain":false},{"metricName":"Fully unaligned length","quality":"Less is better","values":[0,0,1478,706,0,0],"isMain":false},{"metricName":"# partially unaligned contigs","quality":"Less is better","values":[0,1,0,1,2,0],"isMain":false},{"metricName":"Partially unaligned length","quality":"Less is better","values":[0,828,0,1585,2350,0],"isMain":false}]],["Mismatches",[{"metricName":"# mismatches","quality":"Less is better","values":[66,195,96,310,98,33],"isMain":false},{"metricName":"# indels","quality":"Less is better","values":[0,3,1,4,0,0],"isMain":false},{"metricName":"Indels length","quality":"Less is better","values":[0,3,1,6,0,0],"isMain":false},{"metricName":"# mismatches per 100 kbp","quality":"Less is better","values":["1557.34","2427.79","3111.83","2867.72","2785.67","1884.64"],"isMain":true},{"metricName":"# indels per 100 kbp","quality":"Less is better","values":["0.00","37.35","32.41","37.00","0.00","0.00"],"isMain":true},{"metricName":" # indels (<= 5 bp)","quality":"Less is better","values":[0,3,1,4,0,0],"isMain":false},{"metricName":" # indels (> 5 bp)","quality":"Less is better","values":[0,0,0,0,0,0],"isMain":false},{"metricName":"# N's","quality":"Less is better","values":[0,0,0,0,0,0],"isMain":false},{"metricName":"# N's per 100 kbp","quality":"Less is better","values":["0.00","0.00","0.00","0.00","0.00","0.00"],"isMain":true}]],["Statistics without reference",[{"metricName":"# contigs","quality":"Equal","values":[8,7,9,18,6,3],"isMain":true},{"metricName":"# contigs (>= 1000 bp)","quality":"Equal","values":[1,3,0,3,1,1],"isMain":false},{"metricName":"# contigs (>= 5000 bp)","quality":"Equal","values":[0,0,0,0,0,0],"isMain":false},{"metricName":"# contigs (>= 10000 bp)","quality":"Equal","values":[0,0,0,0,0,0],"isMain":false},{"metricName":"# contigs (>= 25000 bp)","quality":"Equal","values":[0,0,0,0,0,0],"isMain":false},{"metricName":"# contigs (>= 50000 bp)","quality":"Equal","values":[0,0,0,0,0,0],"isMain":false},{"metricName":"Largest contig","quality":"More is better","values":[1080,3446,829,2164,2797,1035],"isMain":true},{"metricName":"Total length","quality":"More is better","values":[5159,9214,5378,15221,6164,2115],"isMain":true},{"metricName":"Total length (>= 1000 bp)","quality":"More is better","values":[1080,6391,0,5747,2797,1035],"isMain":true},{"metricName":"Total length (>= 5000 bp)","quality":"More is better","values":[0,0,0,0,0,0],"isMain":false},{"metricName":"Total length (>= 10000 bp)","quality":"More is better","values":[0,0,0,0,0,0],"isMain":true},{"metricName":"Total length (>= 25000 bp)","quality":"More is better","values":[0,0,0,0,0,0],"isMain":false},{"metricName":"Total length (>= 50000 bp)","quality":"More is better","values":[0,0,0,0,0,0],"isMain":true},{"metricName":"N50","quality":"More is better","values":[594,1781,602,767,867,575],"isMain":false},{"metricName":"N75","quality":"More is better","values":[516,992,515,588,623,575],"isMain":false},{"metricName":"L50","quality":"Less is better","values":[4,2,4,6,2,2],"isMain":false},{"metricName":"L75","quality":"Less is better","values":[6,4,7,11,4,2],"isMain":false},{"metricName":"GC (%)","quality":"Equal","values":["41.97","43.22","44.59","44.20","42.10","41.84"],"isMain":false}]],["K-mer-based statistics",[]],["Predicted genes",[]],["Similarity statistics",[]],["Reference statistics",[{"metricName":"Reference length","quality":"Equal","values":[2588569,2588569,2588569,2588569,2588569,2588569],"isMain":false},{"metricName":"Reference fragments","quality":"Equal","values":[58,58,58,58,58,58],"isMain":false},{"metricName":"Reference GC (%)","quality":"Equal","values":["45.49","45.49","45.49","45.49","45.49","45.49"],"isMain":false}]]],"subreferences":[],"subreports":[],"minContig":500},{"date":"10 February 2020, Monday, 19:18:47","assembliesNames":["megahit-103-df","megahit-113-df","megahit-113-ml","megahit-113-ms","megahit-129-df","metaSPAdes"],"referenceName":"GCF\_000234155.1\_Clos\_clos\_2\_1\_49FAA\_V1","order":[0,1,2,3,4,5],"report":[["Genome statistics",[{"metricName":"Genome fraction (%)","quality":"More is better","values":["14.343","14.980","15.945","15.566","14.283","8.957"],"isMain":true},{"metricName":"Duplication ratio","quality":"Less is better","values":["1.022","1.027","1.024","1.036","1.031","1.017"],"isMain":true},{"metricName":"Largest alignment","quality":"More is better","values":[32721,24011,27264,35780,40478,87104],"isMain":true},{"metricName":"Total aligned length","quality":"More is better","values":[795211,831680,886402,872921,797421,494375],"isMain":true},{"metricName":"NA50","quality":"More is better","values":[4087,4242,4582,5253,2875,5091],"isMain":false},{"metricName":"NA75","quality":"More is better","values":[1308,1438,1196,1149,1029,1053],"isMain":false},{"metricName":"NGA50","quality":"More is better","values":[null,null,null,null,null,null],"isMain":true},{"metricName":"LA50","quality":"Less is better","values":[48,43,40,41,49,10],"isMain":false},{"metricName":"LA75","quality":"Less is better","values":[142,136,144,152,178,88],"isMain":false}]],["Reads mapping",[]],["Misassemblies",[{"metricName":"# misassemblies","quality":"Less is better","values":[5,5,4,9,11,1],"isMain":true},{"metricName":" # relocations","quality":"Less is better","values":[0,0,0,0,0,0],"isMain":false},{"metricName":" # translocations","quality":"Less is better","values":[4,5,4,8,10,1],"isMain":false},{"metricName":" # inversions","quality":"Less is better","values":[1,0,0,1,1,0],"isMain":false},{"metricName":" # interspecies translocations","quality":"Less is better","values":[11,19,13,20,32,3],"isMain":false},{"metricName":"# misassembled contigs","quality":"Less is better","values":[5,5,4,9,9,1],"isMain":false},{"metricName":"Misassembled contigs length","quality":"Less is better","values":[6704,9494,13057,52485,20227,860],"isMain":true},{"metricName":"# possibly misassembled contigs","quality":"Less is better","values":[0,0,0,0,0,0],"isMain":false},{"metricName":" # possible misassemblies","quality":"Less is better","values":[0,0,0,0,0,0],"isMain":false},{"metricName":"# local misassemblies","quality":"Less is better","values":[4,4,3,4,7,1],"isMain":false},{"metricName":"# scaffold gap ext. mis.","quality":"Less is better","values":[0,0,0,0,0,0],"isMain":false},{"metricName":"# scaffold gap loc. mis.","quality":"Less is better","values":[0,0,0,0,0,0],"isMain":false},{"metricName":"# unaligned mis. contigs","quality":"Less is better","values":[0,0,2,2,4,0],"isMain":false}]],["Unaligned",[{"metricName":"# fully unaligned contigs","quality":"Less is better","values":[0,1,2,1,2,0],"isMain":false},{"metricName":"Fully unaligned length","quality":"Less is better","values":[0,820,1127,600,1701,0],"isMain":false},{"metricName":"# partially unaligned contigs","quality":"Less is better","values":[1,3,2,4,14,0],"isMain":false},{"metricName":"Partially unaligned length","quality":"Less is better","values":[955,2007,1587,3291,67131,0],"isMain":false}]],["Mismatches",[{"metricName":"# mismatches","quality":"Less is better","values":[4374,7556,7387,8128,6525,2456],"isMain":false},{"metricName":"# indels","quality":"Less is better","values":[25,109,38,53,75,13],"isMain":false},{"metricName":"Indels length","quality":"Less is better","values":[115,547,163,220,236,45],"isMain":false},{"metricName":"# mismatches per 100 kbp","quality":"Less is better","values":["558.93","924.49","849.13","957.04","837.34","502.55"],"isMain":true},{"metricName":"# indels per 100 kbp","quality":"Less is better","values":["3.19","13.34","4.37","6.24","9.62","2.66"],"isMain":true},{"metricName":" # indels (<= 5 bp)","quality":"Less is better","values":[19,95,32,46,69,11],"isMain":false},{"metricName":" # indels (> 5 bp)","quality":"Less is better","values":[6,14,6,7,6,2],"isMain":false},{"metricName":"# N's","quality":"Less is better","values":[0,0,0,0,0,0],"isMain":false},{"metricName":"# N's per 100 kbp","quality":"Less is better","values":["0.00","0.00","0.00","0.00","0.00","0.00"],"isMain":true}]],["Statistics without reference",[{"metricName":"# contigs","quality":"Equal","values":[387,371,433,445,381,259],"isMain":true},{"metricName":"# contigs (>= 1000 bp)","quality":"Equal","values":[195,216,192,180,195,95],"isMain":false},{"metricName":"# contigs (>= 5000 bp)","quality":"Equal","values":[36,38,37,43,33,10],"isMain":false},{"metricName":"# contigs (>= 10000 bp)","quality":"Equal","values":[10,17,21,19,18,7],"isMain":false},{"metricName":"# contigs (>= 25000 bp)","quality":"Equal","values":[1,0,1,1,3,3],"isMain":false},{"metricName":"# contigs (>= 50000 bp)","quality":"Equal","values":[0,0,0,0,0,2],"isMain":false},{"metricName":"Largest contig","quality":"More is better","values":[32721,24011,27264,35867,40526,87104],"isMain":true},{"metricName":"Total length","quality":"More is better","values":[800363,841841,893676,883654,872400,497043],"isMain":true},{"metricName":"Total length (>= 1000 bp)","quality":"More is better","values":[664006,734457,726986,701549,741997,380963],"isMain":true},{"metricName":"Total length (>= 5000 bp)","quality":"More is better","values":[348295,402150,434947,466195,431736,252418],"isMain":false},{"metricName":"Total length (>= 10000 bp)","quality":"More is better","values":[174928,254200,322004,296468,317294,234336],"isMain":true},{"metricName":"Total length (>= 25000 bp)","quality":"More is better","values":[32721,0,27264,35867,96761,163485],"isMain":false},{"metricName":"Total length (>= 50000 bp)","quality":"More is better","values":[0,0,0,0,0,137399],"isMain":true},{"metricName":"N50","quality":"More is better","values":[4087,4243,4587,5492,4959,5091],"isMain":false},{"metricName":"N75","quality":"More is better","values":[1319,1501,1254,1200,1515,1059],"isMain":false},{"metricName":"L50","quality":"Less is better","values":[48,43,40,39,34,10],"isMain":false},{"metricName":"L75","quality":"Less is better","values":[141,133,141,145,124,88],"isMain":false},{"metricName":"GC (%)","quality":"Equal","values":["46.89","46.83","47.52","47.21","46.59","45.58"],"isMain":false}]],["K-mer-based statistics",[]],["Predicted genes",[]],["Similarity statistics",[]],["Reference statistics",[{"metricName":"Reference length","quality":"Equal","values":[5455948,5455948,5455948,5455948,5455948,5455948],"isMain":false},{"metricName":"Reference fragments","quality":"Equal","values":[152,152,152,152,152,152],"isMain":false},{"metricName":"Reference GC (%)","quality":"Equal","values":["48.93","48.93","48.93","48.93","48.93","48.93"],"isMain":false}]]],"subreferences":[],"subreports":[],"minContig":500},{"date":"10 February 2020, Monday, 18:53:08","assembliesNames":["megahit-113-ms","metaSPAdes"],"referenceName":"GCF\_000235445.1\_John\_igna\_ATCC\_51276\_V1","order":[0,1],"report":[["Genome statistics",[{"metricName":"Genome fraction (%)","quality":"More is better","values":["0.086","0.004"],"isMain":true},{"metricName":"Duplication ratio","quality":"Less is better","values":["1.000","1.000"],"isMain":true},{"metricName":"Largest alignment","quality":"More is better","values":[1460,97],"isMain":true},{"metricName":"Total aligned length","quality":"More is better","values":[2310,97],"isMain":true},{"metricName":"NGA50","quality":"More is better","values":[null,null],"isMain":true}]],["Reads mapping",[]],["Misassemblies",[{"metricName":"# misassemblies","quality":"Less is better","values":[0,0],"isMain":true},{"metricName":" # relocations","quality":"Less is better","values":[0,0],"isMain":false},{"metricName":" # translocations","quality":"Less is better","values":[0,0],"isMain":false},{"metricName":" # inversions","quality":"Less is better","values":[0,0],"isMain":false},{"metricName":" # interspecies translocations","quality":"Less is better","values":[1,1],"isMain":false},{"metricName":"# misassembled contigs","quality":"Less is better","values":[0,0],"isMain":false},{"metricName":"Misassembled contigs length","quality":"Less is better","values":[0,0],"isMain":true},{"metricName":"# possibly misassembled contigs","quality":"Less is better","values":[0,0],"isMain":false},{"metricName":" # possible misassemblies","quality":"Less is better","values":[0,0],"isMain":false},{"metricName":"# local misassemblies","quality":"Less is better","values":[0,0],"isMain":false},{"metricName":"# scaffold gap ext. mis.","quality":"Less is better","values":[0,0],"isMain":false},{"metricName":"# scaffold gap loc. mis.","quality":"Less is better","values":[0,0],"isMain":false},{"metricName":"# unaligned mis. contigs","quality":"Less is better","values":[0,0],"isMain":false}]],["Unaligned",[{"metricName":"# fully unaligned contigs","quality":"Less is better","values":[0,0],"isMain":false},{"metricName":"Fully unaligned length","quality":"Less is better","values":[0,0],"isMain":false},{"metricName":"# partially unaligned contigs","quality":"Less is better","values":[1,1],"isMain":false},{"metricName":"Partially unaligned length","quality":"Less is better","values":[2517,699],"isMain":false}]],["Mismatches",[{"metricName":"# mismatches","quality":"Less is better","values":[98,4],"isMain":false},{"metricName":"# indels","quality":"Less is better","values":[0,0],"isMain":false},{"metricName":"Indels length","quality":"Less is better","values":[0,0],"isMain":false},{"metricName":"# mismatches per 100 kbp","quality":"Less is better","values":["4242.42","4123.71"],"isMain":true},{"metricName":"# indels per 100 kbp","quality":"Less is better","values":["0.00","0.00"],"isMain":true},{"metricName":" # indels (<= 5 bp)","quality":"Less is better","values":[0,0],"isMain":false},{"metricName":" # indels (> 5 bp)","quality":"Less is better","values":[0,0],"isMain":false},{"metricName":"# N's","quality":"Less is better","values":[0,0],"isMain":false},{"metricName":"# N's per 100 kbp","quality":"Less is better","values":["0.00","0.00"],"isMain":true}]],["Statistics without reference",[{"metricName":"# contigs","quality":"Equal","values":[2,1],"isMain":true},{"metricName":"# contigs (>= 1000 bp)","quality":"Equal","values":[1,0],"isMain":false},{"metricName":"# contigs (>= 5000 bp)","quality":"Equal","values":[0,0],"isMain":false},{"metricName":"# contigs (>= 10000 bp)","quality":"Equal","values":[0,0],"isMain":false},{"metricName":"# contigs (>= 25000 bp)","quality":"Equal","values":[0,0],"isMain":false},{"metricName":"# contigs (>= 50000 bp)","quality":"Equal","values":[0,0],"isMain":false},{"metricName":"Largest contig","quality":"More is better","values":[3977,796],"isMain":true},{"metricName":"Total length","quality":"More is better","values":[4827,796],"isMain":true},{"metricName":"Total length (>= 1000 bp)","quality":"More is better","values":[3977,0],"isMain":true},{"metricName":"Total length (>= 5000 bp)","quality":"More is better","values":[0,0],"isMain":false},{"metricName":"Total length (>= 10000 bp)","quality":"More is better","values":[0,0],"isMain":true},{"metricName":"Total length (>= 25000 bp)","quality":"More is better","values":[0,0],"isMain":false},{"metricName":"Total length (>= 50000 bp)","quality":"More is better","values":[0,0],"isMain":true},{"metricName":"N50","quality":"More is better","values":[3977,796],"isMain":false},{"metricName":"N75","quality":"More is better","values":[3977,796],"isMain":false},{"metricName":"L50","quality":"Less is better","values":[1,1],"isMain":false},{"metricName":"L75","quality":"Less is better","values":[1,1],"isMain":false},{"metricName":"GC (%)","quality":"Equal","values":["40.94","39.70"],"isMain":false}]],["K-mer-based statistics",[]],["Predicted genes",[]],["Similarity statistics",[]],["Reference statistics",[{"metricName":"Reference length","quality":"Equal","values":[2672402,2672402],"isMain":false},{"metricName":"Reference fragments","quality":"Equal","values":[82,82],"isMain":false},{"metricName":"Reference GC (%)","quality":"Equal","values":["35.74","35.74"],"isMain":false}]]],"subreferences":[],"subreports":[],"minContig":500},{"date":"10 February 2020, Monday, 19:33:38","assembliesNames":["megahit-103-df","megahit-113-df","megahit-113-ml","megahit-113-ms","megahit-129-df","metaSPAdes"],"referenceName":"GCF\_000236865.1\_ASM23686v1","order":[0,1,2,3,4,5],"report":[["Genome statistics",[{"metricName":"Genome fraction (%)","quality":"More is better","values":["3.070","3.687","4.808","5.248","4.094","1.845"],"isMain":true},{"metricName":"Duplication ratio","quality":"Less is better","values":["1.041","1.025","1.014","1.036","1.020","1.037"],"isMain":true},{"metricName":"Largest alignment","quality":"More is better","values":[3539,5467,1963,1594,6116,761],"isMain":true},{"metricName":"Total aligned length","quality":"More is better","values":[72333,86955,113249,123926,96768,43509],"isMain":true},{"metricName":"NA50","quality":"More is better","values":[570,591,576,560,568,564],"isMain":false},{"metricName":"NA75","quality":"More is better","values":[517,528,525,517,520,516],"isMain":false},{"metricName":"NGA50","quality":"More is better","values":[null,null,null,null,null,null],"isMain":true},{"metricName":"LA50","quality":"Less is better","values":[53,56,82,97,70,37],"isMain":false},{"metricName":"LA75","quality":"Less is better","values":[88,97,136,159,119,58],"isMain":false}]],["Reads mapping",[]],["Misassemblies",[{"metricName":"# misassemblies","quality":"Less is better","values":[0,1,1,0,1,0],"isMain":true},{"metricName":" # relocations","quality":"Less is better","values":[0,1,0,0,1,0],"isMain":false},{"metricName":" # translocations","quality":"Less is better","values":[0,0,1,0,0,0],"isMain":false},{"metricName":" # inversions","quality":"Less is better","values":[0,0,0,0,0,0],"isMain":false},{"metricName":" # interspecies translocations","quality":"Less is better","values":[12,6,4,16,9,6],"isMain":false},{"metricName":"# misassembled contigs","quality":"Less is better","values":[0,1,1,0,1,0],"isMain":false},{"metricName":"Misassembled contigs length","quality":"Less is better","values":[0,604,550,0,604,0],"isMain":true},{"metricName":"# possibly misassembled contigs","quality":"Less is better","values":[0,0,0,0,0,0],"isMain":false},{"metricName":" # possible misassemblies","quality":"Less is better","values":[0,0,0,0,0,0],"isMain":false},{"metricName":"# local misassemblies","quality":"Less is better","values":[2,2,0,1,2,0],"isMain":false},{"metricName":"# scaffold gap ext. mis.","quality":"Less is better","values":[0,0,0,0,0,0],"isMain":false},{"metricName":"# scaffold gap loc. mis.","quality":"Less is better","values":[0,0,0,0,0,0],"isMain":false},{"metricName":"# unaligned mis. contigs","quality":"Less is better","values":[0,0,0,1,0,0],"isMain":false}]],["Unaligned",[{"metricName":"# fully unaligned contigs","quality":"Less is better","values":[0,0,1,0,0,0],"isMain":false},{"metricName":"Fully unaligned length","quality":"Less is better","values":[0,0,1167,0,0,0],"isMain":false},{"metricName":"# partially unaligned contigs","quality":"Less is better","values":[1,2,2,4,6,1],"isMain":false},{"metricName":"Partially unaligned length","quality":"Less is better","values":[1478,1919,1592,4225,8452,583],"isMain":false}]],["Mismatches",[{"metricName":"# mismatches","quality":"Less is better","values":[1191,1581,2237,2665,1776,746],"isMain":false},{"metricName":"# indels","quality":"Less is better","values":[0,4,1,0,8,0],"isMain":false},{"metricName":"Indels length","quality":"Less is better","values":[0,4,2,0,8,0],"isMain":false},{"metricName":"# mismatches per 100 kbp","quality":"Less is better","values":["1647.71","1821.13","1976.31","2156.74","1842.46","1717.03"],"isMain":true},{"metricName":"# indels per 100 kbp","quality":"Less is better","values":["0.00","4.61","0.88","0.00","8.30","0.00"],"isMain":true},{"metricName":" # indels (<= 5 bp)","quality":"Less is better","values":[0,4,1,0,8,0],"isMain":false},{"metricName":" # indels (> 5 bp)","quality":"Less is better","values":[0,0,0,0,0,0],"isMain":false},{"metricName":"# N's","quality":"Less is better","values":[0,0,0,0,0,0],"isMain":false},{"metricName":"# N's per 100 kbp","quality":"Less is better","values":["0.00","0.00","0.00","0.00","0.00","0.00"],"isMain":true}]],["Statistics without reference",[{"metricName":"# contigs","quality":"Equal","values":[121,136,188,214,155,78],"isMain":true},{"metricName":"# contigs (>= 1000 bp)","quality":"Equal","values":[2,6,6,8,5,0],"isMain":false},{"metricName":"# contigs (>= 5000 bp)","quality":"Equal","values":[0,1,0,0,1,0],"isMain":false},{"metricName":"# contigs (>= 10000 bp)","quality":"Equal","values":[0,0,0,0,0,0],"isMain":false},{"metricName":"# contigs (>= 25000 bp)","quality":"Equal","values":[0,0,0,0,0,0],"isMain":false},{"metricName":"# contigs (>= 50000 bp)","quality":"Equal","values":[0,0,0,0,0,0],"isMain":false},{"metricName":"Largest contig","quality":"More is better","values":[3542,5635,1973,1705,7836,761],"isMain":true},{"metricName":"Total length","quality":"More is better","values":[76738,90941,117587,132178,106757,45621],"isMain":true},{"metricName":"Total length (>= 1000 bp)","quality":"More is better","values":[5164,11629,8535,11173,15091,0],"isMain":true},{"metricName":"Total length (>= 5000 bp)","quality":"More is better","values":[0,5635,0,0,7836,0],"isMain":false},{"metricName":"Total length (>= 10000 bp)","quality":"More is better","values":[0,0,0,0,0,0],"isMain":true},{"metricName":"Total length (>= 25000 bp)","quality":"More is better","values":[0,0,0,0,0,0],"isMain":false},{"metricName":"Total length (>= 50000 bp)","quality":"More is better","values":[0,0,0,0,0,0],"isMain":true},{"metricName":"N50","quality":"More is better","values":[583,604,591,579,617,583],"isMain":false},{"metricName":"N75","quality":"More is better","values":[537,540,534,531,543,533],"isMain":false},{"metricName":"L50","quality":"Less is better","values":[50,53,79,91,58,36],"isMain":false},{"metricName":"L75","quality":"Less is better","values":[84,93,132,150,104,56],"isMain":false},{"metricName":"GC (%)","quality":"Equal","values":["61.40","61.02","61.71","61.24","61.40","60.11"],"isMain":false}]],["K-mer-based statistics",[]],["Predicted genes",[]],["Similarity statistics",[]],["Reference statistics",[{"metricName":"Reference length","quality":"Equal","values":[2354438,2354438,2354438,2354438,2354438,2354438],"isMain":false},{"metricName":"Reference fragments","quality":"Equal","values":[84,84,84,84,84,84],"isMain":false},{"metricName":"Reference GC (%)","quality":"Equal","values":["62.06","62.06","62.06","62.06","62.06","62.06"],"isMain":false}]]],"subreferences":[],"subreports":[],"minContig":500},{"date":"10 February 2020, Monday, 19:33:58","assembliesNames":["megahit-103-df","megahit-113-df","megahit-113-ml","megahit-113-ms","megahit-129-df","metaSPAdes"],"referenceName":"GCF\_000237085.1\_ASM23708v1","order":[0,1,2,3,4,5],"report":[["Genome statistics",[{"metricName":"Genome fraction (%)","quality":"More is better","values":["14.423","17.993","17.962","20.539","20.442","10.915"],"isMain":true},{"metricName":"Duplication ratio","quality":"Less is better","values":["1.033","1.031","1.021","1.035","1.030","1.044"],"isMain":true},{"metricName":"Largest alignment","quality":"More is better","values":[3169,5983,3422,2985,4833,2705],"isMain":true},{"metricName":"Total aligned length","quality":"More is better","values":[712449,887020,888214,1018782,1010553,538368],"isMain":true},{"metricName":"NA50","quality":"More is better","values":[634,644,637,645,664,599],"isMain":false},{"metricName":"NA75","quality":"More is better","values":[546,540,556,550,554,528],"isMain":false},{"metricName":"NGA50","quality":"More is better","values":[null,null,null,null,null,null],"isMain":true},{"metricName":"LA50","quality":"Less is better","values":[444,545,542,632,598,368],"isMain":false},{"metricName":"LA75","quality":"Less is better","values":[763,951,920,1074,1043,621],"isMain":false}]],["Reads mapping",[]],["Misassemblies",[{"metricName":"# misassemblies","quality":"Less is better","values":[12,21,5,7,19,3],"isMain":true},{"metricName":" # relocations","quality":"Less is better","values":[10,21,4,7,19,3],"isMain":false},{"metricName":" # translocations","quality":"Less is better","values":[0,0,0,0,0,0],"isMain":false},{"metricName":" # inversions","quality":"Less is better","values":[2,0,1,0,0,0],"isMain":false},{"metricName":" # interspecies translocations","quality":"Less is better","values":[62,79,8,54,70,56],"isMain":false},{"metricName":"# misassembled contigs","quality":"Less is better","values":[12,21,5,7,19,3],"isMain":false},{"metricName":"Misassembled contigs length","quality":"Less is better","values":[10493,26843,7885,8958,24937,2921],"isMain":true},{"metricName":"# possibly misassembled contigs","quality":"Less is better","values":[0,1,0,1,0,0],"isMain":false},{"metricName":" # possible misassemblies","quality":"Less is better","values":[0,1,0,1,0,0],"isMain":false},{"metricName":"# local misassemblies","quality":"Less is better","values":[2,3,1,3,4,0],"isMain":false},{"metricName":"# scaffold gap ext. mis.","quality":"Less is better","values":[0,0,0,0,0,0],"isMain":false},{"metricName":"# scaffold gap loc. mis.","quality":"Less is better","values":[0,0,0,0,0,0],"isMain":false},{"metricName":"# unaligned mis. contigs","quality":"Less is better","values":[0,0,0,0,0,0],"isMain":false}]],["Unaligned",[{"metricName":"# fully unaligned contigs","quality":"Less is better","values":[1,1,1,0,1,0],"isMain":false},{"metricName":"Fully unaligned length","quality":"Less is better","values":[682,830,542,0,771,0],"isMain":false},{"metricName":"# partially unaligned contigs","quality":"Less is better","values":[16,26,1,10,20,12],"isMain":false},{"metricName":"Partially unaligned length","quality":"Less is better","values":[15706,45182,623,10449,38535,9589],"isMain":false}]],["Mismatches",[{"metricName":"# mismatches","quality":"Less is better","values":[10825,13170,16957,20411,15286,8619],"isMain":false},{"metricName":"# indels","quality":"Less is better","values":[31,27,40,54,45,18],"isMain":false},{"metricName":"Indels length","quality":"Less is better","values":[46,41,79,109,84,157],"isMain":false},{"metricName":"# mismatches per 100 kbp","quality":"Less is better","values":["1532.39","1494.49","1927.49","2029.10","1526.78","1612.26"],"isMain":true},{"metricName":"# indels per 100 kbp","quality":"Less is better","values":["4.39","3.06","4.55","5.37","4.49","3.37"],"isMain":true},{"metricName":" # indels (<= 5 bp)","quality":"Less is better","values":[30,26,38,50,41,15],"isMain":false},{"metricName":" # indels (> 5 bp)","quality":"Less is better","values":[1,1,2,4,4,3],"isMain":false},{"metricName":"# N's","quality":"Less is better","values":[0,0,0,0,0,0],"isMain":false},{"metricName":"# N's per 100 kbp","quality":"Less is better","values":["0.00","0.00","0.00","0.00","0.00","0.00"],"isMain":true}]],["Statistics without reference",[{"metricName":"# contigs","quality":"Equal","values":[1074,1304,1330,1529,1458,862],"isMain":true},{"metricName":"# contigs (>= 1000 bp)","quality":"Equal","values":[86,127,79,97,161,45],"isMain":false},{"metricName":"# contigs (>= 5000 bp)","quality":"Equal","values":[0,3,0,0,2,0],"isMain":false},{"metricName":"# contigs (>= 10000 bp)","quality":"Equal","values":[0,1,0,0,1,0],"isMain":false},{"metricName":"# contigs (>= 25000 bp)","quality":"Equal","values":[0,0,0,0,0,0],"isMain":false},{"metricName":"# contigs (>= 50000 bp)","quality":"Equal","values":[0,0,0,0,0,0],"isMain":false},{"metricName":"Largest contig","quality":"More is better","values":[3169,13119,3427,3298,18533,2914],"isMain":true},{"metricName":"Total length","quality":"More is better","values":[746287,954229,899641,1051070,1071010,567848],"isMain":true},{"metricName":"Total length (>= 1000 bp)","quality":"More is better","values":[114784,195035,103136,124626,230643,61025],"isMain":true},{"metricName":"Total length (>= 5000 bp)","quality":"More is better","values":[0,28128,0,0,23947,0],"isMain":false},{"metricName":"Total length (>= 10000 bp)","quality":"More is better","values":[0,13119,0,0,18533,0],"isMain":true},{"metricName":"Total length (>= 25000 bp)","quality":"More is better","values":[0,0,0,0,0,0],"isMain":false},{"metricName":"Total length (>= 50000 bp)","quality":"More is better","values":[0,0,0,0,0,0],"isMain":true},{"metricName":"N50","quality":"More is better","values":[666,691,648,664,701,624],"isMain":false},{"metricName":"N75","quality":"More is better","values":[565,573,565,571,581,552],"isMain":false},{"metricName":"L50","quality":"Less is better","values":[417,479,533,607,539,350],"isMain":false},{"metricName":"L75","quality":"Less is better","values":[724,858,907,1036,961,592],"isMain":false},{"metricName":"GC (%)","quality":"Equal","values":["36.45","36.63","36.15","36.16","36.29","36.46"],"isMain":false}]],["K-mer-based statistics",[]],["Predicted genes",[]],["Similarity statistics",[]],["Reference statistics",[{"metricName":"Reference length","quality":"Equal","values":[4897678,4897678,4897678,4897678,4897678,4897678],"isMain":false},{"metricName":"Reference fragments","quality":"Equal","values":[1,1,1,1,1,1],"isMain":false},{"metricName":"Reference GC (%)","quality":"Equal","values":["35.72","35.72","35.72","35.72","35.72","35.72"],"isMain":false}]]],"subreferences":[],"subreports":[],"minContig":500},{"date":"10 February 2020, Monday, 19:34:30","assembliesNames":["megahit-103-df","megahit-113-df","megahit-113-ml","megahit-113-ms","megahit-129-df","metaSPAdes"],"referenceName":"GCF\_000238055.2\_Oribacterium\_sp\_ACB1\_V2","order":[0,1,2,3,4,5],"report":[["Genome statistics",[{"metricName":"Genome fraction (%)","quality":"More is better","values":["98.802","98.921","98.768","98.905","98.941","98.147"],"isMain":true},{"metricName":"Duplication ratio","quality":"Less is better","values":["1.009","1.008","1.053","1.047","1.010","1.007"],"isMain":true},{"metricName":"Largest alignment","quality":"More is better","values":[123969,115384,21043,29497,101549,70440],"isMain":true},{"metricName":"Total aligned length","quality":"More is better","values":[2462259,2461391,2564115,2549160,2466329,2441061],"isMain":true},{"metricName":"NG50","quality":"More is better","values":[23687,39970,6401,7406,36549,24951],"isMain":false},{"metricName":"NG75","quality":"More is better","values":[14442,21537,3807,4272,22741,13876],"isMain":false},{"metricName":"NA50","quality":"More is better","values":[23687,39970,5859,7265,36523,24944],"isMain":false},{"metricName":"NA75","quality":"More is better","values":[14427,20484,3328,3955,22741,14839],"isMain":false},{"metricName":"NGA50","quality":"More is better","values":[23687,39970,6262,7390,36523,24936],"isMain":true},{"metricName":"NGA75","quality":"More is better","values":[14427,20484,3685,4237,22741,13836],"isMain":false},{"metricName":"LG50","quality":"Less is better","values":[35,21,128,109,21,34],"isMain":false},{"metricName":"LG75","quality":"Less is better","values":[68,42,254,218,41,65],"isMain":false},{"metricName":"LA50","quality":"Less is better","values":[35,21,143,117,21,33],"isMain":false},{"metricName":"LA75","quality":"Less is better","values":[68,42,290,238,41,64],"isMain":false},{"metricName":"LGA50","quality":"Less is better","values":[35,21,132,110,21,34],"isMain":true},{"metricName":"LGA75","quality":"Less is better","values":[68,42,262,220,41,65],"isMain":false}]],["Reads mapping",[]],["Misassemblies",[{"metricName":"# misassemblies","quality":"Less is better","values":[0,1,3,2,0,0],"isMain":true},{"metricName":" # relocations","quality":"Less is better","values":[0,0,0,0,0,0],"isMain":false},{"metricName":" # translocations","quality":"Less is better","values":[0,1,2,2,0,0],"isMain":false},{"metricName":" # inversions","quality":"Less is better","values":[0,0,1,0,0,0],"isMain":false},{"metricName":" # interspecies translocations","quality":"Less is better","values":[2,4,7,21,5,0],"isMain":false},{"metricName":"# misassembled contigs","quality":"Less is better","values":[0,1,3,2,0,0],"isMain":false},{"metricName":"Misassembled contigs length","quality":"Less is better","values":[0,2136,27209,22988,0,0],"isMain":true},{"metricName":"# possibly misassembled contigs","quality":"Less is better","values":[0,0,0,0,0,0],"isMain":false},{"metricName":" # possible misassemblies","quality":"Less is better","values":[0,0,0,0,0,0],"isMain":false},{"metricName":"# local misassemblies","quality":"Less is better","values":[1,1,3,4,1,1],"isMain":false},{"metricName":"# scaffold gap ext. mis.","quality":"Less is better","values":[0,0,0,0,0,0],"isMain":false},{"metricName":"# scaffold gap loc. mis.","quality":"Less is better","values":[0,0,0,0,0,0],"isMain":false},{"metricName":"# unaligned mis. contigs","quality":"Less is better","values":[0,0,1,1,0,0],"isMain":false}]],["Unaligned",[{"metricName":"# fully unaligned contigs","quality":"Less is better","values":[1,0,0,0,0,0],"isMain":false},{"metricName":"Fully unaligned length","quality":"Less is better","values":[1093,0,0,0,0,0],"isMain":false},{"metricName":"# partially unaligned contigs","quality":"Less is better","values":[0,1,2,7,2,0],"isMain":false},{"metricName":"Partially unaligned length","quality":"Less is better","values":[0,4895,28478,8074,1909,0],"isMain":false}]],["Mismatches",[{"metricName":"# mismatches","quality":"Less is better","values":[2214,2173,13513,12878,2028,1390],"isMain":false},{"metricName":"# indels","quality":"Less is better","values":[17,22,24,30,32,24],"isMain":false},{"metricName":"Indels length","quality":"Less is better","values":[254,178,196,288,869,133],"isMain":false},{"metricName":"# mismatches per 100 kbp","quality":"Less is better","values":["90.70","88.91","553.78","527.02","82.96","57.32"],"isMain":true},{"metricName":"# indels per 100 kbp","quality":"Less is better","values":["0.70","0.90","0.98","1.23","1.31","0.99"],"isMain":true},{"metricName":" # indels (<= 5 bp)","quality":"Less is better","values":[11,15,20,23,9,20],"isMain":false},{"metricName":" # indels (> 5 bp)","quality":"Less is better","values":[6,7,4,7,23,4],"isMain":false},{"metricName":"# N's","quality":"Less is better","values":[0,0,0,0,0,0],"isMain":false},{"metricName":"# N's per 100 kbp","quality":"Less is better","values":["0.00","0.00","0.00","0.00","0.00","0.00"],"isMain":true}]],["Statistics without reference",[{"metricName":"# contigs","quality":"Equal","values":[177,130,664,571,148,162],"isMain":true},{"metricName":"# contigs (>= 1000 bp)","quality":"Equal","values":[156,109,555,490,126,149],"isMain":false},{"metricName":"# contigs (>= 5000 bp)","quality":"Equal","values":[121,87,184,182,85,118],"isMain":false},{"metricName":"# contigs (>= 10000 bp)","quality":"Equal","values":[88,66,44,59,59,85],"isMain":false},{"metricName":"# contigs (>= 25000 bp)","quality":"Equal","values":[30,35,0,1,37,31],"isMain":false},{"metricName":"# contigs (>= 50000 bp)","quality":"Equal","values":[3,14,0,0,13,4],"isMain":false},{"metricName":"Largest contig","quality":"More is better","values":[123969,115384,21043,29510,101551,70440],"isMain":true},{"metricName":"Total length","quality":"More is better","values":[2464698,2469104,2598256,2567648,2471064,2442467],"isMain":true},{"metricName":"Total length (>= 1000 bp)","quality":"More is better","values":[2449309,2453329,2520396,2511915,2455242,2432558],"isMain":true},{"metricName":"Total length (>= 5000 bp)","quality":"More is better","values":[2352799,2399921,1554646,1689128,2347925,2335923],"isMain":false},{"metricName":"Total length (>= 10000 bp)","quality":"More is better","values":[2101557,2244665,574079,824061,2161105,2097506],"isMain":true},{"metricName":"Total length (>= 25000 bp)","quality":"More is better","values":[1134175,1699549,0,29510,1777134,1179608],"isMain":false},{"metricName":"Total length (>= 50000 bp)","quality":"More is better","values":[252258,950490,0,0,932160,251941],"isMain":true},{"metricName":"N50","quality":"More is better","values":[23687,39970,6062,7300,36549,24959],"isMain":false},{"metricName":"N75","quality":"More is better","values":[14442,21537,3451,4005,22741,14847],"isMain":false},{"metricName":"L50","quality":"Less is better","values":[35,21,138,116,21,33],"isMain":false},{"metricName":"L75","quality":"Less is better","values":[68,42,280,236,41,64],"isMain":false},{"metricName":"GC (%)","quality":"Equal","values":["42.12","42.11","42.23","42.19","42.14","42.13"],"isMain":false}]],["K-mer-based statistics",[]],["Predicted genes",[]],["Similarity statistics",[]],["Reference statistics",[{"metricName":"Reference length","quality":"Equal","values":[2470590,2470590,2470590,2470590,2470590,2470590],"isMain":false},{"metricName":"Reference fragments","quality":"Equal","values":[4,4,4,4,4,4],"isMain":false},{"metricName":"Reference GC (%)","quality":"Equal","values":["42.17","42.17","42.17","42.17","42.17","42.17"],"isMain":false}]]],"subreferences":[],"subreports":[],"minContig":500},{"date":"10 February 2020, Monday, 19:34:58","assembliesNames":["megahit-103-df","megahit-113-df","megahit-113-ml","megahit-113-ms","megahit-129-df","metaSPAdes"],"referenceName":"GCF\_000238075.1\_Oribacterium\_sp\_ACB7\_V1","order":[0,1,2,3,4,5],"report":[["Genome statistics",[{"metricName":"Genome fraction (%)","quality":"More is better","values":["0.431","0.374","1.102","0.963","0.591","0.267"],"isMain":true},{"metricName":"Duplication ratio","quality":"Less is better","values":["1.068","1.216","1.022","1.067","1.035","1.299"],"isMain":true},{"metricName":"Largest alignment","quality":"More is better","values":[754,754,794,710,755,794],"isMain":true},{"metricName":"Total aligned length","quality":"More is better","values":[10847,9417,27729,24222,14863,6707],"isMain":true},{"metricName":"NA50","quality":"More is better","values":[514,529,574,552,549,481],"isMain":false},{"metricName":"NA75","quality":"More is better","values":[null,null,533,506,521,null],"isMain":false},{"metricName":"NGA50","quality":"More is better","values":[null,null,null,null,null,null],"isMain":true},{"metricName":"LA50","quality":"Less is better","values":[14,11,22,23,13,9],"isMain":false},{"metricName":"LA75","quality":"Less is better","values":[null,null,35,36,21,null],"isMain":false}]],["Reads mapping",[]],["Misassemblies",[{"metricName":"# misassemblies","quality":"Less is better","values":[0,0,0,0,0,0],"isMain":true},{"metricName":" # relocations","quality":"Less is better","values":[0,0,0,0,0,0],"isMain":false},{"metricName":" # translocations","quality":"Less is better","values":[0,0,0,0,0,0],"isMain":false},{"metricName":" # inversions","quality":"Less is better","values":[0,0,0,0,0,0],"isMain":false},{"metricName":" # interspecies translocations","quality":"Less is better","values":[5,7,1,5,3,7],"isMain":false},{"metricName":"# misassembled contigs","quality":"Less is better","values":[0,0,0,0,0,0],"isMain":false},{"metricName":"Misassembled contigs length","quality":"Less is better","values":[0,0,0,0,0,0],"isMain":true},{"metricName":"# possibly misassembled contigs","quality":"Less is better","values":[0,0,0,0,0,0],"isMain":false},{"metricName":" # possible misassemblies","quality":"Less is better","values":[0,0,0,0,0,0],"isMain":false},{"metricName":"# local misassemblies","quality":"Less is better","values":[0,0,0,0,0,0],"isMain":false},{"metricName":"# scaffold gap ext. mis.","quality":"Less is better","values":[0,0,0,0,0,0],"isMain":false},{"metricName":"# scaffold gap loc. mis.","quality":"Less is better","values":[0,0,0,0,0,0],"isMain":false},{"metricName":"# unaligned mis. contigs","quality":"Less is better","values":[0,0,0,0,0,0],"isMain":false}]],["Unaligned",[{"metricName":"# fully unaligned contigs","quality":"Less is better","values":[0,0,0,0,0,0],"isMain":false},{"metricName":"Fully unaligned length","quality":"Less is better","values":[0,0,0,0,0,0],"isMain":false},{"metricName":"# partially unaligned contigs","quality":"Less is better","values":[3,2,0,1,1,1],"isMain":false},{"metricName":"Partially unaligned length","quality":"Less is better","values":[4054,1260,0,1443,515,1421],"isMain":false}]],["Mismatches",[{"metricName":"# mismatches","quality":"Less is better","values":[186,149,540,469,250,118],"isMain":false},{"metricName":"# indels","quality":"Less is better","values":[0,0,0,1,2,0],"isMain":false},{"metricName":"Indels length","quality":"Less is better","values":[0,0,0,1,3,0],"isMain":false},{"metricName":"# mismatches per 100 kbp","quality":"Less is better","values":["1714.76","1582.24","1947.42","1936.34","1681.92","1759.36"],"isMain":true},{"metricName":"# indels per 100 kbp","quality":"Less is better","values":["0.00","0.00","0.00","4.13","13.46","0.00"],"isMain":true},{"metricName":" # indels (<= 5 bp)","quality":"Less is better","values":[0,0,0,1,2,0],"isMain":false},{"metricName":" # indels (> 5 bp)","quality":"Less is better","values":[0,0,0,0,0,0],"isMain":false},{"metricName":"# N's","quality":"Less is better","values":[0,0,0,0,0,0],"isMain":false},{"metricName":"# N's per 100 kbp","quality":"Less is better","values":["0.00","0.00","0.00","0.00","0.00","0.00"],"isMain":true}]],["Statistics without reference",[{"metricName":"# contigs","quality":"Equal","values":[22,21,47,46,27,16],"isMain":true},{"metricName":"# contigs (>= 1000 bp)","quality":"Equal","values":[1,0,0,1,0,1],"isMain":false},{"metricName":"# contigs (>= 5000 bp)","quality":"Equal","values":[0,0,0,0,0,0],"isMain":false},{"metricName":"# contigs (>= 10000 bp)","quality":"Equal","values":[0,0,0,0,0,0],"isMain":false},{"metricName":"# contigs (>= 25000 bp)","quality":"Equal","values":[0,0,0,0,0,0],"isMain":false},{"metricName":"# contigs (>= 50000 bp)","quality":"Equal","values":[0,0,0,0,0,0],"isMain":false},{"metricName":"Largest contig","quality":"More is better","values":[2950,862,794,1647,755,1549],"isMain":true},{"metricName":"Total length","quality":"More is better","values":[15641,12708,28349,27279,15904,10136],"isMain":true},{"metricName":"Total length (>= 1000 bp)","quality":"More is better","values":[2950,0,0,1647,0,1549],"isMain":true},{"metricName":"Total length (>= 5000 bp)","quality":"More is better","values":[0,0,0,0,0,0],"isMain":false},{"metricName":"Total length (>= 10000 bp)","quality":"More is better","values":[0,0,0,0,0,0],"isMain":true},{"metricName":"Total length (>= 25000 bp)","quality":"More is better","values":[0,0,0,0,0,0],"isMain":false},{"metricName":"Total length (>= 50000 bp)","quality":"More is better","values":[0,0,0,0,0,0],"isMain":true},{"metricName":"N50","quality":"More is better","values":[636,589,597,573,577,574],"isMain":false},{"metricName":"N75","quality":"More is better","values":[558,538,547,528,537,524],"isMain":false},{"metricName":"L50","quality":"Less is better","values":[8,10,21,21,13,7],"isMain":false},{"metricName":"L75","quality":"Less is better","values":[15,15,34,33,20,12],"isMain":false},{"metricName":"GC (%)","quality":"Equal","values":["42.06","43.79","43.41","42.85","43.89","41.50"],"isMain":false}]],["K-mer-based statistics",[]],["Predicted genes",[]],["Similarity statistics",[]],["Reference statistics",[{"metricName":"Reference length","quality":"Equal","values":[2515190,2515190,2515190,2515190,2515190,2515190],"isMain":false},{"metricName":"Reference fragments","quality":"Equal","values":[22,22,22,22,22,22],"isMain":false},{"metricName":"Reference GC (%)","quality":"Equal","values":["43.32","43.32","43.32","43.32","43.32","43.32"],"isMain":false}]]],"subreferences":[],"subreports":[],"minContig":500},{"date":"10 February 2020, Monday, 19:42:21","assembliesNames":["megahit-103-df","megahit-113-df","megahit-113-ml","megahit-113-ms","megahit-129-df","metaSPAdes"],"referenceName":"GCF\_000238635.1\_Subdol\_sp\_4\_3\_54A2FAA\_V1","order":[0,1,2,3,4,5],"report":[["Genome statistics",[{"metricName":"Genome fraction (%)","quality":"More is better","values":["6.562","9.084","9.370","9.788","9.982","4.308"],"isMain":true},{"metricName":"Duplication ratio","quality":"Less is better","values":["1.035","1.037","1.021","1.022","1.029","1.038"],"isMain":true},{"metricName":"Largest alignment","quality":"More is better","values":[2085,2667,6075,1978,5939,6359],"isMain":true},{"metricName":"Total aligned length","quality":"More is better","values":[277416,384283,395518,413406,421300,182038],"isMain":true},{"metricName":"NA50","quality":"More is better","values":[582,585,591,602,617,583],"isMain":false},{"metricName":"NA75","quality":"More is better","values":[521,511,523,526,537,519],"isMain":false},{"metricName":"NGA50","quality":"More is better","values":[null,null,null,null,null,null],"isMain":true},{"metricName":"LA50","quality":"Less is better","values":[202,308,290,295,273,128],"isMain":false},{"metricName":"LA75","quality":"Less is better","values":[336,519,484,490,466,217],"isMain":false}]],["Reads mapping",[]],["Misassemblies",[{"metricName":"# misassemblies","quality":"Less is better","values":[8,8,3,5,5,1],"isMain":true},{"metricName":" # relocations","quality":"Less is better","values":[1,1,0,1,0,0],"isMain":false},{"metricName":" # translocations","quality":"Less is better","values":[5,7,3,4,5,1],"isMain":false},{"metricName":" # inversions","quality":"Less is better","values":[2,0,0,0,0,0],"isMain":false},{"metricName":" # interspecies translocations","quality":"Less is better","values":[35,46,11,24,30,21],"isMain":false},{"metricName":"# misassembled contigs","quality":"Less is better","values":[7,8,3,5,5,1],"isMain":false},{"metricName":"Misassembled contigs length","quality":"Less is better","values":[6291,5529,2501,6980,3347,1410],"isMain":true},{"metricName":"# possibly misassembled contigs","quality":"Less is better","values":[0,0,0,2,0,0],"isMain":false},{"metricName":" # possible misassemblies","quality":"Less is better","values":[0,0,0,2,0,0],"isMain":false},{"metricName":"# local misassemblies","quality":"Less is better","values":[3,3,1,0,2,0],"isMain":false},{"metricName":"# scaffold gap ext. mis.","quality":"Less is better","values":[0,0,0,0,0,0],"isMain":false},{"metricName":"# scaffold gap loc. mis.","quality":"Less is better","values":[0,0,0,0,0,0],"isMain":false},{"metricName":"# unaligned mis. contigs","quality":"Less is better","values":[0,0,1,0,0,0],"isMain":false}]],["Unaligned",[{"metricName":"# fully unaligned contigs","quality":"Less is better","values":[2,0,1,3,0,0],"isMain":false},{"metricName":"Fully unaligned length","quality":"Less is better","values":[1675,0,550,3359,0,0],"isMain":false},{"metricName":"# partially unaligned contigs","quality":"Less is better","values":[8,9,3,11,2,6],"isMain":false},{"metricName":"Partially unaligned length","quality":"Less is better","values":[6610,61667,30649,13359,8812,5465],"isMain":false}]],["Mismatches",[{"metricName":"# mismatches","quality":"Less is better","values":[4387,6524,8040,8436,7351,3208],"isMain":false},{"metricName":"# indels","quality":"Less is better","values":[25,27,18,11,17,5],"isMain":false},{"metricName":"Indels length","quality":"Less is better","values":[27,34,37,13,21,5],"isMain":false},{"metricName":"# mismatches per 100 kbp","quality":"Less is better","values":["1589.88","1707.92","2040.47","2049.52","1751.23","1770.77"],"isMain":true},{"metricName":"# indels per 100 kbp","quality":"Less is better","values":["9.06","7.07","4.57","2.67","4.05","2.76"],"isMain":true},{"metricName":" # indels (<= 5 bp)","quality":"Less is better","values":[25,27,17,11,17,5],"isMain":false},{"metricName":" # indels (> 5 bp)","quality":"Less is better","values":[0,0,1,0,0,0],"isMain":false},{"metricName":"# N's","quality":"Less is better","values":[0,0,0,0,0,0],"isMain":false},{"metricName":"# N's per 100 kbp","quality":"Less is better","values":["0.00","0.00","0.00","0.00","0.00","0.00"],"isMain":true}]],["Statistics without reference",[{"metricName":"# contigs","quality":"Equal","values":[458,609,626,663,649,296],"isMain":true},{"metricName":"# contigs (>= 1000 bp)","quality":"Equal","values":[16,28,17,22,33,10],"isMain":false},{"metricName":"# contigs (>= 5000 bp)","quality":"Equal","values":[0,3,2,0,2,1],"isMain":false},{"metricName":"# contigs (>= 10000 bp)","quality":"Equal","values":[0,2,1,0,0,0],"isMain":false},{"metricName":"# contigs (>= 25000 bp)","quality":"Equal","values":[0,1,1,0,0,0],"isMain":false},{"metricName":"# contigs (>= 50000 bp)","quality":"Equal","values":[0,0,0,0,0,0],"isMain":false},{"metricName":"Largest contig","quality":"More is better","values":[2163,37773,29039,4811,9727,6359],"isMain":true},{"metricName":"Total length","quality":"More is better","values":[293864,457616,433675,437576,440928,193439],"isMain":true},{"metricName":"Total length (>= 1000 bp)","quality":"More is better","values":[21219,89894,57016,33408,53013,18978],"isMain":true},{"metricName":"Total length (>= 5000 bp)","quality":"More is better","values":[0,58575,35114,0,15666,6359],"isMain":false},{"metricName":"Total length (>= 10000 bp)","quality":"More is better","values":[0,49275,29039,0,0,0],"isMain":true},{"metricName":"Total length (>= 25000 bp)","quality":"More is better","values":[0,37773,29039,0,0,0],"isMain":false},{"metricName":"Total length (>= 50000 bp)","quality":"More is better","values":[0,0,0,0,0,0],"isMain":true},{"metricName":"N50","quality":"More is better","values":[610,666,627,634,634,609],"isMain":false},{"metricName":"N75","quality":"More is better","values":[542,565,555,551,554,543],"isMain":false},{"metricName":"L50","quality":"Less is better","values":[189,205,235,269,254,119],"isMain":false},{"metricName":"L75","quality":"Less is better","values":[317,393,419,454,440,203],"isMain":false},{"metricName":"GC (%)","quality":"Equal","values":["55.80","54.86","55.86","55.87","55.82","56.14"],"isMain":false}]],["K-mer-based statistics",[]],["Predicted genes",[]],["Similarity statistics",[]],["Reference statistics",[{"metricName":"Reference length","quality":"Equal","values":[4205237,4205237,4205237,4205237,4205237,4205237],"isMain":false},{"metricName":"Reference fragments","quality":"Equal","values":[103,103,103,103,103,103],"isMain":false},{"metricName":"Reference GC (%)","quality":"Equal","values":["56.18","56.18","56.18","56.18","56.18","56.18"],"isMain":false}]]],"subreferences":[],"subreports":[],"minContig":500},{"date":"10 February 2020, Monday, 19:42:21","assembliesNames":["megahit-103-df","megahit-113-df","megahit-113-ml","megahit-113-ms","megahit-129-df","metaSPAdes"],"referenceName":"GCF\_000238835.1\_ASM23883v1","order":[0,1,2,3,4,5],"report":[["Genome statistics",[{"metricName":"Genome fraction (%)","quality":"More is better","values":["4.519","5.999","5.654","7.351","6.826","2.978"],"isMain":true},{"metricName":"Duplication ratio","quality":"Less is better","values":["1.016","1.030","1.013","1.026","1.020","1.066"],"isMain":true},{"metricName":"Largest alignment","quality":"More is better","values":[1003,1194,1251,1251,1138,873],"isMain":true},{"metricName":"Total aligned length","quality":"More is better","values":[129084,171433,161569,209857,195406,85074],"isMain":true},{"metricName":"NA50","quality":"More is better","values":[574,574,573,583,580,558],"isMain":false},{"metricName":"NA75","quality":"More is better","values":[524,521,526,525,523,511],"isMain":false},{"metricName":"NGA50","quality":"More is better","values":[null,null,null,null,null,null],"isMain":true},{"metricName":"LA50","quality":"Less is better","values":[101,136,122,159,148,73],"isMain":false},{"metricName":"LA75","quality":"Less is better","values":[161,221,197,259,241,117],"isMain":false}]],["Reads mapping",[]],["Misassemblies",[{"metricName":"# misassemblies","quality":"Less is better","values":[1,0,0,1,2,0],"isMain":true},{"metricName":" # relocations","quality":"Less is better","values":[0,0,0,0,0,0],"isMain":false},{"metricName":" # translocations","quality":"Less is better","values":[0,0,0,1,1,0],"isMain":false},{"metricName":" # inversions","quality":"Less is better","values":[1,0,0,0,1,0],"isMain":false},{"metricName":" # interspecies translocations","quality":"Less is better","values":[8,19,2,14,13,18],"isMain":false},{"metricName":"# misassembled contigs","quality":"Less is better","values":[1,0,0,1,2,0],"isMain":false},{"metricName":"Misassembled contigs length","quality":"Less is better","values":[573,0,0,703,1265,0],"isMain":true},{"metricName":"# possibly misassembled contigs","quality":"Less is better","values":[0,0,0,0,0,0],"isMain":false},{"metricName":" # possible misassemblies","quality":"Less is better","values":[0,0,0,0,0,0],"isMain":false},{"metricName":"# local misassemblies","quality":"Less is better","values":[0,0,0,0,0,0],"isMain":false},{"metricName":"# scaffold gap ext. mis.","quality":"Less is better","values":[0,0,0,0,0,0],"isMain":false},{"metricName":"# scaffold gap loc. mis.","quality":"Less is better","values":[0,0,0,0,0,0],"isMain":false},{"metricName":"# unaligned mis. contigs","quality":"Less is better","values":[0,0,0,0,0,0],"isMain":false}]],["Unaligned",[{"metricName":"# fully unaligned contigs","quality":"Less is better","values":[0,1,0,0,0,0],"isMain":false},{"metricName":"Fully unaligned length","quality":"Less is better","values":[0,503,0,0,0,0],"isMain":false},{"metricName":"# partially unaligned contigs","quality":"Less is better","values":[2,7,1,7,5,4],"isMain":false},{"metricName":"Partially unaligned length","quality":"Less is better","values":[1134,8621,2908,6960,5531,2565],"isMain":false}]],["Mismatches",[{"metricName":"# mismatches","quality":"Less is better","values":[2141,2945,3247,4387,3248,1560],"isMain":false},{"metricName":"# indels","quality":"Less is better","values":[2,6,6,8,7,1],"isMain":false},{"metricName":"Indels length","quality":"Less is better","values":[2,11,6,8,7,1],"isMain":false},{"metricName":"# mismatches per 100 kbp","quality":"Less is better","values":["1660.95","1721.18","2013.39","2092.56","1668.42","1836.44"],"isMain":true},{"metricName":"# indels per 100 kbp","quality":"Less is better","values":["1.55","3.51","3.72","3.82","3.60","1.18"],"isMain":true},{"metricName":" # indels (<= 5 bp)","quality":"Less is better","values":[2,5,6,8,7,1],"isMain":false},{"metricName":" # indels (> 5 bp)","quality":"Less is better","values":[0,1,0,0,0,0],"isMain":false},{"metricName":"# N's","quality":"Less is better","values":[0,0,0,0,0,0],"isMain":false},{"metricName":"# N's per 100 kbp","quality":"Less is better","values":["0.00","0.00","0.00","0.00","0.00","0.00"],"isMain":true}]],["Statistics without reference",[{"metricName":"# contigs","quality":"Equal","values":[223,293,270,352,328,155],"isMain":true},{"metricName":"# contigs (>= 1000 bp)","quality":"Equal","values":[1,9,2,9,4,0],"isMain":false},{"metricName":"# contigs (>= 5000 bp)","quality":"Equal","values":[0,0,0,0,0,0],"isMain":false},{"metricName":"# contigs (>= 10000 bp)","quality":"Equal","values":[0,0,0,0,0,0],"isMain":false},{"metricName":"# contigs (>= 25000 bp)","quality":"Equal","values":[0,0,0,0,0,0],"isMain":false},{"metricName":"# contigs (>= 50000 bp)","quality":"Equal","values":[0,0,0,0,0,0],"isMain":false},{"metricName":"Largest contig","quality":"More is better","values":[1003,3231,3104,3231,3231,993],"isMain":true},{"metricName":"Total length","quality":"More is better","values":[132074,185363,166286,222036,204158,93154],"isMain":true},{"metricName":"Total length (>= 1000 bp)","quality":"More is better","values":[1003,13668,4355,12144,6408,0],"isMain":true},{"metricName":"Total length (>= 5000 bp)","quality":"More is better","values":[0,0,0,0,0,0],"isMain":false},{"metricName":"Total length (>= 10000 bp)","quality":"More is better","values":[0,0,0,0,0,0],"isMain":true},{"metricName":"Total length (>= 25000 bp)","quality":"More is better","values":[0,0,0,0,0,0],"isMain":false},{"metricName":"Total length (>= 50000 bp)","quality":"More is better","values":[0,0,0,0,0,0],"isMain":true},{"metricName":"N50","quality":"More is better","values":[577,589,588,607,592,586],"isMain":false},{"metricName":"N75","quality":"More is better","values":[531,542,535,547,541,534],"isMain":false},{"metricName":"L50","quality":"Less is better","values":[99,122,116,149,140,69],"isMain":false},{"metricName":"L75","quality":"Less is better","values":[159,204,190,246,230,110],"isMain":false},{"metricName":"GC (%)","quality":"Equal","values":["46.32","45.76","45.70","45.68","45.94","45.10"],"isMain":false}]],["K-mer-based statistics",[]],["Predicted genes",[]],["Similarity statistics",[]],["Reference statistics",[{"metricName":"Reference length","quality":"Equal","values":[2852137,2852137,2852137,2852137,2852137,2852137],"isMain":false},{"metricName":"Reference fragments","quality":"Equal","values":[212,212,212,212,212,212],"isMain":false},{"metricName":"Reference GC (%)","quality":"Equal","values":["45.60","45.60","45.60","45.60","45.60","45.60"],"isMain":false}]]],"subreferences":[],"subreports":[],"minContig":500},{"date":"10 February 2020, Monday, 19:42:39","assembliesNames":["megahit-103-df","megahit-113-df","megahit-113-ml","megahit-113-ms","megahit-129-df","metaSPAdes"],"referenceName":"GCF\_000242315.1\_ASM24231v1","order":[0,1,2,3,4,5],"report":[["Genome statistics",[{"metricName":"Genome fraction (%)","quality":"More is better","values":["13.423","16.499","15.077","17.836","18.597","8.083"],"isMain":true},{"metricName":"Duplication ratio","quality":"Less is better","values":["1.034","1.046","1.033","1.038","1.041","1.042"],"isMain":true},{"metricName":"Largest alignment","quality":"More is better","values":[1611,1519,1380,1598,1735,1290],"isMain":true},{"metricName":"Total aligned length","quality":"More is better","values":[396221,487838,448391,528978,550968,238827],"isMain":true},{"metricName":"NA50","quality":"More is better","values":[617,630,609,626,641,583],"isMain":false},{"metricName":"NA75","quality":"More is better","values":[537,527,539,543,554,529],"isMain":false},{"metricName":"NGA50","quality":"More is better","values":[null,null,null,null,null,null],"isMain":true},{"metricName":"LA50","quality":"Less is better","values":[268,342,309,354,347,177],"isMain":false},{"metricName":"LA75","quality":"Less is better","values":[448,584,511,592,592,293],"isMain":false}]],["Reads mapping",[]],["Misassemblies",[{"metricName":"# misassemblies","quality":"Less is better","values":[1,3,2,2,3,2],"isMain":true},{"metricName":" # relocations","quality":"Less is better","values":[1,1,1,1,1,1],"isMain":false},{"metricName":" # translocations","quality":"Less is better","values":[0,1,0,0,1,1],"isMain":false},{"metricName":" # inversions","quality":"Less is better","values":[0,1,1,1,1,0],"isMain":false},{"metricName":" # interspecies translocations","quality":"Less is better","values":[41,60,14,33,50,29],"isMain":false},{"metricName":"# misassembled contigs","quality":"Less is better","values":[1,3,2,2,3,2],"isMain":false},{"metricName":"Misassembled contigs length","quality":"Less is better","values":[911,2110,1616,1497,2141,1504],"isMain":true},{"metricName":"# possibly misassembled contigs","quality":"Less is better","values":[0,0,0,1,0,0],"isMain":false},{"metricName":" # possible misassemblies","quality":"Less is better","values":[0,0,0,1,0,0],"isMain":false},{"metricName":"# local misassemblies","quality":"Less is better","values":[0,0,0,0,1,1],"isMain":false},{"metricName":"# scaffold gap ext. mis.","quality":"Less is better","values":[0,0,0,0,0,0],"isMain":false},{"metricName":"# scaffold gap loc. mis.","quality":"Less is better","values":[0,0,0,0,0,0],"isMain":false},{"metricName":"# unaligned mis. contigs","quality":"Less is better","values":[1,1,1,1,1,0],"isMain":false}]],["Unaligned",[{"metricName":"# fully unaligned contigs","quality":"Less is better","values":[1,1,3,2,2,2],"isMain":false},{"metricName":"Fully unaligned length","quality":"Less is better","values":[656,656,1682,1363,1318,1171],"isMain":false},{"metricName":"# partially unaligned contigs","quality":"Less is better","values":[7,11,1,6,9,9],"isMain":false},{"metricName":"Partially unaligned length","quality":"Less is better","values":[5837,47198,2967,6664,10718,6808],"isMain":false}]],["Mismatches",[{"metricName":"# mismatches","quality":"Less is better","values":[7240,8859,9842,11993,10050,4323],"isMain":false},{"metricName":"# indels","quality":"Less is better","values":[42,43,35,62,61,23],"isMain":false},{"metricName":"Indels length","quality":"Less is better","values":[92,53,65,66,78,31],"isMain":false},{"metricName":"# mismatches per 100 kbp","quality":"Less is better","values":["1835.78","1827.53","2221.79","2288.64","1839.34","1820.23"],"isMain":true},{"metricName":"# indels per 100 kbp","quality":"Less is better","values":["10.65","8.87","7.90","11.83","11.16","9.68"],"isMain":true},{"metricName":" # indels (<= 5 bp)","quality":"Less is better","values":[40,43,34,62,60,23],"isMain":false},{"metricName":" # indels (> 5 bp)","quality":"Less is better","values":[2,0,1,0,1,0],"isMain":false},{"metricName":"# N's","quality":"Less is better","values":[0,0,0,0,0,0],"isMain":false},{"metricName":"# N's per 100 kbp","quality":"Less is better","values":["0.00","0.00","0.00","0.00","0.00","0.00"],"isMain":true}]],["Statistics without reference",[{"metricName":"# contigs","quality":"Equal","values":[629,742,712,829,831,398],"isMain":true},{"metricName":"# contigs (>= 1000 bp)","quality":"Equal","values":[32,60,25,41,64,9],"isMain":false},{"metricName":"# contigs (>= 5000 bp)","quality":"Equal","values":[0,3,1,0,1,0],"isMain":false},{"metricName":"# contigs (>= 10000 bp)","quality":"Equal","values":[0,3,0,0,0,0],"isMain":false},{"metricName":"# contigs (>= 25000 bp)","quality":"Equal","values":[0,0,0,0,0,0],"isMain":false},{"metricName":"# contigs (>= 50000 bp)","quality":"Equal","values":[0,0,0,0,0,0],"isMain":false},{"metricName":"Largest contig","quality":"More is better","values":[2406,17701,5830,2436,5805,2203],"isMain":true},{"metricName":"Total length","quality":"More is better","values":[414279,554969,462465,551787,580736,255409],"isMain":true},{"metricName":"Total length (>= 1000 bp)","quality":"More is better","values":[38226,109540,33253,49965,79798,11109],"isMain":true},{"metricName":"Total length (>= 5000 bp)","quality":"More is better","values":[0,42363,5830,0,5805,0],"isMain":false},{"metricName":"Total length (>= 10000 bp)","quality":"More is better","values":[0,42363,0,0,0,0],"isMain":true},{"metricName":"Total length (>= 25000 bp)","quality":"More is better","values":[0,0,0,0,0,0],"isMain":false},{"metricName":"Total length (>= 50000 bp)","quality":"More is better","values":[0,0,0,0,0,0],"isMain":true},{"metricName":"N50","quality":"More is better","values":[635,699,622,648,668,623],"isMain":false},{"metricName":"N75","quality":"More is better","values":[560,586,555,564,579,549],"isMain":false},{"metricName":"L50","quality":"Less is better","values":[259,268,296,340,328,166],"isMain":false},{"metricName":"L75","quality":"Less is better","values":[433,485,492,568,562,277],"isMain":false},{"metricName":"GC (%)","quality":"Equal","values":["38.17","37.86","38.23","38.08","38.13","38.04"],"isMain":false}]],["K-mer-based statistics",[]],["Predicted genes",[]],["Similarity statistics",[]],["Reference statistics",[{"metricName":"Reference length","quality":"Equal","values":[2938075,2938075,2938075,2938075,2938075,2938075],"isMain":false},{"metricName":"Reference fragments","quality":"Equal","values":[227,227,227,227,227,227],"isMain":false},{"metricName":"Reference GC (%)","quality":"Equal","values":["37.61","37.61","37.61","37.61","37.61","37.61"],"isMain":false}]]],"subreferences":[],"subreports":[],"minContig":500},{"date":"10 February 2020, Monday, 19:42:49","assembliesNames":["megahit-103-df","megahit-113-df","megahit-113-ml","megahit-113-ms","megahit-129-df","metaSPAdes"],"referenceName":"GCF\_000243015.1\_Prev\_macu\_OT\_289\_V1","order":[0,1,2,3,4,5],"report":[["Genome statistics",[{"metricName":"Genome fraction (%)","quality":"More is better","values":["28.521","99.386","98.568","98.612","98.626","98.330"],"isMain":true},{"metricName":"Duplication ratio","quality":"Less is better","values":["1.037","1.046","1.041","1.075","1.092","1.001"],"isMain":true},{"metricName":"Largest alignment","quality":"More is better","values":[2243,156990,225720,86528,155160,424351],"isMain":true},{"metricName":"Total aligned length","quality":"More is better","values":[909850,3244572,3200459,3294951,3353718,3084573],"isMain":true},{"metricName":"NG50","quality":"More is better","values":[null,89934,61284,86788,45685,74445],"isMain":false},{"metricName":"NG75","quality":"More is better","values":[null,56362,45865,60973,27817,48858],"isMain":false},{"metricName":"NA50","quality":"More is better","values":[692,29976,44139,15519,28553,68149],"isMain":false},{"metricName":"NA75","quality":"More is better","values":[557,536,18765,null,10945,40753],"isMain":false},{"metricName":"NGA50","quality":"More is better","values":[null,42820,46922,35653,32972,68149],"isMain":true},{"metricName":"NGA75","quality":"More is better","values":[null,26789,29266,17548,19512,39152],"isMain":false},{"metricName":"LG50","quality":"Less is better","values":[null,10,13,12,24,10],"isMain":false},{"metricName":"LG75","quality":"Less is better","values":[null,21,28,24,47,22],"isMain":false},{"metricName":"LA50","quality":"Less is better","values":[514,40,26,68,40,12],"isMain":false},{"metricName":"LA75","quality":"Less is better","values":[903,252,56,null,87,26],"isMain":false},{"metricName":"LGA50","quality":"Less is better","values":[null,24,21,31,31,12],"isMain":true},{"metricName":"LGA75","quality":"Less is better","values":[null,47,42,62,62,27],"isMain":false}]],["Reads mapping",[]],["Misassemblies",[{"metricName":"# misassemblies","quality":"Less is better","values":[15,19,29,41,40,9],"isMain":true},{"metricName":" # relocations","quality":"Less is better","values":[0,1,5,1,5,0],"isMain":false},{"metricName":" # translocations","quality":"Less is better","values":[6,16,24,39,23,9],"isMain":false},{"metricName":" # inversions","quality":"Less is better","values":[9,2,0,1,12,0],"isMain":false},{"metricName":" # interspecies translocations","quality":"Less is better","values":[79,48,53,112,67,5],"isMain":false},{"metricName":"# misassembled contigs","quality":"Less is better","values":[14,18,24,33,36,6],"isMain":false},{"metricName":"Misassembled contigs length","quality":"Less is better","values":[8158,593738,1111801,592563,574918,771498],"isMain":true},{"metricName":"# possibly misassembled contigs","quality":"Less is better","values":[1,0,2,1,0,0],"isMain":false},{"metricName":" # possible misassemblies","quality":"Less is better","values":[1,0,2,1,0,0],"isMain":false},{"metricName":"# local misassemblies","quality":"Less is better","values":[2,9,11,9,17,1],"isMain":false},{"metricName":"# scaffold gap ext. mis.","quality":"Less is better","values":[0,0,0,0,0,0],"isMain":false},{"metricName":"# scaffold gap loc. mis.","quality":"Less is better","values":[0,0,0,0,0,0],"isMain":false},{"metricName":"# unaligned mis. contigs","quality":"Less is better","values":[0,1,1,5,0,0],"isMain":false}]],["Unaligned",[{"metricName":"# fully unaligned contigs","quality":"Less is better","values":[0,2,3,6,4,0],"isMain":false},{"metricName":"Fully unaligned length","quality":"Less is better","values":[0,2266,2582,6471,2153,0],"isMain":false},{"metricName":"# partially unaligned contigs","quality":"Less is better","values":[35,27,30,58,22,2],"isMain":false},{"metricName":"Partially unaligned length","quality":"Less is better","values":[30513,1013020,368352,1593263,292975,8485],"isMain":false}]],["Mismatches",[{"metricName":"# mismatches","quality":"Less is better","values":[3276,5863,5190,10185,9119,832],"isMain":false},{"metricName":"# indels","quality":"Less is better","values":[3,90,76,214,102,26],"isMain":false},{"metricName":"Indels length","quality":"Less is better","values":[68,433,683,1586,847,130],"isMain":false},{"metricName":"# mismatches per 100 kbp","quality":"Less is better","values":["366.48","188.22","168.00","329.54","295.01","27.00"],"isMain":true},{"metricName":"# indels per 100 kbp","quality":"Less is better","values":["0.34","2.89","2.46","6.92","3.30","0.84"],"isMain":true},{"metricName":" # indels (<= 5 bp)","quality":"Less is better","values":[1,81,62,180,83,21],"isMain":false},{"metricName":" # indels (> 5 bp)","quality":"Less is better","values":[2,9,14,34,19,5],"isMain":false},{"metricName":"# N's","quality":"Less is better","values":[0,0,0,0,0,0],"isMain":false},{"metricName":"# N's per 100 kbp","quality":"Less is better","values":["0.00","0.00","0.00","0.00","0.00","0.00"],"isMain":true}]],["Statistics without reference",[{"metricName":"# contigs","quality":"Equal","values":[1318,334,311,489,629,75],"isMain":true},{"metricName":"# contigs (>= 1000 bp)","quality":"Equal","values":[161,143,128,177,168,70],"isMain":false},{"metricName":"# contigs (>= 5000 bp)","quality":"Equal","values":[0,94,80,121,111,52],"isMain":false},{"metricName":"# contigs (>= 10000 bp)","quality":"Equal","values":[0,75,69,96,89,46],"isMain":false},{"metricName":"# contigs (>= 25000 bp)","quality":"Equal","values":[0,52,45,60,50,38],"isMain":false},{"metricName":"# contigs (>= 50000 bp)","quality":"Equal","values":[0,25,25,32,19,21],"isMain":false},{"metricName":"Largest contig","quality":"More is better","values":[2647,521349,272355,295363,182464,424351],"isMain":true},{"metricName":"Total length","quality":"More is better","values":[957137,4272987,3587245,4921524,3671353,3094608],"isMain":true},{"metricName":"Total length (>= 1000 bp)","quality":"More is better","values":[198904,4157942,3477014,4727486,3406541,3091243],"isMain":true},{"metricName":"Total length (>= 5000 bp)","quality":"More is better","values":[0,4042220,3371820,4606405,3265034,3052582],"isMain":false},{"metricName":"Total length (>= 10000 bp)","quality":"More is better","values":[0,3907343,3296148,4429700,3106064,3019482],"isMain":true},{"metricName":"Total length (>= 25000 bp)","quality":"More is better","values":[0,3533619,2934206,3827868,2434575,2912099],"isMain":false},{"metricName":"Total length (>= 50000 bp)","quality":"More is better","values":[0,2572979,2233941,2833736,1345257,2318217],"isMain":true},{"metricName":"N50","quality":"More is better","values":[714,62299,55948,58913,36858,83173],"isMain":false},{"metricName":"N75","quality":"More is better","values":[577,31284,31310,27824,19188,48858],"isMain":false},{"metricName":"L50","quality":"Less is better","values":[497,18,17,26,31,9],"isMain":false},{"metricName":"L75","quality":"Less is better","values":[871,41,37,55,65,22],"isMain":false},{"metricName":"GC (%)","quality":"Equal","values":["47.90","48.01","47.57","48.38","47.66","47.72"],"isMain":false}]],["K-mer-based statistics",[]],["Predicted genes",[]],["Similarity statistics",[]],["Reference statistics",[{"metricName":"Reference length","quality":"Equal","values":[3134147,3134147,3134147,3134147,3134147,3134147],"isMain":false},{"metricName":"Reference fragments","quality":"Equal","values":[54,54,54,54,54,54],"isMain":false},{"metricName":"Reference GC (%)","quality":"Equal","values":["47.71","47.71","47.71","47.71","47.71","47.71"],"isMain":false}]]],"subreferences":[],"subreports":[],"minContig":500},{"date":"10 February 2020, Monday, 19:43:02","assembliesNames":["megahit-103-df","megahit-113-df","megahit-113-ml","megahit-113-ms","megahit-129-df","metaSPAdes"],"referenceName":"GCF\_000243215.1\_Odor\_lane\_YIT\_12061\_V1","order":[0,1,2,3,4,5],"report":[["Genome statistics",[{"metricName":"Genome fraction (%)","quality":"More is better","values":["98.983","99.349","99.184","99.349","99.321","98.982"],"isMain":true},{"metricName":"Duplication ratio","quality":"Less is better","values":["1.005","1.001","1.003","1.002","1.001","1.000"],"isMain":true},{"metricName":"Largest alignment","quality":"More is better","values":[152045,242913,242899,242807,227171,242827],"isMain":true},{"metricName":"Total aligned length","quality":"More is better","values":[3731307,3732957,3729076,3734122,3730597,3715937],"isMain":true},{"metricName":"NG50","quality":"More is better","values":[31749,118973,118945,116455,135812,135640],"isMain":false},{"metricName":"NG75","quality":"More is better","values":[18493,64465,68366,80424,68515,73317],"isMain":false},{"metricName":"NA50","quality":"More is better","values":[31314,118973,118945,103114,119182,118513],"isMain":false},{"metricName":"NA75","quality":"More is better","values":[17895,61157,60366,50809,61142,67067],"isMain":false},{"metricName":"NGA50","quality":"More is better","values":[31314,118973,118945,103114,119182,118513],"isMain":true},{"metricName":"NGA75","quality":"More is better","values":[17895,61157,59805,58858,61142,67067],"isMain":false},{"metricName":"LG50","quality":"Less is better","values":[37,11,12,11,10,9],"isMain":false},{"metricName":"LG75","quality":"Less is better","values":[76,22,22,21,20,19],"isMain":false},{"metricName":"LA50","quality":"Less is better","values":[38,12,12,13,12,12],"isMain":false},{"metricName":"LA75","quality":"Less is better","values":[77,23,23,27,22,23],"isMain":false},{"metricName":"LGA50","quality":"Less is better","values":[38,12,12,13,12,12],"isMain":true},{"metricName":"LGA75","quality":"Less is better","values":[77,23,24,25,22,23],"isMain":false}]],["Reads mapping",[]],["Misassemblies",[{"metricName":"# misassemblies","quality":"Less is better","values":[7,7,7,9,9,8],"isMain":true},{"metricName":" # relocations","quality":"Less is better","values":[0,0,0,0,0,1],"isMain":false},{"metricName":" # translocations","quality":"Less is better","values":[7,7,7,9,9,7],"isMain":false},{"metricName":" # inversions","quality":"Less is better","values":[0,0,0,0,0,0],"isMain":false},{"metricName":" # interspecies translocations","quality":"Less is better","values":[18,3,9,11,3,1],"isMain":false},{"metricName":"# misassembled contigs","quality":"Less is better","values":[6,6,6,8,8,7],"isMain":false},{"metricName":"Misassembled contigs length","quality":"Less is better","values":[196737,692029,396267,763183,912594,1314786],"isMain":true},{"metricName":"# possibly misassembled contigs","quality":"Less is better","values":[0,0,0,0,0,0],"isMain":false},{"metricName":" # possible misassemblies","quality":"Less is better","values":[0,0,0,0,0,0],"isMain":false},{"metricName":"# local misassemblies","quality":"Less is better","values":[1,1,1,2,1,0],"isMain":false},{"metricName":"# scaffold gap ext. mis.","quality":"Less is better","values":[0,0,0,0,0,0],"isMain":false},{"metricName":"# scaffold gap loc. mis.","quality":"Less is better","values":[0,0,0,0,0,0],"isMain":false},{"metricName":"# unaligned mis. contigs","quality":"Less is better","values":[0,0,0,1,0,0],"isMain":false}]],["Unaligned",[{"metricName":"# fully unaligned contigs","quality":"Less is better","values":[0,0,0,0,0,0],"isMain":false},{"metricName":"Fully unaligned length","quality":"Less is better","values":[0,0,0,0,0,0],"isMain":false},{"metricName":"# partially unaligned contigs","quality":"Less is better","values":[13,2,5,10,0,1],"isMain":false},{"metricName":"Partially unaligned length","quality":"Less is better","values":[12867,1427,8228,211152,0,731],"isMain":false}]],["Mismatches",[{"metricName":"# mismatches","quality":"Less is better","values":[5693,258,441,471,277,515],"isMain":false},{"metricName":"# indels","quality":"Less is better","values":[0,2,6,17,10,37],"isMain":false},{"metricName":"Indels length","quality":"Less is better","values":[0,2,8,66,17,110],"isMain":false},{"metricName":"# mismatches per 100 kbp","quality":"Less is better","values":["153.23","6.92","11.85","12.63","7.43","13.86"],"isMain":true},{"metricName":"# indels per 100 kbp","quality":"Less is better","values":["0.00","0.05","0.16","0.46","0.27","1.00"],"isMain":true},{"metricName":" # indels (<= 5 bp)","quality":"Less is better","values":[0,2,6,14,9,34],"isMain":false},{"metricName":" # indels (> 5 bp)","quality":"Less is better","values":[0,0,0,3,1,3],"isMain":false},{"metricName":"# N's","quality":"Less is better","values":[0,0,0,0,0,0],"isMain":false},{"metricName":"# N's per 100 kbp","quality":"Less is better","values":["0.00","0.00","0.00","0.00","0.00","0.00"],"isMain":true}]],["Statistics without reference",[{"metricName":"# contigs","quality":"Equal","values":[221,73,75,72,66,64],"isMain":true},{"metricName":"# contigs (>= 1000 bp)","quality":"Equal","values":[198,65,66,65,58,59],"isMain":false},{"metricName":"# contigs (>= 5000 bp)","quality":"Equal","values":[145,47,48,50,44,43],"isMain":false},{"metricName":"# contigs (>= 10000 bp)","quality":"Equal","values":[117,43,42,45,40,39],"isMain":false},{"metricName":"# contigs (>= 25000 bp)","quality":"Equal","values":[51,39,39,41,36,35],"isMain":false},{"metricName":"# contigs (>= 50000 bp)","quality":"Equal","values":[12,27,29,31,26,26],"isMain":false},{"metricName":"Largest contig","quality":"More is better","values":[152045,383042,245064,382936,339553,382838],"isMain":true},{"metricName":"Total length","quality":"More is better","values":[3746486,3735991,3742348,3946672,3732689,3716935],"isMain":true},{"metricName":"Total length (>= 1000 bp)","quality":"More is better","values":[3731430,3730825,3735936,3942318,3728313,3712974],"isMain":true},{"metricName":"Total length (>= 5000 bp)","quality":"More is better","values":[3609722,3700419,3703902,3915713,3706500,3688949],"isMain":false},{"metricName":"Total length (>= 10000 bp)","quality":"More is better","values":[3392527,3673189,3660104,3882043,3679166,3661894],"isMain":true},{"metricName":"Total length (>= 25000 bp)","quality":"More is better","values":[2283840,3598564,3607113,3805873,3601312,3579883],"isMain":false},{"metricName":"Total length (>= 50000 bp)","quality":"More is better","values":[891145,3124295,3249992,3429801,3218797,3279447],"isMain":true},{"metricName":"N50","quality":"More is better","values":[31749,118973,118945,107167,135812,135640],"isMain":false},{"metricName":"N75","quality":"More is better","values":[18493,64465,68366,64471,68515,73317],"isMain":false},{"metricName":"L50","quality":"Less is better","values":[37,11,12,12,10,9],"isMain":false},{"metricName":"L75","quality":"Less is better","values":[76,22,22,23,20,19],"isMain":false},{"metricName":"GC (%)","quality":"Equal","values":["40.55","40.53","40.53","40.62","40.53","40.52"],"isMain":false}]],["K-mer-based statistics",[]],["Predicted genes",[]],["Similarity statistics",[]],["Reference statistics",[{"metricName":"Reference length","quality":"Equal","values":[3753461,3753461,3753461,3753461,3753461,3753461],"isMain":false},{"metricName":"Reference fragments","quality":"Equal","values":[42,42,42,42,42,42],"isMain":false},{"metricName":"Reference GC (%)","quality":"Equal","values":["40.55","40.55","40.55","40.55","40.55","40.55"],"isMain":false}]]],"subreferences":[],"subreports":[],"minContig":500},{"date":"10 February 2020, Monday, 19:43:12","assembliesNames":["megahit-103-df","megahit-113-df","megahit-113-ml","megahit-113-ms","megahit-129-df","metaSPAdes"],"referenceName":"GCF\_000245055.1\_ASM24505v1","order":[0,1,2,3,4,5],"report":[["Genome statistics",[{"metricName":"Genome fraction (%)","quality":"More is better","values":["97.851","99.217","99.157","99.176","99.106","98.808"],"isMain":true},{"metricName":"Duplication ratio","quality":"Less is better","values":["1.016","1.004","1.003","1.006","1.004","1.000"],"isMain":true},{"metricName":"Largest alignment","quality":"More is better","values":[32037,254042,257117,250596,250924,257124],"isMain":true},{"metricName":"Total aligned length","quality":"More is better","values":[4546871,4568076,4558860,4573654,4563228,4534324],"isMain":true},{"metricName":"NG50","quality":"More is better","values":[8535,76350,71670,72164,80042,71177],"isMain":false},{"metricName":"NG75","quality":"More is better","values":[4916,43541,43072,42777,46310,42904],"isMain":false},{"metricName":"NA50","quality":"More is better","values":[8535,71610,71321,70170,74135,72168],"isMain":false},{"metricName":"NA75","quality":"More is better","values":[4916,42839,40363,40385,43540,42904],"isMain":false},{"metricName":"NGA50","quality":"More is better","values":[8535,71610,71321,71329,74135,71177],"isMain":true},{"metricName":"NGA75","quality":"More is better","values":[4916,42297,40363,40912,43066,40926],"isMain":false},{"metricName":"LG50","quality":"Less is better","values":[170,19,20,20,21,21],"isMain":false},{"metricName":"LG75","quality":"Less is better","values":[347,39,40,41,39,41],"isMain":false},{"metricName":"LA50","quality":"Less is better","values":[170,20,21,22,21,20],"isMain":false},{"metricName":"LA75","quality":"Less is better","values":[347,40,42,44,40,40],"isMain":false},{"metricName":"LGA50","quality":"Less is better","values":[170,20,21,21,21,21],"isMain":true},{"metricName":"LGA75","quality":"Less is better","values":[347,41,42,43,41,41],"isMain":false}]],["Reads mapping",[]],["Misassemblies",[{"metricName":"# misassemblies","quality":"Less is better","values":[3,9,3,8,4,2],"isMain":true},{"metricName":" # relocations","quality":"Less is better","values":[0,0,1,0,0,0],"isMain":false},{"metricName":" # translocations","quality":"Less is better","values":[2,8,2,8,2,2],"isMain":false},{"metricName":" # inversions","quality":"Less is better","values":[1,1,0,0,2,0],"isMain":false},{"metricName":" # interspecies translocations","quality":"Less is better","values":[66,6,4,9,1,1],"isMain":false},{"metricName":"# misassembled contigs","quality":"Less is better","values":[3,8,3,8,4,2],"isMain":false},{"metricName":"Misassembled contigs length","quality":"Less is better","values":[4908,251926,105818,294406,194436,103833],"isMain":true},{"metricName":"# possibly misassembled contigs","quality":"Less is better","values":[0,0,0,0,0,0],"isMain":false},{"metricName":" # possible misassemblies","quality":"Less is better","values":[0,0,0,0,0,0],"isMain":false},{"metricName":"# local misassemblies","quality":"Less is better","values":[1,3,0,1,1,0],"isMain":false},{"metricName":"# scaffold gap ext. mis.","quality":"Less is better","values":[0,0,0,0,0,0],"isMain":false},{"metricName":"# scaffold gap loc. mis.","quality":"Less is better","values":[0,0,0,0,0,0],"isMain":false},{"metricName":"# unaligned mis. contigs","quality":"Less is better","values":[0,0,0,2,0,0],"isMain":false}]],["Unaligned",[{"metricName":"# fully unaligned contigs","quality":"Less is better","values":[0,0,0,2,0,0],"isMain":false},{"metricName":"Fully unaligned length","quality":"Less is better","values":[0,0,0,1080,0,0],"isMain":false},{"metricName":"# partially unaligned contigs","quality":"Less is better","values":[28,4,3,6,1,1],"isMain":false},{"metricName":"Partially unaligned length","quality":"Less is better","values":[22962,3450,57026,71007,585,765],"isMain":false}]],["Mismatches",[{"metricName":"# mismatches","quality":"Less is better","values":[7612,905,665,1513,977,379],"isMain":false},{"metricName":"# indels","quality":"Less is better","values":[0,21,19,35,26,13],"isMain":false},{"metricName":"Indels length","quality":"Less is better","values":[0,332,432,483,401,21],"isMain":false},{"metricName":"# mismatches per 100 kbp","quality":"Less is better","values":["169.58","19.88","14.62","33.26","21.49","8.36"],"isMain":true},{"metricName":"# indels per 100 kbp","quality":"Less is better","values":["0.00","0.46","0.42","0.77","0.57","0.29"],"isMain":true},{"metricName":" # indels (<= 5 bp)","quality":"Less is better","values":[0,15,10,24,17,13],"isMain":false},{"metricName":" # indels (> 5 bp)","quality":"Less is better","values":[0,6,9,11,9,0],"isMain":false},{"metricName":"# N's","quality":"Less is better","values":[0,0,0,0,0,0],"isMain":false},{"metricName":"# N's per 100 kbp","quality":"Less is better","values":["0.00","0.00","0.00","0.00","0.00","0.00"],"isMain":true}]],["Statistics without reference",[{"metricName":"# contigs","quality":"Equal","values":[873,140,122,153,138,108],"isMain":true},{"metricName":"# contigs (>= 1000 bp)","quality":"Equal","values":[728,103,102,108,99,100],"isMain":false},{"metricName":"# contigs (>= 5000 bp)","quality":"Equal","values":[339,89,90,95,88,86],"isMain":false},{"metricName":"# contigs (>= 10000 bp)","quality":"Equal","values":[128,78,80,81,78,77],"isMain":false},{"metricName":"# contigs (>= 25000 bp)","quality":"Equal","values":[8,58,61,60,56,61],"isMain":false},{"metricName":"# contigs (>= 50000 bp)","quality":"Equal","values":[0,32,31,32,36,34],"isMain":false},{"metricName":"Largest contig","quality":"More is better","values":[32037,254042,257117,250596,250924,257124],"isMain":true},{"metricName":"Total length","quality":"More is better","values":[4585562,4573562,4617814,4649759,4566065,4535114],"isMain":true},{"metricName":"Total length (>= 1000 bp)","quality":"More is better","values":[4487944,4550200,4605531,4621053,4542018,4529243],"isMain":true},{"metricName":"Total length (>= 5000 bp)","quality":"More is better","values":[3403029,4517956,4574972,4593026,4512355,4494249],"isMain":false},{"metricName":"Total length (>= 10000 bp)","quality":"More is better","values":[1916431,4441920,4496777,4485345,4437352,4429557],"isMain":true},{"metricName":"Total length (>= 25000 bp)","quality":"More is better","values":[221154,4099306,4157209,4120698,4039019,4139231],"isMain":false},{"metricName":"Total length (>= 50000 bp)","quality":"More is better","values":[0,3124727,3025059,3069315,3302223,3151403],"isMain":true},{"metricName":"N50","quality":"More is better","values":[8535,76350,71670,71349,80042,72168],"isMain":false},{"metricName":"N75","quality":"More is better","values":[4916,43541,42814,40912,46310,43000],"isMain":false},{"metricName":"L50","quality":"Less is better","values":[170,19,20,21,21,20],"isMain":false},{"metricName":"L75","quality":"Less is better","values":[347,39,41,42,39,40],"isMain":false},{"metricName":"GC (%)","quality":"Equal","values":["65.31","65.34","65.28","65.16","65.33","65.37"],"isMain":false}]],["K-mer-based statistics",[]],["Predicted genes",[]],["Similarity statistics",[]],["Reference statistics",[{"metricName":"Reference length","quality":"Equal","values":[4587315,4587315,4587315,4587315,4587315,4587315],"isMain":false},{"metricName":"Reference fragments","quality":"Equal","values":[38,38,38,38,38,38],"isMain":false},{"metricName":"Reference GC (%)","quality":"Equal","values":["65.29","65.29","65.29","65.29","65.29","65.29"],"isMain":false}]]],"subreferences":[],"subreports":[],"minContig":500},{"date":"10 February 2020, Monday, 19:42:38","assembliesNames":["megahit-103-df","megahit-113-df","megahit-113-ml","megahit-113-ms","megahit-129-df","metaSPAdes"],"referenceName":"GCF\_000259235.1\_ASM25923v1","order":[0,1,2,3,4,5],"report":[["Genome statistics",[{"metricName":"Genome fraction (%)","quality":"More is better","values":["66.958","67.316","75.807","73.195","70.956","25.200"],"isMain":true},{"metricName":"Duplication ratio","quality":"Less is better","values":["1.077","1.125","1.138","1.133","1.153","1.026"],"isMain":true},{"metricName":"Largest alignment","quality":"More is better","values":[6187,15980,6681,5899,6910,9430],"isMain":true},{"metricName":"Total aligned length","quality":"More is better","values":[1154318,1211492,1377683,1325198,1310231,413586],"isMain":true},{"metricName":"NG50","quality":"More is better","values":[815,1200,1027,997,1037,null],"isMain":false},{"metricName":"NG75","quality":"More is better","values":[null,525,648,607,620,null],"isMain":false},{"metricName":"NA50","quality":"More is better","values":[1061,1681,1121,1157,1230,1247],"isMain":false},{"metricName":"NA75","quality":"More is better","values":[725,886,750,770,774,780],"isMain":false},{"metricName":"NGA50","quality":"More is better","values":[803,1136,1007,968,1001,null],"isMain":true},{"metricName":"NGA75","quality":"More is better","values":[null,500,632,587,593,null],"isMain":false},{"metricName":"LG50","quality":"Less is better","values":[566,341,492,481,413,null],"isMain":false},{"metricName":"LG75","quality":"Less is better","values":[null,867,990,996,920,null],"isMain":false},{"metricName":"LA50","quality":"Less is better","values":[333,217,407,369,308,97],"isMain":false},{"metricName":"LA75","quality":"Less is better","values":[666,473,788,729,656,205],"isMain":false},{"metricName":"LGA50","quality":"Less is better","values":[575,355,507,494,433,null],"isMain":true},{"metricName":"LGA75","quality":"Less is better","values":[null,905,1019,1024,960,null],"isMain":false}]],["Reads mapping",[]],["Misassemblies",[{"metricName":"# misassemblies","quality":"Less is better","values":[2,6,10,6,9,0],"isMain":true},{"metricName":" # relocations","quality":"Less is better","values":[2,6,7,5,8,0],"isMain":false},{"metricName":" # translocations","quality":"Less is better","values":[0,0,0,0,0,0],"isMain":false},{"metricName":" # inversions","quality":"Less is better","values":[0,0,3,1,1,0],"isMain":false},{"metricName":" # interspecies translocations","quality":"Less is better","values":[9,16,20,20,17,5],"isMain":false},{"metricName":"# misassembled contigs","quality":"Less is better","values":[2,6,10,6,9,0],"isMain":false},{"metricName":"Misassembled contigs length","quality":"Less is better","values":[2879,13756,23112,10576,20932,0],"isMain":true},{"metricName":"# possibly misassembled contigs","quality":"Less is better","values":[0,0,0,0,0,0],"isMain":false},{"metricName":" # possible misassemblies","quality":"Less is better","values":[0,0,0,0,0,0],"isMain":false},{"metricName":"# local misassemblies","quality":"Less is better","values":[9,10,9,7,4,0],"isMain":false},{"metricName":"# scaffold gap ext. mis.","quality":"Less is better","values":[0,0,0,0,0,0],"isMain":false},{"metricName":"# scaffold gap loc. mis.","quality":"Less is better","values":[0,0,0,0,0,0],"isMain":false},{"metricName":"# unaligned mis. contigs","quality":"Less is better","values":[0,1,0,0,2,1],"isMain":false}]],["Unaligned",[{"metricName":"# fully unaligned contigs","quality":"Less is better","values":[0,0,0,1,1,1],"isMain":false},{"metricName":"Fully unaligned length","quality":"Less is better","values":[0,0,0,987,665,618],"isMain":false},{"metricName":"# partially unaligned contigs","quality":"Less is better","values":[3,10,7,9,12,1],"isMain":false},{"metricName":"Partially unaligned length","quality":"Less is better","values":[4814,11581,7246,8603,17216,1171],"isMain":false}]],["Mismatches",[{"metricName":"# mismatches","quality":"Less is better","values":[11823,18099,20487,20358,18321,4792],"isMain":false},{"metricName":"# indels","quality":"Less is better","values":[99,314,178,148,184,80],"isMain":false},{"metricName":"Indels length","quality":"Less is better","values":[308,1030,623,494,621,276],"isMain":false},{"metricName":"# mismatches per 100 kbp","quality":"Less is better","values":["1098.38","1672.48","1681.11","1730.12","1606.16","1182.90"],"isMain":true},{"metricName":"# indels per 100 kbp","quality":"Less is better","values":["9.20","29.02","14.61","12.58","16.13","19.75"],"isMain":true},{"metricName":" # indels (<= 5 bp)","quality":"Less is better","values":[90,278,153,132,158,71],"isMain":false},{"metricName":" # indels (> 5 bp)","quality":"Less is better","values":[9,36,25,16,26,9],"isMain":false},{"metricName":"# N's","quality":"Less is better","values":[0,0,0,0,0,0],"isMain":false},{"metricName":"# N's per 100 kbp","quality":"Less is better","values":["0.00","0.00","0.00","0.00","0.00","0.00"],"isMain":true}]],["Statistics without reference",[{"metricName":"# contigs","quality":"Equal","values":[1138,912,1320,1245,1147,366],"isMain":true},{"metricName":"# contigs (>= 1000 bp)","quality":"Equal","values":[384,420,519,479,441,141],"isMain":false},{"metricName":"# contigs (>= 5000 bp)","quality":"Equal","values":[4,11,4,3,10,4],"isMain":false},{"metricName":"# contigs (>= 10000 bp)","quality":"Equal","values":[0,1,0,0,0,0],"isMain":false},{"metricName":"# contigs (>= 25000 bp)","quality":"Equal","values":[0,0,0,0,0,0],"isMain":false},{"metricName":"# contigs (>= 50000 bp)","quality":"Equal","values":[0,0,0,0,0,0],"isMain":false},{"metricName":"Largest contig","quality":"More is better","values":[7477,15980,6696,5899,6912,9430],"isMain":true},{"metricName":"Total length","quality":"More is better","values":[1164010,1229031,1394647,1343078,1332785,417558],"isMain":true},{"metricName":"Total length (>= 1000 bp)","quality":"More is better","values":[641407,890045,831608,802228,833032,259228],"isMain":true},{"metricName":"Total length (>= 5000 bp)","quality":"More is better","values":[24261,76763,23610,16920,56867,26219],"isMain":false},{"metricName":"Total length (>= 10000 bp)","quality":"More is better","values":[0,15980,0,0,0,0],"isMain":true},{"metricName":"Total length (>= 25000 bp)","quality":"More is better","values":[0,0,0,0,0,0],"isMain":false},{"metricName":"Total length (>= 50000 bp)","quality":"More is better","values":[0,0,0,0,0,0],"isMain":true},{"metricName":"N50","quality":"More is better","values":[1091,1735,1157,1179,1281,1271],"isMain":false},{"metricName":"N75","quality":"More is better","values":[741,932,769,797,803,799],"isMain":false},{"metricName":"L50","quality":"Less is better","values":[327,209,394,360,293,96],"isMain":false},{"metricName":"L75","quality":"Less is better","values":[655,453,765,710,628,202],"isMain":false},{"metricName":"GC (%)","quality":"Equal","values":["38.87","38.91","38.98","39.02","38.97","37.76"],"isMain":false}]],["K-mer-based statistics",[]],["Predicted genes",[]],["Similarity statistics",[]],["Reference statistics",[{"metricName":"Reference length","quality":"Equal","values":[1607584,1607584,1607584,1607584,1607584,1607584],"isMain":false},{"metricName":"Reference fragments","quality":"Equal","values":[2,2,2,2,2,2],"isMain":false},{"metricName":"Reference GC (%)","quality":"Equal","values":["39.04","39.04","39.04","39.04","39.04","39.04"],"isMain":false}]]],"subreferences":[],"subreports":[],"minContig":500},{"date":"10 February 2020, Monday, 19:42:28","assembliesNames":["megahit-103-df","megahit-113-df","megahit-113-ml","megahit-113-ms","megahit-129-df","metaSPAdes"],"referenceName":"GCF\_000261025.1\_ASM26102v1","order":[0,1,2,3,4,5],"report":[["Genome statistics",[{"metricName":"Genome fraction (%)","quality":"More is better","values":["2.249","2.668","2.660","2.841","4.388","2.955"],"isMain":true},{"metricName":"Duplication ratio","quality":"Less is better","values":["1.027","1.052","1.037","1.033","1.030","1.008"],"isMain":true},{"metricName":"Largest alignment","quality":"More is better","values":[1930,16093,4184,5124,33985,51870],"isMain":true},{"metricName":"Total aligned length","quality":"More is better","values":[61272,72437,72628,77354,118968,79769],"isMain":true},{"metricName":"NA50","quality":"More is better","values":[678,890,725,719,1085,51870],"isMain":false},{"metricName":"NA75","quality":"More is better","values":[536,585,567,545,647,1024],"isMain":false},{"metricName":"NGA50","quality":"More is better","values":[null,null,null,null,null,null],"isMain":true},{"metricName":"LA50","quality":"Less is better","values":[33,16,30,34,17,1],"isMain":false},{"metricName":"LA75","quality":"Less is better","values":[60,43,60,69,56,8],"isMain":false}]],["Reads mapping",[]],["Misassemblies",[{"metricName":"# misassemblies","quality":"Less is better","values":[1,2,0,3,0,0],"isMain":true},{"metricName":" # relocations","quality":"Less is better","values":[1,1,0,1,0,0],"isMain":false},{"metricName":" # translocations","quality":"Less is better","values":[0,0,0,0,0,0],"isMain":false},{"metricName":" # inversions","quality":"Less is better","values":[0,1,0,2,0,0],"isMain":false},{"metricName":" # interspecies translocations","quality":"Less is better","values":[5,7,3,10,11,4],"isMain":false},{"metricName":"# misassembled contigs","quality":"Less is better","values":[1,2,0,3,0,0],"isMain":false},{"metricName":"Misassembled contigs length","quality":"Less is better","values":[531,1801,0,2216,0,0],"isMain":true},{"metricName":"# possibly misassembled contigs","quality":"Less is better","values":[0,0,0,0,0,0],"isMain":false},{"metricName":" # possible misassemblies","quality":"Less is better","values":[0,0,0,0,0,0],"isMain":false},{"metricName":"# local misassemblies","quality":"Less is better","values":[0,1,0,1,1,1],"isMain":false},{"metricName":"# scaffold gap ext. mis.","quality":"Less is better","values":[0,0,0,0,0,0],"isMain":false},{"metricName":"# scaffold gap loc. mis.","quality":"Less is better","values":[0,0,0,0,0,0],"isMain":false},{"metricName":"# unaligned mis. contigs","quality":"Less is better","values":[0,0,0,1,0,0],"isMain":false}]],["Unaligned",[{"metricName":"# fully unaligned contigs","quality":"Less is better","values":[1,0,0,0,0,0],"isMain":false},{"metricName":"Fully unaligned length","quality":"Less is better","values":[767,0,0,0,0,0],"isMain":false},{"metricName":"# partially unaligned contigs","quality":"Less is better","values":[1,0,0,3,3,2],"isMain":false},{"metricName":"Partially unaligned length","quality":"Less is better","values":[859,0,0,7626,2477,1341],"isMain":false}]],["Mismatches",[{"metricName":"# mismatches","quality":"Less is better","values":[895,1301,1601,1530,2251,1840],"isMain":false},{"metricName":"# indels","quality":"Less is better","values":[3,13,16,21,15,17],"isMain":false},{"metricName":"Indels length","quality":"Less is better","values":[3,16,20,21,16,18],"isMain":false},{"metricName":"# mismatches per 100 kbp","quality":"Less is better","values":["1474.44","1806.22","2230.05","1994.78","1900.57","2306.60"],"isMain":true},{"metricName":"# indels per 100 kbp","quality":"Less is better","values":["4.94","18.05","22.29","27.38","12.66","21.31"],"isMain":true},{"metricName":" # indels (<= 5 bp)","quality":"Less is better","values":[3,13,16,21,15,17],"isMain":false},{"metricName":" # indels (> 5 bp)","quality":"Less is better","values":[0,0,0,0,0,0],"isMain":false},{"metricName":"# N's","quality":"Less is better","values":[0,0,0,0,0,0],"isMain":false},{"metricName":"# N's per 100 kbp","quality":"Less is better","values":["0.00","0.00","0.00","0.00","0.00","0.00"],"isMain":true}]],["Statistics without reference",[{"metricName":"# contigs","quality":"Equal","values":[88,75,93,91,108,38],"isMain":true},{"metricName":"# contigs (>= 1000 bp)","quality":"Equal","values":[11,13,13,17,20,9],"isMain":false},{"metricName":"# contigs (>= 5000 bp)","quality":"Equal","values":[0,1,0,2,2,1],"isMain":false},{"metricName":"# contigs (>= 10000 bp)","quality":"Equal","values":[0,1,0,0,1,1],"isMain":false},{"metricName":"# contigs (>= 25000 bp)","quality":"Equal","values":[0,0,0,0,1,1],"isMain":false},{"metricName":"# contigs (>= 50000 bp)","quality":"Equal","values":[0,0,0,0,0,1],"isMain":false},{"metricName":"Largest contig","quality":"More is better","values":[1930,16480,4184,8841,33985,52006],"isMain":true},{"metricName":"Total length","quality":"More is better","values":[63968,75775,74426,86895,124427,81782],"isMain":true},{"metricName":"Total length (>= 1000 bp)","quality":"More is better","values":[14926,36006,23450,37679,66682,62607],"isMain":true},{"metricName":"Total length (>= 5000 bp)","quality":"More is better","values":[0,16480,0,13965,40684,52006],"isMain":false},{"metricName":"Total length (>= 10000 bp)","quality":"More is better","values":[0,16480,0,0,33985,52006],"isMain":true},{"metricName":"Total length (>= 25000 bp)","quality":"More is better","values":[0,0,0,0,33985,52006],"isMain":false},{"metricName":"Total length (>= 50000 bp)","quality":"More is better","values":[0,0,0,0,0,52006],"isMain":true},{"metricName":"N50","quality":"More is better","values":[697,916,727,963,1159,52006],"isMain":false},{"metricName":"N75","quality":"More is better","values":[563,621,584,639,681,1031],"isMain":false},{"metricName":"L50","quality":"Less is better","values":[32,16,30,23,16,1],"isMain":false},{"metricName":"L75","quality":"Less is better","values":[58,41,59,53,54,8],"isMain":false},{"metricName":"GC (%)","quality":"Equal","values":["40.95","41.76","39.60","41.48","42.19","42.35"],"isMain":false}]],["K-mer-based statistics",[]],["Predicted genes",[]],["Similarity statistics",[]],["Reference statistics",[{"metricName":"Reference length","quality":"Equal","values":[2699437,2699437,2699437,2699437,2699437,2699437],"isMain":false},{"metricName":"Reference fragments","quality":"Equal","values":[2,2,2,2,2,2],"isMain":false},{"metricName":"Reference GC (%)","quality":"Equal","values":["43.48","43.48","43.48","43.48","43.48","43.48"],"isMain":false}]]],"subreferences":[],"subreports":[],"minContig":500},{"date":"10 February 2020, Monday, 19:43:05","assembliesNames":["megahit-103-df","megahit-113-df","megahit-113-ml","megahit-113-ms","megahit-129-df","metaSPAdes"],"referenceName":"GCF\_000265365.1\_ASM26536v1","order":[0,1,2,3,4,5],"report":[["Genome statistics",[{"metricName":"Genome fraction (%)","quality":"More is better","values":["26.175","98.082","98.045","98.340","98.077","97.160"],"isMain":true},{"metricName":"Duplication ratio","quality":"Less is better","values":["1.043","1.037","1.034","1.043","1.057","1.001"],"isMain":true},{"metricName":"Largest alignment","quality":"More is better","values":[2379,207726,263235,263293,192186,263142],"isMain":true},{"metricName":"Total aligned length","quality":"More is better","values":[993316,3791905,3780985,3823089,3863832,3630578],"isMain":true},{"metricName":"NG50","quality":"More is better","values":[null,71101,81679,89334,78497,98453],"isMain":false},{"metricName":"NG75","quality":"More is better","values":[null,38003,54636,58331,41285,59578],"isMain":false},{"metricName":"NA50","quality":"More is better","values":[676,67002,81651,57405,62802,103316],"isMain":false},{"metricName":"NA75","quality":"More is better","values":[546,33211,51966,9454,27210,60608],"isMain":false},{"metricName":"NGA50","quality":"More is better","values":[null,71101,81651,63279,67294,98453],"isMain":true},{"metricName":"NGA75","quality":"More is better","values":[null,38003,54636,31075,36596,59578],"isMain":false},{"metricName":"LG50","quality":"Less is better","values":[null,17,14,13,16,12],"isMain":false},{"metricName":"LG75","quality":"Less is better","values":[null,35,28,26,32,23],"isMain":false},{"metricName":"LA50","quality":"Less is better","values":[577,18,14,24,21,11],"isMain":false},{"metricName":"LA75","quality":"Less is better","values":[1018,37,29,73,44,22],"isMain":false},{"metricName":"LGA50","quality":"Less is better","values":[null,17,14,18,19,12],"isMain":true},{"metricName":"LGA75","quality":"Less is better","values":[null,35,28,37,37,23],"isMain":false}]],["Reads mapping",[]],["Misassemblies",[{"metricName":"# misassemblies","quality":"Less is better","values":[8,10,13,26,16,0],"isMain":true},{"metricName":" # relocations","quality":"Less is better","values":[0,9,12,20,12,0],"isMain":false},{"metricName":" # translocations","quality":"Less is better","values":[0,0,0,0,0,0],"isMain":false},{"metricName":" # inversions","quality":"Less is better","values":[8,1,1,6,4,0],"isMain":false},{"metricName":" # interspecies translocations","quality":"Less is better","values":[115,25,19,48,21,1],"isMain":false},{"metricName":"# misassembled contigs","quality":"Less is better","values":[8,10,13,24,15,0],"isMain":false},{"metricName":"Misassembled contigs length","quality":"Less is better","values":[4540,98930,211257,394717,372164,0],"isMain":true},{"metricName":"# possibly misassembled contigs","quality":"Less is better","values":[1,0,0,0,0,0],"isMain":false},{"metricName":" # possible misassemblies","quality":"Less is better","values":[1,0,0,0,0,0],"isMain":false},{"metricName":"# local misassemblies","quality":"Less is better","values":[5,1,5,1,12,0],"isMain":false},{"metricName":"# scaffold gap ext. mis.","quality":"Less is better","values":[0,0,0,0,0,0],"isMain":false},{"metricName":"# scaffold gap loc. mis.","quality":"Less is better","values":[0,0,0,0,0,0],"isMain":false},{"metricName":"# unaligned mis. contigs","quality":"Less is better","values":[1,0,1,0,0,0],"isMain":false}]],["Unaligned",[{"metricName":"# fully unaligned contigs","quality":"Less is better","values":[2,2,3,5,4,0],"isMain":false},{"metricName":"Fully unaligned length","quality":"Less is better","values":[1127,1341,1973,3041,2257,0],"isMain":false},{"metricName":"# partially unaligned contigs","quality":"Less is better","values":[52,14,7,30,8,0],"isMain":false},{"metricName":"Partially unaligned length","quality":"Less is better","values":[45502,20014,6507,680628,169488,0],"isMain":false}]],["Mismatches",[{"metricName":"# mismatches","quality":"Less is better","values":[3577,5618,5766,6975,7602,577],"isMain":false},{"metricName":"# indels","quality":"Less is better","values":[4,52,104,87,57,21],"isMain":false},{"metricName":"Indels length","quality":"Less is better","values":[4,497,644,1122,605,72],"isMain":false},{"metricName":"# mismatches per 100 kbp","quality":"Less is better","values":["365.96","153.39","157.49","189.94","207.57","15.90"],"isMain":true},{"metricName":"# indels per 100 kbp","quality":"Less is better","values":["0.41","1.42","2.84","2.37","1.56","0.58"],"isMain":true},{"metricName":" # indels (<= 5 bp)","quality":"Less is better","values":[4,44,81,55,41,18],"isMain":false},{"metricName":" # indels (> 5 bp)","quality":"Less is better","values":[0,8,23,32,16,3],"isMain":false},{"metricName":"# N's","quality":"Less is better","values":[0,0,0,0,0,0],"isMain":false},{"metricName":"# N's per 100 kbp","quality":"Less is better","values":["0.00","0.00","0.00","0.00","0.00","0.00"],"isMain":true}]],["Statistics without reference",[{"metricName":"# contigs","quality":"Equal","values":[1471,392,368,454,509,98],"isMain":true},{"metricName":"# contigs (>= 1000 bp)","quality":"Equal","values":[188,135,111,130,118,85],"isMain":false},{"metricName":"# contigs (>= 5000 bp)","quality":"Equal","values":[0,80,70,86,78,57],"isMain":false},{"metricName":"# contigs (>= 10000 bp)","quality":"Equal","values":[0,69,56,73,71,48],"isMain":false},{"metricName":"# contigs (>= 25000 bp)","quality":"Equal","values":[0,40,35,47,44,33],"isMain":false},{"metricName":"# contigs (>= 50000 bp)","quality":"Equal","values":[0,29,29,34,27,27],"isMain":false},{"metricName":"Largest contig","quality":"More is better","values":[2659,207726,263235,263293,228664,263142],"isMain":true},{"metricName":"Total length","quality":"More is better","values":[1065797,3819733,3795470,4514588,4042900,3630964],"isMain":true},{"metricName":"Total length (>= 1000 bp)","quality":"More is better","values":[235855,3665638,3637283,4313923,3815199,3621610],"isMain":true},{"metricName":"Total length (>= 5000 bp)","quality":"More is better","values":[0,3546230,3548947,4211871,3721369,3565592],"isMain":false},{"metricName":"Total length (>= 10000 bp)","quality":"More is better","values":[0,3468647,3452242,4117854,3674091,3501764],"isMain":true},{"metricName":"Total length (>= 25000 bp)","quality":"More is better","values":[0,2990341,3109658,3689322,3225078,3235788],"isMain":false},{"metricName":"Total length (>= 50000 bp)","quality":"More is better","values":[0,2580482,2890473,3274310,2596557,3030371],"isMain":true},{"metricName":"N50","quality":"More is better","values":[707,67016,81679,81610,68905,103316],"isMain":false},{"metricName":"N75","quality":"More is better","values":[580,35231,51966,35605,35825,60608],"isMain":false},{"metricName":"L50","quality":"Less is better","values":[551,18,14,18,18,11],"isMain":false},{"metricName":"L75","quality":"Less is better","values":[971,36,29,37,38,22],"isMain":false},{"metricName":"GC (%)","quality":"Equal","values":["56.96","56.61","56.65","55.63","56.47","56.80"],"isMain":false}]],["K-mer-based statistics",[]],["Predicted genes",[]],["Similarity statistics",[]],["Reference statistics",[{"metricName":"Reference length","quality":"Equal","values":[3734239,3734239,3734239,3734239,3734239,3734239],"isMain":false},{"metricName":"Reference fragments","quality":"Equal","values":[1,1,1,1,1,1],"isMain":false},{"metricName":"Reference GC (%)","quality":"Equal","values":["56.65","56.65","56.65","56.65","56.65","56.65"],"isMain":false}]]],"subreferences":[],"subreports":[],"minContig":500},{"date":"10 February 2020, Monday, 19:42:29","assembliesNames":["megahit-103-df","megahit-113-df","megahit-113-ml","megahit-113-ms","megahit-129-df","metaSPAdes"],"referenceName":"GCF\_000270225.1\_ASM27022v1","order":[0,1,2,3,4,5],"report":[["Genome statistics",[{"metricName":"Genome fraction (%)","quality":"More is better","values":["2.279","3.523","2.569","2.451","3.174","0.725"],"isMain":true},{"metricName":"Duplication ratio","quality":"Less is better","values":["1.019","1.013","1.020","1.022","1.042","1.011"],"isMain":true},{"metricName":"Largest alignment","quality":"More is better","values":[10592,22751,8427,11620,13248,7830],"isMain":true},{"metricName":"Total aligned length","quality":"More is better","values":[53331,82921,60293,58063,74649,16954],"isMain":true},{"metricName":"NA50","quality":"More is better","values":[1428,10099,2605,521,2037,1116],"isMain":false},{"metricName":"NA75","quality":"More is better","values":[786,1670,836,null,703,747],"isMain":false},{"metricName":"NGA50","quality":"More is better","values":[null,null,null,null,null,null],"isMain":true},{"metricName":"LA50","quality":"Less is better","values":[9,3,7,38,7,3],"isMain":false},{"metricName":"LA75","quality":"Less is better","values":[22,9,19,null,23,8],"isMain":false}]],["Reads mapping",[]],["Misassemblies",[{"metricName":"# misassemblies","quality":"Less is better","values":[0,1,1,2,5,0],"isMain":true},{"metricName":" # relocations","quality":"Less is better","values":[0,0,1,1,3,0],"isMain":false},{"metricName":" # translocations","quality":"Less is better","values":[0,0,0,1,1,0],"isMain":false},{"metricName":" # inversions","quality":"Less is better","values":[0,1,0,0,1,0],"isMain":false},{"metricName":" # interspecies translocations","quality":"Less is better","values":[3,3,2,9,11,3],"isMain":false},{"metricName":"# misassembled contigs","quality":"Less is better","values":[0,1,1,2,5,0],"isMain":false},{"metricName":"Misassembled contigs length","quality":"Less is better","values":[0,511,532,7208,35562,0],"isMain":true},{"metricName":"# possibly misassembled contigs","quality":"Less is better","values":[0,0,0,0,0,0],"isMain":false},{"metricName":" # possible misassemblies","quality":"Less is better","values":[0,0,0,0,0,0],"isMain":false},{"metricName":"# local misassemblies","quality":"Less is better","values":[0,0,0,0,1,0],"isMain":false},{"metricName":"# scaffold gap ext. mis.","quality":"Less is better","values":[0,0,0,0,0,0],"isMain":false},{"metricName":"# scaffold gap loc. mis.","quality":"Less is better","values":[0,0,0,0,0,0],"isMain":false},{"metricName":"# unaligned mis. contigs","quality":"Less is better","values":[1,0,0,2,0,0],"isMain":false}]],["Unaligned",[{"metricName":"# fully unaligned contigs","quality":"Less is better","values":[0,0,0,0,0,0],"isMain":false},{"metricName":"Fully unaligned length","quality":"Less is better","values":[0,0,0,0,0,0],"isMain":false},{"metricName":"# partially unaligned contigs","quality":"Less is better","values":[1,1,0,5,2,2],"isMain":false},{"metricName":"Partially unaligned length","quality":"Less is better","values":[1499,1521,0,53498,2193,1682],"isMain":false}]],["Mismatches",[{"metricName":"# mismatches","quality":"Less is better","values":[449,889,861,872,1275,154],"isMain":false},{"metricName":"# indels","quality":"Less is better","values":[6,26,27,17,45,1],"isMain":false},{"metricName":"Indels length","quality":"Less is better","values":[35,60,40,46,264,1],"isMain":false},{"metricName":"# mismatches per 100 kbp","quality":"Less is better","values":["841.86","1078.31","1432.40","1520.36","1716.85","908.39"],"isMain":true},{"metricName":"# indels per 100 kbp","quality":"Less is better","values":["11.25","31.54","44.92","29.64","60.59","5.90"],"isMain":true},{"metricName":" # indels (<= 5 bp)","quality":"Less is better","values":[5,23,27,16,38,1],"isMain":false},{"metricName":" # indels (> 5 bp)","quality":"Less is better","values":[1,3,0,1,7,0],"isMain":false},{"metricName":"# N's","quality":"Less is better","values":[0,0,0,0,0,0],"isMain":false},{"metricName":"# N's per 100 kbp","quality":"Less is better","values":["0.00","0.00","0.00","0.00","0.00","0.00"],"isMain":true}]],["Statistics without reference",[{"metricName":"# contigs","quality":"Equal","values":[42,35,44,40,48,13],"isMain":true},{"metricName":"# contigs (>= 1000 bp)","quality":"Equal","values":[17,13,13,14,17,5],"isMain":false},{"metricName":"# contigs (>= 5000 bp)","quality":"Equal","values":[2,3,4,4,3,1],"isMain":false},{"metricName":"# contigs (>= 10000 bp)","quality":"Equal","values":[1,3,0,3,2,0],"isMain":false},{"metricName":"# contigs (>= 25000 bp)","quality":"Equal","values":[0,0,0,1,0,0],"isMain":false},{"metricName":"# contigs (>= 50000 bp)","quality":"Equal","values":[0,0,0,0,0,0],"isMain":false},{"metricName":"Largest contig","quality":"More is better","values":[10596,22751,8427,46774,20093,7834],"isMain":true},{"metricName":"Total length","quality":"More is better","values":[55848,85025,61332,112115,79583,18818],"isMain":true},{"metricName":"Total length (>= 1000 bp)","quality":"More is better","values":[39469,70790,41281,95880,60762,12511],"isMain":true},{"metricName":"Total length (>= 5000 bp)","quality":"More is better","values":[16084,51180,24820,77443,35658,7834],"isMain":false},{"metricName":"Total length (>= 10000 bp)","quality":"More is better","values":[10596,51180,0,71824,30576,0],"isMain":true},{"metricName":"Total length (>= 25000 bp)","quality":"More is better","values":[0,0,0,46774,0,0],"isMain":false},{"metricName":"Total length (>= 50000 bp)","quality":"More is better","values":[0,0,0,0,0,0],"isMain":true},{"metricName":"N50","quality":"More is better","values":[1699,10099,2605,12775,4805,1116],"isMain":false},{"metricName":"N75","quality":"More is better","values":[854,1881,836,2468,1081,903],"isMain":false},{"metricName":"L50","quality":"Less is better","values":[8,3,7,2,4,3],"isMain":false},{"metricName":"L75","quality":"Less is better","values":[20,8,19,6,17,7],"isMain":false},{"metricName":"GC (%)","quality":"Equal","values":["44.51","46.50","46.63","43.26","47.50","44.54"],"isMain":false}]],["K-mer-based statistics",[]],["Predicted genes",[]],["Similarity statistics",[]],["Reference statistics",[{"metricName":"Reference length","quality":"Equal","values":[2339897,2339897,2339897,2339897,2339897,2339897],"isMain":false},{"metricName":"Reference fragments","quality":"Equal","values":[2,2,2,2,2,2],"isMain":false},{"metricName":"Reference GC (%)","quality":"Equal","values":["48.34","48.34","48.34","48.34","48.34","48.34"],"isMain":false}]]],"subreferences":[],"subreports":[],"minContig":500},{"date":"10 February 2020, Monday, 19:42:51","assembliesNames":["megahit-103-df","megahit-113-df","megahit-113-ml","megahit-113-ms","megahit-129-df","metaSPAdes"],"referenceName":"GCF\_000273035.1\_Bact\_dore\_CL02T00C15\_V1","order":[0,1,2,3,4,5],"report":[["Genome statistics",[{"metricName":"Genome fraction (%)","quality":"More is better","values":["9.665","11.667","12.161","12.328","11.044","6.421"],"isMain":true},{"metricName":"Duplication ratio","quality":"Less is better","values":["1.025","1.021","1.026","1.026","1.023","1.025"],"isMain":true},{"metricName":"Largest alignment","quality":"More is better","values":[7676,23014,4613,17982,22010,9054],"isMain":true},{"metricName":"Total aligned length","quality":"More is better","values":[586954,707301,741466,749081,670169,387793],"isMain":true},{"metricName":"NA50","quality":"More is better","values":[820,951,785,805,812,729],"isMain":false},{"metricName":"NA75","quality":"More is better","values":[611,676,601,610,588,588],"isMain":false},{"metricName":"NGA50","quality":"More is better","values":[null,null,null,null,null,null],"isMain":true},{"metricName":"LA50","quality":"Less is better","values":[230,209,318,305,270,176],"isMain":false},{"metricName":"LA75","quality":"Less is better","values":[444,437,593,585,543,327],"isMain":false}]],["Reads mapping",[]],["Misassemblies",[{"metricName":"# misassemblies","quality":"Less is better","values":[3,6,2,2,5,0],"isMain":true},{"metricName":" # relocations","quality":"Less is better","values":[0,1,0,0,2,0],"isMain":false},{"metricName":" # translocations","quality":"Less is better","values":[2,4,2,1,3,0],"isMain":false},{"metricName":" # inversions","quality":"Less is better","values":[1,1,0,1,0,0],"isMain":false},{"metricName":" # interspecies translocations","quality":"Less is better","values":[26,31,13,32,34,16],"isMain":false},{"metricName":"# misassembled contigs","quality":"Less is better","values":[3,6,2,2,4,0],"isMain":false},{"metricName":"Misassembled contigs length","quality":"Less is better","values":[3103,9820,2456,1561,10242,0],"isMain":true},{"metricName":"# possibly misassembled contigs","quality":"Less is better","values":[0,1,0,0,1,0],"isMain":false},{"metricName":" # possible misassemblies","quality":"Less is better","values":[0,1,0,0,1,0],"isMain":false},{"metricName":"# local misassemblies","quality":"Less is better","values":[0,1,1,1,1,1],"isMain":false},{"metricName":"# scaffold gap ext. mis.","quality":"Less is better","values":[0,0,0,0,0,0],"isMain":false},{"metricName":"# scaffold gap loc. mis.","quality":"Less is better","values":[0,0,0,0,0,0],"isMain":false},{"metricName":"# unaligned mis. contigs","quality":"Less is better","values":[1,0,0,1,1,0],"isMain":false}]],["Unaligned",[{"metricName":"# fully unaligned contigs","quality":"Less is better","values":[1,1,2,0,1,0],"isMain":false},{"metricName":"Fully unaligned length","quality":"Less is better","values":[502,825,1446,0,1457,0],"isMain":false},{"metricName":"# partially unaligned contigs","quality":"Less is better","values":[3,7,4,10,12,3],"isMain":false},{"metricName":"Partially unaligned length","quality":"Less is better","values":[2673,8022,3358,9172,63794,2251],"isMain":false}]],["Mismatches",[{"metricName":"# mismatches","quality":"Less is better","values":[7514,9343,14360,14091,9314,5185],"isMain":false},{"metricName":"# indels","quality":"Less is better","values":[40,38,59,55,35,13],"isMain":false},{"metricName":"Indels length","quality":"Less is better","values":[82,82,95,115,120,15],"isMain":false},{"metricName":"# mismatches per 100 kbp","quality":"Less is better","values":["1293.37","1332.24","1964.56","1901.65","1403.06","1343.48"],"isMain":true},{"metricName":"# indels per 100 kbp","quality":"Less is better","values":["6.89","5.42","8.07","7.42","5.27","3.37"],"isMain":true},{"metricName":" # indels (<= 5 bp)","quality":"Less is better","values":[37,35,57,48,32,13],"isMain":false},{"metricName":" # indels (> 5 bp)","quality":"Less is better","values":[3,3,2,7,3,0],"isMain":false},{"metricName":"# N's","quality":"Less is better","values":[0,0,0,0,0,0],"isMain":false},{"metricName":"# N's per 100 kbp","quality":"Less is better","values":["0.00","0.00","0.00","0.00","0.00","0.00"],"isMain":true}]],["Statistics without reference",[{"metricName":"# contigs","quality":"Equal","values":[699,728,919,910,752,499],"isMain":true},{"metricName":"# contigs (>= 1000 bp)","quality":"Equal","values":[142,203,180,174,193,75],"isMain":false},{"metricName":"# contigs (>= 5000 bp)","quality":"Equal","values":[3,7,0,3,5,2],"isMain":false},{"metricName":"# contigs (>= 10000 bp)","quality":"Equal","values":[0,1,0,1,2,0],"isMain":false},{"metricName":"# contigs (>= 25000 bp)","quality":"Equal","values":[0,0,0,0,1,0],"isMain":false},{"metricName":"# contigs (>= 50000 bp)","quality":"Equal","values":[0,0,0,0,0,0],"isMain":false},{"metricName":"Largest contig","quality":"More is better","values":[7676,23014,4613,18115,49839,9054],"isMain":true},{"metricName":"Total length","quality":"More is better","values":[598568,725107,754703,769322,744478,397982],"isMain":true},{"metricName":"Total length (>= 1000 bp)","quality":"More is better","values":[224791,364526,262852,274600,369223,117239],"isMain":true},{"metricName":"Total length (>= 5000 bp)","quality":"More is better","values":[19368,63999,0,30928,92974,16495],"isMain":false},{"metricName":"Total length (>= 10000 bp)","quality":"More is better","values":[0,23014,0,18115,71849,0],"isMain":true},{"metricName":"Total length (>= 25000 bp)","quality":"More is better","values":[0,0,0,0,49839,0],"isMain":false},{"metricName":"Total length (>= 50000 bp)","quality":"More is better","values":[0,0,0,0,0,0],"isMain":true},{"metricName":"N50","quality":"More is better","values":[837,1001,806,831,992,752],"isMain":false},{"metricName":"N75","quality":"More is better","values":[629,699,616,629,675,612],"isMain":false},{"metricName":"L50","quality":"Less is better","values":[224,202,309,297,197,172],"isMain":false},{"metricName":"L75","quality":"Less is better","values":[432,420,578,567,430,319],"isMain":false},{"metricName":"GC (%)","quality":"Equal","values":["41.90","42.27","41.94","41.84","42.12","41.69"],"isMain":false}]],["K-mer-based statistics",[]],["Predicted genes",[]],["Similarity statistics",[]],["Reference statistics",[{"metricName":"Reference length","quality":"Equal","values":[6010738,6010738,6010738,6010738,6010738,6010738],"isMain":false},{"metricName":"Reference fragments","quality":"Equal","values":[95,95,95,95,95,95],"isMain":false},{"metricName":"Reference GC (%)","quality":"Equal","values":["42.15","42.15","42.15","42.15","42.15","42.15"],"isMain":false}]]],"subreferences":[],"subreports":[],"minContig":500},{"date":"10 February 2020, Monday, 19:42:51","assembliesNames":["megahit-103-df","megahit-113-df","megahit-113-ml","megahit-113-ms","megahit-129-df","metaSPAdes"],"referenceName":"GCF\_000273055.1\_Bact\_dore\_CL02T12C06\_V1","order":[0,1,2,3,4,5],"report":[["Genome statistics",[{"metricName":"Genome fraction (%)","quality":"More is better","values":["8.678","9.693","11.719","12.439","11.624","5.512"],"isMain":true},{"metricName":"Duplication ratio","quality":"Less is better","values":["1.023","1.021","1.027","1.031","1.020","1.018"],"isMain":true},{"metricName":"Largest alignment","quality":"More is better","values":[6271,7860,5071,7542,9806,3215],"isMain":true},{"metricName":"Total aligned length","quality":"More is better","values":[524964,586944,712489,757789,702750,332402],"isMain":true},{"metricName":"NA50","quality":"More is better","values":[806,910,786,756,913,783],"isMain":false},{"metricName":"NA75","quality":"More is better","values":[615,656,606,572,661,589],"isMain":false},{"metricName":"NGA50","quality":"More is better","values":[null,null,null,null,null,null],"isMain":true},{"metricName":"LA50","quality":"Less is better","values":[211,209,308,353,250,150],"isMain":false},{"metricName":"LA75","quality":"Less is better","values":[403,406,571,671,484,275],"isMain":false}]],["Reads mapping",[]],["Misassemblies",[{"metricName":"# misassemblies","quality":"Less is better","values":[2,3,3,5,6,1],"isMain":true},{"metricName":" # relocations","quality":"Less is better","values":[0,1,0,1,1,0],"isMain":false},{"metricName":" # translocations","quality":"Less is better","values":[2,2,1,3,5,1],"isMain":false},{"metricName":" # inversions","quality":"Less is better","values":[0,0,2,1,0,0],"isMain":false},{"metricName":" # interspecies translocations","quality":"Less is better","values":[21,21,15,31,24,4],"isMain":false},{"metricName":"# misassembled contigs","quality":"Less is better","values":[2,3,3,5,5,1],"isMain":false},{"metricName":"Misassembled contigs length","quality":"Less is better","values":[2667,4269,4069,4447,9480,742],"isMain":true},{"metricName":"# possibly misassembled contigs","quality":"Less is better","values":[0,0,0,1,0,0],"isMain":false},{"metricName":" # possible misassemblies","quality":"Less is better","values":[0,0,0,1,0,0],"isMain":false},{"metricName":"# local misassemblies","quality":"Less is better","values":[1,3,0,2,1,0],"isMain":false},{"metricName":"# scaffold gap ext. mis.","quality":"Less is better","values":[0,0,0,0,0,0],"isMain":false},{"metricName":"# scaffold gap loc. mis.","quality":"Less is better","values":[0,0,0,0,0,0],"isMain":false},{"metricName":"# unaligned mis. contigs","quality":"Less is better","values":[0,0,1,0,1,0],"isMain":false}]],["Unaligned",[{"metricName":"# fully unaligned contigs","quality":"Less is better","values":[0,0,0,0,0,0],"isMain":false},{"metricName":"Fully unaligned length","quality":"Less is better","values":[0,0,0,0,0,0],"isMain":false},{"metricName":"# partially unaligned contigs","quality":"Less is better","values":[4,7,1,9,5,1],"isMain":false},{"metricName":"Partially unaligned length","quality":"Less is better","values":[3145,6544,2484,65303,8434,769],"isMain":false}]],["Mismatches",[{"metricName":"# mismatches","quality":"Less is better","values":[6998,8127,13455,14839,9659,4386],"isMain":false},{"metricName":"# indels","quality":"Less is better","values":[30,34,55,54,64,14],"isMain":false},{"metricName":"Indels length","quality":"Less is better","values":[73,55,122,90,113,15],"isMain":false},{"metricName":"# mismatches per 100 kbp","quality":"Less is better","values":["1344.61","1397.96","1914.43","1989.15","1385.60","1326.71"],"isMain":true},{"metricName":"# indels per 100 kbp","quality":"Less is better","values":["5.76","5.85","7.83","7.24","9.18","4.23"],"isMain":true},{"metricName":" # indels (<= 5 bp)","quality":"Less is better","values":[26,31,49,52,60,14],"isMain":false},{"metricName":" # indels (> 5 bp)","quality":"Less is better","values":[4,3,6,2,4,0],"isMain":false},{"metricName":"# N's","quality":"Less is better","values":[0,0,0,0,0,0],"isMain":false},{"metricName":"# N's per 100 kbp","quality":"Less is better","values":["0.00","0.00","0.00","0.00","0.00","0.00"],"isMain":true}]],["Statistics without reference",[{"metricName":"# contigs","quality":"Equal","values":[632,648,883,923,772,422],"isMain":true},{"metricName":"# contigs (>= 1000 bp)","quality":"Equal","values":[128,178,175,171,205,68],"isMain":false},{"metricName":"# contigs (>= 5000 bp)","quality":"Equal","values":[1,3,1,3,4,0],"isMain":false},{"metricName":"# contigs (>= 10000 bp)","quality":"Equal","values":[0,0,0,2,0,0],"isMain":false},{"metricName":"# contigs (>= 25000 bp)","quality":"Equal","values":[0,0,0,1,0,0],"isMain":false},{"metricName":"# contigs (>= 50000 bp)","quality":"Equal","values":[0,0,0,0,0,0],"isMain":false},{"metricName":"Largest contig","quality":"More is better","values":[6273,7860,5071,47753,9806,3240],"isMain":true},{"metricName":"Total length","quality":"More is better","values":[535559,600316,724484,834561,719492,337351],"isMain":true},{"metricName":"Total length (>= 1000 bp)","quality":"More is better","values":[198014,278852,251357,326978,328626,98876],"isMain":true},{"metricName":"Total length (>= 5000 bp)","quality":"More is better","values":[6273,20388,5071,67335,31310,0],"isMain":false},{"metricName":"Total length (>= 10000 bp)","quality":"More is better","values":[0,0,0,59776,0,0],"isMain":true},{"metricName":"Total length (>= 25000 bp)","quality":"More is better","values":[0,0,0,47753,0,0],"isMain":false},{"metricName":"Total length (>= 50000 bp)","quality":"More is better","values":[0,0,0,0,0,0],"isMain":true},{"metricName":"N50","quality":"More is better","values":[824,940,812,857,940,798],"isMain":false},{"metricName":"N75","quality":"More is better","values":[628,680,623,640,685,605],"isMain":false},{"metricName":"L50","quality":"Less is better","values":[206,201,299,269,238,146],"isMain":false},{"metricName":"L75","quality":"Less is better","values":[393,391,556,553,463,269],"isMain":false},{"metricName":"GC (%)","quality":"Equal","values":["41.60","41.27","41.86","42.05","41.79","41.77"],"isMain":false}]],["K-mer-based statistics",[]],["Predicted genes",[]],["Similarity statistics",[]],["Reference statistics",[{"metricName":"Reference length","quality":"Equal","values":[5997311,5997311,5997311,5997311,5997311,5997311],"isMain":false},{"metricName":"Reference fragments","quality":"Equal","values":[110,110,110,110,110,110],"isMain":false},{"metricName":"Reference GC (%)","quality":"Equal","values":["42.20","42.20","42.20","42.20","42.20","42.20"],"isMain":false}]]],"subreferences":[],"subreports":[],"minContig":500},{"date":"10 February 2020, Monday, 19:42:47","assembliesNames":["megahit-103-df","megahit-113-df","megahit-113-ml","megahit-113-ms","megahit-129-df","metaSPAdes"],"referenceName":"GCF\_000273135.1\_Bact\_frag\_CL05T12C13\_V1","order":[0,1,2,3,4,5],"report":[["Genome statistics",[{"metricName":"Genome fraction (%)","quality":"More is better","values":["2.221","1.694","3.494","1.998","1.799","0.552"],"isMain":true},{"metricName":"Duplication ratio","quality":"Less is better","values":["1.046","1.065","1.019","1.032","1.033","1.026"],"isMain":true},{"metricName":"Largest alignment","quality":"More is better","values":[4768,6306,47644,4756,2935,1253],"isMain":true},{"metricName":"Total aligned length","quality":"More is better","values":[117521,90069,185080,105832,95559,29308],"isMain":true},{"metricName":"NA50","quality":"More is better","values":[670,726,773,663,699,601],"isMain":false},{"metricName":"NA75","quality":"More is better","values":[545,532,null,540,553,522],"isMain":false},{"metricName":"NGA50","quality":"More is better","values":[null,null,null,null,null,null],"isMain":true},{"metricName":"LA50","quality":"Less is better","values":[66,42,35,62,48,20],"isMain":false},{"metricName":"LA75","quality":"Less is better","values":[119,84,null,110,90,34],"isMain":false}]],["Reads mapping",[]],["Misassemblies",[{"metricName":"# misassemblies","quality":"Less is better","values":[1,1,4,0,0,0],"isMain":true},{"metricName":" # relocations","quality":"Less is better","values":[0,0,1,0,0,0],"isMain":false},{"metricName":" # translocations","quality":"Less is better","values":[0,0,3,0,0,0],"isMain":false},{"metricName":" # inversions","quality":"Less is better","values":[1,1,0,0,0,0],"isMain":false},{"metricName":" # interspecies translocations","quality":"Less is better","values":[19,17,13,8,11,0],"isMain":false},{"metricName":"# misassembled contigs","quality":"Less is better","values":[1,1,2,0,0,0],"isMain":false},{"metricName":"Misassembled contigs length","quality":"Less is better","values":[515,535,89175,0,0,0],"isMain":true},{"metricName":"# possibly misassembled contigs","quality":"Less is better","values":[0,0,0,0,0,0],"isMain":false},{"metricName":" # possible misassemblies","quality":"Less is better","values":[0,0,0,0,0,0],"isMain":false},{"metricName":"# local misassemblies","quality":"Less is better","values":[0,0,2,0,0,0],"isMain":false},{"metricName":"# scaffold gap ext. mis.","quality":"Less is better","values":[0,0,0,0,0,0],"isMain":false},{"metricName":"# scaffold gap loc. mis.","quality":"Less is better","values":[0,0,0,0,0,0],"isMain":false},{"metricName":"# unaligned mis. contigs","quality":"Less is better","values":[0,0,2,0,1,0],"isMain":false}]],["Unaligned",[{"metricName":"# fully unaligned contigs","quality":"Less is better","values":[0,1,0,1,2,0],"isMain":false},{"metricName":"Fully unaligned length","quality":"Less is better","values":[0,598,0,644,1642,0],"isMain":false},{"metricName":"# partially unaligned contigs","quality":"Less is better","values":[7,5,5,2,5,0],"isMain":false},{"metricName":"Partially unaligned length","quality":"Less is better","values":[6032,5757,74737,8524,4531,0],"isMain":false}]],["Mismatches",[{"metricName":"# mismatches","quality":"Less is better","values":[1261,1449,2590,2235,1506,568],"isMain":false},{"metricName":"# indels","quality":"Less is better","values":[11,8,25,12,11,1],"isMain":false},{"metricName":"Indels length","quality":"Less is better","values":[12,10,93,22,12,1],"isMain":false},{"metricName":"# mismatches per 100 kbp","quality":"Less is better","values":["1076.94","1622.66","1405.77","2121.74","1588.00","1950.28"],"isMain":true},{"metricName":"# indels per 100 kbp","quality":"Less is better","values":["9.39","8.96","13.57","11.39","11.60","3.43"],"isMain":true},{"metricName":" # indels (<= 5 bp)","quality":"Less is better","values":[11,8,23,10,11,1],"isMain":false},{"metricName":" # indels (> 5 bp)","quality":"Less is better","values":[0,0,2,2,0,0],"isMain":false},{"metricName":"# N's","quality":"Less is better","values":[0,0,0,0,0,0],"isMain":false},{"metricName":"# N's per 100 kbp","quality":"Less is better","values":["0.00","0.00","0.00","0.00","0.00","0.00"],"isMain":true}]],["Statistics without reference",[{"metricName":"# contigs","quality":"Equal","values":[171,119,124,148,129,47],"isMain":true},{"metricName":"# contigs (>= 1000 bp)","quality":"Equal","values":[17,25,22,18,24,2],"isMain":false},{"metricName":"# contigs (>= 5000 bp)","quality":"Equal","values":[0,2,2,1,0,0],"isMain":false},{"metricName":"# contigs (>= 10000 bp)","quality":"Equal","values":[0,0,2,1,0,0],"isMain":false},{"metricName":"# contigs (>= 25000 bp)","quality":"Equal","values":[0,0,2,0,0,0],"isMain":false},{"metricName":"# contigs (>= 50000 bp)","quality":"Equal","values":[0,0,2,0,0,0],"isMain":false},{"metricName":"Largest contig","quality":"More is better","values":[4768,6334,88346,11560,2935,1293],"isMain":true},{"metricName":"Total length","quality":"More is better","values":[128553,101502,262552,117826,104158,29894],"isMain":true},{"metricName":"Total length (>= 1000 bp)","quality":"More is better","values":[28236,40267,197056,34620,35173,2352],"isMain":true},{"metricName":"Total length (>= 5000 bp)","quality":"More is better","values":[0,11587,165410,11560,0,0],"isMain":false},{"metricName":"Total length (>= 10000 bp)","quality":"More is better","values":[0,0,165410,11560,0,0],"isMain":true},{"metricName":"Total length (>= 25000 bp)","quality":"More is better","values":[0,0,165410,0,0,0],"isMain":false},{"metricName":"Total length (>= 50000 bp)","quality":"More is better","values":[0,0,165410,0,0,0],"isMain":true},{"metricName":"N50","quality":"More is better","values":[705,852,77064,705,796,606],"isMain":false},{"metricName":"N75","quality":"More is better","values":[592,594,1086,596,595,540],"isMain":false},{"metricName":"L50","quality":"Less is better","values":[62,37,2,49,44,20],"isMain":false},{"metricName":"L75","quality":"Less is better","values":[112,73,22,94,82,33],"isMain":false},{"metricName":"GC (%)","quality":"Equal","values":["42.55","41.02","43.39","42.94","43.03","41.47"],"isMain":false}]],["K-mer-based statistics",[]],["Predicted genes",[]],["Similarity statistics",[]],["Reference statistics",[{"metricName":"Reference length","quality":"Equal","values":[5272458,5272458,5272458,5272458,5272458,5272458],"isMain":false},{"metricName":"Reference fragments","quality":"Equal","values":[59,59,59,59,59,59],"isMain":false},{"metricName":"Reference GC (%)","quality":"Equal","values":["43.45","43.45","43.45","43.45","43.45","43.45"],"isMain":false}]]],"subreferences":[],"subreports":[],"minContig":500},{"date":"10 February 2020, Monday, 19:42:43","assembliesNames":["megahit-103-df","megahit-113-df","megahit-113-ml","megahit-113-ms","megahit-129-df","metaSPAdes"],"referenceName":"GCF\_000273155.1\_Bact\_frag\_CL07T12C05\_V1","order":[0,1,2,3,4,5],"report":[["Genome statistics",[{"metricName":"Genome fraction (%)","quality":"More is better","values":["2.755","3.233","1.536","2.786","1.194","0.511"],"isMain":true},{"metricName":"Duplication ratio","quality":"Less is better","values":["1.031","1.015","1.034","1.011","1.028","1.003"],"isMain":true},{"metricName":"Largest alignment","quality":"More is better","values":[8585,53993,41686,90743,11773,8424],"isMain":true},{"metricName":"Total aligned length","quality":"More is better","values":[150238,176495,84032,152019,65206,27782],"isMain":true},{"metricName":"NA50","quality":"More is better","values":[930,26887,3051,90743,1726,3158],"isMain":false},{"metricName":"NA75","quality":"More is better","values":[593,1567,793,1501,676,837],"isMain":false},{"metricName":"NGA50","quality":"More is better","values":[null,null,null,null,null,null],"isMain":true},{"metricName":"LA50","quality":"Less is better","values":[48,3,2,1,8,3],"isMain":false},{"metricName":"LA75","quality":"Less is better","values":[105,8,20,11,24,8],"isMain":false}]],["Reads mapping",[]],["Misassemblies",[{"metricName":"# misassemblies","quality":"Less is better","values":[3,3,3,2,1,0],"isMain":true},{"metricName":" # relocations","quality":"Less is better","values":[0,0,0,2,0,0],"isMain":false},{"metricName":" # translocations","quality":"Less is better","values":[2,2,3,0,1,0],"isMain":false},{"metricName":" # inversions","quality":"Less is better","values":[1,1,0,0,0,0],"isMain":false},{"metricName":" # interspecies translocations","quality":"Less is better","values":[19,6,7,9,2,1],"isMain":false},{"metricName":"# misassembled contigs","quality":"Less is better","values":[2,2,2,2,1,0],"isMain":false},{"metricName":"Misassembled contigs length","quality":"Less is better","values":[2603,2382,2596,13901,528,0],"isMain":true},{"metricName":"# possibly misassembled contigs","quality":"Less is better","values":[0,0,0,0,0,0],"isMain":false},{"metricName":" # possible misassemblies","quality":"Less is better","values":[0,0,0,0,0,0],"isMain":false},{"metricName":"# local misassemblies","quality":"Less is better","values":[0,1,0,1,0,0],"isMain":false},{"metricName":"# scaffold gap ext. mis.","quality":"Less is better","values":[0,0,0,0,0,0],"isMain":false},{"metricName":"# scaffold gap loc. mis.","quality":"Less is better","values":[0,0,0,0,0,0],"isMain":false},{"metricName":"# unaligned mis. contigs","quality":"Less is better","values":[0,1,0,2,0,0],"isMain":false}]],["Unaligned",[{"metricName":"# fully unaligned contigs","quality":"Less is better","values":[1,0,0,2,0,0],"isMain":false},{"metricName":"Fully unaligned length","quality":"Less is better","values":[1278,0,0,1313,0,0],"isMain":false},{"metricName":"# partially unaligned contigs","quality":"Less is better","values":[8,3,4,6,0,1],"isMain":false},{"metricName":"Partially unaligned length","quality":"Less is better","values":[7031,14459,2946,12719,0,821],"isMain":false}]],["Mismatches",[{"metricName":"# mismatches","quality":"Less is better","values":[510,1011,827,938,799,134],"isMain":false},{"metricName":"# indels","quality":"Less is better","values":[5,15,10,13,11,2],"isMain":false},{"metricName":"Indels length","quality":"Less is better","values":[24,39,22,43,16,16],"isMain":false},{"metricName":"# mismatches per 100 kbp","quality":"Less is better","values":["340.94","575.93","991.46","620.07","1232.64","483.32"],"isMain":true},{"metricName":"# indels per 100 kbp","quality":"Less is better","values":["3.34","8.55","11.99","8.59","16.97","7.21"],"isMain":true},{"metricName":" # indels (<= 5 bp)","quality":"Less is better","values":[3,13,9,11,11,1],"isMain":false},{"metricName":" # indels (> 5 bp)","quality":"Less is better","values":[2,2,1,2,0,1],"isMain":false},{"metricName":"# N's","quality":"Less is better","values":[0,0,0,0,0,0],"isMain":false},{"metricName":"# N's per 100 kbp","quality":"Less is better","values":["0.00","0.00","0.00","0.00","0.00","0.00"],"isMain":true}]],["Statistics without reference",[{"metricName":"# contigs","quality":"Equal","values":[167,51,52,51,51,18],"isMain":true},{"metricName":"# contigs (>= 1000 bp)","quality":"Equal","values":[46,15,18,16,14,7],"isMain":false},{"metricName":"# contigs (>= 5000 bp)","quality":"Equal","values":[2,5,1,3,3,1],"isMain":false},{"metricName":"# contigs (>= 10000 bp)","quality":"Equal","values":[0,5,1,2,1,0],"isMain":false},{"metricName":"# contigs (>= 25000 bp)","quality":"Equal","values":[0,4,1,1,0,0],"isMain":false},{"metricName":"# contigs (>= 50000 bp)","quality":"Equal","values":[0,1,0,1,0,0],"isMain":false},{"metricName":"Largest contig","quality":"More is better","values":[8585,53993,41686,90743,11934,8425],"isMain":true},{"metricName":"Total length","quality":"More is better","values":[162604,192650,89170,166978,66637,28635],"isMain":true},{"metricName":"Total length (>= 1000 bp)","quality":"More is better","values":[82261,167446,67190,141601,42486,20965],"isMain":true},{"metricName":"Total length (>= 5000 bp)","quality":"More is better","values":[14011,151922,41686,112643,22709,8425],"isMain":false},{"metricName":"Total length (>= 10000 bp)","quality":"More is better","values":[0,151922,41686,103887,11934,0],"isMain":true},{"metricName":"Total length (>= 25000 bp)","quality":"More is better","values":[0,140788,41686,90743,0,0],"isMain":false},{"metricName":"Total length (>= 50000 bp)","quality":"More is better","values":[0,53993,0,90743,0,0],"isMain":true},{"metricName":"N50","quality":"More is better","values":[1000,26887,3051,90743,1726,3158],"isMain":false},{"metricName":"N75","quality":"More is better","values":[647,11134,1021,2637,704,976],"isMain":false},{"metricName":"L50","quality":"Less is better","values":[46,3,2,1,8,3],"isMain":false},{"metricName":"L75","quality":"Less is better","values":[96,5,18,8,23,8],"isMain":false},{"metricName":"GC (%)","quality":"Equal","values":["42.26","44.09","45.21","43.99","41.44","41.07"],"isMain":false}]],["K-mer-based statistics",[]],["Predicted genes",[]],["Similarity statistics",[]],["Reference statistics",[{"metricName":"Reference length","quality":"Equal","values":[5430227,5430227,5430227,5430227,5430227,5430227],"isMain":false},{"metricName":"Reference fragments","quality":"Equal","values":[35,35,35,35,35,35],"isMain":false},{"metricName":"Reference GC (%)","quality":"Equal","values":["43.62","43.62","43.62","43.62","43.62","43.62"],"isMain":false}]]],"subreferences":[],"subreports":[],"minContig":500},{"date":"10 February 2020, Monday, 19:42:41","assembliesNames":["megahit-103-df","megahit-113-df","megahit-113-ml","megahit-113-ms","megahit-129-df","metaSPAdes"],"referenceName":"GCF\_000273175.1\_Bact\_nord\_CL02T12C05\_V1","order":[0,1,2,3,4,5],"report":[["Genome statistics",[{"metricName":"Genome fraction (%)","quality":"More is better","values":["4.461","4.885","5.317","5.923","5.740","2.213"],"isMain":true},{"metricName":"Duplication ratio","quality":"Less is better","values":["1.052","1.044","1.029","1.039","1.043","1.053"],"isMain":true},{"metricName":"Largest alignment","quality":"More is better","values":[10662,1948,2960,8336,13964,2611],"isMain":true},{"metricName":"Total aligned length","quality":"More is better","values":[255725,279340,304846,339552,328746,126534],"isMain":true},{"metricName":"NA50","quality":"More is better","values":[621,593,592,587,598,555],"isMain":false},{"metricName":"NA75","quality":"More is better","values":[521,510,528,520,504,505],"isMain":false},{"metricName":"NGA50","quality":"More is better","values":[null,null,null,null,null,null],"isMain":true},{"metricName":"LA50","quality":"Less is better","values":[140,208,213,218,223,103],"isMain":false},{"metricName":"LA75","quality":"Less is better","values":[266,356,356,381,403,171],"isMain":false}]],["Reads mapping",[]],["Misassemblies",[{"metricName":"# misassemblies","quality":"Less is better","values":[3,6,3,5,7,2],"isMain":true},{"metricName":" # relocations","quality":"Less is better","values":[0,1,0,1,3,0],"isMain":false},{"metricName":" # translocations","quality":"Less is better","values":[3,5,3,3,4,2],"isMain":false},{"metricName":" # inversions","quality":"Less is better","values":[0,0,0,1,0,0],"isMain":false},{"metricName":" # interspecies translocations","quality":"Less is better","values":[49,65,23,37,63,28],"isMain":false},{"metricName":"# misassembled contigs","quality":"Less is better","values":[3,5,3,5,7,2],"isMain":false},{"metricName":"Misassembled contigs length","quality":"Less is better","values":[4116,8126,2357,3431,6398,1210],"isMain":true},{"metricName":"# possibly misassembled contigs","quality":"Less is better","values":[0,0,0,0,0,0],"isMain":false},{"metricName":" # possible misassemblies","quality":"Less is better","values":[0,0,0,0,0,0],"isMain":false},{"metricName":"# local misassemblies","quality":"Less is better","values":[0,1,3,1,1,0],"isMain":false},{"metricName":"# scaffold gap ext. mis.","quality":"Less is better","values":[0,0,0,0,0,0],"isMain":false},{"metricName":"# scaffold gap loc. mis.","quality":"Less is better","values":[0,0,0,0,0,0],"isMain":false},{"metricName":"# unaligned mis. contigs","quality":"Less is better","values":[0,0,0,0,0,0],"isMain":false}]],["Unaligned",[{"metricName":"# fully unaligned contigs","quality":"Less is better","values":[2,0,4,0,2,0],"isMain":false},{"metricName":"Fully unaligned length","quality":"Less is better","values":[2021,0,3803,0,1853,0],"isMain":false},{"metricName":"# partially unaligned contigs","quality":"Less is better","values":[8,26,4,11,26,11],"isMain":false},{"metricName":"Partially unaligned length","quality":"Less is better","values":[12969,33383,3288,8255,48342,10969],"isMain":false}]],["Mismatches",[{"metricName":"# mismatches","quality":"Less is better","values":[3906,4958,6458,7311,5560,2345],"isMain":false},{"metricName":"# indels","quality":"Less is better","values":[15,20,26,30,15,10],"isMain":false},{"metricName":"Indels length","quality":"Less is better","values":[23,34,33,57,27,24],"isMain":false},{"metricName":"# mismatches per 100 kbp","quality":"Less is better","values":["1534.25","1778.37","2128.14","2162.68","1697.00","1856.73"],"isMain":true},{"metricName":"# indels per 100 kbp","quality":"Less is better","values":["5.89","7.17","8.57","8.87","4.58","7.92"],"isMain":true},{"metricName":" # indels (<= 5 bp)","quality":"Less is better","values":[14,18,25,27,14,9],"isMain":false},{"metricName":" # indels (> 5 bp)","quality":"Less is better","values":[1,2,1,3,1,1],"isMain":false},{"metricName":"# N's","quality":"Less is better","values":[0,0,0,0,0,0],"isMain":false},{"metricName":"# N's per 100 kbp","quality":"Less is better","values":["0.00","0.00","0.00","0.00","0.00","0.00"],"isMain":true}]],["Statistics without reference",[{"metricName":"# contigs","quality":"Equal","values":[367,443,494,532,493,220],"isMain":true},{"metricName":"# contigs (>= 1000 bp)","quality":"Equal","values":[27,45,23,26,44,11],"isMain":false},{"metricName":"# contigs (>= 5000 bp)","quality":"Equal","values":[4,0,0,2,4,0],"isMain":false},{"metricName":"# contigs (>= 10000 bp)","quality":"Equal","values":[1,0,0,0,1,0],"isMain":false},{"metricName":"# contigs (>= 25000 bp)","quality":"Equal","values":[0,0,0,0,0,0],"isMain":false},{"metricName":"# contigs (>= 50000 bp)","quality":"Equal","values":[0,0,0,0,0,0],"isMain":false},{"metricName":"Largest contig","quality":"More is better","values":[10678,3420,2971,8336,16206,2691],"isMain":true},{"metricName":"Total length","quality":"More is better","values":[282745,324361,319212,359654,391879,144010],"isMain":true},{"metricName":"Total length (>= 1000 bp)","quality":"More is better","values":[68937,72604,29910,50610,109498,17126],"isMain":true},{"metricName":"Total length (>= 5000 bp)","quality":"More is better","values":[31450,0,0,14206,39096,0],"isMain":false},{"metricName":"Total length (>= 10000 bp)","quality":"More is better","values":[10678,0,0,0,16206,0],"isMain":true},{"metricName":"Total length (>= 25000 bp)","quality":"More is better","values":[0,0,0,0,0,0],"isMain":false},{"metricName":"Total length (>= 50000 bp)","quality":"More is better","values":[0,0,0,0,0,0],"isMain":true},{"metricName":"N50","quality":"More is better","values":[680,678,606,616,680,607],"isMain":false},{"metricName":"N75","quality":"More is better","values":[568,572,544,544,574,541],"isMain":false},{"metricName":"L50","quality":"Less is better","values":[119,160,203,204,153,88],"isMain":false},{"metricName":"L75","quality":"Less is better","values":[233,291,341,360,309,151],"isMain":false},{"metricName":"GC (%)","quality":"Equal","values":["42.26","41.13","41.45","41.23","42.44","41.27"],"isMain":false}]],["K-mer-based statistics",[]],["Predicted genes",[]],["Similarity statistics",[]],["Reference statistics",[{"metricName":"Reference length","quality":"Equal","values":[5707588,5707588,5707588,5707588,5707588,5707588],"isMain":false},{"metricName":"Reference fragments","quality":"Equal","values":[36,36,36,36,36,36],"isMain":false},{"metricName":"Reference GC (%)","quality":"Equal","values":["40.83","40.83","40.83","40.83","40.83","40.83"],"isMain":false}]]],"subreferences":[],"subreports":[],"minContig":500},{"date":"10 February 2020, Monday, 19:42:50","assembliesNames":["megahit-103-df","megahit-113-df","megahit-113-ml","megahit-113-ms","megahit-129-df","metaSPAdes"],"referenceName":"GCF\_000273195.1\_Bact\_ovat\_CL02T12C04\_V1","order":[0,1,2,3,4,5],"report":[["Genome statistics",[{"metricName":"Genome fraction (%)","quality":"More is better","values":["4.055","3.181","2.791","3.177","2.839","1.083"],"isMain":true},{"metricName":"Duplication ratio","quality":"Less is better","values":["1.007","1.013","1.019","1.021","1.025","1.026"],"isMain":true},{"metricName":"Largest alignment","quality":"More is better","values":[37176,19510,18342,44539,23733,12993],"isMain":true},{"metricName":"Total aligned length","quality":"More is better","values":[320040,251725,221133,252451,224949,87038],"isMain":true},{"metricName":"NA50","quality":"More is better","values":[8244,6135,6480,7983,1753,1989],"isMain":false},{"metricName":"NA75","quality":"More is better","values":[2175,1890,1356,1397,null,714],"isMain":false},{"metricName":"NGA50","quality":"More is better","values":[null,null,null,null,null,null],"isMain":true},{"metricName":"LA50","quality":"Less is better","values":[10,12,11,7,26,13],"isMain":false},{"metricName":"LA75","quality":"Less is better","values":[28,29,26,26,null,34],"isMain":false}]],["Reads mapping",[]],["Misassemblies",[{"metricName":"# misassemblies","quality":"Less is better","values":[1,0,2,1,2,0],"isMain":true},{"metricName":" # relocations","quality":"Less is better","values":[0,0,1,1,0,0],"isMain":false},{"metricName":" # translocations","quality":"Less is better","values":[1,0,0,0,2,0],"isMain":false},{"metricName":" # inversions","quality":"Less is better","values":[0,0,1,0,0,0],"isMain":false},{"metricName":" # interspecies translocations","quality":"Less is better","values":[9,9,8,10,19,4],"isMain":false},{"metricName":"# misassembled contigs","quality":"Less is better","values":[1,0,2,1,2,0],"isMain":false},{"metricName":"Misassembled contigs length","quality":"Less is better","values":[68794,0,25532,45382,43658,0],"isMain":true},{"metricName":"# possibly misassembled contigs","quality":"Less is better","values":[0,0,0,0,0,0],"isMain":false},{"metricName":" # possible misassemblies","quality":"Less is better","values":[0,0,0,0,0,0],"isMain":false},{"metricName":"# local misassemblies","quality":"Less is better","values":[1,0,0,1,0,0],"isMain":false},{"metricName":"# scaffold gap ext. mis.","quality":"Less is better","values":[0,0,0,0,0,0],"isMain":false},{"metricName":"# scaffold gap loc. mis.","quality":"Less is better","values":[0,0,0,0,0,0],"isMain":false},{"metricName":"# unaligned mis. contigs","quality":"Less is better","values":[0,0,0,0,1,0],"isMain":false}]],["Unaligned",[{"metricName":"# fully unaligned contigs","quality":"Less is better","values":[2,0,0,1,0,0],"isMain":false},{"metricName":"Fully unaligned length","quality":"Less is better","values":[1076,0,0,597,0,0],"isMain":false},{"metricName":"# partially unaligned contigs","quality":"Less is better","values":[3,2,4,4,7,2],"isMain":false},{"metricName":"Partially unaligned length","quality":"Less is better","values":[2796,1629,3956,3972,90491,7341],"isMain":false}]],["Mismatches",[{"metricName":"# mismatches","quality":"Less is better","values":[1450,1419,1639,1965,1817,569],"isMain":false},{"metricName":"# indels","quality":"Less is better","values":[16,13,8,12,23,1],"isMain":false},{"metricName":"Indels length","quality":"Less is better","values":[31,16,12,13,32,1],"isMain":false},{"metricName":"# mismatches per 100 kbp","quality":"Less is better","values":["453.78","566.10","745.18","784.75","812.04","666.75"],"isMain":true},{"metricName":"# indels per 100 kbp","quality":"Less is better","values":["5.01","5.19","3.64","4.79","10.28","1.17"],"isMain":true},{"metricName":" # indels (<= 5 bp)","quality":"Less is better","values":[15,13,8,12,23,1],"isMain":false},{"metricName":" # indels (> 5 bp)","quality":"Less is better","values":[1,0,0,0,0,0],"isMain":false},{"metricName":"# N's","quality":"Less is better","values":[0,0,0,0,0,0],"isMain":false},{"metricName":"# N's per 100 kbp","quality":"Less is better","values":["0.00","0.00","0.00","0.00","0.00","0.00"],"isMain":true}]],["Statistics without reference",[{"metricName":"# contigs","quality":"Equal","values":[122,100,102,114,118,61],"isMain":true},{"metricName":"# contigs (>= 1000 bp)","quality":"Equal","values":[47,50,31,34,43,25],"isMain":false},{"metricName":"# contigs (>= 5000 bp)","quality":"Equal","values":[15,15,14,9,12,3],"isMain":false},{"metricName":"# contigs (>= 10000 bp)","quality":"Equal","values":[6,6,6,5,5,1],"isMain":false},{"metricName":"# contigs (>= 25000 bp)","quality":"Equal","values":[1,0,0,2,2,0],"isMain":false},{"metricName":"# contigs (>= 50000 bp)","quality":"Equal","values":[1,0,0,0,1,0],"isMain":false},{"metricName":"Largest contig","quality":"More is better","values":[68794,19510,24946,45382,75678,12993],"isMain":true},{"metricName":"Total length","quality":"More is better","values":[325582,255577,228059,260216,319756,94872],"isMain":true},{"metricName":"Total length (>= 1000 bp)","quality":"More is better","values":[276658,221482,181390,206735,271270,71309],"isMain":true},{"metricName":"Total length (>= 5000 bp)","quality":"More is better","values":[209267,150166,141799,151966,210087,26122],"isMain":false},{"metricName":"Total length (>= 10000 bp)","quality":"More is better","values":[144300,87708,91597,120413,163782,12993],"isMain":true},{"metricName":"Total length (>= 25000 bp)","quality":"More is better","values":[68794,0,0,70861,118615,0],"isMain":false},{"metricName":"Total length (>= 50000 bp)","quality":"More is better","values":[68794,0,0,0,75678,0],"isMain":true},{"metricName":"N50","quality":"More is better","values":[8244,6143,6480,8011,12671,2348],"isMain":false},{"metricName":"N75","quality":"More is better","values":[2502,1905,1734,1528,2699,1021],"isMain":false},{"metricName":"L50","quality":"Less is better","values":[9,12,10,7,5,10],"isMain":false},{"metricName":"L75","quality":"Less is better","values":[26,29,23,25,21,25],"isMain":false},{"metricName":"GC (%)","quality":"Equal","values":["42.68","42.63","42.97","43.08","42.73","43.03"],"isMain":false}]],["K-mer-based statistics",[]],["Predicted genes",[]],["Similarity statistics",[]],["Reference statistics",[{"metricName":"Reference length","quality":"Equal","values":[7880760,7880760,7880760,7880760,7880760,7880760],"isMain":false},{"metricName":"Reference fragments","quality":"Equal","values":[47,47,47,47,47,47],"isMain":false},{"metricName":"Reference GC (%)","quality":"Equal","values":["42.28","42.28","42.28","42.28","42.28","42.28"],"isMain":false}]]],"subreferences":[],"subreports":[],"minContig":500},{"date":"10 February 2020, Monday, 19:42:53","assembliesNames":["megahit-103-df","megahit-113-df","megahit-113-ml","megahit-113-ms","megahit-129-df","metaSPAdes"],"referenceName":"GCF\_000273235.1\_Bact\_saly\_CL02T12C01\_V1","order":[0,1,2,3,4,5],"report":[["Genome statistics",[{"metricName":"Genome fraction (%)","quality":"More is better","values":["5.933","5.130","7.553","8.555","4.696","5.403"],"isMain":true},{"metricName":"Duplication ratio","quality":"Less is better","values":["1.034","1.030","1.034","1.041","1.037","1.027"],"isMain":true},{"metricName":"Largest alignment","quality":"More is better","values":[7619,4145,28507,28593,3577,8237],"isMain":true},{"metricName":"Total aligned length","quality":"More is better","values":[347448,300011,444169,502873,275496,315320],"isMain":true},{"metricName":"NA50","quality":"More is better","values":[866,935,802,813,891,750],"isMain":false},{"metricName":"NA75","quality":"More is better","values":[635,645,589,611,611,581],"isMain":false},{"metricName":"NGA50","quality":"More is better","values":[null,null,null,null,null,null],"isMain":true},{"metricName":"LA50","quality":"Less is better","values":[136,108,185,204,106,161],"isMain":false},{"metricName":"LA75","quality":"Less is better","values":[255,209,358,392,211,293],"isMain":false}]],["Reads mapping",[]],["Misassemblies",[{"metricName":"# misassemblies","quality":"Less is better","values":[1,4,3,5,1,2],"isMain":true},{"metricName":" # relocations","quality":"Less is better","values":[0,0,0,0,0,1],"isMain":false},{"metricName":" # translocations","quality":"Less is better","values":[0,3,2,5,1,1],"isMain":false},{"metricName":" # inversions","quality":"Less is better","values":[1,1,1,0,0,0],"isMain":false},{"metricName":" # interspecies translocations","quality":"Less is better","values":[21,17,17,30,17,13],"isMain":false},{"metricName":"# misassembled contigs","quality":"Less is better","values":[1,3,3,3,1,2],"isMain":false},{"metricName":"Misassembled contigs length","quality":"Less is better","values":[777,7985,2361,7125,1621,13898],"isMain":true},{"metricName":"# possibly misassembled contigs","quality":"Less is better","values":[0,0,0,0,0,0],"isMain":false},{"metricName":" # possible misassemblies","quality":"Less is better","values":[0,0,0,0,0,0],"isMain":false},{"metricName":"# local misassemblies","quality":"Less is better","values":[0,0,0,0,0,1],"isMain":false},{"metricName":"# scaffold gap ext. mis.","quality":"Less is better","values":[0,0,0,0,0,0],"isMain":false},{"metricName":"# scaffold gap loc. mis.","quality":"Less is better","values":[0,0,0,0,0,0],"isMain":false},{"metricName":"# unaligned mis. contigs","quality":"Less is better","values":[0,0,0,0,0,0],"isMain":false}]],["Unaligned",[{"metricName":"# fully unaligned contigs","quality":"Less is better","values":[2,1,1,0,1,0],"isMain":false},{"metricName":"Fully unaligned length","quality":"Less is better","values":[1253,617,523,0,634,0],"isMain":false},{"metricName":"# partially unaligned contigs","quality":"Less is better","values":[3,3,1,8,6,4],"isMain":false},{"metricName":"Partially unaligned length","quality":"Less is better","values":[1926,8661,20802,7612,25960,27996],"isMain":false}]],["Mismatches",[{"metricName":"# mismatches","quality":"Less is better","values":[3935,3542,7296,8507,3487,3733],"isMain":false},{"metricName":"# indels","quality":"Less is better","values":[10,18,32,34,23,12],"isMain":false},{"metricName":"Indels length","quality":"Less is better","values":[78,80,73,156,104,12],"isMain":false},{"metricName":"# mismatches per 100 kbp","quality":"Less is better","values":["1147.20","1194.27","1670.80","1719.80","1284.32","1194.88"],"isMain":true},{"metricName":"# indels per 100 kbp","quality":"Less is better","values":["2.92","6.07","7.33","6.87","8.47","3.84"],"isMain":true},{"metricName":" # indels (<= 5 bp)","quality":"Less is better","values":[6,14,28,26,18,12],"isMain":false},{"metricName":" # indels (> 5 bp)","quality":"Less is better","values":[4,4,4,8,5,0],"isMain":false},{"metricName":"# N's","quality":"Less is better","values":[0,0,0,0,0,0],"isMain":false},{"metricName":"# N's per 100 kbp","quality":"Less is better","values":["0.00","0.00","0.00","0.00","0.00","0.00"],"isMain":true}]],["Statistics without reference",[{"metricName":"# contigs","quality":"Equal","values":[403,325,527,604,298,393],"isMain":true},{"metricName":"# contigs (>= 1000 bp)","quality":"Equal","values":[100,102,97,109,98,72],"isMain":false},{"metricName":"# contigs (>= 5000 bp)","quality":"Equal","values":[1,1,2,1,1,2],"isMain":false},{"metricName":"# contigs (>= 10000 bp)","quality":"Equal","values":[0,0,2,1,1,2],"isMain":false},{"metricName":"# contigs (>= 25000 bp)","quality":"Equal","values":[0,0,1,1,0,1],"isMain":false},{"metricName":"# contigs (>= 50000 bp)","quality":"Equal","values":[0,0,0,0,0,0],"isMain":false},{"metricName":"Largest contig","quality":"More is better","values":[7688,7861,28507,28593,18626,25144],"isMain":true},{"metricName":"Total length","quality":"More is better","values":[357994,314855,472717,522530,308046,348885],"isMain":true},{"metricName":"Total length (>= 1000 bp)","quality":"More is better","values":[148847,162368,180521,183673,172887,130209],"isMain":true},{"metricName":"Total length (>= 5000 bp)","quality":"More is better","values":[7688,7861,50487,28593,18626,38089],"isMain":false},{"metricName":"Total length (>= 10000 bp)","quality":"More is better","values":[0,0,50487,28593,18626,38089],"isMain":true},{"metricName":"Total length (>= 25000 bp)","quality":"More is better","values":[0,0,28507,28593,0,25144],"isMain":false},{"metricName":"Total length (>= 50000 bp)","quality":"More is better","values":[0,0,0,0,0,0],"isMain":true},{"metricName":"N50","quality":"More is better","values":[889,1035,858,849,1102,821],"isMain":false},{"metricName":"N75","quality":"More is better","values":[676,700,639,639,714,651],"isMain":false},{"metricName":"L50","quality":"Less is better","values":[133,98,158,194,80,122],"isMain":false},{"metricName":"L75","quality":"Less is better","values":[248,191,317,373,169,240],"isMain":false},{"metricName":"GC (%)","quality":"Equal","values":["42.17","42.09","42.33","42.13","41.93","42.09"],"isMain":false}]],["K-mer-based statistics",[]],["Predicted genes",[]],["Similarity statistics",[]],["Reference statistics",[{"metricName":"Reference length","quality":"Equal","values":[5781840,5781840,5781840,5781840,5781840,5781840],"isMain":false},{"metricName":"Reference fragments","quality":"Equal","values":[56,56,56,56,56,56],"isMain":false},{"metricName":"Reference GC (%)","quality":"Equal","values":["42.03","42.03","42.03","42.03","42.03","42.03"],"isMain":false}]]],"subreferences":[],"subreports":[],"minContig":500},{"date":"10 February 2020, Monday, 19:42:51","assembliesNames":["megahit-103-df","megahit-113-df","megahit-113-ml","megahit-113-ms","megahit-129-df","metaSPAdes"],"referenceName":"GCF\_000273295.1\_Bact\_vulg\_CL09T03C04\_V1","order":[0,1,2,3,4,5],"report":[["Genome statistics",[{"metricName":"Genome fraction (%)","quality":"More is better","values":["1.533","1.399","3.144","2.958","1.578","0.861"],"isMain":true},{"metricName":"Duplication ratio","quality":"Less is better","values":["1.014","1.047","1.022","1.032","1.021","1.016"],"isMain":true},{"metricName":"Largest alignment","quality":"More is better","values":[10159,3837,3887,5660,10728,6801],"isMain":true},{"metricName":"Total aligned length","quality":"More is better","values":[75524,68875,155587,147842,78271,42366],"isMain":true},{"metricName":"NA50","quality":"More is better","values":[1408,1675,1211,1079,2003,1542],"isMain":false},{"metricName":"NA75","quality":"More is better","values":[874,763,779,701,597,850],"isMain":false},{"metricName":"NGA50","quality":"More is better","values":[null,null,null,null,null,null],"isMain":true},{"metricName":"LA50","quality":"Less is better","values":[14,16,42,38,10,7],"isMain":false},{"metricName":"LA75","quality":"Less is better","values":[33,32,84,83,34,16],"isMain":false}]],["Reads mapping",[]],["Misassemblies",[{"metricName":"# misassemblies","quality":"Less is better","values":[0,1,0,1,0,0],"isMain":true},{"metricName":" # relocations","quality":"Less is better","values":[0,0,0,1,0,0],"isMain":false},{"metricName":" # translocations","quality":"Less is better","values":[0,1,0,0,0,0],"isMain":false},{"metricName":" # inversions","quality":"Less is better","values":[0,0,0,0,0,0],"isMain":false},{"metricName":" # interspecies translocations","quality":"Less is better","values":[1,9,5,5,4,1],"isMain":false},{"metricName":"# misassembled contigs","quality":"Less is better","values":[0,1,0,1,0,0],"isMain":false},{"metricName":"Misassembled contigs length","quality":"Less is better","values":[0,755,0,4004,0,0],"isMain":true},{"metricName":"# possibly misassembled contigs","quality":"Less is better","values":[0,0,0,0,0,0],"isMain":false},{"metricName":" # possible misassemblies","quality":"Less is better","values":[0,0,0,0,0,0],"isMain":false},{"metricName":"# local misassemblies","quality":"Less is better","values":[0,1,0,2,0,0],"isMain":false},{"metricName":"# scaffold gap ext. mis.","quality":"Less is better","values":[0,0,0,0,0,0],"isMain":false},{"metricName":"# scaffold gap loc. mis.","quality":"Less is better","values":[0,0,0,0,0,0],"isMain":false},{"metricName":"# unaligned mis. contigs","quality":"Less is better","values":[0,0,0,1,0,0],"isMain":false}]],["Unaligned",[{"metricName":"# fully unaligned contigs","quality":"Less is better","values":[0,0,0,0,0,0],"isMain":false},{"metricName":"Fully unaligned length","quality":"Less is better","values":[0,0,0,0,0,0],"isMain":false},{"metricName":"# partially unaligned contigs","quality":"Less is better","values":[0,2,2,3,2,1],"isMain":false},{"metricName":"Partially unaligned length","quality":"Less is better","values":[0,1866,1389,3497,14765,511],"isMain":false}]],["Mismatches",[{"metricName":"# mismatches","quality":"Less is better","values":[903,1077,2636,2682,973,415],"isMain":false},{"metricName":"# indels","quality":"Less is better","values":[9,17,24,24,22,3],"isMain":false},{"metricName":"Indels length","quality":"Less is better","values":[17,19,29,29,30,4],"isMain":false},{"metricName":"# mismatches per 100 kbp","quality":"Less is better","values":["1202.06","1570.68","1710.90","1850.20","1258.50","983.30"],"isMain":true},{"metricName":"# indels per 100 kbp","quality":"Less is better","values":["11.98","24.79","15.58","16.56","28.46","7.11"],"isMain":true},{"metricName":" # indels (<= 5 bp)","quality":"Less is better","values":[9,17,24,24,22,3],"isMain":false},{"metricName":" # indels (> 5 bp)","quality":"Less is better","values":[0,0,0,0,0,0],"isMain":false},{"metricName":"# N's","quality":"Less is better","values":[0,0,0,0,0,0],"isMain":false},{"metricName":"# N's per 100 kbp","quality":"Less is better","values":["0.00","0.00","0.00","0.00","0.00","0.00"],"isMain":true}]],["Statistics without reference",[{"metricName":"# contigs","quality":"Equal","values":[61,55,146,141,49,31],"isMain":true},{"metricName":"# contigs (>= 1000 bp)","quality":"Equal","values":[23,28,54,47,20,13],"isMain":false},{"metricName":"# contigs (>= 5000 bp)","quality":"Equal","values":[1,1,0,1,4,2],"isMain":false},{"metricName":"# contigs (>= 10000 bp)","quality":"Equal","values":[1,0,0,0,2,0],"isMain":false},{"metricName":"# contigs (>= 25000 bp)","quality":"Equal","values":[0,0,0,0,0,0],"isMain":false},{"metricName":"# contigs (>= 50000 bp)","quality":"Equal","values":[0,0,0,0,0,0],"isMain":false},{"metricName":"Largest contig","quality":"More is better","values":[10159,5095,3917,5671,16206,6836],"isMain":true},{"metricName":"Total length","quality":"More is better","values":[76169,73651,158850,153112,93706,43392],"isMain":true},{"metricName":"Total length (>= 1000 bp)","quality":"More is better","values":[48261,54914,93925,90978,74944,30434],"isMain":true},{"metricName":"Total length (>= 5000 bp)","quality":"More is better","values":[10159,5095,0,5671,44459,12730],"isMain":false},{"metricName":"Total length (>= 10000 bp)","quality":"More is better","values":[10159,0,0,0,26975,0],"isMain":true},{"metricName":"Total length (>= 25000 bp)","quality":"More is better","values":[0,0,0,0,0,0],"isMain":false},{"metricName":"Total length (>= 50000 bp)","quality":"More is better","values":[0,0,0,0,0,0],"isMain":true},{"metricName":"N50","quality":"More is better","values":[1408,1708,1211,1230,3507,1566],"isMain":false},{"metricName":"N75","quality":"More is better","values":[874,999,799,740,1227,881],"isMain":false},{"metricName":"L50","quality":"Less is better","values":[14,15,42,34,5,6],"isMain":false},{"metricName":"L75","quality":"Less is better","values":[33,29,83,76,16,16],"isMain":false},{"metricName":"GC (%)","quality":"Equal","values":["42.71","43.15","41.58","42.32","44.47","40.81"],"isMain":false}]],["K-mer-based statistics",[]],["Predicted genes",[]],["Similarity statistics",[]],["Reference statistics",[{"metricName":"Reference length","quality":"Equal","values":[4900419,4900419,4900419,4900419,4900419,4900419],"isMain":false},{"metricName":"Reference fragments","quality":"Equal","values":[41,41,41,41,41,41],"isMain":false},{"metricName":"Reference GC (%)","quality":"Equal","values":["41.83","41.83","41.83","41.83","41.83","41.83"],"isMain":false}]]],"subreferences":[],"subreports":[],"minContig":500},{"date":"10 February 2020, Monday, 19:43:11","assembliesNames":["megahit-103-df","megahit-113-df","megahit-113-ml","megahit-113-ms","megahit-129-df","metaSPAdes"],"referenceName":"GCF\_000273315.1\_Bact\_xyla\_CL03T12C04\_V1","order":[0,1,2,3,4,5],"report":[["Genome statistics",[{"metricName":"Genome fraction (%)","quality":"More is better","values":["10.107","11.616","12.965","12.368","13.328","4.709"],"isMain":true},{"metricName":"Duplication ratio","quality":"Less is better","values":["1.028","1.032","1.030","1.033","1.028","1.031"],"isMain":true},{"metricName":"Largest alignment","quality":"More is better","values":[4231,12212,11371,12103,15040,2372],"isMain":true},{"metricName":"Total aligned length","quality":"More is better","values":[617448,709192,793605,757803,814762,287436],"isMain":true},{"metricName":"NA50","quality":"More is better","values":[646,677,649,648,679,641],"isMain":false},{"metricName":"NA75","quality":"More is better","values":[554,553,550,550,550,541],"isMain":false},{"metricName":"NGA50","quality":"More is better","values":[null,null,null,null,null,null],"isMain":true},{"metricName":"LA50","quality":"Less is better","values":[365,367,463,449,427,172],"isMain":false},{"metricName":"LA75","quality":"Less is better","values":[633,674,806,783,794,301],"isMain":false}]],["Reads mapping",[]],["Misassemblies",[{"metricName":"# misassemblies","quality":"Less is better","values":[4,9,7,3,7,1],"isMain":true},{"metricName":" # relocations","quality":"Less is better","values":[2,3,2,1,2,1],"isMain":false},{"metricName":" # translocations","quality":"Less is better","values":[2,5,3,1,5,0],"isMain":false},{"metricName":" # inversions","quality":"Less is better","values":[0,1,2,1,0,0],"isMain":false},{"metricName":" # interspecies translocations","quality":"Less is better","values":[38,53,34,46,49,17],"isMain":false},{"metricName":"# misassembled contigs","quality":"Less is better","values":[4,9,7,3,7,1],"isMain":false},{"metricName":"Misassembled contigs length","quality":"Less is better","values":[3106,23375,18310,2769,6883,1175],"isMain":true},{"metricName":"# possibly misassembled contigs","quality":"Less is better","values":[0,1,0,0,1,0],"isMain":false},{"metricName":" # possible misassemblies","quality":"Less is better","values":[0,1,0,0,1,0],"isMain":false},{"metricName":"# local misassemblies","quality":"Less is better","values":[0,3,1,3,1,0],"isMain":false},{"metricName":"# scaffold gap ext. mis.","quality":"Less is better","values":[0,0,0,0,0,0],"isMain":false},{"metricName":"# scaffold gap loc. mis.","quality":"Less is better","values":[0,0,0,0,0,0],"isMain":false},{"metricName":"# unaligned mis. contigs","quality":"Less is better","values":[0,0,0,0,3,0],"isMain":false}]],["Unaligned",[{"metricName":"# fully unaligned contigs","quality":"Less is better","values":[0,1,0,1,1,1],"isMain":false},{"metricName":"Fully unaligned length","quality":"Less is better","values":[0,553,0,733,519,1166],"isMain":false},{"metricName":"# partially unaligned contigs","quality":"Less is better","values":[9,15,10,6,18,5],"isMain":false},{"metricName":"Partially unaligned length","quality":"Less is better","values":[8944,19781,8020,17374,58542,3141],"isMain":false}]],["Mismatches",[{"metricName":"# mismatches","quality":"Less is better","values":[9525,10927,15797,15455,13492,4254],"isMain":false},{"metricName":"# indels","quality":"Less is better","values":[27,36,41,24,35,10],"isMain":false},{"metricName":"Indels length","quality":"Less is better","values":[69,92,89,71,166,53],"isMain":false},{"metricName":"# mismatches per 100 kbp","quality":"Less is better","values":["1556.09","1553.24","2011.87","2063.45","1671.50","1491.66"],"isMain":true},{"metricName":"# indels per 100 kbp","quality":"Less is better","values":["4.41","5.12","5.22","3.20","4.34","3.51"],"isMain":true},{"metricName":" # indels (<= 5 bp)","quality":"Less is better","values":[26,34,40,23,31,9],"isMain":false},{"metricName":" # indels (> 5 bp)","quality":"Less is better","values":[1,2,1,1,4,1],"isMain":false},{"metricName":"# N's","quality":"Less is better","values":[0,0,0,0,0,0],"isMain":false},{"metricName":"# N's per 100 kbp","quality":"Less is better","values":["0.00","0.00","0.00","0.00","0.00","0.00"],"isMain":true}]],["Statistics without reference",[{"metricName":"# contigs","quality":"Equal","values":[910,974,1162,1107,1085,430],"isMain":true},{"metricName":"# contigs (>= 1000 bp)","quality":"Equal","values":[84,112,82,84,134,35],"isMain":false},{"metricName":"# contigs (>= 5000 bp)","quality":"Equal","values":[0,2,2,2,4,0],"isMain":false},{"metricName":"# contigs (>= 10000 bp)","quality":"Equal","values":[0,2,1,2,4,0],"isMain":false},{"metricName":"# contigs (>= 25000 bp)","quality":"Equal","values":[0,0,0,0,2,0],"isMain":false},{"metricName":"# contigs (>= 50000 bp)","quality":"Equal","values":[0,0,0,0,0,0],"isMain":false},{"metricName":"Largest contig","quality":"More is better","values":[4265,16010,12935,12103,29537,2551],"isMain":true},{"metricName":"Total length","quality":"More is better","values":[638324,746046,816703,792135,888984,298354],"isMain":true},{"metricName":"Total length (>= 1000 bp)","quality":"More is better","values":[109399,186646,121953,135007,268856,47335],"isMain":true},{"metricName":"Total length (>= 5000 bp)","quality":"More is better","values":[0,28222,19351,24194,85966,0],"isMain":false},{"metricName":"Total length (>= 10000 bp)","quality":"More is better","values":[0,28222,12935,24194,85966,0],"isMain":true},{"metricName":"Total length (>= 25000 bp)","quality":"More is better","values":[0,0,0,0,55476,0],"isMain":false},{"metricName":"Total length (>= 50000 bp)","quality":"More is better","values":[0,0,0,0,0,0],"isMain":true},{"metricName":"N50","quality":"More is better","values":[668,719,666,673,754,667],"isMain":false},{"metricName":"N75","quality":"More is better","values":[569,586,568,571,592,560],"isMain":false},{"metricName":"L50","quality":"Less is better","values":[350,337,446,416,340,166],"isMain":false},{"metricName":"L75","quality":"Less is better","values":[611,627,779,738,676,289],"isMain":false},{"metricName":"GC (%)","quality":"Equal","values":["41.62","41.42","41.83","41.69","41.73","40.70"],"isMain":false}]],["K-mer-based statistics",[]],["Predicted genes",[]],["Similarity statistics",[]],["Reference statistics",[{"metricName":"Reference length","quality":"Equal","values":[6056103,6056103,6056103,6056103,6056103,6056103],"isMain":false},{"metricName":"Reference fragments","quality":"Equal","values":[41,41,41,41,41,41],"isMain":false},{"metricName":"Reference GC (%)","quality":"Equal","values":["42.20","42.20","42.20","42.20","42.20","42.20"],"isMain":false}]]],"subreferences":[],"subreports":[],"minContig":500},{"date":"10 February 2020, Monday, 19:43:01","assembliesNames":["megahit-103-df","megahit-113-df","megahit-113-ml","megahit-113-ms","megahit-129-df","metaSPAdes"],"referenceName":"GCF\_000273765.1\_Bact\_frag\_CL05T00C42\_V1","order":[0,1,2,3,4,5],"report":[["Genome statistics",[{"metricName":"Genome fraction (%)","quality":"More is better","values":["1.853","1.559","1.936","2.220","2.013","0.950"],"isMain":true},{"metricName":"Duplication ratio","quality":"Less is better","values":["1.064","1.028","1.048","1.031","1.026","1.023"],"isMain":true},{"metricName":"Largest alignment","quality":"More is better","values":[4725,2494,1614,9872,6306,5069],"isMain":true},{"metricName":"Total aligned length","quality":"More is better","values":[98418,82821,103084,118005,106297,50273],"isMain":true},{"metricName":"NA50","quality":"More is better","values":[696,666,622,692,695,702],"isMain":false},{"metricName":"NA75","quality":"More is better","values":[544,529,534,557,522,564],"isMain":false},{"metricName":"NGA50","quality":"More is better","values":[null,null,null,null,null,null],"isMain":true},{"metricName":"LA50","quality":"Less is better","values":[56,51,69,49,59,21],"isMain":false},{"metricName":"LA75","quality":"Less is better","values":[101,92,119,100,110,42],"isMain":false}]],["Reads mapping",[]],["Misassemblies",[{"metricName":"# misassemblies","quality":"Less is better","values":[1,0,0,1,1,0],"isMain":true},{"metricName":" # relocations","quality":"Less is better","values":[0,0,0,1,0,0],"isMain":false},{"metricName":" # translocations","quality":"Less is better","values":[1,0,0,0,1,0],"isMain":false},{"metricName":" # inversions","quality":"Less is better","values":[0,0,0,0,0,0],"isMain":false},{"metricName":" # interspecies translocations","quality":"Less is better","values":[25,11,15,12,12,2],"isMain":false},{"metricName":"# misassembled contigs","quality":"Less is better","values":[1,0,0,1,1,0],"isMain":false},{"metricName":"Misassembled contigs length","quality":"Less is better","values":[1038,0,0,16558,792,0],"isMain":true},{"metricName":"# possibly misassembled contigs","quality":"Less is better","values":[0,0,0,0,0,0],"isMain":false},{"metricName":" # possible misassemblies","quality":"Less is better","values":[0,0,0,0,0,0],"isMain":false},{"metricName":"# local misassemblies","quality":"Less is better","values":[1,0,1,0,1,0],"isMain":false},{"metricName":"# scaffold gap ext. mis.","quality":"Less is better","values":[0,0,0,0,0,0],"isMain":false},{"metricName":"# scaffold gap loc. mis.","quality":"Less is better","values":[0,0,0,0,0,0],"isMain":false},{"metricName":"# unaligned mis. contigs","quality":"Less is better","values":[0,0,0,0,0,0],"isMain":false}]],["Unaligned",[{"metricName":"# fully unaligned contigs","quality":"Less is better","values":[0,1,1,1,1,0],"isMain":false},{"metricName":"Fully unaligned length","quality":"Less is better","values":[0,1031,506,613,615,0],"isMain":false},{"metricName":"# partially unaligned contigs","quality":"Less is better","values":[8,7,4,5,6,1],"isMain":false},{"metricName":"Partially unaligned length","quality":"Less is better","values":[6134,11136,6536,6741,12348,519],"isMain":false}]],["Mismatches",[{"metricName":"# mismatches","quality":"Less is better","values":[1327,1348,2197,2390,1908,769],"isMain":false},{"metricName":"# indels","quality":"Less is better","values":[6,9,10,19,18,0],"isMain":false},{"metricName":"Indels length","quality":"Less is better","values":[6,10,17,62,30,0],"isMain":false},{"metricName":"# mismatches per 100 kbp","quality":"Less is better","values":["1356.30","1638.13","2149.83","2038.90","1794.95","1533.83"],"isMain":true},{"metricName":"# indels per 100 kbp","quality":"Less is better","values":["6.13","10.94","9.79","16.21","16.93","0.00"],"isMain":true},{"metricName":" # indels (<= 5 bp)","quality":"Less is better","values":[6,9,9,17,17,0],"isMain":false},{"metricName":" # indels (> 5 bp)","quality":"Less is better","values":[0,0,1,2,1,0],"isMain":false},{"metricName":"# N's","quality":"Less is better","values":[0,0,0,0,0,0],"isMain":false},{"metricName":"# N's per 100 kbp","quality":"Less is better","values":["0.00","0.00","0.00","0.00","0.00","0.00"],"isMain":true}]],["Statistics without reference",[{"metricName":"# contigs","quality":"Equal","values":[143,115,157,146,143,65],"isMain":true},{"metricName":"# contigs (>= 1000 bp)","quality":"Equal","values":[17,15,17,20,24,10],"isMain":false},{"metricName":"# contigs (>= 5000 bp)","quality":"Equal","values":[0,0,0,1,1,1],"isMain":false},{"metricName":"# contigs (>= 10000 bp)","quality":"Equal","values":[0,0,0,1,0,0],"isMain":false},{"metricName":"# contigs (>= 25000 bp)","quality":"Equal","values":[0,0,0,0,0,0],"isMain":false},{"metricName":"# contigs (>= 50000 bp)","quality":"Equal","values":[0,0,0,0,0,0],"isMain":false},{"metricName":"Largest contig","quality":"More is better","values":[4725,4533,2899,16558,6334,5069],"isMain":true},{"metricName":"Total length","quality":"More is better","values":[110249,96730,114188,128173,122027,51795],"isMain":true},{"metricName":"Total length (>= 1000 bp)","quality":"More is better","values":[25920,29634,23679,46881,43947,17574],"isMain":true},{"metricName":"Total length (>= 5000 bp)","quality":"More is better","values":[0,0,0,16558,6334,5069],"isMain":false},{"metricName":"Total length (>= 10000 bp)","quality":"More is better","values":[0,0,0,16558,0,0],"isMain":true},{"metricName":"Total length (>= 25000 bp)","quality":"More is better","values":[0,0,0,0,0,0],"isMain":false},{"metricName":"Total length (>= 50000 bp)","quality":"More is better","values":[0,0,0,0,0,0],"isMain":true},{"metricName":"N50","quality":"More is better","values":[747,779,692,759,794,704],"isMain":false},{"metricName":"N75","quality":"More is better","values":[603,610,582,605,612,575],"isMain":false},{"metricName":"L50","quality":"Less is better","values":[52,37,59,40,44,21],"isMain":false},{"metricName":"L75","quality":"Less is better","values":[93,72,104,88,87,41],"isMain":false},{"metricName":"GC (%)","quality":"Equal","values":["42.89","42.27","42.97","42.34","40.29","39.83"],"isMain":false}]],["K-mer-based statistics",[]],["Predicted genes",[]],["Similarity statistics",[]],["Reference statistics",[{"metricName":"Reference length","quality":"Equal","values":[5279416,5279416,5279416,5279416,5279416,5279416],"isMain":false},{"metricName":"Reference fragments","quality":"Equal","values":[25,25,25,25,25,25],"isMain":false},{"metricName":"Reference GC (%)","quality":"Equal","values":["43.49","43.49","43.49","43.49","43.49","43.49"],"isMain":false}]]],"subreferences":[],"subreports":[],"minContig":500},{"date":"10 February 2020, Monday, 19:42:50","assembliesNames":["megahit-103-df","megahit-113-df","megahit-113-ml","megahit-113-ms","megahit-129-df","metaSPAdes"],"referenceName":"GCF\_000296465.1\_Barn\_inte\_YIT\_11860\_V1","order":[0,1,2,3,4,5],"report":[["Genome statistics",[{"metricName":"Genome fraction (%)","quality":"More is better","values":["0.202","0.008","0.340","0.160","0.255","0.277"],"isMain":true},{"metricName":"Duplication ratio","quality":"Less is better","values":["1.097","1.989","1.049","1.212","1.000","1.004"],"isMain":true},{"metricName":"Largest alignment","quality":"More is better","values":[1513,278,5357,1719,4798,2915],"isMain":true},{"metricName":"Total aligned length","quality":"More is better","values":[7042,278,11678,5508,8744,9522],"isMain":true},{"metricName":"NA50","quality":"More is better","values":[1176,278,1432,736,4798,2724],"isMain":false},{"metricName":"NA75","quality":"More is better","values":[869,null,626,493,1440,2659],"isMain":false},{"metricName":"NGA50","quality":"More is better","values":[null,null,null,null,null,null],"isMain":true},{"metricName":"LA50","quality":"Less is better","values":[3,1,2,3,1,2],"isMain":false},{"metricName":"LA75","quality":"Less is better","values":[5,null,5,6,3,3],"isMain":false}]],["Reads mapping",[]],["Misassemblies",[{"metricName":"# misassemblies","quality":"Less is better","values":[0,0,0,0,0,0],"isMain":true},{"metricName":" # relocations","quality":"Less is better","values":[0,0,0,0,0,0],"isMain":false},{"metricName":" # translocations","quality":"Less is better","values":[0,0,0,0,0,0],"isMain":false},{"metricName":" # inversions","quality":"Less is better","values":[0,0,0,0,0,0],"isMain":false},{"metricName":" # interspecies translocations","quality":"Less is better","values":[2,1,1,3,0,0],"isMain":false},{"metricName":"# misassembled contigs","quality":"Less is better","values":[0,0,0,0,0,0],"isMain":false},{"metricName":"Misassembled contigs length","quality":"Less is better","values":[0,0,0,0,0,0],"isMain":true},{"metricName":"# possibly misassembled contigs","quality":"Less is better","values":[0,0,0,0,0,0],"isMain":false},{"metricName":" # possible misassemblies","quality":"Less is better","values":[0,0,0,0,0,0],"isMain":false},{"metricName":"# local misassemblies","quality":"Less is better","values":[0,0,0,0,0,0],"isMain":false},{"metricName":"# scaffold gap ext. mis.","quality":"Less is better","values":[0,0,0,0,0,0],"isMain":false},{"metricName":"# scaffold gap loc. mis.","quality":"Less is better","values":[0,0,0,0,0,0],"isMain":false},{"metricName":"# unaligned mis. contigs","quality":"Less is better","values":[0,0,0,0,0,0],"isMain":false}]],["Unaligned",[{"metricName":"# fully unaligned contigs","quality":"Less is better","values":[0,0,0,0,0,0],"isMain":false},{"metricName":"Fully unaligned length","quality":"Less is better","values":[0,0,0,0,0,0],"isMain":false},{"metricName":"# partially unaligned contigs","quality":"Less is better","values":[0,0,0,0,0,0],"isMain":false},{"metricName":"Partially unaligned length","quality":"Less is better","values":[0,0,0,0,0,0],"isMain":false}]],["Mismatches",[{"metricName":"# mismatches","quality":"Less is better","values":[48,3,148,82,58,31],"isMain":false},{"metricName":"# indels","quality":"Less is better","values":[0,0,0,0,0,0],"isMain":false},{"metricName":"Indels length","quality":"Less is better","values":[0,0,0,0,0,0],"isMain":false},{"metricName":"# mismatches per 100 kbp","quality":"Less is better","values":["691.34","1079.14","1267.34","1488.74","663.31","325.56"],"isMain":true},{"metricName":"# indels per 100 kbp","quality":"Less is better","values":["0.00","0.00","0.00","0.00","0.00","0.00"],"isMain":true},{"metricName":" # indels (<= 5 bp)","quality":"Less is better","values":[0,0,0,0,0,0],"isMain":false},{"metricName":" # indels (> 5 bp)","quality":"Less is better","values":[0,0,0,0,0,0],"isMain":false},{"metricName":"# N's","quality":"Less is better","values":[0,0,0,0,0,0],"isMain":false},{"metricName":"# N's per 100 kbp","quality":"Less is better","values":["0.00","0.00","0.00","0.00","0.00","0.00"],"isMain":true}]],["Statistics without reference",[{"metricName":"# contigs","quality":"Equal","values":[8,1,10,8,5,5],"isMain":true},{"metricName":"# contigs (>= 1000 bp)","quality":"Equal","values":[4,0,3,1,3,3],"isMain":false},{"metricName":"# contigs (>= 5000 bp)","quality":"Equal","values":[0,0,1,0,0,0],"isMain":false},{"metricName":"# contigs (>= 10000 bp)","quality":"Equal","values":[0,0,0,0,0,0],"isMain":false},{"metricName":"# contigs (>= 25000 bp)","quality":"Equal","values":[0,0,0,0,0,0],"isMain":false},{"metricName":"# contigs (>= 50000 bp)","quality":"Equal","values":[0,0,0,0,0,0],"isMain":false},{"metricName":"Largest contig","quality":"More is better","values":[1513,553,5388,1733,4798,2915],"isMain":true},{"metricName":"Total length","quality":"More is better","values":[7616,553,12249,6677,8744,9563],"isMain":true},{"metricName":"Total length (>= 1000 bp)","quality":"More is better","values":[5096,0,8034,1733,7688,8332],"isMain":true},{"metricName":"Total length (>= 5000 bp)","quality":"More is better","values":[0,0,5388,0,0,0],"isMain":false},{"metricName":"Total length (>= 10000 bp)","quality":"More is better","values":[0,0,0,0,0,0],"isMain":true},{"metricName":"Total length (>= 25000 bp)","quality":"More is better","values":[0,0,0,0,0,0],"isMain":false},{"metricName":"Total length (>= 50000 bp)","quality":"More is better","values":[0,0,0,0,0,0],"isMain":true},{"metricName":"N50","quality":"More is better","values":[1176,553,1434,942,4798,2737],"isMain":false},{"metricName":"N75","quality":"More is better","values":[869,553,655,624,1440,2680],"isMain":false},{"metricName":"L50","quality":"Less is better","values":[3,1,2,3,1,2],"isMain":false},{"metricName":"L75","quality":"Less is better","values":[5,1,5,6,3,3],"isMain":false},{"metricName":"GC (%)","quality":"Equal","values":["51.43","34.36","52.77","50.68","53.39","50.20"],"isMain":false}]],["K-mer-based statistics",[]],["Predicted genes",[]],["Similarity statistics",[]],["Reference statistics",[{"metricName":"Reference length","quality":"Equal","values":[3433706,3433706,3433706,3433706,3433706,3433706],"isMain":false},{"metricName":"Reference fragments","quality":"Equal","values":[19,19,19,19,19,19],"isMain":false},{"metricName":"Reference GC (%)","quality":"Equal","values":["43.91","43.91","43.91","43.91","43.91","43.91"],"isMain":false}]]],"subreferences":[],"subreports":[],"minContig":500},{"date":"10 February 2020, Monday, 19:43:19","assembliesNames":["megahit-103-df","megahit-113-df","megahit-113-ml","megahit-113-ms","megahit-129-df","metaSPAdes"],"referenceName":"GCF\_000297695.1\_Bact\_frag\_HMW\_610\_V1","order":[0,1,2,3,4,5],"report":[["Genome statistics",[{"metricName":"Genome fraction (%)","quality":"More is better","values":["28.688","27.046","31.484","28.269","29.948","14.371"],"isMain":true},{"metricName":"Duplication ratio","quality":"Less is better","values":["1.036","1.034","1.039","1.042","1.038","1.031"],"isMain":true},{"metricName":"Largest alignment","quality":"More is better","values":[2771,7831,35486,12580,6830,4495],"isMain":true},{"metricName":"Total aligned length","quality":"More is better","values":[1614801,1519808,1784519,1599331,1690762,802688],"isMain":true},{"metricName":"NA50","quality":"More is better","values":[739,782,719,719,776,673],"isMain":false},{"metricName":"NA75","quality":"More is better","values":[596,615,582,582,587,562],"isMain":false},{"metricName":"NGA50","quality":"More is better","values":[null,null,null,null,null,null],"isMain":true},{"metricName":"LA50","quality":"Less is better","values":[818,717,909,853,854,463],"isMain":false},{"metricName":"LA75","quality":"Less is better","values":[1440,1290,1627,1492,1547,801],"isMain":false}]],["Reads mapping",[]],["Misassemblies",[{"metricName":"# misassemblies","quality":"Less is better","values":[6,4,4,10,10,1],"isMain":true},{"metricName":" # relocations","quality":"Less is better","values":[0,1,0,1,1,0],"isMain":false},{"metricName":" # translocations","quality":"Less is better","values":[3,1,4,6,4,1],"isMain":false},{"metricName":" # inversions","quality":"Less is better","values":[3,2,0,3,5,0],"isMain":false},{"metricName":" # interspecies translocations","quality":"Less is better","values":[68,75,30,69,70,32],"isMain":false},{"metricName":"# misassembled contigs","quality":"Less is better","values":[6,4,4,10,10,1],"isMain":false},{"metricName":"Misassembled contigs length","quality":"Less is better","values":[4985,3239,44790,7574,7585,1160],"isMain":true},{"metricName":"# possibly misassembled contigs","quality":"Less is better","values":[0,1,0,0,0,0],"isMain":false},{"metricName":" # possible misassemblies","quality":"Less is better","values":[0,1,0,0,0,0],"isMain":false},{"metricName":"# local misassemblies","quality":"Less is better","values":[1,0,7,3,4,0],"isMain":false},{"metricName":"# scaffold gap ext. mis.","quality":"Less is better","values":[0,0,0,0,0,0],"isMain":false},{"metricName":"# scaffold gap loc. mis.","quality":"Less is better","values":[0,0,0,0,0,0],"isMain":false},{"metricName":"# unaligned mis. contigs","quality":"Less is better","values":[0,0,0,1,1,0],"isMain":false}]],["Unaligned",[{"metricName":"# fully unaligned contigs","quality":"Less is better","values":[0,1,0,2,0,0],"isMain":false},{"metricName":"Fully unaligned length","quality":"Less is better","values":[0,506,0,1212,0,0],"isMain":false},{"metricName":"# partially unaligned contigs","quality":"Less is better","values":[8,20,4,14,15,7],"isMain":false},{"metricName":"Partially unaligned length","quality":"Less is better","values":[7730,37311,38492,26627,147196,8703],"isMain":false}]],["Mismatches",[{"metricName":"# mismatches","quality":"Less is better","values":[23409,21339,33710,32051,24525,11718],"isMain":false},{"metricName":"# indels","quality":"Less is better","values":[69,73,81,77,77,28],"isMain":false},{"metricName":"Indels length","quality":"Less is better","values":[135,104,203,92,142,105],"isMain":false},{"metricName":"# mismatches per 100 kbp","quality":"Less is better","values":["1477.44","1428.54","1938.60","2052.85","1482.75","1476.34"],"isMain":true},{"metricName":"# indels per 100 kbp","quality":"Less is better","values":["4.35","4.89","4.66","4.93","4.66","3.53"],"isMain":true},{"metricName":" # indels (<= 5 bp)","quality":"Less is better","values":[67,71,77,76,72,25],"isMain":false},{"metricName":" # indels (> 5 bp)","quality":"Less is better","values":[2,2,4,1,5,3],"isMain":false},{"metricName":"# N's","quality":"Less is better","values":[0,0,0,0,0,0],"isMain":false},{"metricName":"# N's per 100 kbp","quality":"Less is better","values":["0.00","0.00","0.00","0.00","0.00","0.00"],"isMain":true}]],["Statistics without reference",[{"metricName":"# contigs","quality":"Equal","values":[2151,1915,2381,2181,2103,1160],"isMain":true},{"metricName":"# contigs (>= 1000 bp)","quality":"Equal","values":[320,382,315,305,425,107],"isMain":false},{"metricName":"# contigs (>= 5000 bp)","quality":"Equal","values":[0,2,3,3,5,0],"isMain":false},{"metricName":"# contigs (>= 10000 bp)","quality":"Equal","values":[0,1,2,2,2,0],"isMain":false},{"metricName":"# contigs (>= 25000 bp)","quality":"Equal","values":[0,0,2,0,2,0],"isMain":false},{"metricName":"# contigs (>= 50000 bp)","quality":"Equal","values":[0,0,0,0,1,0],"isMain":false},{"metricName":"Largest contig","quality":"More is better","values":[3738,16891,42898,12580,89771,4506],"isMain":true},{"metricName":"Total length","quality":"More is better","values":[1648627,1581922,1845558,1655487,1864414,827388],"isMain":true},{"metricName":"Total length (>= 1000 bp)","quality":"More is better","values":[412071,528090,473780,401070,705821,138518],"isMain":true},{"metricName":"Total length (>= 5000 bp)","quality":"More is better","values":[0,24722,84520,28978,154240,0],"isMain":false},{"metricName":"Total length (>= 10000 bp)","quality":"More is better","values":[0,16891,78873,23400,135164,0],"isMain":true},{"metricName":"Total length (>= 25000 bp)","quality":"More is better","values":[0,0,78873,0,135164,0],"isMain":false},{"metricName":"Total length (>= 50000 bp)","quality":"More is better","values":[0,0,0,0,89771,0],"isMain":true},{"metricName":"N50","quality":"More is better","values":[760,822,746,740,871,696],"isMain":false},{"metricName":"N75","quality":"More is better","values":[612,644,603,607,656,580],"isMain":false},{"metricName":"L50","quality":"Less is better","values":[799,673,846,811,670,448],"isMain":false},{"metricName":"L75","quality":"Less is better","values":[1404,1220,1540,1427,1289,776],"isMain":false},{"metricName":"GC (%)","quality":"Equal","values":["43.45","43.45","43.51","43.32","43.67","43.08"],"isMain":false}]],["K-mer-based statistics",[]],["Predicted genes",[]],["Similarity statistics",[]],["Reference statistics",[{"metricName":"Reference length","quality":"Equal","values":[5523030,5523030,5523030,5523030,5523030,5523030],"isMain":false},{"metricName":"Reference fragments","quality":"Equal","values":[42,42,42,42,42,42],"isMain":false},{"metricName":"Reference GC (%)","quality":"Equal","values":["43.31","43.31","43.31","43.31","43.31","43.31"],"isMain":false}]]],"subreferences":[],"subreports":[],"minContig":500},{"date":"10 February 2020, Monday, 19:43:13","assembliesNames":["megahit-103-df","megahit-113-df","megahit-113-ml","megahit-113-ms","megahit-129-df","metaSPAdes"],"referenceName":"GCF\_000297735.1\_Bact\_frag\_HMW\_615\_V1","order":[0,1,2,3,4,5],"report":[["Genome statistics",[{"metricName":"Genome fraction (%)","quality":"More is better","values":["2.288","0.762","1.051","3.601","0.570","0.422"],"isMain":true},{"metricName":"Duplication ratio","quality":"Less is better","values":["1.024","1.069","1.058","1.015","1.052","1.027"],"isMain":true},{"metricName":"Largest alignment","quality":"More is better","values":[15796,11129,12461,62347,5390,3727],"isMain":true},{"metricName":"Total aligned length","quality":"More is better","values":[122803,41511,56992,194057,31419,23088],"isMain":true},{"metricName":"NA50","quality":"More is better","values":[1021,1230,936,34978,805,1235],"isMain":false},{"metricName":"NA75","quality":"More is better","values":[579,593,null,null,null,849],"isMain":false},{"metricName":"NGA50","quality":"More is better","values":[null,null,null,null,null,null],"isMain":true},{"metricName":"LA50","quality":"Less is better","values":[27,7,14,3,12,5],"isMain":false},{"metricName":"LA75","quality":"Less is better","values":[72,21,null,null,null,11],"isMain":false}]],["Reads mapping",[]],["Misassemblies",[{"metricName":"# misassemblies","quality":"Less is better","values":[0,0,1,3,0,0],"isMain":true},{"metricName":" # relocations","quality":"Less is better","values":[0,0,1,3,0,0],"isMain":false},{"metricName":" # translocations","quality":"Less is better","values":[0,0,0,0,0,0],"isMain":false},{"metricName":" # inversions","quality":"Less is better","values":[0,0,0,0,0,0],"isMain":false},{"metricName":" # interspecies translocations","quality":"Less is better","values":[22,11,9,10,5,0],"isMain":false},{"metricName":"# misassembled contigs","quality":"Less is better","values":[0,0,1,2,0,0],"isMain":false},{"metricName":"Misassembled contigs length","quality":"Less is better","values":[0,0,873,125591,0,0],"isMain":true},{"metricName":"# possibly misassembled contigs","quality":"Less is better","values":[0,0,0,0,0,0],"isMain":false},{"metricName":" # possible misassemblies","quality":"Less is better","values":[0,0,0,0,0,0],"isMain":false},{"metricName":"# local misassemblies","quality":"Less is better","values":[0,0,0,1,0,0],"isMain":false},{"metricName":"# scaffold gap ext. mis.","quality":"Less is better","values":[0,0,0,0,0,0],"isMain":false},{"metricName":"# scaffold gap loc. mis.","quality":"Less is better","values":[0,0,0,0,0,0],"isMain":false},{"metricName":"# unaligned mis. contigs","quality":"Less is better","values":[0,1,1,1,1,0],"isMain":false}]],["Unaligned",[{"metricName":"# fully unaligned contigs","quality":"Less is better","values":[1,0,0,1,0,0],"isMain":false},{"metricName":"Fully unaligned length","quality":"Less is better","values":[787,0,0,1371,0,0],"isMain":false},{"metricName":"# partially unaligned contigs","quality":"Less is better","values":[14,5,3,4,4,0],"isMain":false},{"metricName":"Partially unaligned length","quality":"Less is better","values":[10059,4505,22355,74004,10155,0],"isMain":false}]],["Mismatches",[{"metricName":"# mismatches","quality":"Less is better","values":[397,412,835,1220,471,167],"isMain":false},{"metricName":"# indels","quality":"Less is better","values":[3,3,10,15,8,0],"isMain":false},{"metricName":"Indels length","quality":"Less is better","values":[3,5,41,42,53,0],"isMain":false},{"metricName":"# mismatches per 100 kbp","quality":"Less is better","values":["323.88","1009.28","1483.28","632.21","1542.64","738.91"],"isMain":true},{"metricName":"# indels per 100 kbp","quality":"Less is better","values":["2.45","7.35","17.76","7.77","26.20","0.00"],"isMain":true},{"metricName":" # indels (<= 5 bp)","quality":"Less is better","values":[3,3,8,14,7,0],"isMain":false},{"metricName":" # indels (> 5 bp)","quality":"Less is better","values":[0,0,2,1,1,0],"isMain":false},{"metricName":"# N's","quality":"Less is better","values":[0,0,0,0,0,0],"isMain":false},{"metricName":"# N's per 100 kbp","quality":"Less is better","values":["0.00","0.00","0.00","0.00","0.00","0.00"],"isMain":true}]],["Statistics without reference",[{"metricName":"# contigs","quality":"Equal","values":[126,36,41,40,30,19],"isMain":true},{"metricName":"# contigs (>= 1000 bp)","quality":"Equal","values":[31,11,12,14,11,6],"isMain":false},{"metricName":"# contigs (>= 5000 bp)","quality":"Equal","values":[3,1,3,3,2,0],"isMain":false},{"metricName":"# contigs (>= 10000 bp)","quality":"Equal","values":[1,1,2,3,1,0],"isMain":false},{"metricName":"# contigs (>= 25000 bp)","quality":"Equal","values":[0,0,1,3,0,0],"isMain":false},{"metricName":"# contigs (>= 50000 bp)","quality":"Equal","values":[0,0,0,3,0,0],"isMain":false},{"metricName":"Largest contig","quality":"More is better","values":[15796,11129,26373,125035,10007,3736],"isMain":true},{"metricName":"Total length","quality":"More is better","values":[136303,48134,81902,271176,42270,23222],"isMain":true},{"metricName":"Total length (>= 1000 bp)","quality":"More is better","values":[73515,30892,62476,254901,29755,13549],"isMain":true},{"metricName":"Total length (>= 5000 bp)","quality":"More is better","values":[29713,11129,44423,236330,15443,0],"isMain":false},{"metricName":"Total length (>= 10000 bp)","quality":"More is better","values":[15796,11129,38895,236330,10007,0],"isMain":true},{"metricName":"Total length (>= 25000 bp)","quality":"More is better","values":[0,0,26373,236330,0,0],"isMain":false},{"metricName":"Total length (>= 50000 bp)","quality":"More is better","values":[0,0,0,236330,0,0],"isMain":true},{"metricName":"N50","quality":"More is better","values":[1072,1694,5528,61148,1955,1255],"isMain":false},{"metricName":"N75","quality":"More is better","values":[690,804,1065,50147,854,854],"isMain":false},{"metricName":"L50","quality":"Less is better","values":[26,6,3,2,5,5],"isMain":false},{"metricName":"L75","quality":"Less is better","values":[66,17,11,3,14,11],"isMain":false},{"metricName":"GC (%)","quality":"Equal","values":["45.35","45.29","43.58","45.69","46.51","43.81"],"isMain":false}]],["K-mer-based statistics",[]],["Predicted genes",[]],["Similarity statistics",[]],["Reference statistics",[{"metricName":"Reference length","quality":"Equal","values":[5358243,5358243,5358243,5358243,5358243,5358243],"isMain":false},{"metricName":"Reference fragments","quality":"Equal","values":[43,43,43,43,43,43],"isMain":false},{"metricName":"Reference GC (%)","quality":"Equal","values":["43.48","43.48","43.48","43.48","43.48","43.48"],"isMain":false}]]],"subreferences":[],"subreports":[],"minContig":500},{"date":"10 February 2020, Monday, 19:43:05","assembliesNames":["megahit-103-df","megahit-113-df","megahit-113-ml","megahit-113-ms","megahit-129-df","metaSPAdes"],"referenceName":"GCF\_000297755.1\_Bact\_frag\_HMW\_616\_V1","order":[0,1,2,3,4,5],"report":[["Genome statistics",[{"metricName":"Genome fraction (%)","quality":"More is better","values":["3.742","2.960","4.557","4.104","3.587","1.720"],"isMain":true},{"metricName":"Duplication ratio","quality":"Less is better","values":["1.021","1.052","1.024","1.025","1.021","1.033"],"isMain":true},{"metricName":"Largest alignment","quality":"More is better","values":[5490,10166,6374,4430,7227,8225],"isMain":true},{"metricName":"Total aligned length","quality":"More is better","values":[203370,161226,247737,223458,195676,93568],"isMain":true},{"metricName":"NA50","quality":"More is better","values":[758,658,661,694,631,658],"isMain":false},{"metricName":"NA75","quality":"More is better","values":[606,552,558,567,332,548],"isMain":false},{"metricName":"NGA50","quality":"More is better","values":[null,null,null,null,null,null],"isMain":true},{"metricName":"LA50","quality":"Less is better","values":[88,91,138,108,134,46],"isMain":false},{"metricName":"LA75","quality":"Less is better","values":[164,165,246,201,250,88],"isMain":false}]],["Reads mapping",[]],["Misassemblies",[{"metricName":"# misassemblies","quality":"Less is better","values":[2,3,2,4,2,0],"isMain":true},{"metricName":" # relocations","quality":"Less is better","values":[1,0,0,1,0,0],"isMain":false},{"metricName":" # translocations","quality":"Less is better","values":[0,3,2,2,2,0],"isMain":false},{"metricName":" # inversions","quality":"Less is better","values":[1,0,0,1,0,0],"isMain":false},{"metricName":" # interspecies translocations","quality":"Less is better","values":[12,26,11,14,15,9],"isMain":false},{"metricName":"# misassembled contigs","quality":"Less is better","values":[2,3,2,3,2,0],"isMain":false},{"metricName":"Misassembled contigs length","quality":"Less is better","values":[1800,2150,1785,1840,1472,0],"isMain":true},{"metricName":"# possibly misassembled contigs","quality":"Less is better","values":[0,1,0,0,0,0],"isMain":false},{"metricName":" # possible misassemblies","quality":"Less is better","values":[0,1,0,0,0,0],"isMain":false},{"metricName":"# local misassemblies","quality":"Less is better","values":[0,0,0,0,0,0],"isMain":false},{"metricName":"# scaffold gap ext. mis.","quality":"Less is better","values":[0,0,0,0,0,0],"isMain":false},{"metricName":"# scaffold gap loc. mis.","quality":"Less is better","values":[0,0,0,0,0,0],"isMain":false},{"metricName":"# unaligned mis. contigs","quality":"Less is better","values":[0,0,0,0,1,0],"isMain":false}]],["Unaligned",[{"metricName":"# fully unaligned contigs","quality":"Less is better","values":[0,0,1,0,1,0],"isMain":false},{"metricName":"Fully unaligned length","quality":"Less is better","values":[0,0,531,0,740,0],"isMain":false},{"metricName":"# partially unaligned contigs","quality":"Less is better","values":[2,8,4,3,8,3],"isMain":false},{"metricName":"Partially unaligned length","quality":"Less is better","values":[1562,7971,7938,1615,58143,3248],"isMain":false}]],["Mismatches",[{"metricName":"# mismatches","quality":"Less is better","values":[2935,2615,4926,4702,2967,1243],"isMain":false},{"metricName":"# indels","quality":"Less is better","values":[21,17,13,25,15,2],"isMain":false},{"metricName":"Indels length","quality":"Less is better","values":[40,34,16,32,24,2],"isMain":false},{"metricName":"# mismatches per 100 kbp","quality":"Less is better","values":["1448.05","1631.14","1995.81","2114.94","1527.08","1333.98"],"isMain":true},{"metricName":"# indels per 100 kbp","quality":"Less is better","values":["10.36","10.60","5.27","11.24","7.72","2.15"],"isMain":true},{"metricName":" # indels (<= 5 bp)","quality":"Less is better","values":[20,16,13,25,14,2],"isMain":false},{"metricName":" # indels (> 5 bp)","quality":"Less is better","values":[1,1,0,0,1,0],"isMain":false},{"metricName":"# N's","quality":"Less is better","values":[0,0,0,0,0,0],"isMain":false},{"metricName":"# N's per 100 kbp","quality":"Less is better","values":["0.00","0.00","0.00","0.00","0.00","0.00"],"isMain":true}]],["Statistics without reference",[{"metricName":"# contigs","quality":"Equal","values":[253,229,348,301,258,129],"isMain":true},{"metricName":"# contigs (>= 1000 bp)","quality":"Equal","values":[43,25,26,33,41,9],"isMain":false},{"metricName":"# contigs (>= 5000 bp)","quality":"Equal","values":[2,1,2,0,2,1],"isMain":false},{"metricName":"# contigs (>= 10000 bp)","quality":"Equal","values":[0,1,0,0,1,0],"isMain":false},{"metricName":"# contigs (>= 25000 bp)","quality":"Equal","values":[0,0,0,0,1,0],"isMain":false},{"metricName":"# contigs (>= 50000 bp)","quality":"Equal","values":[0,0,0,0,1,0],"isMain":false},{"metricName":"Largest contig","quality":"More is better","values":[5507,10166,6979,4436,51971,8225],"isMain":true},{"metricName":"Total length","quality":"More is better","values":[208594,176548,261214,229467,257295,99484],"isMain":true},{"metricName":"Total length (>= 1000 bp)","quality":"More is better","values":[67374,41889,49463,54275,114363,23142],"isMain":true},{"metricName":"Total length (>= 5000 bp)","quality":"More is better","values":[10833,10166,13353,0,59198,8225],"isMain":false},{"metricName":"Total length (>= 10000 bp)","quality":"More is better","values":[0,10166,0,0,51971,0],"isMain":true},{"metricName":"Total length (>= 25000 bp)","quality":"More is better","values":[0,0,0,0,51971,0],"isMain":false},{"metricName":"Total length (>= 50000 bp)","quality":"More is better","values":[0,0,0,0,51971,0],"isMain":true},{"metricName":"N50","quality":"More is better","values":[778,712,705,718,877,692],"isMain":false},{"metricName":"N75","quality":"More is better","values":[627,590,585,593,639,578],"isMain":false},{"metricName":"L50","quality":"Less is better","values":[85,81,125,106,57,43],"isMain":false},{"metricName":"L75","quality":"Less is better","values":[160,149,228,196,145,83],"isMain":false},{"metricName":"GC (%)","quality":"Equal","values":["44.27","44.51","44.12","44.71","43.95","44.07"],"isMain":false}]],["K-mer-based statistics",[]],["Predicted genes",[]],["Similarity statistics",[]],["Reference statistics",[{"metricName":"Reference length","quality":"Equal","values":[5416584,5416584,5416584,5416584,5416584,5416584],"isMain":false},{"metricName":"Reference fragments","quality":"Equal","values":[25,25,25,25,25,25],"isMain":false},{"metricName":"Reference GC (%)","quality":"Equal","values":["43.37","43.37","43.37","43.37","43.37","43.37"],"isMain":false}]]],"subreferences":[],"subreports":[],"minContig":500},{"date":"10 February 2020, Monday, 19:43:06","assembliesNames":["megahit-103-df","megahit-113-df","megahit-113-ml","megahit-113-ms","megahit-129-df","metaSPAdes"],"referenceName":"GCF\_000304195.1\_Bact\_thet\_CL09T03C10\_V1","order":[0,1,2,3,4,5],"report":[["Genome statistics",[{"metricName":"Genome fraction (%)","quality":"More is better","values":["0.394","0.516","0.466","0.323","0.335","0.253"],"isMain":true},{"metricName":"Duplication ratio","quality":"Less is better","values":["1.025","1.028","1.034","1.082","1.072","1.006"],"isMain":true},{"metricName":"Largest alignment","quality":"More is better","values":[1927,3023,2206,1165,1459,1368],"isMain":true},{"metricName":"Total aligned length","quality":"More is better","values":[20186,26511,24027,16686,17067,12888],"isMain":true},{"metricName":"NA50","quality":"More is better","values":[928,508,703,null,617,805],"isMain":false},{"metricName":"NA75","quality":"More is better","values":[635,null,527,null,77,660],"isMain":false},{"metricName":"NGA50","quality":"More is better","values":[null,null,null,null,null,null],"isMain":true},{"metricName":"LA50","quality":"Less is better","values":[9,23,14,null,13,8],"isMain":false},{"metricName":"LA75","quality":"Less is better","values":[15,null,26,null,27,13],"isMain":false}]],["Reads mapping",[]],["Misassemblies",[{"metricName":"# misassemblies","quality":"Less is better","values":[0,0,0,0,0,0],"isMain":true},{"metricName":" # relocations","quality":"Less is better","values":[0,0,0,0,0,0],"isMain":false},{"metricName":" # translocations","quality":"Less is better","values":[0,0,0,0,0,0],"isMain":false},{"metricName":" # inversions","quality":"Less is better","values":[0,0,0,0,0,0],"isMain":false},{"metricName":" # interspecies translocations","quality":"Less is better","values":[1,7,4,7,7,2],"isMain":false},{"metricName":"# misassembled contigs","quality":"Less is better","values":[0,0,0,0,0,0],"isMain":false},{"metricName":"Misassembled contigs length","quality":"Less is better","values":[0,0,0,0,0,0],"isMain":true},{"metricName":"# possibly misassembled contigs","quality":"Less is better","values":[0,0,0,0,0,0],"isMain":false},{"metricName":" # possible misassemblies","quality":"Less is better","values":[0,0,0,0,0,0],"isMain":false},{"metricName":"# local misassemblies","quality":"Less is better","values":[0,0,0,0,0,0],"isMain":false},{"metricName":"# scaffold gap ext. mis.","quality":"Less is better","values":[0,0,0,0,0,0],"isMain":false},{"metricName":"# scaffold gap loc. mis.","quality":"Less is better","values":[0,0,0,0,0,0],"isMain":false},{"metricName":"# unaligned mis. contigs","quality":"Less is better","values":[0,0,0,0,0,0],"isMain":false}]],["Unaligned",[{"metricName":"# fully unaligned contigs","quality":"Less is better","values":[0,1,1,2,1,0],"isMain":false},{"metricName":"Fully unaligned length","quality":"Less is better","values":[0,505,612,1635,830,0],"isMain":false},{"metricName":"# partially unaligned contigs","quality":"Less is better","values":[0,6,2,3,4,2],"isMain":false},{"metricName":"Partially unaligned length","quality":"Less is better","values":[0,20072,2106,32321,3603,1794],"isMain":false}]],["Mismatches",[{"metricName":"# mismatches","quality":"Less is better","values":[317,317,465,391,282,242],"isMain":false},{"metricName":"# indels","quality":"Less is better","values":[3,2,0,4,4,3],"isMain":false},{"metricName":"Indels length","quality":"Less is better","values":[3,3,0,6,4,3],"isMain":false},{"metricName":"# mismatches per 100 kbp","quality":"Less is better","values":["1578.29","1205.14","1958.31","2372.14","1652.51","1878.15"],"isMain":true},{"metricName":"# indels per 100 kbp","quality":"Less is better","values":["14.94","7.60","0.00","24.27","23.44","23.28"],"isMain":true},{"metricName":" # indels (<= 5 bp)","quality":"Less is better","values":[3,2,0,4,4,3],"isMain":false},{"metricName":" # indels (> 5 bp)","quality":"Less is better","values":[0,0,0,0,0,0],"isMain":false},{"metricName":"# N's","quality":"Less is better","values":[0,0,0,0,0,0],"isMain":false},{"metricName":"# N's per 100 kbp","quality":"Less is better","values":["0.00","0.00","0.00","0.00","0.00","0.00"],"isMain":true}]],["Statistics without reference",[{"metricName":"# contigs","quality":"Equal","values":[24,31,35,31,28,17],"isMain":true},{"metricName":"# contigs (>= 1000 bp)","quality":"Equal","values":[7,13,4,4,4,4],"isMain":false},{"metricName":"# contigs (>= 5000 bp)","quality":"Equal","values":[0,1,0,1,0,0],"isMain":false},{"metricName":"# contigs (>= 10000 bp)","quality":"Equal","values":[0,1,0,1,0,0],"isMain":false},{"metricName":"# contigs (>= 25000 bp)","quality":"Equal","values":[0,0,0,1,0,0],"isMain":false},{"metricName":"# contigs (>= 50000 bp)","quality":"Equal","values":[0,0,0,0,0,0],"isMain":false},{"metricName":"Largest contig","quality":"More is better","values":[1927,16677,2206,29629,2363,1441],"isMain":true},{"metricName":"Total length","quality":"More is better","values":[20592,47621,27270,51792,22720,14753],"isMain":true},{"metricName":"Total length (>= 1000 bp)","quality":"More is better","values":[9288,36089,6849,34419,6284,5094],"isMain":true},{"metricName":"Total length (>= 5000 bp)","quality":"More is better","values":[0,16677,0,29629,0,0],"isMain":false},{"metricName":"Total length (>= 10000 bp)","quality":"More is better","values":[0,16677,0,29629,0,0],"isMain":true},{"metricName":"Total length (>= 25000 bp)","quality":"More is better","values":[0,0,0,29629,0,0],"isMain":false},{"metricName":"Total length (>= 50000 bp)","quality":"More is better","values":[0,0,0,0,0,0],"isMain":true},{"metricName":"N50","quality":"More is better","values":[939,1873,783,29629,779,830],"isMain":false},{"metricName":"N75","quality":"More is better","values":[640,1015,581,711,618,685],"isMain":false},{"metricName":"L50","quality":"Less is better","values":[9,5,12,1,10,7],"isMain":false},{"metricName":"L75","quality":"Less is better","values":[15,13,23,10,19,12],"isMain":false},{"metricName":"GC (%)","quality":"Equal","values":["44.44","43.92","46.48","47.56","45.29","44.62"],"isMain":false}]],["K-mer-based statistics",[]],["Predicted genes",[]],["Similarity statistics",[]],["Reference statistics",[{"metricName":"Reference length","quality":"Equal","values":[5097678,5097678,5097678,5097678,5097678,5097678],"isMain":false},{"metricName":"Reference fragments","quality":"Equal","values":[16,16,16,16,16,16],"isMain":false},{"metricName":"Reference GC (%)","quality":"Equal","values":["42.48","42.48","42.48","42.48","42.48","42.48"],"isMain":false}]]],"subreferences":[],"subreports":[],"minContig":500},{"date":"10 February 2020, Monday, 19:43:07","assembliesNames":["megahit-103-df","megahit-113-df","megahit-113-ml","megahit-113-ms","megahit-129-df","metaSPAdes"],"referenceName":"GCF\_000307265.1\_ASM30726v1","order":[0,1,2,3,4,5],"report":[["Genome statistics",[{"metricName":"Genome fraction (%)","quality":"More is better","values":["3.411","4.123","4.417","5.450","4.583","1.874"],"isMain":true},{"metricName":"Duplication ratio","quality":"Less is better","values":["1.059","1.064","1.020","1.048","1.058","1.077"],"isMain":true},{"metricName":"Largest alignment","quality":"More is better","values":[1052,1245,1149,1058,1413,964],"isMain":true},{"metricName":"Total aligned length","quality":"More is better","values":[105271,127071,136356,168176,141104,57959],"isMain":true},{"metricName":"NA50","quality":"More is better","values":[557,554,565,561,566,550],"isMain":false},{"metricName":"NA75","quality":"More is better","values":[511,505,517,509,512,509],"isMain":false},{"metricName":"NGA50","quality":"More is better","values":[null,null,null,null,null,null],"isMain":true},{"metricName":"LA50","quality":"Less is better","values":[91,111,105,145,119,49],"isMain":false},{"metricName":"LA75","quality":"Less is better","values":[147,181,172,235,195,79],"isMain":false}]],["Reads mapping",[]],["Misassemblies",[{"metricName":"# misassemblies","quality":"Less is better","values":[0,1,0,0,0,0],"isMain":true},{"metricName":" # relocations","quality":"Less is better","values":[0,0,0,0,0,0],"isMain":false},{"metricName":" # translocations","quality":"Less is better","values":[0,1,0,0,0,0],"isMain":false},{"metricName":" # inversions","quality":"Less is better","values":[0,0,0,0,0,0],"isMain":false},{"metricName":" # interspecies translocations","quality":"Less is better","values":[27,33,9,28,32,14],"isMain":false},{"metricName":"# misassembled contigs","quality":"Less is better","values":[0,1,0,0,0,0],"isMain":false},{"metricName":"Misassembled contigs length","quality":"Less is better","values":[0,537,0,0,0,0],"isMain":true},{"metricName":"# possibly misassembled contigs","quality":"Less is better","values":[0,0,0,0,0,0],"isMain":false},{"metricName":" # possible misassemblies","quality":"Less is better","values":[0,0,0,0,0,0],"isMain":false},{"metricName":"# local misassemblies","quality":"Less is better","values":[0,0,1,0,0,1],"isMain":false},{"metricName":"# scaffold gap ext. mis.","quality":"Less is better","values":[0,0,0,0,0,0],"isMain":false},{"metricName":"# scaffold gap loc. mis.","quality":"Less is better","values":[0,0,0,0,0,0],"isMain":false},{"metricName":"# unaligned mis. contigs","quality":"Less is better","values":[1,0,0,0,0,0],"isMain":false}]],["Unaligned",[{"metricName":"# fully unaligned contigs","quality":"Less is better","values":[0,0,0,1,0,0],"isMain":false},{"metricName":"Fully unaligned length","quality":"Less is better","values":[0,0,0,874,0,0],"isMain":false},{"metricName":"# partially unaligned contigs","quality":"Less is better","values":[10,13,4,6,13,1],"isMain":false},{"metricName":"Partially unaligned length","quality":"Less is better","values":[8247,12800,3631,14248,13940,756],"isMain":false}]],["Mismatches",[{"metricName":"# mismatches","quality":"Less is better","values":[1857,2149,2710,3442,2388,1020],"isMain":false},{"metricName":"# indels","quality":"Less is better","values":[7,5,4,2,4,2],"isMain":false},{"metricName":"Indels length","quality":"Less is better","values":[9,5,7,4,4,4],"isMain":false},{"metricName":"# mismatches per 100 kbp","quality":"Less is better","values":["1768.62","1693.07","1992.72","2051.64","1692.37","1768.29"],"isMain":true},{"metricName":"# indels per 100 kbp","quality":"Less is better","values":["6.67","3.94","2.94","1.19","2.83","3.47"],"isMain":true},{"metricName":" # indels (<= 5 bp)","quality":"Less is better","values":[7,5,4,2,4,2],"isMain":false},{"metricName":" # indels (> 5 bp)","quality":"Less is better","values":[0,0,0,0,0,0],"isMain":false},{"metricName":"# N's","quality":"Less is better","values":[0,0,0,0,0,0],"isMain":false},{"metricName":"# N's per 100 kbp","quality":"Less is better","values":["0.00","0.00","0.00","0.00","0.00","0.00"],"isMain":true}]],["Statistics without reference",[{"metricName":"# contigs","quality":"Equal","values":[190,227,234,296,249,104],"isMain":true},{"metricName":"# contigs (>= 1000 bp)","quality":"Equal","values":[7,8,3,5,9,0],"isMain":false},{"metricName":"# contigs (>= 5000 bp)","quality":"Equal","values":[0,0,0,1,0,0],"isMain":false},{"metricName":"# contigs (>= 10000 bp)","quality":"Equal","values":[0,0,0,0,0,0],"isMain":false},{"metricName":"# contigs (>= 25000 bp)","quality":"Equal","values":[0,0,0,0,0,0],"isMain":false},{"metricName":"# contigs (>= 50000 bp)","quality":"Equal","values":[0,0,0,0,0,0],"isMain":false},{"metricName":"Largest contig","quality":"More is better","values":[1693,3332,1247,8135,2610,976],"isMain":true},{"metricName":"Total length","quality":"More is better","values":[119414,147888,142381,191012,163271,62898],"isMain":true},{"metricName":"Total length (>= 1000 bp)","quality":"More is better","values":[8684,13505,3459,15210,14718,0],"isMain":true},{"metricName":"Total length (>= 5000 bp)","quality":"More is better","values":[0,0,0,8135,0,0],"isMain":false},{"metricName":"Total length (>= 10000 bp)","quality":"More is better","values":[0,0,0,0,0,0],"isMain":true},{"metricName":"Total length (>= 25000 bp)","quality":"More is better","values":[0,0,0,0,0,0],"isMain":false},{"metricName":"Total length (>= 50000 bp)","quality":"More is better","values":[0,0,0,0,0,0],"isMain":true},{"metricName":"N50","quality":"More is better","values":[604,612,589,599,622,587],"isMain":false},{"metricName":"N75","quality":"More is better","values":[541,544,528,540,547,539],"isMain":false},{"metricName":"L50","quality":"Less is better","values":[80,92,101,120,101,46],"isMain":false},{"metricName":"L75","quality":"Less is better","values":[133,156,165,204,172,74],"isMain":false},{"metricName":"GC (%)","quality":"Equal","values":["54.96","55.08","54.53","54.52","55.15","53.83"],"isMain":false}]],["K-mer-based statistics",[]],["Predicted genes",[]],["Similarity statistics",[]],["Reference statistics",[{"metricName":"Reference length","quality":"Equal","values":[3078564,3078564,3078564,3078564,3078564,3078564],"isMain":false},{"metricName":"Reference fragments","quality":"Equal","values":[58,58,58,58,58,58],"isMain":false},{"metricName":"Reference GC (%)","quality":"Equal","values":["54.97","54.97","54.97","54.97","54.97","54.97"],"isMain":false}]]],"subreferences":[],"subreports":[],"minContig":500},{"date":"10 February 2020, Monday, 19:43:32","assembliesNames":["megahit-103-df","megahit-113-df","megahit-113-ml","megahit-113-ms","megahit-129-df","metaSPAdes"],"referenceName":"GCF\_000307345.1\_Para\_merd\_CL03T12C32\_V1","order":[0,1,2,3,4,5],"report":[["Genome statistics",[{"metricName":"Genome fraction (%)","quality":"More is better","values":["38.955","43.567","39.762","39.122","45.481","30.019"],"isMain":true},{"metricName":"Duplication ratio","quality":"Less is better","values":["1.043","1.050","1.056","1.054","1.055","1.040"],"isMain":true},{"metricName":"Largest alignment","quality":"More is better","values":[4152,6945,4707,4122,8576,4373],"isMain":true},{"metricName":"Total aligned length","quality":"More is better","values":[1962362,2201920,2028834,1987814,2311084,1503464],"isMain":true},{"metricName":"NG50","quality":"More is better","values":[null,null,null,null,522,null],"isMain":false},{"metricName":"NA50","quality":"More is better","values":[831,907,777,774,914,736],"isMain":false},{"metricName":"NA75","quality":"More is better","values":[630,648,608,600,641,585],"isMain":false},{"metricName":"NGA50","quality":"More is better","values":[null,null,null,null,null,null],"isMain":true},{"metricName":"LG50","quality":"Less is better","values":[null,null,null,null,2422,null],"isMain":false},{"metricName":"LA50","quality":"Less is better","values":[852,867,973,949,936,761],"isMain":false},{"metricName":"LA75","quality":"Less is better","values":[1546,1620,1736,1705,1764,1351],"isMain":false}]],["Reads mapping",[]],["Misassemblies",[{"metricName":"# misassemblies","quality":"Less is better","values":[12,17,6,10,18,4],"isMain":true},{"metricName":" # relocations","quality":"Less is better","values":[1,7,1,0,6,1],"isMain":false},{"metricName":" # translocations","quality":"Less is better","values":[9,9,5,7,9,3],"isMain":false},{"metricName":" # inversions","quality":"Less is better","values":[2,1,0,3,3,0],"isMain":false},{"metricName":" # interspecies translocations","quality":"Less is better","values":[83,120,58,88,124,45],"isMain":false},{"metricName":"# misassembled contigs","quality":"Less is better","values":[12,17,6,10,18,4],"isMain":false},{"metricName":"Misassembled contigs length","quality":"Less is better","values":[10721,20922,5199,7762,21471,5247],"isMain":true},{"metricName":"# possibly misassembled contigs","quality":"Less is better","values":[0,4,1,2,4,1],"isMain":false},{"metricName":" # possible misassemblies","quality":"Less is better","values":[0,4,1,2,4,1],"isMain":false},{"metricName":"# local misassemblies","quality":"Less is better","values":[7,7,4,3,9,1],"isMain":false},{"metricName":"# scaffold gap ext. mis.","quality":"Less is better","values":[0,0,0,0,0,0],"isMain":false},{"metricName":"# scaffold gap loc. mis.","quality":"Less is better","values":[0,0,0,0,0,0],"isMain":false},{"metricName":"# unaligned mis. contigs","quality":"Less is better","values":[1,1,0,1,3,0],"isMain":false}]],["Unaligned",[{"metricName":"# fully unaligned contigs","quality":"Less is better","values":[4,4,8,6,7,2],"isMain":false},{"metricName":"Fully unaligned length","quality":"Less is better","values":[2626,2730,6040,4918,4445,1571],"isMain":false},{"metricName":"# partially unaligned contigs","quality":"Less is better","values":[5,28,7,15,30,7],"isMain":false},{"metricName":"Partially unaligned length","quality":"Less is better","values":[4470,52740,16982,17646,151781,8907],"isMain":false}]],["Mismatches",[{"metricName":"# mismatches","quality":"Less is better","values":[29817,35186,40744,41775,37396,23159],"isMain":false},{"metricName":"# indels","quality":"Less is better","values":[154,218,143,173,189,141],"isMain":false},{"metricName":"Indels length","quality":"Less is better","values":[321,327,481,482,333,311],"isMain":false},{"metricName":"# mismatches per 100 kbp","quality":"Less is better","values":["1556.35","1642.19","2083.55","2171.19","1671.89","1568.65"],"isMain":true},{"metricName":"# indels per 100 kbp","quality":"Less is better","values":["8.04","10.17","7.31","8.99","8.45","9.55"],"isMain":true},{"metricName":" # indels (<= 5 bp)","quality":"Less is better","values":[144,211,131,155,184,135],"isMain":false},{"metricName":" # indels (> 5 bp)","quality":"Less is better","values":[10,7,12,18,5,6],"isMain":false},{"metricName":"# N's","quality":"Less is better","values":[0,0,0,0,0,0],"isMain":false},{"metricName":"# N's per 100 kbp","quality":"Less is better","values":["0.00","0.00","0.00","0.00","0.00","0.00"],"isMain":true}]],["Statistics without reference",[{"metricName":"# contigs","quality":"Equal","values":[2385,2480,2604,2556,2533,2004],"isMain":true},{"metricName":"# contigs (>= 1000 bp)","quality":"Equal","values":[525,730,476,464,813,294],"isMain":false},{"metricName":"# contigs (>= 5000 bp)","quality":"Equal","values":[0,3,1,1,6,1],"isMain":false},{"metricName":"# contigs (>= 10000 bp)","quality":"Equal","values":[0,1,1,0,2,0],"isMain":false},{"metricName":"# contigs (>= 25000 bp)","quality":"Equal","values":[0,0,0,0,1,0],"isMain":false},{"metricName":"# contigs (>= 50000 bp)","quality":"Equal","values":[0,0,0,0,1,0],"isMain":false},{"metricName":"Largest contig","quality":"More is better","values":[4358,19225,13613,6944,97232,5189],"isMain":true},{"metricName":"Total length","quality":"More is better","values":[2005868,2304495,2088029,2051140,2515826,1545962],"isMain":true},{"metricName":"Total length (>= 1000 bp)","quality":"More is better","values":[719337,1080375,635240,623986,1301436,394466],"isMain":true},{"metricName":"Total length (>= 5000 bp)","quality":"More is better","values":[0,36118,13613,6944,144949,5189],"isMain":false},{"metricName":"Total length (>= 10000 bp)","quality":"More is better","values":[0,19225,13613,0,109150,0],"isMain":true},{"metricName":"Total length (>= 25000 bp)","quality":"More is better","values":[0,0,0,0,97232,0],"isMain":false},{"metricName":"Total length (>= 50000 bp)","quality":"More is better","values":[0,0,0,0,97232,0],"isMain":true},{"metricName":"N50","quality":"More is better","values":[855,963,802,805,1025,757],"isMain":false},{"metricName":"N75","quality":"More is better","values":[652,705,632,630,727,611],"isMain":false},{"metricName":"L50","quality":"Less is better","values":[834,804,939,916,770,736],"isMain":false},{"metricName":"L75","quality":"Less is better","values":[1507,1508,1674,1639,1500,1302],"isMain":false},{"metricName":"GC (%)","quality":"Equal","values":["45.41","45.44","45.48","45.53","45.44","45.59"],"isMain":false}]],["K-mer-based statistics",[]],["Predicted genes",[]],["Similarity statistics",[]],["Reference statistics",[{"metricName":"Reference length","quality":"Equal","values":[4918045,4918045,4918045,4918045,4918045,4918045],"isMain":false},{"metricName":"Reference fragments","quality":"Equal","values":[37,37,37,37,37,37],"isMain":false},{"metricName":"Reference GC (%)","quality":"Equal","values":["45.25","45.25","45.25","45.25","45.25","45.25"],"isMain":false}]]],"subreferences":[],"subreports":[],"minContig":500},{"date":"10 February 2020, Monday, 19:43:20","assembliesNames":["megahit-103-df","megahit-113-df","megahit-113-ml","megahit-113-ms","megahit-129-df","metaSPAdes"],"referenceName":"GCF\_000307455.1\_Para\_dist\_CL03T12C09\_V1","order":[0,1,2,3,4,5],"report":[["Genome statistics",[{"metricName":"Genome fraction (%)","quality":"More is better","values":["6.218","7.389","6.701","6.836","7.858","4.935"],"isMain":true},{"metricName":"Duplication ratio","quality":"Less is better","values":["1.017","1.022","1.023","1.029","1.021","1.024"],"isMain":true},{"metricName":"Largest alignment","quality":"More is better","values":[2031,5009,2220,3354,3088,4729],"isMain":true},{"metricName":"Total aligned length","quality":"More is better","values":[316171,375503,341510,348234,399324,250541],"isMain":true},{"metricName":"NA50","quality":"More is better","values":[745,877,701,695,880,692],"isMain":false},{"metricName":"NA75","quality":"More is better","values":[604,640,564,570,631,575],"isMain":false},{"metricName":"NGA50","quality":"More is better","values":[null,null,null,null,null,null],"isMain":true},{"metricName":"LA50","quality":"Less is better","values":[161,150,182,184,167,138],"isMain":false},{"metricName":"LA75","quality":"Less is better","values":[283,281,321,327,308,240],"isMain":false}]],["Reads mapping",[]],["Misassemblies",[{"metricName":"# misassemblies","quality":"Less is better","values":[1,1,1,0,2,0],"isMain":true},{"metricName":" # relocations","quality":"Less is better","values":[1,0,0,0,0,0],"isMain":false},{"metricName":" # translocations","quality":"Less is better","values":[0,1,0,0,1,0],"isMain":false},{"metricName":" # inversions","quality":"Less is better","values":[0,0,1,0,1,0],"isMain":false},{"metricName":" # interspecies translocations","quality":"Less is better","values":[13,24,5,17,20,12],"isMain":false},{"metricName":"# misassembled contigs","quality":"Less is better","values":[1,1,1,0,2,0],"isMain":false},{"metricName":"Misassembled contigs length","quality":"Less is better","values":[1214,1520,506,0,1729,0],"isMain":true},{"metricName":"# possibly misassembled contigs","quality":"Less is better","values":[1,0,0,0,0,0],"isMain":false},{"metricName":" # possible misassemblies","quality":"Less is better","values":[1,0,0,0,0,0],"isMain":false},{"metricName":"# local misassemblies","quality":"Less is better","values":[0,0,1,2,0,0],"isMain":false},{"metricName":"# scaffold gap ext. mis.","quality":"Less is better","values":[0,0,0,0,0,0],"isMain":false},{"metricName":"# scaffold gap loc. mis.","quality":"Less is better","values":[0,0,0,0,0,0],"isMain":false},{"metricName":"# unaligned mis. contigs","quality":"Less is better","values":[0,0,1,0,0,0],"isMain":false}]],["Unaligned",[{"metricName":"# fully unaligned contigs","quality":"Less is better","values":[1,2,0,1,1,0],"isMain":false},{"metricName":"Fully unaligned length","quality":"Less is better","values":[1035,1127,0,703,596,0],"isMain":false},{"metricName":"# partially unaligned contigs","quality":"Less is better","values":[2,6,0,5,6,3],"isMain":false},{"metricName":"Partially unaligned length","quality":"Less is better","values":[3001,6285,0,4623,9031,2678],"isMain":false}]],["Mismatches",[{"metricName":"# mismatches","quality":"Less is better","values":[5019,5873,6907,7388,6478,4123],"isMain":false},{"metricName":"# indels","quality":"Less is better","values":[16,24,30,24,19,5],"isMain":false},{"metricName":"Indels length","quality":"Less is better","values":[81,52,114,27,24,62],"isMain":false},{"metricName":"# mismatches per 100 kbp","quality":"Less is better","values":["1596.46","1572.12","2038.85","2137.66","1630.65","1652.38"],"isMain":true},{"metricName":"# indels per 100 kbp","quality":"Less is better","values":["5.09","6.42","8.86","6.94","4.78","2.00"],"isMain":true},{"metricName":" # indels (<= 5 bp)","quality":"Less is better","values":[15,22,26,24,18,4],"isMain":false},{"metricName":" # indels (> 5 bp)","quality":"Less is better","values":[1,2,4,0,1,1],"isMain":false},{"metricName":"# N's","quality":"Less is better","values":[0,0,0,0,0,0],"isMain":false},{"metricName":"# N's per 100 kbp","quality":"Less is better","values":["0.00","0.00","0.00","0.00","0.00","0.00"],"isMain":true}]],["Statistics without reference",[{"metricName":"# contigs","quality":"Equal","values":[420,435,477,479,470,351],"isMain":true},{"metricName":"# contigs (>= 1000 bp)","quality":"Equal","values":[58,128,55,70,121,33],"isMain":false},{"metricName":"# contigs (>= 5000 bp)","quality":"Equal","values":[0,1,0,0,0,0],"isMain":false},{"metricName":"# contigs (>= 10000 bp)","quality":"Equal","values":[0,0,0,0,0,0],"isMain":false},{"metricName":"# contigs (>= 25000 bp)","quality":"Equal","values":[0,0,0,0,0,0],"isMain":false},{"metricName":"# contigs (>= 50000 bp)","quality":"Equal","values":[0,0,0,0,0,0],"isMain":false},{"metricName":"Largest contig","quality":"More is better","values":[2960,5009,2224,3354,4710,4759],"isMain":true},{"metricName":"Total length","quality":"More is better","values":[323683,389344,346497,360844,415350,258153],"isMain":true},{"metricName":"Total length (>= 1000 bp)","quality":"More is better","values":[78223,181053,70926,91549,174416,45693],"isMain":true},{"metricName":"Total length (>= 5000 bp)","quality":"More is better","values":[0,5009,0,0,0,0],"isMain":false},{"metricName":"Total length (>= 10000 bp)","quality":"More is better","values":[0,0,0,0,0,0],"isMain":true},{"metricName":"Total length (>= 25000 bp)","quality":"More is better","values":[0,0,0,0,0,0],"isMain":false},{"metricName":"Total length (>= 50000 bp)","quality":"More is better","values":[0,0,0,0,0,0],"isMain":true},{"metricName":"N50","quality":"More is better","values":[760,962,718,726,913,708],"isMain":false},{"metricName":"N75","quality":"More is better","values":[618,665,575,599,666,593],"isMain":false},{"metricName":"L50","quality":"Less is better","values":[156,142,180,176,156,133],"isMain":false},{"metricName":"L75","quality":"Less is better","values":[274,267,316,314,290,232],"isMain":false},{"metricName":"GC (%)","quality":"Equal","values":["44.93","45.20","45.48","45.18","45.20","45.46"],"isMain":false}]],["K-mer-based statistics",[]],["Predicted genes",[]],["Similarity statistics",[]],["Reference statistics",[{"metricName":"Reference length","quality":"Equal","values":[5055862,5055862,5055862,5055862,5055862,5055862],"isMain":false},{"metricName":"Reference fragments","quality":"Equal","values":[24,24,24,24,24,24],"isMain":false},{"metricName":"Reference GC (%)","quality":"Equal","values":["45.13","45.13","45.13","45.13","45.13","45.13"],"isMain":false}]]],"subreferences":[],"subreports":[],"minContig":500},{"date":"10 February 2020, Monday, 19:43:42","assembliesNames":["megahit-103-df","megahit-113-df","megahit-113-ml","megahit-113-ms","megahit-129-df","metaSPAdes"],"referenceName":"GCF\_000307475.1\_Paraba\_sp\_D25\_V1","order":[0,1,2,3,4,5],"report":[["Genome statistics",[{"metricName":"Genome fraction (%)","quality":"More is better","values":["5.597","6.620","6.024","6.005","6.409","4.431"],"isMain":true},{"metricName":"Duplication ratio","quality":"Less is better","values":["1.018","1.025","1.015","1.027","1.022","1.020"],"isMain":true},{"metricName":"Largest alignment","quality":"More is better","values":[2957,5329,2941,2383,3020,1794],"isMain":true},{"metricName":"Total aligned length","quality":"More is better","values":[289891,343046,311563,311673,332539,229574],"isMain":true},{"metricName":"NA50","quality":"More is better","values":[811,891,716,733,891,697],"isMain":false},{"metricName":"NA75","quality":"More is better","values":[623,627,590,592,667,567],"isMain":false},{"metricName":"NGA50","quality":"More is better","values":[null,null,null,null,null,null],"isMain":true},{"metricName":"LA50","quality":"Less is better","values":[132,134,159,164,123,120],"isMain":false},{"metricName":"LA75","quality":"Less is better","values":[237,255,281,286,234,214],"isMain":false}]],["Reads mapping",[]],["Misassemblies",[{"metricName":"# misassemblies","quality":"Less is better","values":[1,4,0,0,0,0],"isMain":true},{"metricName":" # relocations","quality":"Less is better","values":[0,0,0,0,0,0],"isMain":false},{"metricName":" # translocations","quality":"Less is better","values":[1,4,0,0,0,0],"isMain":false},{"metricName":" # inversions","quality":"Less is better","values":[0,0,0,0,0,0],"isMain":false},{"metricName":" # interspecies translocations","quality":"Less is better","values":[15,21,8,15,16,6],"isMain":false},{"metricName":"# misassembled contigs","quality":"Less is better","values":[1,4,0,0,0,0],"isMain":false},{"metricName":"Misassembled contigs length","quality":"Less is better","values":[723,3669,0,0,0,0],"isMain":true},{"metricName":"# possibly misassembled contigs","quality":"Less is better","values":[1,1,0,0,0,0],"isMain":false},{"metricName":" # possible misassemblies","quality":"Less is better","values":[1,1,0,0,0,0],"isMain":false},{"metricName":"# local misassemblies","quality":"Less is better","values":[2,0,1,0,1,0],"isMain":false},{"metricName":"# scaffold gap ext. mis.","quality":"Less is better","values":[0,0,0,0,0,0],"isMain":false},{"metricName":"# scaffold gap loc. mis.","quality":"Less is better","values":[0,0,0,0,0,0],"isMain":false},{"metricName":"# unaligned mis. contigs","quality":"Less is better","values":[0,0,0,0,0,0],"isMain":false}]],["Unaligned",[{"metricName":"# fully unaligned contigs","quality":"Less is better","values":[0,0,0,0,0,0],"isMain":false},{"metricName":"Fully unaligned length","quality":"Less is better","values":[0,0,0,0,0,0],"isMain":false},{"metricName":"# partially unaligned contigs","quality":"Less is better","values":[2,3,3,2,3,2],"isMain":false},{"metricName":"Partially unaligned length","quality":"Less is better","values":[1388,3166,2263,1800,2259,1277],"isMain":false}]],["Mismatches",[{"metricName":"# mismatches","quality":"Less is better","values":[4580,5758,6401,6897,5400,3511],"isMain":false},{"metricName":"# indels","quality":"Less is better","values":[21,20,23,21,21,10],"isMain":false},{"metricName":"Indels length","quality":"Less is better","values":[60,42,26,32,50,10],"isMain":false},{"metricName":"# mismatches per 100 kbp","quality":"Less is better","values":["1586.78","1686.68","2060.43","2227.12","1633.78","1536.62"],"isMain":true},{"metricName":"# indels per 100 kbp","quality":"Less is better","values":["7.28","5.86","7.40","6.78","6.35","4.38"],"isMain":true},{"metricName":" # indels (<= 5 bp)","quality":"Less is better","values":[18,19,23,19,20,10],"isMain":false},{"metricName":" # indels (> 5 bp)","quality":"Less is better","values":[3,1,0,2,1,0],"isMain":false},{"metricName":"# N's","quality":"Less is better","values":[0,0,0,0,0,0],"isMain":false},{"metricName":"# N's per 100 kbp","quality":"Less is better","values":["0.00","0.00","0.00","0.00","0.00","0.00"],"isMain":true}]],["Statistics without reference",[{"metricName":"# contigs","quality":"Equal","values":[360,397,419,425,372,317],"isMain":true},{"metricName":"# contigs (>= 1000 bp)","quality":"Equal","values":[79,106,60,46,109,48],"isMain":false},{"metricName":"# contigs (>= 5000 bp)","quality":"Equal","values":[0,1,0,0,0,0],"isMain":false},{"metricName":"# contigs (>= 10000 bp)","quality":"Equal","values":[0,0,0,0,0,0],"isMain":false},{"metricName":"# contigs (>= 25000 bp)","quality":"Equal","values":[0,0,0,0,0,0],"isMain":false},{"metricName":"# contigs (>= 50000 bp)","quality":"Equal","values":[0,0,0,0,0,0],"isMain":false},{"metricName":"Largest contig","quality":"More is better","values":[2962,5329,2954,2383,3029,1820],"isMain":true},{"metricName":"Total length","quality":"More is better","values":[295201,353155,317633,319839,339908,234229],"isMain":true},{"metricName":"Total length (>= 1000 bp)","quality":"More is better","values":[102737,153719,77533,62044,158795,59695],"isMain":true},{"metricName":"Total length (>= 5000 bp)","quality":"More is better","values":[0,5329,0,0,0,0],"isMain":false},{"metricName":"Total length (>= 10000 bp)","quality":"More is better","values":[0,0,0,0,0,0],"isMain":true},{"metricName":"Total length (>= 25000 bp)","quality":"More is better","values":[0,0,0,0,0,0],"isMain":false},{"metricName":"Total length (>= 50000 bp)","quality":"More is better","values":[0,0,0,0,0,0],"isMain":true},{"metricName":"N50","quality":"More is better","values":[821,935,747,746,922,716],"isMain":false},{"metricName":"N75","quality":"More is better","values":[647,661,605,613,682,584],"isMain":false},{"metricName":"L50","quality":"Less is better","values":[129,130,155,161,121,118],"isMain":false},{"metricName":"L75","quality":"Less is better","values":[231,245,275,279,228,209],"isMain":false},{"metricName":"GC (%)","quality":"Equal","values":["45.85","45.83","45.62","45.70","45.72","45.88"],"isMain":false}]],["K-mer-based statistics",[]],["Predicted genes",[]],["Similarity statistics",[]],["Reference statistics",[{"metricName":"Reference length","quality":"Equal","values":[5157107,5157107,5157107,5157107,5157107,5157107],"isMain":false},{"metricName":"Reference fragments","quality":"Equal","values":[36,36,36,36,36,36],"isMain":false},{"metricName":"Reference GC (%)","quality":"Equal","values":["45.07","45.07","45.07","45.07","45.07","45.07"],"isMain":false}]]],"subreferences":[],"subreports":[],"minContig":500},{"date":"10 February 2020, Monday, 19:43:27","assembliesNames":["megahit-103-df","megahit-113-df","megahit-113-ml","megahit-113-ms","megahit-129-df","metaSPAdes"],"referenceName":"GCF\_000307495.1\_Para\_merd\_CL09T00C40\_V1","order":[0,1,2,3,4,5],"report":[["Genome statistics",[{"metricName":"Genome fraction (%)","quality":"More is better","values":["8.360","9.289","9.977","10.175","9.982","6.192"],"isMain":true},{"metricName":"Duplication ratio","quality":"Less is better","values":["1.028","1.032","1.032","1.035","1.038","1.030"],"isMain":true},{"metricName":"Largest alignment","quality":"More is better","values":[3618,3729,8274,15853,3085,3205],"isMain":true},{"metricName":"Total aligned length","quality":"More is better","values":[374554,416208,447883,456380,447621,276734],"isMain":true},{"metricName":"NA50","quality":"More is better","values":[765,781,729,748,798,670],"isMain":false},{"metricName":"NA75","quality":"More is better","values":[581,596,587,578,604,570],"isMain":false},{"metricName":"NGA50","quality":"More is better","values":[null,null,null,null,null,null],"isMain":true},{"metricName":"LA50","quality":"Less is better","values":[182,186,226,216,198,160],"isMain":false},{"metricName":"LA75","quality":"Less is better","values":[328,347,402,395,372,277],"isMain":false}]],["Reads mapping",[]],["Misassemblies",[{"metricName":"# misassemblies","quality":"Less is better","values":[2,1,3,2,3,2],"isMain":true},{"metricName":" # relocations","quality":"Less is better","values":[0,0,0,0,0,0],"isMain":false},{"metricName":" # translocations","quality":"Less is better","values":[2,0,3,2,2,2],"isMain":false},{"metricName":" # inversions","quality":"Less is better","values":[0,1,0,0,1,0],"isMain":false},{"metricName":" # interspecies translocations","quality":"Less is better","values":[27,30,18,24,39,16],"isMain":false},{"metricName":"# misassembled contigs","quality":"Less is better","values":[2,1,3,2,3,2],"isMain":false},{"metricName":"Misassembled contigs length","quality":"Less is better","values":[2247,527,1847,2154,3893,1635],"isMain":true},{"metricName":"# possibly misassembled contigs","quality":"Less is better","values":[0,0,2,0,1,0],"isMain":false},{"metricName":" # possible misassemblies","quality":"Less is better","values":[0,0,2,0,1,0],"isMain":false},{"metricName":"# local misassemblies","quality":"Less is better","values":[2,0,1,1,1,0],"isMain":false},{"metricName":"# scaffold gap ext. mis.","quality":"Less is better","values":[0,0,0,0,0,0],"isMain":false},{"metricName":"# scaffold gap loc. mis.","quality":"Less is better","values":[0,0,0,0,0,0],"isMain":false},{"metricName":"# unaligned mis. contigs","quality":"Less is better","values":[0,0,0,0,0,0],"isMain":false}]],["Unaligned",[{"metricName":"# fully unaligned contigs","quality":"Less is better","values":[0,3,1,2,3,0],"isMain":false},{"metricName":"Fully unaligned length","quality":"Less is better","values":[0,1694,903,1336,1975,0],"isMain":false},{"metricName":"# partially unaligned contigs","quality":"Less is better","values":[7,11,3,4,14,5],"isMain":false},{"metricName":"Partially unaligned length","quality":"Less is better","values":[4953,10098,2224,5792,12147,4151],"isMain":false}]],["Mismatches",[{"metricName":"# mismatches","quality":"Less is better","values":[6407,7900,9722,9945,8850,4900],"isMain":false},{"metricName":"# indels","quality":"Less is better","values":[52,70,53,35,57,26],"isMain":false},{"metricName":"Indels length","quality":"Less is better","values":[142,122,94,52,167,30],"isMain":false},{"metricName":"# mismatches per 100 kbp","quality":"Less is better","values":["1722.77","1911.86","2190.45","2197.07","1992.98","1778.84"],"isMain":true},{"metricName":"# indels per 100 kbp","quality":"Less is better","values":["13.98","16.94","11.94","7.73","12.84","9.44"],"isMain":true},{"metricName":" # indels (<= 5 bp)","quality":"Less is better","values":[49,68,50,34,50,26],"isMain":false},{"metricName":" # indels (> 5 bp)","quality":"Less is better","values":[3,2,3,1,7,0],"isMain":false},{"metricName":"# N's","quality":"Less is better","values":[0,0,0,0,0,0],"isMain":false},{"metricName":"# N's per 100 kbp","quality":"Less is better","values":["0.00","0.00","0.00","0.00","0.00","0.00"],"isMain":true}]],["Statistics without reference",[{"metricName":"# contigs","quality":"Equal","values":[491,522,600,592,554,399],"isMain":true},{"metricName":"# contigs (>= 1000 bp)","quality":"Equal","values":[86,119,89,93,123,47],"isMain":false},{"metricName":"# contigs (>= 5000 bp)","quality":"Equal","values":[0,0,1,1,0,0],"isMain":false},{"metricName":"# contigs (>= 10000 bp)","quality":"Equal","values":[0,0,0,1,0,0],"isMain":false},{"metricName":"# contigs (>= 25000 bp)","quality":"Equal","values":[0,0,0,0,0,0],"isMain":false},{"metricName":"# contigs (>= 50000 bp)","quality":"Equal","values":[0,0,0,0,0,0],"isMain":false},{"metricName":"Largest contig","quality":"More is better","values":[3618,3744,8274,15853,3085,3205],"isMain":true},{"metricName":"Total length","quality":"More is better","values":[387200,438178,461065,475496,474888,287755],"isMain":true},{"metricName":"Total length (>= 1000 bp)","quality":"More is better","values":[114988,168467,117698,137935,178605,58582],"isMain":true},{"metricName":"Total length (>= 5000 bp)","quality":"More is better","values":[0,0,8274,15853,0,0],"isMain":false},{"metricName":"Total length (>= 10000 bp)","quality":"More is better","values":[0,0,0,15853,0,0],"isMain":true},{"metricName":"Total length (>= 25000 bp)","quality":"More is better","values":[0,0,0,0,0,0],"isMain":false},{"metricName":"Total length (>= 50000 bp)","quality":"More is better","values":[0,0,0,0,0,0],"isMain":true},{"metricName":"N50","quality":"More is better","values":[788,843,749,799,860,690],"isMain":false},{"metricName":"N75","quality":"More is better","values":[613,638,615,616,654,589],"isMain":false},{"metricName":"L50","quality":"Less is better","values":[176,175,221,206,187,154],"isMain":false},{"metricName":"L75","quality":"Less is better","values":[315,327,391,376,347,267],"isMain":false},{"metricName":"GC (%)","quality":"Equal","values":["45.71","45.63","45.82","45.72","45.33","45.16"],"isMain":false}]],["K-mer-based statistics",[]],["Predicted genes",[]],["Similarity statistics",[]],["Reference statistics",[{"metricName":"Reference length","quality":"Equal","values":[4448563,4448563,4448563,4448563,4448563,4448563],"isMain":false},{"metricName":"Reference fragments","quality":"Equal","values":[38,38,38,38,38,38],"isMain":false},{"metricName":"Reference GC (%)","quality":"Equal","values":["45.25","45.25","45.25","45.25","45.25","45.25"],"isMain":false}]]],"subreferences":[],"subreports":[],"minContig":500},{"date":"10 February 2020, Monday, 19:43:56","assembliesNames":["megahit-103-df","megahit-113-df","megahit-113-ml","megahit-113-ms","megahit-129-df","metaSPAdes"],"referenceName":"GCF\_000312145.1\_ASM31214v1","order":[0,1,2,3,4,5],"report":[["Genome statistics",[{"metricName":"Genome fraction (%)","quality":"More is better","values":["0.247","0.156","0.370","0.298","0.236","0.078"],"isMain":true},{"metricName":"Duplication ratio","quality":"Less is better","values":["1.198","1.371","1.159","1.238","1.184","1.067"],"isMain":true},{"metricName":"Largest alignment","quality":"More is better","values":[1550,914,2772,2772,914,717],"isMain":true},{"metricName":"Total aligned length","quality":"More is better","values":[9838,6226,14860,12010,9421,3111],"isMain":true},{"metricName":"NA50","quality":"More is better","values":[608,140,548,479,null,483],"isMain":false},{"metricName":"NA75","quality":"More is better","values":[257,null,134,null,null,null],"isMain":false},{"metricName":"NGA50","quality":"More is better","values":[null,null,null,null,null,null],"isMain":true},{"metricName":"LA50","quality":"Less is better","values":[7,13,9,12,null,4],"isMain":false},{"metricName":"LA75","quality":"Less is better","values":[14,null,21,null,null,null],"isMain":false}]],["Reads mapping",[]],["Misassemblies",[{"metricName":"# misassemblies","quality":"Less is better","values":[0,0,1,0,0,0],"isMain":true},{"metricName":" # relocations","quality":"Less is better","values":[0,0,1,0,0,0],"isMain":false},{"metricName":" # translocations","quality":"Less is better","values":[0,0,0,0,0,0],"isMain":false},{"metricName":" # inversions","quality":"Less is better","values":[0,0,0,0,0,0],"isMain":false},{"metricName":" # interspecies translocations","quality":"Less is better","values":[4,8,6,7,10,2],"isMain":false},{"metricName":"# misassembled contigs","quality":"Less is better","values":[0,0,1,0,0,0],"isMain":false},{"metricName":"Misassembled contigs length","quality":"Less is better","values":[0,0,5845,0,0,0],"isMain":true},{"metricName":"# possibly misassembled contigs","quality":"Less is better","values":[0,0,0,0,0,0],"isMain":false},{"metricName":" # possible misassemblies","quality":"Less is better","values":[0,0,0,0,0,0],"isMain":false},{"metricName":"# local misassemblies","quality":"Less is better","values":[0,0,2,0,0,0],"isMain":false},{"metricName":"# scaffold gap ext. mis.","quality":"Less is better","values":[0,0,0,0,0,0],"isMain":false},{"metricName":"# scaffold gap loc. mis.","quality":"Less is better","values":[0,0,0,0,0,0],"isMain":false},{"metricName":"# unaligned mis. contigs","quality":"Less is better","values":[0,0,0,0,1,0],"isMain":false}]],["Unaligned",[{"metricName":"# fully unaligned contigs","quality":"Less is better","values":[1,2,1,0,2,0],"isMain":false},{"metricName":"Fully unaligned length","quality":"Less is better","values":[579,1670,639,0,1063,0],"isMain":false},{"metricName":"# partially unaligned contigs","quality":"Less is better","values":[0,2,2,2,5,1],"isMain":false},{"metricName":"Partially unaligned length","quality":"Less is better","values":[0,1665,1798,3909,67312,838],"isMain":false}]],["Mismatches",[{"metricName":"# mismatches","quality":"Less is better","values":[183,151,412,296,268,70],"isMain":false},{"metricName":"# indels","quality":"Less is better","values":[0,2,5,2,3,0],"isMain":false},{"metricName":"Indels length","quality":"Less is better","values":[0,2,8,2,3,0],"isMain":false},{"metricName":"# mismatches per 100 kbp","quality":"Less is better","values":["1860.13","2424.53","2795.30","2494.31","2844.41","2250.08"],"isMain":true},{"metricName":"# indels per 100 kbp","quality":"Less is better","values":["0.00","32.11","33.92","16.85","31.84","0.00"],"isMain":true},{"metricName":" # indels (<= 5 bp)","quality":"Less is better","values":[0,2,5,2,3,0],"isMain":false},{"metricName":" # indels (> 5 bp)","quality":"Less is better","values":[0,0,0,0,0,0],"isMain":false},{"metricName":"# N's","quality":"Less is better","values":[0,0,0,0,0,0],"isMain":false},{"metricName":"# N's per 100 kbp","quality":"Less is better","values":["0.00","0.00","0.00","0.00","0.00","0.00"],"isMain":true}]],["Statistics without reference",[{"metricName":"# contigs","quality":"Equal","values":[18,17,22,21,22,6],"isMain":true},{"metricName":"# contigs (>= 1000 bp)","quality":"Equal","values":[1,2,3,2,4,1],"isMain":false},{"metricName":"# contigs (>= 5000 bp)","quality":"Equal","values":[0,0,1,1,1,0],"isMain":false},{"metricName":"# contigs (>= 10000 bp)","quality":"Equal","values":[0,0,0,0,1,0],"isMain":false},{"metricName":"# contigs (>= 25000 bp)","quality":"Equal","values":[0,0,0,0,1,0],"isMain":false},{"metricName":"# contigs (>= 50000 bp)","quality":"Equal","values":[0,0,0,0,1,0],"isMain":false},{"metricName":"Largest contig","quality":"More is better","values":[1550,1258,5845,6034,63805,1158],"isMain":true},{"metricName":"Total length","quality":"More is better","values":[12368,11873,19519,18601,79527,4157],"isMain":true},{"metricName":"Total length (>= 1000 bp)","quality":"More is better","values":[1550,2349,8237,7042,68121,1158],"isMain":true},{"metricName":"Total length (>= 5000 bp)","quality":"More is better","values":[0,0,5845,6034,63805,0],"isMain":false},{"metricName":"Total length (>= 10000 bp)","quality":"More is better","values":[0,0,0,0,63805,0],"isMain":true},{"metricName":"Total length (>= 25000 bp)","quality":"More is better","values":[0,0,0,0,63805,0],"isMain":false},{"metricName":"Total length (>= 50000 bp)","quality":"More is better","values":[0,0,0,0,63805,0],"isMain":true},{"metricName":"N50","quality":"More is better","values":[620,647,707,741,63805,619],"isMain":false},{"metricName":"N75","quality":"More is better","values":[562,579,558,552,63805,552],"isMain":false},{"metricName":"L50","quality":"Less is better","values":[7,7,5,5,1,3],"isMain":false},{"metricName":"L75","quality":"Less is better","values":[13,12,13,13,1,5],"isMain":false},{"metricName":"GC (%)","quality":"Equal","values":["58.13","58.65","55.78","54.53","57.88","56.46"],"isMain":false}]],["K-mer-based statistics",[]],["Predicted genes",[]],["Similarity statistics",[]],["Reference statistics",[{"metricName":"Reference length","quality":"Equal","values":[3985109,3985109,3985109,3985109,3985109,3985109],"isMain":false},{"metricName":"Reference fragments","quality":"Equal","values":[41,41,41,41,41,41],"isMain":false},{"metricName":"Reference GC (%)","quality":"Equal","values":["58.40","58.40","58.40","58.40","58.40","58.40"],"isMain":false}]]],"subreferences":[],"subreports":[],"minContig":500},{"date":"10 February 2020, Monday, 19:17:51","assembliesNames":["megahit-113-df","megahit-113-ml","megahit-113-ms"],"referenceName":"GCF\_000312405.1\_ASM31240v1","order":[0,1,2],"report":[["Genome statistics",[{"metricName":"Genome fraction (%)","quality":"More is better","values":[null,"0.029","0.008"],"isMain":true},{"metricName":"Duplication ratio","quality":"Less is better","values":[null,"1.901","1.000"],"isMain":true},{"metricName":"Largest alignment","quality":"More is better","values":[null,576,77],"isMain":true},{"metricName":"Total aligned length","quality":"More is better","values":[null,576,152],"isMain":true},{"metricName":"NA50","quality":"More is better","values":[null,576,null],"isMain":false},{"metricName":"NGA50","quality":"More is better","values":[null,null,null],"isMain":true},{"metricName":"LA50","quality":"Less is better","values":[null,1,null],"isMain":false}]],["Reads mapping",[]],["Misassemblies",[{"metricName":"# misassemblies","quality":"Less is better","values":[null,0,0],"isMain":true},{"metricName":" # relocations","quality":"Less is better","values":[null,0,0],"isMain":false},{"metricName":" # translocations","quality":"Less is better","values":[null,0,0],"isMain":false},{"metricName":" # inversions","quality":"Less is better","values":[null,0,0],"isMain":false},{"metricName":" # interspecies translocations","quality":"Less is better","values":[null,1,1],"isMain":false},{"metricName":"# misassembled contigs","quality":"Less is better","values":[null,0,0],"isMain":false},{"metricName":"Misassembled contigs length","quality":"Less is better","values":[null,0,0],"isMain":true},{"metricName":"# possibly misassembled contigs","quality":"Less is better","values":[null,0,0],"isMain":false},{"metricName":" # possible misassemblies","quality":"Less is better","values":[null,0,0],"isMain":false},{"metricName":"# local misassemblies","quality":"Less is better","values":[null,0,0],"isMain":false},{"metricName":"# scaffold gap ext. mis.","quality":"Less is better","values":[null,0,0],"isMain":false},{"metricName":"# scaffold gap loc. mis.","quality":"Less is better","values":[null,0,0],"isMain":false},{"metricName":"# unaligned mis. contigs","quality":"Less is better","values":[null,0,1],"isMain":false}]],["Unaligned",[{"metricName":"# fully unaligned contigs","quality":"Less is better","values":[2,0,0],"isMain":false},{"metricName":"Fully unaligned length","quality":"Less is better","values":[1316,0,0],"isMain":false},{"metricName":"# partially unaligned contigs","quality":"Less is better","values":[null,0,1],"isMain":false},{"metricName":"Partially unaligned length","quality":"Less is better","values":[null,0,913],"isMain":false}]],["Mismatches",[{"metricName":"# mismatches","quality":"Less is better","values":[null,47,3],"isMain":false},{"metricName":"# indels","quality":"Less is better","values":[null,5,0],"isMain":false},{"metricName":"Indels length","quality":"Less is better","values":[null,35,0],"isMain":false},{"metricName":"# mismatches per 100 kbp","quality":"Less is better","values":[null,"8159.72","1973.68"],"isMain":true},{"metricName":"# indels per 100 kbp","quality":"Less is better","values":[null,"868.06","0.00"],"isMain":true},{"metricName":" # indels (<= 5 bp)","quality":"Less is better","values":[null,4,0],"isMain":false},{"metricName":" # indels (> 5 bp)","quality":"Less is better","values":[null,1,0],"isMain":false},{"metricName":"# N's","quality":"Less is better","values":[0,0,0],"isMain":false},{"metricName":"# N's per 100 kbp","quality":"Less is better","values":["0.00","0.00","0.00"],"isMain":true}]],["Statistics without reference",[{"metricName":"# contigs","quality":"Equal","values":[2,1,1],"isMain":true},{"metricName":"# contigs (>= 1000 bp)","quality":"Equal","values":[0,1,1],"isMain":false},{"metricName":"# contigs (>= 5000 bp)","quality":"Equal","values":[0,0,0],"isMain":false},{"metricName":"# contigs (>= 10000 bp)","quality":"Equal","values":[0,0,0],"isMain":false},{"metricName":"# contigs (>= 25000 bp)","quality":"Equal","values":[0,0,0],"isMain":false},{"metricName":"# contigs (>= 50000 bp)","quality":"Equal","values":[0,0,0],"isMain":false},{"metricName":"Largest contig","quality":"More is better","values":[753,1095,1065],"isMain":true},{"metricName":"Total length","quality":"More is better","values":[1316,1095,1065],"isMain":true},{"metricName":"Total length (>= 1000 bp)","quality":"More is better","values":[0,1095,1065],"isMain":true},{"metricName":"Total length (>= 5000 bp)","quality":"More is better","values":[0,0,0],"isMain":false},{"metricName":"Total length (>= 10000 bp)","quality":"More is better","values":[0,0,0],"isMain":true},{"metricName":"Total length (>= 25000 bp)","quality":"More is better","values":[0,0,0],"isMain":false},{"metricName":"Total length (>= 50000 bp)","quality":"More is better","values":[0,0,0],"isMain":true},{"metricName":"N50","quality":"More is better","values":[753,1095,1065],"isMain":false},{"metricName":"N75","quality":"More is better","values":[563,1095,1065],"isMain":false},{"metricName":"L50","quality":"Less is better","values":[1,1,1],"isMain":false},{"metricName":"L75","quality":"Less is better","values":[2,1,1],"isMain":false},{"metricName":"GC (%)","quality":"Equal","values":["52.28","49.77","36.90"],"isMain":false}]],["K-mer-based statistics",[]],["Predicted genes",[]],["Similarity statistics",[]],["Reference statistics",[{"metricName":"Reference length","quality":"Equal","values":[1979644,1979644,1979644],"isMain":false},{"metricName":"Reference fragments","quality":"Equal","values":[40,40,40],"isMain":false},{"metricName":"Reference GC (%)","quality":"Equal","values":["50.90","50.90","50.90"],"isMain":false}]]],"subreferences":[],"subreports":[],"minContig":500},{"date":"10 February 2020, Monday, 19:43:48","assembliesNames":["megahit-103-df","megahit-113-df","megahit-113-ml","megahit-113-ms","megahit-129-df","metaSPAdes"],"referenceName":"GCF\_000318195.2\_ASM31819v2","order":[0,1,2,3,4,5],"report":[["Genome statistics",[{"metricName":"Genome fraction (%)","quality":"More is better","values":["98.193","98.852","98.529","98.802","99.150","98.251"],"isMain":true},{"metricName":"Duplication ratio","quality":"Less is better","values":["1.009","1.004","1.005","1.008","1.005","1.001"],"isMain":true},{"metricName":"Largest alignment","quality":"More is better","values":[60189,114033,114011,101981,114033,114033],"isMain":true},{"metricName":"Total aligned length","quality":"More is better","values":[2817199,2824216,2819580,2833686,2835263,2803461],"isMain":true},{"metricName":"NG50","quality":"More is better","values":[15446,40121,36438,49064,37359,35425],"isMain":false},{"metricName":"NG75","quality":"More is better","values":[9352,22294,21302,29829,21292,19191],"isMain":false},{"metricName":"NA50","quality":"More is better","values":[15446,36174,34727,25372,34893,34727],"isMain":false},{"metricName":"NA75","quality":"More is better","values":[9396,19593,18241,11133,19181,17490],"isMain":false},{"metricName":"NGA50","quality":"More is better","values":[15446,36174,35425,33692,35425,34727],"isMain":true},{"metricName":"NGA75","quality":"More is better","values":[9352,19866,19345,18820,19427,17584],"isMain":false},{"metricName":"LG50","quality":"Less is better","values":[54,24,25,20,24,26],"isMain":false},{"metricName":"LG75","quality":"Less is better","values":[114,48,51,38,49,53],"isMain":false},{"metricName":"LA50","quality":"Less is better","values":[54,25,28,36,27,27],"isMain":false},{"metricName":"LA75","quality":"Less is better","values":[112,52,58,82,55,57],"isMain":false},{"metricName":"LGA50","quality":"Less is better","values":[54,25,26,28,26,27],"isMain":true},{"metricName":"LGA75","quality":"Less is better","values":[114,51,54,57,53,56],"isMain":false}]],["Reads mapping",[]],["Misassemblies",[{"metricName":"# misassemblies","quality":"Less is better","values":[2,14,5,17,13,9],"isMain":true},{"metricName":" # relocations","quality":"Less is better","values":[0,1,0,0,0,1],"isMain":false},{"metricName":" # translocations","quality":"Less is better","values":[2,13,5,16,13,8],"isMain":false},{"metricName":" # inversions","quality":"Less is better","values":[0,0,0,1,0,0],"isMain":false},{"metricName":" # interspecies translocations","quality":"Less is better","values":[24,13,13,25,14,5],"isMain":false},{"metricName":"# misassembled contigs","quality":"Less is better","values":[2,12,5,17,13,8],"isMain":false},{"metricName":"Misassembled contigs length","quality":"Less is better","values":[5690,306999,31707,296953,226789,188223],"isMain":true},{"metricName":"# possibly misassembled contigs","quality":"Less is better","values":[0,0,0,0,0,0],"isMain":false},{"metricName":" # possible misassemblies","quality":"Less is better","values":[0,0,0,0,0,0],"isMain":false},{"metricName":"# local misassemblies","quality":"Less is better","values":[2,1,0,1,1,0],"isMain":false},{"metricName":"# scaffold gap ext. mis.","quality":"Less is better","values":[0,0,0,0,0,0],"isMain":false},{"metricName":"# scaffold gap loc. mis.","quality":"Less is better","values":[0,0,0,0,0,0],"isMain":false},{"metricName":"# unaligned mis. contigs","quality":"Less is better","values":[0,0,2,2,0,0],"isMain":false}]],["Unaligned",[{"metricName":"# fully unaligned contigs","quality":"Less is better","values":[0,0,0,0,0,0],"isMain":false},{"metricName":"Fully unaligned length","quality":"Less is better","values":[0,0,0,0,0,0],"isMain":false},{"metricName":"# partially unaligned contigs","quality":"Less is better","values":[7,7,10,15,9,2],"isMain":false},{"metricName":"Partially unaligned length","quality":"Less is better","values":[5703,38990,112109,501594,63840,66117],"isMain":false}]],["Mismatches",[{"metricName":"# mismatches","quality":"Less is better","values":[4949,565,607,1467,711,668],"isMain":false},{"metricName":"# indels","quality":"Less is better","values":[2,11,8,32,29,30],"isMain":false},{"metricName":"Indels length","quality":"Less is better","values":[2,110,141,391,447,60],"isMain":false},{"metricName":"# mismatches per 100 kbp","quality":"Less is better","values":["176.75","20.04","21.60","52.07","25.15","23.84"],"isMain":true},{"metricName":"# indels per 100 kbp","quality":"Less is better","values":["0.07","0.39","0.28","1.14","1.03","1.07"],"isMain":true},{"metricName":" # indels (<= 5 bp)","quality":"Less is better","values":[2,8,4,25,19,28],"isMain":false},{"metricName":" # indels (> 5 bp)","quality":"Less is better","values":[0,3,4,7,10,2],"isMain":false},{"metricName":"# N's","quality":"Less is better","values":[0,0,0,0,0,0],"isMain":false},{"metricName":"# N's per 100 kbp","quality":"Less is better","values":["0.00","0.00","0.00","0.00","0.00","0.00"],"isMain":true}]],["Statistics without reference",[{"metricName":"# contigs","quality":"Equal","values":[327,160,178,182,168,163],"isMain":true},{"metricName":"# contigs (>= 1000 bp)","quality":"Equal","values":[258,124,133,136,127,143],"isMain":false},{"metricName":"# contigs (>= 5000 bp)","quality":"Equal","values":[169,101,107,109,103,106],"isMain":false},{"metricName":"# contigs (>= 10000 bp)","quality":"Equal","values":[103,78,84,90,81,83],"isMain":false},{"metricName":"# contigs (>= 25000 bp)","quality":"Equal","values":[21,40,39,43,39,39],"isMain":false},{"metricName":"# contigs (>= 50000 bp)","quality":"Equal","values":[3,14,14,18,15,13],"isMain":false},{"metricName":"Largest contig","quality":"More is better","values":[60216,114033,114011,128298,114033,114033],"isMain":true},{"metricName":"Total length","quality":"More is better","values":[2832037,2868128,2935283,3341688,2904102,2871180],"isMain":true},{"metricName":"Total length (>= 1000 bp)","quality":"More is better","values":[2786315,2843131,2904207,3310838,2874701,2856808],"isMain":true},{"metricName":"Total length (>= 5000 bp)","quality":"More is better","values":[2545641,2788906,2832987,3243788,2814527,2754497],"isMain":false},{"metricName":"Total length (>= 10000 bp)","quality":"More is better","values":[2037967,2607300,2651704,3088868,2638916,2574857],"isMain":true},{"metricName":"Total length (>= 25000 bp)","quality":"More is better","values":[772076,1948370,1860355,2272406,1911142,1837436],"isMain":false},{"metricName":"Total length (>= 50000 bp)","quality":"More is better","values":[161174,981577,958584,1336481,1033886,909449],"isMain":true},{"metricName":"N50","quality":"More is better","values":[15446,37875,35429,41257,36479,35425],"isMain":false},{"metricName":"N75","quality":"More is better","values":[9396,21711,19634,22210,20238,19179],"isMain":false},{"metricName":"L50","quality":"Less is better","values":[54,25,27,26,25,26],"isMain":false},{"metricName":"L75","quality":"Less is better","values":[112,49,54,53,51,54],"isMain":false},{"metricName":"GC (%)","quality":"Equal","values":["43.12","43.18","43.17","43.91","43.14","43.10"],"isMain":false}]],["K-mer-based statistics",[]],["Predicted genes",[]],["Similarity statistics",[]],["Reference statistics",[{"metricName":"Reference length","quality":"Equal","values":[2851590,2851590,2851590,2851590,2851590,2851590],"isMain":false},{"metricName":"Reference fragments","quality":"Equal","values":[166,166,166,166,166,166],"isMain":false},{"metricName":"Reference GC (%)","quality":"Equal","values":["43.16","43.16","43.16","43.16","43.16","43.16"],"isMain":false}]]],"subreferences":[],"subreports":[],"minContig":500},{"date":"10 February 2020, Monday, 19:44:09","assembliesNames":["megahit-103-df","megahit-113-df","megahit-113-ml","megahit-113-ms","megahit-129-df","metaSPAdes"],"referenceName":"GCF\_000331715.1\_ASM33171v1","order":[0,1,2,3,4,5],"report":[["Genome statistics",[{"metricName":"Genome fraction (%)","quality":"More is better","values":["0.198","0.183","0.474","0.727","0.244","0.136"],"isMain":true},{"metricName":"Duplication ratio","quality":"Less is better","values":["1.353","1.277","1.014","1.068","1.019","1.269"],"isMain":true},{"metricName":"Largest alignment","quality":"More is better","values":[630,523,661,754,731,612],"isMain":true},{"metricName":"Total aligned length","quality":"More is better","values":[3397,3153,8142,12667,4193,2342],"isMain":true},{"metricName":"NA50","quality":"More is better","values":[497,276,515,511,525,269],"isMain":false},{"metricName":"NA75","quality":"More is better","values":[null,null,505,250,487,null],"isMain":false},{"metricName":"NGA50","quality":"More is better","values":[null,null,null,null,null,null],"isMain":true},{"metricName":"LA50","quality":"Less is better","values":[5,6,8,15,5,4],"isMain":false},{"metricName":"LA75","quality":"Less is better","values":[null,null,12,24,7,null],"isMain":false}]],["Reads mapping",[]],["Misassemblies",[{"metricName":"# misassemblies","quality":"Less is better","values":[0,0,0,0,0,0],"isMain":true},{"metricName":" # relocations","quality":"Less is better","values":[0,0,0,0,0,0],"isMain":false},{"metricName":" # translocations","quality":"Less is better","values":[0,0,0,0,0,0],"isMain":false},{"metricName":" # inversions","quality":"Less is better","values":[0,0,0,0,0,0],"isMain":false},{"metricName":" # interspecies translocations","quality":"Less is better","values":[3,3,0,7,1,2],"isMain":false},{"metricName":"# misassembled contigs","quality":"Less is better","values":[0,0,0,0,0,0],"isMain":false},{"metricName":"Misassembled contigs length","quality":"Less is better","values":[0,0,0,0,0,0],"isMain":true},{"metricName":"# possibly misassembled contigs","quality":"Less is better","values":[0,0,0,0,0,0],"isMain":false},{"metricName":" # possible misassemblies","quality":"Less is better","values":[0,0,0,0,0,0],"isMain":false},{"metricName":"# local misassemblies","quality":"Less is better","values":[0,0,0,1,0,0],"isMain":false},{"metricName":"# scaffold gap ext. mis.","quality":"Less is better","values":[0,0,0,0,0,0],"isMain":false},{"metricName":"# scaffold gap loc. mis.","quality":"Less is better","values":[0,0,0,0,0,0],"isMain":false},{"metricName":"# unaligned mis. contigs","quality":"Less is better","values":[0,0,0,0,0,0],"isMain":false}]],["Unaligned",[{"metricName":"# fully unaligned contigs","quality":"Less is better","values":[0,0,0,0,0,0],"isMain":false},{"metricName":"Fully unaligned length","quality":"Less is better","values":[0,0,0,0,0,0],"isMain":false},{"metricName":"# partially unaligned contigs","quality":"Less is better","values":[0,1,0,3,1,1],"isMain":false},{"metricName":"Partially unaligned length","quality":"Less is better","values":[0,1348,0,3261,918,527],"isMain":false}]],["Mismatches",[{"metricName":"# mismatches","quality":"Less is better","values":[57,61,157,315,79,49],"isMain":false},{"metricName":"# indels","quality":"Less is better","values":[0,0,0,0,0,0],"isMain":false},{"metricName":"Indels length","quality":"Less is better","values":[0,0,0,0,0,0],"isMain":false},{"metricName":"# mismatches per 100 kbp","quality":"Less is better","values":["1677.95","1934.67","1928.27","2520.60","1884.09","2092.23"],"isMain":true},{"metricName":"# indels per 100 kbp","quality":"Less is better","values":["0.00","0.00","0.00","0.00","0.00","0.00"],"isMain":true},{"metricName":" # indels (<= 5 bp)","quality":"Less is better","values":[0,0,0,0,0,0],"isMain":false},{"metricName":" # indels (> 5 bp)","quality":"Less is better","values":[0,0,0,0,0,0],"isMain":false},{"metricName":"# N's","quality":"Less is better","values":[0,0,0,0,0,0],"isMain":false},{"metricName":"# N's per 100 kbp","quality":"Less is better","values":["0.00","0.00","0.00","0.00","0.00","0.00"],"isMain":true}]],["Statistics without reference",[{"metricName":"# contigs","quality":"Equal","values":[8,8,15,25,8,6],"isMain":true},{"metricName":"# contigs (>= 1000 bp)","quality":"Equal","values":[0,1,0,3,1,0],"isMain":false},{"metricName":"# contigs (>= 5000 bp)","quality":"Equal","values":[0,0,0,0,0,0],"isMain":false},{"metricName":"# contigs (>= 10000 bp)","quality":"Equal","values":[0,0,0,0,0,0],"isMain":false},{"metricName":"# contigs (>= 25000 bp)","quality":"Equal","values":[0,0,0,0,0,0],"isMain":false},{"metricName":"# contigs (>= 50000 bp)","quality":"Equal","values":[0,0,0,0,0,0],"isMain":false},{"metricName":"Largest contig","quality":"More is better","values":[664,1624,663,1871,1156,736],"isMain":true},{"metricName":"Total length","quality":"More is better","values":[4596,5374,8255,16609,5191,3499],"isMain":true},{"metricName":"Total length (>= 1000 bp)","quality":"More is better","values":[0,1624,0,4201,1156,0],"isMain":true},{"metricName":"Total length (>= 5000 bp)","quality":"More is better","values":[0,0,0,0,0,0],"isMain":false},{"metricName":"Total length (>= 10000 bp)","quality":"More is better","values":[0,0,0,0,0,0],"isMain":true},{"metricName":"Total length (>= 25000 bp)","quality":"More is better","values":[0,0,0,0,0,0],"isMain":false},{"metricName":"Total length (>= 50000 bp)","quality":"More is better","values":[0,0,0,0,0,0],"isMain":true},{"metricName":"N50","quality":"More is better","values":[622,544,536,569,569,564],"isMain":false},{"metricName":"N75","quality":"More is better","values":[508,513,512,530,541,506],"isMain":false},{"metricName":"L50","quality":"Less is better","values":[4,3,7,10,4,3],"isMain":false},{"metricName":"L75","quality":"Less is better","values":[6,6,11,17,6,5],"isMain":false},{"metricName":"GC (%)","quality":"Equal","values":["32.16","36.29","34.39","37.77","35.47","33.21"],"isMain":false}]],["K-mer-based statistics",[]],["Predicted genes",[]],["Similarity statistics",[]],["Reference statistics",[{"metricName":"Reference length","quality":"Equal","values":[1719192,1719192,1719192,1719192,1719192,1719192],"isMain":false},{"metricName":"Reference fragments","quality":"Equal","values":[4,4,4,4,4,4],"isMain":false},{"metricName":"Reference GC (%)","quality":"Equal","values":["33.08","33.08","33.08","33.08","33.08","33.08"],"isMain":false}]]],"subreferences":[],"subreports":[],"minContig":500},{"date":"10 February 2020, Monday, 19:44:23","assembliesNames":["megahit-103-df","megahit-113-df","megahit-113-ml","megahit-113-ms","megahit-129-df","metaSPAdes"],"referenceName":"GCF\_000331995.1\_ASM33199v1","order":[0,1,2,3,4,5],"report":[["Genome statistics",[{"metricName":"Genome fraction (%)","quality":"More is better","values":["0.754","1.413","1.217","1.408","1.288","0.499"],"isMain":true},{"metricName":"Duplication ratio","quality":"Less is better","values":["1.074","1.052","1.008","1.029","1.034","1.046"],"isMain":true},{"metricName":"Largest alignment","quality":"More is better","values":[1479,1402,1300,1105,1109,1569],"isMain":true},{"metricName":"Total aligned length","quality":"More is better","values":[22415,42034,36210,41865,38318,14837],"isMain":true},{"metricName":"NA50","quality":"More is better","values":[575,613,584,574,579,544],"isMain":false},{"metricName":"NA75","quality":"More is better","values":[505,524,515,517,527,518],"isMain":false},{"metricName":"NGA50","quality":"More is better","values":[null,null,null,null,null,null],"isMain":true},{"metricName":"LA50","quality":"Less is better","values":[18,28,27,32,30,12],"isMain":false},{"metricName":"LA75","quality":"Less is better","values":[29,48,44,52,48,19],"isMain":false}]],["Reads mapping",[]],["Misassemblies",[{"metricName":"# misassemblies","quality":"Less is better","values":[0,0,0,0,0,0],"isMain":true},{"metricName":" # relocations","quality":"Less is better","values":[0,0,0,0,0,0],"isMain":false},{"metricName":" # translocations","quality":"Less is better","values":[0,0,0,0,0,0],"isMain":false},{"metricName":" # inversions","quality":"Less is better","values":[0,0,0,0,0,0],"isMain":false},{"metricName":" # interspecies translocations","quality":"Less is better","values":[4,6,0,5,5,2],"isMain":false},{"metricName":"# misassembled contigs","quality":"Less is better","values":[0,0,0,0,0,0],"isMain":false},{"metricName":"Misassembled contigs length","quality":"Less is better","values":[0,0,0,0,0,0],"isMain":true},{"metricName":"# possibly misassembled contigs","quality":"Less is better","values":[0,0,0,0,0,0],"isMain":false},{"metricName":" # possible misassemblies","quality":"Less is better","values":[0,0,0,0,0,0],"isMain":false},{"metricName":"# local misassemblies","quality":"Less is better","values":[0,0,0,0,0,0],"isMain":false},{"metricName":"# scaffold gap ext. mis.","quality":"Less is better","values":[0,0,0,0,0,0],"isMain":false},{"metricName":"# scaffold gap loc. mis.","quality":"Less is better","values":[0,0,0,0,0,0],"isMain":false},{"metricName":"# unaligned mis. contigs","quality":"Less is better","values":[0,0,0,0,0,0],"isMain":false}]],["Unaligned",[{"metricName":"# fully unaligned contigs","quality":"Less is better","values":[0,0,0,0,0,0],"isMain":false},{"metricName":"Fully unaligned length","quality":"Less is better","values":[0,0,0,0,0,0],"isMain":false},{"metricName":"# partially unaligned contigs","quality":"Less is better","values":[0,1,0,1,1,0],"isMain":false},{"metricName":"Partially unaligned length","quality":"Less is better","values":[0,520,0,772,1038,0],"isMain":false}]],["Mismatches",[{"metricName":"# mismatches","quality":"Less is better","values":[361,622,731,793,603,258],"isMain":false},{"metricName":"# indels","quality":"Less is better","values":[0,1,2,0,1,1],"isMain":false},{"metricName":"Indels length","quality":"Less is better","values":[0,1,2,0,1,1],"isMain":false},{"metricName":"# mismatches per 100 kbp","quality":"Less is better","values":["1610.53","1479.72","2018.78","1894.18","1573.63","1738.78"],"isMain":true},{"metricName":"# indels per 100 kbp","quality":"Less is better","values":["0.00","2.38","5.52","0.00","2.61","6.74"],"isMain":true},{"metricName":" # indels (<= 5 bp)","quality":"Less is better","values":[0,1,2,0,1,1],"isMain":false},{"metricName":" # indels (> 5 bp)","quality":"Less is better","values":[0,0,0,0,0,0],"isMain":false},{"metricName":"# N's","quality":"Less is better","values":[0,0,0,0,0,0],"isMain":false},{"metricName":"# N's per 100 kbp","quality":"Less is better","values":["0.00","0.00","0.00","0.00","0.00","0.00"],"isMain":true}]],["Statistics without reference",[{"metricName":"# contigs","quality":"Equal","values":[39,67,61,71,65,25],"isMain":true},{"metricName":"# contigs (>= 1000 bp)","quality":"Equal","values":[1,5,1,1,2,1],"isMain":false},{"metricName":"# contigs (>= 5000 bp)","quality":"Equal","values":[0,0,0,0,0,0],"isMain":false},{"metricName":"# contigs (>= 10000 bp)","quality":"Equal","values":[0,0,0,0,0,0],"isMain":false},{"metricName":"# contigs (>= 25000 bp)","quality":"Equal","values":[0,0,0,0,0,0],"isMain":false},{"metricName":"# contigs (>= 50000 bp)","quality":"Equal","values":[0,0,0,0,0,0],"isMain":false},{"metricName":"Largest contig","quality":"More is better","values":[1561,1402,1309,1108,1294,1733],"isMain":true},{"metricName":"Total length","quality":"More is better","values":[24065,44757,36503,43839,40663,15524],"isMain":true},{"metricName":"Total length (>= 1000 bp)","quality":"More is better","values":[1561,6386,1309,1108,2403,1733],"isMain":true},{"metricName":"Total length (>= 5000 bp)","quality":"More is better","values":[0,0,0,0,0,0],"isMain":false},{"metricName":"Total length (>= 10000 bp)","quality":"More is better","values":[0,0,0,0,0,0],"isMain":true},{"metricName":"Total length (>= 25000 bp)","quality":"More is better","values":[0,0,0,0,0,0],"isMain":false},{"metricName":"Total length (>= 50000 bp)","quality":"More is better","values":[0,0,0,0,0,0],"isMain":true},{"metricName":"N50","quality":"More is better","values":[587,639,588,594,602,552],"isMain":false},{"metricName":"N75","quality":"More is better","values":[542,548,521,539,544,540],"isMain":false},{"metricName":"L50","quality":"Less is better","values":[17,27,27,31,28,11],"isMain":false},{"metricName":"L75","quality":"Less is better","values":[28,46,44,50,46,18],"isMain":false},{"metricName":"GC (%)","quality":"Equal","values":["41.10","42.00","42.25","42.13","42.94","42.06"],"isMain":false}]],["K-mer-based statistics",[]],["Predicted genes",[]],["Similarity statistics",[]],["Reference statistics",[{"metricName":"Reference length","quality":"Equal","values":[2974370,2974370,2974370,2974370,2974370,2974370],"isMain":false},{"metricName":"Reference fragments","quality":"Equal","values":[4,4,4,4,4,4],"isMain":false},{"metricName":"Reference GC (%)","quality":"Equal","values":["42.24","42.24","42.24","42.24","42.24","42.24"],"isMain":false}]]],"subreferences":[],"subreports":[],"minContig":500},{"date":"10 February 2020, Monday, 19:44:24","assembliesNames":["megahit-103-df","megahit-113-df","megahit-113-ml","megahit-113-ms","megahit-129-df","metaSPAdes"],"referenceName":"GCF\_000332875.2\_ASM33287v2","order":[0,1,2,3,4,5],"report":[["Genome statistics",[{"metricName":"Genome fraction (%)","quality":"More is better","values":["1.274","3.228","3.026","3.366","1.677","1.692"],"isMain":true},{"metricName":"Duplication ratio","quality":"Less is better","values":["1.045","1.032","1.020","1.041","1.046","1.016"],"isMain":true},{"metricName":"Largest alignment","quality":"More is better","values":[5765,24354,6161,13180,6767,7656],"isMain":true},{"metricName":"Total aligned length","quality":"More is better","values":[35341,89881,83888,94073,46475,46918],"isMain":true},{"metricName":"NA50","quality":"More is better","values":[1810,4162,1345,1195,822,4265],"isMain":false},{"metricName":"NA75","quality":"More is better","values":[776,740,540,568,null,630],"isMain":false},{"metricName":"NGA50","quality":"More is better","values":[null,null,null,null,null,null],"isMain":true},{"metricName":"LA50","quality":"Less is better","values":[6,5,22,24,13,5],"isMain":false},{"metricName":"LA75","quality":"Less is better","values":[14,19,55,62,null,14],"isMain":false}]],["Reads mapping",[]],["Misassemblies",[{"metricName":"# misassemblies","quality":"Less is better","values":[6,7,9,6,7,4],"isMain":true},{"metricName":" # relocations","quality":"Less is better","values":[0,0,2,0,0,1],"isMain":false},{"metricName":" # translocations","quality":"Less is better","values":[6,7,7,5,7,3],"isMain":false},{"metricName":" # inversions","quality":"Less is better","values":[0,0,0,1,0,0],"isMain":false},{"metricName":" # interspecies translocations","quality":"Less is better","values":[7,11,13,19,12,6],"isMain":false},{"metricName":"# misassembled contigs","quality":"Less is better","values":[3,4,7,4,5,3],"isMain":false},{"metricName":"Misassembled contigs length","quality":"Less is better","values":[20498,54676,26196,30955,29400,15350],"isMain":true},{"metricName":"# possibly misassembled contigs","quality":"Less is better","values":[0,0,0,0,0,0],"isMain":false},{"metricName":" # possible misassemblies","quality":"Less is better","values":[0,0,0,0,0,0],"isMain":false},{"metricName":"# local misassemblies","quality":"Less is better","values":[0,2,1,0,1,0],"isMain":false},{"metricName":"# scaffold gap ext. mis.","quality":"Less is better","values":[0,0,0,0,0,0],"isMain":false},{"metricName":"# scaffold gap loc. mis.","quality":"Less is better","values":[0,0,0,0,0,0],"isMain":false},{"metricName":"# unaligned mis. contigs","quality":"Less is better","values":[0,0,0,1,1,1],"isMain":false}]],["Unaligned",[{"metricName":"# fully unaligned contigs","quality":"Less is better","values":[0,0,0,0,0,0],"isMain":false},{"metricName":"Fully unaligned length","quality":"Less is better","values":[0,0,0,0,0,0],"isMain":false},{"metricName":"# partially unaligned contigs","quality":"Less is better","values":[1,4,5,7,5,3],"isMain":false},{"metricName":"Partially unaligned length","quality":"Less is better","values":[1017,12495,15419,16716,16959,5968],"isMain":false}]],["Mismatches",[{"metricName":"# mismatches","quality":"Less is better","values":[453,1063,1354,1473,785,533],"isMain":false},{"metricName":"# indels","quality":"Less is better","values":[9,33,43,34,21,11],"isMain":false},{"metricName":"Indels length","quality":"Less is better","values":[15,71,114,57,42,100],"isMain":false},{"metricName":"# mismatches per 100 kbp","quality":"Less is better","values":["1282.49","1188.04","1614.60","1579.00","1688.54","1136.70"],"isMain":true},{"metricName":"# indels per 100 kbp","quality":"Less is better","values":["25.48","36.88","51.28","36.45","45.17","23.46"],"isMain":true},{"metricName":" # indels (<= 5 bp)","quality":"Less is better","values":[9,32,42,32,19,9],"isMain":false},{"metricName":" # indels (> 5 bp)","quality":"Less is better","values":[0,1,1,2,2,2],"isMain":false},{"metricName":"# N's","quality":"Less is better","values":[0,0,0,0,0,0],"isMain":false},{"metricName":"# N's per 100 kbp","quality":"Less is better","values":["0.00","0.00","0.00","0.00","0.00","0.00"],"isMain":true}]],["Statistics without reference",[{"metricName":"# contigs","quality":"Equal","values":[23,35,64,77,31,22],"isMain":true},{"metricName":"# contigs (>= 1000 bp)","quality":"Equal","values":[8,9,27,26,9,11],"isMain":false},{"metricName":"# contigs (>= 5000 bp)","quality":"Equal","values":[1,5,3,3,4,3],"isMain":false},{"metricName":"# contigs (>= 10000 bp)","quality":"Equal","values":[1,3,1,3,1,0],"isMain":false},{"metricName":"# contigs (>= 25000 bp)","quality":"Equal","values":[0,1,0,0,0,0],"isMain":false},{"metricName":"# contigs (>= 50000 bp)","quality":"Equal","values":[0,0,0,0,0,0],"isMain":false},{"metricName":"Largest contig","quality":"More is better","values":[12866,30272,11877,14212,14428,9725],"isMain":true},{"metricName":"Total length","quality":"More is better","values":[37946,104816,100977,113842,65585,53589],"isMain":true},{"metricName":"Total length (>= 1000 bp)","quality":"More is better","values":[28549,87964,77621,79463,51887,46917],"isMain":true},{"metricName":"Total length (>= 5000 bp)","quality":"More is better","values":[12866,74355,26205,36268,37262,25105],"isMain":false},{"metricName":"Total length (>= 10000 bp)","quality":"More is better","values":[12866,62590,11877,36268,14428,0],"isMain":true},{"metricName":"Total length (>= 25000 bp)","quality":"More is better","values":[0,30272,0,0,0,0],"isMain":false},{"metricName":"Total length (>= 50000 bp)","quality":"More is better","values":[0,0,0,0,0,0],"isMain":true},{"metricName":"N50","quality":"More is better","values":[2817,14338,2142,1603,7130,4570],"isMain":false},{"metricName":"N75","quality":"More is better","values":[1281,3956,1129,866,2448,2464],"isMain":false},{"metricName":"L50","quality":"Less is better","values":[3,3,11,10,4,4],"isMain":false},{"metricName":"L75","quality":"Less is better","values":[8,7,26,33,8,8],"isMain":false},{"metricName":"GC (%)","quality":"Equal","values":["37.46","35.86","35.85","36.72","36.85","36.83"],"isMain":false}]],["K-mer-based statistics",[]],["Predicted genes",[]],["Similarity statistics",[]],["Reference statistics",[{"metricName":"Reference length","quality":"Equal","values":[2771521,2771521,2771521,2771521,2771521,2771521],"isMain":false},{"metricName":"Reference fragments","quality":"Equal","values":[101,101,101,101,101,101],"isMain":false},{"metricName":"Reference GC (%)","quality":"Equal","values":["37.15","37.15","37.15","37.15","37.15","37.15"],"isMain":false}]]],"subreferences":[],"subreports":[],"minContig":500},{"date":"10 February 2020, Monday, 19:45:26","assembliesNames":["megahit-103-df","megahit-113-df","megahit-113-ml","megahit-113-ms","megahit-129-df","metaSPAdes"],"referenceName":"GCF\_000333275.1\_ASM33327v1","order":[0,1,2,3,4,5],"report":[["Genome statistics",[{"metricName":"Genome fraction (%)","quality":"More is better","values":["97.899","98.040","98.060","98.125","98.146","97.976"],"isMain":true},{"metricName":"Duplication ratio","quality":"Less is better","values":["1.004","1.004","1.008","1.006","1.004","1.002"],"isMain":true},{"metricName":"Largest alignment","quality":"More is better","values":[86757,127814,59314,91389,127817,127825],"isMain":true},{"metricName":"Total aligned length","quality":"More is better","values":[7214649,7221109,7257600,7241618,7231079,7206064],"isMain":true},{"metricName":"NG50","quality":"More is better","values":[14062,16463,12336,14364,16089,14856],"isMain":false},{"metricName":"NG75","quality":"More is better","values":[7212,8322,6688,7804,8236,7526],"isMain":false},{"metricName":"NA50","quality":"More is better","values":[13833,15602,11990,13834,15226,14856],"isMain":false},{"metricName":"NA75","quality":"More is better","values":[7212,8203,6490,7236,7963,7582],"isMain":false},{"metricName":"NGA50","quality":"More is better","values":[13532,15288,11918,13577,15010,14470],"isMain":true},{"metricName":"NGA75","quality":"More is better","values":[7001,7900,6331,7136,7811,7265],"isMain":false},{"metricName":"LG50","quality":"Less is better","values":[156,130,173,156,134,146],"isMain":false},{"metricName":"LG75","quality":"Less is better","values":[342,288,371,333,298,326],"isMain":false},{"metricName":"LA50","quality":"Less is better","values":[159,137,177,162,139,147],"isMain":false},{"metricName":"LA75","quality":"Less is better","values":[343,300,380,349,309,322],"isMain":false},{"metricName":"LGA50","quality":"Less is better","values":[164,140,180,164,142,152],"isMain":true},{"metricName":"LGA75","quality":"Less is better","values":[356,309,390,352,317,336],"isMain":false}]],["Reads mapping",[]],["Misassemblies",[{"metricName":"# misassemblies","quality":"Less is better","values":[41,51,56,52,56,21],"isMain":true},{"metricName":" # relocations","quality":"Less is better","values":[0,0,1,0,0,1],"isMain":false},{"metricName":" # translocations","quality":"Less is better","values":[41,51,55,52,56,20],"isMain":false},{"metricName":" # inversions","quality":"Less is better","values":[0,0,0,0,0,0],"isMain":false},{"metricName":" # interspecies translocations","quality":"Less is better","values":[6,11,1,16,10,6],"isMain":false},{"metricName":"# misassembled contigs","quality":"Less is better","values":[37,47,53,49,51,19],"isMain":false},{"metricName":"Misassembled contigs length","quality":"Less is better","values":[634045,1036621,694865,721417,964342,534616],"isMain":true},{"metricName":"# possibly misassembled contigs","quality":"Less is better","values":[0,0,0,0,0,0],"isMain":false},{"metricName":" # possible misassemblies","quality":"Less is better","values":[0,0,0,0,0,0],"isMain":false},{"metricName":"# local misassemblies","quality":"Less is better","values":[2,2,4,4,4,0],"isMain":false},{"metricName":"# scaffold gap ext. mis.","quality":"Less is better","values":[0,0,0,0,0,0],"isMain":false},{"metricName":"# scaffold gap loc. mis.","quality":"Less is better","values":[0,0,0,0,0,0],"isMain":false},{"metricName":"# unaligned mis. contigs","quality":"Less is better","values":[0,0,0,0,0,0],"isMain":false}]],["Unaligned",[{"metricName":"# fully unaligned contigs","quality":"Less is better","values":[0,0,0,1,0,0],"isMain":false},{"metricName":"Fully unaligned length","quality":"Less is better","values":[0,0,0,785,0,0],"isMain":false},{"metricName":"# partially unaligned contigs","quality":"Less is better","values":[3,7,0,10,9,1],"isMain":false},{"metricName":"Partially unaligned length","quality":"Less is better","values":[2297,20437,0,62081,20636,1507],"isMain":false}]],["Mismatches",[{"metricName":"# mismatches","quality":"Less is better","values":[3133,2517,8113,7811,2597,1133],"isMain":false},{"metricName":"# indels","quality":"Less is better","values":[14,36,30,64,61,22],"isMain":false},{"metricName":"Indels length","quality":"Less is better","values":[39,292,295,583,580,103],"isMain":false},{"metricName":"# mismatches per 100 kbp","quality":"Less is better","values":["43.57","34.95","112.64","108.38","36.02","15.74"],"isMain":true},{"metricName":"# indels per 100 kbp","quality":"Less is better","values":["0.19","0.50","0.42","0.89","0.85","0.31"],"isMain":true},{"metricName":" # indels (<= 5 bp)","quality":"Less is better","values":[13,31,23,53,40,21],"isMain":false},{"metricName":" # indels (> 5 bp)","quality":"Less is better","values":[1,5,7,11,21,1],"isMain":false},{"metricName":"# N's","quality":"Less is better","values":[0,0,0,0,0,0],"isMain":false},{"metricName":"# N's per 100 kbp","quality":"Less is better","values":["0.00","0.00","0.00","0.00","0.00","0.00"],"isMain":true}]],["Statistics without reference",[{"metricName":"# contigs","quality":"Equal","values":[902,802,971,886,813,852],"isMain":true},{"metricName":"# contigs (>= 1000 bp)","quality":"Equal","values":[823,737,887,809,746,803],"isMain":false},{"metricName":"# contigs (>= 5000 bp)","quality":"Equal","values":[459,426,474,470,442,451],"isMain":false},{"metricName":"# contigs (>= 10000 bp)","quality":"Equal","values":[239,233,240,246,233,228],"isMain":false},{"metricName":"# contigs (>= 25000 bp)","quality":"Equal","values":[40,57,41,47,54,46],"isMain":false},{"metricName":"# contigs (>= 50000 bp)","quality":"Equal","values":[7,10,2,2,10,8],"isMain":false},{"metricName":"Largest contig","quality":"More is better","values":[86757,127814,59314,113619,127817,127825],"isMain":true},{"metricName":"Total length","quality":"More is better","values":[7220742,7247588,7262958,7311757,7259999,7211338],"isMain":true},{"metricName":"Total length (>= 1000 bp)","quality":"More is better","values":[7162215,7200073,7200835,7252907,7209319,7172772],"isMain":true},{"metricName":"Total length (>= 5000 bp)","quality":"More is better","values":[6221844,6417073,6107662,6371990,6450509,6284019],"isMain":false},{"metricName":"Total length (>= 10000 bp)","quality":"More is better","values":[4641043,5008661,4430070,4750371,4924223,4666053],"isMain":true},{"metricName":"Total length (>= 25000 bp)","quality":"More is better","values":[1532044,2226544,1337159,1610447,2115413,1781220],"isMain":false},{"metricName":"Total length (>= 50000 bp)","quality":"More is better","values":[470239,739987,113273,186257,738535,585865],"isMain":true},{"metricName":"N50","quality":"More is better","values":[14351,16712,12484,14503,16444,15233],"isMain":false},{"metricName":"N75","quality":"More is better","values":[7544,8636,6879,7839,8417,7748],"isMain":false},{"metricName":"L50","quality":"Less is better","values":[152,127,169,155,131,142],"isMain":false},{"metricName":"L75","quality":"Less is better","values":[329,280,362,330,290,313],"isMain":false},{"metricName":"GC (%)","quality":"Equal","values":["67.53","67.51","67.53","67.41","67.50","67.55"],"isMain":false}]],["K-mer-based statistics",[]],["Predicted genes",[]],["Similarity statistics",[]],["Reference statistics",[{"metricName":"Reference length","quality":"Equal","values":[7345116,7345116,7345116,7345116,7345116,7345116],"isMain":false},{"metricName":"Reference fragments","quality":"Equal","values":[929,929,929,929,929,929],"isMain":false},{"metricName":"Reference GC (%)","quality":"Equal","values":["67.52","67.52","67.52","67.52","67.52","67.52"],"isMain":false}]]],"subreferences":[],"subreports":[],"minContig":500},{"date":"10 February 2020, Monday, 19:44:49","assembliesNames":["megahit-103-df","megahit-113-df","megahit-113-ml","megahit-113-ms","megahit-129-df","metaSPAdes"],"referenceName":"GCF\_000333595.1\_ASM33359v1","order":[0,1,2,3,4,5],"report":[["Genome statistics",[{"metricName":"Genome fraction (%)","quality":"More is better","values":["4.388","5.662","6.457","7.477","6.516","2.756"],"isMain":true},{"metricName":"Duplication ratio","quality":"Less is better","values":["1.035","1.031","1.015","1.029","1.025","1.054"],"isMain":true},{"metricName":"Largest alignment","quality":"More is better","values":[1193,1423,1081,1227,1423,979],"isMain":true},{"metricName":"Total aligned length","quality":"More is better","values":[318005,410633,468991,544251,472651,199936],"isMain":true},{"metricName":"NA50","quality":"More is better","values":[568,574,586,582,582,558],"isMain":false},{"metricName":"NA75","quality":"More is better","values":[515,519,535,524,524,517],"isMain":false},{"metricName":"NGA50","quality":"More is better","values":[null,null,null,null,null,null],"isMain":true},{"metricName":"LA50","quality":"Less is better","values":[253,324,341,421,348,164],"isMain":false},{"metricName":"LA75","quality":"Less is better","values":[412,529,553,686,574,263],"isMain":false}]],["Reads mapping",[]],["Misassemblies",[{"metricName":"# misassemblies","quality":"Less is better","values":[12,14,11,13,14,6],"isMain":true},{"metricName":" # relocations","quality":"Less is better","values":[1,1,1,1,1,1],"isMain":false},{"metricName":" # translocations","quality":"Less is better","values":[11,13,9,10,13,5],"isMain":false},{"metricName":" # inversions","quality":"Less is better","values":[0,0,1,2,0,0],"isMain":false},{"metricName":" # interspecies translocations","quality":"Less is better","values":[45,46,4,40,33,30],"isMain":false},{"metricName":"# misassembled contigs","quality":"Less is better","values":[12,14,10,13,14,6],"isMain":false},{"metricName":"Misassembled contigs length","quality":"Less is better","values":[7961,10178,7068,8898,11467,3817],"isMain":true},{"metricName":"# possibly misassembled contigs","quality":"Less is better","values":[0,0,0,0,0,0],"isMain":false},{"metricName":" # possible misassemblies","quality":"Less is better","values":[0,0,0,0,0,0],"isMain":false},{"metricName":"# local misassemblies","quality":"Less is better","values":[1,1,4,4,2,0],"isMain":false},{"metricName":"# scaffold gap ext. mis.","quality":"Less is better","values":[0,0,0,0,0,0],"isMain":false},{"metricName":"# scaffold gap loc. mis.","quality":"Less is better","values":[0,0,0,0,0,0],"isMain":false},{"metricName":"# unaligned mis. contigs","quality":"Less is better","values":[0,0,0,0,0,0],"isMain":false}]],["Unaligned",[{"metricName":"# fully unaligned contigs","quality":"Less is better","values":[0,0,1,0,0,0],"isMain":false},{"metricName":"Fully unaligned length","quality":"Less is better","values":[0,0,636,0,0,0],"isMain":false},{"metricName":"# partially unaligned contigs","quality":"Less is better","values":[10,16,0,14,10,3],"isMain":false},{"metricName":"Partially unaligned length","quality":"Less is better","values":[14879,24175,0,25229,14029,2290],"isMain":false}]],["Mismatches",[{"metricName":"# mismatches","quality":"Less is better","values":[5134,6466,9209,10922,7754,3510],"isMain":false},{"metricName":"# indels","quality":"Less is better","values":[13,15,20,34,17,6],"isMain":false},{"metricName":"Indels length","quality":"Less is better","values":[21,21,36,110,74,45],"isMain":false},{"metricName":"# mismatches per 100 kbp","quality":"Less is better","values":["1616.53","1577.73","1970.38","2018.24","1644.17","1759.62"],"isMain":true},{"metricName":"# indels per 100 kbp","quality":"Less is better","values":["4.09","3.66","4.28","6.28","3.60","3.01"],"isMain":true},{"metricName":" # indels (<= 5 bp)","quality":"Less is better","values":[13,15,19,32,15,4],"isMain":false},{"metricName":" # indels (> 5 bp)","quality":"Less is better","values":[0,0,1,2,2,2],"isMain":false},{"metricName":"# N's","quality":"Less is better","values":[0,0,0,0,0,0],"isMain":false},{"metricName":"# N's per 100 kbp","quality":"Less is better","values":["0.00","0.00","0.00","0.00","0.00","0.00"],"isMain":true}]],["Statistics without reference",[{"metricName":"# contigs","quality":"Equal","values":[544,693,773,910,779,354],"isMain":true},{"metricName":"# contigs (>= 1000 bp)","quality":"Equal","values":[11,20,10,21,23,1],"isMain":false},{"metricName":"# contigs (>= 5000 bp)","quality":"Equal","values":[0,0,0,1,0,0],"isMain":false},{"metricName":"# contigs (>= 10000 bp)","quality":"Equal","values":[0,0,0,1,0,0],"isMain":false},{"metricName":"# contigs (>= 25000 bp)","quality":"Equal","values":[0,0,0,0,0,0],"isMain":false},{"metricName":"# contigs (>= 50000 bp)","quality":"Equal","values":[0,0,0,0,0,0],"isMain":false},{"metricName":"Largest contig","quality":"More is better","values":[4144,3803,1094,12173,3079,1189],"isMain":true},{"metricName":"Total length","quality":"More is better","values":[343474,446826,474975,582236,497353,212589],"isMain":true},{"metricName":"Total length (>= 1000 bp)","quality":"More is better","values":[19798,34703,10528,38070,32335,1189],"isMain":true},{"metricName":"Total length (>= 5000 bp)","quality":"More is better","values":[0,0,0,12173,0,0],"isMain":false},{"metricName":"Total length (>= 10000 bp)","quality":"More is better","values":[0,0,0,12173,0,0],"isMain":true},{"metricName":"Total length (>= 25000 bp)","quality":"More is better","values":[0,0,0,0,0,0],"isMain":false},{"metricName":"Total length (>= 50000 bp)","quality":"More is better","values":[0,0,0,0,0,0],"isMain":true},{"metricName":"N50","quality":"More is better","values":[597,607,595,608,608,584],"isMain":false},{"metricName":"N75","quality":"More is better","values":[540,545,545,545,542,536],"isMain":false},{"metricName":"L50","quality":"Less is better","values":[227,284,336,378,322,156],"isMain":false},{"metricName":"L75","quality":"Less is better","values":[379,479,545,631,540,252],"isMain":false},{"metricName":"GC (%)","quality":"Equal","values":["65.86","66.01","66.73","66.26","66.28","65.74"],"isMain":false}]],["K-mer-based statistics",[]],["Predicted genes",[]],["Similarity statistics",[]],["Reference statistics",[{"metricName":"Reference length","quality":"Equal","values":[7237917,7237917,7237917,7237917,7237917,7237917],"isMain":false},{"metricName":"Reference fragments","quality":"Equal","values":[1289,1289,1289,1289,1289,1289],"isMain":false},{"metricName":"Reference GC (%)","quality":"Equal","values":["67.12","67.12","67.12","67.12","67.12","67.12"],"isMain":false}]]],"subreferences":[],"subreports":[],"minContig":500},{"date":"10 February 2020, Monday, 19:44:48","assembliesNames":["megahit-103-df","megahit-113-df","megahit-113-ml","megahit-113-ms","megahit-129-df","metaSPAdes"],"referenceName":"GCF\_000345685.2\_ASM34568v2","order":[0,1,2,3,4,5],"report":[["Genome statistics",[{"metricName":"Genome fraction (%)","quality":"More is better","values":["88.383","80.944","90.923","90.726","89.576","87.233"],"isMain":true},{"metricName":"Duplication ratio","quality":"Less is better","values":["1.091","1.146","1.163","1.175","1.140","1.027"],"isMain":true},{"metricName":"Largest alignment","quality":"More is better","values":[15165,19308,7009,8903,20394,11658],"isMain":true},{"metricName":"Total aligned length","quality":"More is better","values":[1591601,1531220,1742430,1753376,1684241,1479894],"isMain":true},{"metricName":"NG50","quality":"More is better","values":[1753,2034,1869,2220,2776,2629],"isMain":false},{"metricName":"NG75","quality":"More is better","values":[1059,1056,1274,1421,1710,1249],"isMain":false},{"metricName":"NA50","quality":"More is better","values":[1790,2101,1760,2036,2605,2903],"isMain":false},{"metricName":"NA75","quality":"More is better","values":[1104,1192,1123,1201,1526,1566],"isMain":false},{"metricName":"NGA50","quality":"More is better","values":[1719,1970,1839,2159,2675,2612],"isMain":true},{"metricName":"NGA75","quality":"More is better","values":[1027,1020,1246,1364,1646,1221],"isMain":false},{"metricName":"LG50","quality":"Less is better","values":[294,233,292,253,197,189],"isMain":false},{"metricName":"LG75","quality":"Less is better","values":[598,512,563,487,385,416],"isMain":false},{"metricName":"LA50","quality":"Less is better","values":[281,215,326,286,210,160],"isMain":false},{"metricName":"LA75","quality":"Less is better","values":[567,458,638,564,417,332],"isMain":false},{"metricName":"LGA50","quality":"Less is better","values":[297,241,298,258,202,191],"isMain":true},{"metricName":"LGA75","quality":"Less is better","values":[607,531,573,497,396,423],"isMain":false}]],["Reads mapping",[]],["Misassemblies",[{"metricName":"# misassemblies","quality":"Less is better","values":[8,9,11,13,13,8],"isMain":true},{"metricName":" # relocations","quality":"Less is better","values":[0,0,0,0,1,0],"isMain":false},{"metricName":" # translocations","quality":"Less is better","values":[5,5,10,10,10,8],"isMain":false},{"metricName":" # inversions","quality":"Less is better","values":[3,4,1,3,2,0],"isMain":false},{"metricName":" # interspecies translocations","quality":"Less is better","values":[6,17,18,27,14,6],"isMain":false},{"metricName":"# misassembled contigs","quality":"Less is better","values":[6,9,9,12,12,8],"isMain":false},{"metricName":"Misassembled contigs length","quality":"Less is better","values":[10220,17053,19548,31147,31490,27598],"isMain":true},{"metricName":"# possibly misassembled contigs","quality":"Less is better","values":[0,0,0,0,0,0],"isMain":false},{"metricName":" # possible misassemblies","quality":"Less is better","values":[0,0,0,0,0,0],"isMain":false},{"metricName":"# local misassemblies","quality":"Less is better","values":[2,2,3,5,3,1],"isMain":false},{"metricName":"# scaffold gap ext. mis.","quality":"Less is better","values":[0,0,0,0,0,0],"isMain":false},{"metricName":"# scaffold gap loc. mis.","quality":"Less is better","values":[0,0,0,0,0,0],"isMain":false},{"metricName":"# unaligned mis. contigs","quality":"Less is better","values":[0,1,1,0,0,0],"isMain":false}]],["Unaligned",[{"metricName":"# fully unaligned contigs","quality":"Less is better","values":[0,0,0,0,0,0],"isMain":false},{"metricName":"Fully unaligned length","quality":"Less is better","values":[0,0,0,0,0,0],"isMain":false},{"metricName":"# partially unaligned contigs","quality":"Less is better","values":[1,10,8,6,7,2],"isMain":false},{"metricName":"Partially unaligned length","quality":"Less is better","values":[3290,14238,8154,5645,10313,2254],"isMain":false}]],["Mismatches",[{"metricName":"# mismatches","quality":"Less is better","values":[11279,19913,21031,22074,15820,9164],"isMain":false},{"metricName":"# indels","quality":"Less is better","values":[156,397,157,237,216,207],"isMain":false},{"metricName":"Indels length","quality":"Less is better","values":[303,1110,291,526,640,501],"isMain":false},{"metricName":"# mismatches per 100 kbp","quality":"Less is better","values":["770.64","1485.61","1396.82","1469.27","1066.52","634.39"],"isMain":true},{"metricName":"# indels per 100 kbp","quality":"Less is better","values":["10.66","29.62","10.43","15.77","14.56","14.33"],"isMain":true},{"metricName":" # indels (<= 5 bp)","quality":"Less is better","values":[145,352,147,217,194,186],"isMain":false},{"metricName":" # indels (> 5 bp)","quality":"Less is better","values":[11,45,10,20,22,21],"isMain":false},{"metricName":"# N's","quality":"Less is better","values":[0,0,0,0,0,0],"isMain":false},{"metricName":"# N's per 100 kbp","quality":"Less is better","values":["0.00","0.00","0.00","0.00","0.00","0.00"],"isMain":true}]],["Statistics without reference",[{"metricName":"# contigs","quality":"Equal","values":[1075,933,1186,1088,847,707],"isMain":true},{"metricName":"# contigs (>= 1000 bp)","quality":"Equal","values":[625,541,725,691,598,492],"isMain":false},{"metricName":"# contigs (>= 5000 bp)","quality":"Equal","values":[16,31,13,18,38,50],"isMain":false},{"metricName":"# contigs (>= 10000 bp)","quality":"Equal","values":[1,2,0,0,2,6],"isMain":false},{"metricName":"# contigs (>= 25000 bp)","quality":"Equal","values":[0,0,0,0,0,0],"isMain":false},{"metricName":"# contigs (>= 50000 bp)","quality":"Equal","values":[0,0,0,0,0,0],"isMain":false},{"metricName":"Largest contig","quality":"More is better","values":[15165,19320,7022,8904,20394,11658],"isMain":true},{"metricName":"Total length","quality":"More is better","values":[1599477,1550208,1758889,1770986,1701002,1485459],"isMain":true},{"metricName":"Total length (>= 1000 bp)","quality":"More is better","values":[1269666,1272627,1426318,1483228,1521786,1326937],"isMain":true},{"metricName":"Total length (>= 5000 bp)","quality":"More is better","values":[116738,212853,77103,111822,263305,344765],"isMain":false},{"metricName":"Total length (>= 10000 bp)","quality":"More is better","values":[15165,34517,0,0,31525,67008],"isMain":true},{"metricName":"Total length (>= 25000 bp)","quality":"More is better","values":[0,0,0,0,0,0],"isMain":false},{"metricName":"Total length (>= 50000 bp)","quality":"More is better","values":[0,0,0,0,0,0],"isMain":true},{"metricName":"N50","quality":"More is better","values":[1800,2194,1803,2061,2734,2941],"isMain":false},{"metricName":"N75","quality":"More is better","values":[1122,1246,1165,1260,1601,1586],"isMain":false},{"metricName":"L50","quality":"Less is better","values":[278,208,320,280,205,158],"isMain":false},{"metricName":"L75","quality":"Less is better","values":[560,442,626,552,405,327],"isMain":false},{"metricName":"GC (%)","quality":"Equal","values":["39.37","39.26","39.50","39.45","39.38","39.37"],"isMain":false}]],["K-mer-based statistics",[]],["Predicted genes",[]],["Similarity statistics",[]],["Reference statistics",[{"metricName":"Reference length","quality":"Equal","values":[1655951,1655951,1655951,1655951,1655951,1655951],"isMain":false},{"metricName":"Reference fragments","quality":"Equal","values":[111,111,111,111,111,111],"isMain":false},{"metricName":"Reference GC (%)","quality":"Equal","values":["39.37","39.37","39.37","39.37","39.37","39.37"],"isMain":false}]]],"subreferences":[],"subreports":[],"minContig":500},{"date":"10 February 2020, Monday, 19:45:03","assembliesNames":["megahit-103-df","megahit-113-df","megahit-113-ml","megahit-113-ms","megahit-129-df","metaSPAdes"],"referenceName":"GCF\_000365025.1\_ASM36502v1","order":[0,1,2,3,4,5],"report":[["Genome statistics",[{"metricName":"Genome fraction (%)","quality":"More is better","values":["99.891","99.874","99.796","99.889","99.901","99.816"],"isMain":true},{"metricName":"Duplication ratio","quality":"Less is better","values":["1.007","1.002","1.037","1.028","1.002","1.003"],"isMain":true},{"metricName":"Largest alignment","quality":"More is better","values":[132303,280577,34404,41059,271986,210622],"isMain":true},{"metricName":"Total aligned length","quality":"More is better","values":[2783520,2769527,2861380,2840189,2772130,2771208],"isMain":true},{"metricName":"NG50","quality":"More is better","values":[34593,133713,8967,10864,130439,65826],"isMain":false},{"metricName":"NG75","quality":"More is better","values":[18230,63611,5368,6699,83353,45632],"isMain":false},{"metricName":"NA50","quality":"More is better","values":[34110,105413,8547,10247,119635,65826],"isMain":false},{"metricName":"NA75","quality":"More is better","values":[17878,58509,4979,6341,68810,45632],"isMain":false},{"metricName":"NGA50","quality":"More is better","values":[34167,105413,8938,10538,119635,65826],"isMain":true},{"metricName":"NGA75","quality":"More is better","values":[17897,58509,5354,6680,68810,45632],"isMain":false},{"metricName":"LG50","quality":"Less is better","values":[24,8,96,80,8,13],"isMain":false},{"metricName":"LG75","quality":"Less is better","values":[50,16,198,161,14,26],"isMain":false},{"metricName":"LA50","quality":"Less is better","values":[26,9,102,84,8,13],"isMain":false},{"metricName":"LA75","quality":"Less is better","values":[53,18,212,171,15,26],"isMain":false},{"metricName":"LGA50","quality":"Less is better","values":[25,9,96,81,8,13],"isMain":true},{"metricName":"LGA75","quality":"Less is better","values":[52,18,198,162,15,26],"isMain":false}]],["Reads mapping",[]],["Misassemblies",[{"metricName":"# misassemblies","quality":"Less is better","values":[4,6,4,7,7,3],"isMain":true},{"metricName":" # relocations","quality":"Less is better","values":[0,1,0,0,1,0],"isMain":false},{"metricName":" # translocations","quality":"Less is better","values":[4,5,3,7,6,3],"isMain":false},{"metricName":" # inversions","quality":"Less is better","values":[0,0,1,0,0,0],"isMain":false},{"metricName":" # interspecies translocations","quality":"Less is better","values":[0,1,2,5,1,0],"isMain":false},{"metricName":"# misassembled contigs","quality":"Less is better","values":[3,5,4,6,7,2],"isMain":false},{"metricName":"Misassembled contigs length","quality":"Less is better","values":[121252,669709,32628,53195,686205,131519],"isMain":true},{"metricName":"# possibly misassembled contigs","quality":"Less is better","values":[0,0,0,0,0,0],"isMain":false},{"metricName":" # possible misassemblies","quality":"Less is better","values":[0,0,0,0,0,0],"isMain":false},{"metricName":"# local misassemblies","quality":"Less is better","values":[0,0,0,0,0,0],"isMain":false},{"metricName":"# scaffold gap ext. mis.","quality":"Less is better","values":[0,0,0,0,0,0],"isMain":false},{"metricName":"# scaffold gap loc. mis.","quality":"Less is better","values":[0,0,0,0,0,0],"isMain":false},{"metricName":"# unaligned mis. contigs","quality":"Less is better","values":[0,0,0,0,0,0],"isMain":false}]],["Unaligned",[{"metricName":"# fully unaligned contigs","quality":"Less is better","values":[0,0,0,0,0,0],"isMain":false},{"metricName":"Fully unaligned length","quality":"Less is better","values":[0,0,0,0,0,0],"isMain":false},{"metricName":"# partially unaligned contigs","quality":"Less is better","values":[0,1,0,3,1,0],"isMain":false},{"metricName":"Partially unaligned length","quality":"Less is better","values":[0,530,0,2014,834,0],"isMain":false}]],["Mismatches",[{"metricName":"# mismatches","quality":"Less is better","values":[1685,679,11103,9755,611,526],"isMain":false},{"metricName":"# indels","quality":"Less is better","values":[0,0,9,11,10,3],"isMain":false},{"metricName":"Indels length","quality":"Less is better","values":[0,0,11,28,50,3],"isMain":false},{"metricName":"# mismatches per 100 kbp","quality":"Less is better","values":["60.92","24.55","401.82","352.71","22.09","19.03"],"isMain":true},{"metricName":"# indels per 100 kbp","quality":"Less is better","values":["0.00","0.00","0.33","0.40","0.36","0.11"],"isMain":true},{"metricName":" # indels (<= 5 bp)","quality":"Less is better","values":[0,0,9,10,8,3],"isMain":false},{"metricName":" # indels (> 5 bp)","quality":"Less is better","values":[0,0,0,1,2,0],"isMain":false},{"metricName":"# N's","quality":"Less is better","values":[0,0,0,0,0,0],"isMain":false},{"metricName":"# N's per 100 kbp","quality":"Less is better","values":["0.00","0.00","0.00","0.00","0.00","0.00"],"isMain":true}]],["Statistics without reference",[{"metricName":"# contigs","quality":"Equal","values":[130,42,501,389,38,72],"isMain":true},{"metricName":"# contigs (>= 1000 bp)","quality":"Equal","values":[128,40,462,368,37,71],"isMain":false},{"metricName":"# contigs (>= 5000 bp)","quality":"Equal","values":[103,36,210,216,33,59],"isMain":false},{"metricName":"# contigs (>= 10000 bp)","quality":"Equal","values":[84,34,79,90,31,51],"isMain":false},{"metricName":"# contigs (>= 25000 bp)","quality":"Equal","values":[38,26,4,13,25,35],"isMain":false},{"metricName":"# contigs (>= 50000 bp)","quality":"Equal","values":[15,22,0,0,20,23],"isMain":false},{"metricName":"Largest contig","quality":"More is better","values":[132303,280577,34404,41066,271997,210622],"isMain":true},{"metricName":"Total length","quality":"More is better","values":[2784487,2770623,2864581,2846160,2773429,2771821],"isMain":true},{"metricName":"Total length (>= 1000 bp)","quality":"More is better","values":[2782555,2769488,2837079,2830449,2772537,2771120],"isMain":true},{"metricName":"Total length (>= 5000 bp)","quality":"More is better","values":[2711527,2760778,2142725,2403765,2764125,2739018],"isMain":false},{"metricName":"Total length (>= 10000 bp)","quality":"More is better","values":[2561876,2744326,1228197,1490531,2748580,2679486],"isMain":true},{"metricName":"Total length (>= 25000 bp)","quality":"More is better","values":[1815281,2604176,125402,373382,2653029,2415834],"isMain":false},{"metricName":"Total length (>= 50000 bp)","quality":"More is better","values":[1025764,2452926,0,0,2479348,1968920],"isMain":true},{"metricName":"N50","quality":"More is better","values":[34167,133713,8547,10309,130439,65826],"isMain":false},{"metricName":"N75","quality":"More is better","values":[18116,63611,4979,6451,83353,45632],"isMain":false},{"metricName":"L50","quality":"Less is better","values":[25,8,102,84,8,13],"isMain":false},{"metricName":"L75","quality":"Less is better","values":[51,16,212,170,14,26],"isMain":false},{"metricName":"GC (%)","quality":"Equal","values":["52.47","52.48","52.46","52.45","52.48","52.48"],"isMain":false}]],["K-mer-based statistics",[]],["Predicted genes",[]],["Similarity statistics",[]],["Reference statistics",[{"metricName":"Reference length","quality":"Equal","values":[2768821,2768821,2768821,2768821,2768821,2768821],"isMain":false},{"metricName":"Reference fragments","quality":"Equal","values":[6,6,6,6,6,6],"isMain":false},{"metricName":"Reference GC (%)","quality":"Equal","values":["52.48","52.48","52.48","52.48","52.48","52.48"],"isMain":false}]]],"subreferences":[],"subreports":[],"minContig":500},{"date":"10 February 2020, Monday, 19:45:11","assembliesNames":["megahit-103-df","megahit-113-df","megahit-113-ml","megahit-113-ms","megahit-129-df","metaSPAdes"],"referenceName":"GCF\_000371405.1\_Clos\_clos\_CM201\_V1","order":[0,1,2,3,4,5],"report":[["Genome statistics",[{"metricName":"Genome fraction (%)","quality":"More is better","values":["17.774","12.595","12.388","14.279","12.992","16.927"],"isMain":true},{"metricName":"Duplication ratio","quality":"Less is better","values":["1.006","1.009","1.012","1.010","1.011","1.002"],"isMain":true},{"metricName":"Largest alignment","quality":"More is better","values":[22103,30919,45370,46371,54659,109581],"isMain":true},{"metricName":"Total aligned length","quality":"More is better","values":[997092,707800,697491,802456,729146,946416],"isMain":true},{"metricName":"NA50","quality":"More is better","values":[10267,8801,7630,10670,11227,21509],"isMain":false},{"metricName":"NA75","quality":"More is better","values":[5594,4522,4370,5159,5368,11206],"isMain":false},{"metricName":"NGA50","quality":"More is better","values":[null,null,null,null,null,null],"isMain":true},{"metricName":"LA50","quality":"Less is better","values":[33,25,25,23,19,12],"isMain":false},{"metricName":"LA75","quality":"Less is better","values":[65,52,54,51,40,29],"isMain":false}]],["Reads mapping",[]],["Misassemblies",[{"metricName":"# misassemblies","quality":"Less is better","values":[1,2,1,7,2,0],"isMain":true},{"metricName":" # relocations","quality":"Less is better","values":[0,1,0,1,2,0],"isMain":false},{"metricName":" # translocations","quality":"Less is better","values":[1,1,1,6,0,0],"isMain":false},{"metricName":" # inversions","quality":"Less is better","values":[0,0,0,0,0,0],"isMain":false},{"metricName":" # interspecies translocations","quality":"Less is better","values":[3,9,9,15,14,0],"isMain":false},{"metricName":"# misassembled contigs","quality":"Less is better","values":[1,2,1,5,2,0],"isMain":false},{"metricName":"Misassembled contigs length","quality":"Less is better","values":[9105,25401,3589,26051,17105,0],"isMain":true},{"metricName":"# possibly misassembled contigs","quality":"Less is better","values":[0,0,1,1,0,0],"isMain":false},{"metricName":" # possible misassemblies","quality":"Less is better","values":[0,0,1,1,0,0],"isMain":false},{"metricName":"# local misassemblies","quality":"Less is better","values":[2,0,2,3,7,0],"isMain":false},{"metricName":"# scaffold gap ext. mis.","quality":"Less is better","values":[0,0,0,0,0,0],"isMain":false},{"metricName":"# scaffold gap loc. mis.","quality":"Less is better","values":[0,0,0,0,0,0],"isMain":false},{"metricName":"# unaligned mis. contigs","quality":"Less is better","values":[0,0,1,1,0,0],"isMain":false}]],["Unaligned",[{"metricName":"# fully unaligned contigs","quality":"Less is better","values":[0,0,0,0,1,0],"isMain":false},{"metricName":"Fully unaligned length","quality":"Less is better","values":[0,0,0,0,583,0],"isMain":false},{"metricName":"# partially unaligned contigs","quality":"Less is better","values":[0,7,5,5,8,0],"isMain":false},{"metricName":"Partially unaligned length","quality":"Less is better","values":[0,8386,19507,7961,9700,0],"isMain":false}]],["Mismatches",[{"metricName":"# mismatches","quality":"Less is better","values":[2323,3094,2841,3623,2405,554],"isMain":false},{"metricName":"# indels","quality":"Less is better","values":[42,42,27,40,37,22],"isMain":false},{"metricName":"Indels length","quality":"Less is better","values":[237,134,79,195,120,68],"isMain":false},{"metricName":"# mismatches per 100 kbp","quality":"Less is better","values":["233.97","439.74","410.55","454.22","331.38","58.59"],"isMain":true},{"metricName":"# indels per 100 kbp","quality":"Less is better","values":["4.23","5.97","3.90","5.01","5.10","2.33"],"isMain":true},{"metricName":" # indels (<= 5 bp)","quality":"Less is better","values":[35,39,22,34,31,20],"isMain":false},{"metricName":" # indels (> 5 bp)","quality":"Less is better","values":[7,3,5,6,6,2],"isMain":false},{"metricName":"# N's","quality":"Less is better","values":[0,0,0,0,0,0],"isMain":false},{"metricName":"# N's per 100 kbp","quality":"Less is better","values":["0.00","0.00","0.00","0.00","0.00","0.00"],"isMain":true}]],["Statistics without reference",[{"metricName":"# contigs","quality":"Equal","values":[176,138,136,145,122,88],"isMain":true},{"metricName":"# contigs (>= 1000 bp)","quality":"Equal","values":[147,109,116,111,92,73],"isMain":false},{"metricName":"# contigs (>= 5000 bp)","quality":"Equal","values":[73,51,50,53,46,48],"isMain":false},{"metricName":"# contigs (>= 10000 bp)","quality":"Equal","values":[35,21,17,25,24,33],"isMain":false},{"metricName":"# contigs (>= 25000 bp)","quality":"Equal","values":[0,1,2,2,5,10],"isMain":false},{"metricName":"# contigs (>= 50000 bp)","quality":"Equal","values":[0,0,0,0,1,2],"isMain":false},{"metricName":"Largest contig","quality":"More is better","values":[22105,30919,45370,46378,54659,109581],"isMain":true},{"metricName":"Total length","quality":"More is better","values":[998487,718351,719514,813862,743772,947067],"isMain":true},{"metricName":"Total length (>= 1000 bp)","quality":"More is better","values":[977838,699132,706148,789788,723486,936422],"isMain":true},{"metricName":"Total length (>= 5000 bp)","quality":"More is better","values":[795061,542621,531116,627652,597924,871660],"isMain":false},{"metricName":"Total length (>= 10000 bp)","quality":"More is better","values":[524731,333992,300505,432422,438292,758765],"isMain":true},{"metricName":"Total length (>= 25000 bp)","quality":"More is better","values":[0,30919,79509,73175,169302,432494],"isMain":false},{"metricName":"Total length (>= 50000 bp)","quality":"More is better","values":[0,0,0,0,54659,159768],"isMain":true},{"metricName":"N50","quality":"More is better","values":[10267,9420,8355,10672,11511,21509],"isMain":false},{"metricName":"N75","quality":"More is better","values":[5715,5023,4819,5291,6345,11206],"isMain":false},{"metricName":"L50","quality":"Less is better","values":[33,24,24,23,18,12],"isMain":false},{"metricName":"L75","quality":"Less is better","values":[65,51,52,50,39,29],"isMain":false},{"metricName":"GC (%)","quality":"Equal","values":["49.87","49.73","49.42","48.75","49.64","50.28"],"isMain":false}]],["K-mer-based statistics",[]],["Predicted genes",[]],["Similarity statistics",[]],["Reference statistics",[{"metricName":"Reference length","quality":"Equal","values":[5586179,5586179,5586179,5586179,5586179,5586179],"isMain":false},{"metricName":"Reference fragments","quality":"Equal","values":[47,47,47,47,47,47],"isMain":false},{"metricName":"Reference GC (%)","quality":"Equal","values":["49.16","49.16","49.16","49.16","49.16","49.16"],"isMain":false}]]],"subreferences":[],"subreports":[],"minContig":500},{"date":"10 February 2020, Monday, 19:46:27","assembliesNames":["megahit-103-df","megahit-113-df","megahit-113-ml","megahit-113-ms","megahit-129-df","metaSPAdes"],"referenceName":"GCF\_000371445.1\_Clos\_hath\_12489931\_V1","order":[0,1,2,3,4,5],"report":[["Genome statistics",[{"metricName":"Genome fraction (%)","quality":"More is better","values":["96.912","96.409","96.745","94.948","93.713","81.081"],"isMain":true},{"metricName":"Duplication ratio","quality":"Less is better","values":["1.013","1.073","1.058","1.066","1.056","1.006"],"isMain":true},{"metricName":"Largest alignment","quality":"More is better","values":[125726,129419,85095,183499,109344,250806],"isMain":true},{"metricName":"Total aligned length","quality":"More is better","values":[6719751,7058960,6991998,6912699,6755209,5578331],"isMain":true},{"metricName":"NG50","quality":"More is better","values":[14446,12851,16921,19055,14857,6857],"isMain":false},{"metricName":"NG75","quality":"More is better","values":[5993,6649,10195,9874,7886,1691],"isMain":false},{"metricName":"NA50","quality":"More is better","values":[14744,7002,10740,6596,7496,8220],"isMain":false},{"metricName":"NA75","quality":"More is better","values":[6194,2111,3367,1589,2167,2774],"isMain":false},{"metricName":"NGA50","quality":"More is better","values":[14446,9237,12524,9560,9420,5855],"isMain":true},{"metricName":"NGA75","quality":"More is better","values":[5993,3876,5907,4425,3843,1246],"isMain":false},{"metricName":"LG50","quality":"Less is better","values":[126,145,112,97,125,219],"isMain":false},{"metricName":"LG75","quality":"Less is better","values":[312,332,245,223,287,733],"isMain":false},{"metricName":"LA50","quality":"Less is better","values":[122,257,196,280,250,167],"isMain":false},{"metricName":"LA75","quality":"Less is better","values":[297,774,523,917,760,472],"isMain":false},{"metricName":"LGA50","quality":"Less is better","values":[126,186,149,173,183,242],"isMain":true},{"metricName":"LGA75","quality":"Less is better","values":[312,483,341,440,470,887],"isMain":false}]],["Reads mapping",[]],["Misassemblies",[{"metricName":"# misassemblies","quality":"Less is better","values":[3,17,14,24,17,3],"isMain":true},{"metricName":" # relocations","quality":"Less is better","values":[0,7,5,8,5,0],"isMain":false},{"metricName":" # translocations","quality":"Less is better","values":[3,8,7,11,12,3],"isMain":false},{"metricName":" # inversions","quality":"Less is better","values":[0,2,2,5,0,0],"isMain":false},{"metricName":" # interspecies translocations","quality":"Less is better","values":[5,183,138,193,212,39],"isMain":false},{"metricName":"# misassembled contigs","quality":"Less is better","values":[3,16,14,21,17,3],"isMain":false},{"metricName":"Misassembled contigs length","quality":"Less is better","values":[22142,209199,169930,200389,151495,10100],"isMain":true},{"metricName":"# possibly misassembled contigs","quality":"Less is better","values":[0,2,1,3,2,1],"isMain":false},{"metricName":" # possible misassemblies","quality":"Less is better","values":[0,2,2,5,4,2],"isMain":false},{"metricName":"# local misassemblies","quality":"Less is better","values":[0,21,28,26,32,6],"isMain":false},{"metricName":"# scaffold gap ext. mis.","quality":"Less is better","values":[0,0,0,0,0,0],"isMain":false},{"metricName":"# scaffold gap loc. mis.","quality":"Less is better","values":[0,0,0,0,0,0],"isMain":false},{"metricName":"# unaligned mis. contigs","quality":"Less is better","values":[0,22,15,26,24,8],"isMain":false}]],["Unaligned",[{"metricName":"# fully unaligned contigs","quality":"Less is better","values":[0,6,3,5,9,0],"isMain":false},{"metricName":"Fully unaligned length","quality":"Less is better","values":[0,30094,20937,17354,28208,0],"isMain":false},{"metricName":"# partially unaligned contigs","quality":"Less is better","values":[1,134,100,142,145,35],"isMain":false},{"metricName":"Partially unaligned length","quality":"Less is better","values":[757,852167,896518,1620010,1162157,215052],"isMain":false}]],["Mismatches",[{"metricName":"# mismatches","quality":"Less is better","values":[6077,41252,38486,44207,35106,15096],"isMain":false},{"metricName":"# indels","quality":"Less is better","values":[45,384,440,503,382,219],"isMain":false},{"metricName":"Indels length","quality":"Less is better","values":[356,2149,2954,2534,2556,1670],"isMain":false},{"metricName":"# mismatches per 100 kbp","quality":"Less is better","values":["91.56","624.76","580.85","679.82","546.98","271.85"],"isMain":true},{"metricName":"# indels per 100 kbp","quality":"Less is better","values":["0.68","5.82","6.64","7.74","5.95","3.94"],"isMain":true},{"metricName":" # indels (<= 5 bp)","quality":"Less is better","values":[39,317,370,431,308,168],"isMain":false},{"metricName":" # indels (> 5 bp)","quality":"Less is better","values":[6,67,70,72,74,51],"isMain":false},{"metricName":"# N's","quality":"Less is better","values":[0,0,0,0,0,0],"isMain":false},{"metricName":"# N's per 100 kbp","quality":"Less is better","values":["0.00","0.00","0.00","0.00","0.00","0.00"],"isMain":true}]],["Statistics without reference",[{"metricName":"# contigs","quality":"Equal","values":[1091,1794,1208,1405,1431,1409],"isMain":true},{"metricName":"# contigs (>= 1000 bp)","quality":"Equal","values":[829,1247,968,1123,1133,1038],"isMain":false},{"metricName":"# contigs (>= 5000 bp)","quality":"Equal","values":[357,422,422,448,422,295],"isMain":false},{"metricName":"# contigs (>= 10000 bp)","quality":"Equal","values":[194,214,251,218,209,140],"isMain":false},{"metricName":"# contigs (>= 25000 bp)","quality":"Equal","values":[55,41,53,58,50,19],"isMain":false},{"metricName":"# contigs (>= 50000 bp)","quality":"Equal","values":[8,10,12,16,9,4],"isMain":false},{"metricName":"Largest contig","quality":"More is better","values":[125726,129419,160824,183499,112101,250806],"isMain":true},{"metricName":"Total length","quality":"More is better","values":[6725370,7968430,7925975,8570693,7966635,5800326],"isMain":true},{"metricName":"Total length (>= 1000 bp)","quality":"More is better","values":[6537803,7584607,7754713,8369746,7753657,5534162],"isMain":true},{"metricName":"Total length (>= 5000 bp)","quality":"More is better","values":[5386898,5653749,6424778,6718633,6002871,3875178],"isMain":false},{"metricName":"Total length (>= 10000 bp)","quality":"More is better","values":[4242903,4200030,5204019,5092856,4449854,2777063],"isMain":true},{"metricName":"Total length (>= 25000 bp)","quality":"More is better","values":[2124492,1611467,2216478,2585458,1997207,906954],"isMain":false},{"metricName":"Total length (>= 50000 bp)","quality":"More is better","values":[596516,633322,883466,1246678,681420,443536],"isMain":true},{"metricName":"N50","quality":"More is better","values":[14756,10587,14308,13690,12310,9413],"isMain":false},{"metricName":"N75","quality":"More is better","values":[6194,4231,6986,5716,5073,3447],"isMain":false},{"metricName":"L50","quality":"Less is better","values":[122,194,147,151,167,153],"isMain":false},{"metricName":"L75","quality":"Less is better","values":[297,494,341,394,417,409],"isMain":false},{"metricName":"GC (%)","quality":"Equal","values":["49.35","49.30","49.28","49.01","49.30","49.34"],"isMain":false}]],["K-mer-based statistics",[]],["Predicted genes",[]],["Similarity statistics",[]],["Reference statistics",[{"metricName":"Reference length","quality":"Equal","values":[6848758,6848758,6848758,6848758,6848758,6848758],"isMain":false},{"metricName":"Reference fragments","quality":"Equal","values":[31,31,31,31,31,31],"isMain":false},{"metricName":"Reference GC (%)","quality":"Equal","values":["49.26","49.26","49.26","49.26","49.26","49.26"],"isMain":false}]]],"subreferences":[],"subreports":[],"minContig":500},{"date":"10 February 2020, Monday, 19:46:04","assembliesNames":["megahit-103-df","megahit-113-df","megahit-113-ml","megahit-113-ms","megahit-129-df","metaSPAdes"],"referenceName":"GCF\_000371485.1\_Clos\_clos\_90B1\_V1","order":[0,1,2,3,4,5],"report":[["Genome statistics",[{"metricName":"Genome fraction (%)","quality":"More is better","values":["12.949","13.861","14.141","16.527","16.161","17.409"],"isMain":true},{"metricName":"Duplication ratio","quality":"Less is better","values":["1.004","1.017","1.010","1.012","1.010","1.001"],"isMain":true},{"metricName":"Largest alignment","quality":"More is better","values":[23625,27342,23688,29403,41817,56425],"isMain":true},{"metricName":"Total aligned length","quality":"More is better","values":[718880,775363,786767,921210,898490,963691],"isMain":true},{"metricName":"NA50","quality":"More is better","values":[7551,8460,7345,8218,12841,22355],"isMain":false},{"metricName":"NA75","quality":"More is better","values":[3953,4506,3922,4415,6297,10094],"isMain":false},{"metricName":"NGA50","quality":"More is better","values":[null,null,null,null,null,null],"isMain":true},{"metricName":"LA50","quality":"Less is better","values":[28,32,34,36,22,16],"isMain":false},{"metricName":"LA75","quality":"Less is better","values":[61,62,71,76,49,32],"isMain":false}]],["Reads mapping",[]],["Misassemblies",[{"metricName":"# misassemblies","quality":"Less is better","values":[0,3,1,0,3,0],"isMain":true},{"metricName":" # relocations","quality":"Less is better","values":[0,1,0,0,0,0],"isMain":false},{"metricName":" # translocations","quality":"Less is better","values":[0,2,1,0,3,0],"isMain":false},{"metricName":" # inversions","quality":"Less is better","values":[0,0,0,0,0,0],"isMain":false},{"metricName":" # interspecies translocations","quality":"Less is better","values":[2,12,10,15,16,1],"isMain":false},{"metricName":"# misassembled contigs","quality":"Less is better","values":[0,3,1,0,2,0],"isMain":false},{"metricName":"Misassembled contigs length","quality":"Less is better","values":[0,11490,3020,0,11460,0],"isMain":true},{"metricName":"# possibly misassembled contigs","quality":"Less is better","values":[0,0,0,1,0,0],"isMain":false},{"metricName":" # possible misassemblies","quality":"Less is better","values":[0,0,0,1,0,0],"isMain":false},{"metricName":"# local misassemblies","quality":"Less is better","values":[1,7,4,5,3,2],"isMain":false},{"metricName":"# scaffold gap ext. mis.","quality":"Less is better","values":[0,0,0,0,0,0],"isMain":false},{"metricName":"# scaffold gap loc. mis.","quality":"Less is better","values":[0,0,0,0,0,0],"isMain":false},{"metricName":"# unaligned mis. contigs","quality":"Less is better","values":[0,2,0,1,1,0],"isMain":false}]],["Unaligned",[{"metricName":"# fully unaligned contigs","quality":"Less is better","values":[1,1,1,1,2,0],"isMain":false},{"metricName":"Fully unaligned length","quality":"Less is better","values":[583,598,705,559,1042,0],"isMain":false},{"metricName":"# partially unaligned contigs","quality":"Less is better","values":[1,4,1,11,7,0],"isMain":false},{"metricName":"Partially unaligned length","quality":"Less is better","values":[900,13254,618,35191,42817,0],"isMain":false}]],["Mismatches",[{"metricName":"# mismatches","quality":"Less is better","values":[1605,3740,3233,4481,3289,771],"isMain":false},{"metricName":"# indels","quality":"Less is better","values":[21,52,25,58,55,21],"isMain":false},{"metricName":"Indels length","quality":"Less is better","values":[41,227,143,297,293,111],"isMain":false},{"metricName":"# mismatches per 100 kbp","quality":"Less is better","values":["224.02","487.68","413.21","490.05","367.84","80.04"],"isMain":true},{"metricName":"# indels per 100 kbp","quality":"Less is better","values":["2.93","6.78","3.20","6.34","6.15","2.18"],"isMain":true},{"metricName":" # indels (<= 5 bp)","quality":"Less is better","values":[20,44,17,51,45,17],"isMain":false},{"metricName":" # indels (> 5 bp)","quality":"Less is better","values":[1,8,8,7,10,4],"isMain":false},{"metricName":"# N's","quality":"Less is better","values":[0,0,0,0,0,0],"isMain":false},{"metricName":"# N's per 100 kbp","quality":"Less is better","values":["0.00","0.00","0.00","0.00","0.00","0.00"],"isMain":true}]],["Statistics without reference",[{"metricName":"# contigs","quality":"Equal","values":[153,158,191,215,150,101],"isMain":true},{"metricName":"# contigs (>= 1000 bp)","quality":"Equal","values":[132,122,140,141,100,87],"isMain":false},{"metricName":"# contigs (>= 5000 bp)","quality":"Equal","values":[48,59,52,71,57,49],"isMain":false},{"metricName":"# contigs (>= 10000 bp)","quality":"Equal","values":[22,22,18,26,30,32],"isMain":false},{"metricName":"# contigs (>= 25000 bp)","quality":"Equal","values":[0,1,0,2,7,10],"isMain":false},{"metricName":"# contigs (>= 50000 bp)","quality":"Equal","values":[0,0,0,0,0,1],"isMain":false},{"metricName":"Largest contig","quality":"More is better","values":[23625,27342,23811,29403,42250,56425],"isMain":true},{"metricName":"Total length","quality":"More is better","values":[721037,793955,791609,960837,946720,964640],"isMain":true},{"metricName":"Total length (>= 1000 bp)","quality":"More is better","values":[706955,769030,757909,912943,914018,954493],"isMain":true},{"metricName":"Total length (>= 5000 bp)","quality":"More is better","values":[485085,597309,514299,725111,799495,861377],"isMain":false},{"metricName":"Total length (>= 10000 bp)","quality":"More is better","values":[313260,313694,261084,410828,599265,730751],"isMain":true},{"metricName":"Total length (>= 25000 bp)","quality":"More is better","values":[0,27342,0,54998,229575,346275],"isMain":false},{"metricName":"Total length (>= 50000 bp)","quality":"More is better","values":[0,0,0,0,0,56425],"isMain":true},{"metricName":"N50","quality":"More is better","values":[7551,8880,7345,8702,14616,22358],"isMain":false},{"metricName":"N75","quality":"More is better","values":[3953,5083,3948,5043,7492,10099],"isMain":false},{"metricName":"L50","quality":"Less is better","values":[28,31,34,34,20,16],"isMain":false},{"metricName":"L75","quality":"Less is better","values":[61,59,71,71,43,32],"isMain":false},{"metricName":"GC (%)","quality":"Equal","values":["48.61","49.42","49.02","49.28","49.30","48.58"],"isMain":false}]],["K-mer-based statistics",[]],["Predicted genes",[]],["Similarity statistics",[]],["Reference statistics",[{"metricName":"Reference length","quality":"Equal","values":[5532763,5532763,5532763,5532763,5532763,5532763],"isMain":false},{"metricName":"Reference fragments","quality":"Equal","values":[57,57,57,57,57,57],"isMain":false},{"metricName":"Reference GC (%)","quality":"Equal","values":["49.09","49.09","49.09","49.09","49.09","49.09"],"isMain":false}]]],"subreferences":[],"subreports":[],"minContig":500},{"date":"10 February 2020, Monday, 20:09:11","assembliesNames":["megahit-103-df","megahit-113-df","megahit-113-ml","megahit-113-ms","megahit-129-df","metaSPAdes"],"referenceName":"GCF\_000371505.1\_Clos\_clos\_90A8\_V1","order":[0,1,2,3,4,5],"report":[["Genome statistics",[{"metricName":"Genome fraction (%)","quality":"More is better","values":["3.153","4.138","3.608","3.511","2.580","1.071"],"isMain":true},{"metricName":"Duplication ratio","quality":"Less is better","values":["1.004","1.018","1.024","1.030","1.020","1.005"],"isMain":true},{"metricName":"Largest alignment","quality":"More is better","values":[14759,25597,18222,23424,16923,10185],"isMain":true},{"metricName":"Total aligned length","quality":"More is better","values":[187130,246379,216411,210075,153302,63592],"isMain":true},{"metricName":"NA50","quality":"More is better","values":[3682,4908,4815,3423,4508,4160],"isMain":false},{"metricName":"NA75","quality":"More is better","values":[2270,2127,1795,830,1140,1249],"isMain":false},{"metricName":"NGA50","quality":"More is better","values":[null,null,null,null,null,null],"isMain":true},{"metricName":"LA50","quality":"Less is better","values":[14,13,12,17,9,6],"isMain":false},{"metricName":"LA75","quality":"Less is better","values":[31,31,31,53,28,12],"isMain":false}]],["Reads mapping",[]],["Misassemblies",[{"metricName":"# misassemblies","quality":"Less is better","values":[0,9,4,6,2,0],"isMain":true},{"metricName":" # relocations","quality":"Less is better","values":[0,3,1,0,1,0],"isMain":false},{"metricName":" # translocations","quality":"Less is better","values":[0,6,3,6,1,0],"isMain":false},{"metricName":" # inversions","quality":"Less is better","values":[0,0,0,0,0,0],"isMain":false},{"metricName":" # interspecies translocations","quality":"Less is better","values":[7,15,9,19,10,0],"isMain":false},{"metricName":"# misassembled contigs","quality":"Less is better","values":[0,6,3,5,2,0],"isMain":false},{"metricName":"Misassembled contigs length","quality":"Less is better","values":[0,39596,8530,18995,4343,0],"isMain":true},{"metricName":"# possibly misassembled contigs","quality":"Less is better","values":[0,0,0,1,0,0],"isMain":false},{"metricName":" # possible misassemblies","quality":"Less is better","values":[0,0,0,1,0,0],"isMain":false},{"metricName":"# local misassemblies","quality":"Less is better","values":[1,2,0,1,2,0],"isMain":false},{"metricName":"# scaffold gap ext. mis.","quality":"Less is better","values":[0,0,0,0,0,0],"isMain":false},{"metricName":"# scaffold gap loc. mis.","quality":"Less is better","values":[0,0,0,0,0,0],"isMain":false},{"metricName":"# unaligned mis. contigs","quality":"Less is better","values":[1,0,0,3,2,0],"isMain":false}]],["Unaligned",[{"metricName":"# fully unaligned contigs","quality":"Less is better","values":[0,0,1,1,0,0],"isMain":false},{"metricName":"Fully unaligned length","quality":"Less is better","values":[0,0,6692,699,0,0],"isMain":false},{"metricName":"# partially unaligned contigs","quality":"Less is better","values":[6,5,2,9,4,0],"isMain":false},{"metricName":"Partially unaligned length","quality":"Less is better","values":[9433,6720,1585,33809,22799,0],"isMain":false}]],["Mismatches",[{"metricName":"# mismatches","quality":"Less is better","values":[740,1976,1747,1652,1035,254],"isMain":false},{"metricName":"# indels","quality":"Less is better","values":[9,41,17,15,29,1],"isMain":false},{"metricName":"Indels length","quality":"Less is better","values":[11,331,41,55,72,1],"isMain":false},{"metricName":"# mismatches per 100 kbp","quality":"Less is better","values":["395.95","805.66","817.02","793.98","676.82","400.24"],"isMain":true},{"metricName":"# indels per 100 kbp","quality":"Less is better","values":["4.82","16.72","7.95","7.21","18.96","1.58"],"isMain":true},{"metricName":" # indels (<= 5 bp)","quality":"Less is better","values":[9,31,16,14,25,1],"isMain":false},{"metricName":" # indels (> 5 bp)","quality":"Less is better","values":[0,10,1,1,4,0],"isMain":false},{"metricName":"# N's","quality":"Less is better","values":[0,0,0,0,0,0],"isMain":false},{"metricName":"# N's per 100 kbp","quality":"Less is better","values":["0.00","0.00","0.00","0.00","0.00","0.00"],"isMain":true}]],["Statistics without reference",[{"metricName":"# contigs","quality":"Equal","values":[73,80,87,94,56,33],"isMain":true},{"metricName":"# contigs (>= 1000 bp)","quality":"Equal","values":[50,54,48,49,34,15],"isMain":false},{"metricName":"# contigs (>= 5000 bp)","quality":"Equal","values":[11,13,12,16,9,3],"isMain":false},{"metricName":"# contigs (>= 10000 bp)","quality":"Equal","values":[3,5,5,4,6,1],"isMain":false},{"metricName":"# contigs (>= 25000 bp)","quality":"Equal","values":[0,1,0,0,0,0],"isMain":false},{"metricName":"# contigs (>= 50000 bp)","quality":"Equal","values":[0,0,0,0,0,0],"isMain":false},{"metricName":"Largest contig","quality":"More is better","values":[14767,25612,18222,23424,22085,10185],"isMain":true},{"metricName":"Total length","quality":"More is better","values":[197001,256463,227166,248785,178771,63766],"isMain":true},{"metricName":"Total length (>= 1000 bp)","quality":"More is better","values":[181410,238784,201233,218979,163263,52214],"isMain":true},{"metricName":"Total length (>= 5000 bp)","quality":"More is better","values":[89384,141467,120447,152368,110471,22550],"isMain":false},{"metricName":"Total length (>= 10000 bp)","quality":"More is better","values":[37895,87467,66635,68025,93383,10185],"isMain":true},{"metricName":"Total length (>= 25000 bp)","quality":"More is better","values":[0,25612,0,0,0,0],"isMain":false},{"metricName":"Total length (>= 50000 bp)","quality":"More is better","values":[0,0,0,0,0,0],"isMain":true},{"metricName":"N50","quality":"More is better","values":[3690,6047,6692,6034,11202,4160],"isMain":false},{"metricName":"N75","quality":"More is better","values":[2328,2983,2119,2218,2787,1257],"isMain":false},{"metricName":"L50","quality":"Less is better","values":[14,11,11,12,6,6],"isMain":false},{"metricName":"L75","quality":"Less is better","values":[30,27,27,28,17,12],"isMain":false},{"metricName":"GC (%)","quality":"Equal","values":["49.18","48.50","49.75","47.54","48.37","46.51"],"isMain":false}]],["K-mer-based statistics",[]],["Predicted genes",[]],["Similarity statistics",[]],["Reference statistics",[{"metricName":"Reference length","quality":"Equal","values":[5926778,5926778,5926778,5926778,5926778,5926778],"isMain":false},{"metricName":"Reference fragments","quality":"Equal","values":[100,100,100,100,100,100],"isMain":false},{"metricName":"Reference GC (%)","quality":"Equal","values":["48.88","48.88","48.88","48.88","48.88","48.88"],"isMain":false}]]],"subreferences":[],"subreports":[],"minContig":500},{"date":"10 February 2020, Monday, 20:10:50","assembliesNames":["megahit-103-df","megahit-113-df","megahit-113-ml","megahit-113-ms","megahit-129-df","metaSPAdes"],"referenceName":"GCF\_000371525.1\_Clos\_clos\_90A7\_V1","order":[0,1,2,3,4,5],"report":[["Genome statistics",[{"metricName":"Genome fraction (%)","quality":"More is better","values":["6.898","8.362","9.737","8.682","9.659","3.970"],"isMain":true},{"metricName":"Duplication ratio","quality":"Less is better","values":["1.017","1.042","1.029","1.031","1.025","1.027"],"isMain":true},{"metricName":"Largest alignment","quality":"More is better","values":[2770,11630,6587,11630,11630,11630],"isMain":true},{"metricName":"Total aligned length","quality":"More is better","values":[427464,525789,607212,540953,601056,246034],"isMain":true},{"metricName":"NA50","quality":"More is better","values":[690,731,679,662,747,711],"isMain":false},{"metricName":"NA75","quality":"More is better","values":[582,576,569,541,574,575],"isMain":false},{"metricName":"NGA50","quality":"More is better","values":[null,null,null,null,null,null],"isMain":true},{"metricName":"LA50","quality":"Less is better","values":[230,239,342,305,290,112],"isMain":false},{"metricName":"LA75","quality":"Less is better","values":[401,456,595,559,545,210],"isMain":false}]],["Reads mapping",[]],["Misassemblies",[{"metricName":"# misassemblies","quality":"Less is better","values":[1,10,3,7,4,0],"isMain":true},{"metricName":" # relocations","quality":"Less is better","values":[1,6,0,2,3,0],"isMain":false},{"metricName":" # translocations","quality":"Less is better","values":[0,2,1,1,1,0],"isMain":false},{"metricName":" # inversions","quality":"Less is better","values":[0,2,2,4,0,0],"isMain":false},{"metricName":" # interspecies translocations","quality":"Less is better","values":[15,41,24,30,26,7],"isMain":false},{"metricName":"# misassembled contigs","quality":"Less is better","values":[1,9,3,6,4,0],"isMain":false},{"metricName":"Misassembled contigs length","quality":"Less is better","values":[528,34291,3262,24652,4610,0],"isMain":true},{"metricName":"# possibly misassembled contigs","quality":"Less is better","values":[0,0,0,0,0,0],"isMain":false},{"metricName":" # possible misassemblies","quality":"Less is better","values":[0,0,0,0,0,0],"isMain":false},{"metricName":"# local misassemblies","quality":"Less is better","values":[0,2,3,2,1,0],"isMain":false},{"metricName":"# scaffold gap ext. mis.","quality":"Less is better","values":[0,0,0,0,0,0],"isMain":false},{"metricName":"# scaffold gap loc. mis.","quality":"Less is better","values":[0,0,0,0,0,0],"isMain":false},{"metricName":"# unaligned mis. contigs","quality":"Less is better","values":[0,0,0,1,2,0],"isMain":false}]],["Unaligned",[{"metricName":"# fully unaligned contigs","quality":"Less is better","values":[2,1,3,1,2,0],"isMain":false},{"metricName":"Fully unaligned length","quality":"Less is better","values":[1185,630,2251,595,1681,0],"isMain":false},{"metricName":"# partially unaligned contigs","quality":"Less is better","values":[2,12,7,9,10,0],"isMain":false},{"metricName":"Partially unaligned length","quality":"Less is better","values":[1906,19080,10383,54929,50515,0],"isMain":false}]],["Mismatches",[{"metricName":"# mismatches","quality":"Less is better","values":[7610,9511,13676,12297,11580,4079],"isMain":false},{"metricName":"# indels","quality":"Less is better","values":[33,60,55,57,64,29],"isMain":false},{"metricName":"Indels length","quality":"Less is better","values":[116,203,255,185,296,82],"isMain":false},{"metricName":"# mismatches per 100 kbp","quality":"Less is better","values":["1791.29","1846.85","2280.54","2299.75","1946.76","1668.21"],"isMain":true},{"metricName":"# indels per 100 kbp","quality":"Less is better","values":["7.77","11.65","9.17","10.66","10.76","11.86"],"isMain":true},{"metricName":" # indels (<= 5 bp)","quality":"Less is better","values":[27,52,47,49,54,25],"isMain":false},{"metricName":" # indels (> 5 bp)","quality":"Less is better","values":[6,8,8,8,10,4],"isMain":false},{"metricName":"# N's","quality":"Less is better","values":[0,0,0,0,0,0],"isMain":false},{"metricName":"# N's per 100 kbp","quality":"Less is better","values":["0.00","0.00","0.00","0.00","0.00","0.00"],"isMain":true}]],["Statistics without reference",[{"metricName":"# contigs","quality":"Equal","values":[595,667,857,733,742,321],"isMain":true},{"metricName":"# contigs (>= 1000 bp)","quality":"Equal","values":[68,100,71,88,134,38],"isMain":false},{"metricName":"# contigs (>= 5000 bp)","quality":"Equal","values":[0,3,2,4,5,2],"isMain":false},{"metricName":"# contigs (>= 10000 bp)","quality":"Equal","values":[0,2,0,2,2,1],"isMain":false},{"metricName":"# contigs (>= 25000 bp)","quality":"Equal","values":[0,0,0,1,1,0],"isMain":false},{"metricName":"# contigs (>= 50000 bp)","quality":"Equal","values":[0,0,0,0,0,0],"isMain":false},{"metricName":"Largest contig","quality":"More is better","values":[2770,14017,8198,33133,42250,11684],"isMain":true},{"metricName":"Total length","quality":"More is better","values":[435134,556077,629499,606619,661629,251145],"isMain":true},{"metricName":"Total length (>= 1000 bp)","quality":"More is better","values":[88371,179653,105873,183961,252210,65668],"isMain":true},{"metricName":"Total length (>= 5000 bp)","quality":"More is better","values":[0,33976,14886,66245,72219,18346],"isMain":false},{"metricName":"Total length (>= 10000 bp)","quality":"More is better","values":[0,26348,0,54546,54057,11684],"isMain":true},{"metricName":"Total length (>= 25000 bp)","quality":"More is better","values":[0,0,0,33133,42250,0],"isMain":false},{"metricName":"Total length (>= 50000 bp)","quality":"More is better","values":[0,0,0,0,0,0],"isMain":true},{"metricName":"N50","quality":"More is better","values":[700,781,707,762,848,723],"isMain":false},{"metricName":"N75","quality":"More is better","values":[591,610,593,597,637,594],"isMain":false},{"metricName":"L50","quality":"Less is better","values":[226,212,323,227,221,110],"isMain":false},{"metricName":"L75","quality":"Less is better","values":[395,415,567,454,450,206],"isMain":false},{"metricName":"GC (%)","quality":"Equal","values":["49.60","48.74","49.25","48.87","49.56","48.26"],"isMain":false}]],["K-mer-based statistics",[]],["Predicted genes",[]],["Similarity statistics",[]],["Reference statistics",[{"metricName":"Reference length","quality":"Equal","values":[6158650,6158650,6158650,6158650,6158650,6158650],"isMain":false},{"metricName":"Reference fragments","quality":"Equal","values":[35,35,35,35,35,35],"isMain":false},{"metricName":"Reference GC (%)","quality":"Equal","values":["49.22","49.22","49.22","49.22","49.22","49.22"],"isMain":false}]]],"subreferences":[],"subreports":[],"minContig":500},{"date":"10 February 2020, Monday, 20:25:05","assembliesNames":["megahit-103-df","megahit-113-df","megahit-113-ml","megahit-113-ms","megahit-129-df","metaSPAdes"],"referenceName":"GCF\_000371545.1\_Clos\_clos\_90A6\_V1","order":[0,1,2,3,4,5],"report":[["Genome statistics",[{"metricName":"Genome fraction (%)","quality":"More is better","values":["26.052","29.167","27.470","24.781","27.177","26.881"],"isMain":true},{"metricName":"Duplication ratio","quality":"Less is better","values":["1.019","1.017","1.033","1.040","1.028","1.012"],"isMain":true},{"metricName":"Largest alignment","quality":"More is better","values":[33645,84604,23334,62546,38346,78118],"isMain":true},{"metricName":"Total aligned length","quality":"More is better","values":[1580285,1765053,1683706,1530207,1655953,1620726],"isMain":true},{"metricName":"NA50","quality":"More is better","values":[6522,8375,6685,5534,8948,15464],"isMain":false},{"metricName":"NA75","quality":"More is better","values":[3241,4088,2406,2065,3557,3672],"isMain":false},{"metricName":"NGA50","quality":"More is better","values":[null,null,null,null,null,null],"isMain":true},{"metricName":"LA50","quality":"Less is better","values":[73,53,76,63,52,21],"isMain":false},{"metricName":"LA75","quality":"Less is better","values":[160,128,182,184,126,81],"isMain":false}]],["Reads mapping",[]],["Misassemblies",[{"metricName":"# misassemblies","quality":"Less is better","values":[5,5,8,12,7,1],"isMain":true},{"metricName":" # relocations","quality":"Less is better","values":[0,1,1,1,2,0],"isMain":false},{"metricName":" # translocations","quality":"Less is better","values":[5,4,7,10,5,1],"isMain":false},{"metricName":" # inversions","quality":"Less is better","values":[0,0,0,1,0,0],"isMain":false},{"metricName":" # interspecies translocations","quality":"Less is better","values":[17,20,23,25,35,5],"isMain":false},{"metricName":"# misassembled contigs","quality":"Less is better","values":[4,5,8,11,7,1],"isMain":false},{"metricName":"Misassembled contigs length","quality":"Less is better","values":[17368,27981,30586,23356,94662,1646],"isMain":true},{"metricName":"# possibly misassembled contigs","quality":"Less is better","values":[0,1,0,2,1,0],"isMain":false},{"metricName":" # possible misassemblies","quality":"Less is better","values":[0,1,0,2,1,0],"isMain":false},{"metricName":"# local misassemblies","quality":"Less is better","values":[2,2,3,6,10,2],"isMain":false},{"metricName":"# scaffold gap ext. mis.","quality":"Less is better","values":[0,0,0,0,0,0],"isMain":false},{"metricName":"# scaffold gap loc. mis.","quality":"Less is better","values":[0,0,0,0,0,0],"isMain":false},{"metricName":"# unaligned mis. contigs","quality":"Less is better","values":[1,1,0,0,3,0],"isMain":false}]],["Unaligned",[{"metricName":"# fully unaligned contigs","quality":"Less is better","values":[0,1,0,1,1,0],"isMain":false},{"metricName":"Fully unaligned length","quality":"Less is better","values":[0,1948,0,800,683,0],"isMain":false},{"metricName":"# partially unaligned contigs","quality":"Less is better","values":[6,10,6,11,11,3],"isMain":false},{"metricName":"Partially unaligned length","quality":"Less is better","values":[6106,17708,6829,9478,55569,2024],"isMain":false}]],["Mismatches",[{"metricName":"# mismatches","quality":"Less is better","values":[5148,7197,10068,10817,7135,2800],"isMain":false},{"metricName":"# indels","quality":"Less is better","values":[43,95,43,76,89,20],"isMain":false},{"metricName":"Indels length","quality":"Less is better","values":[136,512,165,446,404,47],"isMain":false},{"metricName":"# mismatches per 100 kbp","quality":"Less is better","values":["331.00","413.31","613.91","731.16","439.77","174.47"],"isMain":true},{"metricName":"# indels per 100 kbp","quality":"Less is better","values":["2.76","5.46","2.62","5.14","5.49","1.25"],"isMain":true},{"metricName":" # indels (<= 5 bp)","quality":"Less is better","values":[37,84,40,64,74,17],"isMain":false},{"metricName":" # indels (> 5 bp)","quality":"Less is better","values":[6,11,3,12,15,3],"isMain":false},{"metricName":"# N's","quality":"Less is better","values":[0,0,0,0,0,0],"isMain":false},{"metricName":"# N's per 100 kbp","quality":"Less is better","values":["0.00","0.00","0.00","0.00","0.00","0.00"],"isMain":true}]],["Statistics without reference",[{"metricName":"# contigs","quality":"Equal","values":[427,370,559,549,384,340],"isMain":true},{"metricName":"# contigs (>= 1000 bp)","quality":"Equal","values":[332,294,348,338,279,249],"isMain":false},{"metricName":"# contigs (>= 5000 bp)","quality":"Equal","values":[103,108,102,71,99,56],"isMain":false},{"metricName":"# contigs (>= 10000 bp)","quality":"Equal","values":[30,43,40,30,42,31],"isMain":false},{"metricName":"# contigs (>= 25000 bp)","quality":"Equal","values":[1,8,0,4,9,16],"isMain":false},{"metricName":"# contigs (>= 50000 bp)","quality":"Equal","values":[0,1,0,1,0,5],"isMain":false},{"metricName":"Largest contig","quality":"More is better","values":[33645,85490,23334,62546,48437,78118],"isMain":true},{"metricName":"Total length","quality":"More is better","values":[1591709,1791126,1700449,1548868,1724444,1625919],"isMain":true},{"metricName":"Total length (>= 1000 bp)","quality":"More is better","values":[1526139,1737887,1550796,1398195,1650978,1558459],"isMain":true},{"metricName":"Total length (>= 5000 bp)","quality":"More is better","values":[977167,1275008,1004671,820784,1246887,1117500],"isMain":false},{"metricName":"Total length (>= 10000 bp)","quality":"More is better","values":[457476,813661,565786,532259,832438,948067],"isMain":true},{"metricName":"Total length (>= 25000 bp)","quality":"More is better","values":[33645,301587,0,145397,292285,723165],"isMain":false},{"metricName":"Total length (>= 50000 bp)","quality":"More is better","values":[0,85490,0,62546,0,327497],"isMain":true},{"metricName":"N50","quality":"More is better","values":[6581,8990,6698,5564,9937,15464],"isMain":false},{"metricName":"N75","quality":"More is better","values":[3270,4206,2686,2218,4300,3688],"isMain":false},{"metricName":"L50","quality":"Less is better","values":[72,52,75,63,45,21],"isMain":false},{"metricName":"L75","quality":"Less is better","values":[157,123,178,180,109,81],"isMain":false},{"metricName":"GC (%)","quality":"Equal","values":["47.76","47.56","47.59","47.79","47.70","47.98"],"isMain":false}]],["K-mer-based statistics",[]],["Predicted genes",[]],["Similarity statistics",[]],["Reference statistics",[{"metricName":"Reference length","quality":"Equal","values":[5970004,5970004,5970004,5970004,5970004,5970004],"isMain":false},{"metricName":"Reference fragments","quality":"Equal","values":[72,72,72,72,72,72],"isMain":false},{"metricName":"Reference GC (%)","quality":"Equal","values":["48.75","48.75","48.75","48.75","48.75","48.75"],"isMain":false}]]],"subreferences":[],"subreports":[],"minContig":500},{"date":"10 February 2020, Monday, 20:25:25","assembliesNames":["megahit-103-df","megahit-113-df","megahit-113-ml","megahit-113-ms","megahit-129-df","metaSPAdes"],"referenceName":"GCF\_000371585.1\_Clos\_clos\_90A3\_V1","order":[0,1,2,3,4,5],"report":[["Genome statistics",[{"metricName":"Genome fraction (%)","quality":"More is better","values":["16.222","18.159","16.024","16.896","20.060","17.983"],"isMain":true},{"metricName":"Duplication ratio","quality":"Less is better","values":["1.006","1.013","1.015","1.023","1.011","1.001"],"isMain":true},{"metricName":"Largest alignment","quality":"More is better","values":[67113,27755,36045,30749,45965,62630],"isMain":true},{"metricName":"Total aligned length","quality":"More is better","values":[891097,1000571,883765,937546,1103411,984193],"isMain":true},{"metricName":"NA50","quality":"More is better","values":[9718,9911,8542,8024,10951,20169],"isMain":false},{"metricName":"NA75","quality":"More is better","values":[4613,5223,4772,3919,4688,10295],"isMain":false},{"metricName":"NGA50","quality":"More is better","values":[null,null,null,null,null,null],"isMain":true},{"metricName":"LA50","quality":"Less is better","values":[25,31,33,39,31,16],"isMain":false},{"metricName":"LA75","quality":"Less is better","values":[57,66,68,82,70,33],"isMain":false}]],["Reads mapping",[]],["Misassemblies",[{"metricName":"# misassemblies","quality":"Less is better","values":[0,4,5,8,10,0],"isMain":true},{"metricName":" # relocations","quality":"Less is better","values":[0,1,0,0,2,0],"isMain":false},{"metricName":" # translocations","quality":"Less is better","values":[0,2,4,6,8,0],"isMain":false},{"metricName":" # inversions","quality":"Less is better","values":[0,1,1,2,0,0],"isMain":false},{"metricName":" # interspecies translocations","quality":"Less is better","values":[4,14,12,24,24,1],"isMain":false},{"metricName":"# misassembled contigs","quality":"Less is better","values":[0,3,4,7,7,0],"isMain":false},{"metricName":"Misassembled contigs length","quality":"Less is better","values":[0,5286,11447,55374,85323,0],"isMain":true},{"metricName":"# possibly misassembled contigs","quality":"Less is better","values":[0,0,0,0,0,0],"isMain":false},{"metricName":" # possible misassemblies","quality":"Less is better","values":[0,0,0,0,0,0],"isMain":false},{"metricName":"# local misassemblies","quality":"Less is better","values":[0,4,3,0,2,0],"isMain":false},{"metricName":"# scaffold gap ext. mis.","quality":"Less is better","values":[0,0,0,0,0,0],"isMain":false},{"metricName":"# scaffold gap loc. mis.","quality":"Less is better","values":[0,0,0,0,0,0],"isMain":false},{"metricName":"# unaligned mis. contigs","quality":"Less is better","values":[0,0,0,2,0,0],"isMain":false}]],["Unaligned",[{"metricName":"# fully unaligned contigs","quality":"Less is better","values":[1,1,1,0,0,0],"isMain":false},{"metricName":"Fully unaligned length","quality":"Less is better","values":[528,518,649,0,0,0],"isMain":false},{"metricName":"# partially unaligned contigs","quality":"Less is better","values":[4,4,4,10,10,1],"isMain":false},{"metricName":"Partially unaligned length","quality":"Less is better","values":[3141,4215,4435,46896,115021,28564],"isMain":false}]],["Mismatches",[{"metricName":"# mismatches","quality":"Less is better","values":[1718,4154,3837,5027,3218,781],"isMain":false},{"metricName":"# indels","quality":"Less is better","values":[21,83,33,70,45,18],"isMain":false},{"metricName":"Indels length","quality":"Less is better","values":[52,400,122,409,290,22],"isMain":false},{"metricName":"# mismatches per 100 kbp","quality":"Less is better","values":["193.71","418.43","438.00","544.23","293.43","79.44"],"isMain":true},{"metricName":"# indels per 100 kbp","quality":"Less is better","values":["2.37","8.36","3.77","7.58","4.10","1.83"],"isMain":true},{"metricName":" # indels (<= 5 bp)","quality":"Less is better","values":[19,75,31,58,34,18],"isMain":false},{"metricName":" # indels (> 5 bp)","quality":"Less is better","values":[2,8,2,12,11,0],"isMain":false},{"metricName":"# N's","quality":"Less is better","values":[0,0,0,0,0,0],"isMain":false},{"metricName":"# N's per 100 kbp","quality":"Less is better","values":["0.00","0.00","0.00","0.00","0.00","0.00"],"isMain":true}]],["Statistics without reference",[{"metricName":"# contigs","quality":"Equal","values":[176,185,194,220,180,96],"isMain":true},{"metricName":"# contigs (>= 1000 bp)","quality":"Equal","values":[137,144,136,151,129,79],"isMain":false},{"metricName":"# contigs (>= 5000 bp)","quality":"Equal","values":[55,74,63,70,65,50],"isMain":false},{"metricName":"# contigs (>= 10000 bp)","quality":"Equal","values":[23,30,20,26,38,34],"isMain":false},{"metricName":"# contigs (>= 25000 bp)","quality":"Equal","values":[3,2,2,3,9,10],"isMain":false},{"metricName":"# contigs (>= 50000 bp)","quality":"Equal","values":[1,0,0,0,1,3],"isMain":false},{"metricName":"Largest contig","quality":"More is better","values":[67572,28125,36050,42744,114511,62630],"isMain":true},{"metricName":"Total length","quality":"More is better","values":[895990,1010267,894128,991658,1223755,1012942],"isMain":true},{"metricName":"Total length (>= 1000 bp)","quality":"More is better","values":[869205,983266,855940,946971,1191707,1000752],"isMain":true},{"metricName":"Total length (>= 5000 bp)","quality":"More is better","values":[667812,806310,648491,742391,1023887,924261],"isMain":false},{"metricName":"Total length (>= 10000 bp)","quality":"More is better","values":[436604,497684,333304,427731,819552,806817],"isMain":true},{"metricName":"Total length (>= 25000 bp)","quality":"More is better","values":[126017,53628,69828,101778,366138,414314],"isMain":false},{"metricName":"Total length (>= 50000 bp)","quality":"More is better","values":[67572,0,0,0,114511,177926],"isMain":true},{"metricName":"N50","quality":"More is better","values":[9718,9911,8542,8573,16433,21174],"isMain":false},{"metricName":"N75","quality":"More is better","values":[4858,5561,4772,4942,8455,11596],"isMain":false},{"metricName":"L50","quality":"Less is better","values":[25,31,33,34,22,15],"isMain":false},{"metricName":"L75","quality":"Less is better","values":[56,65,68,71,49,30],"isMain":false},{"metricName":"GC (%)","quality":"Equal","values":["48.26","48.45","48.80","49.12","48.05","49.01"],"isMain":false}]],["K-mer-based statistics",[]],["Predicted genes",[]],["Similarity statistics",[]],["Reference statistics",[{"metricName":"Reference length","quality":"Equal","values":[5466970,5466970,5466970,5466970,5466970,5466970],"isMain":false},{"metricName":"Reference fragments","quality":"Equal","values":[58,58,58,58,58,58],"isMain":false},{"metricName":"Reference GC (%)","quality":"Equal","values":["49.11","49.11","49.11","49.11","49.11","49.11"],"isMain":false}]]],"subreferences":[],"subreports":[],"minContig":500},{"date":"10 February 2020, Monday, 20:26:01","assembliesNames":["megahit-103-df","megahit-113-df","megahit-113-ml","megahit-113-ms","megahit-129-df","metaSPAdes"],"referenceName":"GCF\_000371605.1\_Clos\_clos\_90A1\_V1","order":[0,1,2,3,4,5],"report":[["Genome statistics",[{"metricName":"Genome fraction (%)","quality":"More is better","values":["13.431","12.624","17.555","15.442","13.740","12.762"],"isMain":true},{"metricName":"Duplication ratio","quality":"Less is better","values":["1.008","1.016","1.020","1.017","1.019","1.003"],"isMain":true},{"metricName":"Largest alignment","quality":"More is better","values":[26605,25817,23875,21634,27753,42628],"isMain":true},{"metricName":"Total aligned length","quality":"More is better","values":[778654,735267,1026892,901187,802039,737278],"isMain":true},{"metricName":"NA50","quality":"More is better","values":[6825,7202,7171,7839,9346,16985],"isMain":false},{"metricName":"NA75","quality":"More is better","values":[3839,2606,3686,4235,3634,8055],"isMain":false},{"metricName":"NGA50","quality":"More is better","values":[null,null,null,null,null,null],"isMain":true},{"metricName":"LA50","quality":"Less is better","values":[38,31,46,41,28,15],"isMain":false},{"metricName":"LA75","quality":"Less is better","values":[74,78,98,82,63,31],"isMain":false}]],["Reads mapping",[]],["Misassemblies",[{"metricName":"# misassemblies","quality":"Less is better","values":[1,5,5,7,7,1],"isMain":true},{"metricName":" # relocations","quality":"Less is better","values":[0,0,0,1,3,0],"isMain":false},{"metricName":" # translocations","quality":"Less is better","values":[0,5,4,6,4,0],"isMain":false},{"metricName":" # inversions","quality":"Less is better","values":[1,0,1,0,0,1],"isMain":false},{"metricName":" # interspecies translocations","quality":"Less is better","values":[8,16,13,25,21,3],"isMain":false},{"metricName":"# misassembled contigs","quality":"Less is better","values":[1,3,4,6,6,1],"isMain":false},{"metricName":"Misassembled contigs length","quality":"Less is better","values":[1121,17797,21793,31920,43986,725],"isMain":true},{"metricName":"# possibly misassembled contigs","quality":"Less is better","values":[0,0,0,0,0,0],"isMain":false},{"metricName":" # possible misassemblies","quality":"Less is better","values":[0,0,0,0,0,0],"isMain":false},{"metricName":"# local misassemblies","quality":"Less is better","values":[1,1,2,4,3,0],"isMain":false},{"metricName":"# scaffold gap ext. mis.","quality":"Less is better","values":[0,0,0,0,0,0],"isMain":false},{"metricName":"# scaffold gap loc. mis.","quality":"Less is better","values":[0,0,0,0,0,0],"isMain":false},{"metricName":"# unaligned mis. contigs","quality":"Less is better","values":[0,1,0,0,2,0],"isMain":false}]],["Unaligned",[{"metricName":"# fully unaligned contigs","quality":"Less is better","values":[0,1,0,0,1,0],"isMain":false},{"metricName":"Fully unaligned length","quality":"Less is better","values":[0,823,0,0,608,0],"isMain":false},{"metricName":"# partially unaligned contigs","quality":"Less is better","values":[3,8,5,15,9,1],"isMain":false},{"metricName":"Partially unaligned length","quality":"Less is better","values":[2797,52854,35767,43775,14423,703],"isMain":false}]],["Mismatches",[{"metricName":"# mismatches","quality":"Less is better","values":[2261,3585,5222,5213,3520,903],"isMain":false},{"metricName":"# indels","quality":"Less is better","values":[35,43,35,63,50,5],"isMain":false},{"metricName":"Indels length","quality":"Less is better","values":[285,329,227,405,308,16],"isMain":false},{"metricName":"# mismatches per 100 kbp","quality":"Less is better","values":["291.80","492.28","515.63","585.16","444.07","122.65"],"isMain":true},{"metricName":"# indels per 100 kbp","quality":"Less is better","values":["4.52","5.90","3.46","7.07","6.31","0.68"],"isMain":true},{"metricName":" # indels (<= 5 bp)","quality":"Less is better","values":[28,32,28,50,38,4],"isMain":false},{"metricName":" # indels (> 5 bp)","quality":"Less is better","values":[7,11,7,13,12,1],"isMain":false},{"metricName":"# N's","quality":"Less is better","values":[0,0,0,0,0,0],"isMain":false},{"metricName":"# N's per 100 kbp","quality":"Less is better","values":["0.00","0.00","0.00","0.00","0.00","0.00"],"isMain":true}]],["Statistics without reference",[{"metricName":"# contigs","quality":"Equal","values":[205,201,273,225,200,125],"isMain":true},{"metricName":"# contigs (>= 1000 bp)","quality":"Equal","values":[146,138,175,145,135,89],"isMain":false},{"metricName":"# contigs (>= 5000 bp)","quality":"Equal","values":[60,39,71,74,50,39],"isMain":false},{"metricName":"# contigs (>= 10000 bp)","quality":"Equal","values":[17,22,25,28,23,25],"isMain":false},{"metricName":"# contigs (>= 25000 bp)","quality":"Equal","values":[1,2,1,0,1,6],"isMain":false},{"metricName":"# contigs (>= 50000 bp)","quality":"Equal","values":[0,0,0,0,0,0],"isMain":false},{"metricName":"Largest contig","quality":"More is better","values":[26622,49720,39277,22970,27753,42734],"isMain":true},{"metricName":"Total length","quality":"More is better","values":[784195,793604,1068476,950144,822866,739314],"isMain":true},{"metricName":"Total length (>= 1000 bp)","quality":"More is better","values":[742511,747092,1000699,896374,779826,713324],"isMain":true},{"metricName":"Total length (>= 5000 bp)","quality":"More is better","values":[528877,499031,725118,718182,572703,605280],"isMain":false},{"metricName":"Total length (>= 10000 bp)","quality":"More is better","values":[228060,375096,384873,393204,374613,499101],"isMain":true},{"metricName":"Total length (>= 25000 bp)","quality":"More is better","values":[26622,75537,39277,0,27753,199634],"isMain":false},{"metricName":"Total length (>= 50000 bp)","quality":"More is better","values":[0,0,0,0,0,0],"isMain":true},{"metricName":"N50","quality":"More is better","values":[6930,8264,8130,8278,9431,16985],"isMain":false},{"metricName":"N75","quality":"More is better","values":[3930,3687,4157,5135,4204,8055],"isMain":false},{"metricName":"L50","quality":"Less is better","values":[38,25,42,37,27,15],"isMain":false},{"metricName":"L75","quality":"Less is better","values":[73,62,88,73,60,31],"isMain":false},{"metricName":"GC (%)","quality":"Equal","values":["48.34","48.33","48.67","48.28","49.05","48.34"],"isMain":false}]],["K-mer-based statistics",[]],["Predicted genes",[]],["Similarity statistics",[]],["Reference statistics",[{"metricName":"Reference length","quality":"Equal","values":[5768952,5768952,5768952,5768952,5768952,5768952],"isMain":false},{"metricName":"Reference fragments","quality":"Equal","values":[47,47,47,47,47,47],"isMain":false},{"metricName":"Reference GC (%)","quality":"Equal","values":["48.73","48.73","48.73","48.73","48.73","48.73"],"isMain":false}]]],"subreferences":[],"subreports":[],"minContig":500},{"date":"10 February 2020, Monday, 20:26:27","assembliesNames":["megahit-103-df","megahit-113-df","megahit-113-ml","megahit-113-ms","megahit-129-df","metaSPAdes"],"referenceName":"GCF\_000371685.1\_Clos\_bolt\_90B3\_V1","order":[0,1,2,3,4,5],"report":[["Genome statistics",[{"metricName":"Genome fraction (%)","quality":"More is better","values":["7.936","9.654","10.090","9.865","10.526","4.303"],"isMain":true},{"metricName":"Duplication ratio","quality":"Less is better","values":["1.023","1.021","1.025","1.033","1.027","1.024"],"isMain":true},{"metricName":"Largest alignment","quality":"More is better","values":[2227,3800,2495,6561,3025,3581],"isMain":true},{"metricName":"Total aligned length","quality":"More is better","values":[518154,628625,660722,648018,689875,280122],"isMain":true},{"metricName":"NA50","quality":"More is better","values":[691,762,690,686,762,666],"isMain":false},{"metricName":"NA75","quality":"More is better","values":[566,596,564,568,600,551],"isMain":false},{"metricName":"NGA50","quality":"More is better","values":[null,null,null,null,null,null],"isMain":true},{"metricName":"LA50","quality":"Less is better","values":[280,290,379,360,328,158],"isMain":false},{"metricName":"LA75","quality":"Less is better","values":[496,530,661,632,595,278],"isMain":false}]],["Reads mapping",[]],["Misassemblies",[{"metricName":"# misassemblies","quality":"Less is better","values":[0,0,3,2,2,0],"isMain":true},{"metricName":" # relocations","quality":"Less is better","values":[0,0,0,0,1,0],"isMain":false},{"metricName":" # translocations","quality":"Less is better","values":[0,0,1,1,0,0],"isMain":false},{"metricName":" # inversions","quality":"Less is better","values":[0,0,2,1,1,0],"isMain":false},{"metricName":" # interspecies translocations","quality":"Less is better","values":[26,26,10,30,28,14],"isMain":false},{"metricName":"# misassembled contigs","quality":"Less is better","values":[0,0,3,2,2,0],"isMain":false},{"metricName":"Misassembled contigs length","quality":"Less is better","values":[0,0,5510,1399,1734,0],"isMain":true},{"metricName":"# possibly misassembled contigs","quality":"Less is better","values":[0,1,0,0,1,0],"isMain":false},{"metricName":" # possible misassemblies","quality":"Less is better","values":[0,1,0,0,1,0],"isMain":false},{"metricName":"# local misassemblies","quality":"Less is better","values":[1,2,4,0,3,0],"isMain":false},{"metricName":"# scaffold gap ext. mis.","quality":"Less is better","values":[0,0,0,0,0,0],"isMain":false},{"metricName":"# scaffold gap loc. mis.","quality":"Less is better","values":[0,0,0,0,0,0],"isMain":false},{"metricName":"# unaligned mis. contigs","quality":"Less is better","values":[0,0,1,1,3,0],"isMain":false}]],["Unaligned",[{"metricName":"# fully unaligned contigs","quality":"Less is better","values":[2,1,0,5,1,1],"isMain":false},{"metricName":"Fully unaligned length","quality":"Less is better","values":[1265,517,0,4347,690,1328],"isMain":false},{"metricName":"# partially unaligned contigs","quality":"Less is better","values":[6,6,3,5,7,6],"isMain":false},{"metricName":"Partially unaligned length","quality":"Less is better","values":[6372,5847,32537,10715,17522,4778],"isMain":false}]],["Mismatches",[{"metricName":"# mismatches","quality":"Less is better","values":[8656,10737,14196,14479,12155,4661],"isMain":false},{"metricName":"# indels","quality":"Less is better","values":[46,57,38,47,45,7],"isMain":false},{"metricName":"Indels length","quality":"Less is better","values":[245,299,176,207,160,88],"isMain":false},{"metricName":"# mismatches per 100 kbp","quality":"Less is better","values":["1683.39","1716.43","2171.21","2265.06","1782.17","1671.71"],"isMain":true},{"metricName":"# indels per 100 kbp","quality":"Less is better","values":["8.95","9.11","5.81","7.35","6.60","2.51"],"isMain":true},{"metricName":" # indels (<= 5 bp)","quality":"Less is better","values":[41,49,34,44,41,5],"isMain":false},{"metricName":" # indels (> 5 bp)","quality":"Less is better","values":[5,8,4,3,4,2],"isMain":false},{"metricName":"# N's","quality":"Less is better","values":[0,0,0,0,0,0],"isMain":false},{"metricName":"# N's per 100 kbp","quality":"Less is better","values":["0.00","0.00","0.00","0.00","0.00","0.00"],"isMain":true}]],["Statistics without reference",[{"metricName":"# contigs","quality":"Equal","values":[727,802,919,910,879,400],"isMain":true},{"metricName":"# contigs (>= 1000 bp)","quality":"Equal","values":[97,140,101,98,162,38],"isMain":false},{"metricName":"# contigs (>= 5000 bp)","quality":"Equal","values":[0,0,2,3,1,0],"isMain":false},{"metricName":"# contigs (>= 10000 bp)","quality":"Equal","values":[0,0,1,0,1,0],"isMain":false},{"metricName":"# contigs (>= 25000 bp)","quality":"Equal","values":[0,0,1,0,0,0],"isMain":false},{"metricName":"# contigs (>= 50000 bp)","quality":"Equal","values":[0,0,0,0,0,0],"isMain":false},{"metricName":"Largest contig","quality":"More is better","values":[2732,3800,25636,6626,10702,3593],"isMain":true},{"metricName":"Total length","quality":"More is better","values":[533772,645310,702711,675598,718933,291526],"isMain":true},{"metricName":"Total length (>= 1000 bp)","quality":"More is better","values":[121093,197844,161685,136609,233561,54067],"isMain":true},{"metricName":"Total length (>= 5000 bp)","quality":"More is better","values":[0,0,34343,18409,10702,0],"isMain":false},{"metricName":"Total length (>= 10000 bp)","quality":"More is better","values":[0,0,25636,0,10702,0],"isMain":true},{"metricName":"Total length (>= 25000 bp)","quality":"More is better","values":[0,0,25636,0,0,0],"isMain":false},{"metricName":"Total length (>= 50000 bp)","quality":"More is better","values":[0,0,0,0,0,0],"isMain":true},{"metricName":"N50","quality":"More is better","values":[716,789,734,720,817,702],"isMain":false},{"metricName":"N75","quality":"More is better","values":[582,620,598,594,626,577],"isMain":false},{"metricName":"L50","quality":"Less is better","values":[271,282,327,338,302,150],"isMain":false},{"metricName":"L75","quality":"Less is better","values":[480,513,595,599,558,265],"isMain":false},{"metricName":"GC (%)","quality":"Equal","values":["49.42","49.12","49.70","49.73","49.48","48.56"],"isMain":false}]],["K-mer-based statistics",[]],["Predicted genes",[]],["Similarity statistics",[]],["Reference statistics",[{"metricName":"Reference length","quality":"Equal","values":[6479660,6479660,6479660,6479660,6479660,6479660],"isMain":false},{"metricName":"Reference fragments","quality":"Equal","values":[25,25,25,25,25,25],"isMain":false},{"metricName":"Reference GC (%)","quality":"Equal","values":["49.39","49.39","49.39","49.39","49.39","49.39"],"isMain":false}]]],"subreferences":[],"subreports":[],"minContig":500},{"date":"10 February 2020, Monday, 20:33:36","assembliesNames":["megahit-103-df","megahit-113-df","megahit-113-ml","megahit-113-ms","megahit-129-df","metaSPAdes"],"referenceName":"GCF\_000371705.1\_Clos\_bolt\_90A9\_V1","order":[0,1,2,3,4,5],"report":[["Genome statistics",[{"metricName":"Genome fraction (%)","quality":"More is better","values":["7.760","9.569","10.345","10.371","10.095","4.104"],"isMain":true},{"metricName":"Duplication ratio","quality":"Less is better","values":["1.020","1.024","1.020","1.027","1.024","1.026"],"isMain":true},{"metricName":"Largest alignment","quality":"More is better","values":[3084,3139,2237,2272,3800,2141],"isMain":true},{"metricName":"Total aligned length","quality":"More is better","values":[497026,613497,664691,665616,649387,262805],"isMain":true},{"metricName":"NA50","quality":"More is better","values":[696,740,702,712,761,672],"isMain":false},{"metricName":"NA75","quality":"More is better","values":[570,581,577,577,600,560],"isMain":false},{"metricName":"NGA50","quality":"More is better","values":[null,null,null,null,null,null],"isMain":true},{"metricName":"LA50","quality":"Less is better","values":[270,310,365,356,304,156],"isMain":false},{"metricName":"LA75","quality":"Less is better","values":[476,559,631,624,551,265],"isMain":false}]],["Reads mapping",[]],["Misassemblies",[{"metricName":"# misassemblies","quality":"Less is better","values":[0,1,0,0,2,0],"isMain":true},{"metricName":" # relocations","quality":"Less is better","values":[0,0,0,0,0,0],"isMain":false},{"metricName":" # translocations","quality":"Less is better","values":[0,0,0,0,2,0],"isMain":false},{"metricName":" # inversions","quality":"Less is better","values":[0,1,0,0,0,0],"isMain":false},{"metricName":" # interspecies translocations","quality":"Less is better","values":[19,32,9,24,21,7],"isMain":false},{"metricName":"# misassembled contigs","quality":"Less is better","values":[0,1,0,0,1,0],"isMain":false},{"metricName":"Misassembled contigs length","quality":"Less is better","values":[0,959,0,0,2584,0],"isMain":true},{"metricName":"# possibly misassembled contigs","quality":"Less is better","values":[1,0,0,2,0,0],"isMain":false},{"metricName":" # possible misassemblies","quality":"Less is better","values":[1,0,0,2,0,0],"isMain":false},{"metricName":"# local misassemblies","quality":"Less is better","values":[1,2,1,3,2,0],"isMain":false},{"metricName":"# scaffold gap ext. mis.","quality":"Less is better","values":[0,0,0,0,0,0],"isMain":false},{"metricName":"# scaffold gap loc. mis.","quality":"Less is better","values":[0,0,0,0,0,0],"isMain":false},{"metricName":"# unaligned mis. contigs","quality":"Less is better","values":[1,0,0,1,0,0],"isMain":false}]],["Unaligned",[{"metricName":"# fully unaligned contigs","quality":"Less is better","values":[0,2,0,2,2,0],"isMain":false},{"metricName":"Fully unaligned length","quality":"Less is better","values":[0,1377,0,1221,1161,0],"isMain":false},{"metricName":"# partially unaligned contigs","quality":"Less is better","values":[6,8,1,4,4,1],"isMain":false},{"metricName":"Partially unaligned length","quality":"Less is better","values":[10936,25842,1252,10157,4642,623],"isMain":false}]],["Mismatches",[{"metricName":"# mismatches","quality":"Less is better","values":[7975,10453,14576,15246,11591,4429],"isMain":false},{"metricName":"# indels","quality":"Less is better","values":[28,64,58,45,54,14],"isMain":false},{"metricName":"Indels length","quality":"Less is better","values":[73,219,152,175,205,18],"isMain":false},{"metricName":"# mismatches per 100 kbp","quality":"Less is better","values":["1614.07","1715.63","2212.76","2308.74","1803.15","1694.70"],"isMain":true},{"metricName":"# indels per 100 kbp","quality":"Less is better","values":["5.67","10.50","8.80","6.81","8.40","5.36"],"isMain":true},{"metricName":" # indels (<= 5 bp)","quality":"Less is better","values":[26,56,52,40,51,14],"isMain":false},{"metricName":" # indels (> 5 bp)","quality":"Less is better","values":[2,8,6,5,3,0],"isMain":false},{"metricName":"# N's","quality":"Less is better","values":[0,0,0,0,0,0],"isMain":false},{"metricName":"# N's per 100 kbp","quality":"Less is better","values":["0.00","0.00","0.00","0.00","0.00","0.00"],"isMain":true}]],["Statistics without reference",[{"metricName":"# contigs","quality":"Equal","values":[689,803,930,909,832,384],"isMain":true},{"metricName":"# contigs (>= 1000 bp)","quality":"Equal","values":[90,136,108,122,146,29],"isMain":false},{"metricName":"# contigs (>= 5000 bp)","quality":"Equal","values":[1,2,0,1,0,0],"isMain":false},{"metricName":"# contigs (>= 10000 bp)","quality":"Equal","values":[0,1,0,1,0,0],"isMain":false},{"metricName":"# contigs (>= 25000 bp)","quality":"Equal","values":[0,0,0,0,0,0],"isMain":false},{"metricName":"# contigs (>= 50000 bp)","quality":"Equal","values":[0,0,0,0,0,0],"isMain":false},{"metricName":"Largest contig","quality":"More is better","values":[6531,10830,2240,12336,3800,2141],"isMain":true},{"metricName":"Total length","quality":"More is better","values":[514838,651220,673389,689689,664313,268820],"isMain":true},{"metricName":"Total length (>= 1000 bp)","quality":"More is better","values":[125304,201671,129185,162751,202667,35096],"isMain":true},{"metricName":"Total length (>= 5000 bp)","quality":"More is better","values":[6531,17350,0,12336,0,0],"isMain":false},{"metricName":"Total length (>= 10000 bp)","quality":"More is better","values":[0,10830,0,12336,0,0],"isMain":true},{"metricName":"Total length (>= 25000 bp)","quality":"More is better","values":[0,0,0,0,0,0],"isMain":false},{"metricName":"Total length (>= 50000 bp)","quality":"More is better","values":[0,0,0,0,0,0],"isMain":true},{"metricName":"N50","quality":"More is better","values":[715,791,712,735,776,690],"isMain":false},{"metricName":"N75","quality":"More is better","values":[586,620,588,605,623,582],"isMain":false},{"metricName":"L50","quality":"Less is better","values":[253,276,360,337,295,153],"isMain":false},{"metricName":"L75","quality":"Less is better","values":[453,510,621,596,535,259],"isMain":false},{"metricName":"GC (%)","quality":"Equal","values":["49.13","49.41","49.81","49.47","49.21","48.72"],"isMain":false}]],["K-mer-based statistics",[]],["Predicted genes",[]],["Similarity statistics",[]],["Reference statistics",[{"metricName":"Reference length","quality":"Equal","values":[6367468,6367468,6367468,6367468,6367468,6367468],"isMain":false},{"metricName":"Reference fragments","quality":"Equal","values":[18,18,18,18,18,18],"isMain":false},{"metricName":"Reference GC (%)","quality":"Equal","values":["49.56","49.56","49.56","49.56","49.56","49.56"],"isMain":false}]]],"subreferences":[],"subreports":[],"minContig":500},{"date":"10 February 2020, Monday, 20:33:44","assembliesNames":["megahit-103-df","megahit-113-df","megahit-113-ml","megahit-113-ms","megahit-129-df","metaSPAdes"],"referenceName":"GCF\_000373885.1\_ASM37388v1","order":[0,1,2,3,4,5],"report":[["Genome statistics",[{"metricName":"Genome fraction (%)","quality":"More is better","values":["1.139","1.453","1.773","2.105","1.474","0.668"],"isMain":true},{"metricName":"Duplication ratio","quality":"Less is better","values":["1.059","1.091","1.022","1.069","1.064","1.111"],"isMain":true},{"metricName":"Largest alignment","quality":"More is better","values":[1777,1348,1782,1254,1329,3134],"isMain":true},{"metricName":"Total aligned length","quality":"More is better","values":[69516,88383,108087,128061,90038,40641],"isMain":true},{"metricName":"NA50","quality":"More is better","values":[543,549,555,547,531,547],"isMain":false},{"metricName":"NA75","quality":"More is better","values":[509,506,511,499,328,502],"isMain":false},{"metricName":"NGA50","quality":"More is better","values":[null,null,null,null,null,null],"isMain":true},{"metricName":"LA50","quality":"Less is better","values":[58,78,90,116,91,31],"isMain":false},{"metricName":"LA75","quality":"Less is better","values":[95,125,146,187,148,53],"isMain":false}]],["Reads mapping",[]],["Misassemblies",[{"metricName":"# misassemblies","quality":"Less is better","values":[1,2,0,4,2,1],"isMain":true},{"metricName":" # relocations","quality":"Less is better","values":[0,1,0,0,1,0],"isMain":false},{"metricName":" # translocations","quality":"Less is better","values":[1,1,0,3,0,1],"isMain":false},{"metricName":" # inversions","quality":"Less is better","values":[0,0,0,1,1,0],"isMain":false},{"metricName":" # interspecies translocations","quality":"Less is better","values":[13,24,4,27,20,11],"isMain":false},{"metricName":"# misassembled contigs","quality":"Less is better","values":[1,2,0,4,2,1],"isMain":false},{"metricName":"Misassembled contigs length","quality":"Less is better","values":[537,1716,0,2338,1524,574],"isMain":true},{"metricName":"# possibly misassembled contigs","quality":"Less is better","values":[0,0,0,0,0,0],"isMain":false},{"metricName":" # possible misassemblies","quality":"Less is better","values":[0,0,0,0,0,0],"isMain":false},{"metricName":"# local misassemblies","quality":"Less is better","values":[0,0,0,0,1,0],"isMain":false},{"metricName":"# scaffold gap ext. mis.","quality":"Less is better","values":[0,0,0,0,0,0],"isMain":false},{"metricName":"# scaffold gap loc. mis.","quality":"Less is better","values":[0,0,0,0,0,0],"isMain":false},{"metricName":"# unaligned mis. contigs","quality":"Less is better","values":[0,0,0,0,0,0],"isMain":false}]],["Unaligned",[{"metricName":"# fully unaligned contigs","quality":"Less is better","values":[1,0,1,0,0,0],"isMain":false},{"metricName":"Fully unaligned length","quality":"Less is better","values":[712,0,1932,0,0,0],"isMain":false},{"metricName":"# partially unaligned contigs","quality":"Less is better","values":[2,3,1,8,6,1],"isMain":false},{"metricName":"Partially unaligned length","quality":"Less is better","values":[2045,2632,5703,11991,19152,578],"isMain":false}]],["Mismatches",[{"metricName":"# mismatches","quality":"Less is better","values":[1273,1605,2283,2613,1641,752],"isMain":false},{"metricName":"# indels","quality":"Less is better","values":[8,2,8,9,2,5],"isMain":false},{"metricName":"Indels length","quality":"Less is better","values":[10,2,8,9,2,5],"isMain":false},{"metricName":"# mismatches per 100 kbp","quality":"Less is better","values":["1836.44","1815.96","2116.24","2040.42","1829.86","1850.39"],"isMain":true},{"metricName":"# indels per 100 kbp","quality":"Less is better","values":["11.54","2.26","7.42","7.03","2.23","12.30"],"isMain":true},{"metricName":" # indels (<= 5 bp)","quality":"Less is better","values":[8,2,8,9,2,5],"isMain":false},{"metricName":" # indels (> 5 bp)","quality":"Less is better","values":[0,0,0,0,0,0],"isMain":false},{"metricName":"# N's","quality":"Less is better","values":[0,0,0,0,0,0],"isMain":false},{"metricName":"# N's per 100 kbp","quality":"Less is better","values":["0.00","0.00","0.00","0.00","0.00","0.00"],"isMain":true}]],["Statistics without reference",[{"metricName":"# contigs","quality":"Equal","values":[126,164,187,233,162,71],"isMain":true},{"metricName":"# contigs (>= 1000 bp)","quality":"Equal","values":[4,4,3,8,5,3],"isMain":false},{"metricName":"# contigs (>= 5000 bp)","quality":"Equal","values":[0,0,1,0,1,0],"isMain":false},{"metricName":"# contigs (>= 10000 bp)","quality":"Equal","values":[0,0,0,0,1,0],"isMain":false},{"metricName":"# contigs (>= 25000 bp)","quality":"Equal","values":[0,0,0,0,0,0],"isMain":false},{"metricName":"# contigs (>= 50000 bp)","quality":"Equal","values":[0,0,0,0,0,0],"isMain":false},{"metricName":"Largest contig","quality":"More is better","values":[1777,1486,6069,4600,12716,3142],"isMain":true},{"metricName":"Total length","quality":"More is better","values":[76180,99038,117932,148894,114539,45712],"isMain":true},{"metricName":"Total length (>= 1000 bp)","quality":"More is better","values":[5849,5069,9831,14177,20905,5390],"isMain":true},{"metricName":"Total length (>= 5000 bp)","quality":"More is better","values":[0,0,6069,0,12716,0],"isMain":false},{"metricName":"Total length (>= 10000 bp)","quality":"More is better","values":[0,0,0,0,12716,0],"isMain":true},{"metricName":"Total length (>= 25000 bp)","quality":"More is better","values":[0,0,0,0,0,0],"isMain":false},{"metricName":"Total length (>= 50000 bp)","quality":"More is better","values":[0,0,0,0,0,0],"isMain":true},{"metricName":"N50","quality":"More is better","values":[569,577,580,601,601,588],"isMain":false},{"metricName":"N75","quality":"More is better","values":[525,530,527,543,539,535],"isMain":false},{"metricName":"L50","quality":"Less is better","values":[54,71,76,97,57,29],"isMain":false},{"metricName":"L75","quality":"Less is better","values":[89,116,130,162,107,49],"isMain":false},{"metricName":"GC (%)","quality":"Equal","values":["45.45","45.37","46.28","45.26","46.69","44.33"],"isMain":false}]],["K-mer-based statistics",[]],["Predicted genes",[]],["Similarity statistics",[]],["Reference statistics",[{"metricName":"Reference length","quality":"Equal","values":[6084759,6084759,6084759,6084759,6084759,6084759],"isMain":false},{"metricName":"Reference fragments","quality":"Equal","values":[407,407,407,407,407,407],"isMain":false},{"metricName":"Reference GC (%)","quality":"Equal","values":["45.69","45.69","45.69","45.69","45.69","45.69"],"isMain":false}]]],"subreferences":[],"subreports":[],"minContig":500},{"date":"10 February 2020, Monday, 20:34:28","assembliesNames":["megahit-103-df","megahit-113-df","megahit-113-ml","megahit-113-ms","megahit-129-df","metaSPAdes"],"referenceName":"GCF\_000374365.1\_ASM37436v1","order":[0,1,2,3,4,5],"report":[["Genome statistics",[{"metricName":"Genome fraction (%)","quality":"More is better","values":["90.600","91.539","87.719","87.848","92.865","86.343"],"isMain":true},{"metricName":"Duplication ratio","quality":"Less is better","values":["1.063","1.054","1.106","1.108","1.061","1.058"],"isMain":true},{"metricName":"Largest alignment","quality":"More is better","values":[9118,12505,7228,6803,19520,11245],"isMain":true},{"metricName":"Total aligned length","quality":"More is better","values":[4696573,4692440,4722896,4732823,4803197,4453964],"isMain":true},{"metricName":"NG50","quality":"More is better","values":[1930,2688,1257,1300,3183,1497],"isMain":false},{"metricName":"NG75","quality":"More is better","values":[1135,1471,838,876,1949,879],"isMain":false},{"metricName":"NA50","quality":"More is better","values":[1955,2626,1265,1252,2375,1606],"isMain":false},{"metricName":"NA75","quality":"More is better","values":[1181,1428,845,840,1158,995],"isMain":false},{"metricName":"NGA50","quality":"More is better","values":[1888,2569,1239,1242,2606,1478],"isMain":true},{"metricName":"NGA75","quality":"More is better","values":[1108,1371,809,826,1462,850],"isMain":false},{"metricName":"LG50","quality":"Less is better","values":[819,556,1293,1208,407,1013],"isMain":false},{"metricName":"LG75","quality":"Less is better","values":[1644,1177,2496,2364,900,2080],"isMain":false},{"metricName":"LA50","quality":"Less is better","values":[784,564,1254,1256,669,890],"isMain":false},{"metricName":"LA75","quality":"Less is better","values":[1559,1190,2415,2444,1462,1779],"isMain":false},{"metricName":"LGA50","quality":"Less is better","values":[829,582,1308,1274,579,1025],"isMain":true},{"metricName":"LGA75","quality":"Less is better","values":[1672,1241,2537,2485,1207,2119],"isMain":false}]],["Reads mapping",[]],["Misassemblies",[{"metricName":"# misassemblies","quality":"Less is better","values":[9,9,14,15,10,7],"isMain":true},{"metricName":" # relocations","quality":"Less is better","values":[0,2,1,1,0,0],"isMain":false},{"metricName":" # translocations","quality":"Less is better","values":[6,6,9,9,9,6],"isMain":false},{"metricName":" # inversions","quality":"Less is better","values":[3,1,4,5,1,1],"isMain":false},{"metricName":" # interspecies translocations","quality":"Less is better","values":[70,109,48,106,96,29],"isMain":false},{"metricName":"# misassembled contigs","quality":"Less is better","values":[9,9,14,15,10,6],"isMain":false},{"metricName":"Misassembled contigs length","quality":"Less is better","values":[13577,20741,33470,29900,47136,26144],"isMain":true},{"metricName":"# possibly misassembled contigs","quality":"Less is better","values":[0,1,3,1,2,0],"isMain":false},{"metricName":" # possible misassemblies","quality":"Less is better","values":[0,1,3,1,2,0],"isMain":false},{"metricName":"# local misassemblies","quality":"Less is better","values":[2,2,4,5,5,1],"isMain":false},{"metricName":"# scaffold gap ext. mis.","quality":"Less is better","values":[0,0,0,0,0,0],"isMain":false},{"metricName":"# scaffold gap loc. mis.","quality":"Less is better","values":[0,0,0,0,0,0],"isMain":false},{"metricName":"# unaligned mis. contigs","quality":"Less is better","values":[2,0,1,1,1,0],"isMain":false}]],["Unaligned",[{"metricName":"# fully unaligned contigs","quality":"Less is better","values":[0,2,4,3,1,0],"isMain":false},{"metricName":"Fully unaligned length","quality":"Less is better","values":[0,2156,5541,2098,646,0],"isMain":false},{"metricName":"# partially unaligned contigs","quality":"Less is better","values":[7,24,9,25,37,8],"isMain":false},{"metricName":"Partially unaligned length","quality":"Less is better","values":[11496,77694,7386,83055,513818,10344],"isMain":false}]],["Mismatches",[{"metricName":"# mismatches","quality":"Less is better","values":[33845,31101,66630,68327,33751,32396],"isMain":false},{"metricName":"# indels","quality":"Less is better","values":[97,105,88,98,134,69],"isMain":false},{"metricName":"Indels length","quality":"Less is better","values":[380,565,508,352,845,333],"isMain":false},{"metricName":"# mismatches per 100 kbp","quality":"Less is better","values":["759.75","690.99","1544.83","1581.87","739.16","763.08"],"isMain":true},{"metricName":"# indels per 100 kbp","quality":"Less is better","values":["2.18","2.33","2.04","2.27","2.93","1.63"],"isMain":true},{"metricName":" # indels (<= 5 bp)","quality":"Less is better","values":[89,94,80,92,117,63],"isMain":false},{"metricName":" # indels (> 5 bp)","quality":"Less is better","values":[8,11,8,6,17,6],"isMain":false},{"metricName":"# N's","quality":"Less is better","values":[0,0,0,0,0,0],"isMain":false},{"metricName":"# N's per 100 kbp","quality":"Less is better","values":["0.00","0.00","0.00","0.00","0.00","0.00"],"isMain":true}]],["Statistics without reference",[{"metricName":"# contigs","quality":"Equal","values":[2997,2415,4181,4129,2470,3276],"isMain":true},{"metricName":"# contigs (>= 1000 bp)","quality":"Equal","values":[1857,1656,1915,1930,1686,1797],"isMain":false},{"metricName":"# contigs (>= 5000 bp)","quality":"Equal","values":[52,138,7,6,144,29],"isMain":false},{"metricName":"# contigs (>= 10000 bp)","quality":"Equal","values":[0,11,0,2,12,1],"isMain":false},{"metricName":"# contigs (>= 25000 bp)","quality":"Equal","values":[0,1,0,1,3,0],"isMain":false},{"metricName":"# contigs (>= 50000 bp)","quality":"Equal","values":[0,0,0,0,2,0],"isMain":false},{"metricName":"Largest contig","quality":"More is better","values":[9118,41267,8321,48640,298133,11593],"isMain":true},{"metricName":"Total length","quality":"More is better","values":[4744686,4821578,4781783,4870828,5359890,4501963],"isMain":true},{"metricName":"Total length (>= 1000 bp)","quality":"More is better","values":[3915083,4274647,3156583,3281635,4797742,3423709],"isMain":true},{"metricName":"Total length (>= 5000 bp)","quality":"More is better","values":[320999,965003,44511,90357,1428408,176632],"isMain":false},{"metricName":"Total length (>= 10000 bp)","quality":"More is better","values":[0,151830,0,64371,575343,11593],"isMain":true},{"metricName":"Total length (>= 25000 bp)","quality":"More is better","values":[0,41267,0,48640,452333,0],"isMain":false},{"metricName":"Total length (>= 50000 bp)","quality":"More is better","values":[0,0,0,0,411048,0],"isMain":true},{"metricName":"N50","quality":"More is better","values":[1987,2727,1284,1312,2946,1623],"isMain":false},{"metricName":"N75","quality":"More is better","values":[1219,1526,869,886,1664,1026],"isMain":false},{"metricName":"L50","quality":"Less is better","values":[775,539,1240,1190,479,880],"isMain":false},{"metricName":"L75","quality":"Less is better","values":[1534,1129,2377,2324,1083,1751],"isMain":false},{"metricName":"GC (%)","quality":"Equal","values":["47.77","47.72","47.82","47.74","47.43","47.86"],"isMain":false}]],["K-mer-based statistics",[]],["Predicted genes",[]],["Similarity statistics",[]],["Reference statistics",[{"metricName":"Reference length","quality":"Equal","values":[4916921,4916921,4916921,4916921,4916921,4916921],"isMain":false},{"metricName":"Reference fragments","quality":"Equal","values":[105,105,105,105,105,105],"isMain":false},{"metricName":"Reference GC (%)","quality":"Equal","values":["47.70","47.70","47.70","47.70","47.70","47.70"],"isMain":false}]]],"subreferences":[],"subreports":[],"minContig":500},{"date":"10 February 2020, Monday, 20:33:57","assembliesNames":["megahit-103-df","megahit-113-df","megahit-113-ml","megahit-113-ms","megahit-129-df","metaSPAdes"],"referenceName":"GCF\_000374505.1\_ASM37450v1","order":[0,1,2,3,4,5],"report":[["Genome statistics",[{"metricName":"Genome fraction (%)","quality":"More is better","values":["27.759","28.259","30.797","30.314","30.555","17.308"],"isMain":true},{"metricName":"Duplication ratio","quality":"Less is better","values":["1.038","1.029","1.037","1.040","1.029","1.032"],"isMain":true},{"metricName":"Largest alignment","quality":"More is better","values":[2096,12656,3383,8491,8475,18513],"isMain":true},{"metricName":"Total aligned length","quality":"More is better","values":[1089826,1104346,1218465,1196361,1198056,676951],"isMain":true},{"metricName":"NA50","quality":"More is better","values":[680,726,687,701,748,643],"isMain":false},{"metricName":"NA75","quality":"More is better","values":[566,581,577,575,597,552],"isMain":false},{"metricName":"NGA50","quality":"More is better","values":[null,null,null,null,null,null],"isMain":true},{"metricName":"LA50","quality":"Less is better","values":[628,561,678,640,600,386],"isMain":false},{"metricName":"LA75","quality":"Less is better","values":[1084,1006,1172,1128,1064,679],"isMain":false}]],["Reads mapping",[]],["Misassemblies",[{"metricName":"# misassemblies","quality":"Less is better","values":[6,6,0,5,5,3],"isMain":true},{"metricName":" # relocations","quality":"Less is better","values":[2,2,0,2,2,2],"isMain":false},{"metricName":" # translocations","quality":"Less is better","values":[2,3,0,2,2,1],"isMain":false},{"metricName":" # inversions","quality":"Less is better","values":[2,1,0,1,1,0],"isMain":false},{"metricName":" # interspecies translocations","quality":"Less is better","values":[59,67,16,44,51,27],"isMain":false},{"metricName":"# misassembled contigs","quality":"Less is better","values":[6,6,0,4,5,3],"isMain":false},{"metricName":"Misassembled contigs length","quality":"Less is better","values":[3832,3883,0,14759,3636,3815],"isMain":true},{"metricName":"# possibly misassembled contigs","quality":"Less is better","values":[0,0,0,0,0,0],"isMain":false},{"metricName":" # possible misassemblies","quality":"Less is better","values":[0,0,0,0,0,0],"isMain":false},{"metricName":"# local misassemblies","quality":"Less is better","values":[3,4,3,2,1,1],"isMain":false},{"metricName":"# scaffold gap ext. mis.","quality":"Less is better","values":[0,0,0,0,0,0],"isMain":false},{"metricName":"# scaffold gap loc. mis.","quality":"Less is better","values":[0,0,0,0,0,0],"isMain":false},{"metricName":"# unaligned mis. contigs","quality":"Less is better","values":[0,0,0,1,0,0],"isMain":false}]],["Unaligned",[{"metricName":"# fully unaligned contigs","quality":"Less is better","values":[3,5,1,2,5,0],"isMain":false},{"metricName":"Fully unaligned length","quality":"Less is better","values":[2018,4998,696,1442,5627,0],"isMain":false},{"metricName":"# partially unaligned contigs","quality":"Less is better","values":[8,18,2,8,14,4],"isMain":false},{"metricName":"Partially unaligned length","quality":"Less is better","values":[8906,24413,1808,8528,14147,6000],"isMain":false}]],["Mismatches",[{"metricName":"# mismatches","quality":"Less is better","values":[15566,14604,22999,23201,16339,9641],"isMain":false},{"metricName":"# indels","quality":"Less is better","values":[34,49,30,47,27,24],"isMain":false},{"metricName":"Indels length","quality":"Less is better","values":[36,54,36,118,28,54],"isMain":false},{"metricName":"# mismatches per 100 kbp","quality":"Less is better","values":["1450.97","1337.21","1932.38","1980.40","1383.67","1441.33"],"isMain":true},{"metricName":"# indels per 100 kbp","quality":"Less is better","values":["3.17","4.49","2.52","4.01","2.29","3.59"],"isMain":true},{"metricName":" # indels (<= 5 bp)","quality":"Less is better","values":[34,49,30,42,27,23],"isMain":false},{"metricName":" # indels (> 5 bp)","quality":"Less is better","values":[0,0,0,5,0,1],"isMain":false},{"metricName":"# N's","quality":"Less is better","values":[0,0,0,0,0,0],"isMain":false},{"metricName":"# N's per 100 kbp","quality":"Less is better","values":["0.00","0.00","0.00","0.00","0.00","0.00"],"isMain":true}]],["Statistics without reference",[{"metricName":"# contigs","quality":"Equal","values":[1569,1478,1725,1658,1585,981],"isMain":true},{"metricName":"# contigs (>= 1000 bp)","quality":"Equal","values":[163,225,168,178,265,80],"isMain":false},{"metricName":"# contigs (>= 5000 bp)","quality":"Equal","values":[0,3,0,4,1,1],"isMain":false},{"metricName":"# contigs (>= 10000 bp)","quality":"Equal","values":[0,1,0,0,0,1],"isMain":false},{"metricName":"# contigs (>= 25000 bp)","quality":"Equal","values":[0,0,0,0,0,0],"isMain":false},{"metricName":"# contigs (>= 50000 bp)","quality":"Equal","values":[0,0,0,0,0,0],"isMain":false},{"metricName":"Largest contig","quality":"More is better","values":[3862,12656,3383,8491,8488,18563],"isMain":true},{"metricName":"Total length","quality":"More is better","values":[1124215,1152881,1236853,1227880,1234973,696457],"isMain":true},{"metricName":"Total length (>= 1000 bp)","quality":"More is better","values":[200639,314557,211164,245590,346247,120603],"isMain":true},{"metricName":"Total length (>= 5000 bp)","quality":"More is better","values":[0,27219,0,28128,8488,18563],"isMain":false},{"metricName":"Total length (>= 10000 bp)","quality":"More is better","values":[0,12656,0,0,0,18563],"isMain":true},{"metricName":"Total length (>= 25000 bp)","quality":"More is better","values":[0,0,0,0,0,0],"isMain":false},{"metricName":"Total length (>= 50000 bp)","quality":"More is better","values":[0,0,0,0,0,0],"isMain":true},{"metricName":"N50","quality":"More is better","values":[700,756,702,723,769,665],"isMain":false},{"metricName":"N75","quality":"More is better","values":[586,615,586,592,618,571],"isMain":false},{"metricName":"L50","quality":"Less is better","values":[609,530,669,621,579,371],"isMain":false},{"metricName":"L75","quality":"Less is better","values":[1049,954,1154,1093,1028,655],"isMain":false},{"metricName":"GC (%)","quality":"Equal","values":["57.70","57.35","58.01","57.38","57.34","57.10"],"isMain":false}]],["K-mer-based statistics",[]],["Predicted genes",[]],["Similarity statistics",[]],["Reference statistics",[{"metricName":"Reference length","quality":"Equal","values":[3864643,3864643,3864643,3864643,3864643,3864643],"isMain":false},{"metricName":"Reference fragments","quality":"Equal","values":[32,32,32,32,32,32],"isMain":false},{"metricName":"Reference GC (%)","quality":"Equal","values":["57.85","57.85","57.85","57.85","57.85","57.85"],"isMain":false}]]],"subreferences":[],"subreports":[],"minContig":500},{"date":"10 February 2020, Monday, 20:33:39","assembliesNames":["megahit-103-df","megahit-113-df","megahit-113-ml","megahit-113-ms","megahit-129-df","metaSPAdes"],"referenceName":"GCF\_000374585.1\_ASM37458v1","order":[0,1,2,3,4,5],"report":[["Genome statistics",[{"metricName":"Genome fraction (%)","quality":"More is better","values":["1.855","2.543","2.881","3.755","2.824","0.968"],"isMain":true},{"metricName":"Duplication ratio","quality":"Less is better","values":["1.065","1.049","1.032","1.071","1.055","1.078"],"isMain":true},{"metricName":"Largest alignment","quality":"More is better","values":[1365,1060,3050,1437,1108,913],"isMain":true},{"metricName":"Total aligned length","quality":"More is better","values":[67522,92104,104696,136133,102384,35065],"isMain":true},{"metricName":"NA50","quality":"More is better","values":[560,549,566,544,559,540],"isMain":false},{"metricName":"NA75","quality":"More is better","values":[513,506,522,503,505,503],"isMain":false},{"metricName":"NGA50","quality":"More is better","values":[null,null,null,null,null,null],"isMain":true},{"metricName":"LA50","quality":"Less is better","values":[57,80,80,122,87,31],"isMain":false},{"metricName":"LA75","quality":"Less is better","values":[91,129,129,197,141,50],"isMain":false}]],["Reads mapping",[]],["Misassemblies",[{"metricName":"# misassemblies","quality":"Less is better","values":[0,0,1,1,0,1],"isMain":true},{"metricName":" # relocations","quality":"Less is better","values":[0,0,0,0,0,1],"isMain":false},{"metricName":" # translocations","quality":"Less is better","values":[0,0,1,1,0,0],"isMain":false},{"metricName":" # inversions","quality":"Less is better","values":[0,0,0,0,0,0],"isMain":false},{"metricName":" # interspecies translocations","quality":"Less is better","values":[14,20,7,37,21,10],"isMain":false},{"metricName":"# misassembled contigs","quality":"Less is better","values":[0,0,1,1,0,1],"isMain":false},{"metricName":"Misassembled contigs length","quality":"Less is better","values":[0,0,669,888,0,532],"isMain":true},{"metricName":"# possibly misassembled contigs","quality":"Less is better","values":[0,0,0,0,0,0],"isMain":false},{"metricName":" # possible misassemblies","quality":"Less is better","values":[0,0,0,0,0,0],"isMain":false},{"metricName":"# local misassemblies","quality":"Less is better","values":[1,0,0,0,1,0],"isMain":false},{"metricName":"# scaffold gap ext. mis.","quality":"Less is better","values":[0,0,0,0,0,0],"isMain":false},{"metricName":"# scaffold gap loc. mis.","quality":"Less is better","values":[0,0,0,0,0,0],"isMain":false},{"metricName":"# unaligned mis. contigs","quality":"Less is better","values":[0,0,0,0,0,0],"isMain":false}]],["Unaligned",[{"metricName":"# fully unaligned contigs","quality":"Less is better","values":[1,0,0,1,0,0],"isMain":false},{"metricName":"Fully unaligned length","quality":"Less is better","values":[540,0,0,589,0,0],"isMain":false},{"metricName":"# partially unaligned contigs","quality":"Less is better","values":[2,7,0,13,7,3],"isMain":false},{"metricName":"Partially unaligned length","quality":"Less is better","values":[1182,7345,0,10776,7838,2240],"isMain":false}]],["Mismatches",[{"metricName":"# mismatches","quality":"Less is better","values":[1231,1615,2151,2988,1810,629],"isMain":false},{"metricName":"# indels","quality":"Less is better","values":[6,3,2,5,3,2],"isMain":false},{"metricName":"Indels length","quality":"Less is better","values":[64,61,2,5,61,2],"isMain":false},{"metricName":"# mismatches per 100 kbp","quality":"Less is better","values":["1834.25","1755.38","2063.76","2199.55","1771.89","1796.12"],"isMain":true},{"metricName":"# indels per 100 kbp","quality":"Less is better","values":["8.94","3.26","1.92","3.68","2.94","5.71"],"isMain":true},{"metricName":" # indels (<= 5 bp)","quality":"Less is better","values":[5,2,2,5,2,2],"isMain":false},{"metricName":" # indels (> 5 bp)","quality":"Less is better","values":[1,1,0,0,1,0],"isMain":false},{"metricName":"# N's","quality":"Less is better","values":[0,0,0,0,0,0],"isMain":false},{"metricName":"# N's per 100 kbp","quality":"Less is better","values":["0.00","0.00","0.00","0.00","0.00","0.00"],"isMain":true}]],["Statistics without reference",[{"metricName":"# contigs","quality":"Equal","values":[121,164,178,252,180,65],"isMain":true},{"metricName":"# contigs (>= 1000 bp)","quality":"Equal","values":[1,7,1,10,9,1],"isMain":false},{"metricName":"# contigs (>= 5000 bp)","quality":"Equal","values":[0,0,0,0,0,0],"isMain":false},{"metricName":"# contigs (>= 10000 bp)","quality":"Equal","values":[0,0,0,0,0,0],"isMain":false},{"metricName":"# contigs (>= 25000 bp)","quality":"Equal","values":[0,0,0,0,0,0],"isMain":false},{"metricName":"# contigs (>= 50000 bp)","quality":"Equal","values":[0,0,0,0,0,0],"isMain":false},{"metricName":"Largest contig","quality":"More is better","values":[1365,2756,3050,1598,2375,1336],"isMain":true},{"metricName":"Total length","quality":"More is better","values":[73169,103869,107594,156899,115641,39987],"isMain":true},{"metricName":"Total length (>= 1000 bp)","quality":"More is better","values":[1365,10184,3050,12279,12635,1336],"isMain":true},{"metricName":"Total length (>= 5000 bp)","quality":"More is better","values":[0,0,0,0,0,0],"isMain":false},{"metricName":"Total length (>= 10000 bp)","quality":"More is better","values":[0,0,0,0,0,0],"isMain":true},{"metricName":"Total length (>= 25000 bp)","quality":"More is better","values":[0,0,0,0,0,0],"isMain":false},{"metricName":"Total length (>= 50000 bp)","quality":"More is better","values":[0,0,0,0,0,0],"isMain":true},{"metricName":"N50","quality":"More is better","values":[584,583,582,589,610,602],"isMain":false},{"metricName":"N75","quality":"More is better","values":[538,535,534,534,543,526],"isMain":false},{"metricName":"L50","quality":"Less is better","values":[54,68,78,106,74,28],"isMain":false},{"metricName":"L75","quality":"Less is better","values":[86,114,126,177,125,46],"isMain":false},{"metricName":"GC (%)","quality":"Equal","values":["47.08","46.73","47.11","46.28","46.35","46.45"],"isMain":false}]],["K-mer-based statistics",[]],["Predicted genes",[]],["Similarity statistics",[]],["Reference statistics",[{"metricName":"Reference length","quality":"Equal","values":[3617816,3617816,3617816,3617816,3617816,3617816],"isMain":false},{"metricName":"Reference fragments","quality":"Equal","values":[47,47,47,47,47,47],"isMain":false},{"metricName":"Reference GC (%)","quality":"Equal","values":["46.86","46.86","46.86","46.86","46.86","46.86"],"isMain":false}]]],"subreferences":[],"subreports":[],"minContig":500},{"date":"10 February 2020, Monday, 20:33:31","assembliesNames":["megahit-103-df","megahit-113-df","megahit-113-ml","megahit-113-ms","megahit-129-df","metaSPAdes"],"referenceName":"GCF\_000379925.1\_ASM37992v1","order":[0,1,2,3,4,5],"report":[["Genome statistics",[{"metricName":"Genome fraction (%)","quality":"More is better","values":["0.122","0.269","0.318","0.473","0.264","0.106"],"isMain":true},{"metricName":"Duplication ratio","quality":"Less is better","values":["1.001","1.080","1.020","1.129","1.082","1.240"],"isMain":true},{"metricName":"Largest alignment","quality":"More is better","values":[947,1294,803,760,999,593],"isMain":true},{"metricName":"Total aligned length","quality":"More is better","values":[3077,6768,8006,11881,6631,2653],"isMain":true},{"metricName":"NA50","quality":"More is better","values":[541,563,548,535,606,527],"isMain":false},{"metricName":"NA75","quality":"More is better","values":[527,514,519,494,546,356],"isMain":false},{"metricName":"NGA50","quality":"More is better","values":[null,null,null,null,null,null],"isMain":true},{"metricName":"LA50","quality":"Less is better","values":[3,5,7,12,5,3],"isMain":false},{"metricName":"LA75","quality":"Less is better","values":[4,8,11,19,8,5],"isMain":false}]],["Reads mapping",[]],["Misassemblies",[{"metricName":"# misassemblies","quality":"Less is better","values":[0,0,0,0,0,0],"isMain":true},{"metricName":" # relocations","quality":"Less is better","values":[0,0,0,0,0,0],"isMain":false},{"metricName":" # translocations","quality":"Less is better","values":[0,0,0,0,0,0],"isMain":false},{"metricName":" # inversions","quality":"Less is better","values":[0,0,0,0,0,0],"isMain":false},{"metricName":" # interspecies translocations","quality":"Less is better","values":[0,2,0,5,2,2],"isMain":false},{"metricName":"# misassembled contigs","quality":"Less is better","values":[0,0,0,0,0,0],"isMain":false},{"metricName":"Misassembled contigs length","quality":"Less is better","values":[0,0,0,0,0,0],"isMain":true},{"metricName":"# possibly misassembled contigs","quality":"Less is better","values":[0,0,0,0,0,0],"isMain":false},{"metricName":" # possible misassemblies","quality":"Less is better","values":[0,0,0,0,0,0],"isMain":false},{"metricName":"# local misassemblies","quality":"Less is better","values":[0,0,0,0,0,0],"isMain":false},{"metricName":"# scaffold gap ext. mis.","quality":"Less is better","values":[0,0,0,0,0,0],"isMain":false},{"metricName":"# scaffold gap loc. mis.","quality":"Less is better","values":[0,0,0,0,0,0],"isMain":false},{"metricName":"# unaligned mis. contigs","quality":"Less is better","values":[0,0,0,0,0,0],"isMain":false}]],["Unaligned",[{"metricName":"# fully unaligned contigs","quality":"Less is better","values":[0,0,0,0,0,0],"isMain":false},{"metricName":"Fully unaligned length","quality":"Less is better","values":[0,0,0,0,0,0],"isMain":false},{"metricName":"# partially unaligned contigs","quality":"Less is better","values":[0,0,0,1,0,0],"isMain":false},{"metricName":"Partially unaligned length","quality":"Less is better","values":[0,0,0,556,0,0],"isMain":false}]],["Mismatches",[{"metricName":"# mismatches","quality":"Less is better","values":[58,105,149,256,98,44],"isMain":false},{"metricName":"# indels","quality":"Less is better","values":[0,0,0,0,0,0],"isMain":false},{"metricName":"Indels length","quality":"Less is better","values":[0,0,0,0,0,0],"isMain":false},{"metricName":"# mismatches per 100 kbp","quality":"Less is better","values":["1884.95","1551.42","1861.10","2154.70","1477.91","1658.50"],"isMain":true},{"metricName":"# indels per 100 kbp","quality":"Less is better","values":["0.00","0.00","0.00","0.00","0.00","0.00"],"isMain":true},{"metricName":" # indels (<= 5 bp)","quality":"Less is better","values":[0,0,0,0,0,0],"isMain":false},{"metricName":" # indels (> 5 bp)","quality":"Less is better","values":[0,0,0,0,0,0],"isMain":false},{"metricName":"# N's","quality":"Less is better","values":[0,0,0,0,0,0],"isMain":false},{"metricName":"# N's per 100 kbp","quality":"Less is better","values":["0.00","0.00","0.00","0.00","0.00","0.00"],"isMain":true}]],["Statistics without reference",[{"metricName":"# contigs","quality":"Equal","values":[5,11,14,24,11,6],"isMain":true},{"metricName":"# contigs (>= 1000 bp)","quality":"Equal","values":[0,1,0,0,0,0],"isMain":false},{"metricName":"# contigs (>= 5000 bp)","quality":"Equal","values":[0,0,0,0,0,0],"isMain":false},{"metricName":"# contigs (>= 10000 bp)","quality":"Equal","values":[0,0,0,0,0,0],"isMain":false},{"metricName":"# contigs (>= 25000 bp)","quality":"Equal","values":[0,0,0,0,0,0],"isMain":false},{"metricName":"# contigs (>= 50000 bp)","quality":"Equal","values":[0,0,0,0,0,0],"isMain":false},{"metricName":"Largest contig","quality":"More is better","values":[947,1294,803,762,999,608],"isMain":true},{"metricName":"Total length","quality":"More is better","values":[3079,7309,8169,13970,7176,3290],"isMain":true},{"metricName":"Total length (>= 1000 bp)","quality":"More is better","values":[0,1294,0,0,0,0],"isMain":true},{"metricName":"Total length (>= 5000 bp)","quality":"More is better","values":[0,0,0,0,0,0],"isMain":false},{"metricName":"Total length (>= 10000 bp)","quality":"More is better","values":[0,0,0,0,0,0],"isMain":true},{"metricName":"Total length (>= 25000 bp)","quality":"More is better","values":[0,0,0,0,0,0],"isMain":false},{"metricName":"Total length (>= 50000 bp)","quality":"More is better","values":[0,0,0,0,0,0],"isMain":true},{"metricName":"N50","quality":"More is better","values":[541,613,552,593,613,553],"isMain":false},{"metricName":"N75","quality":"More is better","values":[527,527,527,526,576,508],"isMain":false},{"metricName":"L50","quality":"Less is better","values":[3,5,7,11,5,3],"isMain":false},{"metricName":"L75","quality":"Less is better","values":[4,8,11,18,8,5],"isMain":false},{"metricName":"GC (%)","quality":"Equal","values":["46.31","46.74","46.19","45.02","46.31","46.87"],"isMain":false}]],["K-mer-based statistics",[]],["Predicted genes",[]],["Similarity statistics",[]],["Reference statistics",[{"metricName":"Reference length","quality":"Equal","values":[2514463,2514463,2514463,2514463,2514463,2514463],"isMain":false},{"metricName":"Reference fragments","quality":"Equal","values":[179,179,179,179,179,179],"isMain":false},{"metricName":"Reference GC (%)","quality":"Equal","values":["45.69","45.69","45.69","45.69","45.69","45.69"],"isMain":false}]]],"subreferences":[],"subreports":[],"minContig":500},{"date":"10 February 2020, Monday, 20:33:38","assembliesNames":["megahit-103-df","megahit-113-df","megahit-113-ml","megahit-113-ms","megahit-129-df","metaSPAdes"],"referenceName":"GCF\_000379965.1\_ASM37996v1","order":[0,1,2,3,4,5],"report":[["Genome statistics",[{"metricName":"Genome fraction (%)","quality":"More is better","values":["0.069","0.066","0.081","0.165","0.120","0.026"],"isMain":true},{"metricName":"Duplication ratio","quality":"Less is better","values":["1.000","1.573","1.284","1.398","1.370","1.642"],"isMain":true},{"metricName":"Largest alignment","quality":"More is better","values":[547,535,664,721,845,537],"isMain":true},{"metricName":"Total aligned length","quality":"More is better","values":[1821,1744,2158,4372,3181,698],"isMain":true},{"metricName":"NA50","quality":"More is better","values":[null,null,616,220,null,161],"isMain":false},{"metricName":"NA75","quality":"More is better","values":[null,null,237,null,null,null],"isMain":false},{"metricName":"NGA50","quality":"More is better","values":[null,null,null,null,null,null],"isMain":true},{"metricName":"LA50","quality":"Less is better","values":[null,null,3,8,null,2],"isMain":false},{"metricName":"LA75","quality":"Less is better","values":[null,null,4,null,null,null],"isMain":false}]],["Reads mapping",[]],["Misassemblies",[{"metricName":"# misassemblies","quality":"Less is better","values":[0,0,0,0,0,0],"isMain":true},{"metricName":" # relocations","quality":"Less is better","values":[0,0,0,0,0,0],"isMain":false},{"metricName":" # translocations","quality":"Less is better","values":[0,0,0,0,0,0],"isMain":false},{"metricName":" # inversions","quality":"Less is better","values":[0,0,0,0,0,0],"isMain":false},{"metricName":" # interspecies translocations","quality":"Less is better","values":[5,8,1,5,6,1],"isMain":false},{"metricName":"# misassembled contigs","quality":"Less is better","values":[0,0,0,0,0,0],"isMain":false},{"metricName":"Misassembled contigs length","quality":"Less is better","values":[0,0,0,0,0,0],"isMain":true},{"metricName":"# possibly misassembled contigs","quality":"Less is better","values":[0,0,0,0,0,0],"isMain":false},{"metricName":" # possible misassemblies","quality":"Less is better","values":[0,0,0,0,0,0],"isMain":false},{"metricName":"# local misassemblies","quality":"Less is better","values":[0,0,0,0,1,0],"isMain":false},{"metricName":"# scaffold gap ext. mis.","quality":"Less is better","values":[0,0,0,0,0,0],"isMain":false},{"metricName":"# scaffold gap loc. mis.","quality":"Less is better","values":[0,0,0,0,0,0],"isMain":false},{"metricName":"# unaligned mis. contigs","quality":"Less is better","values":[0,0,0,0,0,0],"isMain":false}]],["Unaligned",[{"metricName":"# fully unaligned contigs","quality":"Less is better","values":[0,2,0,0,0,0],"isMain":false},{"metricName":"Fully unaligned length","quality":"Less is better","values":[0,1396,0,0,0,0],"isMain":false},{"metricName":"# partially unaligned contigs","quality":"Less is better","values":[5,4,0,2,3,0],"isMain":false},{"metricName":"Partially unaligned length","quality":"Less is better","values":[6013,3102,0,1506,3552,0],"isMain":false}]],["Mismatches",[{"metricName":"# mismatches","quality":"Less is better","values":[50,47,69,145,93,14],"isMain":false},{"metricName":"# indels","quality":"Less is better","values":[0,0,2,0,0,0],"isMain":false},{"metricName":"Indels length","quality":"Less is better","values":[0,0,2,0,0,0],"isMain":false},{"metricName":"# mismatches per 100 kbp","quality":"Less is better","values":["2745.74","2694.95","3197.41","3316.56","2923.61","2005.73"],"isMain":true},{"metricName":"# indels per 100 kbp","quality":"Less is better","values":["0.00","0.00","92.68","0.00","0.00","0.00"],"isMain":true},{"metricName":" # indels (<= 5 bp)","quality":"Less is better","values":[0,0,2,0,0,0],"isMain":false},{"metricName":" # indels (> 5 bp)","quality":"Less is better","values":[0,0,0,0,0,0],"isMain":false},{"metricName":"# N's","quality":"Less is better","values":[0,0,0,0,0,0],"isMain":false},{"metricName":"# N's per 100 kbp","quality":"Less is better","values":["0.00","0.00","0.00","0.00","0.00","0.00"],"isMain":true}]],["Statistics without reference",[{"metricName":"# contigs","quality":"Equal","values":[7,10,4,11,8,2],"isMain":true},{"metricName":"# contigs (>= 1000 bp)","quality":"Equal","values":[2,1,0,1,3,0],"isMain":false},{"metricName":"# contigs (>= 5000 bp)","quality":"Equal","values":[0,0,0,0,0,0],"isMain":false},{"metricName":"# contigs (>= 10000 bp)","quality":"Equal","values":[0,0,0,0,0,0],"isMain":false},{"metricName":"# contigs (>= 25000 bp)","quality":"Equal","values":[0,0,0,0,0,0],"isMain":false},{"metricName":"# contigs (>= 50000 bp)","quality":"Equal","values":[0,0,0,0,0,0],"isMain":false},{"metricName":"Largest contig","quality":"More is better","values":[3283,1151,990,1044,2055,609],"isMain":true},{"metricName":"Total length","quality":"More is better","values":[7834,7242,2771,7619,7911,1146],"isMain":true},{"metricName":"Total length (>= 1000 bp)","quality":"More is better","values":[4434,1151,0,1044,4376,0],"isMain":true},{"metricName":"Total length (>= 5000 bp)","quality":"More is better","values":[0,0,0,0,0,0],"isMain":false},{"metricName":"Total length (>= 10000 bp)","quality":"More is better","values":[0,0,0,0,0,0],"isMain":true},{"metricName":"Total length (>= 25000 bp)","quality":"More is better","values":[0,0,0,0,0,0],"isMain":false},{"metricName":"Total length (>= 50000 bp)","quality":"More is better","values":[0,0,0,0,0,0],"isMain":true},{"metricName":"N50","quality":"More is better","values":[1151,711,664,682,1151,609],"isMain":false},{"metricName":"N75","quality":"More is better","values":[816,591,616,613,754,537],"isMain":false},{"metricName":"L50","quality":"Less is better","values":[2,4,2,5,3,1],"isMain":false},{"metricName":"L75","quality":"Less is better","values":[4,7,3,8,5,2],"isMain":false},{"metricName":"GC (%)","quality":"Equal","values":["38.37","39.44","41.54","40.32","40.79","43.80"],"isMain":false}]],["K-mer-based statistics",[]],["Predicted genes",[]],["Similarity statistics",[]],["Reference statistics",[{"metricName":"Reference length","quality":"Equal","values":[2648549,2648549,2648549,2648549,2648549,2648549],"isMain":false},{"metricName":"Reference fragments","quality":"Equal","values":[19,19,19,19,19,19],"isMain":false},{"metricName":"Reference GC (%)","quality":"Equal","values":["38.36","38.36","38.36","38.36","38.36","38.36"],"isMain":false}]]],"subreferences":[],"subreports":[],"minContig":500},{"date":"10 February 2020, Monday, 20:34:41","assembliesNames":["megahit-103-df","megahit-113-df","megahit-113-ml","megahit-113-ms","megahit-129-df","metaSPAdes"],"referenceName":"GCF\_000381365.1\_Bact\_saly\_DSM\_18765\_V1","order":[0,1,2,3,4,5],"report":[["Genome statistics",[{"metricName":"Genome fraction (%)","quality":"More is better","values":["75.752","79.583","73.396","72.323","81.490","67.866"],"isMain":true},{"metricName":"Duplication ratio","quality":"Less is better","values":["1.051","1.046","1.077","1.080","1.051","1.050"],"isMain":true},{"metricName":"Largest alignment","quality":"More is better","values":[7362,8203,6360,6792,9115,11274],"isMain":true},{"metricName":"Total aligned length","quality":"More is better","values":[4187854,4372616,4155955,4097660,4501201,3742160],"isMain":true},{"metricName":"NG50","quality":"More is better","values":[1082,1391,864,884,1518,817],"isMain":false},{"metricName":"NG75","quality":"More is better","values":[583,769,550,550,842,null],"isMain":false},{"metricName":"NA50","quality":"More is better","values":[1307,1508,1006,1024,1554,1045],"isMain":false},{"metricName":"NA75","quality":"More is better","values":[850,921,718,719,932,739],"isMain":false},{"metricName":"NGA50","quality":"More is better","values":[1063,1288,844,854,1389,797],"isMain":true},{"metricName":"NGA75","quality":"More is better","values":[562,677,529,517,734,null],"isMain":false},{"metricName":"LG50","quality":"Less is better","values":[1507,1114,1977,1893,1029,2014],"isMain":false},{"metricName":"LG75","quality":"Less is better","values":[3171,2397,3902,3795,2208,null],"isMain":false},{"metricName":"LA50","quality":"Less is better","values":[1073,960,1419,1379,950,1221],"isMain":false},{"metricName":"LA75","quality":"Less is better","values":[2079,1933,2660,2608,1933,2301],"isMain":false},{"metricName":"LGA50","quality":"Less is better","values":[1529,1227,2014,1974,1150,2053],"isMain":true},{"metricName":"LGA75","quality":"Less is better","values":[3242,2638,3996,3974,2465,null],"isMain":false}]],["Reads mapping",[]],["Misassemblies",[{"metricName":"# misassemblies","quality":"Less is better","values":[8,10,12,13,9,7],"isMain":true},{"metricName":" # relocations","quality":"Less is better","values":[1,3,3,0,2,2],"isMain":false},{"metricName":" # translocations","quality":"Less is better","values":[2,6,4,6,6,5],"isMain":false},{"metricName":" # inversions","quality":"Less is better","values":[5,1,5,7,1,0],"isMain":false},{"metricName":" # interspecies translocations","quality":"Less is better","values":[89,125,43,102,112,48],"isMain":false},{"metricName":"# misassembled contigs","quality":"Less is better","values":[8,10,12,13,9,7],"isMain":false},{"metricName":"Misassembled contigs length","quality":"Less is better","values":[10932,16204,16148,18808,12330,10273],"isMain":true},{"metricName":"# possibly misassembled contigs","quality":"Less is better","values":[0,1,0,0,0,0],"isMain":false},{"metricName":" # possible misassemblies","quality":"Less is better","values":[0,1,0,0,0,0],"isMain":false},{"metricName":"# local misassemblies","quality":"Less is better","values":[3,2,5,3,2,3],"isMain":false},{"metricName":"# scaffold gap ext. mis.","quality":"Less is better","values":[0,0,0,0,0,0],"isMain":false},{"metricName":"# scaffold gap loc. mis.","quality":"Less is better","values":[0,0,0,0,0,0],"isMain":false},{"metricName":"# unaligned mis. contigs","quality":"Less is better","values":[1,2,0,0,0,0],"isMain":false}]],["Unaligned",[{"metricName":"# fully unaligned contigs","quality":"Less is better","values":[0,4,3,4,3,0],"isMain":false},{"metricName":"Fully unaligned length","quality":"Less is better","values":[0,2464,3320,4649,60286,0],"isMain":false},{"metricName":"# partially unaligned contigs","quality":"Less is better","values":[9,25,6,15,30,6],"isMain":false},{"metricName":"Partially unaligned length","quality":"Less is better","values":[9456,143222,12227,49686,116087,10041],"isMain":false}]],["Mismatches",[{"metricName":"# mismatches","quality":"Less is better","values":[38687,37959,66321,66206,40726,36863],"isMain":false},{"metricName":"# indels","quality":"Less is better","values":[113,134,103,112,161,90],"isMain":false},{"metricName":"Indels length","quality":"Less is better","values":[167,296,163,158,256,146],"isMain":false},{"metricName":"# mismatches per 100 kbp","quality":"Less is better","values":["961.78","898.26","1701.71","1723.96","941.18","1022.92"],"isMain":true},{"metricName":"# indels per 100 kbp","quality":"Less is better","values":["2.81","3.17","2.64","2.92","3.72","2.50"],"isMain":true},{"metricName":" # indels (<= 5 bp)","quality":"Less is better","values":[110,129,100,109,157,86],"isMain":false},{"metricName":" # indels (> 5 bp)","quality":"Less is better","values":[3,5,3,3,4,4],"isMain":false},{"metricName":"# N's","quality":"Less is better","values":[0,0,0,0,0,0],"isMain":false},{"metricName":"# N's per 100 kbp","quality":"Less is better","values":["0.00","0.00","0.00","0.00","0.00","0.00"],"isMain":true}]],["Statistics without reference",[{"metricName":"# contigs","quality":"Equal","values":[3639,3325,4340,4213,3335,3789],"isMain":true},{"metricName":"# contigs (>= 1000 bp)","quality":"Equal","values":[1705,1776,1473,1486,1834,1392],"isMain":false},{"metricName":"# contigs (>= 5000 bp)","quality":"Equal","values":[8,22,3,4,32,5],"isMain":false},{"metricName":"# contigs (>= 10000 bp)","quality":"Equal","values":[0,2,1,1,6,2],"isMain":false},{"metricName":"# contigs (>= 25000 bp)","quality":"Equal","values":[0,2,0,1,2,0],"isMain":false},{"metricName":"# contigs (>= 50000 bp)","quality":"Equal","values":[0,1,0,0,1,0],"isMain":false},{"metricName":"Largest contig","quality":"More is better","values":[7362,74950,10720,27404,56868,12945],"isMain":true},{"metricName":"Total length","quality":"More is better","values":[4235452,4566223,4211836,4201394,4722514,3794333],"isMain":true},{"metricName":"Total length (>= 1000 bp)","quality":"More is better","values":[2861104,3438104,2186596,2272942,3640271,2095212],"isMain":true},{"metricName":"Total length (>= 5000 bp)","quality":"More is better","values":[47028,232229,22230,52445,301346,46881],"isMain":false},{"metricName":"Total length (>= 10000 bp)","quality":"More is better","values":[0,114641,10720,27404,149280,24538],"isMain":true},{"metricName":"Total length (>= 25000 bp)","quality":"More is better","values":[0,114641,0,27404,98992,0],"isMain":false},{"metricName":"Total length (>= 50000 bp)","quality":"More is better","values":[0,74950,0,0,56868,0],"isMain":true},{"metricName":"N50","quality":"More is better","values":[1324,1633,1023,1061,1687,1062],"isMain":false},{"metricName":"N75","quality":"More is better","values":[868,1008,739,751,1041,761],"isMain":false},{"metricName":"L50","quality":"Less is better","values":[1059,866,1394,1319,845,1200],"isMain":false},{"metricName":"L75","quality":"Less is better","values":[2045,1763,2608,2501,1738,2255],"isMain":false},{"metricName":"GC (%)","quality":"Equal","values":["41.82","41.87","41.95","41.93","41.89","41.93"],"isMain":false}]],["K-mer-based statistics",[]],["Predicted genes",[]],["Similarity statistics",[]],["Reference statistics",[{"metricName":"Reference length","quality":"Equal","values":[5309980,5309980,5309980,5309980,5309980,5309980],"isMain":false},{"metricName":"Reference fragments","quality":"Equal","values":[21,21,21,21,21,21],"isMain":false},{"metricName":"Reference GC (%)","quality":"Equal","values":["41.90","41.90","41.90","41.90","41.90","41.90"],"isMain":false}]]],"subreferences":[],"subreports":[],"minContig":500},{"date":"10 February 2020, Monday, 20:33:57","assembliesNames":["megahit-103-df","megahit-113-df","megahit-113-ml","megahit-113-ms","megahit-129-df","metaSPAdes"],"referenceName":"GCF\_000382385.1\_ASM38238v1","order":[0,1,2,3,4,5],"report":[["Genome statistics",[{"metricName":"Genome fraction (%)","quality":"More is better","values":["12.978","12.909","14.714","13.542","15.036","8.898"],"isMain":true},{"metricName":"Duplication ratio","quality":"Less is better","values":["1.060","1.045","1.070","1.084","1.065","1.035"],"isMain":true},{"metricName":"Largest alignment","quality":"More is better","values":[3362,21152,31541,27312,31541,31551],"isMain":true},{"metricName":"Total aligned length","quality":"More is better","values":[444548,438740,511881,470798,516553,302403],"isMain":true},{"metricName":"NA50","quality":"More is better","values":[849,989,792,861,1057,1027],"isMain":false},{"metricName":"NA75","quality":"More is better","values":[604,533,591,585,653,699],"isMain":false},{"metricName":"NGA50","quality":"More is better","values":[null,null,null,null,null,null],"isMain":true},{"metricName":"LA50","quality":"Less is better","values":[177,148,197,184,136,90],"isMain":false},{"metricName":"LA75","quality":"Less is better","values":[342,345,393,371,301,183],"isMain":false}]],["Reads mapping",[]],["Misassemblies",[{"metricName":"# misassemblies","quality":"Less is better","values":[5,10,4,7,15,1],"isMain":true},{"metricName":" # relocations","quality":"Less is better","values":[1,2,0,2,2,0],"isMain":false},{"metricName":" # translocations","quality":"Less is better","values":[3,7,4,4,10,1],"isMain":false},{"metricName":" # inversions","quality":"Less is better","values":[1,1,0,1,3,0],"isMain":false},{"metricName":" # interspecies translocations","quality":"Less is better","values":[31,28,29,40,40,8],"isMain":false},{"metricName":"# misassembled contigs","quality":"Less is better","values":[5,9,4,5,14,1],"isMain":false},{"metricName":"Misassembled contigs length","quality":"Less is better","values":[2985,37898,12200,6639,14852,1412],"isMain":true},{"metricName":"# possibly misassembled contigs","quality":"Less is better","values":[0,0,0,0,0,0],"isMain":false},{"metricName":" # possible misassemblies","quality":"Less is better","values":[0,0,0,0,0,0],"isMain":false},{"metricName":"# local misassemblies","quality":"Less is better","values":[1,0,1,3,5,0],"isMain":false},{"metricName":"# scaffold gap ext. mis.","quality":"Less is better","values":[0,0,0,0,0,0],"isMain":false},{"metricName":"# scaffold gap loc. mis.","quality":"Less is better","values":[0,0,0,0,0,0],"isMain":false},{"metricName":"# unaligned mis. contigs","quality":"Less is better","values":[2,2,2,1,0,0],"isMain":false}]],["Unaligned",[{"metricName":"# fully unaligned contigs","quality":"Less is better","values":[1,1,3,2,2,0],"isMain":false},{"metricName":"Fully unaligned length","quality":"Less is better","values":[518,507,2551,1408,1649,0],"isMain":false},{"metricName":"# partially unaligned contigs","quality":"Less is better","values":[7,11,8,10,6,7],"isMain":false},{"metricName":"Partially unaligned length","quality":"Less is better","values":[9014,99914,9058,34220,7401,9072],"isMain":false}]],["Mismatches",[{"metricName":"# mismatches","quality":"Less is better","values":[5184,5399,9712,8730,7221,3688],"isMain":false},{"metricName":"# indels","quality":"Less is better","values":[26,32,63,58,56,39],"isMain":false},{"metricName":"Indels length","quality":"Less is better","values":[129,126,291,280,261,288],"isMain":false},{"metricName":"# mismatches per 100 kbp","quality":"Less is better","values":["1205.98","1262.78","1992.92","1946.39","1449.95","1251.39"],"isMain":true},{"metricName":"# indels per 100 kbp","quality":"Less is better","values":["6.05","7.48","12.93","12.93","11.24","13.23"],"isMain":true},{"metricName":" # indels (<= 5 bp)","quality":"Less is better","values":[23,31,53,50,47,31],"isMain":false},{"metricName":" # indels (> 5 bp)","quality":"Less is better","values":[3,1,10,8,9,8],"isMain":false},{"metricName":"# N's","quality":"Less is better","values":[0,0,0,0,0,0],"isMain":false},{"metricName":"# N's per 100 kbp","quality":"Less is better","values":["0.00","0.00","0.00","0.00","0.00","0.00"],"isMain":true}]],["Statistics without reference",[{"metricName":"# contigs","quality":"Equal","values":[533,405,610,534,513,298],"isMain":true},{"metricName":"# contigs (>= 1000 bp)","quality":"Equal","values":[125,149,128,144,158,104],"isMain":false},{"metricName":"# contigs (>= 5000 bp)","quality":"Equal","values":[0,3,3,2,3,1],"isMain":false},{"metricName":"# contigs (>= 10000 bp)","quality":"Equal","values":[0,2,2,2,1,1],"isMain":false},{"metricName":"# contigs (>= 25000 bp)","quality":"Equal","values":[0,2,1,2,1,1],"isMain":false},{"metricName":"# contigs (>= 50000 bp)","quality":"Equal","values":[0,1,0,0,0,0],"isMain":false},{"metricName":"Largest contig","quality":"More is better","values":[4495,83634,31541,27338,31541,31551],"isMain":true},{"metricName":"Total length","quality":"More is better","values":[465206,547129,532909,521986,539282,314218],"isMain":true},{"metricName":"Total length (>= 1000 bp)","quality":"More is better","values":[191506,374129,213805,258390,301407,179863],"isMain":true},{"metricName":"Total length (>= 5000 bp)","quality":"More is better","values":[0,120895,47097,53224,44293,31551],"isMain":false},{"metricName":"Total length (>= 10000 bp)","quality":"More is better","values":[0,114910,41877,53224,31541,31551],"isMain":true},{"metricName":"Total length (>= 25000 bp)","quality":"More is better","values":[0,114910,31541,53224,31541,31551],"isMain":false},{"metricName":"Total length (>= 50000 bp)","quality":"More is better","values":[0,83634,0,0,0,0],"isMain":true},{"metricName":"N50","quality":"More is better","values":[880,1546,847,994,1117,1073],"isMain":false},{"metricName":"N75","quality":"More is better","values":[640,839,628,669,705,748],"isMain":false},{"metricName":"L50","quality":"Less is better","values":[169,68,186,147,128,83],"isMain":false},{"metricName":"L75","quality":"Less is better","values":[325,189,372,308,283,168],"isMain":false},{"metricName":"GC (%)","quality":"Equal","values":["43.79","43.78","43.90","44.06","43.64","42.73"],"isMain":false}]],["K-mer-based statistics",[]],["Predicted genes",[]],["Similarity statistics",[]],["Reference statistics",[{"metricName":"Reference length","quality":"Equal","values":[3312100,3312100,3312100,3312100,3312100,3312100],"isMain":false},{"metricName":"Reference fragments","quality":"Equal","values":[63,63,63,63,63,63],"isMain":false},{"metricName":"Reference GC (%)","quality":"Equal","values":["47.39","47.39","47.39","47.39","47.39","47.39"],"isMain":false}]]],"subreferences":[],"subreports":[],"minContig":500},{"date":"10 February 2020, Monday, 20:33:58","assembliesNames":["megahit-103-df","megahit-113-df","megahit-113-ml","megahit-113-ms","megahit-129-df","metaSPAdes"],"referenceName":"GCF\_000382445.1\_Bact\_mass\_DSM\_17679\_V1","order":[0,1,2,3,4,5],"report":[["Genome statistics",[{"metricName":"Genome fraction (%)","quality":"More is better","values":["0.319","0.498","0.425","0.534","0.400","0.290"],"isMain":true},{"metricName":"Duplication ratio","quality":"Less is better","values":["1.135","1.066","1.013","1.084","1.148","1.026"],"isMain":true},{"metricName":"Largest alignment","quality":"More is better","values":[2917,2663,1252,2379,2193,1346],"isMain":true},{"metricName":"Total aligned length","quality":"More is better","values":[14557,22827,19480,24822,18454,13362],"isMain":true},{"metricName":"NA50","quality":"More is better","values":[658,642,760,613,619,728],"isMain":false},{"metricName":"NA75","quality":"More is better","values":[285,null,589,null,501,574],"isMain":false},{"metricName":"NGA50","quality":"More is better","values":[null,null,null,null,null,null],"isMain":true},{"metricName":"LA50","quality":"Less is better","values":[8,13,11,20,11,7],"isMain":false},{"metricName":"LA75","quality":"Less is better","values":[16,null,18,null,21,13],"isMain":false}]],["Reads mapping",[]],["Misassemblies",[{"metricName":"# misassemblies","quality":"Less is better","values":[0,1,0,0,0,0],"isMain":true},{"metricName":" # relocations","quality":"Less is better","values":[0,0,0,0,0,0],"isMain":false},{"metricName":" # translocations","quality":"Less is better","values":[0,0,0,0,0,0],"isMain":false},{"metricName":" # inversions","quality":"Less is better","values":[0,1,0,0,0,0],"isMain":false},{"metricName":" # interspecies translocations","quality":"Less is better","values":[6,10,0,9,9,1],"isMain":false},{"metricName":"# misassembled contigs","quality":"Less is better","values":[0,1,0,0,0,0],"isMain":false},{"metricName":"Misassembled contigs length","quality":"Less is better","values":[0,697,0,0,0,0],"isMain":true},{"metricName":"# possibly misassembled contigs","quality":"Less is better","values":[1,0,0,0,0,0],"isMain":false},{"metricName":" # possible misassemblies","quality":"Less is better","values":[1,0,0,0,0,0],"isMain":false},{"metricName":"# local misassemblies","quality":"Less is better","values":[0,0,0,1,2,0],"isMain":false},{"metricName":"# scaffold gap ext. mis.","quality":"Less is better","values":[0,0,0,0,0,0],"isMain":false},{"metricName":"# scaffold gap loc. mis.","quality":"Less is better","values":[0,0,0,0,0,0],"isMain":false},{"metricName":"# unaligned mis. contigs","quality":"Less is better","values":[0,0,0,0,0,0],"isMain":false}]],["Unaligned",[{"metricName":"# fully unaligned contigs","quality":"Less is better","values":[0,0,0,0,1,0],"isMain":false},{"metricName":"Fully unaligned length","quality":"Less is better","values":[0,0,0,0,684,0],"isMain":false},{"metricName":"# partially unaligned contigs","quality":"Less is better","values":[2,6,0,4,1,1],"isMain":false},{"metricName":"Partially unaligned length","quality":"Less is better","values":[1624,6870,0,12627,786,726],"isMain":false}]],["Mismatches",[{"metricName":"# mismatches","quality":"Less is better","values":[271,420,468,536,254,145],"isMain":false},{"metricName":"# indels","quality":"Less is better","values":[3,3,4,6,4,0],"isMain":false},{"metricName":"Indels length","quality":"Less is better","values":[3,3,6,6,6,0],"isMain":false},{"metricName":"# mismatches per 100 kbp","quality":"Less is better","values":["1861.26","1848.43","2410.01","2198.34","1391.70","1093.51"],"isMain":true},{"metricName":"# indels per 100 kbp","quality":"Less is better","values":["20.60","13.20","20.60","24.61","21.92","0.00"],"isMain":true},{"metricName":" # indels (<= 5 bp)","quality":"Less is better","values":[3,3,4,6,4,0],"isMain":false},{"metricName":" # indels (> 5 bp)","quality":"Less is better","values":[0,0,0,0,0,0],"isMain":false},{"metricName":"# N's","quality":"Less is better","values":[0,0,0,0,0,0],"isMain":false},{"metricName":"# N's per 100 kbp","quality":"Less is better","values":["0.00","0.00","0.00","0.00","0.00","0.00"],"isMain":true}]],["Statistics without reference",[{"metricName":"# contigs","quality":"Equal","values":[20,29,26,34,27,18],"isMain":true},{"metricName":"# contigs (>= 1000 bp)","quality":"Equal","values":[2,11,4,9,7,4],"isMain":false},{"metricName":"# contigs (>= 5000 bp)","quality":"Equal","values":[0,0,0,1,0,0],"isMain":false},{"metricName":"# contigs (>= 10000 bp)","quality":"Equal","values":[0,0,0,0,0,0],"isMain":false},{"metricName":"# contigs (>= 25000 bp)","quality":"Equal","values":[0,0,0,0,0,0],"isMain":false},{"metricName":"# contigs (>= 50000 bp)","quality":"Equal","values":[0,0,0,0,0,0],"isMain":false},{"metricName":"Largest contig","quality":"More is better","values":[2962,4366,1267,8196,2204,1452],"isMain":true},{"metricName":"Total length","quality":"More is better","values":[18154,31102,19662,39061,22419,14333],"isMain":true},{"metricName":"Total length (>= 1000 bp)","quality":"More is better","values":[5922,18881,4608,22526,9393,4969],"isMain":true},{"metricName":"Total length (>= 5000 bp)","quality":"More is better","values":[0,0,0,8196,0,0],"isMain":false},{"metricName":"Total length (>= 10000 bp)","quality":"More is better","values":[0,0,0,0,0,0],"isMain":true},{"metricName":"Total length (>= 25000 bp)","quality":"More is better","values":[0,0,0,0,0,0],"isMain":false},{"metricName":"Total length (>= 50000 bp)","quality":"More is better","values":[0,0,0,0,0,0],"isMain":true},{"metricName":"N50","quality":"More is better","values":[835,1169,802,1114,894,842],"isMain":false},{"metricName":"N75","quality":"More is better","values":[640,719,589,685,613,658],"isMain":false},{"metricName":"L50","quality":"Less is better","values":[6,9,10,7,9,7],"isMain":false},{"metricName":"L75","quality":"Less is better","values":[12,17,18,18,17,12],"isMain":false},{"metricName":"GC (%)","quality":"Equal","values":["43.30","41.14","41.69","47.01","41.64","39.83"],"isMain":false}]],["K-mer-based statistics",[]],["Predicted genes",[]],["Similarity statistics",[]],["Reference statistics",[{"metricName":"Reference length","quality":"Equal","values":[4564584,4564584,4564584,4564584,4564584,4564584],"isMain":false},{"metricName":"Reference fragments","quality":"Equal","values":[55,55,55,55,55,55],"isMain":false},{"metricName":"Reference GC (%)","quality":"Equal","values":["42.69","42.69","42.69","42.69","42.69","42.69"],"isMain":false}]]],"subreferences":[],"subreports":[],"minContig":500},{"date":"10 February 2020, Monday, 20:34:12","assembliesNames":["megahit-103-df","megahit-113-df","megahit-113-ml","megahit-113-ms","megahit-129-df","metaSPAdes"],"referenceName":"GCF\_000382465.1\_Bact\_sp\_HPS0048\_V1","order":[0,1,2,3,4,5],"report":[["Genome statistics",[{"metricName":"Genome fraction (%)","quality":"More is better","values":["2.250","2.852","3.251","3.822","3.271","1.402"],"isMain":true},{"metricName":"Duplication ratio","quality":"Less is better","values":["1.063","1.064","1.032","1.053","1.047","1.066"],"isMain":true},{"metricName":"Largest alignment","quality":"More is better","values":[1806,2462,1736,1676,2460,1996],"isMain":true},{"metricName":"Total aligned length","quality":"More is better","values":[134622,170517,194404,228756,195403,83753],"isMain":true},{"metricName":"NA50","quality":"More is better","values":[572,591,583,575,591,550],"isMain":false},{"metricName":"NA75","quality":"More is better","values":[513,507,523,517,517,506],"isMain":false},{"metricName":"NGA50","quality":"More is better","values":[null,null,null,null,null,null],"isMain":true},{"metricName":"LA50","quality":"Less is better","values":[105,127,139,177,138,69],"isMain":false},{"metricName":"LA75","quality":"Less is better","values":[174,218,231,291,236,115],"isMain":false}]],["Reads mapping",[]],["Misassemblies",[{"metricName":"# misassemblies","quality":"Less is better","values":[3,7,3,2,4,0],"isMain":true},{"metricName":" # relocations","quality":"Less is better","values":[0,1,0,0,1,0],"isMain":false},{"metricName":" # translocations","quality":"Less is better","values":[3,5,3,2,3,0],"isMain":false},{"metricName":" # inversions","quality":"Less is better","values":[0,1,0,0,0,0],"isMain":false},{"metricName":" # interspecies translocations","quality":"Less is better","values":[35,47,17,43,38,24],"isMain":false},{"metricName":"# misassembled contigs","quality":"Less is better","values":[3,6,3,2,3,0],"isMain":false},{"metricName":"Misassembled contigs length","quality":"Less is better","values":[1772,7675,1745,1371,2112,0],"isMain":true},{"metricName":"# possibly misassembled contigs","quality":"Less is better","values":[0,0,0,0,0,0],"isMain":false},{"metricName":" # possible misassemblies","quality":"Less is better","values":[0,0,0,0,0,0],"isMain":false},{"metricName":"# local misassemblies","quality":"Less is better","values":[0,1,0,1,1,0],"isMain":false},{"metricName":"# scaffold gap ext. mis.","quality":"Less is better","values":[0,0,0,0,0,0],"isMain":false},{"metricName":"# scaffold gap loc. mis.","quality":"Less is better","values":[0,0,0,0,0,0],"isMain":false},{"metricName":"# unaligned mis. contigs","quality":"Less is better","values":[1,1,0,0,0,0],"isMain":false}]],["Unaligned",[{"metricName":"# fully unaligned contigs","quality":"Less is better","values":[0,0,0,1,1,0],"isMain":false},{"metricName":"Fully unaligned length","quality":"Less is better","values":[0,0,0,829,556,0],"isMain":false},{"metricName":"# partially unaligned contigs","quality":"Less is better","values":[7,12,2,10,11,8],"isMain":false},{"metricName":"Partially unaligned length","quality":"Less is better","values":[7710,15516,1772,8192,9365,8171],"isMain":false}]],["Mismatches",[{"metricName":"# mismatches","quality":"Less is better","values":[2466,3091,4190,5070,3525,1490],"isMain":false},{"metricName":"# indels","quality":"Less is better","values":[6,8,16,20,9,5],"isMain":false},{"metricName":"Indels length","quality":"Less is better","values":[40,55,28,67,22,28],"isMain":false},{"metricName":"# mismatches per 100 kbp","quality":"Less is better","values":["1836.97","1816.64","2160.52","2223.17","1806.06","1781.38"],"isMain":true},{"metricName":"# indels per 100 kbp","quality":"Less is better","values":["4.47","4.70","8.25","8.77","4.61","5.98"],"isMain":true},{"metricName":" # indels (<= 5 bp)","quality":"Less is better","values":[5,6,15,17,8,3],"isMain":false},{"metricName":" # indels (> 5 bp)","quality":"Less is better","values":[1,2,1,3,1,2],"isMain":false},{"metricName":"# N's","quality":"Less is better","values":[0,0,0,0,0,0],"isMain":false},{"metricName":"# N's per 100 kbp","quality":"Less is better","values":["0.00","0.00","0.00","0.00","0.00","0.00"],"isMain":true}]],["Statistics without reference",[{"metricName":"# contigs","quality":"Equal","values":[228,275,320,390,318,145],"isMain":true},{"metricName":"# contigs (>= 1000 bp)","quality":"Equal","values":[10,21,10,13,20,10],"isMain":false},{"metricName":"# contigs (>= 5000 bp)","quality":"Equal","values":[0,0,0,0,0,0],"isMain":false},{"metricName":"# contigs (>= 10000 bp)","quality":"Equal","values":[0,0,0,0,0,0],"isMain":false},{"metricName":"# contigs (>= 25000 bp)","quality":"Equal","values":[0,0,0,0,0,0],"isMain":false},{"metricName":"# contigs (>= 50000 bp)","quality":"Equal","values":[0,0,0,0,0,0],"isMain":false},{"metricName":"Largest contig","quality":"More is better","values":[2902,3824,1846,2531,2605,2292],"isMain":true},{"metricName":"Total length","quality":"More is better","values":[150465,196614,201956,249269,214362,97293],"isMain":true},{"metricName":"Total length (>= 1000 bp)","quality":"More is better","values":[14862,35139,13628,17251,28718,14803],"isMain":true},{"metricName":"Total length (>= 5000 bp)","quality":"More is better","values":[0,0,0,0,0,0],"isMain":false},{"metricName":"Total length (>= 10000 bp)","quality":"More is better","values":[0,0,0,0,0,0],"isMain":true},{"metricName":"Total length (>= 25000 bp)","quality":"More is better","values":[0,0,0,0,0,0],"isMain":false},{"metricName":"Total length (>= 50000 bp)","quality":"More is better","values":[0,0,0,0,0,0],"isMain":true},{"metricName":"N50","quality":"More is better","values":[619,659,595,611,631,613],"isMain":false},{"metricName":"N75","quality":"More is better","values":[550,567,537,545,558,545],"isMain":false},{"metricName":"L50","quality":"Less is better","values":[92,102,134,163,125,57],"isMain":false},{"metricName":"L75","quality":"Less is better","values":[157,183,223,271,216,99],"isMain":false},{"metricName":"GC (%)","quality":"Equal","values":["40.89","40.69","40.78","40.93","40.93","40.02"],"isMain":false}]],["K-mer-based statistics",[]],["Predicted genes",[]],["Similarity statistics",[]],["Reference statistics",[{"metricName":"Reference length","quality":"Equal","values":[5966298,5966298,5966298,5966298,5966298,5966298],"isMain":false},{"metricName":"Reference fragments","quality":"Equal","values":[83,83,83,83,83,83],"isMain":false},{"metricName":"Reference GC (%)","quality":"Equal","values":["40.39","40.39","40.39","40.39","40.39","40.39"],"isMain":false}]]],"subreferences":[],"subreports":[],"minContig":500},{"date":"10 February 2020, Monday, 20:33:46","assembliesNames":["megahit-103-df","megahit-113-df","megahit-113-ml","megahit-113-ms","megahit-129-df","metaSPAdes"],"referenceName":"GCF\_000383295.1\_ASM38329v1","order":[0,1,2,3,4,5],"report":[["Genome statistics",[{"metricName":"Genome fraction (%)","quality":"More is better","values":["0.495","0.701","0.628","0.935","0.735","0.251"],"isMain":true},{"metricName":"Duplication ratio","quality":"Less is better","values":["1.131","1.113","1.033","1.028","1.045","1.100"],"isMain":true},{"metricName":"Largest alignment","quality":"More is better","values":[665,712,750,798,756,798],"isMain":true},{"metricName":"Total aligned length","quality":"More is better","values":[15766,22264,19917,29682,23401,7950],"isMain":true},{"metricName":"NA50","quality":"More is better","values":[515,519,549,537,531,null],"isMain":false},{"metricName":"NA75","quality":"More is better","values":[385,null,512,471,230,null],"isMain":false},{"metricName":"NGA50","quality":"More is better","values":[null,null,null,null,null,null],"isMain":true},{"metricName":"LA50","quality":"Less is better","values":[17,27,18,32,25,null],"isMain":false},{"metricName":"LA75","quality":"Less is better","values":[27,null,28,51,42,null],"isMain":false}]],["Reads mapping",[]],["Misassemblies",[{"metricName":"# misassemblies","quality":"Less is better","values":[0,0,0,0,0,0],"isMain":true},{"metricName":" # relocations","quality":"Less is better","values":[0,0,0,0,0,0],"isMain":false},{"metricName":" # translocations","quality":"Less is better","values":[0,0,0,0,0,0],"isMain":false},{"metricName":" # inversions","quality":"Less is better","values":[0,0,0,0,0,0],"isMain":false},{"metricName":" # interspecies translocations","quality":"Less is better","values":[6,12,2,6,8,6],"isMain":false},{"metricName":"# misassembled contigs","quality":"Less is better","values":[0,0,0,0,0,0],"isMain":false},{"metricName":"Misassembled contigs length","quality":"Less is better","values":[0,0,0,0,0,0],"isMain":true},{"metricName":"# possibly misassembled contigs","quality":"Less is better","values":[0,0,0,0,0,0],"isMain":false},{"metricName":" # possible misassemblies","quality":"Less is better","values":[0,0,0,0,0,0],"isMain":false},{"metricName":"# local misassemblies","quality":"Less is better","values":[0,0,0,0,0,0],"isMain":false},{"metricName":"# scaffold gap ext. mis.","quality":"Less is better","values":[0,0,0,0,0,0],"isMain":false},{"metricName":"# scaffold gap loc. mis.","quality":"Less is better","values":[0,0,0,0,0,0],"isMain":false},{"metricName":"# unaligned mis. contigs","quality":"Less is better","values":[0,0,0,0,0,0],"isMain":false}]],["Unaligned",[{"metricName":"# fully unaligned contigs","quality":"Less is better","values":[0,0,0,0,0,0],"isMain":false},{"metricName":"Fully unaligned length","quality":"Less is better","values":[0,0,0,0,0,0],"isMain":false},{"metricName":"# partially unaligned contigs","quality":"Less is better","values":[1,5,1,5,5,4],"isMain":false},{"metricName":"Partially unaligned length","quality":"Less is better","values":[1131,6372,653,7590,5927,18703],"isMain":false}]],["Mismatches",[{"metricName":"# mismatches","quality":"Less is better","values":[270,425,373,639,422,132],"isMain":false},{"metricName":"# indels","quality":"Less is better","values":[0,0,0,0,0,0],"isMain":false},{"metricName":"Indels length","quality":"Less is better","values":[0,0,0,0,0,0],"isMain":false},{"metricName":"# mismatches per 100 kbp","quality":"Less is better","values":["1718.10","1910.37","1872.77","2152.82","1808.60","1660.38"],"isMain":true},{"metricName":"# indels per 100 kbp","quality":"Less is better","values":["0.00","0.00","0.00","0.00","0.00","0.00"],"isMain":true},{"metricName":" # indels (<= 5 bp)","quality":"Less is better","values":[0,0,0,0,0,0],"isMain":false},{"metricName":" # indels (> 5 bp)","quality":"Less is better","values":[0,0,0,0,0,0],"isMain":false},{"metricName":"# N's","quality":"Less is better","values":[0,0,0,0,0,0],"isMain":false},{"metricName":"# N's per 100 kbp","quality":"Less is better","values":["0.00","0.00","0.00","0.00","0.00","0.00"],"isMain":true}]],["Statistics without reference",[{"metricName":"# contigs","quality":"Equal","values":[33,46,37,56,45,18],"isMain":true},{"metricName":"# contigs (>= 1000 bp)","quality":"Equal","values":[1,4,0,3,5,2],"isMain":false},{"metricName":"# contigs (>= 5000 bp)","quality":"Equal","values":[0,0,0,0,0,1],"isMain":false},{"metricName":"# contigs (>= 10000 bp)","quality":"Equal","values":[0,0,0,0,0,1],"isMain":false},{"metricName":"# contigs (>= 25000 bp)","quality":"Equal","values":[0,0,0,0,0,0],"isMain":false},{"metricName":"# contigs (>= 50000 bp)","quality":"Equal","values":[0,0,0,0,0,0],"isMain":false},{"metricName":"Largest contig","quality":"More is better","values":[1382,2927,777,3164,2416,16109],"isMain":true},{"metricName":"Total length","quality":"More is better","values":[18910,31122,21224,38089,30306,27445],"isMain":true},{"metricName":"Total length (>= 1000 bp)","quality":"More is better","values":[1382,6465,0,7198,6914,18039],"isMain":true},{"metricName":"Total length (>= 5000 bp)","quality":"More is better","values":[0,0,0,0,0,16109],"isMain":false},{"metricName":"Total length (>= 10000 bp)","quality":"More is better","values":[0,0,0,0,0,16109],"isMain":true},{"metricName":"Total length (>= 25000 bp)","quality":"More is better","values":[0,0,0,0,0,0],"isMain":false},{"metricName":"Total length (>= 50000 bp)","quality":"More is better","values":[0,0,0,0,0,0],"isMain":true},{"metricName":"N50","quality":"More is better","values":[538,598,557,596,612,16109],"isMain":false},{"metricName":"N75","quality":"More is better","values":[517,534,522,539,539,595],"isMain":false},{"metricName":"L50","quality":"Less is better","values":[15,18,17,21,18,1],"isMain":false},{"metricName":"L75","quality":"Less is better","values":[24,31,27,38,31,6],"isMain":false},{"metricName":"GC (%)","quality":"Equal","values":["52.54","53.68","51.78","51.98","54.64","60.35"],"isMain":false}]],["K-mer-based statistics",[]],["Predicted genes",[]],["Similarity statistics",[]],["Reference statistics",[{"metricName":"Reference length","quality":"Equal","values":[3173517,3173517,3173517,3173517,3173517,3173517],"isMain":false},{"metricName":"Reference fragments","quality":"Equal","values":[24,24,24,24,24,24],"isMain":false},{"metricName":"Reference GC (%)","quality":"Equal","values":["53.56","53.56","53.56","53.56","53.56","53.56"],"isMain":false}]]],"subreferences":[],"subreports":[],"minContig":500},{"date":"10 February 2020, Monday, 20:16:49","assembliesNames":["megahit-113-df","megahit-113-ml","megahit-113-ms","megahit-129-df"],"referenceName":"GCF\_000384195.1\_ASM38419v1","order":[0,1,2,3],"report":[["Genome statistics",[{"metricName":"Genome fraction (%)","quality":"More is better","values":["0.026","0.059","0.057","0.092"],"isMain":true},{"metricName":"Duplication ratio","quality":"Less is better","values":["1.048","1.055","1.005","1.100"],"isMain":true},{"metricName":"Largest alignment","quality":"More is better","values":[419,692,570,1193],"isMain":true},{"metricName":"Total aligned length","quality":"More is better","values":[540,1220,1175,2061],"isMain":true},{"metricName":"NA50","quality":"More is better","values":[419,692,91,1193],"isMain":false},{"metricName":"NA75","quality":"More is better","values":[121,528,null,868],"isMain":false},{"metricName":"NGA50","quality":"More is better","values":[null,null,null,null],"isMain":true},{"metricName":"LA50","quality":"Less is better","values":[1,1,3,1],"isMain":false},{"metricName":"LA75","quality":"Less is better","values":[2,2,null,2],"isMain":false}]],["Reads mapping",[]],["Misassemblies",[{"metricName":"# misassemblies","quality":"Less is better","values":[1,0,0,0],"isMain":true},{"metricName":" # relocations","quality":"Less is better","values":[0,0,0,0],"isMain":false},{"metricName":" # translocations","quality":"Less is better","values":[0,0,0,0],"isMain":false},{"metricName":" # inversions","quality":"Less is better","values":[1,0,0,0],"isMain":false},{"metricName":" # interspecies translocations","quality":"Less is better","values":[0,0,1,0],"isMain":false},{"metricName":"# misassembled contigs","quality":"Less is better","values":[1,0,0,0],"isMain":false},{"metricName":"Misassembled contigs length","quality":"Less is better","values":[566,0,0,0],"isMain":true},{"metricName":"# possibly misassembled contigs","quality":"Less is better","values":[0,0,0,0],"isMain":false},{"metricName":" # possible misassemblies","quality":"Less is better","values":[0,0,0,0],"isMain":false},{"metricName":"# local misassemblies","quality":"Less is better","values":[0,0,0,0],"isMain":false},{"metricName":"# scaffold gap ext. mis.","quality":"Less is better","values":[0,0,0,0],"isMain":false},{"metricName":"# scaffold gap loc. mis.","quality":"Less is better","values":[0,0,0,0],"isMain":false},{"metricName":"# unaligned mis. contigs","quality":"Less is better","values":[0,0,0,0],"isMain":false}]],["Unaligned",[{"metricName":"# fully unaligned contigs","quality":"Less is better","values":[0,0,0,0],"isMain":false},{"metricName":"Fully unaligned length","quality":"Less is better","values":[0,0,0,0],"isMain":false},{"metricName":"# partially unaligned contigs","quality":"Less is better","values":[0,0,1,0],"isMain":false},{"metricName":"Partially unaligned length","quality":"Less is better","values":[0,0,1013,0],"isMain":false}]],["Mismatches",[{"metricName":"# mismatches","quality":"Less is better","values":[16,18,25,41],"isMain":false},{"metricName":"# indels","quality":"Less is better","values":[0,0,0,0],"isMain":false},{"metricName":"Indels length","quality":"Less is better","values":[0,0,0,0],"isMain":false},{"metricName":"# mismatches per 100 kbp","quality":"Less is better","values":["2962.96","1475.41","2127.66","2172.76"],"isMain":true},{"metricName":"# indels per 100 kbp","quality":"Less is better","values":["0.00","0.00","0.00","0.00"],"isMain":true},{"metricName":" # indels (<= 5 bp)","quality":"Less is better","values":[0,0,0,0],"isMain":false},{"metricName":" # indels (> 5 bp)","quality":"Less is better","values":[0,0,0,0],"isMain":false},{"metricName":"# N's","quality":"Less is better","values":[0,0,0,0],"isMain":false},{"metricName":"# N's per 100 kbp","quality":"Less is better","values":["0.00","0.00","0.00","0.00"],"isMain":true}]],["Statistics without reference",[{"metricName":"# contigs","quality":"Equal","values":[1,2,3,2],"isMain":true},{"metricName":"# contigs (>= 1000 bp)","quality":"Equal","values":[0,0,1,1],"isMain":false},{"metricName":"# contigs (>= 5000 bp)","quality":"Equal","values":[0,0,0,0],"isMain":false},{"metricName":"# contigs (>= 10000 bp)","quality":"Equal","values":[0,0,0,0],"isMain":false},{"metricName":"# contigs (>= 25000 bp)","quality":"Equal","values":[0,0,0,0],"isMain":false},{"metricName":"# contigs (>= 50000 bp)","quality":"Equal","values":[0,0,0,0],"isMain":false},{"metricName":"Largest contig","quality":"More is better","values":[566,759,1104,1202],"isMain":true},{"metricName":"Total length","quality":"More is better","values":[566,1287,2194,2076],"isMain":true},{"metricName":"Total length (>= 1000 bp)","quality":"More is better","values":[0,0,1104,1202],"isMain":true},{"metricName":"Total length (>= 5000 bp)","quality":"More is better","values":[0,0,0,0],"isMain":false},{"metricName":"Total length (>= 10000 bp)","quality":"More is better","values":[0,0,0,0],"isMain":true},{"metricName":"Total length (>= 25000 bp)","quality":"More is better","values":[0,0,0,0],"isMain":false},{"metricName":"Total length (>= 50000 bp)","quality":"More is better","values":[0,0,0,0],"isMain":true},{"metricName":"N50","quality":"More is better","values":[566,759,1104,1202],"isMain":false},{"metricName":"N75","quality":"More is better","values":[566,528,570,874],"isMain":false},{"metricName":"L50","quality":"Less is better","values":[1,1,1,1],"isMain":false},{"metricName":"L75","quality":"Less is better","values":[1,2,2,2],"isMain":false},{"metricName":"GC (%)","quality":"Equal","values":["38.16","48.48","41.98","56.12"],"isMain":false}]],["K-mer-based statistics",[]],["Predicted genes",[]],["Similarity statistics",[]],["Reference statistics",[{"metricName":"Reference length","quality":"Equal","values":[2052669,2052669,2052669,2052669],"isMain":false},{"metricName":"Reference fragments","quality":"Equal","values":[11,11,11,11],"isMain":false},{"metricName":"Reference GC (%)","quality":"Equal","values":["37.83","37.83","37.83","37.83"],"isMain":false}]]],"subreferences":[],"subreports":[],"minContig":500},{"date":"10 February 2020, Monday, 20:33:45","assembliesNames":["megahit-103-df","megahit-113-df","megahit-113-ml","megahit-113-ms","megahit-129-df","metaSPAdes"],"referenceName":"GCF\_000391985.1\_Ente\_faec\_9731349\_1\_V1","order":[0,1,2,3,4,5],"report":[["Genome statistics",[{"metricName":"Genome fraction (%)","quality":"More is better","values":["6.821","8.596","9.131","10.724","9.734","4.593"],"isMain":true},{"metricName":"Duplication ratio","quality":"Less is better","values":["1.027","1.025","1.021","1.033","1.018","1.042"],"isMain":true},{"metricName":"Largest alignment","quality":"More is better","values":[1778,1900,2050,2064,2295,1946],"isMain":true},{"metricName":"Total aligned length","quality":"More is better","values":[178811,224974,239454,281541,254973,120084],"isMain":true},{"metricName":"NA50","quality":"More is better","values":[583,599,603,594,622,560],"isMain":false},{"metricName":"NA75","quality":"More is better","values":[528,533,540,527,535,515],"isMain":false},{"metricName":"NGA50","quality":"More is better","values":[null,null,null,null,null,null],"isMain":true},{"metricName":"LA50","quality":"Less is better","values":[128,151,163,200,163,87],"isMain":false},{"metricName":"LA75","quality":"Less is better","values":[212,256,270,333,277,147],"isMain":false}]],["Reads mapping",[]],["Misassemblies",[{"metricName":"# misassemblies","quality":"Less is better","values":[2,9,6,5,9,2],"isMain":true},{"metricName":" # relocations","quality":"Less is better","values":[0,0,0,0,0,0],"isMain":false},{"metricName":" # translocations","quality":"Less is better","values":[2,9,6,5,9,2],"isMain":false},{"metricName":" # inversions","quality":"Less is better","values":[0,0,0,0,0,0],"isMain":false},{"metricName":" # interspecies translocations","quality":"Less is better","values":[13,24,5,26,13,16],"isMain":false},{"metricName":"# misassembled contigs","quality":"Less is better","values":[2,9,6,5,8,2],"isMain":false},{"metricName":"Misassembled contigs length","quality":"Less is better","values":[1017,10456,6748,5708,5668,3871],"isMain":true},{"metricName":"# possibly misassembled contigs","quality":"Less is better","values":[0,0,0,0,0,0],"isMain":false},{"metricName":" # possible misassemblies","quality":"Less is better","values":[0,0,0,0,0,0],"isMain":false},{"metricName":"# local misassemblies","quality":"Less is better","values":[0,3,1,0,2,0],"isMain":false},{"metricName":"# scaffold gap ext. mis.","quality":"Less is better","values":[0,0,0,0,0,0],"isMain":false},{"metricName":"# scaffold gap loc. mis.","quality":"Less is better","values":[0,0,0,0,0,0],"isMain":false},{"metricName":"# unaligned mis. contigs","quality":"Less is better","values":[0,0,0,0,0,0],"isMain":false}]],["Unaligned",[{"metricName":"# fully unaligned contigs","quality":"Less is better","values":[0,1,0,2,0,0],"isMain":false},{"metricName":"Fully unaligned length","quality":"Less is better","values":[0,559,0,1088,0,0],"isMain":false},{"metricName":"# partially unaligned contigs","quality":"Less is better","values":[5,8,1,8,4,4],"isMain":false},{"metricName":"Partially unaligned length","quality":"Less is better","values":[4269,7111,577,6174,2692,2735],"isMain":false}]],["Mismatches",[{"metricName":"# mismatches","quality":"Less is better","values":[2630,3381,4475,5638,3838,1900],"isMain":false},{"metricName":"# indels","quality":"Less is better","values":[3,2,4,9,7,3],"isMain":false},{"metricName":"Indels length","quality":"Less is better","values":[5,2,4,9,9,3],"isMain":false},{"metricName":"# mismatches per 100 kbp","quality":"Less is better","values":["1477.74","1507.47","1878.44","2014.97","1511.26","1585.34"],"isMain":true},{"metricName":"# indels per 100 kbp","quality":"Less is better","values":["1.69","0.89","1.68","3.22","2.76","2.50"],"isMain":true},{"metricName":" # indels (<= 5 bp)","quality":"Less is better","values":[3,2,4,9,7,3],"isMain":false},{"metricName":" # indels (> 5 bp)","quality":"Less is better","values":[0,0,0,0,0,0],"isMain":false},{"metricName":"# N's","quality":"Less is better","values":[0,0,0,0,0,0],"isMain":false},{"metricName":"# N's per 100 kbp","quality":"Less is better","values":["0.00","0.00","0.00","0.00","0.00","0.00"],"isMain":true}]],["Statistics without reference",[{"metricName":"# contigs","quality":"Equal","values":[293,353,378,457,393,198],"isMain":true},{"metricName":"# contigs (>= 1000 bp)","quality":"Equal","values":[13,20,16,19,23,11],"isMain":false},{"metricName":"# contigs (>= 5000 bp)","quality":"Equal","values":[0,0,0,0,0,0],"isMain":false},{"metricName":"# contigs (>= 10000 bp)","quality":"Equal","values":[0,0,0,0,0,0],"isMain":false},{"metricName":"# contigs (>= 25000 bp)","quality":"Equal","values":[0,0,0,0,0,0],"isMain":false},{"metricName":"# contigs (>= 50000 bp)","quality":"Equal","values":[0,0,0,0,0,0],"isMain":false},{"metricName":"Largest contig","quality":"More is better","values":[2107,3239,3114,2387,2361,2238],"isMain":true},{"metricName":"Total length","quality":"More is better","values":[187065,237627,243811,296355,261308,127579],"isMain":true},{"metricName":"Total length (>= 1000 bp)","quality":"More is better","values":[18399,31339,22659,25828,31034,15819],"isMain":true},{"metricName":"Total length (>= 5000 bp)","quality":"More is better","values":[0,0,0,0,0,0],"isMain":false},{"metricName":"Total length (>= 10000 bp)","quality":"More is better","values":[0,0,0,0,0,0],"isMain":true},{"metricName":"Total length (>= 25000 bp)","quality":"More is better","values":[0,0,0,0,0,0],"isMain":false},{"metricName":"Total length (>= 50000 bp)","quality":"More is better","values":[0,0,0,0,0,0],"isMain":true},{"metricName":"N50","quality":"More is better","values":[602,628,612,620,637,595],"isMain":false},{"metricName":"N75","quality":"More is better","values":[537,554,549,548,554,533],"isMain":false},{"metricName":"L50","quality":"Less is better","values":[121,138,157,188,158,80],"isMain":false},{"metricName":"L75","quality":"Less is better","values":[203,240,262,315,269,137],"isMain":false},{"metricName":"GC (%)","quality":"Equal","values":["38.08","38.11","38.14","38.28","38.08","37.83"],"isMain":false}]],["K-mer-based statistics",[]],["Predicted genes",[]],["Similarity statistics",[]],["Reference statistics",[{"metricName":"Reference length","quality":"Equal","values":[2609133,2609133,2609133,2609133,2609133,2609133],"isMain":false},{"metricName":"Reference fragments","quality":"Equal","values":[59,59,59,59,59,59],"isMain":false},{"metricName":"Reference GC (%)","quality":"Equal","values":["38.05","38.05","38.05","38.05","38.05","38.05"],"isMain":false}]]],"subreferences":[],"subreports":[],"minContig":500},{"date":"10 February 2020, Monday, 20:34:11","assembliesNames":["megahit-103-df","megahit-113-df","megahit-113-ml","megahit-113-ms","megahit-129-df","metaSPAdes"],"referenceName":"GCF\_000396085.1\_Ente\_faec\_B1933\_V1","order":[0,1,2,3,4,5],"report":[["Genome statistics",[{"metricName":"Genome fraction (%)","quality":"More is better","values":["4.970","6.441","6.396","7.832","7.347","3.440"],"isMain":true},{"metricName":"Duplication ratio","quality":"Less is better","values":["1.031","1.028","1.011","1.028","1.022","1.045"],"isMain":true},{"metricName":"Largest alignment","quality":"More is better","values":[1358,2116,1445,1508,2116,1241],"isMain":true},{"metricName":"Total aligned length","quality":"More is better","values":[161626,209057,207751,254632,239105,111840],"isMain":true},{"metricName":"NA50","quality":"More is better","values":[594,618,604,614,616,575],"isMain":false},{"metricName":"NA75","quality":"More is better","values":[536,537,541,541,541,514],"isMain":false},{"metricName":"NGA50","quality":"More is better","values":[null,null,null,null,null,null],"isMain":true},{"metricName":"LA50","quality":"Less is better","values":[115,140,145,172,159,85],"isMain":false},{"metricName":"LA75","quality":"Less is better","values":[189,237,237,286,266,140],"isMain":false}]],["Reads mapping",[]],["Misassemblies",[{"metricName":"# misassemblies","quality":"Less is better","values":[1,1,2,0,1,1],"isMain":true},{"metricName":" # relocations","quality":"Less is better","values":[0,0,0,0,0,1],"isMain":false},{"metricName":" # translocations","quality":"Less is better","values":[1,1,2,0,1,0],"isMain":false},{"metricName":" # inversions","quality":"Less is better","values":[0,0,0,0,0,0],"isMain":false},{"metricName":" # interspecies translocations","quality":"Less is better","values":[15,19,2,20,15,14],"isMain":false},{"metricName":"# misassembled contigs","quality":"Less is better","values":[1,1,2,0,1,1],"isMain":false},{"metricName":"Misassembled contigs length","quality":"Less is better","values":[679,1296,2169,0,866,542],"isMain":true},{"metricName":"# possibly misassembled contigs","quality":"Less is better","values":[0,0,0,0,0,0],"isMain":false},{"metricName":" # possible misassemblies","quality":"Less is better","values":[0,0,0,0,0,0],"isMain":false},{"metricName":"# local misassemblies","quality":"Less is better","values":[0,0,0,0,0,0],"isMain":false},{"metricName":"# scaffold gap ext. mis.","quality":"Less is better","values":[0,0,0,0,0,0],"isMain":false},{"metricName":"# scaffold gap loc. mis.","quality":"Less is better","values":[0,0,0,0,0,0],"isMain":false},{"metricName":"# unaligned mis. contigs","quality":"Less is better","values":[0,0,0,0,0,0],"isMain":false}]],["Unaligned",[{"metricName":"# fully unaligned contigs","quality":"Less is better","values":[0,0,0,1,1,0],"isMain":false},{"metricName":"Fully unaligned length","quality":"Less is better","values":[0,0,0,660,612,0],"isMain":false},{"metricName":"# partially unaligned contigs","quality":"Less is better","values":[1,5,1,2,3,3],"isMain":false},{"metricName":"Partially unaligned length","quality":"Less is better","values":[1464,6960,777,1923,3741,2810],"isMain":false}]],["Mismatches",[{"metricName":"# mismatches","quality":"Less is better","values":[2552,3110,4094,5260,3761,1882],"isMain":false},{"metricName":"# indels","quality":"Less is better","values":[5,1,8,3,2,2],"isMain":false},{"metricName":"Indels length","quality":"Less is better","values":[47,1,9,3,2,4],"isMain":false},{"metricName":"# mismatches per 100 kbp","quality":"Less is better","values":["1582.88","1488.24","1973.02","2070.30","1577.92","1686.38"],"isMain":true},{"metricName":"# indels per 100 kbp","quality":"Less is better","values":["3.10","0.48","3.86","1.18","0.84","1.79"],"isMain":true},{"metricName":" # indels (<= 5 bp)","quality":"Less is better","values":[4,1,8,3,2,2],"isMain":false},{"metricName":" # indels (> 5 bp)","quality":"Less is better","values":[1,0,0,0,0,0],"isMain":false},{"metricName":"# N's","quality":"Less is better","values":[0,0,0,0,0,0],"isMain":false},{"metricName":"# N's per 100 kbp","quality":"Less is better","values":["0.00","0.00","0.00","0.00","0.00","0.00"],"isMain":true}]],["Statistics without reference",[{"metricName":"# contigs","quality":"Equal","values":[263,326,333,401,374,190],"isMain":true},{"metricName":"# contigs (>= 1000 bp)","quality":"Equal","values":[9,19,9,15,18,5],"isMain":false},{"metricName":"# contigs (>= 5000 bp)","quality":"Equal","values":[0,0,0,0,0,0],"isMain":false},{"metricName":"# contigs (>= 10000 bp)","quality":"Equal","values":[0,0,0,0,0,0],"isMain":false},{"metricName":"# contigs (>= 25000 bp)","quality":"Equal","values":[0,0,0,0,0,0],"isMain":false},{"metricName":"# contigs (>= 50000 bp)","quality":"Equal","values":[0,0,0,0,0,0],"isMain":false},{"metricName":"Largest contig","quality":"More is better","values":[1643,3684,1580,1592,2832,2158],"isMain":true},{"metricName":"Total length","quality":"More is better","values":[167682,221729,210559,263892,247927,119396],"isMain":true},{"metricName":"Total length (>= 1000 bp)","quality":"More is better","values":[10660,26275,10657,18134,23144,6595],"isMain":true},{"metricName":"Total length (>= 5000 bp)","quality":"More is better","values":[0,0,0,0,0,0],"isMain":false},{"metricName":"Total length (>= 10000 bp)","quality":"More is better","values":[0,0,0,0,0,0],"isMain":true},{"metricName":"Total length (>= 25000 bp)","quality":"More is better","values":[0,0,0,0,0,0],"isMain":false},{"metricName":"Total length (>= 50000 bp)","quality":"More is better","values":[0,0,0,0,0,0],"isMain":true},{"metricName":"N50","quality":"More is better","values":[614,647,611,641,639,594],"isMain":false},{"metricName":"N75","quality":"More is better","values":[556,562,550,559,559,537],"isMain":false},{"metricName":"L50","quality":"Less is better","values":[111,129,142,166,152,80],"isMain":false},{"metricName":"L75","quality":"Less is better","values":[183,221,233,277,256,133],"isMain":false},{"metricName":"GC (%)","quality":"Equal","values":["37.86","37.97","37.37","37.76","37.69","37.84"],"isMain":false}]],["K-mer-based statistics",[]],["Predicted genes",[]],["Similarity statistics",[]],["Reference statistics",[{"metricName":"Reference length","quality":"Equal","values":[3244166,3244166,3244166,3244166,3244166,3244166],"isMain":false},{"metricName":"Reference fragments","quality":"Equal","values":[34,34,34,34,34,34],"isMain":false},{"metricName":"Reference GC (%)","quality":"Equal","values":["37.12","37.12","37.12","37.12","37.12","37.12"],"isMain":false}]]],"subreferences":[],"subreports":[],"minContig":500},{"date":"10 February 2020, Monday, 20:33:55","assembliesNames":["megahit-103-df","megahit-113-df","megahit-113-ml","megahit-113-ms","megahit-129-df","metaSPAdes"],"referenceName":"GCF\_000396285.1\_Ente\_faec\_B2867\_V1","order":[0,1,2,3,4,5],"report":[["Genome statistics",[{"metricName":"Genome fraction (%)","quality":"More is better","values":["5.034","6.029","6.986","7.131","7.046","3.436"],"isMain":true},{"metricName":"Duplication ratio","quality":"Less is better","values":["1.027","1.026","1.017","1.026","1.025","1.044"],"isMain":true},{"metricName":"Largest alignment","quality":"More is better","values":[1600,1287,1256,1378,1447,1380],"isMain":true},{"metricName":"Total aligned length","quality":"More is better","values":[164992,197806,230156,234560,231859,112828],"isMain":true},{"metricName":"NA50","quality":"More is better","values":[580,610,601,597,639,576],"isMain":false},{"metricName":"NA75","quality":"More is better","values":[529,534,536,536,550,521],"isMain":false},{"metricName":"NGA50","quality":"More is better","values":[null,null,null,null,null,null],"isMain":true},{"metricName":"LA50","quality":"Less is better","values":[122,135,157,164,148,85],"isMain":false},{"metricName":"LA75","quality":"Less is better","values":[199,227,260,272,251,139],"isMain":false}]],["Reads mapping",[]],["Misassemblies",[{"metricName":"# misassemblies","quality":"Less is better","values":[1,0,0,0,2,0],"isMain":true},{"metricName":" # relocations","quality":"Less is better","values":[0,0,0,0,0,0],"isMain":false},{"metricName":" # translocations","quality":"Less is better","values":[1,0,0,0,1,0],"isMain":false},{"metricName":" # inversions","quality":"Less is better","values":[0,0,0,0,1,0],"isMain":false},{"metricName":" # interspecies translocations","quality":"Less is better","values":[17,18,3,16,18,15],"isMain":false},{"metricName":"# misassembled contigs","quality":"Less is better","values":[1,0,0,0,2,0],"isMain":false},{"metricName":"Misassembled contigs length","quality":"Less is better","values":[661,0,0,0,2164,0],"isMain":true},{"metricName":"# possibly misassembled contigs","quality":"Less is better","values":[0,0,0,0,0,0],"isMain":false},{"metricName":" # possible misassemblies","quality":"Less is better","values":[0,0,0,0,0,0],"isMain":false},{"metricName":"# local misassemblies","quality":"Less is better","values":[1,1,0,1,2,0],"isMain":false},{"metricName":"# scaffold gap ext. mis.","quality":"Less is better","values":[0,0,0,0,0,0],"isMain":false},{"metricName":"# scaffold gap loc. mis.","quality":"Less is better","values":[0,0,0,0,0,0],"isMain":false},{"metricName":"# unaligned mis. contigs","quality":"Less is better","values":[0,0,0,0,0,0],"isMain":false}]],["Unaligned",[{"metricName":"# fully unaligned contigs","quality":"Less is better","values":[1,1,0,0,0,0],"isMain":false},{"metricName":"Fully unaligned length","quality":"Less is better","values":[587,587,0,0,0,0],"isMain":false},{"metricName":"# partially unaligned contigs","quality":"Less is better","values":[2,5,1,6,4,2],"isMain":false},{"metricName":"Partially unaligned length","quality":"Less is better","values":[1211,4836,1180,5119,5038,1074],"isMain":false}]],["Mismatches",[{"metricName":"# mismatches","quality":"Less is better","values":[2639,3029,4530,5011,3544,1847],"isMain":false},{"metricName":"# indels","quality":"Less is better","values":[0,2,2,4,0,3],"isMain":false},{"metricName":"Indels length","quality":"Less is better","values":[0,2,2,61,0,46],"isMain":false},{"metricName":"# mismatches per 100 kbp","quality":"Less is better","values":["1600.69","1533.93","1979.97","2145.47","1535.80","1641.09"],"isMain":true},{"metricName":"# indels per 100 kbp","quality":"Less is better","values":["0.00","1.01","0.87","1.71","0.00","2.67"],"isMain":true},{"metricName":" # indels (<= 5 bp)","quality":"Less is better","values":[0,2,2,3,0,2],"isMain":false},{"metricName":" # indels (> 5 bp)","quality":"Less is better","values":[0,0,0,1,0,1],"isMain":false},{"metricName":"# N's","quality":"Less is better","values":[0,0,0,0,0,0],"isMain":false},{"metricName":"# N's per 100 kbp","quality":"Less is better","values":["0.00","0.00","0.00","0.00","0.00","0.00"],"isMain":true}]],["Statistics without reference",[{"metricName":"# contigs","quality":"Equal","values":[276,316,367,379,352,191],"isMain":true},{"metricName":"# contigs (>= 1000 bp)","quality":"Equal","values":[5,15,10,14,19,2],"isMain":false},{"metricName":"# contigs (>= 5000 bp)","quality":"Equal","values":[0,0,0,0,0,0],"isMain":false},{"metricName":"# contigs (>= 10000 bp)","quality":"Equal","values":[0,0,0,0,0,0],"isMain":false},{"metricName":"# contigs (>= 25000 bp)","quality":"Equal","values":[0,0,0,0,0,0],"isMain":false},{"metricName":"# contigs (>= 50000 bp)","quality":"Equal","values":[0,0,0,0,0,0],"isMain":false},{"metricName":"Largest contig","quality":"More is better","values":[1604,1709,1260,2048,2904,1382],"isMain":true},{"metricName":"Total length","quality":"More is better","values":[171100,207971,233923,244757,241607,118557],"isMain":true},{"metricName":"Total length (>= 1000 bp)","quality":"More is better","values":[6174,18229,11201,17255,24564,2404],"isMain":true},{"metricName":"Total length (>= 5000 bp)","quality":"More is better","values":[0,0,0,0,0,0],"isMain":false},{"metricName":"Total length (>= 10000 bp)","quality":"More is better","values":[0,0,0,0,0,0],"isMain":true},{"metricName":"Total length (>= 25000 bp)","quality":"More is better","values":[0,0,0,0,0,0],"isMain":false},{"metricName":"Total length (>= 50000 bp)","quality":"More is better","values":[0,0,0,0,0,0],"isMain":true},{"metricName":"N50","quality":"More is better","values":[596,632,613,616,666,603],"isMain":false},{"metricName":"N75","quality":"More is better","values":[544,552,545,553,570,540],"isMain":false},{"metricName":"L50","quality":"Less is better","values":[119,130,155,158,141,82],"isMain":false},{"metricName":"L75","quality":"Less is better","values":[194,218,256,263,239,134],"isMain":false},{"metricName":"GC (%)","quality":"Equal","values":["37.40","37.49","37.75","37.56","37.39","37.65"],"isMain":false}]],["K-mer-based statistics",[]],["Predicted genes",[]],["Similarity statistics",[]],["Reference statistics",[{"metricName":"Reference length","quality":"Equal","values":[3275195,3275195,3275195,3275195,3275195,3275195],"isMain":false},{"metricName":"Reference fragments","quality":"Equal","values":[60,60,60,60,60,60],"isMain":false},{"metricName":"Reference GC (%)","quality":"Equal","values":["37.06","37.06","37.06","37.06","37.06","37.06"],"isMain":false}]]],"subreferences":[],"subreports":[],"minContig":500},{"date":"10 February 2020, Monday, 20:34:12","assembliesNames":["megahit-103-df","megahit-113-df","megahit-113-ml","megahit-113-ms","megahit-129-df","metaSPAdes"],"referenceName":"GCF\_000398925.1\_Buty\_pull\_1\_20\_V1","order":[0,1,2,3,4,5],"report":[["Genome statistics",[{"metricName":"Genome fraction (%)","quality":"More is better","values":["6.587","7.162","7.435","8.230","8.551","4.621"],"isMain":true},{"metricName":"Duplication ratio","quality":"Less is better","values":["1.014","1.025","1.026","1.031","1.028","1.022"],"isMain":true},{"metricName":"Largest alignment","quality":"More is better","values":[1600,1969,3950,3290,6187,1111],"isMain":true},{"metricName":"Total aligned length","quality":"More is better","values":[218082,237439,247596,273948,285736,152658],"isMain":true},{"metricName":"NA50","quality":"More is better","values":[637,646,632,649,662,610],"isMain":false},{"metricName":"NA75","quality":"More is better","values":[555,545,542,541,558,534],"isMain":false},{"metricName":"NGA50","quality":"More is better","values":[null,null,null,null,null,null],"isMain":true},{"metricName":"LA50","quality":"Less is better","values":[142,143,153,166,166,103],"isMain":false},{"metricName":"LA75","quality":"Less is better","values":[236,246,261,288,290,172],"isMain":false}]],["Reads mapping",[]],["Misassemblies",[{"metricName":"# misassemblies","quality":"Less is better","values":[4,3,4,6,7,1],"isMain":true},{"metricName":" # relocations","quality":"Less is better","values":[1,2,1,2,2,1],"isMain":false},{"metricName":" # translocations","quality":"Less is better","values":[3,1,3,4,4,0],"isMain":false},{"metricName":" # inversions","quality":"Less is better","values":[0,0,0,0,1,0],"isMain":false},{"metricName":" # interspecies translocations","quality":"Less is better","values":[6,16,7,15,14,6],"isMain":false},{"metricName":"# misassembled contigs","quality":"Less is better","values":[3,3,3,6,7,1],"isMain":false},{"metricName":"Misassembled contigs length","quality":"Less is better","values":[2339,2498,2224,9646,5162,534],"isMain":true},{"metricName":"# possibly misassembled contigs","quality":"Less is better","values":[0,0,0,0,0,0],"isMain":false},{"metricName":" # possible misassemblies","quality":"Less is better","values":[0,0,0,0,0,0],"isMain":false},{"metricName":"# local misassemblies","quality":"Less is better","values":[2,0,0,2,0,0],"isMain":false},{"metricName":"# scaffold gap ext. mis.","quality":"Less is better","values":[0,0,0,0,0,0],"isMain":false},{"metricName":"# scaffold gap loc. mis.","quality":"Less is better","values":[0,0,0,0,0,0],"isMain":false},{"metricName":"# unaligned mis. contigs","quality":"Less is better","values":[0,0,0,0,0,0],"isMain":false}]],["Unaligned",[{"metricName":"# fully unaligned contigs","quality":"Less is better","values":[1,0,0,0,0,0],"isMain":false},{"metricName":"Fully unaligned length","quality":"Less is better","values":[630,0,0,0,0,0],"isMain":false},{"metricName":"# partially unaligned contigs","quality":"Less is better","values":[2,3,1,2,6,2],"isMain":false},{"metricName":"Partially unaligned length","quality":"Less is better","values":[2220,3300,863,5366,10132,1095],"isMain":false}]],["Mismatches",[{"metricName":"# mismatches","quality":"Less is better","values":[3223,3430,5078,5593,4128,2520],"isMain":false},{"metricName":"# indels","quality":"Less is better","values":[4,4,5,15,11,3],"isMain":false},{"metricName":"Indels length","quality":"Less is better","values":[6,4,17,21,11,3],"isMain":false},{"metricName":"# mismatches per 100 kbp","quality":"Less is better","values":["1483.00","1451.55","2069.97","2059.65","1463.14","1652.94"],"isMain":true},{"metricName":"# indels per 100 kbp","quality":"Less is better","values":["1.84","1.69","2.04","5.52","3.90","1.97"],"isMain":true},{"metricName":" # indels (<= 5 bp)","quality":"Less is better","values":[4,4,4,15,11,3],"isMain":false},{"metricName":" # indels (> 5 bp)","quality":"Less is better","values":[0,0,1,0,0,0],"isMain":false},{"metricName":"# N's","quality":"Less is better","values":[0,0,0,0,0,0],"isMain":false},{"metricName":"# N's per 100 kbp","quality":"Less is better","values":["0.00","0.00","0.00","0.00","0.00","0.00"],"isMain":true}]],["Statistics without reference",[{"metricName":"# contigs","quality":"Equal","values":[334,349,375,404,409,243],"isMain":true},{"metricName":"# contigs (>= 1000 bp)","quality":"Equal","values":[16,27,22,34,40,5],"isMain":false},{"metricName":"# contigs (>= 5000 bp)","quality":"Equal","values":[0,0,0,1,1,0],"isMain":false},{"metricName":"# contigs (>= 10000 bp)","quality":"Equal","values":[0,0,0,0,0,0],"isMain":false},{"metricName":"# contigs (>= 25000 bp)","quality":"Equal","values":[0,0,0,0,0,0],"isMain":false},{"metricName":"# contigs (>= 50000 bp)","quality":"Equal","values":[0,0,0,0,0,0],"isMain":false},{"metricName":"Largest contig","quality":"More is better","values":[1600,1981,3950,6626,9727,1134],"isMain":true},{"metricName":"Total length","quality":"More is better","values":[223161,245617,252618,285354,300187,156978],"isMain":true},{"metricName":"Total length (>= 1000 bp)","quality":"More is better","values":[18687,33854,30277,47598,59398,5349],"isMain":true},{"metricName":"Total length (>= 5000 bp)","quality":"More is better","values":[0,0,0,6626,9727,0],"isMain":false},{"metricName":"Total length (>= 10000 bp)","quality":"More is better","values":[0,0,0,0,0,0],"isMain":true},{"metricName":"Total length (>= 25000 bp)","quality":"More is better","values":[0,0,0,0,0,0],"isMain":false},{"metricName":"Total length (>= 50000 bp)","quality":"More is better","values":[0,0,0,0,0,0],"isMain":true},{"metricName":"N50","quality":"More is better","values":[647,682,651,684,699,628],"isMain":false},{"metricName":"N75","quality":"More is better","values":[564,572,554,561,582,549],"isMain":false},{"metricName":"L50","quality":"Less is better","values":[138,136,149,153,151,101],"isMain":false},{"metricName":"L75","quality":"Less is better","values":[230,234,256,270,269,168],"isMain":false},{"metricName":"GC (%)","quality":"Equal","values":["53.42","53.72","54.73","53.83","53.76","53.92"],"isMain":false}]],["K-mer-based statistics",[]],["Predicted genes",[]],["Similarity statistics",[]],["Reference statistics",[{"metricName":"Reference length","quality":"Equal","values":[3299517,3299517,3299517,3299517,3299517,3299517],"isMain":false},{"metricName":"Reference fragments","quality":"Equal","values":[17,17,17,17,17,17],"isMain":false},{"metricName":"Reference GC (%)","quality":"Equal","values":["54.08","54.08","54.08","54.08","54.08","54.08"],"isMain":false}]]],"subreferences":[],"subreports":[],"minContig":500},{"date":"10 February 2020, Monday, 20:35:06","assembliesNames":["megahit-103-df","megahit-113-df","megahit-113-ml","megahit-113-ms","megahit-129-df","metaSPAdes"],"referenceName":"GCF\_000403155.2\_Bact\_thet\_dnLKV9\_V1","order":[0,1,2,3,4,5],"report":[["Genome statistics",[{"metricName":"Genome fraction (%)","quality":"More is better","values":["82.863","80.355","80.514","80.158","85.252","90.960"],"isMain":true},{"metricName":"Duplication ratio","quality":"Less is better","values":["1.043","1.048","1.084","1.080","1.058","1.018"],"isMain":true},{"metricName":"Largest alignment","quality":"More is better","values":[25054,36075,16129,23961,43565,52515],"isMain":true},{"metricName":"Total aligned length","quality":"More is better","values":[5360477,5217274,5398522,5355974,5581423,5744606],"isMain":true},{"metricName":"NG50","quality":"More is better","values":[3638,5188,2390,2842,6022,7299],"isMain":false},{"metricName":"NG75","quality":"More is better","values":[1473,1974,1132,1320,2749,3332],"isMain":false},{"metricName":"NA50","quality":"More is better","values":[4276,6086,2694,3019,5527,7971],"isMain":false},{"metricName":"NA75","quality":"More is better","values":[2250,3168,1522,1539,2584,4111],"isMain":false},{"metricName":"NGA50","quality":"More is better","values":[3603,5039,2344,2617,5143,7131],"isMain":true},{"metricName":"NGA75","quality":"More is better","values":[1446,1826,1092,1132,2196,3306],"isMain":false},{"metricName":"LG50","quality":"Less is better","values":[477,316,735,567,265,246],"isMain":false},{"metricName":"LG75","quality":"Less is better","values":[1133,790,1664,1362,649,564],"isMain":false},{"metricName":"LA50","quality":"Less is better","values":[372,246,598,518,284,216],"isMain":false},{"metricName":"LA75","quality":"Less is better","values":[800,545,1266,1163,671,472],"isMain":false},{"metricName":"LGA50","quality":"Less is better","values":[479,329,749,633,313,246],"isMain":true},{"metricName":"LGA75","quality":"Less is better","values":[1143,829,1706,1525,770,566],"isMain":false}]],["Reads mapping",[]],["Misassemblies",[{"metricName":"# misassemblies","quality":"Less is better","values":[0,7,11,10,10,1],"isMain":true},{"metricName":" # relocations","quality":"Less is better","values":[0,5,3,5,2,0],"isMain":false},{"metricName":" # translocations","quality":"Less is better","values":[0,2,5,3,7,1],"isMain":false},{"metricName":" # inversions","quality":"Less is better","values":[0,0,3,2,1,0],"isMain":false},{"metricName":" # interspecies translocations","quality":"Less is better","values":[18,40,40,65,79,6],"isMain":false},{"metricName":"# misassembled contigs","quality":"Less is better","values":[0,7,11,10,10,1],"isMain":false},{"metricName":"Misassembled contigs length","quality":"Less is better","values":[0,77088,25933,32806,52831,2582],"isMain":true},{"metricName":"# possibly misassembled contigs","quality":"Less is better","values":[0,0,1,1,3,0],"isMain":false},{"metricName":" # possible misassemblies","quality":"Less is better","values":[0,0,1,1,3,0],"isMain":false},{"metricName":"# local misassemblies","quality":"Less is better","values":[2,2,1,6,4,2],"isMain":false},{"metricName":"# scaffold gap ext. mis.","quality":"Less is better","values":[0,0,0,0,0,0],"isMain":false},{"metricName":"# scaffold gap loc. mis.","quality":"Less is better","values":[0,0,0,0,0,0],"isMain":false},{"metricName":"# unaligned mis. contigs","quality":"Less is better","values":[0,0,0,2,2,0],"isMain":false}]],["Unaligned",[{"metricName":"# fully unaligned contigs","quality":"Less is better","values":[0,0,1,2,0,1],"isMain":false},{"metricName":"Fully unaligned length","quality":"Less is better","values":[0,0,859,2452,0,655],"isMain":false},{"metricName":"# partially unaligned contigs","quality":"Less is better","values":[9,21,12,29,34,4],"isMain":false},{"metricName":"Partially unaligned length","quality":"Less is better","values":[8326,60099,31900,188281,294229,2992],"isMain":false}]],["Mismatches",[{"metricName":"# mismatches","quality":"Less is better","values":[26123,33393,51173,51949,33315,19570],"isMain":false},{"metricName":"# indels","quality":"Less is better","values":[211,372,276,294,389,273],"isMain":false},{"metricName":"Indels length","quality":"Less is better","values":[653,1235,837,668,1224,633],"isMain":false},{"metricName":"# mismatches per 100 kbp","quality":"Less is better","values":["507.05","668.40","1022.27","1042.38","628.53","346.05"],"isMain":true},{"metricName":"# indels per 100 kbp","quality":"Less is better","values":["4.10","7.45","5.51","5.90","7.34","4.83"],"isMain":true},{"metricName":" # indels (<= 5 bp)","quality":"Less is better","values":[190,335,252,274,359,254],"isMain":false},{"metricName":" # indels (> 5 bp)","quality":"Less is better","values":[21,37,24,20,30,19],"isMain":false},{"metricName":"# N's","quality":"Less is better","values":[0,0,0,0,0,0],"isMain":false},{"metricName":"# N's per 100 kbp","quality":"Less is better","values":["0.00","0.00","0.00","0.00","0.00","0.00"],"isMain":true}]],["Statistics without reference",[{"metricName":"# contigs","quality":"Equal","values":[1901,1369,2692,2428,1578,1253],"isMain":true},{"metricName":"# contigs (>= 1000 bp)","quality":"Equal","values":[1457,1080,1816,1695,1201,1050],"isMain":false},{"metricName":"# contigs (>= 5000 bp)","quality":"Equal","values":[287,337,165,214,339,380],"isMain":false},{"metricName":"# contigs (>= 10000 bp)","quality":"Equal","values":[64,102,21,31,108,135],"isMain":false},{"metricName":"# contigs (>= 25000 bp)","quality":"Equal","values":[1,9,0,2,13,11],"isMain":false},{"metricName":"# contigs (>= 50000 bp)","quality":"Equal","values":[0,0,0,1,2,1],"isMain":false},{"metricName":"Largest contig","quality":"More is better","values":[25067,38870,19532,99460,100734,52519],"isMain":true},{"metricName":"Total length","quality":"More is better","values":[5381225,5296516,5457859,5575360,5900125,5758460],"isMain":true},{"metricName":"Total length (>= 1000 bp)","quality":"More is better","values":[5059854,5090945,4825864,5047399,5632824,5606977],"isMain":true},{"metricName":"Total length (>= 5000 bp)","quality":"More is better","values":[2306162,3216715,1219851,1792914,3512324,3908922],"isMain":false},{"metricName":"Total length (>= 10000 bp)","quality":"More is better","values":[811307,1614288,269472,544543,1908529,2171497],"isMain":true},{"metricName":"Total length (>= 25000 bp)","quality":"More is better","values":[25067,274478,0,136359,518796,373173],"isMain":false},{"metricName":"Total length (>= 50000 bp)","quality":"More is better","values":[0,0,0,99460,154100,52519],"isMain":true},{"metricName":"N50","quality":"More is better","values":[4288,6331,2738,3322,6553,7971],"isMain":false},{"metricName":"N75","quality":"More is better","values":[2292,3337,1581,1755,3169,4142],"isMain":false},{"metricName":"L50","quality":"Less is better","values":[372,236,586,463,240,216],"isMain":false},{"metricName":"L75","quality":"Less is better","values":[795,524,1239,1044,569,471],"isMain":false},{"metricName":"GC (%)","quality":"Equal","values":["42.83","42.78","42.86","43.00","42.90","42.89"],"isMain":false}]],["K-mer-based statistics",[]],["Predicted genes",[]],["Similarity statistics",[]],["Reference statistics",[{"metricName":"Reference length","quality":"Equal","values":[6217351,6217351,6217351,6217351,6217351,6217351],"isMain":false},{"metricName":"Reference fragments","quality":"Equal","values":[20,20,20,20,20,20],"isMain":false},{"metricName":"Reference GC (%)","quality":"Equal","values":["42.88","42.88","42.88","42.88","42.88","42.88"],"isMain":false}]]],"subreferences":[],"subreports":[],"minContig":500},{"date":"10 February 2020, Monday, 20:34:18","assembliesNames":["megahit-103-df","megahit-113-df","megahit-113-ml","megahit-113-ms","megahit-129-df","metaSPAdes"],"referenceName":"GCF\_000403235.1\_Bact\_vulg\_dnLKV7\_V1","order":[0,1,2,3,4,5],"report":[["Genome statistics",[{"metricName":"Genome fraction (%)","quality":"More is better","values":["7.893","7.807","12.131","11.922","8.287","4.957"],"isMain":true},{"metricName":"Duplication ratio","quality":"Less is better","values":["1.020","1.031","1.040","1.042","1.028","1.022"],"isMain":true},{"metricName":"Largest alignment","quality":"More is better","values":[7109,12633,17882,6542,20325,6599],"isMain":true},{"metricName":"Total aligned length","quality":"More is better","values":[408008,404954,639103,626165,429935,255564],"isMain":true},{"metricName":"NA50","quality":"More is better","values":[1942,2728,1368,1415,2271,2149],"isMain":false},{"metricName":"NA75","quality":"More is better","values":[978,1040,807,785,705,799],"isMain":false},{"metricName":"NGA50","quality":"More is better","values":[null,null,null,null,null,null],"isMain":true},{"metricName":"LA50","quality":"Less is better","values":[66,40,144,141,47,44],"isMain":false},{"metricName":"LA75","quality":"Less is better","values":[142,105,302,293,144,101],"isMain":false}]],["Reads mapping",[]],["Misassemblies",[{"metricName":"# misassemblies","quality":"Less is better","values":[4,3,1,7,5,0],"isMain":true},{"metricName":" # relocations","quality":"Less is better","values":[0,0,0,0,0,0],"isMain":false},{"metricName":" # translocations","quality":"Less is better","values":[3,3,1,4,5,0],"isMain":false},{"metricName":" # inversions","quality":"Less is better","values":[1,0,0,3,0,0],"isMain":false},{"metricName":" # interspecies translocations","quality":"Less is better","values":[12,21,15,26,21,8],"isMain":false},{"metricName":"# misassembled contigs","quality":"Less is better","values":[4,3,1,7,4,0],"isMain":false},{"metricName":"Misassembled contigs length","quality":"Less is better","values":[7557,19669,1845,10784,8087,0],"isMain":true},{"metricName":"# possibly misassembled contigs","quality":"Less is better","values":[0,0,0,0,0,0],"isMain":false},{"metricName":" # possible misassemblies","quality":"Less is better","values":[0,0,0,0,0,0],"isMain":false},{"metricName":"# local misassemblies","quality":"Less is better","values":[0,2,0,1,2,0],"isMain":false},{"metricName":"# scaffold gap ext. mis.","quality":"Less is better","values":[0,0,0,0,0,0],"isMain":false},{"metricName":"# scaffold gap loc. mis.","quality":"Less is better","values":[0,0,0,0,0,0],"isMain":false},{"metricName":"# unaligned mis. contigs","quality":"Less is better","values":[0,0,0,0,0,0],"isMain":false}]],["Unaligned",[{"metricName":"# fully unaligned contigs","quality":"Less is better","values":[0,0,0,0,0,0],"isMain":false},{"metricName":"Fully unaligned length","quality":"Less is better","values":[0,0,0,0,0,0],"isMain":false},{"metricName":"# partially unaligned contigs","quality":"Less is better","values":[2,5,5,9,8,2],"isMain":false},{"metricName":"Partially unaligned length","quality":"Less is better","values":[3589,7373,13121,9460,69147,37296],"isMain":false}]],["Mismatches",[{"metricName":"# mismatches","quality":"Less is better","values":[3680,4191,8855,9328,4178,1920],"isMain":false},{"metricName":"# indels","quality":"Less is better","values":[26,49,38,50,61,26],"isMain":false},{"metricName":"Indels length","quality":"Less is better","values":[141,64,130,77,141,163],"isMain":false},{"metricName":"# mismatches per 100 kbp","quality":"Less is better","values":["912.06","1050.13","1428.02","1530.67","986.26","757.79"],"isMain":true},{"metricName":"# indels per 100 kbp","quality":"Less is better","values":["6.44","12.28","6.13","8.20","14.40","10.26"],"isMain":true},{"metricName":" # indels (<= 5 bp)","quality":"Less is better","values":[22,49,34,49,59,21],"isMain":false},{"metricName":" # indels (> 5 bp)","quality":"Less is better","values":[4,0,4,1,2,5],"isMain":false},{"metricName":"# N's","quality":"Less is better","values":[0,0,0,0,0,0],"isMain":false},{"metricName":"# N's per 100 kbp","quality":"Less is better","values":["0.00","0.00","0.00","0.00","0.00","0.00"],"isMain":true}]],["Statistics without reference",[{"metricName":"# contigs","quality":"Equal","values":[286,244,535,536,236,158],"isMain":true},{"metricName":"# contigs (>= 1000 bp)","quality":"Equal","values":[142,111,240,232,112,85],"isMain":false},{"metricName":"# contigs (>= 5000 bp)","quality":"Equal","values":[3,16,6,3,18,5],"isMain":false},{"metricName":"# contigs (>= 10000 bp)","quality":"Equal","values":[0,4,1,0,6,1],"isMain":false},{"metricName":"# contigs (>= 25000 bp)","quality":"Equal","values":[0,0,1,0,1,1],"isMain":false},{"metricName":"# contigs (>= 50000 bp)","quality":"Equal","values":[0,0,0,0,1,0],"isMain":false},{"metricName":"Largest contig","quality":"More is better","values":[7112,12715,28106,6665,60580,37160],"isMain":true},{"metricName":"Total length","quality":"More is better","values":[415286,418968,658092,644554,504794,296148],"isMain":true},{"metricName":"Total length (>= 1000 bp)","quality":"More is better","values":[317183,331470,453584,438353,421096,245382],"isMain":true},{"metricName":"Total length (>= 5000 bp)","quality":"More is better","values":[19814,122949,59491,19448,214280,60789],"isMain":false},{"metricName":"Total length (>= 10000 bp)","quality":"More is better","values":[0,45435,28106,0,135771,37160],"isMain":true},{"metricName":"Total length (>= 25000 bp)","quality":"More is better","values":[0,0,28106,0,60580,37160],"isMain":false},{"metricName":"Total length (>= 50000 bp)","quality":"More is better","values":[0,0,0,0,60580,0],"isMain":true},{"metricName":"N50","quality":"More is better","values":[2099,3059,1408,1467,4015,2773],"isMain":false},{"metricName":"N75","quality":"More is better","values":[1063,1164,864,829,1591,1391],"isMain":false},{"metricName":"L50","quality":"Less is better","values":[64,38,134,137,27,29],"isMain":false},{"metricName":"L75","quality":"Less is better","values":[137,95,284,282,79,66],"isMain":false},{"metricName":"GC (%)","quality":"Equal","values":["41.67","41.48","42.36","42.17","40.63","41.23"],"isMain":false}]],["K-mer-based statistics",[]],["Predicted genes",[]],["Similarity statistics",[]],["Reference statistics",[{"metricName":"Reference length","quality":"Equal","values":[5111694,5111694,5111694,5111694,5111694,5111694],"isMain":false},{"metricName":"Reference fragments","quality":"Equal","values":[36,36,36,36,36,36],"isMain":false},{"metricName":"Reference GC (%)","quality":"Equal","values":["42.41","42.41","42.41","42.41","42.41","42.41"],"isMain":false}]]],"subreferences":[],"subreports":[],"minContig":500},{"date":"10 February 2020, Monday, 20:35:02","assembliesNames":["megahit-103-df","megahit-113-df","megahit-113-ml","megahit-113-ms","megahit-129-df","metaSPAdes"],"referenceName":"GCF\_000403255.1\_Lach\_bact\_3\_1\_V1","order":[0,1,2,3,4,5],"report":[["Genome statistics",[{"metricName":"Genome fraction (%)","quality":"More is better","values":["95.682","96.103","96.469","96.480","96.416","94.570"],"isMain":true},{"metricName":"Duplication ratio","quality":"Less is better","values":["1.007","1.011","1.013","1.017","1.014","1.006"],"isMain":true},{"metricName":"Largest alignment","quality":"More is better","values":[95007,151785,137989,110765,217793,207713],"isMain":true},{"metricName":"Total aligned length","quality":"More is better","values":[4863226,4899807,4930895,4946078,4931866,4804764],"isMain":true},{"metricName":"NG50","quality":"More is better","values":[33157,46598,26054,34829,47827,48959],"isMain":false},{"metricName":"NG75","quality":"More is better","values":[14227,19134,12587,15698,22493,22336],"isMain":false},{"metricName":"NA50","quality":"More is better","values":[34765,46337,26054,34119,42793,49784],"isMain":false},{"metricName":"NA75","quality":"More is better","values":[15328,17379,13326,14885,17383,27304],"isMain":false},{"metricName":"NGA50","quality":"More is better","values":[32823,45891,25736,34119,42793,46049],"isMain":true},{"metricName":"NGA75","quality":"More is better","values":[13842,16474,12450,14885,17908,22246],"isMain":false},{"metricName":"LG50","quality":"Less is better","values":[50,31,53,47,30,28],"isMain":false},{"metricName":"LG75","quality":"Less is better","values":[106,71,122,100,65,64],"isMain":false},{"metricName":"LA50","quality":"Less is better","values":[48,34,53,49,34,27],"isMain":false},{"metricName":"LA75","quality":"Less is better","values":[98,75,119,106,78,59],"isMain":false},{"metricName":"LGA50","quality":"Less is better","values":[51,35,55,49,34,30],"isMain":true},{"metricName":"LGA75","quality":"Less is better","values":[107,79,126,106,77,67],"isMain":false}]],["Reads mapping",[]],["Misassemblies",[{"metricName":"# misassemblies","quality":"Less is better","values":[6,22,14,19,23,7],"isMain":true},{"metricName":" # relocations","quality":"Less is better","values":[0,2,1,0,0,0],"isMain":false},{"metricName":" # translocations","quality":"Less is better","values":[6,20,13,19,22,7],"isMain":false},{"metricName":" # inversions","quality":"Less is better","values":[0,0,0,0,1,0],"isMain":false},{"metricName":" # interspecies translocations","quality":"Less is better","values":[12,19,11,27,17,4],"isMain":false},{"metricName":"# misassembled contigs","quality":"Less is better","values":[6,20,13,18,21,6],"isMain":false},{"metricName":"Misassembled contigs length","quality":"Less is better","values":[278062,1011231,312944,476538,863377,399187],"isMain":true},{"metricName":"# possibly misassembled contigs","quality":"Less is better","values":[0,0,0,1,0,0],"isMain":false},{"metricName":" # possible misassemblies","quality":"Less is better","values":[0,0,0,2,0,0],"isMain":false},{"metricName":"# local misassemblies","quality":"Less is better","values":[1,3,1,5,5,1],"isMain":false},{"metricName":"# scaffold gap ext. mis.","quality":"Less is better","values":[0,0,0,0,0,0],"isMain":false},{"metricName":"# scaffold gap loc. mis.","quality":"Less is better","values":[0,0,0,0,0,0],"isMain":false},{"metricName":"# unaligned mis. contigs","quality":"Less is better","values":[0,1,1,0,0,1],"isMain":false}]],["Unaligned",[{"metricName":"# fully unaligned contigs","quality":"Less is better","values":[0,0,0,2,2,0],"isMain":false},{"metricName":"Fully unaligned length","quality":"Less is better","values":[0,0,0,1289,1927,0],"isMain":false},{"metricName":"# partially unaligned contigs","quality":"Less is better","values":[4,9,3,14,10,2],"isMain":false},{"metricName":"Partially unaligned length","quality":"Less is better","values":[2753,38691,8212,90441,153348,3319],"isMain":false}]],["Mismatches",[{"metricName":"# mismatches","quality":"Less is better","values":[2872,4249,7163,8793,4786,2301],"isMain":false},{"metricName":"# indels","quality":"Less is better","values":[19,37,55,76,61,93],"isMain":false},{"metricName":"Indels length","quality":"Less is better","values":[66,209,229,209,510,200],"isMain":false},{"metricName":"# mismatches per 100 kbp","quality":"Less is better","values":["59.42","87.53","147.00","180.43","98.27","48.17"],"isMain":true},{"metricName":"# indels per 100 kbp","quality":"Less is better","values":["0.39","0.76","1.13","1.56","1.25","1.95"],"isMain":true},{"metricName":" # indels (<= 5 bp)","quality":"Less is better","values":[17,32,50,70,46,88],"isMain":false},{"metricName":" # indels (> 5 bp)","quality":"Less is better","values":[2,5,5,6,15,5],"isMain":false},{"metricName":"# N's","quality":"Less is better","values":[0,0,0,0,0,0],"isMain":false},{"metricName":"# N's per 100 kbp","quality":"Less is better","values":["0.00","0.00","0.00","0.00","0.00","0.00"],"isMain":true}]],["Statistics without reference",[{"metricName":"# contigs","quality":"Equal","values":[430,364,448,428,401,285],"isMain":true},{"metricName":"# contigs (>= 1000 bp)","quality":"Equal","values":[337,288,349,325,276,248],"isMain":false},{"metricName":"# contigs (>= 5000 bp)","quality":"Equal","values":[180,151,215,189,148,128],"isMain":false},{"metricName":"# contigs (>= 10000 bp)","quality":"Equal","values":[127,102,144,132,103,98],"isMain":false},{"metricName":"# contigs (>= 25000 bp)","quality":"Equal","values":[69,58,59,70,62,60],"isMain":false},{"metricName":"# contigs (>= 50000 bp)","quality":"Equal","values":[20,27,18,19,27,25],"isMain":false},{"metricName":"Largest contig","quality":"More is better","values":[96639,229684,137989,110787,217793,207713],"isMain":true},{"metricName":"Total length","quality":"More is better","values":[4869977,4947365,4944622,5047891,5095315,4809348],"isMain":true},{"metricName":"Total length (>= 1000 bp)","quality":"More is better","values":[4805917,4895816,4877433,4978516,5009912,4783464],"isMain":true},{"metricName":"Total length (>= 5000 bp)","quality":"More is better","values":[4436557,4556127,4554911,4644853,4696891,4517614],"isMain":false},{"metricName":"Total length (>= 10000 bp)","quality":"More is better","values":[4052277,4203201,4045537,4222875,4371741,4303945],"isMain":true},{"metricName":"Total length (>= 25000 bp)","quality":"More is better","values":[3100634,3516465,2694287,3209536,3716514,3696797],"isMain":false},{"metricName":"Total length (>= 50000 bp)","quality":"More is better","values":[1331467,2361300,1281381,1373172,2412154,2389617],"isMain":true},{"metricName":"N50","quality":"More is better","values":[35510,47576,26727,34829,47827,49796],"isMain":false},{"metricName":"N75","quality":"More is better","values":[15677,20631,13341,15698,21890,27926],"isMain":false},{"metricName":"L50","quality":"Less is better","values":[47,30,51,47,30,26],"isMain":false},{"metricName":"L75","quality":"Less is better","values":[97,67,116,100,67,57],"isMain":false},{"metricName":"GC (%)","quality":"Equal","values":["42.08","42.14","42.15","42.25","42.21","41.97"],"isMain":false}]],["K-mer-based statistics",[]],["Predicted genes",[]],["Similarity statistics",[]],["Reference statistics",[{"metricName":"Reference length","quality":"Equal","values":[5051289,5051289,5051289,5051289,5051289,5051289],"isMain":false},{"metricName":"Reference fragments","quality":"Equal","values":[50,50,50,50,50,50],"isMain":false},{"metricName":"Reference GC (%)","quality":"Equal","values":["42.35","42.35","42.35","42.35","42.35","42.35"],"isMain":false}]]],"subreferences":[],"subreports":[],"minContig":500},{"date":"10 February 2020, Monday, 20:34:48","assembliesNames":["megahit-103-df","megahit-113-df","megahit-113-ml","megahit-113-ms","megahit-129-df","metaSPAdes"],"referenceName":"GCF\_000403275.1\_Lach\_bact\_28\_4\_V1","order":[0,1,2,3,4,5],"report":[["Genome statistics",[{"metricName":"Genome fraction (%)","quality":"More is better","values":["95.595","95.680","96.000","95.920","95.987","94.058"],"isMain":true},{"metricName":"Duplication ratio","quality":"Less is better","values":["1.009","1.011","1.027","1.023","1.016","1.004"],"isMain":true},{"metricName":"Largest alignment","quality":"More is better","values":[113603,129395,50382,60933,102485,145129],"isMain":true},{"metricName":"Total aligned length","quality":"More is better","values":[4072947,4079096,4155412,4135510,4110563,3985900],"isMain":true},{"metricName":"NG50","quality":"More is better","values":[24416,37174,12728,21759,34033,40359],"isMain":false},{"metricName":"NG75","quality":"More is better","values":[12546,18285,6579,11156,18076,15954],"isMain":false},{"metricName":"NA50","quality":"More is better","values":[25388,34308,12673,15865,32538,42410],"isMain":false},{"metricName":"NA75","quality":"More is better","values":[13787,16581,6430,7028,14447,19852],"isMain":false},{"metricName":"NGA50","quality":"More is better","values":[24416,33815,12643,17018,32708,40349],"isMain":true},{"metricName":"NGA75","quality":"More is better","values":[12448,16122,6393,8393,15270,15946],"isMain":false},{"metricName":"LG50","quality":"Less is better","values":[51,36,104,62,37,31],"isMain":false},{"metricName":"LG75","quality":"Less is better","values":[111,77,217,132,77,69],"isMain":false},{"metricName":"LA50","quality":"Less is better","values":[48,37,104,84,40,28],"isMain":false},{"metricName":"LA75","quality":"Less is better","values":[103,82,219,186,86,60],"isMain":false},{"metricName":"LGA50","quality":"Less is better","values":[51,38,105,76,39,31],"isMain":true},{"metricName":"LGA75","quality":"Less is better","values":[111,83,222,164,83,70],"isMain":false}]],["Reads mapping",[]],["Misassemblies",[{"metricName":"# misassemblies","quality":"Less is better","values":[2,14,14,13,12,3],"isMain":true},{"metricName":" # relocations","quality":"Less is better","values":[0,3,1,1,1,0],"isMain":false},{"metricName":" # translocations","quality":"Less is better","values":[2,10,12,12,9,3],"isMain":false},{"metricName":" # inversions","quality":"Less is better","values":[0,1,1,0,2,0],"isMain":false},{"metricName":" # interspecies translocations","quality":"Less is better","values":[5,19,21,31,30,3],"isMain":false},{"metricName":"# misassembled contigs","quality":"Less is better","values":[2,14,13,13,12,3],"isMain":false},{"metricName":"Misassembled contigs length","quality":"Less is better","values":[34653,411398,80483,214619,229939,180376],"isMain":true},{"metricName":"# possibly misassembled contigs","quality":"Less is better","values":[0,0,0,0,0,0],"isMain":false},{"metricName":" # possible misassemblies","quality":"Less is better","values":[0,0,0,0,0,0],"isMain":false},{"metricName":"# local misassemblies","quality":"Less is better","values":[1,4,3,4,3,0],"isMain":false},{"metricName":"# scaffold gap ext. mis.","quality":"Less is better","values":[0,0,0,0,0,0],"isMain":false},{"metricName":"# scaffold gap loc. mis.","quality":"Less is better","values":[0,0,0,0,0,0],"isMain":false},{"metricName":"# unaligned mis. contigs","quality":"Less is better","values":[0,1,1,1,0,0],"isMain":false}]],["Unaligned",[{"metricName":"# fully unaligned contigs","quality":"Less is better","values":[0,2,1,1,3,1],"isMain":false},{"metricName":"Fully unaligned length","quality":"Less is better","values":[0,2042,781,582,2037,912],"isMain":false},{"metricName":"# partially unaligned contigs","quality":"Less is better","values":[3,14,7,22,18,1],"isMain":false},{"metricName":"Partially unaligned length","quality":"Less is better","values":[2722,106392,34105,314939,152570,714],"isMain":false}]],["Mismatches",[{"metricName":"# mismatches","quality":"Less is better","values":[3162,4264,11769,11672,4841,2157],"isMain":false},{"metricName":"# indels","quality":"Less is better","values":[26,52,48,76,85,72],"isMain":false},{"metricName":"Indels length","quality":"Less is better","values":[149,205,272,316,544,183],"isMain":false},{"metricName":"# mismatches per 100 kbp","quality":"Less is better","values":["78.32","105.53","290.29","288.14","119.42","54.30"],"isMain":true},{"metricName":"# indels per 100 kbp","quality":"Less is better","values":["0.64","1.29","1.18","1.88","2.10","1.81"],"isMain":true},{"metricName":" # indels (<= 5 bp)","quality":"Less is better","values":[24,47,43,69,68,67],"isMain":false},{"metricName":" # indels (> 5 bp)","quality":"Less is better","values":[2,5,5,7,17,5],"isMain":false},{"metricName":"# N's","quality":"Less is better","values":[0,0,0,0,0,0],"isMain":false},{"metricName":"# N's per 100 kbp","quality":"Less is better","values":["0.00","0.00","0.00","0.00","0.00","0.00"],"isMain":true}]],["Statistics without reference",[{"metricName":"# contigs","quality":"Equal","values":[394,347,611,516,380,261],"isMain":true},{"metricName":"# contigs (>= 1000 bp)","quality":"Equal","values":[298,266,517,412,261,223],"isMain":false},{"metricName":"# contigs (>= 5000 bp)","quality":"Equal","values":[178,142,271,230,156,126],"isMain":false},{"metricName":"# contigs (>= 10000 bp)","quality":"Equal","values":[127,112,141,146,115,98],"isMain":false},{"metricName":"# contigs (>= 25000 bp)","quality":"Equal","values":[50,54,22,49,61,52],"isMain":false},{"metricName":"# contigs (>= 50000 bp)","quality":"Equal","values":[11,17,1,6,20,21],"isMain":false},{"metricName":"Largest contig","quality":"More is better","values":[113603,129395,50392,108763,105763,145129],"isMain":true},{"metricName":"Total length","quality":"More is better","values":[4077774,4193035,4198682,4458739,4272849,3988801],"isMain":true},{"metricName":"Total length (>= 1000 bp)","quality":"More is better","values":[4010701,4136033,4133608,4386839,4189696,3960292],"isMain":true},{"metricName":"Total length (>= 5000 bp)","quality":"More is better","values":[3735568,3860810,3485552,3937162,3967058,3730794],"isMain":false},{"metricName":"Total length (>= 10000 bp)","quality":"More is better","values":[3359299,3650366,2538439,3323180,3678688,3539206],"isMain":true},{"metricName":"Total length (>= 25000 bp)","quality":"More is better","values":[2093951,2675134,745562,1826643,2831335,2835289],"isMain":false},{"metricName":"Total length (>= 50000 bp)","quality":"More is better","values":[749845,1286317,50392,445639,1408936,1710376],"isMain":true},{"metricName":"N50","quality":"More is better","values":[25388,37174,12733,19702,33804,42410],"isMain":false},{"metricName":"N75","quality":"More is better","values":[13936,18433,6785,9694,16896,20807],"isMain":false},{"metricName":"L50","quality":"Less is better","values":[48,36,103,67,38,28],"isMain":false},{"metricName":"L75","quality":"Less is better","values":[102,76,214,149,79,59],"isMain":false},{"metricName":"GC (%)","quality":"Equal","values":["44.96","44.96","44.97","44.80","44.95","44.88"],"isMain":false}]],["K-mer-based statistics",[]],["Predicted genes",[]],["Similarity statistics",[]],["Reference statistics",[{"metricName":"Reference length","quality":"Equal","values":[4223124,4223124,4223124,4223124,4223124,4223124],"isMain":false},{"metricName":"Reference fragments","quality":"Equal","values":[35,35,35,35,35,35],"isMain":false},{"metricName":"Reference GC (%)","quality":"Equal","values":["45.08","45.08","45.08","45.08","45.08","45.08"],"isMain":false}]]],"subreferences":[],"subreports":[],"minContig":500},{"date":"10 February 2020, Monday, 20:34:10","assembliesNames":["megahit-103-df","megahit-113-df","megahit-113-ml","megahit-113-ms","megahit-129-df","metaSPAdes"],"referenceName":"GCF\_000403355.2\_Ente\_caec\_B7\_V1","order":[0,1,2,3,4,5],"report":[["Genome statistics",[{"metricName":"Genome fraction (%)","quality":"More is better","values":["0.142","0.503","0.051","0.564","0.580","0.005"],"isMain":true},{"metricName":"Duplication ratio","quality":"Less is better","values":["1.000","1.000","1.330","1.000","1.028","3.639"],"isMain":true},{"metricName":"Largest alignment","quality":"More is better","values":[1813,5277,909,5277,5277,147],"isMain":true},{"metricName":"Total aligned length","quality":"More is better","values":[4169,14801,1488,16596,17077,147],"isMain":true},{"metricName":"NA50","quality":"More is better","values":[null,null,579,null,null,null],"isMain":false},{"metricName":"NA75","quality":"More is better","values":[null,null,579,null,null,null],"isMain":false},{"metricName":"NGA50","quality":"More is better","values":[null,null,null,null,null,null],"isMain":true},{"metricName":"LA50","quality":"Less is better","values":[null,null,2,null,null,null],"isMain":false},{"metricName":"LA75","quality":"Less is better","values":[null,null,2,null,null,null],"isMain":false}]],["Reads mapping",[]],["Misassemblies",[{"metricName":"# misassemblies","quality":"Less is better","values":[0,0,0,0,0,0],"isMain":true},{"metricName":" # relocations","quality":"Less is better","values":[0,0,0,0,0,0],"isMain":false},{"metricName":" # translocations","quality":"Less is better","values":[0,0,0,0,0,0],"isMain":false},{"metricName":" # inversions","quality":"Less is better","values":[0,0,0,0,0,0],"isMain":false},{"metricName":" # interspecies translocations","quality":"Less is better","values":[1,1,1,3,2,1],"isMain":false},{"metricName":"# misassembled contigs","quality":"Less is better","values":[0,0,0,0,0,0],"isMain":false},{"metricName":"Misassembled contigs length","quality":"Less is better","values":[0,0,0,0,0,0],"isMain":true},{"metricName":"# possibly misassembled contigs","quality":"Less is better","values":[0,0,0,0,0,0],"isMain":false},{"metricName":" # possible misassemblies","quality":"Less is better","values":[0,0,0,0,0,0],"isMain":false},{"metricName":"# local misassemblies","quality":"Less is better","values":[0,0,0,0,0,0],"isMain":false},{"metricName":"# scaffold gap ext. mis.","quality":"Less is better","values":[0,0,0,0,0,0],"isMain":false},{"metricName":"# scaffold gap loc. mis.","quality":"Less is better","values":[0,0,0,0,0,0],"isMain":false},{"metricName":"# unaligned mis. contigs","quality":"Less is better","values":[1,1,0,1,1,0],"isMain":false}]],["Unaligned",[{"metricName":"# fully unaligned contigs","quality":"Less is better","values":[0,0,0,0,0,0],"isMain":false},{"metricName":"Fully unaligned length","quality":"Less is better","values":[0,0,0,0,0,0],"isMain":false},{"metricName":"# partially unaligned contigs","quality":"Less is better","values":[1,1,0,3,1,0],"isMain":false},{"metricName":"Partially unaligned length","quality":"Less is better","values":[15117,41093,0,46588,44631,0],"isMain":false}]],["Mismatches",[{"metricName":"# mismatches","quality":"Less is better","values":[114,520,47,578,597,2],"isMain":false},{"metricName":"# indels","quality":"Less is better","values":[0,15,4,16,32,0],"isMain":false},{"metricName":"Indels length","quality":"Less is better","values":[0,22,4,24,42,0],"isMain":false},{"metricName":"# mismatches per 100 kbp","quality":"Less is better","values":["2734.47","3512.80","3158.60","3481.93","3494.70","1360.54"],"isMain":true},{"metricName":"# indels per 100 kbp","quality":"Less is better","values":["0.00","101.33","268.82","96.39","187.32","0.00"],"isMain":true},{"metricName":" # indels (<= 5 bp)","quality":"Less is better","values":[0,15,4,16,32,0],"isMain":false},{"metricName":" # indels (> 5 bp)","quality":"Less is better","values":[0,0,0,0,0,0],"isMain":false},{"metricName":"# N's","quality":"Less is better","values":[0,0,0,0,0,0],"isMain":false},{"metricName":"# N's per 100 kbp","quality":"Less is better","values":["0.00","0.00","0.00","0.00","0.00","0.00"],"isMain":true}]],["Statistics without reference",[{"metricName":"# contigs","quality":"Equal","values":[1,1,2,3,3,1],"isMain":true},{"metricName":"# contigs (>= 1000 bp)","quality":"Equal","values":[1,1,1,2,1,0],"isMain":false},{"metricName":"# contigs (>= 5000 bp)","quality":"Equal","values":[1,1,0,1,1,0],"isMain":false},{"metricName":"# contigs (>= 10000 bp)","quality":"Equal","values":[1,1,0,1,1,0],"isMain":false},{"metricName":"# contigs (>= 25000 bp)","quality":"Equal","values":[0,1,0,1,1,0],"isMain":false},{"metricName":"# contigs (>= 50000 bp)","quality":"Equal","values":[0,1,0,1,1,0],"isMain":false},{"metricName":"Largest contig","quality":"More is better","values":[19286,55894,1400,60942,60942,535],"isMain":true},{"metricName":"Total length","quality":"More is better","values":[19286,55894,1979,63184,62192,535],"isMain":true},{"metricName":"Total length (>= 1000 bp)","quality":"More is better","values":[19286,55894,1400,62439,60942,0],"isMain":true},{"metricName":"Total length (>= 5000 bp)","quality":"More is better","values":[19286,55894,0,60942,60942,0],"isMain":false},{"metricName":"Total length (>= 10000 bp)","quality":"More is better","values":[19286,55894,0,60942,60942,0],"isMain":true},{"metricName":"Total length (>= 25000 bp)","quality":"More is better","values":[0,55894,0,60942,60942,0],"isMain":false},{"metricName":"Total length (>= 50000 bp)","quality":"More is better","values":[0,55894,0,60942,60942,0],"isMain":true},{"metricName":"N50","quality":"More is better","values":[19286,55894,1400,60942,60942,535],"isMain":false},{"metricName":"N75","quality":"More is better","values":[19286,55894,579,60942,60942,535],"isMain":false},{"metricName":"L50","quality":"Less is better","values":[1,1,1,1,1,1],"isMain":false},{"metricName":"L75","quality":"Less is better","values":[1,1,2,1,1,1],"isMain":false},{"metricName":"GC (%)","quality":"Equal","values":["62.77","65.62","65.34","64.76","65.55","68.79"],"isMain":false}]],["K-mer-based statistics",[]],["Predicted genes",[]],["Similarity statistics",[]],["Reference statistics",[{"metricName":"Reference length","quality":"Equal","values":[2943296,2943296,2943296,2943296,2943296,2943296],"isMain":false},{"metricName":"Reference fragments","quality":"Equal","values":[17,17,17,17,17,17],"isMain":false},{"metricName":"Reference GC (%)","quality":"Equal","values":["64.13","64.13","64.13","64.13","64.13","64.13"],"isMain":false}]]],"subreferences":[],"subreports":[],"minContig":500},{"date":"10 February 2020, Monday, 20:34:48","assembliesNames":["megahit-103-df","megahit-113-df","megahit-113-ml","megahit-113-ms","megahit-129-df","metaSPAdes"],"referenceName":"GCF\_000403435.2\_Osci\_bact\_1\_3\_V1","order":[0,1,2,3,4,5],"report":[["Genome statistics",[{"metricName":"Genome fraction (%)","quality":"More is better","values":["38.572","44.348","42.199","43.090","47.433","30.374"],"isMain":true},{"metricName":"Duplication ratio","quality":"Less is better","values":["1.034","1.034","1.041","1.045","1.035","1.042"],"isMain":true},{"metricName":"Largest alignment","quality":"More is better","values":[5628,5702,2604,2919,4222,2403],"isMain":true},{"metricName":"Total aligned length","quality":"More is better","values":[1738636,1997107,1920048,1961424,2142284,1370157],"isMain":true},{"metricName":"NG50","quality":"More is better","values":[null,null,null,null,512,null],"isMain":false},{"metricName":"NA50","quality":"More is better","values":[722,783,699,715,778,671],"isMain":false},{"metricName":"NA75","quality":"More is better","values":[581,605,573,580,595,562],"isMain":false},{"metricName":"NGA50","quality":"More is better","values":[null,null,null,null,null,null],"isMain":true},{"metricName":"LG50","quality":"Less is better","values":[null,null,null,null,2587,null],"isMain":false},{"metricName":"LA50","quality":"Less is better","values":[936,954,1040,1044,1033,810],"isMain":false},{"metricName":"LA75","quality":"Less is better","values":[1636,1712,1820,1828,1868,1384],"isMain":false}]],["Reads mapping",[]],["Misassemblies",[{"metricName":"# misassemblies","quality":"Less is better","values":[24,25,15,21,26,7],"isMain":true},{"metricName":" # relocations","quality":"Less is better","values":[3,2,0,3,2,1],"isMain":false},{"metricName":" # translocations","quality":"Less is better","values":[21,22,14,16,21,6],"isMain":false},{"metricName":" # inversions","quality":"Less is better","values":[0,1,1,2,3,0],"isMain":false},{"metricName":" # interspecies translocations","quality":"Less is better","values":[103,120,41,55,107,59],"isMain":false},{"metricName":"# misassembled contigs","quality":"Less is better","values":[24,25,15,21,26,6],"isMain":false},{"metricName":"Misassembled contigs length","quality":"Less is better","values":[22211,29489,12576,24313,31576,6622],"isMain":true},{"metricName":"# possibly misassembled contigs","quality":"Less is better","values":[1,0,0,0,1,0],"isMain":false},{"metricName":" # possible misassemblies","quality":"Less is better","values":[1,0,0,0,1,0],"isMain":false},{"metricName":"# local misassemblies","quality":"Less is better","values":[4,4,4,2,2,2],"isMain":false},{"metricName":"# scaffold gap ext. mis.","quality":"Less is better","values":[0,0,0,0,0,0],"isMain":false},{"metricName":"# scaffold gap loc. mis.","quality":"Less is better","values":[0,0,0,0,0,0],"isMain":false},{"metricName":"# unaligned mis. contigs","quality":"Less is better","values":[1,2,0,0,0,0],"isMain":false}]],["Unaligned",[{"metricName":"# fully unaligned contigs","quality":"Less is better","values":[2,4,4,1,3,2],"isMain":false},{"metricName":"Fully unaligned length","quality":"Less is better","values":[1315,3033,4997,948,3451,1787],"isMain":false},{"metricName":"# partially unaligned contigs","quality":"Less is better","values":[24,41,11,7,41,9],"isMain":false},{"metricName":"Partially unaligned length","quality":"Less is better","values":[35164,40930,11682,6663,89998,7548],"isMain":false}]],["Mismatches",[{"metricName":"# mismatches","quality":"Less is better","values":[23597,26192,35369,37565,28850,19508],"isMain":false},{"metricName":"# indels","quality":"Less is better","values":[66,60,78,56,78,49],"isMain":false},{"metricName":"Indels length","quality":"Less is better","values":[135,96,122,84,127,87],"isMain":false},{"metricName":"# mismatches per 100 kbp","quality":"Less is better","values":["1380.88","1333.11","1891.90","1967.83","1372.91","1449.73"],"isMain":true},{"metricName":"# indels per 100 kbp","quality":"Less is better","values":["3.86","3.05","4.17","2.93","3.71","3.64"],"isMain":true},{"metricName":" # indels (<= 5 bp)","quality":"Less is better","values":[64,57,76,54,75,47],"isMain":false},{"metricName":" # indels (> 5 bp)","quality":"Less is better","values":[2,3,2,2,3,2],"isMain":false},{"metricName":"# N's","quality":"Less is better","values":[0,0,0,0,0,0],"isMain":false},{"metricName":"# N's per 100 kbp","quality":"Less is better","values":["0.00","0.00","0.00","0.00","0.00","0.00"],"isMain":true}]],["Statistics without reference",[{"metricName":"# contigs","quality":"Equal","values":[2368,2536,2673,2692,2693,1995],"isMain":true},{"metricName":"# contigs (>= 1000 bp)","quality":"Equal","values":[304,519,333,345,553,162],"isMain":false},{"metricName":"# contigs (>= 5000 bp)","quality":"Equal","values":[3,1,0,0,4,0],"isMain":false},{"metricName":"# contigs (>= 10000 bp)","quality":"Equal","values":[0,0,0,0,2,0],"isMain":false},{"metricName":"# contigs (>= 25000 bp)","quality":"Equal","values":[0,0,0,0,1,0],"isMain":false},{"metricName":"# contigs (>= 50000 bp)","quality":"Equal","values":[0,0,0,0,0,0],"isMain":false},{"metricName":"Largest contig","quality":"More is better","values":[8104,5702,3426,2924,25654,2408],"isMain":true},{"metricName":"Total length","quality":"More is better","values":[1803703,2075106,1963628,2002819,2269187,1411403],"isMain":true},{"metricName":"Total length (>= 1000 bp)","quality":"More is better","values":[408450,695191,420073,437354,801342,203557],"isMain":true},{"metricName":"Total length (>= 5000 bp)","quality":"More is better","values":[19025,5702,0,0,53594,0],"isMain":false},{"metricName":"Total length (>= 10000 bp)","quality":"More is better","values":[0,0,0,0,38319,0],"isMain":true},{"metricName":"Total length (>= 25000 bp)","quality":"More is better","values":[0,0,0,0,25654,0],"isMain":false},{"metricName":"Total length (>= 50000 bp)","quality":"More is better","values":[0,0,0,0,0,0],"isMain":true},{"metricName":"N50","quality":"More is better","values":[753,829,723,732,845,692],"isMain":false},{"metricName":"N75","quality":"More is better","values":[611,638,589,598,644,584],"isMain":false},{"metricName":"L50","quality":"Less is better","values":[881,899,1009,1015,918,785],"isMain":false},{"metricName":"L75","quality":"Less is better","values":[1550,1617,1766,1777,1690,1340],"isMain":false},{"metricName":"GC (%)","quality":"Equal","values":["57.98","57.99","58.15","58.21","57.87","58.00"],"isMain":false}]],["K-mer-based statistics",[]],["Predicted genes",[]],["Similarity statistics",[]],["Reference statistics",[{"metricName":"Reference length","quality":"Equal","values":[4430205,4430205,4430205,4430205,4430205,4430205],"isMain":false},{"metricName":"Reference fragments","quality":"Equal","values":[43,43,43,43,43,43],"isMain":false},{"metricName":"Reference GC (%)","quality":"Equal","values":["58.36","58.36","58.36","58.36","58.36","58.36"],"isMain":false}]]],"subreferences":[],"subreports":[],"minContig":500},{"date":"10 February 2020, Monday, 20:34:25","assembliesNames":["megahit-103-df","megahit-113-df","megahit-113-ml","megahit-113-ms","megahit-129-df","metaSPAdes"],"referenceName":"GCF\_000403455.2\_Dore\_bact\_5\_2\_V1","order":[0,1,2,3,4,5],"report":[["Genome statistics",[{"metricName":"Genome fraction (%)","quality":"More is better","values":["0.125","0.039","0.145","0.215","0.141","0.065"],"isMain":true},{"metricName":"Duplication ratio","quality":"Less is better","values":["1.037","1.000","1.003","1.154","1.078","1.007"],"isMain":true},{"metricName":"Largest alignment","quality":"More is better","values":[1021,900,1561,1135,1359,896],"isMain":true},{"metricName":"Total aligned length","quality":"More is better","values":[7346,2273,8538,12652,8292,3816],"isMain":true},{"metricName":"NA50","quality":"More is better","values":[451,null,630,502,null,703],"isMain":false},{"metricName":"NA75","quality":"More is better","values":[null,null,null,null,null,585],"isMain":false},{"metricName":"NGA50","quality":"More is better","values":[null,null,null,null,null,null],"isMain":true},{"metricName":"LA50","quality":"Less is better","values":[11,null,7,16,null,3],"isMain":false},{"metricName":"LA75","quality":"Less is better","values":[null,null,null,null,null,5],"isMain":false}]],["Reads mapping",[]],["Misassemblies",[{"metricName":"# misassemblies","quality":"Less is better","values":[0,0,0,1,0,0],"isMain":true},{"metricName":" # relocations","quality":"Less is better","values":[0,0,0,0,0,0],"isMain":false},{"metricName":" # translocations","quality":"Less is better","values":[0,0,0,1,0,0],"isMain":false},{"metricName":" # inversions","quality":"Less is better","values":[0,0,0,0,0,0],"isMain":false},{"metricName":" # interspecies translocations","quality":"Less is better","values":[4,1,2,9,7,1],"isMain":false},{"metricName":"# misassembled contigs","quality":"Less is better","values":[0,0,0,1,0,0],"isMain":false},{"metricName":"Misassembled contigs length","quality":"Less is better","values":[0,0,0,955,0,0],"isMain":true},{"metricName":"# possibly misassembled contigs","quality":"Less is better","values":[0,0,0,0,0,0],"isMain":false},{"metricName":" # possible misassemblies","quality":"Less is better","values":[0,0,0,0,0,0],"isMain":false},{"metricName":"# local misassemblies","quality":"Less is better","values":[0,0,0,0,0,0],"isMain":false},{"metricName":"# scaffold gap ext. mis.","quality":"Less is better","values":[0,0,0,0,0,0],"isMain":false},{"metricName":"# scaffold gap loc. mis.","quality":"Less is better","values":[0,0,0,0,0,0],"isMain":false},{"metricName":"# unaligned mis. contigs","quality":"Less is better","values":[0,0,0,0,1,0],"isMain":false}]],["Unaligned",[{"metricName":"# fully unaligned contigs","quality":"Less is better","values":[0,0,0,1,0,0],"isMain":false},{"metricName":"Fully unaligned length","quality":"Less is better","values":[0,0,0,755,0,0],"isMain":false},{"metricName":"# partially unaligned contigs","quality":"Less is better","values":[3,1,2,3,5,1],"isMain":false},{"metricName":"Partially unaligned length","quality":"Less is better","values":[5985,2291,4175,5786,7844,585],"isMain":false}]],["Mismatches",[{"metricName":"# mismatches","quality":"Less is better","values":[115,55,100,269,158,95],"isMain":false},{"metricName":"# indels","quality":"Less is better","values":[0,0,2,1,3,0],"isMain":false},{"metricName":"Indels length","quality":"Less is better","values":[0,0,2,2,3,0],"isMain":false},{"metricName":"# mismatches per 100 kbp","quality":"Less is better","values":["1565.48","2419.71","1170.96","2125.81","1905.22","2489.52"],"isMain":true},{"metricName":"# indels per 100 kbp","quality":"Less is better","values":["0.00","0.00","23.42","7.90","36.18","0.00"],"isMain":true},{"metricName":" # indels (<= 5 bp)","quality":"Less is better","values":[0,0,2,1,3,0],"isMain":false},{"metricName":" # indels (> 5 bp)","quality":"Less is better","values":[0,0,0,0,0,0],"isMain":false},{"metricName":"# N's","quality":"Less is better","values":[0,0,0,0,0,0],"isMain":false},{"metricName":"# N's per 100 kbp","quality":"Less is better","values":["0.00","0.00","0.00","0.00","0.00","0.00"],"isMain":true}]],["Statistics without reference",[{"metricName":"# contigs","quality":"Equal","values":[12,4,11,23,14,6],"isMain":true},{"metricName":"# contigs (>= 1000 bp)","quality":"Equal","values":[4,1,4,4,6,0],"isMain":false},{"metricName":"# contigs (>= 5000 bp)","quality":"Equal","values":[0,0,0,0,0,0],"isMain":false},{"metricName":"# contigs (>= 10000 bp)","quality":"Equal","values":[0,0,0,0,0,0],"isMain":false},{"metricName":"# contigs (>= 25000 bp)","quality":"Equal","values":[0,0,0,0,0,0],"isMain":false},{"metricName":"# contigs (>= 50000 bp)","quality":"Equal","values":[0,0,0,0,0,0],"isMain":false},{"metricName":"Largest contig","quality":"More is better","values":[2906,2427,2455,4289,4010,906],"isMain":true},{"metricName":"Total length","quality":"More is better","values":[13606,4564,12738,21146,16786,4428],"isMain":true},{"metricName":"Total length (>= 1000 bp)","quality":"More is better","values":[8357,2427,7691,8585,12070,0],"isMain":true},{"metricName":"Total length (>= 5000 bp)","quality":"More is better","values":[0,0,0,0,0,0],"isMain":false},{"metricName":"Total length (>= 10000 bp)","quality":"More is better","values":[0,0,0,0,0,0],"isMain":true},{"metricName":"Total length (>= 25000 bp)","quality":"More is better","values":[0,0,0,0,0,0],"isMain":false},{"metricName":"Total length (>= 50000 bp)","quality":"More is better","values":[0,0,0,0,0,0],"isMain":true},{"metricName":"N50","quality":"More is better","values":[2039,2427,1569,782,1559,759],"isMain":false},{"metricName":"N75","quality":"More is better","values":[709,694,735,634,703,697],"isMain":false},{"metricName":"L50","quality":"Less is better","values":[3,1,3,7,4,3],"isMain":false},{"metricName":"L75","quality":"Less is better","values":[7,3,7,14,7,5],"isMain":false},{"metricName":"GC (%)","quality":"Equal","values":["40.78","38.34","44.26","46.20","46.94","43.41"],"isMain":false}]],["K-mer-based statistics",[]],["Predicted genes",[]],["Similarity statistics",[]],["Reference statistics",[{"metricName":"Reference length","quality":"Equal","values":[5887770,5887770,5887770,5887770,5887770,5887770],"isMain":false},{"metricName":"Reference fragments","quality":"Equal","values":[241,241,241,241,241,241],"isMain":false},{"metricName":"Reference GC (%)","quality":"Equal","values":["46.03","46.03","46.03","46.03","46.03","46.03"],"isMain":false}]]],"subreferences":[],"subreports":[],"minContig":500},{"date":"10 February 2020, Monday, 20:34:20","assembliesNames":["megahit-103-df","megahit-113-df","megahit-113-ml","megahit-113-ms","megahit-129-df","metaSPAdes"],"referenceName":"GCF\_000404205.1\_ASM40420v1","order":[0,1,2,3,4,5],"report":[["Genome statistics",[{"metricName":"Genome fraction (%)","quality":"More is better","values":["80.767","84.743","77.642","78.877","87.054","73.661"],"isMain":true},{"metricName":"Duplication ratio","quality":"Less is better","values":["1.052","1.039","1.080","1.081","1.044","1.056"],"isMain":true},{"metricName":"Largest alignment","quality":"More is better","values":[6570,8546,3985,3679,6490,3253],"isMain":true},{"metricName":"Total aligned length","quality":"More is better","values":[1478450,1533577,1458396,1480326,1583798,1347134],"isMain":true},{"metricName":"NG50","quality":"More is better","values":[1021,1319,848,875,1402,813],"isMain":false},{"metricName":"NG75","quality":"More is better","values":[650,754,589,604,831,531],"isMain":false},{"metricName":"NA50","quality":"More is better","values":[1151,1445,948,944,1509,957],"isMain":false},{"metricName":"NA75","quality":"More is better","values":[782,916,687,691,944,687],"isMain":false},{"metricName":"NGA50","quality":"More is better","values":[1005,1289,831,857,1375,798],"isMain":true},{"metricName":"NGA75","quality":"More is better","values":[634,735,571,579,806,509],"isMain":false},{"metricName":"LG50","quality":"Less is better","values":[552,409,691,670,395,703],"isMain":false},{"metricName":"LG75","quality":"Less is better","values":[1095,849,1314,1274,797,1370],"isMain":false},{"metricName":"LA50","quality":"Less is better","values":[435,341,540,550,347,492],"isMain":false},{"metricName":"LA75","quality":"Less is better","values":[828,679,1000,1018,680,914],"isMain":false},{"metricName":"LGA50","quality":"Less is better","values":[557,416,699,688,401,714],"isMain":true},{"metricName":"LGA75","quality":"Less is better","values":[1108,867,1335,1311,812,1403],"isMain":false}]],["Reads mapping",[]],["Misassemblies",[{"metricName":"# misassemblies","quality":"Less is better","values":[3,6,6,8,6,3],"isMain":true},{"metricName":" # relocations","quality":"Less is better","values":[3,4,4,6,5,3],"isMain":false},{"metricName":" # translocations","quality":"Less is better","values":[0,0,0,0,0,0],"isMain":false},{"metricName":" # inversions","quality":"Less is better","values":[0,2,2,2,1,0],"isMain":false},{"metricName":" # interspecies translocations","quality":"Less is better","values":[23,23,5,19,17,21],"isMain":false},{"metricName":"# misassembled contigs","quality":"Less is better","values":[3,6,6,8,6,3],"isMain":false},{"metricName":"Misassembled contigs length","quality":"Less is better","values":[5941,12826,8323,12405,13042,3336],"isMain":true},{"metricName":"# possibly misassembled contigs","quality":"Less is better","values":[0,0,0,0,0,0],"isMain":false},{"metricName":" # possible misassemblies","quality":"Less is better","values":[0,0,0,0,0,0],"isMain":false},{"metricName":"# local misassemblies","quality":"Less is better","values":[2,2,9,5,4,2],"isMain":false},{"metricName":"# scaffold gap ext. mis.","quality":"Less is better","values":[0,0,0,0,0,0],"isMain":false},{"metricName":"# scaffold gap loc. mis.","quality":"Less is better","values":[0,0,0,0,0,0],"isMain":false},{"metricName":"# unaligned mis. contigs","quality":"Less is better","values":[0,0,1,0,0,0],"isMain":false}]],["Unaligned",[{"metricName":"# fully unaligned contigs","quality":"Less is better","values":[0,0,0,0,0,0],"isMain":false},{"metricName":"Fully unaligned length","quality":"Less is better","values":[0,0,0,0,0,0],"isMain":false},{"metricName":"# partially unaligned contigs","quality":"Less is better","values":[2,7,1,5,6,4],"isMain":false},{"metricName":"Partially unaligned length","quality":"Less is better","values":[1239,5335,1480,10569,5655,2545],"isMain":false}]],["Mismatches",[{"metricName":"# mismatches","quality":"Less is better","values":[14676,13595,24095,24807,14399,14291],"isMain":false},{"metricName":"# indels","quality":"Less is better","values":[18,30,30,28,14,23],"isMain":false},{"metricName":"Indels length","quality":"Less is better","values":[39,91,148,113,65,26],"isMain":false},{"metricName":"# mismatches per 100 kbp","quality":"Less is better","values":["1036.61","915.20","1770.40","1794.16","943.59","1106.79"],"isMain":true},{"metricName":"# indels per 100 kbp","quality":"Less is better","values":["1.27","2.02","2.20","2.03","0.92","1.78"],"isMain":true},{"metricName":" # indels (<= 5 bp)","quality":"Less is better","values":[16,29,28,27,11,23],"isMain":false},{"metricName":" # indels (> 5 bp)","quality":"Less is better","values":[2,1,2,1,3,0],"isMain":false},{"metricName":"# N's","quality":"Less is better","values":[0,0,0,0,0,0],"isMain":false},{"metricName":"# N's per 100 kbp","quality":"Less is better","values":["0.00","0.00","0.00","0.00","0.00","0.00"],"isMain":true}]],["Statistics without reference",[{"metricName":"# contigs","quality":"Equal","values":[1403,1228,1602,1620,1238,1470],"isMain":true},{"metricName":"# contigs (>= 1000 bp)","quality":"Equal","values":[578,615,481,487,648,457],"isMain":false},{"metricName":"# contigs (>= 5000 bp)","quality":"Equal","values":[1,7,0,1,8,0],"isMain":false},{"metricName":"# contigs (>= 10000 bp)","quality":"Equal","values":[0,0,0,0,0,0],"isMain":false},{"metricName":"# contigs (>= 25000 bp)","quality":"Equal","values":[0,0,0,0,0,0],"isMain":false},{"metricName":"# contigs (>= 50000 bp)","quality":"Equal","values":[0,0,0,0,0,0],"isMain":false},{"metricName":"Largest contig","quality":"More is better","values":[6571,8560,3985,6435,6490,3259],"isMain":true},{"metricName":"Total length","quality":"More is better","values":[1489934,1548731,1471305,1504923,1599118,1366182],"isMain":true},{"metricName":"Total length (>= 1000 bp)","quality":"More is better","values":[902786,1112114,684049,706446,1179228,655975],"isMain":true},{"metricName":"Total length (>= 5000 bp)","quality":"More is better","values":[6571,42940,0,6435,45231,0],"isMain":false},{"metricName":"Total length (>= 10000 bp)","quality":"More is better","values":[0,0,0,0,0,0],"isMain":true},{"metricName":"Total length (>= 25000 bp)","quality":"More is better","values":[0,0,0,0,0,0],"isMain":false},{"metricName":"Total length (>= 50000 bp)","quality":"More is better","values":[0,0,0,0,0,0],"isMain":true},{"metricName":"N50","quality":"More is better","values":[1166,1475,958,960,1529,971],"isMain":false},{"metricName":"N75","quality":"More is better","values":[793,935,701,711,974,709],"isMain":false},{"metricName":"L50","quality":"Less is better","values":[432,336,534,534,343,485],"isMain":false},{"metricName":"L75","quality":"Less is better","values":[819,667,986,990,669,898],"isMain":false},{"metricName":"GC (%)","quality":"Equal","values":["36.46","36.47","36.68","36.67","36.48","36.53"],"isMain":false}]],["K-mer-based statistics",[]],["Predicted genes",[]],["Similarity statistics",[]],["Reference statistics",[{"metricName":"Reference length","quality":"Equal","values":[1752910,1752910,1752910,1752910,1752910,1752910],"isMain":false},{"metricName":"Reference fragments","quality":"Equal","values":[1,1,1,1,1,1],"isMain":false},{"metricName":"Reference GC (%)","quality":"Equal","values":["36.36","36.36","36.36","36.36","36.36","36.36"],"isMain":false}]]],"subreferences":[],"subreports":[],"minContig":500},{"date":"10 February 2020, Monday, 20:34:40","assembliesNames":["megahit-103-df","megahit-113-df","megahit-113-ml","megahit-113-ms","megahit-129-df","metaSPAdes"],"referenceName":"GCF\_000410615.1\_ASM41061v1","order":[0,1,2,3,4,5],"report":[["Genome statistics",[{"metricName":"Genome fraction (%)","quality":"More is better","values":["15.947","19.768","18.947","20.763","22.094","10.565"],"isMain":true},{"metricName":"Duplication ratio","quality":"Less is better","values":["1.025","1.025","1.021","1.025","1.023","1.030"],"isMain":true},{"metricName":"Largest alignment","quality":"More is better","values":[2049,2049,1417,1677,2090,1297],"isMain":true},{"metricName":"Total aligned length","quality":"More is better","values":[780152,966248,931320,1017602,1081032,515941],"isMain":true},{"metricName":"NA50","quality":"More is better","values":[623,641,627,636,651,596],"isMain":false},{"metricName":"NA75","quality":"More is better","values":[544,550,547,550,553,532],"isMain":false},{"metricName":"NGA50","quality":"More is better","values":[null,null,null,null,null,null],"isMain":true},{"metricName":"LA50","quality":"Less is better","values":[514,612,606,661,679,371],"isMain":false},{"metricName":"LA75","quality":"Less is better","values":[859,1041,1009,1103,1151,614],"isMain":false}]],["Reads mapping",[]],["Misassemblies",[{"metricName":"# misassemblies","quality":"Less is better","values":[7,17,2,6,13,3],"isMain":true},{"metricName":" # relocations","quality":"Less is better","values":[0,1,0,1,0,0],"isMain":false},{"metricName":" # translocations","quality":"Less is better","values":[6,16,0,4,13,3],"isMain":false},{"metricName":" # inversions","quality":"Less is better","values":[1,0,2,1,0,0],"isMain":false},{"metricName":" # interspecies translocations","quality":"Less is better","values":[54,58,2,32,52,35],"isMain":false},{"metricName":"# misassembled contigs","quality":"Less is better","values":[7,17,2,6,13,3],"isMain":false},{"metricName":"Misassembled contigs length","quality":"Less is better","values":[5274,12690,1170,4550,10686,2020],"isMain":true},{"metricName":"# possibly misassembled contigs","quality":"Less is better","values":[0,0,0,0,0,0],"isMain":false},{"metricName":" # possible misassemblies","quality":"Less is better","values":[0,0,0,0,0,0],"isMain":false},{"metricName":"# local misassemblies","quality":"Less is better","values":[0,0,2,1,2,0],"isMain":false},{"metricName":"# scaffold gap ext. mis.","quality":"Less is better","values":[0,0,0,0,0,0],"isMain":false},{"metricName":"# scaffold gap loc. mis.","quality":"Less is better","values":[0,0,0,0,0,0],"isMain":false},{"metricName":"# unaligned mis. contigs","quality":"Less is better","values":[0,0,0,0,0,0],"isMain":false}]],["Unaligned",[{"metricName":"# fully unaligned contigs","quality":"Less is better","values":[0,1,0,1,1,0],"isMain":false},{"metricName":"Fully unaligned length","quality":"Less is better","values":[0,749,0,675,801,0],"isMain":false},{"metricName":"# partially unaligned contigs","quality":"Less is better","values":[8,14,0,7,13,4],"isMain":false},{"metricName":"Partially unaligned length","quality":"Less is better","values":[6056,27278,0,6482,28448,14205],"isMain":false}]],["Mismatches",[{"metricName":"# mismatches","quality":"Less is better","values":[11499,14080,17858,20347,15787,8283],"isMain":false},{"metricName":"# indels","quality":"Less is better","values":[7,17,13,28,19,11],"isMain":false},{"metricName":"Indels length","quality":"Less is better","values":[7,20,13,99,42,12],"isMain":false},{"metricName":"# mismatches per 100 kbp","quality":"Less is better","values":["1482.41","1464.35","1937.73","2014.71","1468.97","1611.79"],"isMain":true},{"metricName":"# indels per 100 kbp","quality":"Less is better","values":["0.90","1.77","1.41","2.77","1.77","2.14"],"isMain":true},{"metricName":" # indels (<= 5 bp)","quality":"Less is better","values":[7,17,13,27,18,11],"isMain":false},{"metricName":" # indels (> 5 bp)","quality":"Less is better","values":[0,0,0,1,1,0],"isMain":false},{"metricName":"# N's","quality":"Less is better","values":[0,0,0,0,0,0],"isMain":false},{"metricName":"# N's per 100 kbp","quality":"Less is better","values":["0.00","0.00","0.00","0.00","0.00","0.00"],"isMain":true}]],["Statistics without reference",[{"metricName":"# contigs","quality":"Equal","values":[1218,1450,1445,1568,1611,836],"isMain":true},{"metricName":"# contigs (>= 1000 bp)","quality":"Equal","values":[54,102,57,65,110,19],"isMain":false},{"metricName":"# contigs (>= 5000 bp)","quality":"Equal","values":[0,1,0,0,1,1],"isMain":false},{"metricName":"# contigs (>= 10000 bp)","quality":"Equal","values":[0,1,0,0,1,1],"isMain":false},{"metricName":"# contigs (>= 25000 bp)","quality":"Equal","values":[0,0,0,0,0,0],"isMain":false},{"metricName":"# contigs (>= 50000 bp)","quality":"Equal","values":[0,0,0,0,0,0],"isMain":false},{"metricName":"Largest contig","quality":"More is better","values":[2049,18689,1426,2807,18878,12626],"isMain":true},{"metricName":"Total length","quality":"More is better","values":[801454,1013434,941046,1041854,1128573,543607],"isMain":true},{"metricName":"Total length (>= 1000 bp)","quality":"More is better","values":[63819,137730,64196,75168,149257,32384],"isMain":true},{"metricName":"Total length (>= 5000 bp)","quality":"More is better","values":[0,18689,0,0,18878,12626],"isMain":false},{"metricName":"Total length (>= 10000 bp)","quality":"More is better","values":[0,18689,0,0,18878,12626],"isMain":true},{"metricName":"Total length (>= 25000 bp)","quality":"More is better","values":[0,0,0,0,0,0],"isMain":false},{"metricName":"Total length (>= 50000 bp)","quality":"More is better","values":[0,0,0,0,0,0],"isMain":true},{"metricName":"N50","quality":"More is better","values":[638,675,634,647,678,622],"isMain":false},{"metricName":"N75","quality":"More is better","values":[557,573,557,565,576,551],"isMain":false},{"metricName":"L50","quality":"Less is better","values":[501,566,600,646,631,344],"isMain":false},{"metricName":"L75","quality":"Less is better","values":[838,977,998,1078,1084,577],"isMain":false},{"metricName":"GC (%)","quality":"Equal","values":["64.20","64.30","64.38","64.25","64.34","64.01"],"isMain":false}]],["K-mer-based statistics",[]],["Predicted genes",[]],["Similarity statistics",[]],["Reference statistics",[{"metricName":"Reference length","quality":"Equal","values":[4864130,4864130,4864130,4864130,4864130,4864130],"isMain":false},{"metricName":"Reference fragments","quality":"Equal","values":[848,848,848,848,848,848],"isMain":false},{"metricName":"Reference GC (%)","quality":"Equal","values":["65.08","65.08","65.08","65.08","65.08","65.08"],"isMain":false}]]],"subreferences":[],"subreports":[],"minContig":500},{"date":"10 February 2020, Monday, 20:34:13","assembliesNames":["megahit-103-df","megahit-113-df","megahit-113-ml","megahit-113-ms","megahit-129-df","metaSPAdes"],"referenceName":"GCF\_000410995.1\_ASM41099v1","order":[0,1,2,3,4,5],"report":[["Genome statistics",[{"metricName":"Genome fraction (%)","quality":"More is better","values":["1.726","2.219","2.149","2.226","2.378","1.446"],"isMain":true},{"metricName":"Duplication ratio","quality":"Less is better","values":["1.071","1.059","1.071","1.100","1.031","1.047"],"isMain":true},{"metricName":"Largest alignment","quality":"More is better","values":[1624,2910,1367,1238,1771,1349],"isMain":true},{"metricName":"Total aligned length","quality":"More is better","values":[36314,46778,45233,47555,49795,30428],"isMain":true},{"metricName":"NA50","quality":"More is better","values":[730,768,623,636,842,704],"isMain":false},{"metricName":"NA75","quality":"More is better","values":[582,554,543,550,595,563],"isMain":false},{"metricName":"NGA50","quality":"More is better","values":[null,null,null,null,null,null],"isMain":true},{"metricName":"LA50","quality":"Less is better","values":[21,23,30,31,23,17],"isMain":false},{"metricName":"LA75","quality":"Less is better","values":[36,43,51,53,41,30],"isMain":false}]],["Reads mapping",[]],["Misassemblies",[{"metricName":"# misassemblies","quality":"Less is better","values":[0,6,3,0,4,1],"isMain":true},{"metricName":" # relocations","quality":"Less is better","values":[0,5,3,0,4,1],"isMain":false},{"metricName":" # translocations","quality":"Less is better","values":[0,0,0,0,0,0],"isMain":false},{"metricName":" # inversions","quality":"Less is better","values":[0,1,0,0,0,0],"isMain":false},{"metricName":" # interspecies translocations","quality":"Less is better","values":[6,11,7,10,7,2],"isMain":false},{"metricName":"# misassembled contigs","quality":"Less is better","values":[0,4,2,0,3,1],"isMain":false},{"metricName":"Misassembled contigs length","quality":"Less is better","values":[0,6191,1447,0,4639,1150],"isMain":true},{"metricName":"# possibly misassembled contigs","quality":"Less is better","values":[0,0,0,0,0,0],"isMain":false},{"metricName":" # possible misassemblies","quality":"Less is better","values":[0,0,0,0,0,0],"isMain":false},{"metricName":"# local misassemblies","quality":"Less is better","values":[0,0,0,0,0,0],"isMain":false},{"metricName":"# scaffold gap ext. mis.","quality":"Less is better","values":[0,0,0,0,0,0],"isMain":false},{"metricName":"# scaffold gap loc. mis.","quality":"Less is better","values":[0,0,0,0,0,0],"isMain":false},{"metricName":"# unaligned mis. contigs","quality":"Less is better","values":[0,1,0,0,0,0],"isMain":false}]],["Unaligned",[{"metricName":"# fully unaligned contigs","quality":"Less is better","values":[1,0,0,0,0,0],"isMain":false},{"metricName":"Fully unaligned length","quality":"Less is better","values":[707,0,0,0,0,0],"isMain":false},{"metricName":"# partially unaligned contigs","quality":"Less is better","values":[0,3,1,2,2,0],"isMain":false},{"metricName":"Partially unaligned length","quality":"Less is better","values":[0,3211,521,1126,1684,0],"isMain":false}]],["Mismatches",[{"metricName":"# mismatches","quality":"Less is better","values":[805,1366,1324,1316,1258,696],"isMain":false},{"metricName":"# indels","quality":"Less is better","values":[11,20,19,8,17,3],"isMain":false},{"metricName":"Indels length","quality":"Less is better","values":[11,36,25,10,31,3],"isMain":false},{"metricName":"# mismatches per 100 kbp","quality":"Less is better","values":["2227.88","2940.29","2943.40","2824.64","2526.71","2299.61"],"isMain":true},{"metricName":"# indels per 100 kbp","quality":"Less is better","values":["30.44","43.05","42.24","17.17","34.14","9.91"],"isMain":true},{"metricName":" # indels (<= 5 bp)","quality":"Less is better","values":[11,19,19,8,16,3],"isMain":false},{"metricName":" # indels (> 5 bp)","quality":"Less is better","values":[0,1,0,0,1,0],"isMain":false},{"metricName":"# N's","quality":"Less is better","values":[0,0,0,0,0,0],"isMain":false},{"metricName":"# N's per 100 kbp","quality":"Less is better","values":["0.00","0.00","0.00","0.00","0.00","0.00"],"isMain":true}]],["Statistics without reference",[{"metricName":"# contigs","quality":"Equal","values":[52,55,70,72,61,43],"isMain":true},{"metricName":"# contigs (>= 1000 bp)","quality":"Equal","values":[6,18,7,7,16,7],"isMain":false},{"metricName":"# contigs (>= 5000 bp)","quality":"Equal","values":[0,0,0,0,0,0],"isMain":false},{"metricName":"# contigs (>= 10000 bp)","quality":"Equal","values":[0,0,0,0,0,0],"isMain":false},{"metricName":"# contigs (>= 25000 bp)","quality":"Equal","values":[0,0,0,0,0,0],"isMain":false},{"metricName":"# contigs (>= 50000 bp)","quality":"Equal","values":[0,0,0,0,0,0],"isMain":false},{"metricName":"Largest contig","quality":"More is better","values":[1624,3092,1371,1345,2129,1546],"isMain":true},{"metricName":"Total length","quality":"More is better","values":[39402,52416,48696,52364,53019,31681],"isMain":true},{"metricName":"Total length (>= 1000 bp)","quality":"More is better","values":[7676,27627,7689,8307,21464,8494],"isMain":true},{"metricName":"Total length (>= 5000 bp)","quality":"More is better","values":[0,0,0,0,0,0],"isMain":false},{"metricName":"Total length (>= 10000 bp)","quality":"More is better","values":[0,0,0,0,0,0],"isMain":true},{"metricName":"Total length (>= 25000 bp)","quality":"More is better","values":[0,0,0,0,0,0],"isMain":false},{"metricName":"Total length (>= 50000 bp)","quality":"More is better","values":[0,0,0,0,0,0],"isMain":true},{"metricName":"N50","quality":"More is better","values":[745,1014,686,741,919,704],"isMain":false},{"metricName":"N75","quality":"More is better","values":[640,697,580,601,701,586],"isMain":false},{"metricName":"L50","quality":"Less is better","values":[21,17,29,29,22,17],"isMain":false},{"metricName":"L75","quality":"Less is better","values":[35,33,48,49,38,29],"isMain":false},{"metricName":"GC (%)","quality":"Equal","values":["37.97","39.27","38.86","39.85","39.36","37.60"],"isMain":false}]],["K-mer-based statistics",[]],["Predicted genes",[]],["Similarity statistics",[]],["Reference statistics",[{"metricName":"Reference length","quality":"Equal","values":[2093275,2093275,2093275,2093275,2093275,2093275],"isMain":false},{"metricName":"Reference fragments","quality":"Equal","values":[7,7,7,7,7,7],"isMain":false},{"metricName":"Reference GC (%)","quality":"Equal","values":["38.93","38.93","38.93","38.93","38.93","38.93"],"isMain":false}]]],"subreferences":[],"subreports":[],"minContig":500},{"date":"10 February 2020, Monday, 20:34:30","assembliesNames":["megahit-103-df","megahit-113-df","megahit-113-ml","megahit-113-ms","megahit-129-df","metaSPAdes"],"referenceName":"GCF\_000411355.1\_Copr\_sp\_HPP0048\_V1","order":[0,1,2,3,4,5],"report":[["Genome statistics",[{"metricName":"Genome fraction (%)","quality":"More is better","values":["3.459","4.002","4.833","4.571","4.746","1.798"],"isMain":true},{"metricName":"Duplication ratio","quality":"Less is better","values":["1.055","1.065","1.027","1.039","1.042","1.064"],"isMain":true},{"metricName":"Largest alignment","quality":"More is better","values":[1237,1876,1391,1637,1330,1054],"isMain":true},{"metricName":"Total aligned length","quality":"More is better","values":[96623,111627,135476,127707,132405,50220],"isMain":true},{"metricName":"NA50","quality":"More is better","values":[581,566,580,567,579,550],"isMain":false},{"metricName":"NA75","quality":"More is better","values":[518,505,528,511,514,507],"isMain":false},{"metricName":"NGA50","quality":"More is better","values":[null,null,null,null,null,null],"isMain":true},{"metricName":"LA50","quality":"Less is better","values":[74,92,99,109,105,43],"isMain":false},{"metricName":"LA75","quality":"Less is better","values":[121,153,162,178,174,70],"isMain":false}]],["Reads mapping",[]],["Misassemblies",[{"metricName":"# misassemblies","quality":"Less is better","values":[1,0,0,1,0,0],"isMain":true},{"metricName":" # relocations","quality":"Less is better","values":[0,0,0,1,0,0],"isMain":false},{"metricName":" # translocations","quality":"Less is better","values":[0,0,0,0,0,0],"isMain":false},{"metricName":" # inversions","quality":"Less is better","values":[1,0,0,0,0,0],"isMain":false},{"metricName":" # interspecies translocations","quality":"Less is better","values":[21,27,8,14,22,12],"isMain":false},{"metricName":"# misassembled contigs","quality":"Less is better","values":[1,0,0,1,0,0],"isMain":false},{"metricName":"Misassembled contigs length","quality":"Less is better","values":[691,0,0,511,0,0],"isMain":true},{"metricName":"# possibly misassembled contigs","quality":"Less is better","values":[0,0,0,0,0,0],"isMain":false},{"metricName":" # possible misassemblies","quality":"Less is better","values":[0,0,0,0,0,0],"isMain":false},{"metricName":"# local misassemblies","quality":"Less is better","values":[0,0,0,0,0,0],"isMain":false},{"metricName":"# scaffold gap ext. mis.","quality":"Less is better","values":[0,0,0,0,0,0],"isMain":false},{"metricName":"# scaffold gap loc. mis.","quality":"Less is better","values":[0,0,0,0,0,0],"isMain":false},{"metricName":"# unaligned mis. contigs","quality":"Less is better","values":[0,0,0,1,0,0],"isMain":false}]],["Unaligned",[{"metricName":"# fully unaligned contigs","quality":"Less is better","values":[0,0,0,1,0,0],"isMain":false},{"metricName":"Fully unaligned length","quality":"Less is better","values":[0,0,0,567,0,0],"isMain":false},{"metricName":"# partially unaligned contigs","quality":"Less is better","values":[1,8,2,4,8,4],"isMain":false},{"metricName":"Partially unaligned length","quality":"Less is better","values":[834,9888,1349,16903,11064,3159],"isMain":false}]],["Mismatches",[{"metricName":"# mismatches","quality":"Less is better","values":[1723,2040,2820,2650,2371,965],"isMain":false},{"metricName":"# indels","quality":"Less is better","values":[5,6,8,7,4,4],"isMain":false},{"metricName":"Indels length","quality":"Less is better","values":[5,6,9,7,4,4],"isMain":false},{"metricName":"# mismatches per 100 kbp","quality":"Less is better","values":["1789.96","1832.08","2096.86","2083.71","1795.27","1928.96"],"isMain":true},{"metricName":"# indels per 100 kbp","quality":"Less is better","values":["5.19","5.39","5.95","5.50","3.03","8.00"],"isMain":true},{"metricName":" # indels (<= 5 bp)","quality":"Less is better","values":[5,6,8,7,4,4],"isMain":false},{"metricName":" # indels (> 5 bp)","quality":"Less is better","values":[0,0,0,0,0,0],"isMain":false},{"metricName":"# N's","quality":"Less is better","values":[0,0,0,0,0,0],"isMain":false},{"metricName":"# N's per 100 kbp","quality":"Less is better","values":["0.00","0.00","0.00","0.00","0.00","0.00"],"isMain":true}]],["Statistics without reference",[{"metricName":"# contigs","quality":"Equal","values":[164,193,225,215,221,92],"isMain":true},{"metricName":"# contigs (>= 1000 bp)","quality":"Equal","values":[2,8,7,10,15,3],"isMain":false},{"metricName":"# contigs (>= 5000 bp)","quality":"Equal","values":[0,0,0,1,0,0],"isMain":false},{"metricName":"# contigs (>= 10000 bp)","quality":"Equal","values":[0,0,0,1,0,0],"isMain":false},{"metricName":"# contigs (>= 25000 bp)","quality":"Equal","values":[0,0,0,0,0,0],"isMain":false},{"metricName":"# contigs (>= 50000 bp)","quality":"Equal","values":[0,0,0,0,0,0],"isMain":false},{"metricName":"Largest contig","quality":"More is better","values":[1316,4135,1419,11896,3682,1118],"isMain":true},{"metricName":"Total length","quality":"More is better","values":[102370,128514,139495,149571,148632,56397],"isMain":true},{"metricName":"Total length (>= 1000 bp)","quality":"More is better","values":[2553,14635,7808,25583,22824,3262],"isMain":true},{"metricName":"Total length (>= 5000 bp)","quality":"More is better","values":[0,0,0,11896,0,0],"isMain":false},{"metricName":"Total length (>= 10000 bp)","quality":"More is better","values":[0,0,0,11896,0,0],"isMain":true},{"metricName":"Total length (>= 25000 bp)","quality":"More is better","values":[0,0,0,0,0,0],"isMain":false},{"metricName":"Total length (>= 50000 bp)","quality":"More is better","values":[0,0,0,0,0,0],"isMain":true},{"metricName":"N50","quality":"More is better","values":[608,616,593,613,625,580],"isMain":false},{"metricName":"N75","quality":"More is better","values":[550,547,539,553,554,538],"isMain":false},{"metricName":"L50","quality":"Less is better","values":[71,76,96,80,87,40],"isMain":false},{"metricName":"L75","quality":"Less is better","values":[115,132,158,144,150,65],"isMain":false},{"metricName":"GC (%)","quality":"Equal","values":["42.04","42.32","42.17","42.06","42.47","42.54"],"isMain":false}]],["K-mer-based statistics",[]],["Predicted genes",[]],["Similarity statistics",[]],["Reference statistics",[{"metricName":"Reference length","quality":"Equal","values":[2782472,2782472,2782472,2782472,2782472,2782472],"isMain":false},{"metricName":"Reference fragments","quality":"Equal","values":[11,11,11,11,11,11],"isMain":false},{"metricName":"Reference GC (%)","quality":"Equal","values":["41.86","41.86","41.86","41.86","41.86","41.86"],"isMain":false}]]],"subreferences":[],"subreports":[],"minContig":500},{"date":"10 February 2020, Monday, 20:34:44","assembliesNames":["megahit-103-df","megahit-113-df","megahit-113-ml","megahit-113-ms","megahit-129-df","metaSPAdes"],"referenceName":"GCF\_000413355.1\_Prev\_oral\_HGA0225\_V1","order":[0,1,2,3,4,5],"report":[["Genome statistics",[{"metricName":"Genome fraction (%)","quality":"More is better","values":["0.053","0.163","0.272","0.431","0.144","0.102"],"isMain":true},{"metricName":"Duplication ratio","quality":"Less is better","values":["1.242","1.256","1.004","1.186","1.333","1.590"],"isMain":true},{"metricName":"Largest alignment","quality":"More is better","values":[694,906,986,943,758,564],"isMain":true},{"metricName":"Total aligned length","quality":"More is better","values":[1501,4637,7736,12278,4684,2897],"isMain":true},{"metricName":"NA50","quality":"More is better","values":[198,null,531,467,109,155],"isMain":false},{"metricName":"NA75","quality":"More is better","values":[null,null,516,null,null,null],"isMain":false},{"metricName":"NGA50","quality":"More is better","values":[null,null,null,null,null,null],"isMain":true},{"metricName":"LA50","quality":"Less is better","values":[3,null,6,17,12,7],"isMain":false},{"metricName":"LA75","quality":"Less is better","values":[null,null,10,null,null,null],"isMain":false}]],["Reads mapping",[]],["Misassemblies",[{"metricName":"# misassemblies","quality":"Less is better","values":[0,0,0,0,0,0],"isMain":true},{"metricName":" # relocations","quality":"Less is better","values":[0,0,0,0,0,0],"isMain":false},{"metricName":" # translocations","quality":"Less is better","values":[0,0,0,0,0,0],"isMain":false},{"metricName":" # inversions","quality":"Less is better","values":[0,0,0,0,0,0],"isMain":false},{"metricName":" # interspecies translocations","quality":"Less is better","values":[2,6,0,8,5,5],"isMain":false},{"metricName":"# misassembled contigs","quality":"Less is better","values":[0,0,0,0,0,0],"isMain":false},{"metricName":"Misassembled contigs length","quality":"Less is better","values":[0,0,0,0,0,0],"isMain":true},{"metricName":"# possibly misassembled contigs","quality":"Less is better","values":[0,0,0,0,0,0],"isMain":false},{"metricName":" # possible misassemblies","quality":"Less is better","values":[0,0,0,0,0,0],"isMain":false},{"metricName":"# local misassemblies","quality":"Less is better","values":[0,0,0,0,0,0],"isMain":false},{"metricName":"# scaffold gap ext. mis.","quality":"Less is better","values":[0,0,0,0,0,0],"isMain":false},{"metricName":"# scaffold gap loc. mis.","quality":"Less is better","values":[0,0,0,0,0,0],"isMain":false},{"metricName":"# unaligned mis. contigs","quality":"Less is better","values":[0,0,0,0,1,0],"isMain":false}]],["Unaligned",[{"metricName":"# fully unaligned contigs","quality":"Less is better","values":[0,0,0,0,0,0],"isMain":false},{"metricName":"Fully unaligned length","quality":"Less is better","values":[0,0,0,0,0,0],"isMain":false},{"metricName":"# partially unaligned contigs","quality":"Less is better","values":[1,3,0,3,3,1],"isMain":false},{"metricName":"Partially unaligned length","quality":"Less is better","values":[535,24646,0,4794,3697,565],"isMain":false}]],["Mismatches",[{"metricName":"# mismatches","quality":"Less is better","values":[30,116,183,293,121,62],"isMain":false},{"metricName":"# indels","quality":"Less is better","values":[0,0,0,0,2,0],"isMain":false},{"metricName":"Indels length","quality":"Less is better","values":[0,0,0,0,2,0],"isMain":false},{"metricName":"# mismatches per 100 kbp","quality":"Less is better","values":["1998.67","2501.62","2365.56","2386.38","2953.38","2140.14"],"isMain":true},{"metricName":"# indels per 100 kbp","quality":"Less is better","values":["0.00","0.00","0.00","0.00","48.82","0.00"],"isMain":true},{"metricName":" # indels (<= 5 bp)","quality":"Less is better","values":[0,0,0,0,2,0],"isMain":false},{"metricName":" # indels (> 5 bp)","quality":"Less is better","values":[0,0,0,0,0,0],"isMain":false},{"metricName":"# N's","quality":"Less is better","values":[0,0,0,0,0,0],"isMain":false},{"metricName":"# N's per 100 kbp","quality":"Less is better","values":["0.00","0.00","0.00","0.00","0.00","0.00"],"isMain":true}]],["Statistics without reference",[{"metricName":"# contigs","quality":"Equal","values":[4,10,13,26,11,9],"isMain":true},{"metricName":"# contigs (>= 1000 bp)","quality":"Equal","values":[0,3,0,2,3,0],"isMain":false},{"metricName":"# contigs (>= 5000 bp)","quality":"Equal","values":[0,1,0,0,0,0],"isMain":false},{"metricName":"# contigs (>= 10000 bp)","quality":"Equal","values":[0,1,0,0,0,0],"isMain":false},{"metricName":"# contigs (>= 25000 bp)","quality":"Equal","values":[0,0,0,0,0,0],"isMain":false},{"metricName":"# contigs (>= 50000 bp)","quality":"Equal","values":[0,0,0,0,0,0],"isMain":false},{"metricName":"Largest contig","quality":"More is better","values":[694,20008,986,2742,1535,708],"isMain":true},{"metricName":"Total length","quality":"More is better","values":[2399,30469,7767,19356,9157,5171],"isMain":true},{"metricName":"Total length (>= 1000 bp)","quality":"More is better","values":[0,25944,0,4716,4294,0],"isMain":true},{"metricName":"Total length (>= 5000 bp)","quality":"More is better","values":[0,20008,0,0,0,0],"isMain":false},{"metricName":"Total length (>= 10000 bp)","quality":"More is better","values":[0,20008,0,0,0,0],"isMain":true},{"metricName":"Total length (>= 25000 bp)","quality":"More is better","values":[0,0,0,0,0,0],"isMain":false},{"metricName":"Total length (>= 50000 bp)","quality":"More is better","values":[0,0,0,0,0,0],"isMain":true},{"metricName":"N50","quality":"More is better","values":[644,20008,531,628,758,564],"isMain":false},{"metricName":"N75","quality":"More is better","values":[561,4624,516,541,628,510],"isMain":false},{"metricName":"L50","quality":"Less is better","values":[2,1,6,9,4,5],"isMain":false},{"metricName":"L75","quality":"Less is better","values":[3,2,10,17,7,7],"isMain":false},{"metricName":"GC (%)","quality":"Equal","values":["48.65","43.89","45.81","44.53","45.78","46.53"],"isMain":false}]],["K-mer-based statistics",[]],["Predicted genes",[]],["Similarity statistics",[]],["Reference statistics",[{"metricName":"Reference length","quality":"Equal","values":[2849142,2849142,2849142,2849142,2849142,2849142],"isMain":false},{"metricName":"Reference fragments","quality":"Equal","values":[21,21,21,21,21,21],"isMain":false},{"metricName":"Reference GC (%)","quality":"Equal","values":["44.54","44.54","44.54","44.54","44.54","44.54"],"isMain":false}]]],"subreferences":[],"subreports":[],"minContig":500},{"date":"10 February 2020, Monday, 20:35:07","assembliesNames":["megahit-103-df","megahit-113-df","megahit-113-ml","megahit-113-ms","megahit-129-df","metaSPAdes"],"referenceName":"GCF\_000422625.1\_ASM42262v1","order":[0,1,2,3,4,5],"report":[["Genome statistics",[{"metricName":"Genome fraction (%)","quality":"More is better","values":["99.475","99.518","99.381","99.404","99.381","99.334"],"isMain":true},{"metricName":"Duplication ratio","quality":"Less is better","values":["1.003","1.002","1.003","1.003","1.003","1.002"],"isMain":true},{"metricName":"Largest alignment","quality":"More is better","values":[142742,307708,442835,443103,357372,442763],"isMain":true},{"metricName":"Total aligned length","quality":"More is better","values":[3001160,2998856,2994200,2993834,2997253,2990379],"isMain":true},{"metricName":"NG50","quality":"More is better","values":[63501,144192,144567,144595,111807,144423],"isMain":false},{"metricName":"NG75","quality":"More is better","values":[41733,76217,74012,106711,75375,84125],"isMain":false},{"metricName":"NA50","quality":"More is better","values":[63501,144192,144567,144595,106003,144423],"isMain":false},{"metricName":"NA75","quality":"More is better","values":[41628,76217,74012,102709,67160,84125],"isMain":false},{"metricName":"NGA50","quality":"More is better","values":[63501,144192,144567,144595,106003,144423],"isMain":true},{"metricName":"NGA75","quality":"More is better","values":[41628,76217,74012,102709,67160,84125],"isMain":false},{"metricName":"LG50","quality":"Less is better","values":[17,8,6,5,8,6],"isMain":false},{"metricName":"LG75","quality":"Less is better","values":[32,15,13,11,16,13],"isMain":false},{"metricName":"LA50","quality":"Less is better","values":[17,8,6,5,8,6],"isMain":false},{"metricName":"LA75","quality":"Less is better","values":[32,15,13,12,17,13],"isMain":false},{"metricName":"LGA50","quality":"Less is better","values":[17,8,6,5,8,6],"isMain":true},{"metricName":"LGA75","quality":"Less is better","values":[32,15,13,12,17,13],"isMain":false}]],["Reads mapping",[]],["Misassemblies",[{"metricName":"# misassemblies","quality":"Less is better","values":[4,5,2,3,6,4],"isMain":true},{"metricName":" # relocations","quality":"Less is better","values":[0,0,0,0,1,0],"isMain":false},{"metricName":" # translocations","quality":"Less is better","values":[4,5,2,3,5,4],"isMain":false},{"metricName":" # inversions","quality":"Less is better","values":[0,0,0,0,0,0],"isMain":false},{"metricName":" # interspecies translocations","quality":"Less is better","values":[7,6,3,4,7,1],"isMain":false},{"metricName":"# misassembled contigs","quality":"Less is better","values":[4,5,2,3,6,3],"isMain":false},{"metricName":"Misassembled contigs length","quality":"Less is better","values":[84577,474605,225021,180195,398290,116876],"isMain":true},{"metricName":"# possibly misassembled contigs","quality":"Less is better","values":[0,0,0,0,0,0],"isMain":false},{"metricName":" # possible misassemblies","quality":"Less is better","values":[0,0,0,0,0,0],"isMain":false},{"metricName":"# local misassemblies","quality":"Less is better","values":[0,1,0,1,1,0],"isMain":false},{"metricName":"# scaffold gap ext. mis.","quality":"Less is better","values":[0,0,0,0,0,0],"isMain":false},{"metricName":"# scaffold gap loc. mis.","quality":"Less is better","values":[0,0,0,0,0,0],"isMain":false},{"metricName":"# unaligned mis. contigs","quality":"Less is better","values":[0,0,0,0,0,0],"isMain":false}]],["Unaligned",[{"metricName":"# fully unaligned contigs","quality":"Less is better","values":[0,0,0,0,1,0],"isMain":false},{"metricName":"Fully unaligned length","quality":"Less is better","values":[0,0,0,0,580,0],"isMain":false},{"metricName":"# partially unaligned contigs","quality":"Less is better","values":[3,5,2,2,4,1],"isMain":false},{"metricName":"Partially unaligned length","quality":"Less is better","values":[1914,3311,1490,1355,22738,724],"isMain":false}]],["Mismatches",[{"metricName":"# mismatches","quality":"Less is better","values":[2082,280,236,357,454,221],"isMain":false},{"metricName":"# indels","quality":"Less is better","values":[0,2,3,3,12,11],"isMain":false},{"metricName":"Indels length","quality":"Less is better","values":[0,39,19,66,124,18],"isMain":false},{"metricName":"# mismatches per 100 kbp","quality":"Less is better","values":["69.54","9.35","7.89","11.93","15.18","7.39"],"isMain":true},{"metricName":"# indels per 100 kbp","quality":"Less is better","values":["0.00","0.07","0.10","0.10","0.40","0.37"],"isMain":true},{"metricName":" # indels (<= 5 bp)","quality":"Less is better","values":[0,0,2,0,7,11],"isMain":false},{"metricName":" # indels (> 5 bp)","quality":"Less is better","values":[0,2,1,3,5,0],"isMain":false},{"metricName":"# N's","quality":"Less is better","values":[0,0,0,0,0,0],"isMain":false},{"metricName":"# N's per 100 kbp","quality":"Less is better","values":["0.00","0.00","0.00","0.00","0.00","0.00"],"isMain":true}]],["Statistics without reference",[{"metricName":"# contigs","quality":"Equal","values":[101,49,48,38,51,40],"isMain":true},{"metricName":"# contigs (>= 1000 bp)","quality":"Equal","values":[85,41,44,35,43,34],"isMain":false},{"metricName":"# contigs (>= 5000 bp)","quality":"Equal","values":[71,32,31,26,35,30],"isMain":false},{"metricName":"# contigs (>= 10000 bp)","quality":"Equal","values":[60,30,29,24,32,29],"isMain":false},{"metricName":"# contigs (>= 25000 bp)","quality":"Equal","values":[39,27,24,21,28,26],"isMain":false},{"metricName":"# contigs (>= 50000 bp)","quality":"Equal","values":[25,21,18,17,22,18],"isMain":false},{"metricName":"Largest contig","quality":"More is better","values":[142746,307708,446771,447467,357372,446627],"isMain":true},{"metricName":"Total length","quality":"More is better","values":[3004422,3003331,3001393,3002241,3023726,2995311],"isMain":true},{"metricName":"Total length (>= 1000 bp)","quality":"More is better","values":[2992147,2997887,2999087,2999796,3018373,2990734],"isMain":true},{"metricName":"Total length (>= 5000 bp)","quality":"More is better","values":[2961087,2973672,2973281,2979080,2999574,2981935],"isMain":false},{"metricName":"Total length (>= 10000 bp)","quality":"More is better","values":[2881679,2962387,2962294,2964110,2981288,2976852],"isMain":true},{"metricName":"Total length (>= 25000 bp)","quality":"More is better","values":[2518024,2914118,2877308,2915259,2911732,2923840],"isMain":false},{"metricName":"Total length (>= 50000 bp)","quality":"More is better","values":[1981574,2694370,2631978,2750418,2672940,2602689],"isMain":true},{"metricName":"N50","quality":"More is better","values":[63501,144192,144567,144595,111807,144423],"isMain":false},{"metricName":"N75","quality":"More is better","values":[41733,76217,74012,106711,75375,84125],"isMain":false},{"metricName":"L50","quality":"Less is better","values":[17,8,6,5,8,6],"isMain":false},{"metricName":"L75","quality":"Less is better","values":[32,15,13,11,16,13],"isMain":false},{"metricName":"GC (%)","quality":"Equal","values":["64.31","64.31","64.31","64.31","64.29","64.31"],"isMain":false}]],["K-mer-based statistics",[]],["Predicted genes",[]],["Similarity statistics",[]],["Reference statistics",[{"metricName":"Reference length","quality":"Equal","values":[3009802,3009802,3009802,3009802,3009802,3009802],"isMain":false},{"metricName":"Reference fragments","quality":"Equal","values":[30,30,30,30,30,30],"isMain":false},{"metricName":"Reference GC (%)","quality":"Equal","values":["64.31","64.31","64.31","64.31","64.31","64.31"],"isMain":false}]]],"subreferences":[],"subreports":[],"minContig":500},{"date":"10 February 2020, Monday, 20:35:01","assembliesNames":["megahit-103-df","megahit-113-df","megahit-113-ml","megahit-113-ms","megahit-129-df","metaSPAdes"],"referenceName":"GCF\_000423925.1\_ASM42392v1","order":[0,1,2,3,4,5],"report":[["Genome statistics",[{"metricName":"Genome fraction (%)","quality":"More is better","values":["0.905","0.929","1.192","1.220","1.003","0.545"],"isMain":true},{"metricName":"Duplication ratio","quality":"Less is better","values":["1.131","1.128","1.019","1.068","1.071","1.123"],"isMain":true},{"metricName":"Largest alignment","quality":"More is better","values":[1229,969,1061,1051,1019,813],"isMain":true},{"metricName":"Total aligned length","quality":"More is better","values":[42516,43638,56057,57415,47132,25611],"isMain":true},{"metricName":"NA50","quality":"More is better","values":[527,530,568,544,532,529],"isMain":false},{"metricName":"NA75","quality":"More is better","values":[444,262,520,407,null,493],"isMain":false},{"metricName":"NGA50","quality":"More is better","values":[null,null,null,null,null,null],"isMain":true},{"metricName":"LA50","quality":"Less is better","values":[41,45,43,55,51,25],"isMain":false},{"metricName":"LA75","quality":"Less is better","values":[65,75,70,90,null,40],"isMain":false}]],["Reads mapping",[]],["Misassemblies",[{"metricName":"# misassemblies","quality":"Less is better","values":[0,0,0,0,0,0],"isMain":true},{"metricName":" # relocations","quality":"Less is better","values":[0,0,0,0,0,0],"isMain":false},{"metricName":" # translocations","quality":"Less is better","values":[0,0,0,0,0,0],"isMain":false},{"metricName":" # inversions","quality":"Less is better","values":[0,0,0,0,0,0],"isMain":false},{"metricName":" # interspecies translocations","quality":"Less is better","values":[19,20,2,20,17,9],"isMain":false},{"metricName":"# misassembled contigs","quality":"Less is better","values":[0,0,0,0,0,0],"isMain":false},{"metricName":"Misassembled contigs length","quality":"Less is better","values":[0,0,0,0,0,0],"isMain":true},{"metricName":"# possibly misassembled contigs","quality":"Less is better","values":[0,0,0,0,0,0],"isMain":false},{"metricName":" # possible misassemblies","quality":"Less is better","values":[0,0,0,0,0,0],"isMain":false},{"metricName":"# local misassemblies","quality":"Less is better","values":[0,1,0,1,0,0],"isMain":false},{"metricName":"# scaffold gap ext. mis.","quality":"Less is better","values":[0,0,0,0,0,0],"isMain":false},{"metricName":"# scaffold gap loc. mis.","quality":"Less is better","values":[0,0,0,0,0,0],"isMain":false},{"metricName":"# unaligned mis. contigs","quality":"Less is better","values":[0,0,0,0,0,0],"isMain":false}]],["Unaligned",[{"metricName":"# fully unaligned contigs","quality":"Less is better","values":[0,0,0,0,0,0],"isMain":false},{"metricName":"Fully unaligned length","quality":"Less is better","values":[0,0,0,0,0,0],"isMain":false},{"metricName":"# partially unaligned contigs","quality":"Less is better","values":[2,4,1,9,7,1],"isMain":false},{"metricName":"Partially unaligned length","quality":"Less is better","values":[1475,6272,542,9723,14090,1238],"isMain":false}]],["Mismatches",[{"metricName":"# mismatches","quality":"Less is better","values":[749,769,1209,1270,871,559],"isMain":false},{"metricName":"# indels","quality":"Less is better","values":[0,0,2,0,0,0],"isMain":false},{"metricName":"Indels length","quality":"Less is better","values":[0,0,2,0,0,0],"isMain":false},{"metricName":"# mismatches per 100 kbp","quality":"Less is better","values":["1761.69","1762.23","2158.04","2215.82","1848.00","2182.66"],"isMain":true},{"metricName":"# indels per 100 kbp","quality":"Less is better","values":["0.00","0.00","3.57","0.00","0.00","0.00"],"isMain":true},{"metricName":" # indels (<= 5 bp)","quality":"Less is better","values":[0,0,2,0,0,0],"isMain":false},{"metricName":" # indels (> 5 bp)","quality":"Less is better","values":[0,0,0,0,0,0],"isMain":false},{"metricName":"# N's","quality":"Less is better","values":[0,0,0,0,0,0],"isMain":false},{"metricName":"# N's per 100 kbp","quality":"Less is better","values":["0.00","0.00","0.00","0.00","0.00","0.00"],"isMain":true}]],["Statistics without reference",[{"metricName":"# contigs","quality":"Equal","values":[83,85,96,108,87,50],"isMain":true},{"metricName":"# contigs (>= 1000 bp)","quality":"Equal","values":[2,3,1,7,7,2],"isMain":false},{"metricName":"# contigs (>= 5000 bp)","quality":"Equal","values":[0,0,0,0,1,0],"isMain":false},{"metricName":"# contigs (>= 10000 bp)","quality":"Equal","values":[0,0,0,0,0,0],"isMain":false},{"metricName":"# contigs (>= 25000 bp)","quality":"Equal","values":[0,0,0,0,0,0],"isMain":false},{"metricName":"# contigs (>= 50000 bp)","quality":"Equal","values":[0,0,0,0,0,0],"isMain":false},{"metricName":"Largest contig","quality":"More is better","values":[1229,4086,1095,2603,8102,1364],"isMain":true},{"metricName":"Total length","quality":"More is better","values":[49542,55509,57606,70912,64553,29991],"isMain":true},{"metricName":"Total length (>= 1000 bp)","quality":"More is better","values":[2396,6522,1095,10267,16919,2450],"isMain":true},{"metricName":"Total length (>= 5000 bp)","quality":"More is better","values":[0,0,0,0,8102,0],"isMain":false},{"metricName":"Total length (>= 10000 bp)","quality":"More is better","values":[0,0,0,0,0,0],"isMain":true},{"metricName":"Total length (>= 25000 bp)","quality":"More is better","values":[0,0,0,0,0,0],"isMain":false},{"metricName":"Total length (>= 50000 bp)","quality":"More is better","values":[0,0,0,0,0,0],"isMain":true},{"metricName":"N50","quality":"More is better","values":[565,601,586,586,627,562],"isMain":false},{"metricName":"N75","quality":"More is better","values":[529,538,528,538,550,526],"isMain":false},{"metricName":"L50","quality":"Less is better","values":[36,34,43,43,29,22],"isMain":false},{"metricName":"L75","quality":"Less is better","values":[59,59,68,74,56,36],"isMain":false},{"metricName":"GC (%)","quality":"Equal","values":["41.14","41.00","41.45","41.77","41.15","41.29"],"isMain":false}]],["K-mer-based statistics",[]],["Predicted genes",[]],["Similarity statistics",[]],["Reference statistics",[{"metricName":"Reference length","quality":"Equal","values":[4699139,4699139,4699139,4699139,4699139,4699139],"isMain":false},{"metricName":"Reference fragments","quality":"Equal","values":[82,82,82,82,82,82],"isMain":false},{"metricName":"Reference GC (%)","quality":"Equal","values":["41.19","41.19","41.19","41.19","41.19","41.19"],"isMain":false}]]],"subreferences":[],"subreports":[],"minContig":500},{"date":"10 February 2020, Monday, 20:35:30","assembliesNames":["megahit-103-df","megahit-113-df","megahit-113-ml","megahit-113-ms","megahit-129-df","metaSPAdes"],"referenceName":"GCF\_000423945.1\_ASM42394v1","order":[0,1,2,3,4,5],"report":[["Genome statistics",[{"metricName":"Genome fraction (%)","quality":"More is better","values":["0.268","0.629","0.093","0.299","0.377","0.018"],"isMain":true},{"metricName":"Duplication ratio","quality":"Less is better","values":["1.471","1.076","1.049","1.226","1.133","1.623"],"isMain":true},{"metricName":"Largest alignment","quality":"More is better","values":[1450,12541,713,3665,6129,717],"isMain":true},{"metricName":"Total aligned length","quality":"More is better","values":[11907,28937,4144,13555,17356,798],"isMain":true},{"metricName":"NA50","quality":"More is better","values":[null,848,628,null,1961,717],"isMain":false},{"metricName":"NA75","quality":"More is better","values":[null,null,526,null,181,null],"isMain":false},{"metricName":"NGA50","quality":"More is better","values":[null,null,null,null,null,null],"isMain":true},{"metricName":"LA50","quality":"Less is better","values":[null,5,4,null,3,1],"isMain":false},{"metricName":"LA75","quality":"Less is better","values":[null,null,6,null,13,null],"isMain":false}]],["Reads mapping",[]],["Misassemblies",[{"metricName":"# misassemblies","quality":"Less is better","values":[0,4,0,0,1,0],"isMain":true},{"metricName":" # relocations","quality":"Less is better","values":[0,3,0,0,1,0],"isMain":false},{"metricName":" # translocations","quality":"Less is better","values":[0,1,0,0,0,0],"isMain":false},{"metricName":" # inversions","quality":"Less is better","values":[0,0,0,0,0,0],"isMain":false},{"metricName":" # interspecies translocations","quality":"Less is better","values":[26,5,1,10,7,1],"isMain":false},{"metricName":"# misassembled contigs","quality":"Less is better","values":[0,2,0,0,1,0],"isMain":false},{"metricName":"Misassembled contigs length","quality":"Less is better","values":[0,44974,0,0,11878,0],"isMain":true},{"metricName":"# possibly misassembled contigs","quality":"Less is better","values":[0,0,0,0,0,0],"isMain":false},{"metricName":" # possible misassemblies","quality":"Less is better","values":[0,0,0,0,0,0],"isMain":false},{"metricName":"# local misassemblies","quality":"Less is better","values":[0,2,1,1,0,0],"isMain":false},{"metricName":"# scaffold gap ext. mis.","quality":"Less is better","values":[0,0,0,0,0,0],"isMain":false},{"metricName":"# scaffold gap loc. mis.","quality":"Less is better","values":[0,0,0,0,0,0],"isMain":false},{"metricName":"# unaligned mis. contigs","quality":"Less is better","values":[0,0,0,1,0,0],"isMain":false}]],["Unaligned",[{"metricName":"# fully unaligned contigs","quality":"Less is better","values":[0,1,0,0,0,0],"isMain":false},{"metricName":"Fully unaligned length","quality":"Less is better","values":[0,640,0,0,0,0],"isMain":false},{"metricName":"# partially unaligned contigs","quality":"Less is better","values":[13,1,0,2,2,0],"isMain":false},{"metricName":"Partially unaligned length","quality":"Less is better","values":[12033,18626,0,40186,3541,0],"isMain":false}]],["Mismatches",[{"metricName":"# mismatches","quality":"Less is better","values":[151,414,98,237,227,20],"isMain":false},{"metricName":"# indels","quality":"Less is better","values":[0,17,6,3,8,0],"isMain":false},{"metricName":"Indels length","quality":"Less is better","values":[0,276,6,7,82,0],"isMain":false},{"metricName":"# mismatches per 100 kbp","quality":"Less is better","values":["1268.16","1482.60","2364.86","1785.85","1356.60","2506.27"],"isMain":true},{"metricName":"# indels per 100 kbp","quality":"Less is better","values":["0.00","60.88","144.79","22.61","47.81","0.00"],"isMain":true},{"metricName":" # indels (<= 5 bp)","quality":"Less is better","values":[0,10,6,3,7,0],"isMain":false},{"metricName":" # indels (> 5 bp)","quality":"Less is better","values":[0,7,0,0,1,0],"isMain":false},{"metricName":"# N's","quality":"Less is better","values":[0,0,0,0,0,0],"isMain":false},{"metricName":"# N's per 100 kbp","quality":"Less is better","values":["0.00","0.00","0.00","0.00","0.00","0.00"],"isMain":true}]],["Statistics without reference",[{"metricName":"# contigs","quality":"Equal","values":[34,9,7,21,15,2],"isMain":true},{"metricName":"# contigs (>= 1000 bp)","quality":"Equal","values":[10,2,0,4,3,0],"isMain":false},{"metricName":"# contigs (>= 5000 bp)","quality":"Equal","values":[0,1,0,1,1,0],"isMain":false},{"metricName":"# contigs (>= 10000 bp)","quality":"Equal","values":[0,1,0,1,1,0],"isMain":false},{"metricName":"# contigs (>= 25000 bp)","quality":"Equal","values":[0,1,0,1,0,0],"isMain":false},{"metricName":"# contigs (>= 50000 bp)","quality":"Equal","values":[0,0,0,0,0,0],"isMain":false},{"metricName":"Largest contig","quality":"More is better","values":[1934,40599,740,38309,11878,720],"isMain":true},{"metricName":"Total length","quality":"More is better","values":[29544,49323,4347,56453,22494,1295],"isMain":true},{"metricName":"Total length (>= 1000 bp)","quality":"More is better","values":[13123,44974,0,46445,15553,0],"isMain":true},{"metricName":"Total length (>= 5000 bp)","quality":"More is better","values":[0,40599,0,38309,11878,0],"isMain":false},{"metricName":"Total length (>= 10000 bp)","quality":"More is better","values":[0,40599,0,38309,11878,0],"isMain":true},{"metricName":"Total length (>= 25000 bp)","quality":"More is better","values":[0,40599,0,38309,0,0],"isMain":false},{"metricName":"Total length (>= 50000 bp)","quality":"More is better","values":[0,0,0,0,0,0],"isMain":true},{"metricName":"N50","quality":"More is better","values":[893,40599,639,38309,11878,720],"isMain":false},{"metricName":"N75","quality":"More is better","values":[678,40599,579,4084,664,575],"isMain":false},{"metricName":"L50","quality":"Less is better","values":[12,1,4,1,1,1],"isMain":false},{"metricName":"L75","quality":"Less is better","values":[22,1,5,2,5,2],"isMain":false},{"metricName":"GC (%)","quality":"Equal","values":["41.59","40.92","42.33","42.44","42.72","47.57"],"isMain":false}]],["K-mer-based statistics",[]],["Predicted genes",[]],["Similarity statistics",[]],["Reference statistics",[{"metricName":"Reference length","quality":"Equal","values":[4438133,4438133,4438133,4438133,4438133,4438133],"isMain":false},{"metricName":"Reference fragments","quality":"Equal","values":[53,53,53,53,53,53],"isMain":false},{"metricName":"Reference GC (%)","quality":"Equal","values":["41.77","41.77","41.77","41.77","41.77","41.77"],"isMain":false}]]],"subreferences":[],"subreports":[],"minContig":500},{"date":"10 February 2020, Monday, 20:35:52","assembliesNames":["megahit-103-df","megahit-113-df","megahit-113-ml","megahit-113-ms","megahit-129-df","metaSPAdes"],"referenceName":"GCF\_000423985.1\_ASM42398v1","order":[0,1,2,3,4,5],"report":[["Genome statistics",[{"metricName":"Genome fraction (%)","quality":"More is better","values":["6.016","9.128","8.529","7.882","7.678","3.804"],"isMain":true},{"metricName":"Duplication ratio","quality":"Less is better","values":["1.023","1.031","1.039","1.044","1.033","1.026"],"isMain":true},{"metricName":"Largest alignment","quality":"More is better","values":[6815,9786,12150,5033,10011,7656],"isMain":true},{"metricName":"Total aligned length","quality":"More is better","values":[283029,430467,406469,375877,363046,179150],"isMain":true},{"metricName":"NA50","quality":"More is better","values":[1202,1917,1020,1122,1335,1172],"isMain":false},{"metricName":"NA75","quality":"More is better","values":[756,925,679,688,673,792],"isMain":false},{"metricName":"NGA50","quality":"More is better","values":[null,null,null,null,null,null],"isMain":true},{"metricName":"LA50","quality":"Less is better","values":[72,63,117,102,84,49],"isMain":false},{"metricName":"LA75","quality":"Less is better","values":[150,151,246,217,200,98],"isMain":false}]],["Reads mapping",[]],["Misassemblies",[{"metricName":"# misassemblies","quality":"Less is better","values":[12,5,10,9,19,4],"isMain":true},{"metricName":" # relocations","quality":"Less is better","values":[2,1,1,3,5,0],"isMain":false},{"metricName":" # translocations","quality":"Less is better","values":[8,3,8,5,12,4],"isMain":false},{"metricName":" # inversions","quality":"Less is better","values":[2,1,1,1,2,0],"isMain":false},{"metricName":" # interspecies translocations","quality":"Less is better","values":[17,23,19,30,35,8],"isMain":false},{"metricName":"# misassembled contigs","quality":"Less is better","values":[9,4,9,8,13,3],"isMain":false},{"metricName":"Misassembled contigs length","quality":"Less is better","values":[14710,16531,10725,11719,34641,13319],"isMain":true},{"metricName":"# possibly misassembled contigs","quality":"Less is better","values":[0,0,0,0,0,0],"isMain":false},{"metricName":" # possible misassemblies","quality":"Less is better","values":[0,0,0,0,0,0],"isMain":false},{"metricName":"# local misassemblies","quality":"Less is better","values":[3,11,6,5,8,2],"isMain":false},{"metricName":"# scaffold gap ext. mis.","quality":"Less is better","values":[0,0,0,0,0,0],"isMain":false},{"metricName":"# scaffold gap loc. mis.","quality":"Less is better","values":[0,0,0,0,0,0],"isMain":false},{"metricName":"# unaligned mis. contigs","quality":"Less is better","values":[1,3,1,0,2,0],"isMain":false}]],["Unaligned",[{"metricName":"# fully unaligned contigs","quality":"Less is better","values":[0,0,0,0,0,0],"isMain":false},{"metricName":"Fully unaligned length","quality":"Less is better","values":[0,0,0,0,0,0],"isMain":false},{"metricName":"# partially unaligned contigs","quality":"Less is better","values":[5,9,8,12,15,3],"isMain":false},{"metricName":"Partially unaligned length","quality":"Less is better","values":[7884,16744,10790,14674,43449,4055],"isMain":false}]],["Mismatches",[{"metricName":"# mismatches","quality":"Less is better","values":[3334,6223,7298,6527,5538,1738],"isMain":false},{"metricName":"# indels","quality":"Less is better","values":[39,109,87,79,107,23],"isMain":false},{"metricName":"Indels length","quality":"Less is better","values":[154,382,333,196,353,97],"isMain":false},{"metricName":"# mismatches per 100 kbp","quality":"Less is better","values":["1191.38","1465.71","1839.66","1780.39","1550.68","982.28"],"isMain":true},{"metricName":"# indels per 100 kbp","quality":"Less is better","values":["13.94","25.67","21.93","21.55","29.96","13.00"],"isMain":true},{"metricName":" # indels (<= 5 bp)","quality":"Less is better","values":[37,94,76,75,94,17],"isMain":false},{"metricName":" # indels (> 5 bp)","quality":"Less is better","values":[2,15,11,4,13,6],"isMain":false},{"metricName":"# N's","quality":"Less is better","values":[0,0,0,0,0,0],"isMain":false},{"metricName":"# N's per 100 kbp","quality":"Less is better","values":["0.00","0.00","0.00","0.00","0.00","0.00"],"isMain":true}]],["Statistics without reference",[{"metricName":"# contigs","quality":"Equal","values":[247,284,397,354,286,159],"isMain":true},{"metricName":"# contigs (>= 1000 bp)","quality":"Equal","values":[102,132,132,131,113,64],"isMain":false},{"metricName":"# contigs (>= 5000 bp)","quality":"Equal","values":[1,12,4,2,9,3],"isMain":false},{"metricName":"# contigs (>= 10000 bp)","quality":"Equal","values":[0,0,1,0,4,0],"isMain":false},{"metricName":"# contigs (>= 25000 bp)","quality":"Equal","values":[0,0,0,0,0,0],"isMain":false},{"metricName":"# contigs (>= 50000 bp)","quality":"Equal","values":[0,0,0,0,0,0],"isMain":false},{"metricName":"Largest contig","quality":"More is better","values":[6826,9947,12203,7011,18829,7656],"isMain":true},{"metricName":"Total length","quality":"More is better","values":[294222,454632,423075,397458,412258,185578],"isMain":true},{"metricName":"Total length (>= 1000 bp)","quality":"More is better","values":[191980,347943,239574,245194,291826,116668],"isMain":true},{"metricName":"Total length (>= 5000 bp)","quality":"More is better","values":[6826,85043,28409,12057,83064,20109],"isMain":false},{"metricName":"Total length (>= 10000 bp)","quality":"More is better","values":[0,0,12203,0,50498,0],"isMain":true},{"metricName":"Total length (>= 25000 bp)","quality":"More is better","values":[0,0,0,0,0,0],"isMain":false},{"metricName":"Total length (>= 50000 bp)","quality":"More is better","values":[0,0,0,0,0,0],"isMain":true},{"metricName":"N50","quality":"More is better","values":[1383,2239,1135,1247,1955,1297],"isMain":false},{"metricName":"N75","quality":"More is better","values":[845,1043,735,761,916,838],"isMain":false},{"metricName":"L50","quality":"Less is better","values":[64,55,106,90,53,43],"isMain":false},{"metricName":"L75","quality":"Less is better","values":[134,126,223,193,132,89],"isMain":false},{"metricName":"GC (%)","quality":"Equal","values":["39.87","39.48","39.99","39.87","39.71","38.72"],"isMain":false}]],["K-mer-based statistics",[]],["Predicted genes",[]],["Similarity statistics",[]],["Reference statistics",[{"metricName":"Reference length","quality":"Equal","values":[4651344,4651344,4651344,4651344,4651344,4651344],"isMain":false},{"metricName":"Reference fragments","quality":"Equal","values":[82,82,82,82,82,82],"isMain":false},{"metricName":"Reference GC (%)","quality":"Equal","values":["39.78","39.78","39.78","39.78","39.78","39.78"],"isMain":false}]]],"subreferences":[],"subreports":[],"minContig":500},{"date":"10 February 2020, Monday, 20:36:16","assembliesNames":["megahit-103-df","megahit-113-df","megahit-113-ml","megahit-113-ms","megahit-129-df","metaSPAdes"],"referenceName":"GCF\_000424005.1\_ASM42400v1","order":[0,1,2,3,4,5],"report":[["Genome statistics",[{"metricName":"Genome fraction (%)","quality":"More is better","values":["14.699","17.924","18.878","20.218","19.926","8.615"],"isMain":true},{"metricName":"Duplication ratio","quality":"Less is better","values":["1.038","1.040","1.032","1.044","1.042","1.040"],"isMain":true},{"metricName":"Largest alignment","quality":"More is better","values":[3758,2942,2942,2942,6028,2942],"isMain":true},{"metricName":"Total aligned length","quality":"More is better","values":[484292,589631,624583,669375,656053,283177],"isMain":true},{"metricName":"NA50","quality":"More is better","values":[627,641,627,627,631,593],"isMain":false},{"metricName":"NA75","quality":"More is better","values":[540,533,532,534,527,525],"isMain":false},{"metricName":"NGA50","quality":"More is better","values":[null,null,null,null,null,null],"isMain":true},{"metricName":"LA50","quality":"Less is better","values":[316,386,416,450,442,200],"isMain":false},{"metricName":"LA75","quality":"Less is better","values":[536,663,706,763,761,337],"isMain":false}]],["Reads mapping",[]],["Misassemblies",[{"metricName":"# misassemblies","quality":"Less is better","values":[3,0,1,1,3,0],"isMain":true},{"metricName":" # relocations","quality":"Less is better","values":[2,0,1,1,2,0],"isMain":false},{"metricName":" # translocations","quality":"Less is better","values":[0,0,0,0,1,0],"isMain":false},{"metricName":" # inversions","quality":"Less is better","values":[1,0,0,0,0,0],"isMain":false},{"metricName":" # interspecies translocations","quality":"Less is better","values":[58,82,36,67,94,33],"isMain":false},{"metricName":"# misassembled contigs","quality":"Less is better","values":[3,0,1,1,3,0],"isMain":false},{"metricName":"Misassembled contigs length","quality":"Less is better","values":[4614,0,591,569,2021,0],"isMain":true},{"metricName":"# possibly misassembled contigs","quality":"Less is better","values":[0,0,0,0,0,0],"isMain":false},{"metricName":" # possible misassemblies","quality":"Less is better","values":[0,0,0,0,0,0],"isMain":false},{"metricName":"# local misassemblies","quality":"Less is better","values":[2,1,2,2,1,0],"isMain":false},{"metricName":"# scaffold gap ext. mis.","quality":"Less is better","values":[0,0,0,0,0,0],"isMain":false},{"metricName":"# scaffold gap loc. mis.","quality":"Less is better","values":[0,0,0,0,0,0],"isMain":false},{"metricName":"# unaligned mis. contigs","quality":"Less is better","values":[1,0,0,0,0,0],"isMain":false}]],["Unaligned",[{"metricName":"# fully unaligned contigs","quality":"Less is better","values":[0,0,7,1,4,0],"isMain":false},{"metricName":"Fully unaligned length","quality":"Less is better","values":[0,0,4705,755,3874,0],"isMain":false},{"metricName":"# partially unaligned contigs","quality":"Less is better","values":[14,33,17,21,34,11],"isMain":false},{"metricName":"Partially unaligned length","quality":"Less is better","values":[11853,37815,22544,28542,51268,9951],"isMain":false}]],["Mismatches",[{"metricName":"# mismatches","quality":"Less is better","values":[7588,9038,12249,13822,10462,4584],"isMain":false},{"metricName":"# indels","quality":"Less is better","values":[24,31,28,28,46,8],"isMain":false},{"metricName":"Indels length","quality":"Less is better","values":[50,136,49,60,99,10],"isMain":false},{"metricName":"# mismatches per 100 kbp","quality":"Less is better","values":["1578.43","1541.77","1983.96","2090.30","1605.36","1627.00"],"isMain":true},{"metricName":"# indels per 100 kbp","quality":"Less is better","values":["4.99","5.29","4.54","4.23","7.06","2.84"],"isMain":true},{"metricName":" # indels (<= 5 bp)","quality":"Less is better","values":[21,28,27,27,42,8],"isMain":false},{"metricName":" # indels (> 5 bp)","quality":"Less is better","values":[3,3,1,1,4,0],"isMain":false},{"metricName":"# N's","quality":"Less is better","values":[0,0,0,0,0,0],"isMain":false},{"metricName":"# N's per 100 kbp","quality":"Less is better","values":["0.00","0.00","0.00","0.00","0.00","0.00"],"isMain":true}]],["Statistics without reference",[{"metricName":"# contigs","quality":"Equal","values":[748,894,970,1040,1002,461],"isMain":true},{"metricName":"# contigs (>= 1000 bp)","quality":"Equal","values":[38,83,57,67,89,23],"isMain":false},{"metricName":"# contigs (>= 5000 bp)","quality":"Equal","values":[0,1,1,1,2,0],"isMain":false},{"metricName":"# contigs (>= 10000 bp)","quality":"Equal","values":[0,0,0,0,1,0],"isMain":false},{"metricName":"# contigs (>= 25000 bp)","quality":"Equal","values":[0,0,0,0,0,0],"isMain":false},{"metricName":"# contigs (>= 50000 bp)","quality":"Equal","values":[0,0,0,0,0,0],"isMain":false},{"metricName":"Largest contig","quality":"More is better","values":[3758,9223,6471,7330,22846,3070],"isMain":true},{"metricName":"Total length","quality":"More is better","values":[510735,647588,664465,719378,734068,302856],"isMain":true},{"metricName":"Total length (>= 1000 bp)","quality":"More is better","values":[54057,116279,80403,94424,138908,31987],"isMain":true},{"metricName":"Total length (>= 5000 bp)","quality":"More is better","values":[0,9223,6471,7330,28874,0],"isMain":false},{"metricName":"Total length (>= 10000 bp)","quality":"More is better","values":[0,0,0,0,22846,0],"isMain":true},{"metricName":"Total length (>= 25000 bp)","quality":"More is better","values":[0,0,0,0,0,0],"isMain":false},{"metricName":"Total length (>= 50000 bp)","quality":"More is better","values":[0,0,0,0,0,0],"isMain":true},{"metricName":"N50","quality":"More is better","values":[653,702,659,668,700,625],"isMain":false},{"metricName":"N75","quality":"More is better","values":[569,583,559,565,584,550],"isMain":false},{"metricName":"L50","quality":"Less is better","values":[298,338,381,408,371,187],"isMain":false},{"metricName":"L75","quality":"Less is better","values":[508,593,656,701,660,317],"isMain":false},{"metricName":"GC (%)","quality":"Equal","values":["39.39","39.37","39.64","39.42","39.35","39.03"],"isMain":false}]],["K-mer-based statistics",[]],["Predicted genes",[]],["Similarity statistics",[]],["Reference statistics",[{"metricName":"Reference length","quality":"Equal","values":[3270570,3270570,3270570,3270570,3270570,3270570],"isMain":false},{"metricName":"Reference fragments","quality":"Equal","values":[17,17,17,17,17,17],"isMain":false},{"metricName":"Reference GC (%)","quality":"Equal","values":["39.50","39.50","39.50","39.50","39.50","39.50"],"isMain":false}]]],"subreferences":[],"subreports":[],"minContig":500},{"date":"10 February 2020, Monday, 20:36:06","assembliesNames":["megahit-103-df","megahit-113-df","megahit-113-ml","megahit-113-ms","megahit-129-df","metaSPAdes"],"referenceName":"GCF\_000424085.1\_ASM42408v1","order":[0,1,2,3,4,5],"report":[["Genome statistics",[{"metricName":"Genome fraction (%)","quality":"More is better","values":["0.265","0.243","0.237","0.408","0.275","0.079"],"isMain":true},{"metricName":"Duplication ratio","quality":"Less is better","values":["1.141","1.142","1.241","1.108","1.147","1.131"],"isMain":true},{"metricName":"Largest alignment","quality":"More is better","values":[2147,1274,1232,3816,1591,1465],"isMain":true},{"metricName":"Total aligned length","quality":"More is better","values":[9499,8711,8959,14614,9833,2815],"isMain":true},{"metricName":"NA50","quality":"More is better","values":[557,null,null,null,null,null],"isMain":false},{"metricName":"NGA50","quality":"More is better","values":[null,null,null,null,null,null],"isMain":true},{"metricName":"LA50","quality":"Less is better","values":[6,null,null,null,null,null],"isMain":false}]],["Reads mapping",[]],["Misassemblies",[{"metricName":"# misassemblies","quality":"Less is better","values":[2,1,1,3,1,0],"isMain":true},{"metricName":" # relocations","quality":"Less is better","values":[0,0,0,0,0,0],"isMain":false},{"metricName":" # translocations","quality":"Less is better","values":[2,1,1,3,1,0],"isMain":false},{"metricName":" # inversions","quality":"Less is better","values":[0,0,0,0,0,0],"isMain":false},{"metricName":" # interspecies translocations","quality":"Less is better","values":[6,7,4,5,7,4],"isMain":false},{"metricName":"# misassembled contigs","quality":"Less is better","values":[2,1,1,3,1,0],"isMain":false},{"metricName":"Misassembled contigs length","quality":"Less is better","values":[1110,515,795,1675,545,0],"isMain":true},{"metricName":"# possibly misassembled contigs","quality":"Less is better","values":[0,0,0,0,0,0],"isMain":false},{"metricName":" # possible misassemblies","quality":"Less is better","values":[0,0,0,0,0,0],"isMain":false},{"metricName":"# local misassemblies","quality":"Less is better","values":[0,0,0,0,0,0],"isMain":false},{"metricName":"# scaffold gap ext. mis.","quality":"Less is better","values":[0,0,0,0,0,0],"isMain":false},{"metricName":"# scaffold gap loc. mis.","quality":"Less is better","values":[0,0,0,0,0,0],"isMain":false},{"metricName":"# unaligned mis. contigs","quality":"Less is better","values":[0,0,0,0,0,0],"isMain":false}]],["Unaligned",[{"metricName":"# fully unaligned contigs","quality":"Less is better","values":[0,0,0,0,1,0],"isMain":false},{"metricName":"Fully unaligned length","quality":"Less is better","values":[0,0,0,0,535,0],"isMain":false},{"metricName":"# partially unaligned contigs","quality":"Less is better","values":[2,4,2,2,2,2],"isMain":false},{"metricName":"Partially unaligned length","quality":"Less is better","values":[2197,28865,17952,22943,15250,21296],"isMain":false}]],["Mismatches",[{"metricName":"# mismatches","quality":"Less is better","values":[168,141,217,373,186,49],"isMain":false},{"metricName":"# indels","quality":"Less is better","values":[3,3,1,5,1,1],"isMain":false},{"metricName":"Indels length","quality":"Less is better","values":[7,5,1,26,1,1],"isMain":false},{"metricName":"# mismatches per 100 kbp","quality":"Less is better","values":["1768.79","1618.83","2553.84","2552.70","1891.78","1741.29"],"isMain":true},{"metricName":"# indels per 100 kbp","quality":"Less is better","values":["31.59","34.44","11.77","34.22","10.17","35.54"],"isMain":true},{"metricName":" # indels (<= 5 bp)","quality":"Less is better","values":[3,3,1,4,1,1],"isMain":false},{"metricName":" # indels (> 5 bp)","quality":"Less is better","values":[0,0,0,1,0,0],"isMain":false},{"metricName":"# N's","quality":"Less is better","values":[0,0,0,0,0,0],"isMain":false},{"metricName":"# N's per 100 kbp","quality":"Less is better","values":["0.00","0.00","0.00","0.00","0.00","0.00"],"isMain":true}]],["Statistics without reference",[{"metricName":"# contigs","quality":"Equal","values":[14,17,16,21,18,5],"isMain":true},{"metricName":"# contigs (>= 1000 bp)","quality":"Equal","values":[3,3,2,2,3,2],"isMain":false},{"metricName":"# contigs (>= 5000 bp)","quality":"Equal","values":[0,2,1,1,1,1],"isMain":false},{"metricName":"# contigs (>= 10000 bp)","quality":"Equal","values":[0,2,1,1,1,1],"isMain":false},{"metricName":"# contigs (>= 25000 bp)","quality":"Equal","values":[0,0,0,0,0,0],"isMain":false},{"metricName":"# contigs (>= 50000 bp)","quality":"Equal","values":[0,0,0,0,0,0],"isMain":false},{"metricName":"Largest contig","quality":"More is better","values":[3022,18012,17934,23573,15404,21811],"isMain":true},{"metricName":"Total length","quality":"More is better","values":[13032,38810,28497,39133,27062,24479],"isMain":true},{"metricName":"Total length (>= 1000 bp)","quality":"More is better","values":[6498,30650,19380,27389,18323,22893],"isMain":true},{"metricName":"Total length (>= 5000 bp)","quality":"More is better","values":[0,29508,17934,23573,15404,21811],"isMain":false},{"metricName":"Total length (>= 10000 bp)","quality":"More is better","values":[0,29508,17934,23573,15404,21811],"isMain":true},{"metricName":"Total length (>= 25000 bp)","quality":"More is better","values":[0,0,0,0,0,0],"isMain":false},{"metricName":"Total length (>= 50000 bp)","quality":"More is better","values":[0,0,0,0,0,0],"isMain":true},{"metricName":"N50","quality":"More is better","values":[839,11496,17934,23573,15404,21811],"isMain":false},{"metricName":"N75","quality":"More is better","values":[566,11496,795,759,652,21811],"isMain":false},{"metricName":"L50","quality":"Less is better","values":[4,2,1,1,1,1],"isMain":false},{"metricName":"L75","quality":"Less is better","values":[8,2,5,5,6,1],"isMain":false},{"metricName":"GC (%)","quality":"Equal","values":["42.17","34.45","35.31","36.63","35.67","33.60"],"isMain":false}]],["K-mer-based statistics",[]],["Predicted genes",[]],["Similarity statistics",[]],["Reference statistics",[{"metricName":"Reference length","quality":"Equal","values":[3581157,3581157,3581157,3581157,3581157,3581157],"isMain":false},{"metricName":"Reference fragments","quality":"Equal","values":[192,192,192,192,192,192],"isMain":false},{"metricName":"Reference GC (%)","quality":"Equal","values":["41.58","41.58","41.58","41.58","41.58","41.58"],"isMain":false}]]],"subreferences":[],"subreports":[],"minContig":500},{"date":"10 February 2020, Monday, 20:36:21","assembliesNames":["megahit-103-df","megahit-113-df","megahit-113-ml","megahit-113-ms","megahit-129-df","metaSPAdes"],"referenceName":"GCF\_000424105.1\_ASM42410v1","order":[0,1,2,3,4,5],"report":[["Genome statistics",[{"metricName":"Genome fraction (%)","quality":"More is better","values":["0.065","0.007","0.031","0.018","0.087","0.021"],"isMain":true},{"metricName":"Duplication ratio","quality":"Less is better","values":["1.008","2.488","1.449","1.000","1.145","1.000"],"isMain":true},{"metricName":"Largest alignment","quality":"More is better","values":[1856,213,736,526,2257,609],"isMain":true},{"metricName":"Total aligned length","quality":"More is better","values":[1856,213,889,526,2488,609],"isMain":true},{"metricName":"NA50","quality":"More is better","values":[1856,null,736,526,2257,609],"isMain":false},{"metricName":"NA75","quality":"More is better","values":[1856,null,null,526,2257,609],"isMain":false},{"metricName":"NGA50","quality":"More is better","values":[null,null,null,null,null,null],"isMain":true},{"metricName":"LA50","quality":"Less is better","values":[1,null,1,1,1,1],"isMain":false},{"metricName":"LA75","quality":"Less is better","values":[1,null,null,1,1,1],"isMain":false}]],["Reads mapping",[]],["Misassemblies",[{"metricName":"# misassemblies","quality":"Less is better","values":[0,0,0,0,0,0],"isMain":true},{"metricName":" # relocations","quality":"Less is better","values":[0,0,0,0,0,0],"isMain":false},{"metricName":" # translocations","quality":"Less is better","values":[0,0,0,0,0,0],"isMain":false},{"metricName":" # inversions","quality":"Less is better","values":[0,0,0,0,0,0],"isMain":false},{"metricName":" # interspecies translocations","quality":"Less is better","values":[0,1,1,0,1,0],"isMain":false},{"metricName":"# misassembled contigs","quality":"Less is better","values":[0,0,0,0,0,0],"isMain":false},{"metricName":"Misassembled contigs length","quality":"Less is better","values":[0,0,0,0,0,0],"isMain":true},{"metricName":"# possibly misassembled contigs","quality":"Less is better","values":[0,0,0,0,0,0],"isMain":false},{"metricName":" # possible misassemblies","quality":"Less is better","values":[0,0,0,0,0,0],"isMain":false},{"metricName":"# local misassemblies","quality":"Less is better","values":[0,0,0,0,0,0],"isMain":false},{"metricName":"# scaffold gap ext. mis.","quality":"Less is better","values":[0,0,0,0,0,0],"isMain":false},{"metricName":"# scaffold gap loc. mis.","quality":"Less is better","values":[0,0,0,0,0,0],"isMain":false},{"metricName":"# unaligned mis. contigs","quality":"Less is better","values":[0,0,0,0,0,0],"isMain":false}]],["Unaligned",[{"metricName":"# fully unaligned contigs","quality":"Less is better","values":[0,0,0,0,0,0],"isMain":false},{"metricName":"Fully unaligned length","quality":"Less is better","values":[0,0,0,0,0,0],"isMain":false},{"metricName":"# partially unaligned contigs","quality":"Less is better","values":[0,0,0,0,0,0],"isMain":false},{"metricName":"Partially unaligned length","quality":"Less is better","values":[0,0,0,0,0,0],"isMain":false}]],["Mismatches",[{"metricName":"# mismatches","quality":"Less is better","values":[49,3,20,12,59,28],"isMain":false},{"metricName":"# indels","quality":"Less is better","values":[0,0,0,0,0,0],"isMain":false},{"metricName":"Indels length","quality":"Less is better","values":[0,0,0,0,0,0],"isMain":false},{"metricName":"# mismatches per 100 kbp","quality":"Less is better","values":["2640.09","1408.45","2249.72","2281.37","2371.38","4597.70"],"isMain":true},{"metricName":"# indels per 100 kbp","quality":"Less is better","values":["0.00","0.00","0.00","0.00","0.00","0.00"],"isMain":true},{"metricName":" # indels (<= 5 bp)","quality":"Less is better","values":[0,0,0,0,0,0],"isMain":false},{"metricName":" # indels (> 5 bp)","quality":"Less is better","values":[0,0,0,0,0,0],"isMain":false},{"metricName":"# N's","quality":"Less is better","values":[0,0,0,0,0,0],"isMain":false},{"metricName":"# N's per 100 kbp","quality":"Less is better","values":["0.00","0.00","0.00","0.00","0.00","0.00"],"isMain":true}]],["Statistics without reference",[{"metricName":"# contigs","quality":"Equal","values":[1,1,2,1,2,1],"isMain":true},{"metricName":"# contigs (>= 1000 bp)","quality":"Equal","values":[1,0,0,0,1,0],"isMain":false},{"metricName":"# contigs (>= 5000 bp)","quality":"Equal","values":[0,0,0,0,0,0],"isMain":false},{"metricName":"# contigs (>= 10000 bp)","quality":"Equal","values":[0,0,0,0,0,0],"isMain":false},{"metricName":"# contigs (>= 25000 bp)","quality":"Equal","values":[0,0,0,0,0,0],"isMain":false},{"metricName":"# contigs (>= 50000 bp)","quality":"Equal","values":[0,0,0,0,0,0],"isMain":false},{"metricName":"Largest contig","quality":"More is better","values":[1871,530,746,526,2272,609],"isMain":true},{"metricName":"Total length","quality":"More is better","values":[1871,530,1288,526,2848,609],"isMain":true},{"metricName":"Total length (>= 1000 bp)","quality":"More is better","values":[1871,0,0,0,2272,0],"isMain":true},{"metricName":"Total length (>= 5000 bp)","quality":"More is better","values":[0,0,0,0,0,0],"isMain":false},{"metricName":"Total length (>= 10000 bp)","quality":"More is better","values":[0,0,0,0,0,0],"isMain":true},{"metricName":"Total length (>= 25000 bp)","quality":"More is better","values":[0,0,0,0,0,0],"isMain":false},{"metricName":"Total length (>= 50000 bp)","quality":"More is better","values":[0,0,0,0,0,0],"isMain":true},{"metricName":"N50","quality":"More is better","values":[1871,530,746,526,2272,609],"isMain":false},{"metricName":"N75","quality":"More is better","values":[1871,530,542,526,2272,609],"isMain":false},{"metricName":"L50","quality":"Less is better","values":[1,1,1,1,1,1],"isMain":false},{"metricName":"L75","quality":"Less is better","values":[1,1,2,1,1,1],"isMain":false},{"metricName":"GC (%)","quality":"Equal","values":["42.70","36.98","39.83","40.68","40.77","38.26"],"isMain":false}]],["K-mer-based statistics",[]],["Predicted genes",[]],["Similarity statistics",[]],["Reference statistics",[{"metricName":"Reference length","quality":"Equal","values":[2869885,2869885,2869885,2869885,2869885,2869885],"isMain":false},{"metricName":"Reference fragments","quality":"Equal","values":[32,32,32,32,32,32],"isMain":false},{"metricName":"Reference GC (%)","quality":"Equal","values":["35.27","35.27","35.27","35.27","35.27","35.27"],"isMain":false}]]],"subreferences":[],"subreports":[],"minContig":500},{"date":"10 February 2020, Monday, 20:37:08","assembliesNames":["megahit-103-df","megahit-113-df","megahit-113-ml","megahit-113-ms","megahit-129-df","metaSPAdes"],"referenceName":"GCF\_000424125.1\_ASM42412v1","order":[0,1,2,3,4,5],"report":[["Genome statistics",[{"metricName":"Genome fraction (%)","quality":"More is better","values":["96.194","99.290","99.590","99.580","99.577","99.454"],"isMain":true},{"metricName":"Duplication ratio","quality":"Less is better","values":["1.020","1.004","1.005","1.009","1.010","1.000"],"isMain":true},{"metricName":"Largest alignment","quality":"More is better","values":[25268,904953,859640,575870,394968,635050],"isMain":true},{"metricName":"Total aligned length","quality":"More is better","values":[4470448,4557068,4573836,4589700,4596148,4550083],"isMain":true},{"metricName":"NG50","quality":"More is better","values":[5057,158273,107076,132389,102696,199253],"isMain":false},{"metricName":"NG75","quality":"More is better","values":[2697,70166,49131,71936,45435,77341],"isMain":false},{"metricName":"NA50","quality":"More is better","values":[5092,158273,93256,96865,85855,199253],"isMain":false},{"metricName":"NA75","quality":"More is better","values":[2749,63374,47597,38877,38877,75475],"isMain":false},{"metricName":"NGA50","quality":"More is better","values":[5057,158273,93256,111924,91306,199253],"isMain":true},{"metricName":"NGA75","quality":"More is better","values":[2677,63374,48927,48299,41557,75475],"isMain":false},{"metricName":"LG50","quality":"Less is better","values":[285,9,10,10,11,8],"isMain":false},{"metricName":"LG75","quality":"Less is better","values":[594,20,27,22,27,17],"isMain":false},{"metricName":"LA50","quality":"Less is better","values":[281,9,11,12,13,8],"isMain":false},{"metricName":"LA75","quality":"Less is better","values":[582,22,29,32,34,17],"isMain":false},{"metricName":"LGA50","quality":"Less is better","values":[285,9,11,10,12,8],"isMain":true},{"metricName":"LGA75","quality":"Less is better","values":[595,22,28,25,32,17],"isMain":false}]],["Reads mapping",[]],["Misassemblies",[{"metricName":"# misassemblies","quality":"Less is better","values":[2,10,8,15,17,3],"isMain":true},{"metricName":" # relocations","quality":"Less is better","values":[0,1,1,5,3,0],"isMain":false},{"metricName":" # translocations","quality":"Less is better","values":[2,9,7,9,13,3],"isMain":false},{"metricName":" # inversions","quality":"Less is better","values":[0,0,0,1,1,0],"isMain":false},{"metricName":" # interspecies translocations","quality":"Less is better","values":[96,12,16,27,22,2],"isMain":false},{"metricName":"# misassembled contigs","quality":"Less is better","values":[2,10,8,15,15,3],"isMain":false},{"metricName":"Misassembled contigs length","quality":"Less is better","values":[5081,605476,547428,572868,939370,140695],"isMain":true},{"metricName":"# possibly misassembled contigs","quality":"Less is better","values":[0,0,0,0,0,0],"isMain":false},{"metricName":" # possible misassemblies","quality":"Less is better","values":[0,0,0,0,0,0],"isMain":false},{"metricName":"# local misassemblies","quality":"Less is better","values":[1,1,1,4,3,0],"isMain":false},{"metricName":"# scaffold gap ext. mis.","quality":"Less is better","values":[0,0,0,0,0,0],"isMain":false},{"metricName":"# scaffold gap loc. mis.","quality":"Less is better","values":[0,0,0,0,0,0],"isMain":false},{"metricName":"# unaligned mis. contigs","quality":"Less is better","values":[0,1,0,2,1,0],"isMain":false}]],["Unaligned",[{"metricName":"# fully unaligned contigs","quality":"Less is better","values":[1,0,2,2,0,0],"isMain":false},{"metricName":"Fully unaligned length","quality":"Less is better","values":[577,0,1440,1042,0,0],"isMain":false},{"metricName":"# partially unaligned contigs","quality":"Less is better","values":[48,6,9,21,10,1],"isMain":false},{"metricName":"Partially unaligned length","quality":"Less is better","values":[38643,8828,12242,339792,116161,1016],"isMain":false}]],["Mismatches",[{"metricName":"# mismatches","quality":"Less is better","values":[6598,622,618,2148,2129,323],"isMain":false},{"metricName":"# indels","quality":"Less is better","values":[4,13,16,72,68,6],"isMain":false},{"metricName":"Indels length","quality":"Less is better","values":[12,376,538,781,1050,14],"isMain":false},{"metricName":"# mismatches per 100 kbp","quality":"Less is better","values":["149.96","13.70","13.57","47.16","46.75","7.10"],"isMain":true},{"metricName":"# indels per 100 kbp","quality":"Less is better","values":["0.09","0.29","0.35","1.58","1.49","0.13"],"isMain":true},{"metricName":" # indels (<= 5 bp)","quality":"Less is better","values":[3,6,6,56,50,5],"isMain":false},{"metricName":" # indels (> 5 bp)","quality":"Less is better","values":[1,7,10,16,18,1],"isMain":false},{"metricName":"# N's","quality":"Less is better","values":[0,0,0,0,0,0],"isMain":false},{"metricName":"# N's per 100 kbp","quality":"Less is better","values":["0.00","0.00","0.00","0.00","0.00","0.00"],"isMain":true}]],["Statistics without reference",[{"metricName":"# contigs","quality":"Equal","values":[1430,107,138,155,170,72],"isMain":true},{"metricName":"# contigs (>= 1000 bp)","quality":"Equal","values":[1082,81,104,107,104,71],"isMain":false},{"metricName":"# contigs (>= 5000 bp)","quality":"Equal","values":[294,64,76,79,82,51],"isMain":false},{"metricName":"# contigs (>= 10000 bp)","quality":"Equal","values":[58,52,61,62,69,46],"isMain":false},{"metricName":"# contigs (>= 25000 bp)","quality":"Equal","values":[1,36,43,44,49,34],"isMain":false},{"metricName":"# contigs (>= 50000 bp)","quality":"Equal","values":[0,24,26,28,24,25],"isMain":false},{"metricName":"Largest contig","quality":"More is better","values":[25268,904953,859640,609940,394968,635058],"isMain":true},{"metricName":"Total length","quality":"More is better","values":[4527631,4569192,4591291,4937374,4717544,4551267],"isMain":true},{"metricName":"Total length (>= 1000 bp)","quality":"More is better","values":[4281243,4553144,4570327,4908143,4679542,4550728],"isMain":true},{"metricName":"Total length (>= 5000 bp)","quality":"More is better","values":[2333793,4511456,4508038,4843452,4627152,4505358],"isMain":false},{"metricName":"Total length (>= 10000 bp)","quality":"More is better","values":[760267,4416225,4391394,4715935,4534952,4463364],"isMain":true},{"metricName":"Total length (>= 25000 bp)","quality":"More is better","values":[25268,4134412,4097826,4411744,4221521,4270063],"isMain":false},{"metricName":"Total length (>= 50000 bp)","quality":"More is better","values":[0,3672698,3430035,3832699,3290529,3954387],"isMain":true},{"metricName":"N50","quality":"More is better","values":[5092,158273,107076,116839,102696,199253],"isMain":false},{"metricName":"N75","quality":"More is better","values":[2751,70166,49131,56920,43786,77341],"isMain":false},{"metricName":"L50","quality":"Less is better","values":[281,9,10,11,11,8],"isMain":false},{"metricName":"L75","quality":"Less is better","values":[581,20,27,26,30,17],"isMain":false},{"metricName":"GC (%)","quality":"Equal","values":["41.69","41.67","41.68","41.69","41.64","41.66"],"isMain":false}]],["K-mer-based statistics",[]],["Predicted genes",[]],["Similarity statistics",[]],["Reference statistics",[{"metricName":"Reference length","quality":"Equal","values":[4573762,4573762,4573762,4573762,4573762,4573762],"isMain":false},{"metricName":"Reference fragments","quality":"Equal","values":[46,46,46,46,46,46],"isMain":false},{"metricName":"Reference GC (%)","quality":"Equal","values":["41.67","41.67","41.67","41.67","41.67","41.67"],"isMain":false}]]],"subreferences":[],"subreports":[],"minContig":500},{"date":"10 February 2020, Monday, 20:36:50","assembliesNames":["megahit-103-df","megahit-113-df","megahit-113-ml","megahit-113-ms","megahit-129-df","metaSPAdes"],"referenceName":"GCF\_000424305.1\_ASM42430v1","order":[0,1,2,3,4,5],"report":[["Genome statistics",[{"metricName":"Genome fraction (%)","quality":"More is better","values":["0.042","0.022","0.211","0.139","0.065","0.025"],"isMain":true},{"metricName":"Duplication ratio","quality":"Less is better","values":["2.040","1.703","1.089","1.299","1.397","1.551"],"isMain":true},{"metricName":"Largest alignment","quality":"More is better","values":[344,452,3536,1631,638,491],"isMain":true},{"metricName":"Total aligned length","quality":"More is better","values":[1237,653,6149,4051,1887,721],"isMain":true},{"metricName":"NA50","quality":"More is better","values":[null,201,576,560,null,230],"isMain":false},{"metricName":"NGA50","quality":"More is better","values":[null,null,null,null,null,null],"isMain":true},{"metricName":"LA50","quality":"Less is better","values":[null,2,3,4,null,2],"isMain":false}]],["Reads mapping",[]],["Misassemblies",[{"metricName":"# misassemblies","quality":"Less is better","values":[1,0,0,0,0,0],"isMain":true},{"metricName":" # relocations","quality":"Less is better","values":[1,0,0,0,0,0],"isMain":false},{"metricName":" # translocations","quality":"Less is better","values":[0,0,0,0,0,0],"isMain":false},{"metricName":" # inversions","quality":"Less is better","values":[0,0,0,0,0,0],"isMain":false},{"metricName":" # interspecies translocations","quality":"Less is better","values":[5,1,3,3,3,1],"isMain":false},{"metricName":"# misassembled contigs","quality":"Less is better","values":[1,0,0,0,0,0],"isMain":false},{"metricName":"Misassembled contigs length","quality":"Less is better","values":[500,0,0,0,0,0],"isMain":true},{"metricName":"# possibly misassembled contigs","quality":"Less is better","values":[0,0,0,1,0,0],"isMain":false},{"metricName":" # possible misassemblies","quality":"Less is better","values":[0,0,0,1,0,0],"isMain":false},{"metricName":"# local misassemblies","quality":"Less is better","values":[0,0,0,0,0,0],"isMain":false},{"metricName":"# scaffold gap ext. mis.","quality":"Less is better","values":[0,0,0,0,0,0],"isMain":false},{"metricName":"# scaffold gap loc. mis.","quality":"Less is better","values":[0,0,0,0,0,0],"isMain":false},{"metricName":"# unaligned mis. contigs","quality":"Less is better","values":[0,0,0,0,0,0],"isMain":false}]],["Unaligned",[{"metricName":"# fully unaligned contigs","quality":"Less is better","values":[0,0,1,0,0,0],"isMain":false},{"metricName":"Fully unaligned length","quality":"Less is better","values":[0,0,727,0,0,0],"isMain":false},{"metricName":"# partially unaligned contigs","quality":"Less is better","values":[1,0,1,1,1,0],"isMain":false},{"metricName":"Partially unaligned length","quality":"Less is better","values":[1808,0,1147,852,1748,0],"isMain":false}]],["Mismatches",[{"metricName":"# mismatches","quality":"Less is better","values":[42,25,123,119,74,26],"isMain":false},{"metricName":"# indels","quality":"Less is better","values":[2,0,2,1,3,1],"isMain":false},{"metricName":"Indels length","quality":"Less is better","values":[2,0,3,1,3,1],"isMain":false},{"metricName":"# mismatches per 100 kbp","quality":"Less is better","values":["3395.31","3828.48","2001.30","2938.27","3923.65","3611.11"],"isMain":true},{"metricName":"# indels per 100 kbp","quality":"Less is better","values":["161.68","0.00","32.54","24.69","159.07","138.89"],"isMain":true},{"metricName":" # indels (<= 5 bp)","quality":"Less is better","values":[2,0,2,1,3,1],"isMain":false},{"metricName":" # indels (> 5 bp)","quality":"Less is better","values":[0,0,0,0,0,0],"isMain":false},{"metricName":"# N's","quality":"Less is better","values":[0,0,0,0,0,0],"isMain":false},{"metricName":"# N's per 100 kbp","quality":"Less is better","values":["0.00","0.00","0.00","0.00","0.00","0.00"],"isMain":true}]],["Statistics without reference",[{"metricName":"# contigs","quality":"Equal","values":[5,2,8,8,5,2],"isMain":true},{"metricName":"# contigs (>= 1000 bp)","quality":"Equal","values":[1,0,2,2,1,0],"isMain":false},{"metricName":"# contigs (>= 5000 bp)","quality":"Equal","values":[0,0,0,0,0,0],"isMain":false},{"metricName":"# contigs (>= 10000 bp)","quality":"Equal","values":[0,0,0,0,0,0],"isMain":false},{"metricName":"# contigs (>= 25000 bp)","quality":"Equal","values":[0,0,0,0,0,0],"isMain":false},{"metricName":"# contigs (>= 50000 bp)","quality":"Equal","values":[0,0,0,0,0,0],"isMain":false},{"metricName":"Largest contig","quality":"More is better","values":[2152,572,3536,1635,2092,608],"isMain":true},{"metricName":"Total length","quality":"More is better","values":[4332,1112,8567,6113,4382,1117],"isMain":true},{"metricName":"Total length (>= 1000 bp)","quality":"More is better","values":[2152,0,4904,2739,2092,0],"isMain":true},{"metricName":"Total length (>= 5000 bp)","quality":"More is better","values":[0,0,0,0,0,0],"isMain":false},{"metricName":"Total length (>= 10000 bp)","quality":"More is better","values":[0,0,0,0,0,0],"isMain":true},{"metricName":"Total length (>= 25000 bp)","quality":"More is better","values":[0,0,0,0,0,0],"isMain":false},{"metricName":"Total length (>= 50000 bp)","quality":"More is better","values":[0,0,0,0,0,0],"isMain":true},{"metricName":"N50","quality":"More is better","values":[566,572,1368,583,655,608],"isMain":false},{"metricName":"N75","quality":"More is better","values":[561,540,617,566,566,509],"isMain":false},{"metricName":"L50","quality":"Less is better","values":[2,1,2,3,2,1],"isMain":false},{"metricName":"L75","quality":"Less is better","values":[3,2,5,6,3,2],"isMain":false},{"metricName":"GC (%)","quality":"Equal","values":["40.54","43.35","39.64","39.21","39.09","38.59"],"isMain":false}]],["K-mer-based statistics",[]],["Predicted genes",[]],["Similarity statistics",[]],["Reference statistics",[{"metricName":"Reference length","quality":"Equal","values":[2911787,2911787,2911787,2911787,2911787,2911787],"isMain":false},{"metricName":"Reference fragments","quality":"Equal","values":[22,22,22,22,22,22],"isMain":false},{"metricName":"Reference GC (%)","quality":"Equal","values":["38.48","38.48","38.48","38.48","38.48","38.48"],"isMain":false}]]],"subreferences":[],"subreports":[],"minContig":500},{"date":"10 February 2020, Monday, 20:37:35","assembliesNames":["megahit-103-df","megahit-113-df","megahit-113-ml","megahit-113-ms","megahit-129-df","metaSPAdes"],"referenceName":"GCF\_000424365.1\_ASM42436v1","order":[0,1,2,3,4,5],"report":[["Genome statistics",[{"metricName":"Genome fraction (%)","quality":"More is better","values":["98.822","99.472","99.215","99.265","99.297","98.714"],"isMain":true},{"metricName":"Duplication ratio","quality":"Less is better","values":["1.006","1.004","1.005","1.005","1.007","1.000"],"isMain":true},{"metricName":"Largest alignment","quality":"More is better","values":[82175,167441,167413,167441,167504,428461],"isMain":true},{"metricName":"Total aligned length","quality":"More is better","values":[4062418,4081870,4072110,4075011,4083792,4037375],"isMain":true},{"metricName":"NG50","quality":"More is better","values":[22521,52897,51154,60949,53135,75219],"isMain":false},{"metricName":"NG75","quality":"More is better","values":[12705,29960,27708,35984,29960,42469],"isMain":false},{"metricName":"NA50","quality":"More is better","values":[22453,47976,49884,45171,44656,66994],"isMain":false},{"metricName":"NA75","quality":"More is better","values":[12701,26448,25918,21242,25847,35692],"isMain":false},{"metricName":"NGA50","quality":"More is better","values":[22453,47976,49884,46160,44656,66620],"isMain":true},{"metricName":"NGA75","quality":"More is better","values":[12657,26448,25918,25070,25951,33274],"isMain":false},{"metricName":"LG50","quality":"Less is better","values":[62,26,27,22,27,13],"isMain":false},{"metricName":"LG75","quality":"Less is better","values":[123,52,54,43,51,30],"isMain":false},{"metricName":"LA50","quality":"Less is better","values":[62,27,29,30,30,14],"isMain":false},{"metricName":"LA75","quality":"Less is better","values":[123,56,59,66,61,33],"isMain":false},{"metricName":"LGA50","quality":"Less is better","values":[62,27,29,27,30,15],"isMain":true},{"metricName":"LGA75","quality":"Less is better","values":[124,56,59,57,60,35],"isMain":false}]],["Reads mapping",[]],["Misassemblies",[{"metricName":"# misassemblies","quality":"Less is better","values":[4,15,12,17,29,12],"isMain":true},{"metricName":" # relocations","quality":"Less is better","values":[0,1,1,0,1,0],"isMain":false},{"metricName":" # translocations","quality":"Less is better","values":[4,14,11,17,27,12],"isMain":false},{"metricName":" # inversions","quality":"Less is better","values":[0,0,0,0,1,0],"isMain":false},{"metricName":" # interspecies translocations","quality":"Less is better","values":[22,10,8,18,9,1],"isMain":false},{"metricName":"# misassembled contigs","quality":"Less is better","values":[4,14,12,16,27,8],"isMain":false},{"metricName":"Misassembled contigs length","quality":"Less is better","values":[43728,663957,618033,515835,1146206,839599],"isMain":true},{"metricName":"# possibly misassembled contigs","quality":"Less is better","values":[0,0,0,0,1,0],"isMain":false},{"metricName":" # possible misassemblies","quality":"Less is better","values":[0,0,0,0,1,0],"isMain":false},{"metricName":"# local misassemblies","quality":"Less is better","values":[0,1,0,3,1,0],"isMain":false},{"metricName":"# scaffold gap ext. mis.","quality":"Less is better","values":[0,0,0,0,0,0],"isMain":false},{"metricName":"# scaffold gap loc. mis.","quality":"Less is better","values":[0,0,0,0,0,0],"isMain":false},{"metricName":"# unaligned mis. contigs","quality":"Less is better","values":[0,0,0,0,0,0],"isMain":false}]],["Unaligned",[{"metricName":"# fully unaligned contigs","quality":"Less is better","values":[0,0,0,0,0,1],"isMain":false},{"metricName":"Fully unaligned length","quality":"Less is better","values":[0,0,0,0,0,500],"isMain":false},{"metricName":"# partially unaligned contigs","quality":"Less is better","values":[10,5,5,12,6,1],"isMain":false},{"metricName":"Partially unaligned length","quality":"Less is better","values":[11185,9951,17241,310807,36977,1133],"isMain":false}]],["Mismatches",[{"metricName":"# mismatches","quality":"Less is better","values":[6329,604,617,852,984,403],"isMain":false},{"metricName":"# indels","quality":"Less is better","values":[0,12,8,17,15,15],"isMain":false},{"metricName":"Indels length","quality":"Less is better","values":[0,118,118,237,391,16],"isMain":false},{"metricName":"# mismatches per 100 kbp","quality":"Less is better","values":["156.59","14.85","15.21","20.99","24.23","9.98"],"isMain":true},{"metricName":"# indels per 100 kbp","quality":"Less is better","values":["0.00","0.29","0.20","0.42","0.37","0.37"],"isMain":true},{"metricName":" # indels (<= 5 bp)","quality":"Less is better","values":[0,10,6,11,7,15],"isMain":false},{"metricName":" # indels (> 5 bp)","quality":"Less is better","values":[0,2,2,6,8,0],"isMain":false},{"metricName":"# N's","quality":"Less is better","values":[0,0,0,0,0,0],"isMain":false},{"metricName":"# N's per 100 kbp","quality":"Less is better","values":["0.00","0.00","0.00","0.00","0.00","0.00"],"isMain":true}]],["Statistics without reference",[{"metricName":"# contigs","quality":"Equal","values":[354,156,168,164,178,102],"isMain":true},{"metricName":"# contigs (>= 1000 bp)","quality":"Equal","values":[311,142,151,149,145,95],"isMain":false},{"metricName":"# contigs (>= 5000 bp)","quality":"Equal","values":[205,108,111,110,108,75],"isMain":false},{"metricName":"# contigs (>= 10000 bp)","quality":"Equal","values":[148,94,97,98,94,68],"isMain":false},{"metricName":"# contigs (>= 25000 bp)","quality":"Equal","values":[49,62,63,59,61,47],"isMain":false},{"metricName":"# contigs (>= 50000 bp)","quality":"Equal","values":[6,27,29,30,31,26],"isMain":false},{"metricName":"Largest contig","quality":"More is better","values":[82175,167441,167413,232668,167504,428461],"isMain":true},{"metricName":"Total length","quality":"More is better","values":[4078264,4095781,4093455,4392292,4125711,4039622],"isMain":true},{"metricName":"Total length (>= 1000 bp)","quality":"More is better","values":[4048983,4085856,4081841,4382472,4105254,4034874],"isMain":true},{"metricName":"Total length (>= 5000 bp)","quality":"More is better","values":[3770129,3995550,3975149,4274876,4002976,3991018],"isMain":false},{"metricName":"Total length (>= 10000 bp)","quality":"More is better","values":[3358958,3890847,3876640,4186128,3902581,3943170],"isMain":true},{"metricName":"Total length (>= 25000 bp)","quality":"More is better","values":[1740889,3349952,3308369,3526452,3338222,3564520],"isMain":false},{"metricName":"Total length (>= 50000 bp)","quality":"More is better","values":[386555,2128397,2151192,2533066,2291533,2900881],"isMain":true},{"metricName":"N50","quality":"More is better","values":[22521,52897,51154,57094,53135,76367],"isMain":false},{"metricName":"N75","quality":"More is better","values":[12737,29960,27708,26754,29903,42659],"isMain":false},{"metricName":"L50","quality":"Less is better","values":[62,26,27,24,27,12],"isMain":false},{"metricName":"L75","quality":"Less is better","values":[122,52,54,51,52,29],"isMain":false},{"metricName":"GC (%)","quality":"Equal","values":["41.66","41.64","41.67","41.67","41.68","41.64"],"isMain":false}]],["K-mer-based statistics",[]],["Predicted genes",[]],["Similarity statistics",[]],["Reference statistics",[{"metricName":"Reference length","quality":"Equal","values":[4089863,4089863,4089863,4089863,4089863,4089863],"isMain":false},{"metricName":"Reference fragments","quality":"Equal","values":[75,75,75,75,75,75],"isMain":false},{"metricName":"Reference GC (%)","quality":"Equal","values":["41.62","41.62","41.62","41.62","41.62","41.62"],"isMain":false}]]],"subreferences":[],"subreports":[],"minContig":500},{"date":"10 February 2020, Monday, 20:37:27","assembliesNames":["megahit-103-df","megahit-113-df","megahit-113-ml","megahit-113-ms","megahit-129-df","metaSPAdes"],"referenceName":"GCF\_000424405.1\_ASM42440v1","order":[0,1,2,3,4,5],"report":[["Genome statistics",[{"metricName":"Genome fraction (%)","quality":"More is better","values":["36.301","40.121","40.692","39.704","38.849","23.891"],"isMain":true},{"metricName":"Duplication ratio","quality":"Less is better","values":["1.044","1.042","1.069","1.077","1.056","1.037"],"isMain":true},{"metricName":"Largest alignment","quality":"More is better","values":[9278,20923,10744,7360,11228,13513],"isMain":true},{"metricName":"Total aligned length","quality":"More is better","values":[1753026,1931502,2011120,1971415,1894571,1144445],"isMain":true},{"metricName":"NA50","quality":"More is better","values":[1360,1962,1073,1102,1683,1224],"isMain":false},{"metricName":"NA75","quality":"More is better","values":[844,1057,715,727,893,794],"isMain":false},{"metricName":"NGA50","quality":"More is better","values":[null,null,null,null,null,null],"isMain":true},{"metricName":"LA50","quality":"Less is better","values":[398,275,575,543,339,278],"isMain":false},{"metricName":"LA75","quality":"Less is better","values":[818,619,1161,1106,735,580],"isMain":false}]],["Reads mapping",[]],["Misassemblies",[{"metricName":"# misassemblies","quality":"Less is better","values":[21,28,30,29,50,5],"isMain":true},{"metricName":" # relocations","quality":"Less is better","values":[3,10,7,7,16,0],"isMain":false},{"metricName":" # translocations","quality":"Less is better","values":[14,16,18,18,31,5],"isMain":false},{"metricName":" # inversions","quality":"Less is better","values":[4,2,5,4,3,0],"isMain":false},{"metricName":" # interspecies translocations","quality":"Less is better","values":[41,54,43,55,74,18],"isMain":false},{"metricName":"# misassembled contigs","quality":"Less is better","values":[21,27,29,28,47,5],"isMain":false},{"metricName":"Misassembled contigs length","quality":"Less is better","values":[40928,89789,49403,45906,141842,7834],"isMain":true},{"metricName":"# possibly misassembled contigs","quality":"Less is better","values":[0,0,1,0,0,0],"isMain":false},{"metricName":" # possible misassemblies","quality":"Less is better","values":[0,0,1,0,0,0],"isMain":false},{"metricName":"# local misassemblies","quality":"Less is better","values":[10,31,14,11,21,5],"isMain":false},{"metricName":"# scaffold gap ext. mis.","quality":"Less is better","values":[0,0,0,0,0,0],"isMain":false},{"metricName":"# scaffold gap loc. mis.","quality":"Less is better","values":[0,0,0,0,0,0],"isMain":false},{"metricName":"# unaligned mis. contigs","quality":"Less is better","values":[3,3,0,0,0,0],"isMain":false}]],["Unaligned",[{"metricName":"# fully unaligned contigs","quality":"Less is better","values":[0,0,1,0,0,0],"isMain":false},{"metricName":"Fully unaligned length","quality":"Less is better","values":[0,0,653,0,0,0],"isMain":false},{"metricName":"# partially unaligned contigs","quality":"Less is better","values":[11,18,14,8,14,7],"isMain":false},{"metricName":"Partially unaligned length","quality":"Less is better","values":[23413,45817,11885,17642,34933,7395],"isMain":false}]],["Mismatches",[{"metricName":"# mismatches","quality":"Less is better","values":[19305,21370,32780,32858,22026,11566],"isMain":false},{"metricName":"# indels","quality":"Less is better","values":[170,300,188,196,243,111],"isMain":false},{"metricName":"Indels length","quality":"Less is better","values":[918,1572,680,1010,1191,712],"isMain":false},{"metricName":"# mismatches per 100 kbp","quality":"Less is better","values":["1140.75","1142.53","1727.95","1775.16","1216.15","1038.45"],"isMain":true},{"metricName":"# indels per 100 kbp","quality":"Less is better","values":["10.05","16.04","9.91","10.59","13.42","9.97"],"isMain":true},{"metricName":" # indels (<= 5 bp)","quality":"Less is better","values":[142,258,161,167,211,91],"isMain":false},{"metricName":" # indels (> 5 bp)","quality":"Less is better","values":[28,42,27,29,32,20],"isMain":false},{"metricName":"# N's","quality":"Less is better","values":[0,0,0,0,0,0],"isMain":false},{"metricName":"# N's per 100 kbp","quality":"Less is better","values":["0.00","0.00","0.00","0.00","0.00","0.00"],"isMain":true}]],["Statistics without reference",[{"metricName":"# contigs","quality":"Equal","values":[1454,1207,1957,1872,1373,1011],"isMain":true},{"metricName":"# contigs (>= 1000 bp)","quality":"Equal","values":[652,664,679,675,662,391],"isMain":false},{"metricName":"# contigs (>= 5000 bp)","quality":"Equal","values":[12,49,7,9,31,11],"isMain":false},{"metricName":"# contigs (>= 10000 bp)","quality":"Equal","values":[0,10,1,1,3,2],"isMain":false},{"metricName":"# contigs (>= 25000 bp)","quality":"Equal","values":[0,0,0,0,0,0],"isMain":false},{"metricName":"# contigs (>= 50000 bp)","quality":"Equal","values":[0,0,0,0,0,0],"isMain":false},{"metricName":"Largest contig","quality":"More is better","values":[9370,20923,10770,11724,18101,13564],"isMain":true},{"metricName":"Total length","quality":"More is better","values":[1789976,1994700,2041269,2012006,1947314,1162272],"isMain":true},{"metricName":"Total length (>= 1000 bp)","quality":"More is better","values":[1223091,1613642,1151043,1177400,1455681,718616],"isMain":true},{"metricName":"Total length (>= 5000 bp)","quality":"More is better","values":[80402,397153,53472,55667,228906,85015],"isMain":false},{"metricName":"Total length (>= 10000 bp)","quality":"More is better","values":[0,129678,10770,11724,39814,25464],"isMain":true},{"metricName":"Total length (>= 25000 bp)","quality":"More is better","values":[0,0,0,0,0,0],"isMain":false},{"metricName":"Total length (>= 50000 bp)","quality":"More is better","values":[0,0,0,0,0,0],"isMain":true},{"metricName":"N50","quality":"More is better","values":[1419,2171,1116,1159,1809,1267],"isMain":false},{"metricName":"N75","quality":"More is better","values":[886,1201,739,763,994,824],"isMain":false},{"metricName":"L50","quality":"Less is better","values":[377,246,556,516,305,269],"isMain":false},{"metricName":"L75","quality":"Less is better","values":[778,557,1120,1056,667,560],"isMain":false},{"metricName":"GC (%)","quality":"Equal","values":["39.14","39.21","39.45","39.47","39.17","38.80"],"isMain":false}]],["K-mer-based statistics",[]],["Predicted genes",[]],["Similarity statistics",[]],["Reference statistics",[{"metricName":"Reference length","quality":"Equal","values":[4661921,4661921,4661921,4661921,4661921,4661921],"isMain":false},{"metricName":"Reference fragments","quality":"Equal","values":[91,91,91,91,91,91],"isMain":false},{"metricName":"Reference GC (%)","quality":"Equal","values":["39.85","39.85","39.85","39.85","39.85","39.85"],"isMain":false}]]],"subreferences":[],"subreports":[],"minContig":500},{"date":"10 February 2020, Monday, 20:38:21","assembliesNames":["megahit-103-df","megahit-113-df","megahit-113-ml","megahit-113-ms","megahit-129-df","metaSPAdes"],"referenceName":"GCF\_000424425.1\_ASM42442v1","order":[0,1,2,3,4,5],"report":[["Genome statistics",[{"metricName":"Genome fraction (%)","quality":"More is better","values":["84.888","80.404","82.279","83.166","86.700","88.872"],"isMain":true},{"metricName":"Duplication ratio","quality":"Less is better","values":["1.057","1.056","1.112","1.108","1.067","1.023"],"isMain":true},{"metricName":"Largest alignment","quality":"More is better","values":[23406,31076,11957,13588,27798,27986],"isMain":true},{"metricName":"Total aligned length","quality":"More is better","values":[3859184,3640964,3926071,3946652,3965928,3912714],"isMain":true},{"metricName":"NG50","quality":"More is better","values":[2696,4441,1981,2385,4857,4398],"isMain":false},{"metricName":"NG75","quality":"More is better","values":[1341,2065,1104,1297,2481,2079],"isMain":false},{"metricName":"NA50","quality":"More is better","values":[2917,4440,2004,2279,4733,4777],"isMain":false},{"metricName":"NA75","quality":"More is better","values":[1690,2348,1170,1250,2481,2531],"isMain":false},{"metricName":"NGA50","quality":"More is better","values":[2654,3891,1859,2157,4369,4303],"isMain":true},{"metricName":"NGA75","quality":"More is better","values":[1290,1527,1009,1131,2202,2014],"isMain":false},{"metricName":"LG50","quality":"Less is better","values":[465,265,671,542,250,276],"isMain":false},{"metricName":"LG75","quality":"Less is better","values":[1020,616,1402,1154,566,637],"isMain":false},{"metricName":"LA50","quality":"Less is better","values":[398,256,649,570,250,239],"isMain":false},{"metricName":"LA75","quality":"Less is better","values":[832,549,1313,1185,553,525],"isMain":false},{"metricName":"LGA50","quality":"Less is better","values":[473,313,721,611,275,280],"isMain":true},{"metricName":"LGA75","quality":"Less is better","values":[1045,737,1505,1300,624,650],"isMain":false}]],["Reads mapping",[]],["Misassemblies",[{"metricName":"# misassemblies","quality":"Less is better","values":[13,34,25,24,36,7],"isMain":true},{"metricName":" # relocations","quality":"Less is better","values":[1,3,1,2,3,2],"isMain":false},{"metricName":" # translocations","quality":"Less is better","values":[8,27,20,19,30,5],"isMain":false},{"metricName":" # inversions","quality":"Less is better","values":[4,4,4,3,3,0],"isMain":false},{"metricName":" # interspecies translocations","quality":"Less is better","values":[32,52,55,69,75,18],"isMain":false},{"metricName":"# misassembled contigs","quality":"Less is better","values":[13,27,24,24,31,7],"isMain":false},{"metricName":"Misassembled contigs length","quality":"Less is better","values":[39594,145610,52953,45570,152898,40182],"isMain":true},{"metricName":"# possibly misassembled contigs","quality":"Less is better","values":[0,1,0,0,0,1],"isMain":false},{"metricName":" # possible misassemblies","quality":"Less is better","values":[0,2,0,0,0,2],"isMain":false},{"metricName":"# local misassemblies","quality":"Less is better","values":[6,12,12,19,23,7],"isMain":false},{"metricName":"# scaffold gap ext. mis.","quality":"Less is better","values":[0,0,0,0,0,0],"isMain":false},{"metricName":"# scaffold gap loc. mis.","quality":"Less is better","values":[0,0,0,0,0,0],"isMain":false},{"metricName":"# unaligned mis. contigs","quality":"Less is better","values":[0,2,3,3,2,0],"isMain":false}]],["Unaligned",[{"metricName":"# fully unaligned contigs","quality":"Less is better","values":[0,0,0,0,0,0],"isMain":false},{"metricName":"Fully unaligned length","quality":"Less is better","values":[0,0,0,0,0,0],"isMain":false},{"metricName":"# partially unaligned contigs","quality":"Less is better","values":[16,18,16,17,26,13],"isMain":false},{"metricName":"Partially unaligned length","quality":"Less is better","values":[21482,172950,87771,155732,102127,13299],"isMain":false}]],["Mismatches",[{"metricName":"# mismatches","quality":"Less is better","values":[23510,25273,43550,44017,24392,14902],"isMain":false},{"metricName":"# indels","quality":"Less is better","values":[296,544,370,390,452,252],"isMain":false},{"metricName":"Indels length","quality":"Less is better","values":[1276,3102,1696,1478,2569,1163],"isMain":false},{"metricName":"# mismatches per 100 kbp","quality":"Less is better","values":["641.81","728.42","1226.60","1226.53","651.98","388.58"],"isMain":true},{"metricName":"# indels per 100 kbp","quality":"Less is better","values":["8.08","15.68","10.42","10.87","12.08","6.57"],"isMain":true},{"metricName":" # indels (<= 5 bp)","quality":"Less is better","values":[255,458,322,345,370,220],"isMain":false},{"metricName":" # indels (> 5 bp)","quality":"Less is better","values":[41,86,48,45,82,32],"isMain":false},{"metricName":"# N's","quality":"Less is better","values":[0,0,0,0,0,0],"isMain":false},{"metricName":"# N's per 100 kbp","quality":"Less is better","values":["0.00","0.00","0.00","0.00","0.00","0.00"],"isMain":true}]],["Statistics without reference",[{"metricName":"# contigs","quality":"Equal","values":[1777,1155,2451,2220,1254,1222],"isMain":true},{"metricName":"# contigs (>= 1000 bp)","quality":"Equal","values":[1262,895,1537,1450,942,989],"isMain":false},{"metricName":"# contigs (>= 5000 bp)","quality":"Equal","values":[138,215,53,87,238,231],"isMain":false},{"metricName":"# contigs (>= 10000 bp)","quality":"Equal","values":[15,47,4,11,53,47],"isMain":false},{"metricName":"# contigs (>= 25000 bp)","quality":"Equal","values":[0,4,1,3,2,2],"isMain":false},{"metricName":"# contigs (>= 50000 bp)","quality":"Equal","values":[0,1,1,0,1,0],"isMain":false},{"metricName":"Largest contig","quality":"More is better","values":[23421,80712,54142,46382,66297,27986],"isMain":true},{"metricName":"Total length","quality":"More is better","values":[3893399,3835420,4037154,4132705,4093469,3938167],"isMain":true},{"metricName":"Total length (>= 1000 bp)","quality":"More is better","values":[3516715,3647671,3378851,3576331,3874023,3767454],"isMain":true},{"metricName":"Total length (>= 5000 bp)","quality":"More is better","values":[1000042,1925577,387374,691815,2099645,1948385],"isMain":false},{"metricName":"Total length (>= 10000 bp)","quality":"More is better","values":[187164,775939,97586,220929,837831,684617],"isMain":true},{"metricName":"Total length (>= 25000 bp)","quality":"More is better","values":[0,185842,54142,119557,94125,53852],"isMain":false},{"metricName":"Total length (>= 50000 bp)","quality":"More is better","values":[0,80712,54142,0,66297,0],"isMain":true},{"metricName":"N50","quality":"More is better","values":[2970,5017,2117,2487,5103,4889],"isMain":false},{"metricName":"N75","quality":"More is better","values":[1733,2815,1254,1429,2741,2599],"isMain":false},{"metricName":"L50","quality":"Less is better","values":[391,214,603,505,228,236],"isMain":false},{"metricName":"L75","quality":"Less is better","values":[814,466,1225,1053,502,515],"isMain":false},{"metricName":"GC (%)","quality":"Equal","values":["39.64","39.73","39.72","39.75","39.67","39.66"],"isMain":false}]],["K-mer-based statistics",[]],["Predicted genes",[]],["Similarity statistics",[]],["Reference statistics",[{"metricName":"Reference length","quality":"Equal","values":[4315150,4315150,4315150,4315150,4315150,4315150],"isMain":false},{"metricName":"Reference fragments","quality":"Equal","values":[68,68,68,68,68,68],"isMain":false},{"metricName":"Reference GC (%)","quality":"Equal","values":["39.67","39.67","39.67","39.67","39.67","39.67"],"isMain":false}]]],"subreferences":[],"subreports":[],"minContig":500},{"date":"10 February 2020, Monday, 20:38:38","assembliesNames":["megahit-103-df","megahit-113-df","megahit-113-ml","megahit-113-ms","megahit-129-df","metaSPAdes"],"referenceName":"GCF\_000424465.1\_ASM42446v1","order":[0,1,2,3,4,5],"report":[["Genome statistics",[{"metricName":"Genome fraction (%)","quality":"More is better","values":["95.312","96.464","96.952","95.868","95.499","95.548"],"isMain":true},{"metricName":"Duplication ratio","quality":"Less is better","values":["1.011","1.028","1.024","1.040","1.031","1.002"],"isMain":true},{"metricName":"Largest alignment","quality":"More is better","values":[84565,98404,101738,110589,101989,130739],"isMain":true},{"metricName":"Total aligned length","quality":"More is better","values":[3243447,3334049,3338180,3349696,3309030,3223697],"isMain":true},{"metricName":"NG50","quality":"More is better","values":[14776,22324,23633,33664,28939,27301],"isMain":false},{"metricName":"NG75","quality":"More is better","values":[6339,12476,11303,20068,16010,14660],"isMain":false},{"metricName":"NA50","quality":"More is better","values":[14776,17802,21034,15511,20112,28002],"isMain":false},{"metricName":"NA75","quality":"More is better","values":[6891,8143,8704,3080,7077,15956],"isMain":false},{"metricName":"NGA50","quality":"More is better","values":[13888,18488,21463,19939,23003,27301],"isMain":true},{"metricName":"NGA75","quality":"More is better","values":[5849,9337,9475,8916,9081,14159],"isMain":false},{"metricName":"LG50","quality":"Less is better","values":[67,47,42,27,35,35],"isMain":false},{"metricName":"LG75","quality":"Less is better","values":[153,96,94,58,74,76],"isMain":false},{"metricName":"LA50","quality":"Less is better","values":[65,61,45,68,49,33],"isMain":false},{"metricName":"LA75","quality":"Less is better","values":[145,135,106,202,124,69],"isMain":false},{"metricName":"LGA50","quality":"Less is better","values":[69,55,44,47,43,35],"isMain":true},{"metricName":"LGA75","quality":"Less is better","values":[157,118,102,107,99,76],"isMain":false}]],["Reads mapping",[]],["Misassemblies",[{"metricName":"# misassemblies","quality":"Less is better","values":[6,12,8,13,13,2],"isMain":true},{"metricName":" # relocations","quality":"Less is better","values":[1,5,2,2,1,0],"isMain":false},{"metricName":" # translocations","quality":"Less is better","values":[5,7,6,11,12,2],"isMain":false},{"metricName":" # inversions","quality":"Less is better","values":[0,0,0,0,0,0],"isMain":false},{"metricName":" # interspecies translocations","quality":"Less is better","values":[10,37,19,55,36,1],"isMain":false},{"metricName":"# misassembled contigs","quality":"Less is better","values":[6,12,7,11,12,2],"isMain":false},{"metricName":"Misassembled contigs length","quality":"Less is better","values":[50058,305687,167192,197361,266496,24122],"isMain":true},{"metricName":"# possibly misassembled contigs","quality":"Less is better","values":[0,0,0,0,0,0],"isMain":false},{"metricName":" # possible misassemblies","quality":"Less is better","values":[0,0,0,0,0,0],"isMain":false},{"metricName":"# local misassemblies","quality":"Less is better","values":[4,8,8,10,11,0],"isMain":false},{"metricName":"# scaffold gap ext. mis.","quality":"Less is better","values":[0,0,0,0,0,0],"isMain":false},{"metricName":"# scaffold gap loc. mis.","quality":"Less is better","values":[0,0,0,0,0,0],"isMain":false},{"metricName":"# unaligned mis. contigs","quality":"Less is better","values":[0,8,1,7,5,0],"isMain":false}]],["Unaligned",[{"metricName":"# fully unaligned contigs","quality":"Less is better","values":[0,0,0,0,0,0],"isMain":false},{"metricName":"Fully unaligned length","quality":"Less is better","values":[0,0,0,0,0,0],"isMain":false},{"metricName":"# partially unaligned contigs","quality":"Less is better","values":[5,26,15,40,20,0],"isMain":false},{"metricName":"Partially unaligned length","quality":"Less is better","values":[22786,225797,77533,748372,324751,0],"isMain":false}]],["Mismatches",[{"metricName":"# mismatches","quality":"Less is better","values":[2670,6726,6207,11897,7236,3421],"isMain":false},{"metricName":"# indels","quality":"Less is better","values":[32,94,108,319,178,76],"isMain":false},{"metricName":"Indels length","quality":"Less is better","values":[373,689,922,1944,1762,171],"isMain":false},{"metricName":"# mismatches per 100 kbp","quality":"Less is better","values":["83.22","207.13","190.19","368.66","225.09","106.36"],"isMain":true},{"metricName":"# indels per 100 kbp","quality":"Less is better","values":["1.00","2.89","3.31","9.89","5.54","2.36"],"isMain":true},{"metricName":" # indels (<= 5 bp)","quality":"Less is better","values":[28,79,91,285,145,72],"isMain":false},{"metricName":" # indels (> 5 bp)","quality":"Less is better","values":[4,15,17,34,33,4],"isMain":false},{"metricName":"# N's","quality":"Less is better","values":[0,0,0,0,0,0],"isMain":false},{"metricName":"# N's per 100 kbp","quality":"Less is better","values":["0.00","0.00","0.00","0.00","0.00","0.00"],"isMain":true}]],["Statistics without reference",[{"metricName":"# contigs","quality":"Equal","values":[546,384,385,384,386,275],"isMain":true},{"metricName":"# contigs (>= 1000 bp)","quality":"Equal","values":[398,311,306,314,291,230],"isMain":false},{"metricName":"# contigs (>= 5000
